# An entropy-based diagnostic framework for characterizing methylation state dynamics during preimplantation development

**DOI:** 10.64898/2026.08.25.746714

**Authors:** Bingjie Hao, Yuxiang Cheng, Zhiyuan Liu

## Abstract

DNA methylation undergoes predictable changes with age, and preimplantation embryos are known to undergo global epigenetic reprogramming. However, the specific fate of age associated methylation signatures during early development has not been systematically quantified. Using published human sperm age associated differentially methylated regions (DMRs) as a feature space, we integrated single cell methylome and transcriptome data to develop the Transgenerational Reset Operator (TRO), a computational framework for profiling preimplantation stages. We found that the morula stage represents the nadir of age associated methylation entropy while retaining high developmental potency, distinguishing it from a simple demethylation endpoint. Dynamical modelling revealed that independent DMR drift fails to recapitulate the morula state, requiring a coordinated, structured correction concentrated in specific DMR subsets and modules with marked directional sensitivity. Independent chromatin accessibility data supported a stage specific methylation–accessibility coupling at morula, albeit with modest effect sizes. Cross species mouse and orthogonal multiomic evidence suggested partial conservation but with weight dependence and heterogeneity. Collectively, our study defines morula as a computational “ground zero” candidate for age associated methylation features and proposes a testable hypothesis of developmental regulation, while emphasizing that matched parental–offspring perturbation experiments are needed to establish causal mechanisms.

## Introduction

DNA methylation constitutes a prominent form of epigenetic regulation, modulating gene expression via methylation modifications at CpG dinucleotides and other genomic loci. To date, extensive studies have established a robust association between DNA methylation patterns and the aging process, as well as with the pathogenesis of age-related disorders. Among the most representative and widely cited contributions in this field is the first-generation “methylation clock” model originally proposed by Hannum and Horvath,^[1, 2]^ which predicts chronological age by measuring DNA methylation levels in whole blood and pan-tissue samples. This model demonstrates high predictive accuracy, with correlation coefficients between predicted and actual age reaching 0.96 and 0.97, respectively. Subsequent second- and third-generation methylation clocks have further elucidated the relationships between methylation profiles, biological age, and aging rates, indirectly corroborating that organismal lifespan is accompanied by corresponding dynamic changes in methylation status. ^[3-5]^Moreover, a statistical investigation of white blood cell methylomes across 421 subjects revealed that 29% of CpG sites undergo significant age-associated methylation alterations, of which 60.5% exhibit hypomethylation and 39.5% show hypermethylation. ^[6]^These findings partially delineate the genomic distribution characteristics of methylation changes with advancing age, supporting the premise that methylation modifications in distinct genomic regions tend to evolve directionally over time.

Building upon this paradigm, we extrapolate the developmental timeline backward and propose the following hypothesis that, given that ontogeny originates from the fusion of paternal sperm and maternal oocyte, and assuming that both parental and offspring genomic methylation levels follow the aforementioned age-dependent accumulation trends, the intergenerational transmission of methylation marks via gametes may involve a partial reprogramming process—namely, the erasure of parentally derived methylation signatures to reset the epigenetic landscape, thereby providing a de novo starting point for the offspring’s subsequent methylation accumulation. Consequently, our investigative focus transitioned from adult somatic tissues to embryonic development. A comprehensive review of the recent literature corroborated that this reprogramming phenomenon has been extensively documented. Prior to fertilization, sperm and oocytes exhibit CpG methylation levels of 80–90% and 40%, respectively. Upon fertilization, the paternal genome undergoes active demethylation mediated by TET3, whereas the maternal genome experiences passive demethylation progressively during cleavage divisions. By the morula-to-blastocyst transition, global methylation reaches its nadir, after which the genome initiates de novo methylation to complete the reprogramming cascade.^[7]^

Cellular physiological processes are governed by information flow according to the central dogma, and gene expression is inherently stochastic. Notably, increased transcriptional noise has been implicated in accelerated organismal senescence. ^[8]^Entropy, as a fundamental concept in information theory, quantifies the uncertainty associated with stochastic events. Importantly, empirical evidence indicates that an elevation in genomic methylation entropy is not necessarily accompanied by a corresponding increase in average methylation. Incorporation of single-site methylation entropy into age-prediction models has yielded lower prediction errors compared to methods utilizing a larger number of CpG sites, further supporting the notion that entropy may constitute a more biologically meaningful explanatory variable for lifespan trajectories. ^[9]^Therefore, in the construction of the mathematical model, we ultimately selected methylation entropy as the descriptor.

Although existing studies and computational models have successfully characterized or fitted the static methylation profiles at various stages of mammalian embryogenesis, these investigations remain largely descriptive and cross-sectional in nature. Our research, by contrast, seeks to elucidate the mechanistic underpinnings of the dynamic transitions between these stage-specific methylation states, with the ultimate goal of exploring the causal interrelationships among cellular methylation states across developmental time from a dynamical systems perspective. ^[10-12]^

Accordingly, our experimental design commenced with foundational static analyses, in which we independently derived a multi-modal methylation entropy metric, performed preliminary statistical assessments of entropy distributions during embryonic development, and examined the association between methylation entropy and developmental potential. Subsequently, we advanced to dynamical modeling, defining a first-layer framework grounded solely in methylation drift theory to characterize the numerical relationships between adjacent DMR (differentially methylated region) state spaces, and conducted calibration analyses to refine the model. Finally, cross-species examination was performed to evaluate the generalizability of our proposed framework. To date, our experiments have successfully uncovered both the static distributional patterns and the dynamical transition pathways of human embryonic methylomes, thereby constructing a diagnostic control framework. Nevertheless, it should be acknowledged that the current model exhibits certain species-specific constraints, and substantial further investigation is required to fully elucidate the ultimate causal chain—an objective that constitutes a primary direction for our future research endeavors.

## Results

We first asked whether the paternal-age-associated methylation signature exhibits a discrete minimum during human preimplantation development. The primary analysis used 204 technically valid methylome profiles from GSE81233, of which 172 represented the seven ordered stages from metaphase-II (MII) oocyte to blastocyst and the remainder represented inner-cell-mass and trophectoderm derivatives. After stage-specific coverage filtering, the age-weighted methylation entropy, denoted S_epi−a_g_e_, declined from 0.446 in MII oocytes to 0.429 in zygotes, 0.375 at the 2-cell stage and 0.324 at the 4-cell stage. This decline was interrupted at the 8-cell stage, where S_epi−a_g_e_ rose to 0.383, before falling to its global minimum of 0.278 at morula and increasing again to 0.313 in blastocysts (Fig.1A). Thus, the morula minimum was not the trivial endpoint of monotonic genome-wide demethylation; it emerged after a transient 8-cell increase and was followed by a rebound during blastocyst formation. Generic methylation entropy showed the same stage ordering at the minimum, but the age-weighted statistic provided the biologically prespecified coordinate used throughout the operator analysis.

**Fig. 1.**
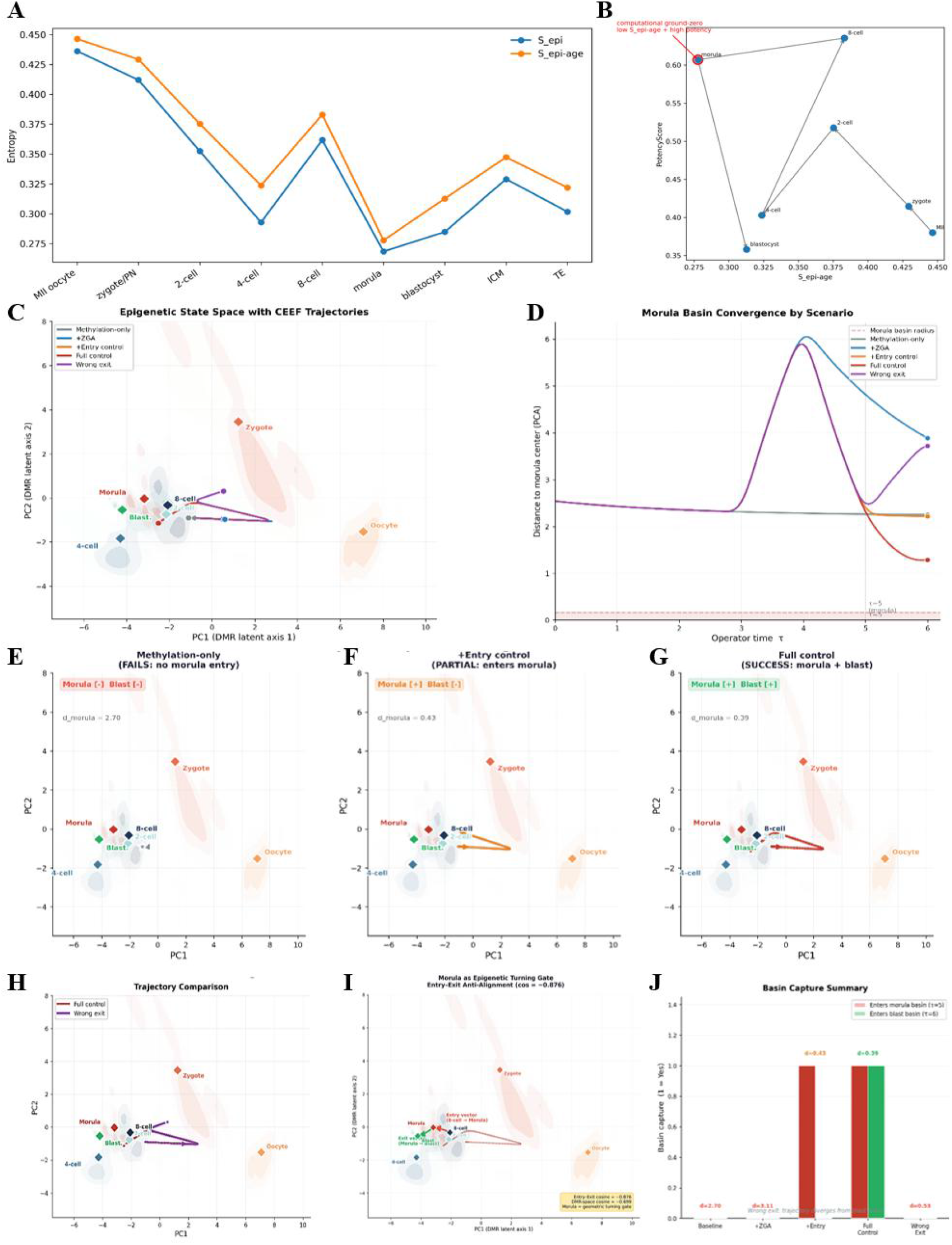
**A**. A stage-restricted minimum of age-associated methylation entropy in human preimplantation development. Stage-level generic methylation entropy and age-weighted methylation entropy in the technically valid GSE81233 cohort. Morula displays the lowest value of both summaries, with an 8-cell rise preceding the minimum and a blastocyst rebound following it. Error bars and sample-level dispersion are as defined in Methods. **B**.Joint epigenetic and transcriptomic state space identifies a high-potency, low-perturbation morula state. Dual-entropy phase map integrating age-associated methylation disorder with transcriptomic state. The 8-cell and morula stages occupy a high-potency region, but morula uniquely combines preserved potency with the lowest age-associated methylation entropy. The map is descriptive and does not imply lineage tracking of individual embryos. **C-D**.COMSOL operator-time landscape separates methylation-only failure from basin entry. Direct COMSOL-derived state-space trajectories and distance-to-morula profiles for methylation-only, ZGA-only, entry-control, full-control and wrong-exit numerical realization. The methylation-only and ZGA-only trajectories fail to enter the morula basin, whereas the complete control enters morula and subsequently approaches blastocyst. Basin boundaries and operator times were prespecified from the empirical latent-state distributions. **E-G**.Progressive COMSOL control reconstructs morula entry and blastocyst completion in the model. Direct comparison of the methylation-only solution, accessibility-gated entry control and full entry-plus-exit control. Entry control is sufficient for morula capture but not terminal progression; the full solution completes both transitions. Curves are numerical trajectories exported from the archived COMSOL 6.4 models. **H**.Wrong-direction exit causes counterfactual trajectory collapse. COMSOL comparison of the complete control and a sign-reversed exit operator. Correct control captures the morula basin and reaches blastocyst, whereas the wrong-exit trajectory diverges and fails terminal basin capture. **I**.COMSOL trajectories identify morula as an epigenetic turning gate in the fitted model. Entry from 8-cell and exit towards blastocyst are strongly anti-aligned in the correction-vector plane (cosine = −0.876) and in DMR space (cosine = −0.699). The direct numerical trajectory bends at the morula basin rather than extending along a single reset axis. **J**.The bar summary reports basin calls for all five simulated scenarios.

The morula call remained stable when we altered the effective DMR universe and sample balance. Morula was selected in 94.3% of bootstrap resamples under the primary specification and in 91.6% when the analysis was restricted to DMRs shared across stages. Varying the stage-level coverage threshold from 10% to 50% did not change the identity of the minimum. Balanced bootstraps with five or eight samples per stage selected morula in 76.8% and 85.9% of resamples, respectively, despite the marked differences in available sample numbers across development. By comparison, shuffling the age weights reduced the morula selection frequency to 81.7%, and random age-DMR subsets reduced it to 69.8%. We therefore interpret the result strictly as a stage-restricted minimum within the externally defined paternal-age-associated DMR feature space, not as evidence of specific erasure of age-associated methylation at morula. Because the datasets do not contain matched sperm and embryos from the same fathers, we interpret this result as an age-DMR-defined computational ground-zero candidate, rather than direct observation of paternal-age erasure in a paired human lineage.

The signed age projection separated low entropy from directional cancellation. Morula had the lowest signed projection among the seven preimplantation stages (0.056), compared with 0.125 at the 8-cell stage and 0.114 at blastocyst. The concordant minima in uncertainty and signed projection argue that morula is not merely a heterogeneous mixture whose positive and negative age-associated loci cancel in aggregate. Instead, the age-informed DMR state is simultaneously low in disorder and displaced away from the larger projections observed immediately before and after compaction. This distinction is central to the life-operator formulation: the candidate ground zero is defined by the joint organization of a prespecified perturbation coordinate, not by a conventional epigenetic clock or by generic methylation loss alone.

A ground-zero state should not be defined by methylation loss at the expense of developmental competence. We therefore analysed single-cell transcriptomes from GSE36552 using global RNA entropy, detected-gene richness and a curated potency-marker score. Global transcriptomic entropy did not peak at morula, and we did not require it to do so. Instead, morula retained a high marker score (0.824), essentially matching the 8-cell value (0.824) and substantially exceeding the blastocyst value (0.436). The composite developmental-potency score was 0.607 at morula, compared with 0.635 at the 8-cell stage and 0.358 in blastocysts. SOX2, KLF4, KLF17, NANOG and TFAP2C were the largest positive contributors to the morula marker signal. Leave-one-marker-out analyses preserved the high-potency 8-cell–morula region, showing that the result was not determined by a single canonical factor.

An independent RNA dataset, GSE44183, reproduced the ordering of this high-potency window: 8-cell ranked first and morula second among the seven tested stages, with potency scores of 0.587 and 0.508, respectively. We therefore regard the RNA evidence as support for preserved potency at morula, not for a uniquely maximal morula transcriptomic state. This distinction matters because methylation entropy and transcriptional diversity describe different aspects of developmental state. Their convergence is informative precisely because the morula minimum in age-associated methylation disorder occurs without a collapse of the molecular programme required for subsequent lineage formation.

Integration of the DNA and RNA coordinates resolved this convergence at the stage level (Fig.1B). Relative to 8-cell, morula moved sharply towards lower age-associated methylation entropy while retaining 95.5% of the maximum observed potency. Relative to blastocyst, morula combined lower age-associated entropy with a 69.5% higher potency score. The independent ground-zero score, obtained by standardizing low perturbation and high potency across stages, ranked morula first,which is consistent with the previously established single-cell transcriptomic atlas of human preimplantation development. ^[13, 14]^The operational Transgenerational Reset Operator (TRO) score, which multiplied the internally anchored reset score by potency preservation, likewise ranked morula first at 0.955. The next highest stages were 4-cell (0.461) and blastocyst (0.447), whereas the 8-cell stage scored 0.376 because its high potency coincided with an elevated perturbation burden. The agreement of two differently constructed rankings makes the ground-zero call less dependent on any single normalization.

We next treated development as an ordered sequence of transitions rather than a collection of stage means. These transitions are defined at the stage level and are coupled distributionally via optimal transport; they do not represent individual embryo lineage tracking, nor does the stage-anchored pseudotime imply equal physical durations between stages. In the standardized TRO state space, the 8-cell-to-morula transition reduced the damage coordinate by 0.105 and increased the internally anchored reset coordinate by 0.624, while incurring only a small reduction in potency (0.028). Its productive reset gain was 0.730 and its gain-to-displacement efficiency was 0.271, the highest of all six adjacent transitions. By contrast, the 4-cell-to-8-cell transition had the largest raw state-space displacement but increased the perturbation burden and therefore had low productive efficiency. The morula-to-blastocyst transition reduced potency by 0.249, increased the age-associated damage coordinate by 0.035 and produced no positive reset gain. These comparisons show why displacement magnitude alone is an inadequate descriptor: the decisive event is the directionally productive movement from the 8-cell state into the morula basin.

DMR-level decomposition showed that this movement was concentrated but not reducible to one locus. The 20 highest-ranked reset-driving DMRs accounted for 60.1% of the positive 8-cell-to-morula contribution, and the top 50 accounted for 91.8%. Nearest-gene annotations included PCDH17, WNT5B, JAKMIP2, ANK2, RASA3, GPANK1 and IRF9, among others. Conventional genomic-context enrichment did not yield a single dominant compartment after multiple-testing correction, and nearest-gene pathway assignments were correspondingly treated as exploratory. The robust conclusion is therefore architectural: a structured subset of age-informative DMRs carries most of the productive reset displacement, while the biological identities assigned to individual DMRs remain hypotheses for experimental follow-up.

The morula state also displayed an entry–exit geometry consistent with a developmental inflection. The entry vector from 8-cell to morula and the exit vector from morula to blastocyst produced a duality score of 0.699, exceeding the 95th percentile of a label-permutation null (0.138; empirical P < 0.001). Bootstrap resampling gave a 95% interval of 0.548–0.812, and all bootstrap estimates were positive. At morula, 78 of 156 DMRs (50%) were fully demethylated, the highest fraction among the seven stages, while the remaining DMRs retained a broad methylation distribution. This yielded the largest bimodality index (1.564), consistent with the interpretation that morula is not a uniformly erased state but a branch point at which one DMR subset reaches a floor while another retains information needed for the ensuing remethylation and differentiation programme.

To test whether the morula state could be predicted from preceding stages, we first fitted each DMR independently and withheld morula transitions. This single-DMR model produced a leave-morula-out RMSE of 0.311, worse than carrying forward the 8-cell mean (RMSE 0.297). Independent locus-wise drift is therefore insufficient to explain the observed morula configuration. Grouping DMRs into coordinated modules improved the withheld-morula RMSE to 0.287, and a low-dimensional latent-state model further reduced it to 0.270. Bootstrap comparisons favoured the latent representation over the independent-DMR model under both DMR resampling (95% interval for the RMSE difference, −0.0486 to −0.0046) and sample/stage resampling (−0.0351 to −0.0217). Paired squared-error tests supported the latent model (P = 0.0067) and, more modestly, the module model (P = 0.0381). Randomizing operator time or module assignments abolished the advantage (empirical P = 0.0050 and P = 0.0299, respectively).

The improvement did not depend convincingly on a privileged optimal-transport coupling. Random target couplings were not consistently worse, indicating that the gain arose mainly from coordinated representation rather than from inferred sample-to-sample correspondence. We therefore use optimal transport as a distribution-aware interpolation device, not as evidence of true embryo lineage tracking. Across broader rollouts, the latent model retained similar performance when morula was withheld (RMSE 0.252; correlation 0.598) and when fitting was restricted to stages through 4-cell (RMSE 0.260; correlation 0.579), compared with the full-stage fit (RMSE 0.247; correlation 0.620). The modest degradation supports a reproducible low-dimensional trajectory, while the remaining error indicates that an important component of morula attraction is absent from methylation-only dynamics.

The failure became explicit when the reduced operator was solved as a time-dependent vector-field model in COMSOL Multiphysics 6.4. Five prespecified scenarios were instantiated from the same initial condition: methylation-only baseline, baseline plus the zygotic-genome-activation component, entry control alone, complete entry-and-exit control and a wrong-direction exit counterfactual. In the two-dimensional correction-vector coordinate system, the methylation-only trajectory remained outside both target basins and was 2.700 standardized units from the morula centre at operator time 5 (Fig.1C-D). Adding the ZGA component alone did not correct the failure and increased the morula distance to 3.111. By contrast, the entry-control trajectory crossed the morula boundary at a distance of 0.430, and the complete-control trajectory entered morula at 0.391 before reaching the blastocyst basin at operator time 6. The wrong-exit trajectory approached the morula neighbourhood but subsequently diverged from blastocyst, ending 5.908 units from the terminal centre. All five scenario models, trajectories and basin calls were generated from archived ‘.mph’ files and corresponding time-series exports.

The COMSOL field solution also exposed a sequential organization in the fitted field that cannot be read from stage means alone (Fig.1E-G). At early operator time, the baseline attraction field organized cleavage-stage motion but lacked a morula-directed attractor. The ZGA reconstruction altered the early trajectory without creating morula capture. An accessibility-gated entry field then deflected the trajectory into the morula neighbourhood, whereas the later exit field was required to rotate the flow towards blastocyst. In the full-control solution, the morula-localized field formed a transient capture region rather than a terminal fixed point. The numerical solution therefore identifies morula as a controlled passage basin: within the fitted dynamics, entry and residence require a stage-restricted correction, and successful continuation requires a distinct exit operator.

The same conclusion held at the distribution level. The observed morula distribution had a basin occupancy of 0.875 under the prespecified 90th-percentile criterion. A deterministic methylation-only rollout achieved 0.378 occupancy when all stages were available but only 0.044 when morula transitions were excluded. Global residual noise, stage-conditioned noise and an affine transition kernel did not rescue the non-leaking prediction, yielding occupancies of 0.023, 0.085 and 0.016, respectively; moment matching yielded zero. An Ornstein–Uhlenbeck process calibrated directly on the observed morula distribution recovered high occupancy, but this constitutes a diagnostic upper bound because it uses the withheld target. These results separate two questions: methylation coordinates describe the observed basin, yet their autonomous pre-morula dynamics do not locate it.

The discrepancy defined a missing attraction vector from the strict pre-morula prediction to the observed morula centre. In standardized latent coordinates, this residual was largest along the third component and had Euclidean norm 2.075. Projecting it back to DMR space produced a sparse-to-distributed control architecture. Within the diagnostic framework, adding the top 10 residual DMRs raised occupancy to 0.600, the top 15 to 0.867 and the top 25 to 0.956; the top 50 reached 1.000. In the fitted model, matched random sets of 25 DMRs had a mean occupancy of 0.108 and a 95th percentile of 0.156, while sign-flipping the selected correction reduced occupancy to zero. Removing the top 25 DMRs still left 0.556 occupancy in the remainder, but removing the top 50 reduced it to 0.178. Thus, the missing attraction is strongly concentrated in a structured core while retaining a weaker distributed component.

Amplitude scans further revealed a nonlinear dose–occupancy relationship within the reconstructed model. Applying 10% of the correction produced 0.133 occupancy, 25% produced 0.422, 50% produced 0.933 and 75% produced 0.978; the full correction reached 1.000. At the module level, the leading module alone generated 0.422 occupancy, and sequential addition of the next most informative modules raised occupancy to 0.600, 0.867 and 0.956. This concordance between DMR- and module-level reconstructions argues that basin entry depends on coordinated direction and amplitude in the diagnostic framework, rather than the indiscriminate addition of methylation variance.

The COMSOL trajectory changed direction sharply at morula. In the correction-vector plane, the entry and exit directions had a cosine similarity of −0.876, while the corresponding DMR-space cosine was −0.699 (Fig.1I). Thus, the trajectory does not simply continue along a single demethylation axis. It is redirected at morula from an entry programme dominated by reset towards an exit programme in the fitted field. The field geometry is consistent with the DMR-level duality score and bootstrap analysis above, but makes the dynamical implication explicit: morula behaves as a turning gate whose entry and exit vectors are strongly anti-aligned.

The gate interpretation was challenged by reversing the exit field while leaving the rest of the model unchanged. The correct full-control solution entered both morula and blastocyst, whereas the wrong-exit solution failed the prespecified basin calls and diverged from the blastocyst centre (Fig.1H,1J). Entry control alone entered morula but failed blastocyst, demonstrating that morula capture and terminal completion are separable conditions.

We next tested the inferred correction as a counterfactual component of the operator rather than merely an error vector. Removing the correction term left the trajectory 2.700 standardized units from the morula centre at operator time 5 and outside the morula basin. Restoring the complete term reduced the distance to 0.391, a 6.90-fold rescue, brought the trajectory into the morula basin and allowed subsequent entry into the blastocyst basin. Entry-only control reached morula but failed to complete the blastocyst transition, whereas a directionally incorrect exit failed terminal progression. In an independent parameterization used for hardening tests, dose escalation moved the predicted state monotonically towards morula; basin entry first occurred at full dose, while a sign-flipped entry vector placed the trajectory 7.291 units from morula. These tests establish necessity, directional sensitivity and stage-specific branch logic inside the fitted operator-time model.

Within the fitted model, structure-matched reconstruction provided a stronger diagnostic test. A top-50 residual-DMR knock-in reduced the morula distance from 2.700 to 0.458 and entered the basin, whereas the 95th percentile of matched random sets remained outside at 0.781 and a shuffled-direction control diverged to 7.291 (Fig.2I-L). The empirical probability that a matched random top-50 set performed at least as well was 0.001. Coupling the correction to public chromatin-accessibility information reduced the distance further to 0.157. Public morula accessibility was elevated for the top 25 residual DMRs relative to matched controls, and the stage-specific methylation–accessibility association was significant only at morula in the tested sequence. Together, these observations support a chromatin-coupled component of the missing attraction, but they do not identify a unique molecular controller.

**Fig. 2.**
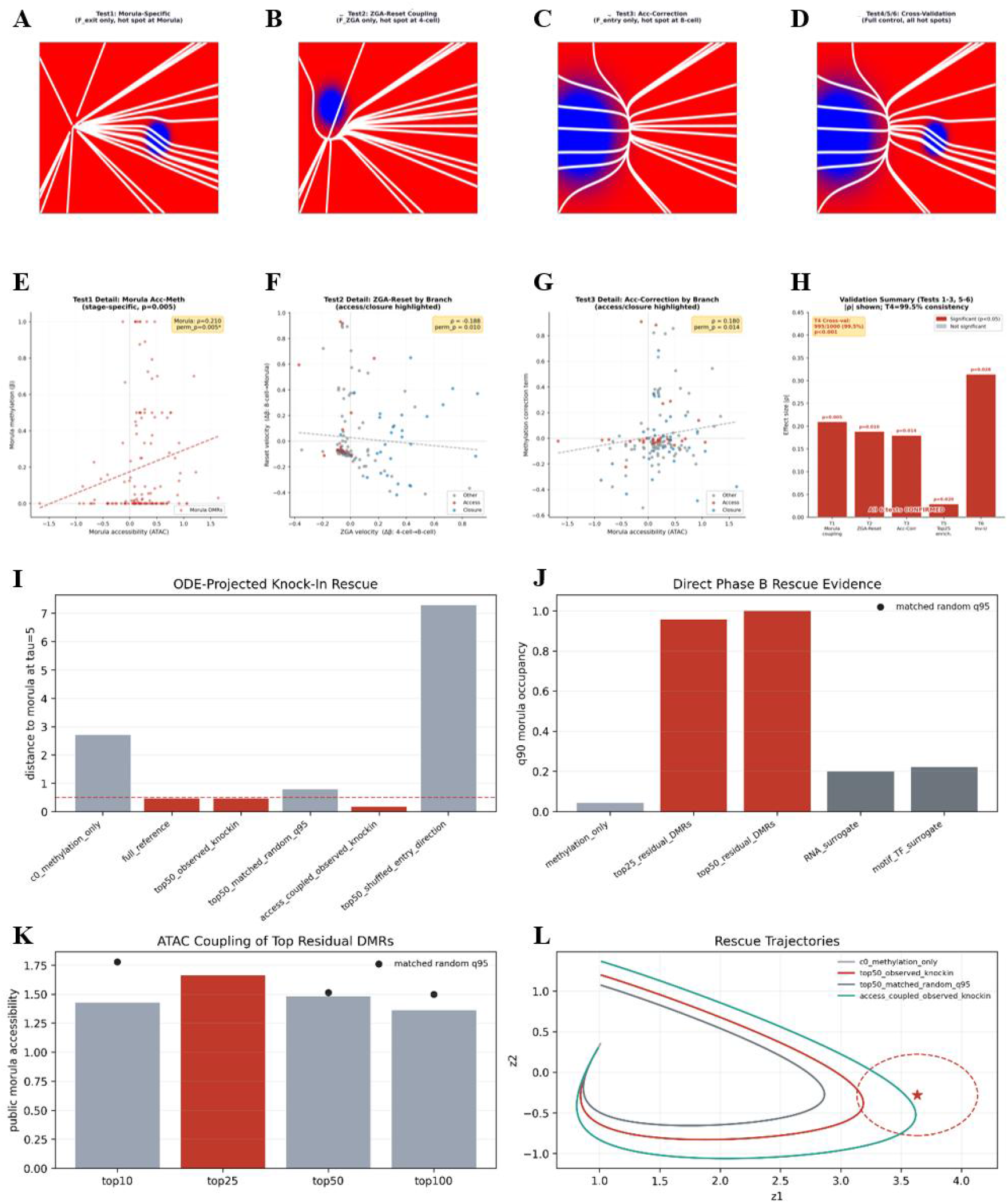
Independent chromatin accessibility supports the stage-restricted model-predicted configuration and structured residual DMRs rescue morula-basin entry. **A–D**.Field representations for morula-specific exit coupling, ZGA-associated reset coupling, accessibility–correction coupling and the full cross-validated control. **E**.Morula accessibility–methylation association (ρ = 0.210; permutation P = 0.005). **F**.Association between ZGA velocity and subsequent reset velocity, with access and closure branches highlighted (ρ = −0.188; permutation P = 0.010). **G**.Association between morula accessibility and the inferred methylation correction term (ρ = 0.180; permutation P = 0.014). **H**.Summary of six prespecified validation tests, including 995 of 1,000 cross-validation resamples with the expected sign, top-25 residual-DMR enrichment and inverted-U coupling. Chromatin data are independent of the methylation profiles used to fit the operator. **I-L**.Top-ranked residual-DMR knock-in and accessibility-coupled rescue are compared with matched-random and shuffled-direction controls. Successful reconstruction in the diagnostic framework requires both the selected DMR structure and its inferred direction. The experiment is computational and should be interpreted as model-implied partial sufficiency.

We next asked whether the configuration inferred from the methylation trajectory was detectable in an independently generated human preimplantation chromatin-accessibility dataset comprising measurements mapped to the same 156 DMRs. Six prespecified tests interrogated distinct predictions of the model (Fig.2A-H). First, the association between accessibility and methylation was stage restricted: the morula coefficient was positive (Spearman ρ = 0.210, permutation P = 0.005), whereas the corresponding associations at the 2-cell, 4-cell and 8-cell stages did not pass the permutation criterion. This localization argues against a fixed accessibility–methylation correlation operating uniformly across cleavage development and instead places the coupling at the predicted transition gate.

Second, accessibility-linked DMR structure connected the entry field to the preceding ZGA-associated transition. ZGA velocity from 4-cell to 8-cell was negatively associated with the subsequent reset velocity from 8-cell to morula (ρ = −0.188, permutation P = 0.010), with the access and closure branches occupying different regions of the joint distribution. Third, morula accessibility was positively associated with the inferred methylation correction term (ρ = 0.180, permutation P = 0.014). These effects are modest in magnitude, as expected for DMR-resolved public data, but their stage, sign and branch specificity match independent predictions made by the operator model rather than merely reproducing the stage-level methylation trajectory.

Three additional stress tests examined whether the association depended on a small number of favourable observations. Cross-validation retained the expected coupling in 995 of 1,000 resamples (99.5%; empirical P < 0.001). The top 25 residual DMRs were enriched relative to the matched background (P = 0.020), and an inverted-U coupling test was also significant (P = 0.028), consistent with the proposition that the most productive control does not arise from indiscriminately maximal accessibility. Taken together, all six prespecified chromatin tests supported the predicted model configuration. Nevertheless, these are associations across independently generated public profiles. They validate the geometry and stage restriction of the inferred control but do not demonstrate that accessibility perturbation directly causes the human morula methylation state.

The exit from ground zero was not the inverse of entry. From morula to blastocyst, approximately one-third of informative DMRs followed a remethylating branch and two-thirds followed a demethylating branch. Supplying the correct direction for both branches reduced prediction error to 0.0177, a 93.9% improvement over baseline. Correct demethylation with incorrect remethylation retained only a 19.2% improvement, whereas correct remethylation with incorrect demethylation performed 5.5% worse than baseline; reversing both branches was 32.8% worse. The marked asymmetry identifies the demethylating branch as especially direction-sensitive while showing that accurate exit requires coordinated bidirectional change.

Chromatin association also changed across the transition. Morula accessibility was the only tested stage-specific accessibility measure associated with the subsequent blastocyst correction (Spearman ρ = −0.173, nominal P = 0.0369, one-sided permutation P = 0.0163); corresponding associations at 2-cell, 4-cell and 8-cell were not significant. This stage restriction mirrors the entry-side observation that the correction becomes accessible near the basin rather than being encoded as a fixed global property across cleavage stages. However, leave-one-out prediction improvement was not significant at the global all-DMR level. We therefore interpret accessibility as a plausible mediator and branch selector, not as a completed causal chain.

Independent human RRBS data from GSE49828 provided directional DNA support. Age-weighted entropy decreased from 0.354 in sperm to 0.303 at morula, a reduction of 0.050, and morula ranked third among embryo-associated stages after MII oocyte and 4-cell. Sparse overlap with the age-DMR panel precluded a stringent replication of the primary stage ordering, but the sperm-to-morula direction was concordant with the proposed reset. In paired mouse parental methylomes from GSE56697, the paternal ICM state was the ground-zero call across four genomic bin sizes (100 kb to 1 Mb), and sperm-to-paternal-2-cell was consistently the most productive early transition. These data show that TRO can be instantiated as a true paired gamete-to-embryo methylome operator in mouse, while not substituting for paired paternal-age data in human embryos.

A gene-matched cross-species analysis yielded a narrower diagnostic signal. Of 153 human genes carrying age-DMR weights, 114 matched the mouse GLEANER methylation matrix. Their age-weighted methylation was lowest at mouse morula (0.302), 0.049 below the next-lowest stage. However, the full human-mouse stage-profile correlation was weak (ρ = 0.143, P = 0.760), morula was the minimum in only 39.0% of matched-gene bootstraps, and the result disappeared under equal weighting. The cross-species result is therefore limited weight-dependent support for a conserved morula-low configuration, rather than evidence for broad conservation of the complete human trajectory (Fig. 3).

**Fig. 3.**
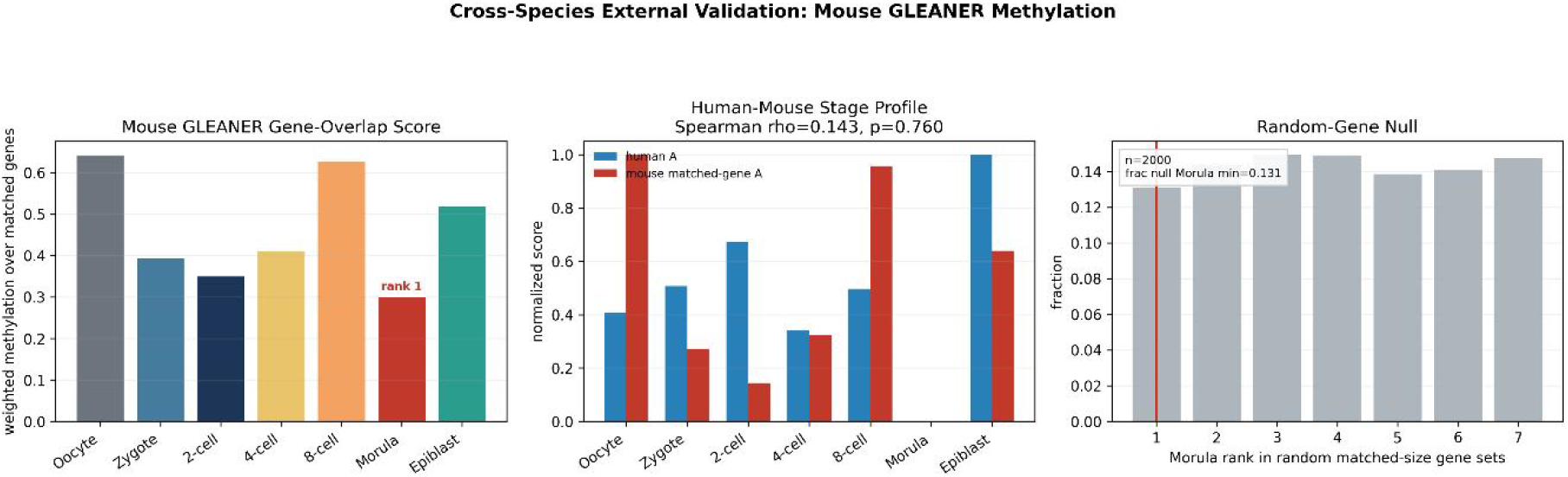
External mouse data provide limited and weight-dependent cross-species consistency, not generalization. Cross-species mouse methylation analysis of genes matched to the human age-DMR anchor. Age-weighted methylation reaches its minimum at mouse morula, but the stage-profile correlation is weak and sensitivity analyses show dependence on the human weighting scheme. The panel is an orthogonal diagnostic rather than paired human validation.

Finally, we assembled seven independent human trophoblast, embryo-model and methylation-machinery contrasts as a prespecified orthogonal evidence matrix. The six trophoblast or embryo contrasts produced an unoriented Fisher-combined P = 0.0317, but the weighted Stouffer result was not significant (P = 0.328). Across all seven contrasts, the corresponding values were 0.0590 and 0.381. The strongest individual DMR-level association arose in the GSE109682 trophoblast contrast (ρ = 0.261, P = 0.0278), whereas several datasets were null or directionally inconsistent. RNA perturbation of human naive ESC/iTSC models identified responsive genes near selected DMRs, but enrichment across the 17 tested genes was not significant against matched random genes in the final conservative analysis. Accordingly, the integrated evidence supports involvement of the trophoblast methylome and responsiveness to methylation/chromatin perturbation, but it does not establish a single-dataset, mechanism-specific causal closure.

Collectively, these results identify morula as a computational ground-zero candidate characterized by minimal age-associated methylation disorder, preserved developmental potency and maximal productive reset efficiency. Coordinated latent dynamics capture more of the transition than independent DMR drift but still fail to generate the observed basin without a structured correction. Counterfactual removal, dose response, sign reversal and matched-random rescue show that this correction is necessary and partially sufficient within the operator-time model, while stage-restricted chromatin associations provide a plausible biological bridge. Independent human and mouse datasets support selected directions and components of the framework, but the decisive experimental test remains a matched perturbation-methylome study capable of identifying the in vivo molecular controller of basin entry.

## Methods

We developed the transgenerational reset operator (TRO) as an interpretable computational framework for representing the joint evolution of age-associated DNA-methylation perturbation and developmental potency during mammalian preimplantation development. The analysis was designed to answer three nested questions. First, does the preimplantation trajectory contain a stage at which age-associated methylation disorder is minimized without a concomitant loss of developmental potency? Second, can this stage be represented as a state reached by an operational reset operator with a measurable transition cost? Third, can the stage-level construction be extended to a distributional, DMR-resolved operator-time model that supports out-of-sample prediction, counterfactual intervention and external perturbational assessment? The workflow therefore proceeded from a transparent stage-level entropy representation to a DMR-level dynamical model, and then to orthogonal validation across independent methylome, transcriptome, chromatin and perturbation datasets.

The primary human developmental ordering was MII oocyte, zygote/pronuclear, 2-cell, 4-cell, 8-cell, morula and blastocyst. Inner-cell-mass and trophectoderm samples were retained in DNA-only summaries when available, but were excluded from the principal DNA–RNA aligned trajectory because they do not form a one-to-one continuation of the seven-stage axis. The central inferential target was a computational reset-basin candidate, defined as a stage with low age-associated methylation entropy and preserved potency, rather than a universal biological zero point. All public datasets were analysed as independent or partially complementary observations. No dataset was treated as a longitudinal record of the same human embryo, and no public-data comparison was interpreted as a matched aged-father-to-embryo experiment.

Age-associated methylation weights were obtained from the supplementary Table S6 associated with GSE102970. The GEO exposure matrix did not contain usable sample-level paternal-age metadata for direct re-estimation of the sperm-age model; consequently, the published DMR and CpG weights were treated as an externally defined feature set. GSE81233 provided the primary single-cell human preimplantation methylome. One file, GSM2986343_scBS-2C-10-1.Cmet.bed.gz, repeatedly failed gzip validation and was excluded before analysis, leaving 204 valid Cmet profiles for the stage-level entropy analyses. The DMR-resolved dynamics matrix comprised 169 samples with sufficient coverage across 156 age-associated DMR clusters.

GSE36552 supplied the primary human preimplantation transcriptome used to estimate transcriptomic entropy, detected-gene richness and potency-marker activity. GSE44183 served as an independent transcriptomic validation dataset; because platform and normalization differed, only within-dataset rankings were compared. GSE49828 provided an independent human RRBS trajectory with sperm and embryonic methylomes and was used as directional, not paired, gamete-to-embryo support. GSE56697 supplied paired mouse parental-allele methylomes. Its paternal branch mapped sperm to paternal embryonic alleles across 2-cell, 4-cell, inner-cell-mass, E6.5 and E7.5 states, while the maternal branch provided an oocyte-to-maternal-allele contrast. These data instantiate the operator on paired mouse parental methylomes but do not establish paired human paternal-age resetting.

Additional analyses evaluated the 156 residual CSB–TRO DMRs across human trophoblast and embryo-model datasets. These included GSE109682, GSE150168, GSE182015, GSE266195, GSE280039 and GSE291172 for methylome state or perturbation contrasts; GSE126958 for TET/DNMT3 machinery perturbation in human embryonic stem cells; GSE247631 for A-485/P300-inhibition and dimethyl-α-ketoglutarate transcriptomic perturbations in a naïve human preimplantation-lineage model; E-MTAB-10096 for embryo transcriptional effects; and E-MTAB-10097 for embryo bisulfite-sequencing analyses. The resulting evidence table was treated as a prespecified, heterogeneous, orthogonal meta-dataset. It was not represented as a single paired experiment.

For sample (i), DMR (j) and stage (g), the observed methylation fraction was denoted by β_ij_ ∈ [0, 1]. Methylation values were coerced to numeric form, joined to metadata by sample identifier and aggregated within developmental stage. A DMR was included in a stage summary when at least 30% of the corresponding stage-specific observations were non-missing. The stage mean was

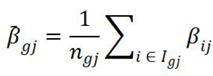

where Igj is the set of valid samples for DMR j at stage g, and ngj = |Igj|. Values entering logarithms were clipped to [ε, 1 − ε], with ε = 10^−6^.

The binary methylation entropy of a methylation fraction p was defined as

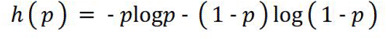

Generic stage-level epigenetic entropy was calculated as

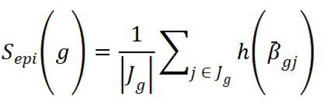

where Jg is the set of DMRs passing the stage-specific coverage criterion. This quantity measures methylation-state uncertainty or mixture and was not interpreted as ageing entropy. To isolate perturbation associated with the externally defined sperm-age signature, we used the absolute published age weight |wj| for each overlapping DMR. Age-associated methylation entropy was

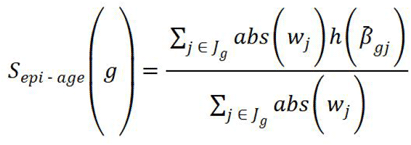

Absolute weights were used in the entropy because the quantity measures the uncertainty carried by age-informative regions, irrespective of the direction of the original age association. Direction was retained separately in the signed age projection,

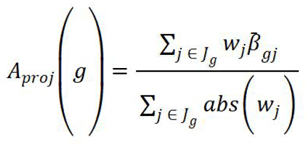

For each cell or sample i, non-negative expression values xik were converted into a compositional distribution over genes,

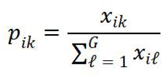

and global transcriptomic Shannon entropy was calculated as

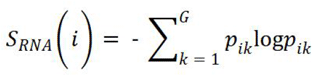

Cell-level values were summarized by stage using the arithmetic mean and sample standard deviation. Global RNA entropy was retained as an independent state descriptor because transcriptional diversity and developmental potency are not interchangeable. In particular, the model did not require morula to maximize global RNA entropy.

Developmental potency was represented by a composite score integrating detected-gene richness with the activity of a curated developmental potency-marker panel. Each component was oriented so that a larger value indicated greater potency, standardized across the aligned stages, and combined using the fixed implementation recorded in the project scripts. The stage-level potency score was denoted by P(g). Robustness was assessed by removing one marker at a time, recomputing the score and repeating the morula-versus-blastocyst comparison. An expanded marker panel and the independent GSE44183 dataset were used as additional sensitivity checks. Cross-dataset comparisons were restricted to ranks or direction because absolute expression scales were not commensurate.

The stage-level operator was defined as the ordered tuple

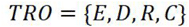

where E is an entropy encoder, D is a damage–potency decomposer, R is a reset map and C is a transition-cost estimator. The encoder maps stage g to

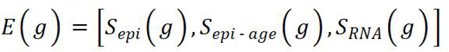

The decomposition was

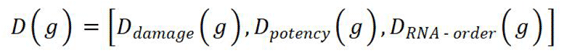

with

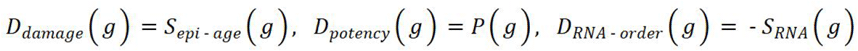

The internal reset score used MII oocyte and morula as operational anchors,

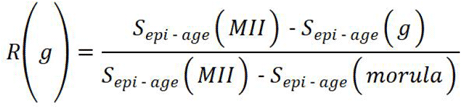

Thus, R(MII) = 0 and R(morula) = 1 by construction. This normalization is an internal developmental score, not a young-versus-aged paternal reset index. Potency preservation was

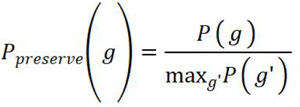

and the composite TRO score was

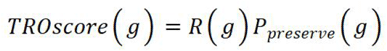

An independent ground-zero score combined low perturbation with preserved potency,

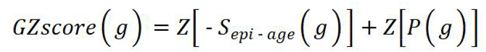

where Z(·) denotes standardization across stages. A computational ground-zero candidate was required to rank first by both GZscore and TROscore; the operational audit also required first rank by the separately calculated bio-age score and concordance of the DNA, RNA and robustness checks.

Each stage was embedded in a standardized four-dimensional state vector,

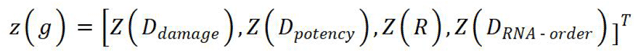

For adjacent stages g → h, transition cost was the Euclidean displacement

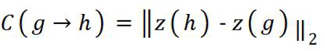

Damage reduction, potency change and reset gain were computed as

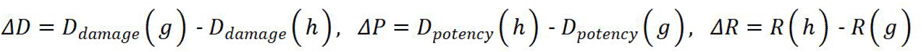

Only beneficial components contributed to productive reset gain,

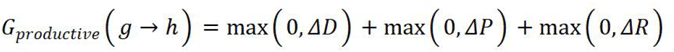

and reset efficiency was

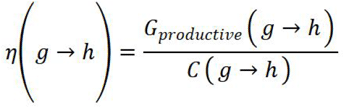

This construction separates the magnitude of a developmental displacement from its biological direction. A large transition is not classified as productive when it increases the perturbation burden or sacrifices potency without compensating reset gain.

Because the available profiles are cross-sectional, developmental operator time τ was defined as a stage-anchored pseudo-time rather than physical time, which has been applied in previous researches. ^[15, 16]^The seven ordered stages were mapped linearly to

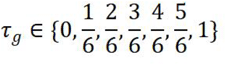

Morula therefore occupies τ = 5/6 and is an intermediate basin candidate rather than a terminal state. Adjacent-stage samples were linked through precomputed entropically regularized optimal-transport couplings. For source states x_a_ and target states x_β_, the cost matrix was Euclidean, Mab = ‖x_a_ − x_β_‖_2_, and uniform empirical masses were used. The coupling π minimized

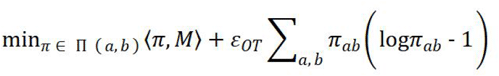

subject to the prescribed source and target marginals. The stage-level implementation used an entropy-regularization parameter of 0.05. Coupling-resolved intermediate states were defined by displacement interpolation,

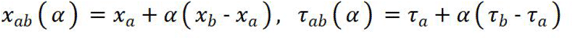

with α ∈ [0, 1]. Intermediate summaries were weighted by π ab. These paths provide a distribution-aware interpolation between observed stages; they do not imply lineage tracking of individual embryos.

For DMR j, each coupled sample pair generated the target velocity

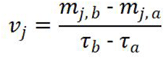

where mj,a and mj,b are the source and target methylation fractions. A transparent weighted ridge model was fitted independently for each DMR,

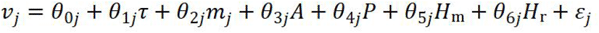

Here, A denotes the age-associated perturbation coordinate, P the potency coordinate, H_m_ the methylation-heterogeneity coordinate and H_r_ the RNA-heterogeneity coordinate. Coefficients were estimated by weighted ridge regression,

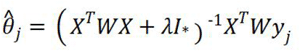

where W contains coupling weights, λ = 50 for the single-DMR baseline and I* leaves the intercept unpenalized. Predictions were clipped to the biologically admissible interval [0, 1]. Strict morula extrapolation excluded every transition containing morula from model fitting; an analogous procedure excluded blastocyst for terminal-stage prediction. Comparators included carry-forward of the 8-cell mean, extrapolation of the previous stage velocity, random target coupling and time/module randomization.

The 156-dimensional DMR state was compressed to coordinated modules and a principal-component latent representation because independent DMR dynamics failed to outperform the 8-cell carry-forward baseline under strict morula exclusion. Let m_i_ ∈ ℝ^156^ be the standardized DMR vector. The latent state was

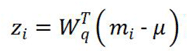

where Wq contains the first q = 3 principal axes. Latent velocities were fitted with ridge-regularized affine dynamics and integrated over operator time. The advanced rollout used λ = 1000, and its local linear stability was summarized from the eigenvalues of the fitted Jacobian. Negative real parts were interpreted as local contraction of the fitted mean-state dynamics, not proof of a physical attractor.

Validation was performed with the entire morula stage withheld. The archived strict-validation run used 200 null replicates, 1,000 bootstrap resamples and random seed 20260525. Bootstrap confidence intervals were calculated for the paired change in DMR-level root-mean-square error relative to the 8-cell baseline. Paired squared-error tests and random-time, random-module and random-coupling controls assessed whether gains depended on coordinated representation, developmental ordering or the specific transport coupling. Because the random-coupling control was not consistently inferior, improvements were attributed to the module/latent representation rather than to a uniquely established sample-level coupling mechanism.

The morula basin was defined in the three-dimensional latent space using the empirical morula distribution. Occupancy thresholds were based on the distribution of distances from morula samples to the morula centre, with the 90th percentile used for the primary basin boundary. Deterministic rollout and several stochastic extensions were evaluated, including global-residual diffusion, stage-conditioned diffusion, empirical transition noise, affine Gaussian kernels and Ornstein–Uhlenbeck diagnostics. The primary SDE had the generic form

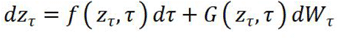

The archived morula-basin run used 200 particles per starting sample, 40 particles per starting sample during calibration, 12 integration steps and seed 20260525. A non-leaking validation repeated the analysis without using morula transitions for fitting or calibration. Models calibrated on the observed morula distribution were reported only as diagnostic upper bounds.

The missing basin-attraction correction was defined as the difference between the observed morula latent centre and its strict pre-morula prediction,

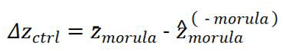

For the final operator-time step Δτ, the equivalent correction velocity was

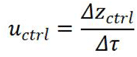

Amplitude scans applied αΔzctrl, with α ranging from zero to one, to quantify the dose–occupancy relationship. The correction was mapped back through latent loadings to rank DMRs and modules, followed by top-K addition, removal, sign-flip and matched-random controls. This procedure measures the composite quantity Bubio required by the fitted dynamics; it does not identify a unique biological controller ubio.

Morula geometry was evaluated by comparing entry and exit directions in DMR space. For stage means μ_8_, μM and μB, the entry and exit vectors were

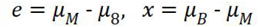

Direction reversal was summarized through a cosine-based entry–exit duality statistic, with significance assessed by permutation and uncertainty by bootstrap resampling of DMRs. Stage-wise methylation distributions were additionally characterized by the fraction of fully demethylated DMRs and a bimodality index. The morula-to-blastocyst transition was decomposed into remethylating and demethylating branches. Correct-direction, wrong-direction and mixed-branch reconstructions quantified whether prediction depended on branch identity rather than merely on correction magnitude.

Chromatin accessibility was evaluated as a candidate explanatory coordinate for the residual correction. For DMR j, the strict correction was

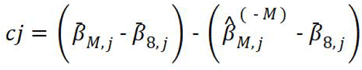

Spearman correlations related quantitative morula accessibility to cj, both globally and within prespecified curvature classes. Empirical significance was obtained by permuting DMR labels, and partial correlations were used where covariate adjustment was available. Accessibility was treated as a candidate control-associated variable. Correlation with the correction term was not interpreted as a causal do-intervention.

Counterfactual simulations used the fitted operator-time system to compare a methylation-only trajectory, the full inferred correction, entry/access-only control and an incorrect exit-sign control. For dynamical state z, the controlled system was represented as

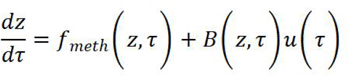

Necessity was defined within the model: removing the inferred control term had to prevent entry into the prespecified morula basin, while the complete control restored basin entry and permitted a valid post-morula exit trajectory. Wrong-sign and branch-restricted controls tested directionality and sufficiency of individual components. These interventions establish model-implied necessity or rescue, not in vivo molecular causality.

For transcriptomic perturbation datasets, gene-level effects were estimated within each study according to its archived design matrix and linked to DMRs through the recorded nearest-gene mapping. CSB-linked genes were compared with expression-matched random gene sets. Empirical one-sided P values were calculated as

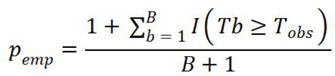

where Tobs is the observed mean absolute or weighted perturbation effect and Tb is the statistic for matched random set b. Weighting analyses used the corresponding CSB residual or age-DMR weights and were reported alongside unweighted analyses.

For independent methylome contrasts, methylation was aggregated over the 156 residual DMR intervals after coordinate and coverage harmonization. Contrast direction was compared with the CSB residual direction by Spearman correlation and sign concordance. The integrated evidence matrix retained assay, system, perturbation status, covered DMR count, effect direction, P value, evidence tier and an explicit usable-claim field. Heterogeneous P values were combined only as unoriented diagnostics using Fisher and weighted Stouffer procedures. Such aggregation tests whether any association is present across heterogeneous contrasts; it cannot establish a common effect direction or causal closure.

Stage-level uncertainty was quantified by nonparametric bootstrap resampling of samples within stage. Unless a module-specific archived command specified otherwise, 1,000 bootstrap iterations were used. The ground-zero frequency was the fraction of iterations in which each stage minimized age-associated methylation entropy. Adjacent-stage differences and potency comparisons were assessed with two-sided Mann–Whitney tests, followed by Benjamini–Hochberg correction within each analysis family. DMR-level association tests used Spearman rank correlation. Empirical nulls were generated by shuffling age weights, sampling random DMR sets matched for relevant properties, randomizing stage time, randomizing module labels or reversing correction signs, as appropriate to the hypothesis.

Robustness analyses included alternative coverage thresholds, common-DMR restriction, balanced bootstrap sampling, shuffled-weight controls, random age-DMR subsets, leave-one-marker-out potency scores, expanded potency-marker panels, genomic-bin sensitivity in paired mouse data, top-K DMR addition and removal, sign-flip controls, matched-random DMR controls, representation sensitivity and strict stage-exclusion prediction. All hypothesis tests were interpreted together with effect magnitude, confidence intervals, null distributions and the stated claim boundary. Exploratory enrichment, nearest-gene mapping, motif overlap, ATAC overlap and histone overlap were treated as mechanism-generating evidence rather than proof that an individual DMR controls the reset state.

The analysis was implemented in Python using archived scripts, metadata, result tables, run summaries and command records in the TRO, CSB–TRO operator-time and causal-chain workspaces. The result-only package can be audited with the supplied check scripts; full methylome regeneration requires restoration of the raw Cmet, RRBS, WGBS, PBAT or sequencing files identified in the manifests. Deterministic analyses used explicitly recorded seeds, including seed 20260525 for the principal latent validation and basin simulations. All stage names, coordinate systems and feature identifiers were held constant within each workflow and translated only through documented mapping tables.

## Discussion

Here, we used an externally defined sperm-age-associated DMR feature space to ask whether age-associated methylation features undergo a structured transition during human preimplantation development. We identified morula as a computational ground-zero candidate that combines the lowest age-DMR-weighted methylation entropy with preserved RNA-derived developmental potency. Importantly, morula was not the stage of maximal potency; rather, the 8-cell and morula stages together formed a high-potency developmental region, within which morula showed the most pronounced reduction in the age-informed methylation coordinate. Age-associated methylation changes in human sperm have been reported across independent cohorts, providing biological motivation for this feature space. ^[17, 18]^Our result does not imply complete erasure of paternal-age information or a biological age-zero state. Instead, it defines a stage-restricted configuration in an externally specified age-DMR space and should be interpreted as a computational reset candidate rather than direct evidence of paternal-age rejuvenation in a matched human lineage.

The morula-associated minimum should not, however, be interpreted simply as a rediscovery of the extensive DNA demethylation that accompanies early embryogenesis. Genome-scale studies established that the human preimplantation methylome undergoes profound reprogramming after fertilization: a major wave of genome-wide demethylation is largely completed by the 2-cell stage, a transient globally hypomethylated state characterizes preimplantation development, and single-cell methylome profiling further revealed that reprogramming reflects a dynamic balance between widespread demethylation and focused de novo remethylation.^[19-21]^ Our age-DMR trajectory is consistent with this broader view of heterogeneous methylome reprogramming but reveals a distinct feature-space organization. Age-DMR-weighted methylation entropy declined through the 4-cell stage, increased again at 8-cell, fell to its minimum at morula and subsequently rebounded in blastocyst, while the signed age-associated projection showed a concordant morula minimum. Thus, the computational reset candidate does not arise at the endpoint of a simple monotonic loss of methylation. We instead view the morula state as a stage-restricted reorganization of an age-informed methylation feature space superimposed on the broader programme of embryonic methylome reprogramming.

The developmental position of this computational state is also notable. Human preimplantation development is accompanied by sequential transcriptional programmes and major shifts in regulatory state. Single-cell transcriptomic studies have described stage-specific co-expression modules from cleavage stages to morula, together with broad conservation but species-specific timing of developmental programmes. ^[13, 14]^At later preimplantation stages, cells pass through intermediate transcriptional states before the establishment of trophectoderm, epiblast and primitive-endoderm identities associated with blastocyst formation. ^[22]^Human embryo compaction likewise represents an active morphogenetic transition involving marked remodelling of cell-surface tension and requirements for contractility and adhesion. ^[23]^Against this background, the location of the age-DMR minimum around morula is consistent with a broader developmental interval in which transcriptional, epigenetic and morphogenetic programmes are being reorganized, rather than with a terminal low-methylation state. We do not infer that lineage specification, compaction mechanics or any individual transcriptional programme causes the methylation configuration identified here; these observations instead place the CSB-TRO turning point within a biologically active developmental window whose regulatory architecture changes across multiple molecular layers.

A key implication of the operator-time analysis is that identifying a developmental state and dynamically reconstructing its formation are not equivalent problems. The morula-associated methylation configuration is readily apparent in the observed stage-level data, yet it was not reproduced by a model that treated DMRs as independent local trajectories: under strict morula exclusion, the single-DMR model performed worse than simply carrying forward the 8-cell methylation state. Introducing coordinated structure improved reconstruction, first through DMR modules and then through a low-dimensional latent representation, indicating that the transition contains reproducible information distributed across groups of loci rather than being reducible to independent locus-wise drift. This gain, however, remained incomplete. Strict pre-morula rollouts still failed to recover the observed morula population basin, suggesting that methylation history contains a coherent low-dimensional developmental trajectory but is insufficient, on its own, to specify the full morula state. Equally importantly, the improvement was not consistently dependent on a privileged optimal-transport coupling, arguing that the principal gain arose from coordinated representation rather than uniquely inferred sample-to-sample correspondence. We therefore interpret operator time as a stage-anchored, distribution-aware reconstruction coordinate rather than as physical time, lineage inference or prospective forecasting. In this sense, its most informative result is not that morula can be predicted from earlier stages, but that the discrepancy between what preceding methylation dynamics can generate and what is actually observed at morula can be quantified.

The target-informed residual required to reconcile the strict pre-morula reconstruction with the observed morula state provides a useful constraint on what is missing from the methylation-only dynamics, but it should not be mistaken for an independently identified biological control signal. By construction, the diagnostic residual is defined from the difference between the observed morula state and its stage-withheld reconstruction; consequently, restoring the full correction necessarily moves the model towards the target and cannot itself constitute independent validation. The informative feature is instead the organization of this discrepancy. When projected back to DMR space, the residual was concentrated in reproducible subsets and modules, progressively improved basin reconstruction as ranked components were added, and lost efficacy when its direction was reversed or when matched random DMR sets were substituted. These observations indicate that failure of the autonomous model is not well described as diffuse stochastic error, but instead contains coordinated direction and amplitude structure. At the same time, because residual ranking and rescue are derived from the same observed target, these counterfactuals establish structure, direction sensitivity and model-internal rescue rather than molecular causality. The fitted data constrain a composite diagnostic control term, but do not uniquely identify the biological input or the molecular operator through which it acts. A complementary geometric analysis showed that entry from 8-cell to morula and exit from morula to blastocyst were strongly anti-aligned, indicating that the developmental trajectory bends around morula rather than continuing along a single demethylation axis. We therefore interpret morula as a pronounced developmental turning point in the age-DMR trajectory, while reserving the stronger notion of a molecular gate for future perturbational tests.

Chromatin accessibility provides one plausible biological bridge between the structured residual and the broader regulatory reorganization of the preimplantation embryo, but the magnitude and design of the present evidence argue against treating it as a complete control mechanism. Human embryo chromatin studies have established that accessibility is extensively remodelled during preimplantation development, including widespread reorganization around zygotic genome activation and coordinated changes in accessible chromatin and transcription. ^[24, 25]^Against this dynamic background, the morula accessibility-methylation association and the positive association between morula accessibility and the strict DMR-level diagnostic correction place part of the missing methylation structure within an independently measured chromatin context. These effects were modest (Spearman’s rho approximately 0.21 and 0.18, respectively), and accessibility produced only a small all-DMR predictive gain. We therefore favour an interpretation in which accessibility marks or contributes to a stage-restricted component of the regulatory environment associated with the morula transition, rather than constituting the sole instructive signal for basin entry. This distinction is particularly important because the chromatin and methylation measurements were obtained from independent public datasets, and no experiment in the present study directly perturbed accessibility while measuring the resulting methylation state at the same DMRs. Accordingly, accessibility-residual coupling is best viewed as a chromatin-associated candidate control component that narrows the biological search space for the missing input, not as causal closure.

The external datasets further reveal both the fragile generality and the evident limitations of the framework. An independent human preimplantation RRBS trajectory showed a concordant decrease in the age-weighted coordinate from sperm towards morula, although incomplete overlap with the age-DMR panel prevented stringent replication of the primary stage ordering.3 In paired mouse parental methylomes, the framework could be instantiated on a genuine gamete-to-embryo, parent-of-origin-resolved trajectory, providing a stronger test of the operator concept while not substituting for human paternal-age data. ^[26]^ Cross-species transfer was nevertheless limited: the mouse morula-low signal depended on the transferred human weighting scheme, the full stage-profile correlation was weak, and the result was not stable under equal weighting. The broader integrated orthogonal evidence matrix was similarly heterogeneous, with supportive, null and directionally inconsistent contrasts and aggregation-sensitive combined evidence. This heterogeneity is biologically informative rather than merely inconvenient. In human donor-oocyte blastocysts, paternal-age-associated methylation differences have been detected in both inner-cell-mass and trophectoderm lineages, demonstrating that paternal-age-related epigenetic differences can remain detectable after the developmental interval in which we identify the computational minimum. ^[27]^Taken together, these observations argue against universal or permanent erasure of paternal-age-associated methylation information. A more defensible interpretation is that CSB-TRO identifies a transient, feature-space-specific developmental reorganization whose magnitude and persistence depend on genomic region, developmental context and biological system.

Several features of the present study define the boundary between the computational framework and a causal model of human epigenetic resetting. The methylation and transcriptomic measurements were obtained from different embryos and were integrated at the developmental-stage level; the resulting joint states therefore represent a stage-conditioned computational fusion rather than paired multi-omic measurements from the same cells. Operator time is an ordered stage-anchored pseudo-time, not physical time or longitudinal observation of individual embryos, and the optimal-transport couplings should accordingly be interpreted as distribution-aware interpolation rather than lineage correspondence. Both the morula-conditioned evaluation basin and the diagnostic residual also use information from the observed target state, so target-calibrated basin recovery and full-residual rescue are model diagnostics rather than independent evidence that the corresponding control operates in vivo. The numerical drift-diffusion and COMSOL representations carry the same boundary: they realize and interrogate the fitted field but do not independently establish a physical force or molecular mechanism. Finally, the external evidence is drawn from heterogeneous human and mouse systems, and no available dataset provides a matched human father-sperm-embryo lineage in which paternal age, candidate regulatory machinery and the same target DMRs are measured together. The most decisive next step is therefore perturbational. A stringent test would alter candidate chromatin- or methylation-regulatory machinery in a developmentally relevant and ethically appropriate embryo or embryo-model system, quantify methylation at the same residual DMRs, and ask prospectively whether the direction and magnitude of morula-like state entry change as predicted by the model. A complementary observational design would link paternal age and sperm methylation to matched embryonic methylomes, allowing inherited signal and developmental reorganization to be separated. Until such data are available, CSB-TRO is best viewed as a framework that converts a stage-restricted age-DMR configuration into quantitative, falsifiable hypotheses about developmental control, rather than as proof of a unique rejuvenation mechanism.

## Supporting information

Supplemental docs

Supplemental environment

Supplemental models

Supplemental other

## Notes

https://www.ncbi.nlm.nih.gov/geo/query/acc.cgi?acc=GSE81233

https://www.ncbi.nlm.nih.gov/geo/query/acc.cgi?acc=GSE102970

https://www.ncbi.nlm.nih.gov/geo/query/acc.cgi?acc=GSE36552

https://www.ncbi.nlm.nih.gov/geo/query/acc.cgi?acc=GSE44183

https://www.ncbi.nlm.nih.gov/geo/query/acc.cgi?acc=GSE49828

https://www.ncbi.nlm.nih.gov/geo/query/acc.cgi?acc=GSE56697

https://www.ncbi.nlm.nih.gov/geo/query/acc.cgi?acc=GSE109682

https://www.ncbi.nlm.nih.gov/geo/query/acc.cgi?acc=GSE126958

https://www.ncbi.nlm.nih.gov/geo/query/acc.cgi?acc=GSE247631

https://github.com/xiangyu-star/Operator

### Competing Interest Statement

The authors have declared no competing interest.

