## Supplemental code for "An entropy-based diagnostic framework for characterizing methylation state dynamics during preimplantation development": 00_project_config.py

```
from pathlib import Path

PROJECT_ROOT = Path(__file__).resolve().parents[1]

DATA_RAW = PROJECT_ROOT / "data_raw"
DATA_PROCESSED = PROJECT_ROOT / "data_processed"
METHYLATION_DIR = DATA_PROCESSED / "methylation_matrix"
RNASEQ_DIR = DATA_PROCESSED / "rnaseq_matrix"
METADATA_DIR = DATA_PROCESSED / "metadata"
RESULTS_DIR = PROJECT_ROOT / "results"
RESULTS_TABLES = RESULTS_DIR / "tables"
RESULTS_FIGURES = RESULTS_DIR / "figures"
LOGS_DIR = PROJECT_ROOT / "logs"

NCBI_GEO_FTP = "https://ftp.ncbi.nlm.nih.gov/geo/series"

GEO_DATASETS = {
    "GSE102970": {
        "series_path": "GSE102nnn/GSE102970",
        "role": "human sperm methylation and male age",
        "priority": 1,
    },
    "GSE81233": {
        "series_path": "GSE81nnn/GSE81233",
        "role": "human preimplantation embryo DNA methylome",
        "priority": 1,
    },
    "GSE155179": {
        "series_path": "GSE155nnn/GSE155179",
        "role": "human oocyte aging scRNA-seq",
        "priority": 2,
    },
    "GSE36552": {
        "series_path": "GSE36nnn/GSE36552",
        "role": "human embryo scRNA-seq",
        "priority": 2,
    },
    "GSE44183": {
        "series_path": "GSE44nnn/GSE44183",
        "role": "human and mouse early embryo scRNA-seq",
        "priority": 2,
    },
}

STAGE_ORDER = [
    "MII oocyte",
    "zygote",
    "2-cell",
    "4-cell",
    "8-cell",
    "morula",
    "blastocyst",
    "ICM",
    "TE",
]

DEFAULT_MIN_NON_MISSING_FRAC = 0.30
DEFAULT_EPS = 1e-6

def ensure_dirs():
    for path in [
        DATA_RAW,
        METHYLATION_DIR,
        RNASEQ_DIR,
        METADATA_DIR,
        RESULTS_TABLES,
        RESULTS_FIGURES,
        LOGS_DIR,
    ]:
        path.mkdir(parents=True, exist_ok=True)
```
