## Supplemental code for "An entropy-based diagnostic framework for characterizing methylation state dynamics during preimplantation development": 01_list_geo_files.py

```
import argparse
import html
import importlib.util
import re
from pathlib import Path
from urllib.parse import urljoin

import pandas as pd
import requests


def load_config():
    cfg_path = Path(__file__).with_name("00_project_config.py")
    spec = importlib.util.spec_from_file_location("project_config", cfg_path)
    cfg = importlib.util.module_from_spec(spec)
    spec.loader.exec_module(cfg)
    return cfg


def parse_apache_links(text):
    links = []
    for href in re.findall(r'href="([^"]+)"', text):
        href = html.unescape(href)
        if href.startswith("?") or href.startswith("/") or href == "../":
            continue
        links.append(href)
    return links


def extension_group(filename):
    lower = filename.lower()
    for ext in [".tar.gz", ".fastq.gz", ".fq.gz", ".bed.gz", ".txt.gz", ".csv.gz", ".tsv.gz"]:
        if lower.endswith(ext):
            return ext
    return Path(lower).suffix


def list_supplementary_files(dataset, info, base_url, timeout=60):
    url = f"{base_url}/{info['series_path']}/suppl/"
    r = requests.get(url, timeout=timeout)
    r.raise_for_status()
    rows = []
    for href in parse_apache_links(r.text):
        full_url = urljoin(url, href)
        filename = href.rstrip("/")
        rows.append(
            {
                "dataset": dataset,
                "role": info.get("role", ""),
                "priority": info.get("priority", ""),
                "filename": filename,
                "extension_group": extension_group(filename),
                "url": full_url,
            }
        )
    return rows


def main():
    cfg = load_config()
    cfg.ensure_dirs()

    parser = argparse.ArgumentParser()
    parser.add_argument("--datasets", nargs="*", default=list(cfg.GEO_DATASETS.keys()))
    parser.add_argument("--out", default=str(cfg.RESULTS_TABLES / "geo_supplementary_files.tsv"))
    args = parser.parse_args()

    all_rows = []
    for dataset in args.datasets:
        info = cfg.GEO_DATASETS[dataset]
        print(f"Listing {dataset}: {info['role']}", flush=True)
        rows = list_supplementary_files(dataset, info, cfg.NCBI_GEO_FTP)
        print(f"  files: {len(rows)}", flush=True)
        all_rows.extend(rows)

    df = pd.DataFrame(all_rows)
    out = Path(args.out)
    out.parent.mkdir(parents=True, exist_ok=True)
    df.to_csv(out, sep="\t", index=False)

    print(f"Wrote: {out}")
    if len(df):
        print(df.groupby(["dataset", "extension_group"]).size().reset_index(name="n").to_string(index=False))


if __name__ == "__main__":
    main()
```
