## Supplemental code for "An entropy-based diagnostic framework for characterizing methylation state dynamics during preimplantation development": 02_core_entropy.py

```
import argparse
import numpy as np
import pandas as pd

EPS = 1e-6


def binary_entropy(p, eps=EPS):
    p = np.asarray(p, dtype=float)
    p = np.clip(p, eps, 1.0 - eps)
    return -(p * np.log(p) + (1.0 - p) * np.log(1.0 - p))


def as_numeric_frame(df):
    return df.apply(pd.to_numeric, errors="coerce")


def ordered_stages(metadata, stage_col="stage", stage_order=None):
    observed = list(pd.Series(metadata[stage_col].dropna().unique()).astype(str))
    if stage_order is None:
        return observed
    ordered = [s for s in stage_order if s in observed]
    ordered.extend([s for s in observed if s not in ordered])
    return ordered


def compute_stage_epi_entropy(
    beta_matrix,
    metadata,
    stage_col="stage",
    stage_order=None,
    min_non_missing_frac=0.30,
    eps=EPS,
):
    beta = as_numeric_frame(beta_matrix)
    meta = metadata.copy()
    meta.index = meta.index.astype(str)
    beta.index = beta.index.astype(str)

    common_samples = meta.index.intersection(beta.index)
    meta = meta.loc[common_samples]
    beta = beta.loc[common_samples]

    rows = []
    for stage in ordered_stages(meta, stage_col, stage_order):
        sample_ids = meta.index[meta[stage_col].astype(str) == str(stage)]
        sub = beta.loc[sample_ids]
        valid = sub.notna().mean(axis=0) >= min_non_missing_frac
        p = sub.loc[:, valid].mean(axis=0, skipna=True)
        h = binary_entropy(p, eps=eps)
        rows.append(
            {
                "stage": stage,
                "n_cells": int(len(sample_ids)),
                "n_regions": int(valid.sum()),
                "s_epi": float(np.nanmean(h)) if len(h) else np.nan,
            }
        )

    return pd.DataFrame(rows)


def compute_age_weighted_epi_entropy(
    beta_matrix,
    metadata,
    age_weights,
    stage_col="stage",
    stage_order=None,
    min_non_missing_frac=0.30,
    eps=EPS,
):
    beta = as_numeric_frame(beta_matrix)
    weights = pd.to_numeric(age_weights, errors="coerce").dropna()
    weights.index = weights.index.astype(str)
    beta.columns = beta.columns.astype(str)

    common_regions = beta.columns.intersection(weights.index)
    if len(common_regions) == 0:
        raise ValueError("No overlapping regions between beta_matrix and age_weights.")

    beta = beta.loc[:, common_regions]
    weights = weights.loc[common_regions].abs()

    meta = metadata.copy()
    meta.index = meta.index.astype(str)
    beta.index = beta.index.astype(str)

    common_samples = meta.index.intersection(beta.index)
    meta = meta.loc[common_samples]
    beta = beta.loc[common_samples]

    rows = []
    for stage in ordered_stages(meta, stage_col, stage_order):
        sample_ids = meta.index[meta[stage_col].astype(str) == str(stage)]
        sub = beta.loc[sample_ids]
        valid = sub.notna().mean(axis=0) >= min_non_missing_frac
        p = sub.loc[:, valid].mean(axis=0, skipna=True)
        w = weights.loc[p.index]
        h = pd.Series(binary_entropy(p, eps=eps), index=p.index)
        denom = float(w.sum())
        s_epi_age = float((w * h).sum() / denom) if denom > 0 else np.nan

        rows.append(
            {
                "stage": stage,
                "n_cells": int(len(sample_ids)),
                "n_regions": int(valid.sum()),
                "weight_sum": denom,
                "s_epi_age": s_epi_age,
            }
        )

    return pd.DataFrame(rows)


def compute_age_projection(
    beta_matrix,
    metadata,
    age_weights,
    stage_col="stage",
    stage_order=None,
    min_non_missing_frac=0.30,
):
    beta = as_numeric_frame(beta_matrix)
    weights = pd.to_numeric(age_weights, errors="coerce").dropna()
    weights.index = weights.index.astype(str)
    beta.columns = beta.columns.astype(str)

    common_regions = beta.columns.intersection(weights.index)
    beta = beta.loc[:, common_regions]
    weights = weights.loc[common_regions]

    meta = metadata.copy()
    meta.index = meta.index.astype(str)
    beta.index = beta.index.astype(str)

    common_samples = meta.index.intersection(beta.index)
    meta = meta.loc[common_samples]
    beta = beta.loc[common_samples]

    rows = []
    for stage in ordered_stages(meta, stage_col, stage_order):
        sample_ids = meta.index[meta[stage_col].astype(str) == str(stage)]
        sub = beta.loc[sample_ids]
        valid = sub.notna().mean(axis=0) >= min_non_missing_frac
        p = sub.loc[:, valid].mean(axis=0, skipna=True)
        w = weights.loc[p.index]
        denom = float(np.abs(w).sum())
        projection = float((w * p).sum() / denom) if denom > 0 else np.nan
        rows.append(
            {
                "stage": stage,
                "n_cells": int(len(sample_ids)),
                "n_regions": int(valid.sum()),
                "age_projection": projection,
            }
        )

    return pd.DataFrame(rows)


def compute_reset_index(stage_df, aged_value, young_value, value_col="s_epi_age", eps=1e-12):
    df = stage_df.copy()
    denom = float(aged_value - young_value)
    if abs(denom) < eps:
        raise ValueError("aged_value - young_value is too close to zero.")
    df["reset_index"] = (float(aged_value) - df[value_col]) / denom
    return df


def compute_rna_entropy(expr, eps=1e-12, normalize=False):
    x = as_numeric_frame(expr).clip(lower=0)
    row_sum = x.sum(axis=1)
    x = x.loc[row_sum > 0]
    p = x.div(x.sum(axis=1), axis=0).clip(lower=eps)
    s = -(p * np.log(p)).sum(axis=1)
    if normalize and p.shape[1] > 1:
        s = s / np.log(p.shape[1])
    return s


def compute_stage_metric_mean(metric, metadata, stage_col="stage", stage_order=None, metric_name="metric"):
    meta = metadata.copy()
    meta.index = meta.index.astype(str)
    metric = pd.Series(metric)
    metric.index = metric.index.astype(str)
    common = meta.index.intersection(metric.index)
    meta = meta.loc[common]
    metric = metric.loc[common]

    rows = []
    for stage in ordered_stages(meta, stage_col, stage_order):
        ids = meta.index[meta[stage_col].astype(str) == str(stage)]
        vals = metric.loc[ids].dropna()
        rows.append(
            {
                "stage": stage,
                "n_cells": int(len(vals)),
                f"{metric_name}_mean": float(vals.mean()) if len(vals) else np.nan,
                f"{metric_name}_sd": float(vals.std(ddof=1)) if len(vals) > 1 else np.nan,
            }
        )
    return pd.DataFrame(rows)


def bootstrap_ground_zero(
    beta_matrix,
    metadata,
    age_weights,
    stage_col="stage",
    stage_order=None,
    n_boot=1000,
    min_non_missing_frac=0.30,
    seed=1,
):
    rng = np.random.default_rng(seed)
    beta = as_numeric_frame(beta_matrix)
    weights = pd.to_numeric(age_weights, errors="coerce").dropna()
    weights.index = weights.index.astype(str)
    beta.columns = beta.columns.astype(str)

    common_regions = beta.columns.intersection(weights.index)
    beta = beta.loc[:, common_regions]
    weights = weights.loc[common_regions].abs()

    meta = metadata.copy()
    meta.index = meta.index.astype(str)
    beta.index = beta.index.astype(str)
    common_samples = meta.index.intersection(beta.index)
    meta = meta.loc[common_samples]
    beta = beta.loc[common_samples]

    stages = ordered_stages(meta, stage_col, stage_order)
    records = []
    winners = []

    for b in range(n_boot):
        stage_values = {}
        for stage in stages:
            ids = np.asarray(meta.index[meta[stage_col].astype(str) == str(stage)])
            if len(ids) == 0:
                continue
            boot_ids = rng.choice(ids, size=len(ids), replace=True)
            sub = beta.loc[boot_ids]
            valid = sub.notna().mean(axis=0) >= min_non_missing_frac
            p = sub.loc[:, valid].mean(axis=0, skipna=True)
            if len(p) == 0:
                continue
            w = weights.loc[p.index]
            h = pd.Series(binary_entropy(p), index=p.index)
            denom = float(w.sum())
            val = float((w * h).sum() / denom) if denom > 0 else np.nan
            stage_values[stage] = val
            records.append({"boot": b, "stage": stage, "s_epi_age": val})

        finite = {k: v for k, v in stage_values.items() if np.isfinite(v)}
        if finite:
            winners.append(min(finite, key=finite.get))

    freq = (
        pd.Series(winners)
        .value_counts()
        .rename_axis("stage")
        .reset_index(name="n_min")
    )
    freq["frequency"] = freq["n_min"] / float(n_boot)
    return freq, pd.DataFrame(records)


def compute_ot_cost(X_source, X_target, reg=0.05):
    import ot

    X_source = np.asarray(X_source, dtype=float)
    X_target = np.asarray(X_target, dtype=float)
    n = X_source.shape[0]
    m = X_target.shape[0]
    a = np.ones(n) / n
    b = np.ones(m) / m
    C = ot.dist(X_source, X_target, metric="euclidean")
    max_c = float(C.max())
    if max_c > 0:
        C = C / max_c
    G = ot.sinkhorn(a, b, C, reg)
    return float(np.sum(G * C))


def self_test():
    beta = pd.DataFrame(
        [
            [0.1, 0.8, 0.5, np.nan],
            [0.2, 0.7, 0.4, 0.9],
            [0.3, 0.4, 0.2, 0.8],
            [0.4, 0.3, 0.2, 0.7],
            [0.1, 0.2, 0.1, 0.6],
            [0.2, 0.2, 0.1, 0.5],
        ],
        index=["c1", "c2", "c3", "c4", "c5", "c6"],
        columns=["r1", "r2", "r3", "r4"],
    )
    meta = pd.DataFrame(
        {"stage": ["MII oocyte", "MII oocyte", "zygote", "zygote", "2-cell", "2-cell"]},
        index=beta.index,
    )
    weights = pd.Series({"r1": 1.0, "r2": -0.5, "r3": 0.25, "r4": 0.1})

    print("stage s_epi")
    print(compute_stage_epi_entropy(beta, meta).to_string(index=False))
    print()
    print("stage s_epi_age")
    age_df = compute_age_weighted_epi_entropy(beta, meta, weights)
    print(age_df.to_string(index=False))
    print()
    print("reset index")
    print(compute_reset_index(age_df, aged_value=0.65, young_value=0.45).to_string(index=False))
    print()
    print("bootstrap")
    freq, _ = bootstrap_ground_zero(beta, meta, weights, n_boot=20, seed=7)
    print(freq.to_string(index=False))


def main():
    parser = argparse.ArgumentParser()
    parser.add_argument("--self-test", action="store_true")
    args = parser.parse_args()
    if args.self_test:
        self_test()


if __name__ == "__main__":
    main()
```
