## Supplemental code for "An entropy-based diagnostic framework for characterizing methylation state dynamics during preimplantation development": 03_download_priority_data.sh

```
#!/usr/bin/env bash
set -euo pipefail

cd /root/autodl-tmp/TRO_Project

mkdir -p data_raw/GSE102970_sperm_methylation
mkdir -p data_raw/GSE81233_embryo_methylation
mkdir -p logs

echo "===== CHECK REMOTE FILE SIZES ====="

urls=(
"https://ftp.ncbi.nlm.nih.gov/geo/series/GSE102nnn/GSE102970/suppl/GSE102970_Exposure_Matrix.csv.gz"
"https://ftp.ncbi.nlm.nih.gov/geo/series/GSE102nnn/GSE102970/suppl/GSE102970_mval_clean.csv.gz"
"https://ftp.ncbi.nlm.nih.gov/geo/series/GSE102nnn/GSE102970/suppl/GSE102970_readme_for_Exposure_Matrix.txt"
"https://ftp.ncbi.nlm.nih.gov/geo/series/GSE81nnn/GSE81233/suppl/GSE81233_RAW.tar"
"https://ftp.ncbi.nlm.nih.gov/geo/series/GSE81nnn/GSE81233/suppl/filelist.txt"
)

for url in "${urls[@]}"; do
  echo
  echo "URL: $url"
  curl -L -I "$url" 2>/dev/null | awk 'BEGIN{IGNORECASE=1} /^HTTP\// || /^content-length:/ || /^content-type:/ || /^last-modified:/ {print}'
done

echo
echo "===== DOWNLOAD GSE102970 SMALL/CLEAN FILES ====="
cd data_raw/GSE102970_sperm_methylation

wget -c https://ftp.ncbi.nlm.nih.gov/geo/series/GSE102nnn/GSE102970/suppl/GSE102970_Exposure_Matrix.csv.gz
wget -c https://ftp.ncbi.nlm.nih.gov/geo/series/GSE102nnn/GSE102970/suppl/GSE102970_mval_clean.csv.gz
wget -c https://ftp.ncbi.nlm.nih.gov/geo/series/GSE102nnn/GSE102970/suppl/GSE102970_readme_for_Exposure_Matrix.txt
wget -c https://ftp.ncbi.nlm.nih.gov/geo/series/GSE102nnn/GSE102970/suppl/filelist.txt

echo
echo "===== DOWNLOAD GSE81233 FILELIST ONLY ====="
cd /root/autodl-tmp/TRO_Project/data_raw/GSE81233_embryo_methylation
wget -c https://ftp.ncbi.nlm.nih.gov/geo/series/GSE81nnn/GSE81233/suppl/filelist.txt

echo
echo "===== LOCAL FILES ====="
cd /root/autodl-tmp/TRO_Project
find data_raw -maxdepth 3 -type f -printf '%p\t%k KB\n' | sort

echo
echo "DONE_DOWNLOAD_PRIORITY"
```
