## Supplemental code for "An entropy-based diagnostic framework for characterizing methylation state dynamics during preimplantation development": 04_inspect_downloaded_data.py

```
from pathlib import Path
import gzip
import pandas as pd

ROOT = Path("/root/autodl-tmp/TRO_Project")
GSE102970 = ROOT / "data_raw" / "GSE102970_sperm_methylation"
GSE81233 = ROOT / "data_raw" / "GSE81233_embryo_methylation"

def file_size(path):
    mb = path.stat().st_size / 1024 / 1024
    return f"{mb:.2f} MB"

def count_gzip_lines(path):
    n = 0
    with gzip.open(path, "rt", errors="replace") as f:
        for n, _ in enumerate(f, start=1):
            pass
    return n

def inspect_gse102970():
    print("===== GSE102970 FILES =====")
    for p in sorted(GSE102970.glob("*")):
        print(f"{p.name}\t{file_size(p)}")

    readme = GSE102970 / "GSE102970_readme_for_Exposure_Matrix.txt"
    print("\n===== EXPOSURE README =====")
    print(readme.read_text(errors="replace")[:4000])

    exposure_path = GSE102970 / "GSE102970_Exposure_Matrix.csv.gz"
    print("\n===== EXPOSURE MATRIX =====")
    exposure = pd.read_csv(exposure_path)
    print("shape:", exposure.shape)
    print("columns:")
    for i, col in enumerate(exposure.columns):
        print(f"  {i}: {col}")
    print("\nhead:")
    print(exposure.head().to_string())

    mval_path = GSE102970 / "GSE102970_mval_clean.csv.gz"
    print("\n===== MVAL MATRIX HEADER =====")
    with gzip.open(mval_path, "rt", errors="replace") as f:
        header = f.readline().rstrip("\n").split(",")
        first_data = f.readline().rstrip("\n").split(",")
    print("n_columns_including_index:", len(header))
    print("first 15 header fields:", header[:15])
    print("first data row first 8 fields:", first_data[:8])

    print("\nCounting mval rows...")
    n_lines = count_gzip_lines(mval_path)
    print("mval text lines including header:", n_lines)
    print("mval CpG rows:", n_lines - 1)
    print("mval sample columns if first column is CpG id:", len(header) - 1)

    mval_sample_cols = [str(x) for x in header[1:]]
    exposure_text = exposure.astype(str)
    print("\n===== SAMPLE OVERLAP HINTS =====")
    for col in exposure.columns:
        vals = set(exposure_text[col].dropna())
        overlap = len(vals.intersection(mval_sample_cols))
        if overlap:
            print(f"exposure column '{col}' overlaps mval sample columns: {overlap}")

    possible_age_cols = [c for c in exposure.columns if "age" in c.lower()]
    print("possible age columns:", possible_age_cols)

def inspect_gse81233_filelist():
    print("\n===== GSE81233 FILELIST PREVIEW =====")
    path = GSE81233 / "filelist.txt"
    lines = path.read_text(errors="replace").splitlines()
    print("n_lines:", len(lines))
    for line in lines[:80]:
        print(line)

    print("\n===== GSE81233 FILELIST KEYWORD COUNTS =====")
    keywords = ["methyl", "CpG", "bed", "cov", "txt", "gz", "MII", "zygote", "2cell", "2-cell", "4cell", "8cell", "morula", "blast", "ICM", "TE"]
    lower_lines = [x.lower() for x in lines]
    for kw in keywords:
        print(f"{kw}: {sum(kw.lower() in x for x in lower_lines)}")

def main():
    inspect_gse102970()
    inspect_gse81233_filelist()

if __name__ == "__main__":
    main()
```
