## Supplemental code for "An entropy-based diagnostic framework for characterizing methylation state dynamics during preimplantation development": 05_inspect_gse102970_age_sources.py

```
from pathlib import Path
import gzip
import re
import pandas as pd
import requests

ROOT = Path("/root/autodl-tmp/TRO_Project")
RAW = ROOT / "data_raw" / "GSE102970_sperm_methylation"
TABLES = ROOT / "results" / "tables"
LOGS = ROOT / "logs"
RAW.mkdir(parents=True, exist_ok=True)
TABLES.mkdir(parents=True, exist_ok=True)
LOGS.mkdir(parents=True, exist_ok=True)

SERIES_MATRIX_URL = "https://ftp.ncbi.nlm.nih.gov/geo/series/GSE102nnn/GSE102970/matrix/GSE102970_series_matrix.txt.gz"
SUPP_XLSX_URL = "https://static-content.springer.com/esm/art%3A10.1038%2Fs41598-020-80857-2/MediaObjects/41598_2020_80857_MOESM1_ESM.xlsx"

def download(url, out):
    out = Path(out)
    if out.exists() and out.stat().st_size > 0:
        print(f"exists: {out} ({out.stat().st_size} bytes)")
        return out
    print(f"downloading: {url}")
    r = requests.get(url, timeout=120)
    r.raise_for_status()
    out.write_bytes(r.content)
    print(f"wrote: {out} ({out.stat().st_size} bytes)")
    return out

def inspect_series_matrix():
    path = download(SERIES_MATRIX_URL, RAW / "GSE102970_series_matrix.txt.gz")
    print("\n===== SERIES MATRIX AGE/BMI/SMOKING SEARCH =====")
    keywords = ["age", "bmi", "smok", "male", "infert", "phenotype", "characteristics", "title", "geo_accession"]
    hits = []
    with gzip.open(path, "rt", errors="replace") as f:
        for line_no, line in enumerate(f, start=1):
            low = line.lower()
            if any(k in low for k in keywords):
                hits.append((line_no, line.rstrip("\n")))
    print(f"hit lines: {len(hits)}")
    for line_no, line in hits[:120]:
        print(f"{line_no}: {line[:500]}")
    pd.DataFrame(hits, columns=["line_no", "line"]).to_csv(
        TABLES / "gse102970_series_matrix_keyword_hits.tsv", sep="\t", index=False
    )

def inspect_supplement_xlsx():
    path = download(SUPP_XLSX_URL, RAW / "Oluwayiose_2021_SciRep_supplement.xlsx")
    print("\n===== SUPPLEMENT XLSX =====")
    xls = pd.ExcelFile(path)
    print("sheet names:")
    for s in xls.sheet_names:
        print(" -", s)

    summary_rows = []
    keyword_rows = []
    keywords = ["age", "cpg", "dmr", "methyl", "beta", "estimate", "effect", "coef", "p", "q", "fdr"]

    for sheet in xls.sheet_names:
        df = pd.read_excel(path, sheet_name=sheet)
        cols = [str(c) for c in df.columns]
        summary_rows.append({
            "sheet": sheet,
            "n_rows": df.shape[0],
            "n_cols": df.shape[1],
            "columns": " | ".join(cols[:30]),
        })

        print(f"\n--- SHEET: {sheet} shape={df.shape} ---")
        print("columns:", cols[:40])
        print(df.head(5).to_string(index=False))

        for col in cols:
            low = col.lower()
            if any(k in low for k in keywords):
                keyword_rows.append({"sheet": sheet, "where": "column", "hit": col})

        text_preview = df.astype(str).head(30)
        for r_idx, row in text_preview.iterrows():
            joined = " | ".join(row.tolist())
            low = joined.lower()
            if any(k in low for k in keywords):
                keyword_rows.append({"sheet": sheet, "where": f"row_{r_idx}", "hit": joined[:1000]})

    pd.DataFrame(summary_rows).to_csv(
        TABLES / "gse102970_supplement_sheet_summary.tsv", sep="\t", index=False
    )
    pd.DataFrame(keyword_rows).to_csv(
        TABLES / "gse102970_supplement_keyword_hits.tsv", sep="\t", index=False
    )

def main():
    inspect_series_matrix()
    inspect_supplement_xlsx()
    print("\nDONE_INSPECT_AGE_SOURCES")

if __name__ == "__main__":
    main()
```
