## Supplemental code for "An entropy-based diagnostic framework for characterizing methylation state dynamics during preimplantation development": 05b_summarize_gse102970_supplement.py

```
from pathlib import Path
import pandas as pd

ROOT = Path("/root/autodl-tmp/TRO_Project")
xlsx = ROOT / "data_raw" / "GSE102970_sperm_methylation" / "Oluwayiose_2021_SciRep_supplement.xlsx"
outdir = ROOT / "results" / "tables"
outdir.mkdir(parents=True, exist_ok=True)

keywords = ["age", "cpg", "dmr", "methyl", "estimate", "effect", "coef", "beta", "fdr", "adj", "p.value", "p-value", "chr", "start", "end"]

xls = pd.ExcelFile(xlsx)
summary = []
hits = []

for sheet in xls.sheet_names:
    df = pd.read_excel(xlsx, sheet_name=sheet)
    cols = [str(c) for c in df.columns]

    summary.append({
        "sheet": sheet,
        "n_rows": df.shape[0],
        "n_cols": df.shape[1],
        "columns": " | ".join(cols)
    })

    for col in cols:
        low = col.lower()
        if any(k in low for k in keywords):
            hits.append({"sheet": sheet, "hit_type": "column", "hit": col})

    preview = df.head(15)
    for idx, row in preview.iterrows():
        joined = " | ".join(str(x) for x in row.tolist())
        low = joined.lower()
        if any(k in low for k in keywords):
            hits.append({"sheet": sheet, "hit_type": f"row_{idx}", "hit": joined[:1000]})

    safe_sheet = sheet.replace(" ", "_").replace("/", "_")
    df.head(20).to_csv(outdir / f"preview_{safe_sheet}.tsv", sep="\t", index=False)

summary_df = pd.DataFrame(summary)
hits_df = pd.DataFrame(hits)

summary_df.to_csv(outdir / "gse102970_supplement_sheet_summary.tsv", sep="\t", index=False)
hits_df.to_csv(outdir / "gse102970_supplement_keyword_hits.tsv", sep="\t", index=False)

print("XLSX:", xlsx)
print("SHEETS:")
print(summary_df[["sheet", "n_rows", "n_cols"]].to_string(index=False))
print()
print("KEYWORD HITS:")
if len(hits_df):
    print(hits_df.head(160).to_string(index=False))
else:
    print("No keyword hits")
print()
print("WROTE:")
print(outdir / "gse102970_supplement_sheet_summary.tsv")
print(outdir / "gse102970_supplement_keyword_hits.tsv")
```
