## Supplemental code for "An entropy-based diagnostic framework for characterizing methylation state dynamics during preimplantation development": 06_extract_gse102970_age_weights.py

```
from pathlib import Path
import gzip
import re
import numpy as np
import pandas as pd

ROOT = Path("/root/autodl-tmp/TRO_Project")
RAW = ROOT / "data_raw" / "GSE102970_sperm_methylation"
META = ROOT / "data_processed" / "metadata"
METH = ROOT / "data_processed" / "methylation_matrix"
TABLES = ROOT / "results" / "tables"

for d in [META, METH, TABLES]:
    d.mkdir(parents=True, exist_ok=True)

XLSX = RAW / "Oluwayiose_2021_SciRep_supplement.xlsx"
MVAL = RAW / "GSE102970_mval_clean.csv.gz"

def find_header_row(raw_df, required):
    for i in range(raw_df.shape[0]):
        values = [str(x).strip() for x in raw_df.iloc[i].tolist()]
        lower = [x.lower() for x in values]
        ok = all(any(req.lower() == v for v in lower) for req in required)
        if ok:
            return i
    raise RuntimeError(f"Could not find header row with required columns: {required}")

def clean_colname(x):
    return str(x).strip()

def read_table_s6():
    raw = pd.read_excel(XLSX, sheet_name="Table S6", header=None)
    header_i = find_header_row(raw, ["cluster_name", "cluster_sites.y", "chr", "start", "end"])
    cols = [clean_colname(x) for x in raw.iloc[header_i].tolist()]
    df = raw.iloc[header_i + 1:].copy()
    df.columns = cols
    df = df.dropna(how="all")

    needed = ["cluster_name", "cluster_sites.y", "chr", "start", "end", "effct_size/5yrs"]
    missing = [c for c in needed if c not in df.columns]
    if missing:
        raise RuntimeError(f"Missing expected Table S6 columns: {missing}. Columns are: {list(df.columns)}")

    df = df[df["cluster_name"].astype(str).str.startswith("cluster_")].copy()
    df["chr"] = df["chr"].astype(str)
    df["start"] = pd.to_numeric(df["start"], errors="coerce").astype("Int64")
    df["end"] = pd.to_numeric(df["end"], errors="coerce").astype("Int64")
    df["age_weight_5yr"] = pd.to_numeric(df["effct_size/5yrs"], errors="coerce")
    df["age_weight_per_year"] = df["age_weight_5yr"] / 5.0
    df["n_cpg_listed"] = df["cluster_sites.y"].astype(str).apply(lambda s: len([x for x in s.split(";") if x.startswith("cg")]))

    keep = [
        "cluster_name",
        "chr",
        "start",
        "end",
        "cluster_sites.y",
        "n_cpg_listed",
        "age_weight_5yr",
        "age_weight_per_year",
    ]
    for c in ["exposure_pvalue.y", "exposure_padjusted.y", "n_sites_in_cluster.y"]:
        if c in df.columns:
            keep.append(c)

    return df[keep].rename(columns={"cluster_sites.y": "cpg_list"})

def explode_cpg_weights(dmr_df):
    rows = []
    for _, r in dmr_df.iterrows():
        cpgs = [x.strip() for x in str(r["cpg_list"]).split(";") if x.strip().startswith("cg")]
        for cpg in cpgs:
            rows.append({
                "cpg_id": cpg,
                "cluster_name": r["cluster_name"],
                "chr": r["chr"],
                "start": r["start"],
                "end": r["end"],
                "age_weight_5yr": r["age_weight_5yr"],
                "age_weight_per_year": r["age_weight_per_year"],
            })
    out = pd.DataFrame(rows)
    out = out.dropna(subset=["cpg_id", "age_weight_5yr"])
    return out

def mval_index():
    ids = []
    with gzip.open(MVAL, "rt", errors="replace") as f:
        header = f.readline().rstrip("\n").split(",")
        samples = header[1:]
        for line in f:
            ids.append(line.split(",", 1)[0])
    return pd.Index(ids, name="cpg_id"), samples

def m_to_beta(m):
    return 1.0 / (1.0 + np.exp2(-m))

def make_age_cpg_matrices(cpg_weights):
    wanted = set(cpg_weights["cpg_id"])
    chunks = []
    for chunk in pd.read_csv(MVAL, compression="gzip", chunksize=50000):
        first = chunk.columns[0]
        chunk = chunk.rename(columns={first: "cpg_id"})
        sub = chunk[chunk["cpg_id"].isin(wanted)].copy()
        if len(sub):
            chunks.append(sub)

    if not chunks:
        raise RuntimeError("No age CpGs found in mval_clean.")

    mval_sub = pd.concat(chunks, axis=0)
    mval_sub = mval_sub.drop_duplicates("cpg_id").set_index("cpg_id")
    beta_sub = mval_sub.apply(pd.to_numeric, errors="coerce").apply(m_to_beta)

    beta_sample_by_cpg = beta_sub.T
    beta_sample_by_cpg.index.name = "sample_id"
    beta_sample_by_cpg.to_csv(METH / "GSE102970_age_cpg_beta_sample_by_cpg.tsv", sep="\t")

    weights = cpg_weights.drop_duplicates("cpg_id").set_index("cpg_id")["age_weight_5yr"]
    common = beta_sample_by_cpg.columns.intersection(weights.index)
    beta_common = beta_sample_by_cpg[common]
    w = weights.loc[common]

    eps = 1e-6
    p = beta_common.clip(eps, 1 - eps)
    h = -(p * np.log(p) + (1 - p) * np.log(1 - p))
    s_epi_age = h.mul(w.abs(), axis=1).sum(axis=1) / w.abs().sum()
    age_projection = beta_common.mul(w, axis=1).sum(axis=1) / w.abs().sum()

    sample_metrics = pd.DataFrame({
        "sample_id": beta_common.index,
        "n_age_cpg": len(common),
        "s_epi_age": s_epi_age.values,
        "age_projection": age_projection.values,
    })
    sample_metrics.to_csv(TABLES / "GSE102970_sperm_age_cpg_sample_metrics.tsv", sep="\t", index=False)

    return {
        "n_rows_mval_subset": beta_sample_by_cpg.shape[0],
        "n_cols_age_cpg": beta_sample_by_cpg.shape[1],
        "n_common_weighted_cpg": len(common),
        "sample_metrics": sample_metrics,
    }

def main():
    print("===== PARSE TABLE S6 AGE-DMR WEIGHTS =====")
    dmr = read_table_s6()
    dmr_out = META / "GSE102970_TableS6_age_dmr_weights.tsv"
    dmr.to_csv(dmr_out, sep="\t", index=False)

    cpg = explode_cpg_weights(dmr)
    cpg_out = META / "GSE102970_TableS6_age_cpg_weights.tsv"
    cpg.to_csv(cpg_out, sep="\t", index=False)

    print("DMR rows:", len(dmr))
    print("CpG weight rows:", len(cpg))
    print("Unique CpGs:", cpg["cpg_id"].nunique())
    print("Positive DMR weights:", int((dmr["age_weight_5yr"] > 0).sum()))
    print("Negative DMR weights:", int((dmr["age_weight_5yr"] < 0).sum()))
    print("DMR output:", dmr_out)
    print("CpG output:", cpg_out)

    print("\n===== CHECK OVERLAP WITH GSE102970 MVAL =====")
    idx, samples = mval_index()
    overlap = idx.intersection(pd.Index(cpg["cpg_id"].unique()))
    print("mval CpGs:", len(idx))
    print("mval samples:", len(samples))
    print("age CpGs overlapping mval:", len(overlap))
    print("overlap fraction:", len(overlap) / max(1, cpg["cpg_id"].nunique()))

    print("\n===== MAKE AGE-CPG BETA SUBSET AND SAMPLE METRICS =====")
    metrics_info = make_age_cpg_matrices(cpg[cpg["cpg_id"].isin(overlap)].copy())
    print("beta sample x CpG shape:", metrics_info["n_rows_mval_subset"], metrics_info["n_cols_age_cpg"])
    print("weighted CpGs used:", metrics_info["n_common_weighted_cpg"])
    print(metrics_info["sample_metrics"].head().to_string(index=False))

    print("\n===== OUTPUT PREVIEW =====")
    print(dmr.head(10).to_string(index=False))
    print("\nDONE_EXTRACT_AGE_WEIGHTS")

if __name__ == "__main__":
    main()
```
