## Supplemental code for "An entropy-based diagnostic framework for characterizing methylation state dynamics during preimplantation development": 07_plan_gse81233_pilot_download.py

```
from pathlib import Path
import re
import pandas as pd
import requests

ROOT = Path("/root/autodl-tmp/TRO_Project")
FILELIST = ROOT / "data_raw" / "GSE81233_embryo_methylation" / "filelist.txt"
TABLES = ROOT / "results" / "tables"
TABLES.mkdir(parents=True, exist_ok=True)

def gsm_group(gsm):
    m = re.match(r"GSM(\d+)", gsm)
    if not m:
        return None
    digits = m.group(1)
    return "GSM" + digits[:4] + "nnn"

def sample_url(filename):
    gsm = filename.split("_", 1)[0]
    group = gsm_group(gsm)
    return f"https://ftp.ncbi.nlm.nih.gov/geo/samples/{group}/{gsm}/suppl/{filename}"

def parse_stage(filename):
    name = filename.lower()
    if ".cmet.bed.gz" not in name:
        return None
    if "-mii-" in name or "-mii" in name:
        return "MII oocyte"
    if "-pn" in name:
        return "zygote/PN"
    if "-2c-" in name:
        return "2-cell"
    if "-4c-" in name:
        return "4-cell"
    if "-8c-" in name:
        return "8-cell"
    if "morula" in name or "morluae" in name or "morlua" in name:
        return "morula"
    if "-bst" in name or "blast" in name:
        return "blastocyst"
    if "-icm" in name:
        return "ICM"
    if "-te" in name:
        return "TE"
    if "-gv" in name:
        return "GV oocyte"
    if "hsp" in name:
        return "sperm"
    return "other"

def read_filelist():
    rows = []
    with FILELIST.open(errors="replace") as f:
        header = f.readline().rstrip("\n").split("\t")
        for line in f:
            parts = line.rstrip("\n").split("\t")
            if len(parts) < 5:
                continue
            kind, name, time, size, typ = parts[:5]
            if kind != "File":
                continue
            size = int(size) if str(size).isdigit() else None
            rows.append({"kind": kind, "filename": name, "time": time, "size_bytes": size, "type": typ})
    df = pd.DataFrame(rows)
    df["size_mb"] = df["size_bytes"] / 1024 / 1024
    df["stage"] = df["filename"].apply(parse_stage)
    df["url"] = df["filename"].apply(sample_url)
    return df

def head_url(url, timeout=30):
    try:
        r = requests.head(url, allow_redirects=True, timeout=timeout)
        return {
            "http_status": r.status_code,
            "content_length": r.headers.get("Content-Length", ""),
            "content_type": r.headers.get("Content-Type", ""),
            "final_url": r.url,
        }
    except Exception as e:
        return {
            "http_status": "ERROR",
            "content_length": "",
            "content_type": "",
            "final_url": str(e),
        }

def main():
    df = read_filelist()
    cmet = df[df["filename"].str.endswith(".Cmet.bed.gz", na=False)].copy()
    cmet = cmet[cmet["stage"].notna()].copy()
    cmet.to_csv(TABLES / "GSE81233_cmet_manifest.tsv", sep="\t", index=False)

    print("===== CMET STAGE SUMMARY =====")
    summary = (
        cmet.groupby("stage")
        .agg(n_files=("filename", "size"),
             min_mb=("size_mb", "min"),
             median_mb=("size_mb", "median"),
             max_mb=("size_mb", "max"),
             total_gb=("size_bytes", lambda x: x.sum() / 1024 / 1024 / 1024))
        .reset_index()
        .sort_values("stage")
    )
    print(summary.to_string(index=False))
    summary.to_csv(TABLES / "GSE81233_cmet_stage_summary.tsv", sep="\t", index=False)

    stage_order = ["MII oocyte", "zygote/PN", "2-cell", "4-cell", "8-cell", "morula", "blastocyst", "ICM", "TE"]
    pilot_rows = []
    for stage in stage_order:
        sub = cmet[cmet["stage"] == stage].sort_values("size_bytes")
        if len(sub) == 0:
            continue
        n = 2 if stage not in ["blastocyst", "ICM", "TE"] else 1
        pilot_rows.append(sub.head(n))
    pilot = pd.concat(pilot_rows, axis=0) if pilot_rows else pd.DataFrame()
    pilot.to_csv(TABLES / "GSE81233_pilot_download_manifest.tsv", sep="\t", index=False)

    print("\n===== PILOT DOWNLOAD CANDIDATES =====")
    print(pilot[["stage", "filename", "size_mb", "url"]].to_string(index=False))
    print("pilot total GB:", pilot["size_bytes"].sum() / 1024 / 1024 / 1024)

    print("\n===== DIRECT URL HEAD TEST =====")
    checks = []
    for _, row in pilot.iterrows():
        info = head_url(row["url"])
        checks.append({**row[["stage", "filename", "size_mb", "url"]].to_dict(), **info})
        print(row["stage"], row["filename"], info["http_status"], info["content_length"], info["content_type"])
    checks_df = pd.DataFrame(checks)
    checks_df.to_csv(TABLES / "GSE81233_pilot_url_head_check.tsv", sep="\t", index=False)

    print("\nWROTE:")
    print(TABLES / "GSE81233_cmet_manifest.tsv")
    print(TABLES / "GSE81233_cmet_stage_summary.tsv")
    print(TABLES / "GSE81233_pilot_download_manifest.tsv")
    print(TABLES / "GSE81233_pilot_url_head_check.tsv")
    print("DONE_PLAN_GSE81233_PILOT")

if __name__ == "__main__":
    main()
```
