## Supplemental code for "An entropy-based diagnostic framework for characterizing methylation state dynamics during preimplantation development": 08_download_gse81233_pilot.sh

```
#!/usr/bin/env bash
set -euo pipefail

cd /root/autodl-tmp/TRO_Project

manifest="results/tables/GSE81233_pilot_download_manifest.tsv"
outdir="data_raw/GSE81233_embryo_methylation/pilot_cmet"
mkdir -p "$outdir" logs

echo "===== START DOWNLOAD $(date) ====="
echo "manifest: $manifest"
echo "outdir: $outdir"

awk -F '\t' '
NR==1 {
  for (i=1; i<=NF; i++) {
    if ($i=="stage") stage_i=i
    if ($i=="filename") filename_i=i
    if ($i=="url") url_i=i
  }
  next
}
{
  print $stage_i "\t" $filename_i "\t" $url_i
}
' "$manifest" | while IFS=$'\t' read -r stage filename url; do
  echo
  echo "===== FILE ====="
  echo "stage: $stage"
  echo "filename: $filename"
  echo "url: $url"

  cd /root/autodl-tmp/TRO_Project/"$outdir"
  wget -c --tries=10 --timeout=60 "$url"
done

cd /root/autodl-tmp/TRO_Project

echo
echo "===== DOWNLOADED FILES ====="
find "$outdir" -type f -name "*.Cmet.bed.gz" -printf '%p\t%k KB\n' | sort

echo
echo "===== DISK ====="
df -h .

echo
echo "===== DONE DOWNLOAD $(date) ====="
```
