## Supplemental code for "An entropy-based diagnostic framework for characterizing methylation state dynamics during preimplantation development": 09_stream_gse81233_age_dmr_aggregation.py

```
from pathlib import Path
import argparse, gzip, os, time
from bisect import bisect_left, bisect_right
import numpy as np
import pandas as pd
import requests

ROOT = Path("/root/autodl-tmp/TRO_Project")
DMR_PATH = ROOT / "data_processed/metadata/GSE102970_TableS6_age_dmr_weights.tsv"
DEFAULT_MANIFEST = ROOT / "results/tables/GSE81233_full_relevant_cmet_manifest.tsv"
LOCAL_DIR = ROOT / "data_raw/GSE81233_embryo_methylation/pilot_cmet"
TABLES = ROOT / "results/tables"

STAGE_ORDER = ["MII oocyte","zygote/PN","2-cell","4-cell","8-cell","morula","blastocyst","ICM","TE"]

def binary_entropy(p, eps=1e-6):
    p = np.clip(np.asarray(p, dtype=float), eps, 1 - eps)
    return -(p*np.log(p) + (1-p)*np.log(1-p))

def load_dmrs():
    dmr = pd.read_csv(DMR_PATH, sep="\t")
    dmr = dmr.dropna(subset=["cluster_name","chr","start","end","age_weight_5yr"]).copy()
    dmr["start"] = dmr["start"].astype(int)
    dmr["end"] = dmr["end"].astype(int)
    dmr["age_weight_5yr"] = pd.to_numeric(dmr["age_weight_5yr"], errors="coerce")
    dmr = dmr.dropna(subset=["age_weight_5yr"]).reset_index(drop=True)

    intervals, starts, max_width = {}, {}, {}
    for idx, r in dmr.iterrows():
        cb = str(r["chr"]).encode()
        intervals.setdefault(cb, []).append((int(r["start"]), int(r["end"]), idx))
    for cb in intervals:
        intervals[cb].sort()
        starts[cb] = [x[0] for x in intervals[cb]]
        max_width[cb] = max(e - s + 1 for s, e, _ in intervals[cb])
    return dmr, intervals, starts, max_width

def sample_id_from_filename(fn):
    return str(fn).split("_", 1)[0]

def open_cmet_stream(url, local_path=None, expected_size=None):
    if local_path and Path(local_path).exists():
        p = Path(local_path)
        if expected_size is None or p.stat().st_size >= int(expected_size) * 0.98:
            return gzip.open(p, "rb"), f"local:{p}"
    r = requests.get(url, stream=True, timeout=(30, 120))
    r.raise_for_status()
    r.raw.decode_content = False
    return gzip.GzipFile(fileobj=r.raw), f"remote:{url}"

def process_sample(row, dmr, intervals, starts, max_width, outdir, force=False, min_total=1):
    filename = row["filename"]
    sample_id = sample_id_from_filename(filename)
    out = outdir / "sample_dmr" / f"{sample_id}.age_dmr.tsv"
    tmp = out.with_suffix(".tmp")
    out.parent.mkdir(parents=True, exist_ok=True)

    if out.exists() and not force:
        print(f"SKIP {sample_id}: exists", flush=True)
        return out

    local_path = LOCAL_DIR / filename
    expected = int(row["size_bytes"]) if not pd.isna(row["size_bytes"]) else None
    met = np.zeros(len(dmr), dtype=np.float64)
    total = np.zeros(len(dmr), dtype=np.float64)
    sites = np.zeros(len(dmr), dtype=np.int32)

    n_lines = n_cg = n_match = 0
    t0 = time.time()
    fh, source = open_cmet_stream(row["url"], local_path, expected)
    print(f"START {sample_id} stage={row['stage']} source={source}", flush=True)

    with fh:
        for raw in fh:
            if not raw or raw.startswith(b"#"):
                continue
            n_lines += 1
            parts = raw.rstrip(b"\n").split(b"\t")
            if len(parts) < 10:
                continue
            typ = parts[9].strip()
            if typ not in (b"CG", b"CpG", b"cg", b"cpg"):
                continue
            n_cg += 1
            cb = parts[0]
            if cb not in intervals:
                continue
            try:
                pos = int(parts[1])
                tot = int(parts[4])
                m = int(parts[5])
            except Exception:
                continue
            if tot < min_total:
                continue

            left = bisect_left(starts[cb], pos - max_width[cb])
            right = bisect_right(starts[cb], pos)
            for k in range(left, right):
                s, e, idx = intervals[cb][k]
                if s <= pos <= e:
                    met[idx] += m
                    total[idx] += tot
                    sites[idx] += 1
                    n_match += 1

            if n_lines % 5000000 == 0:
                dt = time.time() - t0
                print(f"PROGRESS {sample_id}: lines={n_lines} CG={n_cg} matched={n_match} elapsed_min={dt/60:.1f}", flush=True)

    beta = np.where(total > 0, met / total, np.nan)
    res = dmr[["cluster_name","chr","start","end","age_weight_5yr"]].copy()
    res.insert(0, "sample_id", sample_id)
    res.insert(1, "stage", row["stage"])
    res["filename"] = filename
    res["met_reads"] = met
    res["total_reads"] = total
    res["n_cpg_rows_covered"] = sites
    res["beta"] = beta
    res.to_csv(tmp, sep="\t", index=False)
    os.replace(tmp, out)

    covered = int((total > 0).sum())
    print(f"DONE {sample_id}: lines={n_lines} CG={n_cg} matched={n_match} covered_DMR={covered} elapsed_min={(time.time()-t0)/60:.1f}", flush=True)
    return out

def combine(outdir, manifest, min_sample_frac=0.30):
    sample_dir = outdir / "sample_dmr"
    files = sorted(sample_dir.glob("*.age_dmr.tsv"))
    if not files:
        raise RuntimeError("No sample DMR files to combine.")

    all_df = pd.concat([pd.read_csv(f, sep="\t") for f in files], ignore_index=True)
    all_df.to_csv(outdir / "GSE81233_all_sample_age_dmr_long.tsv", sep="\t", index=False)

    beta_mat = all_df.pivot(index="sample_id", columns="cluster_name", values="beta")
    beta_mat.to_csv(outdir / "GSE81233_age_dmr_beta_sample_by_dmr.tsv", sep="\t")

    rows = []
    for stage in STAGE_ORDER:
        sub = all_df[all_df["stage"] == stage]
        n_samples = sub["sample_id"].nunique()
        if n_samples == 0:
            continue
        g = sub.groupby("cluster_name", as_index=False).agg(
            met_reads=("met_reads","sum"),
            total_reads=("total_reads","sum"),
            n_samples_covered=("total_reads", lambda x: int((x > 0).sum())),
            age_weight_5yr=("age_weight_5yr","first"),
        )
        min_cov = max(1, int(np.ceil(min_sample_frac * n_samples)))
        valid = g[(g["total_reads"] > 0) & (g["n_samples_covered"] >= min_cov)].copy()
        p = valid["met_reads"] / valid["total_reads"]
        h = binary_entropy(p)
        w = valid["age_weight_5yr"].abs().to_numpy()
        signed_w = valid["age_weight_5yr"].to_numpy()

        rows.append({
            "stage": stage,
            "n_samples": n_samples,
            "n_regions_valid": len(valid),
            "min_samples_required": min_cov,
            "s_epi": float(np.mean(h)) if len(valid) else np.nan,
            "s_epi_age": float(np.sum(w*h)/np.sum(w)) if len(valid) and np.sum(w)>0 else np.nan,
            "age_projection": float(np.sum(signed_w*p)/np.sum(w)) if len(valid) and np.sum(w)>0 else np.nan,
        })

    metrics = pd.DataFrame(rows)
    metrics["stage"] = pd.Categorical(metrics["stage"], categories=STAGE_ORDER, ordered=True)
    metrics = metrics.sort_values("stage")
    metrics.to_csv(TABLES / "GSE81233_full_stage_epi_age_metrics.tsv", sep="\t", index=False)

    gz = metrics.dropna(subset=["s_epi_age"]).sort_values("s_epi_age").head(1)
    gz.to_csv(TABLES / "GSE81233_full_ground_zero_summary.tsv", sep="\t", index=False)

    print("===== STAGE METRICS =====")
    print(metrics.to_string(index=False))
    print("===== GROUND ZERO =====")
    print(gz.to_string(index=False))
    return metrics

def main():
    ap = argparse.ArgumentParser()
    ap.add_argument("--manifest", default=str(DEFAULT_MANIFEST))
    ap.add_argument("--outdir", default=str(ROOT / "data_processed/methylation_matrix/GSE81233_age_dmr_stream_full"))
    ap.add_argument("--max-samples", type=int, default=None)
    ap.add_argument("--force", action="store_true")
    ap.add_argument("--combine-only", action="store_true")
    ap.add_argument("--require-match", action="store_true")
    args = ap.parse_args()

    outdir = Path(args.outdir)
    outdir.mkdir(parents=True, exist_ok=True)
    (outdir / "sample_dmr").mkdir(parents=True, exist_ok=True)

    manifest = pd.read_csv(args.manifest, sep="\t")
    manifest = manifest[manifest["stage"].isin(STAGE_ORDER)].copy()
    manifest["stage"] = pd.Categorical(manifest["stage"], categories=STAGE_ORDER, ordered=True)
    manifest = manifest.sort_values(["stage","size_bytes"])
    if args.max_samples:
        manifest = manifest.head(args.max_samples)

    dmr, intervals, starts, max_width = load_dmrs()
    print(f"DMR n={len(dmr)} samples_to_process={len(manifest)} outdir={outdir}", flush=True)

    if not args.combine_only:
        for _, row in manifest.iterrows():
            process_sample(row, dmr, intervals, starts, max_width, outdir, force=args.force)

    metrics = combine(outdir, manifest)
    if args.require_match and metrics["n_regions_valid"].fillna(0).max() == 0:
        raise RuntimeError("Validation failed: no age-DMR regions matched.")

if __name__ == "__main__":
    main()
```
