## Supplemental code for "An entropy-based diagnostic framework for characterizing methylation state dynamics during preimplantation development": 09b_stream_gse81233_shard_worker.py

```
from pathlib import Path
import argparse
import importlib.util
import traceback
import pandas as pd

ROOT = Path("/root/autodl-tmp/TRO_Project")
MOD_PATH = ROOT / "scripts/09_stream_gse81233_age_dmr_aggregation.py"

spec = importlib.util.spec_from_file_location("stream_mod", MOD_PATH)
mod = importlib.util.module_from_spec(spec)
spec.loader.exec_module(mod)

def main():
    ap = argparse.ArgumentParser()
    ap.add_argument("--manifest", required=True)
    ap.add_argument("--outdir", required=True)
    ap.add_argument("--n-shards", type=int, required=True)
    ap.add_argument("--shard-index", type=int, required=True)
    ap.add_argument("--force", action="store_true")
    args = ap.parse_args()

    outdir = Path(args.outdir)
    outdir.mkdir(parents=True, exist_ok=True)
    (outdir / "sample_dmr").mkdir(parents=True, exist_ok=True)

    manifest = pd.read_csv(args.manifest, sep="\t")
    manifest = manifest[manifest["stage"].isin(mod.STAGE_ORDER)].copy()
    manifest["stage"] = pd.Categorical(manifest["stage"], categories=mod.STAGE_ORDER, ordered=True)
    manifest = manifest.sort_values(["stage", "size_bytes"]).reset_index(drop=True)

    shard = manifest.iloc[args.shard_index::args.n_shards].copy()
    dmr, intervals, starts, max_width = mod.load_dmrs()

    print(f"SHARD_START index={args.shard_index}/{args.n_shards} samples={len(shard)}", flush=True)

    failed = []
    for _, row in shard.iterrows():
        try:
            mod.process_sample(row, dmr, intervals, starts, max_width, outdir, force=args.force)
        except Exception as e:
            print(f"ERROR sample={row['filename']} stage={row['stage']} error={repr(e)}", flush=True)
            traceback.print_exc()
            failed.append({
                "stage": row["stage"],
                "filename": row["filename"],
                "url": row["url"],
                "error": repr(e),
            })

    if failed:
        fail_path = outdir / f"failed_shard_{args.shard_index}.tsv"
        pd.DataFrame(failed).to_csv(fail_path, sep="\t", index=False)
        print(f"SHARD_FAILED_WRITTEN {fail_path}", flush=True)

    print(f"SHARD_DONE index={args.shard_index}/{args.n_shards}", flush=True)

if __name__ == "__main__":
    main()
```
