## Supplemental code for "An entropy-based diagnostic framework for characterizing methylation state dynamics during preimplantation development": 09c_cache_gse81233_shard_worker.py

```
from pathlib import Path
import argparse
import importlib.util
import subprocess
import traceback
import os
import pandas as pd

ROOT = Path("/root/autodl-tmp/TRO_Project")
MOD_PATH = ROOT / "scripts/09_stream_gse81233_age_dmr_aggregation.py"
CACHE_DIR = ROOT / "data_raw/GSE81233_embryo_methylation/cache_cmet"

spec = importlib.util.spec_from_file_location("stream_mod", MOD_PATH)
mod = importlib.util.module_from_spec(spec)
spec.loader.exec_module(mod)

def download_to_cache(url, filename):
    CACHE_DIR.mkdir(parents=True, exist_ok=True)
    out = CACHE_DIR / filename
    cmd = [
        "wget",
        "-c",
        "--tries=5",
        "--timeout=60",
        "--read-timeout=120",
        "--waitretry=10",
        "-O",
        str(out),
        url,
    ]
    print("DOWNLOAD", filename, flush=True)
    subprocess.run(cmd, check=True)
    return out

def main():
    ap = argparse.ArgumentParser()
    ap.add_argument("--manifest", required=True)
    ap.add_argument("--outdir", required=True)
    ap.add_argument("--n-shards", type=int, required=True)
    ap.add_argument("--shard-index", type=int, required=True)
    ap.add_argument("--force", action="store_true")
    ap.add_argument("--keep-cache", action="store_true")
    args = ap.parse_args()

    outdir = Path(args.outdir)
    outdir.mkdir(parents=True, exist_ok=True)
    (outdir / "sample_dmr").mkdir(parents=True, exist_ok=True)

    manifest = pd.read_csv(args.manifest, sep="\t")
    manifest = manifest[manifest["stage"].isin(mod.STAGE_ORDER)].copy()
    manifest["stage"] = pd.Categorical(manifest["stage"], categories=mod.STAGE_ORDER, ordered=True)
    manifest = manifest.sort_values(["stage", "size_bytes"]).reset_index(drop=True)
    shard = manifest.iloc[args.shard_index::args.n_shards].copy()

    dmr, intervals, starts, max_width = mod.load_dmrs()

    print(f"CACHE_SHARD_START index={args.shard_index}/{args.n_shards} samples={len(shard)}", flush=True)

    failed = []
    for _, row in shard.iterrows():
        sample_id = mod.sample_id_from_filename(row["filename"])
        done = outdir / "sample_dmr" / f"{sample_id}.age_dmr.tsv"
        if done.exists() and not args.force:
            print(f"SKIP {sample_id}: exists", flush=True)
            continue

        cache_path = CACHE_DIR / row["filename"]
        try:
            if not cache_path.exists() or cache_path.stat().st_size < int(row["size_bytes"]) * 0.98:
                cache_path = download_to_cache(row["url"], row["filename"])
            else:
                print(f"CACHE_EXISTS {row['filename']}", flush=True)

            row2 = row.copy()
            # process_sample will prefer local cache only if this file is also in LOCAL_DIR,
            # so temporarily copy path by directly setting module LOCAL_DIR to cache dir.
            old_local = mod.LOCAL_DIR
            mod.LOCAL_DIR = CACHE_DIR
            try:
                mod.process_sample(row2, dmr, intervals, starts, max_width, outdir, force=args.force)
            finally:
                mod.LOCAL_DIR = old_local

            if not args.keep_cache and cache_path.exists():
                cache_path.unlink()
                print(f"CACHE_REMOVED {cache_path.name}", flush=True)

        except Exception as e:
            print(f"ERROR sample={row['filename']} stage={row['stage']} error={repr(e)}", flush=True)
            traceback.print_exc()
            failed.append({
                "stage": row["stage"],
                "filename": row["filename"],
                "url": row["url"],
                "error": repr(e),
            })

    if failed:
        fail_path = outdir / f"failed_cache_shard_{args.shard_index}.tsv"
        pd.DataFrame(failed).to_csv(fail_path, sep="\t", index=False)
        print(f"CACHE_SHARD_FAILED_WRITTEN {fail_path}", flush=True)

    print(f"CACHE_SHARD_DONE index={args.shard_index}/{args.n_shards}", flush=True)

if __name__ == "__main__":
    main()
```
