## Supplemental code for "An entropy-based diagnostic framework for characterizing methylation state dynamics during preimplantation development": 10_finalize_gse81233_valid204.py

```
from pathlib import Path
import importlib.util
import pandas as pd
import matplotlib.pyplot as plt

ROOT = Path("/root/autodl-tmp/TRO_Project")
OUTDIR = ROOT / "data_processed/methylation_matrix/GSE81233_age_dmr_stream_full"
TABLES = ROOT / "results/tables"
FIGS = ROOT / "results/figures"
TABLES.mkdir(parents=True, exist_ok=True)
FIGS.mkdir(parents=True, exist_ok=True)

full = pd.read_csv(TABLES / "GSE81233_full_relevant_cmet_manifest.tsv", sep="\t")
full["sample_id"] = full["filename"].str.split("_").str[0]

sample_dir = OUTDIR / "sample_dmr"
completed = {p.name.replace(".age_dmr.tsv", "") for p in sample_dir.glob("*.age_dmr.tsv")}

valid = full[full["sample_id"].isin(completed)].copy()
missing = full[~full["sample_id"].isin(completed)].copy()

valid.to_csv(TABLES / "GSE81233_valid_cmet_manifest_204.tsv", sep="\t", index=False)

exclusion = missing.copy()
if len(exclusion):
    exclusion["exclusion_reason"] = "gzip_integrity_failed_after_clean_aria2_and_curl_download"
exclusion.to_csv(TABLES / "GSE81233_excluded_corrupt_or_missing_files.tsv", sep="\t", index=False)

counts = valid.groupby("stage").size().reset_index(name="n_valid_samples")
counts.to_csv(TABLES / "GSE81233_valid204_stage_sample_counts.tsv", sep="\t", index=False)

print("completed_n:", len(completed))
print("valid_manifest_n:", len(valid))
print("missing_n:", len(missing))
print("stage counts:")
print(counts.to_string(index=False))
print("\nmissing/excluded:")
print(exclusion[["stage", "sample_id", "filename", "size_mb", "exclusion_reason"]].to_string(index=False) if len(exclusion) else "none")

spec = importlib.util.spec_from_file_location("stream_mod", ROOT / "scripts/09_stream_gse81233_age_dmr_aggregation.py")
mod = importlib.util.module_from_spec(spec)
spec.loader.exec_module(mod)

metrics = mod.combine(OUTDIR, valid)

metrics = pd.read_csv(TABLES / "GSE81233_full_stage_epi_age_metrics.tsv", sep="\t")
metrics.to_csv(TABLES / "GSE81233_valid204_stage_epi_age_metrics.tsv", sep="\t", index=False)

gz = metrics.dropna(subset=["s_epi_age"]).sort_values("s_epi_age").head(1)
gz.to_csv(TABLES / "GSE81233_valid204_ground_zero_summary.tsv", sep="\t", index=False)

x = range(len(metrics))
labels = metrics["stage"].astype(str).tolist()

plt.figure(figsize=(9, 4.8))
plt.plot(x, metrics["s_epi_age"], marker="o", linewidth=2)
plt.xticks(x, labels, rotation=35, ha="right")
plt.ylabel("S_epi-age")
plt.xlabel("Developmental stage")
plt.title("Age-associated epigenetic entropy across human preimplantation stages")
plt.tight_layout()
plt.savefig(FIGS / "GSE81233_valid204_s_epi_age_by_stage.png", dpi=300)
plt.savefig(FIGS / "GSE81233_valid204_s_epi_age_by_stage.pdf")

plt.figure(figsize=(9, 4.8))
plt.plot(x, metrics["s_epi"], marker="o", linewidth=2, label="S_epi")
plt.plot(x, metrics["s_epi_age"], marker="o", linewidth=2, label="S_epi-age")
plt.xticks(x, labels, rotation=35, ha="right")
plt.ylabel("Entropy")
plt.xlabel("Developmental stage")
plt.legend(frameon=False)
plt.tight_layout()
plt.savefig(FIGS / "GSE81233_valid204_epi_vs_epi_age_by_stage.png", dpi=300)
plt.savefig(FIGS / "GSE81233_valid204_epi_vs_epi_age_by_stage.pdf")

print("\nfinal metrics:")
print(metrics.to_string(index=False))
print("\nground zero:")
print(gz.to_string(index=False))
print("\nfigures written:")
print(FIGS / "GSE81233_valid204_s_epi_age_by_stage.png")
print(FIGS / "GSE81233_valid204_epi_vs_epi_age_by_stage.png")
```
