## Supplemental code for "An entropy-based diagnostic framework for characterizing methylation state dynamics during preimplantation development": 11_stats_and_reset_valid204.py

```
from pathlib import Path
import numpy as np
import pandas as pd
import matplotlib.pyplot as plt
import seaborn as sns
from scipy import stats
from statsmodels.stats.multitest import multipletests

ROOT = Path("/root/autodl-tmp/TRO_Project")
OUTDIR = ROOT / "data_processed/methylation_matrix/GSE81233_age_dmr_stream_full"
TABLES = ROOT / "results/tables"
FIGS = ROOT / "results/figures"
TABLES.mkdir(parents=True, exist_ok=True)
FIGS.mkdir(parents=True, exist_ok=True)

STAGE_ORDER = ["MII oocyte", "zygote/PN", "2-cell", "4-cell", "8-cell", "morula", "blastocyst", "ICM", "TE"]

def binary_entropy(p, eps=1e-6):
    p = np.clip(np.asarray(p, dtype=float), eps, 1 - eps)
    return -(p * np.log(p) + (1 - p) * np.log(1 - p))

def cliffs_delta(x, y):
    x = np.asarray(x)
    y = np.asarray(y)
    gt = sum((xi > y).sum() for xi in x)
    lt = sum((xi < y).sum() for xi in x)
    return (gt - lt) / (len(x) * len(y))

rows = []
sample_dir = OUTDIR / "sample_dmr"
for f in sorted(sample_dir.glob("*.age_dmr.tsv")):
    df = pd.read_csv(f, sep="\t")
    sid = df["sample_id"].iloc[0]
    stage = df["stage"].iloc[0]
    valid = df[df["total_reads"] > 0].copy()
    if len(valid) == 0:
        continue
    p = valid["beta"].astype(float).to_numpy()
    h = binary_entropy(p)
    w = valid["age_weight_5yr"].astype(float).to_numpy()
    rows.append({
        "sample_id": sid,
        "stage": stage,
        "n_regions_covered": len(valid),
        "s_epi_sample": float(np.mean(h)),
        "s_epi_age_sample": float(np.sum(np.abs(w) * h) / np.sum(np.abs(w))),
        "age_projection_sample": float(np.sum(w * p) / np.sum(np.abs(w))),
    })

sample_metrics = pd.DataFrame(rows)
sample_metrics["stage"] = pd.Categorical(sample_metrics["stage"], categories=STAGE_ORDER, ordered=True)
sample_metrics = sample_metrics.sort_values(["stage", "sample_id"])
sample_metrics.to_csv(TABLES / "GSE81233_valid204_sample_level_entropy_metrics.tsv", sep="\t", index=False)

summary = sample_metrics.groupby("stage", observed=True).agg(
    n_samples=("sample_id", "count"),
    s_epi_age_mean=("s_epi_age_sample", "mean"),
    s_epi_age_median=("s_epi_age_sample", "median"),
    s_epi_age_sd=("s_epi_age_sample", "std"),
    s_epi_age_sem=("s_epi_age_sample", lambda x: x.std(ddof=1) / np.sqrt(len(x))),
    s_epi_mean=("s_epi_sample", "mean"),
    age_projection_mean=("age_projection_sample", "mean"),
    covered_regions_median=("n_regions_covered", "median"),
).reset_index()
summary.to_csv(TABLES / "GSE81233_valid204_sample_level_stage_summary.tsv", sep="\t", index=False)

groups = [g["s_epi_age_sample"].dropna().to_numpy() for _, g in sample_metrics.groupby("stage", observed=True)]
kw = stats.kruskal(*groups)
pd.DataFrame([{"test": "Kruskal-Wallis", "metric": "s_epi_age_sample", "statistic": kw.statistic, "p_value": kw.pvalue}]).to_csv(
    TABLES / "GSE81233_valid204_kruskal_s_epi_age.tsv", sep="\t", index=False
)

pairs = []
for a, b in zip(STAGE_ORDER[:-1], STAGE_ORDER[1:]):
    x = sample_metrics.loc[sample_metrics["stage"] == a, "s_epi_age_sample"].dropna()
    y = sample_metrics.loc[sample_metrics["stage"] == b, "s_epi_age_sample"].dropna()
    if len(x) and len(y):
        u = stats.mannwhitneyu(x, y, alternative="two-sided")
        pairs.append({"comparison": f"{a} vs {b}", "stage_a": a, "stage_b": b, "n_a": len(x), "n_b": len(y),
                      "u_stat": u.statistic, "p_value": u.pvalue, "cliffs_delta_a_minus_b": cliffs_delta(x, y)})
pair_df = pd.DataFrame(pairs)
pair_df["p_adj_BH"] = multipletests(pair_df["p_value"], method="fdr_bh")[1]
pair_df.to_csv(TABLES / "GSE81233_valid204_adjacent_stage_mannwhitney.tsv", sep="\t", index=False)

agg = pd.read_csv(TABLES / "GSE81233_valid204_stage_epi_age_metrics.tsv", sep="\t")
mii = float(agg.loc[agg["stage"] == "MII oocyte", "s_epi_age"].iloc[0])
ground_stage = agg.sort_values("s_epi_age").iloc[0]["stage"]
ground = float(agg["s_epi_age"].min())
agg["relative_reset_score_internal"] = (mii - agg["s_epi_age"]) / (mii - ground)
agg.to_csv(TABLES / "GSE81233_valid204_internal_reset_score.tsv", sep="\t", index=False)

rng = np.random.default_rng(42)
boot_rows = []
winners = []
for b in range(2000):
    vals = {}
    for stage in STAGE_ORDER:
        x = sample_metrics.loc[sample_metrics["stage"] == stage, "s_epi_age_sample"].dropna().to_numpy()
        if len(x) == 0:
            continue
        vals[stage] = float(rng.choice(x, size=len(x), replace=True).mean())
        boot_rows.append({"boot": b, "stage": stage, "s_epi_age_mean": vals[stage]})
    winners.append(min(vals, key=vals.get))

boot = pd.DataFrame(boot_rows)
ci = boot.groupby("stage").agg(
    boot_mean=("s_epi_age_mean", "mean"),
    ci_low=("s_epi_age_mean", lambda x: np.quantile(x, 0.025)),
    ci_high=("s_epi_age_mean", lambda x: np.quantile(x, 0.975)),
).reset_index()
ci.to_csv(TABLES / "GSE81233_valid204_bootstrap_stage_ci.tsv", sep="\t", index=False)

freq = pd.Series(winners).value_counts().rename_axis("stage").reset_index(name="n_min")
freq["frequency"] = freq["n_min"] / 2000
freq.to_csv(TABLES / "GSE81233_valid204_bootstrap_ground_zero_frequency.tsv", sep="\t", index=False)

sns.set(style="whitegrid", context="talk")

plt.figure(figsize=(10, 5))
sns.boxplot(data=sample_metrics, x="stage", y="s_epi_age_sample", order=STAGE_ORDER, color="#d9e8f5", fliersize=2)
sns.stripplot(data=sample_metrics, x="stage", y="s_epi_age_sample", order=STAGE_ORDER, color="black", alpha=0.45, size=3)
plt.xticks(rotation=35, ha="right")
plt.xlabel("Developmental stage")
plt.ylabel("Sample-level S_epi-age")
plt.tight_layout()
plt.savefig(FIGS / "GSE81233_valid204_sample_level_s_epi_age_boxplot.png", dpi=300)
plt.savefig(FIGS / "GSE81233_valid204_sample_level_s_epi_age_boxplot.pdf")

plt.figure(figsize=(10, 5))
x = np.arange(len(agg))
plt.plot(x, agg["relative_reset_score_internal"], marker="o", linewidth=2)
plt.axhline(1, linestyle="--", color="gray", linewidth=1)
plt.xticks(x, agg["stage"], rotation=35, ha="right")
plt.xlabel("Developmental stage")
plt.ylabel("Internal relative reset score")
plt.tight_layout()
plt.savefig(FIGS / "GSE81233_valid204_internal_reset_score.png", dpi=300)
plt.savefig(FIGS / "GSE81233_valid204_internal_reset_score.pdf")

print("sample metrics:", TABLES / "GSE81233_valid204_sample_level_entropy_metrics.tsv")
print("Kruskal:", kw)
print("ground_stage:", ground_stage)
print("bootstrap ground-zero frequency:")
print(freq.to_string(index=False))
print("internal reset score:")
print(agg[["stage", "s_epi_age", "relative_reset_score_internal"]].to_string(index=False))
```
