## Supplemental code for "An entropy-based diagnostic framework for characterizing methylation state dynamics during preimplantation development": 12_experiment1b_dna_robustness.py

```
from pathlib import Path
import argparse
import json

import numpy as np
import pandas as pd

STAGE_ORDER = [
    "MII oocyte",
    "zygote/PN",
    "2-cell",
    "4-cell",
    "8-cell",
    "morula",
    "blastocyst",
    "ICM",
    "TE",
]


def project_root():
    here = Path(__file__).resolve()
    default_root = here.parents[1]
    server_root = Path("/root/autodl-tmp/TRO_Project")
    if server_root.exists() and str(here).startswith("/root/"):
        return server_root
    return default_root


ROOT = project_root()
TABLES = ROOT / "results" / "tables" if (ROOT / "results" / "tables").exists() else ROOT / "tables"
FIGS = ROOT / "results" / "figures" if (ROOT / "results" / "figures").exists() else ROOT / "figures"
MATRIX_DIR = ROOT / "data_processed" / "methylation_matrix" / "GSE81233_age_dmr_stream_full"
SAMPLE_DMR_DIR = MATRIX_DIR / "sample_dmr"


def binary_entropy(p, eps=1e-6):
    p = np.clip(np.asarray(p, dtype=float), eps, 1 - eps)
    return -(p * np.log(p) + (1 - p) * np.log(1 - p))


def load_sample_dmr(sample_dir):
    files = sorted(Path(sample_dir).glob("*.age_dmr.tsv"))
    if not files:
        raise FileNotFoundError(
            f"No sample DMR files found in {sample_dir}. "
            "Pull or generate data_processed/methylation_matrix/"
            "GSE81233_age_dmr_stream_full/sample_dmr first."
        )
    frames = []
    for f in files:
        usecols = [
            "sample_id",
            "stage",
            "cluster_name",
            "age_weight_5yr",
            "met_reads",
            "total_reads",
            "beta",
        ]
        frames.append(pd.read_csv(f, sep="\t", usecols=usecols))
    df = pd.concat(frames, ignore_index=True)
    df = df[df["stage"].isin(STAGE_ORDER)].copy()
    df["stage"] = pd.Categorical(df["stage"], categories=STAGE_ORDER, ordered=True)
    return df


def aggregate_stage_metrics(long_df, min_sample_frac=0.30, region_set=None, weight_map=None):
    df = long_df
    if region_set is not None:
        df = df[df["cluster_name"].isin(region_set)].copy()
    if weight_map is not None:
        df = df.copy()
        df["age_weight_5yr"] = df["cluster_name"].map(weight_map)

    rows = []
    valid_regions_by_stage = {}
    for stage in STAGE_ORDER:
        sub = df[df["stage"] == stage]
        n_samples = sub["sample_id"].nunique()
        if n_samples == 0:
            continue
        g = sub.groupby("cluster_name", as_index=False).agg(
            met_reads=("met_reads", "sum"),
            total_reads=("total_reads", "sum"),
            n_samples_covered=("total_reads", lambda x: int((x > 0).sum())),
            age_weight_5yr=("age_weight_5yr", "first"),
        )
        min_cov = max(1, int(np.ceil(min_sample_frac * n_samples)))
        valid = g[(g["total_reads"] > 0) & (g["n_samples_covered"] >= min_cov)].copy()
        valid_regions_by_stage[stage] = set(valid["cluster_name"])
        if len(valid) == 0:
            rows.append(
                {
                    "stage": stage,
                    "n_samples": n_samples,
                    "n_regions_valid": 0,
                    "min_samples_required": min_cov,
                    "s_epi": np.nan,
                    "s_epi_age": np.nan,
                    "age_projection": np.nan,
                }
            )
            continue
        p = valid["met_reads"] / valid["total_reads"]
        h = binary_entropy(p)
        signed_w = valid["age_weight_5yr"].astype(float).to_numpy()
        abs_w = np.abs(signed_w)
        rows.append(
            {
                "stage": stage,
                "n_samples": n_samples,
                "n_regions_valid": len(valid),
                "min_samples_required": min_cov,
                "s_epi": float(np.mean(h)),
                "s_epi_age": float(np.sum(abs_w * h) / np.sum(abs_w)),
                "age_projection": float(np.sum(signed_w * p) / np.sum(abs_w)),
            }
        )

    metrics = pd.DataFrame(rows)
    metrics["stage"] = pd.Categorical(metrics["stage"], categories=STAGE_ORDER, ordered=True)
    metrics = metrics.sort_values("stage").reset_index(drop=True)
    return metrics, valid_regions_by_stage


def ground_stage(metrics):
    ok = metrics.dropna(subset=["s_epi_age"])
    if ok.empty:
        return None
    return str(ok.sort_values("s_epi_age").iloc[0]["stage"])


def sample_level_metrics(long_df, min_regions=1, weight_map=None):
    df = long_df
    if weight_map is not None:
        df = df.copy()
        df["age_weight_5yr"] = df["cluster_name"].map(weight_map)
    rows = []
    for (sample_id, stage), sub in df.groupby(["sample_id", "stage"], observed=True):
        valid = sub[sub["total_reads"] > 0].copy()
        if len(valid) < min_regions:
            continue
        p = valid["beta"].astype(float).to_numpy()
        h = binary_entropy(p)
        w = valid["age_weight_5yr"].astype(float).to_numpy()
        rows.append(
            {
                "sample_id": sample_id,
                "stage": str(stage),
                "n_regions_covered": len(valid),
                "s_epi_age_sample": float(np.sum(np.abs(w) * h) / np.sum(np.abs(w))),
            }
        )
    out = pd.DataFrame(rows)
    out["stage"] = pd.Categorical(out["stage"], categories=STAGE_ORDER, ordered=True)
    return out.sort_values(["stage", "sample_id"]).reset_index(drop=True)


def bootstrap_winner_frequency(sample_metrics, n_boot=2000, balanced_n=None, seed=42):
    rng = np.random.default_rng(seed)
    winners = []
    for b in range(n_boot):
        vals = {}
        for stage in STAGE_ORDER:
            x = sample_metrics.loc[sample_metrics["stage"] == stage, "s_epi_age_sample"].dropna().to_numpy()
            if len(x) == 0:
                continue
            n = len(x) if balanced_n is None else min(int(balanced_n), len(x))
            vals[stage] = float(rng.choice(x, size=n, replace=True).mean())
        winners.append(min(vals, key=vals.get))
    freq = pd.Series(winners).value_counts().rename_axis("stage").reset_index(name="n_min")
    freq["frequency"] = freq["n_min"] / n_boot
    return freq.sort_values("frequency", ascending=False).reset_index(drop=True)


def shuffled_weight_control(long_df, n_perm=1000, min_sample_frac=0.30, seed=43):
    rng = np.random.default_rng(seed)
    weights = long_df.drop_duplicates("cluster_name").set_index("cluster_name")["age_weight_5yr"].astype(float)
    clusters = weights.index.to_numpy()
    wvals = weights.to_numpy()
    winners = []
    morula_vals = []
    for i in range(n_perm):
        shuffled = dict(zip(clusters, rng.permutation(wvals)))
        metrics, _ = aggregate_stage_metrics(long_df, min_sample_frac=min_sample_frac, weight_map=shuffled)
        winners.append(ground_stage(metrics))
        morula_vals.append(float(metrics.loc[metrics["stage"] == "morula", "s_epi_age"].iloc[0]))
    freq = pd.Series(winners).value_counts().rename_axis("stage").reset_index(name="n_min")
    freq["frequency"] = freq["n_min"] / n_perm
    return freq.sort_values("frequency", ascending=False).reset_index(drop=True), morula_vals


def random_subset_control(long_df, subset_size, n_perm=1000, min_sample_frac=0.30, seed=44):
    rng = np.random.default_rng(seed)
    clusters = long_df["cluster_name"].drop_duplicates().to_numpy()
    subset_size = min(int(subset_size), len(clusters))
    winners = []
    for i in range(n_perm):
        region_set = set(rng.choice(clusters, size=subset_size, replace=False))
        metrics, _ = aggregate_stage_metrics(long_df, min_sample_frac=min_sample_frac, region_set=region_set)
        winners.append(ground_stage(metrics))
    freq = pd.Series(winners).value_counts().rename_axis("stage").reset_index(name="n_min")
    freq["frequency"] = freq["n_min"] / n_perm
    return freq.sort_values("frequency", ascending=False).reset_index(drop=True)


def plot_summary(summary):
    try:
        import matplotlib.pyplot as plt
    except ImportError:
        print("matplotlib is not installed; skipping summary figure.")
        return

    FIGS.mkdir(parents=True, exist_ok=True)
    plot_df = summary.copy()
    plot_df["morula_frequency"] = pd.to_numeric(plot_df["morula_frequency"], errors="coerce")
    plt.figure(figsize=(10, 4.8))
    colors = ["#2c7fb8" if x == "pass" else "#fdae61" for x in plot_df["conclusion"]]
    plt.bar(plot_df["test"], plot_df["morula_frequency"], color=colors)
    plt.axhline(0.5, color="gray", linestyle="--", linewidth=1)
    plt.xticks(rotation=35, ha="right")
    plt.ylim(0, 1.05)
    plt.ylabel("Morula ground-zero frequency")
    plt.xlabel("")
    plt.tight_layout()
    plt.savefig(FIGS / "Experiment1B_DNA_robustness_summary.png", dpi=300)
    plt.savefig(FIGS / "Experiment1B_DNA_robustness_summary.pdf")


def main():
    ap = argparse.ArgumentParser()
    ap.add_argument("--sample-dir", default=str(SAMPLE_DMR_DIR))
    ap.add_argument("--n-boot", type=int, default=2000)
    ap.add_argument("--n-perm", type=int, default=1000)
    ap.add_argument("--thresholds", default="0.1,0.2,0.3,0.4,0.5")
    args = ap.parse_args()

    TABLES.mkdir(parents=True, exist_ok=True)
    long_df = load_sample_dmr(args.sample_dir)
    long_df.to_csv(TABLES / "Experiment1B_all_sample_age_dmr_long.tsv.gz", sep="\t", index=False, compression="gzip")

    summary_rows = []

    original, valid_by_stage = aggregate_stage_metrics(long_df, min_sample_frac=0.30)
    original.to_csv(TABLES / "Experiment1B_original_valid204_recomputed.tsv", sep="\t", index=False)
    original_sample = sample_level_metrics(long_df)
    original_freq = bootstrap_winner_frequency(original_sample, n_boot=args.n_boot, seed=42)
    original_freq.to_csv(TABLES / "Experiment1B_original_bootstrap_ground_zero_frequency.tsv", sep="\t", index=False)
    summary_rows.append(
        {
            "test": "original valid204",
            "ground_zero_stage": ground_stage(original),
            "morula_frequency": float(original_freq.loc[original_freq["stage"] == "morula", "frequency"].iloc[0]),
            "p_value": np.nan,
            "n_regions": int(original["n_regions_valid"].min()),
            "conclusion": "pass" if ground_stage(original) == "morula" else "check",
        }
    )

    common_regions = set.intersection(*[valid_by_stage[s] for s in STAGE_ORDER if s in valid_by_stage])
    common, _ = aggregate_stage_metrics(long_df, min_sample_frac=0.30, region_set=common_regions)
    common.to_csv(TABLES / "Experiment1B_common_DMR_stage_metrics.tsv", sep="\t", index=False)
    common_sample = sample_level_metrics(long_df[long_df["cluster_name"].isin(common_regions)])
    common_freq = bootstrap_winner_frequency(common_sample, n_boot=args.n_boot, seed=45)
    common_freq.to_csv(TABLES / "Experiment1B_common_DMR_bootstrap_ground_zero_frequency.tsv", sep="\t", index=False)
    summary_rows.append(
        {
            "test": "common DMR",
            "ground_zero_stage": ground_stage(common),
            "morula_frequency": float(common_freq.loc[common_freq["stage"] == "morula", "frequency"].iloc[0])
            if (common_freq["stage"] == "morula").any()
            else 0.0,
            "p_value": np.nan,
            "n_regions": len(common_regions),
            "conclusion": "pass" if ground_stage(common) == "morula" else "check",
        }
    )

    threshold_rows = []
    for t in [float(x) for x in args.thresholds.split(",") if x.strip()]:
        metrics, _ = aggregate_stage_metrics(long_df, min_sample_frac=t)
        metrics.insert(0, "min_sample_frac", t)
        threshold_rows.append(metrics)
        summary_rows.append(
            {
                "test": f"min_frac {t:g}",
                "ground_zero_stage": ground_stage(metrics),
                "morula_frequency": np.nan,
                "p_value": np.nan,
                "n_regions": int(metrics["n_regions_valid"].min()),
                "conclusion": "pass" if ground_stage(metrics) == "morula" else "check",
            }
        )
    pd.concat(threshold_rows, ignore_index=True).to_csv(
        TABLES / "Experiment1B_coverage_threshold_sensitivity.tsv", sep="\t", index=False
    )

    shuffled_freq, shuffled_morula = shuffled_weight_control(long_df, n_perm=args.n_perm, min_sample_frac=0.30)
    shuffled_freq.to_csv(TABLES / "Experiment1B_shuffled_weight_ground_zero_frequency.tsv", sep="\t", index=False)
    shuffled_morula_frequency = (
        float(shuffled_freq.loc[shuffled_freq["stage"] == "morula", "frequency"].iloc[0])
        if (shuffled_freq["stage"] == "morula").any()
        else 0.0
    )
    summary_rows.append(
        {
            "test": "shuffled weights",
            "ground_zero_stage": str(shuffled_freq.iloc[0]["stage"]),
            "morula_frequency": shuffled_morula_frequency,
            "p_value": np.nan,
            "n_regions": int(original["n_regions_valid"].min()),
            "conclusion": "control",
        }
    )

    subset_size = max(5, len(common_regions)) if common_regions else int(original["n_regions_valid"].min())
    subset_freq = random_subset_control(long_df, subset_size=subset_size, n_perm=args.n_perm, min_sample_frac=0.30)
    subset_freq.to_csv(TABLES / "Experiment1B_random_age_DMR_subset_ground_zero_frequency.tsv", sep="\t", index=False)
    subset_morula_frequency = (
        float(subset_freq.loc[subset_freq["stage"] == "morula", "frequency"].iloc[0])
        if (subset_freq["stage"] == "morula").any()
        else 0.0
    )
    summary_rows.append(
        {
            "test": "random age-DMR subset",
            "ground_zero_stage": str(subset_freq.iloc[0]["stage"]),
            "morula_frequency": subset_morula_frequency,
            "p_value": np.nan,
            "n_regions": subset_size,
            "conclusion": "control",
        }
    )

    for n in [5, 8]:
        freq = bootstrap_winner_frequency(original_sample, n_boot=args.n_boot, balanced_n=n, seed=100 + n)
        freq.to_csv(TABLES / f"Experiment1B_balanced_bootstrap_n{n}_ground_zero_frequency.tsv", sep="\t", index=False)
        morula_frequency = (
            float(freq.loc[freq["stage"] == "morula", "frequency"].iloc[0])
            if (freq["stage"] == "morula").any()
            else 0.0
        )
        summary_rows.append(
            {
                "test": f"balanced bootstrap n={n}",
                "ground_zero_stage": str(freq.iloc[0]["stage"]),
                "morula_frequency": morula_frequency,
                "p_value": np.nan,
                "n_regions": int(original["n_regions_valid"].min()),
                "conclusion": "pass" if str(freq.iloc[0]["stage"]) == "morula" else "check",
            }
        )

    summary = pd.DataFrame(summary_rows)
    summary.to_csv(TABLES / "Experiment1B_DNA_robustness_summary.tsv", sep="\t", index=False)
    plot_summary(summary)

    with open(TABLES / "Experiment1B_DNA_robustness_run_info.json", "w", encoding="utf-8") as fh:
        json.dump(
            {
                "root": str(ROOT),
                "sample_dir": str(args.sample_dir),
                "n_samples": int(long_df["sample_id"].nunique()),
                "n_regions_total": int(long_df["cluster_name"].nunique()),
                "n_boot": args.n_boot,
                "n_perm": args.n_perm,
                "note": "random age-DMR subset is an internal resampling control, not a non-age genomic random-region control.",
            },
            fh,
            indent=2,
        )

    print(summary.to_string(index=False))
    print("Wrote:", TABLES / "Experiment1B_DNA_robustness_summary.tsv")
    print("Wrote:", FIGS / "Experiment1B_DNA_robustness_summary.png")


if __name__ == "__main__":
    main()
```
