## Supplemental code for "An entropy-based diagnostic framework for characterizing methylation state dynamics during preimplantation development": 13_experiment2a_gse36552_rna_entropy_potency.py

```
from pathlib import Path
import argparse
import gzip
import re
import urllib.request

import numpy as np
import pandas as pd


STAGE_ORDER = [
    "MII oocyte",
    "zygote/PN",
    "2-cell",
    "4-cell",
    "8-cell",
    "morula",
    "blastocyst",
    "ICM",
    "TE",
]

RNA_STAGE_ORDER = [
    "oocyte",
    "zygote",
    "2-cell",
    "4-cell",
    "8-cell",
    "morula",
    "blastocyst",
]

DNA_TO_RNA_STAGE = {
    "MII oocyte": "oocyte",
    "zygote/PN": "zygote",
    "2-cell": "2-cell",
    "4-cell": "4-cell",
    "8-cell": "8-cell",
    "morula": "morula",
    "blastocyst": "blastocyst",
}

POTENCY_MARKERS = [
    "POU5F1",
    "NANOG",
    "SOX2",
    "KLF4",
    "DPPA3",
    "ZSCAN4",
    "KLF17",
    "TFAP2C",
    "GDF3",
]

SUPP_URL = (
    "https://static-content.springer.com/esm/art%3A10.1038%2Fnsmb.2660/"
    "MediaObjects/41594_2013_BFnsmb2660_MOESM31_ESM.xlsx"
)
SERIES_MATRIX_URL = "https://ftp.ncbi.nlm.nih.gov/geo/series/GSE36nnn/GSE36552/matrix/GSE36552_series_matrix.txt.gz"


def project_root():
    here = Path(__file__).resolve()
    server_root = Path("/root/autodl-tmp/TRO_Project")
    if server_root.exists() and str(here).startswith("/root/"):
        return server_root
    return here.parents[1]


ROOT = project_root()
RAW = ROOT / "data_raw" / "GSE36552_rna"
TABLES = ROOT / "results" / "tables" if (ROOT / "results" / "tables").exists() else ROOT / "tables"
FIGS = ROOT / "results" / "figures" if (ROOT / "results" / "figures").exists() else ROOT / "figures"


def download(url, path):
    path.parent.mkdir(parents=True, exist_ok=True)
    if path.exists() and path.stat().st_size > 1000:
        return path
    print(f"Downloading {url}")
    urllib.request.urlretrieve(url, path)
    return path


def minmax(x):
    x = np.asarray(x, dtype=float)
    lo = np.nanmin(x)
    hi = np.nanmax(x)
    if not np.isfinite(lo) or not np.isfinite(hi) or hi == lo:
        return np.zeros_like(x, dtype=float)
    return (x - lo) / (hi - lo)


def shannon_entropy(values):
    x = np.array(values, dtype=float, copy=True)
    x[~np.isfinite(x)] = 0.0
    x[x < 0] = 0.0
    total = x.sum()
    if total <= 0:
        return np.nan
    p = x / total
    p = p[p > 0]
    return float(-(p * np.log(p)).sum())


def parse_geo_series_matrix(path):
    rows = []
    with gzip.open(path, "rt", encoding="utf-8", errors="replace") as fh:
        sample_ids = None
        titles = None
        sources = None
        characteristics = []
        for line in fh:
            if line.startswith("!Sample_geo_accession"):
                sample_ids = [x.strip('"') for x in line.rstrip("\n").split("\t")[1:]]
            elif line.startswith("!Sample_title"):
                titles = [x.strip('"') for x in line.rstrip("\n").split("\t")[1:]]
            elif line.startswith("!Sample_source_name_ch1"):
                sources = [x.strip('"') for x in line.rstrip("\n").split("\t")[1:]]
            elif line.startswith("!Sample_characteristics_ch1"):
                characteristics.append([x.strip('"') for x in line.rstrip("\n").split("\t")[1:]])
    if sample_ids is None:
        return pd.DataFrame()
    for i, sid in enumerate(sample_ids):
        row = {
            "geo_accession": sid,
            "title": titles[i] if titles and i < len(titles) else "",
            "source_name": sources[i] if sources and i < len(sources) else "",
        }
        for j, vals in enumerate(characteristics):
            row[f"characteristics_{j+1}"] = vals[i] if i < len(vals) else ""
        rows.append(row)
    return pd.DataFrame(rows)


def infer_stage_from_text(text):
    t = str(text).lower()
    t = re.sub(r"[_\-]+", " ", t)
    if "morula" in t:
        return "morula"
    if "blastocyst" in t or "blast" in t:
        return "blastocyst"
    if "zygote" in t or "pronuclear" in t or "pn" in t:
        return "zygote"
    if "oocyte" in t or "mii" in t or re.search(r"\booc", t):
        return "oocyte"
    if re.search(r"\b2\s*cell\b", t) or re.search(r"\b2c\b", t):
        return "2-cell"
    if re.search(r"\b4\s*cell\b", t) or re.search(r"\b4c\b", t):
        return "4-cell"
    if re.search(r"\b8\s*cell\b", t) or re.search(r"\b8c\b", t):
        return "8-cell"
    return None


def read_expression_xlsx(path):
    xls = pd.ExcelFile(path)
    best = None
    best_score = -1
    for sheet in xls.sheet_names:
        preview = pd.read_excel(path, sheet_name=sheet, nrows=8, header=None)
        text = " ".join(preview.fillna("").astype(str).values.ravel()).lower()
        score = text.count("rpkm") + text.count("refseq") + text.count("gene")
        if score > best_score:
            best = sheet
            best_score = score

    raw = pd.read_excel(path, sheet_name=best, header=None)
    header_idx = None
    for i in range(min(20, len(raw))):
        vals = raw.iloc[i].fillna("").astype(str).str.lower().tolist()
        if any("gene" in v or "symbol" in v for v in vals[:8]) and sum(v != "" for v in vals) > 20:
            header_idx = i
            break
    if header_idx is None:
        header_idx = 0

    df = pd.read_excel(path, sheet_name=best, header=header_idx)
    df = df.dropna(axis=1, how="all").dropna(axis=0, how="all")

    gene_col = None
    for c in df.columns:
        lc = str(c).lower()
        if "gene" in lc and ("symbol" in lc or lc.strip() == "gene"):
            gene_col = c
            break
    if gene_col is None:
        for c in df.columns[:10]:
            vals = df[c].dropna().astype(str).head(100)
            if vals.str.match(r"^[A-Za-z0-9_.-]+$").mean() > 0.8:
                gene_col = c
                break
    if gene_col is None:
        raise RuntimeError(f"Could not identify gene column in sheet {best}")

    sample_cols = []
    for c in df.columns:
        if c == gene_col:
            continue
        vals = pd.to_numeric(df[c], errors="coerce")
        if vals.notna().sum() > 1000:
            sample_cols.append(c)

    sample_cols = list(dict.fromkeys(sample_cols))
    expr = df[[gene_col] + sample_cols].copy()
    expr = expr.rename(columns={gene_col: "gene_symbol"})
    expr["gene_symbol"] = expr["gene_symbol"].astype(str).str.strip()
    expr = expr[expr["gene_symbol"].notna() & (expr["gene_symbol"] != "") & (expr["gene_symbol"].str.lower() != "nan")]
    for c in sample_cols:
        expr[c] = pd.to_numeric(expr[c], errors="coerce").fillna(0.0)

    expr = expr.groupby("gene_symbol", as_index=False)[sample_cols].max()
    sample_meta = pd.DataFrame({"sample_name": [str(c) for c in sample_cols]})
    sample_meta["stage"] = sample_meta["sample_name"].map(infer_stage_from_text)
    return expr, sample_meta, best


def add_geo_stage_if_needed(sample_meta, geo_meta):
    if geo_meta.empty:
        return sample_meta
    geo_meta = geo_meta.copy()
    geo_meta["geo_text"] = geo_meta.astype(str).agg(" ".join, axis=1)
    geo_meta["geo_stage"] = geo_meta["geo_text"].map(infer_stage_from_text)
    stages = geo_meta["geo_stage"].dropna().tolist()
    if sample_meta["stage"].notna().sum() == 0 and len(stages) == len(sample_meta):
        sample_meta = sample_meta.copy()
        sample_meta["stage"] = stages
    return sample_meta


def compute_metrics(expr, sample_meta):
    genes = expr["gene_symbol"].astype(str).str.upper()
    matrix = expr.drop(columns=["gene_symbol"])
    sample_cols = matrix.columns.tolist()
    rows = []

    detected = (matrix > 0).sum(axis=0).astype(float)
    marker_present = [m for m in POTENCY_MARKERS if m in set(genes)]
    marker_expr = pd.DataFrame(index=matrix.columns)
    for marker in marker_present:
        idx = np.where(genes == marker)[0]
        vals = matrix.iloc[idx].mean(axis=0)
        marker_expr[marker] = np.log1p(vals)
    if len(marker_expr.columns):
        marker_score_raw = marker_expr.mean(axis=1)
    else:
        marker_score_raw = pd.Series(np.nan, index=matrix.columns)

    detected_score = pd.Series(minmax(detected.values), index=matrix.columns)
    marker_score = pd.Series(minmax(marker_score_raw.values), index=matrix.columns)
    potency = 0.5 * detected_score + 0.5 * marker_score

    for sample in sample_cols:
        stage = sample_meta.loc[sample_meta["sample_name"] == str(sample), "stage"]
        stage = stage.iloc[0] if len(stage) else infer_stage_from_text(sample)
        rows.append(
            {
                "sample_name": str(sample),
                "stage": stage,
                "s_rna": shannon_entropy(matrix[sample].values),
                "detected_genes": int(detected.loc[sample]),
                "detected_gene_score": float(detected_score.loc[sample]),
                "marker_score": float(marker_score.loc[sample]) if np.isfinite(marker_score.loc[sample]) else np.nan,
                "potency_score": float(potency.loc[sample]) if np.isfinite(potency.loc[sample]) else np.nan,
            }
        )
    out = pd.DataFrame(rows)
    out = out[out["stage"].isin(RNA_STAGE_ORDER)].copy()
    out["stage"] = pd.Categorical(out["stage"], categories=RNA_STAGE_ORDER, ordered=True)
    return out.sort_values(["stage", "sample_name"]).reset_index(drop=True), marker_present


def summarize_stage(cell_metrics):
    summary = cell_metrics.groupby("stage", observed=True).agg(
        n_cells=("sample_name", "count"),
        s_rna_mean=("s_rna", "mean"),
        s_rna_median=("s_rna", "median"),
        s_rna_sd=("s_rna", "std"),
        detected_genes_mean=("detected_genes", "mean"),
        marker_score_mean=("marker_score", "mean"),
        potency_score_mean=("potency_score", "mean"),
        potency_score_median=("potency_score", "median"),
    ).reset_index()
    summary["stage"] = summary["stage"].astype(str)
    return summary


def merge_dna_rna(rna_stage):
    dna_path = TABLES / "GSE81233_valid204_internal_reset_score.tsv"
    if not dna_path.exists():
        return pd.DataFrame()
    dna = pd.read_csv(dna_path, sep="\t")
    dna = dna[dna["stage"].isin(DNA_TO_RNA_STAGE)].copy()
    dna["rna_stage"] = dna["stage"].map(DNA_TO_RNA_STAGE)
    merged = dna.merge(rna_stage, left_on="rna_stage", right_on="stage", how="left", suffixes=("_dna", "_rna"))
    out = merged.rename(
        columns={
            "stage_dna": "stage",
            "s_epi_age": "S_epi_age",
            "s_epi": "S_epi",
            "relative_reset_score_internal": "ResetScore",
            "s_rna_mean": "S_RNA",
            "potency_score_mean": "PotencyScore",
        }
    )
    keep = ["stage", "rna_stage", "S_epi_age", "S_epi", "ResetScore", "S_RNA", "PotencyScore", "n_cells"]
    return out[[c for c in keep if c in out.columns]]


def make_figures(rna_stage, dual):
    try:
        import matplotlib.pyplot as plt
    except ImportError:
        print("matplotlib unavailable; skipping figures.")
        return
    FIGS.mkdir(parents=True, exist_ok=True)

    x = np.arange(len(rna_stage))
    plt.figure(figsize=(8, 4.5))
    plt.plot(x, rna_stage["s_rna_mean"], marker="o", linewidth=2)
    plt.xticks(x, rna_stage["stage"], rotation=35, ha="right")
    plt.ylabel("S_RNA")
    plt.xlabel("RNA developmental stage")
    plt.tight_layout()
    plt.savefig(FIGS / "GSE36552_RNA_entropy_by_stage.png", dpi=300)
    plt.savefig(FIGS / "GSE36552_RNA_entropy_by_stage.pdf")

    plt.figure(figsize=(8, 4.5))
    plt.plot(x, rna_stage["potency_score_mean"], marker="o", linewidth=2, color="#2c7fb8")
    plt.xticks(x, rna_stage["stage"], rotation=35, ha="right")
    plt.ylabel("Potency proxy")
    plt.xlabel("RNA developmental stage")
    plt.tight_layout()
    plt.savefig(FIGS / "GSE36552_potency_score_by_stage.png", dpi=300)
    plt.savefig(FIGS / "GSE36552_potency_score_by_stage.pdf")

    if not dual.empty:
        plt.figure(figsize=(6, 5))
        plt.scatter(dual["S_epi_age"], dual["PotencyScore"], s=70)
        for _, r in dual.iterrows():
            plt.annotate(str(r["stage"]), (r["S_epi_age"], r["PotencyScore"]), xytext=(4, 4), textcoords="offset points", fontsize=8)
        plt.xlabel("S_epi-age")
        plt.ylabel("Potency proxy")
        plt.tight_layout()
        plt.savefig(FIGS / "dual_entropy_phase_map.png", dpi=300)
        plt.savefig(FIGS / "dual_entropy_phase_map.pdf")


def main():
    ap = argparse.ArgumentParser()
    ap.add_argument("--force-download", action="store_true")
    args = ap.parse_args()

    RAW.mkdir(parents=True, exist_ok=True)
    TABLES.mkdir(parents=True, exist_ok=True)

    supp = RAW / "Yan_2013_NSMB_supplementary_table_1_RPKM.xlsx"
    series = RAW / "GSE36552_series_matrix.txt.gz"
    if args.force_download:
        for p in [supp, series]:
            if p.exists():
                p.unlink()
    download(SUPP_URL, supp)
    download(SERIES_MATRIX_URL, series)

    geo_meta = parse_geo_series_matrix(series)
    geo_meta.to_csv(TABLES / "GSE36552_geo_sample_metadata.tsv", sep="\t", index=False)

    expr, sample_meta, sheet = read_expression_xlsx(supp)
    sample_meta = add_geo_stage_if_needed(sample_meta, geo_meta)
    sample_meta.to_csv(TABLES / "GSE36552_rna_sample_metadata.tsv", sep="\t", index=False)

    cell_metrics, marker_present = compute_metrics(expr, sample_meta)
    stage_summary = summarize_stage(cell_metrics)
    dual = merge_dna_rna(stage_summary)

    cell_metrics.to_csv(TABLES / "GSE36552_RNA_entropy_potency_cell_metrics.tsv", sep="\t", index=False)
    stage_summary.to_csv(TABLES / "GSE36552_RNA_entropy_potency_by_stage.tsv", sep="\t", index=False)
    dual.to_csv(TABLES / "dual_entropy_stage_table.tsv", sep="\t", index=False)
    make_figures(stage_summary, dual)

    run_info = pd.DataFrame(
        [
            {
                "source": "Yan et al. 2013 Nature Structural & Molecular Biology Supplementary Table 1",
                "geo": "GSE36552",
                "xlsx_sheet": sheet,
                "n_genes": expr.shape[0],
                "n_expression_columns": expr.shape[1] - 1,
                "n_cells_with_stage": len(cell_metrics),
                "markers_found": ",".join(marker_present),
            }
        ]
    )
    run_info.to_csv(TABLES / "GSE36552_RNA_entropy_potency_run_info.tsv", sep="\t", index=False)

    print("RNA stage summary:")
    print(stage_summary.to_string(index=False))
    print("\nDual entropy stage table:")
    print(dual.to_string(index=False))
    print("\nMarkers found:", ", ".join(marker_present))
    print("Wrote:", TABLES / "GSE36552_RNA_entropy_potency_by_stage.tsv")
    print("Wrote:", TABLES / "dual_entropy_stage_table.tsv")


if __name__ == "__main__":
    main()
```
