## Supplemental code for "An entropy-based diagnostic framework for characterizing methylation state dynamics during preimplantation development": 14_experiment2b_rna_potency_robustness.py

```
from pathlib import Path
import argparse
import importlib.util
import math

import numpy as np
import pandas as pd


RNA_STAGE_ORDER = ["oocyte", "zygote", "2-cell", "4-cell", "8-cell", "morula", "blastocyst"]
POTENCY_MARKERS = ["POU5F1", "NANOG", "SOX2", "KLF4", "DPPA3", "ZSCAN4", "KLF17", "TFAP2C", "GDF3"]
PAIRWISE = [
    ("morula", "blastocyst"),
    ("morula", "8-cell"),
    ("morula", "4-cell"),
    ("8-cell", "blastocyst"),
    ("2-cell", "morula"),
]


def project_root():
    here = Path(__file__).resolve()
    server_root = Path("/root/autodl-tmp/TRO_Project")
    if server_root.exists() and str(here).startswith("/root/"):
        return server_root
    return here.parents[1]


ROOT = project_root()
TABLES = ROOT / "results" / "tables" if (ROOT / "results" / "tables").exists() else ROOT / "tables"
FIGS = ROOT / "results" / "figures" if (ROOT / "results" / "figures").exists() else ROOT / "figures"
SCRIPT13 = ROOT / "scripts" / "13_experiment2a_gse36552_rna_entropy_potency.py"


def load_exp2a_module():
    spec = importlib.util.spec_from_file_location("exp2a", SCRIPT13)
    mod = importlib.util.module_from_spec(spec)
    spec.loader.exec_module(mod)
    return mod


def bh_adjust(pvals):
    p = np.asarray(pvals, dtype=float)
    out = np.full(len(p), np.nan)
    ok = np.isfinite(p)
    if ok.sum() == 0:
        return out
    idx = np.where(ok)[0]
    order = idx[np.argsort(p[idx])]
    ranked = p[order]
    m = len(ranked)
    adj = ranked * m / np.arange(1, m + 1)
    adj = np.minimum.accumulate(adj[::-1])[::-1]
    out[order] = np.minimum(adj, 1.0)
    return out


def cliffs_delta(x, y):
    x = np.asarray(x, dtype=float)
    y = np.asarray(y, dtype=float)
    gt = sum((xi > y).sum() for xi in x)
    lt = sum((xi < y).sum() for xi in x)
    return (gt - lt) / (len(x) * len(y))


def mannwhitney(x, y):
    try:
        from scipy import stats

        res = stats.mannwhitneyu(x, y, alternative="two-sided")
        return float(res.statistic), float(res.pvalue)
    except Exception:
        x = np.asarray(x, dtype=float)
        y = np.asarray(y, dtype=float)
        x = x[np.isfinite(x)]
        y = y[np.isfinite(y)]
        n1 = len(x)
        n2 = len(y)
        if n1 == 0 or n2 == 0:
            return np.nan, np.nan
        vals = np.concatenate([x, y])
        ranks = pd.Series(vals).rank(method="average").to_numpy()
        r1 = ranks[:n1].sum()
        u1 = r1 - n1 * (n1 + 1) / 2.0
        mean_u = n1 * n2 / 2.0
        _, counts = np.unique(vals, return_counts=True)
        tie_term = np.sum(counts**3 - counts)
        n = n1 + n2
        var_u = n1 * n2 / 12.0 * ((n + 1) - tie_term / (n * (n - 1))) if n > 1 else np.nan
        if not np.isfinite(var_u) or var_u <= 0:
            return float(u1), np.nan
        z = (u1 - mean_u) / np.sqrt(var_u)
        p = math.erfc(abs(z) / np.sqrt(2.0))
        return float(u1), float(p)


def zscore_by_marker(marker_mat):
    z = marker_mat.copy()
    for c in z.columns:
        vals = z[c].astype(float)
        sd = vals.std(ddof=0)
        if not np.isfinite(sd) or sd == 0:
            z[c] = 0.0
        else:
            z[c] = (vals - vals.mean()) / sd
    return z


def make_marker_matrix():
    mod = load_exp2a_module()
    supp = mod.RAW / "Yan_2013_NSMB_supplementary_table_1_RPKM.xlsx"
    series = mod.RAW / "GSE36552_series_matrix.txt.gz"
    if not supp.exists():
        mod.download(mod.SUPP_URL, supp)
    if not series.exists():
        mod.download(mod.SERIES_MATRIX_URL, series)

    geo_meta = mod.parse_geo_series_matrix(series)
    expr, sample_meta, _ = mod.read_expression_xlsx(supp)
    sample_meta = mod.add_geo_stage_if_needed(sample_meta, geo_meta)

    genes_upper = expr["gene_symbol"].astype(str).str.upper()
    matrix = expr.drop(columns=["gene_symbol"])
    rows = []
    for sample in matrix.columns:
        stage = sample_meta.loc[sample_meta["sample_name"] == str(sample), "stage"]
        stage = stage.iloc[0] if len(stage) else mod.infer_stage_from_text(sample)
        if stage not in RNA_STAGE_ORDER:
            continue
        row = {"sample_name": str(sample), "stage": stage}
        for marker in POTENCY_MARKERS:
            idx = np.where(genes_upper == marker)[0]
            row[marker] = float(np.log1p(matrix.iloc[idx][sample].mean())) if len(idx) else np.nan
        rows.append(row)

    marker = pd.DataFrame(rows)
    marker["stage"] = pd.Categorical(marker["stage"], categories=RNA_STAGE_ORDER, ordered=True)
    marker = marker.sort_values(["stage", "sample_name"]).reset_index(drop=True)
    marker_z = marker[["sample_name", "stage"]].join(zscore_by_marker(marker[POTENCY_MARKERS]))
    return marker, marker_z


def bootstrap_ci(cell_metrics, n_boot=2000, seed=2026):
    rng = np.random.default_rng(seed)
    rows = []
    metrics = ["s_rna", "detected_genes", "detected_gene_score", "marker_score", "potency_score"]
    for stage in RNA_STAGE_ORDER:
        sub = cell_metrics[cell_metrics["stage"] == stage]
        if len(sub) == 0:
            continue
        for metric in metrics:
            x = sub[metric].dropna().to_numpy(dtype=float)
            if len(x) == 0:
                continue
            boots = [float(rng.choice(x, size=len(x), replace=True).mean()) for _ in range(n_boot)]
            rows.append(
                {
                    "stage": stage,
                    "metric": metric,
                    "n_cells": len(x),
                    "mean": float(np.mean(x)),
                    "ci_low": float(np.quantile(boots, 0.025)),
                    "ci_high": float(np.quantile(boots, 0.975)),
                }
            )
    return pd.DataFrame(rows)


def pairwise_tests(cell_metrics):
    rows = []
    metrics = ["s_rna", "detected_genes", "detected_gene_score", "marker_score", "potency_score"]
    for metric in metrics:
        for a, b in PAIRWISE:
            x = cell_metrics.loc[cell_metrics["stage"] == a, metric].dropna().to_numpy(dtype=float)
            y = cell_metrics.loc[cell_metrics["stage"] == b, metric].dropna().to_numpy(dtype=float)
            if len(x) == 0 or len(y) == 0:
                continue
            u, p = mannwhitney(x, y)
            rows.append(
                {
                    "metric": metric,
                    "comparison": f"{a} vs {b}",
                    "stage_a": a,
                    "stage_b": b,
                    "n_a": len(x),
                    "n_b": len(y),
                    "mean_a": float(np.mean(x)),
                    "mean_b": float(np.mean(y)),
                    "median_a": float(np.median(x)),
                    "median_b": float(np.median(y)),
                    "u_stat": u,
                    "p_value": p,
                    "cliffs_delta_a_minus_b": cliffs_delta(x, y),
                }
            )
    out = pd.DataFrame(rows)
    if not out.empty:
        out["p_adj_BH"] = bh_adjust(out["p_value"].to_numpy())
    return out


def stage_components(cell_metrics):
    out = cell_metrics.groupby("stage", observed=True).agg(
        n_cells=("sample_name", "count"),
        s_rna_mean=("s_rna", "mean"),
        detected_genes_mean=("detected_genes", "mean"),
        detected_gene_score_mean=("detected_gene_score", "mean"),
        marker_score_mean=("marker_score", "mean"),
        potency_score_mean=("potency_score", "mean"),
        potency_score_median=("potency_score", "median"),
    ).reset_index()
    out["stage"] = out["stage"].astype(str)
    return out


def marker_stage_summary(marker, marker_z):
    mean = marker.groupby("stage", observed=True)[POTENCY_MARKERS].mean().reset_index()
    mean_z = marker_z.groupby("stage", observed=True)[POTENCY_MARKERS].mean().reset_index()
    mean["stage"] = mean["stage"].astype(str)
    mean_z["stage"] = mean_z["stage"].astype(str)
    long = mean.melt(id_vars="stage", var_name="marker", value_name="log1p_rpkm_mean")
    long_z = mean_z.melt(id_vars="stage", var_name="marker", value_name="zscore_mean")
    return long.merge(long_z, on=["stage", "marker"], how="left"), mean_z


def plot_outputs(components, boot, marker_z_stage):
    try:
        import matplotlib.pyplot as plt
    except ImportError:
        print("matplotlib unavailable; skipping figures.")
        return
    FIGS.mkdir(parents=True, exist_ok=True)

    x = np.arange(len(components))
    plt.figure(figsize=(9, 5))
    plt.plot(x, components["detected_gene_score_mean"], marker="o", label="Detected-gene score")
    plt.plot(x, components["marker_score_mean"], marker="o", label="Marker score")
    plt.plot(x, components["potency_score_mean"], marker="o", linewidth=2.5, label="Combined potency")
    plt.xticks(x, components["stage"], rotation=35, ha="right")
    plt.ylabel("Normalized score")
    plt.xlabel("RNA developmental stage")
    plt.legend(frameon=False)
    plt.tight_layout()
    plt.savefig(FIGS / "GSE36552_potency_component_by_stage.png", dpi=300)
    plt.savefig(FIGS / "GSE36552_potency_component_by_stage.pdf")

    pot = boot[boot["metric"] == "potency_score"].copy()
    pot["stage"] = pd.Categorical(pot["stage"], categories=RNA_STAGE_ORDER, ordered=True)
    pot = pot.sort_values("stage")
    x = np.arange(len(pot))
    yerr = np.vstack([pot["mean"] - pot["ci_low"], pot["ci_high"] - pot["mean"]])
    plt.figure(figsize=(8, 4.8))
    plt.errorbar(x, pot["mean"], yerr=yerr, marker="o", linewidth=2, capsize=4)
    plt.xticks(x, pot["stage"].astype(str), rotation=35, ha="right")
    plt.ylabel("Potency proxy mean with 95% bootstrap CI")
    plt.xlabel("RNA developmental stage")
    plt.tight_layout()
    plt.savefig(FIGS / "GSE36552_potency_bootstrap_ci.png", dpi=300)
    plt.savefig(FIGS / "GSE36552_potency_bootstrap_ci.pdf")

    heat = marker_z_stage.set_index("stage")[POTENCY_MARKERS]
    plt.figure(figsize=(9, 4.8))
    im = plt.imshow(heat.values, aspect="auto", cmap="coolwarm", vmin=-1.5, vmax=1.5)
    plt.colorbar(im, label="Mean marker z-score")
    plt.yticks(np.arange(len(heat.index)), heat.index)
    plt.xticks(np.arange(len(POTENCY_MARKERS)), POTENCY_MARKERS, rotation=45, ha="right")
    plt.tight_layout()
    plt.savefig(FIGS / "GSE36552_marker_heatmap.png", dpi=300)
    plt.savefig(FIGS / "GSE36552_marker_heatmap.pdf")


def main():
    ap = argparse.ArgumentParser()
    ap.add_argument("--n-boot", type=int, default=2000)
    args = ap.parse_args()

    TABLES.mkdir(parents=True, exist_ok=True)
    cell_path = TABLES / "GSE36552_RNA_entropy_potency_cell_metrics.tsv"
    if not cell_path.exists():
        raise FileNotFoundError(f"Missing {cell_path}. Run script 13 first.")

    cell = pd.read_csv(cell_path, sep="\t")
    cell = cell[cell["stage"].isin(RNA_STAGE_ORDER)].copy()
    cell["stage"] = pd.Categorical(cell["stage"], categories=RNA_STAGE_ORDER, ordered=True)

    marker, marker_z = make_marker_matrix()
    components = stage_components(cell)
    boot = bootstrap_ci(cell, n_boot=args.n_boot)
    tests = pairwise_tests(cell)
    marker_long, marker_z_stage = marker_stage_summary(marker, marker_z)

    components.to_csv(TABLES / "GSE36552_potency_component_by_stage.tsv", sep="\t", index=False)
    boot.to_csv(TABLES / "GSE36552_potency_bootstrap_ci.tsv", sep="\t", index=False)
    tests.to_csv(TABLES / "GSE36552_potency_pairwise_tests.tsv", sep="\t", index=False)
    marker.to_csv(TABLES / "GSE36552_marker_log1p_rpkm_by_cell.tsv", sep="\t", index=False)
    marker_z.to_csv(TABLES / "GSE36552_marker_zscore_by_cell.tsv", sep="\t", index=False)
    marker_long.to_csv(TABLES / "GSE36552_marker_score_by_stage.tsv", sep="\t", index=False)
    marker_z_stage.to_csv(TABLES / "GSE36552_marker_zscore_heatmap_matrix.tsv", sep="\t", index=False)
    plot_outputs(components, boot, marker_z_stage)

    focus = tests[tests["metric"].isin(["marker_score", "potency_score", "s_rna"])].copy()
    print("Potency components by stage:")
    print(components.to_string(index=False))
    print("\nBootstrap CI for potency_score:")
    print(boot[boot["metric"] == "potency_score"].to_string(index=False))
    print("\nKey pairwise tests:")
    print(focus.to_string(index=False))
    print("\nMarker z-score stage matrix:")
    print(marker_z_stage.to_string(index=False))
    print("\nWrote:", TABLES / "GSE36552_potency_component_by_stage.tsv")
    print("Wrote:", TABLES / "GSE36552_potency_pairwise_tests.tsv")
    print("Wrote:", FIGS / "GSE36552_marker_heatmap.png")


if __name__ == "__main__":
    main()
```
