## Supplemental code for "An entropy-based diagnostic framework for characterizing methylation state dynamics during preimplantation development": 15_experiment3_tro_composite_and_marker_robustness.py

```
from pathlib import Path
import argparse
import math

import numpy as np
import pandas as pd


STAGE_ORDER = ["MII oocyte", "zygote/PN", "2-cell", "4-cell", "8-cell", "morula", "blastocyst"]
RNA_STAGE_ORDER = ["oocyte", "zygote", "2-cell", "4-cell", "8-cell", "morula", "blastocyst"]
MARKERS_CORE = ["POU5F1", "NANOG", "SOX2", "KLF4", "KLF17", "TFAP2C", "GDF3"]
MARKERS_FULL = ["POU5F1", "NANOG", "SOX2", "KLF4", "DPPA3", "ZSCAN4", "KLF17", "TFAP2C", "GDF3"]


def project_root():
    here = Path(__file__).resolve()
    server_root = Path("/root/autodl-tmp/TRO_Project")
    if server_root.exists() and str(here).startswith("/root/"):
        return server_root
    return here.parents[1]


ROOT = project_root()
TABLES = ROOT / "results" / "tables" if (ROOT / "results" / "tables").exists() else ROOT / "tables"
FIGS = ROOT / "results" / "figures" if (ROOT / "results" / "figures").exists() else ROOT / "figures"


def zscore(x):
    arr = np.asarray(x, dtype=float)
    sd = np.nanstd(arr, ddof=0)
    if not np.isfinite(sd) or sd == 0:
        return np.zeros_like(arr, dtype=float)
    return (arr - np.nanmean(arr)) / sd


def minmax(x):
    arr = np.asarray(x, dtype=float)
    lo = np.nanmin(arr)
    hi = np.nanmax(arr)
    if not np.isfinite(lo) or not np.isfinite(hi) or hi == lo:
        return np.zeros_like(arr, dtype=float)
    return (arr - lo) / (hi - lo)


def bh_adjust(pvals):
    p = np.asarray(pvals, dtype=float)
    out = np.full(len(p), np.nan)
    ok = np.isfinite(p)
    if ok.sum() == 0:
        return out
    idx = np.where(ok)[0]
    order = idx[np.argsort(p[idx])]
    ranked = p[order]
    m = len(ranked)
    adj = ranked * m / np.arange(1, m + 1)
    adj = np.minimum.accumulate(adj[::-1])[::-1]
    out[order] = np.minimum(adj, 1.0)
    return out


def cliffs_delta(x, y):
    x = np.asarray(x, dtype=float)
    y = np.asarray(y, dtype=float)
    gt = sum((xi > y).sum() for xi in x)
    lt = sum((xi < y).sum() for xi in x)
    return (gt - lt) / (len(x) * len(y))


def mannwhitney(x, y):
    try:
        from scipy import stats

        res = stats.mannwhitneyu(x, y, alternative="two-sided")
        return float(res.statistic), float(res.pvalue)
    except Exception:
        x = np.asarray(x, dtype=float)
        y = np.asarray(y, dtype=float)
        x = x[np.isfinite(x)]
        y = y[np.isfinite(y)]
        n1 = len(x)
        n2 = len(y)
        if n1 == 0 or n2 == 0:
            return np.nan, np.nan
        vals = np.concatenate([x, y])
        ranks = pd.Series(vals).rank(method="average").to_numpy()
        r1 = ranks[:n1].sum()
        u1 = r1 - n1 * (n1 + 1) / 2.0
        mean_u = n1 * n2 / 2.0
        _, counts = np.unique(vals, return_counts=True)
        n = n1 + n2
        tie_term = np.sum(counts**3 - counts)
        var_u = n1 * n2 / 12.0 * ((n + 1) - tie_term / (n * (n - 1))) if n > 1 else np.nan
        if not np.isfinite(var_u) or var_u <= 0:
            return float(u1), np.nan
        z = (u1 - mean_u) / np.sqrt(var_u)
        p = math.erfc(abs(z) / np.sqrt(2.0))
        return float(u1), float(p)


def compute_tro_scores():
    dual = pd.read_csv(TABLES / "dual_entropy_stage_table.tsv", sep="\t")
    dual = dual[dual["stage"].isin(STAGE_ORDER)].copy()
    dual["stage"] = pd.Categorical(dual["stage"], categories=STAGE_ORDER, ordered=True)
    dual = dual.sort_values("stage").reset_index(drop=True)

    dual["z_neg_S_epi_age"] = zscore(-dual["S_epi_age"].to_numpy())
    dual["z_PotencyScore"] = zscore(dual["PotencyScore"].to_numpy())
    dual["GZ_score"] = dual["z_neg_S_epi_age"] + dual["z_PotencyScore"]
    dual["PotencyPreserve"] = dual["PotencyScore"] / dual["PotencyScore"].max()
    dual["TRO_score"] = dual["ResetScore"] * dual["PotencyPreserve"]
    dual["GZ_rank"] = dual["GZ_score"].rank(method="min", ascending=False).astype(int)
    dual["TRO_rank"] = dual["TRO_score"].rank(method="min", ascending=False).astype(int)

    keep = [
        "stage",
        "rna_stage",
        "S_epi_age",
        "S_epi",
        "ResetScore",
        "S_RNA",
        "PotencyScore",
        "PotencyPreserve",
        "z_neg_S_epi_age",
        "z_PotencyScore",
        "GZ_score",
        "TRO_score",
        "GZ_rank",
        "TRO_rank",
        "n_cells",
    ]
    out = dual[keep].copy()
    out.to_csv(TABLES / "TRO_composite_score_by_stage.tsv", sep="\t", index=False)
    return out


def leave_one_marker_out(marker_panel):
    cell = pd.read_csv(TABLES / "GSE36552_RNA_entropy_potency_cell_metrics.tsv", sep="\t")
    marker = pd.read_csv(TABLES / "GSE36552_marker_log1p_rpkm_by_cell.tsv", sep="\t")
    merged = cell[["sample_name", "stage", "detected_gene_score"]].merge(marker, on=["sample_name", "stage"], how="inner")
    merged = merged[merged["stage"].isin(RNA_STAGE_ORDER)].copy()
    markers = [m for m in marker_panel if m in merged.columns]

    rows = []
    for removed in ["none"] + markers:
        use_markers = markers if removed == "none" else [m for m in markers if m != removed]
        raw = merged[use_markers].mean(axis=1)
        marker_score = minmax(raw.to_numpy())
        potency = 0.5 * merged["detected_gene_score"].to_numpy(dtype=float) + 0.5 * marker_score
        tmp = merged[["sample_name", "stage"]].copy()
        tmp["marker_score_recomputed"] = marker_score
        tmp["potency_score_recomputed"] = potency

        for metric in ["marker_score_recomputed", "potency_score_recomputed"]:
            m = tmp.loc[tmp["stage"] == "morula", metric].dropna().to_numpy(dtype=float)
            b = tmp.loc[tmp["stage"] == "blastocyst", metric].dropna().to_numpy(dtype=float)
            e = tmp.loc[tmp["stage"] == "8-cell", metric].dropna().to_numpy(dtype=float)
            u_mb, p_mb = mannwhitney(m, b)
            u_m8, p_m8 = mannwhitney(m, e)
            rows.append(
                {
                    "marker_panel": ",".join(markers),
                    "removed_marker": removed,
                    "metric": metric,
                    "n_markers_used": len(use_markers),
                    "morula_mean": float(np.mean(m)),
                    "blastocyst_mean": float(np.mean(b)),
                    "eight_cell_mean": float(np.mean(e)),
                    "morula_vs_blastocyst_u": u_mb,
                    "morula_vs_blastocyst_p": p_mb,
                    "morula_vs_blastocyst_cliffs_delta": cliffs_delta(m, b),
                    "morula_vs_8cell_u": u_m8,
                    "morula_vs_8cell_p": p_m8,
                    "morula_vs_8cell_cliffs_delta": cliffs_delta(m, e),
                    "conclusion": "pass" if np.mean(m) > np.mean(b) else "check",
                }
            )

    out = pd.DataFrame(rows)
    mask = out["metric"] == "potency_score_recomputed"
    out.loc[mask, "morula_vs_blastocyst_p_adj_BH"] = bh_adjust(out.loc[mask, "morula_vs_blastocyst_p"].to_numpy())
    out.loc[~mask, "morula_vs_blastocyst_p_adj_BH"] = bh_adjust(out.loc[~mask, "morula_vs_blastocyst_p"].to_numpy())
    out.to_csv(TABLES / "marker_leave_one_out_summary.tsv", sep="\t", index=False)
    return out


def expanded_marker_panel():
    candidate = [
        "POU5F1", "NANOG", "SOX2", "KLF4", "KLF17", "TFAP2C", "GDF3",
        "PRDM14", "DPPA3", "SALL4", "LIN28A", "UTF1", "ZFP42", "TDGF1", "DNMT3L", "TCL1A",
    ]
    cell = pd.read_csv(TABLES / "GSE36552_RNA_entropy_potency_cell_metrics.tsv", sep="\t")
    marker = pd.read_csv(TABLES / "GSE36552_marker_log1p_rpkm_by_cell.tsv", sep="\t")
    available = [m for m in candidate if m in marker.columns]
    merged = cell[["sample_name", "stage", "detected_gene_score"]].merge(marker, on=["sample_name", "stage"], how="inner")
    if not available:
        return pd.DataFrame()
    raw = merged[available].mean(axis=1)
    merged["expanded_marker_score"] = minmax(raw.to_numpy())
    merged["expanded_potency_score"] = 0.5 * merged["detected_gene_score"].to_numpy(dtype=float) + 0.5 * merged["expanded_marker_score"]
    summary = merged.groupby("stage", observed=True).agg(
        n_cells=("sample_name", "count"),
        expanded_marker_score_mean=("expanded_marker_score", "mean"),
        expanded_potency_score_mean=("expanded_potency_score", "mean"),
        expanded_potency_score_median=("expanded_potency_score", "median"),
    ).reset_index()
    summary["stage"] = pd.Categorical(summary["stage"], categories=RNA_STAGE_ORDER, ordered=True)
    summary = summary.sort_values("stage")
    summary["markers_available"] = ",".join(available)
    summary.to_csv(TABLES / "expanded_marker_panel_potency_score.tsv", sep="\t", index=False)
    return summary


def plot_tro(tro):
    try:
        import matplotlib.pyplot as plt
    except ImportError:
        print("matplotlib unavailable; skipping TRO figures.")
        return
    FIGS.mkdir(parents=True, exist_ok=True)
    x = np.arange(len(tro))

    plt.figure(figsize=(9, 5))
    plt.plot(x, tro["ResetScore"], marker="o", label="ResetScore")
    plt.plot(x, tro["PotencyPreserve"], marker="o", label="PotencyPreserve")
    plt.plot(x, tro["TRO_score"], marker="o", linewidth=2.5, label="TRO score")
    plt.xticks(x, tro["stage"], rotation=35, ha="right")
    plt.ylabel("Score")
    plt.xlabel("Developmental stage")
    plt.legend(frameon=False)
    idx = int(np.where(tro["stage"].astype(str).to_numpy() == "morula")[0][0])
    plt.scatter([idx], [tro.loc[idx, "TRO_score"]], s=120, facecolors="none", edgecolors="red", linewidth=2)
    plt.tight_layout()
    plt.savefig(FIGS / "TRO_composite_score_by_stage.png", dpi=300)
    plt.savefig(FIGS / "TRO_composite_score_by_stage.pdf")

    plt.figure(figsize=(6.8, 5.8))
    plt.scatter(tro["S_epi_age"], tro["PotencyScore"], s=85, zorder=3)
    for i in range(len(tro) - 1):
        plt.annotate(
            "",
            xy=(tro.loc[i + 1, "S_epi_age"], tro.loc[i + 1, "PotencyScore"]),
            xytext=(tro.loc[i, "S_epi_age"], tro.loc[i, "PotencyScore"]),
            arrowprops=dict(arrowstyle="->", color="0.45", lw=1.2),
            zorder=2,
        )
    for _, r in tro.iterrows():
        label = str(r["stage"]).replace("MII oocyte", "MII").replace("zygote/PN", "zygote")
        plt.annotate(label, (r["S_epi_age"], r["PotencyScore"]), xytext=(5, 5), textcoords="offset points", fontsize=8)
    morula = tro[tro["stage"].astype(str) == "morula"].iloc[0]
    plt.scatter([morula["S_epi_age"]], [morula["PotencyScore"]], s=180, facecolors="none", edgecolors="red", linewidth=2.2)
    plt.annotate(
        "computational ground-zero\nlow S_epi-age + high potency",
        (morula["S_epi_age"], morula["PotencyScore"]),
        xytext=(-115, 42),
        textcoords="offset points",
        arrowprops=dict(arrowstyle="->", color="red", lw=1.2),
        fontsize=9,
        color="red",
    )
    plt.xlabel("S_epi-age")
    plt.ylabel("PotencyScore")
    plt.tight_layout()
    plt.savefig(FIGS / "dual_entropy_phase_map_final.png", dpi=300)
    plt.savefig(FIGS / "dual_entropy_phase_map_final.pdf")


def plot_marker_robustness(loo):
    try:
        import matplotlib.pyplot as plt
    except ImportError:
        return
    df = loo[loo["metric"] == "potency_score_recomputed"].copy()
    df["removed_marker"] = pd.Categorical(df["removed_marker"], categories=df["removed_marker"].tolist(), ordered=True)
    x = np.arange(len(df))
    plt.figure(figsize=(10, 4.8))
    plt.plot(x, df["morula_mean"], marker="o", label="morula")
    plt.plot(x, df["blastocyst_mean"], marker="o", label="blastocyst")
    plt.xticks(x, df["removed_marker"], rotation=35, ha="right")
    plt.ylabel("Recomputed potency score")
    plt.xlabel("Removed marker")
    plt.legend(frameon=False)
    plt.tight_layout()
    plt.savefig(FIGS / "marker_leave_one_out_summary.png", dpi=300)
    plt.savefig(FIGS / "marker_leave_one_out_summary.pdf")


def main():
    ap = argparse.ArgumentParser()
    ap.add_argument("--marker-panel", choices=["core", "full"], default="core")
    args = ap.parse_args()
    TABLES.mkdir(parents=True, exist_ok=True)
    FIGS.mkdir(parents=True, exist_ok=True)

    panel = MARKERS_CORE if args.marker_panel == "core" else MARKERS_FULL
    tro = compute_tro_scores()
    loo = leave_one_marker_out(panel)
    expanded = expanded_marker_panel()
    plot_tro(tro)
    plot_marker_robustness(loo)

    print("TRO composite score by stage:")
    print(tro[["stage", "S_epi_age", "ResetScore", "PotencyScore", "PotencyPreserve", "GZ_score", "TRO_score", "GZ_rank", "TRO_rank"]].to_string(index=False))
    print("\nLeave-one-marker-out potency robustness:")
    print(loo[loo["metric"] == "potency_score_recomputed"][
        [
            "removed_marker",
            "morula_mean",
            "blastocyst_mean",
            "eight_cell_mean",
            "morula_vs_blastocyst_p",
            "morula_vs_blastocyst_p_adj_BH",
            "morula_vs_blastocyst_cliffs_delta",
            "conclusion",
        ]
    ].to_string(index=False))
    if not expanded.empty:
        print("\nExpanded marker panel summary:")
        print(expanded.to_string(index=False))
    print("\nWrote:", TABLES / "TRO_composite_score_by_stage.tsv")
    print("Wrote:", TABLES / "marker_leave_one_out_summary.tsv")
    print("Wrote:", TABLES / "expanded_marker_panel_potency_score.tsv")
    print("Wrote:", FIGS / "TRO_composite_score_by_stage.png")
    print("Wrote:", FIGS / "dual_entropy_phase_map_final.png")


if __name__ == "__main__":
    main()
```
