## Supplemental code for "An entropy-based diagnostic framework for characterizing methylation state dynamics during preimplantation development": 16_experiment4_gse44183_external_rna_validation.py

```
from pathlib import Path
import argparse
import gzip
import math
import re
import urllib.request

import numpy as np
import pandas as pd


URL = "https://ftp.ncbi.nlm.nih.gov/geo/series/GSE44nnn/GSE44183/suppl/GSE44183_human_expression_mat.txt.gz"
RNA_STAGE_ORDER = ["oocyte", "pronuclear", "zygote", "2-cell", "4-cell", "8-cell", "morula"]
MARKERS = ["POU5F1", "NANOG", "SOX2", "KLF4", "DPPA3", "ZSCAN4", "KLF17", "TFAP2C", "GDF3"]


def project_root():
    here = Path(__file__).resolve()
    server_root = Path("/root/autodl-tmp/TRO_Project")
    if server_root.exists() and str(here).startswith("/root/"):
        return server_root
    return here.parents[1]


ROOT = project_root()
RAW = ROOT / "data_raw" / "GSE44183_rna"
TABLES = ROOT / "results" / "tables" if (ROOT / "results" / "tables").exists() else ROOT / "tables"
FIGS = ROOT / "results" / "figures" if (ROOT / "results" / "figures").exists() else ROOT / "figures"


def download(url, path):
    path.parent.mkdir(parents=True, exist_ok=True)
    if path.exists() and path.stat().st_size > 1000:
        return path
    print(f"Downloading {url}")
    urllib.request.urlretrieve(url, path)
    return path


def infer_stage(sample):
    s = str(sample).lower().replace("_", " ").replace("-", " ")
    if "morula" in s:
        return "morula"
    if re.search(r"\b8\s*cell\b", s):
        return "8-cell"
    if re.search(r"\b4\s*cell\b", s):
        return "4-cell"
    if re.search(r"\b2\s*cell\b", s):
        return "2-cell"
    if "zygote" in s:
        return "zygote"
    if "pronuclear" in s or "pn" in s:
        return "pronuclear"
    if "oocyte" in s:
        return "oocyte"
    return None


def minmax(x):
    arr = np.asarray(x, dtype=float)
    lo = np.nanmin(arr)
    hi = np.nanmax(arr)
    if not np.isfinite(lo) or not np.isfinite(hi) or hi == lo:
        return np.zeros_like(arr, dtype=float)
    return (arr - lo) / (hi - lo)


def shannon_entropy(values):
    x = np.array(values, dtype=float, copy=True)
    x[~np.isfinite(x)] = 0.0
    x[x < 0] = 0.0
    total = x.sum()
    if total <= 0:
        return np.nan
    p = x / total
    p = p[p > 0]
    return float(-(p * np.log(p)).sum())


def bh_adjust(pvals):
    p = np.asarray(pvals, dtype=float)
    out = np.full(len(p), np.nan)
    ok = np.isfinite(p)
    if ok.sum() == 0:
        return out
    idx = np.where(ok)[0]
    order = idx[np.argsort(p[idx])]
    ranked = p[order]
    m = len(ranked)
    adj = ranked * m / np.arange(1, m + 1)
    adj = np.minimum.accumulate(adj[::-1])[::-1]
    out[order] = np.minimum(adj, 1.0)
    return out


def mannwhitney(x, y):
    try:
        from scipy import stats
        res = stats.mannwhitneyu(x, y, alternative="two-sided")
        return float(res.statistic), float(res.pvalue)
    except Exception:
        x = np.asarray(x, dtype=float)
        y = np.asarray(y, dtype=float)
        x = x[np.isfinite(x)]
        y = y[np.isfinite(y)]
        n1, n2 = len(x), len(y)
        if n1 == 0 or n2 == 0:
            return np.nan, np.nan
        vals = np.concatenate([x, y])
        ranks = pd.Series(vals).rank(method="average").to_numpy()
        r1 = ranks[:n1].sum()
        u1 = r1 - n1 * (n1 + 1) / 2
        mean_u = n1 * n2 / 2
        _, counts = np.unique(vals, return_counts=True)
        n = n1 + n2
        tie_term = np.sum(counts**3 - counts)
        var_u = n1 * n2 / 12 * ((n + 1) - tie_term / (n * (n - 1))) if n > 1 else np.nan
        if not np.isfinite(var_u) or var_u <= 0:
            return float(u1), np.nan
        z = (u1 - mean_u) / np.sqrt(var_u)
        return float(u1), float(math.erfc(abs(z) / np.sqrt(2)))


def cliffs_delta(x, y):
    x = np.asarray(x, dtype=float)
    y = np.asarray(y, dtype=float)
    gt = sum((xi > y).sum() for xi in x)
    lt = sum((xi < y).sum() for xi in x)
    return (gt - lt) / (len(x) * len(y))


def load_matrix(path):
    with gzip.open(path, "rt", encoding="utf-8", errors="replace") as fh:
        expr = pd.read_csv(fh, sep="\t")
    gene_col = expr.columns[0]
    expr = expr.rename(columns={gene_col: "gene_symbol"})
    expr["gene_symbol"] = expr["gene_symbol"].astype(str).str.upper()
    sample_cols = [c for c in expr.columns if c != "gene_symbol"]
    for c in sample_cols:
        expr[c] = pd.to_numeric(expr[c], errors="coerce").fillna(0.0)
    return expr, sample_cols


def compute_cell_metrics(expr, sample_cols):
    genes = expr["gene_symbol"]
    matrix = expr[sample_cols]
    detected = (matrix > 0).sum(axis=0).astype(float)
    detected_score = pd.Series(minmax(detected.to_numpy()), index=sample_cols)
    marker_present = [m for m in MARKERS if m in set(genes)]
    marker_raw = pd.DataFrame(index=sample_cols)
    for m in marker_present:
        idx = np.where(genes == m)[0]
        marker_raw[m] = np.log1p(matrix.iloc[idx].mean(axis=0))
    marker_score = pd.Series(minmax(marker_raw.mean(axis=1).to_numpy()), index=sample_cols)
    potency = 0.5 * detected_score + 0.5 * marker_score

    rows = []
    marker_rows = []
    for sample in sample_cols:
        stage = infer_stage(sample)
        if stage is None:
            continue
        rows.append(
            {
                "sample_name": sample,
                "stage": stage,
                "s_rna": shannon_entropy(matrix[sample].to_numpy()),
                "detected_genes": int(detected.loc[sample]),
                "detected_gene_score": float(detected_score.loc[sample]),
                "marker_score": float(marker_score.loc[sample]),
                "potency_score": float(potency.loc[sample]),
            }
        )
        mr = {"sample_name": sample, "stage": stage}
        for m in marker_present:
            mr[m] = float(marker_raw.loc[sample, m])
        marker_rows.append(mr)
    cell = pd.DataFrame(rows)
    cell["stage"] = pd.Categorical(cell["stage"], categories=RNA_STAGE_ORDER, ordered=True)
    cell = cell.sort_values(["stage", "sample_name"]).reset_index(drop=True)
    marker = pd.DataFrame(marker_rows)
    marker["stage"] = pd.Categorical(marker["stage"], categories=RNA_STAGE_ORDER, ordered=True)
    marker = marker.sort_values(["stage", "sample_name"]).reset_index(drop=True)
    return cell, marker, marker_present


def summarize(cell):
    out = cell.groupby("stage", observed=True).agg(
        n_cells=("sample_name", "count"),
        s_rna_mean=("s_rna", "mean"),
        detected_genes_mean=("detected_genes", "mean"),
        marker_score_mean=("marker_score", "mean"),
        potency_score_mean=("potency_score", "mean"),
        potency_score_median=("potency_score", "median"),
    ).reset_index()
    out["stage"] = out["stage"].astype(str)
    out["potency_rank"] = out["potency_score_mean"].rank(method="min", ascending=False).astype(int)
    return out


def pairwise(cell):
    pairs = [("morula", "8-cell"), ("morula", "4-cell"), ("morula", "2-cell"), ("8-cell", "4-cell")]
    rows = []
    for metric in ["s_rna", "marker_score", "potency_score"]:
        for a, b in pairs:
            x = cell.loc[cell["stage"] == a, metric].dropna().to_numpy(float)
            y = cell.loc[cell["stage"] == b, metric].dropna().to_numpy(float)
            if len(x) == 0 or len(y) == 0:
                continue
            u, p = mannwhitney(x, y)
            rows.append(
                {
                    "metric": metric,
                    "comparison": f"{a} vs {b}",
                    "stage_a": a,
                    "stage_b": b,
                    "n_a": len(x),
                    "n_b": len(y),
                    "mean_a": float(np.mean(x)),
                    "mean_b": float(np.mean(y)),
                    "u_stat": u,
                    "p_value": p,
                    "cliffs_delta_a_minus_b": cliffs_delta(x, y),
                }
            )
    out = pd.DataFrame(rows)
    if not out.empty:
        out["p_adj_BH"] = bh_adjust(out["p_value"].to_numpy())
    return out


def plot(summary):
    try:
        import matplotlib.pyplot as plt
    except ImportError:
        return
    FIGS.mkdir(parents=True, exist_ok=True)
    x = np.arange(len(summary))
    plt.figure(figsize=(8, 4.8))
    plt.plot(x, summary["marker_score_mean"], marker="o", label="Marker score")
    plt.plot(x, summary["potency_score_mean"], marker="o", linewidth=2.5, label="Potency score")
    plt.xticks(x, summary["stage"], rotation=35, ha="right")
    plt.ylabel("Normalized score")
    plt.xlabel("GSE44183 human stage")
    plt.legend(frameon=False)
    plt.tight_layout()
    plt.savefig(FIGS / "GSE44183_external_potency_validation.png", dpi=300)
    plt.savefig(FIGS / "GSE44183_external_potency_validation.pdf")


def main():
    ap = argparse.ArgumentParser()
    ap.add_argument("--force-download", action="store_true")
    args = ap.parse_args()
    RAW.mkdir(parents=True, exist_ok=True)
    TABLES.mkdir(parents=True, exist_ok=True)
    path = RAW / "GSE44183_human_expression_mat.txt.gz"
    if args.force_download and path.exists():
        path.unlink()
    download(URL, path)

    expr, sample_cols = load_matrix(path)
    cell, marker, marker_present = compute_cell_metrics(expr, sample_cols)
    summary = summarize(cell)
    tests = pairwise(cell)
    cell.to_csv(TABLES / "GSE44183_external_potency_cell_metrics.tsv", sep="\t", index=False)
    marker.to_csv(TABLES / "GSE44183_external_marker_log1p_by_cell.tsv", sep="\t", index=False)
    summary.to_csv(TABLES / "GSE44183_external_potency_validation.tsv", sep="\t", index=False)
    tests.to_csv(TABLES / "GSE44183_external_potency_pairwise_tests.tsv", sep="\t", index=False)
    plot(summary)

    print("GSE44183 external potency validation:")
    print(summary.to_string(index=False))
    print("\nPairwise tests:")
    print(tests.to_string(index=False))
    print("\nMarkers found:", ", ".join(marker_present))
    print("Wrote:", TABLES / "GSE44183_external_potency_validation.tsv")
    print("Wrote:", FIGS / "GSE44183_external_potency_validation.png")


if __name__ == "__main__":
    main()
```
