## Supplemental code for "An entropy-based diagnostic framework for characterizing methylation state dynamics during preimplantation development": 17_experiment5_transition_reset_cost.py

```
from pathlib import Path
import argparse

import numpy as np
import pandas as pd


STAGE_ORDER = ["MII oocyte", "zygote/PN", "2-cell", "4-cell", "8-cell", "morula", "blastocyst"]


def project_root():
    here = Path(__file__).resolve()
    server_root = Path("/root/autodl-tmp/TRO_Project")
    if server_root.exists() and str(here).startswith("/root/"):
        return server_root
    return here.parents[1]


ROOT = project_root()
TABLES = ROOT / "results" / "tables" if (ROOT / "results" / "tables").exists() else ROOT / "tables"
FIGS = ROOT / "results" / "figures" if (ROOT / "results" / "figures").exists() else ROOT / "figures"


def zscore(x):
    arr = np.asarray(x, dtype=float)
    sd = np.nanstd(arr, ddof=0)
    if not np.isfinite(sd) or sd == 0:
        return np.zeros_like(arr, dtype=float)
    return (arr - np.nanmean(arr)) / sd


def minmax(x):
    arr = np.asarray(x, dtype=float)
    lo = np.nanmin(arr)
    hi = np.nanmax(arr)
    if not np.isfinite(lo) or not np.isfinite(hi) or hi == lo:
        return np.zeros_like(arr)
    return (arr - lo) / (hi - lo)


def load_state_table():
    path = TABLES / "TRO_composite_score_by_stage.tsv"
    if not path.exists():
        path = TABLES / "dual_entropy_stage_table.tsv"
    df = pd.read_csv(path, sep="\t")
    df = df[df["stage"].isin(STAGE_ORDER)].copy()
    df["stage"] = pd.Categorical(df["stage"], categories=STAGE_ORDER, ordered=True)
    df = df.sort_values("stage").reset_index(drop=True)
    if "TRO_score" not in df.columns:
        df["PotencyPreserve"] = df["PotencyScore"] / df["PotencyScore"].max()
        df["TRO_score"] = df["ResetScore"] * df["PotencyPreserve"]
    return df


def build_state_vectors(df):
    out = df.copy()
    out["damage_proxy"] = out["S_epi_age"]
    out["potency_proxy"] = out["PotencyScore"]
    out["reset_proxy"] = out["ResetScore"]
    out["rna_order_proxy"] = -out["S_RNA"]

    for col in ["damage_proxy", "potency_proxy", "reset_proxy", "rna_order_proxy", "S_epi", "S_RNA"]:
        out[f"z_{col}"] = zscore(out[col].to_numpy())

    out["bio_age_score"] = (
        out["z_damage_proxy"]
        - out["z_potency_proxy"]
        - out["z_reset_proxy"]
    )
    out["bio_youth_score"] = -out["bio_age_score"]
    out["bio_age_rank"] = out["bio_age_score"].rank(method="min", ascending=True).astype(int)
    out.to_csv(TABLES / "TRO_stage_state_vectors.tsv", sep="\t", index=False)
    return out


def transition_costs(state):
    rows = []
    vec_cols = ["z_damage_proxy", "z_potency_proxy", "z_reset_proxy", "z_rna_order_proxy"]
    for i in range(len(state) - 1):
        a = state.iloc[i]
        b = state.iloc[i + 1]
        va = a[vec_cols].to_numpy(dtype=float)
        vb = b[vec_cols].to_numpy(dtype=float)
        delta = vb - va
        damage_reduction = float(a["damage_proxy"] - b["damage_proxy"])
        potency_change = float(b["potency_proxy"] - a["potency_proxy"])
        reset_gain = float(b["reset_proxy"] - a["reset_proxy"])
        rna_entropy_change = float(b["S_RNA"] - a["S_RNA"])
        cost = float(np.linalg.norm(delta))
        productive_reset_gain = max(0.0, damage_reduction) + max(0.0, potency_change) + max(0.0, reset_gain)
        reset_efficiency = productive_reset_gain / cost if cost > 0 else np.nan
        rows.append(
            {
                "transition": f"{a['stage']} -> {b['stage']}",
                "stage_from": a["stage"],
                "stage_to": b["stage"],
                "transition_cost": cost,
                "damage_reduction": damage_reduction,
                "potency_change": potency_change,
                "reset_gain": reset_gain,
                "rna_entropy_change": rna_entropy_change,
                "productive_reset_gain": productive_reset_gain,
                "reset_efficiency": reset_efficiency,
                "delta_damage_z": float(delta[0]),
                "delta_potency_z": float(delta[1]),
                "delta_reset_z": float(delta[2]),
                "delta_rna_order_z": float(delta[3]),
            }
        )
    out = pd.DataFrame(rows)
    out["cost_rank"] = out["transition_cost"].rank(method="min", ascending=False).astype(int)
    out["efficiency_rank"] = out["reset_efficiency"].rank(method="min", ascending=False).astype(int)
    out.to_csv(TABLES / "TRO_stage_transition_cost.tsv", sep="\t", index=False)
    return out


def reset_depth_summary(state, trans):
    mii = state[state["stage"].astype(str) == "MII oocyte"].iloc[0]
    morula = state[state["stage"].astype(str) == "morula"].iloc[0]
    blast = state[state["stage"].astype(str) == "blastocyst"].iloc[0]
    rows = []
    for target_name, target in [("morula", morula), ("blastocyst", blast)]:
        delta_s_epi_age = float(mii["S_epi_age"] - target["S_epi_age"])
        delta_s_epi = float(mii["S_epi"] - target["S_epi"])
        delta_potency = float(target["PotencyScore"] - mii["PotencyScore"])
        delta_reset = float(target["ResetScore"] - mii["ResetScore"])
        rows.append(
            {
                "from_stage": "MII oocyte",
                "to_stage": target_name,
                "delta_S_epi_age_reduction": delta_s_epi_age,
                "delta_S_epi_reduction": delta_s_epi,
                "delta_PotencyScore": delta_potency,
                "delta_ResetScore": delta_reset,
                "relative_S_epi_age_reduction": delta_s_epi_age / float(mii["S_epi_age"]),
                "TRO_score_to_stage": float(target["TRO_score"]),
            }
        )
    out = pd.DataFrame(rows)
    out.to_csv(TABLES / "TRO_reset_depth_summary.tsv", sep="\t", index=False)
    return out


def plot_outputs(state, trans):
    try:
        import matplotlib.pyplot as plt
    except ImportError:
        return
    FIGS.mkdir(parents=True, exist_ok=True)

    x = np.arange(len(trans))
    plt.figure(figsize=(10, 5))
    plt.bar(x, trans["transition_cost"], color="#6baed6")
    plt.xticks(x, trans["transition"], rotation=35, ha="right")
    plt.ylabel("State transition cost")
    plt.xlabel("")
    plt.tight_layout()
    plt.savefig(FIGS / "TRO_stage_transition_cost.png", dpi=300)
    plt.savefig(FIGS / "TRO_stage_transition_cost.pdf")

    plt.figure(figsize=(10, 5))
    plt.plot(x, trans["damage_reduction"], marker="o", label="Damage reduction")
    plt.plot(x, trans["potency_change"], marker="o", label="Potency change")
    plt.plot(x, trans["reset_gain"], marker="o", label="Reset gain")
    plt.axhline(0, color="gray", linewidth=1)
    plt.xticks(x, trans["transition"], rotation=35, ha="right")
    plt.ylabel("Transition component change")
    plt.xlabel("")
    plt.legend(frameon=False)
    plt.tight_layout()
    plt.savefig(FIGS / "TRO_transition_component_changes.png", dpi=300)
    plt.savefig(FIGS / "TRO_transition_component_changes.pdf")

    plt.figure(figsize=(7, 5.5))
    plt.scatter(state["damage_proxy"], state["potency_proxy"], s=90)
    for i in range(len(state) - 1):
        plt.annotate(
            "",
            xy=(state.loc[i + 1, "damage_proxy"], state.loc[i + 1, "potency_proxy"]),
            xytext=(state.loc[i, "damage_proxy"], state.loc[i, "potency_proxy"]),
            arrowprops=dict(arrowstyle="->", color="0.45", lw=1.2),
        )
    for _, r in state.iterrows():
        label = str(r["stage"]).replace("MII oocyte", "MII").replace("zygote/PN", "zygote")
        plt.annotate(label, (r["damage_proxy"], r["potency_proxy"]), xytext=(5, 5), textcoords="offset points", fontsize=8)
    morula = state[state["stage"].astype(str) == "morula"].iloc[0]
    plt.scatter([morula["damage_proxy"]], [morula["potency_proxy"]], s=190, facecolors="none", edgecolors="red", linewidth=2)
    plt.xlabel("Damage proxy (S_epi-age)")
    plt.ylabel("Potency proxy")
    plt.tight_layout()
    plt.savefig(FIGS / "TRO_damage_potency_state_space.png", dpi=300)
    plt.savefig(FIGS / "TRO_damage_potency_state_space.pdf")


def main():
    ap = argparse.ArgumentParser()
    args = ap.parse_args()
    TABLES.mkdir(parents=True, exist_ok=True)
    FIGS.mkdir(parents=True, exist_ok=True)
    df = load_state_table()
    state = build_state_vectors(df)
    trans = transition_costs(state)
    depth = reset_depth_summary(state, trans)
    plot_outputs(state, trans)

    print("TRO stage state vectors:")
    print(state[["stage", "damage_proxy", "potency_proxy", "reset_proxy", "bio_age_score", "bio_youth_score", "bio_age_rank", "TRO_score"]].to_string(index=False))
    print("\nTRO stage transition cost:")
    print(trans[["transition", "transition_cost", "damage_reduction", "potency_change", "reset_gain", "productive_reset_gain", "reset_efficiency", "cost_rank", "efficiency_rank"]].to_string(index=False))
    print("\nReset depth summary:")
    print(depth.to_string(index=False))
    print("\nWrote:", TABLES / "TRO_stage_transition_cost.tsv")
    print("Wrote:", TABLES / "TRO_reset_depth_summary.tsv")
    print("Wrote:", FIGS / "TRO_stage_transition_cost.png")


if __name__ == "__main__":
    main()
```
