## Supplemental code for "An entropy-based diagnostic framework for characterizing methylation state dynamics during preimplantation development": 18_experiment6_full_operational_tro_operator.py

````
from pathlib import Path
import argparse
import json

import numpy as np
import pandas as pd


STAGE_ORDER = ["MII oocyte", "zygote/PN", "2-cell", "4-cell", "8-cell", "morula", "blastocyst"]


def project_root():
    here = Path(__file__).resolve()
    server_root = Path("/root/autodl-tmp/TRO_Project")
    if server_root.exists() and str(here).startswith("/root/"):
        return server_root
    return here.parents[1]


ROOT = project_root()
TABLES = ROOT / "results" / "tables" if (ROOT / "results" / "tables").exists() else ROOT / "tables"
FIGS = ROOT / "results" / "figures" if (ROOT / "results" / "figures").exists() else ROOT / "figures"
NOTES = ROOT / "notes"


def read_table(name):
    path = TABLES / name
    if not path.exists():
        raise FileNotFoundError(path)
    return pd.read_csv(path, sep="\t")


def pass_fail(condition):
    return "pass" if bool(condition) else "check"


def build_stage_output():
    stage = read_table("TRO_stage_state_vectors.tsv")
    score = read_table("TRO_composite_score_by_stage.tsv")
    cols_from_score = ["stage", "GZ_score", "TRO_score", "GZ_rank", "TRO_rank", "PotencyPreserve"]
    merged = stage.merge(score[cols_from_score], on="stage", how="left", suffixes=("", "_score"))
    if "TRO_score_score" in merged.columns:
        merged["TRO_score"] = merged["TRO_score_score"]
    if "GZ_score_score" in merged.columns:
        merged["GZ_score"] = merged["GZ_score_score"]
    if "GZ_rank_score" in merged.columns:
        merged["GZ_rank"] = merged["GZ_rank_score"]
    if "TRO_rank_score" in merged.columns:
        merged["TRO_rank"] = merged["TRO_rank_score"]

    out = pd.DataFrame(
        {
            "stage": merged["stage"].astype(str),
            "E_S_epi": merged["S_epi"],
            "E_S_epi_age": merged["S_epi_age"] if "S_epi_age" in merged.columns else merged["damage_proxy"],
            "E_S_RNA": merged["S_RNA"],
            "D_DamageProxy": merged["damage_proxy"],
            "D_PotencyProxy": merged["potency_proxy"],
            "D_RNAOrderProxy": merged["rna_order_proxy"],
            "R_ResetScore": merged["reset_proxy"],
            "GZ_score": merged["GZ_score"],
            "TRO_score": merged["TRO_score"],
            "PotencyPreserve": merged["PotencyPreserve"],
            "BioAgeScore": merged["bio_age_score"],
            "BioYouthScore": merged["bio_youth_score"],
            "BioAgeRank": merged["bio_age_rank"],
            "GZ_rank": merged["GZ_rank"],
            "TRO_rank": merged["TRO_rank"],
            "operator_decision": np.where(
                (merged["GZ_rank"] == 1) & (merged["TRO_rank"] == 1),
                "computational_ground_zero",
                "non_ground_zero_state",
            ),
        }
    )
    out.to_csv(TABLES / "TRO_operator_stage_output.tsv", sep="\t", index=False)
    return out


def build_transition_output():
    trans = read_table("TRO_stage_transition_cost.tsv")
    out = pd.DataFrame(
        {
            "transition": trans["transition"],
            "stage_from": trans["stage_from"],
            "stage_to": trans["stage_to"],
            "C_transition_cost": trans["transition_cost"],
            "C_cost_rank": trans["cost_rank"],
            "R_damage_reduction": trans["damage_reduction"],
            "R_potency_change": trans["potency_change"],
            "R_reset_gain": trans["reset_gain"],
            "R_productive_reset_gain": trans["productive_reset_gain"],
            "R_reset_efficiency": trans["reset_efficiency"],
            "R_efficiency_rank": trans["efficiency_rank"],
        }
    )
    out["transition_decision"] = np.where(
        out["R_efficiency_rank"] == 1,
        "maximum_productive_reset_transition",
        np.where(out["R_productive_reset_gain"] <= 0, "nonproductive_or_differentiation_transition", "intermediate_transition"),
    )
    out.to_csv(TABLES / "TRO_operator_transition_output.tsv", sep="\t", index=False)
    return out


def collect_evidence(stage_out, trans_out):
    dna_summary = read_table("Experiment1B_DNA_robustness_summary.tsv")
    rna_tests = read_table("GSE36552_potency_pairwise_tests.tsv")
    loo = read_table("marker_leave_one_out_summary.tsv")
    ext = read_table("GSE44183_external_potency_validation.tsv")
    depth = read_table("TRO_reset_depth_summary.tsv")

    morula = stage_out[stage_out["stage"] == "morula"].iloc[0]
    best_transition = trans_out.sort_values("R_reset_efficiency", ascending=False).iloc[0]

    marker_mb = rna_tests[(rna_tests["metric"] == "marker_score") & (rna_tests["comparison"] == "morula vs blastocyst")].iloc[0]
    potency_mb = rna_tests[(rna_tests["metric"] == "potency_score") & (rna_tests["comparison"] == "morula vs blastocyst")].iloc[0]
    loo_pot = loo[loo["metric"] == "potency_score_recomputed"].copy()
    ext_sorted = ext.sort_values("potency_rank")
    balanced_bootstrap = dna_summary[dna_summary["test"].astype(str).str.contains("balanced bootstrap", na=False)]

    evidence = {
        "ground_zero_stage": str(morula["stage"]),
        "stage_decision": str(morula["operator_decision"]),
        "morula_GZ_rank": int(morula["GZ_rank"]),
        "morula_TRO_rank": int(morula["TRO_rank"]),
        "morula_BioAgeRank": int(morula["BioAgeRank"]),
        "morula_TRO_score": float(morula["TRO_score"]),
        "morula_GZ_score": float(morula["GZ_score"]),
        "best_transition": str(best_transition["transition"]),
        "best_transition_decision": str(best_transition["transition_decision"]),
        "best_transition_reset_efficiency": float(best_transition["R_reset_efficiency"]),
        "dna_common_DMR_pass": bool(((dna_summary["test"] == "common DMR") & (dna_summary["ground_zero_stage"] == "morula")).any()),
        "dna_balanced_bootstrap_pass": bool(
            len(balanced_bootstrap) > 0
            and (balanced_bootstrap["ground_zero_stage"] == "morula").all()
            and (balanced_bootstrap["conclusion"] == "pass").all()
        ),
        "rna_morula_vs_blastocyst_marker_BH_p": float(marker_mb["p_adj_BH"]),
        "rna_morula_vs_blastocyst_potency_BH_p": float(potency_mb["p_adj_BH"]),
        "leave_one_marker_out_all_pass": bool((loo_pot["conclusion"] == "pass").all()),
        "leave_one_marker_out_max_BH_p": float(np.nanmax(loo_pot["morula_vs_blastocyst_p_adj_BH"])),
        "external_GSE44183_top_potency_stages": ext_sorted["stage"].head(2).astype(str).tolist(),
        "external_GSE44183_morula_potency_rank": int(ext.loc[ext["stage"] == "morula", "potency_rank"].iloc[0]),
        "reset_depth_MII_to_morula_relative_S_epi_age_reduction": float(
            depth.loc[depth["to_stage"] == "morula", "relative_S_epi_age_reduction"].iloc[0]
        ),
    }

    checks = {
        "GZ_score_morula_rank_1": evidence["morula_GZ_rank"] == 1,
        "TRO_score_morula_rank_1": evidence["morula_TRO_rank"] == 1,
        "BioAgeScore_morula_rank_1": evidence["morula_BioAgeRank"] == 1,
        "best_transition_is_8cell_to_morula": evidence["best_transition"] == "8-cell -> morula",
        "DNA_robustness_common_DMR": evidence["dna_common_DMR_pass"],
        "DNA_robustness_balanced_bootstrap": evidence["dna_balanced_bootstrap_pass"],
        "RNA_potency_morula_gt_blastocyst": evidence["rna_morula_vs_blastocyst_potency_BH_p"] < 0.05,
        "marker_leave_one_out_all_pass": evidence["leave_one_marker_out_all_pass"],
        "external_GSE44183_morula_top2": evidence["external_GSE44183_morula_potency_rank"] <= 2,
    }
    evidence["checks"] = checks
    evidence["all_core_checks_pass"] = all(checks.values())
    return evidence


def write_schema(evidence):
    NOTES.mkdir(parents=True, exist_ok=True)
    text = f"""# Operational Transgenerational Reset Operator (TRO) schema

## Definition

The operational TRO is defined as:

```text
TRO = {{E, D, R, C}}
```

where:

- `E`: Entropy Encoder.
- `D`: Damage-Potency Decomposer.
- `R`: Reset Operator.
- `C`: Cost Estimator.

## E: Entropy Encoder

Input: stage-level multi-omics state.

Output:

```text
E(g) = [S_epi, S_epi-age, S_RNA]
```

`S_epi-age` is used as the age-associated methylation damage proxy.

## D: Damage-Potency Decomposer

Output:

```text
D(g) = [DamageProxy(g), PotencyProxy(g), RNAOrderProxy(g)]
```

where:

```text
DamageProxy(g) = S_epi-age(g)
PotencyProxy(g) = PotencyScore(g)
RNAOrderProxy(g) = -S_RNA(g)
```

## R: Reset Operator

Output:

```text
R(g) = ResetScore(g)
```

and transition-level productive reset gain:

```text
ProductiveResetGain(g -> h)
  = max(0, DamageReduction)
  + max(0, PotencyChange)
  + max(0, ResetGain)
```

## C: Cost Estimator

State transition cost is defined on standardized state vectors:

```text
z(g) = [z(DamageProxy), z(PotencyProxy), z(ResetScore), z(RNAOrderProxy)]
C(g -> h) = ||z(h) - z(g)||_2
```

Reset efficiency:

```text
ResetEfficiency(g -> h) = ProductiveResetGain(g -> h) / C(g -> h)
```

## Ground-Zero Decision

A stage is classified as computational ground zero when it ranks first by both:

```text
GZ_score = Z[-S_epi-age] + Z[PotencyScore]
TRO_score = ResetScore * PotencyPreserve
```

Current result:

```text
ground_zero_stage = {evidence["ground_zero_stage"]}
GZ_rank = {evidence["morula_GZ_rank"]}
TRO_rank = {evidence["morula_TRO_rank"]}
BioAgeRank = {evidence["morula_BioAgeRank"]}
best_transition = {evidence["best_transition"]}
all_core_checks_pass = {evidence["all_core_checks_pass"]}
```

## Interpretation

The completed operational TRO identifies morula as the computational ground-zero state and identifies the 8-cell to morula transition as the most efficient productive reset transition.
"""
    path = NOTES / "TRO_operator_schema.md"
    path.write_text(text, encoding="utf-8")
    return path


def plot_operator(stage_out, trans_out):
    try:
        import matplotlib.pyplot as plt
    except ImportError:
        return
    FIGS.mkdir(parents=True, exist_ok=True)
    fig, ax = plt.subplots(figsize=(9, 5.2))
    ax.axis("off")
    boxes = {
        "E\nEntropy Encoder\nS_epi-age, S_RNA": (0.12, 0.62),
        "D\nDamage-Potency\nDamageProxy, PotencyProxy": (0.38, 0.62),
        "R\nReset Operator\nResetScore, ResetGain": (0.64, 0.62),
        "C\nCost Estimator\nTransitionCost, Efficiency": (0.38, 0.25),
        "Decision\nmorula = ground-zero\n8-cell -> morula = reset transition": (0.68, 0.25),
    }
    for label, (x, y) in boxes.items():
        ax.text(
            x,
            y,
            label,
            ha="center",
            va="center",
            fontsize=10,
            bbox=dict(boxstyle="round,pad=0.45", facecolor="#f0f4f8", edgecolor="#4c78a8", linewidth=1.5),
        )
    arrows = [
        ((0.22, 0.62), (0.30, 0.62)),
        ((0.48, 0.62), (0.56, 0.62)),
        ((0.64, 0.55), (0.45, 0.32)),
        ((0.50, 0.25), (0.58, 0.25)),
    ]
    for xytext, xy in arrows:
        ax.annotate("", xy=xy, xytext=xytext, arrowprops=dict(arrowstyle="->", lw=1.5, color="#4c78a8"))
    ax.set_title("Operational Transgenerational Reset Operator (TRO)", fontsize=13, pad=18)
    plt.tight_layout()
    plt.savefig(FIGS / "TRO_operator_diagram.png", dpi=300)
    plt.savefig(FIGS / "TRO_operator_diagram.pdf")


def main():
    ap = argparse.ArgumentParser()
    args = ap.parse_args()
    TABLES.mkdir(parents=True, exist_ok=True)
    FIGS.mkdir(parents=True, exist_ok=True)

    stage_out = build_stage_output()
    trans_out = build_transition_output()
    evidence = collect_evidence(stage_out, trans_out)

    with open(TABLES / "TRO_operator_summary.json", "w", encoding="utf-8") as fh:
        json.dump(evidence, fh, ensure_ascii=False, indent=2)
    schema_path = write_schema(evidence)
    plot_operator(stage_out, trans_out)

    print("Operational TRO stage output:")
    print(stage_out.to_string(index=False))
    print("\nOperational TRO transition output:")
    print(trans_out.to_string(index=False))
    print("\nTRO evidence summary:")
    print(json.dumps(evidence, ensure_ascii=False, indent=2))
    print("\nWrote:", TABLES / "TRO_operator_stage_output.tsv")
    print("Wrote:", TABLES / "TRO_operator_transition_output.tsv")
    print("Wrote:", TABLES / "TRO_operator_summary.json")
    print("Wrote:", schema_path)
    print("Wrote:", FIGS / "TRO_operator_diagram.png")
    print("Wrote:", FIGS / "TRO_operator_diagram.pdf")


if __name__ == "__main__":
    main()
````
