## Supplemental code for "An entropy-based diagnostic framework for characterizing methylation state dynamics during preimplantation development": 19_experiment7_age_dmr_specificity_matched_control.py

```
from pathlib import Path
import argparse
import gzip
import json
import os
import time
from bisect import bisect_left, bisect_right

import numpy as np
import pandas as pd
import requests


STAGE_ORDER = [
    "MII oocyte",
    "zygote/PN",
    "2-cell",
    "4-cell",
    "8-cell",
    "morula",
    "blastocyst",
    "ICM",
    "TE",
]

HG19_CHR_SIZES = {
    "chr1": 249250621,
    "chr2": 243199373,
    "chr3": 198022430,
    "chr4": 191154276,
    "chr5": 180915260,
    "chr6": 171115067,
    "chr7": 159138663,
    "chr8": 146364022,
    "chr9": 141213431,
    "chr10": 135534747,
    "chr11": 135006516,
    "chr12": 133851895,
    "chr13": 115169878,
    "chr14": 107349540,
    "chr15": 102531392,
    "chr16": 90354753,
    "chr17": 81195210,
    "chr18": 78077248,
    "chr19": 59128983,
    "chr20": 63025520,
    "chr21": 48129895,
    "chr22": 51304566,
    "chrX": 155270560,
    "chrY": 59373566,
}


def project_root():
    here = Path(__file__).resolve()
    server_root = Path("/root/autodl-tmp/TRO_Project")
    if server_root.exists() and str(here).startswith("/root/"):
        return server_root
    return here.parents[1]


ROOT = project_root()
TABLES = ROOT / "results" / "tables" if (ROOT / "results" / "tables").exists() else ROOT / "tables"
FIGS = ROOT / "results" / "figures" if (ROOT / "results" / "figures").exists() else ROOT / "figures"
NOTES = ROOT / "notes"
DMR_PATH = (
    ROOT / "data_processed" / "metadata" / "GSE102970_TableS6_age_dmr_weights.tsv"
    if (ROOT / "data_processed" / "metadata" / "GSE102970_TableS6_age_dmr_weights.tsv").exists()
    else ROOT / "metadata" / "GSE102970_TableS6_age_dmr_weights.tsv"
)
DEFAULT_MANIFEST = TABLES / "GSE81233_valid_cmet_manifest_204.tsv"
LOCAL_CMET_DIR = ROOT / "data_raw" / "GSE81233_embryo_methylation" / "pilot_cmet"
OUTDIR = ROOT / "data_processed" / "methylation_matrix" / "GSE81233_matched_non_age_dmr_controls"


def binary_entropy(p, eps=1e-6):
    p = np.clip(np.asarray(p, dtype=float), eps, 1 - eps)
    return -(p * np.log(p) + (1 - p) * np.log(1 - p))


def load_age_dmr():
    dmr = pd.read_csv(DMR_PATH, sep="\t")
    dmr = dmr.dropna(subset=["cluster_name", "chr", "start", "end", "age_weight_5yr"]).copy()
    dmr["start"] = dmr["start"].astype(int)
    dmr["end"] = dmr["end"].astype(int)
    dmr["age_weight_5yr"] = pd.to_numeric(dmr["age_weight_5yr"], errors="coerce")
    dmr = dmr.dropna(subset=["age_weight_5yr"]).reset_index(drop=True)
    dmr["width"] = dmr["end"] - dmr["start"] + 1
    return dmr


def overlaps_any(chrom, start, end, forbidden):
    intervals = forbidden.get(chrom, [])
    for s, e in intervals:
        if start <= e and end >= s:
            return True
    return False


def make_matched_controls(age_dmr, n_controls=20, seed=777, max_tries=2000):
    rng = np.random.default_rng(seed)
    forbidden = {}
    for _, r in age_dmr.iterrows():
        forbidden.setdefault(str(r["chr"]), []).append((int(r["start"]), int(r["end"])))

    rows = []
    for control_idx in range(int(n_controls)):
        used = {chrom: list(vals) for chrom, vals in forbidden.items()}
        for age_idx, r in age_dmr.iterrows():
            chrom = str(r["chr"])
            width = int(r["width"])
            chrom_size = HG19_CHR_SIZES.get(chrom)
            if chrom_size is None or chrom_size <= width + 1000:
                raise ValueError(f"Unsupported chromosome for matching: {chrom}")

            chosen = None
            for _ in range(max_tries):
                start = int(rng.integers(1, chrom_size - width))
                end = start + width - 1
                if not overlaps_any(chrom, start, end, used):
                    chosen = (start, end)
                    used.setdefault(chrom, []).append(chosen)
                    break
            if chosen is None:
                raise RuntimeError(f"Could not place matched control for {chrom}:{r['start']}-{r['end']}")

            rows.append(
                {
                    "control_set": f"matched_non_age_{control_idx:03d}",
                    "matched_age_cluster": r["cluster_name"],
                    "control_cluster": f"matched_non_age_{control_idx:03d}__{r['cluster_name']}",
                    "chr": chrom,
                    "start": chosen[0],
                    "end": chosen[1],
                    "width": width,
                    "age_weight_5yr": float(r["age_weight_5yr"]),
                }
            )
    return pd.DataFrame(rows)


def build_interval_index(region_df):
    intervals, starts, max_width = {}, {}, {}
    for idx, r in region_df.iterrows():
        cb = str(r["chr"]).encode()
        intervals.setdefault(cb, []).append((int(r["start"]), int(r["end"]), idx))
    for cb in intervals:
        intervals[cb].sort()
        starts[cb] = [x[0] for x in intervals[cb]]
        max_width[cb] = max(e - s + 1 for s, e, _ in intervals[cb])
    return intervals, starts, max_width


def sample_id_from_filename(fn):
    return str(fn).split("_", 1)[0]


def open_cmet_stream(row):
    filename = row["filename"]
    local_path = LOCAL_CMET_DIR / filename
    expected = int(row["size_bytes"]) if not pd.isna(row.get("size_bytes", np.nan)) else None
    if local_path.exists() and (expected is None or local_path.stat().st_size >= expected * 0.98):
        return gzip.open(local_path, "rb"), f"local:{local_path}"

    r = requests.get(row["url"], stream=True, timeout=(30, 120))
    r.raise_for_status()
    r.raw.decode_content = False
    return gzip.GzipFile(fileobj=r.raw), f"remote:{row['url']}"


def process_sample(row, regions, intervals, starts, max_width, outdir, force=False, progress_lines=1000000):
    sample_id = row["sample_id"] if "sample_id" in row and not pd.isna(row["sample_id"]) else sample_id_from_filename(row["filename"])
    out = outdir / "sample_matched_dmr" / f"{sample_id}.matched_dmr.tsv"
    tmp = out.with_suffix(".tmp")
    out.parent.mkdir(parents=True, exist_ok=True)
    if out.exists() and not force:
        print(f"SKIP {sample_id}: exists", flush=True)
        return out

    met = np.zeros(len(regions), dtype=np.float64)
    total = np.zeros(len(regions), dtype=np.float64)
    sites = np.zeros(len(regions), dtype=np.int32)

    n_lines = n_cg = n_match = 0
    t0 = time.time()
    fh, source = open_cmet_stream(row)
    print(f"START {sample_id} stage={row['stage']} source={source}", flush=True)

    with fh:
        for raw in fh:
            if not raw or raw.startswith(b"#"):
                continue
            n_lines += 1
            parts = raw.rstrip(b"\n").split(b"\t")
            if len(parts) < 10:
                continue
            typ = parts[9].strip()
            if typ not in (b"CG", b"CpG", b"cg", b"cpg"):
                continue
            n_cg += 1
            cb = parts[0]
            if cb not in intervals:
                continue
            try:
                pos = int(parts[1])
                tot = int(parts[4])
                m = int(parts[5])
            except Exception:
                continue
            if tot <= 0:
                continue

            left = bisect_left(starts[cb], pos - max_width[cb])
            right = bisect_right(starts[cb], pos)
            for k in range(left, right):
                s, e, idx = intervals[cb][k]
                if s <= pos <= e:
                    met[idx] += m
                    total[idx] += tot
                    sites[idx] += 1
                    n_match += 1

            if progress_lines > 0 and n_lines % int(progress_lines) == 0:
                print(
                    f"PROGRESS {sample_id}: lines={n_lines} CG={n_cg} matched={n_match} elapsed_min={(time.time()-t0)/60:.1f}",
                    flush=True,
                )

    beta = np.where(total > 0, met / total, np.nan)
    res = regions[
        ["region_type", "control_set", "cluster_name", "matched_age_cluster", "chr", "start", "end", "width", "age_weight_5yr"]
    ].copy()
    res.insert(0, "sample_id", sample_id)
    res.insert(1, "stage", row["stage"])
    res["met_reads"] = met
    res["total_reads"] = total
    res["n_cpg_rows_covered"] = sites
    res["beta"] = beta
    res.to_csv(tmp, sep="\t", index=False)
    os.replace(tmp, out)
    print(
        f"DONE {sample_id}: CG={n_cg} matched={n_match} covered_regions={(total > 0).sum()} elapsed_min={(time.time()-t0)/60:.1f}",
        flush=True,
    )
    return out


def aggregate_stage_metrics(long_df, min_sample_frac=0.30):
    rows = []
    for (region_type, control_set), df in long_df.groupby(["region_type", "control_set"], observed=True):
        for stage in STAGE_ORDER:
            sub = df[df["stage"] == stage]
            n_samples = sub["sample_id"].nunique()
            if n_samples == 0:
                continue
            g = sub.groupby("cluster_name", as_index=False).agg(
                met_reads=("met_reads", "sum"),
                total_reads=("total_reads", "sum"),
                n_samples_covered=("total_reads", lambda x: int((x > 0).sum())),
                age_weight_5yr=("age_weight_5yr", "first"),
            )
            min_cov = max(1, int(np.ceil(float(min_sample_frac) * n_samples)))
            valid = g[(g["total_reads"] > 0) & (g["n_samples_covered"] >= min_cov)].copy()
            if len(valid) == 0:
                s_epi = s_epi_age = np.nan
            else:
                p = valid["met_reads"] / valid["total_reads"]
                h = binary_entropy(p)
                w = valid["age_weight_5yr"].astype(float).abs()
                s_epi = float(np.mean(h))
                s_epi_age = float(np.sum(w * h) / np.sum(w)) if np.sum(w) > 0 else np.nan
            rows.append(
                {
                    "region_type": region_type,
                    "control_set": control_set,
                    "stage": stage,
                    "n_samples": n_samples,
                    "n_regions_valid": len(valid),
                    "min_samples_required": min_cov,
                    "s_epi": s_epi,
                    "s_epi_age_weighted": s_epi_age,
                }
            )
    out = pd.DataFrame(rows)
    out["stage"] = pd.Categorical(out["stage"], categories=STAGE_ORDER, ordered=True)
    return out.sort_values(["region_type", "control_set", "stage"]).reset_index(drop=True)


def summarize_specificity(stage_metrics):
    rows = []
    for (region_type, control_set), df in stage_metrics.groupby(["region_type", "control_set"], observed=True):
        ok = df.dropna(subset=["s_epi_age_weighted"]).copy()
        if ok.empty:
            continue
        ground = ok.sort_values("s_epi_age_weighted").iloc[0]
        morula = ok[ok["stage"] == "morula"].iloc[0]
        non_morula_min = ok[ok["stage"] != "morula"]["s_epi_age_weighted"].min()
        mii = ok[ok["stage"] == "MII oocyte"]["s_epi_age_weighted"]
        reset_depth = float(mii.iloc[0] - morula["s_epi_age_weighted"]) if len(mii) else np.nan
        rows.append(
            {
                "region_type": region_type,
                "control_set": control_set,
                "ground_zero_stage": str(ground["stage"]),
                "morula_is_minimum": str(ground["stage"]) == "morula",
                "morula_s_epi_age_weighted": float(morula["s_epi_age_weighted"]),
                "morula_gap_to_next_lowest": float(non_morula_min - morula["s_epi_age_weighted"]),
                "MII_to_morula_reset_depth": reset_depth,
                "morula_n_regions_valid": int(morula["n_regions_valid"]),
            }
        )
    summary = pd.DataFrame(rows)

    age = summary[summary["region_type"] == "age_DMR"].iloc[0]
    controls = summary[summary["region_type"] == "matched_non_age_DMR"].copy()
    p_gap = (1 + int((controls["morula_gap_to_next_lowest"] >= age["morula_gap_to_next_lowest"]).sum())) / (len(controls) + 1)
    p_depth = (1 + int((controls["MII_to_morula_reset_depth"] >= age["MII_to_morula_reset_depth"]).sum())) / (len(controls) + 1)
    comparison = {
        "age_DMR_ground_zero_stage": str(age["ground_zero_stage"]),
        "age_DMR_morula_gap_to_next_lowest": float(age["morula_gap_to_next_lowest"]),
        "age_DMR_MII_to_morula_reset_depth": float(age["MII_to_morula_reset_depth"]),
        "matched_control_n": int(len(controls)),
        "matched_control_morula_min_frequency": float(controls["morula_is_minimum"].mean()),
        "matched_control_morula_gap_mean": float(controls["morula_gap_to_next_lowest"].mean()),
        "matched_control_reset_depth_mean": float(controls["MII_to_morula_reset_depth"].mean()),
        "empirical_p_control_gap_ge_age_gap": float(p_gap),
        "empirical_p_control_depth_ge_age_depth": float(p_depth),
    }
    comparison["conclusion"] = (
        "age_DMR_specificity_supported"
        if p_gap <= 0.10 or p_depth <= 0.10
        else "morula_minimum_broader_methylation_reprogramming"
    )
    return summary, comparison


def plot_specificity(summary, comparison):
    try:
        import matplotlib.pyplot as plt
    except ImportError:
        print("matplotlib is not installed; skipping figure.")
        return

    FIGS.mkdir(parents=True, exist_ok=True)
    controls = summary[summary["region_type"] == "matched_non_age_DMR"].copy()
    age = summary[summary["region_type"] == "age_DMR"].iloc[0]

    fig, axes = plt.subplots(1, 2, figsize=(11, 4.5))
    axes[0].hist(controls["morula_gap_to_next_lowest"], bins=12, color="#a6bddb", edgecolor="white")
    axes[0].axvline(age["morula_gap_to_next_lowest"], color="#d95f02", linewidth=2, label="age-DMR")
    axes[0].set_xlabel("Morula gap to next-lowest stage")
    axes[0].set_ylabel("Matched control sets")
    axes[0].legend(frameon=False)

    axes[1].hist(controls["MII_to_morula_reset_depth"], bins=12, color="#b2df8a", edgecolor="white")
    axes[1].axvline(age["MII_to_morula_reset_depth"], color="#d95f02", linewidth=2, label="age-DMR")
    axes[1].set_xlabel("MII-to-morula reset depth")
    axes[1].set_ylabel("Matched control sets")
    axes[1].legend(frameon=False)

    fig.suptitle(f"Matched non-age-DMR specificity control: {comparison['conclusion']}")
    plt.tight_layout()
    plt.savefig(FIGS / "Experiment7_age_DMR_specificity_matched_control.png", dpi=300)
    plt.savefig(FIGS / "Experiment7_age_DMR_specificity_matched_control.pdf")
    plt.close()


def write_note(comparison):
    NOTES.mkdir(parents=True, exist_ok=True)
    text = f"""# Experiment 7: age-DMR specificity matched-control

## Purpose

This experiment tests whether the morula minimum in age-associated methylation entropy is specific to sperm age-DMR regions, or whether similar morula minima are commonly observed in matched non-age-DMR genomic windows.

## Design

For each age-DMR, matched non-age-DMR windows were sampled on the same chromosome with the same genomic width. Age-DMR weights were transferred to the matched windows to keep the weight distribution fixed. GSE81233 Cmet files were then rescanned to compute stage-level weighted methylation entropy.

## Main comparison

- age_DMR_ground_zero_stage = {comparison['age_DMR_ground_zero_stage']}
- age_DMR_morula_gap_to_next_lowest = {comparison['age_DMR_morula_gap_to_next_lowest']:.6f}
- age_DMR_MII_to_morula_reset_depth = {comparison['age_DMR_MII_to_morula_reset_depth']:.6f}
- matched_control_n = {comparison['matched_control_n']}
- matched_control_morula_min_frequency = {comparison['matched_control_morula_min_frequency']:.4f}
- matched_control_morula_gap_mean = {comparison['matched_control_morula_gap_mean']:.6f}
- matched_control_reset_depth_mean = {comparison['matched_control_reset_depth_mean']:.6f}
- empirical_p_control_gap_ge_age_gap = {comparison['empirical_p_control_gap_ge_age_gap']:.4f}
- empirical_p_control_depth_ge_age_depth = {comparison['empirical_p_control_depth_ge_age_depth']:.4f}

## Conclusion

{comparison['conclusion']}

If the conclusion is age_DMR_specificity_supported, the age-DMR signal is stronger than matched non-age controls. If the conclusion is morula_minimum_broader_methylation_reprogramming, the correct interpretation is that age-DMR weighted entropy captures a broader methylation reprogramming window at morula rather than a strictly age-DMR-only phenomenon.
"""
    path = NOTES / "Experiment7_age_DMR_specificity_matched_control_interpretation.md"
    path.write_text(text, encoding="utf-8")
    return path


def main():
    ap = argparse.ArgumentParser()
    ap.add_argument("--manifest", default=str(DEFAULT_MANIFEST))
    ap.add_argument("--n-controls", type=int, default=20)
    ap.add_argument("--max-samples", type=int, default=None)
    ap.add_argument("--samples-per-stage", type=int, default=None)
    ap.add_argument("--stages", default=None, help="Comma-separated stage list for pilot runs.")
    ap.add_argument("--min-sample-frac", type=float, default=0.30)
    ap.add_argument("--progress-lines", type=int, default=1000000)
    ap.add_argument("--force", action="store_true")
    ap.add_argument("--combine-only", action="store_true")
    args = ap.parse_args()

    OUTDIR.mkdir(parents=True, exist_ok=True)
    TABLES.mkdir(parents=True, exist_ok=True)

    manifest = pd.read_csv(args.manifest, sep="\t")
    manifest = manifest[manifest["stage"].isin(STAGE_ORDER)].copy()
    if args.stages:
        keep_stages = [x.strip() for x in args.stages.split(",") if x.strip()]
        manifest = manifest[manifest["stage"].isin(keep_stages)].copy()
    manifest["stage"] = pd.Categorical(manifest["stage"], categories=STAGE_ORDER, ordered=True)
    manifest["size_bytes"] = pd.to_numeric(manifest["size_bytes"], errors="coerce")
    manifest = manifest.sort_values(["stage", "size_bytes", "sample_id"]).reset_index(drop=True)
    if args.samples_per_stage:
        manifest = (
            manifest.groupby("stage", observed=True, group_keys=False)
            .head(int(args.samples_per_stage))
            .reset_index(drop=True)
        )
    if args.max_samples:
        manifest = manifest.head(int(args.max_samples))

    age_dmr = load_age_dmr()
    controls = make_matched_controls(age_dmr, n_controls=args.n_controls)
    age_regions = age_dmr.rename(columns={"cluster_name": "matched_age_cluster"}).copy()
    age_regions["region_type"] = "age_DMR"
    age_regions["control_set"] = "age_DMR"
    age_regions["cluster_name"] = age_regions["matched_age_cluster"]
    age_regions = age_regions[["region_type", "control_set", "cluster_name", "matched_age_cluster", "chr", "start", "end", "width", "age_weight_5yr"]]

    control_regions = controls.copy()
    control_regions["region_type"] = "matched_non_age_DMR"
    control_regions = control_regions.rename(columns={"control_cluster": "cluster_name"})
    control_regions = control_regions[
        ["region_type", "control_set", "cluster_name", "matched_age_cluster", "chr", "start", "end", "width", "age_weight_5yr"]
    ]
    regions = pd.concat([age_regions, control_regions], ignore_index=True)
    regions.to_csv(TABLES / "Experiment7_matched_non_age_DMR_regions.tsv", sep="\t", index=False)

    if not args.combine_only:
        intervals, starts, max_width = build_interval_index(regions)
        print(
            f"Experiment7 regions={len(regions)} age_regions={len(age_regions)} controls={args.n_controls} samples={len(manifest)}",
            flush=True,
        )
        for _, row in manifest.iterrows():
            process_sample(
                row,
                regions,
                intervals,
                starts,
                max_width,
                OUTDIR,
                force=args.force,
                progress_lines=args.progress_lines,
            )

    files = sorted((OUTDIR / "sample_matched_dmr").glob("*.matched_dmr.tsv"))
    if not files:
        raise RuntimeError("No sample matched-DMR files found.")
    long_df = pd.concat([pd.read_csv(f, sep="\t") for f in files], ignore_index=True)
    long_df = long_df[long_df["sample_id"].isin(set(manifest["sample_id"].astype(str)))].copy()
    long_df.to_csv(TABLES / "Experiment7_all_sample_matched_DMR_long.tsv.gz", sep="\t", index=False, compression="gzip")

    stage_metrics = aggregate_stage_metrics(long_df, min_sample_frac=args.min_sample_frac)
    summary, comparison = summarize_specificity(stage_metrics)

    stage_metrics.to_csv(TABLES / "Experiment7_age_DMR_specificity_stage_metrics.tsv", sep="\t", index=False)
    summary.to_csv(TABLES / "Experiment7_age_DMR_specificity_summary.tsv", sep="\t", index=False)
    with open(TABLES / "Experiment7_age_DMR_specificity_comparison.json", "w", encoding="utf-8") as fh:
        json.dump(comparison, fh, ensure_ascii=False, indent=2)
    note_path = write_note(comparison)
    plot_specificity(summary, comparison)

    print("Experiment 7 age-DMR specificity comparison:")
    print(json.dumps(comparison, ensure_ascii=False, indent=2))
    print("Wrote:", TABLES / "Experiment7_matched_non_age_DMR_regions.tsv")
    print("Wrote:", TABLES / "Experiment7_age_DMR_specificity_stage_metrics.tsv")
    print("Wrote:", TABLES / "Experiment7_age_DMR_specificity_summary.tsv")
    print("Wrote:", TABLES / "Experiment7_age_DMR_specificity_comparison.json")
    print("Wrote:", note_path)
    print("Wrote:", FIGS / "Experiment7_age_DMR_specificity_matched_control.png")
    print("Wrote:", FIGS / "Experiment7_age_DMR_specificity_matched_control.pdf")


if __name__ == "__main__":
    main()
```
