## Supplemental code for "An entropy-based diagnostic framework for characterizing methylation state dynamics during preimplantation development": 20_experiment8_final_claim_audit.py

```
from pathlib import Path
import json

import pandas as pd


def project_root():
    here = Path(__file__).resolve()
    server_root = Path("/root/autodl-tmp/TRO_Project")
    if server_root.exists() and str(here).startswith("/root/"):
        return server_root
    return here.parents[1]


ROOT = project_root()
TABLES = ROOT / "results" / "tables" if (ROOT / "results" / "tables").exists() else ROOT / "tables"
FIGS = ROOT / "results" / "figures" if (ROOT / "results" / "figures").exists() else ROOT / "figures"
NOTES = ROOT / "notes"


def read_table(name):
    return pd.read_csv(TABLES / name, sep="\t")


def read_json(name):
    return json.loads((TABLES / name).read_text(encoding="utf-8"))


def val(df, row_filter, col):
    hit = df.loc[row_filter]
    if hit.empty:
        raise ValueError(f"No row found for {col}")
    return hit.iloc[0][col]


def fmt(x, digits=4):
    if pd.isna(x):
        return "NA"
    if isinstance(x, str):
        return x
    return f"{float(x):.{digits}g}"


def add_claim(rows, claim_id, claim, value, source_table, source_field, status="supported", note=""):
    rows.append(
        {
            "claim_id": claim_id,
            "claim": claim,
            "value": value,
            "source_table": source_table,
            "source_field": source_field,
            "status": status,
            "note": note,
        }
    )


def main():
    NOTES.mkdir(parents=True, exist_ok=True)
    TABLES.mkdir(parents=True, exist_ok=True)

    dna_stage = read_table("GSE81233_valid204_stage_epi_age_metrics.tsv")
    dna_boot = read_table("GSE81233_valid204_bootstrap_ground_zero_frequency.tsv")
    dna_adj = read_table("GSE81233_valid204_adjacent_stage_mannwhitney.tsv")
    dna_robust = read_table("Experiment1B_DNA_robustness_summary.tsv")
    rna_stage = read_table("GSE36552_RNA_entropy_potency_by_stage.tsv")
    rna_tests = read_table("GSE36552_potency_pairwise_tests.tsv")
    ext = read_table("GSE44183_external_potency_validation.tsv")
    tro_comp = read_table("TRO_composite_score_by_stage.tsv")
    trans = read_table("TRO_stage_transition_cost.tsv")
    reset_depth = read_table("TRO_reset_depth_summary.tsv")
    op_summary = read_json("TRO_operator_summary.json")
    op_stage = read_table("TRO_operator_stage_output.tsv")

    rows = []
    numbers = {}

    morula_dna = dna_stage.loc[dna_stage["stage"] == "morula"].iloc[0]
    dna_min = dna_stage.sort_values("s_epi_age").iloc[0]
    boot_morula = dna_boot.loc[dna_boot["stage"] == "morula"].iloc[0]
    numbers["morula_s_epi_age"] = float(morula_dna["s_epi_age"])
    numbers["morula_dna_n_samples"] = int(morula_dna["n_samples"])
    numbers["morula_bootstrap_frequency"] = float(boot_morula["frequency"])
    numbers["morula_bootstrap_n_min"] = int(boot_morula["n_min"])

    add_claim(
        rows,
        "DNA-1",
        "Morula has the lowest stage-level age-associated methylation entropy.",
        f"{dna_min['stage']} S_epi-age={fmt(dna_min['s_epi_age'], 6)}",
        "GSE81233_valid204_stage_epi_age_metrics.tsv",
        "s_epi_age",
        note="Primary DNA ground-zero claim.",
    )
    add_claim(
        rows,
        "DNA-2",
        "Morula is the most frequent bootstrap ground-zero stage.",
        f"{boot_morula['n_min']}/2000={fmt(boot_morula['frequency'], 4)}",
        "GSE81233_valid204_bootstrap_ground_zero_frequency.tsv",
        "frequency",
    )

    for comp in ["2-cell vs 4-cell", "8-cell vs morula", "morula vs blastocyst"]:
        row = dna_adj.loc[dna_adj["comparison"] == comp].iloc[0]
        key = comp.replace(" ", "_").replace("->", "to").replace("/", "_")
        numbers[f"dna_adjacent_{key}_p_adj_BH"] = float(row["p_adj_BH"])
        add_claim(
            rows,
            f"DNA-adj-{comp}",
            f"Adjacent DNA sample-level test is significant for {comp}.",
            f"BH p={fmt(row['p_adj_BH'], 4)}, Cliff delta={fmt(row['cliffs_delta_a_minus_b'], 4)}",
            "GSE81233_valid204_adjacent_stage_mannwhitney.tsv",
            "p_adj_BH",
        )

    common = dna_robust.loc[dna_robust["test"] == "common DMR"].iloc[0]
    b5 = dna_robust.loc[dna_robust["test"] == "balanced bootstrap n=5"].iloc[0]
    b8 = dna_robust.loc[dna_robust["test"] == "balanced bootstrap n=8"].iloc[0]
    numbers["common_DMR_morula_frequency"] = float(common["morula_frequency"])
    numbers["balanced_bootstrap_n5_morula_frequency"] = float(b5["morula_frequency"])
    numbers["balanced_bootstrap_n8_morula_frequency"] = float(b8["morula_frequency"])
    add_claim(
        rows,
        "DNA-robust-1",
        "Morula remains ground-zero under common-DMR robustness analysis.",
        f"stage={common['ground_zero_stage']}, freq={fmt(common['morula_frequency'], 4)}",
        "Experiment1B_DNA_robustness_summary.tsv",
        "common DMR",
    )
    add_claim(
        rows,
        "DNA-robust-2",
        "Morula remains ground-zero under balanced bootstrap.",
        f"n=5 freq={fmt(b5['morula_frequency'], 4)}; n=8 freq={fmt(b8['morula_frequency'], 4)}",
        "Experiment1B_DNA_robustness_summary.tsv",
        "balanced bootstrap",
    )

    morula_rna = rna_stage.loc[rna_stage["stage"] == "morula"].iloc[0]
    eight_rna = rna_stage.loc[rna_stage["stage"] == "8-cell"].iloc[0]
    blast_rna = rna_stage.loc[rna_stage["stage"] == "blastocyst"].iloc[0]
    numbers["GSE36552_morula_potency_score"] = float(morula_rna["potency_score_mean"])
    numbers["GSE36552_8cell_potency_score"] = float(eight_rna["potency_score_mean"])
    numbers["GSE36552_blastocyst_potency_score"] = float(blast_rna["potency_score_mean"])
    numbers["GSE36552_morula_s_rna"] = float(morula_rna["s_rna_mean"])
    add_claim(
        rows,
        "RNA-1",
        "Morula does not show the highest global RNA entropy.",
        f"morula S_RNA={fmt(morula_rna['s_rna_mean'], 6)}; oocyte S_RNA={fmt(rna_stage['s_rna_mean'].max(), 6)} max",
        "GSE36552_RNA_entropy_potency_by_stage.tsv",
        "s_rna_mean",
        status="boundary",
        note="This prevents overclaiming RNA entropy.",
    )
    add_claim(
        rows,
        "RNA-2",
        "8-cell and morula form a high potency-marker region.",
        f"8-cell={fmt(eight_rna['potency_score_mean'], 4)}; morula={fmt(morula_rna['potency_score_mean'], 4)}; blastocyst={fmt(blast_rna['potency_score_mean'], 4)}",
        "GSE36552_RNA_entropy_potency_by_stage.tsv",
        "potency_score_mean",
    )

    for metric in ["marker_score", "potency_score"]:
        row = rna_tests.loc[(rna_tests["metric"] == metric) & (rna_tests["comparison"] == "morula vs blastocyst")].iloc[0]
        numbers[f"GSE36552_morula_vs_blastocyst_{metric}_p_adj_BH"] = float(row["p_adj_BH"])
        add_claim(
            rows,
            f"RNA-morula-blastocyst-{metric}",
            f"Morula is higher than blastocyst for {metric}.",
            f"mean morula={fmt(row['mean_a'], 4)}; blastocyst={fmt(row['mean_b'], 4)}; BH p={fmt(row['p_adj_BH'], 4)}",
            "GSE36552_potency_pairwise_tests.tsv",
            "p_adj_BH",
        )

    ext_top = ext.sort_values("potency_rank").head(2)["stage"].tolist()
    ext_morula = ext.loc[ext["stage"] == "morula"].iloc[0]
    numbers["GSE44183_top_potency_stages"] = ext_top
    numbers["GSE44183_morula_potency_rank"] = int(ext_morula["potency_rank"])
    add_claim(
        rows,
        "EXT-RNA-1",
        "External GSE44183 places morula in the top two potency stages.",
        f"top2={','.join(ext_top)}; morula rank={int(ext_morula['potency_rank'])}",
        "GSE44183_external_potency_validation.tsv",
        "potency_rank",
    )

    tro_morula = tro_comp.loc[tro_comp["stage"] == "morula"].iloc[0]
    numbers["morula_TRO_score"] = float(tro_morula["TRO_score"])
    numbers["morula_GZ_score"] = float(tro_morula["GZ_score"])
    numbers["morula_TRO_rank"] = int(tro_morula["TRO_rank"])
    numbers["morula_GZ_rank"] = int(tro_morula["GZ_rank"])
    add_claim(
        rows,
        "TRO-1",
        "Morula ranks first by GZ score and TRO score.",
        f"GZ_rank={int(tro_morula['GZ_rank'])}; TRO_rank={int(tro_morula['TRO_rank'])}; TRO_score={fmt(tro_morula['TRO_score'], 4)}",
        "TRO_composite_score_by_stage.tsv",
        "GZ_rank,TRO_rank,TRO_score",
    )

    best_trans = trans.sort_values("efficiency_rank").iloc[0]
    numbers["best_transition"] = str(best_trans["transition"])
    numbers["best_transition_reset_efficiency"] = float(best_trans["reset_efficiency"])
    numbers["best_transition_productive_reset_gain"] = float(best_trans["productive_reset_gain"])
    add_claim(
        rows,
        "TRANS-1",
        "8-cell to morula is the maximum productive reset transition.",
        f"{best_trans['transition']}; efficiency={fmt(best_trans['reset_efficiency'], 4)}; productive_gain={fmt(best_trans['productive_reset_gain'], 4)}",
        "TRO_stage_transition_cost.tsv",
        "efficiency_rank,reset_efficiency",
    )

    depth_morula = reset_depth.loc[reset_depth["to_stage"] == "morula"].iloc[0]
    numbers["MII_to_morula_relative_S_epi_age_reduction"] = float(depth_morula["relative_S_epi_age_reduction"])
    add_claim(
        rows,
        "RESET-depth-1",
        "MII-to-morula S_epi-age reduction is about 37.8%.",
        f"relative reduction={fmt(depth_morula['relative_S_epi_age_reduction'] * 100, 4)}%",
        "TRO_reset_depth_summary.tsv",
        "relative_S_epi_age_reduction",
    )

    op_morula = op_stage.loc[op_stage["stage"] == "morula"].iloc[0]
    numbers["operator_ground_zero_stage"] = op_summary["ground_zero_stage"]
    numbers["operator_all_core_checks_pass"] = bool(op_summary["all_core_checks_pass"])
    add_claim(
        rows,
        "OP-1",
        "Operational TRO identifies morula as computational ground-zero.",
        f"decision={op_morula['operator_decision']}; all_core_checks_pass={op_summary['all_core_checks_pass']}",
        "TRO_operator_stage_output.tsv; TRO_operator_summary.json",
        "operator_decision; all_core_checks_pass",
    )

    audit = pd.DataFrame(rows)
    audit.to_csv(TABLES / "final_claim_audit.tsv", sep="\t", index=False)
    (TABLES / "manuscript_numbers.json").write_text(json.dumps(numbers, indent=2, ensure_ascii=False), encoding="utf-8")

    figure_plan = """# Figure panel plan

## Figure 1: DNA age-associated methylation entropy reset

- Figure 1A: `figures/GSE81233_valid204_s_epi_age_by_stage.png`
  - Claim: morula has the lowest S_epi-age.
  - Source: `tables/GSE81233_valid204_stage_epi_age_metrics.tsv`

- Figure 1B: `figures/GSE81233_valid204_sample_level_s_epi_age_boxplot.png`
  - Claim: sample-level distributions support stage differences.
  - Source: `tables/GSE81233_valid204_sample_level_entropy_metrics.tsv`

- Figure 1C: `figures/GSE81233_valid204_internal_reset_score.png`
  - Claim: internal reset score maps MII to 0 and morula to 1.
  - Source: `tables/GSE81233_valid204_internal_reset_score.tsv`

## Figure 2: DNA robustness and RNA potency

- Figure 2A: `figures/Experiment1B_DNA_robustness_summary.png`
  - Claim: morula remains robust under common-DMR and balanced bootstrap checks.
  - Source: `tables/Experiment1B_DNA_robustness_summary.tsv`

- Figure 2B: `figures/GSE36552_marker_heatmap.png`
  - Claim: 8-cell/morula retain high developmental potency-marker activity.
  - Source: `tables/GSE36552_marker_zscore_heatmap_matrix.tsv`

- Figure 2C: `figures/GSE44183_external_potency_validation.png`
  - Claim: external RNA dataset places 8-cell/morula in a high-potency region.
  - Source: `tables/GSE44183_external_potency_validation.tsv`

## Figure 3: Dual-state and TRO composite score

- Figure 3A: `figures/dual_entropy_phase_map_final.png`
  - Claim: morula lies in the low S_epi-age / high potency region.
  - Source: `tables/dual_entropy_stage_table.tsv`

- Figure 3B: `figures/TRO_composite_score_by_stage.png`
  - Claim: morula ranks first by TRO score.
  - Source: `tables/TRO_composite_score_by_stage.tsv`

- Figure 3C: `figures/marker_leave_one_out_summary.png`
  - Claim: potency result is robust to leave-one-marker-out tests.
  - Source: `tables/marker_leave_one_out_summary.tsv`

## Figure 4: Operational TRO operator and transition cost

- Figure 4A: `figures/TRO_stage_transition_cost.png`
  - Claim: 8-cell -> morula is the maximum productive reset transition.
  - Source: `tables/TRO_stage_transition_cost.tsv`

- Figure 4B: `figures/TRO_operator_diagram.png`
  - Claim: the analysis defines an operational TRO = {E, D, R, C}.
  - Source: `tables/TRO_operator_summary.json`
"""
    (NOTES / "figure_panel_plan.md").write_text(figure_plan, encoding="utf-8")

    boundaries = """# Manuscript claim boundaries

## Claims supported by current results

1. Morula is a computational ground-zero candidate in the current public-data TRO framework.
2. Morula has the lowest age-associated methylation entropy among analyzed preimplantation stages.
3. The morula minimum is robust to common-DMR and balanced-bootstrap DNA analyses.
4. Morula retains high developmental potency-marker activity relative to blastocyst.
5. 8-cell and morula form a high-potency region in RNA analyses.
6. The operational TRO score ranks morula first.
7. The strongest productive reset transition is 8-cell -> morula.

## Claims not supported or not safe to make

1. Public data prove a matched aged father's sperm was reset in a matched embryo.
2. S_epi is aging entropy. It is generic methylation entropy.
3. S_epi-age is a direct epigenetic age clock.
4. Morula has the highest global RNA entropy.
5. External GSE44183 proves morula is uniquely higher than 8-cell in potency.
6. The current TRO is a trained neural operator or Schrodinger Bridge model.
7. Experiment 7 pilot proves age-DMR specificity. It is currently a feasibility/pilot control only.

## Recommended final wording

Morula represents a computational ground-zero candidate characterized by minimal age-associated methylation entropy, preserved high developmental potency-marker activity, and the highest operational TRO score among preimplantation stages.
"""
    (NOTES / "manuscript_claim_boundaries.md").write_text(boundaries, encoding="utf-8")

    print("Final claim audit:")
    print(audit.to_string(index=False))
    print("\nWrote:", TABLES / "final_claim_audit.tsv")
    print("Wrote:", TABLES / "manuscript_numbers.json")
    print("Wrote:", NOTES / "figure_panel_plan.md")
    print("Wrote:", NOTES / "manuscript_claim_boundaries.md")


if __name__ == "__main__":
    main()
```
