## Supplemental code for "An entropy-based diagnostic framework for characterizing methylation state dynamics during preimplantation development": 21_experiment9_age_dmr_specificity_boundary_control.py

````
from pathlib import Path
import json

import numpy as np
import pandas as pd


STAGE_ORDER = [
    "MII oocyte",
    "zygote/PN",
    "2-cell",
    "4-cell",
    "8-cell",
    "morula",
    "blastocyst",
    "ICM",
    "TE",
]


def root_dir():
    return Path(__file__).resolve().parents[1]


ROOT = root_dir()
TABLES = ROOT / "tables"
FIGS = ROOT / "figures"
NOTES = ROOT / "notes"


def stage_min_info(df, metric):
    d = df.dropna(subset=[metric]).copy()
    d["stage"] = pd.Categorical(d["stage"], categories=STAGE_ORDER, ordered=True)
    d = d.sort_values("stage")
    ranked = d.sort_values(metric).reset_index(drop=True)
    ground = str(ranked.loc[0, "stage"])
    ground_value = float(ranked.loc[0, metric])
    next_value = float(ranked.loc[1, metric])
    morula_value = float(d.loc[d["stage"].astype(str) == "morula", metric].iloc[0])
    non_morula = d[d["stage"].astype(str) != "morula"]
    next_lowest_non_morula = float(non_morula[metric].min())
    morula_rank = int(ranked.index[ranked["stage"].astype(str) == "morula"][0] + 1)
    return {
        "metric": metric,
        "ground_zero_stage": ground,
        "ground_zero_value": ground_value,
        "next_lowest_value": next_value,
        "morula_value": morula_value,
        "morula_rank": morula_rank,
        "morula_gap_to_next_non_morula": next_lowest_non_morula - morula_value,
    }


def bootstrap_stage_winner(sample_df, metric, n_boot=2000, seed=20260521):
    rng = np.random.default_rng(seed)
    winners = []
    for _ in range(n_boot):
        vals = {}
        for stage in STAGE_ORDER:
            x = sample_df.loc[sample_df["stage"] == stage, metric].dropna().to_numpy()
            if len(x) == 0:
                continue
            vals[stage] = float(rng.choice(x, size=len(x), replace=True).mean())
        if vals:
            winners.append(min(vals, key=vals.get))
    out = pd.Series(winners).value_counts().rename_axis("stage").reset_index(name="n_min")
    out["frequency"] = out["n_min"] / n_boot
    out["metric"] = metric
    return out[["metric", "stage", "n_min", "frequency"]].sort_values(
        ["metric", "frequency"], ascending=[True, False]
    )


def main():
    stage = pd.read_csv(TABLES / "GSE81233_valid204_stage_epi_age_metrics.tsv", sep="\t")
    sample = pd.read_csv(TABLES / "GSE81233_valid204_sample_level_entropy_metrics.tsv", sep="\t")

    metric_rows = []
    metric_rows.append(stage_min_info(stage, "s_epi_age"))
    metric_rows.append(stage_min_info(stage, "s_epi"))
    metric_df = pd.DataFrame(metric_rows)

    boot = pd.concat(
        [
            bootstrap_stage_winner(sample, "s_epi_age_sample", n_boot=2000, seed=1201),
            bootstrap_stage_winner(sample, "s_epi_sample", n_boot=2000, seed=1202),
        ],
        ignore_index=True,
    )

    original_freq = pd.read_csv(TABLES / "Experiment1B_original_bootstrap_ground_zero_frequency.tsv", sep="\t")
    shuffled_freq = pd.read_csv(TABLES / "Experiment1B_shuffled_weight_ground_zero_frequency.tsv", sep="\t")
    random_freq = pd.read_csv(TABLES / "Experiment1B_random_age_DMR_subset_ground_zero_frequency.tsv", sep="\t")

    def morula_freq(df):
        hit = df.loc[df["stage"].astype(str) == "morula", "frequency"]
        return float(hit.iloc[0]) if len(hit) else 0.0

    original_morula_freq = morula_freq(original_freq)
    shuffled_morula_freq = morula_freq(shuffled_freq)
    random_morula_freq = morula_freq(random_freq)
    generic_boot_freq = morula_freq(boot[boot["metric"] == "s_epi_sample"])

    age_info = metric_df.loc[metric_df["metric"] == "s_epi_age"].iloc[0].to_dict()
    generic_info = metric_df.loc[metric_df["metric"] == "s_epi"].iloc[0].to_dict()

    specificity_rows = [
        {
            "control": "age_DMR_true_weights",
            "morula_ground_zero_frequency": original_morula_freq,
            "morula_rank": int(age_info["morula_rank"]),
            "morula_gap_to_next_non_morula": float(age_info["morula_gap_to_next_non_morula"]),
            "interpretation": "primary_age_weighted_signal",
        },
        {
            "control": "generic_unweighted_entropy_on_age_DMR_regions",
            "morula_ground_zero_frequency": generic_boot_freq,
            "morula_rank": int(generic_info["morula_rank"]),
            "morula_gap_to_next_non_morula": float(generic_info["morula_gap_to_next_non_morula"]),
            "interpretation": "broader_methylation_reprogramming_component",
        },
        {
            "control": "shuffled_age_weights",
            "morula_ground_zero_frequency": shuffled_morula_freq,
            "morula_rank": np.nan,
            "morula_gap_to_next_non_morula": np.nan,
            "interpretation": "weight_specificity_control",
        },
        {
            "control": "random_age_DMR_subset",
            "morula_ground_zero_frequency": random_morula_freq,
            "morula_rank": np.nan,
            "morula_gap_to_next_non_morula": np.nan,
            "interpretation": "region_subset_control",
        },
    ]
    specificity = pd.DataFrame(specificity_rows)

    enrichment_vs_shuffled = original_morula_freq / shuffled_morula_freq if shuffled_morula_freq else np.inf
    enrichment_vs_random = original_morula_freq / random_morula_freq if random_morula_freq else np.inf
    gap_gain_over_generic = float(age_info["morula_gap_to_next_non_morula"]) - float(
        generic_info["morula_gap_to_next_non_morula"]
    )

    if (
        str(age_info["ground_zero_stage"]) == "morula"
        and original_morula_freq > shuffled_morula_freq
        and original_morula_freq > random_morula_freq
        and float(age_info["morula_gap_to_next_non_morula"]) > 0
    ):
        conclusion = "age_weighting_strengthens_a_broader_morula_methylation_reprogramming_minimum"
    else:
        conclusion = "age_DMR_specificity_not_established"

    summary = {
        "age_DMR_ground_zero_stage": str(age_info["ground_zero_stage"]),
        "age_DMR_morula_rank": int(age_info["morula_rank"]),
        "age_DMR_morula_gap_to_next_non_morula": float(age_info["morula_gap_to_next_non_morula"]),
        "generic_S_epi_ground_zero_stage": str(generic_info["ground_zero_stage"]),
        "generic_S_epi_morula_rank": int(generic_info["morula_rank"]),
        "generic_S_epi_morula_gap_to_next_non_morula": float(generic_info["morula_gap_to_next_non_morula"]),
        "age_DMR_true_weight_morula_frequency": original_morula_freq,
        "generic_S_epi_bootstrap_morula_frequency": generic_boot_freq,
        "shuffled_weight_morula_frequency": shuffled_morula_freq,
        "random_age_DMR_subset_morula_frequency": random_morula_freq,
        "age_vs_shuffled_frequency_ratio": enrichment_vs_shuffled,
        "age_vs_random_subset_frequency_ratio": enrichment_vs_random,
        "age_weighted_gap_gain_over_generic_S_epi": gap_gain_over_generic,
        "conclusion": conclusion,
        "claim_boundary": (
            "The morula minimum is not exclusive proof of age-DMR specificity. "
            "It is best interpreted as age-weighted methylation entropy strengthening "
            "a broader methylation reprogramming minimum at morula."
        ),
    }

    TABLES.mkdir(exist_ok=True)
    FIGS.mkdir(exist_ok=True)
    NOTES.mkdir(exist_ok=True)

    metric_df.to_csv(TABLES / "Experiment9_age_DMR_specificity_metric_minima.tsv", sep="\t", index=False)
    boot.to_csv(TABLES / "Experiment9_age_DMR_vs_generic_bootstrap_frequency.tsv", sep="\t", index=False)
    specificity.to_csv(TABLES / "Experiment9_age_DMR_specificity_boundary_summary.tsv", sep="\t", index=False)
    (TABLES / "Experiment9_age_DMR_specificity_boundary_summary.json").write_text(
        json.dumps(summary, indent=2, ensure_ascii=False), encoding="utf-8"
    )

    note = f"""# Experiment 9 age-DMR specificity boundary control

This control asks whether the morula minimum is exclusive to age-DMR weighting or reflects a broader methylation reprogramming minimum.

## Result

- True age-DMR weighted entropy ranks morula as the minimum.
- Generic unweighted methylation entropy on the same age-DMR regions also has a morula-associated low point.
- Shuffled weights and random age-DMR subsets still often select morula, although less strongly than the true age-weighted analysis.

## Interpretation

The result should not be phrased as pure age-DMR specificity. The stable wording is:

```text
Age-DMR weighted entropy strengthens a broader methylation reprogramming minimum at morula.
```

This supports the computational ground-zero model while avoiding an overclaim that morula minimum is exclusively caused by age-DMR identity.

## Key values

```json
{json.dumps(summary, indent=2, ensure_ascii=False)}
```
"""
    (NOTES / "Experiment9_age_DMR_specificity_boundary_control.md").write_text(note, encoding="utf-8")

    try:
        import matplotlib.pyplot as plt

        plot_df = specificity.copy()
        fig, ax = plt.subplots(figsize=(8.5, 4.6))
        colors = ["#2b8cbe", "#7bccc4", "#fdae61", "#f46d43"]
        ax.bar(plot_df["control"], plot_df["morula_ground_zero_frequency"], color=colors)
        ax.set_ylabel("Morula ground-zero frequency")
        ax.set_ylim(0, 1.05)
        ax.axhline(0.5, color="gray", linestyle="--", linewidth=1)
        ax.set_title("Age-DMR specificity boundary control")
        ax.tick_params(axis="x", rotation=25)
        for label in ax.get_xticklabels():
            label.set_horizontalalignment("right")
        fig.tight_layout()
        fig.savefig(FIGS / "Experiment9_age_DMR_specificity_boundary_control.png", dpi=300)
        fig.savefig(FIGS / "Experiment9_age_DMR_specificity_boundary_control.pdf")
        plt.close(fig)
    except Exception as exc:
        print(f"Skipping figure because plotting failed: {exc}")

    print("Experiment 9 age-DMR specificity boundary control:")
    print(json.dumps(summary, indent=2, ensure_ascii=False))
    print("Wrote:", TABLES / "Experiment9_age_DMR_specificity_boundary_summary.tsv")
    print("Wrote:", TABLES / "Experiment9_age_DMR_specificity_boundary_summary.json")
    print("Wrote:", NOTES / "Experiment9_age_DMR_specificity_boundary_control.md")


if __name__ == "__main__":
    main()
````
