## Supplemental code for "An entropy-based diagnostic framework for characterizing methylation state dynamics during preimplantation development": 22_experiment10_gse49828_independent_dna_validation.py

````
from pathlib import Path
import argparse
import gzip
import json
import shutil
import urllib.request

import numpy as np
import pandas as pd


ROOT = Path(__file__).resolve().parents[1]
TABLES = ROOT / "tables"
FIGS = ROOT / "figures"
NOTES = ROOT / "notes"
RAW = ROOT / "data_raw" / "GSE49828_independent_dna"
BASE = "https://ftp.ncbi.nlm.nih.gov/geo/series/GSE49nnn/GSE49828/suppl"

STAGE_ORDER = ["MII oocyte", "zygote/PN", "2-cell", "4-cell", "8-cell", "morula", "ICM", "TE"]


SELECTED = [
    ("MII oocyte", "GSE49828_RRBS_MII_Oocyte1_methylation_calling.bed.txt.gz"),
    ("MII oocyte", "GSE49828_RRBS_MII_Oocyte2_methylation_calling.bed.txt.gz"),
    ("zygote/PN", "GSE49828_RRBS_Zygote1_methylation_calling.bed.txt.gz"),
    ("zygote/PN", "GSE49828_RRBS_Zygote2_methylation_calling.bed.txt.gz"),
    ("2-cell", "GSE49828_RRBS_2-cell1_methylation_calling.bed.txt.gz"),
    ("2-cell", "GSE49828_RRBS_2-cell2_methylation_calling.bed.txt.gz"),
    ("4-cell", "GSE49828_RRBS_4-cell1_methylation_calling.bed.txt.gz"),
    ("4-cell", "GSE49828_RRBS_4-cell2_methylation_calling.bed.txt.gz"),
    ("8-cell", "GSE49828_RRBS_8-cell1_methylation_calling.bed.txt.gz"),
    ("8-cell", "GSE49828_RRBS_8-cell2_methylation_calling.bed.txt.gz"),
    ("8-cell", "GSE49828_RRBS_8-cell3_methylation_calling.bed.txt.gz"),
    ("morula", "GSE49828_RRBS_Morula1_methylation_calling.bed.txt.gz"),
    ("morula", "GSE49828_RRBS_Morula2_methylation_calling.bed.txt.gz"),
    ("morula", "GSE49828_RRBS_Morula3_methylation_calling.bed.txt.gz"),
    ("ICM", "GSE49828_RRBS_ICM1_methylation_calling.bed.txt.gz"),
    ("ICM", "GSE49828_RRBS_ICM2_methylation_calling.bed.txt.gz"),
    ("ICM", "GSE49828_RRBS_ICM3_methylation_calling.bed.txt.gz"),
    ("TE", "GSE49828_RRBS_TE1_methylation_calling.bed.txt.gz"),
    ("TE", "GSE49828_RRBS_TE2_methylation_calling.bed.txt.gz"),
    ("TE", "GSE49828_RRBS_TE3_methylation_calling.bed.txt.gz"),
]


def entropy(p, eps=1e-6):
    p = np.clip(np.asarray(p, dtype=float), eps, 1 - eps)
    return -(p * np.log(p) + (1 - p) * np.log(1 - p))


def validate_gzip(path):
    try:
        with gzip.open(path, "rb") as fh:
            while fh.read(1024 * 1024):
                pass
        return True
    except Exception:
        return False


def filter_selected(stage_filter):
    if not stage_filter:
        return SELECTED
    wanted = {x.strip() for x in stage_filter.split(",") if x.strip()}
    return [(stage, name) for stage, name in SELECTED if stage in wanted]


def download_selected(selected, force=False):
    RAW.mkdir(parents=True, exist_ok=True)
    rows = []
    for stage, name in selected:
        url = f"{BASE}/{name}"
        out = RAW / name
        status = "exists"
        if force or not out.exists() or out.stat().st_size == 0 or not validate_gzip(out):
            tmp = out.with_suffix(out.suffix + ".part")
            if tmp.exists():
                tmp.unlink()
            print(f"Downloading {url}")
            urllib.request.urlretrieve(url, tmp)
            shutil.move(str(tmp), str(out))
            status = "downloaded"
        ok = validate_gzip(out)
        rows.append(
            {
                "stage": stage,
                "filename": name,
                "url": url,
                "local_path": str(out),
                "size_bytes": out.stat().st_size if out.exists() else 0,
                "gzip_ok": ok,
                "status": status,
            }
        )
    manifest = pd.DataFrame(rows)
    manifest.to_csv(TABLES / "GSE49828_independent_dna_download_manifest.tsv", sep="\t", index=False)
    return manifest


def load_regions():
    reg = pd.read_csv(ROOT / "metadata" / "GSE102970_TableS6_age_dmr_weights.tsv", sep="\t")
    reg = reg[["cluster_name", "chr", "start", "end", "age_weight_5yr"]].copy()
    reg["start"] = reg["start"].astype(int)
    reg["end"] = reg["end"].astype(int)
    return reg


def build_region_index(reg):
    by_chr = {}
    for chrom, sub in reg.groupby("chr"):
        sub = sub.sort_values("start").reset_index(drop=True)
        by_chr[chrom] = {
            "starts": sub["start"].to_numpy(),
            "ends": sub["end"].to_numpy(),
            "clusters": sub["cluster_name"].to_numpy(),
        }
    return by_chr


def find_region(index, chrom, pos):
    if chrom not in index:
        return None
    starts = index[chrom]["starts"]
    ends = index[chrom]["ends"]
    i = np.searchsorted(starts, pos, side="right") - 1
    if i >= 0 and pos <= ends[i]:
        return str(index[chrom]["clusters"][i])
    return None


def process_file(stage, path, regions, index):
    reg_lookup = regions.set_index("cluster_name")
    met = {name: 0.0 for name in regions["cluster_name"]}
    total = {name: 0.0 for name in regions["cluster_name"]}
    n_cpg = {name: 0 for name in regions["cluster_name"]}
    cg_lines = 0
    matched = 0
    with gzip.open(path, "rt", errors="replace") as fh:
        header = next(fh, "")
        for line in fh:
            parts = line.rstrip("\n").split("\t")
            if len(parts) < 10:
                continue
            typ = parts[9]
            if typ != "CpG":
                continue
            chrom = parts[0]
            try:
                pos = int(parts[1])
                t = float(parts[4])
                m = float(parts[5])
            except ValueError:
                continue
            cg_lines += 1
            cluster = find_region(index, chrom, pos)
            if cluster is None or t <= 0:
                continue
            matched += 1
            met[cluster] += m
            total[cluster] += t
            n_cpg[cluster] += 1

    rows = []
    for cluster, t in total.items():
        if t <= 0:
            continue
        beta = met[cluster] / t
        rows.append(
            {
                "stage": stage,
                "cluster_name": cluster,
                "age_weight_5yr": float(reg_lookup.loc[cluster, "age_weight_5yr"]),
                "met_reads": met[cluster],
                "total_reads": t,
                "beta": beta,
                "n_cpg_observed": n_cpg[cluster],
            }
        )
    return pd.DataFrame(rows), {"stage": stage, "cg_lines": cg_lines, "matched_cpg": matched, "covered_regions": len(rows)}


def combine_metrics(manifest):
    regions = load_regions()
    index = build_region_index(regions)
    per_sample = []
    qc = []
    for _, row in manifest.iterrows():
        if not bool(row["gzip_ok"]):
            continue
        path = Path(row["local_path"])
        if not path.exists():
            continue
        print(f"Processing {row['stage']}: {path.name}")
        sample_df, sample_qc = process_file(str(row["stage"]), path, regions, index)
        sample_df["filename"] = path.name
        per_sample.append(sample_df)
        sample_qc["filename"] = path.name
        qc.append(sample_qc)

    if not per_sample:
        raise SystemExit("No valid GSE49828 files were available for processing.")

    long_df = pd.concat(per_sample, ignore_index=True)
    long_df.to_csv(TABLES / "GSE49828_age_DMR_region_values.tsv", sep="\t", index=False)
    pd.DataFrame(qc).to_csv(TABLES / "GSE49828_independent_dna_processing_qc.tsv", sep="\t", index=False)

    stage_rows = []
    for stage in STAGE_ORDER:
        sub = long_df[long_df["stage"] == stage]
        if sub.empty:
            continue
        g = sub.groupby("cluster_name", as_index=False).agg(
            met_reads=("met_reads", "sum"),
            total_reads=("total_reads", "sum"),
            age_weight_5yr=("age_weight_5yr", "first"),
        )
        g = g[g["total_reads"] > 0].copy()
        p = g["met_reads"] / g["total_reads"]
        h = entropy(p)
        w = g["age_weight_5yr"].astype(float).to_numpy()
        stage_rows.append(
            {
                "stage": stage,
                "n_regions_valid": len(g),
                "s_epi": float(np.mean(h)),
                "s_epi_age": float(np.sum(np.abs(w) * h) / np.sum(np.abs(w))),
                "age_projection": float(np.sum(w * p) / np.sum(np.abs(w))),
            }
        )
    metrics = pd.DataFrame(stage_rows)
    metrics["stage"] = pd.Categorical(metrics["stage"], categories=STAGE_ORDER, ordered=True)
    metrics = metrics.sort_values("stage").reset_index(drop=True)
    metrics.to_csv(TABLES / "GSE49828_independent_DNA_validation_stage_metrics.tsv", sep="\t", index=False)

    ranked = metrics.dropna(subset=["s_epi_age"]).sort_values("s_epi_age").reset_index(drop=True)
    ground = str(ranked.loc[0, "stage"]) if len(ranked) else None
    morula_rank = int(ranked.index[ranked["stage"].astype(str) == "morula"][0] + 1) if "morula" in set(ranked["stage"].astype(str)) else None
    top3 = ranked["stage"].astype(str).head(3).tolist()
    summary = {
        "dataset": "GSE49828",
        "validation_type": "independent_human_RRBS_DNA_methylation",
        "ground_zero_stage_by_s_epi_age": ground,
        "morula_rank_by_s_epi_age": morula_rank,
        "top3_lowest_s_epi_age_stages": top3,
        "supports_morula_or_adjacent_low_age_entropy": bool(morula_rank is not None and morula_rank <= 3),
        "claim_boundary": "GSE49828 is an independent RRBS dataset with sparse age-DMR overlap; use as directional validation only.",
    }
    (TABLES / "GSE49828_independent_DNA_validation_summary.json").write_text(
        json.dumps(summary, indent=2, ensure_ascii=False), encoding="utf-8"
    )

    note = f"""# GSE49828 independent DNA methylation validation

This analysis uses GSE49828 processed RRBS methylation-calling files as an independent DNA methylation validation set.

The validation is directional because GSE49828 is RRBS and has sparse overlap with the GSE102970 age-DMR regions.

```json
{json.dumps(summary, indent=2, ensure_ascii=False)}
```
"""
    (NOTES / "GSE49828_independent_DNA_validation.md").write_text(note, encoding="utf-8")
    print("GSE49828 independent DNA validation:")
    print(json.dumps(summary, indent=2, ensure_ascii=False))
    print("Wrote:", TABLES / "GSE49828_independent_DNA_validation_stage_metrics.tsv")
    print("Wrote:", TABLES / "GSE49828_independent_DNA_validation_summary.json")


def main():
    ap = argparse.ArgumentParser()
    ap.add_argument("--download", action="store_true", help="Download selected GSE49828 processed RRBS files.")
    ap.add_argument("--force-download", action="store_true")
    ap.add_argument("--combine-only", action="store_true", help="Only combine/process already downloaded files.")
    ap.add_argument(
        "--stages",
        default="",
        help="Comma-separated subset of stages, e.g. 'MII oocyte,zygote/PN,2-cell,4-cell,8-cell,morula'.",
    )
    args = ap.parse_args()

    TABLES.mkdir(exist_ok=True)
    FIGS.mkdir(exist_ok=True)
    NOTES.mkdir(exist_ok=True)
    RAW.mkdir(parents=True, exist_ok=True)

    selected = filter_selected(args.stages)

    if args.download:
        manifest = download_selected(selected, force=args.force_download)
    else:
        rows = []
        for stage, name in selected:
            path = RAW / name
            rows.append(
                {
                    "stage": stage,
                    "filename": name,
                    "url": f"{BASE}/{name}",
                    "local_path": str(path),
                    "size_bytes": path.stat().st_size if path.exists() else 0,
                    "gzip_ok": validate_gzip(path) if path.exists() else False,
                    "status": "local" if path.exists() else "missing",
                }
            )
        manifest = pd.DataFrame(rows)
        manifest.to_csv(TABLES / "GSE49828_independent_dna_download_manifest.tsv", sep="\t", index=False)

    combine_metrics(manifest)


if __name__ == "__main__":
    main()
````
