## Supplemental code for "An entropy-based diagnostic framework for characterizing methylation state dynamics during preimplantation development": 23_experiment11_tro_interpretability_analysis.py

````
from pathlib import Path
import gzip
import json
import math
import re
import shutil
import urllib.request

import numpy as np
import pandas as pd


ROOT = Path(__file__).resolve().parents[1]
TABLES = ROOT / "tables"
FIGS = ROOT / "figures"
NOTES = ROOT / "notes"
META = ROOT / "metadata"

REFGENE_URL = "https://hgdownload.soe.ucsc.edu/goldenPath/hg19/database/refGene.txt.gz"
CPG_URL = "https://hgdownload.soe.ucsc.edu/goldenPath/hg19/database/cpgIslandExt.txt.gz"


STAGE_ORDER = [
    "MII oocyte",
    "zygote/PN",
    "2-cell",
    "4-cell",
    "8-cell",
    "morula",
    "blastocyst",
    "ICM",
    "TE",
]


GENE_SET_KEYWORDS = {
    "embryonic_development": {
        "NANOG",
        "POU5F1",
        "SOX2",
        "DPPA3",
        "GDF3",
        "KLF4",
        "KLF17",
        "ZSCAN4",
        "TFAP2C",
        "PRDM14",
        "SALL4",
        "LIN28A",
        "TDGF1",
        "UTF1",
        "ZFP42",
    },
    "chromatin_and_epigenetic_regulation": {
        "DNMT3L",
        "DNMT3A",
        "DNMT3B",
        "TET1",
        "TET2",
        "TET3",
        "EZH2",
        "SUZ12",
        "EED",
        "KDM1A",
        "KDM5B",
        "KMT2D",
        "SMARCA4",
        "ARID1A",
        "CHD1",
        "CHD4",
        "HDAC1",
        "HDAC2",
    },
    "cell_fate_and_stemness": {
        "POU5F1",
        "NANOG",
        "SOX2",
        "KLF4",
        "KLF17",
        "TFAP2C",
        "GDF3",
        "SALL4",
        "PRDM14",
        "LIN28A",
        "UTF1",
        "ZFP42",
    },
    "wnt_tgf_beta_hippo_signaling": {
        "WNT3",
        "WNT5A",
        "CTNNB1",
        "TCF7L1",
        "BMP4",
        "BMP7",
        "NODAL",
        "LEFTY1",
        "LEFTY2",
        "SMAD2",
        "SMAD3",
        "SMAD4",
        "YAP1",
        "TAZ",
        "TEAD4",
    },
    "cell_cycle_and_DNA_repair": {
        "MKI67",
        "PCNA",
        "MCM2",
        "MCM3",
        "MCM4",
        "MCM5",
        "MCM6",
        "BRCA1",
        "BRCA2",
        "RAD51",
        "ATM",
        "ATR",
        "TP53",
    },
}


def download_if_missing(url, dest):
    dest = Path(dest)
    if dest.exists() and dest.stat().st_size > 0:
        return dest
    dest.parent.mkdir(parents=True, exist_ok=True)
    tmp = dest.with_suffix(dest.suffix + ".part")
    if tmp.exists():
        tmp.unlink()
    print(f"Downloading {url}")
    urllib.request.urlretrieve(url, tmp)
    shutil.move(str(tmp), str(dest))
    return dest


def binary_entropy(p, eps=1e-6):
    p = np.clip(np.asarray(p, dtype=float), eps, 1 - eps)
    return -(p * np.log(p) + (1 - p) * np.log(1 - p))


def load_refgene():
    path = download_if_missing(REFGENE_URL, META / "hg19_refGene.txt.gz")
    cols = [
        "bin",
        "name",
        "chrom",
        "strand",
        "txStart",
        "txEnd",
        "cdsStart",
        "cdsEnd",
        "exonCount",
        "exonStarts",
        "exonEnds",
        "score",
        "name2",
        "cdsStartStat",
        "cdsEndStat",
        "exonFrames",
    ]
    genes = pd.read_csv(path, sep="\t", names=cols)
    genes = genes[genes["chrom"].str.startswith("chr")].copy()
    genes = genes.dropna(subset=["name2"])
    genes["txStart"] = genes["txStart"].astype(int)
    genes["txEnd"] = genes["txEnd"].astype(int)
    return genes


def load_cpg_islands():
    path = download_if_missing(CPG_URL, META / "hg19_cpgIslandExt.txt.gz")
    cols = [
        "bin",
        "chrom",
        "chromStart",
        "chromEnd",
        "name",
        "length",
        "cpgNum",
        "gcNum",
        "perCpg",
        "perGc",
        "obsExp",
    ]
    cpg = pd.read_csv(path, sep="\t", names=cols)
    cpg["chromStart"] = cpg["chromStart"].astype(int)
    cpg["chromEnd"] = cpg["chromEnd"].astype(int)
    return cpg


def parse_exons(row):
    starts = [int(x) for x in str(row["exonStarts"]).rstrip(",").split(",") if x != ""]
    ends = [int(x) for x in str(row["exonEnds"]).rstrip(",").split(",") if x != ""]
    return list(zip(starts, ends))


def overlap(a_start, a_end, b_start, b_end):
    return a_start < b_end and b_start < a_end


def annotate_region(chrom, start, end, genes_by_chr, cpg_by_chr):
    mid = int((start + end) / 2)
    gene_sub = genes_by_chr.get(chrom)
    nearest_gene = ""
    nearest_dist = np.inf
    gene_context = "intergenic"
    promoter_hit = False
    exon_hit = False
    intron_hit = False

    if gene_sub is not None and len(gene_sub) > 0:
        for _, g in gene_sub.iterrows():
            tx_start = int(g["txStart"])
            tx_end = int(g["txEnd"])
            tss = tx_start if g["strand"] == "+" else tx_end
            dist = 0 if tx_start <= mid <= tx_end else min(abs(mid - tx_start), abs(mid - tx_end))
            if dist < nearest_dist:
                nearest_dist = dist
                nearest_gene = str(g["name2"])
            if abs(mid - tss) <= 2000:
                promoter_hit = True
                nearest_gene = str(g["name2"]) if not nearest_gene else nearest_gene
            if overlap(start, end, tx_start, tx_end):
                exons = parse_exons(g)
                if any(overlap(start, end, ex_s, ex_e) for ex_s, ex_e in exons):
                    exon_hit = True
                else:
                    intron_hit = True

    if promoter_hit:
        gene_context = "promoter"
    elif exon_hit:
        gene_context = "exon"
    elif intron_hit:
        gene_context = "intron"

    cpg_context = "non_CpG_island"
    cpg_sub = cpg_by_chr.get(chrom)
    if cpg_sub is not None and len(cpg_sub) > 0:
        for _, c in cpg_sub.iterrows():
            c_start = int(c["chromStart"])
            c_end = int(c["chromEnd"])
            if overlap(start, end, c_start, c_end):
                cpg_context = "CpG_island"
                break
            dist = min(abs(mid - c_start), abs(mid - c_end))
            if dist <= 2000 and cpg_context != "CpG_island":
                cpg_context = "CpG_shore"
            elif dist <= 4000 and cpg_context not in {"CpG_island", "CpG_shore"}:
                cpg_context = "CpG_shelf"

    if nearest_dist is np.inf:
        nearest_dist = np.nan

    return nearest_gene, nearest_dist, gene_context, cpg_context


def stage_region_metrics(long_df):
    rows = []
    for (stage, cluster), sub in long_df.groupby(["stage", "cluster_name"], observed=True):
        total = sub["total_reads"].sum()
        if total <= 0:
            continue
        met = sub["met_reads"].sum()
        beta = met / total
        h = float(binary_entropy([beta])[0])
        rows.append(
            {
                "stage": stage,
                "cluster_name": cluster,
                "met_reads": met,
                "total_reads": total,
                "beta": beta,
                "entropy": h,
                "age_weight_5yr": float(sub["age_weight_5yr"].dropna().iloc[0]),
                "n_samples_covered": int((sub["total_reads"] > 0).sum()),
            }
        )
    return pd.DataFrame(rows)


def contribution_table():
    long_df = pd.read_csv(TABLES / "Experiment1B_all_sample_age_dmr_long.tsv.gz", sep="\t")
    reg = pd.read_csv(META / "GSE102970_TableS6_age_dmr_weights.tsv", sep="\t")
    stage_reg = stage_region_metrics(long_df)
    wide = stage_reg.pivot_table(
        index="cluster_name",
        columns="stage",
        values=["entropy", "beta", "total_reads", "n_samples_covered"],
        aggfunc="first",
    )
    wide.columns = [f"{a}_{b}" for a, b in wide.columns]
    wide = wide.reset_index()
    out = reg.merge(wide, on="cluster_name", how="left")

    for stage in ["MII oocyte", "8-cell", "morula", "blastocyst"]:
        h_col = f"entropy_{stage}"
        if h_col not in out.columns:
            out[h_col] = np.nan
    abs_w = out["age_weight_5yr"].abs()
    out["delta_H_MII_to_morula"] = out["entropy_MII oocyte"] - out["entropy_morula"]
    out["delta_H_8cell_to_morula"] = out["entropy_8-cell"] - out["entropy_morula"]
    out["contribution_MII_to_morula"] = abs_w * out["delta_H_MII_to_morula"]
    out["contribution_8cell_to_morula"] = abs_w * out["delta_H_8cell_to_morula"]
    out["abs_contribution_8cell_to_morula"] = out["contribution_8cell_to_morula"].abs()

    genes = load_refgene()
    cpg = load_cpg_islands()
    genes_by_chr = {chrom: sub for chrom, sub in genes.groupby("chrom")}
    cpg_by_chr = {chrom: sub for chrom, sub in cpg.groupby("chrom")}

    annotations = []
    for _, row in out.iterrows():
        annotations.append(
            annotate_region(
                str(row["chr"]),
                int(row["start"]),
                int(row["end"]),
                genes_by_chr,
                cpg_by_chr,
            )
        )
    ann = pd.DataFrame(
        annotations,
        columns=["nearest_gene", "nearest_gene_distance_bp", "gene_context", "cpg_context"],
    )
    out = pd.concat([out, ann], axis=1)
    out["reset_driver_rank_8cell_to_morula"] = out["contribution_8cell_to_morula"].rank(
        ascending=False, method="min", na_option="bottom"
    )
    out["reset_driver_rank_MII_to_morula"] = out["contribution_MII_to_morula"].rank(
        ascending=False, method="min", na_option="bottom"
    )
    out = out.sort_values("reset_driver_rank_8cell_to_morula")
    return out


def hypergeom_sf(k, n, K, N):
    # P[X >= k] for Hypergeometric(N, K, n), computed in log space.
    if k <= 0:
        return 1.0
    max_i = min(K, n)
    vals = []
    for i in range(k, max_i + 1):
        if n - i > N - K:
            continue
        vals.append(
            math.lgamma(K + 1)
            - math.lgamma(i + 1)
            - math.lgamma(K - i + 1)
            + math.lgamma(N - K + 1)
            - math.lgamma(n - i + 1)
            - math.lgamma(N - K - n + i + 1)
            - (
                math.lgamma(N + 1)
                - math.lgamma(n + 1)
                - math.lgamma(N - n + 1)
            )
        )
    if not vals:
        return 1.0
    m = max(vals)
    return float(min(1.0, math.exp(m) * sum(math.exp(v - m) for v in vals)))


def bh_adjust(pvals):
    p = np.asarray(pvals, dtype=float)
    order = np.argsort(p)
    ranked = np.empty_like(p)
    prev = 1.0
    n = len(p)
    for i in range(n - 1, -1, -1):
        idx = order[i]
        val = min(prev, p[idx] * n / (i + 1))
        ranked[idx] = val
        prev = val
    return ranked


def annotation_summary(contrib, top_n=50):
    top = contrib.nsmallest(top_n, "reset_driver_rank_8cell_to_morula")
    all_rows = []
    for category_col in ["gene_context", "cpg_context"]:
        all_counts = contrib[category_col].value_counts()
        top_counts = top[category_col].value_counts()
        for cat in sorted(set(all_counts.index) | set(top_counts.index)):
            all_n = int(all_counts.get(cat, 0))
            top_k = int(top_counts.get(cat, 0))
            all_rows.append(
                {
                    "annotation_type": category_col,
                    "category": cat,
                    "top_count": top_k,
                    "top_fraction": top_k / len(top),
                    "background_count": all_n,
                    "background_fraction": all_n / len(contrib),
                    "p_value": hypergeom_sf(top_k, len(top), all_n, len(contrib)),
                }
            )
    out = pd.DataFrame(all_rows)
    out["p_adj_BH"] = bh_adjust(out["p_value"].fillna(1.0).to_numpy())
    return out.sort_values(["annotation_type", "p_value"])


def keyword_enrichment(contrib, top_n=50):
    genes = set(str(x).upper() for x in contrib["nearest_gene"].dropna() if str(x).strip())
    top = contrib.nsmallest(top_n, "reset_driver_rank_8cell_to_morula")
    top_genes = set(str(x).upper() for x in top["nearest_gene"].dropna() if str(x).strip())
    rows = []
    for term, gene_set in GENE_SET_KEYWORDS.items():
        gs = {g.upper() for g in gene_set}
        bg_hits = genes & gs
        top_hits = top_genes & gs
        p = hypergeom_sf(len(top_hits), len(top_genes), len(bg_hits), len(genes)) if genes and top_genes else 1.0
        rows.append(
            {
                "pathway_or_function": term,
                "top_gene_count": len(top_hits),
                "background_gene_count": len(bg_hits),
                "p_value": p,
                "representative_genes": ",".join(sorted(top_hits)),
            }
        )
    out = pd.DataFrame(rows)
    out["FDR"] = bh_adjust(out["p_value"].fillna(1.0).to_numpy())
    return out.sort_values(["p_value", "pathway_or_function"])


def marker_contribution():
    marker = pd.read_csv(TABLES / "GSE36552_marker_zscore_heatmap_matrix.tsv", sep="\t")
    morula = marker[marker["stage"] == "morula"].iloc[0].drop(labels=["stage"])
    vals = morula.astype(float)
    positive = vals.clip(lower=0)
    denom = positive.sum()
    if denom <= 0:
        contrib = vals / vals.abs().sum()
    else:
        contrib = positive / denom
    out = pd.DataFrame(
        {
            "marker": vals.index,
            "morula_zscore": vals.values,
            "positive_fraction_of_morula_marker_signal": contrib.values,
        }
    ).sort_values("positive_fraction_of_morula_marker_signal", ascending=False)
    loo = pd.read_csv(TABLES / "marker_leave_one_out_summary.tsv", sep="\t")
    potency_loo = loo[loo["metric"] == "potency_score_recomputed"].copy()
    out = out.merge(
        potency_loo[["removed_marker", "morula_mean", "blastocyst_mean", "morula_vs_blastocyst_p_adj_BH", "conclusion"]],
        left_on="marker",
        right_on="removed_marker",
        how="left",
    ).drop(columns=["removed_marker"])
    return out


def draw_bar_png(rows, label_col, value_col, title, out_png, color=(43, 140, 190), max_items=15):
    try:
        from PIL import Image, ImageDraw, ImageFont
    except Exception:
        return False
    data = rows.head(max_items).copy()
    w, h = 1200, 760
    img = Image.new("RGB", (w, h), "white")
    draw = ImageDraw.Draw(img)
    try:
        font_title = ImageFont.truetype("arial.ttf", 30)
        font = ImageFont.truetype("arial.ttf", 18)
        font_small = ImageFont.truetype("arial.ttf", 15)
    except Exception:
        font_title = font = font_small = None
    draw.text((40, 25), title, fill="black", font=font_title)
    left, top, plot_w, bar_h, gap = 330, 90, 790, 28, 12
    vals = data[value_col].astype(float).fillna(0)
    max_val = max(vals.max(), 1e-9)
    for i, (_, row) in enumerate(data.iterrows()):
        y = top + i * (bar_h + gap)
        label = str(row[label_col])[:36]
        val = float(row[value_col]) if pd.notna(row[value_col]) else 0.0
        bw = int(plot_w * max(val, 0) / max_val)
        draw.text((25, y + 4), label, fill="black", font=font_small)
        draw.rectangle((left, y, left + bw, y + bar_h), fill=color)
        draw.rectangle((left, y, left + plot_w, y + bar_h), outline="black")
        draw.text((left + bw + 8, y + 4), f"{val:.4g}", fill="black", font=font_small)
    img.save(out_png)
    return True


def main():
    TABLES.mkdir(exist_ok=True)
    FIGS.mkdir(exist_ok=True)
    NOTES.mkdir(exist_ok=True)

    contrib = contribution_table()
    contrib.to_csv(TABLES / "TRO_interpretability_DMR_contribution_ranking.tsv", sep="\t", index=False)
    contrib.nsmallest(20, "reset_driver_rank_8cell_to_morula").to_csv(
        TABLES / "TRO_interpretability_top20_reset_driving_DMRs.tsv", sep="\t", index=False
    )
    contrib.nsmallest(50, "reset_driver_rank_8cell_to_morula").to_csv(
        TABLES / "TRO_interpretability_top50_reset_driving_DMRs.tsv", sep="\t", index=False
    )
    ann = annotation_summary(contrib, top_n=50)
    ann.to_csv(TABLES / "TRO_interpretability_DMR_annotation_enrichment.tsv", sep="\t", index=False)
    enrich = keyword_enrichment(contrib, top_n=50)
    enrich.to_csv(TABLES / "TRO_interpretability_reset_DMR_gene_function_enrichment.tsv", sep="\t", index=False)
    markers = marker_contribution()
    markers.to_csv(TABLES / "TRO_interpretability_potency_marker_contribution.tsv", sep="\t", index=False)

    draw_bar_png(
        contrib.nsmallest(15, "reset_driver_rank_8cell_to_morula"),
        "cluster_name",
        "contribution_8cell_to_morula",
        "Top reset-driving age-DMRs: 8-cell to morula",
        FIGS / "TRO_interpretability_top_reset_driving_DMRs.png",
    )
    draw_bar_png(
        markers.sort_values("positive_fraction_of_morula_marker_signal", ascending=False),
        "marker",
        "positive_fraction_of_morula_marker_signal",
        "Morula potency-marker contribution",
        FIGS / "TRO_interpretability_potency_marker_contribution.png",
        color=(123, 204, 196),
        max_items=10,
    )

    top_genes = [g for g in contrib.nsmallest(20, "reset_driver_rank_8cell_to_morula")["nearest_gene"].dropna().astype(str) if g]
    top_context = ann.head(8).to_dict(orient="records")
    summary = {
        "analysis": "TRO interpretability analysis",
        "top_reset_driving_DMR_table": "TRO_interpretability_top20_reset_driving_DMRs.tsv",
        "top_reset_driving_genes": top_genes[:20],
        "top_annotation_rows": top_context,
        "top_marker_contributors": markers.head(5)[["marker", "morula_zscore", "positive_fraction_of_morula_marker_signal"]].to_dict(orient="records"),
        "interpretation": (
            "The morula ground-zero call is explained by explicit DMR-level entropy reductions "
            "from 8-cell to morula plus marker-robust preservation of developmental potency. "
            "Nearest-gene and annotation enrichment are exploratory because DMR-to-gene mapping "
            "is based on hg19 nearest transcript annotation."
        ),
    }
    (TABLES / "TRO_interpretability_summary.json").write_text(
        json.dumps(summary, indent=2, ensure_ascii=False), encoding="utf-8"
    )

    note = f"""# TRO interpretability analysis

This analysis explains why morula is selected as the computational ground-zero candidate.

## DMR contribution

For each age-DMR, the main reset-driving contribution was computed as:

```text
abs(age_weight) * (H_8-cell - H_morula)
```

Positive values indicate age-DMR entropy reduction from 8-cell to morula.

Main output:

```text
tables/TRO_interpretability_DMR_contribution_ranking.tsv
tables/TRO_interpretability_top20_reset_driving_DMRs.tsv
tables/TRO_interpretability_top50_reset_driving_DMRs.tsv
```

## Gene and region annotation

DMRs were mapped to hg19 RefGene nearest genes and annotated as promoter, exon, intron, or intergenic, with CpG island/shore/shelf context.

Main output:

```text
tables/TRO_interpretability_DMR_annotation_enrichment.tsv
tables/TRO_interpretability_reset_DMR_gene_function_enrichment.tsv
```

## Potency marker contribution

Morula marker contribution was estimated from positive morula marker z-scores, and interpreted together with leave-one-marker-out robustness.

Main output:

```text
tables/TRO_interpretability_potency_marker_contribution.tsv
tables/marker_leave_one_out_summary.tsv
```

## Summary

```json
{json.dumps(summary, indent=2, ensure_ascii=False)}
```

## Claim boundary

This is a biological interpretability analysis, not a black-box SHAP/LIME analysis. It explains the explicit TRO components by DMR-level entropy reduction, genomic annotation, exploratory nearest-gene function, and potency-marker robustness.
"""
    (NOTES / "TRO_interpretability_analysis.md").write_text(note, encoding="utf-8")

    print("TRO interpretability analysis completed.")
    print(json.dumps(summary, indent=2, ensure_ascii=False))


if __name__ == "__main__":
    main()
````
