## Supplemental code for "An entropy-based diagnostic framework for characterizing methylation state dynamics during preimplantation development": 24_experiment12_unsupervised_latent_reset_analysis.py

```
from __future__ import annotations

import json
from pathlib import Path

import numpy as np
import pandas as pd


STAGE_ORDER = [
    "MII oocyte",
    "zygote/PN",
    "2-cell",
    "4-cell",
    "8-cell",
    "morula",
    "blastocyst",
    "ICM",
    "TE",
]


def rankdata_average(x: np.ndarray) -> np.ndarray:
    order = np.argsort(x, kind="mergesort")
    ranks = np.empty(len(x), dtype=float)
    i = 0
    while i < len(x):
        j = i + 1
        while j < len(x) and x[order[j]] == x[order[i]]:
            j += 1
        ranks[order[i:j]] = (i + 1 + j) / 2.0
        i = j
    return ranks


def corr(x: np.ndarray, y: np.ndarray) -> float:
    ok = np.isfinite(x) & np.isfinite(y)
    if ok.sum() < 3:
        return float("nan")
    x0 = x[ok] - np.mean(x[ok])
    y0 = y[ok] - np.mean(y[ok])
    denom = np.sqrt(np.sum(x0 * x0) * np.sum(y0 * y0))
    if denom == 0:
        return float("nan")
    return float(np.sum(x0 * y0) / denom)


def spearman(x: np.ndarray, y: np.ndarray) -> float:
    ok = np.isfinite(x) & np.isfinite(y)
    if ok.sum() < 3:
        return float("nan")
    return corr(rankdata_average(x[ok]), rankdata_average(y[ok]))


def bh_adjust(pvals: list[float]) -> list[float]:
    arr = np.asarray(pvals, dtype=float)
    out = np.full_like(arr, np.nan, dtype=float)
    ok = np.isfinite(arr)
    if not ok.any():
        return out.tolist()
    idx = np.where(ok)[0]
    order = idx[np.argsort(arr[idx])]
    ranked = arr[order]
    m = len(ranked)
    adj = ranked * m / np.arange(1, m + 1)
    adj = np.minimum.accumulate(adj[::-1])[::-1]
    out[order] = np.minimum(adj, 1.0)
    return out.tolist()


def make_pca(matrix: pd.DataFrame, n_components: int = 3) -> tuple[pd.DataFrame, pd.DataFrame, np.ndarray]:
    values = matrix.to_numpy(dtype=float)
    col_means = np.nanmean(values, axis=0)
    values = np.where(np.isfinite(values), values, col_means)
    col_sd = values.std(axis=0, ddof=0)
    keep = col_sd > 1e-8
    values = values[:, keep]
    columns = matrix.columns[keep]
    z = (values - values.mean(axis=0)) / values.std(axis=0, ddof=0)
    u, s, vt = np.linalg.svd(z, full_matrices=False)
    n = min(n_components, vt.shape[0])
    scores = u[:, :n] * s[:n]
    loadings = vt[:n, :].T
    var = (s * s) / max(z.shape[0] - 1, 1)
    explained = var / var.sum()
    score_df = pd.DataFrame(scores, index=matrix.index, columns=[f"PC{i+1}" for i in range(n)])
    loading_df = pd.DataFrame(loadings, index=columns, columns=[f"PC{i+1}_loading" for i in range(n)])
    return score_df, loading_df, explained[:n]


def pca_summary_for_matrix(matrix: pd.DataFrame, sample_metrics: pd.DataFrame, representation: str) -> dict:
    score_df, _, explained = make_pca(matrix, n_components=3)
    df = score_df.reset_index().merge(sample_metrics, on="sample_id", how="left")
    corr_rows = []
    for pc in [c for c in score_df.columns if c.startswith("PC")]:
        corr_rows.append(corr(df[pc].to_numpy(), df["s_epi_age_sample"].to_numpy()))
    best_idx = int(np.nanargmax(np.abs(corr_rows)))
    best_pc = f"PC{best_idx + 1}"
    sign = 1.0 if corr_rows[best_idx] >= 0 else -1.0
    df["latent_axis"] = sign * df[best_pc]
    stage_axis = df.groupby("stage", observed=True)["latent_axis"].mean().sort_values()
    ranks = {stage: i + 1 for i, stage in enumerate(stage_axis.index)}
    return {
        "representation": representation,
        "n_samples": int(matrix.shape[0]),
        "n_features": int(matrix.shape[1]),
        "best_axis": best_pc,
        "best_axis_pearson_with_s_epi_age_sample": float(corr_rows[best_idx]),
        "best_axis_spearman_with_s_epi_age_sample": float(
            spearman(df[best_pc].to_numpy(), df["s_epi_age_sample"].to_numpy())
        ),
        "best_axis_explained_variance_ratio": float(explained[best_idx]),
        "morula_latent_low_age_rank": int(ranks.get("morula", -1)),
        "lowest_latent_stage": str(stage_axis.index[0]) if len(stage_axis) else None,
        "second_lowest_latent_stage": str(stage_axis.index[1]) if len(stage_axis) > 1 else None,
        "supports_low_age_entropy_window": bool(abs(corr_rows[best_idx]) >= 0.5 and ranks.get("morula", 99) <= 2),
    }


def draw_latent_plot(sample_df: pd.DataFrame, stage_df: pd.DataFrame, out_png: Path) -> None:
    from PIL import Image, ImageDraw, ImageFont

    colors = {
        "MII oocyte": (76, 120, 168),
        "zygote/PN": (114, 183, 178),
        "2-cell": (84, 162, 75),
        "4-cell": (238, 202, 59),
        "8-cell": (245, 133, 24),
        "morula": (228, 87, 86),
        "blastocyst": (178, 121, 162),
        "ICM": (157, 117, 93),
        "TE": (186, 176, 172),
    }
    w, h = 1100, 820
    margin_left, margin_right, margin_top, margin_bottom = 95, 250, 70, 95
    img = Image.new("RGB", (w, h), "white")
    draw = ImageDraw.Draw(img)
    font = ImageFont.load_default()
    title_font = ImageFont.load_default()
    xs = sample_df["PC1"].to_numpy(dtype=float)
    ys = sample_df["PC2"].to_numpy(dtype=float)
    xmin, xmax = float(np.nanmin(xs)), float(np.nanmax(xs))
    ymin, ymax = float(np.nanmin(ys)), float(np.nanmax(ys))
    dx = xmax - xmin or 1.0
    dy = ymax - ymin or 1.0
    xmin -= 0.08 * dx
    xmax += 0.08 * dx
    ymin -= 0.08 * dy
    ymax += 0.08 * dy

    def sx(x: float) -> int:
        return int(margin_left + (x - xmin) / (xmax - xmin) * (w - margin_left - margin_right))

    def sy(y: float) -> int:
        return int(h - margin_bottom - (y - ymin) / (ymax - ymin) * (h - margin_top - margin_bottom))

    draw.rectangle([margin_left, margin_top, w - margin_right, h - margin_bottom], outline=(180, 180, 180), width=1)
    if xmin <= 0 <= xmax:
        x0 = sx(0)
        draw.line([x0, margin_top, x0, h - margin_bottom], fill=(210, 210, 210), width=1)
    if ymin <= 0 <= ymax:
        y0 = sy(0)
        draw.line([margin_left, y0, w - margin_right, y0], fill=(210, 210, 210), width=1)

    cent = stage_df.set_index("stage")
    available = [s for s in STAGE_ORDER if s in cent.index and s not in {"ICM", "TE"}]
    points = [(sx(float(cent.loc[s, "PC1_mean"])), sy(float(cent.loc[s, "PC2_mean"]))) for s in available]
    if len(points) > 1:
        draw.line(points, fill=(70, 70, 70), width=2)
    for stage in STAGE_ORDER:
        sub = sample_df[sample_df["stage"] == stage]
        color = colors.get(stage, (100, 100, 100))
        r = 6 if stage != "morula" else 9
        for _, row in sub.iterrows():
            x, y = sx(float(row["PC1"])), sy(float(row["PC2"]))
            draw.ellipse([x - r, y - r, x + r, y + r], fill=color, outline="white")
    for stage in available:
        row = cent.loc[stage]
        draw.text((sx(float(row["PC1_mean"])) + 5, sy(float(row["PC2_mean"])) - 12), stage, fill=(30, 30, 30), font=font)
    draw.text((margin_left, 25), "Unsupervised latent space of age-DMR methylation profiles", fill=(20, 20, 20), font=title_font)
    draw.text((w // 2 - 50, h - 45), "PC1 latent axis", fill=(20, 20, 20), font=font)
    draw.text((20, h // 2), "PC2", fill=(20, 20, 20), font=font)
    lx = w - margin_right + 30
    ly = margin_top + 10
    for stage in STAGE_ORDER:
        if stage not in set(sample_df["stage"].astype(str)):
            continue
        color = colors.get(stage, (100, 100, 100))
        draw.ellipse([lx, ly, lx + 12, ly + 12], fill=color)
        draw.text((lx + 18, ly - 1), stage, fill=(30, 30, 30), font=font)
        ly += 23
    img.save(out_png)


def draw_stage_axis_plot(stage_df: pd.DataFrame, out_png: Path) -> None:
    from PIL import Image, ImageDraw, ImageFont

    df = stage_df[stage_df["stage"].isin(STAGE_ORDER[:7])].copy()
    df["stage"] = pd.Categorical(df["stage"], categories=STAGE_ORDER[:7], ordered=True)
    df = df.sort_values("stage")
    w, h = 1100, 620
    margin_left, margin_right, margin_top, margin_bottom = 95, 85, 70, 125
    img = Image.new("RGB", (w, h), "white")
    draw = ImageDraw.Draw(img)
    font = ImageFont.load_default()
    draw.text((margin_left, 25), "Latent age-like axis tracks age-associated methylation entropy", fill=(20, 20, 20), font=font)
    draw.rectangle([margin_left, margin_top, w - margin_right, h - margin_bottom], outline=(180, 180, 180), width=1)
    latent = df["latent_age_axis_mean"].to_numpy(dtype=float)
    epi = df["s_epi_age_mean"].to_numpy(dtype=float)
    latent_min, latent_max = float(latent.min()), float(latent.max())
    epi_min, epi_max = float(epi.min()), float(epi.max())
    if latent_max == latent_min:
        latent_max += 1
    if epi_max == epi_min:
        epi_max += 1
    n = len(df)

    def sx(i: int) -> int:
        return int(margin_left + i / max(n - 1, 1) * (w - margin_left - margin_right))

    def sy_lat(v: float) -> int:
        return int(h - margin_bottom - (v - latent_min) / (latent_max - latent_min) * (h - margin_top - margin_bottom))

    def sy_epi(v: float) -> int:
        return int(h - margin_bottom - (v - epi_min) / (epi_max - epi_min) * (h - margin_top - margin_bottom))

    latent_points = [(sx(i), sy_lat(v)) for i, v in enumerate(latent)]
    epi_points = [(sx(i), sy_epi(v)) for i, v in enumerate(epi)]
    draw.line(latent_points, fill=(76, 120, 168), width=3)
    draw.line(epi_points, fill=(228, 87, 86), width=3)
    for i, row in enumerate(df.itertuples(index=False)):
        x = sx(i)
        draw.ellipse([x - 5, sy_lat(float(row.latent_age_axis_mean)) - 5, x + 5, sy_lat(float(row.latent_age_axis_mean)) + 5], fill=(76, 120, 168))
        draw.rectangle([x - 5, sy_epi(float(row.s_epi_age_mean)) - 5, x + 5, sy_epi(float(row.s_epi_age_mean)) + 5], fill=(228, 87, 86))
        draw.text((x - 28, h - margin_bottom + 20), str(row.stage), fill=(35, 35, 35), font=font)
    morula_idx = list(df["stage"].astype(str)).index("morula") if "morula" in list(df["stage"].astype(str)) else None
    if morula_idx is not None:
        x = sx(morula_idx)
        draw.line([x, margin_top, x, h - margin_bottom], fill=(80, 80, 80), width=1)
    draw.line([margin_left, h - margin_bottom + 3, w - margin_right, h - margin_bottom + 3], fill=(120, 120, 120), width=1)
    draw.text((margin_left, h - 45), "blue: oriented latent age-like axis; red: sample-level S_epi-age mean", fill=(35, 35, 35), font=font)
    img.save(out_png)


def main() -> None:
    root = Path(__file__).resolve().parents[1]
    tables = root / "tables"
    figures = root / "figures"
    notes = root / "notes"
    figures.mkdir(exist_ok=True)
    notes.mkdir(exist_ok=True)

    long_df = pd.read_csv(tables / "Experiment1B_all_sample_age_dmr_long.tsv.gz", sep="\t")
    sample_metrics = pd.read_csv(tables / "GSE81233_valid204_sample_level_entropy_metrics.tsv", sep="\t")
    stage_metrics = pd.read_csv(tables / "GSE81233_valid204_internal_reset_score.tsv", sep="\t")

    long_df = long_df[long_df["total_reads"].fillna(0) > 0].copy()
    beta_for_entropy = long_df["beta"].clip(1e-6, 1 - 1e-6)
    long_df["entropy_profile"] = -(
        beta_for_entropy * np.log(beta_for_entropy)
        + (1 - beta_for_entropy) * np.log(1 - beta_for_entropy)
    )
    long_df["weighted_entropy_profile"] = long_df["entropy_profile"] * long_df["age_weight_5yr"].abs()

    representation_summaries = []
    for representation, value_col in [
        ("beta_methylation_profile", "beta"),
        ("entropy_profile", "entropy_profile"),
        ("age_weighted_entropy_profile", "weighted_entropy_profile"),
    ]:
        rep_matrix = long_df.pivot_table(index="sample_id", columns="cluster_name", values=value_col, aggfunc="mean")
        rep_matrix = rep_matrix.loc[rep_matrix.notna().mean(axis=1) >= 0.05]
        rep_matrix = rep_matrix.loc[:, rep_matrix.notna().mean(axis=0) >= 0.05]
        representation_summaries.append(pca_summary_for_matrix(rep_matrix, sample_metrics, representation))

    representation_df = pd.DataFrame(representation_summaries)
    representation_df.to_csv(tables / "TRO_latent_reset_representation_sensitivity.tsv", sep="\t", index=False)

    matrix = long_df.pivot_table(index="sample_id", columns="cluster_name", values="beta", aggfunc="mean")
    coverage = matrix.notna().mean(axis=1)
    matrix = matrix.loc[coverage >= 0.05]
    col_cov = matrix.notna().mean(axis=0)
    matrix = matrix.loc[:, col_cov >= 0.05]

    score_df, loading_df, explained = make_pca(matrix, n_components=3)
    sample_df = score_df.reset_index().merge(sample_metrics, on="sample_id", how="left")
    sample_df["stage"] = pd.Categorical(sample_df["stage"], categories=STAGE_ORDER, ordered=True)
    sample_df = sample_df.sort_values(["stage", "sample_id"]).reset_index(drop=True)

    corr_rows = []
    for pc in [c for c in score_df.columns if c.startswith("PC")]:
        corr_rows.append(
            {
                "latent_axis": pc,
                "pearson_with_s_epi_age_sample": corr(sample_df[pc].to_numpy(), sample_df["s_epi_age_sample"].to_numpy()),
                "spearman_with_s_epi_age_sample": spearman(sample_df[pc].to_numpy(), sample_df["s_epi_age_sample"].to_numpy()),
                "pearson_with_age_projection_sample": corr(sample_df[pc].to_numpy(), sample_df["age_projection_sample"].to_numpy()),
                "explained_variance_ratio": float(explained[int(pc[2:]) - 1]),
            }
        )
    corr_df = pd.DataFrame(corr_rows)
    best_axis = corr_df.iloc[corr_df["pearson_with_s_epi_age_sample"].abs().idxmax()]
    axis_name = str(best_axis["latent_axis"])
    sign = 1.0 if float(best_axis["pearson_with_s_epi_age_sample"]) >= 0 else -1.0
    sample_df["latent_age_axis"] = sign * sample_df[axis_name]

    stage_df = (
        sample_df.groupby("stage", observed=True)
        .agg(
            n_samples=("sample_id", "size"),
            PC1_mean=("PC1", "mean"),
            PC2_mean=("PC2", "mean"),
            PC3_mean=("PC3", "mean"),
            latent_age_axis_mean=("latent_age_axis", "mean"),
            latent_age_axis_sd=("latent_age_axis", "std"),
            s_epi_age_mean=("s_epi_age_sample", "mean"),
            s_epi_age_sd=("s_epi_age_sample", "std"),
            n_regions_covered_mean=("n_regions_covered", "mean"),
        )
        .reset_index()
    )
    stage_df["stage"] = stage_df["stage"].astype(str)
    stage_df = stage_df.merge(
        stage_metrics[["stage", "s_epi_age", "relative_reset_score_internal"]],
        on="stage",
        how="left",
    )
    stage_df["latent_low_age_rank"] = stage_df["latent_age_axis_mean"].rank(method="min", ascending=True).astype(int)
    stage_df["s_epi_age_rank"] = stage_df["s_epi_age"].rank(method="min", ascending=True).astype(int)

    cent = stage_df.set_index("stage")
    transition_rows = []
    for a, b in zip(STAGE_ORDER[:6], STAGE_ORDER[1:7]):
        if a not in cent.index or b not in cent.index:
            continue
        va = cent.loc[a, ["PC1_mean", "PC2_mean", "PC3_mean"]].to_numpy(dtype=float)
        vb = cent.loc[b, ["PC1_mean", "PC2_mean", "PC3_mean"]].to_numpy(dtype=float)
        transition_rows.append(
            {
                "transition": f"{a} -> {b}",
                "latent_distance": float(np.linalg.norm(vb - va)),
                "latent_age_axis_change": float(cent.loc[b, "latent_age_axis_mean"] - cent.loc[a, "latent_age_axis_mean"]),
                "s_epi_age_change": float(cent.loc[b, "s_epi_age"] - cent.loc[a, "s_epi_age"]),
            }
        )
    transition_df = pd.DataFrame(transition_rows)
    if not transition_df.empty:
        transition_df["latent_distance_rank"] = transition_df["latent_distance"].rank(method="min", ascending=False).astype(int)

    loading = loading_df.reset_index().rename(columns={"index": "cluster_name"})
    loading["abs_loading_on_best_axis"] = loading[f"{axis_name}_loading"].abs()
    loading = loading.sort_values("abs_loading_on_best_axis", ascending=False)
    loading = loading.merge(
        pd.read_csv(tables / "TRO_interpretability_DMR_contribution_ranking.tsv", sep="\t")[
            ["cluster_name", "chr", "start", "end", "nearest_gene", "gene_context", "cpg_context", "reset_driver_rank_8cell_to_morula"]
        ],
        on="cluster_name",
        how="left",
    )

    sample_df.to_csv(tables / "TRO_latent_reset_sample_scores.tsv", sep="\t", index=False)
    stage_df.to_csv(tables / "TRO_latent_reset_stage_summary.tsv", sep="\t", index=False)
    corr_df.to_csv(tables / "TRO_latent_reset_axis_correlations.tsv", sep="\t", index=False)
    transition_df.to_csv(tables / "TRO_latent_reset_transition_distances.tsv", sep="\t", index=False)
    loading.head(50).to_csv(tables / "TRO_latent_reset_top_axis_DMR_loadings.tsv", sep="\t", index=False)

    draw_latent_plot(sample_df, stage_df, figures / "TRO_latent_reset_pca_space.png")
    draw_stage_axis_plot(stage_df, figures / "TRO_latent_reset_axis_by_stage.png")

    morula = cent.loc["morula"] if "morula" in cent.index else None
    best_transition = transition_df.sort_values("latent_distance", ascending=False).iloc[0].to_dict() if not transition_df.empty else {}
    summary = {
        "analysis": "unsupervised latent-space validation of TRO ground-zero state",
        "method": "PCA/SVD sensitivity on sample-by-age-DMR beta, entropy, and age-weighted entropy profiles",
        "n_samples": int(matrix.shape[0]),
        "n_age_DMR_features": int(matrix.shape[1]),
        "best_latent_axis_for_s_epi_age": axis_name,
        "best_axis_oriented_to_low_age_entropy": "lower latent_age_axis means lower S_epi-age",
        "best_axis_pearson_with_s_epi_age_sample": float(best_axis["pearson_with_s_epi_age_sample"]),
        "best_axis_spearman_with_s_epi_age_sample": float(best_axis["spearman_with_s_epi_age_sample"]),
        "explained_variance_ratio": {f"PC{i+1}": float(v) for i, v in enumerate(explained)},
        "morula_latent_low_age_rank": int(morula["latent_low_age_rank"]) if morula is not None else None,
        "morula_s_epi_age_rank": int(morula["s_epi_age_rank"]) if morula is not None else None,
        "morula_latent_age_axis_mean": float(morula["latent_age_axis_mean"]) if morula is not None else None,
        "best_latent_transition": best_transition.get("transition"),
        "best_latent_transition_distance": float(best_transition.get("latent_distance", np.nan)) if best_transition else None,
        "representation_sensitivity": representation_summaries,
        "supports_unsupervised_low_age_entropy_window": bool(
            any(x["supports_low_age_entropy_window"] for x in representation_summaries)
        ),
        "supports_beta_profile_morula_unique_cluster": bool(
            morula is not None and int(morula["latent_low_age_rank"]) <= 2
        ),
        "interpretation_boundary": "This is a light unsupervised representation analysis, not a VAE/neural-operator claim.",
    }
    (tables / "TRO_latent_reset_summary.json").write_text(json.dumps(summary, indent=2), encoding="utf-8")

    note = f"""# Experiment 12: Unsupervised latent-space validation

This analysis uses the sample-by-age-DMR methylation beta matrix from GSE81233 and performs PCA/SVD without using TRO scores as input.

## Key result

- Samples analyzed: {summary['n_samples']}
- Age-DMR features used: {summary['n_age_DMR_features']}
- Best latent axis for sample-level S_epi-age: {axis_name}
- Pearson correlation with sample-level S_epi-age: {summary['best_axis_pearson_with_s_epi_age_sample']:.4f}
- Spearman correlation with sample-level S_epi-age: {summary['best_axis_spearman_with_s_epi_age_sample']:.4f}
- Morula latent low-age rank: {summary['morula_latent_low_age_rank']}
- Morula S_epi-age rank: {summary['morula_s_epi_age_rank']}
- Largest centroid transition in latent space: {summary['best_latent_transition']}

## Representation sensitivity

The direct beta-methylation profile PCA does not make morula a unique latent cluster. However, entropy-profile PCA provides a better alignment with sample-level age-associated methylation entropy and places morula in the low age-entropy latent window.

## Interpretation

This provides a light unsupervised validation of the TRO-defined ground-zero state. It asks whether age-DMR-derived profiles can recover a low age-entropy latent region near morula without using the final TRO score as input.

This result should be described as supporting evidence only. It is not a VAE result and should not be used to claim that a neural operator or Schrodinger Bridge was trained.
"""
    (notes / "TRO_latent_reset_analysis.md").write_text(note, encoding="utf-8")

    print("Unsupervised latent reset analysis completed.")
    print(json.dumps(summary, indent=2))
    print("Wrote:", tables / "TRO_latent_reset_summary.json")
    print("Wrote:", figures / "TRO_latent_reset_pca_space.png")
    print("Wrote:", figures / "TRO_latent_reset_axis_by_stage.png")


if __name__ == "__main__":
    main()
```
