## Supplemental code for "An entropy-based diagnostic framework for characterizing methylation state dynamics during preimplantation development": 25_experiment13_parental_age_residual_tro_validation.py

```
from pathlib import Path
import csv
import gzip
import json
import math
import urllib.request

import numpy as np
import pandas as pd


ROOT = Path(__file__).resolve().parents[1]
TABLES = ROOT / "tables"
FIGS = ROOT / "figures"
NOTES = ROOT / "notes"
META = ROOT / "metadata"

GSE273723_MATRIX_URL = (
    "https://ftp.ncbi.nlm.nih.gov/geo/series/GSE273nnn/GSE273723/"
    "matrix/GSE273723_series_matrix.txt.gz"
)

AGE_CPG_WEIGHTS = META / "GSE102970_TableS6_age_cpg_weights.tsv"
CACHE_SELECTED_MATRIX = TABLES / "GSE273723_selected_age_DMR_cpg_beta_matrix.tsv.gz"

AGE_GROUP_ORDER = ["Young", "Middle", "Old"]
AGE_GROUP_CODE = {name: i for i, name in enumerate(AGE_GROUP_ORDER)}


def clean_token(value):
    if value is None:
        return ""
    return str(value).strip().strip('"')


def parse_tsv_line(line):
    return [clean_token(x) for x in next(csv.reader([line], delimiter="\t"))]


def download_and_cache_selected_matrix(target_cpgs, force=False):
    if CACHE_SELECTED_MATRIX.exists() and not force:
        return pd.read_csv(CACHE_SELECTED_MATRIX, sep="\t")

    rows = []
    header = None
    n_scanned = 0

    with urllib.request.urlopen(GSE273723_MATRIX_URL, timeout=120) as response:
        with gzip.GzipFile(fileobj=response) as gz:
            in_table = False
            for raw in gz:
                line = raw.decode("utf-8", errors="replace").rstrip("\n")
                if line.startswith("!series_matrix_table_begin"):
                    in_table = True
                    continue
                if not in_table:
                    continue
                if line.startswith("!series_matrix_table_end"):
                    break
                parts = parse_tsv_line(line)
                if not parts:
                    continue
                if header is None:
                    header = parts
                    continue
                n_scanned += 1
                if parts[0] in target_cpgs:
                    rows.append(parts)
                if n_scanned % 100000 == 0:
                    print(f"Scanned {n_scanned} CpG rows; matched {len(rows)} target CpGs")

    if header is None:
        raise RuntimeError("Could not find matrix header in GSE273723 series matrix.")

    matrix = pd.DataFrame(rows, columns=header)
    sample_cols = [c for c in matrix.columns if c != "ID_REF"]
    for col in sample_cols:
        matrix[col] = pd.to_numeric(matrix[col], errors="coerce")

    matrix.to_csv(CACHE_SELECTED_MATRIX, sep="\t", index=False, compression="gzip")
    return matrix


def fetch_geo_metadata():
    sample_titles = []
    sample_accessions = []
    paternal_groups = []
    source_names = []
    data_rows = []

    with urllib.request.urlopen(GSE273723_MATRIX_URL, timeout=120) as response:
        with gzip.GzipFile(fileobj=response) as gz:
            for raw in gz:
                line = raw.decode("utf-8", errors="replace").rstrip("\n")
                if line.startswith("!series_matrix_table_begin"):
                    break
                parts = parse_tsv_line(line)
                if not parts:
                    continue
                key, values = parts[0], parts[1:]
                if key == "!Sample_title":
                    sample_titles = values
                elif key == "!Sample_geo_accession":
                    sample_accessions = values
                elif key == "!Sample_source_name_ch1":
                    source_names = values
                elif key == "!Sample_characteristics_ch1" and values and "paternal age group:" in values[0]:
                    paternal_groups = [v.split(":", 1)[-1].strip() for v in values]
                elif key == "!Sample_data_row_count":
                    data_rows = values

    n = len(sample_accessions)
    if not (sample_titles and paternal_groups and n):
        raise RuntimeError("Missing sample metadata fields in GSE273723.")

    meta = pd.DataFrame(
        {
            "sample_id": sample_accessions,
            "sample_title": sample_titles[:n],
            "source_name": source_names[:n] if source_names else ["Placenta"] * n,
            "paternal_age_group": paternal_groups[:n],
            "paternal_age_group_code": [AGE_GROUP_CODE.get(g, np.nan) for g in paternal_groups[:n]],
            "data_row_count": data_rows[:n] if data_rows else [""] * n,
        }
    )
    return meta


def shannon_beta(beta):
    x = np.asarray(beta, dtype=float)
    x = np.clip(x, 1e-9, 1 - 1e-9)
    return -(x * np.log(x) + (1 - x) * np.log(1 - x))


def mann_whitney_u(x, y):
    x = np.asarray(x, dtype=float)
    y = np.asarray(y, dtype=float)
    x = x[np.isfinite(x)]
    y = y[np.isfinite(y)]
    if len(x) == 0 or len(y) == 0:
        return np.nan, np.nan, np.nan
    values = np.concatenate([x, y])
    order = np.argsort(values, kind="mergesort")
    ranks = np.empty(len(values), dtype=float)
    ranks[order] = np.arange(1, len(values) + 1)
    # Average ranks for ties.
    sorted_values = values[order]
    start = 0
    while start < len(values):
        end = start + 1
        while end < len(values) and sorted_values[end] == sorted_values[start]:
            end += 1
        if end - start > 1:
            avg = (start + 1 + end) / 2
            ranks[order[start:end]] = avg
        start = end
    n1, n2 = len(x), len(y)
    u1 = ranks[:n1].sum() - n1 * (n1 + 1) / 2
    mean_u = n1 * n2 / 2
    sd_u = math.sqrt(n1 * n2 * (n1 + n2 + 1) / 12)
    if sd_u == 0:
        p = np.nan
    else:
        z = (abs(u1 - mean_u) - 0.5) / sd_u
        p = math.erfc(z / math.sqrt(2))
    cliffs = (sum(a > b for a in x for b in y) - sum(a < b for a in x for b in y)) / (n1 * n2)
    return float(u1), float(p), float(cliffs)


def zscore(values):
    arr = np.asarray(values, dtype=float)
    mu = np.nanmean(arr)
    sd = np.nanstd(arr)
    if not np.isfinite(sd) or sd == 0:
        return np.zeros_like(arr)
    return (arr - mu) / sd


def bh_adjust(p_values):
    p = np.asarray(p_values, dtype=float)
    out = np.full(len(p), np.nan)
    finite = np.where(np.isfinite(p))[0]
    if len(finite) == 0:
        return out
    order = finite[np.argsort(p[finite])]
    ranked = p[order] * len(order) / np.arange(1, len(order) + 1)
    ranked = np.minimum.accumulate(ranked[::-1])[::-1]
    out[order] = np.minimum(ranked, 1.0)
    return out


def make_plot(sample_metrics, cpg_metrics):
    FIGS.mkdir(exist_ok=True)
    try:
        import matplotlib.pyplot as plt
    except ModuleNotFoundError:
        return make_svg_plot(sample_metrics, cpg_metrics)

    fig, axes = plt.subplots(1, 2, figsize=(12, 4.8))

    metric = "signed_age_DMR_residual_score"
    grouped = [
        sample_metrics.loc[sample_metrics["paternal_age_group"] == g, metric].dropna().values
        for g in AGE_GROUP_ORDER
    ]
    axes[0].boxplot(grouped, labels=AGE_GROUP_ORDER, showfliers=False)
    for i, vals in enumerate(grouped, start=1):
        if len(vals):
            axes[0].scatter(
                np.full(len(vals), i) + np.linspace(-0.06, 0.06, len(vals)),
                vals,
                s=22,
                alpha=0.75,
            )
    axes[0].set_title("Placenta residual signal at sperm age-DMR CpGs")
    axes[0].set_xlabel("Paternal age group")
    axes[0].set_ylabel("Signed residual score")

    top = cpg_metrics.sort_values("escape_contribution", ascending=False).head(20).copy()
    y = np.arange(len(top))
    axes[1].barh(y, top["escape_contribution"].values, color="#4C78A8")
    axes[1].set_yticks(y)
    axes[1].set_yticklabels(top["cpg_id"].values, fontsize=7)
    axes[1].invert_yaxis()
    axes[1].set_title("Top placenta residual CpGs aligned with sperm age weights")
    axes[1].set_xlabel("Escape contribution")

    fig.tight_layout()
    out_png = FIGS / "Experiment13_parental_age_residual_TRO_validation.png"
    out_pdf = FIGS / "Experiment13_parental_age_residual_TRO_validation.pdf"
    fig.savefig(out_png, dpi=220)
    fig.savefig(out_pdf)
    plt.close(fig)
    return [out_png, out_pdf]


def make_svg_plot(sample_metrics, cpg_metrics):
    metric = "signed_age_DMR_residual_score"
    values = sample_metrics[metric].dropna().to_numpy(dtype=float)
    if len(values) == 0:
        values = np.array([0.0])
    vmin, vmax = float(np.min(values)), float(np.max(values))
    if vmax == vmin:
        vmax = vmin + 1.0

    def y_scale(v):
        return 370 - (float(v) - vmin) / (vmax - vmin) * 250

    def esc(text):
        return (
            str(text)
            .replace("&", "&amp;")
            .replace("<", "&lt;")
            .replace(">", "&gt;")
            .replace('"', "&quot;")
        )

    colors = {"Young": "#4C78A8", "Middle": "#F58518", "Old": "#E45756"}
    elems = [
        '<svg xmlns="http://www.w3.org/2000/svg" width="1200" height="520" viewBox="0 0 1200 520">',
        '<rect width="1200" height="520" fill="white"/>',
        '<text x="40" y="34" font-family="Arial" font-size="20" font-weight="700">Experiment 13: parental-age residual TRO validation</text>',
        '<text x="40" y="62" font-family="Arial" font-size="13" fill="#555">Left: sample residual score by paternal age group. Right: top CpGs aligned with sperm age-DMR direction.</text>',
        '<line x1="70" y1="370" x2="530" y2="370" stroke="#333"/>',
        '<line x1="70" y1="120" x2="70" y2="370" stroke="#333"/>',
        f'<text x="24" y="124" font-family="Arial" font-size="11">{vmax:.4g}</text>',
        f'<text x="24" y="373" font-family="Arial" font-size="11">{vmin:.4g}</text>',
    ]
    for idx, group in enumerate(AGE_GROUP_ORDER):
        sub = sample_metrics[sample_metrics["paternal_age_group"] == group][metric].dropna().to_numpy(dtype=float)
        x0 = 150 + idx * 140
        if len(sub):
            q1, med, q3 = np.percentile(sub, [25, 50, 75])
            ymin, ymax = np.min(sub), np.max(sub)
            elems.append(f'<line x1="{x0}" y1="{y_scale(ymin):.2f}" x2="{x0}" y2="{y_scale(ymax):.2f}" stroke="{colors[group]}" stroke-width="2"/>')
            elems.append(f'<rect x="{x0-28}" y="{y_scale(q3):.2f}" width="56" height="{max(1, y_scale(q1)-y_scale(q3)):.2f}" fill="{colors[group]}" opacity="0.28" stroke="{colors[group]}"/>')
            elems.append(f'<line x1="{x0-32}" y1="{y_scale(med):.2f}" x2="{x0+32}" y2="{y_scale(med):.2f}" stroke="{colors[group]}" stroke-width="3"/>')
            for j, val in enumerate(sub):
                jitter = ((j % 7) - 3) * 4
                elems.append(f'<circle cx="{x0+jitter}" cy="{y_scale(val):.2f}" r="3" fill="{colors[group]}" opacity="0.75"/>')
        elems.append(f'<text x="{x0-28}" y="397" font-family="Arial" font-size="13">{group}</text>')
    elems.append('<text x="165" y="432" font-family="Arial" font-size="14">Paternal age group</text>')
    elems.append('<text transform="translate(18 320) rotate(-90)" font-family="Arial" font-size="14">Signed residual score</text>')

    top = cpg_metrics.sort_values("escape_contribution", ascending=False).head(15).copy()
    max_bar = float(top["escape_contribution"].max()) if len(top) else 1.0
    if max_bar <= 0 or not np.isfinite(max_bar):
        max_bar = 1.0
    elems.append('<line x1="665" y1="390" x2="1140" y2="390" stroke="#333"/>')
    elems.append('<text x="665" y="104" font-family="Arial" font-size="15" font-weight="700">Top residual CpGs</text>')
    for i, (_, row) in enumerate(top.iterrows()):
        y = 126 + i * 17
        width = 400 * float(row["escape_contribution"]) / max_bar
        elems.append(f'<text x="665" y="{y+10}" font-family="Arial" font-size="10">{esc(row["cpg_id"])}</text>')
        elems.append(f'<rect x="760" y="{y}" width="{width:.2f}" height="11" fill="#4C78A8" opacity="0.85"/>')
    elems.append('<text x="790" y="430" font-family="Arial" font-size="14">Escape contribution</text>')
    elems.append("</svg>")

    out_svg = FIGS / "Experiment13_parental_age_residual_TRO_validation.svg"
    out_svg.write_text("\n".join(elems), encoding="utf-8")
    return [out_svg]


def main():
    import argparse

    parser = argparse.ArgumentParser()
    parser.add_argument("--force-download", action="store_true")
    args = parser.parse_args()

    TABLES.mkdir(exist_ok=True)
    FIGS.mkdir(exist_ok=True)
    NOTES.mkdir(exist_ok=True)

    weights = pd.read_csv(AGE_CPG_WEIGHTS, sep="\t")
    weights = weights.dropna(subset=["cpg_id", "age_weight_per_year"]).copy()
    weights["age_weight_per_year"] = pd.to_numeric(weights["age_weight_per_year"], errors="coerce")
    weights = weights.dropna(subset=["age_weight_per_year"])
    target_cpgs = set(weights["cpg_id"].astype(str))

    print(f"Target sperm age-DMR CpGs: {len(target_cpgs)}")
    meta = fetch_geo_metadata()
    meta = meta[meta["paternal_age_group"].isin(AGE_GROUP_ORDER)].copy()
    meta.to_csv(TABLES / "GSE273723_parental_age_placenta_metadata.tsv", sep="\t", index=False)
    print("GSE273723 sample counts:")
    print(meta["paternal_age_group"].value_counts().reindex(AGE_GROUP_ORDER).to_string())

    matrix = download_and_cache_selected_matrix(target_cpgs, force=args.force_download)
    matrix = matrix.rename(columns={"ID_REF": "cpg_id"})
    matrix = weights.merge(matrix, on="cpg_id", how="inner")
    sample_cols = [c for c in matrix.columns if c.startswith("GSM")]
    sample_cols = [c for c in sample_cols if c in set(meta["sample_id"])]
    matrix = matrix[["cpg_id", "cluster_name", "chr", "start", "end", "age_weight_per_year"] + sample_cols]
    matrix.to_csv(TABLES / "GSE273723_sperm_age_DMR_overlap_beta_matrix.tsv.gz", sep="\t", index=False, compression="gzip")

    w = matrix["age_weight_per_year"].to_numpy(dtype=float)
    abs_w = np.abs(w)
    abs_w_sum = abs_w.sum()
    sign_w = np.sign(w)

    sample_rows = []
    for sample in sample_cols:
        beta = matrix[sample].to_numpy(dtype=float)
        ok = np.isfinite(beta)
        if ok.sum() == 0:
            continue
        residual = np.nansum(sign_w[ok] * beta[ok] * abs_w[ok]) / np.nansum(abs_w[ok])
        weighted_beta = np.nansum(w[ok] * beta[ok]) / np.nansum(abs_w[ok])
        weighted_entropy = np.nansum(abs_w[ok] * shannon_beta(beta[ok])) / np.nansum(abs_w[ok])
        sample_rows.append(
            {
                "sample_id": sample,
                "n_age_DMR_CpGs_observed": int(ok.sum()),
                "weighted_signed_beta_score": weighted_beta,
                "signed_age_DMR_residual_score": residual,
                "weighted_age_DMR_entropy": weighted_entropy,
            }
        )

    sample_metrics = pd.DataFrame(sample_rows).merge(meta, on="sample_id", how="left")
    sample_metrics["signed_age_DMR_residual_z"] = zscore(sample_metrics["signed_age_DMR_residual_score"])
    sample_metrics["weighted_age_DMR_entropy_z"] = zscore(sample_metrics["weighted_age_DMR_entropy"])
    sample_metrics.to_csv(TABLES / "GSE273723_parental_age_residual_sample_metrics.tsv", sep="\t", index=False)

    stage_rows = []
    for group in AGE_GROUP_ORDER:
        sub = sample_metrics[sample_metrics["paternal_age_group"] == group]
        stage_rows.append(
            {
                "paternal_age_group": group,
                "n_samples": len(sub),
                "signed_age_DMR_residual_mean": sub["signed_age_DMR_residual_score"].mean(),
                "signed_age_DMR_residual_median": sub["signed_age_DMR_residual_score"].median(),
                "weighted_age_DMR_entropy_mean": sub["weighted_age_DMR_entropy"].mean(),
                "weighted_signed_beta_mean": sub["weighted_signed_beta_score"].mean(),
            }
        )
    group_metrics = pd.DataFrame(stage_rows)
    group_metrics.to_csv(TABLES / "GSE273723_parental_age_residual_group_metrics.tsv", sep="\t", index=False)

    young = sample_metrics[sample_metrics["paternal_age_group"] == "Young"]
    old = sample_metrics[sample_metrics["paternal_age_group"] == "Old"]
    test_rows = []
    for metric in ["signed_age_DMR_residual_score", "weighted_age_DMR_entropy", "weighted_signed_beta_score"]:
        u, p, cliffs = mann_whitney_u(old[metric].values, young[metric].values)
        test_rows.append(
            {
                "metric": metric,
                "comparison": "Old vs Young paternal age",
                "n_old": len(old),
                "n_young": len(young),
                "old_mean": old[metric].mean(),
                "young_mean": young[metric].mean(),
                "old_minus_young": old[metric].mean() - young[metric].mean(),
                "u_stat": u,
                "p_value": p,
                "cliffs_delta_old_minus_young": cliffs,
            }
        )
    tests = pd.DataFrame(test_rows)
    tests["p_adj_BH"] = bh_adjust(tests["p_value"].values)
    tests.to_csv(TABLES / "GSE273723_parental_age_residual_group_tests.tsv", sep="\t", index=False)

    cpg_rows = []
    for _, row in matrix.iterrows():
        vals = {}
        for group in AGE_GROUP_ORDER:
            ids = meta.loc[meta["paternal_age_group"] == group, "sample_id"].tolist()
            present = [x for x in ids if x in matrix.columns]
            vals[group] = pd.to_numeric(row[present], errors="coerce").astype(float).mean()
        delta = vals["Old"] - vals["Young"]
        direction = np.sign(row["age_weight_per_year"])
        aligned = np.sign(delta) == direction if np.isfinite(delta) and direction != 0 else False
        cpg_rows.append(
            {
                "cpg_id": row["cpg_id"],
                "cluster_name": row["cluster_name"],
                "chr": row["chr"],
                "start": row["start"],
                "end": row["end"],
                "sperm_age_weight_per_year": row["age_weight_per_year"],
                "young_beta_mean": vals["Young"],
                "middle_beta_mean": vals["Middle"],
                "old_beta_mean": vals["Old"],
                "old_minus_young_beta": delta,
                "aligned_with_sperm_age_direction": bool(aligned),
                "escape_contribution": abs(row["age_weight_per_year"]) * max(direction * delta, 0)
                if np.isfinite(delta)
                else np.nan,
                "absolute_delta": abs(delta) if np.isfinite(delta) else np.nan,
            }
        )
    cpg_metrics = pd.DataFrame(cpg_rows)
    cpg_metrics = cpg_metrics.sort_values("escape_contribution", ascending=False)
    cpg_metrics.to_csv(TABLES / "GSE273723_parental_age_residual_CpG_escape_ranking.tsv", sep="\t", index=False)
    cpg_metrics.head(50).to_csv(
        TABLES / "GSE273723_top50_sperm_age_DMR_placenta_escape_CpGs.tsv", sep="\t", index=False
    )

    aligned_fraction = float(cpg_metrics["aligned_with_sperm_age_direction"].mean())
    mean_signed_delta = float(
        np.nanmean(np.sign(cpg_metrics["sperm_age_weight_per_year"]) * cpg_metrics["old_minus_young_beta"])
    )

    figure_paths = make_plot(sample_metrics, cpg_metrics)

    residual_test = tests.loc[tests["metric"] == "signed_age_DMR_residual_score"].iloc[0]
    if residual_test["old_minus_young"] > 0 and residual_test["p_adj_BH"] < 0.05:
        conclusion = "offspring_placenta_retains_directional_paternal_age_DMR_residual"
    elif residual_test["old_minus_young"] > 0:
        conclusion = "directional_but_not_formally_significant_paternal_age_DMR_residual"
    else:
        conclusion = "no_directional_paternal_age_DMR_residual_detected"

    summary = {
        "dataset": "GSE273723",
        "source": GSE273723_MATRIX_URL,
        "biological_scope": "offspring placenta methylation, not preimplantation embryo",
        "target_sperm_age_DMR_CpGs": int(len(target_cpgs)),
        "observed_target_CpGs_in_EPIC_matrix": int(len(matrix)),
        "sample_counts": meta["paternal_age_group"].value_counts().reindex(AGE_GROUP_ORDER).fillna(0).astype(int).to_dict(),
        "old_minus_young_signed_residual": float(residual_test["old_minus_young"]),
        "old_vs_young_signed_residual_BH_p": float(residual_test["p_adj_BH"]),
        "old_vs_young_signed_residual_cliffs_delta": float(residual_test["cliffs_delta_old_minus_young"]),
        "CpG_direction_alignment_fraction": aligned_fraction,
        "mean_sign_weighted_old_minus_young_beta": mean_signed_delta,
        "top_escape_CpGs": cpg_metrics.head(10)["cpg_id"].tolist(),
        "conclusion": conclusion,
        "claim_boundary": (
            "This is a transgenerational residual validation using placenta methylation. "
            "It does not prove sperm-to-preimplantation-embryo paired reset."
        ),
    }
    with open(TABLES / "GSE273723_parental_age_residual_TRO_summary.json", "w", encoding="utf-8") as fh:
        json.dump(summary, fh, indent=2)

    note = f"""# Experiment 13: parental-age residual TRO validation

This experiment tests whether CpGs derived from paternal age-associated sperm DMRs show a directional residual signal in offspring placenta methylation.

Dataset: GSE273723, processed GEO series matrix. Biological scope: offspring placenta, not preimplantation embryo.

Core result:

- Observed sperm age-DMR CpGs in placenta EPIC matrix: {len(matrix)}
- Old vs young paternal-age signed residual difference: {summary['old_minus_young_signed_residual']:.6g}
- BH-adjusted p value: {summary['old_vs_young_signed_residual_BH_p']:.6g}
- CpG direction-alignment fraction: {summary['CpG_direction_alignment_fraction']:.3f}
- Conclusion code: `{summary['conclusion']}`

Interpretation boundary:

This can support a transgenerational residual-signal layer of TRO, because the input variable is paternal age group and the output is offspring placenta methylation at sperm age-DMR CpGs. It should not be described as a direct paired sperm-to-embryo reset operator.

Generated outputs:

- `GSE273723_parental_age_placenta_metadata.tsv`
- `GSE273723_parental_age_residual_sample_metrics.tsv`
- `GSE273723_parental_age_residual_group_metrics.tsv`
- `GSE273723_parental_age_residual_group_tests.tsv`
- `GSE273723_parental_age_residual_CpG_escape_ranking.tsv`
- `GSE273723_top50_sperm_age_DMR_placenta_escape_CpGs.tsv`
- `GSE273723_parental_age_residual_TRO_summary.json`
- `Experiment13_parental_age_residual_TRO_validation` figure (`png/pdf` if matplotlib is installed; otherwise `svg`)
"""
    (NOTES / "Experiment13_parental_age_residual_TRO_validation.md").write_text(note, encoding="utf-8")

    print("Experiment 13 parental-age residual TRO validation:")
    print(json.dumps(summary, indent=2))
    for path in [
        TABLES / "GSE273723_parental_age_placenta_metadata.tsv",
        TABLES / "GSE273723_parental_age_residual_sample_metrics.tsv",
        TABLES / "GSE273723_parental_age_residual_group_metrics.tsv",
        TABLES / "GSE273723_parental_age_residual_group_tests.tsv",
        TABLES / "GSE273723_parental_age_residual_CpG_escape_ranking.tsv",
        TABLES / "GSE273723_top50_sperm_age_DMR_placenta_escape_CpGs.tsv",
        TABLES / "GSE273723_parental_age_residual_TRO_summary.json",
        *figure_paths,
        NOTES / "Experiment13_parental_age_residual_TRO_validation.md",
    ]:
        print(f"Wrote: {path}")


if __name__ == "__main__":
    main()
```
