## Supplemental code for "An entropy-based diagnostic framework for characterizing methylation state dynamics during preimplantation development": 26_experiment14_gse56697_paired_paternal_gamete_to_embryo_operator.py

```
from pathlib import Path
import gzip
import json
import math
import urllib.request

import numpy as np
import pandas as pd


ROOT = Path(__file__).resolve().parents[1]
RAW = ROOT / "data_raw" / "GSE56697_parental_methylome"
TABLES = ROOT / "tables"
FIGS = ROOT / "figures"
NOTES = ROOT / "notes"

BASE = "https://ftp.ncbi.nlm.nih.gov/geo/samples"

SAMPLES = [
    {
        "stage": "sperm",
        "role": "paternal_gamete_input",
        "gsm": "GSM1386020",
        "file": "GSM1386020_sperm_mc_CG_plus.bed.gz",
        "url": f"{BASE}/GSM1386nnn/GSM1386020/suppl/GSM1386020_sperm_mc_CG_plus.bed.gz",
    },
    {
        "stage": "2-cell paternal",
        "role": "paternal_embryo_output",
        "gsm": "GSM1386021",
        "file": "GSM1386021_2cell_mc_CG_paternal_plus.bed.gz",
        "url": f"{BASE}/GSM1386nnn/GSM1386021/suppl/GSM1386021_2cell_mc_CG_paternal_plus.bed.gz",
    },
    {
        "stage": "4-cell paternal",
        "role": "paternal_embryo_output",
        "gsm": "GSM1386022",
        "file": "GSM1386022_4cell_mc_CG_paternal_plus.bed.gz",
        "url": f"{BASE}/GSM1386nnn/GSM1386022/suppl/GSM1386022_4cell_mc_CG_paternal_plus.bed.gz",
    },
    {
        "stage": "ICM paternal",
        "role": "paternal_embryo_output",
        "gsm": "GSM1386023",
        "file": "GSM1386023_ICM_mc_CG_paternal_plus.bed.gz",
        "url": f"{BASE}/GSM1386nnn/GSM1386023/suppl/GSM1386023_ICM_mc_CG_paternal_plus.bed.gz",
    },
    {
        "stage": "E6.5 paternal",
        "role": "paternal_embryo_output",
        "gsm": "GSM1386024",
        "file": "GSM1386024_E65_mc_CG_paternal_plus.bed.gz",
        "url": f"{BASE}/GSM1386nnn/GSM1386024/suppl/GSM1386024_E65_mc_CG_paternal_plus.bed.gz",
    },
    {
        "stage": "E7.5 paternal",
        "role": "paternal_embryo_output",
        "gsm": "GSM1386025",
        "file": "GSM1386025_E75_mc_CG_paternal_plus.bed.gz",
        "url": f"{BASE}/GSM1386nnn/GSM1386025/suppl/GSM1386025_E75_mc_CG_paternal_plus.bed.gz",
    },
]

STAGE_ORDER = [x["stage"] for x in SAMPLES]
EMBRYO_STAGES = [x["stage"] for x in SAMPLES if x["role"] == "paternal_embryo_output"]


def ensure_dirs():
    for path in [RAW, TABLES, FIGS, NOTES]:
        path.mkdir(parents=True, exist_ok=True)


def download_if_missing(sample):
    dest = RAW / sample["file"]
    if dest.exists() and dest.stat().st_size > 1000:
        return dest

    print(f"Downloading {sample['file']}")
    req = urllib.request.Request(sample["url"], headers={"User-Agent": "TRO_Project/1.0"})
    with urllib.request.urlopen(req, timeout=180) as response, open(dest, "wb") as out:
        total = response.headers.get("Content-Length")
        total = int(total) if total else None
        done = 0
        while True:
            chunk = response.read(1024 * 1024)
            if not chunk:
                break
            out.write(chunk)
            done += len(chunk)
            if total and done % (25 * 1024 * 1024) < 1024 * 1024:
                print(f"  {sample['file']}: {done / total:.1%}")
    return dest


def parse_beta_and_weight(parts):
    if len(parts) < 4:
        return None, None
    try:
        beta = float(parts[3])
    except ValueError:
        return None, None
    if beta > 1.0:
        beta = beta / 100.0
    beta = max(0.0, min(1.0, beta))

    weight = 1.0
    if len(parts) >= 6:
        try:
            c_count = float(parts[-2])
            t_count = float(parts[-1])
            cov = c_count + t_count
            if cov > 0:
                weight = cov
        except ValueError:
            pass
    return beta, weight


def summarize_bed_by_window(path, bin_size=500_000):
    sums = {}
    weights = {}
    counts = {}
    n_lines = 0

    with gzip.open(path, "rt", encoding="utf-8", errors="replace") as handle:
        for line in handle:
            if not line or line.startswith("#"):
                continue
            parts = line.rstrip("\n").split("\t")
            if len(parts) < 4:
                continue
            chrom = parts[0]
            if chrom.startswith("chrM") or chrom in {"M", "MT"}:
                continue
            try:
                start = int(float(parts[1]))
            except ValueError:
                continue
            beta, weight = parse_beta_and_weight(parts)
            if beta is None:
                continue
            key = f"{chrom}:{(start // bin_size) * bin_size}"
            sums[key] = sums.get(key, 0.0) + beta * weight
            weights[key] = weights.get(key, 0.0) + weight
            counts[key] = counts.get(key, 0) + 1
            n_lines += 1
            if n_lines % 5_000_000 == 0:
                print(f"  parsed {path.name}: {n_lines:,} CpGs")

    rows = []
    for key, total_weight in weights.items():
        if total_weight <= 0:
            continue
        rows.append(
            {
                "window": key,
                "beta": sums[key] / total_weight,
                "coverage_weight": total_weight,
                "n_cpg": counts[key],
            }
        )
    return pd.DataFrame(rows)


def entropy01(values):
    x = np.clip(np.asarray(values, dtype=float), 1e-12, 1 - 1e-12)
    return -(x * np.log(x) + (1 - x) * np.log(1 - x))


def zscore(values):
    x = np.asarray(values, dtype=float)
    sd = x.std(ddof=0)
    if sd == 0 or not np.isfinite(sd):
        return np.zeros_like(x)
    return (x - x.mean()) / sd


def run(bin_size=500_000, min_cpg_per_window=3):
    ensure_dirs()

    manifest_rows = []
    matrices = {}
    for sample in SAMPLES:
        path = download_if_missing(sample)
        print(f"Summarizing {sample['stage']} from {path.name}")
        df = summarize_bed_by_window(path, bin_size=bin_size)
        df = df[df["n_cpg"] >= min_cpg_per_window].copy()
        matrices[sample["stage"]] = df.set_index("window")["beta"]
        manifest_rows.append(
            {
                **sample,
                "local_path": str(path),
                "size_bytes": path.stat().st_size,
                "windows_after_filter": int(len(df)),
            }
        )

    common = set(matrices[STAGE_ORDER[0]].index)
    for stage in STAGE_ORDER[1:]:
        common &= set(matrices[stage].index)
    common = sorted(common)
    if len(common) < 100:
        raise RuntimeError(f"Too few common windows after filtering: {len(common)}")

    matrix = pd.DataFrame({"window": common})
    for stage in STAGE_ORDER:
        matrix[stage] = matrices[stage].loc[common].values

    sperm = matrix["sperm"].to_numpy(dtype=float)
    embryo_means = {stage: float(matrix[stage].mean()) for stage in EMBRYO_STAGES}
    min_embryo_mean = min(embryo_means.values())
    sperm_mean = float(matrix["sperm"].mean())
    denom = sperm_mean - min_embryo_mean

    stage_rows = []
    for stage in STAGE_ORDER:
        x = matrix[stage].to_numpy(dtype=float)
        mean_beta = float(np.nanmean(x))
        l1 = float(np.nanmean(np.abs(x - sperm)))
        l2 = float(np.linalg.norm(x - sperm) / math.sqrt(len(x)))
        demethylation = float(np.nanmean(np.maximum(sperm - x, 0.0)))
        hypermethylation = float(np.nanmean(np.maximum(x - sperm, 0.0)))
        reset_score = 0.0 if stage == "sperm" else ((sperm_mean - mean_beta) / denom if denom else np.nan)
        stage_rows.append(
            {
                "stage": stage,
                "role": next(s["role"] for s in SAMPLES if s["stage"] == stage),
                "n_common_windows": len(common),
                "mean_methylation": mean_beta,
                "methylation_entropy": float(np.nanmean(entropy01(x))),
                "distance_to_sperm_L1": l1,
                "distance_to_sperm_L2": l2,
                "demethylation_depth_from_sperm": demethylation,
                "hypermethylation_from_sperm": hypermethylation,
                "paternal_reset_score": reset_score,
            }
        )

    stage_metrics = pd.DataFrame(stage_rows)
    stage_metrics["low_methylation_rank"] = stage_metrics["mean_methylation"].rank(method="min", ascending=True).astype(int)
    stage_metrics["reset_score_rank"] = stage_metrics["paternal_reset_score"].rank(method="min", ascending=False).astype(int)

    transitions = []
    for a, b in zip(STAGE_ORDER[:-1], STAGE_ORDER[1:]):
        va = matrix[a].to_numpy(dtype=float)
        vb = matrix[b].to_numpy(dtype=float)
        cost = float(np.linalg.norm(zscore(vb) - zscore(va)) / math.sqrt(len(common)))
        mean_a = float(np.nanmean(va))
        mean_b = float(np.nanmean(vb))
        methylation_drop = mean_a - mean_b
        transitions.append(
            {
                "transition": f"{a} -> {b}",
                "stage_from": a,
                "stage_to": b,
                "transition_cost_zL2": cost,
                "mean_methylation_from": mean_a,
                "mean_methylation_to": mean_b,
                "methylation_drop": methylation_drop,
                "productive_demethylation_gain": max(methylation_drop, 0.0),
                "reset_efficiency": max(methylation_drop, 0.0) / cost if cost > 0 else np.nan,
            }
        )
    transition_metrics = pd.DataFrame(transitions)
    transition_metrics["cost_rank"] = transition_metrics["transition_cost_zL2"].rank(method="min", ascending=False).astype(int)
    transition_metrics["efficiency_rank"] = transition_metrics["reset_efficiency"].rank(method="min", ascending=False).astype(int)

    embryo_stage_metrics = stage_metrics[stage_metrics["stage"].isin(EMBRYO_STAGES)].copy()
    ground_zero_stage = embryo_stage_metrics.sort_values(["mean_methylation", "distance_to_sperm_L1"]).iloc[0]["stage"]
    best_transition = transition_metrics.sort_values("reset_efficiency", ascending=False).iloc[0]["transition"]

    summary = {
        "dataset": "GSE56697",
        "source": "Programming and inheritance of parental DNA methylomes in mammals",
        "operator_type": "paired_paternal_gamete_to_embryo_methylome_operator",
        "species": "Mus musculus",
        "paternal_gamete_input": "DBA/2J sperm methylome",
        "paternal_embryo_outputs": EMBRYO_STAGES,
        "bin_size_bp": bin_size,
        "min_cpg_per_window": min_cpg_per_window,
        "n_common_windows": len(common),
        "ground_zero_stage_by_min_paternal_methylation": ground_zero_stage,
        "best_demethylation_transition": best_transition,
        "sperm_mean_methylation": sperm_mean,
        "minimum_embryo_mean_methylation": float(embryo_stage_metrics["mean_methylation"].min()),
        "max_paternal_reset_score": float(embryo_stage_metrics["paternal_reset_score"].max()),
        "claim_boundary": (
            "This is a paired mouse paternal-gamete-to-paternal-embryo methylome operator pilot, "
            "not a human paternal-age paired embryo experiment."
        ),
    }

    manifest = pd.DataFrame(manifest_rows)
    manifest.to_csv(TABLES / "GSE56697_paired_paternal_operator_manifest.tsv", sep="\t", index=False)
    matrix.to_csv(TABLES / "GSE56697_paired_paternal_operator_window_matrix.tsv.gz", sep="\t", index=False, compression="gzip")
    stage_metrics.to_csv(TABLES / "GSE56697_paired_paternal_operator_stage_metrics.tsv", sep="\t", index=False)
    transition_metrics.to_csv(TABLES / "GSE56697_paired_paternal_operator_transition_metrics.tsv", sep="\t", index=False)
    with open(TABLES / "GSE56697_paired_paternal_operator_summary.json", "w", encoding="utf-8") as handle:
        json.dump(summary, handle, indent=2, ensure_ascii=False)

    make_figure(stage_metrics, transition_metrics)
    write_note(summary, stage_metrics, transition_metrics)

    print("GSE56697 paired paternal gamete-to-embryo reset operator summary:")
    print(json.dumps(summary, indent=2, ensure_ascii=False))
    print(stage_metrics.to_string(index=False))
    print(transition_metrics.to_string(index=False))


def make_figure(stage_metrics, transition_metrics):
    try:
        import matplotlib.pyplot as plt
    except ModuleNotFoundError:
        make_svg_figure(stage_metrics, transition_metrics)
        return

    order = STAGE_ORDER
    stage_metrics = stage_metrics.set_index("stage").loc[order].reset_index()

    fig, axes = plt.subplots(1, 2, figsize=(12, 4.8))
    ax = axes[0]
    ax.plot(stage_metrics["stage"], stage_metrics["mean_methylation"], marker="o", label="mean paternal methylation")
    ax.plot(stage_metrics["stage"], stage_metrics["paternal_reset_score"], marker="s", label="paternal reset score")
    ax.set_xticklabels(stage_metrics["stage"], rotation=35, ha="right")
    ax.set_ylabel("Value")
    ax.set_title("Paired paternal gamete-to-embryo state")
    ax.legend(frameon=False)

    ax = axes[1]
    ax.barh(transition_metrics["transition"], transition_metrics["reset_efficiency"], color="#4C78A8")
    ax.invert_yaxis()
    ax.set_xlabel("Demethylation gain / transition cost")
    ax.set_title("Reset transition efficiency")

    fig.tight_layout()
    fig.savefig(FIGS / "GSE56697_paired_paternal_reset_operator.png", dpi=220)
    fig.savefig(FIGS / "GSE56697_paired_paternal_reset_operator.pdf")
    plt.close(fig)


def make_svg_figure(stage_metrics, transition_metrics):
    order = STAGE_ORDER
    stage_metrics = stage_metrics.set_index("stage").loc[order].reset_index()
    width, height = 1100, 520
    left, top = 85, 55
    plot_w, plot_h = 460, 330
    right_left = 660

    vals = list(stage_metrics["mean_methylation"]) + list(stage_metrics["paternal_reset_score"])
    y_min, y_max = min(vals), max(vals)
    pad = (y_max - y_min) * 0.08 if y_max > y_min else 0.1
    y_min, y_max = y_min - pad, y_max + pad

    def sx(i):
        return left + i * (plot_w / (len(order) - 1))

    def sy(v):
        return top + plot_h - (float(v) - y_min) / (y_max - y_min) * plot_h

    methyl_points = " ".join(f"{sx(i):.1f},{sy(v):.1f}" for i, v in enumerate(stage_metrics["mean_methylation"]))
    reset_points = " ".join(f"{sx(i):.1f},{sy(v):.1f}" for i, v in enumerate(stage_metrics["paternal_reset_score"]))

    bars = []
    max_eff = float(transition_metrics["reset_efficiency"].max())
    for i, row in transition_metrics.iterrows():
        y = top + 30 + i * 44
        bar_w = 330 * (float(row["reset_efficiency"]) / max_eff) if max_eff > 0 else 0
        bars.append(
            f'<text x="{right_left}" y="{y + 16}" font-size="12">{row["transition"]}</text>'
            f'<rect x="{right_left + 210}" y="{y}" width="{bar_w:.1f}" height="20" fill="#4C78A8"/>'
            f'<text x="{right_left + 215 + bar_w:.1f}" y="{y + 15}" font-size="11">{row["reset_efficiency"]:.3f}</text>'
        )

    labels = []
    for i, stage in enumerate(order):
        labels.append(
            f'<text transform="translate({sx(i):.1f},{top + plot_h + 72}) rotate(-35)" '
            f'font-size="12" text-anchor="end">{stage}</text>'
        )

    svg = f'''<svg xmlns="http://www.w3.org/2000/svg" width="{width}" height="{height}" viewBox="0 0 {width} {height}">
<rect width="100%" height="100%" fill="white"/>
<text x="{left}" y="28" font-size="18" font-weight="bold">GSE56697 paired paternal gamete-to-embryo reset operator</text>
<line x1="{left}" y1="{top}" x2="{left}" y2="{top + plot_h}" stroke="#333"/>
<line x1="{left}" y1="{top + plot_h}" x2="{left + plot_w}" y2="{top + plot_h}" stroke="#333"/>
<polyline points="{methyl_points}" fill="none" stroke="#D55E00" stroke-width="3"/>
<polyline points="{reset_points}" fill="none" stroke="#0072B2" stroke-width="3"/>
{''.join(f'<circle cx="{sx(i):.1f}" cy="{sy(v):.1f}" r="4" fill="#D55E00"/>' for i, v in enumerate(stage_metrics["mean_methylation"]))}
{''.join(f'<rect x="{sx(i)-4:.1f}" y="{sy(v)-4:.1f}" width="8" height="8" fill="#0072B2"/>' for i, v in enumerate(stage_metrics["paternal_reset_score"]))}
{''.join(labels)}
<text x="{left}" y="{top + plot_h + 110}" font-size="12" fill="#D55E00">mean paternal methylation</text>
<text x="{left + 210}" y="{top + plot_h + 110}" font-size="12" fill="#0072B2">paternal reset score</text>
<text x="{right_left}" y="28" font-size="18" font-weight="bold">Reset transition efficiency</text>
{''.join(bars)}
</svg>
'''
    (FIGS / "GSE56697_paired_paternal_reset_operator.svg").write_text(svg, encoding="utf-8")


def write_note(summary, stage_metrics, transition_metrics):
    gz = summary["ground_zero_stage_by_min_paternal_methylation"]
    best = summary["best_demethylation_transition"]
    text = f"""# Experiment 14: Paired paternal gamete-to-embryo reset operator pilot

Dataset: GSE56697, mouse parental methylome MethylC-Seq.

This experiment constructs a true paired-direction operator layer from paternal gamete to paternal embryo allele states:

`DBA/2J sperm methylome -> paternal allele methylome in 2-cell, 4-cell, ICM, E6.5, and E7.5 embryos`.

The analysis summarizes CpG methylation into {summary['bin_size_bp']:,} bp genomic windows and keeps windows observed across all paternal states.

Key result:

- common windows: {summary['n_common_windows']}
- lowest paternal embryo methylation state: {gz}
- most efficient demethylation transition: {best}

Interpretation:

This is stronger than the earlier human stage-level TRO prototype because it uses a real parental gamete input and paternal embryo output trajectory. It still is not a human paternal-age paired experiment, so it should be described as a mouse paired paternal-genome reset-operator validation rather than direct proof of aged human sperm reset.

"""
    NOTES.joinpath("Experiment14_GSE56697_paired_paternal_reset_operator.md").write_text(text, encoding="utf-8")


if __name__ == "__main__":
    run()
```
