## Supplemental code for "An entropy-based diagnostic framework for characterizing methylation state dynamics during preimplantation development": 28_experiment16_gse56697_maternal_paternal_branch_contrast.py

```
from pathlib import Path
import gzip
import json
import math
import urllib.request

import numpy as np
import pandas as pd


ROOT = Path(__file__).resolve().parents[1]
RAW = ROOT / "data_raw" / "GSE56697_parental_methylome"
TABLES = ROOT / "tables"
FIGS = ROOT / "figures"
NOTES = ROOT / "notes"

BIN_SIZE = 500_000
MIN_CPG_PER_WINDOW = 3
BASE = "https://ftp.ncbi.nlm.nih.gov/geo/samples/GSM1386nnn"

PATERNAL = [
    ("sperm", "gamete_input", "GSM1386020_sperm_mc_CG_plus.bed.gz"),
    ("2-cell paternal", "embryo_output", "GSM1386021_2cell_mc_CG_paternal_plus.bed.gz"),
    ("4-cell paternal", "embryo_output", "GSM1386022_4cell_mc_CG_paternal_plus.bed.gz"),
    ("ICM paternal", "embryo_output", "GSM1386023_ICM_mc_CG_paternal_plus.bed.gz"),
    ("E6.5 paternal", "embryo_output", "GSM1386024_E65_mc_CG_paternal_plus.bed.gz"),
    ("E7.5 paternal", "embryo_output", "GSM1386025_E75_mc_CG_paternal_plus.bed.gz"),
]

MATERNAL = [
    ("oocyte", "gamete_input", "GSM1386019_oocyte_mc_CG_plus.bed.gz"),
    ("2-cell maternal", "embryo_output", "GSM1386021_2cell_mc_CG_maternal_plus.bed.gz"),
    ("4-cell maternal", "embryo_output", "GSM1386022_4cell_mc_CG_maternal_plus.bed.gz"),
    ("ICM maternal", "embryo_output", "GSM1386023_ICM_mc_CG_maternal_plus.bed.gz"),
    ("E6.5 maternal", "embryo_output", "GSM1386024_E65_mc_CG_maternal_plus.bed.gz"),
    ("E7.5 maternal", "embryo_output", "GSM1386025_E75_mc_CG_maternal_plus.bed.gz"),
]


def ensure_dirs():
    for path in [RAW, TABLES, FIGS, NOTES]:
        path.mkdir(parents=True, exist_ok=True)


def url_for(file_name):
    gsm = file_name.split("_", 1)[0]
    return f"{BASE}/{gsm}/suppl/{file_name}"


def download_if_missing(file_name):
    path = RAW / file_name
    if path.exists() and path.stat().st_size > 1000:
        return path
    url = url_for(file_name)
    print(f"Downloading {file_name}")
    req = urllib.request.Request(url, headers={"User-Agent": "TRO_Project/1.0"})
    with urllib.request.urlopen(req, timeout=180) as response, open(path, "wb") as out:
        while True:
            chunk = response.read(1024 * 1024)
            if not chunk:
                break
            out.write(chunk)
    return path


def parse_beta_and_weight(parts):
    if len(parts) < 4:
        return None, None
    try:
        beta = float(parts[3])
    except ValueError:
        return None, None
    if beta > 1.0:
        beta = beta / 100.0
    beta = max(0.0, min(1.0, beta))
    weight = 1.0
    if len(parts) >= 6:
        try:
            cov = float(parts[-2]) + float(parts[-1])
            if cov > 0:
                weight = cov
        except ValueError:
            pass
    return beta, weight


def summarize(path):
    sums = {}
    weights = {}
    counts = {}
    n = 0
    with gzip.open(path, "rt", encoding="utf-8", errors="replace") as handle:
        for line in handle:
            if not line or line.startswith("#"):
                continue
            parts = line.rstrip("\n").split("\t")
            if len(parts) < 4:
                continue
            chrom = parts[0]
            if chrom.startswith("chrM") or chrom in {"M", "MT"}:
                continue
            try:
                start = int(float(parts[1]))
            except ValueError:
                continue
            beta, weight = parse_beta_and_weight(parts)
            if beta is None:
                continue
            key = f"{chrom}:{(start // BIN_SIZE) * BIN_SIZE}"
            sums[key] = sums.get(key, 0.0) + beta * weight
            weights[key] = weights.get(key, 0.0) + weight
            counts[key] = counts.get(key, 0) + 1
            n += 1
            if n % 5_000_000 == 0:
                print(f"  parsed {path.name}: {n:,} CpGs")
    rows = []
    for key, weight in weights.items():
        cpg = counts[key]
        if weight > 0 and cpg >= MIN_CPG_PER_WINDOW:
            rows.append((key, sums[key] / weight, weight, cpg))
    return pd.DataFrame(rows, columns=["window", "beta", "coverage_weight", "n_cpg"])


def zscore(values):
    x = np.asarray(values, dtype=float)
    sd = x.std(ddof=0)
    if sd == 0 or not np.isfinite(sd):
        return np.zeros_like(x)
    return (x - x.mean()) / sd


def compute_branch(branch_name, samples):
    tables = {}
    for stage, _, file_name in samples:
        path = download_if_missing(file_name)
        print(f"Summarizing {branch_name} {stage}: {file_name}")
        tables[stage] = summarize(path).set_index("window")["beta"]

    stages = [s[0] for s in samples]
    input_stage = stages[0]
    embryo_stages = stages[1:]
    common = set(tables[input_stage].index)
    for stage in stages[1:]:
        common &= set(tables[stage].index)
    common = sorted(common)
    if len(common) < 100:
        raise RuntimeError(f"Too few common windows for {branch_name}: {len(common)}")

    matrix = pd.DataFrame({"window": common})
    for stage in stages:
        matrix[stage] = tables[stage].loc[common].values

    input_vec = matrix[input_stage].to_numpy(dtype=float)
    input_mean = float(np.nanmean(input_vec))
    embryo_means = {stage: float(np.nanmean(matrix[stage])) for stage in embryo_stages}
    minimum_stage = min(embryo_means, key=embryo_means.get)
    minimum_mean = embryo_means[minimum_stage]
    denom = input_mean - minimum_mean

    stage_rows = []
    for stage in stages:
        x = matrix[stage].to_numpy(dtype=float)
        mean_methylation = float(np.nanmean(x))
        reset_score = 0.0 if stage == input_stage else ((input_mean - mean_methylation) / denom if denom else np.nan)
        stage_rows.append(
            {
                "branch": branch_name,
                "stage": stage,
                "role": "gamete_input" if stage == input_stage else "embryo_output",
                "n_common_windows": len(common),
                "mean_methylation": mean_methylation,
                "distance_to_gamete_L1": float(np.nanmean(np.abs(x - input_vec))),
                "distance_to_gamete_L2": float(np.linalg.norm(x - input_vec) / math.sqrt(len(common))),
                "branch_reset_score": reset_score,
            }
        )

    transition_rows = []
    for a, b in zip(stages[:-1], stages[1:]):
        va = matrix[a].to_numpy(dtype=float)
        vb = matrix[b].to_numpy(dtype=float)
        cost = float(np.linalg.norm(zscore(vb) - zscore(va)) / math.sqrt(len(common)))
        drop = float(np.nanmean(va) - np.nanmean(vb))
        transition_rows.append(
            {
                "branch": branch_name,
                "transition": f"{a} -> {b}",
                "transition_cost_zL2": cost,
                "methylation_drop": drop,
                "productive_demethylation_gain": max(drop, 0.0),
                "reset_efficiency": max(drop, 0.0) / cost if cost > 0 else np.nan,
            }
        )

    stage_metrics = pd.DataFrame(stage_rows)
    transition_metrics = pd.DataFrame(transition_rows)
    summary = {
        "branch": branch_name,
        "gamete_input": input_stage,
        "n_common_windows": int(len(common)),
        "input_mean_methylation": input_mean,
        "minimum_embryo_stage": minimum_stage,
        "minimum_embryo_mean_methylation": float(minimum_mean),
        "best_demethylation_transition": str(transition_metrics.sort_values("reset_efficiency", ascending=False).iloc[0]["transition"]),
    }
    return stage_metrics, transition_metrics, summary


def make_svg(stage_all, summary_table):
    branch_stages = {
        "paternal": [s[0] for s in PATERNAL],
        "maternal": [s[0] for s in MATERNAL],
    }
    width, height = 1100, 520
    left, top, plot_w, plot_h = 90, 55, 760, 320
    values = stage_all["mean_methylation"].tolist()
    y_min, y_max = min(values), max(values)
    pad = (y_max - y_min) * 0.08
    y_min -= pad
    y_max += pad

    def sx(i):
        return left + i * (plot_w / 5)

    def sy(v):
        return top + plot_h - (float(v) - y_min) / (y_max - y_min) * plot_h

    lines = []
    for branch, color in [("paternal", "#0072B2"), ("maternal", "#D55E00")]:
        sub = stage_all[stage_all["branch"] == branch].set_index("stage").loc[branch_stages[branch]].reset_index()
        points = " ".join(f"{sx(i):.1f},{sy(v):.1f}" for i, v in enumerate(sub["mean_methylation"]))
        lines.append(f'<polyline points="{points}" fill="none" stroke="{color}" stroke-width="3"/>')
        lines.extend(f'<circle cx="{sx(i):.1f}" cy="{sy(v):.1f}" r="4" fill="{color}"/>' for i, v in enumerate(sub["mean_methylation"]))

    labels = ["gamete", "2-cell", "4-cell", "ICM", "E6.5", "E7.5"]
    label_svg = "".join(f'<text x="{sx(i):.1f}" y="{top + plot_h + 35}" font-size="13" text-anchor="middle">{label}</text>' for i, label in enumerate(labels))
    table = []
    x0, y0 = 875, 90
    for i, row in summary_table.iterrows():
        y = y0 + i * 95
        table.append(f'<text x="{x0}" y="{y}" font-size="14" font-weight="bold">{row["branch"]}</text>')
        table.append(f'<text x="{x0}" y="{y + 25}" font-size="12">input: {row["gamete_input"]}</text>')
        table.append(f'<text x="{x0}" y="{y + 45}" font-size="12">minimum: {row["minimum_embryo_stage"]}</text>')
        table.append(f'<text x="{x0}" y="{y + 65}" font-size="12">best: {row["best_demethylation_transition"]}</text>')

    svg = f'''<svg xmlns="http://www.w3.org/2000/svg" width="{width}" height="{height}" viewBox="0 0 {width} {height}">
<rect width="100%" height="100%" fill="white"/>
<text x="{left}" y="28" font-size="20" font-weight="bold">GSE56697 maternal/oocyte versus paternal/sperm methylome reset branches</text>
<line x1="{left}" y1="{top}" x2="{left}" y2="{top + plot_h}" stroke="#333"/>
<line x1="{left}" y1="{top + plot_h}" x2="{left + plot_w}" y2="{top + plot_h}" stroke="#333"/>
<text x="16" y="{top + 160}" font-size="14" transform="rotate(-90 16,{top + 160})">mean methylation</text>
{''.join(lines)}
{label_svg}
<line x1="{left}" y1="{top + plot_h + 70}" x2="{left + 40}" y2="{top + plot_h + 70}" stroke="#0072B2" stroke-width="4"/>
<text x="{left + 50}" y="{top + plot_h + 75}" font-size="13">paternal branch: sperm -> paternal allele</text>
<line x1="{left + 310}" y1="{top + plot_h + 70}" x2="{left + 350}" y2="{top + plot_h + 70}" stroke="#D55E00" stroke-width="4"/>
<text x="{left + 360}" y="{top + plot_h + 75}" font-size="13">maternal branch: oocyte -> maternal allele</text>
<text x="{x0}" y="55" font-size="16" font-weight="bold">Branch summaries</text>
{''.join(table)}
</svg>
'''
    (FIGS / "GSE56697_maternal_paternal_branch_contrast.svg").write_text(svg, encoding="utf-8")


def write_note(summary_table):
    text = """# Experiment 16: GSE56697 maternal/oocyte versus paternal/sperm branch contrast

This experiment adds the optional parental-allele contrast requested for the paired GSE56697 operator.

It compares:

- paternal branch: sperm -> paternal allele embryo methylomes
- maternal branch: oocyte -> maternal allele embryo methylomes

The purpose is not to force both parental branches to show the same reset pattern. Instead, the analysis tests whether the paternal reset operator is a parental-allele-specific trajectory rather than an artifact of using only embryo stage averages.

"""
    header = "| " + " | ".join(summary_table.columns) + " |\n"
    divider = "| " + " | ".join(["---"] * len(summary_table.columns)) + " |\n"
    rows = []
    for _, row in summary_table.iterrows():
        rows.append("| " + " | ".join(str(row[col]) for col in summary_table.columns) + " |")
    text += header + divider + "\n".join(rows)
    text += "\n\nClaim boundary: this is a mouse parental-allele methylome contrast and does not prove human paternal-age paired reset.\n"
    (NOTES / "Experiment16_GSE56697_maternal_paternal_branch_contrast.md").write_text(text, encoding="utf-8")


def main():
    ensure_dirs()
    paternal_stage, paternal_transition, paternal_summary = compute_branch("paternal", PATERNAL)
    maternal_stage, maternal_transition, maternal_summary = compute_branch("maternal", MATERNAL)

    stage_all = pd.concat([paternal_stage, maternal_stage], ignore_index=True)
    transition_all = pd.concat([paternal_transition, maternal_transition], ignore_index=True)
    summary_table = pd.DataFrame([paternal_summary, maternal_summary])
    summary = {
        "dataset": "GSE56697",
        "analysis": "maternal/oocyte versus paternal/sperm branch contrast",
        "bin_size_bp": BIN_SIZE,
        "min_cpg_per_window": MIN_CPG_PER_WINDOW,
        "branch_summaries": summary_table.to_dict(orient="records"),
        "claim_boundary": "Mouse parental-allele branch contrast; not human paternal-age paired reset proof.",
    }

    stage_all.to_csv(TABLES / "GSE56697_maternal_paternal_branch_stage_metrics.tsv", sep="\t", index=False)
    transition_all.to_csv(TABLES / "GSE56697_maternal_paternal_branch_transition_metrics.tsv", sep="\t", index=False)
    summary_table.to_csv(TABLES / "GSE56697_maternal_paternal_branch_summary.tsv", sep="\t", index=False)
    with open(TABLES / "GSE56697_maternal_paternal_branch_summary.json", "w", encoding="utf-8") as handle:
        json.dump(summary, handle, indent=2, ensure_ascii=False)

    make_svg(stage_all, summary_table)
    write_note(summary_table)

    print("GSE56697 maternal/paternal branch contrast summary:")
    print(json.dumps(summary, indent=2, ensure_ascii=False))
    print(stage_all.to_string(index=False))
    print(transition_all.to_string(index=False))


if __name__ == "__main__":
    main()
```
