## Supplemental code for "An entropy-based diagnostic framework for characterizing methylation state dynamics during preimplantation development": 30_experiment18_dmr_mechanistic_interpretability.py

````
from pathlib import Path
import json
import math
import html

import pandas as pd


ROOT = Path(__file__).resolve().parents[1]
TABLES = ROOT / "tables"
FIGS = ROOT / "figures"
NOTES = ROOT / "notes"


THEME_KEYWORDS = {
    "development_and_patterning": [
        "development",
        "regionalization",
        "pattern",
        "morphogenesis",
        "organogenesis",
        "cell fate",
        "commitment",
    ],
    "wnt_cadherin_signaling": [
        "wnt",
        "cadherin",
        "cell-cell adhesion",
        "adhesion",
    ],
    "chromatin_epigenetic_regulation": [
        "chromatin",
        "histone",
        "methylation",
        "dna methylation",
        "epigen",
    ],
    "stemness_differentiation": [
        "stem cell",
        "differentiation",
        "pluripot",
        "blastocyst",
        "embryo",
    ],
}


def read_json(path):
    return json.loads(Path(path).read_text(encoding="utf-8"))


def theme_for_term(term_name):
    text = str(term_name).lower()
    hits = []
    for theme, keywords in THEME_KEYWORDS.items():
        if any(k in text for k in keywords):
            hits.append(theme)
    return ";".join(hits) if hits else "other"


def contribution_summary(top):
    positive = top["contribution_8cell_to_morula"].clip(lower=0)
    total_positive = positive.sum()
    if total_positive <= 0:
        top["positive_contribution_fraction"] = 0.0
        top["cumulative_positive_contribution_fraction"] = 0.0
        return top
    top["positive_contribution_fraction"] = positive / total_positive
    top["cumulative_positive_contribution_fraction"] = top["positive_contribution_fraction"].cumsum()
    return top


def simplify_dmr_table(contrib):
    cols = [
        "cluster_name",
        "chr",
        "start",
        "end",
        "age_weight_5yr",
        "entropy_8-cell",
        "entropy_morula",
        "delta_H_8cell_to_morula",
        "contribution_8cell_to_morula",
        "positive_contribution_fraction",
        "cumulative_positive_contribution_fraction",
        "nearest_gene",
        "nearest_gene_distance_bp",
        "gene_context",
        "cpg_context",
        "reset_driver_rank_8cell_to_morula",
    ]
    keep = [c for c in cols if c in contrib.columns]
    out = contrib[keep].copy()
    out["reset_interpretation"] = out["delta_H_8cell_to_morula"].apply(
        lambda x: "entropy_decreases_8cell_to_morula" if pd.notna(x) and x > 0 else "not_reset_decreasing"
    )
    return out


def synthesize_pathways(gp):
    if gp.empty:
        return pd.DataFrame()
    gp = gp.copy()
    gp["biological_theme"] = gp["term_name"].apply(theme_for_term)
    gp = gp[gp["biological_theme"] != "other"].copy()
    if gp.empty:
        return gp
    gp["minus_log10_p"] = gp["p_value_gprofiler"].apply(
        lambda p: -math.log10(float(p)) if pd.notna(p) and float(p) > 0 else None
    )
    cols = [
        "biological_theme",
        "source",
        "term_id",
        "term_name",
        "p_value_gprofiler",
        "minus_log10_p",
        "intersection_size",
        "representative_genes",
    ]
    gp = gp[[c for c in cols if c in gp.columns]]
    return gp.sort_values(["p_value_gprofiler", "biological_theme"]).head(50)


def annotation_mechanism_summary(top, background, ann):
    rows = []
    for col in ["gene_context", "cpg_context"]:
        for category in sorted(set(background[col].dropna().astype(str))):
            top_count = int((top[col].astype(str) == category).sum())
            bg_count = int((background[col].astype(str) == category).sum())
            rows.append(
                {
                    "annotation_type": col,
                    "category": category,
                    "top_reset_driving_count": top_count,
                    "top_reset_driving_fraction": top_count / len(top),
                    "all_age_DMR_count": bg_count,
                    "all_age_DMR_fraction": bg_count / len(background),
                    "interpretation": (
                        "enriched_or_common_regulatory_context"
                        if top_count / len(top) >= bg_count / len(background)
                        else "depleted_relative_to_background"
                    ),
                }
            )
    out = pd.DataFrame(rows)
    if not ann.empty:
        ann2 = ann.rename(
            columns={
                "top_count": "previous_top_count",
                "background_count": "previous_background_count",
                "p_value": "hypergeom_p_value",
                "p_adj_BH": "hypergeom_BH_FDR",
            }
        )
        out = out.merge(
            ann2[["annotation_type", "category", "hypergeom_p_value", "hypergeom_BH_FDR"]],
            on=["annotation_type", "category"],
            how="left",
        )
    return out.sort_values(["annotation_type", "top_reset_driving_fraction"], ascending=[True, False])


def svg_text(parts, x, y, text, size=16, weight="400", fill="#111"):
    parts.append(
        f'<text x="{x}" y="{y}" font-family="Arial" font-size="{size}" font-weight="{weight}" fill="{fill}">{html.escape(str(text))}</text>'
    )


def draw_mechanism_svg(top, pathway, summary, out_path):
    width, height = 1480, 940
    parts = [
        f'<svg xmlns="http://www.w3.org/2000/svg" width="{width}" height="{height}" viewBox="0 0 {width} {height}">',
        '<rect width="100%" height="100%" fill="white"/>',
    ]
    svg_text(parts, 42, 48, "DMR-level mechanism map for TRO-defined morula ground-zero", 28, "700")
    svg_text(parts, 42, 78, "Top age-DMR entropy drops from 8-cell to morula are mapped to genes, regulatory context, and developmental pathways.", 16, "400", "#444")

    # Left panel: top DMRs
    parts.append('<rect x="40" y="115" width="430" height="720" rx="8" fill="#f7fbff" stroke="#9ecae1"/>')
    svg_text(parts, 65, 150, "Top reset-driving DMRs", 21, "700", "#08519c")
    max_val = max(float(top["contribution_8cell_to_morula"].clip(lower=0).max()), 1e-9)
    y = 182
    for _, row in top.head(12).iterrows():
        val = max(float(row["contribution_8cell_to_morula"]), 0.0)
        bar = int(210 * val / max_val)
        label = f'{row["cluster_name"]} / {row["nearest_gene"]}'
        svg_text(parts, 65, y, label[:36], 13)
        parts.append(f'<rect x="245" y="{y-13}" width="{bar}" height="13" fill="#3182bd"/>')
        svg_text(parts, 245 + bar + 8, y, f"{val:.3g}", 12, "400", "#333")
        y += 36
    top20_frac = float(top.head(20)["positive_contribution_fraction"].sum())
    svg_text(parts, 65, 650, f"Top 20 explain {top20_frac:.1%} of positive", 16, "700")
    svg_text(parts, 65, 674, "8-cell -> morula reset-driving contribution", 16, "700")
    svg_text(parts, 65, 725, "Formula:", 15, "700", "#333")
    svg_text(parts, 65, 752, "abs(age_weight) * (H_8-cell - H_morula)", 15, "400", "#333")

    # Middle panel: annotations
    parts.append('<rect x="525" y="115" width="390" height="720" rx="8" fill="#f7fcf5" stroke="#a1d99b"/>')
    svg_text(parts, 550, 150, "Genomic context", 21, "700", "#238b45")
    context = summary.copy()
    y = 190
    for _, row in context.head(10).iterrows():
        label = f'{row["annotation_type"].replace("_", " ")}: {row["category"]}'
        svg_text(parts, 550, y, label[:34], 14)
        svg_text(parts, 815, y, f'{row["top_reset_driving_fraction"]:.0%}', 14, "700", "#238b45")
        y += 43
    svg_text(parts, 550, 675, "Interpretation", 16, "700")
    svg_text(parts, 550, 702, "Reset-driving DMRs are annotated against", 14, "400", "#333")
    svg_text(parts, 550, 726, "promoter/exon/intron/intergenic and", 14, "400", "#333")
    svg_text(parts, 550, 750, "CpG island/shore/shelf contexts.", 14, "400", "#333")

    # Right panel: pathway themes
    parts.append('<rect x="970" y="115" width="465" height="720" rx="8" fill="#fff7ec" stroke="#fdae6b"/>')
    svg_text(parts, 995, 150, "Pathway / biological themes", 21, "700", "#a63603")
    y = 190
    if pathway.empty:
        svg_text(parts, 995, y, "No theme-matched g:Profiler terms", 15)
    else:
        for _, row in pathway.head(11).iterrows():
            name = row["term_name"]
            p = row["p_value_gprofiler"]
            genes = row.get("representative_genes", "")
            svg_text(parts, 995, y, str(name)[:45], 14, "700")
            svg_text(parts, 995, y + 21, f'p={float(p):.2g}; genes: {str(genes)[:48]}', 12, "400", "#555")
            y += 55
    svg_text(parts, 995, 765, "Claim boundary", 15, "700")
    svg_text(parts, 995, 790, "Exploratory nearest-gene interpretation,", 13, "400", "#333")
    svg_text(parts, 995, 812, "not causal proof of mechanism.", 13, "400", "#333")

    # Flow arrows
    parts.append('<path d="M470 475 C500 475, 500 475, 525 475" stroke="#666" stroke-width="3" fill="none" marker-end="url(#arrow)"/>')
    parts.append('<path d="M915 475 C945 475, 945 475, 970 475" stroke="#666" stroke-width="3" fill="none" marker-end="url(#arrow)"/>')
    parts.append(
        '<defs><marker id="arrow" markerWidth="10" markerHeight="10" refX="8" refY="3" orient="auto"><path d="M0,0 L0,6 L9,3 z" fill="#666"/></marker></defs>'
    )
    svg_text(parts, 40, 890, "Conclusion: morula is explained by low age-DMR entropy plus preserved potency, with reset-driving DMRs linked to developmental/signaling gene neighborhoods.", 17, "700")
    parts.append("</svg>")
    out_path.write_text("\n".join(parts), encoding="utf-8")


def main():
    TABLES.mkdir(exist_ok=True)
    FIGS.mkdir(exist_ok=True)
    NOTES.mkdir(exist_ok=True)

    contrib = pd.read_csv(TABLES / "TRO_interpretability_DMR_contribution_ranking.tsv", sep="\t")
    gp = pd.read_csv(TABLES / "TRO_interpretability_gprofiler_enrichment_top50_DMR_genes.tsv", sep="\t")
    ann = pd.read_csv(TABLES / "TRO_interpretability_DMR_annotation_enrichment.tsv", sep="\t")
    op = read_json(TABLES / "TRO_operator_summary.json")

    contrib = contrib.sort_values("reset_driver_rank_8cell_to_morula").copy()
    contrib = contribution_summary(contrib)
    top50 = simplify_dmr_table(contrib.head(50))
    top50.to_csv(TABLES / "TRO_DMR_mechanistic_top50_reset_drivers.tsv", sep="\t", index=False)

    pathway = synthesize_pathways(gp)
    pathway.to_csv(TABLES / "TRO_DMR_mechanistic_pathway_synthesis.tsv", sep="\t", index=False)

    ann_summary = annotation_mechanism_summary(contrib.head(50), contrib, ann)
    ann_summary.to_csv(TABLES / "TRO_DMR_mechanistic_annotation_summary.tsv", sep="\t", index=False)

    draw_mechanism_svg(
        top50,
        pathway,
        ann_summary,
        FIGS / "TRO_DMR_mechanistic_interpretability_map.svg",
    )

    top_genes = [str(x) for x in top50["nearest_gene"].head(20).tolist()]
    developmental_terms = pathway[pathway["biological_theme"].str.contains("development|wnt|stemness", na=False)].head(10)
    summary = {
        "analysis": "DMR-level mechanistic interpretability of TRO-defined morula ground-zero",
        "input_tables": [
            "TRO_interpretability_DMR_contribution_ranking.tsv",
            "TRO_interpretability_gprofiler_enrichment_top50_DMR_genes.tsv",
            "TRO_interpretability_DMR_annotation_enrichment.tsv",
        ],
        "top20_positive_contribution_fraction": float(top50.head(20)["positive_contribution_fraction"].sum()),
        "top50_positive_contribution_fraction": float(top50["positive_contribution_fraction"].sum()),
        "top_reset_driving_genes": top_genes,
        "top_pathway_terms": developmental_terms.to_dict(orient="records"),
        "morula_TRO_score": op.get("morula_TRO_score"),
        "main_interpretation": (
            "The morula TRO maximum is traceable to a ranked subset of age-associated DMRs whose entropy "
            "contribution drops from 8-cell to morula. These DMRs map near genes and ontology terms related "
            "to developmental patterning, cadherin/WNT signaling, and cell-fate regulation, while potency "
            "marker robustness supports preservation of developmental competence."
        ),
        "claim_boundary": (
            "Nearest-gene and pathway enrichment are biological interpretability supports. They do not prove "
            "direct causal regulation by each DMR."
        ),
    }
    (TABLES / "TRO_DMR_mechanistic_interpretability_summary.json").write_text(
        json.dumps(summary, indent=2, ensure_ascii=False), encoding="utf-8"
    )

    note = f"""# Experiment18: DMR mechanistic interpretability

This experiment strengthens the biological interpretation of the TRO-defined morula ground-zero state.

## Question

Which age-associated DMRs drive the 8-cell to morula decrease in age-weighted methylation entropy, and what biological gene neighborhoods do they point to?

## Main ranking

Reset-driving contribution:

```text
abs(age_weight) * (H_8-cell - H_morula)
```

Positive values mean the DMR's entropy contribution is lower at morula than at 8-cell.

## Key output

```text
tables/TRO_DMR_mechanistic_top50_reset_drivers.tsv
tables/TRO_DMR_mechanistic_pathway_synthesis.tsv
tables/TRO_DMR_mechanistic_annotation_summary.tsv
tables/TRO_DMR_mechanistic_interpretability_summary.json
figures/TRO_DMR_mechanistic_interpretability_map.svg
```

## Core interpretation

{summary["main_interpretation"]}

## Claim boundary

{summary["claim_boundary"]}

## Manuscript-ready wording

```text
The morula-stage TRO maximum was decomposed to DMR-level entropy contributions. A ranked subset of paternal age-associated DMRs showed marked entropy-contribution loss from 8-cell to morula, and the nearest-gene/pathway synthesis linked these regions to developmental patterning, cadherin/WNT signaling, and cell-fate regulatory neighborhoods. This supports a mechanistic interpretation of morula as a low age-associated methylation entropy and developmentally competent ground-zero candidate, while remaining an exploratory DMR-to-gene annotation analysis.
```
"""
    (NOTES / "Experiment18_DMR_mechanistic_interpretability.md").write_text(note, encoding="utf-8")

    print("Experiment18 DMR mechanistic interpretability completed.")
    print(json.dumps(summary, indent=2, ensure_ascii=False))


if __name__ == "__main__":
    main()
````
