## Supplemental code for "An entropy-based diagnostic framework for characterizing methylation state dynamics during preimplantation development": 31_experiment19_gse49828_gamete_to_embryo_directional_validation.py

````
from pathlib import Path
import argparse
import gzip
import json
import shutil
import urllib.request

import numpy as np
import pandas as pd


ROOT = Path(__file__).resolve().parents[1]
TABLES = ROOT / "tables"
FIGURES = ROOT / "figures"
NOTES = ROOT / "notes"
RAW = ROOT / "data_raw" / "GSE49828_independent_dna"
BASE = "https://ftp.ncbi.nlm.nih.gov/geo/series/GSE49nnn/GSE49828/suppl"

STAGE_ORDER = ["sperm", "MII oocyte", "zygote/PN", "2-cell", "4-cell", "8-cell", "morula"]

FILES = [
    ("sperm", "GSE49828_RRBS_Sperm1_methylation_calling.bed.txt.gz"),
    ("sperm", "GSE49828_RRBS_Sperm2_methylation_calling.bed.txt.gz"),
    ("sperm", "GSE49828_RRBS_Sperm3_methylation_calling.bed.txt.gz"),
    ("sperm", "GSE49828_RRBS_Sperm4_methylation_calling.bed.txt.gz"),
    ("MII oocyte", "GSE49828_RRBS_MII_Oocyte1_methylation_calling.bed.txt.gz"),
    ("MII oocyte", "GSE49828_RRBS_MII_Oocyte2_methylation_calling.bed.txt.gz"),
    ("zygote/PN", "GSE49828_RRBS_Zygote1_methylation_calling.bed.txt.gz"),
    ("zygote/PN", "GSE49828_RRBS_Zygote2_methylation_calling.bed.txt.gz"),
    ("2-cell", "GSE49828_RRBS_2-cell1_methylation_calling.bed.txt.gz"),
    ("2-cell", "GSE49828_RRBS_2-cell2_methylation_calling.bed.txt.gz"),
    ("4-cell", "GSE49828_RRBS_4-cell1_methylation_calling.bed.txt.gz"),
    ("4-cell", "GSE49828_RRBS_4-cell2_methylation_calling.bed.txt.gz"),
    ("8-cell", "GSE49828_RRBS_8-cell1_methylation_calling.bed.txt.gz"),
    ("8-cell", "GSE49828_RRBS_8-cell2_methylation_calling.bed.txt.gz"),
    ("8-cell", "GSE49828_RRBS_8-cell3_methylation_calling.bed.txt.gz"),
    ("morula", "GSE49828_RRBS_Morula1_methylation_calling.bed.txt.gz"),
    ("morula", "GSE49828_RRBS_Morula2_methylation_calling.bed.txt.gz"),
    ("morula", "GSE49828_RRBS_Morula3_methylation_calling.bed.txt.gz"),
]


def entropy(p, eps=1e-6):
    p = np.clip(np.asarray(p, dtype=float), eps, 1 - eps)
    return -(p * np.log(p) + (1 - p) * np.log(1 - p))


def validate_gzip(path):
    if not path.exists() or path.stat().st_size == 0:
        return False
    try:
        with gzip.open(path, "rb") as fh:
            while fh.read(1024 * 1024):
                pass
        return True
    except Exception:
        return False


def download_file(name, force=False):
    RAW.mkdir(parents=True, exist_ok=True)
    out = RAW / name
    url = f"{BASE}/{name}"
    if not force and validate_gzip(out):
        return {"filename": name, "url": url, "local_path": str(out), "status": "exists", "gzip_ok": True, "size_bytes": out.stat().st_size}

    tmp = out.with_suffix(out.suffix + ".part")
    if tmp.exists():
        tmp.unlink()
    print(f"Downloading {url}")
    urllib.request.urlretrieve(url, tmp)
    shutil.move(str(tmp), str(out))
    ok = validate_gzip(out)
    return {"filename": name, "url": url, "local_path": str(out), "status": "downloaded", "gzip_ok": ok, "size_bytes": out.stat().st_size}


def load_regions():
    reg = pd.read_csv(ROOT / "metadata" / "GSE102970_TableS6_age_dmr_weights.tsv", sep="\t")
    reg = reg[["cluster_name", "chr", "start", "end", "age_weight_5yr"]].copy()
    reg["start"] = reg["start"].astype(int)
    reg["end"] = reg["end"].astype(int)
    return reg


def build_region_index(regions):
    index = {}
    for chrom, sub in regions.groupby("chr"):
        sub = sub.sort_values("start").reset_index(drop=True)
        index[chrom] = {
            "starts": sub["start"].to_numpy(),
            "ends": sub["end"].to_numpy(),
            "clusters": sub["cluster_name"].to_numpy(),
        }
    return index


def find_region(index, chrom, pos):
    if chrom not in index:
        return None
    starts = index[chrom]["starts"]
    i = np.searchsorted(starts, pos, side="right") - 1
    if i >= 0 and pos <= index[chrom]["ends"][i]:
        return str(index[chrom]["clusters"][i])
    return None


def process_file(stage, path, regions, index):
    reg_lookup = regions.set_index("cluster_name")
    met = {name: 0.0 for name in regions["cluster_name"]}
    total = {name: 0.0 for name in regions["cluster_name"]}
    n_cpg = {name: 0 for name in regions["cluster_name"]}
    cg_lines = 0
    matched_cpg = 0

    with gzip.open(path, "rt", errors="replace") as fh:
        _ = next(fh, "")
        for line in fh:
            parts = line.rstrip("\n").split("\t")
            if len(parts) < 10 or parts[9] != "CpG":
                continue
            try:
                chrom = parts[0]
                pos = int(parts[1])
                t = float(parts[4])
                m = float(parts[5])
            except ValueError:
                continue
            cg_lines += 1
            if t <= 0:
                continue
            cluster = find_region(index, chrom, pos)
            if cluster is None:
                continue
            matched_cpg += 1
            met[cluster] += m
            total[cluster] += t
            n_cpg[cluster] += 1

    rows = []
    for cluster, t in total.items():
        if t <= 0:
            continue
        rows.append(
            {
                "stage": stage,
                "sample": path.name,
                "cluster_name": cluster,
                "age_weight_5yr": float(reg_lookup.loc[cluster, "age_weight_5yr"]),
                "met_reads": met[cluster],
                "total_reads": t,
                "beta": met[cluster] / t,
                "n_cpg_observed": n_cpg[cluster],
            }
        )
    qc = {
        "stage": stage,
        "sample": path.name,
        "cg_lines": cg_lines,
        "matched_cpg": matched_cpg,
        "covered_regions": len(rows),
    }
    return pd.DataFrame(rows), qc


def stage_metrics(long_df):
    rows = []
    for stage in STAGE_ORDER:
        sub = long_df[long_df["stage"] == stage]
        if sub.empty:
            continue
        g = sub.groupby("cluster_name", as_index=False).agg(
            met_reads=("met_reads", "sum"),
            total_reads=("total_reads", "sum"),
            age_weight_5yr=("age_weight_5yr", "first"),
        )
        g = g[g["total_reads"] > 0].copy()
        p = g["met_reads"] / g["total_reads"]
        h = entropy(p)
        w = g["age_weight_5yr"].astype(float).to_numpy()
        rows.append(
            {
                "stage": stage,
                "n_regions_valid": len(g),
                "s_epi": float(np.mean(h)),
                "s_epi_age": float(np.sum(np.abs(w) * h) / np.sum(np.abs(w))),
                "age_projection": float(np.sum(w * p) / np.sum(np.abs(w))),
            }
        )
    metrics = pd.DataFrame(rows)
    metrics["stage"] = pd.Categorical(metrics["stage"], categories=STAGE_ORDER, ordered=True)
    metrics = metrics.sort_values("stage").reset_index(drop=True)

    sperm_val = float(metrics.loc[metrics["stage"].astype(str) == "sperm", "s_epi_age"].iloc[0])
    embryo = metrics[metrics["stage"].astype(str) != "sperm"].copy()
    min_embryo_val = float(embryo["s_epi_age"].min())
    denom = sperm_val - min_embryo_val
    if abs(denom) < 1e-12:
        metrics["directional_reset_score_from_sperm"] = np.nan
    else:
        metrics["directional_reset_score_from_sperm"] = (sperm_val - metrics["s_epi_age"]) / denom
    return metrics


def transition_metrics(metrics):
    rows = []
    usable = metrics.dropna(subset=["s_epi_age"]).copy()
    usable["stage"] = usable["stage"].astype(str)
    for a, b in zip(STAGE_ORDER[:-1], STAGE_ORDER[1:]):
        if a not in set(usable["stage"]) or b not in set(usable["stage"]):
            continue
        va = float(usable.loc[usable["stage"] == a, "s_epi_age"].iloc[0])
        vb = float(usable.loc[usable["stage"] == b, "s_epi_age"].iloc[0])
        rows.append(
            {
                "transition": f"{a} -> {b}",
                "stage_from": a,
                "stage_to": b,
                "delta_s_epi_age_reduction": va - vb,
                "absolute_step_change": abs(va - vb),
            }
        )
    out = pd.DataFrame(rows)
    if not out.empty:
        out["reduction_rank"] = out["delta_s_epi_age_reduction"].rank(ascending=False, method="min").astype(int)
    return out


def write_svg(metrics, summary, out):
    m = metrics.copy()
    m["stage"] = m["stage"].astype(str)
    width, height = 980, 540
    left, right, top, bottom = 90, 40, 60, 95
    xs = np.linspace(left, width - right, len(m))
    yvals = m["s_epi_age"].to_numpy(dtype=float)
    ymin, ymax = float(np.nanmin(yvals)), float(np.nanmax(yvals))
    pad = max((ymax - ymin) * 0.15, 0.01)
    ymin -= pad
    ymax += pad

    def yscale(v):
        return top + (ymax - v) / (ymax - ymin) * (height - top - bottom)

    pts = [(float(x), float(yscale(v))) for x, v in zip(xs, yvals)]
    poly = " ".join(f"{x:.1f},{y:.1f}" for x, y in pts)
    circles = []
    for x, y, stage, val in zip(xs, [p[1] for p in pts], m["stage"], yvals):
        fill = "#9c2f2f" if stage == "sperm" else ("#1f7a4c" if stage == "morula" else "#2d5f9a")
        circles.append(f'<circle cx="{x:.1f}" cy="{y:.1f}" r="7" fill="{fill}" stroke="white" stroke-width="2"/>')
        circles.append(f'<text x="{x:.1f}" y="{height - 55}" text-anchor="middle" font-size="13">{stage}</text>')
        circles.append(f'<text x="{x:.1f}" y="{y - 14:.1f}" text-anchor="middle" font-size="12">{val:.3f}</text>')

    y_ticks = []
    for tv in np.linspace(ymin, ymax, 5):
        y = yscale(tv)
        y_ticks.append(f'<line x1="{left}" x2="{width-right}" y1="{y:.1f}" y2="{y:.1f}" stroke="#e6e6e6"/>')
        y_ticks.append(f'<text x="{left-12}" y="{y+4:.1f}" text-anchor="end" font-size="12">{tv:.3f}</text>')

    svg = f'''<svg xmlns="http://www.w3.org/2000/svg" width="{width}" height="{height}" viewBox="0 0 {width} {height}">
  <rect width="100%" height="100%" fill="white"/>
  <text x="{left}" y="30" font-size="22" font-weight="700">GSE49828 human gamete-to-embryo directional age-DMR entropy validation</text>
  {''.join(y_ticks)}
  <line x1="{left}" x2="{width-right}" y1="{height-bottom}" y2="{height-bottom}" stroke="#222"/>
  <line x1="{left}" x2="{left}" y1="{top}" y2="{height-bottom}" stroke="#222"/>
  <polyline points="{poly}" fill="none" stroke="#315f9a" stroke-width="3"/>
  {''.join(circles)}
  <text x="22" y="{height/2}" transform="rotate(-90 22,{height/2})" text-anchor="middle" font-size="15">S_epi-age</text>
  <text x="{left}" y="{height-20}" font-size="13" fill="#444">Boundary: independent human RRBS directional validation, not strict matched parental gamete-to-embryo proof.</text>
  <text x="{width-right}" y="62" text-anchor="end" font-size="13" fill="#444">lowest embryo stage: {summary["lowest_embryo_stage_by_s_epi_age"]}</text>
</svg>
'''
    out.write_text(svg, encoding="utf-8")


def main():
    ap = argparse.ArgumentParser()
    ap.add_argument("--download-sperm", action="store_true", help="Download missing GSE49828 sperm RRBS files.")
    ap.add_argument("--force-download", action="store_true")
    ap.add_argument("--max-sperm", type=int, default=4)
    args = ap.parse_args()

    TABLES.mkdir(exist_ok=True)
    FIGURES.mkdir(exist_ok=True)
    NOTES.mkdir(exist_ok=True)
    RAW.mkdir(parents=True, exist_ok=True)

    selected = []
    sperm_seen = 0
    manifest_rows = []
    for stage, name in FILES:
        if stage == "sperm":
            sperm_seen += 1
            if sperm_seen > args.max_sperm:
                continue
            if args.download_sperm:
                row = download_file(name, force=args.force_download)
            else:
                path = RAW / name
                row = {
                    "filename": name,
                    "url": f"{BASE}/{name}",
                    "local_path": str(path),
                    "status": "local" if path.exists() else "missing",
                    "gzip_ok": validate_gzip(path),
                    "size_bytes": path.stat().st_size if path.exists() else 0,
                }
        else:
            path = RAW / name
            row = {
                "filename": name,
                "url": f"{BASE}/{name}",
                "local_path": str(path),
                "status": "local" if path.exists() else "missing",
                "gzip_ok": validate_gzip(path),
                "size_bytes": path.stat().st_size if path.exists() else 0,
            }
        row["stage"] = stage
        manifest_rows.append(row)
        if row["gzip_ok"]:
            selected.append((stage, Path(row["local_path"])))

    manifest = pd.DataFrame(manifest_rows)
    manifest.to_csv(TABLES / "GSE49828_gamete_to_embryo_directional_manifest.tsv", sep="\t", index=False)
    if "sperm" not in {stage for stage, _ in selected}:
        raise SystemExit("No valid sperm RRBS file is available. Re-run with --download-sperm.")

    regions = load_regions()
    index = build_region_index(regions)
    per_file = []
    qc_rows = []
    for stage, path in selected:
        print(f"Processing {stage}: {path.name}")
        df, qc = process_file(stage, path, regions, index)
        per_file.append(df)
        qc_rows.append(qc)

    long_df = pd.concat(per_file, ignore_index=True)
    long_df.to_csv(TABLES / "GSE49828_gamete_to_embryo_age_DMR_region_values.tsv", sep="\t", index=False)
    pd.DataFrame(qc_rows).to_csv(TABLES / "GSE49828_gamete_to_embryo_directional_qc.tsv", sep="\t", index=False)

    metrics = stage_metrics(long_df)
    metrics.to_csv(TABLES / "GSE49828_gamete_to_embryo_directional_stage_metrics.tsv", sep="\t", index=False)
    transitions = transition_metrics(metrics)
    transitions.to_csv(TABLES / "GSE49828_gamete_to_embryo_directional_transition_metrics.tsv", sep="\t", index=False)

    ranked_embryo = metrics[metrics["stage"].astype(str) != "sperm"].sort_values("s_epi_age").reset_index(drop=True)
    sperm_s = float(metrics.loc[metrics["stage"].astype(str) == "sperm", "s_epi_age"].iloc[0])
    morula_s = float(metrics.loc[metrics["stage"].astype(str) == "morula", "s_epi_age"].iloc[0])
    morula_rank = int(ranked_embryo.index[ranked_embryo["stage"].astype(str) == "morula"][0] + 1)
    summary = {
        "dataset": "GSE49828",
        "validation_type": "human_gamete_to_embryo_directional_RRBS_age_DMR_entropy",
        "strict_pairing": False,
        "sperm_s_epi_age": sperm_s,
        "morula_s_epi_age": morula_s,
        "sperm_to_morula_delta_s_epi_age_reduction": sperm_s - morula_s,
        "lowest_embryo_stage_by_s_epi_age": str(ranked_embryo.loc[0, "stage"]),
        "morula_rank_among_embryo_stages": morula_rank,
        "top3_lowest_embryo_stages": ranked_embryo["stage"].astype(str).head(3).tolist(),
        "supports_morula_or_adjacent_low_age_entropy_window": bool(morula_rank <= 3),
        "claim_boundary": "GSE49828 includes human gamete and embryo RRBS methylomes but is not a strict matched parental gamete-to-embryo dataset; use as directional human validation only.",
    }
    (TABLES / "GSE49828_gamete_to_embryo_directional_summary.json").write_text(
        json.dumps(summary, indent=2, ensure_ascii=False), encoding="utf-8"
    )
    write_svg(metrics, summary, FIGURES / "GSE49828_gamete_to_embryo_directional_validation.svg")

    note = f"""# Experiment19: GSE49828 human gamete-to-embryo directional validation

This experiment adds human sperm RRBS methylomes to the previous GSE49828 independent DNA validation and compares
age-DMR weighted methylation entropy across sperm, MII oocyte, and early embryo stages.

This is not a strict paired parental gamete-to-embryo proof because GSE49828 does not link a specific sperm donor to a
specific embryo trajectory. It is a directional human validation of whether gamete-to-embryo development enters a low
age-associated methylation entropy window near morula or adjacent stages.

```json
{json.dumps(summary, indent=2, ensure_ascii=False)}
```
"""
    (NOTES / "Experiment19_GSE49828_gamete_to_embryo_directional_validation.md").write_text(note, encoding="utf-8")

    print("GSE49828 human gamete-to-embryo directional validation:")
    print(json.dumps(summary, indent=2, ensure_ascii=False))
    print("Wrote:", TABLES / "GSE49828_gamete_to_embryo_directional_stage_metrics.tsv")
    print("Wrote:", TABLES / "GSE49828_gamete_to_embryo_directional_summary.json")
    print("Wrote:", FIGURES / "GSE49828_gamete_to_embryo_directional_validation.svg")


if __name__ == "__main__":
    main()
````
