## Supplemental code for "An entropy-based diagnostic framework for characterizing methylation state dynamics during preimplantation development": check_env.sh

```
#!/usr/bin/env bash
set -u

{
echo "===== DISK LOCATION CHECK ====="
pwd
df -h .
df -h /root/autodl-tmp || true
df -h /autodl-pub || true
df -h /autodl-pub/data || true
du -sh /root/autodl-tmp/TRO_Project
ls -ld /root/autodl-tmp /autodl-pub /autodl-pub/data 2>/dev/null || true
readlink -f /root/autodl-tmp

echo
echo "===== NETWORK CHECK ====="
curl -I https://ftp.ncbi.nlm.nih.gov/geo/series/GSE81nnn/GSE81233/ 2>/dev/null | head || true
curl -I https://ftp.ncbi.nlm.nih.gov/geo/series/GSE102nnn/GSE102970/ 2>/dev/null | head || true

echo
echo "===== PYTHON PACKAGE CHECK ====="
python - <<'PY'
mods = ["numpy", "pandas", "scipy", "statsmodels", "matplotlib", "seaborn", "sklearn", "pyarrow", "requests"]
for m in mods:
    try:
        mod = __import__(m)
        print(f"{m}: OK {getattr(mod, '__version__', '')}")
    except Exception as e:
        print(f"{m}: MISSING ({e})")
PY

echo
echo "===== CONDA INFO ====="
conda info --envs || true
} 2>&1 | tee logs/check_env.txt
```
