## Supplemental code for "An entropy-based diagnostic framework for characterizing methylation state dynamics during preimplantation development": check_results_package.py

```
from pathlib import Path
import json

import pandas as pd


def main():
    root = Path(__file__).resolve().parents[1]
    tables = root / "tables"
    figures = root / "figures"

    required_tables = [
        "GSE81233_valid204_stage_epi_age_metrics.tsv",
        "Experiment1B_DNA_robustness_summary.tsv",
        "GSE36552_RNA_entropy_potency_by_stage.tsv",
        "GSE36552_potency_pairwise_tests.tsv",
        "TRO_composite_score_by_stage.tsv",
        "marker_leave_one_out_summary.tsv",
        "GSE44183_external_potency_validation.tsv",
        "TRO_stage_transition_cost.tsv",
        "TRO_operator_stage_output.tsv",
        "TRO_operator_transition_output.tsv",
        "TRO_operator_summary.json",
        "final_claim_audit.tsv",
        "manuscript_numbers.json",
        "Experiment9_age_DMR_specificity_boundary_summary.tsv",
        "Experiment9_age_DMR_specificity_boundary_summary.json",
        "GSE49828_independent_DNA_validation_stage_metrics.tsv",
        "GSE49828_independent_DNA_validation_summary.json",
        "TRO_interpretability_DMR_contribution_ranking.tsv",
        "TRO_interpretability_top20_reset_driving_DMRs.tsv",
        "TRO_interpretability_top50_reset_driving_DMRs.tsv",
        "TRO_interpretability_DMR_annotation_enrichment.tsv",
        "TRO_interpretability_reset_DMR_gene_function_enrichment.tsv",
        "TRO_interpretability_potency_marker_contribution.tsv",
        "TRO_interpretability_summary.json",
        "TRO_latent_reset_sample_scores.tsv",
        "TRO_latent_reset_stage_summary.tsv",
        "TRO_latent_reset_axis_correlations.tsv",
        "TRO_latent_reset_transition_distances.tsv",
        "TRO_latent_reset_representation_sensitivity.tsv",
        "TRO_latent_reset_top_axis_DMR_loadings.tsv",
        "TRO_latent_reset_summary.json",
        "GSE273723_parental_age_placenta_metadata.tsv",
        "GSE273723_parental_age_residual_sample_metrics.tsv",
        "GSE273723_parental_age_residual_group_metrics.tsv",
        "GSE273723_parental_age_residual_group_tests.tsv",
        "GSE273723_parental_age_residual_CpG_escape_ranking.tsv",
        "GSE273723_top50_sperm_age_DMR_placenta_escape_CpGs.tsv",
        "GSE273723_parental_age_residual_TRO_summary.json",
        "GSE56697_paired_paternal_operator_stage_metrics.tsv",
        "GSE56697_paired_paternal_operator_transition_metrics.tsv",
        "GSE56697_paired_paternal_operator_summary.json",
        "GSE56697_paired_paternal_operator_robustness_by_bins.tsv",
        "GSE56697_paired_paternal_operator_robustness_stage_metrics.tsv",
        "GSE56697_paired_paternal_operator_robustness_transition_metrics.tsv",
        "GSE56697_paired_paternal_operator_robustness_summary.json",
        "GSE56697_maternal_paternal_branch_stage_metrics.tsv",
        "GSE56697_maternal_paternal_branch_transition_metrics.tsv",
        "GSE56697_maternal_paternal_branch_summary.tsv",
        "GSE56697_maternal_paternal_branch_summary.json",
        "TRO_interpretability_gprofiler_enrichment_top50_DMR_genes.tsv",
        "TRO_publication_evidence_ladder.tsv",
        "TRO_publication_synthesis_summary.json",
        "TRO_DMR_mechanistic_top50_reset_drivers.tsv",
        "TRO_DMR_mechanistic_pathway_synthesis.tsv",
        "TRO_DMR_mechanistic_annotation_summary.tsv",
        "TRO_DMR_mechanistic_interpretability_summary.json",
        "GSE49828_gamete_to_embryo_directional_stage_metrics.tsv",
        "GSE49828_gamete_to_embryo_directional_transition_metrics.tsv",
        "GSE49828_gamete_to_embryo_directional_summary.json",
    ]

    required_figures = [
        "GSE81233_valid204_s_epi_age_by_stage.png",
        "Experiment1B_DNA_robustness_summary.png",
        "GSE36552_marker_heatmap.png",
        "dual_entropy_phase_map_final.png",
        "TRO_composite_score_by_stage.png",
        "TRO_stage_transition_cost.png",
        "TRO_operator_diagram.png",
        "Experiment9_age_DMR_specificity_boundary_control.png",
        "GSE49828_independent_DNA_validation.png",
        "TRO_interpretability_top_reset_driving_DMRs.png",
        "TRO_interpretability_potency_marker_contribution.png",
        "TRO_latent_reset_pca_space.png",
        "TRO_latent_reset_axis_by_stage.png",
        "Experiment13_parental_age_residual_TRO_validation.svg",
        "GSE56697_paired_paternal_reset_operator.svg",
        "GSE56697_paired_paternal_operator_robustness_by_bins.svg",
        "GSE56697_maternal_paternal_branch_contrast.svg",
        "TRO_publication_evidence_ladder.svg",
        "TRO_DMR_mechanistic_interpretability_map.svg",
        "GSE49828_gamete_to_embryo_directional_validation.svg",
    ]

    required_notes = [
        "data_manifest.md",
        "reproducibility_notes.md",
        "manuscript_claim_boundaries.md",
        "TRO_operator_schema.md",
        "Experiment9_age_DMR_specificity_boundary_control.md",
        "GSE49828_independent_DNA_validation.md",
        "TRO_interpretability_analysis.md",
        "TRO_latent_reset_analysis.md",
        "Experiment13_parental_age_residual_TRO_validation.md",
        "Experiment14_GSE56697_paired_paternal_reset_operator.md",
        "Experiment15_GSE56697_paired_operator_robustness.md",
        "Experiment16_GSE56697_maternal_paternal_branch_contrast.md",
        "TRO_publication_synthesis_and_claim_hierarchy.md",
        "Experiment18_DMR_mechanistic_interpretability.md",
        "Experiment19_GSE49828_gamete_to_embryo_directional_validation.md",
    ]

    required_root_files = [
        "README.md",
        "METHODS_REPRODUCIBILITY.md",
        "environment.yml",
        "run_all_results_only.cmd",
    ]

    missing = []
    for name in required_tables:
        if not (tables / name).exists():
            missing.append(str(tables / name))
    for name in required_figures:
        if not (figures / name).exists():
            missing.append(str(figures / name))
    for name in required_notes:
        if not (root / "notes" / name).exists():
            missing.append(str(root / "notes" / name))
    for name in required_root_files:
        if not (root / name).exists():
            missing.append(str(root / name))

    if missing:
        raise SystemExit("Missing required files:\n" + "\n".join(missing))

    summary = json.loads((tables / "TRO_operator_summary.json").read_text(encoding="utf-8"))
    if summary["ground_zero_stage"] != "morula":
        raise SystemExit("Unexpected ground_zero_stage: " + str(summary["ground_zero_stage"]))
    if summary["all_core_checks_pass"] is not True:
        raise SystemExit("all_core_checks_pass is not true")

    stage = pd.read_csv(tables / "TRO_operator_stage_output.tsv", sep="\t")
    morula = stage.loc[stage["stage"] == "morula"].iloc[0]
    for col in ["GZ_rank", "TRO_rank", "BioAgeRank"]:
        if int(morula[col]) != 1:
            raise SystemExit(f"morula {col} is not 1: {morula[col]}")

    audit = pd.read_csv(tables / "final_claim_audit.tsv", sep="\t")
    required_claims = {"DNA-1", "DNA-2", "RNA-2", "TRO-1", "TRANS-1", "OP-1"}
    observed_claims = set(audit["claim_id"].astype(str))
    missing_claims = required_claims - observed_claims
    if missing_claims:
        raise SystemExit("Missing audit claims: " + ", ".join(sorted(missing_claims)))

    specificity = json.loads((tables / "Experiment9_age_DMR_specificity_boundary_summary.json").read_text(encoding="utf-8"))
    if specificity["age_DMR_ground_zero_stage"] != "morula":
        raise SystemExit("Experiment9 age_DMR_ground_zero_stage is not morula")
    if specificity["conclusion"] != "age_weighting_strengthens_a_broader_morula_methylation_reprogramming_minimum":
        raise SystemExit("Experiment9 conclusion changed: " + specificity["conclusion"])

    independent_dna = json.loads((tables / "GSE49828_independent_DNA_validation_summary.json").read_text(encoding="utf-8"))
    if independent_dna["supports_morula_or_adjacent_low_age_entropy"] is not True:
        raise SystemExit("GSE49828 independent DNA validation no longer supports morula/adjacent low entropy")

    interpretability = json.loads((tables / "TRO_interpretability_summary.json").read_text(encoding="utf-8"))
    if interpretability["analysis"] != "TRO interpretability analysis":
        raise SystemExit("Unexpected interpretability analysis summary")
    marker_table = pd.read_csv(tables / "TRO_interpretability_potency_marker_contribution.tsv", sep="\t")
    if marker_table.empty:
        raise SystemExit("TRO interpretability marker contribution table is empty")

    latent = json.loads((tables / "TRO_latent_reset_summary.json").read_text(encoding="utf-8"))
    if latent["analysis"] != "unsupervised latent-space validation of TRO ground-zero state":
        raise SystemExit("Unexpected latent reset summary")
    if latent["supports_unsupervised_low_age_entropy_window"] is not True:
        raise SystemExit("Latent reset analysis no longer supports a low age-entropy window")

    residual = json.loads((tables / "GSE273723_parental_age_residual_TRO_summary.json").read_text(encoding="utf-8"))
    if residual["dataset"] != "GSE273723":
        raise SystemExit("Unexpected parental-age residual dataset")
    if residual["biological_scope"] != "offspring placenta methylation, not preimplantation embryo":
        raise SystemExit("Parental-age residual claim boundary changed")
    if residual["observed_target_CpGs_in_EPIC_matrix"] <= 0:
        raise SystemExit("No sperm age-DMR CpGs found in GSE273723 residual validation")

    paired = json.loads((tables / "GSE56697_paired_paternal_operator_summary.json").read_text(encoding="utf-8"))
    if paired["dataset"] != "GSE56697":
        raise SystemExit("Unexpected paired paternal operator dataset")
    if paired["ground_zero_stage_by_min_paternal_methylation"] != "ICM paternal":
        raise SystemExit("Unexpected GSE56697 paternal ground-zero stage: " + paired["ground_zero_stage_by_min_paternal_methylation"])
    if paired["best_demethylation_transition"] != "sperm -> 2-cell paternal":
        raise SystemExit("Unexpected GSE56697 paternal best transition: " + paired["best_demethylation_transition"])

    paired_robust = json.loads((tables / "GSE56697_paired_paternal_operator_robustness_summary.json").read_text(encoding="utf-8"))
    if paired_robust["ground_zero_stable_across_bin_sizes"] is not True:
        raise SystemExit("GSE56697 paired paternal ground-zero is not stable across bin sizes")
    if paired_robust["best_transition_stable_across_bin_sizes"] is not True:
        raise SystemExit("GSE56697 paired paternal best transition is not stable across bin sizes")

    branch = json.loads((tables / "GSE56697_maternal_paternal_branch_summary.json").read_text(encoding="utf-8"))
    if branch["dataset"] != "GSE56697":
        raise SystemExit("Unexpected GSE56697 branch contrast dataset")
    branch_names = {row["branch"] for row in branch["branch_summaries"]}
    if branch_names != {"paternal", "maternal"}:
        raise SystemExit("GSE56697 branch contrast does not contain both parental branches")

    synthesis = json.loads((tables / "TRO_publication_synthesis_summary.json").read_text(encoding="utf-8"))
    if synthesis["main_human_claim"] != "Human age-DMR entropy identifies morula as a computational ground-zero candidate.":
        raise SystemExit("Unexpected publication synthesis human claim")
    if "human paired paternal-age gamete-to-embryo reset proof" not in synthesis["not_claimed"]:
        raise SystemExit("Publication synthesis missing human paired-reset claim boundary")
    gp = pd.read_csv(tables / "TRO_interpretability_gprofiler_enrichment_top50_DMR_genes.tsv", sep="\t")
    if gp.empty:
        raise SystemExit("g:Profiler enrichment table is empty")

    mechanism = json.loads((tables / "TRO_DMR_mechanistic_interpretability_summary.json").read_text(encoding="utf-8"))
    if mechanism["analysis"] != "DMR-level mechanistic interpretability of TRO-defined morula ground-zero":
        raise SystemExit("Unexpected DMR mechanistic interpretability summary")
    if mechanism["top20_positive_contribution_fraction"] <= 0:
        raise SystemExit("DMR mechanistic top20 contribution fraction is not positive")
    if not mechanism["top_pathway_terms"]:
        raise SystemExit("DMR mechanistic pathway synthesis is empty")

    gse49828_gamete = json.loads((tables / "GSE49828_gamete_to_embryo_directional_summary.json").read_text(encoding="utf-8"))
    if gse49828_gamete["validation_type"] != "human_gamete_to_embryo_directional_RRBS_age_DMR_entropy":
        raise SystemExit("Unexpected GSE49828 gamete-to-embryo validation type")
    if gse49828_gamete["strict_pairing"] is not False:
        raise SystemExit("GSE49828 gamete validation claim boundary changed")
    if gse49828_gamete["supports_morula_or_adjacent_low_age_entropy_window"] is not True:
        raise SystemExit("GSE49828 gamete validation no longer supports low age-entropy window")

    print("Result-package integrity check passed.")
    print("ground_zero_stage =", summary["ground_zero_stage"])
    print("morula_TRO_score =", summary["morula_TRO_score"])
    print("all_core_checks_pass =", summary["all_core_checks_pass"])


if __name__ == "__main__":
    main()
```
