## Supplemental code for "An entropy-based diagnostic framework for characterizing methylation state dynamics during preimplantation development": retry_GSM2986343_reliable.sh

```
#!/usr/bin/env bash
set -euo pipefail

cd /root/autodl-tmp/TRO_Project

CACHE="data_raw/GSE81233_embryo_methylation/cache_cmet"
FILE="GSM2986343_scBS-2C-10-1.Cmet.bed.gz"
URL="https://ftp.ncbi.nlm.nih.gov/geo/samples/GSM2986nnn/GSM2986343/suppl/GSM2986343_scBS-2C-10-1.Cmet.bed.gz"

mkdir -p "$CACHE"

echo "===== CLEAN BAD FILE ====="
rm -f "$CACHE/$FILE" "$CACHE/$FILE.aria2" "$CACHE/$FILE.tmp"

echo "===== CURL SINGLE STREAM DOWNLOAD ====="
curl -L --http1.1 \
  --retry 20 \
  --retry-delay 10 \
  --connect-timeout 30 \
  --speed-time 120 \
  --speed-limit 10240 \
  -o "$CACHE/$FILE.tmp" \
  "$URL"

mv "$CACHE/$FILE.tmp" "$CACHE/$FILE"

echo "===== FILE SIZE ====="
ls -lh "$CACHE/$FILE"

echo "===== GZIP TEST ====="
gzip -t "$CACHE/$FILE"

echo "===== PARSE ONLY GSM2986343 ====="
/root/miniconda3/envs/tro/bin/python scripts/09d_aria2_cache_gse81233_shard_worker.py \
  --manifest results/tables/GSE81233_retry_GSM2986343_only.tsv \
  --outdir data_processed/methylation_matrix/GSE81233_age_dmr_stream_full \
  --n-shards 1 \
  --shard-index 0 \
  --connections 1

echo "===== FINAL COUNT ====="
find data_processed/methylation_matrix/GSE81233_age_dmr_stream_full/sample_dmr -name "*.age_dmr.tsv" | wc -l
```
