## Supplemental code for "An entropy-based diagnostic framework for characterizing methylation state dynamics during preimplantation development": csb_tro_adjacent_specificity.py

```
from __future__ import annotations

import json
from pathlib import Path

import numpy as np
import pandas as pd


OUT = Path("E:/实验更新_5_22/csb_tro_dynamics")
STAGE_ORDER = ["MII oocyte", "zygote/PN", "2-cell", "4-cell", "8-cell", "morula", "blastocyst"]
STATE_COLS = ["A", "Hm", "P", "Hr"]
EPSILON = 0.075
LAMBDA_A = 1.65
LAMBDA_P = 0.95
POTENCY_THRESHOLD_QUANTILE = 0.60


def stage_agnostic_p_min(particles: pd.DataFrame) -> float:
    return float(particles["P"].quantile(POTENCY_THRESHOLD_QUANTILE))


def sinkhorn(a: np.ndarray, b: np.ndarray, cost: np.ndarray, n_iter: int = 1000) -> np.ndarray:
    k = np.maximum(np.exp(-cost / EPSILON), 1e-300)
    u = np.ones_like(a)
    v = np.ones_like(b)
    for _ in range(n_iter):
        u_new = a / np.maximum(k @ v, 1e-300)
        v_new = b / np.maximum(k.T @ u_new, 1e-300)
        if np.max(np.abs(u_new - u)) < 1e-10 and np.max(np.abs(v_new - v)) < 1e-10:
            u, v = u_new, v_new
            break
        u, v = u_new, v_new
    pi = (u[:, None] * k) * v[None, :]
    return pi / max(pi.sum(), 1e-300)


def transition(x: np.ndarray, y: np.ndarray, p_min: float) -> dict[str, float]:
    diff = y[None, :, :] - x[:, None, :]
    movement = np.sum(diff * diff, axis=2)
    age = np.maximum(y[None, :, 0] - x[:, None, 0], 0.0) ** 2
    potency = np.broadcast_to(np.maximum(p_min - y[None, :, 2], 0.0) ** 2, movement.shape)
    cost = movement + LAMBDA_A * age + LAMBDA_P * potency
    pi = sinkhorn(np.full(x.shape[0], 1 / x.shape[0]), np.full(y.shape[0], 1 / y.shape[0]), cost)
    return {
        "mean_transport_A": float(np.sum(pi * diff[:, :, 0])),
        "mean_transport_P": float(np.sum(pi * diff[:, :, 2])),
        "mean_squared_displacement": float(np.sum(pi * movement)),
    }


def summarize_stages(particles: pd.DataFrame) -> pd.DataFrame:
    sm = particles.groupby("stage", as_index=False).agg(A_mean=("A", "mean"), P_mean=("P", "mean"))
    sm["stage_order"] = sm["stage"].map({s: i for i, s in enumerate(STAGE_ORDER)})
    sm["reset_score_A_minus_P"] = sm["A_mean"] - sm["P_mean"]
    sm["reset_rank_lowest_is_1"] = sm["reset_score_A_minus_P"].rank(method="min", ascending=True).astype(int)
    return sm.sort_values("stage_order")


def adjacent_metrics(particles: pd.DataFrame) -> tuple[pd.DataFrame, dict]:
    sm = summarize_stages(particles)
    p_min = stage_agnostic_p_min(particles)
    rows = []
    for s0, s1 in zip(STAGE_ORDER[:-1], STAGE_ORDER[1:]):
        x = particles[particles["stage"].eq(s0)][STATE_COLS].to_numpy(dtype=float)
        y = particles[particles["stage"].eq(s1)][STATE_COLS].to_numpy(dtype=float)
        tr = transition(x, y, p_min)
        dest = sm[sm["stage"].eq(s1)].iloc[0]
        tr.update(
            {
                "from_stage": s0,
                "to_stage": s1,
                "destination_A_mean": float(dest["A_mean"]),
                "destination_P_mean": float(dest["P_mean"]),
                "destination_reset_score_A_minus_P": float(dest["reset_score_A_minus_P"]),
                "destination_reset_rank_lowest_is_1": int(dest["reset_rank_lowest_is_1"]),
            }
        )
        tr["reset_entry_score"] = -tr["mean_transport_A"] + max(dest["P_mean"] - p_min, 0.0)
        rows.append(tr)
    df = pd.DataFrame(rows)
    df["A_drop_rank_largest_is_1"] = (-df["mean_transport_A"]).rank(method="min", ascending=False).astype(int)
    df["reset_entry_rank_largest_is_1"] = df["reset_entry_score"].rank(method="min", ascending=False).astype(int)
    key = df[df["from_stage"].eq("8-cell")].iloc[0]
    summary = {
        "transition_tested": "8-cell -> morula",
        "A_drop_rank_largest_is_1": int(key["A_drop_rank_largest_is_1"]),
        "reset_entry_rank_largest_is_1": int(key["reset_entry_rank_largest_is_1"]),
        "destination_reset_rank_lowest_is_1": int(key["destination_reset_rank_lowest_is_1"]),
        "mean_transport_A": float(key["mean_transport_A"]),
        "mean_transport_P": float(key["mean_transport_P"]),
        "reset_entry_score": float(key["reset_entry_score"]),
        "p_min": p_min,
        "p_min_definition": f"stage-agnostic global fused-particle P quantile q={POTENCY_THRESHOLD_QUANTILE}; not derived from morula",
    }
    return df, summary


def main() -> None:
    rng = np.random.default_rng(20260524)
    particles = pd.read_csv(OUT / "CSB_TRO_fused_product_particles.tsv", sep="\t")
    particles = particles[particles["stage"].isin(STAGE_ORDER)].copy()
    observed_df, observed = adjacent_metrics(particles)

    groups = {s: df.reset_index(drop=True) for s, df in particles.groupby("stage")}
    rows = []
    for i in range(500):
        sampled = []
        for stage in STAGE_ORDER:
            df = groups[stage]
            idx = rng.integers(0, len(df), size=len(df))
            sampled.append(df.iloc[idx].copy())
        boot = pd.concat(sampled, ignore_index=True)
        _, m = adjacent_metrics(boot)
        m["iteration"] = i
        rows.append(m)
    boot_df = pd.DataFrame(rows)
    summary = {
        "observed": observed,
        "bootstrap_n": int(len(boot_df)),
        "fraction_8cell_morula_largest_A_drop": float(np.mean(boot_df["A_drop_rank_largest_is_1"].eq(1))),
        "fraction_8cell_morula_largest_reset_entry_score": float(np.mean(boot_df["reset_entry_rank_largest_is_1"].eq(1))),
        "fraction_8cell_morula_destination_reset_rank1": float(np.mean(boot_df["destination_reset_rank_lowest_is_1"].eq(1))),
        "A_transport_ci025_median_ci975": [float(x) for x in np.quantile(boot_df["mean_transport_A"], [0.025, 0.5, 0.975])],
        "reset_entry_score_ci025_median_ci975": [float(x) for x in np.quantile(boot_df["reset_entry_score"], [0.025, 0.5, 0.975])],
        "interpretation": (
            "This replaces the non-informative arbitrary stage-order test. The question is not whether any forced "
            "predecessor can decrease A into morula, but whether the biologically adjacent 8-cell -> morula transition "
            "is the strongest reset-basin entry among canonical adjacent developmental transitions."
        ),
    }

    observed_df.to_csv(OUT / "CSB_TRO_adjacent_transition_specificity_observed.tsv", sep="\t", index=False)
    boot_df.to_csv(OUT / "CSB_TRO_adjacent_transition_specificity_bootstrap.tsv", sep="\t", index=False)
    (OUT / "CSB_TRO_adjacent_transition_specificity_summary.json").write_text(json.dumps(summary, indent=2, ensure_ascii=False), encoding="utf-8")
    note = f"""# CSB-TRO adjacent transition specificity

Date: 2026-05-24

This test replaces the arbitrary random-stage-order control, which is not biologically well-posed because morula is the global low-A stage.

## Question

Among canonical adjacent developmental transitions, is 8-cell -> morula the strongest entry into the low-A/high-P reset basin?

## Observed

- A-drop rank, rank 1 = largest A decrease: {observed["A_drop_rank_largest_is_1"]}
- Reset-entry rank, rank 1 = strongest: {observed["reset_entry_rank_largest_is_1"]}
- Destination reset rank, rank 1 = lowest A-P destination: {observed["destination_reset_rank_lowest_is_1"]}
- 8-cell -> morula mean transport A: {observed["mean_transport_A"]:.6f}
- 8-cell -> morula reset-entry score: {observed["reset_entry_score"]:.6f}

## Bootstrap

- n = {summary["bootstrap_n"]}
- Fraction largest A drop: {summary["fraction_8cell_morula_largest_A_drop"]:.3f}
- Fraction strongest reset-entry score: {summary["fraction_8cell_morula_largest_reset_entry_score"]:.3f}
- Fraction destination reset rank 1: {summary["fraction_8cell_morula_destination_reset_rank1"]:.3f}

## Interpretation

The earlier all-random-order test should be treated as a negative-control limitation, not a decision test. This adjacent-transition test answers the biologically constrained specificity question.
"""
    (OUT / "CSB_TRO_adjacent_transition_specificity_interpretation.md").write_text(note, encoding="utf-8")
    print(json.dumps(summary, indent=2, ensure_ascii=True))


if __name__ == "__main__":
    main()
```
