## Supplemental code for "An entropy-based diagnostic framework for characterizing methylation state dynamics during preimplantation development": csb_tro_bayesian_bootstrap_validation.py

```
from __future__ import annotations

import json
from pathlib import Path

import numpy as np
import pandas as pd


ROOT = Path(__file__).resolve().parents[1]
RESULTS = ROOT / "results"
INPUT = ROOT / "input_tables"

STAGE_ORDER = ["MII oocyte", "zygote/PN", "2-cell", "4-cell", "8-cell", "morula", "blastocyst"]
STATE_COLS = ["A", "Hm", "P", "Hr"]
DMR_STAGES = STAGE_ORDER


def weighted_quantile(values: np.ndarray, weights: np.ndarray, q: float) -> float:
    order = np.argsort(values)
    v = values[order]
    w = weights[order]
    cdf = np.cumsum(w) / np.sum(w)
    return float(v[np.searchsorted(cdf, q, side="left")])


def min_rank(series: pd.Series, stage: str, ascending: bool = True) -> int:
    return int(series.rank(method="min", ascending=ascending)[stage])


def stage_weighted_means(particles: pd.DataFrame, rng: np.random.Generator) -> pd.DataFrame:
    rows = []
    for stage in STAGE_ORDER:
        x = particles.loc[particles["stage"].eq(stage), STATE_COLS].to_numpy(dtype=float)
        weights = rng.dirichlet(np.ones(len(x)))
        mean = np.average(x, axis=0, weights=weights)
        rows.append({"stage": stage, **dict(zip(STATE_COLS, mean))})
    return pd.DataFrame(rows)


def dmr_weighted_scores(dmr: pd.DataFrame, rng: np.random.Generator) -> pd.Series:
    entropy_cols = [f"entropy_{stage}" for stage in DMR_STAGES]
    entropy = dmr[entropy_cols].to_numpy(dtype=float)
    age_weights = dmr["age_weight_per_year"].abs().to_numpy(dtype=float)
    age_weights = np.maximum(age_weights, 1e-12)
    feature_weights = rng.dirichlet(np.ones(len(dmr))) * age_weights
    score = np.average(entropy, axis=0, weights=feature_weights)
    return pd.Series(score, index=DMR_STAGES)


def run_bayesian_bootstrap(n_iter: int = 2000) -> tuple[pd.DataFrame, dict]:
    rng = np.random.default_rng(20260524)
    particles = pd.read_csv(RESULTS / "CSB_TRO_fused_product_particles.tsv", sep="\t")
    dmr = pd.read_csv(INPUT / "TRO_interpretability_DMR_contribution_ranking.tsv", sep="\t")

    for col in ["age_weight_per_year", *[f"entropy_{stage}" for stage in DMR_STAGES]]:
        dmr[col] = pd.to_numeric(dmr[col], errors="coerce")
        dmr[col] = dmr[col].fillna(float(dmr[col].median()))

    p_global = particles["P"].to_numpy(dtype=float)
    p_min_fixed = float(np.quantile(p_global, 0.60))

    rows = []
    for i in range(n_iter):
        stage_means = stage_weighted_means(particles, rng).set_index("stage")
        dmr_scores = dmr_weighted_scores(dmr, rng)

        # Stage-agnostic threshold. It is fixed from all fused particles and not derived from morula.
        eligible = stage_means["P"] >= p_min_fixed
        if eligible.any():
            reset_candidate = str(stage_means.loc[eligible, "A"].idxmin())
        else:
            reset_candidate = str((stage_means["A"] - stage_means["P"]).idxmin())

        dmr_min_stage = str(dmr_scores.idxmin())
        rows.append(
            {
                "iteration": i,
                "p_min_fixed_q60": p_min_fixed,
                "particle_reset_candidate": reset_candidate,
                "particle_morula_A_rank1": min_rank(stage_means["A"], "morula", ascending=True),
                "particle_morula_P_rank_high": min_rank(stage_means["P"], "morula", ascending=False),
                "particle_morula_reset_rank": min_rank(stage_means["A"] - stage_means["P"], "morula", ascending=True),
                "particle_morula_A": float(stage_means.loc["morula", "A"]),
                "particle_morula_P": float(stage_means.loc["morula", "P"]),
                "particle_8cell_A": float(stage_means.loc["8-cell", "A"]),
                "particle_8cell_to_morula_A_drop": float(stage_means.loc["8-cell", "A"] - stage_means.loc["morula", "A"]),
                "dmr_min_stage": dmr_min_stage,
                "dmr_morula_rank1": min_rank(dmr_scores, "morula", ascending=True),
                "dmr_morula_score": float(dmr_scores["morula"]),
                "dmr_8cell_score": float(dmr_scores["8-cell"]),
                "dmr_8cell_to_morula_drop": float(dmr_scores["8-cell"] - dmr_scores["morula"]),
                "joint_particle_and_dmr_morula": bool(reset_candidate == "morula" and dmr_min_stage == "morula"),
            }
        )

    df = pd.DataFrame(rows)
    summary = {
        "model": "CSB-TRO Bayesian bootstrap posterior validation",
        "date": "2026-05-24",
        "n_iter": n_iter,
        "p_min_fixed_q60": p_min_fixed,
        "p_min_definition": "stage-agnostic global fused-particle P quantile q=0.60; not derived from morula",
        "particle_posterior": {
            "Pr_reset_candidate_morula": float(df["particle_reset_candidate"].eq("morula").mean()),
            "Pr_morula_A_rank1": float(df["particle_morula_A_rank1"].eq(1).mean()),
            "Pr_morula_P_top2": float((df["particle_morula_P_rank_high"] <= 2).mean()),
            "Pr_morula_reset_rank1": float(df["particle_morula_reset_rank"].eq(1).mean()),
            "Pr_8cell_to_morula_A_drop_positive": float((df["particle_8cell_to_morula_A_drop"] > 0).mean()),
            "median_8cell_to_morula_A_drop": float(df["particle_8cell_to_morula_A_drop"].median()),
        },
        "dmr_posterior": {
            "Pr_DMR_min_stage_morula": float(df["dmr_min_stage"].eq("morula").mean()),
            "Pr_DMR_morula_rank1": float(df["dmr_morula_rank1"].eq(1).mean()),
            "Pr_DMR_8cell_to_morula_drop_positive": float((df["dmr_8cell_to_morula_drop"] > 0).mean()),
            "median_DMR_8cell_to_morula_drop": float(df["dmr_8cell_to_morula_drop"].median()),
        },
        "joint_posterior": {
            "Pr_particle_reset_and_DMR_min_both_morula": float(df["joint_particle_and_dmr_morula"].mean())
        },
        "interpretation": (
            "Bayesian bootstrap resampling gives posterior-style stability estimates for the reset-basin call. "
            "The analysis bootstraps fused stage particles and age-DMR feature weights without using a morula-derived "
            "training constraint. This supports local generalization/stability, not broad supervised forecasting."
        ),
    }
    return df, summary


def main() -> None:
    df, summary = run_bayesian_bootstrap()
    df.to_csv(RESULTS / "CSB_TRO_bayesian_bootstrap_posterior_validation.tsv", sep="\t", index=False)
    (RESULTS / "CSB_TRO_bayesian_bootstrap_posterior_validation_summary.json").write_text(
        json.dumps(summary, indent=2, ensure_ascii=False),
        encoding="utf-8",
    )

    note = f"""# CSB-TRO Bayesian Bootstrap Posterior Validation

Date: 2026-05-24

## Design

This analysis resamples fused stage particles and age-DMR feature weights with Bayesian bootstrap weights. The reset candidate is defined after resampling as the stage with minimum particle-level `A` among stages satisfying a stage-agnostic potency threshold:

`P_min = q60(P)` over all fused particles.

No morula-derived threshold or morula training constraint is used.

## Main posterior-style readouts

- Iterations: {summary["n_iter"]}
- Pr(particle reset candidate = morula): {summary["particle_posterior"]["Pr_reset_candidate_morula"]:.3f}
- Pr(particle morula A rank 1): {summary["particle_posterior"]["Pr_morula_A_rank1"]:.3f}
- Pr(particle morula P top 2): {summary["particle_posterior"]["Pr_morula_P_top2"]:.3f}
- Pr(particle morula reset rank 1): {summary["particle_posterior"]["Pr_morula_reset_rank1"]:.3f}
- Pr(particle 8-cell to morula A drop positive): {summary["particle_posterior"]["Pr_8cell_to_morula_A_drop_positive"]:.3f}
- Pr(DMR minimum stage = morula): {summary["dmr_posterior"]["Pr_DMR_min_stage_morula"]:.3f}
- Pr(DMR morula rank 1): {summary["dmr_posterior"]["Pr_DMR_morula_rank1"]:.3f}
- Pr(DMR 8-cell to morula drop positive): {summary["dmr_posterior"]["Pr_DMR_8cell_to_morula_drop_positive"]:.3f}
- Pr(particle reset and DMR minimum both morula): {summary["joint_posterior"]["Pr_particle_reset_and_DMR_min_both_morula"]:.3f}

## Interpretation

This is a posterior-style stability analysis, not a claim of high-accuracy supervised forecasting. It supports the statement that the morula reset-basin call is stable under particle and DMR uncertainty, while broad stage-level distributional forecasting remains limited.
"""
    (RESULTS / "CSB_TRO_bayesian_bootstrap_posterior_validation_interpretation.md").write_text(note, encoding="utf-8")
    print(json.dumps(summary, indent=2, ensure_ascii=True))


if __name__ == "__main__":
    main()
```
