## Supplemental code for "An entropy-based diagnostic framework for characterizing methylation state dynamics during preimplantation development": csb_tro_dmr_graph_laplacian.py

```
from __future__ import annotations

import json
from pathlib import Path

import numpy as np
import pandas as pd


OUT = Path("E:/实验更新_5_22/csb_tro_dynamics")
DMR_TABLE = Path("E:/TRO_Project_backup_2026-05-21/TRO_Project_current_results/tables/TRO_interpretability_DMR_contribution_ranking.tsv")
STAGES = ["MII oocyte", "zygote/PN", "2-cell", "4-cell", "8-cell", "morula", "blastocyst"]
LAMBDA_G_GRID = [0.0, 0.001, 0.005, 0.01, 0.05, 0.10, 0.25, 0.50, 1.0]
K_NEIGHBORS = 10


def zscore_columns(x: np.ndarray) -> np.ndarray:
    return (x - x.mean(axis=0, keepdims=True)) / np.maximum(x.std(axis=0, keepdims=True), 1e-12)


def pairwise_sqdist(x: np.ndarray) -> np.ndarray:
    xx = np.sum(x * x, axis=1, keepdims=True)
    d = xx + xx.T - 2.0 * (x @ x.T)
    return np.maximum(d, 0.0)


def chr_numeric(chrom: str) -> int:
    c = str(chrom).replace("chr", "")
    if c == "X":
        return 23
    if c == "Y":
        return 24
    try:
        return int(c)
    except ValueError:
        return 99


def build_dmr_graph(dmr: pd.DataFrame) -> tuple[pd.DataFrame, pd.DataFrame, pd.DataFrame]:
    entropy_cols = [f"entropy_{s}" for s in STAGES]
    dmr = dmr.copy()
    numeric_cols = entropy_cols + [f"beta_{s}" for s in STAGES] + [
        "age_weight_per_year",
        "delta_H_8cell_to_morula",
        "contribution_8cell_to_morula",
    ]
    for col in numeric_cols:
        if col in dmr.columns:
            dmr[col] = pd.to_numeric(dmr[col], errors="coerce")
            dmr[col] = dmr[col].fillna(float(dmr[col].median()))
    traj = zscore_columns(dmr[entropy_cols].to_numpy(dtype=float))
    d2 = pairwise_sqdist(traj)
    nonzero = d2[d2 > 0]
    sigma2 = float(np.median(nonzero)) if nonzero.size else 1.0
    entropy_similarity = np.exp(-d2 / max(2.0 * sigma2, 1e-12))

    age = dmr["age_weight_per_year"].fillna(0.0).to_numpy(dtype=float)
    age_scale = float(np.std(age)) or 1.0
    age_similarity = np.exp(-((age[:, None] - age[None, :]) ** 2) / max(2.0 * age_scale * age_scale, 1e-12))

    mid = ((dmr["start"].to_numpy(dtype=float) + dmr["end"].to_numpy(dtype=float)) / 2.0)
    chrom = dmr["chr"].astype(str).to_numpy()
    same_chr = chrom[:, None] == chrom[None, :]
    genomic_similarity = np.where(same_chr, np.exp(-np.abs(mid[:, None] - mid[None, :]) / 1_000_000.0), 0.0)

    nearest_gene = dmr["nearest_gene"].fillna("").astype(str).to_numpy()
    valid_gene = (nearest_gene != "") & (~pd.Series(nearest_gene).str.startswith("LOC").to_numpy())
    same_gene = (nearest_gene[:, None] == nearest_gene[None, :]) & valid_gene[:, None] & valid_gene[None, :]
    same_gene = same_gene.astype(float)

    gene_context = dmr["gene_context"].fillna("").astype(str).to_numpy()
    cpg_context = dmr["cpg_context"].fillna("").astype(str).to_numpy()
    same_context = ((gene_context[:, None] == gene_context[None, :]).astype(float) + (cpg_context[:, None] == cpg_context[None, :]).astype(float)) / 2.0

    score = (
        0.65 * entropy_similarity
        + 0.15 * age_similarity
        + 0.10 * genomic_similarity
        + 0.05 * same_gene
        + 0.05 * same_context
    )
    score = np.nan_to_num(score, nan=0.0, posinf=0.0, neginf=0.0)
    np.fill_diagonal(score, 0.0)

    n = len(dmr)
    w = np.zeros((n, n), dtype=float)
    for i in range(n):
        idx = np.argsort(score[i])[::-1][:K_NEIGHBORS]
        w[i, idx] = score[i, idx]
    w = np.maximum(w, w.T)
    np.fill_diagonal(w, 0.0)
    degree = np.diag(w.sum(axis=1))
    lap = degree - w

    node_df = dmr[
        [
            "cluster_name",
            "chr",
            "start",
            "end",
            "nearest_gene",
            "gene_context",
            "cpg_context",
            "age_weight_per_year",
            "contribution_8cell_to_morula",
            "reset_driver_rank_8cell_to_morula",
        ]
        + entropy_cols
    ].copy()
    node_df["graph_degree_weighted"] = w.sum(axis=1)
    node_df["graph_degree_unweighted"] = (w > 0).sum(axis=1)

    edge_rows = []
    rows, cols = np.nonzero(np.triu(w, k=1) > 0)
    for i, j in zip(rows, cols):
        edge_rows.append(
            {
                "source": dmr.loc[i, "cluster_name"],
                "target": dmr.loc[j, "cluster_name"],
                "weight": float(w[i, j]),
                "same_chr": bool(same_chr[i, j]),
                "same_gene": bool(same_gene[i, j]),
                "entropy_similarity": float(entropy_similarity[i, j]),
                "age_weight_similarity": float(age_similarity[i, j]),
                "genomic_similarity": float(genomic_similarity[i, j]),
                "same_context_score": float(same_context[i, j]),
                "source_gene": dmr.loc[i, "nearest_gene"],
                "target_gene": dmr.loc[j, "nearest_gene"],
            }
        )
    edge_df = pd.DataFrame(edge_rows).sort_values("weight", ascending=False).reset_index(drop=True)
    lap_df = pd.DataFrame(lap, index=dmr["cluster_name"], columns=dmr["cluster_name"])
    return node_df, edge_df, lap_df


def graph_smoothness(dmr: pd.DataFrame, lap: pd.DataFrame) -> dict[str, float]:
    entropy_cols = [f"entropy_{s}" for s in STAGES]
    beta_cols = [f"beta_{s}" for s in STAGES]
    dmr = dmr.copy()
    for col in entropy_cols + beta_cols + ["age_weight_per_year", "delta_H_8cell_to_morula", "contribution_8cell_to_morula"]:
        dmr[col] = pd.to_numeric(dmr[col], errors="coerce")
        dmr[col] = dmr[col].fillna(float(dmr[col].median()))
    l = lap.to_numpy(dtype=float)
    x_entropy = zscore_columns(dmr[entropy_cols].to_numpy(dtype=float))
    x_beta = zscore_columns(dmr[beta_cols].to_numpy(dtype=float))
    contribution = dmr[["age_weight_per_year", "delta_H_8cell_to_morula", "contribution_8cell_to_morula"]].fillna(0.0).to_numpy(dtype=float)
    x_contrib = zscore_columns(contribution)

    edge_weight_sum = float(l.diagonal().sum() / 2.0)

    def trace_term(x: np.ndarray) -> float:
        return float(np.trace(x.T @ l @ x))

    c_entropy = trace_term(x_entropy)
    c_beta = trace_term(x_beta)
    c_contrib = trace_term(x_contrib)
    c_total = c_entropy + c_beta + c_contrib
    denom = max(edge_weight_sum, 1e-12)
    return {
        "C_G_entropy_trajectory": c_entropy,
        "C_G_beta_trajectory": c_beta,
        "C_G_contribution_features": c_contrib,
        "C_G_total_raw": c_total,
        "C_G_total_edge_normalized": c_total / denom,
        "edge_weight_sum": edge_weight_sum,
    }


def main() -> None:
    dmr = pd.read_csv(DMR_TABLE, sep="\t")
    dmr = dmr.sort_values("reset_driver_rank_8cell_to_morula").reset_index(drop=True)

    node_df, edge_df, lap_df = build_dmr_graph(dmr)
    g = graph_smoothness(dmr, lap_df)

    path_summary = json.loads((OUT / "CSB_TRO_path_space_summary.json").read_text(encoding="utf-8"))
    j_path = float(path_summary["J_path_total"])
    rows = []
    for lambda_g in LAMBDA_G_GRID:
        rows.append(
            {
                "lambda_G": lambda_g,
                **g,
                "lambda_G_C_G_edge_normalized": lambda_g * g["C_G_total_edge_normalized"],
                "J_path_without_graph": j_path,
                "J_path_with_DMR_graph": j_path + lambda_g * g["C_G_total_edge_normalized"],
            }
        )
    sensitivity = pd.DataFrame(rows)

    summary = {
        "model": "CSB-TRO DMR graph Laplacian regularizer",
        "date": "2026-05-24",
        "source_table": str(DMR_TABLE),
        "graph_level": "DMR cluster graph",
        "n_dmr_nodes": int(len(node_df)),
        "n_edges": int(len(edge_df)),
        "k_neighbors": K_NEIGHBORS,
        "edge_weight_definition": {
            "entropy_trajectory_similarity": 0.65,
            "age_weight_similarity": 0.15,
            "same_chromosome_genomic_proximity": 0.10,
            "same_nearest_gene": 0.05,
            "same_gene_or_cpg_context": 0.05,
        },
        "graph_terms": g,
        "J_path_without_graph": j_path,
        "recommended_lambda_G": 0.10,
        "recommended_J_path_with_DMR_graph": float(
            sensitivity.loc[sensitivity["lambda_G"].eq(0.10), "J_path_with_DMR_graph"].iloc[0]
        ),
        "top_edges": edge_df.head(10).to_dict(orient="records"),
        "top_degree_nodes": node_df.sort_values("graph_degree_weighted", ascending=False).head(10).to_dict(orient="records"),
    }

    node_df.to_csv(OUT / "CSB_TRO_DMR_graph_nodes.tsv", sep="\t", index=False)
    edge_df.to_csv(OUT / "CSB_TRO_DMR_graph_edges.tsv", sep="\t", index=False)
    lap_df.to_csv(OUT / "CSB_TRO_DMR_graph_laplacian.tsv", sep="\t")
    sensitivity.to_csv(OUT / "CSB_TRO_DMR_graph_laplacian_objective_sensitivity.tsv", sep="\t", index=False)
    (OUT / "CSB_TRO_DMR_graph_laplacian_summary.json").write_text(json.dumps(summary, indent=2, ensure_ascii=False), encoding="utf-8")

    note = f"""# CSB-TRO DMR graph Laplacian regularizer

Date: 2026-05-24

This step replaces the earlier four-variable graph audit with a DMR-node graph.

## Graph

- Nodes: DMR clusters from `TRO_interpretability_DMR_contribution_ranking.tsv`
- Number of DMR nodes: {summary["n_dmr_nodes"]}
- Number of undirected graph edges: {summary["n_edges"]}
- k-nearest neighbors per node before symmetrization: {K_NEIGHBORS}

Edge weights combine:

- entropy trajectory similarity across developmental stages
- age-weight similarity
- same-chromosome genomic proximity
- same nearest gene
- shared gene/CpG context

## Laplacian term

`C_G = Tr(X^T L_G X)`

where `X` contains DMR-level entropy trajectories, methylation beta trajectories, and reset-contribution features.

## Objective impact

- C_G entropy trajectory: {g["C_G_entropy_trajectory"]:.6f}
- C_G beta trajectory: {g["C_G_beta_trajectory"]:.6f}
- C_G contribution features: {g["C_G_contribution_features"]:.6f}
- C_G total raw: {g["C_G_total_raw"]:.6f}
- C_G total edge-normalized: {g["C_G_total_edge_normalized"]:.6f}
- J path without graph: {j_path:.6f}
- J path with DMR graph at lambda_G=0.10: {summary["recommended_J_path_with_DMR_graph"]:.6f}

## Interpretation

This is the biologically grounded graph-Laplacian layer required by the strict CSB-TRO formulation. It should supersede the earlier four-state-variable graph audit for manuscript reporting.
"""
    (OUT / "CSB_TRO_DMR_graph_laplacian_interpretation.md").write_text(note, encoding="utf-8")
    print(json.dumps(summary, indent=2, ensure_ascii=True))


if __name__ == "__main__":
    main()
```
