## Supplemental code for "An entropy-based diagnostic framework for characterizing methylation state dynamics during preimplantation development": csb_tro_fokker_planck_export.py

```
from __future__ import annotations

import json
from pathlib import Path

import numpy as np
import pandas as pd


OUT = Path("E:/实验更新_5_22/csb_tro_dynamics")
STATE_COLS = ["A", "Hm", "P", "Hr"]
VEL_COLS = ["vA", "vHm", "vP", "vHr"]
STAGE_ORDER = ["MII oocyte", "zygote/PN", "2-cell", "4-cell", "8-cell", "morula", "blastocyst"]
RIDGE_ALPHA = 1e-4


def ridge_fit(x: np.ndarray, y: np.ndarray, alpha: float = RIDGE_ALPHA) -> tuple[np.ndarray, float]:
    xtx = x.T @ x
    penalty = np.eye(xtx.shape[0]) * alpha
    penalty[0, 0] = 0.0
    beta = np.linalg.solve(xtx + penalty, x.T @ y)
    pred = x @ beta
    ss_res = float(np.sum((y - pred) ** 2))
    ss_tot = float(np.sum((y - y.mean()) ** 2))
    r2 = 1.0 - ss_res / max(ss_tot, 1e-12)
    return beta, r2


def fit_transition_drifts(velocity: pd.DataFrame) -> tuple[pd.DataFrame, pd.DataFrame]:
    rows = []
    pred_rows = []
    feature_cols = ["intercept"] + STATE_COLS
    for transition_order, df in velocity.groupby("transition_order"):
        df = df.copy()
        x = np.column_stack([np.ones(len(df)), df[STATE_COLS].to_numpy(dtype=float)])
        for vel_col, state_col in zip(VEL_COLS, STATE_COLS):
            beta, r2 = ridge_fit(x, df[vel_col].to_numpy(dtype=float))
            rows.append(
                {
                    "transition_order": int(transition_order),
                    "from_stage": str(df["stage"].iloc[0]),
                    "to_stage": str(df["to_stage"].iloc[0]),
                    "drift_component": f"b_{state_col}",
                    "r2": r2,
                    **{f"coef_{name}": float(value) for name, value in zip(feature_cols, beta)},
                }
            )
            pred_rows.append(
                pd.DataFrame(
                    {
                        "particle_id": df["particle_id"].to_numpy(),
                        "transition_order": int(transition_order),
                        "from_stage": df["stage"].to_numpy(),
                        "to_stage": df["to_stage"].to_numpy(),
                        "drift_component": f"b_{state_col}",
                        "observed_velocity": df[vel_col].to_numpy(dtype=float),
                        "predicted_drift": x @ beta,
                    }
                )
            )
    return pd.DataFrame(rows), pd.concat(pred_rows, ignore_index=True)


def diffusion_from_couplings(couplings: pd.DataFrame) -> pd.DataFrame:
    rows = []
    delta_cols = ["delta_A", "delta_Hm", "delta_P", "delta_Hr"]
    for (order, from_stage, to_stage), df in couplings.groupby(["transition_order", "from_stage", "to_stage"]):
        p = df["probability"].to_numpy(dtype=float)
        p = p / max(p.sum(), 1e-300)
        delta_matrix = df[delta_cols].to_numpy(dtype=float)
        mean_vector = np.sum(p[:, None] * delta_matrix, axis=0)
        centered = delta_matrix - mean_vector[None, :]
        sigma = centered.T @ (centered * p[:, None])
        row = {
            "transition_order": int(order),
            "from_stage": from_stage,
            "to_stage": to_stage,
            "probability_sum": float(p.sum()),
        }
        d_values = []
        for i, (delta_col, state_col) in enumerate(zip(delta_cols, STATE_COLS)):
            mean = float(mean_vector[i])
            var = float(sigma[i, i])
            diffusivity = 0.5 * var
            row[f"mean_delta_{state_col}"] = mean
            row[f"variance_delta_{state_col}"] = var
            row[f"Sigma_{state_col}_{state_col}"] = var
            row[f"D_{state_col}"] = diffusivity
            d_values.append(diffusivity)
        for i, state_i in enumerate(STATE_COLS):
            for j, state_j in enumerate(STATE_COLS):
                row[f"Sigma_{state_i}_{state_j}"] = float(sigma[i, j])
        row["D_isotropic_4d_mean"] = float(np.mean(d_values))
        row["D_isotropic_AP_mean"] = float(np.mean([row["D_A"], row["D_P"]]))
        rows.append(row)
    return pd.DataFrame(rows).sort_values("transition_order").reset_index(drop=True)


def main() -> None:
    velocity = pd.read_csv(OUT / "CSB_TRO_path_space_velocity_field.tsv", sep="\t")
    couplings = pd.read_csv(OUT / "CSB_TRO_path_space_transition_couplings.tsv", sep="\t")

    drift_coef, drift_pred = fit_transition_drifts(velocity)
    diffusion = diffusion_from_couplings(couplings)

    summary = {
        "model": "CSB-TRO Fokker-Planck export",
        "date": "2026-05-24",
        "drift_model": "piecewise-in-time linear ridge drift b_k(z)=beta_0k + B_k z, fitted to path-space conditional velocities",
        "diffusion_model": "transition-wise full diffusion matrix Sigma_k from coupling second moments, with diagonal D_i = 0.5 Sigma_ii for Delta t = 1",
        "fokker_planck_form": "partial_t p_t(z) = -div_z(b(z,t) p_t(z)) + 0.5 sum_ij Sigma_ij(t) partial_{z_i z_j} p_t(z)",
        "reduced_AP_form": "partial_t p_t(A,P) = -partial_A(b_A p_t) - partial_P(b_P p_t) + D_A partial_AA p_t + D_P partial_PP p_t",
        "mean_drift_r2_by_component": {
            comp: float(val) for comp, val in drift_coef.groupby("drift_component")["r2"].mean().items()
        },
        "mean_diffusion": {
            "D_A": float(diffusion["D_A"].mean()),
            "D_Hm": float(diffusion["D_Hm"].mean()),
            "D_P": float(diffusion["D_P"].mean()),
            "D_Hr": float(diffusion["D_Hr"].mean()),
            "D_isotropic_4d_mean": float(diffusion["D_isotropic_4d_mean"].mean()),
            "D_isotropic_AP_mean": float(diffusion["D_isotropic_AP_mean"].mean()),
        },
        "key_transition_8cell_to_morula_diffusion": diffusion[diffusion["from_stage"].eq("8-cell")].iloc[0].to_dict(),
    }

    drift_coef.to_csv(OUT / "CSB_TRO_fokker_planck_drift_coefficients.tsv", sep="\t", index=False)
    drift_pred.to_csv(OUT / "CSB_TRO_fokker_planck_drift_predictions.tsv", sep="\t", index=False)
    diffusion.to_csv(OUT / "CSB_TRO_fokker_planck_diffusion_estimates.tsv", sep="\t", index=False)
    (OUT / "CSB_TRO_fokker_planck_summary.json").write_text(json.dumps(summary, indent=2, ensure_ascii=False), encoding="utf-8")

    note = f"""# CSB-TRO Fokker-Planck export

Date: 2026-05-24

This step converts the learned Markov path-space velocity field into a piecewise continuous drift and diffusion representation.

## Drift

For each transition interval:

`b_k(z) = beta_0,k + B_k z`

where `z = [A, Hm, P, Hr]`.

The coefficients are fitted by ridge regression to the conditional CSB velocities.

## Diffusion

For each transition and state dimension:

`Sigma_k = Cov_pi(delta z)`

and the diagonal shorthand:

`D_i,k = 0.5 Sigma_ii,k`

with `Delta t = 1`, giving the Fokker-Planck form:

`partial_t p_t(z) = -div_z(b(z,t) p_t(z)) + 0.5 sum_ij Sigma_ij(t) partial_{{z_i z_j}} p_t(z)`

Reduced A-P visualization form:

`partial_t p_t(A,P) = -partial_A(b_A p_t) - partial_P(b_P p_t) + D_A partial_AA p_t + D_P partial_PP p_t`

## Mean diffusion

- D_A mean: {summary["mean_diffusion"]["D_A"]:.6f}
- D_Hm mean: {summary["mean_diffusion"]["D_Hm"]:.6f}
- D_P mean: {summary["mean_diffusion"]["D_P"]:.6f}
- D_Hr mean: {summary["mean_diffusion"]["D_Hr"]:.6f}
- D isotropic A-P mean: {summary["mean_diffusion"]["D_isotropic_AP_mean"]:.6f}

## Mean drift R2

{pd.Series(summary["mean_drift_r2_by_component"]).to_string()}

## Interpretation

This file set is the PDE export layer. It does not prove the biological model by itself; it expresses the learned CSB-TRO distributional flow as a Fokker-Planck-compatible drift-diffusion system for visualization and downstream numerical simulation.
"""
    (OUT / "CSB_TRO_fokker_planck_equation.md").write_text(note, encoding="utf-8")
    print(json.dumps(summary, indent=2, ensure_ascii=True))


if __name__ == "__main__":
    main()
```
