## Supplemental code for "An entropy-based diagnostic framework for characterizing methylation state dynamics during preimplantation development": csb_tro_full_stage_distribution_prediction.py

```
from __future__ import annotations

import json
from pathlib import Path

import numpy as np
import pandas as pd


ROOT = Path(__file__).resolve().parents[1]
RESULTS = ROOT / "results"

STAGE_ORDER = ["MII oocyte", "zygote/PN", "2-cell", "4-cell", "8-cell", "morula", "blastocyst"]
STATE_COLS = ["A", "Hm", "P", "Hr"]


def pairwise_sqdist(x: np.ndarray, y: np.ndarray) -> np.ndarray:
    x2 = np.sum(x * x, axis=1)[:, None]
    y2 = np.sum(y * y, axis=1)[None, :]
    return np.maximum(x2 + y2 - 2.0 * x @ y.T, 0.0)


def sinkhorn_coupling(x: np.ndarray, y: np.ndarray, epsilon_scale: float = 0.20, n_iter: int = 1000) -> np.ndarray:
    n, m = len(x), len(y)
    a = np.full(n, 1.0 / n)
    b = np.full(m, 1.0 / m)
    cost = pairwise_sqdist(x, y)
    positive = cost[cost > 0]
    base = float(np.median(positive)) if len(positive) else 1.0
    epsilon = max(epsilon_scale * base, 1e-3)
    kernel = np.exp(-cost / epsilon)
    kernel = np.maximum(kernel, 1e-300)
    u = np.ones(n)
    v = np.ones(m)
    for _ in range(n_iter):
        u = a / (kernel @ v + 1e-300)
        v = b / (kernel.T @ u + 1e-300)
    pi = (u[:, None] * kernel) * v[None, :]
    pi /= pi.sum()
    return pi


def sample_geodesic_distribution(
    x_prev: np.ndarray,
    x_next: np.ndarray,
    n_samples: int,
    rng: np.random.Generator,
    alpha: float = 0.5,
) -> np.ndarray:
    pi = sinkhorn_coupling(x_prev, x_next)
    flat = pi.ravel()
    flat /= flat.sum()
    choices = rng.choice(len(flat), size=n_samples, replace=True, p=flat)
    i, j = np.unravel_index(choices, pi.shape)
    return (1.0 - alpha) * x_prev[i] + alpha * x_next[j]


def energy_distance(x: np.ndarray, y: np.ndarray, rng: np.random.Generator, max_n: int = 260) -> float:
    if len(x) > max_n:
        x = x[rng.choice(len(x), size=max_n, replace=False)]
    if len(y) > max_n:
        y = y[rng.choice(len(y), size=max_n, replace=False)]
    xy = np.linalg.norm(x[:, None, :] - y[None, :, :], axis=2).mean()
    xx = np.linalg.norm(x[:, None, :] - x[None, :, :], axis=2).mean()
    yy = np.linalg.norm(y[:, None, :] - y[None, :, :], axis=2).mean()
    return float(2.0 * xy - xx - yy)


def rbf_mmd2(x: np.ndarray, y: np.ndarray, rng: np.random.Generator, max_n: int = 260) -> float:
    if len(x) > max_n:
        x = x[rng.choice(len(x), size=max_n, replace=False)]
    if len(y) > max_n:
        y = y[rng.choice(len(y), size=max_n, replace=False)]
    pooled = np.vstack([x, y])
    d2 = pairwise_sqdist(pooled, pooled)
    positive = d2[d2 > 0]
    gamma = 1.0 / max(float(np.median(positive)), 1e-6) if len(positive) else 1.0
    kxx = np.exp(-gamma * pairwise_sqdist(x, x)).mean()
    kyy = np.exp(-gamma * pairwise_sqdist(y, y)).mean()
    kxy = np.exp(-gamma * pairwise_sqdist(x, y)).mean()
    return float(kxx + kyy - 2.0 * kxy)


def draw_rows(x: np.ndarray, n: int, rng: np.random.Generator) -> np.ndarray:
    return x[rng.integers(0, len(x), size=n)]


def full_stage_leave_one_out(n_repeats: int = 80) -> tuple[pd.DataFrame, pd.DataFrame, dict]:
    rng = np.random.default_rng(20260524)
    particles = pd.read_csv(RESULTS / "CSB_TRO_fused_product_particles.tsv", sep="\t")
    by_stage = {stage: particles.loc[particles["stage"].eq(stage), STATE_COLS].to_numpy(dtype=float) for stage in STAGE_ORDER}
    global_mean = particles[STATE_COLS].mean().to_numpy(dtype=float)

    rows = []
    pred_rows = []
    for k in range(1, len(STAGE_ORDER) - 1):
        prev_stage = STAGE_ORDER[k - 1]
        target_stage = STAGE_ORDER[k]
        next_stage = STAGE_ORDER[k + 1]
        x_prev = by_stage[prev_stage]
        x_obs = by_stage[target_stage]
        x_next = by_stage[next_stage]
        n = len(x_obs)

        for repeat in range(n_repeats):
            pred_csb = sample_geodesic_distribution(x_prev, x_next, n, rng, alpha=0.5)
            pred_naive = 0.5 * draw_rows(x_prev, n, rng) + 0.5 * draw_rows(x_next, n, rng)
            pred_prev = draw_rows(x_prev, n, rng)
            pred_next = draw_rows(x_next, n, rng)
            pred_global = np.tile(global_mean, (n, 1))

            candidates = {
                "csb_ot_geodesic": pred_csb,
                "naive_random_midpoint": pred_naive,
                "static_previous_stage": pred_prev,
                "static_next_stage": pred_next,
                "static_global_mean": pred_global,
            }
            metrics = {}
            for name, pred in candidates.items():
                metrics[f"energy_{name}"] = energy_distance(x_obs, pred, rng)
                metrics[f"mmd2_{name}"] = rbf_mmd2(x_obs, pred, rng)
            rows.append(
                {
                    "repeat": repeat,
                    "heldout_stage": target_stage,
                    "prev_stage": prev_stage,
                    "next_stage": next_stage,
                    **metrics,
                }
            )
            if repeat == 0:
                for idx, vec in enumerate(pred_csb):
                    pred_rows.append(
                        {
                            "heldout_stage": target_stage,
                            "prediction_method": "csb_ot_geodesic",
                            "pred_particle_id": f"{target_stage.replace(' ', '_').replace('/', '_')}:pred:{idx:04d}",
                            **dict(zip(STATE_COLS, vec)),
                        }
                    )

    df = pd.DataFrame(rows)
    for baseline in ["naive_random_midpoint", "static_previous_stage", "static_next_stage", "static_global_mean"]:
        df[f"csb_beats_{baseline}_energy"] = df["energy_csb_ot_geodesic"] < df[f"energy_{baseline}"]
        df[f"csb_beats_{baseline}_mmd2"] = df["mmd2_csb_ot_geodesic"] < df[f"mmd2_{baseline}"]

    stage_summary_rows = []
    for stage, g in df.groupby("heldout_stage", sort=False):
        row = {
            "heldout_stage": stage,
            "n_repeats": int(len(g)),
            "median_energy_csb_ot_geodesic": float(g["energy_csb_ot_geodesic"].median()),
            "median_energy_naive_random_midpoint": float(g["energy_naive_random_midpoint"].median()),
            "median_energy_static_previous_stage": float(g["energy_static_previous_stage"].median()),
            "median_energy_static_next_stage": float(g["energy_static_next_stage"].median()),
            "median_energy_static_global_mean": float(g["energy_static_global_mean"].median()),
            "median_mmd2_csb_ot_geodesic": float(g["mmd2_csb_ot_geodesic"].median()),
        }
        for baseline in ["naive_random_midpoint", "static_previous_stage", "static_next_stage", "static_global_mean"]:
            row[f"fraction_csb_beats_{baseline}_energy"] = float(g[f"csb_beats_{baseline}_energy"].mean())
            row[f"fraction_csb_beats_{baseline}_mmd2"] = float(g[f"csb_beats_{baseline}_mmd2"].mean())
        stage_summary_rows.append(row)
    stage_summary = pd.DataFrame(stage_summary_rows)

    summary = {
        "model": "CSB-TRO full-stage distribution prediction",
        "date": "2026-05-24",
        "task": "leave-one-stage-out full empirical distribution prediction",
        "method": "entropic OT/CSB geodesic midpoint between neighboring observed stage distributions",
        "n_internal_heldout_stages": int(stage_summary.shape[0]),
        "n_repeats_per_stage": n_repeats,
        "fraction_stage_median_csb_beats_naive_midpoint": float((stage_summary["median_energy_csb_ot_geodesic"] < stage_summary["median_energy_naive_random_midpoint"]).mean()),
        "fraction_stage_median_csb_beats_previous": float((stage_summary["median_energy_csb_ot_geodesic"] < stage_summary["median_energy_static_previous_stage"]).mean()),
        "fraction_stage_median_csb_beats_next": float((stage_summary["median_energy_csb_ot_geodesic"] < stage_summary["median_energy_static_next_stage"]).mean()),
        "fraction_stage_median_csb_beats_global_mean": float((stage_summary["median_energy_csb_ot_geodesic"] < stage_summary["median_energy_static_global_mean"]).mean()),
        "overall_repeat_fraction_csb_beats_naive_midpoint": float(df["csb_beats_naive_random_midpoint_energy"].mean()),
        "overall_repeat_fraction_csb_beats_previous": float(df["csb_beats_static_previous_stage_energy"].mean()),
        "overall_repeat_fraction_csb_beats_next": float(df["csb_beats_static_next_stage_energy"].mean()),
        "overall_repeat_fraction_csb_beats_global_mean": float(df["csb_beats_static_global_mean_energy"].mean()),
        "interpretation": (
            "This implements full held-out stage distribution prediction rather than stage-mean forecasting. "
            "It is a leave-one-stage-out interpolation task using the neighboring stage distributions, so it tests "
            "whether the learned distributional geometry improves over simple baselines. It is still not prospective "
            "prediction of a future stage from only earlier stages."
        ),
    }
    return df, stage_summary, pd.DataFrame(pred_rows), summary


def main() -> None:
    repeat_df, stage_summary, pred_particles, summary = full_stage_leave_one_out()
    repeat_df.to_csv(RESULTS / "CSB_TRO_full_stage_distribution_prediction_repeats.tsv", sep="\t", index=False)
    stage_summary.to_csv(RESULTS / "CSB_TRO_full_stage_distribution_prediction_stage_summary.tsv", sep="\t", index=False)
    pred_particles.to_csv(RESULTS / "CSB_TRO_full_stage_distribution_prediction_particles.tsv", sep="\t", index=False)
    (RESULTS / "CSB_TRO_full_stage_distribution_prediction_summary.json").write_text(
        json.dumps(summary, indent=2, ensure_ascii=False),
        encoding="utf-8",
    )

    note = f"""# CSB-TRO Full-Stage Distribution Prediction

Date: 2026-05-24

## Task

Leave one internal developmental stage out and predict its full empirical distribution in `z = [A, Hm, P, Hr]` from the neighboring observed stage distributions.

## Method

For each held-out stage, an entropic OT/CSB coupling is fitted between the previous and next observed stages. The predicted held-out distribution is sampled from the midpoint geodesic induced by that coupling. This is compared with naive random midpoint interpolation, previous-stage, next-stage, and global-mean baselines by energy distance and RBF MMD.

## Summary

- Held-out internal stages: {summary["n_internal_heldout_stages"]}
- Repeats per stage: {summary["n_repeats_per_stage"]}
- Fraction of stages where CSB median energy beats naive midpoint: {summary["fraction_stage_median_csb_beats_naive_midpoint"]:.3f}
- Fraction of stages where CSB median energy beats previous-stage baseline: {summary["fraction_stage_median_csb_beats_previous"]:.3f}
- Fraction of stages where CSB median energy beats next-stage baseline: {summary["fraction_stage_median_csb_beats_next"]:.3f}
- Fraction of stages where CSB median energy beats global-mean baseline: {summary["fraction_stage_median_csb_beats_global_mean"]:.3f}

## Interpretation

This is a full distribution prediction task, but it is still a leave-one-stage-out interpolation task using both neighboring stages. It should be reported separately from prospective early-to-morula forecasting.
"""
    (RESULTS / "CSB_TRO_full_stage_distribution_prediction_interpretation.md").write_text(note, encoding="utf-8")
    print(json.dumps(summary, indent=2, ensure_ascii=True))


if __name__ == "__main__":
    main()
```
