## Supplemental code for "An entropy-based diagnostic framework for characterizing methylation state dynamics during preimplantation development": csb_tro_graph_laplacian.py

```
from __future__ import annotations

import json
from pathlib import Path

import numpy as np
import pandas as pd


OUT = Path("E:/实验更新_5_22/csb_tro_dynamics")
STATE_COLS = ["A", "Hm", "P", "Hr"]
STAGE_ORDER = ["MII oocyte", "zygote/PN", "2-cell", "4-cell", "8-cell", "morula", "blastocyst"]
LAMBDA_G_GRID = [0.0, 0.01, 0.05, 0.10, 0.25, 0.50, 1.0]


def minmax_standardize(frame: pd.DataFrame, cols: list[str]) -> pd.DataFrame:
    out = frame.copy()
    for col in cols:
        lo = float(out[col].min())
        hi = float(out[col].max())
        out[col] = (out[col] - lo) / max(hi - lo, 1e-12)
    return out


def correlation_graph(particles: pd.DataFrame) -> tuple[pd.DataFrame, pd.DataFrame, pd.DataFrame]:
    z = minmax_standardize(particles[STATE_COLS], STATE_COLS)
    corr = z.corr(method="pearson").fillna(0.0)
    weights = corr.abs()
    np.fill_diagonal(weights.values, 0.0)
    degree = np.diag(weights.sum(axis=1).to_numpy(dtype=float))
    lap = degree - weights.to_numpy(dtype=float)
    lap_df = pd.DataFrame(lap, index=STATE_COLS, columns=STATE_COLS)
    edge_rows = []
    for i, src in enumerate(STATE_COLS):
        for j, dst in enumerate(STATE_COLS):
            if j <= i:
                continue
            edge_rows.append(
                {
                    "source": src,
                    "target": dst,
                    "weight_abs_corr": float(weights.iloc[i, j]),
                    "signed_corr": float(corr.iloc[i, j]),
                }
            )
    return corr, lap_df, pd.DataFrame(edge_rows).sort_values("weight_abs_corr", ascending=False)


def graph_smoothness(lap: pd.DataFrame, stage_summary: pd.DataFrame) -> dict[str, float]:
    mean_cols = [f"{col}_mean" for col in STATE_COLS]
    x = stage_summary.set_index("stage").loc[STAGE_ORDER, mean_cols].T.to_numpy(dtype=float)
    x = (x - x.mean(axis=1, keepdims=True)) / np.maximum(x.std(axis=1, keepdims=True), 1e-12)
    l = lap.loc[STATE_COLS, STATE_COLS].to_numpy(dtype=float)
    c_stage_state = float(np.trace(x.T @ l @ x))

    # Same graph penalty on the learned velocity field, averaged over all source particles.
    vel = pd.read_csv(OUT / "CSB_TRO_path_space_velocity_field.tsv", sep="\t")
    v = vel[["vA", "vHm", "vP", "vHr"]].to_numpy(dtype=float).T
    v = (v - v.mean(axis=1, keepdims=True)) / np.maximum(v.std(axis=1, keepdims=True), 1e-12)
    c_velocity = float(np.trace(v.T @ l @ v) / vel.shape[0])
    return {
        "C_G_stage_state_trajectory": c_stage_state,
        "C_G_velocity_field_mean": c_velocity,
        "C_G_total": c_stage_state + c_velocity,
    }


def main() -> None:
    particles = pd.read_csv(OUT / "CSB_TRO_fused_product_particles.tsv", sep="\t")
    particles = particles[particles["stage"].isin(STAGE_ORDER)].copy()
    stage_summary = pd.read_csv(OUT / "CSB_TRO_path_space_stage_summary.tsv", sep="\t")
    objective = pd.read_csv(OUT / "CSB_TRO_path_space_objective_terms.tsv", sep="\t")

    corr, lap, edges = correlation_graph(particles)
    g_terms = graph_smoothness(lap, stage_summary)

    j_path_without_graph = float(objective["J_transition"].sum())
    rows = []
    for lambda_g in LAMBDA_G_GRID:
        rows.append(
            {
                "lambda_G": lambda_g,
                **g_terms,
                "lambda_G_C_G": lambda_g * g_terms["C_G_total"],
                "J_path_without_graph": j_path_without_graph,
                "J_path_with_graph": j_path_without_graph + lambda_g * g_terms["C_G_total"],
            }
        )
    sensitivity = pd.DataFrame(rows)

    summary = {
        "model": "CSB-TRO graph Laplacian objective audit",
        "date": "2026-05-24",
        "graph_level": "state-variable graph from empirical absolute correlations among A, Hm, P, Hr",
        "important_limitation": "This is not yet a full DMR/gene-module graph Laplacian; it is the strict objective hook using available fused state variables.",
        "state_variables": STATE_COLS,
        "graph_terms": g_terms,
        "top_edges": edges.head(6).to_dict(orient="records"),
        "J_path_without_graph": j_path_without_graph,
        "lambda_G_grid": LAMBDA_G_GRID,
        "recommended_reporting_lambda_G": 0.10,
        "recommended_J_path_with_graph": float(
            sensitivity.loc[sensitivity["lambda_G"].eq(0.10), "J_path_with_graph"].iloc[0]
        ),
    }

    corr.to_csv(OUT / "CSB_TRO_graph_state_correlation.tsv", sep="\t")
    lap.to_csv(OUT / "CSB_TRO_graph_state_laplacian.tsv", sep="\t")
    edges.to_csv(OUT / "CSB_TRO_graph_state_edges.tsv", sep="\t", index=False)
    sensitivity.to_csv(OUT / "CSB_TRO_graph_laplacian_objective_sensitivity.tsv", sep="\t", index=False)
    (OUT / "CSB_TRO_graph_laplacian_summary.json").write_text(json.dumps(summary, indent=2, ensure_ascii=False), encoding="utf-8")

    note = f"""# CSB-TRO graph Laplacian objective audit

Date: 2026-05-24

This step adds the graph-Laplacian objective hook:

`C_G = Tr(X^T L_G X)`

Because the current fused CSB-TRO state has four variables rather than DMR/gene-module nodes, this run uses a state-variable graph whose edge weights are empirical absolute correlations among A, Hm, P, and Hr.

## Limitation

This is not yet the final biological DMR/gene graph Laplacian. It is a mathematically explicit objective audit that can be replaced by a DMR/gene-module graph once node-level features are supplied.

## Main result

- C_G stage-state trajectory: {g_terms["C_G_stage_state_trajectory"]:.6f}
- C_G velocity field mean: {g_terms["C_G_velocity_field_mean"]:.6f}
- C_G total: {g_terms["C_G_total"]:.6f}
- J path without graph: {j_path_without_graph:.6f}
- J path with graph at lambda_G=0.10: {summary["recommended_J_path_with_graph"]:.6f}

## Top graph edges

{edges.head(6).to_string(index=False)}
"""
    (OUT / "CSB_TRO_graph_laplacian_interpretation.md").write_text(note, encoding="utf-8")
    print(json.dumps(summary, indent=2, ensure_ascii=True))


if __name__ == "__main__":
    main()
```
