## Supplemental code for "An entropy-based diagnostic framework for characterizing methylation state dynamics during preimplantation development": csb_tro_limitations_closure.py

```
from __future__ import annotations

import itertools
import json
from pathlib import Path

import numpy as np
import pandas as pd


OUT = Path("E:/实验更新_5_22/csb_tro_dynamics")
STAGE_ORDER = ["MII oocyte", "zygote/PN", "2-cell", "4-cell", "8-cell", "morula", "blastocyst"]
RNA_STAGE = {
    "MII oocyte": "oocyte",
    "zygote/PN": "zygote",
    "2-cell": "2-cell",
    "4-cell": "4-cell",
    "8-cell": "8-cell",
    "morula": "morula",
    "blastocyst": "blastocyst",
}
STATE_COLS = ["A", "Hm", "P", "Hr"]
EPSILON = 0.075
LAMBDA_A = 1.65
LAMBDA_P = 0.95
POTENCY_THRESHOLD_QUANTILE = 0.60


def stage_agnostic_p_min(particles: pd.DataFrame) -> float:
    return float(particles["P"].quantile(POTENCY_THRESHOLD_QUANTILE))


def sinkhorn(a: np.ndarray, b: np.ndarray, cost: np.ndarray, epsilon: float = EPSILON, n_iter: int = 1200) -> np.ndarray:
    k = np.exp(-cost / max(epsilon, 1e-9))
    k = np.maximum(k, 1e-300)
    u = np.ones_like(a)
    v = np.ones_like(b)
    for _ in range(n_iter):
        u_new = a / np.maximum(k @ v, 1e-300)
        v_new = b / np.maximum(k.T @ u_new, 1e-300)
        if np.max(np.abs(u_new - u)) < 1e-10 and np.max(np.abs(v_new - v)) < 1e-10:
            u, v = u_new, v_new
            break
        u, v = u_new, v_new
    pi = (u[:, None] * k) * v[None, :]
    total = pi.sum()
    return pi / total if total > 0 else pi


def cost_terms(x: np.ndarray, y: np.ndarray, p_min: float) -> tuple[np.ndarray, np.ndarray]:
    diff = y[None, :, :] - x[:, None, :]
    movement = np.sum(diff * diff, axis=2)
    age = np.maximum(y[None, :, 0] - x[:, None, 0], 0.0) ** 2
    potency = np.broadcast_to(np.maximum(p_min - y[None, :, 2], 0.0) ** 2, movement.shape)
    return movement + LAMBDA_A * age + LAMBDA_P * potency, diff


def transition_objective(x: np.ndarray, y: np.ndarray, p_min: float) -> dict[str, float]:
    cost, diff = cost_terms(x, y, p_min)
    a = np.full(x.shape[0], 1.0 / x.shape[0])
    b = np.full(y.shape[0], 1.0 / y.shape[0])
    pi = sinkhorn(a, b, cost)
    independent = np.full_like(pi, 1.0 / pi.size)
    mask = pi > 0
    kl = float(np.sum(pi[mask] * np.log(pi[mask] / independent[mask])))
    return {
        "J_transition": float(np.sum(pi * cost) + EPSILON * kl),
        "mean_transport_A": float(np.sum(pi * diff[:, :, 0])),
        "mean_transport_P": float(np.sum(pi * diff[:, :, 2])),
        "mean_squared_displacement": float(np.sum(pi * np.sum(diff * diff, axis=2))),
        "epsilon_KL": EPSILON * kl,
    }


def stage_summary(particles: pd.DataFrame) -> pd.DataFrame:
    sm = particles.groupby("stage", as_index=False).agg(
        n_particles=("particle_id", "count"),
        A_mean=("A", "mean"),
        Hm_mean=("Hm", "mean"),
        P_mean=("P", "mean"),
        Hr_mean=("Hr", "mean"),
    )
    sm["stage_order"] = sm["stage"].map({s: i for i, s in enumerate(STAGE_ORDER)})
    sm["A_rank_lowest_is_1"] = sm["A_mean"].rank(method="min", ascending=True).astype(int)
    sm["P_rank_highest_is_1"] = sm["P_mean"].rank(method="min", ascending=False).astype(int)
    sm["reset_score_A_minus_P"] = sm["A_mean"] - sm["P_mean"]
    sm["reset_rank_lowest_is_1"] = sm["reset_score_A_minus_P"].rank(method="min", ascending=True).astype(int)
    return sm.sort_values("stage_order").reset_index(drop=True)


def build_fused_particles(dna: pd.DataFrame, rna: pd.DataFrame, scheme: str, seed: int, max_particles: int = 420) -> pd.DataFrame:
    rng = np.random.default_rng(seed)
    rows = []
    for stage in STAGE_ORDER:
        d = dna[dna["stage"].eq(stage)].reset_index(drop=True)
        r = rna[rna["stage"].eq(RNA_STAGE[stage])].reset_index(drop=True)
        if len(d) == 0 or len(r) == 0:
            continue
        n = min(max_particles, len(d) * len(r))
        if scheme == "full_or_random_product":
            if len(d) * len(r) <= max_particles:
                pairs = [(i, j) for i in range(len(d)) for j in range(len(r))]
            else:
                pairs = [(int(rng.integers(len(d))), int(rng.integers(len(r)))) for _ in range(n)]
        elif scheme == "rank_matched_lowA_highP":
            di = d.sort_values("A", ascending=True).index.to_numpy()
            rj = r.sort_values("P", ascending=False).index.to_numpy()
            pairs = [(int(di[k % len(di)]), int(rj[k % len(rj)])) for k in range(n)]
        elif scheme == "rank_opposed_lowA_lowP":
            di = d.sort_values("A", ascending=True).index.to_numpy()
            rj = r.sort_values("P", ascending=True).index.to_numpy()
            pairs = [(int(di[k % len(di)]), int(rj[k % len(rj)])) for k in range(n)]
        elif scheme == "bootstrap_independent_product":
            pairs = [(int(rng.integers(len(d))), int(rng.integers(len(r)))) for _ in range(n)]
        else:
            raise ValueError(scheme)
        for k, (i, j) in enumerate(pairs):
            dr = d.iloc[i]
            rr = r.iloc[j]
            rows.append(
                {
                    "particle_id": f"{scheme}:{stage}:{seed}:{k}",
                    "stage": stage,
                    "A": float(dr["A"]),
                    "Hm": float(dr["Hm"]),
                    "P": float(rr["P"]),
                    "Hr": float(rr["Hr"]),
                }
            )
    return pd.DataFrame(rows)


def fusion_sensitivity() -> tuple[pd.DataFrame, dict]:
    dna = pd.read_csv(OUT / "CSB_TRO_state_samples.tsv", sep="\t")
    rna = pd.read_csv(OUT / "CSB_TRO_RNA_GSE36552_cell_states.tsv", sep="\t")
    dna = dna[dna["stage"].isin(STAGE_ORDER)].copy()
    rna = rna[rna["stage"].isin(set(RNA_STAGE.values()))].copy()
    schemes = {
        "full_or_random_product": range(20260524, 20260554),
        "bootstrap_independent_product": range(20260600, 20260630),
        "rank_matched_lowA_highP": [20260701],
        "rank_opposed_lowA_lowP": [20260702],
    }
    rows = []
    for scheme, seeds in schemes.items():
        for seed in seeds:
            fused = build_fused_particles(dna, rna, scheme, seed)
            sm = stage_summary(fused)
            p_min = stage_agnostic_p_min(fused)
            x = fused[fused["stage"].eq("8-cell")][STATE_COLS].to_numpy(dtype=float)
            y = fused[fused["stage"].eq("morula")][STATE_COLS].to_numpy(dtype=float)
            trans = transition_objective(x, y, p_min)
            morula = sm[sm["stage"].eq("morula")].iloc[0]
            rows.append(
                {
                    "scheme": scheme,
                    "seed": seed,
                    "n_particles": int(len(fused)),
                    "morula_A_mean": float(morula["A_mean"]),
                    "morula_P_mean": float(morula["P_mean"]),
                    "morula_A_rank_lowest_is_1": int(morula["A_rank_lowest_is_1"]),
                    "morula_P_rank_highest_is_1": int(morula["P_rank_highest_is_1"]),
                    "morula_reset_rank_lowest_is_1": int(morula["reset_rank_lowest_is_1"]),
                    "transport_8cell_to_morula_A": trans["mean_transport_A"],
                    "transport_8cell_to_morula_P": trans["mean_transport_P"],
                    "J_8cell_to_morula": trans["J_transition"],
                }
            )
    df = pd.DataFrame(rows)
    summary = {
        "n_runs": int(len(df)),
        "schemes": sorted(df["scheme"].unique().tolist()),
        "fraction_morula_A_rank1": float(np.mean(df["morula_A_rank_lowest_is_1"].eq(1))),
        "fraction_morula_P_rank_top2": float(np.mean(df["morula_P_rank_highest_is_1"].le(2))),
        "fraction_morula_reset_rank1": float(np.mean(df["morula_reset_rank_lowest_is_1"].eq(1))),
        "fraction_8cell_to_morula_A_negative": float(np.mean(df["transport_8cell_to_morula_A"] < 0)),
        "by_scheme": df.groupby("scheme").agg(
            n=("seed", "count"),
            fraction_A_rank1=("morula_A_rank_lowest_is_1", lambda s: float(np.mean(s.eq(1)))),
            fraction_P_rank_top2=("morula_P_rank_highest_is_1", lambda s: float(np.mean(s.le(2)))),
            fraction_A_transport_negative=("transport_8cell_to_morula_A", lambda s: float(np.mean(s < 0))),
            A_transport_median=("transport_8cell_to_morula_A", "median"),
        ).reset_index().to_dict(orient="records"),
    }
    return df, summary


def order_path_objective_test(particles: pd.DataFrame) -> tuple[pd.DataFrame, pd.DataFrame, dict]:
    sm = stage_summary(particles)
    p_min = stage_agnostic_p_min(particles)
    states = {s: particles[particles["stage"].eq(s)][STATE_COLS].to_numpy(dtype=float) for s in STAGE_ORDER}
    pair_rows = []
    pair_obj = {}
    for s0 in STAGE_ORDER:
        for s1 in STAGE_ORDER:
            if s0 == s1:
                continue
            obj = transition_objective(states[s0], states[s1], p_min)
            pair_obj[(s0, s1)] = obj
            pair_rows.append({"from_stage": s0, "to_stage": s1, **obj})
    pair_df = pd.DataFrame(pair_rows)

    order_rows = []
    for perm in itertools.permutations(STAGE_ORDER):
        j = 0.0
        enters_morula_from = None
        enter_morula_A = np.nan
        for a, b in zip(perm[:-1], perm[1:]):
            item = pair_obj[(a, b)]
            j += item["J_transition"]
            if b == "morula":
                enters_morula_from = a
                enter_morula_A = item["mean_transport_A"]
        order_rows.append(
            {
                "stage_order": " -> ".join(perm),
                "J_path_order": j,
                "enters_morula_from": enters_morula_from,
                "enter_morula_A": enter_morula_A,
                "is_canonical": list(perm) == STAGE_ORDER,
            }
        )
    order_df = pd.DataFrame(order_rows).sort_values("J_path_order").reset_index(drop=True)
    canonical = order_df[order_df["is_canonical"]].iloc[0]
    summary = {
        "n_orders": int(len(order_df)),
        "canonical_J_path_order": float(canonical["J_path_order"]),
        "canonical_rank_lowest_J_is_1": int(canonical.name + 1),
        "canonical_percentile_low_J": float((canonical.name + 1) / len(order_df)),
        "p_random_order_J_less_or_equal_canonical": float(np.mean(order_df["J_path_order"] <= canonical["J_path_order"])),
        "best_order": str(order_df.iloc[0]["stage_order"]),
        "best_order_J": float(order_df.iloc[0]["J_path_order"]),
        "median_order_J": float(order_df["J_path_order"].median()),
        "orders_entering_morula_from_8cell": int(order_df["enters_morula_from"].eq("8-cell").sum()),
        "median_J_when_entering_morula_from_8cell": float(order_df.loc[order_df["enters_morula_from"].eq("8-cell"), "J_path_order"].median()),
    }
    return pair_df, order_df, summary


def global_multimarginal_bridge(particles: pd.DataFrame) -> tuple[pd.DataFrame, pd.DataFrame, dict]:
    sm = stage_summary(particles)
    p_min = stage_agnostic_p_min(particles)
    stage_frames = [particles[particles["stage"].eq(s)].reset_index(drop=True) for s in STAGE_ORDER]
    arrays = [df[STATE_COLS].to_numpy(dtype=float) for df in stage_frames]
    kernels = []
    diffs = []
    for x, y in zip(arrays[:-1], arrays[1:]):
        c, d = cost_terms(x, y, p_min)
        kernels.append(np.maximum(np.exp(-c / EPSILON), 1e-300))
        diffs.append(d)

    n_time = len(STAGE_ORDER)
    targets = [np.full(len(df), 1.0 / len(df)) for df in stage_frames]
    scales = [np.ones(len(df)) for df in stage_frames]
    def forward_backward(current_scales: list[np.ndarray]) -> tuple[list[np.ndarray], list[np.ndarray], float, list[np.ndarray]]:
        alpha = [None] * n_time
        beta = [None] * n_time
        alpha[0] = current_scales[0].copy()
        for t in range(1, n_time):
            alpha[t] = current_scales[t] * (alpha[t - 1] @ kernels[t - 1])
        beta[-1] = np.ones_like(scales[-1])
        for t in range(n_time - 2, -1, -1):
            beta[t] = kernels[t] @ (current_scales[t + 1] * beta[t + 1])
        z = float(np.sum(alpha[-1]))
        marginals = [alpha[t] * beta[t] / max(z, 1e-300) for t in range(n_time)]
        return alpha, beta, z, marginals

    errors = []
    for it in range(250):
        # Coordinate IPF is slower than simultaneous updates but is much more stable here.
        for t in range(n_time):
            _, _, _, marginals = forward_backward(scales)
            scales[t] *= targets[t] / np.maximum(marginals[t], 1e-300)
            # This centering does not change the normalized path law, because every path
            # contains exactly one node from each time slice.
            scales[t] /= max(float(np.exp(np.mean(np.log(np.maximum(scales[t], 1e-300))))), 1e-300)
        _, _, _, marginals = forward_backward(scales)
        max_err = 0.0
        for t in range(n_time):
            max_err = max(max_err, float(np.max(np.abs(marginals[t] - targets[t]))))
        errors.append(max_err)
        if max_err < 1e-10:
            break

    # Recompute messages after final update.
    alpha, beta, z, _ = forward_backward(scales)

    transition_rows = []
    velocity_rows = []
    max_pair_mass_error = 0.0
    for t, (s0, s1) in enumerate(zip(STAGE_ORDER[:-1], STAGE_ORDER[1:])):
        pair = alpha[t][:, None] * kernels[t] * (scales[t + 1] * beta[t + 1])[None, :] / max(z, 1e-300)
        pair /= pair.sum()
        max_pair_mass_error = max(max_pair_mass_error, abs(float(pair.sum()) - 1.0))
        d = diffs[t]
        transition_rows.append(
            {
                "transition_order": t,
                "from_stage": s0,
                "to_stage": s1,
                "mean_transport_A": float(np.sum(pair * d[:, :, 0])),
                "mean_transport_Hm": float(np.sum(pair * d[:, :, 1])),
                "mean_transport_P": float(np.sum(pair * d[:, :, 2])),
                "mean_transport_Hr": float(np.sum(pair * d[:, :, 3])),
                "mean_squared_displacement": float(np.sum(pair * np.sum(d * d, axis=2))),
            }
        )
        y_cond = (pair @ arrays[t + 1]) / np.maximum(pair.sum(axis=1, keepdims=True), 1e-300)
        vel = y_cond - arrays[t]
        vdf = stage_frames[t][["particle_id", "stage", "A", "Hm", "P", "Hr"]].copy()
        vdf["to_stage"] = s1
        vdf["vA"] = vel[:, 0]
        vdf["vHm"] = vel[:, 1]
        vdf["vP"] = vel[:, 2]
        vdf["vHr"] = vel[:, 3]
        velocity_rows.append(vdf)

    trans_df = pd.DataFrame(transition_rows)
    vel_df = pd.concat(velocity_rows, ignore_index=True)
    summary = {
        "solver": "global multi-marginal Markov Schrödinger bridge by iterative proportional fitting",
        "n_iterations": int(len(errors)),
        "final_max_marginal_error": float(errors[-1]),
        "max_pair_mass_error": float(max_pair_mass_error),
        "epsilon": EPSILON,
        "lambda_A": LAMBDA_A,
        "lambda_P": LAMBDA_P,
        "key_transition_8cell_to_morula": trans_df[trans_df["from_stage"].eq("8-cell")].iloc[0].to_dict(),
        "key_transition_morula_to_blastocyst": trans_df[trans_df["from_stage"].eq("morula")].iloc[0].to_dict(),
    }
    return trans_df, vel_df, summary


def main() -> None:
    particles = pd.read_csv(OUT / "CSB_TRO_fused_product_particles.tsv", sep="\t")
    particles = particles[particles["stage"].isin(STAGE_ORDER)].copy()

    fusion_df, fusion_summary = fusion_sensitivity()
    pair_df, order_df, order_summary = order_path_objective_test(particles)
    global_trans, global_vel, global_summary = global_multimarginal_bridge(particles)

    summary = {
        "model": "CSB-TRO limitation closure experiments",
        "date": "2026-05-24",
        "unpaired_DNA_RNA_closure": fusion_summary,
        "stage_order_closure": order_summary,
        "global_multimarginal_bridge": global_summary,
        "interpretation": (
            "Fusion sensitivity addresses the unpaired DNA/RNA limitation by showing whether the morula basin survives "
            "independent product, bootstrap, and adversarial rank-pairing schemes. The all-order path objective addresses "
            "the random-order concern by comparing the full biological path objective against all 7! stage permutations. "
            "The multi-marginal IPF solver upgrades the bridge from a sequence of independently reported pairwise couplings "
            "to a globally scaled Markov path measure satisfying all empirical stage marginals."
        ),
        "p_min_definition": f"stage-agnostic global fused-particle P quantile q={POTENCY_THRESHOLD_QUANTILE}; not derived from morula",
    }

    fusion_df.to_csv(OUT / "CSB_TRO_limitfix_fusion_sensitivity.tsv", sep="\t", index=False)
    pair_df.to_csv(OUT / "CSB_TRO_limitfix_all_directed_transition_objectives.tsv", sep="\t", index=False)
    order_df.to_csv(OUT / "CSB_TRO_limitfix_all_stage_order_objectives.tsv", sep="\t", index=False)
    global_trans.to_csv(OUT / "CSB_TRO_global_multimarginal_transition_summary.tsv", sep="\t", index=False)
    global_vel.to_csv(OUT / "CSB_TRO_global_multimarginal_velocity_field.tsv", sep="\t", index=False)
    (OUT / "CSB_TRO_limitations_closure_summary.json").write_text(json.dumps(summary, indent=2, ensure_ascii=False), encoding="utf-8")

    note = f"""# CSB-TRO limitation closure experiments

Date: 2026-05-24

## Issue 1: DNA/RNA are not paired single cells

Resolution: run fusion sensitivity under independent product, bootstrap product, rank-matched low-A/high-P, and rank-opposed low-A/low-P schemes.

- Runs: {fusion_summary["n_runs"]}
- Fraction morula A rank 1: {fusion_summary["fraction_morula_A_rank1"]:.3f}
- Fraction morula P rank top 2: {fusion_summary["fraction_morula_P_rank_top2"]:.3f}
- Fraction morula reset rank 1: {fusion_summary["fraction_morula_reset_rank1"]:.3f}
- Fraction 8-cell -> morula A transport negative: {fusion_summary["fraction_8cell_to_morula_A_negative"]:.3f}

## Issue 2: random stage order did not prove 8-cell -> morula uniqueness

Resolution: compare the full path objective of the biological stage order against all 7! stage permutations.

- Number of orders: {order_summary["n_orders"]}
- Canonical path objective rank, rank 1 = lowest: {order_summary["canonical_rank_lowest_J_is_1"]}
- Canonical low-objective percentile: {order_summary["canonical_percentile_low_J"]:.4f}
- p(random order J <= canonical J): {order_summary["p_random_order_J_less_or_equal_canonical"]:.6f}
- Best order: {order_summary["best_order"]}

## Issue 3: pairwise bridge vs global path-space bridge

Resolution: solve a global multi-marginal Markov Schrödinger bridge using iterative proportional fitting over all observed stage marginals.

- IPF iterations: {global_summary["n_iterations"]}
- Final max marginal error: {global_summary["final_max_marginal_error"]:.3e}
- 8-cell -> morula mean transport A: {global_summary["key_transition_8cell_to_morula"]["mean_transport_A"]:.6f}
- 8-cell -> morula mean transport P: {global_summary["key_transition_8cell_to_morula"]["mean_transport_P"]:.6f}

## Interpretation

These experiments do not create true paired DNA/RNA cells, but they show whether the CSB-TRO conclusion is robust to plausible and adversarial fusion assumptions. The stage-order issue is addressed at the whole-path objective level rather than by asking whether any forced predecessor can decrease A into morula. The global multi-marginal solver is the strictest discrete Markov path-space bridge implemented so far.
"""
    (OUT / "CSB_TRO_limitations_closure_interpretation.md").write_text(note, encoding="utf-8")
    print(json.dumps(summary, indent=2, ensure_ascii=True))


if __name__ == "__main__":
    main()
```
