## Supplemental code for "An entropy-based diagnostic framework for characterizing methylation state dynamics during preimplantation development": csb_tro_path_space.py

```
from __future__ import annotations

import json
from pathlib import Path

import numpy as np
import pandas as pd


OUT = Path("E:/实验更新_5_22/csb_tro_dynamics")
STAGE_ORDER = ["MII oocyte", "zygote/PN", "2-cell", "4-cell", "8-cell", "morula", "blastocyst"]
STATE_COLS = ["A", "Hm", "P", "Hr"]

LAMBDA_A = 1.65
LAMBDA_P = 0.95
EPSILON = 0.075
SINKHORN_ITER = 3500
POTENCY_THRESHOLD_QUANTILE = 0.60


def stage_agnostic_p_min(particles: pd.DataFrame) -> float:
    """Prevents circularity: P_min is not derived from morula."""
    return float(particles["P"].quantile(POTENCY_THRESHOLD_QUANTILE))


def sinkhorn(a: np.ndarray, b: np.ndarray, kernel: np.ndarray, n_iter: int = SINKHORN_ITER) -> np.ndarray:
    k = np.maximum(kernel, 1e-300)
    u = np.ones_like(a)
    v = np.ones_like(b)
    for _ in range(n_iter):
        u_new = a / np.maximum(k @ v, 1e-300)
        v_new = b / np.maximum(k.T @ u_new, 1e-300)
        if np.max(np.abs(u_new - u)) < 1e-11 and np.max(np.abs(v_new - v)) < 1e-11:
            u, v = u_new, v_new
            break
        u, v = u_new, v_new
    pi = (u[:, None] * k) * v[None, :]
    total = pi.sum()
    return pi / total if total > 0 else pi


def transition_cost_terms(x: np.ndarray, y: np.ndarray, p_min: float) -> dict[str, np.ndarray]:
    diff = y[None, :, :] - x[:, None, :]
    squared_displacement = np.sum(diff * diff, axis=2)
    age_increase = np.maximum(y[None, :, 0] - x[:, None, 0], 0.0) ** 2
    potency_below = np.broadcast_to(np.maximum(p_min - y[None, :, 2], 0.0) ** 2, squared_displacement.shape)
    constrained_cost = squared_displacement + LAMBDA_A * age_increase + LAMBDA_P * potency_below
    return {
        "diff": diff,
        "squared_displacement": squared_displacement,
        "age_increase_penalty": age_increase,
        "potency_below_threshold_penalty": potency_below,
        "constrained_cost": constrained_cost,
    }


def coupling_objective(pi: np.ndarray, terms: dict[str, np.ndarray], epsilon: float) -> dict[str, float]:
    n_from, n_to = pi.shape
    independent = np.full_like(pi, 1.0 / (n_from * n_to), dtype=float)
    mask = pi > 0
    kl_to_independent = float(np.sum(pi[mask] * np.log(pi[mask] / independent[mask])))
    movement_cost = float(np.sum(pi * terms["squared_displacement"]))
    c_age = float(np.sum(pi * terms["age_increase_penalty"]))
    c_potency = float(np.sum(pi * terms["potency_below_threshold_penalty"]))
    constrained_transport_cost = movement_cost + LAMBDA_A * c_age + LAMBDA_P * c_potency
    entropy_regularized_objective = constrained_transport_cost + epsilon * kl_to_independent
    return {
        "movement_cost": movement_cost,
        "C_A_age_increase": c_age,
        "lambda_A_C_A": LAMBDA_A * c_age,
        "C_P_potency_below": c_potency,
        "lambda_P_C_P": LAMBDA_P * c_potency,
        "KL_pi_to_independent_reference": kl_to_independent,
        "epsilon_KL": epsilon * kl_to_independent,
        "constrained_transport_cost": constrained_transport_cost,
        "J_transition": entropy_regularized_objective,
    }


def build_transition(
    particles: pd.DataFrame,
    from_stage: str,
    to_stage: str,
    p_min: float,
) -> tuple[pd.DataFrame, pd.DataFrame, pd.DataFrame]:
    xdf = particles[particles["stage"].eq(from_stage)].reset_index(drop=True).copy()
    ydf = particles[particles["stage"].eq(to_stage)].reset_index(drop=True).copy()
    x = xdf[STATE_COLS].to_numpy(dtype=float)
    y = ydf[STATE_COLS].to_numpy(dtype=float)
    a = np.full(len(xdf), 1.0 / len(xdf))
    b = np.full(len(ydf), 1.0 / len(ydf))
    terms = transition_cost_terms(x, y, p_min=p_min)
    kernel = np.exp(-terms["constrained_cost"] / EPSILON)
    pi = sinkhorn(a, b, kernel)

    diff = terms["diff"]
    y_cond = (pi @ y) / np.maximum(pi.sum(axis=1, keepdims=True), 1e-12)
    velocity = y_cond - x
    velocity_df = xdf[["particle_id", "stage", "A", "Hm", "P", "Hr"]].copy()
    velocity_df["to_stage"] = to_stage
    velocity_df["vA"] = velocity[:, 0]
    velocity_df["vHm"] = velocity[:, 1]
    velocity_df["vP"] = velocity[:, 2]
    velocity_df["vHr"] = velocity[:, 3]

    obj = coupling_objective(pi, terms, epsilon=EPSILON)
    obj.update(
        {
            "from_stage": from_stage,
            "to_stage": to_stage,
            "n_from_particles": int(len(xdf)),
            "n_to_particles": int(len(ydf)),
            "mean_transport_A": float(np.sum(pi * diff[:, :, 0])),
            "mean_transport_Hm": float(np.sum(pi * diff[:, :, 1])),
            "mean_transport_P": float(np.sum(pi * diff[:, :, 2])),
            "mean_transport_Hr": float(np.sum(pi * diff[:, :, 3])),
            "row_marginal_max_abs_error": float(np.max(np.abs(pi.sum(axis=1) - a))),
            "col_marginal_max_abs_error": float(np.max(np.abs(pi.sum(axis=0) - b))),
        }
    )
    objective_df = pd.DataFrame([obj])

    from_i, to_j = np.nonzero(pi > 1e-14)
    coupling_df = pd.DataFrame(
        {
            "from_stage": from_stage,
            "to_stage": to_stage,
            "from_particle_id": xdf.loc[from_i, "particle_id"].to_numpy(),
            "to_particle_id": ydf.loc[to_j, "particle_id"].to_numpy(),
            "probability": pi[from_i, to_j],
            "delta_A": diff[from_i, to_j, 0],
            "delta_Hm": diff[from_i, to_j, 1],
            "delta_P": diff[from_i, to_j, 2],
            "delta_Hr": diff[from_i, to_j, 3],
            "squared_displacement": terms["squared_displacement"][from_i, to_j],
            "age_increase_penalty": terms["age_increase_penalty"][from_i, to_j],
            "potency_below_threshold_penalty": terms["potency_below_threshold_penalty"][from_i, to_j],
            "constrained_cost": terms["constrained_cost"][from_i, to_j],
        }
    )
    return coupling_df, objective_df, velocity_df


def stage_summary(particles: pd.DataFrame) -> pd.DataFrame:
    summary = particles.groupby("stage", as_index=False).agg(
        n_particles=("particle_id", "count"),
        A_mean=("A", "mean"),
        Hm_mean=("Hm", "mean"),
        P_mean=("P", "mean"),
        Hr_mean=("Hr", "mean"),
    )
    summary["stage_order"] = summary["stage"].map({s: i for i, s in enumerate(STAGE_ORDER)})
    summary["A_rank_lowest_is_1"] = summary["A_mean"].rank(method="min", ascending=True).astype(int)
    summary["P_rank_highest_is_1"] = summary["P_mean"].rank(method="min", ascending=False).astype(int)
    summary["reset_score_A_minus_P"] = summary["A_mean"] - summary["P_mean"]
    summary["reset_rank_lowest_is_1"] = summary["reset_score_A_minus_P"].rank(method="min", ascending=True).astype(int)
    return summary.sort_values("stage_order").reset_index(drop=True)


def main() -> None:
    particles = pd.read_csv(OUT / "CSB_TRO_fused_product_particles.tsv", sep="\t")
    particles = particles[particles["stage"].isin(STAGE_ORDER)].copy()
    particles["stage_order"] = particles["stage"].map({s: i for i, s in enumerate(STAGE_ORDER)})
    particles = particles.sort_values(["stage_order", "particle_id"]).reset_index(drop=True)

    summary_df = stage_summary(particles)
    p_min = stage_agnostic_p_min(particles)

    couplings = []
    objectives = []
    velocities = []
    for from_stage, to_stage in zip(STAGE_ORDER[:-1], STAGE_ORDER[1:]):
        coupling_df, objective_df, velocity_df = build_transition(particles, from_stage, to_stage, p_min=p_min)
        coupling_df["transition_order"] = STAGE_ORDER.index(from_stage)
        objective_df["transition_order"] = STAGE_ORDER.index(from_stage)
        velocity_df["transition_order"] = STAGE_ORDER.index(from_stage)
        couplings.append(coupling_df)
        objectives.append(objective_df)
        velocities.append(velocity_df)

    coupling_all = pd.concat(couplings, ignore_index=True)
    objective_all = pd.concat(objectives, ignore_index=True).sort_values("transition_order").reset_index(drop=True)
    velocity_all = pd.concat(velocities, ignore_index=True)

    total = {
        "model": "CSB-TRO Markov path-space bridge",
        "date": "2026-05-24",
        "state_vector": STATE_COLS,
        "stage_order": STAGE_ORDER,
        "path_space_form": "P*(z0,...,zK) = p0(z0) prod_k P*(z_{k+1}|z_k), with transition kernels induced by constrained entropic couplings.",
        "implementation_level": "discrete empirical Markov path-space Schrödinger bridge approximation",
        "epsilon": EPSILON,
        "lambda_A": LAMBDA_A,
        "lambda_P": LAMBDA_P,
        "p_min": p_min,
        "p_min_definition": f"stage-agnostic global fused-particle P quantile q={POTENCY_THRESHOLD_QUANTILE}; not derived from morula",
        "n_particles": int(len(particles)),
        "n_transitions": int(objective_all.shape[0]),
        "J_path_total": float(objective_all["J_transition"].sum()),
        "movement_cost_total": float(objective_all["movement_cost"].sum()),
        "C_A_total": float(objective_all["C_A_age_increase"].sum()),
        "lambda_A_C_A_total": float(objective_all["lambda_A_C_A"].sum()),
        "C_P_total": float(objective_all["C_P_potency_below"].sum()),
        "lambda_P_C_P_total": float(objective_all["lambda_P_C_P"].sum()),
        "KL_to_independent_reference_total": float(objective_all["KL_pi_to_independent_reference"].sum()),
        "epsilon_KL_total": float(objective_all["epsilon_KL"].sum()),
        "max_row_marginal_error": float(objective_all["row_marginal_max_abs_error"].max()),
        "max_col_marginal_error": float(objective_all["col_marginal_max_abs_error"].max()),
    }
    morula = summary_df[summary_df["stage"].eq("morula")].iloc[0]
    total.update(
        {
            "morula_A_rank_lowest_is_1": int(morula["A_rank_lowest_is_1"]),
            "morula_P_rank_highest_is_1": int(morula["P_rank_highest_is_1"]),
            "morula_reset_rank_lowest_is_1": int(morula["reset_rank_lowest_is_1"]),
            "morula_A_mean": float(morula["A_mean"]),
            "morula_P_mean": float(morula["P_mean"]),
        }
    )

    coupling_all.to_csv(OUT / "CSB_TRO_path_space_transition_couplings.tsv", sep="\t", index=False)
    objective_all.to_csv(OUT / "CSB_TRO_path_space_objective_terms.tsv", sep="\t", index=False)
    velocity_all.to_csv(OUT / "CSB_TRO_path_space_velocity_field.tsv", sep="\t", index=False)
    summary_df.to_csv(OUT / "CSB_TRO_path_space_stage_summary.tsv", sep="\t", index=False)
    (OUT / "CSB_TRO_path_space_summary.json").write_text(json.dumps(total, indent=2, ensure_ascii=False), encoding="utf-8")

    note = f"""# CSB-TRO Markov path-space bridge

Date: 2026-05-24

This step upgrades the fused pairwise bridge into an explicit Markov path-space bridge over developmental stage distributions.

## Path-space form

`P*(z0,...,zK) = p0(z0) prod_k P*(z_{{k+1}} | z_k)`

Each transition kernel is induced by a constrained entropic coupling between adjacent empirical stage distributions.

## Objective

For each transition, the implemented discrete objective is:

`J_k = E_pi[||y-x||^2] + lambda_A C_A + lambda_P C_P + epsilon KL(pi || p_k otimes p_{{k+1}})`

The path objective is the Markov sum:

`J_path = sum_k J_k`

## Main totals

- Total path objective J: {total["J_path_total"]:.6f}
- Movement cost total: {total["movement_cost_total"]:.6f}
- lambda_A C_A total: {total["lambda_A_C_A_total"]:.6f}
- lambda_P C_P total: {total["lambda_P_C_P_total"]:.6f}
- epsilon KL total: {total["epsilon_KL_total"]:.6f}
- Max row marginal error: {total["max_row_marginal_error"]:.3e}
- Max column marginal error: {total["max_col_marginal_error"]:.3e}

## Biological readout

- Morula A rank, rank 1 = lowest: {total["morula_A_rank_lowest_is_1"]}
- Morula P rank, rank 1 = highest: {total["morula_P_rank_highest_is_1"]}
- Morula reset score A-P rank, rank 1 = lowest: {total["morula_reset_rank_lowest_is_1"]}

## Interpretation

This is still a discrete empirical Markov approximation, but it now has an explicit path-space distribution and an auditable objective decomposition. The next strict-math upgrades are graph Laplacian regularization and a continuous drift/Fokker-Planck export.
"""
    (OUT / "CSB_TRO_path_space_interpretation.md").write_text(note, encoding="utf-8")
    print(json.dumps(total, indent=2, ensure_ascii=True))


if __name__ == "__main__":
    main()
```
