## Supplemental code for "An entropy-based diagnostic framework for characterizing methylation state dynamics during preimplantation development": csb_tro_prediction_validation.py

```
from __future__ import annotations

import json
from pathlib import Path

import numpy as np
import pandas as pd


ROOT = Path(__file__).resolve().parents[1]
RESULTS = ROOT / "results"
INPUT = ROOT / "input_tables"
STAGE_ORDER = ["MII oocyte", "zygote/PN", "2-cell", "4-cell", "8-cell", "morula", "blastocyst"]
STATE_COLS = ["A", "Hm", "P", "Hr"]
DMR_STAGES = ["MII oocyte", "zygote/PN", "2-cell", "4-cell", "8-cell", "morula", "blastocyst"]


def energy_distance(x: np.ndarray, y: np.ndarray, rng: np.random.Generator, max_n: int = 240) -> float:
    if len(x) > max_n:
        x = x[rng.choice(len(x), size=max_n, replace=False)]
    if len(y) > max_n:
        y = y[rng.choice(len(y), size=max_n, replace=False)]
    xy = np.linalg.norm(x[:, None, :] - y[None, :, :], axis=2).mean()
    xx = np.linalg.norm(x[:, None, :] - x[None, :, :], axis=2).mean()
    yy = np.linalg.norm(y[:, None, :] - y[None, :, :], axis=2).mean()
    return float(2 * xy - xx - yy)


def stage_distribution_prediction(particles: pd.DataFrame) -> tuple[pd.DataFrame, dict]:
    rng = np.random.default_rng(20260524)
    rows = []
    by_stage = {s: particles[particles["stage"].eq(s)][STATE_COLS].to_numpy(dtype=float) for s in STAGE_ORDER}
    for k in range(1, len(STAGE_ORDER) - 1):
        prev_stage = STAGE_ORDER[k - 1]
        target_stage = STAGE_ORDER[k]
        next_stage = STAGE_ORDER[k + 1]
        prev_x = by_stage[prev_stage]
        next_x = by_stage[next_stage]
        obs = by_stage[target_stage]
        n = len(obs)
        prev_sample = prev_x[rng.integers(0, len(prev_x), size=n)]
        next_sample = next_x[rng.integers(0, len(next_x), size=n)]
        pred_interp = 0.5 * prev_sample + 0.5 * next_sample
        pred_prev = prev_x[rng.integers(0, len(prev_x), size=n)]
        pred_global_mean = np.tile(particles[STATE_COLS].mean().to_numpy(dtype=float), (n, 1))
        rows.append(
            {
                "heldout_stage": target_stage,
                "prev_stage": prev_stage,
                "next_stage": next_stage,
                "energy_dynamic_neighbor_interpolation": energy_distance(obs, pred_interp, rng),
                "energy_static_previous_stage": energy_distance(obs, pred_prev, rng),
                "energy_static_global_mean": energy_distance(obs, pred_global_mean, rng),
            }
        )
    df = pd.DataFrame(rows)
    df["dynamic_beats_previous"] = df["energy_dynamic_neighbor_interpolation"] < df["energy_static_previous_stage"]
    df["dynamic_beats_global_mean"] = df["energy_dynamic_neighbor_interpolation"] < df["energy_static_global_mean"]
    summary = {
        "n_leave_one_internal_stage_tests": int(len(df)),
        "fraction_dynamic_beats_previous_stage": float(df["dynamic_beats_previous"].mean()),
        "fraction_dynamic_beats_global_mean": float(df["dynamic_beats_global_mean"].mean()),
        "median_energy_dynamic": float(df["energy_dynamic_neighbor_interpolation"].median()),
        "median_energy_previous": float(df["energy_static_previous_stage"].median()),
        "median_energy_global_mean": float(df["energy_static_global_mean"].median()),
    }
    return df, summary


def early_to_morula_forecast(stage_summary: pd.DataFrame) -> tuple[pd.DataFrame, dict]:
    early = ["MII oocyte", "zygote/PN", "2-cell", "4-cell", "8-cell"]
    target = "morula"
    ordered = stage_summary.set_index("stage").loc[STAGE_ORDER]
    t = np.arange(len(STAGE_ORDER), dtype=float)
    early_idx = np.array([STAGE_ORDER.index(s) for s in early], dtype=float)
    target_idx = float(STAGE_ORDER.index(target))
    rows = []
    pred_linear = {}
    pred_quad = {}
    for col in ["A_mean", "Hm_mean", "P_mean", "Hr_mean"]:
        y = ordered.loc[early, col].to_numpy(dtype=float)
        beta_linear = np.polyfit(early_idx, y, deg=1)
        beta_quad = np.polyfit(early_idx, y, deg=2)
        pred_linear[col] = float(np.polyval(beta_linear, target_idx))
        pred_quad[col] = float(np.polyval(beta_quad, target_idx))
    observed = ordered.loc[target, ["A_mean", "Hm_mean", "P_mean", "Hr_mean"]].to_dict()
    previous = ordered.loc["8-cell", ["A_mean", "Hm_mean", "P_mean", "Hr_mean"]].to_dict()

    def vec(d: dict) -> np.ndarray:
        return np.array([d["A_mean"], d["Hm_mean"], d["P_mean"], d["Hr_mean"]], dtype=float)

    obs_v = vec(observed)
    linear_v = vec(pred_linear)
    quad_v = vec(pred_quad)
    prev_v = vec(previous)
    rows.extend(
        [
            {
                "forecast_model": "linear_trend_from_pre_morula",
                **{k.replace("_mean", "_pred"): v for k, v in pred_linear.items()},
                "euclidean_error_to_observed_morula": float(np.linalg.norm(linear_v - obs_v)),
            },
            {
                "forecast_model": "quadratic_trend_from_pre_morula",
                **{k.replace("_mean", "_pred"): v for k, v in pred_quad.items()},
                "euclidean_error_to_observed_morula": float(np.linalg.norm(quad_v - obs_v)),
            },
            {
                "forecast_model": "static_previous_8cell_baseline",
                **{k.replace("_mean", "_pred"): v for k, v in previous.items()},
                "euclidean_error_to_observed_morula": float(np.linalg.norm(prev_v - obs_v)),
            },
        ]
    )
    df = pd.DataFrame(rows)
    best = df.sort_values("euclidean_error_to_observed_morula").iloc[0]
    summary = {
        "target": target,
        "best_forecast_model": str(best["forecast_model"]),
        "best_error": float(best["euclidean_error_to_observed_morula"]),
        "linear_error": float(df.loc[df["forecast_model"].eq("linear_trend_from_pre_morula"), "euclidean_error_to_observed_morula"].iloc[0]),
        "quadratic_error": float(df.loc[df["forecast_model"].eq("quadratic_trend_from_pre_morula"), "euclidean_error_to_observed_morula"].iloc[0]),
        "previous_8cell_error": float(df.loc[df["forecast_model"].eq("static_previous_8cell_baseline"), "euclidean_error_to_observed_morula"].iloc[0]),
    }
    return df, summary


def dmr_split_validation(n_iter: int = 500) -> tuple[pd.DataFrame, dict]:
    rng = np.random.default_rng(20260524)
    dmr = pd.read_csv(INPUT / "TRO_interpretability_DMR_contribution_ranking.tsv", sep="\t")
    entropy_cols = [f"entropy_{s}" for s in DMR_STAGES]
    for col in ["age_weight_per_year", *entropy_cols]:
        dmr[col] = pd.to_numeric(dmr[col], errors="coerce")
        dmr[col] = dmr[col].fillna(float(dmr[col].median()))
    weights = dmr["age_weight_per_year"].abs().to_numpy(dtype=float)
    entropy = dmr[entropy_cols].to_numpy(dtype=float)

    def stage_scores(indices: np.ndarray) -> pd.Series:
        w = weights[indices]
        x = entropy[indices]
        score = np.average(x, axis=0, weights=np.maximum(w, 1e-12))
        return pd.Series(score, index=DMR_STAGES)

    rows = []
    n = len(dmr)
    for i in range(n_iter):
        perm = rng.permutation(n)
        train = perm[: n // 2]
        test = perm[n // 2 :]
        train_scores = stage_scores(train)
        test_scores = stage_scores(test)
        train_min = str(train_scores.idxmin())
        test_min = str(test_scores.idxmin())
        train_reset_rank = int(train_scores.rank(method="min", ascending=True)["morula"])
        test_reset_rank = int(test_scores.rank(method="min", ascending=True)["morula"])
        rows.append(
            {
                "iteration": i,
                "train_min_stage": train_min,
                "test_min_stage": test_min,
                "train_morula_rank_lowest_is_1": train_reset_rank,
                "test_morula_rank_lowest_is_1": test_reset_rank,
                "test_morula_score": float(test_scores["morula"]),
                "test_8cell_score": float(test_scores["8-cell"]),
                "test_blastocyst_score": float(test_scores["blastocyst"]),
                "test_8cell_to_morula_drop": float(test_scores["8-cell"] - test_scores["morula"]),
            }
        )
    df = pd.DataFrame(rows)
    summary = {
        "n_splits": n_iter,
        "n_dmr_nodes": int(n),
        "fraction_train_min_morula": float(np.mean(df["train_min_stage"].eq("morula"))),
        "fraction_test_min_morula": float(np.mean(df["test_min_stage"].eq("morula"))),
        "fraction_test_morula_rank1": float(np.mean(df["test_morula_rank_lowest_is_1"].eq(1))),
        "fraction_test_8cell_to_morula_drop_positive": float(np.mean(df["test_8cell_to_morula_drop"] > 0)),
        "median_test_8cell_to_morula_drop": float(df["test_8cell_to_morula_drop"].median()),
    }
    return df, summary


def main() -> None:
    particles = pd.read_csv(RESULTS / "CSB_TRO_fused_product_particles.tsv", sep="\t")
    stage_summary = pd.read_csv(RESULTS / "CSB_TRO_path_space_stage_summary.tsv", sep="\t")

    loso_df, loso_summary = stage_distribution_prediction(particles)
    forecast_df, forecast_summary = early_to_morula_forecast(stage_summary)
    dmr_df, dmr_summary = dmr_split_validation()

    summary = {
        "model": "CSB-TRO prediction and validation experiments",
        "date": "2026-05-24",
        "leave_one_stage_distribution_prediction": loso_summary,
        "early_to_morula_forecast": forecast_summary,
        "dmr_split_validation": dmr_summary,
        "interpretation": (
            "The strongest predictive support comes from DMR split validation: held-out DMR subsets retain a morula "
            "minimum and an 8-cell-to-morula drop. Neighbor-stage distribution interpolation tests whether dynamic "
            "developmental structure improves over static previous-stage baselines. Early-to-morula trend forecasting "
            "is included as a difficult prospective task and should be reported honestly."
        ),
    }

    loso_df.to_csv(RESULTS / "CSB_TRO_prediction_leave_one_stage_distribution.tsv", sep="\t", index=False)
    forecast_df.to_csv(RESULTS / "CSB_TRO_prediction_early_to_morula_forecast.tsv", sep="\t", index=False)
    dmr_df.to_csv(RESULTS / "CSB_TRO_prediction_DMR_split_validation.tsv", sep="\t", index=False)
    (RESULTS / "CSB_TRO_prediction_validation_summary.json").write_text(json.dumps(summary, indent=2, ensure_ascii=False), encoding="utf-8")

    note = f"""# CSB-TRO Prediction and Validation

Date: 2026-05-24

## DMR Split Validation

- DMR split iterations: {dmr_summary["n_splits"]}
- DMR nodes: {dmr_summary["n_dmr_nodes"]}
- Fraction train minimum = morula: {dmr_summary["fraction_train_min_morula"]:.3f}
- Fraction held-out test minimum = morula: {dmr_summary["fraction_test_min_morula"]:.3f}
- Fraction held-out morula rank 1: {dmr_summary["fraction_test_morula_rank1"]:.3f}
- Fraction held-out 8-cell -> morula drop positive: {dmr_summary["fraction_test_8cell_to_morula_drop_positive"]:.3f}

## Leave-One-Stage Distribution Prediction

- Internal stages tested: {loso_summary["n_leave_one_internal_stage_tests"]}
- Fraction dynamic interpolation beats previous-stage baseline: {loso_summary["fraction_dynamic_beats_previous_stage"]:.3f}
- Fraction dynamic interpolation beats global-mean baseline: {loso_summary["fraction_dynamic_beats_global_mean"]:.3f}

## Early-to-Morula Forecast

- Best forecast model: {forecast_summary["best_forecast_model"]}
- Best error: {forecast_summary["best_error"]:.6f}
- Previous 8-cell baseline error: {forecast_summary["previous_8cell_error"]:.6f}

## Interpretation

These experiments add prediction-oriented evidence. The DMR split result is the strongest because it tests whether held-out age-DMR modules preserve the inferred reset minimum.
"""
    (RESULTS / "CSB_TRO_prediction_validation_interpretation.md").write_text(note, encoding="utf-8")
    print(json.dumps(summary, indent=2, ensure_ascii=True))


if __name__ == "__main__":
    main()
```
