## Supplemental code for "An entropy-based diagnostic framework for characterizing methylation state dynamics during preimplantation development": csb_tro_robustness.py

```
from __future__ import annotations

import json
from pathlib import Path

import numpy as np
import pandas as pd


OUT = Path("E:/实验更新_5_22/csb_tro_dynamics")
STAGE_ORDER = ["MII oocyte", "zygote/PN", "2-cell", "4-cell", "8-cell", "morula", "blastocyst"]
STATE_COLS = ["A", "Hm", "P", "Hr"]
POTENCY_THRESHOLD_QUANTILE = 0.60


def stage_agnostic_p_min(particles: pd.DataFrame) -> float:
    return float(particles["P"].quantile(POTENCY_THRESHOLD_QUANTILE))


def sinkhorn(a: np.ndarray, b: np.ndarray, cost: np.ndarray, epsilon: float, n_iter: int = 1200) -> np.ndarray:
    k = np.exp(-cost / max(epsilon, 1e-9))
    k = np.maximum(k, 1e-300)
    u = np.ones_like(a)
    v = np.ones_like(b)
    for _ in range(n_iter):
        u_new = a / np.maximum(k @ v, 1e-300)
        v_new = b / np.maximum(k.T @ u_new, 1e-300)
        if np.max(np.abs(u_new - u)) < 1e-10 and np.max(np.abs(v_new - v)) < 1e-10:
            u, v = u_new, v_new
            break
        u, v = u_new, v_new
    pi = (u[:, None] * k) * v[None, :]
    total = pi.sum()
    return pi / total if total > 0 else pi


def transition_metrics(x: np.ndarray, y: np.ndarray, p_min: float, epsilon: float = 0.075) -> dict[str, float]:
    diff = x[:, None, :] - y[None, :, :]
    base = np.sum(diff * diff, axis=2)
    age_increase = np.maximum(y[None, :, 0] - x[:, None, 0], 0.0) ** 2
    potency_low = np.maximum(p_min - y[None, :, 2], 0.0) ** 2
    cost = base + 1.65 * age_increase + 0.95 * potency_low
    a = np.full(x.shape[0], 1.0 / x.shape[0])
    b = np.full(y.shape[0], 1.0 / y.shape[0])
    pi = sinkhorn(a, b, cost, epsilon=epsilon)
    delta = y[None, :, :] - x[:, None, :]
    return {
        "mean_transport_A": float(np.sum(pi * delta[:, :, 0])),
        "mean_transport_Hm": float(np.sum(pi * delta[:, :, 1])),
        "mean_transport_P": float(np.sum(pi * delta[:, :, 2])),
        "mean_transport_Hr": float(np.sum(pi * delta[:, :, 3])),
        "mean_squared_displacement": float(np.sum(pi * np.sum(delta * delta, axis=2))),
    }


def stage_summary(particles: pd.DataFrame, stage_col: str = "stage") -> pd.DataFrame:
    summary = particles.groupby(stage_col, as_index=False).agg(
        n_particles=("particle_id", "count"),
        A_mean=("A", "mean"),
        Hm_mean=("Hm", "mean"),
        P_mean=("P", "mean"),
        Hr_mean=("Hr", "mean"),
    )
    summary = summary.rename(columns={stage_col: "stage"})
    summary["reset_score_A_minus_P"] = summary["A_mean"] - summary["P_mean"]
    summary["A_rank_lowest_is_1"] = summary["A_mean"].rank(method="min", ascending=True).astype(int)
    summary["P_rank_highest_is_1"] = summary["P_mean"].rank(method="min", ascending=False).astype(int)
    summary["reset_rank_lowest_is_1"] = summary["reset_score_A_minus_P"].rank(method="min", ascending=True).astype(int)
    summary["stage_order"] = summary["stage"].map({s: i for i, s in enumerate(STAGE_ORDER)})
    return summary.sort_values("stage_order").reset_index(drop=True)


def morula_metrics(summary: pd.DataFrame) -> dict[str, float]:
    row = summary[summary["stage"].eq("morula")].iloc[0]
    return {
        "morula_A_mean": float(row["A_mean"]),
        "morula_P_mean": float(row["P_mean"]),
        "morula_reset_score_A_minus_P": float(row["reset_score_A_minus_P"]),
        "morula_A_rank_lowest_is_1": int(row["A_rank_lowest_is_1"]),
        "morula_P_rank_highest_is_1": int(row["P_rank_highest_is_1"]),
        "morula_reset_rank_lowest_is_1": int(row["reset_rank_lowest_is_1"]),
    }


def empirical_p(null_values: np.ndarray, observed: float, direction: str) -> float:
    if direction == "le":
        count = int(np.sum(null_values <= observed))
    elif direction == "ge":
        count = int(np.sum(null_values >= observed))
    else:
        raise ValueError(direction)
    return float((count + 1) / (len(null_values) + 1))


def label_permutation(particles: pd.DataFrame, observed: dict[str, float], n: int, rng: np.random.Generator) -> tuple[pd.DataFrame, dict]:
    labels = particles["stage"].to_numpy(copy=True)
    rows = []
    for i in range(n):
        shuffled = particles.copy()
        shuffled["perm_stage"] = rng.permutation(labels)
        metrics = morula_metrics(stage_summary(shuffled, stage_col="perm_stage"))
        metrics["iteration"] = i
        rows.append(metrics)
    null = pd.DataFrame(rows)
    result = {
        "n": n,
        "p_morula_A_low_or_lower": empirical_p(null["morula_A_mean"].to_numpy(), observed["morula_A_mean"], "le"),
        "p_morula_P_high_or_higher": empirical_p(null["morula_P_mean"].to_numpy(), observed["morula_P_mean"], "ge"),
        "p_morula_reset_score_low_or_lower": empirical_p(
            null["morula_reset_score_A_minus_P"].to_numpy(), observed["morula_reset_score_A_minus_P"], "le"
        ),
        "fraction_A_rank1": float(np.mean(null["morula_A_rank_lowest_is_1"].eq(1))),
        "fraction_P_rank_top2": float(np.mean(null["morula_P_rank_highest_is_1"].le(2))),
        "fraction_reset_rank1": float(np.mean(null["morula_reset_rank_lowest_is_1"].eq(1))),
    }
    return null, result


def bootstrap_by_stage(particles: pd.DataFrame, observed_bridge: dict[str, float], n: int, rng: np.random.Generator) -> tuple[pd.DataFrame, dict]:
    rows = []
    stage_groups = {stage: df.reset_index(drop=True) for stage, df in particles.groupby("stage")}
    for i in range(n):
        parts = []
        for stage in STAGE_ORDER:
            df = stage_groups[stage]
            idx = rng.integers(0, len(df), size=len(df))
            parts.append(df.iloc[idx].copy())
        boot = pd.concat(parts, ignore_index=True)
        sm = stage_summary(boot)
        metrics = morula_metrics(sm)
        p_min = stage_agnostic_p_min(boot)
        x = boot[boot["stage"].eq("8-cell")][STATE_COLS].to_numpy(dtype=float)
        y = boot[boot["stage"].eq("morula")][STATE_COLS].to_numpy(dtype=float)
        bridge = transition_metrics(x, y, p_min=p_min)
        metrics.update({f"bridge_{k}": v for k, v in bridge.items()})
        metrics["iteration"] = i
        rows.append(metrics)
    boot_df = pd.DataFrame(rows)

    def ci(col: str) -> list[float]:
        q = np.quantile(boot_df[col].to_numpy(dtype=float), [0.025, 0.5, 0.975])
        return [float(x) for x in q]

    result = {
        "n": n,
        "morula_A_mean_ci025_median_ci975": ci("morula_A_mean"),
        "morula_P_mean_ci025_median_ci975": ci("morula_P_mean"),
        "morula_reset_score_ci025_median_ci975": ci("morula_reset_score_A_minus_P"),
        "bridge_8cell_to_morula_A_ci025_median_ci975": ci("bridge_mean_transport_A"),
        "bridge_8cell_to_morula_P_ci025_median_ci975": ci("bridge_mean_transport_P"),
        "fraction_morula_A_rank1": float(np.mean(boot_df["morula_A_rank_lowest_is_1"].eq(1))),
        "fraction_morula_P_rank_top2": float(np.mean(boot_df["morula_P_rank_highest_is_1"].le(2))),
        "fraction_8cell_to_morula_A_negative": float(np.mean(boot_df["bridge_mean_transport_A"] < 0)),
        "observed_8cell_to_morula_A": float(observed_bridge["mean_transport_A"]),
        "observed_8cell_to_morula_P": float(observed_bridge["mean_transport_P"]),
    }
    return boot_df, result


def random_stage_order(particles: pd.DataFrame, observed_bridge: dict[str, float], n: int, rng: np.random.Generator) -> tuple[pd.DataFrame, dict]:
    rows = []
    by_stage = {stage: particles[particles["stage"].eq(stage)][STATE_COLS].to_numpy(dtype=float) for stage in STAGE_ORDER}
    p_min = stage_agnostic_p_min(particles)
    for i in range(n):
        order = list(rng.permutation(STAGE_ORDER))
        if order.index("morula") == 0:
            continue
        prev_stage = order[order.index("morula") - 1]
        bridge = transition_metrics(by_stage[prev_stage], by_stage["morula"], p_min=p_min)
        rows.append({"iteration": i, "previous_stage": prev_stage, **bridge})
    null = pd.DataFrame(rows)
    result = {
        "n_orders_with_morula_not_first": int(len(null)),
        "p_entering_morula_A_drop_as_large": empirical_p(
            null["mean_transport_A"].to_numpy(), float(observed_bridge["mean_transport_A"]), "le"
        ),
        "p_entering_morula_P_change_as_low_or_lower": empirical_p(
            null["mean_transport_P"].to_numpy(), float(observed_bridge["mean_transport_P"]), "le"
        ),
        "fraction_entering_morula_A_negative": float(np.mean(null["mean_transport_A"] < 0)),
        "previous_stage_counts": {str(k): int(v) for k, v in null["previous_stage"].value_counts().sort_index().items()},
    }
    return null, result


def main() -> None:
    rng = np.random.default_rng(20260524)
    particles = pd.read_csv(OUT / "CSB_TRO_fused_product_particles.tsv", sep="\t")
    bridges = pd.read_csv(OUT / "CSB_TRO_fused_transition_bridges.tsv", sep="\t")
    particles = particles[particles["stage"].isin(STAGE_ORDER)].copy()

    observed_summary = stage_summary(particles)
    observed = morula_metrics(observed_summary)
    observed_bridge = bridges[(bridges["from_stage"].eq("8-cell")) & (bridges["to_stage"].eq("morula"))].iloc[0].to_dict()

    label_null, label_result = label_permutation(particles, observed, n=2000, rng=rng)
    bootstrap_df, bootstrap_result = bootstrap_by_stage(particles, observed_bridge, n=500, rng=rng)
    order_null, order_result = random_stage_order(particles, observed_bridge, n=1000, rng=rng)

    summary = {
        "model": "CSB-TRO fused product-distribution robustness",
        "date": "2026-05-24",
        "observed_morula": observed,
        "observed_8cell_to_morula": {
            "mean_transport_A": float(observed_bridge["mean_transport_A"]),
            "mean_transport_P": float(observed_bridge["mean_transport_P"]),
            "mean_transport_Hm": float(observed_bridge["mean_transport_Hm"]),
            "mean_transport_Hr": float(observed_bridge["mean_transport_Hr"]),
        },
        "stage_label_permutation": label_result,
        "within_stage_bootstrap": bootstrap_result,
        "random_stage_order": order_result,
        "interpretation": (
            "Small label-permutation p-values support that the morula low-A/high-P basin is not explained by arbitrary "
            "stage labels. Bootstrap fractions test stability under empirical resampling. Random-stage-order p-values "
            "test whether the observed 8-cell-to-morula entry is unusually age-decreasing among non-biological orders."
        ),
        "p_min_definition": f"stage-agnostic global fused-particle P quantile q={POTENCY_THRESHOLD_QUANTILE}; not derived from morula",
    }

    label_null.to_csv(OUT / "CSB_TRO_fused_label_permutation_null.tsv", sep="\t", index=False)
    bootstrap_df.to_csv(OUT / "CSB_TRO_fused_bootstrap_null.tsv", sep="\t", index=False)
    order_null.to_csv(OUT / "CSB_TRO_fused_random_stage_order_null.tsv", sep="\t", index=False)
    observed_summary.to_csv(OUT / "CSB_TRO_fused_robustness_observed_stage_summary.tsv", sep="\t", index=False)
    (OUT / "CSB_TRO_fused_robustness_summary.json").write_text(json.dumps(summary, indent=2, ensure_ascii=False), encoding="utf-8")

    note = f"""# CSB-TRO fused robustness checks

Date: 2026-05-24

## Tests

1. Stage-label permutation, n={label_result["n"]}
2. Within-stage bootstrap of fused particles, n={bootstrap_result["n"]}
3. Random stage order entering morula, usable orders={order_result["n_orders_with_morula_not_first"]}

## Main readout

- Observed morula A rank, rank 1 = lowest: {observed["morula_A_rank_lowest_is_1"]}
- Observed morula P rank, rank 1 = highest: {observed["morula_P_rank_highest_is_1"]}
- Observed morula reset score A-P rank, rank 1 = lowest: {observed["morula_reset_rank_lowest_is_1"]}
- Stage-label permutation p(A as low or lower): {label_result["p_morula_A_low_or_lower"]:.6f}
- Stage-label permutation p(P as high or higher): {label_result["p_morula_P_high_or_higher"]:.6f}
- Stage-label permutation p(A-P as low or lower): {label_result["p_morula_reset_score_low_or_lower"]:.6f}
- Bootstrap fraction morula A rank 1: {bootstrap_result["fraction_morula_A_rank1"]:.3f}
- Bootstrap fraction morula P rank top 2: {bootstrap_result["fraction_morula_P_rank_top2"]:.3f}
- Bootstrap fraction 8-cell -> morula A transport negative: {bootstrap_result["fraction_8cell_to_morula_A_negative"]:.3f}
- Random-order p(entering morula A drop as large): {order_result["p_entering_morula_A_drop_as_large"]:.6f}

## Interpretation

These checks support the fused CSB-TRO claim if morula remains low-A/high-P under bootstrap and if arbitrary relabeling or non-biological orderings rarely reproduce the same basin/entry pattern.
"""
    (OUT / "CSB_TRO_fused_robustness_interpretation.md").write_text(note, encoding="utf-8")
    print(json.dumps(summary, indent=2, ensure_ascii=False))


if __name__ == "__main__":
    main()
```
