## Supplemental code for "An entropy-based diagnostic framework for characterizing methylation state dynamics during preimplantation development": csb_tro_soft_marginal_uncertainty.py

```
from __future__ import annotations

import json
from pathlib import Path

import numpy as np
import pandas as pd


OUT = Path("E:/实验更新_5_22/csb_tro_dynamics")
STAGE_ORDER = ["MII oocyte", "zygote/PN", "2-cell", "4-cell", "8-cell", "morula", "blastocyst"]
STATE_COLS = ["A", "Hm", "P", "Hr"]
RHO_GRID = [0.0, 0.1, 0.5, 1.0, 2.0, 5.0]


def energy_distance(x: np.ndarray, y: np.ndarray, rng: np.random.Generator, max_n: int = 180) -> float:
    if len(x) > max_n:
        x = x[rng.choice(len(x), size=max_n, replace=False)]
    if len(y) > max_n:
        y = y[rng.choice(len(y), size=max_n, replace=False)]
    xy = np.linalg.norm(x[:, None, :] - y[None, :, :], axis=2).mean()
    xx = np.linalg.norm(x[:, None, :] - x[None, :, :], axis=2).mean()
    yy = np.linalg.norm(y[:, None, :] - y[None, :, :], axis=2).mean()
    return float(2 * xy - xx - yy)


def stage_uncertainty(particles: pd.DataFrame, n_boot: int = 400) -> tuple[pd.DataFrame, dict]:
    rng = np.random.default_rng(20260524)
    rows = []
    for stage in STAGE_ORDER:
        df = particles[particles["stage"].eq(stage)].reset_index(drop=True)
        observed = df[STATE_COLS].to_numpy(dtype=float)
        distances = []
        mean_rows = []
        for i in range(n_boot):
            idx = rng.integers(0, len(df), size=len(df))
            boot = observed[idx]
            distances.append(energy_distance(observed, boot, rng))
            mean_rows.append(boot.mean(axis=0))
        means = np.vstack(mean_rows)
        row = {
            "stage": stage,
            "n_particles": int(len(df)),
            "soft_marginal_D_energy_median": float(np.median(distances)),
            "soft_marginal_D_energy_q95": float(np.quantile(distances, 0.95)),
        }
        for j, col in enumerate(STATE_COLS):
            row[f"{col}_mean_boot_sd"] = float(means[:, j].std())
        rows.append(row)
    unc = pd.DataFrame(rows)
    path_summary = json.loads((OUT / "CSB_TRO_path_space_summary.json").read_text(encoding="utf-8"))
    j_path = float(path_summary["J_path_total"])
    total_d = float(unc["soft_marginal_D_energy_q95"].sum())
    sensitivity = []
    for rho in RHO_GRID:
        sensitivity.append(
            {
                "rho": rho,
                "J_path_without_soft_marginal": j_path,
                "soft_marginal_penalty_sum_q95": total_d,
                "rho_soft_marginal_penalty": rho * total_d,
                "J_with_soft_marginal_audit": j_path + rho * total_d,
            }
        )
    summary = {
        "model": "CSB-TRO soft marginal uncertainty audit",
        "date": "2026-05-24",
        "distance": "bootstrap energy distance between observed empirical stage distribution and bootstrap empirical distribution",
        "n_bootstrap": n_boot,
        "J_path_without_soft_marginal": j_path,
        "soft_marginal_penalty_sum_q95": total_d,
        "largest_uncertainty_stage": str(unc.sort_values("soft_marginal_D_energy_q95", ascending=False).iloc[0]["stage"]),
        "stage_uncertainty": unc.to_dict(orient="records"),
        "interpretation": (
            "This does not replace the hard-marginal bridge; it quantifies an uncertainty-aware soft-marginal penalty "
            "that can be included as sum_k rho_k D(p_tk, p_hat_k). It prevents overclaiming that empirical stage "
            "distributions are noise-free."
        ),
    }
    return unc, pd.DataFrame(sensitivity), summary


def main() -> None:
    particles = pd.read_csv(OUT / "CSB_TRO_fused_product_particles.tsv", sep="\t")
    particles = particles[particles["stage"].isin(STAGE_ORDER)].copy()
    uncertainty, sensitivity, summary = stage_uncertainty(particles)
    uncertainty.to_csv(OUT / "CSB_TRO_soft_marginal_stage_uncertainty.tsv", sep="\t", index=False)
    sensitivity.to_csv(OUT / "CSB_TRO_soft_marginal_objective_sensitivity.tsv", sep="\t", index=False)
    (OUT / "CSB_TRO_soft_marginal_uncertainty_summary.json").write_text(
        json.dumps(summary, indent=2, ensure_ascii=False), encoding="utf-8"
    )
    note = f"""# CSB-TRO soft marginal uncertainty audit

Date: 2026-05-24

This experiment addresses the fact that empirical stage distributions are noisy finite-sample estimates.

Instead of claiming only hard constraints `p_tk = p_hat_k`, the uncertainty-aware objective can be written as:

`J = KL(P||Q) + sum_k rho_k D(p_tk, p_hat_k) + lambda_A C_A + lambda_P C_P + lambda_G Omega_G`

where `D` is represented here by bootstrap energy distance.

## Main result

- Bootstrap replicates per stage: {summary["n_bootstrap"]}
- J path without soft marginal penalty: {summary["J_path_without_soft_marginal"]:.6f}
- Sum of stage q95 uncertainty penalties: {summary["soft_marginal_penalty_sum_q95"]:.6f}
- Largest uncertainty stage: {summary["largest_uncertainty_stage"]}

## Interpretation

This is an uncertainty audit and objective extension. It should be reported as a guard against overfitting small empirical stage distributions.
"""
    (OUT / "CSB_TRO_soft_marginal_uncertainty_interpretation.md").write_text(note, encoding="utf-8")
    print(json.dumps(summary, indent=2, ensure_ascii=True))


if __name__ == "__main__":
    main()
```
