## Supplemental code for "An entropy-based diagnostic framework for characterizing methylation state dynamics during preimplantation development": csb_tro_static_dynamic_gain.py

```
from __future__ import annotations

import json
from pathlib import Path

import numpy as np
import pandas as pd


ROOT = Path(__file__).resolve().parents[1]
RESULTS = ROOT / "results"
STATE_COLS = ["A", "Hm", "P", "Hr"]


def main() -> None:
    particles = pd.read_csv(RESULTS / "CSB_TRO_fused_product_particles.tsv", sep="\t")
    stage_summary = pd.read_csv(RESULTS / "CSB_TRO_path_space_stage_summary.tsv", sep="\t")
    transitions = pd.read_csv(RESULTS / "CSB_TRO_global_multimarginal_transition_summary.tsv", sep="\t")
    velocity = pd.read_csv(RESULTS / "CSB_TRO_global_multimarginal_velocity_field.tsv", sep="\t")
    path_summary = json.loads((RESULTS / "CSB_TRO_path_space_summary.json").read_text(encoding="utf-8"))
    dmr_graph = json.loads((RESULTS / "CSB_TRO_DMR_graph_laplacian_summary.json").read_text(encoding="utf-8"))
    fp = json.loads((RESULTS / "CSB_TRO_fokker_planck_summary.json").read_text(encoding="utf-8"))

    a0 = float(particles["A"].quantile(0.25))
    p0 = float(particles["P"].quantile(0.60))

    vel = velocity.copy()
    vel["A_next"] = vel["A"] + vel["vA"]
    vel["P_next"] = vel["P"] + vel["vP"]
    vel["in_reset_basin_now"] = (vel["A"] <= a0) & (vel["P"] >= p0)
    vel["in_reset_basin_next"] = (vel["A_next"] <= a0) & (vel["P_next"] >= p0)
    vel["enters_reset_basin"] = (~vel["in_reset_basin_now"]) & vel["in_reset_basin_next"]
    vel["leaves_reset_basin"] = vel["in_reset_basin_now"] & (~vel["in_reset_basin_next"])

    transition_basin = (
        vel.groupby(["stage", "to_stage"], as_index=False)
        .agg(
            n_particles=("particle_id", "count"),
            fraction_in_basin_now=("in_reset_basin_now", "mean"),
            fraction_in_basin_next=("in_reset_basin_next", "mean"),
            fraction_enters_basin=("enters_reset_basin", "mean"),
            fraction_leaves_basin=("leaves_reset_basin", "mean"),
            mean_vA=("vA", "mean"),
            mean_vP=("vP", "mean"),
        )
        .rename(columns={"stage": "from_stage"})
    )

    static = stage_summary[["stage", "n_particles", "A_mean", "P_mean", "reset_score_A_minus_P"]].copy()
    static["static_A_rank_lowest_is_1"] = static["A_mean"].rank(method="min", ascending=True).astype(int)
    static["static_P_rank_highest_is_1"] = static["P_mean"].rank(method="min", ascending=False).astype(int)
    static["static_reset_rank_lowest_is_1"] = static["reset_score_A_minus_P"].rank(method="min", ascending=True).astype(int)

    morula_static = static[static["stage"].eq("morula")].iloc[0]
    key_entry = transitions[transitions["from_stage"].eq("8-cell")].iloc[0]
    key_exit = transitions[transitions["from_stage"].eq("morula")].iloc[0]
    key_basin_entry = transition_basin[transition_basin["from_stage"].eq("8-cell")].iloc[0]
    key_basin_exit = transition_basin[transition_basin["from_stage"].eq("morula")].iloc[0]

    capability_rows = [
        {
            "capability": "stage reset ranking",
            "static_TRO": "yes",
            "CSB_TRO": "yes",
            "empirical_readout": f"morula static reset rank={int(morula_static['static_reset_rank_lowest_is_1'])}",
        },
        {
            "capability": "path-space objective",
            "static_TRO": "no",
            "CSB_TRO": "yes",
            "empirical_readout": f"J_path={path_summary['J_path_total']:.6f}; max marginal error={path_summary['max_row_marginal_error']:.2e}",
        },
        {
            "capability": "directed transition transport",
            "static_TRO": "no",
            "CSB_TRO": "yes",
            "empirical_readout": f"8-cell->morula dA={key_entry['mean_transport_A']:.6f}; morula->blastocyst dA={key_exit['mean_transport_A']:.6f}",
        },
        {
            "capability": "velocity field",
            "static_TRO": "no",
            "CSB_TRO": "yes",
            "empirical_readout": f"8-cell->morula mean vA={key_basin_entry['mean_vA']:.6f}; mean vP={key_basin_entry['mean_vP']:.6f}",
        },
        {
            "capability": "reset basin entry/exit",
            "static_TRO": "no",
            "CSB_TRO": "yes",
            "empirical_readout": f"entry fraction={key_basin_entry['fraction_enters_basin']:.3f}; morula exit fraction={key_basin_exit['fraction_leaves_basin']:.3f}",
        },
        {
            "capability": "DMR graph regularized objective",
            "static_TRO": "no",
            "CSB_TRO": "yes",
            "empirical_readout": f"DMR nodes={dmr_graph['n_dmr_nodes']}; edges={dmr_graph['n_edges']}; C_G={dmr_graph['graph_terms']['C_G_total_edge_normalized']:.6f}",
        },
        {
            "capability": "Fokker-Planck export",
            "static_TRO": "no",
            "CSB_TRO": "yes",
            "empirical_readout": f"mean diffusion D_A={fp['mean_diffusion']['D_A']:.6f}; D_P={fp['mean_diffusion']['D_P']:.6f}",
        },
    ]
    capability = pd.DataFrame(capability_rows)

    summary = {
        "model": "CSB-TRO static-vs-dynamic gain analysis",
        "date": "2026-05-24",
        "reset_basin_definition": {
            "A_threshold_q25": a0,
            "P_threshold_q60": p0,
            "definition": "B_reset = {z: A <= q25(A), P >= q60(P)} over fused particles",
        },
        "static_morula": {
            "A_rank_lowest_is_1": int(morula_static["static_A_rank_lowest_is_1"]),
            "P_rank_highest_is_1": int(morula_static["static_P_rank_highest_is_1"]),
            "reset_rank_lowest_is_1": int(morula_static["static_reset_rank_lowest_is_1"]),
        },
        "dynamic_gain": {
            "J_path_total": float(path_summary["J_path_total"]),
            "max_marginal_error": float(path_summary["max_row_marginal_error"]),
            "entry_8cell_to_morula_mean_transport_A": float(key_entry["mean_transport_A"]),
            "exit_morula_to_blastocyst_mean_transport_A": float(key_exit["mean_transport_A"]),
            "entry_8cell_to_morula_fraction_enters_basin": float(key_basin_entry["fraction_enters_basin"]),
            "exit_morula_to_blastocyst_fraction_leaves_basin": float(key_basin_exit["fraction_leaves_basin"]),
            "DMR_graph_nodes": int(dmr_graph["n_dmr_nodes"]),
            "DMR_graph_edges": int(dmr_graph["n_edges"]),
        },
        "interpretation": (
            "Static TRO identifies a stage-level reset score. CSB-TRO adds a path-space objective, directed transports, "
            "a velocity field, reset-basin entry/exit readouts, a DMR graph regularizer, and a Fokker-Planck-compatible "
            "drift-diffusion export."
        ),
    }

    capability.to_csv(RESULTS / "CSB_TRO_static_vs_dynamic_capability_table.tsv", sep="\t", index=False)
    transition_basin.to_csv(RESULTS / "CSB_TRO_dynamic_reset_basin_transition_table.tsv", sep="\t", index=False)
    (RESULTS / "CSB_TRO_static_dynamic_gain_summary.json").write_text(json.dumps(summary, indent=2, ensure_ascii=False), encoding="utf-8")

    note = f"""# Static TRO vs CSB-TRO Dynamic Gain

Date: 2026-05-24

## Main Point

Static TRO can rank stages. CSB-TRO adds directed path-space dynamics.

## Reset Basin

`B_reset = {{z: A <= q25(A), P >= q60(P)}}`

- A threshold: {a0:.6f}
- P threshold: {p0:.6f}

## Dynamic Gain

- Path objective J: {summary["dynamic_gain"]["J_path_total"]:.6f}
- 8-cell -> morula dA: {summary["dynamic_gain"]["entry_8cell_to_morula_mean_transport_A"]:.6f}
- Morula -> blastocyst dA: {summary["dynamic_gain"]["exit_morula_to_blastocyst_mean_transport_A"]:.6f}
- 8-cell -> morula reset-basin entry fraction: {summary["dynamic_gain"]["entry_8cell_to_morula_fraction_enters_basin"]:.3f}
- Morula -> blastocyst reset-basin leaving fraction: {summary["dynamic_gain"]["exit_morula_to_blastocyst_fraction_leaves_basin"]:.3f}
- DMR graph: {summary["dynamic_gain"]["DMR_graph_nodes"]} nodes, {summary["dynamic_gain"]["DMR_graph_edges"]} edges

## Interpretation

This analysis answers what CSB-TRO contributes beyond static TRO: path, transport, velocity, basin entry/exit, graph regularization, and PDE-compatible drift-diffusion terms.
"""
    (RESULTS / "CSB_TRO_static_dynamic_gain_interpretation.md").write_text(note, encoding="utf-8")
    print(json.dumps(summary, indent=2, ensure_ascii=True))


if __name__ == "__main__":
    main()
```
