## Supplemental code for "An entropy-based diagnostic framework for characterizing methylation state dynamics during preimplantation development": csb_tro_time_reference_sensitivity.py

```
from __future__ import annotations

import json
from pathlib import Path

import numpy as np
import pandas as pd


OUT = Path("E:/实验更新_5_22/csb_tro_dynamics")
STAGE_ORDER = ["MII oocyte", "zygote/PN", "2-cell", "4-cell", "8-cell", "morula", "blastocyst"]
STATE_COLS = ["A", "Hm", "P", "Hr"]
EPSILON = 0.075
LAMBDA_A = 1.65
LAMBDA_P = 0.95
POTENCY_THRESHOLD_QUANTILE = 0.60

TIME_GRIDS = {
    "unit_stage_time": [0, 1, 2, 3, 4, 5, 6],
    "approx_human_hours": [0, 24, 44, 68, 88, 112, 144],
    "normalized_developmental_time": [0.0, 0.17, 0.31, 0.47, 0.61, 0.78, 1.0],
}
REFERENCE_MODES = [
    "brownian_zero",
    "stage_mean_developmental_drift",
    "linear_time_developmental_drift",
]


def stage_agnostic_p_min(particles: pd.DataFrame) -> float:
    return float(particles["P"].quantile(POTENCY_THRESHOLD_QUANTILE))


def sinkhorn(a: np.ndarray, b: np.ndarray, cost: np.ndarray, epsilon: float = EPSILON, n_iter: int = 1500) -> np.ndarray:
    k = np.maximum(np.exp(-cost / max(epsilon, 1e-9)), 1e-300)
    u = np.ones_like(a)
    v = np.ones_like(b)
    for _ in range(n_iter):
        u_new = a / np.maximum(k @ v, 1e-300)
        v_new = b / np.maximum(k.T @ u_new, 1e-300)
        if np.max(np.abs(u_new - u)) < 1e-10 and np.max(np.abs(v_new - v)) < 1e-10:
            u, v = u_new, v_new
            break
        u, v = u_new, v_new
    pi = (u[:, None] * k) * v[None, :]
    return pi / max(pi.sum(), 1e-300)


def stage_summary(particles: pd.DataFrame) -> pd.DataFrame:
    sm = particles.groupby("stage", as_index=False).agg(
        n_particles=("particle_id", "count"),
        A_mean=("A", "mean"),
        Hm_mean=("Hm", "mean"),
        P_mean=("P", "mean"),
        Hr_mean=("Hr", "mean"),
    )
    sm["stage_order"] = sm["stage"].map({s: i for i, s in enumerate(STAGE_ORDER)})
    sm["A_rank_lowest_is_1"] = sm["A_mean"].rank(method="min", ascending=True).astype(int)
    sm["P_rank_highest_is_1"] = sm["P_mean"].rank(method="min", ascending=False).astype(int)
    sm["reset_score_A_minus_P"] = sm["A_mean"] - sm["P_mean"]
    sm["reset_rank_lowest_is_1"] = sm["reset_score_A_minus_P"].rank(method="min", ascending=True).astype(int)
    return sm.sort_values("stage_order").reset_index(drop=True)


def reference_displacement(
    reference_mode: str,
    x: np.ndarray,
    from_stage: str,
    to_stage: str,
    stage_means: pd.DataFrame,
    linear_coef: np.ndarray,
    dt: float,
) -> np.ndarray:
    if reference_mode == "brownian_zero":
        return np.zeros((1, 1, len(STATE_COLS)))
    if reference_mode == "stage_mean_developmental_drift":
        m0 = stage_means.loc[stage_means["stage"].eq(from_stage), STATE_COLS].to_numpy(dtype=float)[0]
        m1 = stage_means.loc[stage_means["stage"].eq(to_stage), STATE_COLS].to_numpy(dtype=float)[0]
        return (m1 - m0)[None, None, :]
    if reference_mode == "linear_time_developmental_drift":
        drift = linear_coef[:, 0] * dt
        return drift[None, None, :]
    raise ValueError(reference_mode)


def fit_linear_time_drift(stage_means: pd.DataFrame, times: list[float]) -> np.ndarray:
    t = np.asarray(times, dtype=float)
    x = np.column_stack([t, np.ones_like(t)])
    coefs = []
    ordered = stage_means.set_index("stage").loc[STAGE_ORDER, STATE_COLS]
    for col in STATE_COLS:
        beta = np.linalg.lstsq(x, ordered[col].to_numpy(dtype=float), rcond=None)[0]
        coefs.append(beta)
    return np.vstack(coefs)


def transition_metrics(
    x: np.ndarray,
    y: np.ndarray,
    p_min: float,
    dt: float,
    q_disp: np.ndarray,
) -> dict[str, float]:
    diff = y[None, :, :] - x[:, None, :]
    residual = diff - q_disp
    movement = np.sum(residual * residual, axis=2) / max(dt, 1e-12)
    age = np.maximum(y[None, :, 0] - x[:, None, 0], 0.0) ** 2
    potency = np.broadcast_to(np.maximum(p_min - y[None, :, 2], 0.0) ** 2, movement.shape)
    cost = movement + LAMBDA_A * age + LAMBDA_P * potency
    pi = sinkhorn(np.full(x.shape[0], 1 / x.shape[0]), np.full(y.shape[0], 1 / y.shape[0]), cost)
    independent = np.full_like(pi, 1.0 / pi.size)
    mask = pi > 0
    kl = float(np.sum(pi[mask] * np.log(pi[mask] / independent[mask])))
    return {
        "J_transition": float(np.sum(pi * cost) + EPSILON * kl),
        "mean_transport_A": float(np.sum(pi * diff[:, :, 0])),
        "mean_transport_P": float(np.sum(pi * diff[:, :, 2])),
        "drift_A_per_time": float(np.sum(pi * diff[:, :, 0]) / max(dt, 1e-12)),
        "drift_P_per_time": float(np.sum(pi * diff[:, :, 2]) / max(dt, 1e-12)),
    }


def run_sensitivity(particles: pd.DataFrame) -> tuple[pd.DataFrame, dict]:
    p_min = stage_agnostic_p_min(particles)
    base_summary = stage_summary(particles)
    stage_means = base_summary.rename(
        columns={"A_mean": "A", "Hm_mean": "Hm", "P_mean": "P", "Hr_mean": "Hr"}
    )[["stage", *STATE_COLS]]
    by_stage = {s: particles[particles["stage"].eq(s)][STATE_COLS].to_numpy(dtype=float) for s in STAGE_ORDER}
    rows = []
    for time_name, times in TIME_GRIDS.items():
        linear_coef = fit_linear_time_drift(stage_means, times)
        dts = np.diff(np.asarray(times, dtype=float))
        # Normalize biological-hour dt so objective scale is comparable while drift still records real-time units.
        objective_dts = dts / np.median(dts)
        for reference_mode in REFERENCE_MODES:
            j_total = 0.0
            key = {}
            for idx, (from_stage, to_stage) in enumerate(zip(STAGE_ORDER[:-1], STAGE_ORDER[1:])):
                dt_for_objective = float(objective_dts[idx])
                q_disp = reference_displacement(
                    reference_mode,
                    by_stage[from_stage],
                    from_stage,
                    to_stage,
                    stage_means,
                    linear_coef,
                    dt=float(dts[idx]),
                )
                metrics = transition_metrics(
                    by_stage[from_stage],
                    by_stage[to_stage],
                    p_min=p_min,
                    dt=dt_for_objective,
                    q_disp=q_disp,
                )
                j_total += metrics["J_transition"]
                if from_stage == "8-cell" and to_stage == "morula":
                    key = metrics
            morula = base_summary[base_summary["stage"].eq("morula")].iloc[0]
            rows.append(
                {
                    "time_grid": time_name,
                    "reference_mode": reference_mode,
                    "J_path": j_total,
                    "morula_A_rank_lowest_is_1": int(morula["A_rank_lowest_is_1"]),
                    "morula_P_rank_highest_is_1": int(morula["P_rank_highest_is_1"]),
                    "morula_reset_rank_lowest_is_1": int(morula["reset_rank_lowest_is_1"]),
                    "transport_8cell_to_morula_A": key["mean_transport_A"],
                    "transport_8cell_to_morula_P": key["mean_transport_P"],
                    "drift_8cell_to_morula_A_per_scaled_time": key["drift_A_per_time"],
                    "drift_8cell_to_morula_P_per_scaled_time": key["drift_P_per_time"],
                }
            )
    df = pd.DataFrame(rows)
    summary = {
        "n_sensitivity_runs": int(len(df)),
        "p_min": p_min,
        "p_min_definition": f"stage-agnostic global fused-particle P quantile q={POTENCY_THRESHOLD_QUANTILE}; not derived from morula",
        "fraction_morula_A_rank1": float(np.mean(df["morula_A_rank_lowest_is_1"].eq(1))),
        "fraction_morula_P_rank_top2": float(np.mean(df["morula_P_rank_highest_is_1"].le(2))),
        "fraction_morula_reset_rank1": float(np.mean(df["morula_reset_rank_lowest_is_1"].eq(1))),
        "fraction_8cell_to_morula_A_negative": float(np.mean(df["transport_8cell_to_morula_A"] < 0)),
        "J_path_min": float(df["J_path"].min()),
        "J_path_max": float(df["J_path"].max()),
    }
    return df, summary


def main() -> None:
    particles = pd.read_csv(OUT / "CSB_TRO_fused_product_particles.tsv", sep="\t")
    particles = particles[particles["stage"].isin(STAGE_ORDER)].copy()
    sensitivity, summary = run_sensitivity(particles)
    sensitivity.to_csv(OUT / "CSB_TRO_time_reference_sensitivity.tsv", sep="\t", index=False)
    (OUT / "CSB_TRO_time_reference_sensitivity_summary.json").write_text(
        json.dumps(summary, indent=2, ensure_ascii=False), encoding="utf-8"
    )
    note = f"""# CSB-TRO time-scale and reference-process sensitivity

Date: 2026-05-24

This experiment addresses two identifiability concerns: developmental stages are not equally spaced in real time, and the Schrödinger bridge depends on the reference process Q.

## Design

- Potency threshold is stage-agnostic: global P quantile q={POTENCY_THRESHOLD_QUANTILE}
- Time grids: {", ".join(TIME_GRIDS.keys())}
- Reference processes: {", ".join(REFERENCE_MODES)}

## Main result

- Sensitivity runs: {summary["n_sensitivity_runs"]}
- Fraction morula A rank 1: {summary["fraction_morula_A_rank1"]:.3f}
- Fraction morula P rank top 2: {summary["fraction_morula_P_rank_top2"]:.3f}
- Fraction morula reset rank 1: {summary["fraction_morula_reset_rank1"]:.3f}
- Fraction 8-cell -> morula A transport negative: {summary["fraction_8cell_to_morula_A_negative"]:.3f}

## Interpretation

The model should be reported as a minimum-relative-entropy dynamics under specified reference processes. Stability across these time/reference settings reduces, but does not eliminate, identifiability concerns.
"""
    (OUT / "CSB_TRO_time_reference_sensitivity_interpretation.md").write_text(note, encoding="utf-8")
    print(json.dumps(summary, indent=2, ensure_ascii=True))


if __name__ == "__main__":
    main()
```
