## Supplemental code for "An entropy-based diagnostic framework for characterizing methylation state dynamics during preimplantation development": make_csb_tro_paper_figures.py

```
from __future__ import annotations

from pathlib import Path

import numpy as np
import pandas as pd

import matplotlib

matplotlib.use("Agg")
import matplotlib.pyplot as plt
from matplotlib.patches import Rectangle


ROOT = Path(__file__).resolve().parents[1]
RESULTS = ROOT / "results"
FIGURES = ROOT / "figures"
FIGURES.mkdir(exist_ok=True)

STAGE_ORDER = ["MII oocyte", "zygote/PN", "2-cell", "4-cell", "8-cell", "morula", "blastocyst"]
STAGE_LABELS = ["MII", "zygote", "2-cell", "4-cell", "8-cell", "morula", "blastocyst"]
COLORS = {
    "MII oocyte": "#6b7280",
    "zygote/PN": "#2563eb",
    "2-cell": "#0891b2",
    "4-cell": "#059669",
    "8-cell": "#84cc16",
    "morula": "#dc2626",
    "blastocyst": "#7c3aed",
}


def style_axes(ax: plt.Axes) -> None:
    ax.spines["top"].set_visible(False)
    ax.spines["right"].set_visible(False)
    ax.spines["left"].set_color("#374151")
    ax.spines["bottom"].set_color("#374151")
    ax.tick_params(colors="#374151", labelsize=9)
    ax.grid(True, axis="y", color="#e5e7eb", linewidth=0.8)
    ax.set_axisbelow(True)


def savefig(fig: plt.Figure, stem: str) -> None:
    fig.tight_layout()
    fig.savefig(FIGURES / f"{stem}.png", dpi=350, bbox_inches="tight")
    fig.savefig(FIGURES / f"{stem}.pdf", bbox_inches="tight")
    fig.savefig(FIGURES / f"{stem}.svg", bbox_inches="tight")
    plt.close(fig)


def figure_1_static_tro_stage_score() -> None:
    df = pd.read_csv(RESULTS / "CSB_TRO_path_space_stage_summary.tsv", sep="\t")
    df["stage"] = pd.Categorical(df["stage"], STAGE_ORDER, ordered=True)
    df = df.sort_values("stage")
    df["reset_score"] = df["P_mean"] - df["A_mean"]
    df["reset_rank"] = df["reset_score"].rank(method="min", ascending=False).astype(int)

    fig, ax1 = plt.subplots(figsize=(8.2, 4.8))
    x = np.arange(len(df))
    bars = ax1.bar(
        x,
        df["reset_score"],
        color=[COLORS[s] for s in df["stage"]],
        edgecolor="#111827",
        linewidth=0.6,
        alpha=0.88,
    )
    ax1.axhline(0, color="#111827", linewidth=0.9)
    ax1.set_ylabel("Static reset score (P - A)", fontsize=11)
    ax1.set_xticks(x)
    ax1.set_xticklabels(STAGE_LABELS, rotation=25, ha="right")
    style_axes(ax1)
    ax1.grid(True, axis="y", color="#e5e7eb")

    ax2 = ax1.twinx()
    ax2.plot(x, df["A_mean"], "-o", color="#b91c1c", linewidth=2.0, markersize=5, label="A mean")
    ax2.plot(x, df["P_mean"], "-o", color="#047857", linewidth=2.0, markersize=5, label="P mean")
    ax2.set_ylabel("Stage mean", fontsize=11)
    ax2.spines["top"].set_visible(False)
    ax2.tick_params(colors="#374151", labelsize=9)

    morula_idx = list(df["stage"].astype(str)).index("morula")
    ax1.annotate(
        "post hoc\nrank 1",
        xy=(morula_idx, df.iloc[morula_idx]["reset_score"]),
        xytext=(morula_idx - 0.65, df["reset_score"].max() + 0.07),
        arrowprops=dict(arrowstyle="->", color="#dc2626", lw=1.2),
        color="#dc2626",
        fontsize=10,
        fontweight="bold",
        ha="center",
    )
    ax1.set_title("Static TRO stage score identifies morula post hoc", fontsize=13, fontweight="bold")
    ax2.legend(frameon=False, loc="upper left", bbox_to_anchor=(0.02, 0.96), fontsize=9)
    savefig(fig, "Figure1_static_TRO_stage_score")


def figure_2_velocity_field() -> None:
    points = pd.read_csv(RESULTS / "CSB_TRO_fused_product_particles.tsv", sep="\t")
    vel = pd.read_csv(RESULTS / "CSB_TRO_global_multimarginal_velocity_field.tsv", sep="\t")
    stage_mean = points.groupby("stage", as_index=False)[["A", "P"]].mean()
    stage_mean["stage"] = pd.Categorical(stage_mean["stage"], STAGE_ORDER, ordered=True)
    stage_mean = stage_mean.sort_values("stage")

    fig, ax = plt.subplots(figsize=(7.2, 6.2))
    rng = np.random.default_rng(20260524)
    sample_idx = rng.choice(len(vel), size=min(320, len(vel)), replace=False)
    draw = vel.iloc[sample_idx]
    ax.scatter(points["A"], points["P"], s=8, color="#9ca3af", alpha=0.18, linewidth=0)
    ax.quiver(
        draw["A"],
        draw["P"],
        draw["vA"],
        draw["vP"],
        angles="xy",
        scale_units="xy",
        scale=1.0,
        width=0.0022,
        color="#374151",
        alpha=0.33,
    )
    ax.plot(stage_mean["A"], stage_mean["P"], color="#111827", linewidth=1.3, alpha=0.7)
    for _, row in stage_mean.iterrows():
        stage = str(row["stage"])
        ax.scatter(row["A"], row["P"], s=110, color=COLORS[stage], edgecolor="white", linewidth=1.5, zorder=5)
        ax.text(row["A"] + 0.012, row["P"] + 0.012, stage.replace("MII oocyte", "MII"), fontsize=9, color=COLORS[stage], fontweight="bold")

    a0 = points["A"].quantile(0.25)
    p0 = points["P"].quantile(0.60)
    ax.add_patch(Rectangle((points["A"].min() - 0.02, p0), a0 - points["A"].min() + 0.02, points["P"].max() - p0 + 0.02, fill=False, lw=1.6, ls="--", ec="#dc2626"))
    ax.text(points["A"].min() + 0.005, p0 + 0.03, "reset basin\nlow A / high P", color="#dc2626", fontsize=10, fontweight="bold")
    ax.set_xlabel("A: age-associated epigenetic perturbation", fontsize=11)
    ax.set_ylabel("P: developmental potency", fontsize=11)
    ax.set_title("CSB-TRO A-P velocity field shows entry into a reset basin", fontsize=13, fontweight="bold")
    style_axes(ax)
    savefig(fig, "Figure2_CSB_TRO_AP_velocity_field")


def figure_3_transport_barplot() -> None:
    tr = pd.read_csv(RESULTS / "CSB_TRO_global_multimarginal_transition_summary.tsv", sep="\t")
    key = tr[tr["from_stage"].isin(["8-cell", "morula"])].copy()
    key["transition"] = key["from_stage"] + " -> " + key["to_stage"]
    metrics = ["mean_transport_A", "mean_transport_P"]
    plot_df = key.melt(id_vars="transition", value_vars=metrics, var_name="metric", value_name="mean_transport")
    plot_df["metric"] = plot_df["metric"].map({"mean_transport_A": "dA", "mean_transport_P": "dP"})

    fig, ax = plt.subplots(figsize=(7.6, 4.6))
    x = np.arange(len(key))
    width = 0.34
    vals_a = key["mean_transport_A"].to_numpy()
    vals_p = key["mean_transport_P"].to_numpy()
    ax.bar(x - width / 2, vals_a, width, color="#b91c1c", label="dA", edgecolor="#111827", linewidth=0.5)
    ax.bar(x + width / 2, vals_p, width, color="#047857", label="dP", edgecolor="#111827", linewidth=0.5)
    ax.axhline(0, color="#111827", linewidth=0.9)
    ax.set_xticks(x)
    ax.set_xticklabels(key["transition"], fontsize=10)
    ax.set_ylabel("Mean CSB transport", fontsize=11)
    ax.set_title("Directional transport into and out of morula", fontsize=13, fontweight="bold")
    ax.legend(frameon=False, fontsize=10)
    style_axes(ax)
    for xpos, val in zip(x - width / 2, vals_a):
        ax.text(xpos, val + (0.025 if val >= 0 else -0.055), f"{val:.3f}", ha="center", va="bottom" if val >= 0 else "top", fontsize=9)
    for xpos, val in zip(x + width / 2, vals_p):
        ax.text(xpos, val + (0.025 if val >= 0 else -0.055), f"{val:.3f}", ha="center", va="bottom" if val >= 0 else "top", fontsize=9)
    savefig(fig, "Figure3_transport_barplot")


def figure_4_reset_basin_entry_exit() -> None:
    basin = pd.read_csv(RESULTS / "CSB_TRO_dynamic_reset_basin_transition_table.tsv", sep="\t")
    basin["transition"] = basin["from_stage"] + " -> " + basin["to_stage"]
    basin["from_stage"] = pd.Categorical(basin["from_stage"], STAGE_ORDER[:-1], ordered=True)
    basin = basin.sort_values("from_stage")
    x = np.arange(len(basin))

    fig, ax = plt.subplots(figsize=(8.4, 4.8))
    ax.plot(x, basin["fraction_enters_basin"], "-o", color="#dc2626", lw=2.2, label="enters reset basin")
    ax.plot(x, basin["fraction_leaves_basin"], "-o", color="#2563eb", lw=2.2, label="leaves reset basin")
    ax.fill_between(x, basin["fraction_enters_basin"], color="#fecaca", alpha=0.35)
    ax.set_xticks(x)
    ax.set_xticklabels(basin["transition"], rotation=28, ha="right")
    ax.set_ylim(-0.02, max(0.58, basin[["fraction_enters_basin", "fraction_leaves_basin"]].max().max() + 0.08))
    ax.set_ylabel("Fraction of source particles", fontsize=11)
    ax.set_title("Reset basin entry and exit are dynamic CSB-TRO readouts", fontsize=13, fontweight="bold")
    ax.legend(frameon=False, fontsize=10, loc="upper left")
    style_axes(ax)
    savefig(fig, "Figure4_reset_basin_entry_exit")


def figure_5_dmr_split_validation() -> None:
    dmr = pd.read_csv(RESULTS / "CSB_TRO_prediction_DMR_split_validation.tsv", sep="\t")
    summary = pd.read_json(RESULTS / "CSB_TRO_prediction_validation_summary.json")
    fig, axes = plt.subplots(1, 2, figsize=(9.0, 4.5), gridspec_kw={"width_ratios": [1.05, 1.25]})

    cats = ["test min = morula", "test morula rank 1", "8-cell -> morula drop"]
    vals = [
        dmr["test_min_stage"].eq("morula").mean(),
        dmr["test_morula_rank_lowest_is_1"].eq(1).mean(),
        (dmr["test_8cell_to_morula_drop"] > 0).mean(),
    ]
    axes[0].bar(np.arange(len(vals)), vals, color=["#dc2626", "#ef4444", "#f97316"], edgecolor="#111827", linewidth=0.6)
    axes[0].set_xticks(np.arange(len(vals)))
    axes[0].set_xticklabels(cats, rotation=28, ha="right")
    axes[0].set_ylim(0, 1.08)
    axes[0].set_ylabel("Fraction across 500 DMR splits", fontsize=10)
    axes[0].set_title("Held-out DMR validation", fontsize=12, fontweight="bold")
    for i, v in enumerate(vals):
        axes[0].text(i, v + 0.025, f"{v:.3f}", ha="center", fontsize=9, fontweight="bold")
    style_axes(axes[0])

    axes[1].hist(dmr["test_8cell_to_morula_drop"], bins=28, color="#7c3aed", alpha=0.85, edgecolor="white")
    axes[1].axvline(0, color="#111827", lw=1.0)
    axes[1].axvline(dmr["test_8cell_to_morula_drop"].median(), color="#f97316", lw=2.0, label=f"median={dmr['test_8cell_to_morula_drop'].median():.3f}")
    axes[1].set_xlabel("Held-out DMR entropy drop: 8-cell - morula", fontsize=10)
    axes[1].set_ylabel("Split count", fontsize=10)
    axes[1].set_title("Morula minimum persists in held-out DMRs", fontsize=12, fontweight="bold")
    axes[1].legend(frameon=False, fontsize=9)
    style_axes(axes[1])
    fig.suptitle("DMR split validation supports the reset-basin signal", fontsize=13, fontweight="bold", y=1.02)
    savefig(fig, "Figure5_DMR_split_validation")


def main() -> None:
    figure_1_static_tro_stage_score()
    figure_2_velocity_field()
    figure_3_transport_barplot()
    figure_4_reset_basin_entry_exit()
    figure_5_dmr_split_validation()
    manifest = pd.DataFrame(
        {
            "figure": [
                "Figure1_static_TRO_stage_score",
                "Figure2_CSB_TRO_AP_velocity_field",
                "Figure3_transport_barplot",
                "Figure4_reset_basin_entry_exit",
                "Figure5_DMR_split_validation",
            ],
            "purpose": [
                "Static TRO stage score",
                "CSB-TRO A-P velocity field",
                "Key transport barplot",
                "Reset basin entry-exit",
                "DMR split validation",
            ],
        }
    )
    manifest.to_csv(FIGURES / "figure_manifest.tsv", sep="\t", index=False)
    print(f"Wrote figures to {FIGURES}")


if __name__ == "__main__":
    main()
```
