## Supplemental code for "An entropy-based diagnostic framework for characterizing methylation state dynamics during preimplantation development": run_advanced_latent_dynamics.py

```
from __future__ import annotations

import argparse
import json
import math
from pathlib import Path

import numpy as np
import pandas as pd


BASE = Path(r"E:\CSB_TRO_operator_time_DMR_dynamics_2026-05-25")
RESULTS = BASE / "results"
FIGURES = BASE / "figures"
DOCS = BASE / "docs"

STAGES = ["MII oocyte", "zygote/PN", "2-cell", "4-cell", "8-cell", "morula", "blastocyst"]
TAU = {stage: i / (len(STAGES) - 1) for i, stage in enumerate(STAGES)}


def rmse(a, b) -> float:
    a = np.asarray(a, dtype=float)
    b = np.asarray(b, dtype=float)
    return float(np.sqrt(np.mean((a - b) ** 2)))


def corr(a, b) -> float:
    a = np.asarray(a, dtype=float)
    b = np.asarray(b, dtype=float)
    aa = a - a.mean()
    bb = b - b.mean()
    den = math.sqrt(float((aa @ aa) * (bb @ bb)))
    return float((aa @ bb) / den) if den else float("nan")


def ridge(x: np.ndarray, y: np.ndarray, w: np.ndarray, lam: float) -> np.ndarray:
    sw = np.sqrt(np.maximum(w, 1e-12) / np.mean(np.maximum(w, 1e-12)))
    xw = x * sw[:, None]
    yw = y * sw[:, None] if y.ndim == 2 else y * sw
    penalty = lam * np.eye(x.shape[1])
    penalty[0, 0] = 0.0
    return np.linalg.solve(xw.T @ xw + penalty, xw.T @ yw)


def load_inputs():
    matrix = pd.read_csv(RESULTS / "CSB_TRO_DMR_state_matrix.tsv", sep="\t").set_index("sample_id")
    ann = pd.read_csv(RESULTS / "CSB_TRO_sample_tau_annotation.tsv", sep="\t").set_index("sample_id")
    pairs = pd.read_csv(RESULTS / "CSB_TRO_OT_sample_transition_couplings.tsv", sep="\t")
    pairs = pairs[pairs["from_sample_id"].isin(matrix.index) & pairs["to_sample_id"].isin(matrix.index)].copy()
    pairs = pairs[np.abs(pairs["delta_tau"].to_numpy(dtype=float)) > 1e-9].copy()
    return matrix, ann, pairs


def stage_ids(ann: pd.DataFrame, stage: str) -> list[str]:
    return list(ann.index[ann["stage"] == stage])


def fit_latent_basis(matrix: pd.DataFrame, q: int = 3):
    mu = matrix.mean(axis=0).to_numpy(dtype=float)
    sd = matrix.std(axis=0).to_numpy(dtype=float) + 1e-6
    x = (matrix.to_numpy(dtype=float) - mu) / sd
    _, _, vt = np.linalg.svd(x, full_matrices=False)
    components = vt[:q].T
    scores = x @ components
    score_df = pd.DataFrame(scores, index=matrix.index, columns=[f"PC{i + 1}" for i in range(q)])
    return mu, sd, components, score_df


def decode_latent(z: np.ndarray, mu: np.ndarray, sd: np.ndarray, components: np.ndarray) -> np.ndarray:
    z = np.asarray(z, dtype=float)
    return np.clip((z @ components.T) * sd + mu, 0.0, 1.0)


def fit_autonomous_latent_operator(
    score_df: pd.DataFrame,
    pairs: pd.DataFrame,
    train_stages: set[str] | None,
    q: int,
    lam: float,
):
    train = pairs.copy()
    if train_stages is not None:
        train = train[train["from_stage"].isin(train_stages) & train["to_stage"].isin(train_stages)].copy()
    zf = score_df.loc[train["from_sample_id"]].to_numpy(dtype=float)
    zt = score_df.loc[train["to_sample_id"]].to_numpy(dtype=float)
    tau = train["from_tau"].to_numpy(dtype=float)
    y = (zt - zf) / train["delta_tau"].to_numpy(dtype=float)[:, None]
    x = np.column_stack([np.ones(len(train)), tau, zf])
    coef = ridge(x, y, train["sample_coupling_weight"].to_numpy(dtype=float), lam)
    residual = y - x @ coef
    cov = np.cov(residual.T) + np.eye(q) * 1e-8
    return coef, cov, train


def drift(z: np.ndarray, tau: float, coef: np.ndarray) -> np.ndarray:
    z = np.asarray(z, dtype=float)
    x = np.concatenate([[1.0, tau], z])
    return x @ coef


def rollout_latent(
    z0: np.ndarray,
    tau0: float,
    target_taus: list[float],
    coef: np.ndarray,
    residual_cov: np.ndarray | None = None,
    rng: np.random.Generator | None = None,
) -> dict[float, np.ndarray]:
    z = np.asarray(z0, dtype=float).copy()
    tau = float(tau0)
    out = {}
    for target_tau in target_taus:
        dt = float(target_tau - tau)
        if dt < -1e-9:
            raise ValueError("target_taus must be increasing")
        if dt > 1e-12:
            z = z + dt * drift(z, tau, coef)
            if residual_cov is not None and rng is not None:
                z = z + rng.multivariate_normal(np.zeros(len(z)), residual_cov * dt)
        tau = float(target_tau)
        out[tau] = z.copy()
    return out


def rbf_mmd(x: np.ndarray, y: np.ndarray) -> float:
    x = np.asarray(x, dtype=float)
    y = np.asarray(y, dtype=float)
    both = np.vstack([x, y])
    d2 = ((both[:, None, :] - both[None, :, :]) ** 2).sum(axis=2)
    med = np.median(d2[d2 > 0]) if np.any(d2 > 0) else 1.0
    gamma = 1.0 / (2.0 * med + 1e-12)
    kxx = np.exp(-gamma * ((x[:, None, :] - x[None, :, :]) ** 2).sum(axis=2)).mean()
    kyy = np.exp(-gamma * ((y[:, None, :] - y[None, :, :]) ** 2).sum(axis=2)).mean()
    kxy = np.exp(-gamma * ((x[:, None, :] - y[None, :, :]) ** 2).sum(axis=2)).mean()
    return float(max(kxx + kyy - 2 * kxy, 0.0))


def sliced_wasserstein(x: np.ndarray, y: np.ndarray, rng: np.random.Generator, n_proj: int = 128) -> float:
    x = np.asarray(x, dtype=float)
    y = np.asarray(y, dtype=float)
    vals = []
    for _ in range(n_proj):
        v = rng.normal(size=x.shape[1])
        v = v / (np.linalg.norm(v) + 1e-12)
        px = np.sort(x @ v)
        py = np.sort(y @ v)
        n = max(len(px), len(py))
        q = np.linspace(0, 1, n)
        pxq = np.quantile(px, q)
        pyq = np.quantile(py, q)
        vals.append(np.mean(np.abs(pxq - pyq)))
    return float(np.mean(vals))


def covariance_error(x: np.ndarray, y: np.ndarray) -> float:
    cx = np.cov(np.asarray(x, dtype=float).T)
    cy = np.cov(np.asarray(y, dtype=float).T)
    return float(np.linalg.norm(cx - cy, ord="fro"))


def basin_scores(pred_z: np.ndarray, obs_z: np.ndarray, morula_z: np.ndarray) -> dict[str, float]:
    center = morula_z.mean(axis=0)
    morula_dist = np.linalg.norm(morula_z - center, axis=1)
    threshold = float(np.quantile(morula_dist, 0.9))
    pred_dist = np.linalg.norm(pred_z - center, axis=1)
    obs_dist = np.linalg.norm(obs_z - center, axis=1)
    return {
        "basin_threshold_morula_q90": threshold,
        "pred_basin_occupancy": float(np.mean(pred_dist <= threshold)),
        "observed_basin_occupancy": float(np.mean(obs_dist <= threshold)),
        "pred_mean_distance_to_morula_centroid": float(pred_dist.mean()),
        "observed_mean_distance_to_morula_centroid": float(obs_dist.mean()),
    }


def distribution_metrics(pred_z, obs_z, pred_dmr, obs_dmr, morula_z, rng) -> dict[str, float]:
    out = {
        "latent_mean_rmse": rmse(pred_z.mean(axis=0), obs_z.mean(axis=0)),
        "latent_mmd_rbf": rbf_mmd(pred_z, obs_z),
        "latent_sliced_wasserstein": sliced_wasserstein(pred_z, obs_z, rng),
        "latent_covariance_frobenius": covariance_error(pred_z, obs_z),
        "dmr_mean_rmse": rmse(pred_dmr.mean(axis=0), obs_dmr.mean(axis=0)),
        "dmr_correlation": corr(pred_dmr.mean(axis=0), obs_dmr.mean(axis=0)),
        "dmr_distributional_rmse": rmse(pred_dmr, obs_dmr.mean(axis=0)[None, :]),
    }
    out.update(basin_scores(pred_z, obs_z, morula_z))
    return out


def run_multistep_rollouts(matrix, ann, score_df, mu, sd, components, coef, cov, rng, stochastic_n: int, train_protocol: str):
    rows = []
    pred_rows = []
    morula_z = score_df.loc[stage_ids(ann, "morula")].to_numpy(dtype=float)
    for start_stage in ["MII oocyte", "zygote/PN", "2-cell", "4-cell", "8-cell"]:
        target_stages = [s for s in STAGES if TAU[s] >= TAU[start_stage]]
        target_taus = [TAU[s] for s in target_stages]
        start_samples = stage_ids(ann, start_stage)
        det_by_stage = {stage: [] for stage in target_stages}
        sto_by_stage = {stage: [] for stage in target_stages}
        for sample in start_samples:
            z0 = score_df.loc[sample].to_numpy(dtype=float)
            det = rollout_latent(z0, TAU[start_stage], target_taus, coef)
            for stage in target_stages:
                det_by_stage[stage].append(det[TAU[stage]])
            for _ in range(stochastic_n):
                sto = rollout_latent(z0, TAU[start_stage], target_taus, coef, cov, rng)
                for stage in target_stages:
                    sto_by_stage[stage].append(sto[TAU[stage]])
        for stage in target_stages:
            obs_ids = stage_ids(ann, stage)
            obs_z = score_df.loc[obs_ids].to_numpy(dtype=float)
            obs_dmr = matrix.loc[obs_ids].to_numpy(dtype=float)
            for mode, by_stage in [("deterministic", det_by_stage), ("stochastic_residual", sto_by_stage)]:
                pred_z = np.vstack(by_stage[stage])
                pred_dmr = decode_latent(pred_z, mu, sd, components)
                metrics = distribution_metrics(pred_z, obs_z, pred_dmr, obs_dmr, morula_z, rng)
                rows.append({
                    "train_protocol": train_protocol,
                    "rollout_mode": mode,
                    "start_stage": start_stage,
                    "target_stage": stage,
                    "start_tau": TAU[start_stage],
                    "target_tau": TAU[stage],
                    "n_predicted_particles": len(pred_z),
                    "n_observed_samples": len(obs_z),
                    **metrics,
                })
            det_mean = np.vstack(det_by_stage[stage]).mean(axis=0)
            pred_rows.append({
                "train_protocol": train_protocol,
                "start_stage": start_stage,
                "target_stage": stage,
                **{f"pred_PC{i + 1}_mean": float(det_mean[i]) for i in range(det_mean.shape[0])},
                **{f"obs_PC{i + 1}_mean": float(obs_z.mean(axis=0)[i]) for i in range(obs_z.shape[1])},
            })
    return pd.DataFrame(rows), pd.DataFrame(pred_rows)


def jacobian_stability(coef: np.ndarray) -> tuple[pd.DataFrame, pd.DataFrame]:
    q = coef.shape[1]
    jac = coef[2 : 2 + q, :].T
    eig = np.linalg.eigvals(jac)
    jac_rows = []
    for i in range(q):
        for j in range(q):
            jac_rows.append({"row_dPC_dtau": f"dPC{i + 1}", "col_PC": f"PC{j + 1}", "jacobian_value": float(jac[i, j])})
    eig_rows = []
    for i, val in enumerate(eig):
        eig_rows.append({
            "eigen_index": i + 1,
            "real_part": float(np.real(val)),
            "imag_part": float(np.imag(val)),
            "stability_class": "contracting" if np.real(val) < 0 else "expanding",
        })
    return pd.DataFrame(jac_rows), pd.DataFrame(eig_rows)


def perturbation_analysis(matrix, ann, score_df, mu, sd, components, coef, loadings, rng):
    start_stage = "8-cell"
    target_stage = "morula"
    start_samples = stage_ids(ann, start_stage)
    obs = matrix.loc[stage_ids(ann, target_stage)].mean(axis=0).to_numpy(dtype=float)
    base_preds = []
    per_sample_z = []
    for sample in start_samples:
        z0 = score_df.loc[sample].to_numpy(dtype=float)
        zt = rollout_latent(z0, TAU[start_stage], [TAU[target_stage]], coef)[TAU[target_stage]]
        per_sample_z.append(zt)
        base_preds.append(decode_latent(zt[None, :], mu, sd, components)[0])
    base_pred = np.vstack(base_preds).mean(axis=0)
    base_rmse = rmse(base_pred, obs)
    rows = [{"perturbation": "none", "target_stage": target_stage, "morula_rmse": base_rmse, "delta_rmse_vs_unperturbed": 0.0}]

    for pc in range(coef.shape[1]):
        for direction, mult in [("increase", 1.0), ("decrease", -1.0)]:
            preds = []
            scale = score_df[f"PC{pc + 1}"].std()
            for sample in start_samples:
                z0 = score_df.loc[sample].to_numpy(dtype=float).copy()
                z0[pc] += mult * scale
                zt = rollout_latent(z0, TAU[start_stage], [TAU[target_stage]], coef)[TAU[target_stage]]
                preds.append(decode_latent(zt[None, :], mu, sd, components)[0])
            score = rmse(np.vstack(preds).mean(axis=0), obs)
            rows.append({
                "perturbation": f"PC{pc + 1}_{direction}_1sd_at_8cell",
                "target_stage": target_stage,
                "morula_rmse": score,
                "delta_rmse_vs_unperturbed": score - base_rmse,
            })

    ranking = pd.DataFrame({
        "cluster_name": matrix.columns,
        "latent_loading_norm": np.linalg.norm(loadings, axis=1),
    }).sort_values("latent_loading_norm", ascending=False)
    for n_remove in [10, 25, 50]:
        keep = np.ones(matrix.shape[1], dtype=bool)
        top = ranking.head(n_remove)["cluster_name"].tolist()
        idx = [list(matrix.columns).index(c) for c in top]
        keep[idx] = False
        score = rmse(base_pred[keep], obs[keep])
        rows.append({
            "perturbation": f"remove_top{n_remove}_latent_loading_DMRs_from_score",
            "target_stage": target_stage,
            "morula_rmse": score,
            "delta_rmse_vs_unperturbed": score - base_rmse,
        })
    return pd.DataFrame(rows), ranking


def svg_rollout(path: Path, metrics: pd.DataFrame):
    det = metrics[(metrics["rollout_mode"] == "deterministic") & (metrics["train_protocol"] == "full_all_transitions")].copy()
    start = "MII oocyte"
    rows = det[det["start_stage"] == start]
    width, height = 880, 430
    left, right, top, bottom = 80, 30, 45, 95
    vals = rows["dmr_mean_rmse"].to_numpy(dtype=float)
    vmax = max(vals) * 1.2 if len(vals) else 1.0
    gap = (width - left - right) / len(rows)
    out = [
        f'<svg xmlns="http://www.w3.org/2000/svg" width="{width}" height="{height}" viewBox="0 0 {width} {height}">',
        '<rect width="100%" height="100%" fill="white"/>',
        f'<text x="{left}" y="28" font-family="Arial" font-size="20" font-weight="700">Latent autonomous multi-step rollout from MII oocyte</text>',
        f'<line x1="{left}" y1="{height-bottom}" x2="{width-right}" y2="{height-bottom}" stroke="#222"/>',
        f'<line x1="{left}" y1="{top}" x2="{left}" y2="{height-bottom}" stroke="#222"/>',
    ]
    for i, row in enumerate(rows.itertuples()):
        x = left + i * gap + gap * 0.2
        h = row.dmr_mean_rmse / vmax * (height - top - bottom)
        y = height - bottom - h
        fill = "#3b76a8" if row.target_stage == "morula" else "#777777"
        out.append(f'<rect x="{x:.2f}" y="{y:.2f}" width="{gap * 0.55:.2f}" height="{h:.2f}" fill="{fill}"/>')
        out.append(f'<text x="{x+gap*0.275:.2f}" y="{y-6:.2f}" text-anchor="middle" font-family="Arial" font-size="11">{row.dmr_mean_rmse:.3f}</text>')
        out.append(f'<text x="{x+gap*0.275:.2f}" y="{height-bottom+18}" text-anchor="end" transform="rotate(-30 {x+gap*0.275:.2f} {height-bottom+18})" font-family="Arial" font-size="11">{row.target_stage}</text>')
    out.append(f'<text x="{left}" y="{height-25}" font-family="Arial" font-size="12">Metric: DMR mean-state RMSE after continuous latent operator-time integration.</text>')
    out.append("</svg>")
    path.write_text("\n".join(out), encoding="utf-8")


def write_doc(path: Path, metrics: pd.DataFrame, eig: pd.DataFrame, perturb: pd.DataFrame):
    full_metrics = metrics[metrics["train_protocol"] == "full_all_transitions"]
    morula = full_metrics[(full_metrics["rollout_mode"] == "deterministic") & (full_metrics["start_stage"] == "8-cell") & (full_metrics["target_stage"] == "morula")].iloc[0]
    full = full_metrics[(full_metrics["rollout_mode"] == "deterministic") & (full_metrics["start_stage"] == "MII oocyte") & (full_metrics["target_stage"] == "blastocyst")].iloc[0]
    strict = metrics[(metrics["train_protocol"] == "leave_morula_transitions") & (metrics["rollout_mode"] == "deterministic") & (metrics["start_stage"] == "8-cell") & (metrics["target_stage"] == "morula")]
    lines = [
        "# Advanced latent operator-time dynamics",
        "",
        "This package upgrades the prior leave-morula-out point prediction into an autonomous latent-state dynamics analysis. The fitted drift uses only latent state and stage-anchored operator time, so it can be integrated across multiple stages without supplying future-stage summary variables.",
        "",
        f"- 8-cell to morula deterministic rollout DMR mean RMSE: {morula.dmr_mean_rmse:.4f}",
        f"- 8-cell to morula latent MMD: {morula.latent_mmd_rbf:.4f}",
        f"- 8-cell to morula predicted basin occupancy: {morula.pred_basin_occupancy:.4f}",
        f"- MII to blastocyst deterministic rollout DMR mean RMSE: {full.dmr_mean_rmse:.4f}",
        "",
        "Jacobian eigenvalues of the autonomous latent drift:",
    ]
    if len(strict):
        lines.insert(10, f"- strict leave-morula autonomous 8-cell to morula DMR mean RMSE: {strict.iloc[0].dmr_mean_rmse:.4f}")
    for row in eig.itertuples():
        lines.append(f"- eigen {row.eigen_index}: real={row.real_part:.4f}, imag={row.imag_part:.4f}, class={row.stability_class}")
    lines.extend(["", "Largest perturbation effects on morula prediction:"])
    top = perturb.sort_values("delta_rmse_vs_unperturbed", ascending=False).head(6)
    for row in top.itertuples():
        lines.append(f"- {row.perturbation}: delta RMSE={row.delta_rmse_vs_unperturbed:.4f}")
    lines.extend([
        "",
        "Interpretation boundary: this is still stage-anchored operator time, not real longitudinal tracking of the same embryo. The stochastic residual simulation is a low-dimensional residual diffusion approximation, not a full Fokker-Planck population PDE.",
    ])
    path.write_text("\n".join(lines) + "\n", encoding="utf-8")


def main():
    parser = argparse.ArgumentParser()
    parser.add_argument("--q", type=int, default=3)
    parser.add_argument("--lambda", dest="lam", type=float, default=1000.0)
    parser.add_argument("--stochastic-n", type=int, default=20)
    parser.add_argument("--seed", type=int, default=20260525)
    args = parser.parse_args()

    RESULTS.mkdir(parents=True, exist_ok=True)
    FIGURES.mkdir(parents=True, exist_ok=True)
    DOCS.mkdir(parents=True, exist_ok=True)
    rng = np.random.default_rng(args.seed)

    matrix, ann, pairs = load_inputs()
    mu, sd, components, score_df = fit_latent_basis(matrix, q=args.q)
    protocols = [
        ("full_all_transitions", None),
        ("leave_morula_transitions", set(stage for stage in STAGES if stage != "morula")),
        ("early_to_4cell_transitions", {"MII oocyte", "zygote/PN", "2-cell", "4-cell"}),
    ]
    metric_parts = []
    prediction_parts = []
    coef_parts = []
    cov_parts = []
    fitted = {}
    for protocol_name, train_stages in protocols:
        coef, cov, train = fit_autonomous_latent_operator(score_df, pairs, train_stages=train_stages, q=args.q, lam=args.lam)
        fitted[protocol_name] = (coef, cov, len(train))
        metrics_part, predictions_part = run_multistep_rollouts(matrix, ann, score_df, mu, sd, components, coef, cov, rng, args.stochastic_n, protocol_name)
        metric_parts.append(metrics_part)
        prediction_parts.append(predictions_part)
        coef_df = pd.DataFrame(coef, index=["intercept", "tau", *[f"PC{i + 1}" for i in range(args.q)]], columns=[f"dPC{i + 1}_dtau" for i in range(args.q)]).reset_index().rename(columns={"index": "feature"})
        coef_df.insert(0, "train_protocol", protocol_name)
        coef_parts.append(coef_df)
        cov_df = pd.DataFrame(cov, index=[f"PC{i + 1}" for i in range(args.q)], columns=[f"PC{i + 1}" for i in range(args.q)]).reset_index().rename(columns={"index": "row_PC"})
        cov_df.insert(0, "train_protocol", protocol_name)
        cov_parts.append(cov_df)
    rollout_metrics = pd.concat(metric_parts, ignore_index=True)
    rollout_predictions = pd.concat(prediction_parts, ignore_index=True)
    coef, cov, train_n = fitted["full_all_transitions"]
    jac, eig = jacobian_stability(coef)
    perturb, dmr_rank = perturbation_analysis(matrix, ann, score_df, mu, sd, components, coef, components, rng)

    rollout_metrics.to_csv(RESULTS / "CSB_TRO_latent_multistep_rollout_metrics.tsv", sep="\t", index=False)
    rollout_predictions.to_csv(RESULTS / "CSB_TRO_latent_multistep_rollout_stage_predictions.tsv", sep="\t", index=False)
    pd.concat(coef_parts, ignore_index=True).to_csv(RESULTS / "CSB_TRO_latent_autonomous_velocity_coefficients.tsv", sep="\t", index=False)
    pd.concat(cov_parts, ignore_index=True).to_csv(RESULTS / "CSB_TRO_latent_residual_diffusion_covariance.tsv", sep="\t", index=False)
    jac.to_csv(RESULTS / "CSB_TRO_latent_jacobian.tsv", sep="\t", index=False)
    eig.to_csv(RESULTS / "CSB_TRO_latent_stability_eigenvalues.tsv", sep="\t", index=False)
    perturb.to_csv(RESULTS / "CSB_TRO_latent_counterfactual_perturbations.tsv", sep="\t", index=False)
    dmr_rank.to_csv(RESULTS / "CSB_TRO_latent_loading_DMR_ranking.tsv", sep="\t", index=False)
    score_df.reset_index().rename(columns={"index": "sample_id"}).to_csv(RESULTS / "CSB_TRO_latent_autonomous_scores.tsv", sep="\t", index=False)
    pd.DataFrame(components, index=matrix.columns, columns=[f"PC{i + 1}_loading" for i in range(args.q)]).reset_index().rename(columns={"index": "cluster_name"}).to_csv(RESULTS / "CSB_TRO_latent_autonomous_loadings.tsv", sep="\t", index=False)

    svg_rollout(FIGURES / "CSB_TRO_latent_multistep_rollout_rmse.svg", rollout_metrics)
    write_doc(DOCS / "CSB_TRO_advanced_latent_dynamics_interpretation.md", rollout_metrics, eig, perturb)
    manifest = {
        "q": args.q,
        "ridge_lambda": args.lam,
        "stochastic_particles_per_start_sample": args.stochastic_n,
        "seed": args.seed,
        "n_training_pairs_by_protocol": {name: int(fitted[name][2]) for name, _ in protocols},
        "outputs": [
            str(RESULTS / "CSB_TRO_latent_multistep_rollout_metrics.tsv"),
            str(RESULTS / "CSB_TRO_latent_residual_diffusion_covariance.tsv"),
            str(RESULTS / "CSB_TRO_latent_stability_eigenvalues.tsv"),
            str(RESULTS / "CSB_TRO_latent_counterfactual_perturbations.tsv"),
            str(FIGURES / "CSB_TRO_latent_multistep_rollout_rmse.svg"),
            str(DOCS / "CSB_TRO_advanced_latent_dynamics_interpretation.md"),
        ],
    }
    (RESULTS / "CSB_TRO_advanced_latent_dynamics_manifest.json").write_text(json.dumps(manifest, indent=2), encoding="utf-8")
    print(json.dumps({
        "manifest": manifest,
        "morula_from_8cell": rollout_metrics[(rollout_metrics["rollout_mode"] == "deterministic") & (rollout_metrics["start_stage"] == "8-cell") & (rollout_metrics["target_stage"] == "morula")].to_dict(orient="records"),
        "stability": eig.to_dict(orient="records"),
        "top_perturbations": perturb.sort_values("delta_rmse_vs_unperturbed", ascending=False).head(5).to_dict(orient="records"),
    }, indent=2))


if __name__ == "__main__":
    main()
```
