## Supplemental code for "An entropy-based diagnostic framework for characterizing methylation state dynamics during preimplantation development": run_basin_residual_control_field.py

```
from __future__ import annotations

import argparse
import json
import sys
from pathlib import Path

import numpy as np
import pandas as pd


BASE = Path(r"E:\CSB_TRO_operator_time_DMR_dynamics_2026-05-25")
CODE = BASE / "code"
RESULTS = BASE / "results"
FIGURES = BASE / "figures"
DOCS = BASE / "docs"
sys.path.insert(0, str(CODE))

from run_morula_basin_sde import (  # noqa: E402
    TAU,
    basin_definition,
    corr,
    decode_latent,
    distribution_metrics,
    euler_rollout_batch,
    fit_latent_basis,
    fit_operator,
    load_inputs,
    rmse,
    stage_ids,
)


PRE_MORULA_STAGES = ["MII oocyte", "zygote/PN", "2-cell", "4-cell", "8-cell"]


def encode_delta_to_latent(delta_beta: np.ndarray, sd: np.ndarray, components: np.ndarray) -> np.ndarray:
    return (np.asarray(delta_beta, dtype=float) / sd) @ components


def decode_delta_to_dmr(delta_z: np.ndarray, sd: np.ndarray, components: np.ndarray) -> np.ndarray:
    return (np.asarray(delta_z, dtype=float) @ components.T) * sd


def simulate_strict_pre_morula(score_df, ann, coef, n_steps):
    z8 = score_df.loc[stage_ids(ann, "8-cell")].to_numpy(dtype=float)
    return euler_rollout_batch(
        z8,
        TAU["8-cell"],
        TAU["morula"],
        coef,
        np.random.default_rng(1),
        n_steps=n_steps,
    )


def load_optional_table(path: Path) -> pd.DataFrame:
    if path.exists():
        return pd.read_csv(path, sep="\t")
    return pd.DataFrame()


def build_dmr_residual_table(matrix, metadata, modules, components, sd, residual_z, obs_delta_dmr, model_delta_dmr):
    latent_delta_dmr = decode_delta_to_dmr(residual_z, sd, components)
    rows = pd.DataFrame({
        "cluster_name": matrix.columns,
        "latent_residual_delta_beta": latent_delta_dmr,
        "observed_minus_strict_pred_delta_beta": obs_delta_dmr,
        "strict_model_delta_beta_8cell_to_morula": model_delta_dmr,
        "abs_latent_residual_delta_beta": np.abs(latent_delta_dmr),
        "signed_latent_residual_direction": np.sign(latent_delta_dmr),
        "PC_loading_norm": np.linalg.norm(components, axis=1),
    })
    for i in range(components.shape[1]):
        rows[f"PC{i + 1}_loading"] = components[:, i]
        rows[f"residual_contribution_PC{i + 1}"] = residual_z[i] * components[:, i]
    rows["residual_alignment_score"] = rows["latent_residual_delta_beta"] * rows["observed_minus_strict_pred_delta_beta"]
    rows["abs_residual_alignment_score"] = np.abs(rows["residual_alignment_score"])
    rows = rows.merge(modules, on="cluster_name", how="left")
    rows = rows.merge(metadata, on="cluster_name", how="left")
    rows = rows.sort_values("abs_latent_residual_delta_beta", ascending=False).reset_index(drop=True)
    rows["basin_residual_rank"] = np.arange(1, len(rows) + 1)
    return rows


def module_summary(dmr_table: pd.DataFrame, residual_z: np.ndarray, sd: np.ndarray, components: np.ndarray):
    rows = []
    total_abs = dmr_table["abs_latent_residual_delta_beta"].sum()
    for module_id, sub in dmr_table.groupby("module_id", dropna=False):
        mask = np.asarray(dmr_table.index.isin(sub.index), dtype=bool)
        delta = np.zeros(len(dmr_table), dtype=float)
        delta[mask] = dmr_table.loc[mask, "latent_residual_delta_beta"].to_numpy(dtype=float)
        z_part = encode_delta_to_latent(delta, sd, components)
        rows.append({
            "module_id": module_id if pd.notna(module_id) else "unassigned",
            "n_DMRs": int(len(sub)),
            "sum_abs_residual_delta_beta": float(sub["abs_latent_residual_delta_beta"].sum()),
            "fraction_abs_residual_delta_beta": float(sub["abs_latent_residual_delta_beta"].sum() / total_abs) if total_abs else 0.0,
            "mean_signed_residual_delta_beta": float(sub["latent_residual_delta_beta"].mean()),
            "module_latent_residual_norm": float(np.linalg.norm(z_part)),
            "module_latent_residual_cosine_to_missing_direction": cosine(z_part, residual_z),
            "top_DMR": str(sub.sort_values("abs_latent_residual_delta_beta", ascending=False).iloc[0]["cluster_name"]),
        })
    return pd.DataFrame(rows).sort_values(["fraction_abs_residual_delta_beta", "module_latent_residual_norm"], ascending=False)


def cosine(a, b) -> float:
    a = np.asarray(a, dtype=float)
    b = np.asarray(b, dtype=float)
    den = np.linalg.norm(a) * np.linalg.norm(b)
    return float((a @ b) / den) if den > 0 else float("nan")


def overlap_summary(candidate: pd.DataFrame, named_tables: dict[str, pd.DataFrame], universe_n: int):
    rows = []
    cand = set(candidate["cluster_name"])
    for name, table in named_tables.items():
        if table.empty or "cluster_name" not in table.columns:
            continue
        target = set(table["cluster_name"])
        k = len(cand & target)
        rows.append({
            "candidate_set": "top_basin_residual_DMRs",
            "comparison_set": name,
            "candidate_n": len(cand),
            "comparison_n": len(target),
            "overlap_n": k,
            "overlap_fraction_of_candidate": float(k / len(cand)) if cand else 0.0,
            "overlap_fraction_of_comparison": float(k / len(target)) if target else 0.0,
            "expected_overlap_random": float(len(cand) * len(target) / universe_n) if universe_n else float("nan"),
        })
    return pd.DataFrame(rows)


def control_metrics(score_df, ann, matrix, mu, sd, components, strict_pred_z, obs_z, obs_dmr, basin, residual_z, dmr_table, rng):
    rows = []
    controls = []
    base_meta = {
        "control_family": "latent_oracle",
        "control_name": "full_missing_basin_direction",
        "uses_morula_residual": "yes_diagnostic",
        "n_control_DMRs": matrix.shape[1],
        "latent_control_cosine_to_missing_direction": 1.0,
    }
    for alpha in [0.0, 0.25, 0.5, 0.75, 1.0, 1.25]:
        controls.append((f"latent_oracle_alpha_{alpha:g}", strict_pred_z + alpha * residual_z[None, :], {**base_meta, "control_alpha": alpha}))

    for n in [10, 25, 50, 100]:
        selected = dmr_table.head(n)
        delta = np.zeros(matrix.shape[1], dtype=float)
        selected_idx = [list(matrix.columns).index(c) for c in selected["cluster_name"]]
        delta[selected_idx] = dmr_table.set_index("cluster_name").loc[selected["cluster_name"], "latent_residual_delta_beta"].to_numpy(dtype=float)
        z_part = encode_delta_to_latent(delta, sd, components)
        controls.append((
            f"top{n}_DMR_restricted_residual_control",
            strict_pred_z + z_part[None, :],
            {
                "control_family": "DMR_restricted",
                "control_name": f"top{n}_DMR_restricted_residual_control",
                "uses_morula_residual": "yes_diagnostic",
                "control_alpha": 1.0,
                "n_control_DMRs": n,
                "latent_control_norm": float(np.linalg.norm(z_part)),
                "latent_control_cosine_to_missing_direction": cosine(z_part, residual_z),
            },
        ))

    module_order = (
        module_summary(dmr_table, residual_z, sd, components)
        .sort_values("module_latent_residual_cosine_to_missing_direction", ascending=False)
        ["module_id"]
        .tolist()
    )
    for k in [1, 2, 3, 5]:
        selected_modules = module_order[:k]
        sub = dmr_table[dmr_table["module_id"].isin(selected_modules)]
        delta = np.zeros(matrix.shape[1], dtype=float)
        selected_idx = [list(matrix.columns).index(c) for c in sub["cluster_name"]]
        delta[selected_idx] = dmr_table.set_index("cluster_name").loc[sub["cluster_name"], "latent_residual_delta_beta"].to_numpy(dtype=float)
        z_part = encode_delta_to_latent(delta, sd, components)
        controls.append((
            f"top{k}_module_restricted_residual_control",
            strict_pred_z + z_part[None, :],
            {
                "control_family": "module_restricted",
                "control_name": f"top{k}_module_restricted_residual_control",
                "uses_morula_residual": "yes_diagnostic",
                "control_alpha": 1.0,
                "n_control_DMRs": int(len(sub)),
                "control_modules": ",".join(map(str, selected_modules)),
                "latent_control_norm": float(np.linalg.norm(z_part)),
                "latent_control_cosine_to_missing_direction": cosine(z_part, residual_z),
            },
        ))

    for name, pred_z, meta in controls:
        pred_dmr = decode_latent(pred_z, mu, sd, components)
        metrics = distribution_metrics(pred_z, obs_z, pred_dmr, obs_dmr, basin, rng)
        rows.append({
            "control_model": name,
            "dmr_mean_rmse_vs_observed_morula": rmse(pred_dmr.mean(axis=0), obs_dmr.mean(axis=0)),
            "dmr_correlation_vs_observed_morula": corr(pred_dmr.mean(axis=0), obs_dmr.mean(axis=0)),
            **meta,
            **metrics,
        })
    return pd.DataFrame(rows)


def write_bed(path: Path, dmr_table: pd.DataFrame, n: int):
    cols = ["chr", "start", "end", "cluster_name", "abs_latent_residual_delta_beta", "signed_latent_residual_direction"]
    bed = dmr_table.dropna(subset=["chr", "start", "end"]).head(n).copy()
    bed["score"] = (1000 * bed["abs_latent_residual_delta_beta"] / bed["abs_latent_residual_delta_beta"].max()).round(0).astype(int)
    bed["strand"] = "."
    out = bed[["chr", "start", "end", "cluster_name", "score", "strand"]]
    out.to_csv(path, sep="\t", index=False, header=False)


def make_svg(path: Path, module_df: pd.DataFrame):
    rows = module_df.head(12).copy()
    width, height = 920, 430
    left, right, top, bottom = 80, 30, 45, 105
    plot_w = width - left - right
    plot_h = height - top - bottom
    vmax = rows["fraction_abs_residual_delta_beta"].max() * 1.15 if len(rows) else 1
    gap = plot_w / max(len(rows), 1)
    bar_w = gap * 0.62
    out = [
        f'<svg xmlns="http://www.w3.org/2000/svg" width="{width}" height="{height}" viewBox="0 0 {width} {height}">',
        '<rect width="100%" height="100%" fill="white"/>',
        f'<text x="{left}" y="28" font-family="Arial" font-size="19" font-weight="700">DMR modules carrying the missing morula basin direction</text>',
        f'<line x1="{left}" y1="{height-bottom}" x2="{width-right}" y2="{height-bottom}" stroke="#222"/>',
        f'<line x1="{left}" y1="{top}" x2="{left}" y2="{height-bottom}" stroke="#222"/>',
    ]
    for i, row in enumerate(rows.itertuples()):
        x = left + i * gap + (gap - bar_w) / 2
        val = float(row.fraction_abs_residual_delta_beta)
        h = val / vmax * plot_h
        y = height - bottom - h
        fill = "#2f6f8f" if row.module_latent_residual_cosine_to_missing_direction >= 0 else "#9b6b35"
        out.append(f'<rect x="{x:.2f}" y="{y:.2f}" width="{bar_w:.2f}" height="{h:.2f}" fill="{fill}"/>')
        out.append(f'<text x="{x+bar_w/2:.2f}" y="{y-5:.2f}" text-anchor="middle" font-family="Arial" font-size="10">{val:.3f}</text>')
        out.append(f'<text x="{x+bar_w/2:.2f}" y="{height-bottom+18}" text-anchor="end" transform="rotate(-40 {x+bar_w/2:.2f} {height-bottom+18})" font-family="Arial" font-size="11">{row.module_id}</text>')
    out.append(f'<text x="{left}" y="{height-24}" font-family="Arial" font-size="12">Height: fraction of absolute decoded residual. Blue modules align positively with the missing latent direction.</text>')
    out.append("</svg>")
    path.write_text("\n".join(out), encoding="utf-8")


def write_doc(path: Path, residual_summary: dict, module_df: pd.DataFrame, control_df: pd.DataFrame, overlap_df: pd.DataFrame):
    best_control = control_df.sort_values("pred_basin_occupancy_q90", ascending=False).iloc[0]
    top_modules = module_df.head(5)
    lines = [
        "# Basin residual control-field diagnostic",
        "",
        "This analysis treats the strict non-leaking methylation-only failure as a missing control-field problem. It defines the missing morula basin direction in latent space and decodes that direction back to DMR and module space.",
        "",
        f"Observed morula center: {', '.join(f'{v:.4f}' for v in residual_summary['observed_morula_center'])}",
        f"Strict pre-morula predicted center: {', '.join(f'{v:.4f}' for v in residual_summary['strict_predicted_morula_center'])}",
        f"Missing basin residual vector: {', '.join(f'{v:.4f}' for v in residual_summary['missing_basin_residual_vector'])}",
        f"Residual norm: {residual_summary['missing_basin_residual_norm']:.4f}",
        "",
        "Top modules carrying the decoded residual:",
    ]
    for row in top_modules.itertuples():
        lines.append(
            f"- {row.module_id}: fraction_abs_residual={row.fraction_abs_residual_delta_beta:.3f}, "
            f"cosine={row.module_latent_residual_cosine_to_missing_direction:.3f}, top_DMR={row.top_DMR}"
        )
    lines.extend(["", "Diagnostic control results:"])
    for row in control_df.sort_values("pred_basin_occupancy_q90", ascending=False).head(8).itertuples():
        lines.append(
            f"- {row.control_model}: occupancy_q90={row.pred_basin_occupancy_q90:.3f}, "
            f"DMR_mean_RMSE={row.dmr_mean_rmse_vs_observed_morula:.4f}, cosine={getattr(row, 'latent_control_cosine_to_missing_direction', float('nan')):.3f}"
        )
    if len(overlap_df):
        lines.extend(["", "Overlap with existing dynamic DMR sets:"])
        for row in overlap_df.itertuples():
            lines.append(
                f"- {row.comparison_set}: overlap={row.overlap_n}/{row.candidate_n}, expected_random={row.expected_overlap_random:.1f}"
            )
    lines.extend([
        "",
        f"Best diagnostic control occupancy is {best_control.pred_basin_occupancy_q90:.3f}. These controls use the observed morula residual and are therefore not strict predictors. Their purpose is to identify candidate DMR modules and genomic regions that could carry a missing regulatory/chromatin basin-attraction field.",
        "",
        "Interpretation boundary: this does not prove that the listed DMRs are causal or that the hidden field is already measured. It provides a prioritized target set for RNA, ATAC, histone-mark, TF motif, DNMT/TET, and nearby-gene annotation.",
    ])
    path.write_text("\n".join(lines) + "\n", encoding="utf-8")


def main():
    parser = argparse.ArgumentParser()
    parser.add_argument("--q", type=int, default=3)
    parser.add_argument("--lambda", dest="lam", type=float, default=1000.0)
    parser.add_argument("--n-steps", type=int, default=12)
    parser.add_argument("--top-bed", type=int, default=100)
    parser.add_argument("--seed", type=int, default=20260525)
    args = parser.parse_args()

    RESULTS.mkdir(parents=True, exist_ok=True)
    FIGURES.mkdir(parents=True, exist_ok=True)
    DOCS.mkdir(parents=True, exist_ok=True)

    matrix, ann, pairs = load_inputs()
    mu, sd, components, score_df = fit_latent_basis(matrix, args.q)
    coef, cov, train = fit_operator(score_df, pairs, train_stages=set(PRE_MORULA_STAGES), lam=args.lam)

    strict_pred_z = simulate_strict_pre_morula(score_df, ann, coef, args.n_steps)
    strict_center = strict_pred_z.mean(axis=0)
    obs_z = score_df.loc[stage_ids(ann, "morula")].to_numpy(dtype=float)
    obs_center = obs_z.mean(axis=0)
    residual_z = obs_center - strict_center

    obs_dmr = matrix.loc[stage_ids(ann, "morula")].to_numpy(dtype=float)
    dmr8 = matrix.loc[stage_ids(ann, "8-cell")].to_numpy(dtype=float).mean(axis=0)
    strict_pred_dmr = decode_latent(strict_pred_z, mu, sd, components)
    obs_delta_dmr = obs_dmr.mean(axis=0) - strict_pred_dmr.mean(axis=0)
    model_delta_dmr = strict_pred_dmr.mean(axis=0) - dmr8
    basin = basin_definition(obs_z)

    metadata = pd.read_csv(RESULTS / "CSB_TRO_DMR_metadata.tsv", sep="\t")
    modules = pd.read_csv(RESULTS / "CSB_TRO_DMR_module_assignments.tsv", sep="\t")
    dmr_table = build_dmr_residual_table(matrix, metadata, modules, components, sd, residual_z, obs_delta_dmr, model_delta_dmr)
    module_df = module_summary(dmr_table, residual_z, sd, components)
    controls = control_metrics(score_df, ann, matrix, mu, sd, components, strict_pred_z, obs_z, obs_dmr, basin, residual_z, dmr_table, np.random.default_rng(args.seed))

    named = {
        "top100_morula_entry_DMRs": load_optional_table(RESULTS / "CSB_TRO_top100_morula_entry_DMRs.tsv"),
        "top100_dynamic_reset_DMRs": load_optional_table(RESULTS / "CSB_TRO_top100_dynamic_reset_DMRs.tsv"),
        "top100_blastocyst_exit_DMRs": load_optional_table(RESULTS / "CSB_TRO_top100_blastocyst_exit_DMRs.tsv"),
        "top100_latent_loading_DMRs": load_optional_table(RESULTS / "CSB_TRO_latent_loading_DMR_ranking.tsv").head(100),
    }
    overlap = overlap_summary(dmr_table.head(100), named, universe_n=matrix.shape[1])

    residual_summary = {
        "q": args.q,
        "ridge_lambda": args.lam,
        "n_steps": args.n_steps,
        "n_pre_morula_training_pairs": int(len(train)),
        "observed_morula_center": obs_center.tolist(),
        "strict_predicted_morula_center": strict_center.tolist(),
        "missing_basin_residual_vector": residual_z.tolist(),
        "missing_basin_residual_norm": float(np.linalg.norm(residual_z)),
        "decoded_DMR_residual_norm": float(np.linalg.norm(dmr_table["latent_residual_delta_beta"].to_numpy(dtype=float))),
        "strict_predicted_basin_occupancy_q90": float(np.mean(np.linalg.norm(strict_pred_z - obs_center[None, :], axis=1) <= basin["radius_q90"])),
        "observed_basin_occupancy_q90": float(basin["observed_occupancy_q90"]),
    }

    dmr_table.to_csv(RESULTS / "CSB_TRO_basin_residual_DMR_ranking.tsv", sep="\t", index=False)
    module_df.to_csv(RESULTS / "CSB_TRO_basin_residual_module_summary.tsv", sep="\t", index=False)
    controls.to_csv(RESULTS / "CSB_TRO_basin_residual_control_metrics.tsv", sep="\t", index=False)
    overlap.to_csv(RESULTS / "CSB_TRO_basin_residual_overlap_summary.tsv", sep="\t", index=False)
    pd.DataFrame([residual_summary]).to_csv(RESULTS / "CSB_TRO_basin_residual_direction_summary.tsv", sep="\t", index=False)
    write_bed(RESULTS / "CSB_TRO_basin_residual_topDMRs.bed", dmr_table, args.top_bed)
    make_svg(FIGURES / "CSB_TRO_basin_residual_module_contributions.svg", module_df)
    write_doc(DOCS / "CSB_TRO_basin_residual_control_field_interpretation.md", residual_summary, module_df, controls, overlap)

    manifest = {
        **residual_summary,
        "outputs": [
            str(RESULTS / "CSB_TRO_basin_residual_DMR_ranking.tsv"),
            str(RESULTS / "CSB_TRO_basin_residual_module_summary.tsv"),
            str(RESULTS / "CSB_TRO_basin_residual_control_metrics.tsv"),
            str(RESULTS / "CSB_TRO_basin_residual_overlap_summary.tsv"),
            str(RESULTS / "CSB_TRO_basin_residual_topDMRs.bed"),
            str(FIGURES / "CSB_TRO_basin_residual_module_contributions.svg"),
            str(DOCS / "CSB_TRO_basin_residual_control_field_interpretation.md"),
        ],
    }
    (RESULTS / "CSB_TRO_basin_residual_control_field_manifest.json").write_text(json.dumps(manifest, indent=2), encoding="utf-8")
    print(json.dumps({
        "summary": residual_summary,
        "top_modules": module_df.head(8).to_dict(orient="records"),
        "top_controls": controls.sort_values("pred_basin_occupancy_q90", ascending=False).head(8).to_dict(orient="records"),
        "overlap": overlap.to_dict(orient="records"),
    }, indent=2))


if __name__ == "__main__":
    main()
```
