## Supplemental code for "An entropy-based diagnostic framework for characterizing methylation state dynamics during preimplantation development": run_bifurcation_like_basin_entry_scan.py

```
from __future__ import annotations

import json
import sys
from pathlib import Path

import numpy as np
import pandas as pd


BASE = Path(r"E:\CSB_TRO_operator_time_DMR_dynamics_2026-05-25")
CODE = BASE / "code"
RESULTS = BASE / "results"
FIGURES = BASE / "figures"
DOCS = BASE / "docs"
sys.path.insert(0, str(CODE))

from run_basin_residual_control_field import PRE_MORULA_STAGES, cosine, simulate_strict_pre_morula  # noqa: E402
from run_morula_basin_sde import (  # noqa: E402
    basin_definition,
    decode_latent,
    distribution_metrics,
    fit_latent_basis,
    fit_operator,
    load_inputs,
    stage_ids,
)


OUT_ALPHA = RESULTS / "CSB_TRO_alpha_bifurcation_scan.tsv"
OUT_MODULE = RESULTS / "CSB_TRO_module_specific_bifurcation_scan.tsv"
OUT_LOCAL = RESULTS / "CSB_TRO_local_jacobian_eigenvalues.tsv"
OUT_SUMMARY = RESULTS / "CSB_TRO_bifurcation_like_scan_summary.tsv"
OUT_MANIFEST = RESULTS / "CSB_TRO_bifurcation_like_scan_manifest.json"
OUT_ALPHA_SVG = FIGURES / "CSB_TRO_alpha_bifurcation_scan.svg"
OUT_MODULE_SVG = FIGURES / "CSB_TRO_module_specific_bifurcation_scan.svg"
OUT_DOC = DOCS / "CSB_TRO_bifurcation_like_basin_entry_summary.md"

PRIORITY_MODULES = ["M05", "M01", "M12", "M02", "M10"]
MODULE_COMBOS = [
    ("M05", ["M05"]),
    ("M01", ["M01"]),
    ("M12", ["M12"]),
    ("M02", ["M02"]),
    ("M10", ["M10"]),
    ("M05+M01", ["M05", "M01"]),
    ("M05+M01+M12", ["M05", "M01", "M12"]),
    ("M05+M01+M12+M02", ["M05", "M01", "M12", "M02"]),
    ("M05+M01+M12+M02+M10", ["M05", "M01", "M12", "M02", "M10"]),
]


def alpha_values() -> list[float]:
    return [round(float(x), 2) for x in np.arange(0.0, 1.5001, 0.05)]


def cov_stats(z: np.ndarray) -> dict[str, float]:
    cov = np.cov(np.asarray(z, dtype=float).T)
    eig = np.linalg.eigvalsh(cov)
    eig = np.maximum(eig, 0.0)
    return {
        "latent_variance_trace": float(np.trace(cov)),
        "covariance_eigen_min": float(eig.min()),
        "covariance_eigen_max": float(eig.max()),
        "covariance_anisotropy": float(eig.max() / (eig.min() + 1e-12)),
    }


def radial_stats(strict_z: np.ndarray, pred_z: np.ndarray, center: np.ndarray) -> dict[str, float]:
    before = np.linalg.norm(strict_z - center[None, :], axis=1)
    after = np.linalg.norm(pred_z - center[None, :], axis=1)
    return {
        "mean_radial_distance_before": float(before.mean()),
        "mean_radial_distance_after": float(after.mean()),
        "radial_distance_reduction": float(before.mean() - after.mean()),
        "fraction_particles_moved_closer": float(np.mean(after < before)),
    }


def evaluate_scan_row(
    scan_type: str,
    control_name: str,
    alpha: float,
    control_z: np.ndarray,
    strict_z: np.ndarray,
    obs_z: np.ndarray,
    obs_dmr: np.ndarray,
    mu: np.ndarray,
    sd: np.ndarray,
    components: np.ndarray,
    basin: dict[str, object],
    residual_z: np.ndarray,
    rng: np.random.Generator,
    extra: dict[str, object] | None = None,
) -> dict[str, object]:
    pred_z = strict_z + alpha * control_z[None, :]
    pred_dmr = decode_latent(pred_z, mu, sd, components)
    row = {
        "scan_type": scan_type,
        "control_name": control_name,
        "alpha": float(alpha),
        "control_PC1": float(alpha * control_z[0]),
        "control_PC2": float(alpha * control_z[1]),
        "control_PC3": float(alpha * control_z[2]),
        "control_norm": float(np.linalg.norm(alpha * control_z)),
        "control_cosine_to_measured_correction": cosine(control_z, residual_z),
        "PC3_negative_pull_recovery": float(-(alpha * control_z[2]) / (-residual_z[2])) if residual_z[2] < 0 else float("nan"),
    }
    row.update(distribution_metrics(pred_z, obs_z, pred_dmr, obs_dmr, basin, rng))
    row.update(cov_stats(pred_z))
    row.update(radial_stats(strict_z, pred_z, np.asarray(basin["center"], dtype=float)))
    if extra:
        row.update(extra)
    return row


def module_control_vectors() -> dict[str, np.ndarray]:
    basis = pd.read_csv(RESULTS / "CSB_TRO_missing_control_term_module_basis.tsv", sep="\t")
    vecs = {}
    for row in basis.itertuples():
        vecs[str(row.module_id)] = np.array([row.latent_control_PC1, row.latent_control_PC2, row.latent_control_PC3], dtype=float)
    return vecs


def threshold_summary(df: pd.DataFrame, target: float) -> dict[str, object]:
    out = {}
    for name, sub in df.groupby("control_name"):
        sub = sub.sort_values("alpha")
        occ = sub["pred_basin_occupancy_q90"].to_numpy(dtype=float)
        alphas = sub["alpha"].to_numpy(dtype=float)
        reached = sub.loc[sub["pred_basin_occupancy_q90"] >= target]
        alpha_target = float(reached.iloc[0]["alpha"]) if len(reached) else float("nan")
        slope = np.diff(occ) / np.diff(alphas) if len(alphas) > 1 else np.array([float("nan")])
        max_i = int(np.nanargmax(slope)) if np.any(np.isfinite(slope)) else 0
        out[name] = {
            "alpha_to_observed_occupancy": alpha_target,
            "max_local_slope": float(slope[max_i]) if len(slope) else float("nan"),
            "alpha_at_max_slope_left": float(alphas[max_i]) if len(alphas) else float("nan"),
            "occupancy_at_alpha_0": float(occ[0]) if len(occ) else float("nan"),
            "occupancy_at_alpha_1": float(sub.iloc[(sub["alpha"] - 1.0).abs().argsort()].iloc[0]["pred_basin_occupancy_q90"]) if len(sub) else float("nan"),
        }
    return out


def linear_jacobian(coef: np.ndarray) -> pd.DataFrame:
    # fit_operator uses x=[1,tau,z1,z2,z3], so dz/dtau = intercept + tau term + z @ A.
    jac = coef[2:, :]
    eig = np.linalg.eigvals(jac)
    rows = []
    for i, val in enumerate(eig, start=1):
        rows.append(
            {
                "model": "methylation_only_affine_operator",
                "eigen_index": i,
                "real": float(np.real(val)),
                "imag": float(np.imag(val)),
                "note": "Constant measured correction changes basin entry in this diagnostic scan but does not change the affine methylation-only Jacobian.",
            }
        )
    return pd.DataFrame(rows)


def make_alpha_svg(path: Path, df: pd.DataFrame, title: str) -> None:
    sub = df[df["control_name"] == "measured_full_correction"].sort_values("alpha")
    width, height = 860, 430
    left, right, top, bottom = 70, 30, 45, 70
    plot_w = width - left - right
    plot_h = height - top - bottom
    xmin, xmax = float(sub["alpha"].min()), float(sub["alpha"].max())
    points = []
    for row in sub.itertuples():
        x = left + (row.alpha - xmin) / (xmax - xmin) * plot_w
        y = height - bottom - float(row.pred_basin_occupancy_q90) * plot_h
        points.append((x, y))
    lines = [
        f'<svg xmlns="http://www.w3.org/2000/svg" width="{width}" height="{height}" viewBox="0 0 {width} {height}">',
        '<rect width="100%" height="100%" fill="white"/>',
        f'<text x="{left}" y="28" font-family="Arial" font-size="18" font-weight="700">{title}</text>',
        f'<line x1="{left}" y1="{height-bottom}" x2="{width-right}" y2="{height-bottom}" stroke="#222"/>',
        f'<line x1="{left}" y1="{top}" x2="{left}" y2="{height-bottom}" stroke="#222"/>',
    ]
    for tick in [0, 0.25, 0.5, 0.75, 0.875, 1.0]:
        y = height - bottom - tick * plot_h
        color = "#b23b3b" if abs(tick - 0.875) < 1e-6 else "#dddddd"
        lines.append(f'<line x1="{left}" y1="{y:.2f}" x2="{width-right}" y2="{y:.2f}" stroke="{color}" stroke-dasharray="4 4"/>')
        lines.append(f'<text x="{left-8}" y="{y+4:.2f}" text-anchor="end" font-family="Arial" font-size="11">{tick:.3g}</text>')
    lines.append('<polyline points="' + " ".join(f"{x:.2f},{y:.2f}" for x, y in points) + '" fill="none" stroke="#245c7a" stroke-width="2.5"/>')
    for x, y in points:
        lines.append(f'<circle cx="{x:.2f}" cy="{y:.2f}" r="2.8" fill="#245c7a"/>')
    lines.append(f'<text x="{left}" y="{height-22}" font-family="Arial" font-size="12">Occupancy q90 vs alpha. Red guide is observed morula occupancy.</text>')
    lines.append("</svg>")
    path.write_text("\n".join(lines), encoding="utf-8")


def make_module_svg(path: Path, df: pd.DataFrame) -> None:
    chosen = ["M05", "M05+M01", "M05+M01+M12", "M05+M01+M12+M02", "M05+M01+M12+M02+M10"]
    colors = ["#7a3b2e", "#b06b2f", "#6f8f3a", "#245c7a", "#5d4c8c"]
    width, height = 930, 460
    left, right, top, bottom = 80, 160, 45, 70
    plot_w = width - left - right
    plot_h = height - top - bottom
    lines = [
        f'<svg xmlns="http://www.w3.org/2000/svg" width="{width}" height="{height}" viewBox="0 0 {width} {height}">',
        '<rect width="100%" height="100%" fill="white"/>',
        f'<text x="{left}" y="28" font-family="Arial" font-size="18" font-weight="700">Module-specific bifurcation-like scan</text>',
        f'<line x1="{left}" y1="{height-bottom}" x2="{width-right}" y2="{height-bottom}" stroke="#222"/>',
        f'<line x1="{left}" y1="{top}" x2="{left}" y2="{height-bottom}" stroke="#222"/>',
    ]
    for tick in [0, 0.25, 0.5, 0.75, 0.875, 1.0]:
        y = height - bottom - tick * plot_h
        color = "#b23b3b" if abs(tick - 0.875) < 1e-6 else "#dddddd"
        lines.append(f'<line x1="{left}" y1="{y:.2f}" x2="{width-right}" y2="{y:.2f}" stroke="{color}" stroke-dasharray="4 4"/>')
        lines.append(f'<text x="{left-8}" y="{y+4:.2f}" text-anchor="end" font-family="Arial" font-size="11">{tick:.3g}</text>')
    xmin, xmax = float(df["alpha"].min()), float(df["alpha"].max())
    for idx, name in enumerate(chosen):
        sub = df[df["control_name"] == name].sort_values("alpha")
        pts = []
        for row in sub.itertuples():
            x = left + (row.alpha - xmin) / (xmax - xmin) * plot_w
            y = height - bottom - float(row.pred_basin_occupancy_q90) * plot_h
            pts.append((x, y))
        color = colors[idx]
        lines.append('<polyline points="' + " ".join(f"{x:.2f},{y:.2f}" for x, y in pts) + f'" fill="none" stroke="{color}" stroke-width="2.3"/>')
        lines.append(f'<rect x="{width-right+15}" y="{top+idx*24}" width="12" height="12" fill="{color}"/>')
        lines.append(f'<text x="{width-right+33}" y="{top+idx*24+11}" font-family="Arial" font-size="12">{name}</text>')
    lines.append(f'<text x="{left}" y="{height-22}" font-family="Arial" font-size="12">Diagnostic module controls are derived from measured residual module decomposition.</text>')
    lines.append("</svg>")
    path.write_text("\n".join(lines), encoding="utf-8")


def write_doc(alpha_df: pd.DataFrame, module_df: pd.DataFrame, summary_rows: pd.DataFrame) -> None:
    full = summary_rows[summary_rows["control_name"] == "measured_full_correction"].iloc[0]
    module_top = summary_rows[summary_rows["scan_type"] == "module_specific"].sort_values("occupancy_at_alpha_1", ascending=False).head(5)
    lines = [
        "# Bifurcation-like Basin Entry Scan",
        "",
        "Status: `completed`",
        "",
        "This analysis treats the measured missing correction term as a diagnostic control parameter and asks whether morula basin entry shows threshold-like behavior.",
        "",
        "Important boundary: this supports a bifurcation-like or tipping-like basin-entry interpretation, not a proof of a strict saddle-node bifurcation.",
        "",
        "## Full Correction Scan",
        "",
        f"- Occupancy at alpha=0: {full.occupancy_at_alpha_0:.3f}",
        f"- Occupancy at alpha=1: {full.occupancy_at_alpha_1:.3f}",
        f"- First alpha reaching observed q90 occupancy: {full.alpha_to_observed_occupancy:.2f}",
        f"- Steepest local occupancy slope begins near alpha: {full.alpha_at_max_slope_left:.2f}",
        "",
        "## Module Scan",
        "",
    ]
    for row in module_top.itertuples():
        lines.append(
            f"- {row.control_name}: occupancy alpha=1 is {row.occupancy_at_alpha_1:.3f}; "
            f"first alpha to observed occupancy is {row.alpha_to_observed_occupancy if np.isfinite(row.alpha_to_observed_occupancy) else 'not_reached'}"
        )
    lines += [
        "",
        "## Jacobian Boundary",
        "",
        "The current methylation operator is affine in latent z, and the measured correction is added as a constant control vector. Therefore the diagnostic alpha scan changes endpoint basin entry but does not by itself change the affine methylation-only Jacobian. A future biological u_bio(z,tau) or basin-coupled control model is needed to test true local stability changes.",
    ]
    OUT_DOC.write_text("\n".join(lines) + "\n", encoding="utf-8")


def main() -> None:
    RESULTS.mkdir(parents=True, exist_ok=True)
    FIGURES.mkdir(parents=True, exist_ok=True)
    DOCS.mkdir(parents=True, exist_ok=True)

    matrix, ann, pairs = load_inputs()
    mu, sd, components, score_df = fit_latent_basis(matrix, 3)
    coef, _, _ = fit_operator(score_df, pairs, train_stages=set(PRE_MORULA_STAGES), lam=1000.0)
    strict_z = simulate_strict_pre_morula(score_df, ann, coef, n_steps=12)
    obs_z = score_df.loc[stage_ids(ann, "morula")].to_numpy(dtype=float)
    obs_dmr = matrix.loc[stage_ids(ann, "morula")].to_numpy(dtype=float)
    basin = basin_definition(obs_z)
    residual_z = obs_z.mean(axis=0) - strict_z.mean(axis=0)

    rng = np.random.default_rng(20260526)
    alpha_rows = [
        evaluate_scan_row(
            "full_measured_correction",
            "measured_full_correction",
            alpha,
            residual_z,
            strict_z,
            obs_z,
            obs_dmr,
            mu,
            sd,
            components,
            basin,
            residual_z,
            rng,
        )
        for alpha in alpha_values()
    ]
    alpha_df = pd.DataFrame(alpha_rows)

    vecs = module_control_vectors()
    module_rows = []
    for combo_name, modules in MODULE_COMBOS:
        control = np.sum([vecs[m] for m in modules if m in vecs], axis=0)
        for alpha in alpha_values():
            module_rows.append(
                evaluate_scan_row(
                    "module_specific",
                    combo_name,
                    alpha,
                    control,
                    strict_z,
                    obs_z,
                    obs_dmr,
                    mu,
                    sd,
                    components,
                    basin,
                    residual_z,
                    rng,
                    {"modules": ",".join(modules), "n_modules": len(modules)},
                )
            )
    module_df = pd.DataFrame(module_rows)

    alpha_thresholds = threshold_summary(alpha_df, float(basin["observed_occupancy_q90"]))
    module_thresholds = threshold_summary(module_df, float(basin["observed_occupancy_q90"]))
    summary_rows = []
    for name, rec in {**alpha_thresholds, **module_thresholds}.items():
        scan_type = "full_measured_correction" if name in alpha_thresholds else "module_specific"
        summary_rows.append({"scan_type": scan_type, "control_name": name, **rec})
    summary_df = pd.DataFrame(summary_rows)

    jac = linear_jacobian(coef)
    alpha_df.to_csv(OUT_ALPHA, sep="\t", index=False)
    module_df.to_csv(OUT_MODULE, sep="\t", index=False)
    jac.to_csv(OUT_LOCAL, sep="\t", index=False)
    summary_df.to_csv(OUT_SUMMARY, sep="\t", index=False)
    make_alpha_svg(OUT_ALPHA_SVG, alpha_df, "Full correction alpha scan")
    make_module_svg(OUT_MODULE_SVG, module_df)
    write_doc(alpha_df, module_df, summary_df)

    manifest = {
        "status": "completed",
        "interpretation_boundary": "bifurcation-like/tipping-like basin-entry support; not strict saddle-node proof",
        "outputs": [str(OUT_ALPHA), str(OUT_MODULE), str(OUT_LOCAL), str(OUT_SUMMARY), str(OUT_ALPHA_SVG), str(OUT_MODULE_SVG), str(OUT_DOC)],
    }
    OUT_MANIFEST.write_text(json.dumps(manifest, indent=2) + "\n", encoding="utf-8")
    print(json.dumps({"status": "completed", "summary": summary_df.to_dict(orient="records")[:10]}, indent=2))


if __name__ == "__main__":
    main()
```
