## Supplemental code for "An entropy-based diagnostic framework for characterizing methylation state dynamics during preimplantation development": run_biological_control_augmented_dynamics.py

````
from __future__ import annotations

import argparse
import json
import sys
from pathlib import Path

import numpy as np
import pandas as pd


BASE = Path(r"E:\CSB_TRO_operator_time_DMR_dynamics_2026-05-25")
CODE = BASE / "code"
RESULTS = BASE / "results"
FIGURES = BASE / "figures"
DOCS = BASE / "docs"
sys.path.insert(0, str(CODE))

from run_basin_residual_control_field import PRE_MORULA_STAGES, cosine, simulate_strict_pre_morula  # noqa: E402
from run_morula_basin_sde import (  # noqa: E402
    basin_definition,
    decode_latent,
    distribution_metrics,
    fit_latent_basis,
    fit_operator,
    load_inputs,
    stage_ids,
)


def ridge_beta(x: np.ndarray, y: np.ndarray, lam: float) -> np.ndarray:
    return np.linalg.solve(x.T @ x + lam * np.eye(x.shape[1]), x.T @ y)


def build_control_vector(feature_df: pd.DataFrame, beta_mode: str, residual_z: np.ndarray, lam: float, rng: np.random.Generator):
    rows = feature_df.copy()
    rows["u"] = pd.to_numeric(rows["control_value_z"], errors="coerce").fillna(0.0)
    direction = rows[["latent_control_PC1", "latent_control_PC2", "latent_control_PC3"]].to_numpy(dtype=float)
    x = direction * rows["u"].to_numpy(dtype=float)[:, None]
    if beta_mode == "unit":
        beta = np.ones(len(rows), dtype=float)
    elif beta_mode == "ridge_to_measured_correction":
        beta = ridge_beta(x.T, residual_z, lam)
    elif beta_mode == "random_signed":
        beta = rng.choice([-1.0, 1.0], size=len(rows))
    elif beta_mode == "shuffle_u_ridge_to_measured_correction":
        rows["u"] = rng.permutation(rows["u"].to_numpy(dtype=float))
        x = direction * rows["u"].to_numpy(dtype=float)[:, None]
        beta = ridge_beta(x.T, residual_z, lam)
    else:
        raise ValueError(f"Unknown beta mode: {beta_mode}")
    control = (x * beta[:, None]).sum(axis=0)
    rows["beta"] = beta
    rows["weighted_PC1"] = x[:, 0] * beta
    rows["weighted_PC2"] = x[:, 1] * beta
    rows["weighted_PC3"] = x[:, 2] * beta
    return control, rows


def evaluate_control(name, pred_z, obs_z, obs_dmr, mu, sd, components, basin, residual_z, meta, seed):
    pred_dmr = decode_latent(pred_z, mu, sd, components)
    metrics = distribution_metrics(pred_z, obs_z, pred_dmr, obs_dmr, basin, np.random.default_rng(seed))
    return {
        "control_model": name,
        "direction_cosine_to_measured_correction": cosine(meta["control_vector"], residual_z),
        "PC1_control": float(meta["control_vector"][0]),
        "PC2_control": float(meta["control_vector"][1]),
        "PC3_control": float(meta["control_vector"][2]),
        "PC3_negative_pull_recovered": float(-meta["control_vector"][2] / (-residual_z[2])) if residual_z[2] < 0 else float("nan"),
        "control_norm": float(np.linalg.norm(meta["control_vector"])),
        **{k: v for k, v in meta.items() if k != "control_vector"},
        **metrics,
    }


def feature_models(features: pd.DataFrame):
    models = []
    for modality, sub in features.groupby("control_modality"):
        if len(sub):
            leakage_text = ",".join(map(str, sub["leakage_status"]))
            if "uses_morula_methylation_residual" in leakage_text or "residual" in leakage_text:
                unit_status = "measured_residual_diagnostic_not_nonleaking"
            elif modality in {"RNA", "ATAC", "histone", "motif_activity"} or "user_supplied_external_feature" in leakage_text:
                unit_status = "external_feature_defined"
            else:
                unit_status = "internal_proxy_feature_defined_not_external_omics"
            models.append((f"{modality}_unit_beta", sub.copy(), "unit", unit_status))
            models.append((f"{modality}_ridge_beta_diagnostic", sub.copy(), "ridge_to_measured_correction", "uses_morula_methylation_residual_for_beta"))
    external = features[features["control_modality"].isin(["RNA", "ATAC", "histone", "motif_activity"])].copy()
    if len(external):
        models.append(("combined_external_unit_beta", external, "unit", "external_feature_defined"))
        models.append(("combined_external_ridge_beta_diagnostic", external, "ridge_to_measured_correction", "uses_morula_methylation_residual_for_beta"))
    all_non_missing = features.copy()
    models.append(("all_available_unit_beta", all_non_missing, "unit", "includes_internal_proxy_features"))
    models.append(("all_available_ridge_beta_diagnostic", all_non_missing, "ridge_to_measured_correction", "uses_morula_methylation_residual_for_beta"))
    return models


def make_svg(path: Path, metrics: pd.DataFrame):
    rows = metrics.sort_values("pred_basin_occupancy_q90", ascending=False).head(14)
    width, height = 980, 450
    left, right, top, bottom = 80, 25, 45, 140
    plot_w = width - left - right
    plot_h = height - top - bottom
    gap = plot_w / max(len(rows), 1)
    bar_w = gap * 0.65
    out = [
        f'<svg xmlns="http://www.w3.org/2000/svg" width="{width}" height="{height}" viewBox="0 0 {width} {height}">',
        '<rect width="100%" height="100%" fill="white"/>',
        f'<text x="{left}" y="28" font-family="Arial" font-size="18" font-weight="700">Candidate biological control models: occupancy rescue</text>',
        f'<line x1="{left}" y1="{height-bottom}" x2="{width-right}" y2="{height-bottom}" stroke="#222"/>',
        f'<line x1="{left}" y1="{top}" x2="{left}" y2="{height-bottom}" stroke="#222"/>',
    ]
    for tick in [0, 0.25, 0.5, 0.75, 0.875, 1.0]:
        y = height - bottom - tick * plot_h
        color = "#b33" if abs(tick - 0.875) < 1e-6 else "#ddd"
        out.append(f'<line x1="{left}" y1="{y:.2f}" x2="{width-right}" y2="{y:.2f}" stroke="{color}" stroke-dasharray="4 4"/>')
        out.append(f'<text x="{left-8}" y="{y+4:.2f}" text-anchor="end" font-family="Arial" font-size="11">{tick:.3g}</text>')
    for i, row in enumerate(rows.itertuples()):
        x = left + i * gap + (gap - bar_w) / 2
        val = float(row.pred_basin_occupancy_q90)
        h = val * plot_h
        y = height - bottom - h
        fill = "#2f6f8f" if row.validation_status == "feature_defined" else "#b56b2a"
        out.append(f'<rect x="{x:.2f}" y="{y:.2f}" width="{bar_w:.2f}" height="{h:.2f}" fill="{fill}"/>')
        out.append(f'<text x="{x+bar_w/2:.2f}" y="{y-5:.2f}" text-anchor="middle" font-family="Arial" font-size="10">{val:.3f}</text>')
        label = row.control_model.replace("_", " ")
        out.append(f'<text x="{x+bar_w/2:.2f}" y="{height-bottom+18}" text-anchor="end" transform="rotate(-45 {x+bar_w/2:.2f} {height-bottom+18})" font-family="Arial" font-size="9">{label}</text>')
    out.append(f'<text x="{left}" y="{height-20}" font-family="Arial" font-size="12">Blue: feature-defined beta. Orange: diagnostic beta fit to measured correction, not non-leaking.</text>')
    out.append("</svg>")
    path.write_text("\n".join(out), encoding="utf-8")


def write_doc(path: Path, metrics: pd.DataFrame, features: pd.DataFrame):
    lines = [
        "# Biological control-augmented dynamics",
        "",
        "This analysis evaluates candidate module-level biological control features in the form:",
        "",
        "```text",
        "dz/dtau = f_meth(z,tau) + sum_m beta_m u_m(tau) b_m",
        "```",
        "",
        "Current feature modalities and leakage status:",
    ]
    for modality, sub in features.groupby("control_modality"):
        lines.append(f"- {modality}: n={len(sub)}, statuses={','.join(sorted(set(map(str, sub['leakage_status']))))}")
    lines.extend(["", "Top model results:"])
    for row in metrics.sort_values("pred_basin_occupancy_q90", ascending=False).head(10).itertuples():
        lines.append(
            f"- {row.control_model}: occupancy={row.pred_basin_occupancy_q90:.3f}, cosine={row.direction_cosine_to_measured_correction:.3f}, "
            f"PC3_recovery={row.PC3_negative_pull_recovered:.3f}, status={row.validation_status}"
        )
    lines.extend([
        "",
        "Interpretation boundary: models whose beta was fit to the measured correction are diagnostic upper bounds. A real u_bio result requires external feature-defined controls that improve occupancy without using morula methylation to define beta or direction.",
    ])
    path.write_text("\n".join(lines) + "\n", encoding="utf-8")


def main():
    parser = argparse.ArgumentParser()
    parser.add_argument("--feature-table", default=str(RESULTS / "CSB_TRO_module_bio_features.tsv"))
    parser.add_argument("--q", type=int, default=3)
    parser.add_argument("--lambda", dest="lam", type=float, default=1000.0)
    parser.add_argument("--beta-lambda", type=float, default=1e-3)
    parser.add_argument("--n-random", type=int, default=200)
    parser.add_argument("--seed", type=int, default=20260525)
    args = parser.parse_args()

    RESULTS.mkdir(parents=True, exist_ok=True)
    FIGURES.mkdir(parents=True, exist_ok=True)
    DOCS.mkdir(parents=True, exist_ok=True)

    features = pd.read_csv(args.feature_table, sep="\t")
    matrix, ann, pairs = load_inputs()
    mu, sd, components, score_df = fit_latent_basis(matrix, args.q)
    coef, _, _ = fit_operator(score_df, pairs, train_stages=set(PRE_MORULA_STAGES), lam=args.lam)
    strict_pred_z = simulate_strict_pre_morula(score_df, ann, coef, n_steps=12)
    obs_z = score_df.loc[stage_ids(ann, "morula")].to_numpy(dtype=float)
    obs_dmr = matrix.loc[stage_ids(ann, "morula")].to_numpy(dtype=float)
    basin = basin_definition(obs_z)
    residual_z = obs_z.mean(axis=0) - strict_pred_z.mean(axis=0)

    rng = np.random.default_rng(args.seed)
    metric_rows = []
    coef_parts = []
    baseline_meta = {"control_vector": np.zeros(3), "validation_status": "baseline", "n_features": 0, "beta_mode": "none", "feature_modality": "none"}
    metric_rows.append(evaluate_control("methylation_only_strict_baseline", strict_pred_z, obs_z, obs_dmr, mu, sd, components, basin, residual_z, baseline_meta, args.seed))

    upper_meta = {"control_vector": residual_z, "validation_status": "measured_residual_upper_bound", "n_features": 0, "beta_mode": "measured", "feature_modality": "measured_residual"}
    metric_rows.append(evaluate_control("measured_missing_correction_upper_bound", strict_pred_z + residual_z[None, :], obs_z, obs_dmr, mu, sd, components, basin, residual_z, upper_meta, args.seed + 1))

    for name, sub, beta_mode, status in feature_models(features):
        control, coef_df = build_control_vector(sub, beta_mode, residual_z, args.beta_lambda, rng)
        metric_rows.append(evaluate_control(
            name,
            strict_pred_z + control[None, :],
            obs_z,
            obs_dmr,
            mu,
            sd,
            components,
            basin,
            residual_z,
            {
                "control_vector": control,
                "validation_status": status,
                "n_features": int(len(sub)),
                "beta_mode": beta_mode,
                "feature_modality": ",".join(sorted(set(map(str, sub["control_modality"])))),
            },
            args.seed + len(metric_rows) + 10,
        ))
        coef_df.insert(0, "control_model", name)
        coef_parts.append(coef_df)
        if beta_mode == "unit":
            metric_rows.append(evaluate_control(
                name + "_sign_flip",
                strict_pred_z - control[None, :],
                obs_z,
                obs_dmr,
                mu,
                sd,
                components,
                basin,
                residual_z,
                {
                    "control_vector": -control,
                    "validation_status": "sign_flip_control",
                    "n_features": int(len(sub)),
                    "beta_mode": beta_mode,
                    "feature_modality": ",".join(sorted(set(map(str, sub["control_modality"])))),
                },
                args.seed + len(metric_rows) + 100,
            ))

    # Random feature-value control for available features.
    random_rows = []
    base_features = features.copy()
    for i in range(args.n_random):
        sub = base_features.copy()
        sub["control_value_z"] = rng.permutation(sub["control_value_z"].fillna(0.0).to_numpy(dtype=float))
        control, _ = build_control_vector(sub, "unit", residual_z, args.beta_lambda, rng)
        row = evaluate_control(
            "random_shuffled_feature_unit_beta",
            strict_pred_z + control[None, :],
            obs_z,
            obs_dmr,
            mu,
            sd,
            components,
            basin,
            residual_z,
            {
                "control_vector": control,
                "validation_status": "random_feature_value_control",
                "n_features": int(len(sub)),
                "beta_mode": "unit",
                "feature_modality": "all_shuffled",
                "random_iter": i,
            },
            args.seed + 1000 + i,
        )
        random_rows.append(row)
    metrics = pd.DataFrame(metric_rows)
    random_metrics = pd.DataFrame(random_rows)
    coef_all = pd.concat(coef_parts, ignore_index=True) if coef_parts else pd.DataFrame()

    metrics.to_csv(RESULTS / "CSB_TRO_bio_control_occupancy_metrics.tsv", sep="\t", index=False)
    random_metrics.to_csv(RESULTS / "CSB_TRO_bio_control_matched_random_features.tsv", sep="\t", index=False)
    coef_all.to_csv(RESULTS / "CSB_TRO_biological_control_coefficients.tsv", sep="\t", index=False)
    metrics[["control_model", "direction_cosine_to_measured_correction", "PC1_control", "PC2_control", "PC3_control", "PC3_negative_pull_recovered", "pred_basin_occupancy_q90", "validation_status"]].to_csv(
        RESULTS / "CSB_TRO_bio_control_direction_alignment.tsv", sep="\t", index=False
    )
    make_svg(FIGURES / "CSB_TRO_bio_control_occupancy.svg", metrics)
    write_doc(DOCS / "CSB_TRO_bio_control_interpretation.md", metrics, features)
    manifest = {
        "feature_table": args.feature_table,
        "n_features": int(len(features)),
        "n_random": args.n_random,
        "outputs": [
            str(RESULTS / "CSB_TRO_bio_control_occupancy_metrics.tsv"),
            str(RESULTS / "CSB_TRO_bio_control_direction_alignment.tsv"),
            str(RESULTS / "CSB_TRO_biological_control_coefficients.tsv"),
            str(RESULTS / "CSB_TRO_bio_control_matched_random_features.tsv"),
            str(FIGURES / "CSB_TRO_bio_control_occupancy.svg"),
            str(DOCS / "CSB_TRO_bio_control_interpretation.md"),
        ],
    }
    (RESULTS / "CSB_TRO_bio_control_manifest.json").write_text(json.dumps(manifest, indent=2), encoding="utf-8")
    print(json.dumps({
        "manifest": manifest,
        "top_models": metrics.sort_values("pred_basin_occupancy_q90", ascending=False).head(10).to_dict(orient="records"),
        "random_feature_unit_beta": random_metrics["pred_basin_occupancy_q90"].describe().to_dict(),
    }, indent=2))


if __name__ == "__main__":
    main()
````
