## Supplemental code for "An entropy-based diagnostic framework for characterizing methylation state dynamics during preimplantation development": run_branch_bound_biological_control_candidates.py

```
from __future__ import annotations

import json
import sys
from pathlib import Path

import numpy as np
import pandas as pd


BASE = Path(r"E:\CSB_TRO_operator_time_DMR_dynamics_2026-05-25")
CODE = BASE / "code"
RESULTS = BASE / "results"
FIGURES = BASE / "figures"
DOCS = BASE / "docs"
sys.path.insert(0, str(CODE))

from run_basin_residual_control_field import PRE_MORULA_STAGES, cosine, simulate_strict_pre_morula  # noqa: E402
from run_morula_basin_sde import (  # noqa: E402
    basin_definition,
    decode_latent,
    distribution_metrics,
    fit_latent_basis,
    fit_operator,
    load_inputs,
    stage_ids,
)


BASIS = RESULTS / "CSB_TRO_missing_control_term_module_basis.tsv"
CHROM_OVERLAP = RESULTS / "CSB_TRO_module_chromatin_overlap.tsv"
TF_FEATURES = RESULTS / "CSB_TRO_module_TF_activity_control_panel_features.tsv"
RNA_LINKED = RESULTS / "CSB_TRO_module_linked_RNA_delta_8cell_to_morula_priority_features.tsv"
RNA_LONG = RESULTS / "CSB_TRO_GSE36552_gene_stage_expression_long.tsv"

OUT_FEATURES = RESULTS / "CSB_TRO_branch_bound_biological_control_features.tsv"
OUT_ALPHA = RESULTS / "CSB_TRO_branch_bound_biological_control_alpha_scan.tsv"
OUT_SUMMARY = RESULTS / "CSB_TRO_branch_bound_biological_control_summary.tsv"
OUT_MANIFEST = RESULTS / "CSB_TRO_branch_bound_biological_control_manifest.json"
OUT_SVG = FIGURES / "CSB_TRO_branch_bound_biological_control_alpha_scan.svg"
OUT_DOC = DOCS / "CSB_TRO_branch_bound_biological_control_summary.md"


MODULES = ["M05", "M01", "M12", "M02", "M10"]
CLOSURE = ["M05", "M01", "M12"]
ACCESS = ["M02", "M10"]


PROGRAMS = {
    "closure_histone_repression_program": {
        "positive": ["EZH2", "EED", "SUZ12", "RBBP4", "RBBP7", "AEBP2", "JARID2", "HDAC1", "HDAC2", "HDAC3", "CHD4", "MBD3", "MTA1", "MTA2"],
        "negative": ["EP300", "CREBBP", "KAT2A", "KAT2B", "KAT5", "KAT6A", "KAT6B", "KDM6A", "KDM6B"],
        "branch": "closure",
        "interpretation": "PRC2/HDAC/NuRD gain minus acetylation/K27-demethylase gain; surrogate for H3K27ac loss/H3K27me3 gain.",
    },
    "access_promoter_activation_program": {
        "positive": ["KMT2A", "KMT2B", "KMT2C", "KMT2D", "SETD1A", "SETD1B", "WDR5", "ASH2L", "RBBP5", "DPY30", "EP300", "CREBBP", "KLF4", "KLF5", "POU5F1", "SOX2", "NANOG"],
        "negative": ["EZH2", "EED", "SUZ12", "HDAC1", "HDAC2", "HDAC3"],
        "branch": "access",
        "interpretation": "H3K4/H3K27ac/ZGA-TF activation minus repression; surrogate for promoter accessibility.",
    },
    "metabolic_onecarbon_sam_program": {
        "positive": ["MAT2A", "MAT2B", "MTHFD1", "MTHFD1L", "MTHFD2", "MTR", "MTRR", "SHMT1", "SHMT2", "DNMT1", "DNMT3A", "DNMT3B", "TET1", "TET2", "TET3"],
        "negative": [],
        "branch": "closure",
        "interpretation": "One-carbon/SAM/DNMT-TET surrogate; upstream metabolic control candidate, not direct histone state.",
    },
    "mitochondrial_oxphos_energy_program": {
        "positive": ["NDUFA1", "NDUFA2", "NDUFA3", "NDUFA4", "NDUFA5", "NDUFB1", "NDUFB2", "NDUFS1", "NDUFS2", "SDHA", "SDHB", "UQCRC1", "UQCRC2", "COX4I1", "COX5A", "ATP5F1A", "ATP5F1B", "TFAM", "POLG"],
        "negative": [],
        "branch": "access",
        "interpretation": "Mitochondrial/OXPHOS support surrogate; upstream access/promoter activity candidate.",
    },
}


def zscore(x: np.ndarray) -> np.ndarray:
    x = np.asarray(x, dtype=float)
    sd = np.nanstd(x, ddof=1)
    if not np.isfinite(sd) or sd == 0:
        return np.zeros_like(x)
    return (x - np.nanmean(x)) / sd


def basis() -> pd.DataFrame:
    b = pd.read_csv(BASIS, sep="\t")
    return b[b["module_id"].isin(MODULES)][["module_id", "latent_control_PC1", "latent_control_PC2", "latent_control_PC3", "n_DMRs", "ridge_weight"]].copy()


def feature_rows(name: str, values: dict[str, float], branch: str, modality: str, status: str, description: str, standardize_within_branch: bool = True) -> pd.DataFrame:
    b = basis()
    b["raw_value"] = b["module_id"].map(values).fillna(0.0).astype(float)
    active = CLOSURE if branch == "closure" else ACCESS if branch == "access" else MODULES
    b.loc[~b["module_id"].isin(active), "raw_value"] = 0.0
    b["control_value_z"] = 0.0
    mask = b["module_id"].isin(active)
    if standardize_within_branch:
        b.loc[mask, "control_value_z"] = zscore(b.loc[mask, "raw_value"].to_numpy(dtype=float)) if int(mask.sum()) > 1 else b.loc[mask, "raw_value"]
    else:
        b.loc[mask, "control_value_z"] = b.loc[mask, "raw_value"]
    b["feature_set"] = name
    b["branch"] = branch
    b["control_modality"] = modality
    b["biological_status"] = status
    b["description"] = description
    return b


def chromatin_candidates() -> list[pd.DataFrame]:
    overlap = pd.read_csv(CHROM_OVERLAP, sep="\t")
    rows = []
    for track_id in ["ATAC_8cell_3pn", "ATAC_8cell_2pn"]:
        sub = overlap[(overlap["track_id"] == track_id) & (overlap["analysis_status"] == "completed")]
        vals = dict(zip(sub["module_id"], pd.to_numeric(sub["target_overlap_fraction"], errors="coerce").fillna(0.0) - pd.to_numeric(sub["background_overlap_fraction"], errors="coerce").fillna(0.0)))
        rows.append(
            feature_rows(
                f"closure_ATAC_loss_{track_id}",
                {m: -vals.get(m, 0.0) for m in MODULES},
                "closure",
                "ATAC",
                "chromatin_proxy_boundary_no_morula_ATAC",
                "-8-cell ATAC target-background enrichment on M05/M01/M12; closure proxy until morula ATAC or histone tracks are available.",
            )
        )
        rows.append(
            feature_rows(
                f"access_promoter_ATAC_{track_id}",
                vals,
                "access",
                "ATAC",
                "real_ATAC_promoter_access_proxy_no_morula_stage",
                "8-cell ATAC target-background enrichment on promoter-like M02/M10; access branch candidate.",
            )
        )
    return rows


def tf_candidates() -> list[pd.DataFrame]:
    if not TF_FEATURES.exists():
        return []
    tf = pd.read_csv(TF_FEATURES, sep="\t")
    out = []
    for fs in ["q05_zero_filled_zscore_rebuilt", "q05_sparse_flipped_activity_sign", "all_motifs_zero_filled_zscore"]:
        sub = tf[tf["feature_set"] == fs].copy()
        if sub.empty:
            continue
        vals = dict(zip(sub["module_id"], pd.to_numeric(sub["control_value_z"], errors="coerce").fillna(0.0)))
        out.append(
            feature_rows(
                f"access_TF_activity_{fs}",
                vals,
                "access",
                "motif_TF_activity",
                "promoter_TF_surrogate",
                "M02/M10-bound motif x TF activity restricted to access branch; exploratory, not complete u_bio.",
            )
        )
    return out


def linked_rna_candidates() -> list[pd.DataFrame]:
    if not RNA_LINKED.exists():
        return []
    rna = pd.read_csv(RNA_LINKED, sep="\t")
    vals = dict(zip(rna["module_id"], pd.to_numeric(rna["control_value_z"], errors="coerce").fillna(0.0)))
    return [
        feature_rows(
            "closure_linked_RNA_delta_surrogate",
            vals,
            "closure",
            "nearest_gene_RNA",
            "weak_surrogate_gene_linked_not_histone",
            "Nearest-gene RNA delta restricted to closure branch; weak surrogate only.",
        ),
        feature_rows(
            "access_linked_RNA_delta_surrogate",
            vals,
            "access",
            "nearest_gene_RNA",
            "weak_surrogate_gene_linked_not_histone",
            "Nearest-gene RNA delta restricted to access branch; weak surrogate only.",
        ),
    ]


def program_score(program: dict) -> float:
    expr = pd.read_csv(RNA_LONG, sep="\t")
    expr["gene_id"] = expr["gene_id"].astype(str).str.upper()
    piv = expr.pivot_table(index="gene_id", columns="stage", values="expression_mean_RPKM", aggfunc="mean")
    if "8-cell" not in piv.columns or "morula" not in piv.columns:
        return 0.0
    delta = np.log2(pd.to_numeric(piv["morula"], errors="coerce").fillna(0.0) + 1.0) - np.log2(pd.to_numeric(piv["8-cell"], errors="coerce").fillna(0.0) + 1.0)
    pos = [g.upper() for g in program["positive"] if g.upper() in delta.index]
    neg = [g.upper() for g in program["negative"] if g.upper() in delta.index]
    pos_score = float(delta.loc[pos].mean()) if pos else 0.0
    neg_score = float(delta.loc[neg].mean()) if neg else 0.0
    return pos_score - neg_score


def program_candidates() -> list[pd.DataFrame]:
    out = []
    for name, spec in PROGRAMS.items():
        score = program_score(spec)
        branch = str(spec["branch"])
        active = CLOSURE if branch == "closure" else ACCESS
        vals = {m: score for m in active}
        out.append(
            feature_rows(
                name,
                vals,
                branch,
                "RNA_program",
                "surrogate_program_not_direct_histone_track",
                str(spec["interpretation"]) + f" GSE36552 morula-8cell score={score:.3f}.",
                standardize_within_branch=False,
            )
        )
    return out


def composite_candidates(parts: list[pd.DataFrame]) -> list[pd.DataFrame]:
    by_name = {str(p["feature_set"].iloc[0]): p for p in parts if len(p)}
    out = []
    combos = [
        ("biocontrol_ATAC_closure_plus_ATAC_access", ["closure_ATAC_loss_ATAC_8cell_3pn", "access_promoter_ATAC_ATAC_8cell_3pn"], "ATAC_branch_proxy"),
        ("biocontrol_ATAC_closure_plus_TF_access", ["closure_ATAC_loss_ATAC_8cell_3pn", "access_TF_activity_q05_sparse_flipped_activity_sign"], "ATAC_TF_branch_surrogate"),
        ("biocontrol_program_closure_plus_ATAC_access", ["closure_histone_repression_program", "access_promoter_ATAC_ATAC_8cell_3pn"], "program_ATAC_branch_surrogate"),
        ("biocontrol_program_closure_plus_program_access", ["closure_histone_repression_program", "access_promoter_activation_program"], "RNA_program_branch_surrogate"),
        ("biocontrol_ATAC_TF_RNA_composite", ["closure_ATAC_loss_ATAC_8cell_3pn", "access_promoter_ATAC_ATAC_8cell_3pn", "access_TF_activity_q05_sparse_flipped_activity_sign"], "ATAC_TF_branch_surrogate"),
    ]
    for name, keys, status in combos:
        tabs = [by_name[k] for k in keys if k in by_name]
        if not tabs:
            continue
        base = basis()
        base["raw_value"] = 0.0
        base["control_value_z"] = 0.0
        descriptions = []
        modalities = []
        for tab in tabs:
            sub = tab.set_index("module_id")
            base["control_value_z"] += base["module_id"].map(sub["control_value_z"]).fillna(0.0).astype(float)
            descriptions.append(str(tab["description"].iloc[0]))
            modalities.append(str(tab["control_modality"].iloc[0]))
        base["feature_set"] = name
        base["branch"] = "dual"
        base["control_modality"] = "+".join(sorted(set(modalities)))
        base["biological_status"] = status
        base["description"] = " + ".join(descriptions)
        out.append(base)
    return out


def build_features() -> pd.DataFrame:
    parts = []
    parts.extend(chromatin_candidates())
    parts.extend(tf_candidates())
    parts.extend(linked_rna_candidates())
    parts.extend(program_candidates())
    parts.extend(composite_candidates(parts))
    out = pd.concat(parts, ignore_index=True)
    out.to_csv(OUT_FEATURES, sep="\t", index=False)
    return out


def latent_context():
    matrix, ann, pairs = load_inputs()
    mu, sd, components, score_df = fit_latent_basis(matrix, 3)
    coef, _, _ = fit_operator(score_df, pairs, train_stages=set(PRE_MORULA_STAGES), lam=1000.0)
    strict_z = simulate_strict_pre_morula(score_df, ann, coef, n_steps=12)
    obs_z = score_df.loc[stage_ids(ann, "morula")].to_numpy(dtype=float)
    obs_dmr = matrix.loc[stage_ids(ann, "morula")].to_numpy(dtype=float)
    basin = basin_definition(obs_z)
    obs_dmr = matrix.loc[stage_ids(ann, "morula")].to_numpy(dtype=float)
    residual_z = obs_z.mean(axis=0) - strict_z.mean(axis=0)
    return mu, sd, components, strict_z, obs_z, obs_dmr, basin, residual_z


def alpha_values() -> list[float]:
    return [round(float(x), 2) for x in np.arange(0.0, 2.5001, 0.05)]


def evaluate(features: pd.DataFrame) -> pd.DataFrame:
    mu, sd, components, strict_z, obs_z, obs_dmr, basin, residual_z = latent_context()
    rng = np.random.default_rng(20260527)
    rows = []
    for feature_set, sub in features.groupby("feature_set"):
        dirs = sub[["latent_control_PC1", "latent_control_PC2", "latent_control_PC3"]].to_numpy(dtype=float)
        u = pd.to_numeric(sub["control_value_z"], errors="coerce").fillna(0.0).to_numpy(dtype=float)
        control = (dirs * u[:, None]).sum(axis=0)
        meta = sub.iloc[0]
        for alpha in alpha_values():
            for suffix, sign in [("", 1.0), ("_sign_flip", -1.0)]:
                vec = sign * alpha * control
                pred = strict_z + vec[None, :]
                pred_dmr = decode_latent(pred, mu, sd, components)
                rows.append(
                    {
                        "feature_set": feature_set,
                        "control_model": feature_set + suffix,
                        "alpha": alpha,
                        "sign_status": "correct_orientation" if sign > 0 else "sign_flip_control",
                        "branch": meta["branch"],
                        "control_modality": meta["control_modality"],
                        "biological_status": meta["biological_status"],
                        "PC1_control": float(vec[0]),
                        "PC2_control": float(vec[1]),
                        "PC3_control": float(vec[2]),
                        "direction_cosine_to_measured_correction": cosine(vec, residual_z) if np.linalg.norm(vec) else np.nan,
                        "PC3_negative_pull_recovered": float(-vec[2] / (-residual_z[2])) if residual_z[2] < 0 else np.nan,
                        **distribution_metrics(pred, obs_z, pred_dmr, obs_dmr, basin, rng),
                    }
                )
    out = pd.DataFrame(rows)
    out.to_csv(OUT_ALPHA, sep="\t", index=False)
    return out


def summarize(alpha: pd.DataFrame) -> pd.DataFrame:
    rows = []
    target = 0.875
    for feature_set, sub in alpha[alpha["sign_status"] == "correct_orientation"].groupby("feature_set"):
        sub = sub.sort_values("alpha")
        best = sub.sort_values(["pred_basin_occupancy_q90", "direction_cosine_to_measured_correction"], ascending=False).iloc[0]
        at1 = sub.iloc[(sub["alpha"] - 1.0).abs().argsort()].iloc[0]
        reached = sub[sub["pred_basin_occupancy_q90"] >= target]
        flip = alpha[(alpha["feature_set"] == feature_set) & (alpha["sign_status"] == "sign_flip_control")]
        flip_best = flip.sort_values("pred_basin_occupancy_q90", ascending=False).iloc[0] if len(flip) else None
        rows.append(
            {
                "feature_set": feature_set,
                "branch": best["branch"],
                "control_modality": best["control_modality"],
                "biological_status": best["biological_status"],
                "max_occupancy": float(best["pred_basin_occupancy_q90"]),
                "alpha_at_max": float(best["alpha"]),
                "occupancy_at_alpha_1": float(at1["pred_basin_occupancy_q90"]),
                "cosine_at_alpha_1": float(at1["direction_cosine_to_measured_correction"]),
                "PC3_recovery_at_alpha_1": float(at1["PC3_negative_pull_recovered"]),
                "alpha_to_observed_occupancy": float(reached.iloc[0]["alpha"]) if len(reached) else np.nan,
                "max_sign_flip_occupancy": float(flip_best["pred_basin_occupancy_q90"]) if flip_best is not None else np.nan,
            }
        )
    out = pd.DataFrame(rows).sort_values(["biological_status", "max_occupancy", "cosine_at_alpha_1"], ascending=[True, False, False])
    out.to_csv(OUT_SUMMARY, sep="\t", index=False)
    return out


def make_svg(summary: pd.DataFrame) -> None:
    rows = summary.sort_values("max_occupancy", ascending=False).head(12)
    width, height = 1000, 440
    left, right, top, bottom = 85, 25, 45, 150
    plot_w = width - left - right
    plot_h = height - top - bottom
    gap = plot_w / max(len(rows), 1)
    bar_w = gap * 0.62
    lines = [
        f'<svg xmlns="http://www.w3.org/2000/svg" width="{width}" height="{height}" viewBox="0 0 {width} {height}">',
        '<rect width="100%" height="100%" fill="white"/>',
        f'<text x="{left}" y="28" font-family="Arial" font-size="18" font-weight="700">Branch-bound biological control candidates</text>',
        f'<line x1="{left}" y1="{height-bottom}" x2="{width-right}" y2="{height-bottom}" stroke="#222"/>',
        f'<line x1="{left}" y1="{top}" x2="{left}" y2="{height-bottom}" stroke="#222"/>',
    ]
    for tick in [0.044, 0.3, 0.5, 0.875, 1.0]:
        y = height - bottom - tick * plot_h
        color = "#b23b3b" if tick in [0.044, 0.875] else "#dddddd"
        lines.append(f'<line x1="{left}" y1="{y:.2f}" x2="{width-right}" y2="{y:.2f}" stroke="{color}" stroke-dasharray="4 4"/>')
        lines.append(f'<text x="{left-8}" y="{y+4:.2f}" text-anchor="end" font-family="Arial" font-size="10">{tick:.3g}</text>')
    for i, row in enumerate(rows.itertuples()):
        x = left + i * gap + (gap - bar_w) / 2
        val = float(row.max_occupancy)
        y = height - bottom - val * plot_h
        color = "#2c6f5a" if "ATAC" in row.control_modality else "#6f4d8b" if "RNA" in row.control_modality else "#777777"
        lines.append(f'<rect x="{x:.2f}" y="{y:.2f}" width="{bar_w:.2f}" height="{val * plot_h:.2f}" fill="{color}"/>')
        lines.append(f'<text x="{x+bar_w/2:.2f}" y="{y-5:.2f}" text-anchor="middle" font-family="Arial" font-size="10">{val:.3f}</text>')
        label = row.feature_set.replace("_", " ")[:42]
        lines.append(f'<text x="{x+bar_w/2:.2f}" y="{height-bottom+18}" text-anchor="end" transform="rotate(-45 {x+bar_w/2:.2f} {height-bottom+18})" font-family="Arial" font-size="9">{label}</text>')
    lines.append("</svg>")
    OUT_SVG.write_text("\n".join(lines), encoding="utf-8")


def write_doc(summary: pd.DataFrame) -> None:
    lines = [
        "# Branch-Bound Biological Control Candidates",
        "",
        "Status: `completed_with_available_surrogates_no_histone_tracks`",
        "",
        "Goal: start replacing the dual-branch proxy with biologically named closure/access variables while explicitly preserving boundaries.",
        "",
        "## Top Candidates",
        "",
    ]
    for row in summary.sort_values("max_occupancy", ascending=False).head(12).itertuples():
        lines.append(
            f"- {row.feature_set}: status={row.biological_status}; max_occ={row.max_occupancy:.3f}; "
            f"occ@1={row.occupancy_at_alpha_1:.3f}; cosine@1={row.cosine_at_alpha_1:.3f}; "
            f"PC3@1={row.PC3_recovery_at_alpha_1:.3f}; signflip_max={row.max_sign_flip_occupancy:.3f}"
        )
    lines += [
        "",
        "## Boundary",
        "",
        "No H3K27ac/H3K4me3/H3K27me3 track is currently analysis-ready, so none of these should be called final histone u_bio. The strongest current candidates are branch-bound ATAC/TF/RNA surrogates. The final replacement still requires direct histone or chromatin-state tracks.",
    ]
    OUT_DOC.write_text("\n".join(lines) + "\n", encoding="utf-8")


def main() -> None:
    RESULTS.mkdir(parents=True, exist_ok=True)
    FIGURES.mkdir(parents=True, exist_ok=True)
    DOCS.mkdir(parents=True, exist_ok=True)
    features = build_features()
    alpha = evaluate(features)
    summary = summarize(alpha)
    make_svg(summary)
    write_doc(summary)
    OUT_MANIFEST.write_text(
        json.dumps(
            {
                "status": "completed_with_available_surrogates_no_histone_tracks",
                "outputs": [str(OUT_FEATURES), str(OUT_ALPHA), str(OUT_SUMMARY), str(OUT_SVG), str(OUT_DOC)],
                "boundary": "surrogate candidates are not final histone u_bio",
            },
            indent=2,
        )
        + "\n",
        encoding="utf-8",
    )
    print(json.dumps({"status": "completed", "top": summary.sort_values("max_occupancy", ascending=False).head(8).to_dict(orient="records")}, indent=2))


if __name__ == "__main__":
    main()
```
