## Supplemental code for "An entropy-based diagnostic framework for characterizing methylation state dynamics during preimplantation development": run_build_biological_control_features.py

````
from __future__ import annotations

import argparse
import json
from pathlib import Path

import numpy as np
import pandas as pd


BASE = Path(r"E:\CSB_TRO_operator_time_DMR_dynamics_2026-05-25")
RESULTS = BASE / "results"
DOCS = BASE / "docs"


PRIORITY_MODULES = ["M05", "M01", "M12", "M02", "M10", "M09", "M08", "M06"]


def read_tsv(path: Path) -> pd.DataFrame:
    return pd.read_csv(path, sep="\t") if path.exists() else pd.DataFrame()


def zscore(x: pd.Series) -> pd.Series:
    x = pd.to_numeric(x, errors="coerce").astype(float)
    sd = x.std()
    if not np.isfinite(sd) or sd == 0:
        return x * 0.0
    return (x - x.mean()) / sd


def parse_stage_columns(df: pd.DataFrame):
    stage_aliases = {
        "8-cell": ["8-cell", "8cell", "X8_cell", "eight_cell"],
        "morula": ["morula", "Morula"],
        "blastocyst": ["blastocyst", "Blastocyst"],
    }
    out = {}
    lower_map = {c.lower().replace(" ", "_"): c for c in df.columns}
    for stage, aliases in stage_aliases.items():
        for alias in aliases:
            key = alias.lower().replace(" ", "_")
            if key in lower_map:
                out[stage] = lower_map[key]
                break
    return out


def load_gene_links(path: str) -> pd.DataFrame:
    if not path:
        nearest = read_tsv(RESULTS / "CSB_TRO_residual_DMR_nearest_genes.tsv")
        if nearest.empty:
            return pd.DataFrame()
        return nearest.rename(columns={"nearest_gene": "gene_id"})
    df = read_tsv(Path(path))
    if "gene_name" in df.columns and "gene_id" not in df.columns:
        df = df.rename(columns={"gene_name": "gene_id"})
    if "nearest_gene" in df.columns and "gene_id" not in df.columns:
        df = df.rename(columns={"nearest_gene": "gene_id"})
    return df


def module_gene_links(dmr: pd.DataFrame, gene_links: pd.DataFrame) -> pd.DataFrame:
    if gene_links.empty or "cluster_name" not in gene_links.columns or "gene_id" not in gene_links.columns:
        return pd.DataFrame(columns=["module_id", "cluster_name", "gene_id", "link_type", "distance_to_TSS"])
    keep = [c for c in ["cluster_name", "gene_id", "distance_to_TSS", "gene_type", "link_type"] if c in gene_links.columns]
    links = gene_links[keep].copy()
    links = links.merge(dmr[["cluster_name", "module_id", "basin_residual_rank", "abs_latent_residual_delta_beta"]], on="cluster_name", how="inner")
    if "link_type" not in links.columns:
        links["link_type"] = "nearest_gene"
    return links


def build_internal_features(dmr: pd.DataFrame, module_basis: pd.DataFrame) -> pd.DataFrame:
    dmr = dmr.copy()
    dmr["cpg_per_100bp"] = pd.to_numeric(dmr["n_cpg_target"], errors="coerce") / (pd.to_numeric(dmr["width"], errors="coerce") / 100.0)
    rows = []
    for module_id, sub in dmr.groupby("module_id"):
        basis = module_basis[module_basis["module_id"] == module_id]
        if basis.empty:
            continue
        b = basis.iloc[0]
        rows.extend([
            {
                "module_id": module_id,
                "candidate_control": f"internal_genomic_cpg_density_{module_id}",
                "control_modality": "internal_genomic_proxy",
                "control_value": float(sub["cpg_per_100bp"].replace([np.inf, -np.inf], np.nan).mean()),
                "control_stage_window": "static",
                "leakage_status": "non_morula_methylation_but_not_external_omics",
                "interpretation": "Mean CpG density of residual DMRs in this module; useful as a genomic covariate, not a biological activity signal.",
            },
            {
                "module_id": module_id,
                "candidate_control": f"internal_age_weight_abs_{module_id}",
                "control_modality": "internal_methylation_prior_proxy",
                "control_value": float(pd.to_numeric(sub["age_weight_5yr"], errors="coerce").abs().mean()),
                "control_stage_window": "static",
                "leakage_status": "internal_methylation_prior_not_external_omics",
                "interpretation": "Mean absolute age-DMR weight; not a non-leaking external biological control.",
            },
            {
                "module_id": module_id,
                "candidate_control": f"measured_residual_module_weight_{module_id}",
                "control_modality": "measured_residual_diagnostic",
                "control_value": float(abs(b.get("ridge_weight", 0.0))),
                "control_stage_window": "8-cell_to_morula",
                "leakage_status": "uses_morula_methylation_residual",
                "interpretation": "Diagnostic upper-bound feature derived from measured correction term, not a valid u_bio predictor.",
            },
        ])
    feat = pd.DataFrame(rows)
    feat["control_value_z"] = feat.groupby("control_modality")["control_value"].transform(zscore)
    return feat


def build_rna_features(gene_links: pd.DataFrame, rna_path: str) -> pd.DataFrame:
    if not rna_path:
        return pd.DataFrame()
    rna = read_tsv(Path(rna_path))
    if rna.empty:
        return pd.DataFrame()
    gene_col = "gene_id" if "gene_id" in rna.columns else ("gene_name" if "gene_name" in rna.columns else rna.columns[0])
    stage_cols = parse_stage_columns(rna)
    if "8-cell" not in stage_cols or "morula" not in stage_cols:
        raise ValueError("RNA matrix must include gene_id/gene_name and stage columns for at least 8-cell and morula.")
    rna = rna.rename(columns={gene_col: "gene_id"})
    rna["rna_delta_8cell_to_morula"] = pd.to_numeric(rna[stage_cols["morula"]], errors="coerce") - pd.to_numeric(rna[stage_cols["8-cell"]], errors="coerce")
    if "blastocyst" in stage_cols:
        rna["rna_delta_morula_to_blastocyst"] = pd.to_numeric(rna[stage_cols["blastocyst"]], errors="coerce") - pd.to_numeric(rna[stage_cols["morula"]], errors="coerce")
    joined = gene_links.merge(rna, on="gene_id", how="inner")
    rows = []
    for module_id, sub in joined.groupby("module_id"):
        rows.append({
            "module_id": module_id,
            "candidate_control": f"RNA_delta_8cell_to_morula_{module_id}",
            "control_modality": "RNA",
            "control_value": float(sub["rna_delta_8cell_to_morula"].mean()),
            "control_stage_window": "8-cell_to_morula",
            "leakage_status": "methylation_non_leaking_external_morula_allowed",
            "interpretation": "Mean linked-gene RNA expression change from 8-cell to morula.",
            "linked_genes": ",".join(sorted(set(map(str, sub["gene_id"])))[:50]),
        })
        if "rna_delta_morula_to_blastocyst" in sub.columns:
            rows.append({
                "module_id": module_id,
                "candidate_control": f"RNA_delta_morula_to_blastocyst_{module_id}",
                "control_modality": "RNA",
                "control_value": float(sub["rna_delta_morula_to_blastocyst"].mean()),
                "control_stage_window": "morula_to_blastocyst",
                "leakage_status": "methylation_non_leaking_external_morula_allowed",
                "interpretation": "Mean linked-gene RNA expression change from morula to blastocyst.",
                "linked_genes": ",".join(sorted(set(map(str, sub["gene_id"])))[:50]),
            })
    feat = pd.DataFrame(rows)
    if len(feat):
        feat["control_value_z"] = feat.groupby("control_modality")["control_value"].transform(zscore)
    return feat


def build_track_features(dmr: pd.DataFrame, track_summary_path: str, modality: str) -> pd.DataFrame:
    if not track_summary_path:
        return pd.DataFrame()
    tab = read_tsv(Path(track_summary_path))
    required = {"module_id", "control_value"}
    if not required.issubset(tab.columns):
        raise ValueError(f"{modality} feature table must contain module_id and control_value.")
    tab = tab.copy()
    if "candidate_control" not in tab.columns:
        tab["candidate_control"] = modality + "_" + tab["module_id"].astype(str)
    tab["control_modality"] = modality
    if "control_stage_window" not in tab.columns:
        tab["control_stage_window"] = "user_supplied"
    if "leakage_status" not in tab.columns:
        tab["leakage_status"] = "user_supplied_external_feature"
    if "interpretation" not in tab.columns:
        tab["interpretation"] = f"User-supplied {modality} module activity feature."
    tab["control_value_z"] = tab.groupby("control_modality")["control_value"].transform(zscore)
    return tab


def write_doc(path: Path, features: pd.DataFrame, missing: list[str]):
    lines = [
        "# Biological control features",
        "",
        "This table defines candidate module-level u_bio features for the interpretable control term:",
        "",
        "```text",
        "B u_bio(tau) = sum_m beta_m u_m(tau) b_m",
        "```",
        "",
        "Current feature modalities:",
    ]
    for modality, sub in features.groupby("control_modality"):
        statuses = ",".join(sorted(set(map(str, sub["leakage_status"]))))
        lines.append(f"- {modality}: n={len(sub)}, leakage_status={statuses}")
    if missing:
        lines.extend(["", "Missing optional external inputs:"])
        for item in missing:
            lines.append(f"- {item}")
    lines.extend([
        "",
        "Interpretation boundary: internal genomic and measured-residual features are useful for dry-run and upper-bound checks, but they are not proof of a real external biological control variable. RNA/ATAC/histone/motif inputs are required for u_bio identification.",
    ])
    path.write_text("\n".join(lines) + "\n", encoding="utf-8")


def main():
    parser = argparse.ArgumentParser()
    parser.add_argument("--gene-links", default="", help="Optional TSV with cluster_name,gene_id[,distance_to_TSS]. Defaults to residual nearest-gene table if present.")
    parser.add_argument("--rna-matrix", default="", help="Optional gene x stage RNA matrix with gene_id/gene_name and 8-cell,morula[,blastocyst] columns.")
    parser.add_argument("--atac-features", default="", help="Optional module feature TSV with module_id,control_value.")
    parser.add_argument("--histone-features", default="", help="Optional module feature TSV with module_id,control_value.")
    parser.add_argument("--motif-features", default="", help="Optional module feature TSV with module_id,control_value.")
    args = parser.parse_args()

    RESULTS.mkdir(parents=True, exist_ok=True)
    DOCS.mkdir(parents=True, exist_ok=True)

    dmr = read_tsv(RESULTS / "CSB_TRO_basin_residual_DMR_ranking.tsv")
    module_basis = read_tsv(RESULTS / "CSB_TRO_missing_control_term_module_basis.tsv")
    if dmr.empty or module_basis.empty:
        raise FileNotFoundError("Run residual control and missing-control decomposition scripts before building biological features.")

    gene_links = module_gene_links(dmr, load_gene_links(args.gene_links))
    features = [build_internal_features(dmr, module_basis)]
    if len(gene_links):
        gene_links.to_csv(RESULTS / "CSB_TRO_module_gene_links.tsv", sep="\t", index=False)
    else:
        pd.DataFrame(columns=["module_id", "cluster_name", "gene_id", "link_type", "distance_to_TSS"]).to_csv(RESULTS / "CSB_TRO_module_gene_links.tsv", sep="\t", index=False)

    for feat in [
        build_rna_features(gene_links, args.rna_matrix),
        build_track_features(dmr, args.atac_features, "ATAC"),
        build_track_features(dmr, args.histone_features, "histone"),
        build_track_features(dmr, args.motif_features, "motif_activity"),
    ]:
        if len(feat):
            features.append(feat)
    all_features = pd.concat(features, ignore_index=True, sort=False)
    all_features = all_features.merge(module_basis[["module_id", "n_DMRs", "latent_control_PC1", "latent_control_PC2", "latent_control_PC3", "latent_control_norm", "ridge_weight"]], on="module_id", how="left")
    all_features["candidate_control_direction_PC1"] = all_features["control_value_z"].fillna(0.0) * all_features["latent_control_PC1"]
    all_features["candidate_control_direction_PC2"] = all_features["control_value_z"].fillna(0.0) * all_features["latent_control_PC2"]
    all_features["candidate_control_direction_PC3"] = all_features["control_value_z"].fillna(0.0) * all_features["latent_control_PC3"]

    all_features.to_csv(RESULTS / "CSB_TRO_module_bio_features.tsv", sep="\t", index=False)
    all_features[all_features["control_modality"] == "RNA"].to_csv(RESULTS / "CSB_TRO_module_RNA_activity.tsv", sep="\t", index=False)
    all_features[all_features["control_modality"] == "ATAC"].to_csv(RESULTS / "CSB_TRO_module_ATAC_activity.tsv", sep="\t", index=False)
    all_features[all_features["control_modality"] == "histone"].to_csv(RESULTS / "CSB_TRO_module_histone_activity.tsv", sep="\t", index=False)
    all_features[all_features["control_modality"] == "motif_activity"].to_csv(RESULTS / "CSB_TRO_module_motif_activity.tsv", sep="\t", index=False)

    missing = []
    if not args.rna_matrix:
        missing.append("RNA expression matrix for linked genes")
    if not args.atac_features:
        missing.append("ATAC module activity features")
    if not args.histone_features:
        missing.append("histone module activity features")
    if not args.motif_features:
        missing.append("motif enrichment/activity features")
    write_doc(DOCS / "CSB_TRO_module_bio_features_interpretation.md", all_features, missing)
    manifest = {
        "gene_links": args.gene_links or "default_nearest_gene_if_available",
        "rna_matrix": args.rna_matrix or None,
        "atac_features": args.atac_features or None,
        "histone_features": args.histone_features or None,
        "motif_features": args.motif_features or None,
        "n_features": int(len(all_features)),
        "modalities": sorted(set(all_features["control_modality"])),
        "outputs": [
            str(RESULTS / "CSB_TRO_module_bio_features.tsv"),
            str(RESULTS / "CSB_TRO_module_gene_links.tsv"),
            str(RESULTS / "CSB_TRO_module_RNA_activity.tsv"),
            str(RESULTS / "CSB_TRO_module_ATAC_activity.tsv"),
            str(RESULTS / "CSB_TRO_module_histone_activity.tsv"),
            str(RESULTS / "CSB_TRO_module_motif_activity.tsv"),
            str(DOCS / "CSB_TRO_module_bio_features_interpretation.md"),
        ],
    }
    (RESULTS / "CSB_TRO_module_bio_features_manifest.json").write_text(json.dumps(manifest, indent=2), encoding="utf-8")
    print(json.dumps(manifest, indent=2))


if __name__ == "__main__":
    main()
````
