## Supplemental code for "An entropy-based diagnostic framework for characterizing methylation state dynamics during preimplantation development": run_chromatin_overlap_for_m02_klf.py

```
from __future__ import annotations

import gzip
import json
from pathlib import Path

import numpy as np
import pandas as pd


BASE = Path(r"E:\CSB_TRO_operator_time_DMR_dynamics_2026-05-25")
EXTERNAL = BASE / "external"
RESULTS = BASE / "results"
DOCS = BASE / "docs"

TARGETS = RESULTS / "CSB_TRO_M02_KLF4_KLF5_residual_DMR_targets.tsv"
Q05_FEATURES = RESULTS / "CSB_TRO_motif_TF_activity_matched_bg_q05_control_features.tsv"
MODULE_BASIS = RESULTS / "CSB_TRO_missing_control_term_module_basis.tsv"

OUT_TARGET_OVERLAP = RESULTS / "CSB_TRO_M02_KLF4_KLF5_ATAC_overlap.tsv"
OUT_MODULE_FEATURES = RESULTS / "CSB_TRO_M02_KLF4_KLF5_ATAC_gated_control_features.tsv"
OUT_DOC = DOCS / "CSB_TRO_M02_KLF4_KLF5_ATAC_overlap.md"
OUT_MANIFEST = RESULTS / "CSB_TRO_M02_KLF4_KLF5_ATAC_overlap_manifest.json"

ATAC_TRACKS = {
    "GSE101571_8cell_2pn_ATAC": EXTERNAL / "atac" / "GSE101571_8cell_2pn_peaks.bed.gz",
    "GSE101571_8cell_3pn_ATAC": EXTERNAL / "atac" / "GSE101571_8cell_3pn_peaks.bed.gz",
    "GSE101571_icm_2pn_ATAC": EXTERNAL / "atac" / "GSE101571_icm_2pn_peaks.bed.gz",
    "GSE101571_icm_3pn_ATAC": EXTERNAL / "atac" / "GSE101571_icm_3pn_peaks.bed.gz",
}


def read_bed(path: Path) -> pd.DataFrame:
    opener = gzip.open if path.suffix == ".gz" else open
    rows = []
    with opener(path, "rt", encoding="utf-8", errors="replace") as handle:
        for line in handle:
            if not line.strip() or line.startswith("#"):
                continue
            parts = line.rstrip("\n").split("\t")
            if len(parts) < 3:
                continue
            rows.append({"chr": parts[0], "start": int(float(parts[1])), "end": int(float(parts[2])), "name": parts[3] if len(parts) > 3 else ""})
    return pd.DataFrame(rows)


def overlap_count(row, by_chr: dict[str, pd.DataFrame]) -> tuple[int, str]:
    sub = by_chr.get(row["chr"])
    if sub is None or sub.empty:
        return 0, ""
    hits = sub[(sub["start"] < int(row["end"])) & (sub["end"] > int(row["start"]))]
    names = ",".join(hits["name"].astype(str).head(10).tolist()) if "name" in hits.columns else ""
    return int(len(hits)), names


def zscore(x: pd.Series) -> pd.Series:
    y = pd.to_numeric(x, errors="coerce").astype(float)
    sd = y.std()
    if not np.isfinite(sd) or sd == 0:
        return y * 0.0
    return (y - y.mean()) / sd


def main() -> None:
    targets = pd.read_csv(TARGETS, sep="\t")
    target_rows = targets[["cluster_name", "module_id", "chr", "start", "end", "TFs", "basin_residual_rank"]].copy()
    for label, path in ATAC_TRACKS.items():
        peaks = read_bed(path)
        by_chr = {c: sub for c, sub in peaks.groupby("chr")}
        counts = []
        names = []
        for _, row in target_rows.iterrows():
            c, n = overlap_count(row, by_chr)
            counts.append(c)
            names.append(n)
        target_rows[f"{label}_overlap_count"] = counts
        target_rows[f"{label}_overlap"] = [c > 0 for c in counts]
        target_rows[f"{label}_overlap_names"] = names
    target_rows["any_8cell_ATAC_overlap"] = target_rows[["GSE101571_8cell_2pn_ATAC_overlap", "GSE101571_8cell_3pn_ATAC_overlap"]].any(axis=1)
    target_rows["any_icm_ATAC_overlap"] = target_rows[["GSE101571_icm_2pn_ATAC_overlap", "GSE101571_icm_3pn_ATAC_overlap"]].any(axis=1)
    target_rows.to_csv(OUT_TARGET_OVERLAP, sep="\t", index=False)

    motif_features = pd.read_csv(Q05_FEATURES, sep="\t")
    basis = pd.read_csv(MODULE_BASIS, sep="\t")
    overlap_fraction = float(target_rows["any_8cell_ATAC_overlap"].mean()) if len(target_rows) else 0.0
    features = motif_features.copy()
    features["control_value"] = pd.to_numeric(features["control_value"], errors="coerce").fillna(0.0)
    features.loc[features["module_id"] == "M02", "control_value"] *= overlap_fraction
    features["control_value_z"] = zscore(features["control_value"])
    features["candidate_control"] = "ATAC_gated_" + features["candidate_control"].astype(str)
    features["control_modality"] = "ATAC"
    features["leakage_status"] = "methylation_non_leaking_motif_TF_ATAC_gated_no_morula_ATAC"
    features["interpretation"] = "M02 KLF4/KLF5 motif x TF activity gated by GSE101571 8-cell ATAC overlap fraction; no morula ATAC is available."
    for pc in ["PC1", "PC2", "PC3"]:
        features[f"candidate_control_direction_{pc}"] = features["control_value_z"] * pd.to_numeric(features[f"latent_control_{pc}"], errors="coerce")
    features.to_csv(OUT_MODULE_FEATURES, sep="\t", index=False)

    lines = [
        "# M02 KLF4/KLF5 ATAC overlap",
        "",
        "GSE101571 ATAC peaks were used as chromatin accessibility support. These data include 8-cell and ICM, not morula, so this is pre-morula/future-accessibility support rather than direct morula ATAC validation.",
        "",
        f"M02 KLF4/KLF5 target DMRs: {len(target_rows)}",
        f"Any 8-cell ATAC overlap fraction: {overlap_fraction:.3f}",
        f"Any ICM ATAC overlap fraction: {float(target_rows['any_icm_ATAC_overlap'].mean()) if len(target_rows) else 0.0:.3f}",
        "",
        "Track overlap fractions:",
    ]
    for label in ATAC_TRACKS:
        lines.append(f"- {label}: {float(target_rows[f'{label}_overlap'].mean()):.3f}")
    OUT_DOC.write_text("\n".join(lines) + "\n", encoding="utf-8")
    OUT_MANIFEST.write_text(
        json.dumps(
            {
                "targets": str(TARGETS),
                "atac_tracks": {k: str(v) for k, v in ATAC_TRACKS.items()},
                "n_targets": int(len(target_rows)),
                "any_8cell_ATAC_overlap_fraction": overlap_fraction,
                "outputs": [str(OUT_TARGET_OVERLAP), str(OUT_MODULE_FEATURES), str(OUT_DOC)],
                "boundary": "GSE101571 has 8-cell and ICM ATAC, not morula ATAC.",
            },
            indent=2,
        )
        + "\n",
        encoding="utf-8",
    )
    print(json.dumps({"n_targets": int(len(target_rows)), "any_8cell_ATAC_overlap_fraction": overlap_fraction}, indent=2))


if __name__ == "__main__":
    main()
```
