## Supplemental code for "An entropy-based diagnostic framework for characterizing methylation state dynamics during preimplantation development": run_composite_biocontrol_proxy_alpha_scan.py

```
from __future__ import annotations

import json
import sys
from pathlib import Path

import numpy as np
import pandas as pd


BASE = Path(r"E:\CSB_TRO_operator_time_DMR_dynamics_2026-05-25")
CODE = BASE / "code"
RESULTS = BASE / "results"
FIGURES = BASE / "figures"
DOCS = BASE / "docs"
sys.path.insert(0, str(CODE))

from run_basin_residual_control_field import PRE_MORULA_STAGES, cosine, simulate_strict_pre_morula  # noqa: E402
from run_morula_basin_sde import (  # noqa: E402
    basin_definition,
    decode_latent,
    distribution_metrics,
    fit_latent_basis,
    fit_operator,
    load_inputs,
    stage_ids,
)


PRIORITY_MODULES = ["M05", "M01", "M12", "M02", "M10"]
CORE_MODULES = ["M05", "M01", "M12", "M02"]

RNA = RESULTS / "CSB_TRO_module_linked_RNA_delta_8cell_to_morula_priority_features.tsv"
CHROM = RESULTS / "CSB_TRO_chromatin_gated_TF_activity.tsv"
MOTIF = RESULTS / "CSB_TRO_module_TF_activity_control_panel_features.tsv"
BASIS = RESULTS / "CSB_TRO_missing_control_term_module_basis.tsv"
REGION = RESULTS / "CSB_TRO_residual_module_region_composition.tsv"

OUT_FEATURES = RESULTS / "CSB_TRO_composite_biocontrol_proxy_features.tsv"
OUT_ALPHA = RESULTS / "CSB_TRO_composite_biocontrol_alpha_scan.tsv"
OUT_SUMMARY = RESULTS / "CSB_TRO_composite_biocontrol_proxy_summary.tsv"
OUT_MANIFEST = RESULTS / "CSB_TRO_composite_biocontrol_proxy_manifest.json"
OUT_SVG = FIGURES / "CSB_TRO_composite_biocontrol_alpha_scan.svg"
OUT_DOC = DOCS / "CSB_TRO_composite_biocontrol_proxy_summary.md"


def zscore(x: np.ndarray) -> np.ndarray:
    x = np.asarray(x, dtype=float)
    sd = np.nanstd(x, ddof=1)
    if not np.isfinite(sd) or sd == 0:
        return np.zeros_like(x)
    return (x - np.nanmean(x)) / sd


def module_basis() -> pd.DataFrame:
    b = pd.read_csv(BASIS, sep="\t")
    return b[b["module_id"].isin(PRIORITY_MODULES)][
        ["module_id", "n_DMRs", "latent_control_PC1", "latent_control_PC2", "latent_control_PC3", "latent_control_norm", "ridge_weight"]
    ].copy()


def base_table() -> pd.DataFrame:
    return module_basis()[["module_id"]].copy()


def feature_rows(name: str, values: dict[str, float], modality: str, leakage: str, description: str) -> pd.DataFrame:
    tab = base_table()
    tab["raw_value"] = tab["module_id"].map(values).fillna(0.0).astype(float)
    tab["control_value_z"] = zscore(tab["raw_value"].to_numpy(dtype=float))
    tab["feature_set"] = name
    tab["control_modality"] = modality
    tab["leakage_status"] = leakage
    tab["description"] = description
    return tab.merge(module_basis(), on="module_id", how="left")


def load_available_scores() -> dict[str, pd.DataFrame]:
    out = {}
    if RNA.exists():
        rna = pd.read_csv(RNA, sep="\t")
        out["RNA_nearest_delta"] = feature_rows(
            "RNA_nearest_delta",
            dict(zip(rna["module_id"], pd.to_numeric(rna["control_value_z"], errors="coerce").fillna(0.0))),
            "RNA",
            "methylation_non_leaking_nearest_TSS_RNA",
            "Nearest-TSS RNA delta z-score. Known to be coarse but retained as available external signal.",
        )
    if CHROM.exists():
        chrom = pd.read_csv(CHROM, sep="\t")
        for fs in ["ATAC_8cell_2pn_chromatin_only", "ATAC_8cell_3pn_chromatin_only", "ATAC_ICM_2pn_chromatin_only", "ATAC_ICM_3pn_chromatin_only"]:
            sub = chrom[chrom["feature_set"] == fs]
            if len(sub):
                vals = dict(zip(sub["module_id"], pd.to_numeric(sub["control_value_z"], errors="coerce").fillna(0.0)))
                out[fs] = feature_rows(
                    fs,
                    vals,
                    "ATAC",
                    "methylation_non_leaking_ATAC_overlap",
                    "ATAC target-background enrichment z-score.",
                )
                out[fs + "_inverse"] = feature_rows(
                    fs + "_inverse",
                    {k: -v for k, v in vals.items()},
                    "ATAC_inverse",
                    "orientation_corrected_proxy_uses_measured_direction_boundary",
                    "Inverse ATAC orientation because ATAC-only control was anti-aligned; proxy only, not final u_bio.",
                )
    if MOTIF.exists():
        motif = pd.read_csv(MOTIF, sep="\t")
        for fs in ["q05_sparse_flipped_activity_sign", "q05_zero_filled_zscore_rebuilt"]:
            sub = motif[motif["feature_set"] == fs]
            if len(sub):
                out[fs] = feature_rows(
                    fs,
                    dict(zip(sub["module_id"], pd.to_numeric(sub["control_value_z"], errors="coerce").fillna(0.0))),
                    "motif_TF",
                    "methylation_non_leaking_motif_TF_expression",
                    "Module motif x TF expression signal.",
                )
    if REGION.exists():
        reg = pd.read_csv(REGION, sep="\t")
        out["region_state_prior"] = feature_rows(
            "region_state_prior",
            dict(zip(reg["module_id"], pd.to_numeric(reg["intergenic_fraction"], errors="coerce").fillna(0.0))),
            "region_prior",
            "static_genomic_proxy_not_external_dynamic",
            "Static distal/intergenic proxy; useful for module hypothesis but not a dynamic u_bio.",
        )
    return out


def combine_features(name: str, parts: list[pd.DataFrame], modality: str, leakage: str, description: str) -> pd.DataFrame:
    vals = {m: 0.0 for m in PRIORITY_MODULES}
    for part in parts:
        for row in part.itertuples():
            vals[str(row.module_id)] += float(row.control_value_z)
    return feature_rows(name, vals, modality, leakage, description)


def build_feature_panel() -> pd.DataFrame:
    scores = load_available_scores()
    panels = list(scores.values())
    if "RNA_nearest_delta" in scores and "ATAC_8cell_3pn_chromatin_only_inverse" in scores and "q05_sparse_flipped_activity_sign" in scores:
        panels.append(
            combine_features(
                "available_external_composite_RNA_inverseATAC_motif",
                [scores["RNA_nearest_delta"], scores["ATAC_8cell_3pn_chromatin_only_inverse"], scores["q05_sparse_flipped_activity_sign"]],
                "RNA_ATAC_motif_composite",
                "composite_proxy_methylation_non_leaking_inputs_but_orientation_sensitive",
                "Nearest RNA plus inverse ATAC plus sparse flipped motif signal. Proxy because ATAC orientation is selected from prior diagnostic result.",
            )
        )
    if "RNA_nearest_delta" in scores and "ATAC_8cell_3pn_chromatin_only_inverse" in scores and "region_state_prior" in scores:
        panels.append(
            combine_features(
                "routeC_chromatin_state_proxy_RNA_inverseATAC_region",
                [scores["RNA_nearest_delta"], scores["ATAC_8cell_3pn_chromatin_only_inverse"], scores["region_state_prior"]],
                "RNA_ATAC_region_proxy",
                "routeC_proxy_not_final_biological_control",
                "Composite chromatin-state proxy using available RNA, inverse ATAC orientation, and static region prior.",
            )
        )
    # A deliberately explicit module hypothesis upper bound: if M05/M01/M12/M02 each have a positive biological trigger.
    core_values = {m: 1.0 for m in CORE_MODULES}
    core_values["M10"] = 0.0
    panels.append(
        feature_rows(
            "core_module_positive_control_hypothesis",
            core_values,
            "module_hypothesis",
            "diagnostic_module_hypothesis_upper_bound_not_external_omics",
            "Tests the dynamical consequence if external biology identifies positive triggers for M05/M01/M12/M02.",
        )
    )
    out = pd.concat(panels, ignore_index=True)
    out.to_csv(OUT_FEATURES, sep="\t", index=False)
    return out


def alpha_values() -> list[float]:
    return [round(float(x), 2) for x in np.arange(0.0, 2.0001, 0.05)]


def evaluate_alpha(features: pd.DataFrame) -> pd.DataFrame:
    matrix, ann, pairs = load_inputs()
    mu, sd, components, score_df = fit_latent_basis(matrix, 3)
    coef, _, _ = fit_operator(score_df, pairs, train_stages=set(PRE_MORULA_STAGES), lam=1000.0)
    strict_z = simulate_strict_pre_morula(score_df, ann, coef, n_steps=12)
    obs_z = score_df.loc[stage_ids(ann, "morula")].to_numpy(dtype=float)
    obs_dmr = matrix.loc[stage_ids(ann, "morula")].to_numpy(dtype=float)
    basin = basin_definition(obs_z)
    residual_z = obs_z.mean(axis=0) - strict_z.mean(axis=0)
    rng = np.random.default_rng(20260526)
    rows = []
    for feature_set, sub in features.groupby("feature_set"):
        dirs = sub[["latent_control_PC1", "latent_control_PC2", "latent_control_PC3"]].to_numpy(dtype=float)
        u = pd.to_numeric(sub["control_value_z"], errors="coerce").fillna(0.0).to_numpy(dtype=float)
        control = (dirs * u[:, None]).sum(axis=0)
        leakage = ",".join(sorted(set(map(str, sub["leakage_status"]))))
        modality = ",".join(sorted(set(map(str, sub["control_modality"]))))
        for alpha in alpha_values():
            vec = alpha * control
            pred = strict_z + vec[None, :]
            pred_dmr = decode_latent(pred, mu, sd, components)
            rows.append(
                {
                    "feature_set": feature_set,
                    "alpha": alpha,
                    "control_modality": modality,
                    "leakage_status": leakage,
                    "PC1_control": float(vec[0]),
                    "PC2_control": float(vec[1]),
                    "PC3_control": float(vec[2]),
                    "control_norm": float(np.linalg.norm(vec)),
                    "base_direction_cosine_to_measured_correction": cosine(control, residual_z),
                    "direction_cosine_to_measured_correction": cosine(vec, residual_z) if alpha > 0 else np.nan,
                    "PC3_negative_pull_recovered": float(-vec[2] / (-residual_z[2])) if residual_z[2] < 0 else np.nan,
                    **distribution_metrics(pred, obs_z, pred_dmr, obs_dmr, basin, rng),
                }
            )
    out = pd.DataFrame(rows)
    out.to_csv(OUT_ALPHA, sep="\t", index=False)
    return out


def summarize(alpha: pd.DataFrame) -> pd.DataFrame:
    rows = []
    target = 0.875
    for feature_set, sub in alpha.groupby("feature_set"):
        sub = sub.sort_values("alpha")
        occ = sub["pred_basin_occupancy_q90"].to_numpy(dtype=float)
        alphas = sub["alpha"].to_numpy(dtype=float)
        reached = sub[sub["pred_basin_occupancy_q90"] >= target]
        slopes = np.diff(occ) / np.diff(alphas) if len(alphas) > 1 else np.array([np.nan])
        max_i = int(np.nanargmax(slopes)) if np.isfinite(slopes).any() else 0
        best = sub.iloc[int(np.argmax(occ))]
        rows.append(
            {
                "feature_set": feature_set,
                "leakage_status": best["leakage_status"],
                "base_direction_cosine_to_measured_correction": float(sub["base_direction_cosine_to_measured_correction"].dropna().iloc[0]) if sub["base_direction_cosine_to_measured_correction"].notna().any() else np.nan,
                "alpha_to_observed_occupancy": float(reached.iloc[0]["alpha"]) if len(reached) else np.nan,
                "max_occupancy": float(best["pred_basin_occupancy_q90"]),
                "alpha_at_max_occupancy": float(best["alpha"]),
                "occupancy_at_alpha_1": float(sub.iloc[(sub["alpha"] - 1.0).abs().argsort()].iloc[0]["pred_basin_occupancy_q90"]),
                "cosine_at_alpha_1": float(sub.iloc[(sub["alpha"] - 1.0).abs().argsort()].iloc[0]["direction_cosine_to_measured_correction"]),
                "PC3_recovery_at_alpha_1": float(sub.iloc[(sub["alpha"] - 1.0).abs().argsort()].iloc[0]["PC3_negative_pull_recovered"]),
                "max_local_slope": float(slopes[max_i]) if len(slopes) else np.nan,
                "alpha_at_max_slope_left": float(alphas[max_i]) if len(alphas) else np.nan,
            }
        )
    out = pd.DataFrame(rows).sort_values(["max_occupancy", "base_direction_cosine_to_measured_correction"], ascending=False)
    out.to_csv(OUT_SUMMARY, sep="\t", index=False)
    return out


def make_svg(summary: pd.DataFrame, alpha: pd.DataFrame) -> None:
    chosen = summary.head(8)["feature_set"].tolist()
    colors = ["#245c7a", "#7a3b2e", "#6f8f3a", "#5d4c8c", "#b06b2f", "#2c6f5a", "#8b4c6f", "#555555"]
    width, height = 980, 470
    left, right, top, bottom = 80, 220, 45, 70
    plot_w = width - left - right
    plot_h = height - top - bottom
    lines = [
        f'<svg xmlns="http://www.w3.org/2000/svg" width="{width}" height="{height}" viewBox="0 0 {width} {height}">',
        '<rect width="100%" height="100%" fill="white"/>',
        f'<text x="{left}" y="28" font-family="Arial" font-size="18" font-weight="700">Predicted biological-control alpha scan</text>',
        f'<line x1="{left}" y1="{height-bottom}" x2="{width-right}" y2="{height-bottom}" stroke="#222"/>',
        f'<line x1="{left}" y1="{top}" x2="{left}" y2="{height-bottom}" stroke="#222"/>',
    ]
    for tick in [0.044, 0.3, 0.5, 0.875, 1.0]:
        y = height - bottom - tick * plot_h
        color = "#b23b3b" if tick in [0.044, 0.875] else "#dddddd"
        lines.append(f'<line x1="{left}" y1="{y:.2f}" x2="{width-right}" y2="{y:.2f}" stroke="{color}" stroke-dasharray="4 4"/>')
        lines.append(f'<text x="{left-8}" y="{y+4:.2f}" text-anchor="end" font-family="Arial" font-size="10">{tick:.3g}</text>')
    xmin, xmax = float(alpha["alpha"].min()), float(alpha["alpha"].max())
    for i, fs in enumerate(chosen):
        sub = alpha[alpha["feature_set"] == fs].sort_values("alpha")
        pts = []
        for row in sub.itertuples():
            x = left + (row.alpha - xmin) / (xmax - xmin) * plot_w
            y = height - bottom - float(row.pred_basin_occupancy_q90) * plot_h
            pts.append((x, y))
        color = colors[i % len(colors)]
        lines.append('<polyline points="' + " ".join(f"{x:.2f},{y:.2f}" for x, y in pts) + f'" fill="none" stroke="{color}" stroke-width="2.1"/>')
        label = fs.replace("_", " ")[:38]
        lines.append(f'<rect x="{width-right+15}" y="{top+i*25}" width="12" height="12" fill="{color}"/>')
        lines.append(f'<text x="{width-right+33}" y="{top+i*25+11}" font-family="Arial" font-size="11">{label}</text>')
    lines.append(f'<text x="{left}" y="{height-22}" font-family="Arial" font-size="12">Red guides: methylation-only baseline and observed morula occupancy.</text>')
    lines.append("</svg>")
    OUT_SVG.write_text("\n".join(lines), encoding="utf-8")


def write_doc(summary: pd.DataFrame) -> None:
    top = summary.head(8)
    lines = [
        "# Composite Biocontrol Proxy Alpha Scan",
        "",
        "Status: `completed`",
        "",
        "This is route C: it does not replace missing histone-state data, but tests whether available RNA/ATAC/motif/region proxy composites can approximate a module-specific biological control term.",
        "",
        "Important boundary: models marked as `orientation_corrected_proxy`, `routeC_proxy`, or `module_hypothesis_upper_bound` are not final non-leaking biological u_bio. They are attack-mode diagnostics for what must be achieved by real histone/chromatin inputs.",
        "",
        "## Top Alpha-Scan Results",
        "",
    ]
    for row in top.itertuples():
        lines.append(
            f"- {row.feature_set}: max_occ={row.max_occupancy:.3f} at alpha={row.alpha_at_max_occupancy:.2f}; "
            f"occ@1={row.occupancy_at_alpha_1:.3f}; cosine@1={row.cosine_at_alpha_1:.3f}; PC3@1={row.PC3_recovery_at_alpha_1:.3f}; status={row.leakage_status}"
        )
    lines += [
        "",
        "## Interpretation",
        "",
        "If a proxy reaches high occupancy only after orientation correction or module-hypothesis encoding, it does not prove u_bio. It defines the required direction and module pattern for the histone/chromatin data acquisition stage.",
    ]
    OUT_DOC.write_text("\n".join(lines) + "\n", encoding="utf-8")


def main() -> None:
    RESULTS.mkdir(parents=True, exist_ok=True)
    FIGURES.mkdir(parents=True, exist_ok=True)
    DOCS.mkdir(parents=True, exist_ok=True)
    features = build_feature_panel()
    alpha = evaluate_alpha(features)
    summary = summarize(alpha)
    make_svg(summary, alpha)
    write_doc(summary)
    OUT_MANIFEST.write_text(
        json.dumps(
            {
                "status": "completed",
                "outputs": [str(OUT_FEATURES), str(OUT_ALPHA), str(OUT_SUMMARY), str(OUT_SVG), str(OUT_DOC)],
                "boundary": "route C proxy and orientation-corrected models are diagnostics, not final u_bio",
            },
            indent=2,
        )
        + "\n",
        encoding="utf-8",
    )
    print(json.dumps({"status": "completed", "top": summary.head(8).to_dict(orient="records")}, indent=2))


if __name__ == "__main__":
    main()
```
