## Supplemental code for "An entropy-based diagnostic framework for characterizing methylation state dynamics during preimplantation development": run_dmr_operator_time_dynamics.py

```
from __future__ import annotations

import csv
import gzip
import json
import math
import os
from collections import defaultdict
from pathlib import Path

import numpy as np
import pandas as pd


BASE = Path(r"E:\CSB_TRO_operator_time_DMR_dynamics_2026-05-25")
RESULTS = BASE / "results"
FIGURES = BASE / "figures"
DOCS = BASE / "docs"
META = BASE / "metadata"

SRC_DYN = Path(r"E:\实验更新_5_22\csb_tro_dynamics")
SRC_REGIONS = Path(r"E:\实验更新_5_22\processed\GSE81233_strong_controls\sample_region_metrics")
SRC_OPERATOR = Path(r"E:\CSB_TRO_operator_time_2026-05-25\results")

STAGES = ["MII oocyte", "zygote/PN", "2-cell", "4-cell", "8-cell", "morula", "blastocyst"]
TAU = {stage: i / (len(STAGES) - 1) for i, stage in enumerate(STAGES)}
STAGE_ORDER = {stage: i for i, stage in enumerate(STAGES)}
DT = 1.0 / (len(STAGES) - 1)
FEATURES = ["intercept", "tau", "m_j", "A", "P", "Hm", "Hr"]


def ensure_dirs() -> None:
    for path in [RESULTS, FIGURES, DOCS, META]:
        path.mkdir(parents=True, exist_ok=True)


def fnum(x, default=np.nan) -> float:
    try:
        if x is None or x == "":
            return default
        return float(x)
    except (TypeError, ValueError):
        return default


def read_tsv(path: Path):
    with open(path, "r", newline="", encoding="utf-8") as handle:
        yield from csv.DictReader(handle, delimiter="\t")


def write_json(path: Path, obj) -> None:
    with open(path, "w", encoding="utf-8") as handle:
        json.dump(obj, handle, indent=2, ensure_ascii=False)


def safe_corr(a: np.ndarray, b: np.ndarray) -> float:
    mask = np.isfinite(a) & np.isfinite(b)
    if mask.sum() < 3:
        return float("nan")
    aa = a[mask] - np.mean(a[mask])
    bb = b[mask] - np.mean(b[mask])
    den = math.sqrt(float(np.dot(aa, aa) * np.dot(bb, bb)))
    return float(np.dot(aa, bb) / den) if den else float("nan")


def rmse(a: np.ndarray, b: np.ndarray) -> float:
    mask = np.isfinite(a) & np.isfinite(b)
    if not mask.any():
        return float("nan")
    return float(np.sqrt(np.mean((a[mask] - b[mask]) ** 2)))


def load_state_samples() -> pd.DataFrame:
    state = pd.read_csv(SRC_DYN / "CSB_TRO_state_samples.tsv", sep="\t")
    state = state[state["stage"].isin(STAGES)].copy()
    state["tau"] = state["stage"].map(TAU)
    state["stage_order"] = state["stage"].map(STAGE_ORDER)
    return state


def load_particles() -> pd.DataFrame:
    path = SRC_OPERATOR / "CSB_TRO_operator_time_particles.tsv"
    if not path.exists():
        path = SRC_DYN / "CSB_TRO_fused_product_particles.tsv"
    particles = pd.read_csv(path, sep="\t")
    particles = particles[particles["stage"].isin(STAGES)].copy()
    particles["tau"] = particles["stage"].map(TAU)
    return particles


def build_dmr_matrix(state: pd.DataFrame) -> tuple[pd.DataFrame, pd.DataFrame, pd.DataFrame]:
    wanted_samples = set(state["sample_id"].astype(str))
    records = []
    meta_records = {}
    coverage_records = []
    region_files = sorted(SRC_REGIONS.glob("*.regions.tsv.gz"))
    for path in region_files:
        with gzip.open(path, "rt", encoding="utf-8") as handle:
            reader = csv.DictReader(handle, delimiter="\t")
            for row in reader:
                if row["region_type"] != "age_DMR":
                    continue
                sample_id = row["sample_id"]
                if sample_id not in wanted_samples:
                    continue
                cluster = row["cluster_name"]
                total_reads = fnum(row["total_reads"], 0.0)
                beta = fnum(row["beta"], np.nan)
                beta_observed = np.isfinite(beta) and total_reads > 0
                records.append({
                    "sample_id": sample_id,
                    "stage": row["stage"],
                    "cluster_name": cluster,
                    "beta": beta if beta_observed else np.nan,
                    "total_reads": total_reads,
                    "met_reads": fnum(row["met_reads"], np.nan),
                    "observed": int(beta_observed),
                })
                if cluster not in meta_records:
                    meta_records[cluster] = {
                        "cluster_name": cluster,
                        "region_type": row["region_type"],
                        "control_set": row["control_set"],
                        "matched_age_cluster": row["matched_age_cluster"],
                        "chr": row["chr"],
                        "start": int(fnum(row["start"], 0)),
                        "end": int(fnum(row["end"], 0)),
                        "width": int(fnum(row["width"], 0)),
                        "n_cpg_target": int(fnum(row["n_cpg_target"], 0)),
                        "age_weight_5yr": fnum(row["age_weight_5yr"], np.nan),
                    }
    long = pd.DataFrame.from_records(records)
    if long.empty:
        raise RuntimeError("No age_DMR region metrics were found for CSB-TRO samples.")
    matrix_raw = long.pivot_table(index="sample_id", columns="cluster_name", values="beta", aggfunc="mean")
    observed = long.pivot_table(index="sample_id", columns="cluster_name", values="observed", aggfunc="max").fillna(0)
    meta = pd.DataFrame.from_dict(meta_records, orient="index").reset_index(drop=True)
    meta = meta.sort_values(["chr", "start", "end", "cluster_name"])
    clusters = list(meta["cluster_name"])
    matrix_raw = matrix_raw.reindex(columns=clusters)
    observed = observed.reindex(index=matrix_raw.index, columns=clusters).fillna(0)

    sample_stage = state.drop_duplicates("sample_id").set_index("sample_id")["stage"].to_dict()
    matrix = matrix_raw.copy()
    for cluster in clusters:
        global_median = float(matrix[cluster].median()) if np.isfinite(matrix[cluster].median()) else 0.5
        for stage in STAGES:
            idx = [s for s in matrix.index if sample_stage.get(s) == stage]
            if not idx:
                continue
            stage_median = matrix.loc[idx, cluster].median()
            fill_value = float(stage_median) if np.isfinite(stage_median) else global_median
            matrix.loc[idx, cluster] = matrix.loc[idx, cluster].fillna(fill_value)
        matrix[cluster] = matrix[cluster].fillna(global_median)
    matrix = matrix.sort_index()
    observed = observed.reindex(index=matrix.index, columns=clusters).fillna(0)

    cov = observed.mean(axis=1).rename("observed_region_fraction").reset_index()
    cov["n_observed_regions"] = observed.sum(axis=1).astype(int).values
    cov["n_regions"] = len(clusters)
    coverage_records = cov

    matrix_out = matrix.reset_index()
    matrix_out.to_csv(RESULTS / "CSB_TRO_DMR_state_matrix.tsv", sep="\t", index=False)
    meta.to_csv(RESULTS / "CSB_TRO_DMR_metadata.tsv", sep="\t", index=False)
    coverage_records.to_csv(META / "CSB_TRO_DMR_matrix_coverage.tsv", sep="\t", index=False)
    return matrix, meta, observed


def write_sample_tau_annotation(state: pd.DataFrame, observed: pd.DataFrame) -> pd.DataFrame:
    cov = pd.DataFrame({
        "sample_id": observed.index,
        "n_observed_regions": observed.sum(axis=1).astype(int).values,
        "observed_region_fraction": observed.mean(axis=1).values,
    })
    ann = state[["sample_id", "stage", "stage_order", "tau", "A", "P", "Hm", "Hr", "n_age_regions_used"]].drop_duplicates("sample_id")
    ann = ann.merge(cov, on="sample_id", how="left")
    ann = ann.sort_values(["stage_order", "sample_id"])
    ann.to_csv(RESULTS / "CSB_TRO_sample_tau_annotation.tsv", sep="\t", index=False)
    return ann


def aggregate_sample_couplings(particles: pd.DataFrame) -> pd.DataFrame:
    pmap = particles.set_index("particle_id")[["dna_sample_id", "stage"]].to_dict("index")
    accum = defaultdict(lambda: {"weight": 0.0, "n_particle_couplings": 0})
    with open(SRC_DYN / "CSB_TRO_path_space_transition_couplings.tsv", "r", newline="", encoding="utf-8") as handle:
        for row in csv.DictReader(handle, delimiter="\t"):
            fp = pmap.get(row["from_particle_id"])
            tp = pmap.get(row["to_particle_id"])
            if fp is None or tp is None:
                continue
            from_stage = row["from_stage"]
            to_stage = row["to_stage"]
            if from_stage not in STAGES or to_stage not in STAGES:
                continue
            key = (from_stage, to_stage, fp["dna_sample_id"], tp["dna_sample_id"])
            accum[key]["weight"] += fnum(row["probability"], 0.0)
            accum[key]["n_particle_couplings"] += 1
    rows = []
    for (from_stage, to_stage, from_sample, to_sample), val in accum.items():
        rows.append({
            "from_stage": from_stage,
            "to_stage": to_stage,
            "from_sample_id": from_sample,
            "to_sample_id": to_sample,
            "from_tau": TAU[from_stage],
            "to_tau": TAU[to_stage],
            "delta_tau": TAU[to_stage] - TAU[from_stage],
            "sample_coupling_weight": val["weight"],
            "n_particle_couplings": val["n_particle_couplings"],
        })
    pairs = pd.DataFrame(rows)
    pairs = pairs[pairs["delta_tau"] > 0].copy()
    pairs = pairs.sort_values(["from_tau", "from_sample_id", "to_sample_id"])
    pairs.to_csv(RESULTS / "CSB_TRO_OT_sample_transition_couplings.tsv", sep="\t", index=False)
    return pairs


def write_transition_training_pairs(pairs: pd.DataFrame, matrix: pd.DataFrame, clusters: list[str]) -> pd.DataFrame:
    valid_samples = set(matrix.index)
    pairs = pairs[pairs["from_sample_id"].isin(valid_samples) & pairs["to_sample_id"].isin(valid_samples)].copy()
    long_rows = []
    out_path = RESULTS / "CSB_TRO_OT_transition_training_pairs.tsv"
    with open(out_path, "w", newline="", encoding="utf-8") as handle:
        fields = [
            "from_stage", "to_stage", "from_sample_id", "to_sample_id", "from_tau", "to_tau",
            "delta_tau", "sample_coupling_weight", "cluster_name", "m_from", "m_to", "velocity_target",
        ]
        writer = csv.DictWriter(handle, fieldnames=fields, delimiter="\t")
        writer.writeheader()
        for _, pair in pairs.iterrows():
            vf = matrix.loc[pair["from_sample_id"], clusters].to_numpy(dtype=float)
            vt = matrix.loc[pair["to_sample_id"], clusters].to_numpy(dtype=float)
            vel = (vt - vf) / float(pair["delta_tau"])
            for cluster, m_from, m_to, v in zip(clusters, vf, vt, vel):
                writer.writerow({
                    "from_stage": pair["from_stage"],
                    "to_stage": pair["to_stage"],
                    "from_sample_id": pair["from_sample_id"],
                    "to_sample_id": pair["to_sample_id"],
                    "from_tau": pair["from_tau"],
                    "to_tau": pair["to_tau"],
                    "delta_tau": pair["delta_tau"],
                    "sample_coupling_weight": pair["sample_coupling_weight"],
                    "cluster_name": cluster,
                    "m_from": m_from,
                    "m_to": m_to,
                    "velocity_target": v,
                })
    return pairs


def fit_velocity_model(
    pairs: pd.DataFrame,
    matrix: pd.DataFrame,
    ann: pd.DataFrame,
    clusters: list[str],
    excluded_stages: set[str] | None = None,
    random_tau: bool = False,
    random_to: bool = False,
    seed: int = 13,
    ridge_lambda: float = 50.0,
) -> pd.DataFrame:
    rng = np.random.default_rng(seed)
    train = pairs.copy()
    if excluded_stages:
        train = train[~train["from_stage"].isin(excluded_stages) & ~train["to_stage"].isin(excluded_stages)].copy()
    if random_to:
        shuffled = train["to_sample_id"].to_numpy().copy()
        rng.shuffle(shuffled)
        train["to_sample_id"] = shuffled
        train = train[train["to_sample_id"].isin(matrix.index)].copy()
    sample_features = ann.set_index("sample_id")[["A", "P", "Hm", "Hr"]].astype(float)
    n = len(train)
    if n < 5:
        raise RuntimeError("Too few transition pairs for velocity model fitting.")
    tau = train["from_tau"].to_numpy(dtype=float)
    if random_tau:
        tau = tau.copy()
        rng.shuffle(tau)
    aux = sample_features.loc[train["from_sample_id"]].to_numpy(dtype=float)
    weights = np.maximum(train["sample_coupling_weight"].to_numpy(dtype=float), 1e-12)
    weight_scale = weights / np.mean(weights)

    coeff_rows = []
    for cluster in clusters:
        m_from = matrix.loc[train["from_sample_id"], cluster].to_numpy(dtype=float)
        m_to = matrix.loc[train["to_sample_id"], cluster].to_numpy(dtype=float)
        y = (m_to - m_from) / train["delta_tau"].to_numpy(dtype=float)
        x = np.column_stack([np.ones(n), tau, m_from, aux])
        sw = np.sqrt(weight_scale)
        xw = x * sw[:, None]
        yw = y * sw
        penalty = ridge_lambda * np.eye(x.shape[1])
        penalty[0, 0] = 0.0
        beta = np.linalg.solve(xw.T @ xw + penalty, xw.T @ yw)
        pred = x @ beta
        coeff_rows.append({
            "cluster_name": cluster,
            **{f"coef_{name}": beta[i] for i, name in enumerate(FEATURES)},
            "train_n_pairs": n,
            "train_weight_sum": float(weights.sum()),
            "train_rmse": rmse(pred, y),
            "train_correlation": safe_corr(pred, y),
            "ridge_lambda": ridge_lambda,
        })
    return pd.DataFrame(coeff_rows)


def predict_stage(
    model: pd.DataFrame,
    matrix: pd.DataFrame,
    ann: pd.DataFrame,
    from_stage: str,
    target_stage: str,
    clusters: list[str],
    label: str,
) -> tuple[pd.DataFrame, dict]:
    sample_features = ann.set_index("sample_id")[["stage", "A", "P", "Hm", "Hr"]]
    from_samples = list(sample_features[sample_features["stage"] == from_stage].index)
    target_samples = list(sample_features[sample_features["stage"] == target_stage].index)
    dt = TAU[target_stage] - TAU[from_stage]
    coef = model.set_index("cluster_name").loc[clusters]
    rows = []
    pred_matrix = []
    for sample in from_samples:
        aux = sample_features.loc[sample, ["A", "P", "Hm", "Hr"]].astype(float).to_numpy()
        current = matrix.loc[sample, clusters].to_numpy(dtype=float)
        pred = []
        for j, cluster in enumerate(clusters):
            c = coef.loc[cluster, [f"coef_{name}" for name in FEATURES]].to_numpy(dtype=float)
            x = np.array([1.0, TAU[from_stage], current[j], *aux], dtype=float)
            v = float(x @ c)
            pred.append(min(1.0, max(0.0, current[j] + dt * v)))
        pred = np.array(pred)
        pred_matrix.append(pred)
        for cluster, value in zip(clusters, pred):
            rows.append({
                "prediction_label": label,
                "from_stage": from_stage,
                "target_stage": target_stage,
                "from_sample_id": sample,
                "cluster_name": cluster,
                "predicted_beta": value,
            })
    pred_df = pd.DataFrame(rows)
    pred_mean = np.mean(np.vstack(pred_matrix), axis=0)
    obs_mean = matrix.loc[target_samples, clusters].mean(axis=0).to_numpy(dtype=float)
    from_mean = matrix.loc[from_samples, clusters].mean(axis=0).to_numpy(dtype=float)
    age_weights = coef.index.to_series().map(lambda c: 1.0).to_numpy(dtype=float)
    metrics = {
        "prediction_label": label,
        "from_stage": from_stage,
        "target_stage": target_stage,
        "n_from_samples": len(from_samples),
        "n_target_samples": len(target_samples),
        "rmse_predicted_vs_observed_stage_mean": rmse(pred_mean, obs_mean),
        "rmse_from_baseline_vs_observed_stage_mean": rmse(from_mean, obs_mean),
        "correlation_predicted_vs_observed_stage_mean": safe_corr(pred_mean, obs_mean),
        "mean_beta_predicted": float(np.mean(pred_mean)),
        "mean_beta_observed": float(np.mean(obs_mean)),
        "mean_beta_from_stage": float(np.mean(from_mean)),
    }
    return pred_df, metrics


def stage_means(matrix: pd.DataFrame, ann: pd.DataFrame, clusters: list[str]) -> pd.DataFrame:
    rows = []
    sample_stage = ann.set_index("sample_id")["stage"]
    for stage in STAGES:
        samples = list(sample_stage[sample_stage == stage].index)
        if not samples:
            continue
        vals = matrix.loc[samples, clusters].mean(axis=0)
        for cluster in clusters:
            rows.append({"stage": stage, "tau": TAU[stage], "cluster_name": cluster, "mean_beta": vals[cluster]})
    df = pd.DataFrame(rows)
    df.to_csv(RESULTS / "CSB_TRO_DMR_stage_mean_trajectory.tsv", sep="\t", index=False)
    return df


def dynamic_dmr_ranking(stage_mean: pd.DataFrame, model: pd.DataFrame, meta: pd.DataFrame, clusters: list[str]) -> pd.DataFrame:
    pivot = stage_mean.pivot(index="cluster_name", columns="stage", values="mean_beta").reindex(index=clusters)
    rank_rows = []
    coef = model.set_index("cluster_name")
    for cluster in clusters:
        vals = pivot.loc[cluster]
        velocities = {}
        for a, b in zip(STAGES[:-1], STAGES[1:]):
            velocities[f"velocity_{a}_to_{b}".replace(" ", "_").replace("/", "_")] = (vals[b] - vals[a]) / DT
        entry = velocities["velocity_8-cell_to_morula"]
        exit_v = velocities["velocity_morula_to_blastocyst"]
        acc = (vals["blastocyst"] - 2.0 * vals["morula"] + vals["8-cell"]) / (DT * DT)
        max_velocity = max(abs(v) for v in velocities.values())
        pred_importance = abs(float(coef.loc[cluster, "coef_m_j"]))
        reset_score = abs(entry) + 0.5 * abs(acc) + 0.25 * pred_importance
        row = {
            "cluster_name": cluster,
            "max_velocity": max_velocity,
            "morula_entry_velocity": entry,
            "blastocyst_exit_velocity": exit_v,
            "morula_acceleration": acc,
            "prediction_importance_abs_coef_m_j": pred_importance,
            "reset_dynamic_score": reset_score,
            "beta_8cell": vals["8-cell"],
            "beta_morula": vals["morula"],
            "beta_blastocyst": vals["blastocyst"],
            **velocities,
        }
        rank_rows.append(row)
    ranking = pd.DataFrame(rank_rows)
    ranking = ranking.merge(meta, on="cluster_name", how="left")
    ranking = ranking.sort_values("reset_dynamic_score", ascending=False)
    ranking["dynamic_rank"] = np.arange(1, len(ranking) + 1)
    ranking.to_csv(RESULTS / "CSB_TRO_dynamic_DMR_ranking.tsv", sep="\t", index=False)
    ranking.head(100).to_csv(RESULTS / "CSB_TRO_top100_dynamic_reset_DMRs.tsv", sep="\t", index=False)
    ranking.sort_values("morula_entry_velocity", key=lambda s: np.abs(s), ascending=False).head(100).to_csv(
        RESULTS / "CSB_TRO_top100_morula_entry_DMRs.tsv", sep="\t", index=False
    )
    ranking.sort_values("blastocyst_exit_velocity", key=lambda s: np.abs(s), ascending=False).head(100).to_csv(
        RESULTS / "CSB_TRO_top100_blastocyst_exit_DMRs.tsv", sep="\t", index=False
    )
    return ranking


def null_and_ablation(
    pairs: pd.DataFrame,
    matrix: pd.DataFrame,
    ann: pd.DataFrame,
    clusters: list[str],
    main_metrics: list[dict],
    ranking: pd.DataFrame,
) -> tuple[pd.DataFrame, pd.DataFrame]:
    rows = []
    for m in main_metrics:
        rows.append({"experiment": "main_model", **{k: v for k, v in m.items() if not k.endswith("_vector")}})
    for label, kwargs in [
        ("random_tau_null", {"random_tau": True, "seed": 101}),
        ("random_coupling_null", {"random_to": True, "seed": 202}),
    ]:
        model = fit_velocity_model(pairs, matrix, ann, clusters, excluded_stages={"morula"}, **kwargs)
        _, met = predict_stage(model, matrix, ann, "8-cell", "morula", clusters, label)
        rows.append({"experiment": label, **met})
    # Low-dimensional/no-DMR baseline: carry forward the previous stage mean velocity, 4-cell -> 8-cell, to morula.
    sample_stage = ann.set_index("sample_id")["stage"]
    s4 = list(sample_stage[sample_stage == "4-cell"].index)
    s8 = list(sample_stage[sample_stage == "8-cell"].index)
    sm = list(sample_stage[sample_stage == "morula"].index)
    four = matrix.loc[s4, clusters].mean(axis=0).to_numpy(dtype=float)
    eight = matrix.loc[s8, clusters].mean(axis=0).to_numpy(dtype=float)
    morula = matrix.loc[sm, clusters].mean(axis=0).to_numpy(dtype=float)
    pred_low = np.clip(eight + (eight - four), 0.0, 1.0)
    rows.append({
        "experiment": "stage_mean_previous_velocity_baseline",
        "prediction_label": "stage_mean_previous_velocity_baseline",
        "from_stage": "8-cell",
        "target_stage": "morula",
        "n_from_samples": len(s8),
        "n_target_samples": len(sm),
        "rmse_predicted_vs_observed_stage_mean": rmse(pred_low, morula),
        "rmse_from_baseline_vs_observed_stage_mean": rmse(eight, morula),
        "correlation_predicted_vs_observed_stage_mean": safe_corr(pred_low, morula),
        "mean_beta_predicted": float(np.mean(pred_low)),
        "mean_beta_observed": float(np.mean(morula)),
        "mean_beta_from_stage": float(np.mean(eight)),
    })
    ablation = pd.DataFrame(rows)
    ablation.to_csv(RESULTS / "CSB_TRO_ablation_results.tsv", sep="\t", index=False)

    top = list(ranking.head(20)["cluster_name"])
    cf_rows = []
    for top_n in [10, 20, 50]:
        top_clusters = list(ranking.head(top_n)["cluster_name"])
        obs = matrix.loc[sm, clusters].mean(axis=0).to_numpy(dtype=float)
        start = matrix.loc[s8, clusters].mean(axis=0).to_numpy(dtype=float)
        full_pred = np.array(main_metrics[0]["predicted_morula_mean_vector"])
        cf = full_pred.copy()
        idx = [clusters.index(c) for c in top_clusters if c in clusters]
        cf[idx] = start[idx]
        cf_rows.append({
            "counterfactual": f"fix_top{top_n}_dynamic_DMR_at_8cell_mean",
            "top_n": top_n,
            "target_stage": "morula",
            "rmse_full_prediction": rmse(full_pred, obs),
            "rmse_counterfactual": rmse(cf, obs),
            "delta_rmse_counterfactual_minus_full": rmse(cf, obs) - rmse(full_pred, obs),
            "mean_abs_shift_suppressed": float(np.mean(np.abs(full_pred[idx] - start[idx]))) if idx else float("nan"),
        })
    cf = pd.DataFrame(cf_rows)
    cf.to_csv(RESULTS / "CSB_TRO_counterfactual_topDMR_results.tsv", sep="\t", index=False)
    return ablation, cf


def svg_bar(path: Path, labels: list[str], values: list[float], title: str, ylabel: str) -> None:
    width, height = 820, 460
    left, top, bottom, right = 90, 50, 110, 30
    vmax = max(values) if values else 1.0
    vmax = vmax * 1.15 if vmax > 0 else 1.0
    bar_w = (width - left - right) / max(len(values), 1) * 0.68
    gap = (width - left - right) / max(len(values), 1)
    out = [
        f'<svg xmlns="http://www.w3.org/2000/svg" width="{width}" height="{height}" viewBox="0 0 {width} {height}">',
        '<rect width="100%" height="100%" fill="white"/>',
        f'<text x="{left}" y="30" font-family="Arial" font-size="20" font-weight="700">{title}</text>',
        f'<line x1="{left}" y1="{height-bottom}" x2="{width-right}" y2="{height-bottom}" stroke="#222"/>',
        f'<line x1="{left}" y1="{top}" x2="{left}" y2="{height-bottom}" stroke="#222"/>',
        f'<text x="22" y="{height/2}" transform="rotate(-90 22 {height/2})" font-family="Arial" font-size="13">{ylabel}</text>',
    ]
    for i, (label, value) in enumerate(zip(labels, values)):
        x = left + i * gap + gap * 0.16
        h = (value / vmax) * (height - top - bottom)
        y = height - bottom - h
        out.append(f'<rect x="{x:.2f}" y="{y:.2f}" width="{bar_w:.2f}" height="{h:.2f}" fill="#4777b3"/>')
        out.append(f'<text x="{x + bar_w/2:.2f}" y="{y-6:.2f}" text-anchor="middle" font-family="Arial" font-size="11">{value:.3f}</text>')
        out.append(f'<text x="{x + bar_w/2:.2f}" y="{height-bottom+18}" text-anchor="end" transform="rotate(-35 {x + bar_w/2:.2f} {height-bottom+18})" font-family="Arial" font-size="11">{label}</text>')
    out.append("</svg>")
    path.write_text("\n".join(out), encoding="utf-8")


def svg_scatter(path: Path, obs: np.ndarray, pred: np.ndarray, title: str) -> None:
    width, height = 560, 540
    left, top, bottom, right = 70, 45, 65, 30
    vals = np.concatenate([obs, pred])
    lo, hi = float(np.min(vals)), float(np.max(vals))
    pad = (hi - lo) * 0.08 or 0.1
    lo, hi = max(0.0, lo - pad), min(1.0, hi + pad)
    def xmap(x): return left + (x - lo) / (hi - lo) * (width - left - right)
    def ymap(y): return height - bottom - (y - lo) / (hi - lo) * (height - top - bottom)
    out = [
        f'<svg xmlns="http://www.w3.org/2000/svg" width="{width}" height="{height}" viewBox="0 0 {width} {height}">',
        '<rect width="100%" height="100%" fill="white"/>',
        f'<text x="{left}" y="28" font-family="Arial" font-size="18" font-weight="700">{title}</text>',
        f'<line x1="{left}" y1="{height-bottom}" x2="{width-right}" y2="{height-bottom}" stroke="#222"/>',
        f'<line x1="{left}" y1="{top}" x2="{left}" y2="{height-bottom}" stroke="#222"/>',
        f'<line x1="{xmap(lo)}" y1="{ymap(lo)}" x2="{xmap(hi)}" y2="{ymap(hi)}" stroke="#999" stroke-dasharray="5,5"/>',
    ]
    for x, y in zip(obs, pred):
        out.append(f'<circle cx="{xmap(x):.2f}" cy="{ymap(y):.2f}" r="3" fill="#4777b3" opacity="0.72"/>')
    out.append(f'<text x="{width/2}" y="{height-18}" text-anchor="middle" font-family="Arial" font-size="13">observed target-stage DMR mean beta</text>')
    out.append(f'<text x="20" y="{height/2}" transform="rotate(-90 20 {height/2})" text-anchor="middle" font-family="Arial" font-size="13">predicted DMR mean beta</text>')
    out.append("</svg>")
    path.write_text("\n".join(out), encoding="utf-8")


def write_docs(summary: dict) -> None:
    text = f"""# CSB-TRO DMR-level predictive operator-time dynamics

This workspace upgrades the stage-level operator-time trajectory into a DMR-level predictive dynamics experiment. The model is still stage-anchored pseudo-time, not true longitudinal tracking of the same embryo.

## What was built

- A sample x age-DMR methylation matrix from GSE81233 region metrics.
- Sample-level tau annotations for the CSB-TRO developmental stages.
- Sample-level OT transition training pairs aggregated from particle-level path-space couplings.
- A transparent ridge baseline velocity model for each DMR: v_j = beta0 + beta1 tau + beta2 m_j + beta3 A + beta4 P + beta5 Hm + beta6 Hr.
- Forward prediction tests for morula and blastocyst.
- Dynamic DMR ranking, null/ablation baselines, and in silico top-DMR fixation sensitivity.

## Key scale

- Samples in DMR matrix: {summary['n_samples']}
- age-DMR dimensions: {summary['n_dmrs']}
- sample-level OT transition pairs: {summary['n_sample_pairs']}
- long DMR transition rows: {summary['n_sample_pairs'] * summary['n_dmrs']}

## Predictive validation result

- Strict leave-morula-out prediction RMSE: {summary['morula_prediction_rmse']:.4f}
- Strict leave-blastocyst-out prediction RMSE: {summary['blastocyst_prediction_rmse']:.4f}
- Operator-fit morula RMSE when the 8-cell -> morula transition is included: {summary['operator_fit_morula_rmse']:.4f}
- Operator-fit blastocyst RMSE when the morula -> blastocyst transition is included: {summary['operator_fit_blastocyst_rmse']:.4f}

## Interpretation

The model now tests whether DMR-level methylation state at an earlier developmental operator time can predict the next reset-basin state. The current baseline can represent observed operator-time transitions when the relevant transition is included. However, strict leave-morula-out forward prediction does not yet beat the simple 8-cell baseline, so morula emergence is not solved as an out-of-sample predictive problem.

This is a stronger experimental system than trajectory visualization, but it remains a baseline predictive dynamics model. It should not be described as a full stochastic differential equation or solved Fokker-Planck model yet.

## Next validation priority

The next strongest additions are external RNA/motif annotation of top dynamic DMRs, then adjacent-stage ATAC/H3K27ac validation where processed signal exists.
"""
    (DOCS / "CSB_TRO_DMR_dynamics_interpretation.md").write_text(text, encoding="utf-8")


def main() -> None:
    ensure_dirs()
    state = load_state_samples()
    particles = load_particles()
    matrix, meta, observed = build_dmr_matrix(state)
    clusters = list(meta["cluster_name"])
    ann = write_sample_tau_annotation(state, observed)
    pairs = aggregate_sample_couplings(particles)
    pairs = write_transition_training_pairs(pairs, matrix, clusters)

    full_model = fit_velocity_model(pairs, matrix, ann, clusters)
    full_model.to_csv(RESULTS / "CSB_TRO_velocity_model_coefficients.tsv", sep="\t", index=False)
    write_json(RESULTS / "CSB_TRO_velocity_model_metadata.json", {
        "model": "per-DMR weighted ridge regression",
        "features": FEATURES,
        "ridge_lambda": 50.0,
        "equation": "v_j = beta0 + beta1*tau + beta2*m_j + beta3*A + beta4*P + beta5*Hm + beta6*Hr",
    })

    fit_morula_pred, fit_morula_metrics = predict_stage(full_model, matrix, ann, "8-cell", "morula", clusters, "operator_fit_morula_transition_included")
    fit_blast_pred, fit_blast_metrics = predict_stage(full_model, matrix, ann, "morula", "blastocyst", clusters, "operator_fit_blastocyst_transition_included")
    fit_morula_pred.to_csv(RESULTS / "CSB_TRO_forward_prediction_morula_operator_fit.tsv", sep="\t", index=False)
    fit_blast_pred.to_csv(RESULTS / "CSB_TRO_forward_prediction_blastocyst_operator_fit.tsv", sep="\t", index=False)

    morula_model = fit_velocity_model(pairs, matrix, ann, clusters, excluded_stages={"morula"})
    morula_pred, morula_metrics = predict_stage(morula_model, matrix, ann, "8-cell", "morula", clusters, "leave_morula_out")
    blast_model = fit_velocity_model(pairs, matrix, ann, clusters, excluded_stages={"blastocyst"})
    blast_pred, blast_metrics = predict_stage(blast_model, matrix, ann, "morula", "blastocyst", clusters, "leave_blastocyst_out")
    morula_pred.to_csv(RESULTS / "CSB_TRO_forward_prediction_morula.tsv", sep="\t", index=False)
    blast_pred.to_csv(RESULTS / "CSB_TRO_forward_prediction_blastocyst.tsv", sep="\t", index=False)

    stage_mean = stage_means(matrix, ann, clusters)
    ranking = dynamic_dmr_ranking(stage_mean, full_model, meta, clusters)

    sample_stage = ann.set_index("sample_id")["stage"]
    obs_morula = matrix.loc[list(sample_stage[sample_stage == "morula"].index), clusters].mean(axis=0).to_numpy(dtype=float)
    pred_morula_mean = morula_pred.pivot_table(index="from_sample_id", columns="cluster_name", values="predicted_beta").reindex(columns=clusters).mean(axis=0).to_numpy(dtype=float)
    obs_blast = matrix.loc[list(sample_stage[sample_stage == "blastocyst"].index), clusters].mean(axis=0).to_numpy(dtype=float)
    pred_blast_mean = blast_pred.pivot_table(index="from_sample_id", columns="cluster_name", values="predicted_beta").reindex(columns=clusters).mean(axis=0).to_numpy(dtype=float)
    morula_metrics["predicted_morula_mean_vector"] = pred_morula_mean.tolist()
    metrics_table = pd.DataFrame([
        {k: v for k, v in morula_metrics.items() if k != "predicted_morula_mean_vector"},
        blast_metrics,
        fit_morula_metrics,
        fit_blast_metrics,
    ])
    metrics_table.to_csv(RESULTS / "CSB_TRO_forward_prediction_metrics.tsv", sep="\t", index=False)
    write_json(RESULTS / "CSB_TRO_forward_prediction_metrics.json", {
        "leave_morula_out": {k: v for k, v in morula_metrics.items() if k != "predicted_morula_mean_vector"},
        "leave_blastocyst_out": blast_metrics,
        "operator_fit_morula_transition_included": fit_morula_metrics,
        "operator_fit_blastocyst_transition_included": fit_blast_metrics,
    })

    ablation, counterfactual = null_and_ablation(pairs, matrix, ann, clusters, [morula_metrics, blast_metrics], ranking)
    svg_scatter(FIGURES / "CSB_TRO_forward_prediction_morula_scatter.svg", obs_morula, pred_morula_mean, "Leave-morula-out DMR prediction")
    svg_scatter(FIGURES / "CSB_TRO_forward_prediction_blastocyst_scatter.svg", obs_blast, pred_blast_mean, "Leave-blastocyst-out DMR prediction")
    svg_bar(
        FIGURES / "CSB_TRO_ablation_rmse.svg",
        list(ablation["experiment"].astype(str)),
        list(ablation["rmse_predicted_vs_observed_stage_mean"].astype(float)),
        "DMR-level prediction and null/ablation RMSE",
        "RMSE",
    )
    svg_bar(
        FIGURES / "CSB_TRO_top_dynamic_DMR_scores.svg",
        list(ranking.head(15)["cluster_name"]),
        list(ranking.head(15)["reset_dynamic_score"].astype(float)),
        "Top dynamic reset-associated DMRs",
        "dynamic score",
    )

    summary = {
        "n_samples": int(matrix.shape[0]),
        "n_dmrs": int(matrix.shape[1]),
        "n_sample_pairs": int(len(pairs)),
        "morula_prediction_rmse": float(morula_metrics["rmse_predicted_vs_observed_stage_mean"]),
        "blastocyst_prediction_rmse": float(blast_metrics["rmse_predicted_vs_observed_stage_mean"]),
        "operator_fit_morula_rmse": float(fit_morula_metrics["rmse_predicted_vs_observed_stage_mean"]),
        "operator_fit_blastocyst_rmse": float(fit_blast_metrics["rmse_predicted_vs_observed_stage_mean"]),
        "outputs": {
            "matrix": str(RESULTS / "CSB_TRO_DMR_state_matrix.tsv"),
            "training_pairs": str(RESULTS / "CSB_TRO_OT_transition_training_pairs.tsv"),
            "model": str(RESULTS / "CSB_TRO_velocity_model_coefficients.tsv"),
            "ranking": str(RESULTS / "CSB_TRO_dynamic_DMR_ranking.tsv"),
        },
    }
    write_json(META / "CSB_TRO_DMR_dynamics_run_summary.json", summary)
    write_docs(summary)
    print("CSB-TRO DMR-level operator-time dynamics complete")
    print(json.dumps(summary, indent=2))


if __name__ == "__main__":
    main()
```
