## Supplemental code for "An entropy-based diagnostic framework for characterizing methylation state dynamics during preimplantation development": run_dual_branch_chromatin_state_control.py

```
from __future__ import annotations

import json
import sys
from pathlib import Path

import numpy as np
import pandas as pd


BASE = Path(r"E:\CSB_TRO_operator_time_DMR_dynamics_2026-05-25")
CODE = BASE / "code"
RESULTS = BASE / "results"
FIGURES = BASE / "figures"
DOCS = BASE / "docs"
sys.path.insert(0, str(CODE))

from run_basin_residual_control_field import PRE_MORULA_STAGES, cosine, simulate_strict_pre_morula  # noqa: E402
from run_morula_basin_sde import (  # noqa: E402
    basin_definition,
    decode_latent,
    distribution_metrics,
    fit_latent_basis,
    fit_operator,
    load_inputs,
    stage_ids,
)


CONTRIB = RESULTS / "CSB_TRO_inverse_ATAC_module_contributions.tsv"
BASIS = RESULTS / "CSB_TRO_missing_control_term_module_basis.tsv"

OUT_BRANCH = RESULTS / "CSB_TRO_dual_branch_chromatin_state_vectors.tsv"
OUT_GRID = RESULTS / "CSB_TRO_dual_branch_chromatin_state_beta_grid.tsv"
OUT_SUMMARY = RESULTS / "CSB_TRO_dual_branch_chromatin_state_summary.tsv"
OUT_MANIFEST = RESULTS / "CSB_TRO_dual_branch_chromatin_state_manifest.json"
OUT_SVG = FIGURES / "CSB_TRO_dual_branch_chromatin_state_control.svg"
OUT_DOC = DOCS / "CSB_TRO_dual_branch_chromatin_state_control_summary.md"


CLOSURE_MODULES = ["M05", "M01", "M12"]
ACCESS_MODULES = ["M02", "M10"]
FULL_CORE = ["M05", "M01", "M12", "M02"]


def latent_context():
    matrix, ann, pairs = load_inputs()
    mu, sd, components, score_df = fit_latent_basis(matrix, 3)
    coef, _, _ = fit_operator(score_df, pairs, train_stages=set(PRE_MORULA_STAGES), lam=1000.0)
    strict_z = simulate_strict_pre_morula(score_df, ann, coef, n_steps=12)
    obs_z = score_df.loc[stage_ids(ann, "morula")].to_numpy(dtype=float)
    obs_dmr = matrix.loc[stage_ids(ann, "morula")].to_numpy(dtype=float)
    basin = basin_definition(obs_z)
    residual_z = obs_z.mean(axis=0) - strict_z.mean(axis=0)
    return mu, sd, components, strict_z, obs_z, obs_dmr, basin, residual_z


def module_vectors(feature_set: str) -> dict[str, np.ndarray]:
    tab = pd.read_csv(CONTRIB, sep="\t")
    sub = tab[tab["feature_set"] == feature_set].copy()
    return {
        str(r.module_id): np.asarray([float(r.module_PC1_contribution), float(r.module_PC2_contribution), float(r.module_PC3_contribution)])
        for r in sub.itertuples()
    }


def branch_table() -> pd.DataFrame:
    rows = []
    basis = pd.read_csv(BASIS, sep="\t")
    residual_modules = {
        "measured_M05_M01_M12": np.sum(
            basis[basis["module_id"].isin(CLOSURE_MODULES)][["latent_control_PC1", "latent_control_PC2", "latent_control_PC3"]].to_numpy(dtype=float),
            axis=0,
        ),
        "measured_M02_M10": np.sum(
            basis[basis["module_id"].isin(ACCESS_MODULES)][["latent_control_PC1", "latent_control_PC2", "latent_control_PC3"]].to_numpy(dtype=float),
            axis=0,
        ),
    }
    for fs in ["ATAC_8cell_2pn_chromatin_only_inverse", "ATAC_8cell_3pn_chromatin_only_inverse"]:
        vecs = module_vectors(fs)
        inv_closure = sum((vecs[m] for m in CLOSURE_MODULES), start=np.zeros(3))
        inv_access = sum((vecs[m] for m in ACCESS_MODULES), start=np.zeros(3))
        raw_access = -inv_access
        raw_core_access = -sum((vecs[m] for m in ["M02"]), start=np.zeros(3))
        branches = {
            "closure_inverse_ATAC_M05_M01_M12": inv_closure,
            "access_inverse_ATAC_M02_M10": inv_access,
            "access_raw_ATAC_M02_M10": raw_access,
            "access_raw_ATAC_M02_only": raw_core_access,
            "naive_inverse_all_priority": inv_closure + inv_access,
            "closure_plus_raw_access": inv_closure + raw_access,
            "closure_plus_raw_M02": inv_closure + raw_core_access,
        }
        for name, vec in branches.items():
            rows.append(
                {
                    "source_feature_set": fs,
                    "branch": name,
                    "PC1": float(vec[0]),
                    "PC2": float(vec[1]),
                    "PC3": float(vec[2]),
                    "norm": float(np.linalg.norm(vec)),
                }
            )
        for name, vec in residual_modules.items():
            rows.append(
                {
                    "source_feature_set": fs,
                    "branch": name,
                    "PC1": float(vec[0]),
                    "PC2": float(vec[1]),
                    "PC3": float(vec[2]),
                    "norm": float(np.linalg.norm(vec)),
                }
            )
    out = pd.DataFrame(rows)
    out.to_csv(OUT_BRANCH, sep="\t", index=False)
    return out


def beta_values(start: float, stop: float, step: float) -> list[float]:
    return [round(float(x), 2) for x in np.arange(start, stop + 1e-9, step)]


def evaluate_grid(branches: pd.DataFrame) -> pd.DataFrame:
    mu, sd, components, strict_z, obs_z, obs_dmr, basin, residual_z = latent_context()
    rng = np.random.default_rng(20260526)
    rows = []
    baseline_dmr = decode_latent(strict_z, mu, sd, components)
    rows.append(
        {
            "source_feature_set": "baseline",
            "model": "methylation_only_baseline",
            "beta_closure": 0.0,
            "beta_access": 0.0,
            "access_orientation": "none",
            "PC1_control": 0.0,
            "PC2_control": 0.0,
            "PC3_control": 0.0,
            "direction_cosine_to_measured_correction": np.nan,
            "PC3_negative_pull_recovered": 0.0,
            **distribution_metrics(strict_z, obs_z, baseline_dmr, obs_dmr, basin, rng),
        }
    )
    for fs, sub in branches.groupby("source_feature_set"):
        if fs == "baseline":
            continue
        def get(branch: str) -> np.ndarray:
            r = sub[sub["branch"] == branch].iloc[0]
            return np.asarray([float(r.PC1), float(r.PC2), float(r.PC3)])

        closure = get("closure_inverse_ATAC_M05_M01_M12")
        inv_access = get("access_inverse_ATAC_M02_M10")
        raw_access = get("access_raw_ATAC_M02_M10")
        raw_m02 = get("access_raw_ATAC_M02_only")
        access_options = {
            "none": np.zeros(3),
            "inverse_M02_M10": inv_access,
            "raw_M02_M10": raw_access,
            "raw_M02_only": raw_m02,
        }
        for access_name, access_vec in access_options.items():
            for beta_c in beta_values(0.0, 2.5, 0.1):
                access_grid = [0.0] if access_name == "none" else beta_values(-1.5, 1.5, 0.1)
                for beta_a in access_grid:
                    vec = beta_c * closure + beta_a * access_vec
                    pred = strict_z + vec[None, :]
                    pred_dmr = decode_latent(pred, mu, sd, components)
                    rows.append(
                        {
                            "source_feature_set": fs,
                            "model": "dual_branch_grid",
                            "beta_closure": beta_c,
                            "beta_access": beta_a,
                            "access_orientation": access_name,
                            "PC1_control": float(vec[0]),
                            "PC2_control": float(vec[1]),
                            "PC3_control": float(vec[2]),
                            "control_norm": float(np.linalg.norm(vec)),
                            "direction_cosine_to_measured_correction": cosine(vec, residual_z) if np.linalg.norm(vec) else np.nan,
                            "PC3_negative_pull_recovered": float(-vec[2] / (-residual_z[2])) if residual_z[2] < 0 else np.nan,
                            **distribution_metrics(pred, obs_z, pred_dmr, obs_dmr, basin, rng),
                        }
                    )
    out = pd.DataFrame(rows)
    out.to_csv(OUT_GRID, sep="\t", index=False)
    return out


def summarize(grid: pd.DataFrame) -> pd.DataFrame:
    rows = []
    obs_occ = 0.875
    for keys, sub in grid[grid["model"] != "methylation_only_baseline"].groupby(["source_feature_set", "access_orientation"]):
        sub = sub.copy()
        best = sub.sort_values(
            ["pred_basin_occupancy_q90", "direction_cosine_to_measured_correction", "PC3_negative_pull_recovered"],
            ascending=False,
        ).iloc[0]
        near_unit = sub[(sub["beta_closure"] - 1.0).abs() < 1e-9]
        near_unit_best = near_unit.sort_values(["pred_basin_occupancy_q90", "direction_cosine_to_measured_correction"], ascending=False).iloc[0] if len(near_unit) else best
        reached = sub[sub["pred_basin_occupancy_q90"] >= obs_occ].sort_values(["beta_closure", "beta_access"])
        rows.append(
            {
                "source_feature_set": keys[0],
                "access_orientation": keys[1],
                "max_occupancy": float(best["pred_basin_occupancy_q90"]),
                "beta_closure_at_max": float(best["beta_closure"]),
                "beta_access_at_max": float(best["beta_access"]),
                "cosine_at_max": float(best["direction_cosine_to_measured_correction"]),
                "PC3_recovery_at_max": float(best["PC3_negative_pull_recovered"]),
                "occupancy_best_at_beta_closure_1": float(near_unit_best["pred_basin_occupancy_q90"]),
                "beta_access_best_at_beta_closure_1": float(near_unit_best["beta_access"]),
                "cosine_best_at_beta_closure_1": float(near_unit_best["direction_cosine_to_measured_correction"]),
                "alpha_to_observed_beta_closure": float(reached.iloc[0]["beta_closure"]) if len(reached) else np.nan,
                "beta_access_at_first_observed": float(reached.iloc[0]["beta_access"]) if len(reached) else np.nan,
            }
        )
    out = pd.DataFrame(rows).sort_values(["max_occupancy", "cosine_at_max"], ascending=False)
    out.to_csv(OUT_SUMMARY, sep="\t", index=False)
    return out


def make_svg(summary: pd.DataFrame) -> None:
    rows = summary.head(10)
    width, height = 940, 430
    left, right, top, bottom = 85, 25, 45, 135
    plot_w = width - left - right
    plot_h = height - top - bottom
    gap = plot_w / max(len(rows), 1)
    bar_w = gap * 0.62
    lines = [
        f'<svg xmlns="http://www.w3.org/2000/svg" width="{width}" height="{height}" viewBox="0 0 {width} {height}">',
        '<rect width="100%" height="100%" fill="white"/>',
        f'<text x="{left}" y="28" font-family="Arial" font-size="18" font-weight="700">Dual-branch chromatin-state control</text>',
        f'<line x1="{left}" y1="{height-bottom}" x2="{width-right}" y2="{height-bottom}" stroke="#222"/>',
        f'<line x1="{left}" y1="{top}" x2="{left}" y2="{height-bottom}" stroke="#222"/>',
    ]
    for tick in [0.044, 0.5, 0.875, 1.0]:
        y = height - bottom - tick * plot_h
        color = "#b23b3b" if tick in [0.044, 0.875] else "#dddddd"
        lines.append(f'<line x1="{left}" y1="{y:.2f}" x2="{width-right}" y2="{y:.2f}" stroke="{color}" stroke-dasharray="4 4"/>')
        lines.append(f'<text x="{left-8}" y="{y+4:.2f}" text-anchor="end" font-family="Arial" font-size="10">{tick:.3g}</text>')
    for i, row in enumerate(rows.itertuples()):
        x = left + i * gap + (gap - bar_w) / 2
        val = float(row.max_occupancy)
        y = height - bottom - val * plot_h
        color = "#2c6f5a" if row.access_orientation == "none" else "#6f4d8b"
        lines.append(f'<rect x="{x:.2f}" y="{y:.2f}" width="{bar_w:.2f}" height="{val * plot_h:.2f}" fill="{color}"/>')
        lines.append(f'<text x="{x+bar_w/2:.2f}" y="{y-5:.2f}" text-anchor="middle" font-family="Arial" font-size="10">{val:.3f}</text>')
        label = f"{row.source_feature_set.replace('_chromatin_only_inverse','').replace('ATAC_','')} {row.access_orientation}".replace("_", " ")
        lines.append(f'<text x="{x+bar_w/2:.2f}" y="{height-bottom+18}" text-anchor="end" transform="rotate(-45 {x+bar_w/2:.2f} {height-bottom+18})" font-family="Arial" font-size="9">{label}</text>')
    lines.append(f'<text x="{left}" y="{height-22}" font-family="Arial" font-size="12">Green: closure branch only. Purple: closure plus accessibility branch grid.</text>')
    lines.append("</svg>")
    OUT_SVG.write_text("\n".join(lines), encoding="utf-8")


def write_doc(branches: pd.DataFrame, summary: pd.DataFrame) -> None:
    lines = [
        "# Dual-Branch Chromatin-State Control",
        "",
        "Status: `completed`",
        "",
        "This formalizes the inverse ATAC result into two candidate control branches:",
        "",
        "- closure branch: inverse-ATAC signal on M05/M01/M12",
        "- accessibility branch: M02/M10 accessibility signal tested in raw and inverse orientations",
        "",
        "## Branch Vectors",
        "",
    ]
    for row in branches[branches["branch"].isin(["closure_inverse_ATAC_M05_M01_M12", "access_inverse_ATAC_M02_M10", "access_raw_ATAC_M02_M10"])].itertuples():
        lines.append(f"- {row.source_feature_set} / {row.branch}: PC1={row.PC1:.3f}, PC2={row.PC2:.3f}, PC3={row.PC3:.3f}, norm={row.norm:.3f}")
    lines += ["", "## Beta Grid Summary", ""]
    for row in summary.head(10).itertuples():
        lines.append(
            f"- {row.source_feature_set} + {row.access_orientation}: max_occ={row.max_occupancy:.3f} "
            f"at beta_closure={row.beta_closure_at_max:.2f}, beta_access={row.beta_access_at_max:.2f}; "
            f"cosine={row.cosine_at_max:.3f}; PC3={row.PC3_recovery_at_max:.3f}; "
            f"first_observed_beta_closure={row.alpha_to_observed_beta_closure:.2f}, beta_access={row.beta_access_at_first_observed:.2f}"
        )
    lines += [
        "",
        "## Mechanistic Interpretation",
        "",
        "The basin-entry component is dominated by a closure-like M05/M01/M12 branch. M02/M10 accessibility is not a required positive branch for morula basin entry in this model; when included naively with the same inverse-ATAC orientation, it partly cancels the PC3-negative pull. This separates the missing correction into a primary closure/histone-state branch and a secondary promoter-accessibility branch that must be signed and weighted separately.",
        "",
        "This result gives a concrete histone target: H3K27ac/H3K4me3/H3K27me3 data should first be asked whether it supports M05/M01/M12 closure or repressive/poised-state gain, not whether it globally matches all residual modules.",
    ]
    OUT_DOC.write_text("\n".join(lines) + "\n", encoding="utf-8")


def main() -> None:
    RESULTS.mkdir(parents=True, exist_ok=True)
    FIGURES.mkdir(parents=True, exist_ok=True)
    DOCS.mkdir(parents=True, exist_ok=True)
    branches = branch_table()
    grid = evaluate_grid(branches)
    summary = summarize(grid)
    make_svg(summary)
    write_doc(branches, summary)
    OUT_MANIFEST.write_text(
        json.dumps(
            {
                "status": "completed",
                "outputs": [str(OUT_BRANCH), str(OUT_GRID), str(OUT_SUMMARY), str(OUT_SVG), str(OUT_DOC)],
                "interpretation": "primary basin-entry branch is M05/M01/M12 closure-like chromatin-state control",
            },
            indent=2,
        )
        + "\n",
        encoding="utf-8",
    )
    print(json.dumps({"status": "completed", "top": summary.head(8).to_dict(orient="records")}, indent=2))


if __name__ == "__main__":
    main()
```
