## Supplemental code for "An entropy-based diagnostic framework for characterizing methylation state dynamics during preimplantation development": run_dual_branch_robustness_controls.py

```
from __future__ import annotations

import json
import sys
from itertools import combinations, product
from pathlib import Path

import numpy as np
import pandas as pd


BASE = Path(r"E:\CSB_TRO_operator_time_DMR_dynamics_2026-05-25")
CODE = BASE / "code"
RESULTS = BASE / "results"
FIGURES = BASE / "figures"
DOCS = BASE / "docs"
sys.path.insert(0, str(CODE))

from run_basin_residual_control_field import PRE_MORULA_STAGES, cosine, simulate_strict_pre_morula  # noqa: E402
from run_morula_basin_sde import (  # noqa: E402
    basin_definition,
    decode_latent,
    distribution_metrics,
    fit_latent_basis,
    fit_operator,
    load_inputs,
    stage_ids,
)


CONTRIB = RESULTS / "CSB_TRO_inverse_ATAC_module_contributions.tsv"

OUT_ABLATION = RESULTS / "CSB_TRO_dual_branch_ablation.tsv"
OUT_SIGN = RESULTS / "CSB_TRO_dual_branch_sign_control.tsv"
OUT_RANDOM = RESULTS / "CSB_TRO_dual_branch_random_partition_control.tsv"
OUT_EXACT = RESULTS / "CSB_TRO_dual_branch_exact_partition_sign_control.tsv"
OUT_SUMMARY = RESULTS / "CSB_TRO_dual_branch_robustness_summary.tsv"
OUT_MANIFEST = RESULTS / "CSB_TRO_dual_branch_robustness_manifest.json"
OUT_SVG = FIGURES / "CSB_TRO_dual_branch_robustness_controls.svg"
OUT_DOC = DOCS / "CSB_TRO_dual_branch_ablation_summary.md"


PRIORITY_MODULES = ["M05", "M01", "M12", "M02", "M10"]
CLOSURE_MODULES = ["M05", "M01", "M12"]
ACCESS_MODULES = ["M02", "M10"]
SOURCE_FEATURE_SET = "ATAC_8cell_3pn_chromatin_only_inverse"
DEFAULT_BETA_CLOSURE = 1.0
DEFAULT_BETA_ACCESS = 1.5


def latent_context():
    matrix, ann, pairs = load_inputs()
    mu, sd, components, score_df = fit_latent_basis(matrix, 3)
    coef, _, _ = fit_operator(score_df, pairs, train_stages=set(PRE_MORULA_STAGES), lam=1000.0)
    strict_z = simulate_strict_pre_morula(score_df, ann, coef, n_steps=12)
    obs_z = score_df.loc[stage_ids(ann, "morula")].to_numpy(dtype=float)
    obs_dmr = matrix.loc[stage_ids(ann, "morula")].to_numpy(dtype=float)
    basin = basin_definition(obs_z)
    residual_z = obs_z.mean(axis=0) - strict_z.mean(axis=0)
    return mu, sd, components, strict_z, obs_z, obs_dmr, basin, residual_z


def module_vectors(source_feature_set: str = SOURCE_FEATURE_SET) -> dict[str, np.ndarray]:
    tab = pd.read_csv(CONTRIB, sep="\t")
    sub = tab[tab["feature_set"] == source_feature_set].copy()
    if sub.empty:
        raise FileNotFoundError(f"No inverse ATAC module contributions found for {source_feature_set}")
    return {
        str(r.module_id): np.asarray([float(r.module_PC1_contribution), float(r.module_PC2_contribution), float(r.module_PC3_contribution)])
        for r in sub.itertuples()
    }


def raw_access(vecs: dict[str, np.ndarray], modules: list[str] = ACCESS_MODULES) -> np.ndarray:
    # Access branch uses the opposite sign of inverse ATAC for M02/M10.
    return -sum((vecs[m] for m in modules), start=np.zeros(3))


def closure(vecs: dict[str, np.ndarray], modules: list[str] = CLOSURE_MODULES) -> np.ndarray:
    return sum((vecs[m] for m in modules), start=np.zeros(3))


def pc_recoveries(vec: np.ndarray, residual_z: np.ndarray) -> dict[str, float]:
    out = {}
    for i, pc in enumerate(["PC1", "PC2", "PC3"]):
        denom = residual_z[i]
        out[f"{pc}_recovery"] = float(vec[i] / denom) if np.isfinite(denom) and abs(denom) > 1e-12 else np.nan
    out["PC3_negative_pull_recovered"] = float(-vec[2] / (-residual_z[2])) if residual_z[2] < 0 else np.nan
    out["PC1_negative_pull_recovered"] = float(-vec[0] / (-residual_z[0])) if residual_z[0] < 0 else np.nan
    return out


def evaluate_vec(label: str, vec: np.ndarray, status: str, context, rng: np.random.Generator, extra: dict | None = None) -> dict:
    mu, sd, components, strict_z, obs_z, obs_dmr, basin, residual_z = context
    pred = strict_z + vec[None, :]
    pred_dmr = decode_latent(pred, mu, sd, components)
    metrics = distribution_metrics(pred, obs_z, pred_dmr, obs_dmr, basin, rng)
    center_error = float(metrics["pred_mean_distance_to_morula_centroid"] - metrics["observed_mean_distance_to_morula_centroid"])
    row = {
        "model": label,
        "validation_status": status,
        "PC1_control": float(vec[0]),
        "PC2_control": float(vec[1]),
        "PC3_control": float(vec[2]),
        "control_norm": float(np.linalg.norm(vec)),
        "direction_cosine_to_measured_correction": cosine(vec, residual_z) if np.linalg.norm(vec) else np.nan,
        **pc_recoveries(vec, residual_z),
        "center_error_vs_observed": center_error,
        **metrics,
    }
    if extra:
        row.update(extra)
    return row


def run_ablation(context, vecs: dict[str, np.ndarray]) -> pd.DataFrame:
    rng = np.random.default_rng(20260526)
    rows = []
    base_c = closure(vecs)
    base_a = raw_access(vecs)
    full = DEFAULT_BETA_CLOSURE * base_c + DEFAULT_BETA_ACCESS * base_a
    rows.append(evaluate_vec("full_dual_branch", full, "full_model", context, rng, {"included_modules": ",".join(PRIORITY_MODULES), "removed_module": ""}))
    rows.append(evaluate_vec("closure_branch_only", DEFAULT_BETA_CLOSURE * base_c, "branch_only", context, rng, {"included_modules": ",".join(CLOSURE_MODULES), "removed_module": "M02,M10"}))
    rows.append(evaluate_vec("access_branch_only", DEFAULT_BETA_ACCESS * base_a, "branch_only", context, rng, {"included_modules": ",".join(ACCESS_MODULES), "removed_module": "M05,M01,M12"}))
    for module_id in PRIORITY_MODULES:
        c_modules = [m for m in CLOSURE_MODULES if m != module_id]
        a_modules = [m for m in ACCESS_MODULES if m != module_id]
        vec = DEFAULT_BETA_CLOSURE * closure(vecs, c_modules) + DEFAULT_BETA_ACCESS * raw_access(vecs, a_modules)
        rows.append(
            evaluate_vec(
                f"remove_{module_id}",
                vec,
                "leave_one_module_out",
                context,
                rng,
                {"included_modules": ",".join(c_modules + a_modules), "removed_module": module_id},
            )
        )
        rows.append(
            evaluate_vec(
                f"remove_{module_id}_sign_flip",
                -vec,
                "leave_one_module_out_sign_flip",
                context,
                rng,
                {"included_modules": ",".join(c_modules + a_modules), "removed_module": module_id},
            )
        )
    rows.append(evaluate_vec("full_dual_branch_sign_flip", -full, "sign_flip_control", context, rng, {"included_modules": ",".join(PRIORITY_MODULES), "removed_module": ""}))
    out = pd.DataFrame(rows)
    out.to_csv(OUT_ABLATION, sep="\t", index=False)
    return out


def run_sign_controls(context, vecs: dict[str, np.ndarray]) -> pd.DataFrame:
    rng = np.random.default_rng(20260526)
    c = closure(vecs)
    a_raw = raw_access(vecs)
    a_inverse = -a_raw
    naive_inverse = c + a_inverse
    naive_raw = -c + a_raw
    rows = []
    tests = [
        ("correct_closure_correct_access", DEFAULT_BETA_CLOSURE * c + DEFAULT_BETA_ACCESS * a_raw, "correct_dual_branch"),
        ("wrong_closure_correct_access", -DEFAULT_BETA_CLOSURE * c + DEFAULT_BETA_ACCESS * a_raw, "wrong_closure_sign"),
        ("correct_closure_wrong_access", DEFAULT_BETA_CLOSURE * c - DEFAULT_BETA_ACCESS * a_raw, "wrong_access_sign"),
        ("wrong_closure_wrong_access", -DEFAULT_BETA_CLOSURE * c - DEFAULT_BETA_ACCESS * a_raw, "both_signs_wrong"),
        ("naive_inverse_all_modules", naive_inverse, "naive_same_inverse_sign"),
        ("naive_raw_all_modules", naive_raw, "naive_same_raw_sign"),
        ("closure_only_correct_sign", DEFAULT_BETA_CLOSURE * c, "closure_only"),
        ("access_only_correct_sign", DEFAULT_BETA_ACCESS * a_raw, "access_only"),
    ]
    for label, vec, status in tests:
        rows.append(evaluate_vec(label, vec, status, context, rng))
    out = pd.DataFrame(rows)
    out.to_csv(OUT_SIGN, sep="\t", index=False)
    return out


def run_random_partition(context, vecs: dict[str, np.ndarray], n_random: int = 1000) -> pd.DataFrame:
    rng = np.random.default_rng(20260526)
    eval_rng = np.random.default_rng(20260527)
    rows = []
    true_vec = DEFAULT_BETA_CLOSURE * closure(vecs) + DEFAULT_BETA_ACCESS * raw_access(vecs)
    rows.append(
        evaluate_vec(
            "true_dual_branch_partition",
            true_vec,
            "true_partition",
            context,
            eval_rng,
            {"closure_modules": ",".join(CLOSURE_MODULES), "access_modules": ",".join(ACCESS_MODULES), "random_id": -1},
        )
    )
    modules = np.asarray(PRIORITY_MODULES)
    for i in range(n_random):
        closure_modules = list(rng.choice(modules, size=3, replace=False))
        access_modules = [m for m in PRIORITY_MODULES if m not in closure_modules]
        vec = DEFAULT_BETA_CLOSURE * closure(vecs, closure_modules) + DEFAULT_BETA_ACCESS * raw_access(vecs, access_modules)
        rows.append(
            evaluate_vec(
                f"random_partition_{i:04d}",
                vec,
                "random_3closure_2access_partition",
                context,
                eval_rng,
                {"closure_modules": ",".join(closure_modules), "access_modules": ",".join(access_modules), "random_id": i},
            )
        )
    out = pd.DataFrame(rows)
    out.to_csv(OUT_RANDOM, sep="\t", index=False)
    return out


def run_exact_partition_sign_control(context, vecs: dict[str, np.ndarray]) -> pd.DataFrame:
    eval_rng = np.random.default_rng(20260528)
    rows = []
    true_closure = tuple(CLOSURE_MODULES)
    true_access = tuple(ACCESS_MODULES)
    true_vec = DEFAULT_BETA_CLOSURE * closure(vecs, list(true_closure)) + DEFAULT_BETA_ACCESS * raw_access(vecs, list(true_access))
    rows.append(
        evaluate_vec(
            "true_dual_branch_partition",
            true_vec,
            "true_partition_correct_sign",
            context,
            eval_rng,
            {
                "closure_modules": ",".join(true_closure),
                "access_modules": ",".join(true_access),
                "closure_sign": 1,
                "access_sign": 1,
            },
        )
    )
    seen = set()
    for closure_modules in combinations(PRIORITY_MODULES, 3):
        access_modules = tuple(m for m in PRIORITY_MODULES if m not in closure_modules)
        for closure_sign, access_sign in product([-1, 1], repeat=2):
            key = (closure_modules, access_modules, closure_sign, access_sign)
            if key in seen:
                continue
            seen.add(key)
            vec = (
                closure_sign * DEFAULT_BETA_CLOSURE * closure(vecs, list(closure_modules))
                + access_sign * DEFAULT_BETA_ACCESS * raw_access(vecs, list(access_modules))
            )
            rows.append(
                evaluate_vec(
                    f"exact_partition_{len(rows):03d}",
                    vec,
                    "exact_partition_with_branch_signs",
                    context,
                    eval_rng,
                    {
                        "closure_modules": ",".join(closure_modules),
                        "access_modules": ",".join(access_modules),
                        "closure_sign": closure_sign,
                        "access_sign": access_sign,
                    },
                )
            )
    out = pd.DataFrame(rows)
    out.to_csv(OUT_EXACT, sep="\t", index=False)
    return out


def summarize(ablation: pd.DataFrame, sign: pd.DataFrame, random: pd.DataFrame, exact: pd.DataFrame) -> pd.DataFrame:
    rows = []
    full = ablation[ablation["model"] == "full_dual_branch"].iloc[0]
    closure_only = ablation[ablation["model"] == "closure_branch_only"].iloc[0]
    access_only = ablation[ablation["model"] == "access_branch_only"].iloc[0]
    rows.extend(
        [
            {"summary_item": "full_dual_branch_occupancy", "value": float(full["pred_basin_occupancy_q90"]), "detail": "beta_closure=1.0,beta_access=1.5"},
            {"summary_item": "full_dual_branch_cosine", "value": float(full["direction_cosine_to_measured_correction"]), "detail": "direction alignment"},
            {"summary_item": "closure_only_occupancy", "value": float(closure_only["pred_basin_occupancy_q90"]), "detail": "M05/M01/M12 only"},
            {"summary_item": "access_only_occupancy", "value": float(access_only["pred_basin_occupancy_q90"]), "detail": "M02/M10 only"},
        ]
    )
    for module_id in PRIORITY_MODULES:
        rec = ablation[ablation["model"] == f"remove_{module_id}"].iloc[0]
        rows.append(
            {
                "summary_item": f"delta_occupancy_remove_{module_id}",
                "value": float(full["pred_basin_occupancy_q90"] - rec["pred_basin_occupancy_q90"]),
                "detail": f"full minus remove_{module_id}",
            }
        )
    correct = sign[sign["model"] == "correct_closure_correct_access"].iloc[0]
    for model in ["wrong_closure_correct_access", "correct_closure_wrong_access", "wrong_closure_wrong_access", "naive_inverse_all_modules", "naive_raw_all_modules"]:
        rec = sign[sign["model"] == model].iloc[0]
        rows.append(
            {
                "summary_item": f"sign_control_delta_vs_correct_{model}",
                "value": float(correct["pred_basin_occupancy_q90"] - rec["pred_basin_occupancy_q90"]),
                "detail": "positive means correct dual-branch sign is better",
            }
        )
    true = random[random["validation_status"] == "true_partition"].iloc[0]
    rnd = random[random["validation_status"] == "random_3closure_2access_partition"].copy()
    for metric in ["pred_basin_occupancy_q90", "direction_cosine_to_measured_correction", "PC3_negative_pull_recovered"]:
        val = float(true[metric])
        percentile = float((rnd[metric] <= val).mean())
        p_ge = float((rnd[metric] >= val).mean())
        rows.append({"summary_item": f"true_partition_percentile_{metric}", "value": percentile, "detail": f"empirical p_ge={p_ge:.4f}; true={val:.4f}"})
    exact_true = exact[exact["validation_status"] == "true_partition_correct_sign"].iloc[0]
    exact_null = exact[exact["validation_status"] == "exact_partition_with_branch_signs"].copy()
    for metric in ["pred_basin_occupancy_q90", "direction_cosine_to_measured_correction", "PC3_negative_pull_recovered"]:
        val = float(exact_true[metric])
        percentile = float((exact_null[metric] <= val).mean())
        p_ge = float((exact_null[metric] >= val).mean())
        rows.append({"summary_item": f"true_exact_partition_sign_percentile_{metric}", "value": percentile, "detail": f"exact branch-sign null p_ge={p_ge:.4f}; true={val:.4f}; n={len(exact_null)}"})
    out = pd.DataFrame(rows)
    out.to_csv(OUT_SUMMARY, sep="\t", index=False)
    return out


def make_svg(ablation: pd.DataFrame, sign: pd.DataFrame, random: pd.DataFrame) -> None:
    rows = pd.concat(
        [
            ablation[ablation["model"].isin(["full_dual_branch", "closure_branch_only", "access_branch_only", "remove_M05", "remove_M01", "remove_M12", "remove_M02", "remove_M10"])],
            sign[sign["model"].isin(["wrong_closure_correct_access", "correct_closure_wrong_access", "naive_inverse_all_modules", "naive_raw_all_modules"])],
        ],
        ignore_index=True,
    )
    width, height = 1050, 460
    left, right, top, bottom = 85, 30, 45, 150
    plot_w = width - left - right
    plot_h = height - top - bottom
    gap = plot_w / max(len(rows), 1)
    bar_w = gap * 0.62
    lines = [
        f'<svg xmlns="http://www.w3.org/2000/svg" width="{width}" height="{height}" viewBox="0 0 {width} {height}">',
        '<rect width="100%" height="100%" fill="white"/>',
        f'<text x="{left}" y="28" font-family="Arial" font-size="18" font-weight="700">Dual-branch robustness controls</text>',
        f'<line x1="{left}" y1="{height-bottom}" x2="{width-right}" y2="{height-bottom}" stroke="#222"/>',
        f'<line x1="{left}" y1="{top}" x2="{left}" y2="{height-bottom}" stroke="#222"/>',
    ]
    rnd = random[random["validation_status"] == "random_3closure_2access_partition"]
    q95 = float(rnd["pred_basin_occupancy_q90"].quantile(0.95))
    for tick in [0.044, q95, 0.875, 1.0]:
        y = height - bottom - tick * plot_h
        color = "#b23b3b" if tick in [0.044, 0.875] else "#888888" if abs(tick - q95) < 1e-9 else "#dddddd"
        lines.append(f'<line x1="{left}" y1="{y:.2f}" x2="{width-right}" y2="{y:.2f}" stroke="{color}" stroke-dasharray="4 4"/>')
        lines.append(f'<text x="{left-8}" y="{y+4:.2f}" text-anchor="end" font-family="Arial" font-size="10">{tick:.3g}</text>')
    for i, row in enumerate(rows.itertuples()):
        x = left + i * gap + (gap - bar_w) / 2
        val = float(row.pred_basin_occupancy_q90)
        y = height - bottom - val * plot_h
        color = "#2c6f5a" if "full" in row.model or "closure" in row.model else "#6f4d8b" if "remove" in row.model else "#8a8a8a"
        lines.append(f'<rect x="{x:.2f}" y="{y:.2f}" width="{bar_w:.2f}" height="{val * plot_h:.2f}" fill="{color}"/>')
        lines.append(f'<text x="{x+bar_w/2:.2f}" y="{y-5:.2f}" text-anchor="middle" font-family="Arial" font-size="10">{val:.3f}</text>')
        label = row.model.replace("_", " ")
        lines.append(f'<text x="{x+bar_w/2:.2f}" y="{height-bottom+18}" text-anchor="end" transform="rotate(-45 {x+bar_w/2:.2f} {height-bottom+18})" font-family="Arial" font-size="9">{label}</text>')
    lines.append(f'<text x="{left}" y="{height-22}" font-family="Arial" font-size="12">Gray guide: random partition q95. Red guides: methylation baseline and observed morula occupancy.</text>')
    lines.append("</svg>")
    OUT_SVG.write_text("\n".join(lines), encoding="utf-8")


def write_doc(summary: pd.DataFrame, ablation: pd.DataFrame, sign: pd.DataFrame) -> None:
    full = ablation[ablation["model"] == "full_dual_branch"].iloc[0]
    lines = [
        "# Dual-Branch Ablation, Sign, and Random Controls",
        "",
        "Status: `completed`",
        "",
        "This tests whether the M05/M01/M12 closure branch plus the signed M02/M10 access branch is a stable structure rather than a parameter accident.",
        "",
        "## Main Model",
        "",
        f"- full dual-branch occupancy={full.pred_basin_occupancy_q90:.3f}, cosine={full.direction_cosine_to_measured_correction:.3f}, PC3_recovery={full.PC3_negative_pull_recovered:.3f}, center_error={full.center_error_vs_observed:.3f}",
        "",
        "## Ablation",
        "",
    ]
    for model in ["closure_branch_only", "access_branch_only", "remove_M05", "remove_M01", "remove_M12", "remove_M02", "remove_M10"]:
        row = ablation[ablation["model"] == model].iloc[0]
        lines.append(
            f"- {model}: occupancy={row.pred_basin_occupancy_q90:.3f}, cosine={row.direction_cosine_to_measured_correction:.3f}, "
            f"PC1={row.PC1_negative_pull_recovered:.3f}, PC3={row.PC3_negative_pull_recovered:.3f}, center_error={row.center_error_vs_observed:.3f}"
        )
    lines += ["", "## Sign Controls", ""]
    for model in ["correct_closure_correct_access", "wrong_closure_correct_access", "correct_closure_wrong_access", "wrong_closure_wrong_access", "naive_inverse_all_modules", "naive_raw_all_modules"]:
        row = sign[sign["model"] == model].iloc[0]
        lines.append(
            f"- {model}: occupancy={row.pred_basin_occupancy_q90:.3f}, cosine={row.direction_cosine_to_measured_correction:.3f}, PC3={row.PC3_negative_pull_recovered:.3f}"
        )
    lines += ["", "## Summary Stats", ""]
    for row in summary.itertuples():
        lines.append(f"- {row.summary_item}: {row.value:.4f} ({row.detail})")
    lines += [
        "",
        "## Interpretation",
        "",
        "The critical test is whether correct branch sign beats wrong-sign and random partition controls. If it does, the dual-branch model is not merely an ATAC proxy with tuned amplitude; it is a structured chromatin-state control hypothesis that can be directly replaced by histone tracks when available.",
    ]
    OUT_DOC.write_text("\n".join(lines) + "\n", encoding="utf-8")


def main() -> None:
    RESULTS.mkdir(parents=True, exist_ok=True)
    FIGURES.mkdir(parents=True, exist_ok=True)
    DOCS.mkdir(parents=True, exist_ok=True)
    context = latent_context()
    vecs = module_vectors()
    ablation = run_ablation(context, vecs)
    sign = run_sign_controls(context, vecs)
    random = run_random_partition(context, vecs, n_random=1000)
    exact = run_exact_partition_sign_control(context, vecs)
    summary = summarize(ablation, sign, random, exact)
    make_svg(ablation, sign, random)
    write_doc(summary, ablation, sign)
    OUT_MANIFEST.write_text(
        json.dumps(
            {
                "status": "completed",
                "source_feature_set": SOURCE_FEATURE_SET,
                "beta_closure": DEFAULT_BETA_CLOSURE,
                "beta_access": DEFAULT_BETA_ACCESS,
                "outputs": [str(OUT_ABLATION), str(OUT_SIGN), str(OUT_RANDOM), str(OUT_EXACT), str(OUT_SUMMARY), str(OUT_SVG), str(OUT_DOC)],
            },
            indent=2,
        )
        + "\n",
        encoding="utf-8",
    )
    print(json.dumps({"status": "completed", "summary": summary.to_dict(orient="records")}, indent=2))


if __name__ == "__main__":
    main()
```
