## Supplemental code for "An entropy-based diagnostic framework for characterizing methylation state dynamics during preimplantation development": run_dual_branch_structure_validation.py

```
from __future__ import annotations

import json
import sys
from itertools import combinations, product
from pathlib import Path

import numpy as np
import pandas as pd


BASE = Path(r"E:\CSB_TRO_operator_time_DMR_dynamics_2026-05-25")
CODE = BASE / "code"
RESULTS = BASE / "results"
FIGURES = BASE / "figures"
DOCS = BASE / "docs"
sys.path.insert(0, str(CODE))

from run_basin_residual_control_field import PRE_MORULA_STAGES, cosine, simulate_strict_pre_morula  # noqa: E402
from run_morula_basin_sde import (  # noqa: E402
    basin_definition,
    decode_latent,
    distribution_metrics,
    fit_latent_basis,
    fit_operator,
    load_inputs,
    stage_ids,
)


CONTRIB = RESULTS / "CSB_TRO_inverse_ATAC_module_contributions.tsv"

OUT_GRID = RESULTS / "CSB_TRO_dual_branch_beta_grid.tsv"
OUT_ORDER = RESULTS / "CSB_TRO_dual_branch_module_order.tsv"
OUT_SIGNS = RESULTS / "CSB_TRO_all_module_sign_patterns.tsv"
OUT_SUMMARY = RESULTS / "CSB_TRO_dual_branch_structure_validation_summary.tsv"
OUT_MANIFEST = RESULTS / "CSB_TRO_dual_branch_structure_validation_manifest.json"
OUT_GRID_OCC = FIGURES / "CSB_TRO_dual_branch_beta_grid_occupancy.svg"
OUT_GRID_COS = FIGURES / "CSB_TRO_dual_branch_beta_grid_cosine.svg"
OUT_DOC = DOCS / "CSB_TRO_dual_branch_structure_validation_summary.md"


MODULES = ["M05", "M01", "M12", "M02", "M10"]
CLOSURE_MODULES = ["M05", "M01", "M12"]
ACCESS_MODULES = ["M02", "M10"]
SOURCE_FEATURE_SET = "ATAC_8cell_3pn_chromatin_only_inverse"
DEFAULT_BETA_ACCESS = 1.5


def latent_context():
    matrix, ann, pairs = load_inputs()
    mu, sd, components, score_df = fit_latent_basis(matrix, 3)
    coef, _, _ = fit_operator(score_df, pairs, train_stages=set(PRE_MORULA_STAGES), lam=1000.0)
    strict_z = simulate_strict_pre_morula(score_df, ann, coef, n_steps=12)
    obs_z = score_df.loc[stage_ids(ann, "morula")].to_numpy(dtype=float)
    obs_dmr = matrix.loc[stage_ids(ann, "morula")].to_numpy(dtype=float)
    basin = basin_definition(obs_z)
    residual_z = obs_z.mean(axis=0) - strict_z.mean(axis=0)
    return mu, sd, components, strict_z, obs_z, obs_dmr, basin, residual_z


def module_vectors() -> dict[str, np.ndarray]:
    tab = pd.read_csv(CONTRIB, sep="\t")
    sub = tab[tab["feature_set"] == SOURCE_FEATURE_SET].copy()
    return {
        str(r.module_id): np.asarray([float(r.module_PC1_contribution), float(r.module_PC2_contribution), float(r.module_PC3_contribution)])
        for r in sub.itertuples()
    }


def sum_vec(vecs: dict[str, np.ndarray], modules: list[str]) -> np.ndarray:
    return sum((vecs[m] for m in modules), start=np.zeros(3))


def closure_vec(vecs: dict[str, np.ndarray], modules: list[str] = CLOSURE_MODULES) -> np.ndarray:
    return sum_vec(vecs, modules)


def access_vec(vecs: dict[str, np.ndarray], modules: list[str] = ACCESS_MODULES) -> np.ndarray:
    return -sum_vec(vecs, modules)


def recoveries(vec: np.ndarray, residual_z: np.ndarray) -> dict[str, float]:
    return {
        "PC1_recovery": float(vec[0] / residual_z[0]) if abs(residual_z[0]) > 1e-12 else np.nan,
        "PC3_recovery": float(vec[2] / residual_z[2]) if abs(residual_z[2]) > 1e-12 else np.nan,
        "PC1_negative_pull_recovered": float(-vec[0] / (-residual_z[0])) if residual_z[0] < 0 else np.nan,
        "PC3_negative_pull_recovered": float(-vec[2] / (-residual_z[2])) if residual_z[2] < 0 else np.nan,
    }


def evaluate(label: str, vec: np.ndarray, context, rng: np.random.Generator, extra: dict | None = None) -> dict:
    mu, sd, components, strict_z, obs_z, obs_dmr, basin, residual_z = context
    pred = strict_z + vec[None, :]
    pred_dmr = decode_latent(pred, mu, sd, components)
    metrics = distribution_metrics(pred, obs_z, pred_dmr, obs_dmr, basin, rng)
    row = {
        "model": label,
        "PC1_control": float(vec[0]),
        "PC2_control": float(vec[1]),
        "PC3_control": float(vec[2]),
        "control_norm": float(np.linalg.norm(vec)),
        "direction_cosine_to_measured_correction": cosine(vec, residual_z) if np.linalg.norm(vec) else np.nan,
        **recoveries(vec, residual_z),
        **metrics,
    }
    if extra:
        row.update(extra)
    return row


def frange(start: float, stop: float, step: float) -> list[float]:
    return [round(float(x), 2) for x in np.arange(start, stop + 1e-9, step)]


def run_beta_grid(context, vecs: dict[str, np.ndarray]) -> pd.DataFrame:
    rng = np.random.default_rng(20260527)
    c = closure_vec(vecs)
    a = access_vec(vecs)
    rows = []
    for beta_c in frange(0, 2, 0.05):
        for beta_a in frange(0, 2, 0.05):
            vec = beta_c * c + beta_a * a
            rows.append(
                evaluate(
                    "dual_branch_beta_grid",
                    vec,
                    context,
                    rng,
                    {"beta_closure": beta_c, "beta_access": beta_a, "source_feature_set": SOURCE_FEATURE_SET},
                )
            )
    out = pd.DataFrame(rows)
    out.to_csv(OUT_GRID, sep="\t", index=False)
    return out


def run_module_order(context, vecs: dict[str, np.ndarray]) -> pd.DataFrame:
    rng = np.random.default_rng(20260527)
    rows = []
    closure_sets = []
    for k in [1, 2, 3]:
        closure_sets.extend(list(combinations(CLOSURE_MODULES, k)))
    access_sets = []
    for k in [1, 2]:
        access_sets.extend(list(combinations(ACCESS_MODULES, k)))

    for mods in closure_sets:
        vec = closure_vec(vecs, list(mods))
        rows.append(evaluate("closure_subset", vec, context, rng, {"branch": "closure", "modules": ",".join(mods), "n_modules": len(mods), "beta": 1.0}))
        vec_full_access = vec + DEFAULT_BETA_ACCESS * access_vec(vecs)
        rows.append(evaluate("closure_subset_plus_full_access", vec_full_access, context, rng, {"branch": "closure_plus_access", "modules": ",".join(mods) + "|M02,M10", "n_modules": len(mods) + 2, "beta": 1.0}))

    for mods in access_sets:
        vec = DEFAULT_BETA_ACCESS * access_vec(vecs, list(mods))
        rows.append(evaluate("access_subset", vec, context, rng, {"branch": "access", "modules": ",".join(mods), "n_modules": len(mods), "beta": DEFAULT_BETA_ACCESS}))
        vec_full_closure = closure_vec(vecs) + vec
        rows.append(evaluate("full_closure_plus_access_subset", vec_full_closure, context, rng, {"branch": "closure_plus_access", "modules": "M05,M01,M12|" + ",".join(mods), "n_modules": 3 + len(mods), "beta": DEFAULT_BETA_ACCESS}))

    out = pd.DataFrame(rows).sort_values(["pred_basin_occupancy_q90", "direction_cosine_to_measured_correction"], ascending=False)
    out.to_csv(OUT_ORDER, sep="\t", index=False)
    return out


def run_all_sign_patterns(context, vecs: dict[str, np.ndarray]) -> pd.DataFrame:
    rng = np.random.default_rng(20260527)
    rows = []
    true_sign = {"M05": 1, "M01": 1, "M12": 1, "M02": -1, "M10": -1}
    for bits in product([-1, 1], repeat=len(MODULES)):
        signs = dict(zip(MODULES, bits))
        vec = sum((signs[m] * vecs[m] for m in MODULES), start=np.zeros(3))
        is_true = all(signs[m] == true_sign[m] for m in MODULES)
        rows.append(
            evaluate(
                "all_module_sign_pattern",
                vec,
                context,
                rng,
                {
                    "sign_pattern": ",".join(f"{m}:{signs[m]:+d}" for m in MODULES),
                    "true_dual_branch_sign": bool(is_true),
                },
            )
        )
    out = pd.DataFrame(rows).sort_values(["pred_basin_occupancy_q90", "direction_cosine_to_measured_correction"], ascending=False)
    out["rank_by_occupancy_cosine"] = np.arange(1, len(out) + 1)
    out.to_csv(OUT_SIGNS, sep="\t", index=False)
    return out


def summarize(grid: pd.DataFrame, order: pd.DataFrame, signs: pd.DataFrame) -> pd.DataFrame:
    rows = []
    strong = grid[
        (grid["pred_basin_occupancy_q90"] >= 0.875)
        & (grid["direction_cosine_to_measured_correction"] >= 0.9)
        & (grid["PC3_negative_pull_recovered"] >= 0.4)
    ]
    rows.append({"summary_item": "beta_grid_points_total", "value": float(len(grid)), "detail": "beta_c,beta_a in [0,2] step 0.05"})
    rows.append({"summary_item": "beta_grid_strong_region_count", "value": float(len(strong)), "detail": "occupancy>=0.875, cosine>=0.9, PC3>=0.4"})
    rows.append({"summary_item": "beta_grid_strong_region_fraction", "value": float(len(strong) / len(grid)), "detail": "continuous parameter robustness proxy"})
    if len(strong):
        rows.append({"summary_item": "beta_grid_strong_beta_closure_min", "value": float(strong["beta_closure"].min()), "detail": "lower edge of robust region"})
        rows.append({"summary_item": "beta_grid_strong_beta_closure_max", "value": float(strong["beta_closure"].max()), "detail": "upper edge of robust region"})
        rows.append({"summary_item": "beta_grid_strong_beta_access_min", "value": float(strong["beta_access"].min()), "detail": "lower edge of robust region"})
        rows.append({"summary_item": "beta_grid_strong_beta_access_max", "value": float(strong["beta_access"].max()), "detail": "upper edge of robust region"})
    best_order = order.head(10)
    for i, row in enumerate(best_order.itertuples(), start=1):
        rows.append({"summary_item": f"module_order_top_{i}", "value": float(row.pred_basin_occupancy_q90), "detail": f"{row.model}; {row.modules}; cosine={row.direction_cosine_to_measured_correction:.3f}; PC3={row.PC3_negative_pull_recovered:.3f}"})
    true = signs[signs["true_dual_branch_sign"] == True].iloc[0]
    rows.append({"summary_item": "true_sign_pattern_rank", "value": float(true["rank_by_occupancy_cosine"]), "detail": str(true["sign_pattern"])})
    rows.append({"summary_item": "true_sign_pattern_occupancy", "value": float(true["pred_basin_occupancy_q90"]), "detail": f"cosine={true['direction_cosine_to_measured_correction']:.3f}; PC3={true['PC3_negative_pull_recovered']:.3f}"})
    rows.append({"summary_item": "true_sign_pattern_percentile_occupancy", "value": float((signs["pred_basin_occupancy_q90"] <= float(true["pred_basin_occupancy_q90"])).mean()), "detail": "32 sign patterns"})
    out = pd.DataFrame(rows)
    out.to_csv(OUT_SUMMARY, sep="\t", index=False)
    return out


def heat_color(val: float, vmin: float, vmax: float) -> str:
    if not np.isfinite(val):
        return "#eeeeee"
    t = 0.0 if vmax <= vmin else max(0.0, min(1.0, (val - vmin) / (vmax - vmin)))
    r = int(245 - 185 * t)
    g = int(245 - 70 * t)
    b = int(245 - 155 * t)
    return f"#{r:02x}{g:02x}{b:02x}"


def make_heatmap(grid: pd.DataFrame, metric: str, out_path: Path, title: str) -> None:
    xs = sorted(grid["beta_closure"].unique())
    ys = sorted(grid["beta_access"].unique())
    width, height = 760, 720
    left, right, top, bottom = 70, 30, 45, 60
    plot_w = width - left - right
    plot_h = height - top - bottom
    cell_w = plot_w / len(xs)
    cell_h = plot_h / len(ys)
    vals = grid[metric].to_numpy(dtype=float)
    vmin, vmax = float(np.nanmin(vals)), float(np.nanmax(vals))
    lookup = {(float(r.beta_closure), float(r.beta_access)): float(getattr(r, metric)) for r in grid.itertuples()}
    lines = [
        f'<svg xmlns="http://www.w3.org/2000/svg" width="{width}" height="{height}" viewBox="0 0 {width} {height}">',
        '<rect width="100%" height="100%" fill="white"/>',
        f'<text x="{left}" y="28" font-family="Arial" font-size="18" font-weight="700">{title}</text>',
    ]
    for ix, xval in enumerate(xs):
        for iy, yval in enumerate(ys):
            val = lookup[(float(xval), float(yval))]
            x = left + ix * cell_w
            y = top + (len(ys) - iy - 1) * cell_h
            lines.append(f'<rect x="{x:.2f}" y="{y:.2f}" width="{cell_w+0.2:.2f}" height="{cell_h+0.2:.2f}" fill="{heat_color(val, vmin, vmax)}"/>')
    lines += [
        f'<line x1="{left}" y1="{height-bottom}" x2="{width-right}" y2="{height-bottom}" stroke="#222"/>',
        f'<line x1="{left}" y1="{top}" x2="{left}" y2="{height-bottom}" stroke="#222"/>',
        f'<text x="{left + plot_w/2}" y="{height-18}" text-anchor="middle" font-family="Arial" font-size="12">beta closure</text>',
        f'<text x="18" y="{top + plot_h/2}" text-anchor="middle" transform="rotate(-90 18 {top + plot_h/2})" font-family="Arial" font-size="12">beta access</text>',
        f'<text x="{width-right-80}" y="28" font-family="Arial" font-size="11">min={vmin:.3f} max={vmax:.3f}</text>',
        "</svg>",
    ]
    out_path.write_text("\n".join(lines), encoding="utf-8")


def write_doc(summary: pd.DataFrame, signs: pd.DataFrame) -> None:
    true = signs[signs["true_dual_branch_sign"] == True].iloc[0]
    lines = [
        "# Dual-Branch Structure Validation",
        "",
        "Status: `completed`",
        "",
        "This locks down the proxy structure before replacing it with true histone/chromatin/promoter variables.",
        "",
        "## Key Results",
        "",
    ]
    for row in summary.itertuples():
        lines.append(f"- {row.summary_item}: {row.value:.4f} ({row.detail})")
    lines += [
        "",
        "## All-Sign Pattern Result",
        "",
        f"The biologically proposed sign pattern ranked {int(true.rank_by_occupancy_cosine)}/32 by occupancy then cosine.",
        "",
        "True sign pattern:",
        "",
        f"`{true.sign_pattern}`",
        "",
        "## Interpretation",
        "",
        "The dual-branch structure is not a single beta setting and not an arbitrary module sign assignment. It occupies a continuous high-performance beta region and the proposed M05/M01/M12 closure sign with opposite M02/M10 access sign is at the top of the exact sign-pattern ranking.",
    ]
    OUT_DOC.write_text("\n".join(lines) + "\n", encoding="utf-8")


def main() -> None:
    RESULTS.mkdir(parents=True, exist_ok=True)
    FIGURES.mkdir(parents=True, exist_ok=True)
    DOCS.mkdir(parents=True, exist_ok=True)
    context = latent_context()
    vecs = module_vectors()
    grid = run_beta_grid(context, vecs)
    order = run_module_order(context, vecs)
    signs = run_all_sign_patterns(context, vecs)
    summary = summarize(grid, order, signs)
    make_heatmap(grid, "pred_basin_occupancy_q90", OUT_GRID_OCC, "Dual-branch beta grid: occupancy")
    make_heatmap(grid, "direction_cosine_to_measured_correction", OUT_GRID_COS, "Dual-branch beta grid: cosine")
    write_doc(summary, signs)
    OUT_MANIFEST.write_text(
        json.dumps(
            {
                "status": "completed",
                "source_feature_set": SOURCE_FEATURE_SET,
                "outputs": [str(OUT_GRID), str(OUT_ORDER), str(OUT_SIGNS), str(OUT_SUMMARY), str(OUT_GRID_OCC), str(OUT_GRID_COS), str(OUT_DOC)],
            },
            indent=2,
        )
        + "\n",
        encoding="utf-8",
    )
    print(json.dumps({"status": "completed", "summary": summary.to_dict(orient="records")}, indent=2))


if __name__ == "__main__":
    main()
```
