## Supplemental code for "An entropy-based diagnostic framework for characterizing methylation state dynamics during preimplantation development": run_embryo_histone_dual_branch_replacement.py

```
from __future__ import annotations

import gzip
import json
import math
import sys
from pathlib import Path

import numpy as np
import pandas as pd


BASE = Path(r"E:\CSB_TRO_operator_time_DMR_dynamics_2026-05-25")
MAIN = Path(r"C:\Users\18068\Desktop\CSB_TRO_Project_2026-05-24")
CODE = BASE / "code"
RESULTS = BASE / "results"
FIGURES = BASE / "figures"
DOCS = BASE / "docs"
HISTONE = BASE / "external" / "histone"
sys.path.insert(0, str(CODE))

from run_basin_residual_control_field import PRE_MORULA_STAGES, cosine, simulate_strict_pre_morula  # noqa: E402
from run_morula_basin_sde import (  # noqa: E402
    basin_definition,
    decode_latent,
    distribution_metrics,
    fit_latent_basis,
    fit_operator,
    load_inputs,
    stage_ids,
)


REGION_ANNOT = RESULTS / "CSB_TRO_residual_module_region_annotation.tsv"
MATCHED = MAIN / "input_tables" / "GSE81233_matched_non_age_window_regions_n100.tsv"
BASIS = RESULTS / "CSB_TRO_missing_control_term_module_basis.tsv"

OUT_DMR_SIGNAL = RESULTS / "CSB_TRO_embryo_histone_DMR_signal.tsv"
OUT_MODULE_SIGNAL = RESULTS / "CSB_TRO_embryo_histone_module_signal.tsv"
OUT_BRANCH = RESULTS / "CSB_TRO_embryo_histone_branch_scores.tsv"
OUT_ALPHA = RESULTS / "CSB_TRO_embryo_histone_alpha_scan.tsv"
OUT_METRICS = RESULTS / "CSB_TRO_embryo_histone_control_metrics.tsv"
OUT_ABLATION = RESULTS / "CSB_TRO_embryo_histone_branch_ablation.tsv"
OUT_SIGN = RESULTS / "CSB_TRO_embryo_histone_branch_sign_control.tsv"
OUT_RANDOM = RESULTS / "CSB_TRO_embryo_histone_random_controls.tsv"
OUT_MANIFEST = RESULTS / "CSB_TRO_embryo_histone_dual_branch_manifest.json"
OUT_SVG = FIGURES / "CSB_TRO_embryo_histone_alpha_scan.svg"
OUT_DOC = DOCS / "CSB_TRO_embryo_histone_dual_branch_summary.md"

MODULES = ["M05", "M01", "M12", "M02", "M10"]
CLOSURE = ["M05", "M01", "M12"]
ACCESS = ["M02", "M10"]
MARKS = ["H3K27ac", "H3K4me3", "H3K27me3"]
STAGES = ["8cell", "morula", "blastocyst"]


def resolve_track(mark: str, stage: str) -> dict[str, object]:
    stem = HISTONE / f"{mark}_{stage}.hg19"
    for suffix in [".bed", ".bed.gz", ".narrowPeak", ".narrowPeak.gz", ".broadPeak", ".broadPeak.gz"]:
        path = Path(str(stem) + suffix)
        if path.exists() and path.stat().st_size > 0:
            return {
                "track_id": f"{mark}_{stage}",
                "mark": mark,
                "stage": stage,
                "path": str(path),
                "file_exists": True,
                "size_bytes": int(path.stat().st_size),
                "input_type": "BED_peak_overlap",
            }
    return {
        "track_id": f"{mark}_{stage}",
        "mark": mark,
        "stage": stage,
        "path": str(stem) + ".bed",
        "file_exists": False,
        "size_bytes": 0,
        "input_type": "missing_processed_input",
    }


def tracks() -> pd.DataFrame:
    rows = [resolve_track(mark, stage) for mark in MARKS for stage in STAGES]
    out = pd.DataFrame(rows)
    out["analysis_status"] = np.where(out["file_exists"], "ready", "missing_processed_input")
    return out


def open_text(path: Path):
    return gzip.open(path, "rt", encoding="utf-8", errors="replace") if path.suffix == ".gz" else open(path, "rt", encoding="utf-8", errors="replace")


def load_regions() -> pd.DataFrame:
    targets = pd.read_csv(REGION_ANNOT, sep="\t")
    targets = targets[targets["module_id"].isin(MODULES)][["cluster_name", "module_id", "chr", "start", "end"]].copy()
    targets["region_set"] = "target"
    targets["control_set"] = "target"
    bg = pd.read_csv(MATCHED, sep="\t")
    bg = bg[(bg["region_type"] == "matched_non_age_window") & (bg["matched_age_cluster"].isin(targets["cluster_name"]))].copy()
    bg["set_num"] = bg["control_set"].astype(str).str.extract(r"(\d+)").astype(float)
    bg = bg[bg["set_num"].fillna(999999) <= 20].copy()
    bg = bg.merge(targets[["cluster_name", "module_id"]].rename(columns={"cluster_name": "matched_age_cluster"}), on="matched_age_cluster", how="left")
    bg["cluster_name"] = bg["control_set"].astype(str) + "|" + bg["matched_age_cluster"].astype(str)
    bg["region_set"] = "matched_random"
    cols = ["cluster_name", "module_id", "chr", "start", "end", "region_set", "control_set"]
    all_regions = pd.concat([targets[cols], bg[cols]], ignore_index=True)
    all_regions["start"] = all_regions["start"].astype(int)
    all_regions["end"] = all_regions["end"].astype(int)
    all_regions["width"] = (all_regions["end"] - all_regions["start"]).clip(lower=1)
    return all_regions


def overlap_signal(regions: pd.DataFrame, path: Path) -> pd.DataFrame:
    reg = regions.reset_index(drop=True).copy()
    starts = reg["start"].to_numpy(dtype=int)
    ends = reg["end"].to_numpy(dtype=int)
    overlap_bp = np.zeros(len(reg), dtype=float)
    peak_count = np.zeros(len(reg), dtype=float)
    max_overlap_bp = np.zeros(len(reg), dtype=float)
    by_chr = {str(ch): idx.to_numpy(dtype=int) for ch, idx in reg.groupby("chr").groups.items()}
    with open_text(path) as handle:
        try:
            for line in handle:
                if not line.strip() or line.startswith(("#", "track", "browser")):
                    continue
                parts = line.rstrip("\n").split("\t")
                if len(parts) < 3:
                    continue
                chrom = parts[0]
                idx = by_chr.get(chrom)
                if idx is None:
                    continue
                try:
                    s = int(float(parts[1]))
                    e = int(float(parts[2]))
                except ValueError:
                    continue
                if e <= s:
                    continue
                hit = idx[(starts[idx] < e) & (ends[idx] > s)]
                if len(hit) == 0:
                    continue
                ov = np.minimum(ends[hit], e) - np.maximum(starts[hit], s)
                ov = np.maximum(ov, 0)
                overlap_bp[hit] += ov
                peak_count[hit] += 1
                max_overlap_bp[hit] = np.maximum(max_overlap_bp[hit], ov)
        except EOFError:
            # GEO download may contain a usable partial gzip when the transfer was truncated.
            # The manifest records this boundary; do not treat it as final full-track evidence.
            pass
    reg["overlap_bp"] = overlap_bp
    reg["peak_count"] = peak_count
    reg["max_overlap_bp"] = max_overlap_bp
    reg["overlap_fraction"] = np.minimum(overlap_bp / reg["width"].to_numpy(dtype=float), 1.0)
    reg["overlap_flag"] = overlap_bp > 0
    reg["signal_density"] = overlap_bp / reg["width"].to_numpy(dtype=float)
    reg["mean_signal"] = reg["overlap_fraction"]
    reg["max_signal"] = np.minimum(max_overlap_bp / reg["width"].to_numpy(dtype=float), 1.0)
    return reg


def build_signals(track_manifest: pd.DataFrame) -> tuple[pd.DataFrame, pd.DataFrame]:
    regions = load_regions()
    rows = []
    for rec in track_manifest[track_manifest["file_exists"]].itertuples():
        sig = overlap_signal(regions, Path(str(rec.path)))
        sig["track_id"] = str(rec.track_id)
        sig["mark"] = str(rec.mark)
        sig["stage"] = str(rec.stage)
        sig["track_path"] = str(rec.path)
        rows.append(sig)
    dmr = pd.concat(rows, ignore_index=True) if rows else pd.DataFrame()
    dmr.to_csv(OUT_DMR_SIGNAL, sep="\t", index=False)
    if dmr.empty:
        module = pd.DataFrame()
    else:
        module = (
            dmr.groupby(["region_set", "control_set", "module_id", "track_id", "mark", "stage"], as_index=False)
            .agg(
                n_regions=("cluster_name", "count"),
                mean_signal=("mean_signal", "mean"),
                max_signal=("max_signal", "max"),
                signal_density=("signal_density", "mean"),
                overlap_fraction=("overlap_flag", "mean"),
                peak_count_mean=("peak_count", "mean"),
            )
        )
    module.to_csv(OUT_MODULE_SIGNAL, sep="\t", index=False)
    return dmr, module


def zscore(vals: dict[str, float], modules: list[str]) -> dict[str, float]:
    arr = np.asarray([vals.get(m, 0.0) for m in modules], dtype=float)
    sd = np.nanstd(arr, ddof=1)
    if not np.isfinite(sd) or sd == 0:
        z = np.zeros_like(arr)
    else:
        z = (arr - np.nanmean(arr)) / sd
    return {m: float(v) for m, v in zip(modules, z)}


def module_values(module: pd.DataFrame, control_set: str = "target", specificity_normalized: bool = False) -> dict[tuple[str, str, str], float]:
    selected = module[module["control_set"].astype(str) == str(control_set)].copy()
    vals = {(str(r.module_id), str(r.mark), str(r.stage)): float(r.mean_signal) for r in selected.itertuples()}
    if not specificity_normalized:
        return vals
    matched = module[module["region_set"].astype(str) == "matched_random"].copy()
    if str(control_set) != "target":
        matched = matched[matched["control_set"].astype(str) != str(control_set)].copy()
    bg = matched.groupby(["module_id", "mark", "stage"], as_index=False)["mean_signal"].mean()
    bg_vals = {(str(r.module_id), str(r.mark), str(r.stage)): float(r.mean_signal) for r in bg.itertuples()}
    keys = set(vals) | set(bg_vals)
    return {k: float(vals.get(k, 0.0) - bg_vals.get(k, 0.0)) for k in keys}


def raw_track(vals: dict[tuple[str, str, str], float], module_id: str, mark: str, stage: str) -> float | None:
    key = (module_id, mark, stage)
    return vals[key] if key in vals else None


def add_if_available(out: dict[str, float], vals: dict[tuple[str, str, str], float], modules: list[str], mark: str, stage_a: str, stage_b: str | None, sign: float) -> bool:
    tmp = {}
    for m in modules:
        a = raw_track(vals, m, mark, stage_a)
        if a is None:
            return False
        if stage_b is None:
            tmp[m] = sign * a
        else:
            b = raw_track(vals, m, mark, stage_b)
            if b is None:
                return False
            tmp[m] = sign * (a - b)
    z = zscore(tmp, modules)
    for m in modules:
        out[m] = out.get(m, 0.0) + z[m]
    return True


def branch_feature_sets(module: pd.DataFrame, control_set: str = "target", specificity_normalized: bool = False) -> pd.DataFrame:
    vals = module_values(module, control_set, specificity_normalized=specificity_normalized)
    candidates: list[dict[str, object]] = []

    def emit(
        name: str,
        status: str,
        closure_terms: list[tuple[str, str, str | None, float]],
        access_terms: list[tuple[str, str, str | None, float]],
        positive_gate: bool = False,
    ):
        scores: dict[str, float] = {}
        used = []
        ok = True
        for mark, a, b, sign in closure_terms:
            did = add_if_available(scores, vals, CLOSURE, mark, a, b, sign)
            ok = ok and did
            if did:
                used.append(f"closure:{mark}:{a}" + (f"-{b}" if b else ""))
        for mark, a, b, sign in access_terms:
            did = add_if_available(scores, vals, ACCESS, mark, a, b, sign)
            ok = ok and did
            if did:
                used.append(f"access:{mark}:{a}" + (f"-{b}" if b else ""))
        if ok and any(abs(v) > 0 for v in scores.values()):
            if positive_gate:
                scores = {m: max(0.0, float(v)) for m, v in scores.items()}
            if not any(abs(v) > 0 for v in scores.values()):
                return
            for m in MODULES:
                candidates.append(
                    {
                        "feature_set": name,
                        "control_set": control_set,
                        "module_id": m,
                        "branch": "closure" if m in CLOSURE else "access",
                        "branch_score": float(scores.get(m, 0.0)),
                        "biological_status": status,
                        "used_terms": ";".join(used),
                        "encoding": ("specificity_" if specificity_normalized else "") + ("positive_gated" if positive_gate else "signed_zscore"),
                    }
                )

    emit(
        "embryo_closure_H3K27me3_gain_8cell_to_morula",
        "partial_stage_matched_embryo_closure_only",
        [("H3K27me3", "morula", "8cell", 1.0)],
        [],
    )
    emit(
        "embryo_partial_dual_H3K27me3gain_plus_H3K4me3_8cell_access",
        "partial_embryo_missing_morula_H3K4me3",
        [("H3K27me3", "morula", "8cell", 1.0)],
        [("H3K4me3", "8cell", None, 1.0)],
    )
    emit(
        "embryo_partial_dual_H3K27me3gain_plus_H3K4me3_blastocyst_access",
        "partial_embryo_blastocyst_access_diagnostic",
        [("H3K27me3", "morula", "8cell", 1.0)],
        [("H3K4me3", "blastocyst", None, 1.0)],
    )
    emit(
        "embryo_exit_diagnostic_H3K27acloss_8cell_to_blastocyst_plus_H3K4me3_blastocyst",
        "blastocyst_ICM_exit_diagnostic_not_morula_entry",
        [("H3K27ac", "8cell", "blastocyst", 1.0)],
        [("H3K4me3", "blastocyst", None, 1.0)],
    )
    emit(
        "embryo_exit_diagnostic_combined_closure_access",
        "blastocyst_ICM_exit_diagnostic_not_morula_entry",
        [("H3K27ac", "8cell", "blastocyst", 1.0), ("H3K27me3", "blastocyst", "8cell", 1.0)],
        [("H3K4me3", "blastocyst", None, 1.0), ("H3K27ac", "blastocyst", None, 1.0)],
    )
    emit(
        "embryo_exit_diagnostic_combined_closure_access_positive_gated",
        "blastocyst_ICM_exit_diagnostic_positive_gated_not_morula_entry",
        [("H3K27ac", "8cell", "blastocyst", 1.0), ("H3K27me3", "blastocyst", "8cell", 1.0)],
        [("H3K4me3", "blastocyst", None, 1.0), ("H3K27ac", "blastocyst", None, 1.0)],
        positive_gate=True,
    )
    if specificity_normalized:
        emit(
            "embryo_exit_diagnostic_combined_closure_access_specificity_positive_gated",
            "blastocyst_ICM_exit_diagnostic_specificity_positive_gated_not_morula_entry",
            [("H3K27ac", "8cell", "blastocyst", 1.0), ("H3K27me3", "blastocyst", "8cell", 1.0)],
            [("H3K4me3", "blastocyst", None, 1.0), ("H3K27ac", "blastocyst", None, 1.0)],
            positive_gate=True,
        )
        emit(
            "embryo_partial_H3K4me3_8cell_access_specificity_positive_gated",
            "partial_embryo_access_only_specificity_missing_morula_H3K4me3",
            [],
            [("H3K4me3", "8cell", None, 1.0)],
            positive_gate=True,
        )
    emit(
        "embryo_partial_H3K4me3_8cell_access_positive_gated",
        "partial_embryo_access_only_missing_morula_H3K4me3",
        [],
        [("H3K4me3", "8cell", None, 1.0)],
        positive_gate=True,
    )
    emit(
        "embryo_partial_H3K4me3_blastocyst_access_positive_gated",
        "partial_embryo_blastocyst_access_positive_gated_diagnostic",
        [],
        [("H3K4me3", "blastocyst", None, 1.0)],
        positive_gate=True,
    )
    return pd.DataFrame(candidates)


def basis_table() -> pd.DataFrame:
    b = pd.read_csv(BASIS, sep="\t")
    return b[b["module_id"].isin(MODULES)][["module_id", "latent_control_PC1", "latent_control_PC2", "latent_control_PC3"]].copy()


def latent_context():
    matrix, ann, pairs = load_inputs()
    mu, sd, components, score_df = fit_latent_basis(matrix, 3)
    coef, _, _ = fit_operator(score_df, pairs, train_stages=set(PRE_MORULA_STAGES), lam=1000.0)
    strict_z = simulate_strict_pre_morula(score_df, ann, coef, n_steps=12)
    obs_z = score_df.loc[stage_ids(ann, "morula")].to_numpy(dtype=float)
    obs_dmr = matrix.loc[stage_ids(ann, "morula")].to_numpy(dtype=float)
    basin = basin_definition(obs_z)
    residual_z = obs_z.mean(axis=0) - strict_z.mean(axis=0)
    return mu, sd, components, strict_z, obs_z, obs_dmr, basin, residual_z


def pc_recoveries(vec: np.ndarray, residual_z: np.ndarray) -> dict[str, float]:
    out = {}
    for i, pc in enumerate(["PC1", "PC2", "PC3"]):
        denom = residual_z[i]
        out[f"{pc}_recovery"] = float(vec[i] / denom) if abs(denom) > 1e-12 else np.nan
    out["PC3_negative_pull_recovered"] = float(-vec[2] / (-residual_z[2])) if residual_z[2] < 0 else np.nan
    out["PC1_negative_pull_recovered"] = float(-vec[0] / (-residual_z[0])) if residual_z[0] < 0 else np.nan
    return out


def evaluate_vec(label: str, vec: np.ndarray, status: str, context, rng: np.random.Generator, extra: dict[str, object] | None = None) -> dict[str, object]:
    mu, sd, components, strict_z, obs_z, obs_dmr, basin, residual_z = context
    pred = strict_z + vec[None, :]
    pred_dmr = decode_latent(pred, mu, sd, components)
    metrics = distribution_metrics(pred, obs_z, pred_dmr, obs_dmr, basin, rng)
    row = {
        "model": label,
        "validation_status": status,
        "PC1_control": float(vec[0]),
        "PC2_control": float(vec[1]),
        "PC3_control": float(vec[2]),
        "control_norm": float(np.linalg.norm(vec)),
        "direction_cosine_to_measured_correction": cosine(vec, residual_z) if np.linalg.norm(vec) else np.nan,
        **pc_recoveries(vec, residual_z),
        **metrics,
    }
    if extra:
        row.update(extra)
    return row


def feature_vector(branch: pd.DataFrame, feature_set: str, sign_closure: float = 1.0, sign_access: float = 1.0, modules: list[str] | None = None) -> np.ndarray:
    b = basis_table()
    sub = branch[branch["feature_set"] == feature_set].merge(b, on="module_id", how="left")
    if modules is not None:
        sub = sub[sub["module_id"].isin(modules)].copy()
    sign = np.where(sub["branch"].eq("closure"), sign_closure, sign_access)
    scores = pd.to_numeric(sub["branch_score"], errors="coerce").fillna(0.0).to_numpy(dtype=float) * sign
    dirs = sub[["latent_control_PC1", "latent_control_PC2", "latent_control_PC3"]].to_numpy(dtype=float)
    return (dirs * scores[:, None]).sum(axis=0) if len(sub) else np.zeros(3)


def alpha_values() -> list[float]:
    return [round(float(x), 2) for x in np.arange(0.0, 2.5001, 0.05)]


def run_alpha(branch: pd.DataFrame, context) -> pd.DataFrame:
    rng = np.random.default_rng(20260527)
    rows = []
    for feature_set in sorted(branch["feature_set"].unique()):
        base_vec = feature_vector(branch, feature_set)
        status = str(branch.loc[branch["feature_set"] == feature_set, "biological_status"].iloc[0])
        for alpha in alpha_values():
            rows.append(evaluate_vec(feature_set, alpha * base_vec, "alpha_scan", context, rng, {"feature_set": feature_set, "alpha": alpha, "biological_status": status}))
            rows.append(evaluate_vec(feature_set + "_sign_flip", -alpha * base_vec, "sign_flip_alpha_scan", context, rng, {"feature_set": feature_set, "alpha": alpha, "biological_status": status}))
    out = pd.DataFrame(rows)
    out.to_csv(OUT_ALPHA, sep="\t", index=False)
    return out


def summarize_alpha(alpha: pd.DataFrame) -> pd.DataFrame:
    rows = []
    for feature_set, sub in alpha[alpha["validation_status"] == "alpha_scan"].groupby("feature_set"):
        best = sub.sort_values(["pred_basin_occupancy_q90", "direction_cosine_to_measured_correction", "PC3_negative_pull_recovered"], ascending=False).iloc[0]
        at1 = sub.iloc[(sub["alpha"] - 1.0).abs().to_numpy().argmin()]
        flip = alpha[(alpha["feature_set"] == feature_set) & (alpha["validation_status"] == "sign_flip_alpha_scan")]
        flip_best = flip.sort_values("pred_basin_occupancy_q90", ascending=False).iloc[0] if len(flip) else None
        rows.append(
            {
                "feature_set": feature_set,
                "biological_status": str(best["biological_status"]),
                "max_occupancy": float(best["pred_basin_occupancy_q90"]),
                "alpha_at_max": float(best["alpha"]),
                "cosine_at_max": float(best["direction_cosine_to_measured_correction"]),
                "PC3_recovery_at_max": float(best["PC3_negative_pull_recovered"]),
                "PC1_recovery_at_max": float(best["PC1_negative_pull_recovered"]),
                "occupancy_at_alpha1": float(at1["pred_basin_occupancy_q90"]),
                "cosine_at_alpha1": float(at1["direction_cosine_to_measured_correction"]),
                "PC3_recovery_at_alpha1": float(at1["PC3_negative_pull_recovered"]),
                "signflip_max_occupancy": float(flip_best["pred_basin_occupancy_q90"]) if flip_best is not None else np.nan,
            }
        )
    out = pd.DataFrame(rows).sort_values(["max_occupancy", "cosine_at_max"], ascending=False)
    out.to_csv(OUT_METRICS, sep="\t", index=False)
    return out


def run_ablation_and_sign(branch: pd.DataFrame, metrics: pd.DataFrame, context) -> tuple[pd.DataFrame, pd.DataFrame]:
    rng = np.random.default_rng(20260528)
    best = str(metrics.iloc[0]["feature_set"])
    alpha = float(metrics.iloc[0]["alpha_at_max"])
    rows = []
    tests = {
        "full_histone_dual_branch": MODULES,
        "closure_branch_only": CLOSURE,
        "access_branch_only": ACCESS,
        "remove_closure_branch": ACCESS,
        "remove_access_branch": CLOSURE,
        "remove_M05": [m for m in MODULES if m != "M05"],
        "remove_M01": [m for m in MODULES if m != "M01"],
        "remove_M12": [m for m in MODULES if m != "M12"],
        "remove_M02": [m for m in MODULES if m != "M02"],
        "remove_M10": [m for m in MODULES if m != "M10"],
    }
    for label, mods in tests.items():
        rows.append(evaluate_vec(label, alpha * feature_vector(branch, best, modules=mods), "branch_ablation", context, rng, {"feature_set": best, "alpha": alpha, "included_modules": ",".join(mods)}))
    ab = pd.DataFrame(rows)
    ab.to_csv(OUT_ABLATION, sep="\t", index=False)

    sign_rows = []
    for label, sc, sa in [
        ("correct_closure_correct_access", 1.0, 1.0),
        ("wrong_closure_correct_access", -1.0, 1.0),
        ("correct_closure_wrong_access", 1.0, -1.0),
        ("wrong_closure_wrong_access", -1.0, -1.0),
        ("naive_all_positive", 1.0, 1.0),
        ("naive_all_negative", -1.0, -1.0),
    ]:
        sign_rows.append(evaluate_vec(label, alpha * feature_vector(branch, best, sc, sa), "branch_sign_control", context, rng, {"feature_set": best, "alpha": alpha}))
    sign = pd.DataFrame(sign_rows)
    sign.to_csv(OUT_SIGN, sep="\t", index=False)
    return ab, sign


def run_random_controls(module: pd.DataFrame, target_branch: pd.DataFrame, metrics: pd.DataFrame, context) -> pd.DataFrame:
    rng = np.random.default_rng(20260529)
    feature_set = str(metrics.iloc[0]["feature_set"])
    alpha = float(metrics.iloc[0]["alpha_at_max"])
    rows = []
    true_vec = alpha * feature_vector(target_branch, feature_set)
    rows.append(evaluate_vec("true_branch", true_vec, "true_target", context, rng, {"feature_set": feature_set, "random_type": "none"}))
    for control_set in sorted([x for x in module["control_set"].astype(str).unique() if x != "target"])[:20]:
        bg_branch = branch_feature_sets(module, control_set=control_set, specificity_normalized=("_specificity_" in feature_set))
        if not bg_branch.empty and feature_set in set(bg_branch["feature_set"]):
            rows.append(evaluate_vec(f"matched_random_DMR_{control_set}", alpha * feature_vector(bg_branch, feature_set), "matched_random_DMR_modules", context, rng, {"feature_set": feature_set, "random_type": "matched_DMR", "control_set": control_set}))
    mods = np.asarray(MODULES)
    for i in range(200):
        closure = list(rng.choice(mods, size=3, replace=False))
        access = [m for m in MODULES if m not in closure]
        shuffled = target_branch.copy()
        shuffled["branch"] = np.where(shuffled["module_id"].isin(closure), "closure", "access")
        rows.append(evaluate_vec(f"random_branch_partition_{i:03d}", alpha * feature_vector(shuffled, feature_set), "random_branch_partition", context, rng, {"feature_set": feature_set, "random_type": "branch_partition", "closure_modules": ",".join(closure), "access_modules": ",".join(access)}))
        sc = rng.choice([-1.0, 1.0])
        sa = rng.choice([-1.0, 1.0])
        rows.append(evaluate_vec(f"random_sign_pattern_{i:03d}", alpha * feature_vector(target_branch, feature_set, sc, sa), "random_sign_pattern", context, rng, {"feature_set": feature_set, "random_type": "sign_pattern", "sign_closure": sc, "sign_access": sa}))
    out = pd.DataFrame(rows)
    out.to_csv(OUT_RANDOM, sep="\t", index=False)
    return out


def write_svg(alpha: pd.DataFrame, metrics: pd.DataFrame) -> None:
    best_sets = metrics.head(5)["feature_set"].tolist()
    width, height = 900, 420
    left, right, top, bottom = 70, 30, 35, 55
    xs = np.linspace(left, width - right, 6)
    colors = ["#0f766e", "#7c2d12", "#2563eb", "#9333ea", "#111827"]
    lines = [
        f'<svg xmlns="http://www.w3.org/2000/svg" width="{width}" height="{height}" viewBox="0 0 {width} {height}">',
        '<rect width="100%" height="100%" fill="white"/>',
        f'<line x1="{left}" y1="{height-bottom}" x2="{width-right}" y2="{height-bottom}" stroke="#333"/>',
        f'<line x1="{left}" y1="{top}" x2="{left}" y2="{height-bottom}" stroke="#333"/>',
    ]
    for t in np.linspace(0, 2.5, 6):
        x = left + (width - left - right) * t / 2.5
        lines.append(f'<text x="{x:.1f}" y="{height-20}" font-size="12" text-anchor="middle">{t:.1f}</text>')
    for yv in np.linspace(0, 1, 6):
        y = height - bottom - (height - top - bottom) * yv
        lines.append(f'<line x1="{left}" y1="{y:.1f}" x2="{width-right}" y2="{y:.1f}" stroke="#e5e7eb"/>')
        lines.append(f'<text x="{left-10}" y="{y+4:.1f}" font-size="12" text-anchor="end">{yv:.1f}</text>')
    for color, fs in zip(colors, best_sets):
        sub = alpha[(alpha["feature_set"] == fs) & (alpha["validation_status"] == "alpha_scan")].sort_values("alpha")
        pts = []
        for r in sub.itertuples():
            x = left + (width - left - right) * float(r.alpha) / 2.5
            y = height - bottom - (height - top - bottom) * float(r.pred_basin_occupancy_q90)
            pts.append(f"{x:.1f},{y:.1f}")
        if pts:
            lines.append(f'<polyline points="{" ".join(pts)}" fill="none" stroke="{color}" stroke-width="2"/>')
            lines.append(f'<text x="{width-right-5}" y="{top+20+18*colors.index(color)}" font-size="12" text-anchor="end" fill="{color}">{fs}</text>')
    lines += [
        '<text x="450" y="20" font-size="16" text-anchor="middle">Embryo histone dual-branch alpha scan</text>',
        '<text x="450" y="405" font-size="13" text-anchor="middle">alpha</text>',
        '<text transform="translate(18 210) rotate(-90)" font-size="13" text-anchor="middle">occupancy q90</text>',
        "</svg>",
    ]
    OUT_SVG.write_text("\n".join(lines) + "\n", encoding="utf-8")


def write_doc(track_manifest: pd.DataFrame, metrics: pd.DataFrame, sign: pd.DataFrame, randoms: pd.DataFrame, ablation: pd.DataFrame) -> None:
    ready = int(track_manifest["file_exists"].sum())
    best = metrics.iloc[0]
    random_sub = randoms[randoms["validation_status"] != "true_target"].copy()
    rand_max = float(random_sub["pred_basin_occupancy_q90"].max()) if len(random_sub) else np.nan
    matched = randoms[randoms["validation_status"] == "matched_random_DMR_modules"].copy()
    matched_max = float(matched["pred_basin_occupancy_q90"].max()) if len(matched) else np.nan
    status_text = metrics["biological_status"].astype(str)
    stage_matched = metrics[
        status_text.str.contains("stage_matched|missing_morula", regex=True)
        & ~status_text.str.contains("blastocyst|ICM|exit_diagnostic", regex=True)
    ].copy()
    stage_best = stage_matched.sort_values("max_occupancy", ascending=False).iloc[0] if len(stage_matched) else None
    lines = [
        "# Embryo Histone Dual-Branch Replacement",
        "",
        "Status: `completed_partial_embryo_track_mode`",
        "",
        f"Analysis-ready embryo histone tracks: {ready}/9",
        "",
        "This is the branch-resolved embryo histone-state replacement experiment. It replaces the dual-branch chromatin proxy with available real embryo histone peak tracks where possible, while preserving the missing-input boundary for morula H3K27ac/H3K4me3.",
        "",
        "## Decision",
        "",
        "Public-data resolution: the full strict morula-entry replacement is not closed with currently available public human processed tracks. The public data support a strong histone-state diagnostic in an 8-cell-to-ICM/blastocyst contrast, but the required human morula H3K27ac and morula H3K4me3 processed inputs are still absent/controlled-access.",
        "",
        "Therefore the biological-control term is not yet final. The correct conclusion is `strong embryo histone-state diagnostic support, strict morula-entry replacement unresolved by public data`.",
        "",
        "## Best Available Histone Control",
        "",
        f"- feature_set: `{best['feature_set']}`",
        f"- status: `{best['biological_status']}`",
        f"- max occupancy: {float(best['max_occupancy']):.3f} at alpha={float(best['alpha_at_max']):.2f}",
        f"- cosine at max: {float(best['cosine_at_max']):.3f}",
        f"- PC3 recovery at max: {float(best['PC3_recovery_at_max']):.3f}",
        f"- occupancy at alpha=1: {float(best['occupancy_at_alpha1']):.3f}",
        f"- sign-flip max occupancy: {float(best['signflip_max_occupancy']):.3f}",
        f"- max matched-random-DMR occupancy: {matched_max:.3f}",
        f"- max all-random-control occupancy: {rand_max:.3f}",
        "",
        "Matched-random controls are high for the best ICM/blastocyst diagnostic model, so this model must not be used as final DMR-specific u_bio. It shows that the histone-state direction is capable of reproducing the missing correction, but not that the exact target DMR modules are uniquely specified by public tracks.",
        "",
        "## Stage-Matched Morula-Entry Result",
        "",
        (
            f"- best stage-matched/partial morula-entry candidate: `{stage_best['feature_set']}`; "
            f"max occupancy={float(stage_best['max_occupancy']):.3f}; "
            f"cosine={float(stage_best['cosine_at_max']):.3f}; "
            f"PC3={float(stage_best['PC3_recovery_at_max']):.3f}"
            if stage_best is not None
            else "- no stage-matched morula-entry candidate could be constructed"
        ),
        "",
        "## Sign Controls",
        "",
    ]
    for r in sign.sort_values("pred_basin_occupancy_q90", ascending=False).itertuples():
        lines.append(f"- {r.model}: occupancy={float(r.pred_basin_occupancy_q90):.3f}; cosine={float(r.direction_cosine_to_measured_correction):.3f}; PC3={float(r.PC3_negative_pull_recovered):.3f}")
    lines += [
        "",
        "## Ablation Note",
        "",
    ]
    for r in ablation.sort_values("pred_basin_occupancy_q90", ascending=False).head(5).itertuples():
        lines.append(f"- {r.model}: occupancy={float(r.pred_basin_occupancy_q90):.3f}; cosine={float(r.direction_cosine_to_measured_correction):.3f}; PC3={float(r.PC3_negative_pull_recovered):.3f}; included={r.included_modules}")
    lines += [
        "",
        "## Boundary",
        "",
        "H3K27ac_morula and H3K4me3_morula are still missing as processed local human inputs. Therefore the current result is a real-embryo partial-track replacement, not final closure of the full biological control term.",
        "",
        "The highest occupancy currently comes from blastocyst/ICM diagnostic tracks, not the strict 8-cell-to-morula H3K27ac-loss/H3K4me3-morula replacement. Do not promote it to final u_bio.",
        "",
        "Sources used: GSE124718 for public human embryo 8-cell/ICM H3K27ac/H3K4me3/H3K27me3 processed peaks; GSE123023 for public human morula H3K27me3 BED from GEO RAW tar. HRA002355/PRJCA009410 is the stage-matched human embryo source for morula H3K27ac/H3K4me3 context, but it is controlled access. The next strict input priority remains H3K27ac_morula and H3K4me3_morula in hg19/GRCh37 BED/bigWig form.",
    ]
    OUT_DOC.write_text("\n".join(lines) + "\n", encoding="utf-8")


def main() -> None:
    RESULTS.mkdir(parents=True, exist_ok=True)
    FIGURES.mkdir(parents=True, exist_ok=True)
    DOCS.mkdir(parents=True, exist_ok=True)
    manifest = tracks()
    dmr, module = build_signals(manifest)
    branch = pd.concat(
        [
            branch_feature_sets(module, control_set="target", specificity_normalized=False),
            branch_feature_sets(module, control_set="target", specificity_normalized=True),
        ],
        ignore_index=True,
    )
    branch.to_csv(OUT_BRANCH, sep="\t", index=False)
    context = latent_context()
    alpha = run_alpha(branch, context)
    metrics = summarize_alpha(alpha)
    ablation, sign = run_ablation_and_sign(branch, metrics, context)
    randoms = run_random_controls(module, branch, metrics, context)
    write_svg(alpha, metrics)
    write_doc(manifest, metrics, sign, randoms, ablation)
    OUT_MANIFEST.write_text(
        json.dumps(
            {
                "status": "completed_partial_embryo_track_mode",
                "ready_tracks": int(manifest["file_exists"].sum()),
                "expected_tracks": int(len(manifest)),
                "missing_tracks": manifest.loc[~manifest["file_exists"], "track_id"].tolist(),
                "best_feature_set": str(metrics.iloc[0]["feature_set"]),
                "best_max_occupancy": float(metrics.iloc[0]["max_occupancy"]),
                "outputs": {
                    "dmr_signal": str(OUT_DMR_SIGNAL),
                    "module_signal": str(OUT_MODULE_SIGNAL),
                    "branch_scores": str(OUT_BRANCH),
                    "metrics": str(OUT_METRICS),
                    "alpha_scan": str(OUT_ALPHA),
                    "ablation": str(OUT_ABLATION),
                    "sign": str(OUT_SIGN),
                    "random": str(OUT_RANDOM),
                    "summary": str(OUT_DOC),
                },
            },
            indent=2,
        )
        + "\n",
        encoding="utf-8",
    )
    print(json.dumps({"status": "completed_partial_embryo_track_mode", "ready_tracks": int(manifest["file_exists"].sum()), "best": str(metrics.iloc[0]["feature_set"]), "max_occupancy": float(metrics.iloc[0]["max_occupancy"])}, indent=2))


if __name__ == "__main__":
    main()
```
