## Supplemental code for "An entropy-based diagnostic framework for characterizing methylation state dynamics during preimplantation development": run_export_m02_klf_target_bed.py

```
from __future__ import annotations

import json
from pathlib import Path

import pandas as pd


BASE = Path(r"E:\CSB_TRO_operator_time_DMR_dynamics_2026-05-25")
RESULTS = BASE / "results"
EXTERNAL = BASE / "external"
DOCS = BASE / "docs"

DMR_RANKING = RESULTS / "CSB_TRO_basin_residual_DMR_ranking.tsv"
HITS = RESULTS / "CSB_TRO_JASPAR_pwm_matched_background_hits.tsv"
OUT_BED = EXTERNAL / "histone" / "M02_KLF4_KLF5_residual_DMR_targets.hg19.bed"
OUT_TSV = RESULTS / "CSB_TRO_M02_KLF4_KLF5_residual_DMR_targets.tsv"
OUT_DOC = DOCS / "CSB_TRO_M02_KLF4_KLF5_target_regions.md"
OUT_MANIFEST = RESULTS / "CSB_TRO_M02_KLF4_KLF5_target_regions_manifest.json"


def main() -> None:
    EXTERNAL.joinpath("histone").mkdir(parents=True, exist_ok=True)
    DOCS.mkdir(parents=True, exist_ok=True)
    dmr = pd.read_csv(DMR_RANKING, sep="\t")
    hits = pd.read_csv(HITS, sep="\t")
    hits = hits[(hits["module_id"] == "M02") & (hits["TF"].isin(["KLF4", "KLF5"]))].copy()
    hit_summary = (
        hits.groupby("cluster_name", as_index=False)
        .agg(
            TFs=("TF", lambda x: ",".join(sorted(set(map(str, x))))),
            motif_ids=("motif_id", lambda x: ",".join(sorted(set(map(str, x))))),
            max_pwm_score=("score", "max"),
        )
    )
    targets = dmr.merge(hit_summary, on="cluster_name", how="inner")
    targets = targets.sort_values(["basin_residual_rank", "cluster_name"])
    targets.to_csv(OUT_TSV, sep="\t", index=False)
    bed = targets[["chr", "start", "end", "cluster_name", "basin_residual_rank", "TFs"]].copy()
    bed["name"] = bed["cluster_name"] + "|" + bed["TFs"]
    bed[["chr", "start", "end", "name", "basin_residual_rank"]].to_csv(OUT_BED, sep="\t", index=False, header=False)

    lines = [
        "# M02 KLF4/KLF5 residual DMR targets",
        "",
        "These hg19 regions are the M02 residual DMRs with matched-background JASPAR KLF4 and/or KLF5 motif hits.",
        "",
        f"Target DMRs: {len(targets)}",
        "",
        "Top targets:",
    ]
    for row in targets.head(20).itertuples(index=False):
        lines.append(f"- {row.cluster_name} {row.chr}:{int(row.start)}-{int(row.end)} TFs={row.TFs} residual_rank={row.basin_residual_rank}")
    OUT_DOC.write_text("\n".join(lines) + "\n", encoding="utf-8")
    OUT_MANIFEST.write_text(
        json.dumps(
            {
                "genome_build": "hg19/GRCh37",
                "n_targets": int(len(targets)),
                "outputs": [str(OUT_BED), str(OUT_TSV), str(OUT_DOC)],
            },
            indent=2,
        )
        + "\n",
        encoding="utf-8",
    )
    print(json.dumps({"n_targets": int(len(targets)), "bed": str(OUT_BED)}, indent=2))


if __name__ == "__main__":
    main()
```
