## Supplemental code for "An entropy-based diagnostic framework for characterizing methylation state dynamics during preimplantation development": run_external_biocontrol_data_integration.py

```
from __future__ import annotations

import argparse
import gzip
import json
import re
from pathlib import Path

import numpy as np
import pandas as pd


BASE = Path(r"E:\CSB_TRO_operator_time_DMR_dynamics_2026-05-25")
EXTERNAL = BASE / "external"
RESULTS = BASE / "results"
DOCS = BASE / "docs"

DMR_RANKING = RESULTS / "CSB_TRO_basin_residual_DMR_ranking.tsv"
MODULE_BASIS = RESULTS / "CSB_TRO_missing_control_term_module_basis.tsv"

OUT_INPUT_MANIFEST = RESULTS / "CSB_TRO_real_u_bio_input_manifest.tsv"
OUT_AUDIT = RESULTS / "CSB_TRO_genome_build_audit.tsv"
OUT_CHR_SUMMARY = RESULTS / "CSB_TRO_DMR_coordinate_chr_summary.tsv"
OUT_TSS = RESULTS / "CSB_TRO_external_gene_TSS.tsv"
OUT_GENE_ANNOT = RESULTS / "CSB_TRO_residual_DMR_gene_annotation.tsv"
OUT_MODULE_LINKS = RESULTS / "CSB_TRO_residual_module_gene_links.tsv"
OUT_MODULE_LINK_SUMMARY = RESULTS / "CSB_TRO_residual_module_gene_link_summary.tsv"
OUT_REGION_ANNOT = RESULTS / "CSB_TRO_residual_module_region_annotation.tsv"
OUT_FEATURE_TEMPLATE = RESULTS / "CSB_TRO_external_biocontrol_feature_template.tsv"
OUT_DOC = DOCS / "CSB_TRO_external_biocontrol_interpretation.md"
OUT_MANIFEST = RESULTS / "CSB_TRO_external_biocontrol_manifest.json"

PRIORITY_MODULES = ["M05", "M01", "M12", "M02", "M10"]

HG19_LENGTHS = {
    "chr1": 249250621,
    "chr2": 243199373,
    "chr3": 198022430,
    "chr4": 191154276,
    "chr5": 180915260,
    "chr6": 171115067,
    "chr7": 159138663,
    "chr8": 146364022,
    "chr9": 141213431,
    "chr10": 135534747,
    "chr11": 135006516,
    "chr12": 133851895,
    "chr13": 115169878,
    "chr14": 107349540,
    "chr15": 102531392,
    "chr16": 90354753,
    "chr17": 81195210,
    "chr18": 78077248,
    "chr19": 59128983,
    "chr20": 63025520,
    "chr21": 48129895,
    "chr22": 51304566,
    "chrX": 155270560,
    "chrY": 59373566,
}

HG38_LENGTHS = {
    "chr1": 248956422,
    "chr2": 242193529,
    "chr3": 198295559,
    "chr4": 190214555,
    "chr5": 181538259,
    "chr6": 170805979,
    "chr7": 159345973,
    "chr8": 145138636,
    "chr9": 138394717,
    "chr10": 133797422,
    "chr11": 135086622,
    "chr12": 133275309,
    "chr13": 114364328,
    "chr14": 107043718,
    "chr15": 101991189,
    "chr16": 90338345,
    "chr17": 83257441,
    "chr18": 80373285,
    "chr19": 58617616,
    "chr20": 64444167,
    "chr21": 46709983,
    "chr22": 50818468,
    "chrX": 156040895,
    "chrY": 57227415,
}


def read_tsv(path: Path) -> pd.DataFrame:
    return pd.read_csv(path, sep="\t") if path.exists() else pd.DataFrame()


def normalize_chr(value: str) -> str:
    v = str(value)
    return v if v.startswith("chr") else f"chr{v}"


def parse_attrs(attr: str) -> dict[str, str]:
    out = {}
    for part in str(attr).split(";"):
        part = part.strip()
        if not part:
            continue
        m = re.match(r'([^ ]+) "?([^"]+)"?$', part)
        if m:
            out[m.group(1)] = m.group(2)
    return out


def gtf_to_tss(path: Path) -> pd.DataFrame:
    if not path.exists():
        return pd.DataFrame(columns=["chr", "tss", "gene_id", "gene_name", "gene_type", "strand"])
    rows = []
    opener = gzip.open if path.suffix == ".gz" else open
    with opener(path, "rt", encoding="utf-8", errors="replace") as handle:
        for line in handle:
            if not line.strip() or line.startswith("#"):
                continue
            fields = line.rstrip("\n").split("\t")
            if len(fields) < 9 or fields[2] != "gene":
                continue
            chrom, start, end, strand, attr = fields[0], fields[3], fields[4], fields[6], fields[8]
            attrs = parse_attrs(attr)
            tss = int(start) if strand != "-" else int(end)
            rows.append(
                {
                    "chr": normalize_chr(chrom),
                    "tss": tss,
                    "gene_id": attrs.get("gene_id", attrs.get("ID", "")),
                    "gene_name": attrs.get("gene_name", attrs.get("Name", attrs.get("gene_id", ""))),
                    "gene_type": attrs.get("gene_type", attrs.get("gene_biotype", attrs.get("biotype", "NA"))),
                    "strand": strand,
                }
            )
    return pd.DataFrame(rows)


def first_existing(paths: list[Path]) -> Path:
    for path in paths:
        if path.exists():
            return path
    return paths[0]


def build_input_manifest(external: Path) -> pd.DataFrame:
    gtf_path = first_existing(
        [
            external / "annotations" / "gencode.gtf",
            external / "annotations" / "gencode.gtf.gz",
            external / "annotations" / "gencode.v19.annotation.gtf.gz",
        ]
    )
    rows = [
        ("genome_build", external / "genome_build.txt", "required", "Set exact coordinate build: hg19/GRCh37 or hg38/GRCh38."),
        ("gencode_gtf", gtf_path, "required_for_gene_linking", "GENCODE/RefSeq gene model in the same build as DMRs."),
        ("gene_tss", external / "annotations" / "gene_tss.tsv", "optional_if_gtf_present", "TSV with chr,tss,gene_id/gene_name,gene_type."),
        ("cpg_islands", external / "annotations" / "cpg_islands.bed", "optional", "CpG island/shore/shelf annotation in same build."),
        ("repeatmasker", external / "annotations" / "repeatmasker.bed", "optional", "RepeatMasker or simple repeat BED in same build."),
        ("rna_gene_matrix", external / "rna" / "gene_stage_matrix.tsv", "required_for_module_linked_RNA", "Gene x stage expression with 8-cell,morula[,blastocyst]."),
        ("jaspar_meme", external / "motif" / "jaspar_vertebrates.meme", "required_for_motif_TF", "Motif database for module-specific motif enrichment."),
        ("motif_enrichment", external / "motif" / "module_motif_enrichment.tsv", "optional_if_precomputed", "module_id,TF,motif_score/enrichment/NES/pvalue."),
        ("atac_features", external / "atac" / "module_ATAC_features.tsv", "optional", "module_id,control_value feature table or stage-specific overlaps."),
        ("histone_features", external / "histone" / "module_histone_features.tsv", "optional", "module_id,control_value for H3K27ac/H3K4me3/H3K27me3."),
    ]
    manifest = pd.DataFrame(rows, columns=["input_name", "expected_path", "priority", "description"])
    manifest["exists"] = manifest["expected_path"].map(lambda p: Path(p).exists())
    return manifest


def coordinate_audit(dmr: pd.DataFrame, genome_build_text: str) -> tuple[pd.DataFrame, pd.DataFrame]:
    d = dmr.copy()
    d["chr"] = d["chr"].map(normalize_chr)
    d["start"] = pd.to_numeric(d["start"], errors="coerce")
    d["end"] = pd.to_numeric(d["end"], errors="coerce")
    chr_summary = (
        d.groupby("chr", as_index=False)
        .agg(n_DMRs=("cluster_name", "count"), min_start=("start", "min"), max_end=("end", "max"))
        .sort_values("chr")
    )
    for label, lengths in [("hg19", HG19_LENGTHS), ("hg38", HG38_LENGTHS)]:
        chr_summary[f"{label}_chr_known"] = chr_summary["chr"].isin(lengths)
        chr_summary[f"{label}_chr_length"] = chr_summary["chr"].map(lengths)
        chr_summary[f"{label}_within_length"] = chr_summary["max_end"] <= chr_summary[f"{label}_chr_length"]
    rows = []
    chr_style = "chr_prefixed" if d["chr"].astype(str).str.startswith("chr").all() else "mixed_or_unprefixed"
    build_status = {}
    for label in ["hg19", "hg38"]:
        known = bool(chr_summary[f"{label}_chr_known"].all())
        within = bool(chr_summary[f"{label}_within_length"].fillna(False).all())
        build_status[label] = known and within
        rows.append(
            {
                "audit_item": f"{label}_coordinate_bounds",
                "status": "pass" if known and within else "fail",
                "detail": f"all_chr_known={known}; all_DMRs_within_chr_lengths={within}",
            }
        )
    declared = genome_build_text.splitlines()[0].strip() if genome_build_text.strip() else "UNKNOWN"
    if build_status["hg19"] and not build_status["hg38"]:
        discr_status = "hg19_supported_by_bounds"
        discr_detail = "At least one DMR chromosome max coordinate exceeds hg38 but all DMRs fit hg19. Treat coordinates as hg19/GRCh37 unless source metadata proves otherwise."
    elif build_status["hg38"] and not build_status["hg19"]:
        discr_status = "hg38_supported_by_bounds"
        discr_detail = "At least one DMR coordinate pattern supports hg38 over hg19. Confirm with source metadata before overlap."
    elif build_status["hg19"] and build_status["hg38"]:
        discr_status = "ambiguous"
        discr_detail = "DMR coordinate bounds are compatible with both hg19 and hg38; origin metadata or liftOver/spot-check is required before external overlap."
    else:
        discr_status = "fail"
        discr_detail = "DMR coordinates do not cleanly fit hg19 or hg38 chromosome bounds; inspect source files before external overlap."
    rows.extend(
        [
            {"audit_item": "declared_genome_build", "status": "unknown" if declared.upper() == "UNKNOWN" else "declared", "detail": declared},
            {"audit_item": "chr_naming_style", "status": "pass" if chr_style == "chr_prefixed" else "warning", "detail": chr_style},
            {"audit_item": "build_discrimination", "status": discr_status, "detail": discr_detail},
        ]
    )
    return pd.DataFrame(rows), chr_summary


def nearest_gene_links(dmr: pd.DataFrame, tss: pd.DataFrame) -> pd.DataFrame:
    columns = [
        "cluster_name",
        "module_id",
        "chr",
        "start",
        "end",
        "nearest_gene",
        "gene_id",
        "distance_to_TSS",
        "abs_distance_to_TSS",
        "gene_type",
        "link_type",
        "promoter_2kb",
        "promoter_5kb",
        "gene_body",
        "intergenic",
        "residual_contribution",
        "module_basis_weight",
    ]
    if tss.empty:
        return pd.DataFrame(columns=columns)
    tss = tss.copy()
    tss["chr"] = tss["chr"].map(normalize_chr)
    tss["tss"] = pd.to_numeric(tss["tss"], errors="coerce")
    by_chr = {c: sub.dropna(subset=["tss"]) for c, sub in tss.groupby("chr")}
    basis = read_tsv(MODULE_BASIS)
    weight_map = basis.set_index("module_id")["ridge_weight"].to_dict() if not basis.empty and "ridge_weight" in basis.columns else {}
    rows = []
    for row in dmr.itertuples(index=False):
        sub = by_chr.get(normalize_chr(row.chr))
        if sub is None or sub.empty:
            continue
        mid = 0.5 * (float(row.start) + float(row.end))
        dist = sub["tss"].to_numpy(dtype=float) - mid
        i = int(np.argmin(np.abs(dist)))
        hit = sub.iloc[i]
        abs_dist = float(abs(dist[i]))
        rows.append(
            {
                "cluster_name": row.cluster_name,
                "module_id": row.module_id,
                "chr": normalize_chr(row.chr),
                "start": int(row.start),
                "end": int(row.end),
                "nearest_gene": hit.get("gene_name", hit.get("gene_id", "")),
                "gene_id": hit.get("gene_id", ""),
                "distance_to_TSS": float(dist[i]),
                "abs_distance_to_TSS": abs_dist,
                "gene_type": hit.get("gene_type", "NA"),
                "link_type": "nearest_TSS",
                "promoter_2kb": abs_dist <= 2000,
                "promoter_5kb": abs_dist <= 5000,
                "gene_body": False,
                "intergenic": abs_dist > 5000,
                "residual_contribution": float(getattr(row, "abs_latent_residual_delta_beta", np.nan)),
                "module_basis_weight": float(weight_map.get(row.module_id, np.nan)),
            }
        )
    return pd.DataFrame(rows, columns=columns)


def feature_template(module_ids: list[str]) -> pd.DataFrame:
    rows = []
    for module_id in module_ids:
        for modality in ["RNA", "motif_activity", "ATAC", "histone"]:
            rows.append(
                {
                    "module_id": module_id,
                    "candidate_control": f"{modality}_external_{module_id}",
                    "control_modality": modality,
                    "control_value": np.nan,
                    "control_stage_window": "8-cell_to_morula",
                    "leakage_status": "user_supplied_external_feature",
                    "interpretation": "Fill from external module-linked data; do not use morula methylation residual to define the value.",
                }
            )
    return pd.DataFrame(rows)


def module_gene_link_summary(links: pd.DataFrame) -> pd.DataFrame:
    cols = [
        "module_id",
        "n_DMRs_linked",
        "n_unique_nearest_genes",
        "n_promoter_2kb",
        "n_promoter_5kb",
        "median_abs_distance_to_TSS",
        "top_nearest_genes",
    ]
    if links.empty:
        return pd.DataFrame(columns=cols)
    rows = []
    for module_id, sub in links.groupby("module_id"):
        genes = [g for g in sub.sort_values("abs_distance_to_TSS")["nearest_gene"].astype(str).tolist() if g]
        rows.append(
            {
                "module_id": module_id,
                "n_DMRs_linked": int(len(sub)),
                "n_unique_nearest_genes": int(sub["nearest_gene"].nunique()),
                "n_promoter_2kb": int(pd.to_numeric(sub["promoter_2kb"], errors="coerce").fillna(False).sum()),
                "n_promoter_5kb": int(pd.to_numeric(sub["promoter_5kb"], errors="coerce").fillna(False).sum()),
                "median_abs_distance_to_TSS": float(pd.to_numeric(sub["abs_distance_to_TSS"], errors="coerce").median()),
                "top_nearest_genes": ",".join(genes[:20]),
            }
        )
    out = pd.DataFrame(rows)
    if "module_id" in out.columns:
        out["priority_module"] = out["module_id"].isin(PRIORITY_MODULES)
        out = out.sort_values(["priority_module", "module_id"], ascending=[False, True])
    return out[cols + (["priority_module"] if "priority_module" in out.columns else [])]


def write_doc(path: Path, audit: pd.DataFrame, manifest: pd.DataFrame, n_links: int) -> None:
    declared = audit.loc[audit["audit_item"] == "declared_genome_build", "detail"].iloc[0]
    discr = audit.loc[audit["audit_item"] == "build_discrimination"].iloc[0]
    if discr["status"] == "hg19_supported_by_bounds":
        boundary = "Coordinate bounds support hg19/GRCh37 over hg38/GRCh38 because at least one DMR chromosome maximum exceeds hg38 chromosome length while all DMRs fit hg19. Use hg19/GRCh37 external annotations unless source metadata proves otherwise."
    elif discr["status"] == "hg38_supported_by_bounds":
        boundary = "Coordinate bounds support hg38/GRCh38 over hg19/GRCh37. Confirm source metadata before external overlap."
    else:
        boundary = "Coordinate bounds alone cannot distinguish hg19 from hg38 for these DMRs. Do not run external GTF/BED/peak overlap until the source build is declared or validated by origin metadata/liftOver/spot checks."
    lines = [
        "# External biocontrol data integration",
        "",
        "This stage prepares the coordinate and input layer for real external biological controls.",
        "",
        f"Declared genome build: {declared}",
        "",
        "Genome build audit:",
    ]
    for row in audit.itertuples(index=False):
        lines.append(f"- {row.audit_item}: {row.status}; {row.detail}")
    lines.extend(["", "External input status:"])
    for row in manifest.itertuples(index=False):
        status = "present" if row.exists else "missing"
        lines.append(f"- {row.input_name}: {status}; {row.expected_path}")
    lines.extend(
        [
            "",
            f"Current nearest-gene/TSS links generated: {n_links}",
            "",
            f"Interpretation boundary: {boundary}",
        ]
    )
    path.write_text("\n".join(lines) + "\n", encoding="utf-8")


def main() -> None:
    parser = argparse.ArgumentParser()
    parser.add_argument("--external-dir", default=str(EXTERNAL))
    parser.add_argument("--genome-build", default="")
    parser.add_argument("--gtf", default="")
    parser.add_argument("--gene-tss", default="")
    parser.add_argument("--top-n", type=int, default=0, help="Use 0 to audit all DMRs; positive values restrict downstream top-N outputs.")
    args = parser.parse_args()

    external = Path(args.external_dir)
    external.mkdir(parents=True, exist_ok=True)
    (external / "annotations").mkdir(parents=True, exist_ok=True)
    for sub in ["rna", "motif", "atac", "histone"]:
        (external / sub).mkdir(parents=True, exist_ok=True)

    genome_build_path = external / "genome_build.txt"
    if not genome_build_path.exists():
        genome_build_path.write_text("UNKNOWN\n# Set to hg19/GRCh37 or hg38/GRCh38 after audit.\n", encoding="ascii")
    genome_build_text = args.genome_build or genome_build_path.read_text(encoding="utf-8", errors="replace")

    dmr = read_tsv(DMR_RANKING)
    if dmr.empty:
        raise FileNotFoundError(f"Missing DMR ranking: {DMR_RANKING}")
    if args.top_n and args.top_n > 0:
        dmr = dmr.head(args.top_n).copy()
    else:
        dmr = dmr.copy()

    input_manifest = build_input_manifest(external)
    audit, chr_summary = coordinate_audit(dmr, genome_build_text)

    tss = pd.DataFrame()
    tss_path = Path(args.gene_tss) if args.gene_tss else external / "annotations" / "gene_tss.tsv"
    gtf_candidates = [
        Path(args.gtf) if args.gtf else None,
        external / "annotations" / "gencode.gtf",
        external / "annotations" / "gencode.gtf.gz",
        external / "annotations" / "gencode.v19.annotation.gtf.gz",
    ]
    gtf_path = next((p for p in gtf_candidates if p is not None and p.exists()), external / "annotations" / "gencode.gtf")
    if tss_path.exists():
        tss = read_tsv(tss_path)
    elif gtf_path.exists():
        tss = gtf_to_tss(gtf_path)
        tss.to_csv(OUT_TSS, sep="\t", index=False)
        tss.to_csv(external / "annotations" / "gene_tss.tsv", sep="\t", index=False)
    else:
        pd.DataFrame(columns=["chr", "tss", "gene_id", "gene_name", "gene_type", "strand"]).to_csv(OUT_TSS, sep="\t", index=False)

    gene_links = nearest_gene_links(dmr, tss)
    gene_links.to_csv(OUT_GENE_ANNOT, sep="\t", index=False)
    gene_links.to_csv(OUT_MODULE_LINKS, sep="\t", index=False)
    module_gene_link_summary(gene_links).to_csv(OUT_MODULE_LINK_SUMMARY, sep="\t", index=False)

    region_cols = [
        "module_id",
        "cluster_name",
        "chr",
        "start",
        "end",
        "promoter_2kb",
        "promoter_5kb",
        "gene_body",
        "intergenic",
        "CpG_island",
        "CpG_shore",
        "CpG_shelf",
        "repeat_class",
        "repeat_family",
        "residual_contribution",
        "module_basis_weight",
    ]
    if gene_links.empty:
        pd.DataFrame(columns=region_cols).to_csv(OUT_REGION_ANNOT, sep="\t", index=False)
    else:
        region = gene_links.copy()
        region["CpG_island"] = np.nan
        region["CpG_shore"] = np.nan
        region["CpG_shelf"] = np.nan
        region["repeat_class"] = ""
        region["repeat_family"] = ""
        region[[c for c in region_cols if c in region.columns]].to_csv(OUT_REGION_ANNOT, sep="\t", index=False)

    feature_template(PRIORITY_MODULES).to_csv(OUT_FEATURE_TEMPLATE, sep="\t", index=False)
    input_manifest.to_csv(OUT_INPUT_MANIFEST, sep="\t", index=False)
    audit.to_csv(OUT_AUDIT, sep="\t", index=False)
    chr_summary.to_csv(OUT_CHR_SUMMARY, sep="\t", index=False)

    write_doc(OUT_DOC, audit, input_manifest, len(gene_links))
    OUT_MANIFEST.write_text(
        json.dumps(
            {
                "external_dir": str(external),
                "genome_build_file": str(genome_build_path),
                "top_n": args.top_n,
                "gtf": str(gtf_path) if gtf_path.exists() else None,
                "gene_tss": str(tss_path) if tss_path.exists() else None,
                "outputs": [
                    str(OUT_INPUT_MANIFEST),
                    str(OUT_AUDIT),
                    str(OUT_CHR_SUMMARY),
                    str(OUT_GENE_ANNOT),
                    str(OUT_MODULE_LINKS),
                    str(OUT_MODULE_LINK_SUMMARY),
                    str(OUT_REGION_ANNOT),
                    str(OUT_FEATURE_TEMPLATE),
                    str(OUT_DOC),
                ],
            },
            indent=2,
        )
        + "\n",
        encoding="utf-8",
    )

    print(
        json.dumps(
            {
                "declared_build": audit.loc[audit["audit_item"] == "declared_genome_build", "detail"].iloc[0],
                "audit": audit.to_dict(orient="records"),
                "n_gene_links": int(len(gene_links)),
                "missing_required": input_manifest[(~input_manifest["exists"]) & input_manifest["priority"].str.contains("required", na=False)]["input_name"].tolist(),
            },
            indent=2,
        )
    )


if __name__ == "__main__":
    main()
```
