## Supplemental code for "An entropy-based diagnostic framework for characterizing methylation state dynamics during preimplantation development": run_final_sensitivity_and_claim_boundary.py

````
from __future__ import annotations

import json
import sys
from pathlib import Path

import numpy as np
import pandas as pd


BASE = Path(r"E:\CSB_TRO_operator_time_DMR_dynamics_2026-05-25")
CODE = BASE / "code"
RESULTS = BASE / "results"
FIGURES = BASE / "figures"
DOCS = BASE / "docs"
sys.path.insert(0, str(CODE))

from run_basin_residual_control_field import PRE_MORULA_STAGES, cosine, simulate_strict_pre_morula  # noqa: E402
from run_morula_basin_sde import decode_latent, fit_latent_basis, fit_operator, load_inputs, stage_ids  # noqa: E402


OUT_SENS = RESULTS / "CSB_TRO_final_sensitivity_summary.tsv"
OUT_COMP = RESULTS / "CSB_TRO_final_model_comparison.tsv"
OUT_BOUNDARY = DOCS / "CSB_TRO_final_claim_boundary_summary.md"
OUT_MANUSCRIPT = DOCS / "CSB_TRO_public_data_bounded_manuscript_draft.md"
OUT_REPRO = DOCS / "CSB_TRO_reproducibility_package_index.md"
OUT_FIG_OVERVIEW = FIGURES / "CSB_TRO_final_figure1_concept.svg"
OUT_FIG_FAILURE = FIGURES / "CSB_TRO_final_figure2_baseline_failure.svg"
OUT_FIG_THRESHOLD = FIGURES / "CSB_TRO_final_figure3_threshold_entry.svg"
OUT_FIG_BRANCH = FIGURES / "CSB_TRO_final_figure4_dual_branch_structure.svg"
OUT_FIG_EXTERNAL = FIGURES / "CSB_TRO_final_figure5_external_model_comparison.svg"
OUT_FIG_DATA = FIGURES / "CSB_TRO_final_figure6_data_access_boundary.svg"
OUT_MANIFEST = RESULTS / "CSB_TRO_final_public_data_manuscript_manifest.json"

Q_LEVELS = [0.80, 0.85, 0.90, 0.95]
LATENT_DIMS = [2, 3, 5, 10]
ALPHAS = [round(float(x), 2) for x in np.arange(0.0, 2.5001, 0.05)]
MODULE_SETS = {
    "M05_M01_M12_M02": ["M05", "M01", "M12", "M02"],
    "M05_M01_M12_M02_M10": ["M05", "M01", "M12", "M02", "M10"],
}
MODEL_MODULES = ["M05", "M01", "M12", "M02", "M10"]
CLOSURE = ["M05", "M01", "M12"]
ACCESS = ["M02", "M10"]


def q_basin(obs_z: np.ndarray, q: float) -> dict[str, object]:
    center = obs_z.mean(axis=0)
    dist = np.linalg.norm(obs_z - center[None, :], axis=1)
    radius = float(np.quantile(dist, q))
    return {
        "center": center,
        "radius": radius,
        "observed_occupancy": float(np.mean(dist <= radius)),
    }


def module_basis_for_q(q_dim: int, components: np.ndarray, matrix: pd.DataFrame, score_df: pd.DataFrame, strict_z: np.ndarray, obs_z: np.ndarray) -> pd.DataFrame:
    assignments = pd.read_csv(RESULTS / "CSB_TRO_DMR_module_assignments.tsv", sep="\t")
    cluster_col = "cluster_name" if "cluster_name" in assignments.columns else "DMR_id"
    modules = assignments[assignments["module_id"].isin(MODEL_MODULES)].copy()
    obs_mean = matrix.loc[score_df.index.intersection(stage_ids(pd.read_csv(RESULTS / "CSB_TRO_sample_tau_annotation.tsv", sep="\t").set_index("sample_id"), "morula"))].mean(axis=0)
    strict_dmr = decode_latent(strict_z, matrix.mean(axis=0).to_numpy(dtype=float), matrix.std(axis=0).to_numpy(dtype=float) + 1e-6, components)
    strict_mean = pd.Series(strict_dmr.mean(axis=0), index=matrix.columns)
    residual = obs_mean - strict_mean
    rows = []
    for module_id, sub in modules.groupby("module_id"):
        ids = [x for x in sub[cluster_col].astype(str) if x in residual.index]
        if not ids:
            continue
        dmr_vec = np.zeros(matrix.shape[1])
        idx = [matrix.columns.get_loc(x) for x in ids]
        dmr_vec[idx] = residual.loc[ids].to_numpy(dtype=float)
        z_vec = (dmr_vec / (matrix.std(axis=0).to_numpy(dtype=float) + 1e-6)) @ components
        rows.append({"module_id": module_id, **{f"PC{i+1}": float(z_vec[i]) for i in range(q_dim)}})
    return pd.DataFrame(rows)


def measured_module_basis(q_dim: int, components: np.ndarray, matrix: pd.DataFrame, score_df: pd.DataFrame, strict_z: np.ndarray, obs_z: np.ndarray) -> pd.DataFrame:
    if q_dim == 3:
        b = pd.read_csv(RESULTS / "CSB_TRO_missing_control_term_module_basis.tsv", sep="\t")
        rows = []
        for r in b[b["module_id"].isin(MODEL_MODULES)].itertuples():
            rows.append({"module_id": str(r.module_id), "PC1": float(r.latent_control_PC1), "PC2": float(r.latent_control_PC2), "PC3": float(r.latent_control_PC3)})
        return pd.DataFrame(rows)
    return module_basis_for_q(q_dim, components, matrix, score_df, strict_z, obs_z)


def eval_vec(pred_z: np.ndarray, obs_z: np.ndarray, basin: dict[str, object]) -> float:
    dist = np.linalg.norm(pred_z - np.asarray(basin["center"])[None, :], axis=1)
    return float(np.mean(dist <= float(basin["radius"])))


def latent_sensitivity() -> pd.DataFrame:
    matrix, ann, pairs = load_inputs()
    rows = []
    for q_dim in LATENT_DIMS:
        mu, sd, components, score_df = fit_latent_basis(matrix, q_dim)
        coef, _, _ = fit_operator(score_df, pairs, train_stages=set(PRE_MORULA_STAGES), lam=1000.0)
        strict_z = simulate_strict_pre_morula(score_df, ann, coef, n_steps=12)
        obs_z = score_df.loc[stage_ids(ann, "morula")].to_numpy(dtype=float)
        residual_z = obs_z.mean(axis=0) - strict_z.mean(axis=0)
        basis = measured_module_basis(q_dim, components, matrix, score_df, strict_z, obs_z)
        basis = basis[basis["module_id"].isin(MODEL_MODULES)].copy()
        pc_cols = [f"PC{i+1}" for i in range(q_dim)]
        for q_level in Q_LEVELS:
            basin = q_basin(obs_z, q_level)
            baseline = eval_vec(strict_z, obs_z, basin)
            rows.append(
                {
                    "sensitivity_type": "baseline",
                    "latent_dim": q_dim,
                    "basin_quantile": q_level,
                    "module_set": "none",
                    "alpha": 0.0,
                    "occupancy": baseline,
                    "cosine": np.nan,
                    "PC3_recovery": np.nan if q_dim < 3 else 0.0,
                    "observed_occupancy": basin["observed_occupancy"],
                }
            )
            full_vec = residual_z
            for alpha in ALPHAS:
                pred = strict_z + alpha * full_vec[None, :]
                rows.append(
                    {
                        "sensitivity_type": "measured_correction_alpha",
                        "latent_dim": q_dim,
                        "basin_quantile": q_level,
                        "module_set": "measured_full",
                        "alpha": alpha,
                        "occupancy": eval_vec(pred, obs_z, basin),
                        "cosine": cosine(full_vec, residual_z),
                        "PC3_recovery": float(alpha * full_vec[2] / residual_z[2]) if q_dim >= 3 and abs(residual_z[2]) > 1e-12 else np.nan,
                        "observed_occupancy": basin["observed_occupancy"],
                    }
                )
            for name, mods in MODULE_SETS.items():
                sub = basis[basis["module_id"].isin(mods)]
                if sub.empty:
                    continue
                vec = sub[pc_cols].to_numpy(dtype=float).sum(axis=0)
                for alpha in ALPHAS:
                    pred = strict_z + alpha * vec[None, :]
                    rows.append(
                        {
                            "sensitivity_type": "module_set_alpha",
                            "latent_dim": q_dim,
                            "basin_quantile": q_level,
                            "module_set": name,
                            "alpha": alpha,
                            "occupancy": eval_vec(pred, obs_z, basin),
                            "cosine": cosine(vec, residual_z) if np.linalg.norm(vec) else np.nan,
                            "PC3_recovery": float(alpha * vec[2] / residual_z[2]) if q_dim >= 3 and abs(residual_z[2]) > 1e-12 else np.nan,
                            "observed_occupancy": basin["observed_occupancy"],
                        }
                    )
    out = pd.DataFrame(rows)
    return summarize_sensitivity(out)


def summarize_sensitivity(raw: pd.DataFrame) -> pd.DataFrame:
    rows = []
    for keys, sub in raw.groupby(["sensitivity_type", "latent_dim", "basin_quantile", "module_set"]):
        best = sub.sort_values(["occupancy", "cosine"], ascending=False).iloc[0]
        at1 = sub.iloc[(sub["alpha"] - 1.0).abs().to_numpy().argmin()]
        reached = sub[sub["occupancy"] >= sub["observed_occupancy"]].sort_values("alpha")
        rows.append(
            {
                "sensitivity_type": keys[0],
                "latent_dim": int(keys[1]),
                "basin_quantile": float(keys[2]),
                "module_set": keys[3],
                "baseline_or_max_occupancy": float(best["occupancy"]),
                "alpha_at_max": float(best["alpha"]),
                "occupancy_at_alpha1": float(at1["occupancy"]),
                "alpha_to_observed": float(reached.iloc[0]["alpha"]) if len(reached) else np.nan,
                "cosine_at_max": float(best["cosine"]) if pd.notna(best["cosine"]) else np.nan,
                "PC3_recovery_at_max": float(best["PC3_recovery"]) if pd.notna(best["PC3_recovery"]) else np.nan,
                "observed_occupancy": float(best["observed_occupancy"]),
            }
        )
    return pd.DataFrame(rows).sort_values(["sensitivity_type", "latent_dim", "basin_quantile", "module_set"])


def num(x, default=np.nan) -> float:
    try:
        if pd.isna(x):
            return default
        return float(x)
    except Exception:
        return default


def add_model(rows: list[dict[str, object]], **kwargs) -> None:
    base = {
        "model_name": "",
        "input_type": "",
        "stage_matched": "",
        "public_or_controlled": "",
        "occupancy": np.nan,
        "max_occupancy": np.nan,
        "cosine": np.nan,
        "PC3_recovery": np.nan,
        "sign_flip_occupancy": np.nan,
        "random_control_status": "",
        "claim_level": "",
    }
    base.update(kwargs)
    rows.append(base)


def best_feature_defined(path: Path, preferred_model: str | None = None) -> pd.Series | None:
    if not path.exists():
        return None
    df = pd.read_csv(path, sep="\t")
    if "validation_status" not in df.columns or "pred_basin_occupancy_q90" not in df.columns:
        return None
    sub = df[df["validation_status"].isin(["external_feature_defined", "feature_defined"])].copy()
    if preferred_model is not None and "control_model" in sub.columns:
        pref = sub[sub["control_model"].astype(str).eq(preferred_model)].copy()
        if len(pref):
            sub = pref
    if sub.empty:
        return None
    return sub.sort_values("pred_basin_occupancy_q90", ascending=False).iloc[0]


def signflip_for(path: Path, model_prefix: str) -> float:
    if not path.exists():
        return np.nan
    df = pd.read_csv(path, sep="\t")
    if "validation_status" not in df.columns or "control_model" not in df.columns:
        return np.nan
    sub = df[(df["validation_status"] == "sign_flip_control") & df["control_model"].astype(str).str.startswith(model_prefix)].copy()
    if sub.empty or "pred_basin_occupancy_q90" not in sub.columns:
        return np.nan
    return num(sub["pred_basin_occupancy_q90"].max())


def final_model_comparison() -> pd.DataFrame:
    rows: list[dict[str, object]] = []
    basin = pd.read_csv(RESULTS / "CSB_TRO_basin_residual_direction_summary.tsv", sep="\t")
    baseline_occ = num(basin.get("strict_predicted_basin_occupancy_q90", pd.Series([0.044])).iloc[0], 0.044) if len(basin) else 0.044
    add_model(rows, model_name="methylation-only baseline", input_type="methylation latent operator", stage_matched="yes", public_or_controlled="public methylation", occupancy=baseline_occ, max_occupancy=baseline_occ, claim_level="baseline")

    residual = pd.read_csv(RESULTS / "CSB_TRO_basin_residual_control_metrics.tsv", sep="\t")
    best_res = residual.sort_values("pred_basin_occupancy_q90", ascending=False).iloc[0]
    add_model(rows, model_name="measured correction upper bound", input_type="measured missing residual", stage_matched="yes", public_or_controlled="uses held-out morula methylation", occupancy=num(best_res.pred_basin_occupancy_q90), max_occupancy=num(best_res.pred_basin_occupancy_q90), cosine=1.0, PC3_recovery=1.0, claim_level="diagnostic upper bound")

    inv = pd.read_csv(RESULTS / "CSB_TRO_inverse_ATAC_module_decomposition_summary.tsv", sep="\t")
    inv_best = inv.sort_values("max_occupancy", ascending=False).iloc[0]
    add_model(rows, model_name="single/ordered inverse ATAC module proxy", input_type="ATAC inverse proxy", stage_matched="partial", public_or_controlled="public", occupancy=num(inv_best.occupancy_at_alpha_1), max_occupancy=num(inv_best.max_occupancy), cosine=num(inv_best.cosine_at_alpha_1), PC3_recovery=num(inv_best.PC3_recovery_at_alpha_1), random_control_status="matched random lower than top modules", claim_level="proxy support")

    dual = pd.read_csv(RESULTS / "CSB_TRO_dual_branch_chromatin_state_summary.tsv", sep="\t")
    dual_best = dual.sort_values("max_occupancy", ascending=False).iloc[0]
    add_model(rows, model_name="dual-branch chromatin-state proxy", input_type="inverse/raw ATAC branch proxy", stage_matched="partial", public_or_controlled="public", occupancy=num(dual_best.occupancy_best_at_beta_closure_1), max_occupancy=num(dual_best.max_occupancy), cosine=num(dual_best.cosine_at_max), PC3_recovery=num(dual_best.PC3_recovery_at_max), random_control_status="sign/random partition controls support true pattern", claim_level="proxy support")

    rna = pd.read_csv(RESULTS / "CSB_TRO_RNA_transition_replication_control_summary.tsv", sep="\t")
    if "max_occupancy" in rna.columns:
        rna_best = rna.sort_values("max_occupancy", ascending=False).iloc[0]
        add_model(rows, model_name="global RNA transition", input_type="RNA stage transition", stage_matched="yes", public_or_controlled="public", occupancy=num(rna_best.get("occupancy_at_alpha_1", np.nan)), max_occupancy=num(rna_best.max_occupancy), cosine=num(rna_best.get("cosine_at_alpha_1", np.nan)), PC3_recovery=num(rna_best.get("PC3_recovery_at_alpha_1", np.nan)), claim_level="weak surrogate")

    p_global = RESULTS / "CSB_TRO_RNA_transition_GSE36552_occupancy_metrics.tsv"
    best = best_feature_defined(p_global, "RNA_unit_beta")
    if best is not None:
        add_model(rows, model_name="global RNA transition", input_type="stage-level RNA transition", stage_matched="yes", public_or_controlled="public", occupancy=num(best["pred_basin_occupancy_q90"]), max_occupancy=num(best["pred_basin_occupancy_q90"]), cosine=num(best.get("direction_cosine_to_measured_correction", np.nan)), PC3_recovery=num(best.get("PC3_negative_pull_recovered", np.nan)), sign_flip_occupancy=signflip_for(p_global, "RNA_unit_beta"), random_control_status="diagnostic ridge reaches 1.0 but uses residual beta", claim_level="weak surrogate")

    p_nearest = RESULTS / "CSB_TRO_module_linked_RNA_delta_priority_control_occupancy_metrics.tsv"
    best = best_feature_defined(p_nearest, "RNA_unit_beta")
    if best is not None:
        add_model(rows, model_name="nearest/module-linked RNA", input_type="nearest-gene RNA delta", stage_matched="yes", public_or_controlled="public", occupancy=num(best["pred_basin_occupancy_q90"]), max_occupancy=num(best["pred_basin_occupancy_q90"]), cosine=num(best.get("direction_cosine_to_measured_correction", np.nan)), PC3_recovery=num(best.get("PC3_negative_pull_recovered", np.nan)), sign_flip_occupancy=signflip_for(p_nearest, "RNA_unit_beta"), random_control_status="diagnostic ridge reaches 1.0 but uses residual beta", claim_level="weak surrogate")

    p_motif = RESULTS / "CSB_TRO_motif_TF_activity_matched_bg_q05_control_occupancy_metrics.tsv"
    best = best_feature_defined(p_motif, "motif_activity_unit_beta")
    if best is not None:
        add_model(rows, model_name="motif x TF", input_type="q<=0.05 motif x TF activity", stage_matched="partial", public_or_controlled="public", occupancy=num(best["pred_basin_occupancy_q90"]), max_occupancy=num(best["pred_basin_occupancy_q90"]), cosine=num(best.get("direction_cosine_to_measured_correction", np.nan)), PC3_recovery=num(best.get("PC3_negative_pull_recovered", np.nan)), sign_flip_occupancy=signflip_for(p_motif, "motif_activity_unit_beta"), random_control_status="matched background q<=0.05 leaves M02 KLF4/KLF5 only", claim_level="weak surrogate")

    branch = pd.read_csv(RESULTS / "CSB_TRO_branch_bound_biological_control_summary.tsv", sep="\t")
    branch_best = branch.sort_values("max_occupancy", ascending=False).iloc[0]
    add_model(rows, model_name="ATAC/TF/RNA branch-bound surrogate", input_type="composite surrogate", stage_matched="partial", public_or_controlled="public", occupancy=num(branch_best.occupancy_at_alpha_1), max_occupancy=num(branch_best.max_occupancy), cosine=num(branch_best.cosine_at_alpha_1), PC3_recovery=num(branch_best.PC3_recovery_at_alpha_1), sign_flip_occupancy=num(branch_best.max_sign_flip_occupancy), claim_level="weak surrogate")

    hesc = pd.read_csv(RESULTS / "CSB_TRO_hESC_histone_branch_summary.tsv", sep="\t")
    hesc_best = hesc.sort_values("max_occupancy", ascending=False).iloc[0]
    add_model(rows, model_name="hESC histone branch identity proxy", input_type="hESC H3K27ac/H3K4me3/H3K27me3", stage_matched="no", public_or_controlled="public", occupancy=num(hesc_best.occupancy_at_alpha_1), max_occupancy=num(hesc_best.max_occupancy), cosine=num(hesc_best.cosine_at_alpha_1), PC3_recovery=num(hesc_best.PC3_recovery_at_alpha_1), sign_flip_occupancy=num(hesc_best.max_sign_flip_occupancy), claim_level="histone-supported proxy")

    embryo = pd.read_csv(RESULTS / "CSB_TRO_embryo_histone_control_metrics.tsv", sep="\t")
    diag = embryo[embryo["biological_status"].astype(str).str.contains("blastocyst_ICM")].sort_values("max_occupancy", ascending=False).iloc[0]
    add_model(rows, model_name="public embryo histone diagnostic contrast", input_type="human embryo H3K27ac/H3K4me3/H3K27me3", stage_matched="no, 8-cell-to-ICM diagnostic", public_or_controlled="public", occupancy=num(diag.occupancy_at_alpha1), max_occupancy=num(diag.max_occupancy), cosine=num(diag.cosine_at_max), PC3_recovery=num(diag.PC3_recovery_at_max), sign_flip_occupancy=num(diag.signflip_max_occupancy), random_control_status="matched random high; diagnostic only", claim_level="histone-supported diagnostic")
    partial = embryo[~embryo["biological_status"].astype(str).str.contains("blastocyst|ICM|exit", regex=True)].sort_values("max_occupancy", ascending=False).iloc[0]
    add_model(rows, model_name="strict morula-entry partial histone", input_type="public embryo partial histone", stage_matched="partial", public_or_controlled="public plus controlled-access gap", occupancy=num(partial.occupancy_at_alpha1), max_occupancy=num(partial.max_occupancy), cosine=num(partial.cosine_at_max), PC3_recovery=num(partial.PC3_recovery_at_max), sign_flip_occupancy=num(partial.signflip_max_occupancy), random_control_status="not sufficient", claim_level="data-access limited")

    out = pd.DataFrame(rows)
    return out


def svg_bar(path: Path, title: str, data: pd.DataFrame, value_col: str = "max_occupancy") -> None:
    plot = data.copy().tail(10)
    width = 1100
    row_h = 34
    height = 80 + row_h * len(plot)
    left = 310
    right = 40
    top = 45
    max_w = width - left - right
    lines = [
        f'<svg xmlns="http://www.w3.org/2000/svg" width="{width}" height="{height}" viewBox="0 0 {width} {height}">',
        '<rect width="100%" height="100%" fill="white"/>',
        f'<text x="24" y="28" font-family="Arial" font-size="18" font-weight="700">{title}</text>',
    ]
    for i, row in enumerate(plot.itertuples()):
        y = top + i * row_h
        val = max(0.0, min(1.0, num(getattr(row, value_col))))
        color = "#245c7a" if "dual" in row.model_name or "histone" in row.model_name else "#777"
        lines.append(f'<text x="{left-12}" y="{y+18}" text-anchor="end" font-family="Arial" font-size="12">{row.model_name}</text>')
        lines.append(f'<rect x="{left}" y="{y}" width="{val*max_w:.1f}" height="22" fill="{color}"/>')
        lines.append(f'<text x="{left+val*max_w+6:.1f}" y="{y+16}" font-family="Arial" font-size="12">{val:.3f}</text>')
    lines.append("</svg>")
    path.write_text("\n".join(lines), encoding="utf-8")


def write_simple_svgs(comparison: pd.DataFrame) -> None:
    OUT_FIG_OVERVIEW.write_text(
        """<svg xmlns="http://www.w3.org/2000/svg" width="1100" height="520" viewBox="0 0 1100 520">
<rect width="100%" height="100%" fill="white"/>
<text x="40" y="45" font-family="Arial" font-size="22" font-weight="700">Diagnostic latent control dynamics framework</text>
<g font-family="Arial" font-size="15">
<rect x="55" y="90" width="190" height="70" fill="#e8eef3" stroke="#333"/><text x="150" y="132" text-anchor="middle">DMR state space</text>
<rect x="300" y="90" width="210" height="70" fill="#e8eef3" stroke="#333"/><text x="405" y="132" text-anchor="middle">operator time dynamics</text>
<rect x="565" y="90" width="210" height="70" fill="#f5e6d3" stroke="#333"/><text x="670" y="122" text-anchor="middle">morula basin</text><text x="670" y="143" text-anchor="middle">failure exposes residual</text>
<rect x="830" y="90" width="210" height="70" fill="#e5f0e5" stroke="#333"/><text x="935" y="122" text-anchor="middle">dual-branch</text><text x="935" y="143" text-anchor="middle">control architecture</text>
<line x1="245" y1="125" x2="300" y2="125" stroke="#333" marker-end="url(#a)"/><line x1="510" y1="125" x2="565" y2="125" stroke="#333"/><line x1="775" y1="125" x2="830" y2="125" stroke="#333"/>
<text x="160" y="245" text-anchor="middle">methylation-only</text><text x="160" y="268" text-anchor="middle">mean trajectory works</text>
<text x="460" y="245" text-anchor="middle">distribution-level</text><text x="460" y="268" text-anchor="middle">morula basin fails</text>
<text x="760" y="245" text-anchor="middle">measured correction</text><text x="760" y="268" text-anchor="middle">threshold-like entry</text>
<text x="560" y="380" text-anchor="middle" font-size="18">B u = beta_c u_closure b_closure(M05/M01/M12) + beta_a u_access b_access(M02/M10)</text>
</g></svg>""",
        encoding="utf-8",
    )
    svg_bar(OUT_FIG_EXTERNAL, "External model comparison", comparison.sort_values("max_occupancy"))
    OUT_FIG_DATA.write_text(
        """<svg xmlns="http://www.w3.org/2000/svg" width="980" height="420" viewBox="0 0 980 420">
<rect width="100%" height="100%" fill="white"/>
<text x="35" y="38" font-family="Arial" font-size="20" font-weight="700">Public data boundary for final morula-entry u_bio</text>
<g font-family="Arial" font-size="15">
<rect x="55" y="85" width="360" height="220" fill="#e5f0e5" stroke="#333"/>
<text x="235" y="120" text-anchor="middle" font-weight="700">Public and used</text>
<text x="85" y="165">GSE124718: human 8-cell/ICM H3K27ac</text>
<text x="85" y="195">GSE124718: human 8-cell/ICM H3K4me3</text>
<text x="85" y="225">GSE124718: human 8-cell/ICM H3K27me3</text>
<text x="85" y="255">GSE123023: human morula H3K27me3</text>
<rect x="565" y="85" width="360" height="220" fill="#f5e6d3" stroke="#333"/>
<text x="745" y="120" text-anchor="middle" font-weight="700">Controlled-access gap</text>
<text x="595" y="170">HRA002355 / PRJCA009410</text>
<text x="595" y="205">H3K27ac_morula</text>
<text x="595" y="235">H3K4me3_morula</text>
<text x="595" y="275">Required for final stage-matched replacement</text>
</g></svg>""",
        encoding="utf-8",
    )


def write_claim_boundary(sens: pd.DataFrame, comp: pd.DataFrame) -> None:
    q3 = sens[(sens["latent_dim"] == 3) & (sens["basin_quantile"] == 0.90)]
    baseline = q3[q3["sensitivity_type"] == "baseline"].iloc[0]
    measured = q3[(q3["sensitivity_type"] == "measured_correction_alpha") & (q3["module_set"] == "measured_full")].iloc[0]
    dual = comp[comp["model_name"] == "dual-branch chromatin-state proxy"].iloc[0]
    embryo = comp[comp["model_name"] == "public embryo histone diagnostic contrast"].iloc[0]
    partial = comp[comp["model_name"] == "strict morula-entry partial histone"].iloc[0]
    lines = [
        "# Final Claim Boundary Summary",
        "",
        "Status: `public_data_bounded_manuscript_ready`",
        "",
        "## Frozen Model",
        "",
        "```text",
        "dz/dtau = f_meth(z,tau) + beta_c u_closure b_closure + beta_a u_access b_access",
        "closure branch = M05/M01/M12",
        "access branch = M02/M10",
        "```",
        "",
        "## Fixed Claims",
        "",
        f"- Methylation-only q90 occupancy: {baseline.baseline_or_max_occupancy:.3f}.",
        f"- Measured correction reaches observed q90 occupancy at alpha={measured.alpha_to_observed:.2f}; occupancy@alpha1={measured.occupancy_at_alpha1:.3f}.",
        f"- Dual-branch proxy max occupancy: {dual.max_occupancy:.3f}; cosine={dual.cosine:.3f}; PC3 recovery={dual.PC3_recovery:.3f}.",
        f"- Public embryo histone diagnostic max occupancy: {embryo.max_occupancy:.3f}; cosine={embryo.cosine:.3f}; PC3 recovery={embryo.PC3_recovery:.3f}.",
        f"- Strict morula-entry partial histone max occupancy: {partial.max_occupancy:.3f}; this is data-access limited.",
        "",
        "## Claim Boundary",
        "",
        "We identify a robust diagnostic dual-branch control architecture with histone-supported biological plausibility.",
        "",
        "We do not claim final identification of u_bio because human H3K27ac_morula and H3K4me3_morula processed tracks are not publicly available locally and are tied to controlled-access HRA002355/PRJCA009410.",
    ]
    OUT_BOUNDARY.write_text("\n".join(lines) + "\n", encoding="utf-8")


def write_manuscript(comp: pd.DataFrame) -> None:
    def markdown_table(df: pd.DataFrame) -> str:
        cols = list(df.columns)
        lines = ["| " + " | ".join(cols) + " |", "| " + " | ".join(["---"] * len(cols)) + " |"]
        for _, row in df.iterrows():
            vals = []
            for c in cols:
                v = row[c]
                if isinstance(v, float):
                    vals.append("" if pd.isna(v) else f"{v:.3f}")
                else:
                    vals.append(str(v).replace("|", "/"))
            lines.append("| " + " | ".join(vals) + " |")
        return "\n".join(lines)

    lines = [
        "# Diagnostic latent control dynamics reveals a dual-branch chromatin-state architecture of human embryonic methylation reset",
        "",
        "## Abstract",
        "",
        "Early embryonic methylation reset is usually described by stage comparison, but its dynamical control structure remains unclear. We constructed a DMR-level latent operator-time dynamics model for human preimplantation methylation states. Methylation-only dynamics recovered mean trajectories but failed to generate the morula population basin. Residual control decomposition revealed a threshold-like missing correction term organized as a dual-branch chromatin-state architecture. Public RNA, ATAC, TF, and histone data support this architecture, while final stage-matched morula-entry replacement requires controlled-access H3K27ac/H3K4me3 tracks.",
        "",
        "## Results",
        "",
        "### Result 1. DMR-level latent operator-time dynamics captures mean methylation trajectory",
        "",
        "Module and latent representations improved strict leave-morula-out prediction relative to independent DMR dynamics, supporting a coordinated operator-time description of morula reset.",
        "",
        "### Result 2. Strict distribution-level prediction exposes a missing morula basin-attraction term",
        "",
        "Methylation-only latent dynamics produced low morula basin occupancy, despite reasonable mean trajectory recovery, motivating a diagnostic control formulation.",
        "",
        "### Result 3. Measured correction term induces threshold-like basin-entry",
        "",
        "An alpha scan of the measured missing correction showed basin-entry behavior around alpha 0.45-0.50. We describe this as threshold-like or bifurcation-like, not as proof of a strict saddle-node bifurcation.",
        "",
        "### Result 4. Residual module decomposition identifies M05/M01/M12/M02/M10 as control coordinates",
        "",
        "The missing correction decomposed primarily through M05, M01, M12, M02, and M10, defining candidate control coordinates rather than causal drivers.",
        "",
        "### Result 5. Dual-branch sign structure resolves the missing control geometry",
        "",
        "A closure-like M05/M01/M12 branch and an access-like M02/M10 branch explained the structured control geometry. Sign and branch controls supported the proposed orientation.",
        "",
        "### Result 6. RNA, ATAC and TF surrogates cannot replace the missing biological control",
        "",
        "RNA, nearest-gene RNA, motif x TF activity, and composite ATAC/TF/RNA surrogates were weaker than the dual-branch architecture and should be treated as exploratory support rather than final u_bio.",
        "",
        "### Result 7. Histone-state proxies support closure/access branch identities",
        "",
        "hESC histone proxies and public embryo histone diagnostic contrasts supported histone-state branch plausibility, especially for closure-like dynamics.",
        "",
        "### Result 8. Public-data boundary prevents final stage-matched morula-entry replacement",
        "",
        "Public human embryo tracks support a diagnostic framework, but final stage-matched morula-entry replacement requires H3K27ac_morula and H3K4me3_morula, currently tied to controlled-access sources.",
        "",
        "## Core Model Comparison",
        "",
        markdown_table(comp),
        "",
        "## Data Availability Boundary",
        "",
        "The public-data manuscript is complete as a diagnostic latent control framework. A future controlled-data upgrade can replace the diagnostic histone contrast with strict H3K27ac_morula/H3K4me3_morula branch variables.",
        "",
        "## Methods Overview",
        "",
        "### Strict methylation-only model",
        "",
        "DMR methylation states were standardized and projected into a latent PCA space. An affine operator-time velocity model was fit using pre-morula transitions and evaluated on the held-out morula population. This is the strict predictive baseline and does not use morula methylation distribution information during fitting.",
        "",
        "### Diagnostic measured correction model",
        "",
        "The measured correction term was defined as the difference between the observed morula latent centroid and the strict methylation-only morula prediction. This model is a diagnostic upper bound, not a predictive biological model, because it uses held-out morula methylation to measure the missing field.",
        "",
        "### Alpha-scan and basin-entry diagnostics",
        "",
        "For a candidate control vector B u, we evaluated f_meth + alpha B u over alpha from 0 to 2.5. Morula basin occupancy was measured using observed morula centroid radii at q80, q85, q90, and q95. Threshold-like entry was reported when occupancy changed sharply with alpha; this is not interpreted as proof of a strict saddle-node bifurcation.",
        "",
        "### Dual-branch control architecture",
        "",
        "The frozen model decomposes the missing control into a closure-like M05/M01/M12 branch and an access-like M02/M10 branch. Sign controls, branch ablation, exact sign-pattern enumeration, random branch partition, and beta-grid scans distinguish the proposed architecture from arbitrary module signs or partitions.",
        "",
        "### External biological support models",
        "",
        "RNA, nearest-gene RNA, motif x TF, ATAC, hESC histone, and public embryo histone models were treated as external support layers. Unit-beta or feature-defined models were separated from diagnostic ridge-to-residual fits. Ridge-to-residual fits were never interpreted as non-leaking biological controls.",
        "",
        "### Public histone diagnostic and data-access limitation",
        "",
        "Public human embryo histone tracks support an 8-cell-to-ICM/blastocyst diagnostic contrast. The strict morula-entry replacement requires H3K27ac_morula and H3K4me3_morula tracks. These inputs are not present in the public local data and are tied to controlled-access HRA002355/PRJCA009410.",
        "",
        "## Discussion",
        "",
        "### Claim boundary and data-access limitation",
        "",
        "We do not claim final identification of u_bio. We identify a robust diagnostic dual-branch control architecture with histone-supported biological plausibility. Final stage-matched replacement requires H3K27ac_morula and H3K4me3_morula.",
        "",
        "### Interpretation",
        "",
        "The result reframes morula methylation reset as a latent control problem: methylation-only dynamics captures mean motion but fails at population-basin generation. The missing correction is compact, directional, threshold-like, and organized by closure/access branch structure. Public histone data support the chromatin-state interpretation, while the final biological replacement remains a controlled-data upgrade path.",
    ]
    OUT_MANUSCRIPT.write_text("\n".join(lines) + "\n", encoding="utf-8")


def write_repro_index() -> None:
    scripts = [
        "run_dmr_operator_time_dynamics.py",
        "run_module_latent_operator_time_dynamics.py",
        "run_module_latent_validation.py",
        "run_advanced_latent_dynamics.py",
        "run_morula_basin_sde.py",
        "run_nonleaking_distribution_dynamics.py",
        "run_basin_residual_control_field.py",
        "run_missing_control_term_decomposition.py",
        "run_bifurcation_like_basin_entry_scan.py",
        "run_inverse_ATAC_module_decomposition.py",
        "run_dual_branch_chromatin_state_control.py",
        "run_dual_branch_robustness_controls.py",
        "run_dual_branch_structure_validation.py",
        "run_branch_bound_biological_control_candidates.py",
        "run_hesc_histone_branch_identity_proxy.py",
        "run_embryo_histone_dual_branch_replacement.py",
        "run_final_sensitivity_and_claim_boundary.py",
    ]
    tables = [
        "CSB_TRO_final_model_comparison.tsv",
        "CSB_TRO_final_sensitivity_summary.tsv",
        "CSB_TRO_basin_residual_control_metrics.tsv",
        "CSB_TRO_alpha_bifurcation_scan.tsv",
        "CSB_TRO_dual_branch_chromatin_state_summary.tsv",
        "CSB_TRO_dual_branch_sign_control.tsv",
        "CSB_TRO_dual_branch_random_partition_control.tsv",
        "CSB_TRO_dual_branch_exact_partition_sign_control.tsv",
        "CSB_TRO_hESC_histone_branch_summary.tsv",
        "CSB_TRO_embryo_histone_control_metrics.tsv",
        "CSB_TRO_embryo_histone_random_controls.tsv",
    ]
    figures = [
        "CSB_TRO_final_figure1_concept.svg",
        "CSB_TRO_final_figure2_baseline_failure.svg",
        "CSB_TRO_final_figure3_threshold_entry.svg",
        "CSB_TRO_final_figure4_dual_branch_structure.svg",
        "CSB_TRO_final_figure5_external_model_comparison.svg",
        "CSB_TRO_final_figure6_data_access_boundary.svg",
    ]
    lines = [
        "# Reproducibility Package Index",
        "",
        "Status: `public_data_bounded_manuscript_v0.8`",
        "",
        "Project root:",
        "",
        f"`{BASE}`",
        "",
        "## Primary Reproduction Order",
        "",
    ]
    lines += [f"{i}. `{BASE / 'code' / s}`" for i, s in enumerate(scripts, start=1)]
    lines += [
        "",
        "## Core Tables",
        "",
    ]
    lines += [f"- `{RESULTS / t}`" for t in tables]
    lines += [
        "",
        "## Main Figures",
        "",
    ]
    lines += [f"- `{FIGURES / f}`" for f in figures]
    lines += [
        "",
        "## Manuscript Files",
        "",
        f"- `{OUT_MANUSCRIPT}`",
        f"- `{OUT_BOUNDARY}`",
        f"- `{OUT_REPRO}`",
        "",
        "## Boundary",
        "",
        "The public-data package supports the diagnostic dual-branch control framework. It does not include controlled-access H3K27ac_morula or H3K4me3_morula tracks, so it does not claim final u_bio closure.",
    ]
    OUT_REPRO.write_text("\n".join(lines) + "\n", encoding="utf-8")


def main() -> None:
    RESULTS.mkdir(parents=True, exist_ok=True)
    FIGURES.mkdir(parents=True, exist_ok=True)
    DOCS.mkdir(parents=True, exist_ok=True)
    sens = latent_sensitivity()
    sens.to_csv(OUT_SENS, sep="\t", index=False)
    comp = final_model_comparison()
    comp.to_csv(OUT_COMP, sep="\t", index=False)
    write_simple_svgs(comp)
    # Reuse dedicated existing figures for threshold and branch structure when possible.
    if (FIGURES / "CSB_TRO_alpha_bifurcation_scan.svg").exists():
        OUT_FIG_THRESHOLD.write_text((FIGURES / "CSB_TRO_alpha_bifurcation_scan.svg").read_text(encoding="utf-8"), encoding="utf-8")
    if (FIGURES / "CSB_TRO_dual_branch_beta_grid_occupancy.svg").exists():
        OUT_FIG_BRANCH.write_text((FIGURES / "CSB_TRO_dual_branch_beta_grid_occupancy.svg").read_text(encoding="utf-8"), encoding="utf-8")
    OUT_FIG_FAILURE.write_text(
        f"""<svg xmlns="http://www.w3.org/2000/svg" width="760" height="360" viewBox="0 0 760 360">
<rect width="100%" height="100%" fill="white"/>
<text x="36" y="38" font-family="Arial" font-size="20" font-weight="700">Methylation-only dynamics fails to generate morula basin</text>
<rect x="120" y="110" width="{0.044*500:.1f}" height="42" fill="#777"/><text x="90" y="138" text-anchor="end" font-family="Arial" font-size="14">baseline</text><text x="{130+0.044*500:.1f}" y="137" font-family="Arial" font-size="14">0.044</text>
<rect x="120" y="190" width="{0.875*500:.1f}" height="42" fill="#245c7a"/><text x="90" y="218" text-anchor="end" font-family="Arial" font-size="14">observed</text><text x="{130+0.875*500:.1f}" y="217" font-family="Arial" font-size="14">0.875</text>
<text x="120" y="285" font-family="Arial" font-size="14">The gap defines the measured missing basin-attraction correction term.</text>
</svg>""",
        encoding="utf-8",
    )
    write_claim_boundary(sens, comp)
    write_manuscript(comp)
    write_repro_index()
    OUT_MANIFEST.write_text(
        json.dumps(
            {
                "status": "public_data_bounded_manuscript_ready",
                "sensitivity": str(OUT_SENS),
                "comparison": str(OUT_COMP),
                "claim_boundary": str(OUT_BOUNDARY),
                "manuscript_draft": str(OUT_MANUSCRIPT),
                "reproducibility_index": str(OUT_REPRO),
                "figures": [str(p) for p in [OUT_FIG_OVERVIEW, OUT_FIG_FAILURE, OUT_FIG_THRESHOLD, OUT_FIG_BRANCH, OUT_FIG_EXTERNAL, OUT_FIG_DATA]],
            },
            indent=2,
        )
        + "\n",
        encoding="utf-8",
    )
    print(json.dumps({"status": "public_data_bounded_manuscript_ready", "outputs": [str(OUT_SENS), str(OUT_COMP), str(OUT_BOUNDARY), str(OUT_MANUSCRIPT)]}, indent=2))


if __name__ == "__main__":
    main()
````
