## Supplemental code for "An entropy-based diagnostic framework for characterizing methylation state dynamics during preimplantation development": run_gse36552_gene_stage_matrix.py

```
from __future__ import annotations

import json
import re
import zipfile
from pathlib import Path
from xml.etree import ElementTree as ET

import pandas as pd


BASE = Path(r"E:\CSB_TRO_operator_time_DMR_dynamics_2026-05-25")
MAIN = Path(r"C:\Users\18068\Desktop\CSB_TRO_Project_2026-05-24")
EXTERNAL = BASE / "external"
RESULTS = BASE / "results"
DOCS = BASE / "docs"

XLSX = EXTERNAL / "rna" / "GSE36552_Yan2013_SuppTable1_known_refseq_gene_RPKM.xlsx"
SAMPLE_META = MAIN / "input_tables" / "GSE36552_rna_sample_metadata.tsv"
GENE_LINKS = RESULTS / "CSB_TRO_residual_module_gene_links.tsv"

OUT_MATRIX = EXTERNAL / "rna" / "gene_stage_matrix.tsv"
OUT_LONG = RESULTS / "CSB_TRO_GSE36552_gene_stage_expression_long.tsv"
OUT_LINKS = RESULTS / "CSB_TRO_residual_module_gene_links_symbol.tsv"
OUT_MATCH_SUMMARY = RESULTS / "CSB_TRO_module_linked_RNA_gene_match_summary.tsv"
OUT_DOC = DOCS / "CSB_TRO_GSE36552_gene_stage_matrix.md"
OUT_MANIFEST = RESULTS / "CSB_TRO_GSE36552_gene_stage_matrix_manifest.json"

NS = {"m": "http://schemas.openxmlformats.org/spreadsheetml/2006/main"}


def col_index(cell_ref: str) -> int:
    letters = re.sub(r"[^A-Z]", "", cell_ref.upper())
    out = 0
    for ch in letters:
        out = out * 26 + (ord(ch) - ord("A") + 1)
    return out - 1


def shared_strings(z: zipfile.ZipFile) -> list[str]:
    root = ET.fromstring(z.read("xl/sharedStrings.xml"))
    vals = []
    for si in root.findall("m:si", NS):
        texts = []
        for t in si.iter("{http://schemas.openxmlformats.org/spreadsheetml/2006/main}t"):
            texts.append(t.text or "")
        vals.append("".join(texts))
    return vals


def parse_xlsx_rows(path: Path):
    with zipfile.ZipFile(path) as z:
        strings = shared_strings(z)
        for _, elem in ET.iterparse(z.open("xl/worksheets/sheet1.xml"), events=("end",)):
            if not elem.tag.endswith("}row"):
                continue
            row = {}
            for cell in elem:
                if not cell.tag.endswith("}c"):
                    continue
                ref = cell.attrib.get("r", "")
                val_node = cell.find("m:v", NS)
                val = "" if val_node is None else val_node.text
                if cell.attrib.get("t") == "s" and val not in ("", None):
                    val = strings[int(val)]
                row[col_index(ref)] = val
            yield int(elem.attrib.get("r", "0")), row
            elem.clear()


def main() -> None:
    EXTERNAL.joinpath("rna").mkdir(parents=True, exist_ok=True)
    RESULTS.mkdir(parents=True, exist_ok=True)
    DOCS.mkdir(parents=True, exist_ok=True)

    sample_meta = pd.read_csv(SAMPLE_META, sep="\t")
    sample_stage = dict(zip(sample_meta["sample_name"], sample_meta["stage"]))

    iterator = parse_xlsx_rows(XLSX)
    _, header_row = next(iterator)
    headers = [header_row.get(i, "") for i in range(max(header_row) + 1)]
    sample_cols = [(i, h, sample_stage[h]) for i, h in enumerate(headers) if h in sample_stage]

    records = []
    for _, row in iterator:
        gene = str(row.get(0, "")).strip()
        if not gene or gene.lower() in {"gene_id", "average"}:
            continue
        stage_vals: dict[str, list[float]] = {}
        for idx, sample, stage in sample_cols:
            raw = row.get(idx, "")
            try:
                value = float(raw)
            except (TypeError, ValueError):
                continue
            stage_vals.setdefault(stage, []).append(value)
        rec = {"gene_id": gene, "gene_name": gene}
        for stage, vals in stage_vals.items():
            rec[stage] = sum(vals) / len(vals) if vals else None
            rec[f"n_{stage}"] = len(vals)
        records.append(rec)

    matrix = pd.DataFrame(records)
    stage_order = ["oocyte", "zygote", "2-cell", "4-cell", "8-cell", "morula", "blastocyst"]
    cols = ["gene_id", "gene_name"] + [c for s in stage_order for c in (s, f"n_{s}") if c in matrix.columns]
    matrix = matrix[cols]
    matrix.to_csv(OUT_MATRIX, sep="\t", index=False)

    long_rows = []
    for stage in stage_order:
        if stage in matrix.columns:
            sub = matrix[["gene_id", stage]].copy()
            sub["stage"] = stage
            sub = sub.rename(columns={stage: "expression_mean_RPKM"})
            long_rows.append(sub[["gene_id", "stage", "expression_mean_RPKM"]])
    pd.concat(long_rows, ignore_index=True).to_csv(OUT_LONG, sep="\t", index=False)

    links = pd.read_csv(GENE_LINKS, sep="\t")
    symbol_links = links.copy()
    symbol_links["gene_id"] = symbol_links["nearest_gene"]
    symbol_links.to_csv(OUT_LINKS, sep="\t", index=False)

    matrix_genes = set(matrix["gene_id"].astype(str))
    match = symbol_links.copy()
    match["has_GSE36552_RNA"] = match["gene_id"].astype(str).isin(matrix_genes)
    summary = (
        match.groupby("module_id", as_index=False)
        .agg(
            n_linked_genes=("gene_id", "count"),
            n_RNA_matched=("has_GSE36552_RNA", "sum"),
            matched_genes=("gene_id", lambda x: ",".join([g for g in map(str, x) if g in matrix_genes][:30])),
        )
    )
    summary["match_fraction"] = summary["n_RNA_matched"] / summary["n_linked_genes"]
    summary.to_csv(OUT_MATCH_SUMMARY, sep="\t", index=False)

    lines = [
        "# GSE36552 gene-stage matrix",
        "",
        "Source: Yan et al. 2013 supplementary table 1 known RefSeq gene RPKM matrix.",
        "",
        f"Genes parsed: {len(matrix)}",
        f"Sample columns matched to local stage metadata: {len(sample_cols)}",
        "",
        "Outputs:",
        f"- {OUT_MATRIX}",
        f"- {OUT_LINKS}",
        f"- {OUT_MATCH_SUMMARY}",
    ]
    OUT_DOC.write_text("\n".join(lines) + "\n", encoding="utf-8")
    OUT_MANIFEST.write_text(
        json.dumps(
            {
                "source_xlsx": str(XLSX),
                "sample_metadata": str(SAMPLE_META),
                "n_genes": int(len(matrix)),
                "n_sample_columns_matched": int(len(sample_cols)),
                "outputs": [str(OUT_MATRIX), str(OUT_LONG), str(OUT_LINKS), str(OUT_MATCH_SUMMARY), str(OUT_DOC)],
            },
            indent=2,
        )
        + "\n",
        encoding="utf-8",
    )
    print(json.dumps({"n_genes": int(len(matrix)), "n_sample_columns_matched": int(len(sample_cols)), "output": str(OUT_MATRIX)}, indent=2))


if __name__ == "__main__":
    main()
```
