## Supplemental code for "An entropy-based diagnostic framework for characterizing methylation state dynamics during preimplantation development": run_hesc_histone_branch_identity_proxy.py

```
from __future__ import annotations

import gzip
import json
import math
import sys
from pathlib import Path

import numpy as np
import pandas as pd


BASE = Path(r"E:\CSB_TRO_operator_time_DMR_dynamics_2026-05-25")
MAIN = Path(r"C:\Users\18068\Desktop\CSB_TRO_Project_2026-05-24")
CODE = BASE / "code"
RESULTS = BASE / "results"
FIGURES = BASE / "figures"
DOCS = BASE / "docs"
HESC = BASE / "external" / "histone_proxy_hesc"
sys.path.insert(0, str(CODE))

from run_basin_residual_control_field import PRE_MORULA_STAGES, cosine, simulate_strict_pre_morula  # noqa: E402
from run_morula_basin_sde import (  # noqa: E402
    basin_definition,
    decode_latent,
    distribution_metrics,
    fit_latent_basis,
    fit_operator,
    load_inputs,
    stage_ids,
)


REGION_ANNOT = RESULTS / "CSB_TRO_residual_module_region_annotation.tsv"
MATCHED = MAIN / "input_tables" / "GSE81233_matched_non_age_window_regions_n100.tsv"
BASIS = RESULTS / "CSB_TRO_missing_control_term_module_basis.tsv"

OUT_OVERLAP = RESULTS / "CSB_TRO_hESC_histone_branch_overlap.tsv"
OUT_FEATURES = RESULTS / "CSB_TRO_hESC_histone_branch_control_features.tsv"
OUT_ALPHA = RESULTS / "CSB_TRO_hESC_histone_branch_alpha_scan.tsv"
OUT_SUMMARY = RESULTS / "CSB_TRO_hESC_histone_branch_summary.tsv"
OUT_MANIFEST = RESULTS / "CSB_TRO_hESC_histone_branch_manifest.json"
OUT_SVG = FIGURES / "CSB_TRO_hESC_histone_branch_summary.svg"
OUT_DOC = DOCS / "CSB_TRO_hESC_histone_branch_identity_proxy_summary.md"


MODULES = ["M05", "M01", "M12", "M02", "M10"]
CLOSURE = ["M05", "M01", "M12"]
ACCESS = ["M02", "M10"]

TRACKS = [
    ("hESC_naive_H3K27ac", "naive", "H3K27ac", HESC / "GSM1272764_WIBR3_n_H3K27ac_peaks.bed.gz"),
    ("hESC_naive_H3K27me3", "naive", "H3K27me3", HESC / "GSM1272763_WIBR3_n_H3K27me3_peaks.bed.gz"),
    ("hESC_naive_H3K4me3", "naive", "H3K4me3", HESC / "GSM1272762_WIBR3_n_H3K4me3_peaks.bed.gz"),
    ("hESC_primed_H3K27ac", "primed", "H3K27ac", HESC / "GSM1272770_WIBR3_p_H3K27ac_peaks.bed.gz"),
    ("hESC_primed_H3K27me3", "primed", "H3K27me3", HESC / "GSM1272769_WIBR3_p_H3K27me3_peaks.bed.gz"),
    ("hESC_primed_H3K4me3", "primed", "H3K4me3", HESC / "GSM1272768_WIBR3_p_H3K4me3_peaks.bed.gz"),
]


def read_bed(path: Path) -> pd.DataFrame:
    opener = gzip.open if path.suffix == ".gz" else open
    rows = []
    with opener(path, "rt", encoding="utf-8", errors="replace") as handle:
        for line in handle:
            if not line.strip() or line.startswith(("#", "track", "browser")):
                continue
            parts = line.rstrip("\n").split("\t")
            if len(parts) < 3:
                continue
            try:
                rows.append({"chr": parts[0], "start": int(float(parts[1])), "end": int(float(parts[2]))})
            except ValueError:
                continue
    return pd.DataFrame(rows)


def overlap_flags(regions: pd.DataFrame, peaks: pd.DataFrame) -> np.ndarray:
    flags = np.zeros(len(regions), dtype=bool)
    if peaks.empty or regions.empty:
        return flags
    by_chr = {c: sub[["start", "end"]].to_numpy(dtype=int) for c, sub in peaks.groupby("chr")}
    for i, row in enumerate(regions.itertuples()):
        arr = by_chr.get(str(row.chr))
        if arr is not None:
            flags[i] = bool(np.any((arr[:, 0] < int(row.end)) & (arr[:, 1] > int(row.start))))
    return flags


def log_comb(n: int, k: int) -> float:
    if k < 0 or k > n:
        return float("-inf")
    return math.lgamma(n + 1) - math.lgamma(k + 1) - math.lgamma(n - k + 1)


def fisher_greater(a: int, b: int, c: int, d: int) -> float:
    n = a + b + c + d
    row1 = a + b
    col1 = a + c
    lo = max(0, row1 - (n - col1))
    hi = min(row1, col1)
    denom = log_comb(n, row1)
    vals = [log_comb(col1, x) + log_comb(n - col1, row1 - x) - denom for x in range(max(a, lo), hi + 1)]
    if not vals:
        return 1.0
    m = max(vals)
    return float(min(1.0, math.exp(m) * sum(math.exp(v - m) for v in vals)))


def odds_ratio(a: int, b: int, c: int, d: int) -> float:
    return float(((a + 0.5) * (d + 0.5)) / ((b + 0.5) * (c + 0.5)))


def fdr_bh(p: pd.Series) -> pd.Series:
    x = pd.to_numeric(p, errors="coerce").fillna(1.0).to_numpy(dtype=float)
    order = np.argsort(x)
    q = np.empty_like(x)
    prev = 1.0
    n = len(x)
    for rank, idx in enumerate(order[::-1], start=1):
        true_rank = n - rank + 1
        prev = min(prev, x[idx] * n / true_rank)
        q[idx] = prev
    return pd.Series(np.minimum(q, 1.0), index=p.index)


def targets_bg() -> tuple[pd.DataFrame, pd.DataFrame]:
    annot = pd.read_csv(REGION_ANNOT, sep="\t")
    targets = annot[annot["module_id"].isin(MODULES)][["cluster_name", "module_id", "chr", "start", "end"]].copy()
    bg = pd.read_csv(MATCHED, sep="\t")
    bg = bg[(bg["region_type"] == "matched_non_age_window") & (bg["matched_age_cluster"].isin(targets["cluster_name"]))].copy()
    bg["set_num"] = bg["control_set"].astype(str).str.extract(r"(\d+)").astype(float)
    bg = bg[bg["set_num"].fillna(999999) <= 20].copy()
    bg = bg.merge(targets[["cluster_name", "module_id"]].rename(columns={"cluster_name": "matched_age_cluster"}), on="matched_age_cluster", how="left")
    bg["cluster_name"] = bg["control_set"].astype(str) + "|" + bg["matched_age_cluster"].astype(str)
    return targets, bg[["cluster_name", "module_id", "chr", "start", "end", "matched_age_cluster", "control_set"]].copy()


def overlap_analysis() -> pd.DataFrame:
    targets, bg = targets_bg()
    rows = []
    for track_id, cell_state, mark, path in TRACKS:
        peaks = read_bed(path)
        t = targets.copy()
        b = bg.copy()
        t["overlap"] = overlap_flags(t, peaks)
        b["overlap"] = overlap_flags(b, peaks)
        for module_id in MODULES:
            ts = t[t["module_id"] == module_id]
            bs = b[b["module_id"] == module_id]
            a = int(ts["overlap"].sum())
            b_no = int(len(ts) - a)
            c = int(bs["overlap"].sum())
            d = int(len(bs) - c)
            rows.append(
                {
                    "track_id": track_id,
                    "cell_state": cell_state,
                    "mark": mark,
                    "module_id": module_id,
                    "target_n": int(len(ts)),
                    "target_overlap_n": a,
                    "target_overlap_fraction": float(a / len(ts)) if len(ts) else 0.0,
                    "background_n": int(len(bs)),
                    "background_overlap_n": c,
                    "background_overlap_fraction": float(c / len(bs)) if len(bs) else 0.0,
                    "target_minus_background": float(a / len(ts)) - float(c / len(bs)) if len(ts) and len(bs) else 0.0,
                    "odds_ratio": odds_ratio(a, b_no, c, d),
                    "fisher_p_greater": fisher_greater(a, b_no, c, d),
                    "track_path": str(path),
                }
            )
    out = pd.DataFrame(rows)
    out["fisher_q_BH"] = fdr_bh(out["fisher_p_greater"])
    out.to_csv(OUT_OVERLAP, sep="\t", index=False)
    return out


def zscore(x: np.ndarray) -> np.ndarray:
    x = np.asarray(x, dtype=float)
    sd = np.nanstd(x, ddof=1)
    if not np.isfinite(sd) or sd == 0:
        return np.zeros_like(x)
    return (x - np.nanmean(x)) / sd


def basis() -> pd.DataFrame:
    b = pd.read_csv(BASIS, sep="\t")
    return b[b["module_id"].isin(MODULES)][["module_id", "latent_control_PC1", "latent_control_PC2", "latent_control_PC3", "n_DMRs", "ridge_weight"]].copy()


def feature_table(overlap: pd.DataFrame) -> pd.DataFrame:
    piv = overlap.pivot_table(index="module_id", columns=["cell_state", "mark"], values="target_minus_background", aggfunc="mean")
    rows = []
    candidates = {}
    # Biologically named hESC proxy candidates. These are not embryo-stage controls.
    candidates["hESC_proxy_closure_H3K27ac_low_naive"] = {m: -float(piv.loc[m, ("naive", "H3K27ac")]) if (m in piv.index and ("naive", "H3K27ac") in piv.columns) else 0.0 for m in CLOSURE}
    candidates["hESC_proxy_closure_H3K27me3_high_naive"] = {m: float(piv.loc[m, ("naive", "H3K27me3")]) if (m in piv.index and ("naive", "H3K27me3") in piv.columns) else 0.0 for m in CLOSURE}
    candidates["hESC_proxy_closure_H3K27ac_loss_primed_to_naive"] = {
        m: float(piv.loc[m, ("primed", "H3K27ac")] - piv.loc[m, ("naive", "H3K27ac")]) if m in piv.index else 0.0 for m in CLOSURE
    }
    candidates["hESC_proxy_access_H3K4me3_naive"] = {m: float(piv.loc[m, ("naive", "H3K4me3")]) if m in piv.index else 0.0 for m in ACCESS}
    candidates["hESC_proxy_access_H3K27ac_naive"] = {m: float(piv.loc[m, ("naive", "H3K27ac")]) if m in piv.index else 0.0 for m in ACCESS}
    candidates["hESC_proxy_dual_H3K27acloss_H3K4me3"] = {}
    candidates["hESC_proxy_dual_H3K27acloss_H3K4me3"].update(candidates["hESC_proxy_closure_H3K27ac_loss_primed_to_naive"])
    candidates["hESC_proxy_dual_H3K27acloss_H3K4me3"].update(candidates["hESC_proxy_access_H3K4me3_naive"])
    candidates["hESC_proxy_dual_H3K27aclow_H3K27me3_H3K4me3"] = {}
    for m in CLOSURE:
        candidates["hESC_proxy_dual_H3K27aclow_H3K27me3_H3K4me3"][m] = candidates["hESC_proxy_closure_H3K27ac_low_naive"].get(m, 0.0) + candidates["hESC_proxy_closure_H3K27me3_high_naive"].get(m, 0.0)
    for m in ACCESS:
        candidates["hESC_proxy_dual_H3K27aclow_H3K27me3_H3K4me3"][m] = candidates["hESC_proxy_access_H3K4me3_naive"].get(m, 0.0) + candidates["hESC_proxy_access_H3K27ac_naive"].get(m, 0.0)

    b = basis()
    for name, vals in candidates.items():
        tab = b.copy()
        tab["feature_set"] = name
        tab["raw_value"] = tab["module_id"].map(vals).fillna(0.0).astype(float)
        active = [m for m in MODULES if abs(vals.get(m, 0.0)) > 0 or ("dual" in name and m in MODULES)]
        tab["control_value_z"] = 0.0
        for group in [CLOSURE, ACCESS]:
            mask = tab["module_id"].isin(group) & tab["module_id"].isin(active)
            if int(mask.sum()) > 1:
                tab.loc[mask, "control_value_z"] = zscore(tab.loc[mask, "raw_value"].to_numpy(dtype=float))
            elif int(mask.sum()) == 1:
                tab.loc[mask, "control_value_z"] = tab.loc[mask, "raw_value"]
        tab["biological_status"] = "human_hESC_histone_state_proxy_not_embryo_morula"
        tab["control_modality"] = "hESC_histone_BED"
        tab["description"] = "Public human naive/primed hESC histone peaks from GSE52617; tests biological identity but is not final embryo u_bio."
        rows.append(tab)
    out = pd.concat(rows, ignore_index=True)
    out.to_csv(OUT_FEATURES, sep="\t", index=False)
    return out


def latent_context():
    matrix, ann, pairs = load_inputs()
    mu, sd, components, score_df = fit_latent_basis(matrix, 3)
    coef, _, _ = fit_operator(score_df, pairs, train_stages=set(PRE_MORULA_STAGES), lam=1000.0)
    strict_z = simulate_strict_pre_morula(score_df, ann, coef, n_steps=12)
    obs_z = score_df.loc[stage_ids(ann, "morula")].to_numpy(dtype=float)
    obs_dmr = matrix.loc[stage_ids(ann, "morula")].to_numpy(dtype=float)
    basin = basin_definition(obs_z)
    residual_z = obs_z.mean(axis=0) - strict_z.mean(axis=0)
    return mu, sd, components, strict_z, obs_z, obs_dmr, basin, residual_z


def alpha_values() -> list[float]:
    return [round(float(x), 2) for x in np.arange(0.0, 2.5001, 0.05)]


def evaluate(features: pd.DataFrame) -> pd.DataFrame:
    mu, sd, components, strict_z, obs_z, obs_dmr, basin, residual_z = latent_context()
    rng = np.random.default_rng(20260527)
    rows = []
    for feature_set, sub in features.groupby("feature_set"):
        dirs = sub[["latent_control_PC1", "latent_control_PC2", "latent_control_PC3"]].to_numpy(dtype=float)
        u = pd.to_numeric(sub["control_value_z"], errors="coerce").fillna(0.0).to_numpy(dtype=float)
        control = (dirs * u[:, None]).sum(axis=0)
        for alpha in alpha_values():
            for sign, status in [(1.0, "correct_orientation"), (-1.0, "sign_flip_control")]:
                vec = sign * alpha * control
                pred = strict_z + vec[None, :]
                pred_dmr = decode_latent(pred, mu, sd, components)
                rows.append(
                    {
                        "feature_set": feature_set,
                        "alpha": alpha,
                        "sign_status": status,
                        "PC1_control": float(vec[0]),
                        "PC2_control": float(vec[1]),
                        "PC3_control": float(vec[2]),
                        "direction_cosine_to_measured_correction": cosine(vec, residual_z) if np.linalg.norm(vec) else np.nan,
                        "PC3_negative_pull_recovered": float(-vec[2] / (-residual_z[2])) if residual_z[2] < 0 else np.nan,
                        **distribution_metrics(pred, obs_z, pred_dmr, obs_dmr, basin, rng),
                    }
                )
    out = pd.DataFrame(rows)
    out.to_csv(OUT_ALPHA, sep="\t", index=False)
    return out


def summarize(alpha: pd.DataFrame, overlap: pd.DataFrame) -> pd.DataFrame:
    rows = []
    for feature_set, sub in alpha[alpha["sign_status"] == "correct_orientation"].groupby("feature_set"):
        sub = sub.sort_values("alpha")
        best = sub.sort_values(["pred_basin_occupancy_q90", "direction_cosine_to_measured_correction"], ascending=False).iloc[0]
        at1 = sub.iloc[(sub["alpha"] - 1.0).abs().argsort()].iloc[0]
        flip = alpha[(alpha["feature_set"] == feature_set) & (alpha["sign_status"] == "sign_flip_control")].sort_values("pred_basin_occupancy_q90", ascending=False).iloc[0]
        rows.append(
            {
                "feature_set": feature_set,
                "max_occupancy": float(best["pred_basin_occupancy_q90"]),
                "alpha_at_max": float(best["alpha"]),
                "occupancy_at_alpha_1": float(at1["pred_basin_occupancy_q90"]),
                "cosine_at_alpha_1": float(at1["direction_cosine_to_measured_correction"]),
                "PC3_recovery_at_alpha_1": float(at1["PC3_negative_pull_recovered"]),
                "max_sign_flip_occupancy": float(flip["pred_basin_occupancy_q90"]),
            }
        )
    out = pd.DataFrame(rows).sort_values(["max_occupancy", "cosine_at_alpha_1"], ascending=False)
    out.to_csv(OUT_SUMMARY, sep="\t", index=False)
    return out


def make_svg(summary: pd.DataFrame) -> None:
    rows = summary.head(10)
    width, height = 920, 410
    left, right, top, bottom = 85, 25, 45, 135
    plot_w = width - left - right
    plot_h = height - top - bottom
    gap = plot_w / max(len(rows), 1)
    bar_w = gap * 0.62
    lines = [
        f'<svg xmlns="http://www.w3.org/2000/svg" width="{width}" height="{height}" viewBox="0 0 {width} {height}">',
        '<rect width="100%" height="100%" fill="white"/>',
        f'<text x="{left}" y="28" font-family="Arial" font-size="18" font-weight="700">hESC histone-state branch identity proxy</text>',
        f'<line x1="{left}" y1="{height-bottom}" x2="{width-right}" y2="{height-bottom}" stroke="#222"/>',
        f'<line x1="{left}" y1="{top}" x2="{left}" y2="{height-bottom}" stroke="#222"/>',
    ]
    for tick in [0.044, 0.3, 0.5, 0.875, 1.0]:
        y = height - bottom - tick * plot_h
        color = "#b23b3b" if tick in [0.044, 0.875] else "#dddddd"
        lines.append(f'<line x1="{left}" y1="{y:.2f}" x2="{width-right}" y2="{y:.2f}" stroke="{color}" stroke-dasharray="4 4"/>')
        lines.append(f'<text x="{left-8}" y="{y+4:.2f}" text-anchor="end" font-family="Arial" font-size="10">{tick:.3g}</text>')
    for i, row in enumerate(rows.itertuples()):
        x = left + i * gap + (gap - bar_w) / 2
        val = float(row.max_occupancy)
        y = height - bottom - val * plot_h
        lines.append(f'<rect x="{x:.2f}" y="{y:.2f}" width="{bar_w:.2f}" height="{val * plot_h:.2f}" fill="#6f4d8b"/>')
        lines.append(f'<text x="{x+bar_w/2:.2f}" y="{y-5:.2f}" text-anchor="middle" font-family="Arial" font-size="10">{val:.3f}</text>')
        label = row.feature_set.replace("_", " ")[:40]
        lines.append(f'<text x="{x+bar_w/2:.2f}" y="{height-bottom+18}" text-anchor="end" transform="rotate(-45 {x+bar_w/2:.2f} {height-bottom+18})" font-family="Arial" font-size="9">{label}</text>')
    lines.append("</svg>")
    OUT_SVG.write_text("\n".join(lines), encoding="utf-8")


def write_doc(summary: pd.DataFrame, overlap: pd.DataFrame) -> None:
    lines = [
        "# hESC Histone-State Branch Identity Proxy",
        "",
        "Status: `completed_human_hESC_proxy_not_embryo_morula`",
        "",
        "These tracks are public human naive/primed hESC histone peak BEDs from GSE52617. They are useful for testing biological branch identity, but they are not embryo morula histone tracks and must not be called final u_bio.",
        "",
        "## Top Dynamics",
        "",
    ]
    for row in summary.head(10).itertuples():
        lines.append(
            f"- {row.feature_set}: max_occ={row.max_occupancy:.3f}; occ@1={row.occupancy_at_alpha_1:.3f}; "
            f"cosine@1={row.cosine_at_alpha_1:.3f}; PC3@1={row.PC3_recovery_at_alpha_1:.3f}; signflip={row.max_sign_flip_occupancy:.3f}"
        )
    lines += ["", "## Strongest Module Histone Overlaps", ""]
    best = overlap.sort_values("odds_ratio", ascending=False).head(12)
    for row in best.itertuples():
        lines.append(
            f"- {row.module_id} {row.track_id}: target={row.target_overlap_fraction:.3f}, bg={row.background_overlap_fraction:.3f}, OR={row.odds_ratio:.2f}, q={row.fisher_q_BH:.3g}"
        )
    lines += ["", "## Boundary", "", "This is a biological-state proxy layer. Final biologically interpretable control still requires embryo-stage H3K27ac/H3K4me3/H3K27me3 or equivalent chromatin-state tracks."]
    OUT_DOC.write_text("\n".join(lines) + "\n", encoding="utf-8")


def main() -> None:
    RESULTS.mkdir(parents=True, exist_ok=True)
    FIGURES.mkdir(parents=True, exist_ok=True)
    DOCS.mkdir(parents=True, exist_ok=True)
    overlap = overlap_analysis()
    features = feature_table(overlap)
    alpha = evaluate(features)
    summary = summarize(alpha, overlap)
    make_svg(summary)
    write_doc(summary, overlap)
    OUT_MANIFEST.write_text(
        json.dumps(
            {
                "status": "completed_human_hESC_proxy_not_embryo_morula",
                "outputs": [str(OUT_OVERLAP), str(OUT_FEATURES), str(OUT_ALPHA), str(OUT_SUMMARY), str(OUT_SVG), str(OUT_DOC)],
                "boundary": "human hESC histone proxy, not final embryo u_bio",
            },
            indent=2,
        )
        + "\n",
        encoding="utf-8",
    )
    print(json.dumps({"status": "completed", "top": summary.head(8).to_dict(orient="records")}, indent=2))


if __name__ == "__main__":
    main()
```
