## Supplemental code for "An entropy-based diagnostic framework for characterizing methylation state dynamics during preimplantation development": run_histone_data_access_audit.py

````
from __future__ import annotations

import json
from pathlib import Path

import pandas as pd


BASE = Path(r"E:\CSB_TRO_operator_time_DMR_dynamics_2026-05-25")
EXTERNAL = BASE / "external"
RESULTS = BASE / "results"
DOCS = BASE / "docs"
HISTONE = EXTERNAL / "histone"

OUT_MANIFEST = HISTONE / "histone_peak_manifest.tsv"
OUT_AUDIT = RESULTS / "CSB_TRO_histone_data_access_audit.tsv"
OUT_JSON = RESULTS / "CSB_TRO_histone_data_access_audit.json"
OUT_DOC = DOCS / "CSB_TRO_histone_data_access_and_overlap_plan.md"


EXPECTED_TRACKS = [
    {
        "track_id": "H3K27ac_8cell",
        "mark": "H3K27ac",
        "stage": "8-cell",
        "path": HISTONE / "H3K27ac_8cell.hg19.bed",
        "required_for": "8cell_to_morula_histone_transition",
    },
    {
        "track_id": "H3K27ac_morula",
        "mark": "H3K27ac",
        "stage": "morula",
        "path": HISTONE / "H3K27ac_morula.hg19.bed",
        "required_for": "8cell_to_morula_histone_transition",
    },
    {
        "track_id": "H3K27ac_blastocyst",
        "mark": "H3K27ac",
        "stage": "blastocyst",
        "path": HISTONE / "H3K27ac_blastocyst.hg19.bed",
        "required_for": "morula_to_blastocyst_histone_transition",
    },
    {
        "track_id": "H3K4me3_8cell",
        "mark": "H3K4me3",
        "stage": "8-cell",
        "path": HISTONE / "H3K4me3_8cell.hg19.bed",
        "required_for": "8cell_to_morula_histone_transition",
    },
    {
        "track_id": "H3K4me3_morula",
        "mark": "H3K4me3",
        "stage": "morula",
        "path": HISTONE / "H3K4me3_morula.hg19.bed",
        "required_for": "8cell_to_morula_histone_transition",
    },
    {
        "track_id": "H3K4me3_blastocyst",
        "mark": "H3K4me3",
        "stage": "blastocyst",
        "path": HISTONE / "H3K4me3_blastocyst.hg19.bed",
        "required_for": "morula_to_blastocyst_histone_transition",
    },
]


DATASETS = [
    {
        "dataset": "human_early_embryo_histone_acetylation",
        "reported_accessions": "PRJCA009410; HRA002355; CRA006815",
        "source_url": "https://www.nature.com/articles/s41421-022-00514-y",
        "expected_marks": "H3K27ac; H3K4me3",
        "expected_stages": "8-cell; morula; blastocyst",
        "local_status": "manifest_prepared",
        "access_boundary": (
            "This step requires processed peak BED files or controlled-access raw data "
            "downloaded outside the script. The audit does not treat missing files as "
            "negative biological evidence."
        ),
    },
]


def file_status(path: Path) -> dict[str, object]:
    gz_path = Path(str(path) + ".gz")
    if path.exists():
        return {"exists": True, "resolved_path": str(path), "size_bytes": path.stat().st_size}
    if gz_path.exists():
        return {"exists": True, "resolved_path": str(gz_path), "size_bytes": gz_path.stat().st_size}
    return {"exists": False, "resolved_path": str(path), "size_bytes": 0}


def main() -> None:
    HISTONE.mkdir(parents=True, exist_ok=True)
    RESULTS.mkdir(parents=True, exist_ok=True)
    DOCS.mkdir(parents=True, exist_ok=True)

    manifest_rows = []
    for row in EXPECTED_TRACKS:
        status = file_status(Path(row["path"]))
        manifest_rows.append(
            {
                "track_id": row["track_id"],
                "mark": row["mark"],
                "stage": row["stage"],
                "genome_build": "hg19/GRCh37",
                "expected_path": str(row["path"]),
                "resolved_path": status["resolved_path"],
                "file_exists": status["exists"],
                "size_bytes": status["size_bytes"],
                "required_for": row["required_for"],
                "format": "BED3+ optionally gzip-compressed",
            }
        )
    manifest = pd.DataFrame(manifest_rows)
    manifest.to_csv(OUT_MANIFEST, sep="\t", index=False)

    complete_8m = bool(
        manifest.loc[
            manifest["track_id"].isin(["H3K27ac_8cell", "H3K27ac_morula", "H3K4me3_8cell", "H3K4me3_morula"]),
            "file_exists",
        ].all()
    )
    complete_any = bool(manifest["file_exists"].any())
    audit = pd.DataFrame(
        [
            {
                **DATASETS[0],
                "expected_local_tracks": int(len(manifest)),
                "available_local_tracks": int(manifest["file_exists"].sum()),
                "any_histone_overlap_runnable": complete_any,
                "full_8cell_morula_transition_runnable": complete_8m,
                "manifest": str(OUT_MANIFEST),
            }
        ]
    )
    audit.to_csv(OUT_AUDIT, sep="\t", index=False)
    OUT_JSON.write_text(
        json.dumps(
            {
                "manifest": str(OUT_MANIFEST),
                "available_local_tracks": int(manifest["file_exists"].sum()),
                "expected_local_tracks": int(len(manifest)),
                "any_histone_overlap_runnable": complete_any,
                "full_8cell_morula_transition_runnable": complete_8m,
                "missing_tracks": manifest.loc[~manifest["file_exists"], "track_id"].tolist(),
                "boundary": DATASETS[0]["access_boundary"],
            },
            indent=2,
        )
        + "\n",
        encoding="utf-8",
    )

    lines = [
        "# Histone Data Access Audit and Overlap Plan",
        "",
        "Goal: test whether M02 KLF4/KLF5 residual DMR targets overlap human early-embryo H3K27ac/H3K4me3 peaks at 8-cell, morula, and blastocyst stages.",
        "",
        "## Access Boundary",
        "",
        "Source/accession note: the human early-embryo histone acetylation study reports H3K27ac/H3K4me3 dynamics and associated accessions `PRJCA009410`, `HRA002355`, and `CRA006815`. Local overlap requires processed BED peaks or raw-data-derived peaks in the same genome build.",
        "",
        DATASETS[0]["access_boundary"],
        "",
        "## Local Track Status",
        "",
        f"- Expected histone tracks: {len(manifest)}",
        f"- Available local tracks: {int(manifest['file_exists'].sum())}",
        f"- Any overlap runnable: {str(complete_any)}",
        f"- Full 8-cell to morula transition runnable: {str(complete_8m)}",
        "",
        "Expected local files are listed in:",
        "",
        f"`{OUT_MANIFEST}`",
        "",
        "## Required Next Input",
        "",
        "Place BED3+ peak files at the manifest paths, preferably in hg19/GRCh37 coordinates:",
        "",
    ]
    for row in manifest_rows:
        lines.append(f"- `{row['expected_path']}`")
    lines += [
        "",
        "After files are present, run:",
        "",
        "```text",
        "python code\\run_histone_overlap_for_m02_klf.py",
        "```",
        "",
        "Missing histone files should be reported as `not_run_missing_histone_input`, not as absence of H3K27ac/H3K4me3 support.",
    ]
    OUT_DOC.write_text("\n".join(lines) + "\n", encoding="utf-8")
    print(
        json.dumps(
            {
                "expected_tracks": int(len(manifest)),
                "available_tracks": int(manifest["file_exists"].sum()),
                "full_8cell_morula_transition_runnable": complete_8m,
                "manifest": str(OUT_MANIFEST),
            },
            indent=2,
        )
    )


if __name__ == "__main__":
    main()
````
