## Supplemental code for "An entropy-based diagnostic framework for characterizing methylation state dynamics during preimplantation development": run_histone_data_acquisition_manifest.py

````
from __future__ import annotations

import json
from pathlib import Path

import pandas as pd


BASE = Path(r"E:\CSB_TRO_operator_time_DMR_dynamics_2026-05-25")
EXTERNAL = BASE / "external"
HISTONE = EXTERNAL / "histone"
RESULTS = BASE / "results"
DOCS = BASE / "docs"

OUT_TSV = HISTONE / "histone_data_acquisition_manifest.tsv"
OUT_JSON = RESULTS / "CSB_TRO_histone_data_acquisition_manifest.json"
OUT_DOC = DOCS / "CSB_TRO_histone_data_acquisition_plan.md"

MARKS = ["H3K27ac", "H3K4me3", "H3K27me3"]
STAGES = ["8cell", "morula", "blastocyst"]


def file_status(path: Path) -> tuple[bool, str, int]:
    gz = Path(str(path) + ".gz")
    if path.exists():
        return True, str(path), int(path.stat().st_size)
    if gz.exists():
        return True, str(gz), int(gz.stat().st_size)
    return False, str(path), 0


def main() -> None:
    HISTONE.mkdir(parents=True, exist_ok=True)
    RESULTS.mkdir(parents=True, exist_ok=True)
    DOCS.mkdir(parents=True, exist_ok=True)

    rows = []
    for mark in MARKS:
        for stage in STAGES:
            bed = HISTONE / f"{mark}_{stage}.hg19.bed"
            bw = HISTONE / f"{mark}_{stage}.hg19.bw"
            bed_ok, bed_resolved, bed_size = file_status(bed)
            bw_ok, bw_resolved, bw_size = file_status(bw)
            rows.append(
                {
                    "track_id": f"{mark}_{stage}",
                    "mark": mark,
                    "stage": stage,
                    "preferred_input": "processed_peak_BED_or_broadPeak",
                    "alternate_input": "bigWig_signal_track",
                    "genome_build": "hg19/GRCh37",
                    "expected_bed_path": str(bed),
                    "resolved_bed_path": bed_resolved,
                    "bed_exists": bed_ok,
                    "bed_size_bytes": bed_size,
                    "expected_bigwig_path": str(bw),
                    "resolved_bigwig_path": bw_resolved,
                    "bigwig_exists": bw_ok,
                    "bigwig_size_bytes": bw_size,
                    "raw_data_fallback": "FASTQ/BAM requires reproducible alignment, QC, duplicate handling, and mark-specific peak/signal calling.",
                    "status": "ready" if bed_ok or bw_ok else "missing_processed_input",
                }
            )
    manifest = pd.DataFrame(rows)
    manifest.to_csv(OUT_TSV, sep="\t", index=False)

    source_rows = [
        {
            "source": "Cell Discovery 2023 human early embryo H3K27ac/H3K4me3",
            "accessions": "PRJCA009410; HRA002355",
            "usefulness": "highest_priority_human_stage_matched_histone_source",
            "current_boundary": "human raw data appears controlled-access; no local processed BED/bigWig found",
        },
        {
            "source": "DevOmics",
            "accessions": "database resource",
            "usefulness": "possible processed/promoter-level epigenomic signal entry point",
            "current_boundary": "need concrete downloadable track/table URLs before use",
        },
        {
            "source": "CRA006815",
            "accessions": "CRA006815",
            "usefulness": "related early embryo histone acetylation raw-data source",
            "current_boundary": "not a direct local human processed BED input; treat as raw-data fallback only",
        },
    ]
    status = {
        "expected_tracks": int(len(manifest)),
        "tracks_with_bed": int(manifest["bed_exists"].sum()),
        "tracks_with_bigwig": int(manifest["bigwig_exists"].sum()),
        "ready_tracks": int((manifest["bed_exists"] | manifest["bigwig_exists"]).sum()),
        "manifest": str(OUT_TSV),
        "sources": source_rows,
    }
    OUT_JSON.write_text(json.dumps(status, indent=2) + "\n", encoding="utf-8")

    lines = [
        "# Histone Data Acquisition Plan",
        "",
        "Goal: obtain processed or signal-level H3K27ac/H3K4me3/H3K27me3 inputs for M05/M01/M12/M02/M10 residual module control analysis.",
        "",
        "## Current Local Status",
        "",
        f"- Expected histone tracks: {len(manifest)}",
        f"- Tracks with BED/broadPeak: {int(manifest['bed_exists'].sum())}",
        f"- Tracks with bigWig signal: {int(manifest['bigwig_exists'].sum())}",
        f"- Ready tracks: {int((manifest['bed_exists'] | manifest['bigwig_exists']).sum())}",
        "",
        "## Accepted Inputs",
        "",
        "Preferred: processed peak files in BED3+, narrowPeak, or broadPeak form.",
        "",
        "Alternate: bigWig signal tracks. These can support mean/max signal over DMRs and stage-delta module scores, but need a bigWig summarization tool such as `bigWigAverageOverBed` or `pyBigWig`.",
        "",
        "Raw fallback: FASTQ/BAM can be used only after reproducible preprocessing. H3K27ac/H3K4me3 and H3K27me3 should not be peak-called with identical assumptions because H3K27me3 is broad-domain-like.",
        "",
        "## Source Status",
        "",
    ]
    for row in source_rows:
        lines.append(f"- {row['source']}: {row['usefulness']}; boundary: {row['current_boundary']}.")
    lines += [
        "",
        "## Decision Rule",
        "",
        "Do not interpret missing local files as negative histone evidence. Once any BED/bigWig is available at the manifest paths, rerun:",
        "",
        "```text",
        "python code\\run_residual_module_histone_state_control.py",
        "```",
    ]
    OUT_DOC.write_text("\n".join(lines) + "\n", encoding="utf-8")
    print(json.dumps(status, indent=2))


if __name__ == "__main__":
    main()
````
