## Supplemental code for "An entropy-based diagnostic framework for characterizing methylation state dynamics during preimplantation development": run_histone_overlap_for_m02_klf.py

```
from __future__ import annotations

import gzip
import json
from pathlib import Path

import pandas as pd


BASE = Path(r"E:\CSB_TRO_operator_time_DMR_dynamics_2026-05-25")
EXTERNAL = BASE / "external"
RESULTS = BASE / "results"
DOCS = BASE / "docs"
HISTONE = EXTERNAL / "histone"

TARGETS = RESULTS / "CSB_TRO_M02_KLF4_KLF5_residual_DMR_targets.tsv"
MANIFEST = HISTONE / "histone_peak_manifest.tsv"

OUT_OVERLAP = RESULTS / "CSB_TRO_M02_KLF4_KLF5_histone_overlap.tsv"
OUT_SUMMARY = RESULTS / "CSB_TRO_M02_KLF4_KLF5_histone_overlap_summary.tsv"
OUT_JSON = RESULTS / "CSB_TRO_M02_KLF4_KLF5_histone_overlap_manifest.json"
OUT_DOC = DOCS / "CSB_TRO_M02_KLF4_KLF5_histone_overlap_summary.md"


def read_bed(path: Path) -> pd.DataFrame:
    opener = gzip.open if path.suffix == ".gz" else open
    rows = []
    with opener(path, "rt", encoding="utf-8", errors="replace") as handle:
        for line in handle:
            if not line.strip() or line.startswith(("#", "track", "browser")):
                continue
            parts = line.rstrip("\n").split("\t")
            if len(parts) < 3:
                continue
            try:
                start = int(float(parts[1]))
                end = int(float(parts[2]))
            except ValueError:
                continue
            rows.append(
                {
                    "chr": parts[0],
                    "start": start,
                    "end": end,
                    "name": parts[3] if len(parts) > 3 else f"{parts[0]}:{start}-{end}",
                }
            )
    return pd.DataFrame(rows)


def overlap_count(row: pd.Series, by_chr: dict[str, pd.DataFrame]) -> tuple[int, str]:
    sub = by_chr.get(str(row["chr"]))
    if sub is None or sub.empty:
        return 0, ""
    hits = sub[(sub["start"] < int(row["end"])) & (sub["end"] > int(row["start"]))]
    names = ",".join(hits["name"].astype(str).head(10).tolist())
    return int(len(hits)), names


def write_not_run(reason: str, missing_tracks: list[str]) -> None:
    pd.DataFrame(
        [
            {
                "analysis_status": "not_run_missing_histone_input",
                "reason": reason,
                "missing_tracks": ",".join(missing_tracks),
                "required_manifest": str(MANIFEST),
                "targets": str(TARGETS),
            }
        ]
    ).to_csv(OUT_SUMMARY, sep="\t", index=False)
    pd.DataFrame(
        [
            {
                "analysis_status": "not_run_missing_histone_input",
                "cluster_name": "",
                "module_id": "M02",
                "reason": reason,
            }
        ]
    ).to_csv(OUT_OVERLAP, sep="\t", index=False)
    OUT_JSON.write_text(
        json.dumps(
            {
                "analysis_status": "not_run_missing_histone_input",
                "reason": reason,
                "missing_tracks": missing_tracks,
                "manifest": str(MANIFEST),
            },
            indent=2,
        )
        + "\n",
        encoding="utf-8",
    )
    OUT_DOC.write_text(
        "\n".join(
            [
                "# M02 KLF4/KLF5 Histone Overlap",
                "",
                "Status: `not_run_missing_histone_input`",
                "",
                reason,
                "",
                "Missing tracks:",
                "",
                *[f"- `{track}`" for track in missing_tracks],
                "",
                "This is an input availability boundary, not negative H3K27ac/H3K4me3 evidence.",
            ]
        )
        + "\n",
        encoding="utf-8",
    )


def transition_flag(df: pd.DataFrame, mark: str, start_stage: str, end_stage: str) -> pd.Series:
    start_col = f"{mark}_{start_stage}_overlap"
    end_col = f"{mark}_{end_stage}_overlap"
    if start_col not in df.columns or end_col not in df.columns:
        return pd.Series([False] * len(df), index=df.index)
    return (~df[start_col].astype(bool)) & df[end_col].astype(bool)


def main() -> None:
    if not MANIFEST.exists():
        write_not_run("Histone peak manifest is missing. Run code\\run_histone_data_access_audit.py first.", [str(MANIFEST)])
        print(json.dumps({"analysis_status": "not_run_missing_histone_input", "reason": "missing manifest"}, indent=2))
        return
    if not TARGETS.exists():
        write_not_run("M02 KLF4/KLF5 target table is missing. Run code\\run_export_m02_klf_target_bed.py first.", [str(TARGETS)])
        print(json.dumps({"analysis_status": "not_run_missing_histone_input", "reason": "missing targets"}, indent=2))
        return

    manifest = pd.read_csv(MANIFEST, sep="\t")
    available = manifest[manifest["file_exists"].astype(bool)].copy()
    if available.empty:
        write_not_run(
            "No local H3K27ac/H3K4me3 peak BED files are present at the manifest paths.",
            manifest["track_id"].astype(str).tolist(),
        )
        print(json.dumps({"analysis_status": "not_run_missing_histone_input", "available_tracks": 0}, indent=2))
        return

    targets = pd.read_csv(TARGETS, sep="\t")
    out = targets[["cluster_name", "module_id", "chr", "start", "end", "TFs", "basin_residual_rank"]].copy()
    track_summaries = []
    for rec in available.to_dict(orient="records"):
        track_id = str(rec["track_id"])
        path = Path(str(rec["resolved_path"]))
        peaks = read_bed(path)
        by_chr = {chrom: sub for chrom, sub in peaks.groupby("chr")}
        counts = []
        names = []
        for _, row in out.iterrows():
            count, hit_names = overlap_count(row, by_chr)
            counts.append(count)
            names.append(hit_names)
        out[f"{track_id}_overlap_count"] = counts
        out[f"{track_id}_overlap"] = [count > 0 for count in counts]
        out[f"{track_id}_overlap_names"] = names
        track_summaries.append(
            {
                "track_id": track_id,
                "mark": rec["mark"],
                "stage": rec["stage"],
                "n_targets": int(len(out)),
                "n_overlap": int(sum(count > 0 for count in counts)),
                "overlap_fraction": float(sum(count > 0 for count in counts) / len(out)) if len(out) else 0.0,
                "n_peaks": int(len(peaks)),
                "path": str(path),
            }
        )

    for mark in sorted(available["mark"].astype(str).unique()):
        out[f"{mark}_8cell_to_morula_gain"] = transition_flag(out, mark, "8cell", "morula")
        out[f"{mark}_morula_to_blastocyst_gain"] = transition_flag(out, mark, "morula", "blastocyst")

    out.to_csv(OUT_OVERLAP, sep="\t", index=False)

    summary = pd.DataFrame(track_summaries)
    if not summary.empty:
        gain_rows = []
        for mark in sorted(available["mark"].astype(str).unique()):
            for transition in ["8cell_to_morula", "morula_to_blastocyst"]:
                col = f"{mark}_{transition}_gain"
                if col in out.columns:
                    gain_rows.append(
                        {
                            "track_id": f"{mark}_{transition}_gain",
                            "mark": mark,
                            "stage": transition,
                            "n_targets": int(len(out)),
                            "n_overlap": int(out[col].sum()),
                            "overlap_fraction": float(out[col].mean()) if len(out) else 0.0,
                            "n_peaks": "",
                            "path": "derived_transition_gain",
                        }
                    )
        if gain_rows:
            summary = pd.concat([summary, pd.DataFrame(gain_rows)], ignore_index=True)
    summary.to_csv(OUT_SUMMARY, sep="\t", index=False)

    status = {
        "analysis_status": "completed",
        "n_targets": int(len(out)),
        "available_tracks": int(len(available)),
        "outputs": [str(OUT_OVERLAP), str(OUT_SUMMARY), str(OUT_DOC)],
    }
    OUT_JSON.write_text(json.dumps(status, indent=2) + "\n", encoding="utf-8")

    lines = [
        "# M02 KLF4/KLF5 Histone Overlap Summary",
        "",
        "Status: `completed`",
        "",
        f"Target DMRs: {len(out)}",
        f"Available histone tracks: {len(available)}",
        "",
        "## Track Overlap Fractions",
        "",
    ]
    for row in summary.to_dict(orient="records"):
        lines.append(f"- {row['track_id']}: {float(row['overlap_fraction']):.3f} ({row['n_overlap']}/{row['n_targets']})")
    lines += [
        "",
        "Interpretation boundary: these overlaps are only valid for the local BED files listed in the manifest and their genome build. If the files were lifted or filtered upstream, that preprocessing must be reported with the result.",
    ]
    OUT_DOC.write_text("\n".join(lines) + "\n", encoding="utf-8")
    print(json.dumps(status, indent=2))


if __name__ == "__main__":
    main()
```
