## Supplemental code for "An entropy-based diagnostic framework for characterizing methylation state dynamics during preimplantation development": run_inverse_ATAC_module_decomposition.py

```
from __future__ import annotations

import json
import sys
from itertools import combinations
from pathlib import Path

import numpy as np
import pandas as pd


BASE = Path(r"E:\CSB_TRO_operator_time_DMR_dynamics_2026-05-25")
CODE = BASE / "code"
RESULTS = BASE / "results"
FIGURES = BASE / "figures"
DOCS = BASE / "docs"
sys.path.insert(0, str(CODE))

from run_basin_residual_control_field import PRE_MORULA_STAGES, cosine, simulate_strict_pre_morula  # noqa: E402
from run_morula_basin_sde import (  # noqa: E402
    basin_definition,
    decode_latent,
    distribution_metrics,
    fit_latent_basis,
    fit_operator,
    load_inputs,
    stage_ids,
)


PRIORITY_MODULES = ["M05", "M01", "M12", "M02", "M10"]
CORE_ORDER = ["M05", "M01", "M12", "M02"]
ATAC_FEATURES = ["ATAC_8cell_2pn_chromatin_only", "ATAC_8cell_3pn_chromatin_only"]

CHROM_FEATURES = RESULTS / "CSB_TRO_chromatin_gated_TF_activity.tsv"
BASIS = RESULTS / "CSB_TRO_missing_control_term_module_basis.tsv"
OVERLAP = RESULTS / "CSB_TRO_module_chromatin_overlap.tsv"
REGION = RESULTS / "CSB_TRO_residual_module_region_composition.tsv"

OUT_CONTRIB = RESULTS / "CSB_TRO_inverse_ATAC_module_contributions.tsv"
OUT_PANEL = RESULTS / "CSB_TRO_inverse_ATAC_module_decomposition_panel.tsv"
OUT_RANDOM = RESULTS / "CSB_TRO_inverse_ATAC_module_matched_random.tsv"
OUT_SUMMARY = RESULTS / "CSB_TRO_inverse_ATAC_module_decomposition_summary.tsv"
OUT_MANIFEST = RESULTS / "CSB_TRO_inverse_ATAC_module_decomposition_manifest.json"
OUT_SVG = FIGURES / "CSB_TRO_inverse_ATAC_module_decomposition.svg"
OUT_DOC = DOCS / "CSB_TRO_inverse_ATAC_module_decomposition_summary.md"


def alpha_values() -> list[float]:
    return [round(float(x), 2) for x in np.arange(0.0, 2.5001, 0.05)]


def load_latent_context():
    matrix, ann, pairs = load_inputs()
    mu, sd, components, score_df = fit_latent_basis(matrix, 3)
    coef, _, _ = fit_operator(score_df, pairs, train_stages=set(PRE_MORULA_STAGES), lam=1000.0)
    strict_z = simulate_strict_pre_morula(score_df, ann, coef, n_steps=12)
    obs_z = score_df.loc[stage_ids(ann, "morula")].to_numpy(dtype=float)
    obs_dmr = matrix.loc[stage_ids(ann, "morula")].to_numpy(dtype=float)
    basin = basin_definition(obs_z)
    residual_z = obs_z.mean(axis=0) - strict_z.mean(axis=0)
    return matrix, mu, sd, components, strict_z, obs_z, obs_dmr, basin, residual_z


def zscore(x: np.ndarray) -> np.ndarray:
    x = np.asarray(x, dtype=float)
    sd = np.nanstd(x, ddof=1)
    if not np.isfinite(sd) or sd == 0:
        return np.zeros_like(x)
    return (x - np.nanmean(x)) / sd


def feature_table() -> pd.DataFrame:
    feat = pd.read_csv(CHROM_FEATURES, sep="\t")
    basis = pd.read_csv(BASIS, sep="\t")
    overlap = pd.read_csv(OVERLAP, sep="\t")
    region = pd.read_csv(REGION, sep="\t") if REGION.exists() else pd.DataFrame()
    rows = []
    for fs in ATAC_FEATURES:
        sub = feat[feat["feature_set"] == fs].copy()
        if sub.empty:
            continue
        for row in sub.itertuples():
            module_id = str(row.module_id)
            direction = np.asarray([float(row.latent_control_PC1), float(row.latent_control_PC2), float(row.latent_control_PC3)])
            raw_u = float(row.control_value_z)
            inv_u = -raw_u
            vec = direction * inv_u
            rec = {
                "feature_set": fs + "_inverse",
                "source_feature_set": fs,
                "module_id": module_id,
                "raw_ATAC_control_z": raw_u,
                "inverse_ATAC_control_z": inv_u,
                "module_PC1_contribution": float(vec[0]),
                "module_PC2_contribution": float(vec[1]),
                "module_PC3_contribution": float(vec[2]),
                "module_control_norm": float(np.linalg.norm(vec)),
            }
            ov = overlap[(overlap["track_id"] == fs.replace("_chromatin_only", "").replace("ATAC_", "ATAC_")) & (overlap["module_id"] == module_id)]
            if len(ov):
                r = ov.iloc[0]
                rec.update(
                    {
                        "target_overlap_fraction": float(r["target_overlap_fraction"]),
                        "background_overlap_fraction": float(r["background_overlap_fraction"]),
                        "overlap_odds_ratio": float(r["overlap_odds_ratio"]),
                        "fisher_q_BH": float(r["fisher_q_BH"]),
                    }
                )
            if len(region):
                rg = region[region["module_id"] == module_id]
                if len(rg):
                    rec["promoter_2kb_fraction"] = float(rg.iloc[0]["promoter_2kb_fraction"])
                    rec["intergenic_fraction"] = float(rg.iloc[0]["intergenic_fraction"])
            bw = basis[basis["module_id"] == module_id]
            if len(bw):
                rec["measured_ridge_weight"] = float(bw.iloc[0]["ridge_weight"])
            rows.append(rec)
    out = pd.DataFrame(rows)
    out.to_csv(OUT_CONTRIB, sep="\t", index=False)
    return out


def evaluate_panel(contrib: pd.DataFrame):
    _, mu, sd, components, strict_z, obs_z, obs_dmr, basin, residual_z = load_latent_context()
    rng = np.random.default_rng(20260526)
    rows = []
    random_rows = []

    baseline_dmr = decode_latent(strict_z, mu, sd, components)
    rows.append(
        {
            "feature_set": "methylation_only_baseline",
            "panel_model": "methylation_only_baseline",
            "included_modules": "",
            "n_modules": 0,
            "alpha": 0.0,
            "validation_status": "baseline",
            "PC1_control": 0.0,
            "PC2_control": 0.0,
            "PC3_control": 0.0,
            "direction_cosine_to_measured_correction": np.nan,
            "PC3_negative_pull_recovered": 0.0,
            **distribution_metrics(strict_z, obs_z, baseline_dmr, obs_dmr, basin, rng),
        }
    )

    for feature_set, sub in contrib.groupby("feature_set"):
        module_vecs = {
            str(r.module_id): np.asarray([float(r.module_PC1_contribution), float(r.module_PC2_contribution), float(r.module_PC3_contribution)])
            for r in sub.itertuples()
        }
        panels: list[tuple[str, list[str], str]] = []
        for m in PRIORITY_MODULES:
            panels.append((f"single_{m}", [m], "single_module"))
        cumulative: list[str] = []
        for m in CORE_ORDER:
            cumulative.append(m)
            panels.append((f"cumulative_{'+'.join(cumulative)}", list(cumulative), "core_order_cumulative"))
        panels.append(("all_priority_modules", list(PRIORITY_MODULES), "all_modules"))
        for m in PRIORITY_MODULES:
            panels.append((f"leave_one_out_remove_{m}", [x for x in PRIORITY_MODULES if x != m], "leave_one_module_out"))
        for a, b in combinations(CORE_ORDER, 2):
            panels.append((f"pair_{a}+{b}", [a, b], "core_pair"))

        for panel_name, modules, status in panels:
            base_vec = sum((module_vecs.get(m, np.zeros(3)) for m in modules), start=np.zeros(3))
            for alpha in alpha_values():
                vec = alpha * base_vec
                pred = strict_z + vec[None, :]
                pred_dmr = decode_latent(pred, mu, sd, components)
                rows.append(
                    {
                        "feature_set": feature_set,
                        "panel_model": panel_name,
                        "included_modules": ",".join(modules),
                        "n_modules": len(modules),
                        "alpha": alpha,
                        "validation_status": status,
                        "PC1_control": float(vec[0]),
                        "PC2_control": float(vec[1]),
                        "PC3_control": float(vec[2]),
                        "direction_cosine_to_measured_correction": cosine(vec, residual_z) if alpha > 0 else np.nan,
                        "PC3_negative_pull_recovered": float(-vec[2] / (-residual_z[2])) if residual_z[2] < 0 else np.nan,
                        **distribution_metrics(pred, obs_z, pred_dmr, obs_dmr, basin, rng),
                    }
                )

        rng_perm = np.random.default_rng(20260526)
        modules = list(PRIORITY_MODULES)
        original_vals = np.asarray([module_vecs[m] for m in modules])
        for i in range(500):
            perm = rng_perm.permutation(len(modules))
            permuted_vecs = {m: original_vals[perm[j]] for j, m in enumerate(modules)}
            base_vec = sum((permuted_vecs[m] for m in CORE_ORDER), start=np.zeros(3))
            for alpha in [1.0, 1.5, 2.0]:
                vec = alpha * base_vec
                pred = strict_z + vec[None, :]
                pred_dmr = decode_latent(pred, mu, sd, components)
                random_rows.append(
                    {
                        "feature_set": feature_set,
                        "random_id": i,
                        "random_model": "permute_module_control_vectors_core_order",
                        "alpha": alpha,
                        "direction_cosine_to_measured_correction": cosine(vec, residual_z),
                        **distribution_metrics(pred, obs_z, pred_dmr, obs_dmr, basin, rng),
                    }
                )

    panel = pd.DataFrame(rows)
    random = pd.DataFrame(random_rows)
    panel.to_csv(OUT_PANEL, sep="\t", index=False)
    random.to_csv(OUT_RANDOM, sep="\t", index=False)
    return panel, random


def summarize(panel: pd.DataFrame, random: pd.DataFrame) -> pd.DataFrame:
    rows = []
    obs_occ = 0.875
    for (feature_set, panel_model), sub in panel[panel["validation_status"] != "baseline"].groupby(["feature_set", "panel_model"]):
        sub = sub.sort_values("alpha")
        best = sub.iloc[int(np.argmax(sub["pred_basin_occupancy_q90"].to_numpy(dtype=float)))]
        at1 = sub.iloc[(sub["alpha"] - 1.0).abs().argsort()].iloc[0]
        reached = sub[sub["pred_basin_occupancy_q90"] >= obs_occ]
        rnd = random[(random["feature_set"] == feature_set) & (random["alpha"] == float(at1["alpha"]))]
        rnd_p = np.nan
        if len(rnd):
            rnd_p = float((rnd["pred_basin_occupancy_q90"] >= float(at1["pred_basin_occupancy_q90"])).mean())
        rows.append(
            {
                "feature_set": feature_set,
                "panel_model": panel_model,
                "validation_status": str(at1["validation_status"]),
                "included_modules": str(at1["included_modules"]),
                "max_occupancy": float(best["pred_basin_occupancy_q90"]),
                "alpha_at_max_occupancy": float(best["alpha"]),
                "occupancy_at_alpha_1": float(at1["pred_basin_occupancy_q90"]),
                "cosine_at_alpha_1": float(at1["direction_cosine_to_measured_correction"]),
                "PC3_recovery_at_alpha_1": float(at1["PC3_negative_pull_recovered"]),
                "alpha_to_observed_occupancy": float(reached.iloc[0]["alpha"]) if len(reached) else np.nan,
                "matched_random_p_at_alpha_1": rnd_p,
            }
        )
    out = pd.DataFrame(rows).sort_values(["max_occupancy", "occupancy_at_alpha_1", "cosine_at_alpha_1"], ascending=False)
    out.to_csv(OUT_SUMMARY, sep="\t", index=False)
    return out


def make_svg(summary: pd.DataFrame) -> None:
    rows = summary[
        (summary["validation_status"].isin(["single_module", "core_order_cumulative", "leave_one_module_out", "all_modules"]))
        & (summary["feature_set"] == "ATAC_8cell_3pn_chromatin_only_inverse")
    ].head(14)
    width, height = 980, 470
    left, right, top, bottom = 100, 30, 45, 150
    plot_w = width - left - right
    plot_h = height - top - bottom
    gap = plot_w / max(len(rows), 1)
    bar_w = gap * 0.62
    lines = [
        f'<svg xmlns="http://www.w3.org/2000/svg" width="{width}" height="{height}" viewBox="0 0 {width} {height}">',
        '<rect width="100%" height="100%" fill="white"/>',
        f'<text x="{left}" y="28" font-family="Arial" font-size="18" font-weight="700">Inverse ATAC module decomposition</text>',
        f'<line x1="{left}" y1="{height-bottom}" x2="{width-right}" y2="{height-bottom}" stroke="#222"/>',
        f'<line x1="{left}" y1="{top}" x2="{left}" y2="{height-bottom}" stroke="#222"/>',
    ]
    for tick in [0.044, 0.3, 0.5, 0.875, 1.0]:
        y = height - bottom - tick * plot_h
        color = "#b23b3b" if tick in [0.044, 0.875] else "#dddddd"
        lines.append(f'<line x1="{left}" y1="{y:.2f}" x2="{width-right}" y2="{y:.2f}" stroke="{color}" stroke-dasharray="4 4"/>')
        lines.append(f'<text x="{left-8}" y="{y+4:.2f}" text-anchor="end" font-family="Arial" font-size="10">{tick:.3g}</text>')
    for i, row in enumerate(rows.itertuples()):
        x = left + i * gap + (gap - bar_w) / 2
        val = float(row.max_occupancy)
        y = height - bottom - val * plot_h
        color = "#2c6f5a" if "M02" in str(row.included_modules) else "#8a8a8a"
        lines.append(f'<rect x="{x:.2f}" y="{y:.2f}" width="{bar_w:.2f}" height="{val * plot_h:.2f}" fill="{color}"/>')
        lines.append(f'<text x="{x+bar_w/2:.2f}" y="{y-5:.2f}" text-anchor="middle" font-family="Arial" font-size="10">{val:.3f}</text>')
        label = row.panel_model.replace("_", " ")
        lines.append(f'<text x="{x+bar_w/2:.2f}" y="{height-bottom+18}" text-anchor="end" transform="rotate(-45 {x+bar_w/2:.2f} {height-bottom+18})" font-family="Arial" font-size="9">{label}</text>')
    lines.append(f'<text x="{left}" y="{height-22}" font-family="Arial" font-size="12">Green bars include M02; red guides mark methylation baseline and observed morula occupancy.</text>')
    lines.append("</svg>")
    OUT_SVG.write_text("\n".join(lines), encoding="utf-8")


def write_doc(contrib: pd.DataFrame, summary: pd.DataFrame) -> None:
    best = summary.head(12)
    contrib_sorted = contrib.sort_values(["feature_set", "module_control_norm"], ascending=[True, False])
    lines = [
        "# Inverse ATAC Module Decomposition",
        "",
        "Status: `completed`",
        "",
        "Question: whether the inverse ATAC proxy explains the full M05/M01/M12/M02 correction program, or only a local chromatin branch.",
        "",
        "## Module-Level Inference",
        "",
    ]
    for feature_set, sub in contrib_sorted.groupby("feature_set"):
        lines.append(f"### {feature_set}")
        for row in sub.itertuples():
            lines.append(
                f"- {row.module_id}: inverse_u={row.inverse_ATAC_control_z:.3f}, PC3={row.module_PC3_contribution:.3f}, "
                f"norm={row.module_control_norm:.3f}, overlap={getattr(row, 'target_overlap_fraction', np.nan):.3f}, "
                f"bg={getattr(row, 'background_overlap_fraction', np.nan):.3f}, OR={getattr(row, 'overlap_odds_ratio', np.nan):.3f}"
            )
        lines.append("")
    lines += ["## Dynamics Decomposition", ""]
    for row in best.itertuples():
        lines.append(
            f"- {row.feature_set} / {row.panel_model}: modules={row.included_modules}; "
            f"max_occ={row.max_occupancy:.3f} at alpha={row.alpha_at_max_occupancy:.2f}; "
            f"occ@1={row.occupancy_at_alpha_1:.3f}; cosine@1={row.cosine_at_alpha_1:.3f}; "
            f"PC3@1={row.PC3_recovery_at_alpha_1:.3f}; random_p@1={row.matched_random_p_at_alpha_1:.3f}"
        )
    lines += [
        "",
        "## Mechanistic Boundary",
        "",
        "The inverse ATAC proxy is strongest where residual modules are promoter-like/accessibility-linked, especially M02 and M10. It does not supply direct support for the distal/intergenic M05/M01 arm or the M12 promoter-state arm. Therefore inverse ATAC should be treated as a signed chromatin-state benchmark for an accessibility-loss branch, not as the complete biological control term.",
        "",
        "Immediate implication: the missing biological control should be split into at least two layers: an accessibility-loss/promoter branch that inverse ATAC approximates, and a histone-state branch needed to explain M05/M01/M12.",
    ]
    OUT_DOC.write_text("\n".join(lines) + "\n", encoding="utf-8")


def main() -> None:
    RESULTS.mkdir(parents=True, exist_ok=True)
    FIGURES.mkdir(parents=True, exist_ok=True)
    DOCS.mkdir(parents=True, exist_ok=True)
    contrib = feature_table()
    panel, random = evaluate_panel(contrib)
    summary = summarize(panel, random)
    make_svg(summary)
    write_doc(contrib, summary)
    OUT_MANIFEST.write_text(
        json.dumps(
            {
                "status": "completed",
                "boundary": "inverse ATAC is an orientation-corrected diagnostic proxy, not final u_bio",
                "outputs": [str(OUT_CONTRIB), str(OUT_PANEL), str(OUT_RANDOM), str(OUT_SUMMARY), str(OUT_SVG), str(OUT_DOC)],
            },
            indent=2,
        )
        + "\n",
        encoding="utf-8",
    )
    print(json.dumps({"status": "completed", "top": summary.head(8).to_dict(orient="records")}, indent=2))


if __name__ == "__main__":
    main()
```
