## Supplemental code for "An entropy-based diagnostic framework for characterizing methylation state dynamics during preimplantation development": run_jaspar_pwm_matched_background_scan.py

```
from __future__ import annotations

import json
import math
import sys
import time
from pathlib import Path

import numpy as np
import pandas as pd


BASE = Path(r"E:\CSB_TRO_operator_time_DMR_dynamics_2026-05-25")
MAIN = Path(r"C:\Users\18068\Desktop\CSB_TRO_Project_2026-05-24")
CODE = BASE / "code"
EXTERNAL = BASE / "external"
RESULTS = BASE / "results"
DOCS = BASE / "docs"
sys.path.insert(0, str(CODE))

from run_jaspar_pwm_module_scan import (  # noqa: E402
    JASPAR_MEME,
    fetch_ucsc_sequence,
    max_pwm_score,
    parse_meme,
    pwm_log_odds,
)


DMR_RANKING = RESULTS / "CSB_TRO_basin_residual_DMR_ranking.tsv"
MATCHED_REGIONS = MAIN / "input_tables" / "GSE81233_matched_non_age_window_regions_n100.tsv"
TF_DELTA = RESULTS / "CSB_TRO_GSE36552_TF_expression_delta.tsv"

OUT_SEQ = RESULTS / "CSB_TRO_priority_module_matched_background_hg19_sequences.tsv"
OUT_MOTIF = EXTERNAL / "motif" / "module_motif_enrichment.tsv"
OUT_HITS = RESULTS / "CSB_TRO_JASPAR_pwm_matched_background_hits.tsv"
OUT_DOC = DOCS / "CSB_TRO_JASPAR_pwm_matched_background_scan.md"
OUT_MANIFEST = RESULTS / "CSB_TRO_JASPAR_pwm_matched_background_scan_manifest.json"

PRIORITY_MODULES = ["M05", "M01", "M12", "M02", "M10"]
EMBRYO_TF_CANDIDATES = {
    "DUX4",
    "DUXA",
    "TPRX1",
    "TPRX2",
    "LEUTX",
    "ZSCAN4",
    "ZSCAN4C",
    "ZSCAN4D",
    "KLF17",
    "TFAP2C",
    "POU5F1",
    "SOX2",
    "NANOG",
    "TEAD4",
    "GATA3",
    "CDX2",
    "KLF4",
    "KLF5",
    "DPPA2",
    "DPPA4",
    "OTX2",
    "PRDM14",
}


def fisher_pvalue(a: int, b: int, c: int, d: int) -> float:
    def log_choose(n, k):
        if k < 0 or k > n:
            return float("-inf")
        return math.lgamma(n + 1) - math.lgamma(k + 1) - math.lgamma(n - k + 1)

    n = a + b + c + d
    row1 = a + b
    col1 = a + c
    max_x = min(row1, col1)
    logs = []
    for x in range(a, max_x + 1):
        logs.append(log_choose(col1, x) + log_choose(n - col1, row1 - x) - log_choose(n, row1))
    m = max(logs)
    return float(sum(math.exp(v - m) for v in logs) * math.exp(m))


def bh_qvalues(pvals: list[float]) -> list[float]:
    n = len(pvals)
    order = np.argsort(pvals)
    q = np.empty(n)
    prev = 1.0
    for rank, idx in enumerate(order[::-1], start=1):
        i = n - rank + 1
        val = min(prev, pvals[idx] * n / i)
        q[idx] = val
        prev = val
    return q.tolist()


def load_or_fetch_sequences(n_sets: int) -> pd.DataFrame:
    if OUT_SEQ.exists():
        seqs = pd.read_csv(OUT_SEQ, sep="\t")
        if seqs["control_set"].nunique() >= n_sets + 1:
            return seqs

    dmr = pd.read_csv(DMR_RANKING, sep="\t")
    fg = dmr[dmr["module_id"].isin(PRIORITY_MODULES)].copy()
    bg = pd.read_csv(MATCHED_REGIONS, sep="\t")
    bg = bg[bg["region_type"] == "matched_non_age_window"].copy()
    control_sets = sorted(bg["control_set"].unique())[:n_sets]
    bg = bg[(bg["control_set"].isin(control_sets)) & (bg["matched_age_cluster"].isin(fg["cluster_name"]))].copy()

    rows = []
    total = len(fg) + len(bg)
    combined = []
    for row in fg.itertuples(index=False):
        combined.append(
            {
                "set_type": "foreground",
                "control_set": "age_DMR",
                "matched_age_cluster": row.cluster_name,
                "cluster_name": row.cluster_name,
                "module_id": row.module_id,
                "chr": row.chr,
                "start": int(row.start),
                "end": int(row.end),
            }
        )
    for row in bg.itertuples(index=False):
        module = fg.loc[fg["cluster_name"] == row.matched_age_cluster, "module_id"].iloc[0]
        combined.append(
            {
                "set_type": "matched_background",
                "control_set": row.control_set,
                "matched_age_cluster": row.matched_age_cluster,
                "cluster_name": row.cluster_name,
                "module_id": module,
                "chr": row.chr,
                "start": int(row.start),
                "end": int(row.end),
            }
        )
    for i, item in enumerate(combined, 1):
        seq = fetch_ucsc_sequence(item["chr"], item["start"], item["end"])
        item["sequence"] = seq
        item["width"] = item["end"] - item["start"]
        item["gc_fraction"] = (seq.count("G") + seq.count("C")) / len(seq) if seq else np.nan
        rows.append(item)
        if i % 50 == 0:
            print(f"Fetched {i}/{total} matched-background sequences")
            time.sleep(0.2)
    out = pd.DataFrame(rows)
    out.to_csv(OUT_SEQ, sep="\t", index=False)
    return out


def main() -> None:
    n_sets = 10
    quantile = 0.95
    RESULTS.mkdir(parents=True, exist_ok=True)
    DOCS.mkdir(parents=True, exist_ok=True)
    EXTERNAL.joinpath("motif").mkdir(parents=True, exist_ok=True)

    seqs = load_or_fetch_sequences(n_sets)
    tf_delta = pd.read_csv(TF_DELTA, sep="\t")
    expressed_tfs = set(tf_delta["TF"].astype(str).str.upper())
    tf_delta["TF_upper"] = tf_delta["TF"].astype(str).str.upper()
    top_delta_tfs = set(
        tf_delta.dropna(subset=["delta_8cell_to_morula_zscore"])
        .assign(abs_delta=lambda x: pd.to_numeric(x["delta_8cell_to_morula_zscore"], errors="coerce").abs())
        .sort_values("abs_delta", ascending=False)
        .head(150)["TF_upper"]
    )
    target_tfs = expressed_tfs & (top_delta_tfs | EMBRYO_TF_CANDIDATES)
    motifs = [m for m in parse_meme(JASPAR_MEME) if m["TF"] in target_tfs]

    rows = []
    hit_rows = []
    for mi, motif in enumerate(motifs, 1):
        lod = pwm_log_odds(motif["pwm"])
        scores = []
        for row in seqs.itertuples(index=False):
            scores.append(max_pwm_score(str(row.sequence), lod))
        score_series = pd.Series(scores, index=seqs.index)
        for module_id in PRIORITY_MODULES:
            fg_mask = (seqs["module_id"] == module_id) & (seqs["set_type"] == "foreground")
            bg_mask = (seqs["module_id"] == module_id) & (seqs["set_type"] == "matched_background")
            fg_scores = score_series[fg_mask]
            bg_scores = score_series[bg_mask]
            if len(fg_scores) == 0 or len(bg_scores) == 0:
                continue
            threshold = float(np.quantile(bg_scores, quantile))
            fg_hits = fg_scores >= threshold
            bg_hits = bg_scores >= threshold
            a = int(fg_hits.sum())
            b = int(len(fg_hits) - a)
            c = int(bg_hits.sum())
            d = int(len(bg_hits) - c)
            if a == 0 and c == 0:
                continue
            p = fisher_pvalue(a, b, c, d)
            log_or = math.log2(((a + 0.5) / (b + 0.5)) / ((c + 0.5) / (d + 0.5)))
            rows.append(
                {
                    "module_id": module_id,
                    "TF": motif["TF"],
                    "motif_database": f"JASPAR2024_CORE_vertebrates_PWM_matched_bg_q{quantile}",
                    "motif_id": motif["motif_id"],
                    "motif_name": motif["motif_name"],
                    "log_odds_ratio": log_or,
                    "pvalue": p,
                    "motif_hit_count": a,
                    "foreground_nonhit_count": b,
                    "background_hit_count": c,
                    "background_nonhit_count": d,
                    "threshold": threshold,
                    "n_matched_background_sets": n_sets,
                }
            )
            if a:
                for idx in fg_scores[fg_hits].index:
                    hit_rows.append(
                        {
                            "module_id": module_id,
                            "cluster_name": seqs.loc[idx, "cluster_name"],
                            "TF": motif["TF"],
                            "motif_id": motif["motif_id"],
                            "score": float(score_series.loc[idx]),
                            "threshold": threshold,
                        }
                    )
        if mi % 100 == 0:
            print(f"Scanned {mi}/{len(motifs)} expressed motifs with matched background")

    enrich = pd.DataFrame(rows)
    if len(enrich):
        enrich["qvalue"] = bh_qvalues(enrich["pvalue"].astype(float).tolist())
        enrich = enrich.sort_values(["qvalue", "pvalue", "module_id"])
    enrich.to_csv(OUT_MOTIF, sep="\t", index=False)
    pd.DataFrame(hit_rows).to_csv(OUT_HITS, sep="\t", index=False)

    OUT_DOC.write_text(
        "\n".join(
            [
                "# JASPAR PWM matched-background scan",
                "",
                "This scan uses per-DMR matched non-age windows from GSE81233 as background for priority residual modules.",
                "",
                f"Priority module sequences: {int((seqs['set_type'] == 'foreground').sum())}",
                f"Matched background sequences: {int((seqs['set_type'] == 'matched_background').sum())}",
                f"Matched control sets per DMR: {n_sets}",
                f"Expressed motifs scanned: {len(motifs)}",
                f"Enrichment rows: {len(enrich)}",
                "",
                "Boundary: this remains a Python PWM burden approximation. HOMER/FIMO with explicit GC/CpG matching is still preferred for final claims.",
            ]
        )
        + "\n",
        encoding="utf-8",
    )
    OUT_MANIFEST.write_text(
        json.dumps(
            {
                "matched_regions": str(MATCHED_REGIONS),
                "n_sets": n_sets,
                "threshold_quantile": quantile,
                "n_sequences": int(len(seqs)),
                "n_motifs": int(len(motifs)),
                "n_enrichment_rows": int(len(enrich)),
                "outputs": [str(OUT_SEQ), str(OUT_MOTIF), str(OUT_HITS), str(OUT_DOC)],
            },
            indent=2,
        )
        + "\n",
        encoding="utf-8",
    )
    print(json.dumps({"n_sequences": int(len(seqs)), "n_motifs": int(len(motifs)), "n_enrichment_rows": int(len(enrich))}, indent=2))


if __name__ == "__main__":
    main()
```
