## Supplemental code for "An entropy-based diagnostic framework for characterizing methylation state dynamics during preimplantation development": run_jaspar_pwm_module_scan.py

```
from __future__ import annotations

import json
import math
import random
import re
import time
from pathlib import Path
from urllib.request import urlopen
from xml.etree import ElementTree as ET

import numpy as np
import pandas as pd


BASE = Path(r"E:\CSB_TRO_operator_time_DMR_dynamics_2026-05-25")
EXTERNAL = BASE / "external"
RESULTS = BASE / "results"
DOCS = BASE / "docs"

DMR_RANKING = RESULTS / "CSB_TRO_basin_residual_DMR_ranking.tsv"
JASPAR_MEME = EXTERNAL / "motif" / "jaspar_vertebrates.meme"
OUT_SEQ = RESULTS / "CSB_TRO_residual_DMR_hg19_sequences.tsv"
OUT_MOTIF = EXTERNAL / "motif" / "module_motif_enrichment.tsv"
OUT_HITS = RESULTS / "CSB_TRO_JASPAR_pwm_DMR_hits.tsv"
OUT_DOC = DOCS / "CSB_TRO_JASPAR_pwm_module_scan.md"
OUT_MANIFEST = RESULTS / "CSB_TRO_JASPAR_pwm_module_scan_manifest.json"

PRIORITY_MODULES = ["M05", "M01", "M12", "M02", "M10"]
NUC_INDEX = {"A": 0, "C": 1, "G": 2, "T": 3}


def fetch_ucsc_sequence(chrom: str, start0: int, end0: int, retries: int = 3) -> str:
    # UCSC DAS uses 1-based inclusive coordinates.
    start1 = int(start0) + 1
    end1 = int(end0)
    url = f"https://genome.ucsc.edu/cgi-bin/das/hg19/dna?segment={chrom}:{start1},{end1}"
    for attempt in range(retries):
        try:
            with urlopen(url, timeout=30) as handle:
                xml = handle.read()
            root = ET.fromstring(xml)
            dna = "".join(root.itertext())
            dna = re.sub(r"[^ACGTacgt]", "", dna).upper()
            return dna
        except Exception:
            if attempt == retries - 1:
                raise
            time.sleep(1 + attempt)
    return ""


def load_or_fetch_sequences() -> pd.DataFrame:
    if OUT_SEQ.exists():
        return pd.read_csv(OUT_SEQ, sep="\t")
    dmr = pd.read_csv(DMR_RANKING, sep="\t")
    rows = []
    for i, row in enumerate(dmr.itertuples(index=False), 1):
        seq = fetch_ucsc_sequence(row.chr, int(row.start), int(row.end))
        rows.append(
            {
                "cluster_name": row.cluster_name,
                "module_id": row.module_id,
                "chr": row.chr,
                "start": int(row.start),
                "end": int(row.end),
                "width": int(row.end) - int(row.start),
                "sequence": seq,
                "gc_fraction": (seq.count("G") + seq.count("C")) / len(seq) if seq else np.nan,
            }
        )
        if i % 25 == 0:
            print(f"Fetched {i} sequences")
    out = pd.DataFrame(rows)
    out.to_csv(OUT_SEQ, sep="\t", index=False)
    return out


def parse_meme(path: Path) -> list[dict]:
    motifs = []
    current = None
    matrix = []
    width = None
    with path.open("r", encoding="utf-8", errors="replace") as handle:
        for line in handle:
            line = line.strip()
            if line.startswith("MOTIF "):
                if current and matrix:
                    current["pwm"] = np.array(matrix, dtype=float)
                    current["width"] = width or len(matrix)
                    motifs.append(current)
                parts = line.split(maxsplit=2)
                motif_id = parts[1]
                tf = parts[2] if len(parts) > 2 else motif_id
                current = {"motif_id": motif_id, "TF": re.split(r"::|\\s+", tf)[0].upper(), "motif_name": tf}
                matrix = []
                width = None
            elif line.startswith("letter-probability matrix"):
                m = re.search(r"w=\\s*(\\d+)", line)
                width = int(m.group(1)) if m else None
            elif current and re.match(r"^[0-9.eE+\- ]+$", line):
                vals = [float(x) for x in line.split()]
                if len(vals) == 4:
                    matrix.append(vals)
    if current and matrix:
        current["pwm"] = np.array(matrix, dtype=float)
        current["width"] = width or len(matrix)
        motifs.append(current)
    return motifs


def revcomp(seq: str) -> str:
    return seq.translate(str.maketrans("ACGT", "TGCA"))[::-1]


def pwm_log_odds(pwm: np.ndarray) -> np.ndarray:
    pwm = np.clip(pwm, 1e-4, 1.0)
    pwm = pwm / pwm.sum(axis=1, keepdims=True)
    return np.log2(pwm / 0.25)


def max_pwm_score(seq: str, lod: np.ndarray) -> float:
    w = lod.shape[0]
    if len(seq) < w:
        return float("-inf")
    best = float("-inf")
    for strand in [seq, revcomp(seq)]:
        for i in range(0, len(strand) - w + 1):
            kmer = strand[i : i + w]
            if any(ch not in NUC_INDEX for ch in kmer):
                continue
            score = sum(lod[j, NUC_INDEX[ch]] for j, ch in enumerate(kmer))
            if score > best:
                best = score
    return best


def threshold_for_motif(lod: np.ndarray, rng: random.Random, n: int = 1500, quantile: float = 0.995) -> float:
    w = lod.shape[0]
    scores = []
    alphabet = "ACGT"
    for _ in range(n):
        seq = "".join(rng.choice(alphabet) for _ in range(w))
        scores.append(max_pwm_score(seq, lod))
    return float(np.quantile(scores, quantile))


def fisher_pvalue(a: int, b: int, c: int, d: int) -> float:
    # One-sided enrichment p-value P[X >= a] for fixed margins.
    def log_choose(n, k):
        if k < 0 or k > n:
            return float("-inf")
        return math.lgamma(n + 1) - math.lgamma(k + 1) - math.lgamma(n - k + 1)

    n = a + b + c + d
    row1 = a + b
    col1 = a + c
    min_x = max(0, row1 - (n - col1))
    max_x = min(row1, col1)
    logs = []
    for x in range(a, max_x + 1):
        logs.append(log_choose(col1, x) + log_choose(n - col1, row1 - x) - log_choose(n, row1))
    m = max(logs)
    return float(sum(math.exp(v - m) for v in logs) * math.exp(m))


def bh_qvalues(pvals: list[float]) -> list[float]:
    n = len(pvals)
    order = np.argsort(pvals)
    q = np.empty(n)
    prev = 1.0
    for rank, idx in enumerate(order[::-1], start=1):
        i = n - rank + 1
        val = min(prev, pvals[idx] * n / i)
        q[idx] = val
        prev = val
    return q.tolist()


def main() -> None:
    RESULTS.mkdir(parents=True, exist_ok=True)
    DOCS.mkdir(parents=True, exist_ok=True)
    EXTERNAL.joinpath("motif").mkdir(parents=True, exist_ok=True)

    seqs = load_or_fetch_sequences()
    motifs = parse_meme(JASPAR_MEME)
    # Keep motifs whose TF has RNA expression delta. This reduces runtime and focuses on testable TF controls.
    tf_delta = pd.read_csv(RESULTS / "CSB_TRO_GSE36552_TF_expression_delta.tsv", sep="\t")
    expressed_tfs = set(tf_delta["TF"].astype(str).str.upper())
    motifs = [m for m in motifs if m["TF"] in expressed_tfs]

    rng = random.Random(20260525)
    rows = []
    hit_rows = []
    for mi, motif in enumerate(motifs, 1):
        lod = pwm_log_odds(motif["pwm"])
        threshold = threshold_for_motif(lod, rng)
        hit_map = {}
        for row in seqs.itertuples(index=False):
            score = max_pwm_score(str(row.sequence), lod)
            hit = bool(score >= threshold)
            hit_map[row.cluster_name] = hit
            if hit:
                hit_rows.append(
                    {
                        "cluster_name": row.cluster_name,
                        "module_id": row.module_id,
                        "TF": motif["TF"],
                        "motif_id": motif["motif_id"],
                        "max_log_odds_score": score,
                        "threshold": threshold,
                    }
                )
        for module_id in PRIORITY_MODULES:
            foreground = seqs[seqs["module_id"] == module_id]
            background = seqs[seqs["module_id"] != module_id]
            a = int(sum(hit_map[x] for x in foreground["cluster_name"]))
            b = int(len(foreground) - a)
            c = int(sum(hit_map[x] for x in background["cluster_name"]))
            d = int(len(background) - c)
            if a == 0 and c == 0:
                continue
            p = fisher_pvalue(a, b, c, d)
            log_or = math.log2(((a + 0.5) / (b + 0.5)) / ((c + 0.5) / (d + 0.5)))
            rows.append(
                {
                    "module_id": module_id,
                    "TF": motif["TF"],
                    "motif_database": "JASPAR2024_CORE_vertebrates_PWM_lightscan",
                    "motif_id": motif["motif_id"],
                    "motif_name": motif["motif_name"],
                    "log_odds_ratio": log_or,
                    "pvalue": p,
                    "motif_hit_count": a,
                    "foreground_nonhit_count": b,
                    "background_hit_count": c,
                    "background_nonhit_count": d,
                    "threshold": threshold,
                }
            )
        if mi % 100 == 0:
            print(f"Scanned {mi}/{len(motifs)} expressed motifs")

    enrich = pd.DataFrame(rows)
    if len(enrich):
        enrich["qvalue"] = bh_qvalues(enrich["pvalue"].astype(float).tolist())
        enrich = enrich.sort_values(["qvalue", "pvalue", "module_id"])
    enrich.to_csv(OUT_MOTIF, sep="\t", index=False)
    pd.DataFrame(hit_rows).to_csv(OUT_HITS, sep="\t", index=False)

    lines = [
        "# JASPAR PWM module scan",
        "",
        "This is a lightweight Python PWM scan against UCSC hg19 DMR sequences. It is a first-pass motif burden screen, not a replacement for HOMER/FIMO with matched GC/CpG background.",
        "",
        f"DMR sequences: {len(seqs)}",
        f"Expressed JASPAR motifs scanned: {len(motifs)}",
        f"Module motif enrichment rows: {len(enrich)}",
    ]
    OUT_DOC.write_text("\n".join(lines) + "\n", encoding="utf-8")
    OUT_MANIFEST.write_text(
        json.dumps(
            {
                "sequence_source": "UCSC DAS hg19",
                "jaspar_meme": str(JASPAR_MEME),
                "n_DMR_sequences": int(len(seqs)),
                "n_expressed_motifs_scanned": int(len(motifs)),
                "n_enrichment_rows": int(len(enrich)),
                "outputs": [str(OUT_SEQ), str(OUT_MOTIF), str(OUT_HITS), str(OUT_DOC)],
                "boundary": "First-pass PWM burden only; follow with HOMER/FIMO and matched GC/CpG background.",
            },
            indent=2,
        )
        + "\n",
        encoding="utf-8",
    )
    print(json.dumps({"n_sequences": int(len(seqs)), "n_motifs": int(len(motifs)), "n_enrichment_rows": int(len(enrich))}, indent=2))


if __name__ == "__main__":
    main()
```
