## Supplemental code for "An entropy-based diagnostic framework for characterizing methylation state dynamics during preimplantation development": run_missing_control_term_decomposition.py

````
from __future__ import annotations

import argparse
import json
import sys
from pathlib import Path

import numpy as np
import pandas as pd


BASE = Path(r"E:\CSB_TRO_operator_time_DMR_dynamics_2026-05-25")
CODE = BASE / "code"
RESULTS = BASE / "results"
FIGURES = BASE / "figures"
DOCS = BASE / "docs"
sys.path.insert(0, str(CODE))

from run_basin_residual_control_field import (  # noqa: E402
    PRE_MORULA_STAGES,
    cosine,
    encode_delta_to_latent,
    simulate_strict_pre_morula,
)
from run_morula_basin_sde import (  # noqa: E402
    TAU,
    basin_definition,
    decode_latent,
    distribution_metrics,
    fit_latent_basis,
    fit_operator,
    load_inputs,
    stage_ids,
)


def control_from_ids(matrix, sd, components, dmr_table, ids):
    delta = np.zeros(matrix.shape[1], dtype=float)
    by_id = dmr_table.set_index("cluster_name")
    idx = [list(matrix.columns).index(c) for c in ids]
    delta[idx] = by_id.loc[ids, "latent_residual_delta_beta"].to_numpy(dtype=float)
    return encode_delta_to_latent(delta, sd, components), delta


def rmse_vec(a, b):
    return float(np.sqrt(np.mean((np.asarray(a) - np.asarray(b)) ** 2)))


def solve_ridge_basis(basis: np.ndarray, target: np.ndarray, lam: float = 1e-3):
    # basis columns are candidate controls in latent space.
    x = np.asarray(basis, dtype=float)
    y = np.asarray(target, dtype=float)
    return np.linalg.solve(x.T @ x + lam * np.eye(x.shape[1]), x.T @ y)


def evaluate(name, pred_z, obs_z, obs_dmr, mu, sd, components, basin, meta, seed):
    pred_dmr = decode_latent(pred_z, mu, sd, components)
    metrics = distribution_metrics(pred_z, obs_z, pred_dmr, obs_dmr, basin, np.random.default_rng(seed))
    return {"control_term_model": name, **meta, **metrics}


def make_module_basis(matrix, sd, components, dmr_table):
    rows = []
    basis = []
    module_ids = []
    for module_id, sub in dmr_table.groupby("module_id"):
        ids = sub["cluster_name"].tolist()
        z, _ = control_from_ids(matrix, sd, components, dmr_table, ids)
        rows.append({
            "module_id": module_id,
            "n_DMRs": len(ids),
            "latent_control_PC1": float(z[0]),
            "latent_control_PC2": float(z[1]),
            "latent_control_PC3": float(z[2]),
            "latent_control_norm": float(np.linalg.norm(z)),
        })
        basis.append(z)
        module_ids.append(module_id)
    return pd.DataFrame(rows), np.vstack(basis).T, module_ids


def greedy_modules(matrix, sd, components, dmr_table, residual_z, max_modules=8):
    remaining = list(dmr_table["module_id"].dropna().unique())
    selected = []
    rows = []
    current = np.zeros_like(residual_z)
    for step in range(1, max_modules + 1):
        best = None
        for module_id in remaining:
            ids = dmr_table[dmr_table["module_id"] == module_id]["cluster_name"].tolist()
            z, _ = control_from_ids(matrix, sd, components, dmr_table, ids)
            cand = current + z
            score = rmse_vec(cand, residual_z)
            if best is None or score < best["rmse"]:
                best = {"module_id": module_id, "z": z, "rmse": score, "cand": cand}
        selected.append(best["module_id"])
        remaining.remove(best["module_id"])
        current = best["cand"]
        rows.append({
            "step": step,
            "selected_module": best["module_id"],
            "selected_modules": ",".join(map(str, selected)),
            "latent_reconstruction_rmse": best["rmse"],
            "latent_reconstruction_cosine": cosine(current, residual_z),
            "latent_control_norm": float(np.linalg.norm(current)),
            "PC1": float(current[0]),
            "PC2": float(current[1]),
            "PC3": float(current[2]),
        })
    return pd.DataFrame(rows)


def make_svg(path: Path, amplitude_df: pd.DataFrame):
    rows = amplitude_df.copy()
    width, height = 820, 390
    left, right, top, bottom = 70, 25, 40, 70
    plot_w = width - left - right
    plot_h = height - top - bottom
    out = [
        f'<svg xmlns="http://www.w3.org/2000/svg" width="{width}" height="{height}" viewBox="0 0 {width} {height}">',
        '<rect width="100%" height="100%" fill="white"/>',
        f'<text x="{left}" y="26" font-family="Arial" font-size="18" font-weight="700">Missing control term amplitude scan</text>',
        f'<line x1="{left}" y1="{height-bottom}" x2="{width-right}" y2="{height-bottom}" stroke="#222"/>',
        f'<line x1="{left}" y1="{top}" x2="{left}" y2="{height-bottom}" stroke="#222"/>',
    ]
    for tick in [0, 0.25, 0.5, 0.75, 0.875, 1.0]:
        y = height - bottom - tick * plot_h
        color = "#b33" if abs(tick - 0.875) < 1e-6 else "#ddd"
        out.append(f'<line x1="{left}" y1="{y:.2f}" x2="{width-right}" y2="{y:.2f}" stroke="{color}" stroke-dasharray="4 4"/>')
        out.append(f'<text x="{left-8}" y="{y+4:.2f}" text-anchor="end" font-family="Arial" font-size="11">{tick:.3g}</text>')
    pts = []
    xmin, xmax = rows["control_amplitude_alpha"].min(), rows["control_amplitude_alpha"].max()
    for row in rows.itertuples():
        x = left + (row.control_amplitude_alpha - xmin) / (xmax - xmin) * plot_w
        y = height - bottom - float(row.pred_basin_occupancy_q90) * plot_h
        pts.append((x, y))
    out.append('<polyline points="' + " ".join(f"{x:.2f},{y:.2f}" for x, y in pts) + '" fill="none" stroke="#2f6f8f" stroke-width="2.5"/>')
    for x, y in pts:
        out.append(f'<circle cx="{x:.2f}" cy="{y:.2f}" r="3.5" fill="#2f6f8f"/>')
    for alpha in rows["control_amplitude_alpha"]:
        x = left + (alpha - xmin) / (xmax - xmin) * plot_w
        out.append(f'<text x="{x:.2f}" y="{height-bottom+18}" text-anchor="middle" font-family="Arial" font-size="10">{alpha:g}</text>')
    out.append(f'<text x="{left}" y="{height-20}" font-family="Arial" font-size="12">x-axis: fraction of measured correction term added during 8-cell to morula. Red guide: observed occupancy 0.875.</text>')
    out.append("</svg>")
    path.write_text("\n".join(out), encoding="utf-8")


def write_doc(path: Path, summary: dict, amplitude_df: pd.DataFrame, module_fit: pd.DataFrame, greedy_eval: pd.DataFrame):
    best_amp = amplitude_df.iloc[(amplitude_df["pred_basin_occupancy_q90"] - summary["observed_basin_occupancy_q90"]).abs().argsort()].iloc[0]
    top_module = module_fit.sort_values("abs_ridge_weight", ascending=False).head(5)
    lines = [
        "# Missing control-term decomposition",
        "",
        "This analysis formalizes the distinction in the current model state: we have not discovered a biological u_bio yet, but we have measured the correction term that such a u_bio must explain.",
        "",
        "Model form:",
        "",
        "```text",
        "dz_meth/dtau = f_meth(z, tau) + B u_bio(tau)",
        "```",
        "",
        f"Measured correction vector over 8-cell to morula: {', '.join(f'{v:.4f}' for v in summary['missing_control_delta_z'])}",
        f"Equivalent correction velocity: {', '.join(f'{v:.4f}' for v in summary['missing_control_velocity'])}",
        f"Baseline strict occupancy: {summary['strict_predicted_basin_occupancy_q90']:.3f}",
        f"Observed occupancy target: {summary['observed_basin_occupancy_q90']:.3f}",
        "",
        f"Closest amplitude to observed occupancy: alpha={best_amp.control_amplitude_alpha:.3f}, occupancy={best_amp.pred_basin_occupancy_q90:.3f}",
        "",
        "Largest module weights in ridge reconstruction of the correction term:",
    ]
    for row in top_module.itertuples():
        lines.append(f"- {row.module_id}: weight={row.ridge_weight:.4f}, abs_weight={row.abs_ridge_weight:.4f}, n_DMRs={row.n_DMRs}")
    lines.extend(["", "Greedy module reconstruction:"])
    for row in greedy_eval.head(8).itertuples():
        lines.append(f"- step {row.step}, modules={row.selected_modules}: occupancy={row.pred_basin_occupancy_q90:.3f}, cosine={row.latent_reconstruction_cosine:.3f}")
    lines.extend([
        "",
        "Interpretation boundary: this is not a non-leaking biological mechanism yet. It defines the correction term that external RNA, ATAC, histone, motif, or chromatin variables must predict in the next stage.",
    ])
    path.write_text("\n".join(lines) + "\n", encoding="utf-8")


def main():
    parser = argparse.ArgumentParser()
    parser.add_argument("--q", type=int, default=3)
    parser.add_argument("--lambda", dest="lam", type=float, default=1000.0)
    parser.add_argument("--n-steps", type=int, default=12)
    parser.add_argument("--seed", type=int, default=20260525)
    args = parser.parse_args()

    RESULTS.mkdir(parents=True, exist_ok=True)
    FIGURES.mkdir(parents=True, exist_ok=True)
    DOCS.mkdir(parents=True, exist_ok=True)

    matrix, ann, pairs = load_inputs()
    mu, sd, components, score_df = fit_latent_basis(matrix, args.q)
    coef, _, train = fit_operator(score_df, pairs, train_stages=set(PRE_MORULA_STAGES), lam=args.lam)
    strict_pred_z = simulate_strict_pre_morula(score_df, ann, coef, args.n_steps)
    obs_z = score_df.loc[stage_ids(ann, "morula")].to_numpy(dtype=float)
    obs_dmr = matrix.loc[stage_ids(ann, "morula")].to_numpy(dtype=float)
    basin = basin_definition(obs_z)

    pred_center = strict_pred_z.mean(axis=0)
    obs_center = obs_z.mean(axis=0)
    residual_z = obs_center - pred_center
    dt = TAU["morula"] - TAU["8-cell"]
    control_velocity = residual_z / dt

    dmr_table = pd.read_csv(RESULTS / "CSB_TRO_basin_residual_DMR_ranking.tsv", sep="\t")
    module_basis_df, basis, module_ids = make_module_basis(matrix, sd, components, dmr_table)
    weights = solve_ridge_basis(basis, residual_z)
    module_fit = module_basis_df.copy()
    module_fit["ridge_weight"] = weights
    module_fit["abs_ridge_weight"] = np.abs(weights)
    module_fit["weighted_PC1"] = module_fit["latent_control_PC1"] * module_fit["ridge_weight"]
    module_fit["weighted_PC2"] = module_fit["latent_control_PC2"] * module_fit["ridge_weight"]
    module_fit["weighted_PC3"] = module_fit["latent_control_PC3"] * module_fit["ridge_weight"]
    reconstructed_z = basis @ weights

    rows = []
    for alpha in [0.0, 0.1, 0.25, 0.5, 0.75, 0.875, 1.0, 1.125, 1.25]:
        pred_z = strict_pred_z + alpha * residual_z[None, :]
        rows.append(evaluate(
            f"measured_control_alpha_{alpha:g}",
            pred_z,
            obs_z,
            obs_dmr,
            mu,
            sd,
            components,
            basin,
            {
                "control_amplitude_alpha": alpha,
                "control_term_type": "measured_total_correction",
                "uses_morula_methylation_residual": "yes_diagnostic",
                "latent_control_norm": float(np.linalg.norm(alpha * residual_z)),
                "latent_control_cosine_to_missing_control": cosine(alpha * residual_z, residual_z) if alpha != 0 else 0.0,
            },
            args.seed + int(alpha * 1000),
        ))
    amplitude_df = pd.DataFrame(rows)

    greedy = greedy_modules(matrix, sd, components, dmr_table, residual_z, max_modules=8)
    greedy_rows = []
    for row in greedy.itertuples():
        pred_z = strict_pred_z + np.asarray([row.PC1, row.PC2, row.PC3])[None, :]
        greedy_rows.append(evaluate(
            f"greedy_module_step_{row.step}",
            pred_z,
            obs_z,
            obs_dmr,
            mu,
            sd,
            components,
            basin,
            {
                "step": row.step,
                "selected_modules": row.selected_modules,
                "latent_reconstruction_rmse": row.latent_reconstruction_rmse,
                "latent_reconstruction_cosine": row.latent_reconstruction_cosine,
                "latent_control_norm": row.latent_control_norm,
                "uses_morula_methylation_residual": "yes_diagnostic",
            },
            args.seed + 100 + row.step,
        ))
    greedy_eval = pd.DataFrame(greedy_rows)

    recon_eval = pd.DataFrame([evaluate(
        "ridge_module_reconstruction",
        strict_pred_z + reconstructed_z[None, :],
        obs_z,
        obs_dmr,
        mu,
        sd,
        components,
        basin,
        {
            "latent_reconstruction_rmse": rmse_vec(reconstructed_z, residual_z),
            "latent_reconstruction_cosine": cosine(reconstructed_z, residual_z),
            "latent_control_norm": float(np.linalg.norm(reconstructed_z)),
            "uses_morula_methylation_residual": "yes_diagnostic",
        },
        args.seed + 999,
    )])

    summary = {
        "q": args.q,
        "ridge_lambda": args.lam,
        "n_pre_morula_training_pairs": int(len(train)),
        "strict_predicted_center": pred_center.tolist(),
        "observed_morula_center": obs_center.tolist(),
        "missing_control_delta_z": residual_z.tolist(),
        "missing_control_velocity": control_velocity.tolist(),
        "missing_control_norm": float(np.linalg.norm(residual_z)),
        "delta_tau_8cell_to_morula": float(dt),
        "strict_predicted_basin_occupancy_q90": float(np.mean(np.linalg.norm(strict_pred_z - obs_center[None, :], axis=1) <= basin["radius_q90"])),
        "observed_basin_occupancy_q90": float(basin["observed_occupancy_q90"]),
        "module_ridge_reconstruction_rmse": rmse_vec(reconstructed_z, residual_z),
        "module_ridge_reconstruction_cosine": cosine(reconstructed_z, residual_z),
    }

    pd.DataFrame([summary]).to_csv(RESULTS / "CSB_TRO_missing_control_term_summary.tsv", sep="\t", index=False)
    amplitude_df.to_csv(RESULTS / "CSB_TRO_missing_control_term_amplitude_scan.tsv", sep="\t", index=False)
    module_fit.sort_values("abs_ridge_weight", ascending=False).to_csv(RESULTS / "CSB_TRO_missing_control_term_module_basis.tsv", sep="\t", index=False)
    greedy_eval.to_csv(RESULTS / "CSB_TRO_missing_control_term_greedy_modules.tsv", sep="\t", index=False)
    recon_eval.to_csv(RESULTS / "CSB_TRO_missing_control_term_reconstruction_metrics.tsv", sep="\t", index=False)
    make_svg(FIGURES / "CSB_TRO_missing_control_term_amplitude_scan.svg", amplitude_df)
    write_doc(DOCS / "CSB_TRO_missing_control_term_interpretation.md", summary, amplitude_df, module_fit, greedy_eval)
    manifest = {
        **summary,
        "outputs": [
            str(RESULTS / "CSB_TRO_missing_control_term_summary.tsv"),
            str(RESULTS / "CSB_TRO_missing_control_term_amplitude_scan.tsv"),
            str(RESULTS / "CSB_TRO_missing_control_term_module_basis.tsv"),
            str(RESULTS / "CSB_TRO_missing_control_term_greedy_modules.tsv"),
            str(RESULTS / "CSB_TRO_missing_control_term_reconstruction_metrics.tsv"),
            str(FIGURES / "CSB_TRO_missing_control_term_amplitude_scan.svg"),
            str(DOCS / "CSB_TRO_missing_control_term_interpretation.md"),
        ],
    }
    (RESULTS / "CSB_TRO_missing_control_term_manifest.json").write_text(json.dumps(manifest, indent=2), encoding="utf-8")
    print(json.dumps({
        "summary": summary,
        "amplitude_scan": amplitude_df[["control_amplitude_alpha", "pred_basin_occupancy_q90", "pred_mean_distance_to_morula_centroid", "dmr_mean_rmse"]].to_dict(orient="records"),
        "top_module_basis": module_fit.sort_values("abs_ridge_weight", ascending=False).head(8).to_dict(orient="records"),
        "greedy_modules": greedy_eval[["step", "selected_modules", "pred_basin_occupancy_q90", "latent_reconstruction_cosine"]].head(8).to_dict(orient="records"),
    }, indent=2))


if __name__ == "__main__":
    main()
````
