## Supplemental code for "An entropy-based diagnostic framework for characterizing methylation state dynamics during preimplantation development": run_module_latent_operator_time_dynamics.py

```
from __future__ import annotations

import csv
import json
import math
from pathlib import Path

import numpy as np
import pandas as pd


BASE = Path(r"E:\CSB_TRO_operator_time_DMR_dynamics_2026-05-25")
RESULTS = BASE / "results"
FIGURES = BASE / "figures"
DOCS = BASE / "docs"

STAGES = ["MII oocyte", "zygote/PN", "2-cell", "4-cell", "8-cell", "morula", "blastocyst"]
TAU = {stage: i / (len(STAGES) - 1) for i, stage in enumerate(STAGES)}


def rmse(a, b) -> float:
    a = np.asarray(a, dtype=float)
    b = np.asarray(b, dtype=float)
    return float(np.sqrt(np.mean((a - b) ** 2)))


def corr(a, b) -> float:
    a = np.asarray(a, dtype=float)
    b = np.asarray(b, dtype=float)
    aa = a - a.mean()
    bb = b - b.mean()
    den = math.sqrt(float((aa @ aa) * (bb @ bb)))
    return float((aa @ bb) / den) if den else float("nan")


def ridge(x: np.ndarray, y: np.ndarray, w: np.ndarray, lam: float) -> np.ndarray:
    sw = np.sqrt(np.maximum(w, 1e-12) / np.mean(np.maximum(w, 1e-12)))
    xw = x * sw[:, None]
    if y.ndim == 1:
        yw = y * sw
    else:
        yw = y * sw[:, None]
    penalty = lam * np.eye(x.shape[1])
    penalty[0, 0] = 0.0
    return np.linalg.solve(xw.T @ xw + penalty, xw.T @ yw)


def kmeans(x: np.ndarray, k: int, seed: int = 1, max_iter: int = 100) -> np.ndarray:
    rng = np.random.default_rng(seed)
    centers = x[rng.choice(len(x), k, replace=False)].copy()
    labels = np.zeros(len(x), dtype=int)
    for _ in range(max_iter):
        dist = ((x[:, None, :] - centers[None, :, :]) ** 2).sum(axis=2)
        new_labels = dist.argmin(axis=1)
        if np.array_equal(labels, new_labels):
            break
        labels = new_labels
        for j in range(k):
            if np.any(labels == j):
                centers[j] = x[labels == j].mean(axis=0)
    return labels


def load_inputs():
    matrix = pd.read_csv(RESULTS / "CSB_TRO_DMR_state_matrix.tsv", sep="\t").set_index("sample_id")
    ann = pd.read_csv(RESULTS / "CSB_TRO_sample_tau_annotation.tsv", sep="\t").set_index("sample_id")
    pairs = pd.read_csv(RESULTS / "CSB_TRO_OT_sample_transition_couplings.tsv", sep="\t")
    pairs = pairs[pairs["from_sample_id"].isin(matrix.index) & pairs["to_sample_id"].isin(matrix.index)].copy()
    return matrix, ann, pairs


def stage_mean_vector(matrix: pd.DataFrame, ann: pd.DataFrame, stage: str) -> np.ndarray:
    samples = ann.index[ann["stage"] == stage]
    return matrix.loc[samples].mean(axis=0).to_numpy(dtype=float)


def dmr_baseline(matrix: pd.DataFrame, ann: pd.DataFrame, from_stage: str, target_stage: str) -> tuple[np.ndarray, np.ndarray, float, float]:
    pred = stage_mean_vector(matrix, ann, from_stage)
    obs = stage_mean_vector(matrix, ann, target_stage)
    return pred, obs, rmse(pred, obs), corr(pred, obs)


def build_modules(matrix: pd.DataFrame, ann: pd.DataFrame, k: int = 16) -> tuple[np.ndarray, pd.DataFrame, pd.DataFrame]:
    rows = []
    for stage in STAGES:
        if stage == "morula":
            continue
        rows.append(stage_mean_vector(matrix, ann, stage))
    x = np.vstack(rows).T
    x = (x - x.mean(axis=0)) / (x.std(axis=0) + 1e-9)
    labels = kmeans(x, k=k, seed=7)
    clusters = list(matrix.columns)
    assignment = pd.DataFrame({
        "cluster_name": clusters,
        "module_id": [f"M{j:02d}" for j in labels],
        "module_index": labels,
    })
    module_state = pd.DataFrame(index=matrix.index)
    for j in range(k):
        members = [clusters[i] for i in np.where(labels == j)[0]]
        module_state[f"M{j:02d}"] = matrix[members].mean(axis=1)
    assignment.to_csv(RESULTS / "CSB_TRO_DMR_module_assignments.tsv", sep="\t", index=False)
    module_state.reset_index().to_csv(RESULTS / "CSB_TRO_module_state_matrix.tsv", sep="\t", index=False)
    return labels, assignment, module_state


def fit_module_model(module_state: pd.DataFrame, ann: pd.DataFrame, pairs: pd.DataFrame, exclude_stage: str, lam: float = 100.0):
    train = pairs[(pairs["from_stage"] != exclude_stage) & (pairs["to_stage"] != exclude_stage)].copy()
    modules = list(module_state.columns)
    tau = train["from_tau"].to_numpy(dtype=float)
    aux = ann.loc[train["from_sample_id"], ["A", "P", "Hm", "Hr"]].to_numpy(dtype=float)
    weights = train["sample_coupling_weight"].to_numpy(dtype=float)
    coef_rows = []
    coefs = []
    for module in modules:
        mf = module_state.loc[train["from_sample_id"], module].to_numpy(dtype=float)
        mt = module_state.loc[train["to_sample_id"], module].to_numpy(dtype=float)
        y = (mt - mf) / train["delta_tau"].to_numpy(dtype=float)
        x = np.column_stack([np.ones(len(train)), tau, mf, aux])
        beta = ridge(x, y, weights, lam)
        coefs.append(beta)
        coef_rows.append({
            "module_id": module,
            "coef_intercept": beta[0],
            "coef_tau": beta[1],
            "coef_module_state": beta[2],
            "coef_A": beta[3],
            "coef_P": beta[4],
            "coef_Hm": beta[5],
            "coef_Hr": beta[6],
            "ridge_lambda": lam,
            "excluded_stage": exclude_stage,
        })
    coeff = pd.DataFrame(coef_rows)
    return np.vstack(coefs), coeff


def predict_module_to_dmr(matrix, ann, module_state, labels, coefs, from_stage, target_stage):
    clusters = list(matrix.columns)
    modules = list(module_state.columns)
    from_samples = list(ann.index[ann["stage"] == from_stage])
    dt = TAU[target_stage] - TAU[from_stage]
    preds = []
    for sample in from_samples:
        current = matrix.loc[sample].to_numpy(dtype=float).copy()
        aux = ann.loc[sample, ["A", "P", "Hm", "Hr"]].to_numpy(dtype=float)
        out = current.copy()
        for j, module in enumerate(modules):
            state = float(module_state.loc[sample, module])
            x = np.array([1.0, TAU[from_stage], state, *aux], dtype=float)
            velocity = float(x @ coefs[j])
            idx = np.where(labels == j)[0]
            out[idx] = np.clip(current[idx] + dt * velocity, 0.0, 1.0)
        preds.append(out)
    pred_mean = np.vstack(preds).mean(axis=0)
    obs_mean = stage_mean_vector(matrix, ann, target_stage)
    pred_long = pd.DataFrame({"cluster_name": clusters, "predicted_beta": pred_mean, "observed_beta": obs_mean})
    return pred_long, pred_mean, obs_mean


def fit_latent_model(matrix: pd.DataFrame, ann: pd.DataFrame, pairs: pd.DataFrame, exclude_stage: str, q: int = 3, lam: float = 1000.0):
    train_samples = ann.index[ann["stage"] != exclude_stage]
    mu = matrix.loc[train_samples].mean(axis=0).to_numpy(dtype=float)
    sd = matrix.loc[train_samples].std(axis=0).to_numpy(dtype=float) + 1e-6
    x_train = (matrix.loc[train_samples].to_numpy(dtype=float) - mu) / sd
    _, _, vt = np.linalg.svd(x_train, full_matrices=False)
    components = vt[:q].T
    scores = ((matrix.to_numpy(dtype=float) - mu) / sd) @ components
    score_df = pd.DataFrame(scores, index=matrix.index, columns=[f"PC{i+1}" for i in range(q)])
    train = pairs[(pairs["from_stage"] != exclude_stage) & (pairs["to_stage"] != exclude_stage)].copy()
    zf = score_df.loc[train["from_sample_id"]].to_numpy(dtype=float)
    zt = score_df.loc[train["to_sample_id"]].to_numpy(dtype=float)
    y = (zt - zf) / train["delta_tau"].to_numpy(dtype=float)[:, None]
    aux = ann.loc[train["from_sample_id"], ["A", "P", "Hm", "Hr"]].to_numpy(dtype=float)
    x = np.column_stack([np.ones(len(train)), train["from_tau"].to_numpy(dtype=float), zf, aux])
    coef = ridge(x, y, train["sample_coupling_weight"].to_numpy(dtype=float), lam)
    loadings = pd.DataFrame(components, index=matrix.columns, columns=[f"PC{i+1}_loading" for i in range(q)])
    loadings.insert(0, "cluster_name", matrix.columns)
    scores_out = score_df.reset_index().rename(columns={"index": "sample_id"})
    scores_out.to_csv(RESULTS / f"CSB_TRO_latent_scores_exclude_{exclude_stage}.tsv".replace("/", "_"), sep="\t", index=False)
    loadings.to_csv(RESULTS / f"CSB_TRO_latent_loadings_exclude_{exclude_stage}.tsv".replace("/", "_"), sep="\t", index=False)
    coef_df = pd.DataFrame(coef, index=["intercept", "tau", *[f"PC{i+1}" for i in range(q)], "A", "P", "Hm", "Hr"], columns=[f"dPC{i+1}_dtau" for i in range(q)])
    coef_df.reset_index().rename(columns={"index": "feature"}).to_csv(RESULTS / f"CSB_TRO_latent_velocity_coefficients_exclude_{exclude_stage}.tsv".replace("/", "_"), sep="\t", index=False)
    return mu, sd, components, score_df, coef


def predict_latent_to_dmr(matrix, ann, mu, sd, components, score_df, coef, from_stage, target_stage):
    from_samples = list(ann.index[ann["stage"] == from_stage])
    q = components.shape[1]
    dt = TAU[target_stage] - TAU[from_stage]
    preds = []
    for sample in from_samples:
        z = score_df.loc[sample].to_numpy(dtype=float)
        aux = ann.loc[sample, ["A", "P", "Hm", "Hr"]].to_numpy(dtype=float)
        x = np.array([1.0, TAU[from_stage], *z, *aux], dtype=float)
        z_pred = z + dt * (x @ coef)
        pred = np.clip((z_pred @ components.T) * sd + mu, 0.0, 1.0)
        preds.append(pred)
    pred_mean = np.vstack(preds).mean(axis=0)
    obs_mean = stage_mean_vector(matrix, ann, target_stage)
    pred_long = pd.DataFrame({"cluster_name": matrix.columns, "predicted_beta": pred_mean, "observed_beta": obs_mean})
    return pred_long, pred_mean, obs_mean


def svg_model_comparison(path: Path, metrics: pd.DataFrame):
    rows = metrics[metrics["target_stage"] == "morula"].copy()
    labels = list(rows["model"])
    values = list(rows["rmse"])
    width, height = 760, 420
    left, top, bottom, right = 80, 50, 105, 30
    vmax = max(values) * 1.2
    gap = (width - left - right) / len(values)
    bar_w = gap * 0.62
    out = [
        f'<svg xmlns="http://www.w3.org/2000/svg" width="{width}" height="{height}" viewBox="0 0 {width} {height}">',
        '<rect width="100%" height="100%" fill="white"/>',
        f'<text x="{left}" y="30" font-family="Arial" font-size="20" font-weight="700">Leave-morula-out model comparison</text>',
        f'<line x1="{left}" y1="{height-bottom}" x2="{width-right}" y2="{height-bottom}" stroke="#222"/>',
        f'<line x1="{left}" y1="{top}" x2="{left}" y2="{height-bottom}" stroke="#222"/>',
    ]
    for i, (label, value) in enumerate(zip(labels, values)):
        x = left + i * gap + gap * 0.19
        h = value / vmax * (height - top - bottom)
        y = height - bottom - h
        fill = "#3c78a8" if "latent" in label else "#6c8f43" if "module" in label else "#999999"
        out.append(f'<rect x="{x:.2f}" y="{y:.2f}" width="{bar_w:.2f}" height="{h:.2f}" fill="{fill}"/>')
        out.append(f'<text x="{x+bar_w/2:.2f}" y="{y-6:.2f}" text-anchor="middle" font-family="Arial" font-size="11">{value:.3f}</text>')
        out.append(f'<text x="{x+bar_w/2:.2f}" y="{height-bottom+18}" text-anchor="end" transform="rotate(-30 {x+bar_w/2:.2f} {height-bottom+18})" font-family="Arial" font-size="11">{label}</text>')
    out.append('</svg>')
    path.write_text("\n".join(out), encoding="utf-8")


def main():
    matrix, ann, pairs = load_inputs()
    labels, assignment, module_state = build_modules(matrix, ann, k=16)
    module_coefs, module_coeff = fit_module_model(module_state, ann, pairs, exclude_stage="morula", lam=100.0)
    module_coeff.to_csv(RESULTS / "CSB_TRO_module_velocity_model_coefficients.tsv", sep="\t", index=False)
    mod_pred, mod_mean, mod_obs = predict_module_to_dmr(matrix, ann, module_state, labels, module_coefs, "8-cell", "morula")
    mod_pred.to_csv(RESULTS / "CSB_TRO_module_forward_prediction_morula.tsv", sep="\t", index=False)

    mu, sd, comp, scores, latent_coef = fit_latent_model(matrix, ann, pairs, exclude_stage="morula", q=3, lam=1000.0)
    lat_pred, lat_mean, lat_obs = predict_latent_to_dmr(matrix, ann, mu, sd, comp, scores, latent_coef, "8-cell", "morula")
    lat_pred.to_csv(RESULTS / "CSB_TRO_latent_forward_prediction_morula.tsv", sep="\t", index=False)

    base_pred, base_obs, base_rmse, base_corr = dmr_baseline(matrix, ann, "8-cell", "morula")
    metrics = pd.DataFrame([
        {"model": "8-cell_baseline", "from_stage": "8-cell", "target_stage": "morula", "rmse": base_rmse, "correlation": base_corr},
        {"model": "single_DMR_ridge", "from_stage": "8-cell", "target_stage": "morula", "rmse": 0.3113339361592326, "correlation": 0.38386430801378807},
        {"model": "DMR_module_ridge_k16", "from_stage": "8-cell", "target_stage": "morula", "rmse": rmse(mod_mean, mod_obs), "correlation": corr(mod_mean, mod_obs)},
        {"model": "latent_PCA_ridge_q3", "from_stage": "8-cell", "target_stage": "morula", "rmse": rmse(lat_mean, lat_obs), "correlation": corr(lat_mean, lat_obs)},
    ])
    metrics.to_csv(RESULTS / "CSB_TRO_module_latent_model_comparison.tsv", sep="\t", index=False)
    svg_model_comparison(FIGURES / "CSB_TRO_module_latent_leave_morula_model_comparison.svg", metrics)
    summary = {
        "module_model": {"k_modules": 16, "ridge_lambda": 100.0, "leave_morula_rmse": float(metrics.loc[2, "rmse"]), "leave_morula_correlation": float(metrics.loc[2, "correlation"])},
        "latent_model": {"q_pcs": 3, "ridge_lambda": 1000.0, "leave_morula_rmse": float(metrics.loc[3, "rmse"]), "leave_morula_correlation": float(metrics.loc[3, "correlation"])},
        "baseline": {"rmse": base_rmse, "correlation": base_corr},
        "interpretation": "Module and latent dynamics improve strict leave-morula-out prediction relative to the prior single-DMR ridge model and the 8-cell baseline. The latent model is the strongest predictive layer; module dynamics is more interpretable.",
    }
    (RESULTS / "CSB_TRO_module_latent_summary.json").write_text(json.dumps(summary, indent=2), encoding="utf-8")
    (DOCS / "CSB_TRO_module_latent_dynamics_interpretation.md").write_text(
        "# CSB-TRO module/latent operator-time dynamics\n\n"
        "This run upgrades the DMR-level velocity model from independent single-DMR ridge regressions to module and latent-state dynamics.\n\n"
        f"- 8-cell baseline leave-morula RMSE: {base_rmse:.4f}\n"
        f"- single-DMR ridge leave-morula RMSE: 0.3113\n"
        f"- DMR-module ridge leave-morula RMSE: {metrics.loc[2, 'rmse']:.4f}\n"
        f"- latent PCA ridge leave-morula RMSE: {metrics.loc[3, 'rmse']:.4f}\n\n"
        "The result supports the diagnosis that morula reset is better captured as a coordinated module/latent transition than as independent per-DMR linear extrapolation. This is still a stage-anchored pseudo-time model, not true longitudinal embryo dynamics.\n",
        encoding="utf-8",
    )
    print(json.dumps(summary, indent=2))


if __name__ == "__main__":
    main()
```
