## Supplemental code for "An entropy-based diagnostic framework for characterizing methylation state dynamics during preimplantation development": run_module_latent_validation.py

```
from __future__ import annotations

import argparse
import json
import math
from pathlib import Path

import numpy as np
import pandas as pd


BASE = Path(r"E:\CSB_TRO_operator_time_DMR_dynamics_2026-05-25")
RESULTS = BASE / "results"
FIGURES = BASE / "figures"
DOCS = BASE / "docs"

STAGES = ["MII oocyte", "zygote/PN", "2-cell", "4-cell", "8-cell", "morula", "blastocyst"]
TAU = {stage: i / (len(STAGES) - 1) for i, stage in enumerate(STAGES)}
EXCLUDE_STAGE = "morula"
FROM_STAGE = "8-cell"
TARGET_STAGE = "morula"


def rmse(a, b) -> float:
    a = np.asarray(a, dtype=float)
    b = np.asarray(b, dtype=float)
    return float(np.sqrt(np.mean((a - b) ** 2)))


def corr(a, b) -> float:
    a = np.asarray(a, dtype=float)
    b = np.asarray(b, dtype=float)
    aa = a - a.mean()
    bb = b - b.mean()
    den = math.sqrt(float((aa @ aa) * (bb @ bb)))
    return float((aa @ bb) / den) if den else float("nan")


def ridge(x: np.ndarray, y: np.ndarray, w: np.ndarray, lam: float) -> np.ndarray:
    sw = np.sqrt(np.maximum(w, 1e-12) / np.mean(np.maximum(w, 1e-12)))
    xw = x * sw[:, None]
    yw = y * sw if y.ndim == 1 else y * sw[:, None]
    penalty = lam * np.eye(x.shape[1])
    penalty[0, 0] = 0.0
    return np.linalg.solve(xw.T @ xw + penalty, xw.T @ yw)


def kmeans(x: np.ndarray, k: int, seed: int = 1, max_iter: int = 100) -> np.ndarray:
    rng = np.random.default_rng(seed)
    centers = x[rng.choice(len(x), k, replace=False)].copy()
    labels = np.zeros(len(x), dtype=int)
    for _ in range(max_iter):
        dist = ((x[:, None, :] - centers[None, :, :]) ** 2).sum(axis=2)
        new_labels = dist.argmin(axis=1)
        if np.array_equal(labels, new_labels):
            break
        labels = new_labels
        for j in range(k):
            if np.any(labels == j):
                centers[j] = x[labels == j].mean(axis=0)
    return labels


def load_inputs():
    matrix = pd.read_csv(RESULTS / "CSB_TRO_DMR_state_matrix.tsv", sep="\t").set_index("sample_id")
    ann = pd.read_csv(RESULTS / "CSB_TRO_sample_tau_annotation.tsv", sep="\t").set_index("sample_id")
    pairs = pd.read_csv(RESULTS / "CSB_TRO_OT_sample_transition_couplings.tsv", sep="\t")
    pairs = pairs[pairs["from_sample_id"].isin(matrix.index) & pairs["to_sample_id"].isin(matrix.index)].copy()
    return matrix, ann, pairs


def stage_sample_ids(ann: pd.DataFrame, stage: str) -> list[str]:
    return list(ann.index[ann["stage"] == stage])


def stage_mean_vector(matrix: pd.DataFrame, ann: pd.DataFrame, stage: str) -> np.ndarray:
    return matrix.loc[stage_sample_ids(ann, stage)].mean(axis=0).to_numpy(dtype=float)


def build_modules(matrix: pd.DataFrame, ann: pd.DataFrame, k: int = 16, seed: int = 7) -> tuple[np.ndarray, pd.DataFrame]:
    rows = [stage_mean_vector(matrix, ann, stage) for stage in STAGES if stage != EXCLUDE_STAGE]
    x = np.vstack(rows).T
    x = (x - x.mean(axis=0)) / (x.std(axis=0) + 1e-9)
    labels = kmeans(x, k=k, seed=seed)
    return labels, module_state_from_labels(matrix, labels)


def module_state_from_labels(matrix: pd.DataFrame, labels: np.ndarray) -> pd.DataFrame:
    module_state = pd.DataFrame(index=matrix.index)
    for j in sorted(np.unique(labels)):
        members = [matrix.columns[i] for i in np.where(labels == j)[0]]
        module_state[f"M{j:02d}"] = matrix[members].mean(axis=1)
    return module_state


def training_pairs(pairs: pd.DataFrame, exclude_stage: str = EXCLUDE_STAGE) -> pd.DataFrame:
    train = pairs[(pairs["from_stage"] != exclude_stage) & (pairs["to_stage"] != exclude_stage)].copy()
    train = train[np.abs(train["delta_tau"].to_numpy(dtype=float)) > 1e-9].copy()
    return train


def fit_module_model(module_state: pd.DataFrame, ann: pd.DataFrame, pairs: pd.DataFrame, lam: float = 100.0) -> np.ndarray:
    train = training_pairs(pairs)
    modules = list(module_state.columns)
    tau = train["from_tau"].to_numpy(dtype=float)
    aux = ann.loc[train["from_sample_id"], ["A", "P", "Hm", "Hr"]].to_numpy(dtype=float)
    weights = train["sample_coupling_weight"].to_numpy(dtype=float)
    coefs = []
    for module in modules:
        mf = module_state.loc[train["from_sample_id"], module].to_numpy(dtype=float)
        mt = module_state.loc[train["to_sample_id"], module].to_numpy(dtype=float)
        y = (mt - mf) / train["delta_tau"].to_numpy(dtype=float)
        x = np.column_stack([np.ones(len(train)), tau, mf, aux])
        coefs.append(ridge(x, y, weights, lam))
    return np.vstack(coefs)


def predict_module_samples(
    matrix: pd.DataFrame,
    ann: pd.DataFrame,
    module_state: pd.DataFrame,
    labels: np.ndarray,
    coefs: np.ndarray,
) -> tuple[np.ndarray, np.ndarray, np.ndarray]:
    from_samples = stage_sample_ids(ann, FROM_STAGE)
    target_samples = stage_sample_ids(ann, TARGET_STAGE)
    dt = TAU[TARGET_STAGE] - TAU[FROM_STAGE]
    preds = []
    for sample in from_samples:
        current = matrix.loc[sample].to_numpy(dtype=float).copy()
        aux = ann.loc[sample, ["A", "P", "Hm", "Hr"]].to_numpy(dtype=float)
        out = current.copy()
        for j, module in enumerate(module_state.columns):
            state = float(module_state.loc[sample, module])
            x = np.array([1.0, TAU[FROM_STAGE], state, *aux], dtype=float)
            velocity = float(x @ coefs[j])
            idx = np.where(labels == j)[0]
            out[idx] = np.clip(current[idx] + dt * velocity, 0.0, 1.0)
        preds.append(out)
    return np.vstack(preds), matrix.loc[target_samples].to_numpy(dtype=float), matrix.loc[from_samples].to_numpy(dtype=float)


def fit_latent_model(matrix: pd.DataFrame, ann: pd.DataFrame, pairs: pd.DataFrame, q: int = 3, lam: float = 1000.0):
    train_samples = ann.index[ann["stage"] != EXCLUDE_STAGE]
    mu = matrix.loc[train_samples].mean(axis=0).to_numpy(dtype=float)
    sd = matrix.loc[train_samples].std(axis=0).to_numpy(dtype=float) + 1e-6
    x_train = (matrix.loc[train_samples].to_numpy(dtype=float) - mu) / sd
    _, _, vt = np.linalg.svd(x_train, full_matrices=False)
    components = vt[:q].T
    scores = ((matrix.to_numpy(dtype=float) - mu) / sd) @ components
    score_df = pd.DataFrame(scores, index=matrix.index, columns=[f"PC{i + 1}" for i in range(q)])
    train = training_pairs(pairs)
    zf = score_df.loc[train["from_sample_id"]].to_numpy(dtype=float)
    zt = score_df.loc[train["to_sample_id"]].to_numpy(dtype=float)
    y = (zt - zf) / train["delta_tau"].to_numpy(dtype=float)[:, None]
    aux = ann.loc[train["from_sample_id"], ["A", "P", "Hm", "Hr"]].to_numpy(dtype=float)
    x = np.column_stack([np.ones(len(train)), train["from_tau"].to_numpy(dtype=float), zf, aux])
    coef = ridge(x, y, train["sample_coupling_weight"].to_numpy(dtype=float), lam)
    return mu, sd, components, score_df, coef


def predict_latent_samples(matrix: pd.DataFrame, ann: pd.DataFrame, latent_fit) -> tuple[np.ndarray, np.ndarray, np.ndarray]:
    mu, sd, components, score_df, coef = latent_fit
    from_samples = stage_sample_ids(ann, FROM_STAGE)
    target_samples = stage_sample_ids(ann, TARGET_STAGE)
    dt = TAU[TARGET_STAGE] - TAU[FROM_STAGE]
    preds = []
    for sample in from_samples:
        z = score_df.loc[sample].to_numpy(dtype=float)
        aux = ann.loc[sample, ["A", "P", "Hm", "Hr"]].to_numpy(dtype=float)
        x = np.array([1.0, TAU[FROM_STAGE], *z, *aux], dtype=float)
        z_pred = z + dt * (x @ coef)
        preds.append(np.clip((z_pred @ components.T) * sd + mu, 0.0, 1.0))
    return np.vstack(preds), matrix.loc[target_samples].to_numpy(dtype=float), matrix.loc[from_samples].to_numpy(dtype=float)


def vector_metrics(pred_samples: np.ndarray, obs_samples: np.ndarray) -> tuple[np.ndarray, np.ndarray, float, float]:
    pred_mean = pred_samples.mean(axis=0)
    obs_mean = obs_samples.mean(axis=0)
    return pred_mean, obs_mean, rmse(pred_mean, obs_mean), corr(pred_mean, obs_mean)


def permute_tau_by_sample(ann: pd.DataFrame, pairs: pd.DataFrame, rng: np.random.Generator) -> pd.DataFrame:
    shuffled = ann["tau"].copy()
    eligible = ann.index[ann["stage"] != EXCLUDE_STAGE]
    shuffled.loc[eligible] = rng.permutation(shuffled.loc[eligible].to_numpy(dtype=float))
    out = pairs.copy()
    out["from_tau"] = shuffled.loc[out["from_sample_id"]].to_numpy(dtype=float)
    out["to_tau"] = shuffled.loc[out["to_sample_id"]].to_numpy(dtype=float)
    out["delta_tau"] = out["to_tau"] - out["from_tau"]
    return out


def permute_couplings(pairs: pd.DataFrame, rng: np.random.Generator) -> pd.DataFrame:
    out = pairs.copy()
    for _, idx in out.groupby(["from_stage", "to_stage"]).groups.items():
        idx = np.array(list(idx))
        out.loc[idx, "to_sample_id"] = rng.permutation(out.loc[idx, "to_sample_id"].to_numpy())
    return out


def random_module_labels(labels: np.ndarray, rng: np.random.Generator) -> np.ndarray:
    return rng.permutation(labels)


def paired_signflip_p(diff: np.ndarray, rng: np.random.Generator, n_perm: int = 20000) -> tuple[float, float]:
    diff = np.asarray(diff, dtype=float)
    observed = float(diff.mean())
    signs = rng.choice([-1.0, 1.0], size=(n_perm, len(diff)))
    null = (signs * diff[None, :]).mean(axis=1)
    p_one_sided_better = float((np.sum(null >= observed) + 1) / (n_perm + 1))
    p_two_sided = float((np.sum(np.abs(null) >= abs(observed)) + 1) / (n_perm + 1))
    return p_one_sided_better, p_two_sided


def bootstrap_ci(values: list[float]) -> tuple[float, float, float]:
    arr = np.asarray(values, dtype=float)
    return float(np.mean(arr)), float(np.quantile(arr, 0.025)), float(np.quantile(arr, 0.975))


def make_observed_predictions(matrix: pd.DataFrame, ann: pd.DataFrame, pairs: pd.DataFrame):
    labels, module_state = build_modules(matrix, ann, k=16, seed=7)
    module_coefs = fit_module_model(module_state, ann, pairs, lam=100.0)
    module_pred, obs_samples, base_pred_samples = predict_module_samples(matrix, ann, module_state, labels, module_coefs)
    latent_fit = fit_latent_model(matrix, ann, pairs, q=3, lam=1000.0)
    latent_pred, _, _ = predict_latent_samples(matrix, ann, latent_fit)
    return {
        "8-cell_baseline": {"pred_samples": base_pred_samples, "obs_samples": obs_samples},
        "DMR_module_ridge_k16": {"pred_samples": module_pred, "obs_samples": obs_samples, "labels": labels},
        "latent_PCA_ridge_q3": {"pred_samples": latent_pred, "obs_samples": obs_samples},
    }


def observed_metric_table(predictions: dict) -> pd.DataFrame:
    rows = []
    for model, data in predictions.items():
        pred_mean, obs_mean, model_rmse, model_corr = vector_metrics(data["pred_samples"], data["obs_samples"])
        rows.append({
            "model": model,
            "from_stage": FROM_STAGE,
            "target_stage": TARGET_STAGE,
            "rmse": model_rmse,
            "correlation": model_corr,
            "n_from_samples": data["pred_samples"].shape[0],
            "n_target_samples": data["obs_samples"].shape[0],
            "n_dmrs": data["pred_samples"].shape[1],
        })
    return pd.DataFrame(rows)


def run_null_models(matrix, ann, pairs, observed, n_null: int, rng: np.random.Generator) -> pd.DataFrame:
    rows = []
    base_rmse = float(observed.loc[observed["model"] == "8-cell_baseline", "rmse"].iloc[0])
    labels, _ = build_modules(matrix, ann, k=16, seed=7)
    for i in range(n_null):
        for null_type, null_pairs in [
            ("random_tau_null", permute_tau_by_sample(ann, pairs, rng)),
            ("random_coupling_null", permute_couplings(pairs, rng)),
        ]:
            labels_obs, module_state = build_modules(matrix, ann, k=16, seed=7)
            module_coefs = fit_module_model(module_state, ann, null_pairs, lam=100.0)
            mod_pred, obs_samples, _ = predict_module_samples(matrix, ann, module_state, labels_obs, module_coefs)
            _, _, model_rmse, model_corr = vector_metrics(mod_pred, obs_samples)
            rows.append({"null_type": null_type, "iteration": i + 1, "model": "DMR_module_ridge_k16", "rmse": model_rmse, "correlation": model_corr, "delta_rmse_vs_baseline": model_rmse - base_rmse})

            latent_fit = fit_latent_model(matrix, ann, null_pairs, q=3, lam=1000.0)
            lat_pred, obs_samples, _ = predict_latent_samples(matrix, ann, latent_fit)
            _, _, model_rmse, model_corr = vector_metrics(lat_pred, obs_samples)
            rows.append({"null_type": null_type, "iteration": i + 1, "model": "latent_PCA_ridge_q3", "rmse": model_rmse, "correlation": model_corr, "delta_rmse_vs_baseline": model_rmse - base_rmse})

        rand_labels = random_module_labels(labels, rng)
        rand_module_state = module_state_from_labels(matrix, rand_labels)
        rand_coefs = fit_module_model(rand_module_state, ann, pairs, lam=100.0)
        rand_pred, obs_samples, _ = predict_module_samples(matrix, ann, rand_module_state, rand_labels, rand_coefs)
        _, _, model_rmse, model_corr = vector_metrics(rand_pred, obs_samples)
        rows.append({"null_type": "random_module_null", "iteration": i + 1, "model": "DMR_module_ridge_k16", "rmse": model_rmse, "correlation": model_corr, "delta_rmse_vs_baseline": model_rmse - base_rmse})
    return pd.DataFrame(rows)


def run_bootstraps(predictions: dict, n_boot: int, rng: np.random.Generator) -> tuple[pd.DataFrame, pd.DataFrame]:
    raw = []
    models = list(predictions)
    n_dmrs = predictions[models[0]]["pred_samples"].shape[1]
    n_pred = predictions[models[0]]["pred_samples"].shape[0]
    n_obs = predictions[models[0]]["obs_samples"].shape[0]
    for i in range(n_boot):
        dmr_idx = rng.integers(0, n_dmrs, size=n_dmrs)
        pred_idx = rng.integers(0, n_pred, size=n_pred)
        obs_idx = rng.integers(0, n_obs, size=n_obs)
        for model, data in predictions.items():
            pred_mean = data["pred_samples"].mean(axis=0)[dmr_idx]
            obs_mean = data["obs_samples"].mean(axis=0)[dmr_idx]
            raw.append({"bootstrap_type": "DMR_bootstrap", "iteration": i + 1, "model": model, "rmse": rmse(pred_mean, obs_mean), "correlation": corr(pred_mean, obs_mean)})

            pred_mean = data["pred_samples"][pred_idx].mean(axis=0)
            obs_mean = data["obs_samples"][obs_idx].mean(axis=0)
            raw.append({"bootstrap_type": "stage_sample_bootstrap", "iteration": i + 1, "model": model, "rmse": rmse(pred_mean, obs_mean), "correlation": corr(pred_mean, obs_mean)})
    raw_df = pd.DataFrame(raw)
    base = raw_df[raw_df["model"] == "8-cell_baseline"][["bootstrap_type", "iteration", "rmse"]].rename(columns={"rmse": "baseline_rmse"})
    raw_df = raw_df.merge(base, on=["bootstrap_type", "iteration"], how="left")
    raw_df["delta_rmse_vs_baseline"] = raw_df["rmse"] - raw_df["baseline_rmse"]

    summary_rows = []
    for (bootstrap_type, model), sub in raw_df.groupby(["bootstrap_type", "model"], sort=False):
        for metric in ["rmse", "delta_rmse_vs_baseline", "correlation"]:
            mean, lo, hi = bootstrap_ci(list(sub[metric]))
            summary_rows.append({
                "bootstrap_type": bootstrap_type,
                "model": model,
                "metric": metric,
                "mean": mean,
                "ci95_low": lo,
                "ci95_high": hi,
                "n_bootstrap": n_boot,
            })
    return pd.DataFrame(summary_rows), raw_df


def validation_summary(observed: pd.DataFrame, ci: pd.DataFrame, paired: pd.DataFrame, null_summary: pd.DataFrame) -> pd.DataFrame:
    rows = []
    for _, row in observed.iterrows():
        model = row["model"]
        out = row.to_dict()
        for boot_type in ["DMR_bootstrap", "stage_sample_bootstrap"]:
            rmse_ci = ci[(ci["bootstrap_type"] == boot_type) & (ci["model"] == model) & (ci["metric"] == "rmse")]
            delta_ci = ci[(ci["bootstrap_type"] == boot_type) & (ci["model"] == model) & (ci["metric"] == "delta_rmse_vs_baseline")]
            if len(rmse_ci):
                prefix = boot_type.replace("_bootstrap", "")
                out[f"{prefix}_rmse_ci95_low"] = float(rmse_ci["ci95_low"].iloc[0])
                out[f"{prefix}_rmse_ci95_high"] = float(rmse_ci["ci95_high"].iloc[0])
            if len(delta_ci):
                prefix = boot_type.replace("_bootstrap", "")
                out[f"{prefix}_delta_rmse_vs_baseline_ci95_low"] = float(delta_ci["ci95_low"].iloc[0])
                out[f"{prefix}_delta_rmse_vs_baseline_ci95_high"] = float(delta_ci["ci95_high"].iloc[0])
        sq = paired[(paired["model"] == model) & (paired["paired_metric"] == "squared_error")]
        if len(sq):
            out["paired_sq_error_mean_improvement_vs_baseline"] = float(sq["mean_improvement_vs_baseline"].iloc[0])
            out["paired_sq_error_p_one_sided_model_better"] = float(sq["signflip_p_one_sided_model_better"].iloc[0])
        model_nulls = null_summary[null_summary["model"] == model]
        if len(model_nulls):
            out["null_controls"] = ";".join(
                f"{r.null_type}:p={r.empirical_p_null_rmse_le_observed:.4g}" for r in model_nulls.itertuples()
            )
        rows.append(out)
    return pd.DataFrame(rows)


def paired_error_tests(predictions: dict, rng: np.random.Generator) -> tuple[pd.DataFrame, pd.DataFrame]:
    base_pred = predictions["8-cell_baseline"]["pred_samples"].mean(axis=0)
    obs = predictions["8-cell_baseline"]["obs_samples"].mean(axis=0)
    base_abs = np.abs(base_pred - obs)
    base_sq = (base_pred - obs) ** 2
    per_dmr = pd.DataFrame({"cluster_name": pd.read_csv(RESULTS / "CSB_TRO_DMR_state_matrix.tsv", sep="\t", nrows=0).columns[1:], "observed_beta": obs, "baseline_predicted_beta": base_pred, "baseline_abs_error": base_abs, "baseline_sq_error": base_sq})
    rows = []
    for model in ["DMR_module_ridge_k16", "latent_PCA_ridge_q3"]:
        pred = predictions[model]["pred_samples"].mean(axis=0)
        abs_err = np.abs(pred - obs)
        sq_err = (pred - obs) ** 2
        per_dmr[f"{model}_predicted_beta"] = pred
        per_dmr[f"{model}_abs_error"] = abs_err
        per_dmr[f"{model}_sq_error"] = sq_err
        per_dmr[f"{model}_abs_error_improvement_vs_baseline"] = base_abs - abs_err
        per_dmr[f"{model}_sq_error_improvement_vs_baseline"] = base_sq - sq_err
        for metric_name, diff in [("absolute_error", base_abs - abs_err), ("squared_error", base_sq - sq_err)]:
            p_one, p_two = paired_signflip_p(diff, rng)
            rows.append({
                "model": model,
                "paired_metric": metric_name,
                "mean_improvement_vs_baseline": float(np.mean(diff)),
                "median_improvement_vs_baseline": float(np.median(diff)),
                "fraction_DMRs_improved": float(np.mean(diff > 0)),
                "n_dmrs": len(diff),
                "signflip_p_one_sided_model_better": p_one,
                "signflip_p_two_sided": p_two,
            })
    return pd.DataFrame(rows), per_dmr


def empirical_null_summary(observed: pd.DataFrame, nulls: pd.DataFrame) -> pd.DataFrame:
    rows = []
    for (null_type, model), sub in nulls.groupby(["null_type", "model"], sort=False):
        obs_rmse = float(observed.loc[observed["model"] == model, "rmse"].iloc[0])
        rows.append({
            "null_type": null_type,
            "model": model,
            "observed_rmse": obs_rmse,
            "null_rmse_mean": float(sub["rmse"].mean()),
            "null_rmse_ci95_low": float(sub["rmse"].quantile(0.025)),
            "null_rmse_ci95_high": float(sub["rmse"].quantile(0.975)),
            "empirical_p_null_rmse_le_observed": float((np.sum(sub["rmse"].to_numpy() <= obs_rmse) + 1) / (len(sub) + 1)),
            "n_null": int(len(sub)),
        })
    return pd.DataFrame(rows)


def svg_validation(path: Path, observed: pd.DataFrame, ci: pd.DataFrame, null_summary: pd.DataFrame):
    rows = observed.copy()
    width, height = 900, 500
    left, right, top, bottom = 90, 30, 55, 135
    values = rows["rmse"].to_numpy(dtype=float)
    vmax = max(values.max(), null_summary["null_rmse_ci95_high"].max()) * 1.12
    plot_h = height - top - bottom
    gap = (width - left - right) / len(rows)
    bar_w = gap * 0.42
    colors = {"8-cell_baseline": "#777777", "DMR_module_ridge_k16": "#5b8f58", "latent_PCA_ridge_q3": "#3b76a8"}
    out = [
        f'<svg xmlns="http://www.w3.org/2000/svg" width="{width}" height="{height}" viewBox="0 0 {width} {height}">',
        '<rect width="100%" height="100%" fill="white"/>',
        f'<text x="{left}" y="32" font-family="Arial" font-size="20" font-weight="700">Strict leave-morula-out validation</text>',
        f'<line x1="{left}" y1="{height-bottom}" x2="{width-right}" y2="{height-bottom}" stroke="#222"/>',
        f'<line x1="{left}" y1="{top}" x2="{left}" y2="{height-bottom}" stroke="#222"/>',
        f'<text x="18" y="{top+plot_h/2}" transform="rotate(-90 18 {top+plot_h/2})" font-family="Arial" font-size="13">RMSE</text>',
    ]
    for tick in np.linspace(0, vmax, 5):
        y = height - bottom - tick / vmax * plot_h
        out.append(f'<line x1="{left-4}" y1="{y:.2f}" x2="{left}" y2="{y:.2f}" stroke="#222"/>')
        out.append(f'<text x="{left-8}" y="{y+4:.2f}" text-anchor="end" font-family="Arial" font-size="11">{tick:.2f}</text>')
    for i, row in rows.iterrows():
        model = row["model"]
        x = left + i * gap + gap * 0.29
        h = row["rmse"] / vmax * plot_h
        y = height - bottom - h
        out.append(f'<rect x="{x:.2f}" y="{y:.2f}" width="{bar_w:.2f}" height="{h:.2f}" fill="{colors.get(model, "#999")}"/>')
        dmr_ci = ci[(ci["bootstrap_type"] == "DMR_bootstrap") & (ci["model"] == model) & (ci["metric"] == "rmse")]
        if len(dmr_ci):
            lo = float(dmr_ci["ci95_low"].iloc[0])
            hi = float(dmr_ci["ci95_high"].iloc[0])
            ylo = height - bottom - lo / vmax * plot_h
            yhi = height - bottom - hi / vmax * plot_h
            cx = x + bar_w / 2
            out.append(f'<line x1="{cx:.2f}" y1="{yhi:.2f}" x2="{cx:.2f}" y2="{ylo:.2f}" stroke="#111" stroke-width="1.5"/>')
            out.append(f'<line x1="{cx-7:.2f}" y1="{yhi:.2f}" x2="{cx+7:.2f}" y2="{yhi:.2f}" stroke="#111"/>')
            out.append(f'<line x1="{cx-7:.2f}" y1="{ylo:.2f}" x2="{cx+7:.2f}" y2="{ylo:.2f}" stroke="#111"/>')
        out.append(f'<text x="{x+bar_w/2:.2f}" y="{y-7:.2f}" text-anchor="middle" font-family="Arial" font-size="11">{row["rmse"]:.3f}</text>')
        out.append(f'<text x="{x+bar_w/2:.2f}" y="{height-bottom+18}" text-anchor="end" transform="rotate(-32 {x+bar_w/2:.2f} {height-bottom+18})" font-family="Arial" font-size="11">{model}</text>')
    out.append(f'<text x="{left}" y="{height-30}" font-family="Arial" font-size="12">Error bars: DMR bootstrap 95% CI. Null model summaries are reported in the TSV outputs.</text>')
    out.append("</svg>")
    path.write_text("\n".join(out), encoding="utf-8")


def write_interpretation(path: Path, observed: pd.DataFrame, ci: pd.DataFrame, paired: pd.DataFrame, null_summary: pd.DataFrame):
    base_rmse = float(observed.loc[observed["model"] == "8-cell_baseline", "rmse"].iloc[0])
    module_rmse = float(observed.loc[observed["model"] == "DMR_module_ridge_k16", "rmse"].iloc[0])
    latent_rmse = float(observed.loc[observed["model"] == "latent_PCA_ridge_q3", "rmse"].iloc[0])
    lines = [
        "# CSB-TRO module/latent validation interpretation",
        "",
        "This validation package keeps the strict leave-morula-out design: transitions involving morula are excluded from training, and morula is predicted from the 8-cell state in stage-anchored developmental operator time.",
        "",
        f"- 8-cell baseline RMSE: {base_rmse:.4f}",
        f"- DMR-module ridge RMSE: {module_rmse:.4f}",
        f"- latent PCA ridge RMSE: {latent_rmse:.4f}",
        "",
        "The paired DMR error tests compare per-DMR prediction errors against the 8-cell baseline using sign-flip paired permutation tests. The bootstrap tables report both DMR bootstrap and stage/sample bootstrap confidence intervals.",
        "",
        "Null models test whether comparable performance can be obtained after disrupting operator-time labels, sample-level OT coupling, or module membership. The empirical p-value is the fraction of null RMSE values less than or equal to the observed RMSE, with a +1 correction.",
        "",
        "Recommended wording: A DMR-level single-feature velocity model was insufficient for strict leave-morula-out prediction, whereas module-level and latent-state operator-time dynamics improved prediction of the morula methylation reset-basin. The validation package evaluates whether this improvement persists under DMR-level paired errors, bootstrap uncertainty, and randomized time/coupling/module controls.",
        "",
        "Caveat: this is a stage-anchored developmental operator-time model, not true longitudinal tracking of the same embryo. In silico sensitivity results should not be described as strong causal evidence.",
        "",
        "Key paired tests:",
    ]
    for _, row in paired.iterrows():
        lines.append(f"- {row['model']} {row['paired_metric']}: mean improvement {row['mean_improvement_vs_baseline']:.6f}, one-sided p={row['signflip_p_one_sided_model_better']:.4g}")
    lines.extend(["", "Key null summaries:"])
    for _, row in null_summary.iterrows():
        lines.append(f"- {row['model']} / {row['null_type']}: observed RMSE {row['observed_rmse']:.4f}, null mean {row['null_rmse_mean']:.4f}, empirical p={row['empirical_p_null_rmse_le_observed']:.4g}")
    path.write_text("\n".join(lines) + "\n", encoding="utf-8")


def main():
    parser = argparse.ArgumentParser()
    parser.add_argument("--n-null", type=int, default=200)
    parser.add_argument("--n-bootstrap", type=int, default=1000)
    parser.add_argument("--seed", type=int, default=20260525)
    args = parser.parse_args()

    RESULTS.mkdir(parents=True, exist_ok=True)
    FIGURES.mkdir(parents=True, exist_ok=True)
    DOCS.mkdir(parents=True, exist_ok=True)
    rng = np.random.default_rng(args.seed)

    matrix, ann, pairs = load_inputs()
    predictions = make_observed_predictions(matrix, ann, pairs)
    observed = observed_metric_table(predictions)

    nulls = run_null_models(matrix, ann, pairs, observed, args.n_null, rng)
    nulls.to_csv(RESULTS / "CSB_TRO_module_latent_null_models.tsv", sep="\t", index=False)
    null_summary = empirical_null_summary(observed, nulls)
    null_summary.to_csv(RESULTS / "CSB_TRO_module_latent_null_summary.tsv", sep="\t", index=False)

    ci, boot_raw = run_bootstraps(predictions, args.n_bootstrap, rng)
    ci.to_csv(RESULTS / "CSB_TRO_module_latent_bootstrap_CI.tsv", sep="\t", index=False)
    boot_raw.to_csv(RESULTS / "CSB_TRO_module_latent_bootstrap_raw.tsv", sep="\t", index=False)

    paired, per_dmr = paired_error_tests(predictions, rng)
    paired.to_csv(RESULTS / "CSB_TRO_module_latent_paired_error_tests.tsv", sep="\t", index=False)
    per_dmr.to_csv(RESULTS / "CSB_TRO_module_latent_per_DMR_errors.tsv", sep="\t", index=False)

    summary = validation_summary(observed, ci, paired, null_summary)
    summary.to_csv(RESULTS / "CSB_TRO_module_latent_validation_summary.tsv", sep="\t", index=False)

    svg_validation(FIGURES / "CSB_TRO_module_latent_validation_rmse.svg", observed, ci, null_summary)
    write_interpretation(DOCS / "CSB_TRO_module_latent_validation_interpretation.md", observed, ci, paired, null_summary)

    manifest = {
        "n_null": args.n_null,
        "n_bootstrap": args.n_bootstrap,
        "seed": args.seed,
        "outputs": [
            str(RESULTS / "CSB_TRO_module_latent_validation_summary.tsv"),
            str(RESULTS / "CSB_TRO_module_latent_bootstrap_CI.tsv"),
            str(RESULTS / "CSB_TRO_module_latent_null_models.tsv"),
            str(RESULTS / "CSB_TRO_module_latent_paired_error_tests.tsv"),
            str(FIGURES / "CSB_TRO_module_latent_validation_rmse.svg"),
            str(DOCS / "CSB_TRO_module_latent_validation_interpretation.md"),
        ],
    }
    (RESULTS / "CSB_TRO_module_latent_validation_manifest.json").write_text(json.dumps(manifest, indent=2), encoding="utf-8")
    print(json.dumps({
        "observed": summary.to_dict(orient="records"),
        "paired_tests": paired.to_dict(orient="records"),
        "null_summary": null_summary.to_dict(orient="records"),
    }, indent=2))


if __name__ == "__main__":
    main()
```
