## Supplemental code for "An entropy-based diagnostic framework for characterizing methylation state dynamics during preimplantation development": run_module_linked_rna_control_variants.py

```
from __future__ import annotations

import json
from pathlib import Path

import pandas as pd


BASE = Path(r"E:\CSB_TRO_operator_time_DMR_dynamics_2026-05-25")
RESULTS = BASE / "results"
DOCS = BASE / "docs"

SOURCE = RESULTS / "CSB_TRO_module_linked_RNA_activity.tsv"
PRIORITY_MODULES = ["M05", "M01", "M12", "M02", "M10"]


def zscore(s: pd.Series) -> pd.Series:
    x = pd.to_numeric(s, errors="coerce").astype(float)
    sd = x.std()
    if not sd or pd.isna(sd):
        return x * 0.0
    return (x - x.mean()) / sd


def write_variant(name: str, sign: float, interpretation: str) -> str:
    tab = pd.read_csv(SOURCE, sep="\t")
    tab = tab[(tab["module_id"].isin(PRIORITY_MODULES)) & (tab["control_stage_window"] == "8-cell_to_morula")].copy()
    tab["control_value"] = pd.to_numeric(tab["control_value"], errors="coerce") * sign
    tab["control_value_z"] = zscore(tab["control_value"])
    tab["candidate_control"] = name + "_" + tab["module_id"].astype(str)
    tab["leakage_status"] = "methylation_non_leaking_gene_linked_RNA_nearest_TSS"
    tab["interpretation"] = interpretation
    for pc in ["PC1", "PC2", "PC3"]:
        tab[f"candidate_control_direction_{pc}"] = tab["control_value_z"] * pd.to_numeric(tab[f"latent_control_{pc}"], errors="coerce")
    out = RESULTS / f"CSB_TRO_{name}_features.tsv"
    tab.to_csv(out, sep="\t", index=False)
    return str(out)


def main() -> None:
    outputs = {
        "RNA_delta_8cell_to_morula_priority_modules": write_variant(
            "module_linked_RNA_delta_8cell_to_morula_priority",
            1.0,
            "Nearest-TSS linked gene expression change from 8-cell to morula for priority residual modules.",
        ),
        "RNA_repression_8cell_to_morula_priority_modules": write_variant(
            "module_linked_RNA_repression_8cell_to_morula_priority",
            -1.0,
            "Nearest-TSS linked gene expression repression activity, defined as negative 8-cell to morula RNA change for priority residual modules.",
        ),
    }
    doc = [
        "# Module-linked RNA control variants",
        "",
        "This creates focused nearest-TSS gene-linked RNA feature tables for priority residual modules only.",
        "",
        "Variants:",
        "- RNA delta: ΔRNA(8-cell to morula)",
        "- RNA repression: -ΔRNA(8-cell to morula)",
        "",
        "The repression variant is a biologically defined external variable; it does not use morula methylation residuals to fit beta.",
    ]
    (DOCS / "CSB_TRO_module_linked_RNA_control_variants.md").write_text("\n".join(doc) + "\n", encoding="utf-8")
    print(json.dumps(outputs, indent=2))


if __name__ == "__main__":
    main()
```
