## Supplemental code for "An entropy-based diagnostic framework for characterizing methylation state dynamics during preimplantation development": run_module_tf_activity_control_panel.py

```
from __future__ import annotations

import json
import sys
from pathlib import Path

import numpy as np
import pandas as pd


BASE = Path(r"E:\CSB_TRO_operator_time_DMR_dynamics_2026-05-25")
CODE = BASE / "code"
RESULTS = BASE / "results"
FIGURES = BASE / "figures"
DOCS = BASE / "docs"
sys.path.insert(0, str(CODE))

from run_basin_residual_control_field import PRE_MORULA_STAGES, cosine, simulate_strict_pre_morula  # noqa: E402
from run_morula_basin_sde import (  # noqa: E402
    basin_definition,
    decode_latent,
    distribution_metrics,
    fit_latent_basis,
    fit_operator,
    load_inputs,
    stage_ids,
)


MOTIF_ROWS = RESULTS / "CSB_TRO_residual_module_motif_TF_activity_matched_bg.tsv"
OLD_Q05_FEATURES = RESULTS / "CSB_TRO_motif_TF_activity_matched_bg_q05_control_features.tsv"
MODULE_BASIS = RESULTS / "CSB_TRO_missing_control_term_module_basis.tsv"

OUT_FEATURES = RESULTS / "CSB_TRO_module_TF_activity_control_panel_features.tsv"
OUT_METRICS = RESULTS / "CSB_TRO_module_TF_activity_control_panel_metrics.tsv"
OUT_RANDOM = RESULTS / "CSB_TRO_module_TF_activity_control_panel_random.tsv"
OUT_SUMMARY = RESULTS / "CSB_TRO_module_TF_activity_control_panel_summary.tsv"
OUT_MANIFEST = RESULTS / "CSB_TRO_module_TF_activity_control_panel_manifest.json"
OUT_SVG = FIGURES / "CSB_TRO_module_TF_activity_control_panel.svg"
OUT_DOC = DOCS / "CSB_TRO_module_TF_activity_control_panel_summary.md"

PRIORITY_MODULES = ["M05", "M01", "M12", "M02", "M10"]


def zscore_values(x: np.ndarray) -> np.ndarray:
    x = np.asarray(x, dtype=float)
    sd = np.nanstd(x, ddof=1)
    if not np.isfinite(sd) or sd == 0:
        return np.zeros_like(x)
    return (x - np.nanmean(x)) / sd


def basis_table() -> pd.DataFrame:
    basis = pd.read_csv(MODULE_BASIS, sep="\t")
    return basis[["module_id", "n_DMRs", "latent_control_PC1", "latent_control_PC2", "latent_control_PC3", "latent_control_norm", "ridge_weight"]].copy()


def dense_module_features(rows: pd.DataFrame, feature_set: str, selector: pd.Series, description: str, zero_fill: bool = True) -> pd.DataFrame:
    basis = basis_table()
    sub = rows[selector].copy()
    if len(sub):
        module = (
            sub.groupby("module_id", as_index=False)
            .agg(
                control_value=("tf_activity", "sum"),
                n_TF=("TF", "nunique"),
                top_TFs=("TF", lambda s: ",".join(list(dict.fromkeys(map(str, s)))[:20])),
                min_qvalue=("qvalue", "min"),
                max_log_odds_ratio=("log_odds_ratio", "max"),
            )
        )
    else:
        module = pd.DataFrame(columns=["module_id", "control_value", "n_TF", "top_TFs", "min_qvalue", "max_log_odds_ratio"])
    if zero_fill:
        module = basis[["module_id"]].merge(module, on="module_id", how="left")
        module["control_value"] = pd.to_numeric(module["control_value"], errors="coerce").fillna(0.0)
        module["n_TF"] = pd.to_numeric(module["n_TF"], errors="coerce").fillna(0).astype(int)
        module["top_TFs"] = module["top_TFs"].fillna("")
    module = module[module["module_id"].isin(PRIORITY_MODULES)].copy()
    module["control_value_z"] = zscore_values(module["control_value"].to_numpy(dtype=float))
    module["feature_set"] = feature_set
    module["description"] = description
    module["control_modality"] = "motif_TF_activity"
    module["leakage_status"] = "methylation_non_leaking_motif_x_TF_expression"
    module = module.merge(basis, on="module_id", how="left")
    return module


def sparse_features(rows: pd.DataFrame, feature_set: str, selector: pd.Series, polarity: float, description: str) -> pd.DataFrame:
    basis = basis_table()
    sub = rows[selector].copy()
    if not len(sub):
        return pd.DataFrame()
    module = (
        sub.groupby("module_id", as_index=False)
        .agg(
            control_value=("tf_activity", "sum"),
            n_TF=("TF", "nunique"),
            top_TFs=("TF", lambda s: ",".join(list(dict.fromkeys(map(str, s)))[:20])),
            min_qvalue=("qvalue", "min"),
            max_log_odds_ratio=("log_odds_ratio", "max"),
        )
    )
    module = module[module["module_id"].isin(PRIORITY_MODULES)].copy()
    module["control_value_z"] = polarity * np.sign(pd.to_numeric(module["control_value"], errors="coerce").fillna(0.0))
    module["feature_set"] = feature_set
    module["description"] = description
    module["control_modality"] = "motif_TF_activity"
    module["leakage_status"] = "methylation_non_leaking_sparse_motif_x_TF_expression"
    module = module.merge(basis, on="module_id", how="left")
    return module


def old_q05_features() -> pd.DataFrame:
    if not OLD_Q05_FEATURES.exists():
        return pd.DataFrame()
    old = pd.read_csv(OLD_Q05_FEATURES, sep="\t")
    old["feature_set"] = "old_q05_zero_filled_zscore"
    old["description"] = "Previous q<=0.05 feature encoding; retained as direct comparability control."
    old["control_modality"] = "motif_TF_activity"
    old["min_qvalue"] = np.nan
    old["max_log_odds_ratio"] = np.nan
    return old


def build_feature_panel() -> pd.DataFrame:
    rows = pd.read_csv(MOTIF_ROWS, sep="\t")
    rows["qvalue"] = pd.to_numeric(rows["qvalue"], errors="coerce").fillna(1.0)
    rows["log_odds_ratio"] = pd.to_numeric(rows["log_odds_ratio"], errors="coerce").fillna(0.0)
    rows["tf_activity"] = pd.to_numeric(rows["tf_activity"], errors="coerce").fillna(0.0)
    rows["module_id"] = rows["module_id"].astype(str)
    panel = []
    old = old_q05_features()
    if len(old):
        panel.append(old)
    panel.append(
        dense_module_features(
            rows,
            "all_motifs_zero_filled_zscore",
            rows["log_odds_ratio"].notna(),
            "All scanned motifs, module sums, zero-filled across priority modules and z-scored.",
            zero_fill=True,
        )
    )
    panel.append(
        dense_module_features(
            rows,
            "q20_zero_filled_zscore",
            rows["qvalue"] <= 0.20,
            "Relaxed q<=0.20 motif x TF activity, zero-filled and z-scored.",
            zero_fill=True,
        )
    )
    panel.append(
        dense_module_features(
            rows,
            "q10_zero_filled_zscore",
            rows["qvalue"] <= 0.10,
            "Relaxed q<=0.10 motif x TF activity, zero-filled and z-scored.",
            zero_fill=True,
        )
    )
    panel.append(
        dense_module_features(
            rows,
            "q05_zero_filled_zscore_rebuilt",
            rows["qvalue"] <= 0.05,
            "Rebuilt q<=0.05 motif x TF activity, zero-filled and z-scored.",
            zero_fill=True,
        )
    )
    panel.append(
        sparse_features(
            rows,
            "q05_sparse_activity_sign",
            rows["qvalue"] <= 0.05,
            polarity=1.0,
            description="Only modules with q<=0.05 motif hits; sign follows motif x TF activity.",
        )
    )
    panel.append(
        sparse_features(
            rows,
            "q05_sparse_flipped_activity_sign",
            rows["qvalue"] <= 0.05,
            polarity=-1.0,
            description="Only modules with q<=0.05 motif hits; activity sign flipped as orientation sensitivity control.",
        )
    )
    out = pd.concat([p for p in panel if len(p)], ignore_index=True)
    keep = [
        "feature_set",
        "module_id",
        "control_value",
        "control_value_z",
        "n_TF",
        "top_TFs",
        "min_qvalue",
        "max_log_odds_ratio",
        "n_DMRs",
        "latent_control_PC1",
        "latent_control_PC2",
        "latent_control_PC3",
        "latent_control_norm",
        "ridge_weight",
        "control_modality",
        "leakage_status",
        "description",
    ]
    return out[[c for c in keep if c in out.columns]].copy()


def evaluate_feature_set(name: str, sub: pd.DataFrame, strict_z: np.ndarray, obs_z: np.ndarray, obs_dmr: np.ndarray, mu, sd, components, basin, residual_z, rng) -> list[dict[str, object]]:
    rows = []
    u = pd.to_numeric(sub["control_value_z"], errors="coerce").fillna(0.0).to_numpy(dtype=float)
    dirs = sub[["latent_control_PC1", "latent_control_PC2", "latent_control_PC3"]].to_numpy(dtype=float)
    control = (dirs * u[:, None]).sum(axis=0)
    for model_name, vec, status in [
        (name, control, "feature_defined"),
        (name + "_sign_flip", -control, "sign_flip_control"),
    ]:
        pred_z = strict_z + vec[None, :]
        pred_dmr = decode_latent(pred_z, mu, sd, components)
        row = {
            "feature_set": name,
            "control_model": model_name,
            "validation_status": status,
            "n_modules": int(len(sub)),
            "nonzero_modules": int(np.sum(np.abs(u) > 1e-12)),
            "modules": ",".join(sub.loc[np.abs(u) > 1e-12, "module_id"].astype(str).tolist()),
            "top_TFs": ";".join([f"{r.module_id}:{r.top_TFs}" for r in sub.itertuples() if str(getattr(r, "top_TFs", ""))]),
            "PC1_control": float(vec[0]),
            "PC2_control": float(vec[1]),
            "PC3_control": float(vec[2]),
            "control_norm": float(np.linalg.norm(vec)),
            "direction_cosine_to_measured_correction": cosine(vec, residual_z),
            "PC3_negative_pull_recovered": float(-vec[2] / (-residual_z[2])) if residual_z[2] < 0 else float("nan"),
            **distribution_metrics(pred_z, obs_z, pred_dmr, obs_dmr, basin, rng),
        }
        rows.append(row)
    return rows


def random_controls(features: pd.DataFrame, strict_z, obs_z, obs_dmr, mu, sd, components, basin, residual_z) -> pd.DataFrame:
    rng = np.random.default_rng(20260526)
    rows = []
    for feature_set, sub in features.groupby("feature_set"):
        u0 = pd.to_numeric(sub["control_value_z"], errors="coerce").fillna(0.0).to_numpy(dtype=float)
        dirs = sub[["latent_control_PC1", "latent_control_PC2", "latent_control_PC3"]].to_numpy(dtype=float)
        for i in range(200):
            u = rng.permutation(u0)
            vec = (dirs * u[:, None]).sum(axis=0)
            pred_z = strict_z + vec[None, :]
            pred_dmr = decode_latent(pred_z, mu, sd, components)
            rows.append(
                {
                    "feature_set": feature_set,
                    "random_iter": i,
                    "PC1_control": float(vec[0]),
                    "PC2_control": float(vec[1]),
                    "PC3_control": float(vec[2]),
                    "direction_cosine_to_measured_correction": cosine(vec, residual_z),
                    **distribution_metrics(pred_z, obs_z, pred_dmr, obs_dmr, basin, rng),
                }
            )
    return pd.DataFrame(rows)


def make_svg(metrics: pd.DataFrame, random_df: pd.DataFrame) -> None:
    rows = metrics[metrics["validation_status"] == "feature_defined"].sort_values("pred_basin_occupancy_q90", ascending=False)
    width, height = 960, 430
    left, right, top, bottom = 80, 30, 45, 150
    plot_w = width - left - right
    plot_h = height - top - bottom
    gap = plot_w / max(len(rows), 1)
    bar_w = gap * 0.58
    lines = [
        f'<svg xmlns="http://www.w3.org/2000/svg" width="{width}" height="{height}" viewBox="0 0 {width} {height}">',
        '<rect width="100%" height="100%" fill="white"/>',
        f'<text x="{left}" y="28" font-family="Arial" font-size="18" font-weight="700">Module TF activity control panel</text>',
        f'<line x1="{left}" y1="{height-bottom}" x2="{width-right}" y2="{height-bottom}" stroke="#222"/>',
        f'<line x1="{left}" y1="{top}" x2="{left}" y2="{height-bottom}" stroke="#222"/>',
    ]
    for tick in [0, 0.044, 0.2, 0.222, 0.5, 0.875, 1.0]:
        y = height - bottom - tick * plot_h
        color = "#b23b3b" if tick in [0.044, 0.875] else "#dddddd"
        lines.append(f'<line x1="{left}" y1="{y:.2f}" x2="{width-right}" y2="{y:.2f}" stroke="{color}" stroke-dasharray="4 4"/>')
        lines.append(f'<text x="{left-8}" y="{y+4:.2f}" text-anchor="end" font-family="Arial" font-size="10">{tick:.3g}</text>')
    for i, row in enumerate(rows.itertuples()):
        x = left + i * gap + (gap - bar_w) / 2
        val = float(row.pred_basin_occupancy_q90)
        y = height - bottom - val * plot_h
        h = val * plot_h
        q95 = random_df[random_df["feature_set"] == row.feature_set]["pred_basin_occupancy_q90"].quantile(0.95)
        yq = height - bottom - float(q95) * plot_h if np.isfinite(q95) else height - bottom
        lines.append(f'<line x1="{x:.2f}" y1="{yq:.2f}" x2="{x+bar_w:.2f}" y2="{yq:.2f}" stroke="#111" stroke-width="2"/>')
        lines.append(f'<rect x="{x:.2f}" y="{y:.2f}" width="{bar_w:.2f}" height="{h:.2f}" fill="#245c7a"/>')
        lines.append(f'<text x="{x+bar_w/2:.2f}" y="{y-5:.2f}" text-anchor="middle" font-family="Arial" font-size="10">{val:.3f}</text>')
        label = row.feature_set.replace("_", " ")
        lines.append(f'<text x="{x+bar_w/2:.2f}" y="{height-bottom+18}" text-anchor="end" transform="rotate(-45 {x+bar_w/2:.2f} {height-bottom+18})" font-family="Arial" font-size="9">{label}</text>')
    lines.append(f'<text x="{left}" y="{height-22}" font-family="Arial" font-size="12">Black tick on each bar: shuffled module-value q95. Red guides: baseline and observed morula occupancy.</text>')
    lines.append("</svg>")
    OUT_SVG.write_text("\n".join(lines), encoding="utf-8")


def write_doc(summary: pd.DataFrame, metrics: pd.DataFrame) -> None:
    best = summary.sort_values("pred_basin_occupancy_q90", ascending=False).iloc[0]
    sparse = summary[summary["feature_set"].str.contains("sparse", regex=False)]
    lines = [
        "# Module TF Activity Control Panel",
        "",
        "Status: `completed`",
        "",
        "Goal: test whether module-level motif x TF expression activity for M05/M01/M12/M02/M10 robustly explains the measured morula correction direction.",
        "",
        "## Main Result",
        "",
        f"Best feature-defined model: `{best.feature_set}`",
        f"- occupancy_q90: {best.pred_basin_occupancy_q90:.3f}",
        f"- cosine: {best.direction_cosine_to_measured_correction:.3f}",
        f"- PC3 recovery: {best.PC3_negative_pull_recovered:.3f}",
        f"- shuffled q95 occupancy: {best.random_q95_occupancy:.3f}",
        "",
        "## Encoding Sensitivity",
        "",
        "The panel explicitly separates zero-filled z-scored module encodings from sparse encodings that include only modules with significant motif evidence.",
    ]
    if len(sparse):
        for row in sparse.itertuples():
            lines.append(
                f"- {row.feature_set}: occupancy={row.pred_basin_occupancy_q90:.3f}, cosine={row.direction_cosine_to_measured_correction:.3f}, PC3_recovery={row.PC3_negative_pull_recovered:.3f}"
            )
    lines += [
        "",
        "Interpretation boundary: if rescue is strong only in zero-filled/z-scored encodings and not sparse significant-motif encodings, M02-KLF4/KLF5 remains exploratory and needs independent FIMO/HOMER plus chromatin-state validation.",
    ]
    OUT_DOC.write_text("\n".join(lines) + "\n", encoding="utf-8")


def main() -> None:
    RESULTS.mkdir(parents=True, exist_ok=True)
    FIGURES.mkdir(parents=True, exist_ok=True)
    DOCS.mkdir(parents=True, exist_ok=True)

    features = build_feature_panel()
    features.to_csv(OUT_FEATURES, sep="\t", index=False)

    matrix, ann, pairs = load_inputs()
    mu, sd, components, score_df = fit_latent_basis(matrix, 3)
    coef, _, _ = fit_operator(score_df, pairs, train_stages=set(PRE_MORULA_STAGES), lam=1000.0)
    strict_z = simulate_strict_pre_morula(score_df, ann, coef, n_steps=12)
    obs_z = score_df.loc[stage_ids(ann, "morula")].to_numpy(dtype=float)
    obs_dmr = matrix.loc[stage_ids(ann, "morula")].to_numpy(dtype=float)
    basin = basin_definition(obs_z)
    residual_z = obs_z.mean(axis=0) - strict_z.mean(axis=0)

    rng = np.random.default_rng(20260526)
    metric_rows = []
    for feature_set, sub in features.groupby("feature_set"):
        metric_rows.extend(evaluate_feature_set(feature_set, sub, strict_z, obs_z, matrix.loc[stage_ids(ann, "morula")].to_numpy(dtype=float), mu, sd, components, basin, residual_z, rng))
    metrics = pd.DataFrame(metric_rows)
    random_df = random_controls(features, strict_z, obs_z, matrix.loc[stage_ids(ann, "morula")].to_numpy(dtype=float), mu, sd, components, basin, residual_z)
    metrics.to_csv(OUT_METRICS, sep="\t", index=False)
    random_df.to_csv(OUT_RANDOM, sep="\t", index=False)

    defined = metrics[metrics["validation_status"] == "feature_defined"].copy()
    random_summary = (
        random_df.groupby("feature_set", as_index=False)
        .agg(random_mean_occupancy=("pred_basin_occupancy_q90", "mean"), random_q95_occupancy=("pred_basin_occupancy_q90", lambda x: float(np.quantile(x, 0.95))), random_max_occupancy=("pred_basin_occupancy_q90", "max"))
    )
    summary = defined.merge(random_summary, on="feature_set", how="left").sort_values("pred_basin_occupancy_q90", ascending=False)
    summary.to_csv(OUT_SUMMARY, sep="\t", index=False)
    make_svg(metrics, random_df)
    write_doc(summary, metrics)
    OUT_MANIFEST.write_text(
        json.dumps(
            {
                "status": "completed",
                "outputs": [str(OUT_FEATURES), str(OUT_METRICS), str(OUT_RANDOM), str(OUT_SUMMARY), str(OUT_SVG), str(OUT_DOC)],
                "boundary": "panel tests encoding sensitivity; independent motif scanner and chromatin-state validation still required",
            },
            indent=2,
        )
        + "\n",
        encoding="utf-8",
    )
    print(json.dumps({"status": "completed", "summary": summary.head(10).to_dict(orient="records")}, indent=2))


if __name__ == "__main__":
    main()
```
