## Supplemental code for "An entropy-based diagnostic framework for characterizing methylation state dynamics during preimplantation development": run_morula_basin_sde.py

```
from __future__ import annotations

import argparse
import json
import math
from pathlib import Path

import numpy as np
import pandas as pd


BASE = Path(r"E:\CSB_TRO_operator_time_DMR_dynamics_2026-05-25")
RESULTS = BASE / "results"
FIGURES = BASE / "figures"
DOCS = BASE / "docs"

STAGES = ["MII oocyte", "zygote/PN", "2-cell", "4-cell", "8-cell", "morula", "blastocyst"]
TAU = {stage: i / (len(STAGES) - 1) for i, stage in enumerate(STAGES)}


def rmse(a, b) -> float:
    a = np.asarray(a, dtype=float)
    b = np.asarray(b, dtype=float)
    return float(np.sqrt(np.mean((a - b) ** 2)))


def corr(a, b) -> float:
    a = np.asarray(a, dtype=float)
    b = np.asarray(b, dtype=float)
    aa = a - a.mean()
    bb = b - b.mean()
    den = math.sqrt(float((aa @ aa) * (bb @ bb)))
    return float((aa @ bb) / den) if den else float("nan")


def ridge(x: np.ndarray, y: np.ndarray, w: np.ndarray, lam: float) -> np.ndarray:
    sw = np.sqrt(np.maximum(w, 1e-12) / np.mean(np.maximum(w, 1e-12)))
    xw = x * sw[:, None]
    yw = y * sw[:, None] if y.ndim == 2 else y * sw
    penalty = lam * np.eye(x.shape[1])
    penalty[0, 0] = 0.0
    return np.linalg.solve(xw.T @ xw + penalty, xw.T @ yw)


def load_inputs():
    matrix = pd.read_csv(RESULTS / "CSB_TRO_DMR_state_matrix.tsv", sep="\t").set_index("sample_id")
    ann = pd.read_csv(RESULTS / "CSB_TRO_sample_tau_annotation.tsv", sep="\t").set_index("sample_id")
    pairs = pd.read_csv(RESULTS / "CSB_TRO_OT_sample_transition_couplings.tsv", sep="\t")
    pairs = pairs[pairs["from_sample_id"].isin(matrix.index) & pairs["to_sample_id"].isin(matrix.index)].copy()
    pairs = pairs[np.abs(pairs["delta_tau"].to_numpy(dtype=float)) > 1e-9].copy()
    return matrix, ann, pairs


def stage_ids(ann: pd.DataFrame, stage: str) -> list[str]:
    return list(ann.index[ann["stage"] == stage])


def fit_latent_basis(matrix: pd.DataFrame, q: int):
    mu = matrix.mean(axis=0).to_numpy(dtype=float)
    sd = matrix.std(axis=0).to_numpy(dtype=float) + 1e-6
    x = (matrix.to_numpy(dtype=float) - mu) / sd
    _, _, vt = np.linalg.svd(x, full_matrices=False)
    components = vt[:q].T
    scores = x @ components
    score_df = pd.DataFrame(scores, index=matrix.index, columns=[f"PC{i + 1}" for i in range(q)])
    return mu, sd, components, score_df


def decode_latent(z: np.ndarray, mu: np.ndarray, sd: np.ndarray, components: np.ndarray) -> np.ndarray:
    return np.clip((np.asarray(z, dtype=float) @ components.T) * sd + mu, 0.0, 1.0)


def fit_operator(score_df: pd.DataFrame, pairs: pd.DataFrame, train_stages: set[str] | None, lam: float):
    train = pairs.copy()
    if train_stages is not None:
        train = train[train["from_stage"].isin(train_stages) & train["to_stage"].isin(train_stages)].copy()
    zf = score_df.loc[train["from_sample_id"]].to_numpy(dtype=float)
    zt = score_df.loc[train["to_sample_id"]].to_numpy(dtype=float)
    dt = train["delta_tau"].to_numpy(dtype=float)[:, None]
    y = (zt - zf) / dt
    x = np.column_stack([np.ones(len(train)), train["from_tau"].to_numpy(dtype=float), zf])
    coef = ridge(x, y, train["sample_coupling_weight"].to_numpy(dtype=float), lam)
    residual = y - x @ coef
    cov = regularize_cov(np.cov(residual.T), residual.shape[1])
    train = train.copy()
    for i in range(residual.shape[1]):
        train[f"resid_PC{i + 1}"] = residual[:, i]
    return coef, cov, train


def regularize_cov(cov: np.ndarray, q: int) -> np.ndarray:
    cov = np.asarray(cov, dtype=float)
    if cov.ndim == 0:
        cov = np.eye(q) * float(cov)
    return cov + np.eye(q) * 1e-8


def drift(z: np.ndarray, tau: float, coef: np.ndarray) -> np.ndarray:
    x = np.concatenate([[1.0, tau], np.asarray(z, dtype=float)])
    return x @ coef


def drift_batch(z: np.ndarray, tau: float, coef: np.ndarray) -> np.ndarray:
    z = np.asarray(z, dtype=float)
    x = np.column_stack([np.ones(len(z)), np.full(len(z), float(tau)), z])
    return x @ coef


def euler_rollout(
    z0: np.ndarray,
    tau0: float,
    tau1: float,
    coef: np.ndarray,
    rng: np.random.Generator,
    n_steps: int,
    diffusion_cov: np.ndarray | None = None,
    diffusion_scale: float = 1.0,
    basin_center: np.ndarray | None = None,
    kappa: float = 0.0,
) -> np.ndarray:
    z = np.asarray(z0, dtype=float).copy()
    tau = float(tau0)
    dt = (float(tau1) - float(tau0)) / float(n_steps)
    for _ in range(n_steps):
        v = drift(z, tau, coef)
        if basin_center is not None and kappa > 0:
            v = v + kappa * (basin_center - z)
        z = z + dt * v
        if diffusion_cov is not None and diffusion_scale > 0:
            z = z + rng.multivariate_normal(np.zeros(len(z)), diffusion_cov * dt * diffusion_scale)
        tau += dt
    return z


def euler_rollout_batch(
    z0: np.ndarray,
    tau0: float,
    tau1: float,
    coef: np.ndarray,
    rng: np.random.Generator,
    n_steps: int,
    diffusion_cov: np.ndarray | None = None,
    diffusion_scale: float = 1.0,
    basin_center: np.ndarray | None = None,
    kappa: float = 0.0,
) -> np.ndarray:
    z = np.asarray(z0, dtype=float).copy()
    tau = float(tau0)
    dt = (float(tau1) - float(tau0)) / float(n_steps)
    for _ in range(n_steps):
        v = drift_batch(z, tau, coef)
        if basin_center is not None and kappa > 0:
            v = v + kappa * (basin_center[None, :] - z)
        z = z + dt * v
        if diffusion_cov is not None and diffusion_scale > 0:
            z = z + rng.multivariate_normal(
                np.zeros(z.shape[1]),
                diffusion_cov * dt * diffusion_scale,
                size=len(z),
            )
        tau += dt
    return z


def rbf_mmd(x: np.ndarray, y: np.ndarray) -> float:
    x = np.asarray(x, dtype=float)
    y = np.asarray(y, dtype=float)
    both = np.vstack([x, y])
    d2 = ((both[:, None, :] - both[None, :, :]) ** 2).sum(axis=2)
    med = np.median(d2[d2 > 0]) if np.any(d2 > 0) else 1.0
    gamma = 1.0 / (2.0 * med + 1e-12)
    kxx = np.exp(-gamma * ((x[:, None, :] - x[None, :, :]) ** 2).sum(axis=2)).mean()
    kyy = np.exp(-gamma * ((y[:, None, :] - y[None, :, :]) ** 2).sum(axis=2)).mean()
    kxy = np.exp(-gamma * ((x[:, None, :] - y[None, :, :]) ** 2).sum(axis=2)).mean()
    return float(max(kxx + kyy - 2 * kxy, 0.0))


def sliced_wasserstein(x: np.ndarray, y: np.ndarray, rng: np.random.Generator, n_proj: int = 128) -> float:
    x = np.asarray(x, dtype=float)
    y = np.asarray(y, dtype=float)
    vals = []
    for _ in range(n_proj):
        v = rng.normal(size=x.shape[1])
        v = v / (np.linalg.norm(v) + 1e-12)
        px = np.sort(x @ v)
        py = np.sort(y @ v)
        n = max(len(px), len(py))
        q = np.linspace(0, 1, n)
        vals.append(np.mean(np.abs(np.quantile(px, q) - np.quantile(py, q))))
    return float(np.mean(vals))


def covariance_error(x: np.ndarray, y: np.ndarray) -> float:
    return float(np.linalg.norm(np.cov(np.asarray(x).T) - np.cov(np.asarray(y).T), ord="fro"))


def basin_definition(morula_z: np.ndarray) -> dict[str, object]:
    center = morula_z.mean(axis=0)
    dist = np.linalg.norm(morula_z - center, axis=1)
    return {
        "center": center,
        "radius_q90": float(np.quantile(dist, 0.90)),
        "radius_q95": float(np.quantile(dist, 0.95)),
        "observed_occupancy_q90": float(np.mean(dist <= np.quantile(dist, 0.90))),
        "observed_occupancy_q95": float(np.mean(dist <= np.quantile(dist, 0.95))),
        "observed_mean_distance": float(dist.mean()),
    }


def metric_sample(x: np.ndarray, rng: np.random.Generator, max_n: int = 1000) -> np.ndarray:
    x = np.asarray(x, dtype=float)
    if len(x) <= max_n:
        return x
    idx = rng.choice(len(x), size=max_n, replace=False)
    return x[idx]


def distribution_metrics(pred_z, obs_z, pred_dmr, obs_dmr, basin, rng) -> dict[str, float]:
    center = basin["center"]
    pred_dist = np.linalg.norm(pred_z - center, axis=1)
    obs_dist = np.linalg.norm(obs_z - center, axis=1)
    pred_z_metric = metric_sample(pred_z, rng)
    pred_dmr_metric = metric_sample(pred_dmr, rng)
    return {
        "latent_mean_rmse": rmse(pred_z.mean(axis=0), obs_z.mean(axis=0)),
        "latent_mmd_rbf": rbf_mmd(pred_z_metric, obs_z),
        "latent_sliced_wasserstein": sliced_wasserstein(pred_z_metric, obs_z, rng),
        "latent_covariance_frobenius": covariance_error(pred_z_metric, obs_z),
        "dmr_mean_rmse": rmse(pred_dmr.mean(axis=0), obs_dmr.mean(axis=0)),
        "dmr_correlation": corr(pred_dmr.mean(axis=0), obs_dmr.mean(axis=0)),
        "dmr_distributional_rmse": rmse(pred_dmr_metric, obs_dmr.mean(axis=0)[None, :]),
        "basin_radius_q90": float(basin["radius_q90"]),
        "basin_radius_q95": float(basin["radius_q95"]),
        "pred_basin_occupancy_q90": float(np.mean(pred_dist <= basin["radius_q90"])),
        "pred_basin_occupancy_q95": float(np.mean(pred_dist <= basin["radius_q95"])),
        "observed_basin_occupancy_q90": float(np.mean(obs_dist <= basin["radius_q90"])),
        "observed_basin_occupancy_q95": float(np.mean(obs_dist <= basin["radius_q95"])),
        "pred_mean_distance_to_morula_centroid": float(pred_dist.mean()),
        "observed_mean_distance_to_morula_centroid": float(obs_dist.mean()),
    }


def residual_cov_for_stage(train: pd.DataFrame, q: int, from_stage: str, to_stage: str):
    resid_cols = [f"resid_PC{i + 1}" for i in range(q)]
    exact = train[(train["from_stage"] == from_stage) & (train["to_stage"] == to_stage)]
    if len(exact) >= q + 2:
        return regularize_cov(np.cov(exact[resid_cols].to_numpy(dtype=float).T), q), "exact_8cell_to_morula_residual"
    fallback = train[train["from_stage"] == "4-cell"]
    if len(fallback) >= q + 2:
        return regularize_cov(np.cov(fallback[resid_cols].to_numpy(dtype=float).T), q), "fallback_4cell_to_8cell_residual"
    return regularize_cov(np.cov(train[resid_cols].to_numpy(dtype=float).T), q), "fallback_all_training_residual"


def endpoint_noise_cov(score_df: pd.DataFrame, pairs: pd.DataFrame, coef: np.ndarray, q: int):
    sub = pairs[(pairs["from_stage"] == "8-cell") & (pairs["to_stage"] == "morula")].copy()
    if len(sub) < q + 2:
        return None
    residuals = []
    for row in sub.itertuples():
        z0 = score_df.loc[row.from_sample_id].to_numpy(dtype=float)
        z1 = score_df.loc[row.to_sample_id].to_numpy(dtype=float)
        zdet = z0 + float(row.delta_tau) * drift(z0, float(row.from_tau), coef)
        residuals.append(z1 - zdet)
    return regularize_cov(np.cov(np.vstack(residuals).T), q)


def simulate_particles(
    score_df: pd.DataFrame,
    ann: pd.DataFrame,
    coef: np.ndarray,
    rng: np.random.Generator,
    n_particles_per_start: int,
    n_steps: int,
    diffusion_cov: np.ndarray | None = None,
    diffusion_scale: float = 1.0,
    basin_center: np.ndarray | None = None,
    kappa: float = 0.0,
    endpoint_cov: np.ndarray | None = None,
):
    start_samples = stage_ids(ann, "8-cell")
    start_z = score_df.loc[start_samples].to_numpy(dtype=float)
    z0 = np.repeat(start_z, n_particles_per_start, axis=0)
    pred = euler_rollout_batch(
        z0,
        TAU["8-cell"],
        TAU["morula"],
        coef,
        rng,
        n_steps=n_steps,
        diffusion_cov=diffusion_cov,
        diffusion_scale=diffusion_scale,
        basin_center=basin_center,
        kappa=kappa,
    )
    if endpoint_cov is not None:
        pred = pred + rng.multivariate_normal(np.zeros(pred.shape[1]), endpoint_cov, size=len(pred))
    meta = pd.DataFrame({
        "particle_id": np.arange(len(pred), dtype=int),
        "start_sample_id": np.repeat(start_samples, n_particles_per_start),
        "replicate": np.tile(np.arange(n_particles_per_start, dtype=int), len(start_samples)),
    })
    return pred, meta


def calibrate_ou(score_df, ann, coef, cov, basin, target_occ, seed, n_steps, n_calib):
    center = basin["center"]
    grid = []
    for kappa in [0.0, 1.0, 2.0, 4.0, 6.0, 8.0, 12.0, 16.0, 24.0, 32.0]:
        for scale in [0.0, 0.05, 0.10, 0.25, 0.50, 1.00, 1.50, 2.00]:
            rng = np.random.default_rng(seed + int(kappa * 1000) + int(scale * 100))
            pred_z, _ = simulate_particles(
                score_df,
                ann,
                coef,
                rng,
                n_particles_per_start=n_calib,
                n_steps=n_steps,
                diffusion_cov=cov if scale > 0 else None,
                diffusion_scale=scale,
                basin_center=center,
                kappa=kappa,
            )
            dist = np.linalg.norm(pred_z - center, axis=1)
            occ = float(np.mean(dist <= basin["radius_q90"]))
            mean_dist = float(dist.mean())
            objective = abs(occ - target_occ) + 0.05 * abs(mean_dist - basin["observed_mean_distance"])
            grid.append({
                "kappa": kappa,
                "diffusion_scale": scale,
                "calibration_occupancy_q90": occ,
                "calibration_mean_distance": mean_dist,
                "calibration_objective": objective,
            })
    return pd.DataFrame(grid).sort_values("calibration_objective").iloc[0].to_dict()


def local_stability(score_df: pd.DataFrame, ann: pd.DataFrame, pairs: pd.DataFrame, coef: np.ndarray, basin: dict, protocol: str):
    center = basin["center"]
    rows = []
    sample_sets = {
        "morula_samples": stage_ids(ann, "morula"),
        "morula_plus_blastocyst_samples": stage_ids(ann, "morula") + stage_ids(ann, "blastocyst"),
    }
    for set_name, ids in sample_sets.items():
        radial = []
        tangential = []
        norm_v = []
        dist_vals = []
        for sample in ids:
            z = score_df.loc[sample].to_numpy(dtype=float)
            to_center = center - z
            dist = np.linalg.norm(to_center)
            if dist < 1e-12:
                continue
            u = to_center / dist
            v = drift(z, TAU["morula"], coef)
            rv = float(v @ u)
            tv = float(np.linalg.norm(v - rv * u))
            radial.append(rv)
            tangential.append(tv)
            norm_v.append(float(np.linalg.norm(v)))
            dist_vals.append(float(dist))
        rows.append({
            "train_protocol": protocol,
            "neighborhood": set_name,
            "n_samples": len(radial),
            "mean_distance_to_centroid": float(np.mean(dist_vals)) if dist_vals else float("nan"),
            "mean_inward_radial_drift": float(np.mean(radial)) if radial else float("nan"),
            "mean_tangential_drift": float(np.mean(tangential)) if tangential else float("nan"),
            "contraction_score": float(np.mean(np.asarray(radial) / (np.asarray(norm_v) + 1e-12))) if radial else float("nan"),
            "local_jacobian_max_real_eigenvalue": float(np.max(np.real(np.linalg.eigvals(coef[2 : 2 + coef.shape[1], :].T)))),
        })

    empirical = pairs[(pairs["from_stage"] == "8-cell") & (pairs["to_stage"] == "morula")]
    radial = []
    tangential = []
    for row in empirical.itertuples():
        z0 = score_df.loc[row.from_sample_id].to_numpy(dtype=float)
        z1 = score_df.loc[row.to_sample_id].to_numpy(dtype=float)
        midpoint = 0.5 * (z0 + z1)
        to_center = center - midpoint
        dist = np.linalg.norm(to_center)
        if dist < 1e-12:
            continue
        u = to_center / dist
        v = (z1 - z0) / float(row.delta_tau)
        rv = float(v @ u)
        radial.append(rv)
        tangential.append(float(np.linalg.norm(v - rv * u)))
    rows.append({
        "train_protocol": protocol,
        "neighborhood": "empirical_8cell_to_morula_OT_pairs_fullfit_reference",
        "n_samples": len(radial),
        "mean_distance_to_centroid": float("nan"),
        "mean_inward_radial_drift": float(np.mean(radial)) if radial else float("nan"),
        "mean_tangential_drift": float(np.mean(tangential)) if tangential else float("nan"),
        "contraction_score": float(np.mean(np.asarray(radial) / (np.sqrt(np.asarray(radial) ** 2 + np.asarray(tangential) ** 2) + 1e-12))) if radial else float("nan"),
        "local_jacobian_max_real_eigenvalue": float("nan"),
    })
    return pd.DataFrame(rows)


def make_svg(path: Path, metrics: pd.DataFrame):
    rows = metrics.copy()
    order = [
        "deterministic",
        "stochastic_global_residual",
        "stochastic_stage_conditioned",
        "empirical_8cell_morula_noise_fullfit",
        "basin_ou_calibrated",
    ]
    rows["mode_order"] = rows["sde_mode"].map({m: i for i, m in enumerate(order)}).fillna(99)
    rows = rows.sort_values(["train_protocol", "mode_order"])
    width, height = 980, 460
    left, right, top, bottom = 85, 25, 45, 145
    plot_w = width - left - right
    plot_h = height - top - bottom
    ymax = 1.0
    bar_w = plot_w / max(len(rows), 1) * 0.68
    gap = plot_w / max(len(rows), 1)
    out = [
        f'<svg xmlns="http://www.w3.org/2000/svg" width="{width}" height="{height}" viewBox="0 0 {width} {height}">',
        '<rect width="100%" height="100%" fill="white"/>',
        f'<text x="{left}" y="28" font-family="Arial" font-size="19" font-weight="700">Morula basin occupancy after 8-cell to morula latent SDE rollout</text>',
        f'<line x1="{left}" y1="{height-bottom}" x2="{width-right}" y2="{height-bottom}" stroke="#222"/>',
        f'<line x1="{left}" y1="{top}" x2="{left}" y2="{height-bottom}" stroke="#222"/>',
    ]
    for tick in [0, 0.25, 0.5, 0.75, 0.875, 1.0]:
        y = height - bottom - tick / ymax * plot_h
        color = "#b33" if abs(tick - 0.875) < 1e-6 else "#ddd"
        out.append(f'<line x1="{left}" y1="{y:.2f}" x2="{width-right}" y2="{y:.2f}" stroke="{color}" stroke-dasharray="4 4"/>')
        out.append(f'<text x="{left-8}" y="{y+4:.2f}" text-anchor="end" font-family="Arial" font-size="11">{tick:.3g}</text>')
    for i, row in enumerate(rows.itertuples()):
        x = left + i * gap + (gap - bar_w) / 2
        val = min(max(float(row.pred_basin_occupancy_q90), 0.0), 1.0)
        h = val / ymax * plot_h
        y = height - bottom - h
        fill = "#2f6f8f" if row.calibration_uses_morula_distribution == "no" else "#b56b2a"
        out.append(f'<rect x="{x:.2f}" y="{y:.2f}" width="{bar_w:.2f}" height="{h:.2f}" fill="{fill}"/>')
        out.append(f'<text x="{x+bar_w/2:.2f}" y="{y-5:.2f}" text-anchor="middle" font-family="Arial" font-size="10">{val:.3f}</text>')
        label = f"{row.train_protocol}|{row.sde_mode}".replace("_", " ")
        out.append(f'<text x="{x+bar_w/2:.2f}" y="{height-bottom+18}" text-anchor="end" transform="rotate(-45 {x+bar_w/2:.2f} {height-bottom+18})" font-family="Arial" font-size="10">{label}</text>')
    out.append(f'<text x="{left}" y="{height-20}" font-family="Arial" font-size="12">Red guide: observed morula q90 basin occupancy = 0.875. Orange bars use morula distribution calibration.</text>')
    out.append("</svg>")
    path.write_text("\n".join(out), encoding="utf-8")


def write_doc(path: Path, metrics: pd.DataFrame, stability: pd.DataFrame, basin: dict, calib: pd.DataFrame):
    show = metrics.sort_values(["train_protocol", "calibration_uses_morula_distribution", "sde_mode"])
    lines = [
        "# Morula basin SDE analysis",
        "",
        "This analysis keeps the developmental axis as stage-anchored operator time. It does not represent true longitudinal tracking of the same embryo.",
        "",
        f"Morula latent basin center was defined as the observed morula centroid. The q90 radius is {basin['radius_q90']:.4f}, giving observed q90 occupancy {basin['observed_occupancy_q90']:.3f}.",
        "",
        "8-cell to morula distribution-level results:",
    ]
    for row in show.itertuples():
        lines.append(
            f"- {row.train_protocol} / {row.sde_mode}: occupancy_q90={row.pred_basin_occupancy_q90:.3f}, "
            f"DMR_mean_RMSE={row.dmr_mean_rmse:.4f}, latent_MMD={row.latent_mmd_rbf:.4f}, "
            f"mean_dist={row.pred_mean_distance_to_morula_centroid:.4f}, uses_morula_calibration={row.calibration_uses_morula_distribution}"
        )
    lines.extend(["", "Local stability summary:"])
    for row in stability.itertuples():
        lines.append(
            f"- {row.train_protocol} / {row.neighborhood}: inward_radial={row.mean_inward_radial_drift:.4f}, "
            f"tangential={row.mean_tangential_drift:.4f}, contraction_score={row.contraction_score:.4f}, "
            f"max_real_eigen={row.local_jacobian_max_real_eigenvalue:.4f}"
        )
    if len(calib):
        lines.extend(["", "Basin correction calibration:"])
        for row in calib.itertuples():
            lines.append(
                f"- {row.train_protocol}: kappa={row.kappa:.4f}, diffusion_scale={row.diffusion_scale:.4f}, "
                f"calibration_occupancy_q90={row.calibration_occupancy_q90:.3f}, objective={row.calibration_objective:.4f}"
            )
    lines.extend([
        "",
        "Interpretation boundary: deterministic and residual/stage-conditioned SDE modes are forward latent simulations under the fitted operator. The basin-OU mode is a calibrated low-dimensional basin transition model when it uses the morula centroid/radius, so it must not be described as strict leave-morula-out extrapolation. In silico perturbations and basin corrections should be described as sensitivity/calibration analyses, not causal mechanism proof.",
    ])
    path.write_text("\n".join(lines) + "\n", encoding="utf-8")


def main():
    parser = argparse.ArgumentParser()
    parser.add_argument("--q", type=int, default=3)
    parser.add_argument("--lambda", dest="lam", type=float, default=1000.0)
    parser.add_argument("--particles-per-start", type=int, default=200)
    parser.add_argument("--calibration-particles-per-start", type=int, default=40)
    parser.add_argument("--n-steps", type=int, default=12)
    parser.add_argument("--seed", type=int, default=20260525)
    args = parser.parse_args()

    RESULTS.mkdir(parents=True, exist_ok=True)
    FIGURES.mkdir(parents=True, exist_ok=True)
    DOCS.mkdir(parents=True, exist_ok=True)

    matrix, ann, pairs = load_inputs()
    mu, sd, components, score_df = fit_latent_basis(matrix, args.q)
    obs_ids = stage_ids(ann, "morula")
    obs_z = score_df.loc[obs_ids].to_numpy(dtype=float)
    obs_dmr = matrix.loc[obs_ids].to_numpy(dtype=float)
    basin = basin_definition(obs_z)

    protocols = [
        ("full_all_transitions", None),
        ("leave_morula_transitions", set(stage for stage in STAGES if stage != "morula")),
    ]
    metrics_rows = []
    particle_parts = []
    diffusion_rows = []
    stability_parts = []
    calibration_rows = []

    for p_i, (protocol, train_stages) in enumerate(protocols):
        coef, global_cov, train = fit_operator(score_df, pairs, train_stages=train_stages, lam=args.lam)
        stage_cov, stage_cov_source = residual_cov_for_stage(train, args.q, "8-cell", "morula")
        endpoint_cov = endpoint_noise_cov(score_df, pairs, coef, args.q) if protocol == "full_all_transitions" else None

        diffusion_rows.append({"train_protocol": protocol, "diffusion_model": "global_residual_velocity", "source": "all_training_pairs", **cov_to_row(global_cov)})
        diffusion_rows.append({"train_protocol": protocol, "diffusion_model": "stage_conditioned_velocity", "source": stage_cov_source, **cov_to_row(stage_cov)})
        if endpoint_cov is not None:
            diffusion_rows.append({"train_protocol": protocol, "diffusion_model": "empirical_8cell_morula_endpoint", "source": "fullfit_8cell_to_morula_pairs", **cov_to_row(endpoint_cov)})

        stability_parts.append(local_stability(score_df, ann, pairs, coef, basin, protocol))

        calib = calibrate_ou(
            score_df,
            ann,
            coef,
            stage_cov,
            basin,
            basin["observed_occupancy_q90"],
            args.seed + p_i * 10000,
            args.n_steps,
            args.calibration_particles_per_start,
        )
        calib["train_protocol"] = protocol
        calibration_rows.append(calib)

        modes = [
            ("deterministic", None, 0.0, None, 0.0, None, "no"),
            ("stochastic_global_residual", global_cov, 1.0, None, 0.0, None, "no"),
            ("stochastic_stage_conditioned", stage_cov, 1.0, None, 0.0, None, "no"),
            ("basin_ou_calibrated", stage_cov if calib["diffusion_scale"] > 0 else None, float(calib["diffusion_scale"]), basin["center"], float(calib["kappa"]), None, "yes"),
        ]
        if endpoint_cov is not None:
            modes.insert(3, ("empirical_8cell_morula_noise_fullfit", None, 0.0, None, 0.0, endpoint_cov, "yes"))

        for m_i, (mode, cov, scale, center, kappa, end_cov, uses_morula) in enumerate(modes):
            rng = np.random.default_rng(args.seed + p_i * 1000 + m_i)
            reps = 1 if mode == "deterministic" else args.particles_per_start
            pred_z, meta = simulate_particles(
                score_df,
                ann,
                coef,
                rng,
                n_particles_per_start=reps,
                n_steps=args.n_steps,
                diffusion_cov=cov,
                diffusion_scale=scale,
                basin_center=center,
                kappa=kappa,
                endpoint_cov=end_cov,
            )
            pred_dmr = decode_latent(pred_z, mu, sd, components)
            local_rng = np.random.default_rng(args.seed + p_i * 1000 + m_i + 50000)
            met = distribution_metrics(pred_z, obs_z, pred_dmr, obs_dmr, basin, local_rng)
            metrics_rows.append({
                "train_protocol": protocol,
                "sde_mode": mode,
                "start_stage": "8-cell",
                "target_stage": "morula",
                "n_start_samples": len(stage_ids(ann, "8-cell")),
                "particles_per_start": reps,
                "n_predicted_particles": len(pred_z),
                "calibration_uses_morula_distribution": uses_morula,
                "kappa_to_morula_centroid": float(kappa),
                "diffusion_scale": float(scale),
                **met,
            })

            center_vec = basin["center"]
            dist = np.linalg.norm(pred_z - center_vec, axis=1)
            part = pd.concat([meta, pd.DataFrame(pred_z, columns=[f"PC{i + 1}" for i in range(args.q)])], axis=1)
            part.insert(0, "sde_mode", mode)
            part.insert(0, "train_protocol", protocol)
            part["distance_to_morula_centroid"] = dist
            part["inside_basin_q90"] = dist <= basin["radius_q90"]
            part["inside_basin_q95"] = dist <= basin["radius_q95"]
            particle_parts.append(part)

    metrics = pd.DataFrame(metrics_rows)
    particles = pd.concat(particle_parts, ignore_index=True)
    diffusion = pd.DataFrame(diffusion_rows)
    stability = pd.concat(stability_parts, ignore_index=True)
    calibration = pd.DataFrame(calibration_rows)

    metrics.to_csv(RESULTS / "CSB_TRO_morula_basin_sde_metrics.tsv", sep="\t", index=False)
    particles.to_csv(RESULTS / "CSB_TRO_morula_basin_sde_particles.tsv", sep="\t", index=False)
    diffusion.to_csv(RESULTS / "CSB_TRO_morula_basin_sde_diffusion.tsv", sep="\t", index=False)
    stability.to_csv(RESULTS / "CSB_TRO_morula_basin_local_stability.tsv", sep="\t", index=False)
    calibration.to_csv(RESULTS / "CSB_TRO_morula_basin_sde_calibration.tsv", sep="\t", index=False)

    make_svg(FIGURES / "CSB_TRO_morula_basin_sde_occupancy.svg", metrics)
    write_doc(DOCS / "CSB_TRO_morula_basin_sde_interpretation.md", metrics, stability, basin, calibration)

    manifest = {
        "q": args.q,
        "ridge_lambda": args.lam,
        "particles_per_start": args.particles_per_start,
        "calibration_particles_per_start": args.calibration_particles_per_start,
        "n_steps": args.n_steps,
        "seed": args.seed,
        "morula_basin": {k: (v.tolist() if isinstance(v, np.ndarray) else v) for k, v in basin.items()},
        "outputs": [
            str(RESULTS / "CSB_TRO_morula_basin_local_stability.tsv"),
            str(RESULTS / "CSB_TRO_morula_basin_sde_metrics.tsv"),
            str(RESULTS / "CSB_TRO_morula_basin_sde_particles.tsv"),
            str(RESULTS / "CSB_TRO_morula_basin_sde_diffusion.tsv"),
            str(FIGURES / "CSB_TRO_morula_basin_sde_occupancy.svg"),
            str(DOCS / "CSB_TRO_morula_basin_sde_interpretation.md"),
        ],
    }
    (RESULTS / "CSB_TRO_morula_basin_sde_manifest.json").write_text(json.dumps(manifest, indent=2), encoding="utf-8")
    print(json.dumps({
        "morula_basin": manifest["morula_basin"],
        "metrics": metrics.to_dict(orient="records"),
        "calibration": calibration.to_dict(orient="records"),
    }, indent=2))


def cov_to_row(cov: np.ndarray) -> dict[str, float]:
    out = {}
    for i in range(cov.shape[0]):
        for j in range(cov.shape[1]):
            out[f"cov_PC{i + 1}_PC{j + 1}"] = float(cov[i, j])
    return out


if __name__ == "__main__":
    main()
```
