## Supplemental code for "An entropy-based diagnostic framework for characterizing methylation state dynamics during preimplantation development": run_nonleaking_distribution_dynamics.py

```
from __future__ import annotations

import argparse
import json
import sys
from pathlib import Path

import numpy as np
import pandas as pd


BASE = Path(r"E:\CSB_TRO_operator_time_DMR_dynamics_2026-05-25")
CODE = BASE / "code"
RESULTS = BASE / "results"
FIGURES = BASE / "figures"
DOCS = BASE / "docs"
sys.path.insert(0, str(CODE))

from run_morula_basin_sde import (  # noqa: E402
    STAGES,
    TAU,
    basin_definition,
    corr,
    decode_latent,
    distribution_metrics,
    euler_rollout_batch,
    fit_latent_basis,
    fit_operator,
    load_inputs,
    regularize_cov,
    ridge,
    rmse,
    stage_ids,
)


PRE_MORULA_STAGES = ["MII oocyte", "zygote/PN", "2-cell", "4-cell", "8-cell"]


def project_psd(cov: np.ndarray, min_eig: float = 1e-6) -> np.ndarray:
    cov = 0.5 * (np.asarray(cov, dtype=float) + np.asarray(cov, dtype=float).T)
    vals, vecs = np.linalg.eigh(cov)
    vals = np.maximum(vals, min_eig)
    return (vecs * vals) @ vecs.T


def fit_affine_kernel(score_df: pd.DataFrame, pairs: pd.DataFrame, train_stages: set[str], lam: float):
    train = pairs[pairs["from_stage"].isin(train_stages) & pairs["to_stage"].isin(train_stages)].copy()
    zf = score_df.loc[train["from_sample_id"]].to_numpy(dtype=float)
    zt = score_df.loc[train["to_sample_id"]].to_numpy(dtype=float)
    x = np.column_stack([np.ones(len(train)), train["from_tau"].to_numpy(dtype=float), zf])
    beta = ridge(x, zt, train["sample_coupling_weight"].to_numpy(dtype=float), lam)
    residual = zt - x @ beta
    cov = regularize_cov(np.cov(residual.T), residual.shape[1])
    return beta, cov, train


def apply_affine_kernel(z0: np.ndarray, tau0: float, beta: np.ndarray, rng: np.random.Generator, cov: np.ndarray | None, reps: int):
    z0_rep = np.repeat(z0, reps, axis=0)
    x = np.column_stack([np.ones(len(z0_rep)), np.full(len(z0_rep), tau0), z0_rep])
    pred = x @ beta
    if cov is not None:
        pred = pred + rng.multivariate_normal(np.zeros(pred.shape[1]), cov, size=len(pred))
    return pred


def stage_moments(score_df: pd.DataFrame, ann: pd.DataFrame, stages: list[str]):
    rows = []
    covs = {}
    for stage in stages:
        z = score_df.loc[stage_ids(ann, stage)].to_numpy(dtype=float)
        rows.append({"stage": stage, "tau": TAU[stage], **{f"mean_PC{i + 1}": z.mean(axis=0)[i] for i in range(z.shape[1])}})
        covs[stage] = regularize_cov(np.cov(z.T), z.shape[1])
    return pd.DataFrame(rows), covs


def extrapolate_stage_moments_from(score_df: pd.DataFrame, ann: pd.DataFrame, q: int, target_tau: float, stages: list[str], degree: int | None = None):
    moment_df, covs = stage_moments(score_df, ann, stages)
    tau = moment_df["tau"].to_numpy(dtype=float)
    degree = min(2 if degree is None else degree, len(tau) - 1)
    center = []
    for i in range(q):
        y = moment_df[f"mean_PC{i + 1}"].to_numpy(dtype=float)
        coef = np.polyfit(tau, y, deg=degree)
        center.append(float(np.polyval(coef, target_tau)))
    center = np.asarray(center, dtype=float)

    cov_stack = np.stack([covs[s] for s in stages])
    cov_pred = np.zeros((q, q), dtype=float)
    for i in range(q):
        for j in range(q):
            y = cov_stack[:, i, j]
            coef = np.polyfit(tau, y, deg=1)
            cov_pred[i, j] = np.polyval(coef, target_tau)
    cov_pred = project_psd(cov_pred)
    return center, cov_pred, moment_df


def extrapolate_stage_moments(score_df: pd.DataFrame, ann: pd.DataFrame, q: int, target_tau: float):
    return extrapolate_stage_moments_from(score_df, ann, q, target_tau, PRE_MORULA_STAGES, degree=2)


def simulate_from_8cell(score_df, ann, coef, rng, reps, n_steps, cov=None, scale=1.0, center=None, kappa=0.0):
    start_samples = stage_ids(ann, "8-cell")
    z0 = score_df.loc[start_samples].to_numpy(dtype=float)
    z0_rep = np.repeat(z0, reps, axis=0)
    pred = euler_rollout_batch(
        z0_rep,
        TAU["8-cell"],
        TAU["morula"],
        coef,
        rng,
        n_steps=n_steps,
        diffusion_cov=cov,
        diffusion_scale=scale,
        basin_center=center,
        kappa=kappa,
    )
    meta = pd.DataFrame({
        "particle_id": np.arange(len(pred), dtype=int),
        "start_sample_id": np.repeat(start_samples, reps),
        "replicate": np.tile(np.arange(reps, dtype=int), len(start_samples)),
    })
    return pred, meta


def validate_ou_on_4cell_to_8cell(score_df, ann, coef, cov, center_8, obs8_z, seed, n_steps, reps):
    z4 = score_df.loc[stage_ids(ann, "4-cell")].to_numpy(dtype=float)
    grid = []
    for kappa in [0.0, 0.5, 1.0, 2.0, 4.0, 8.0, 12.0, 16.0, 24.0, 32.0]:
        for scale in [0.0, 0.05, 0.10, 0.25, 0.50, 1.00, 1.50, 2.00]:
            rng = np.random.default_rng(seed + int(kappa * 1000) + int(scale * 100))
            z0 = np.repeat(z4, reps, axis=0)
            pred = euler_rollout_batch(
                z0,
                TAU["4-cell"],
                TAU["8-cell"],
                coef,
                rng,
                n_steps=n_steps,
                diffusion_cov=cov if scale > 0 else None,
                diffusion_scale=scale,
                basin_center=center_8,
                kappa=kappa,
            )
            objective = rmse(pred.mean(axis=0), obs8_z.mean(axis=0)) + 0.05 * np.linalg.norm(np.cov(pred.T) - np.cov(obs8_z.T), ord="fro")
            grid.append({
                "kappa": kappa,
                "diffusion_scale": scale,
                "validation_4cell_to_8cell_latent_mean_rmse": rmse(pred.mean(axis=0), obs8_z.mean(axis=0)),
                "validation_4cell_to_8cell_cov_error": float(np.linalg.norm(np.cov(pred.T) - np.cov(obs8_z.T), ord="fro")),
                "validation_objective": float(objective),
            })
    return pd.DataFrame(grid).sort_values("validation_objective").iloc[0].to_dict()


def calibrate_morula_upper_bound(score_df, ann, coef, cov, observed_basin, seed, n_steps, reps):
    center = observed_basin["center"]
    target = observed_basin["observed_occupancy_q90"]
    grid = []
    for kappa in [0.0, 1.0, 2.0, 4.0, 8.0, 12.0, 16.0, 24.0, 32.0, 48.0, 64.0]:
        for scale in [0.0, 0.05, 0.10, 0.25, 0.50, 1.00, 1.50, 2.00]:
            pred, _ = simulate_from_8cell(
                score_df,
                ann,
                coef,
                np.random.default_rng(seed + int(kappa * 1000) + int(scale * 100)),
                reps,
                n_steps,
                cov=cov if scale > 0 else None,
                scale=scale,
                center=center,
                kappa=kappa,
            )
            dist = np.linalg.norm(pred - center, axis=1)
            occ = float(np.mean(dist <= observed_basin["radius_q90"]))
            mean_dist = float(dist.mean())
            objective = abs(occ - target) + 0.05 * abs(mean_dist - observed_basin["observed_mean_distance"])
            grid.append({
                "upper_bound_kappa": kappa,
                "upper_bound_diffusion_scale": scale,
                "upper_bound_calibration_occupancy_q90": occ,
                "upper_bound_calibration_mean_distance": mean_dist,
                "upper_bound_calibration_objective": float(objective),
            })
    return pd.DataFrame(grid).sort_values("upper_bound_calibration_objective").iloc[0].to_dict()


def particle_table(protocol: str, model: str, pred_z: np.ndarray, meta: pd.DataFrame, morula_basin: dict):
    center = morula_basin["center"]
    dist = np.linalg.norm(pred_z - center, axis=1)
    part = pd.concat([meta.reset_index(drop=True), pd.DataFrame(pred_z, columns=[f"PC{i + 1}" for i in range(pred_z.shape[1])])], axis=1)
    part.insert(0, "model", model)
    part.insert(0, "protocol", protocol)
    part["distance_to_observed_morula_centroid"] = dist
    part["inside_observed_morula_q90"] = dist <= morula_basin["radius_q90"]
    return part


def metric_row(protocol, model, leakage, pred_z, obs_z, pred_dmr, obs_dmr, basin, rng, extra=None):
    row = {
        "protocol": protocol,
        "model": model,
        "uses_morula_distribution_in_training": leakage,
        "n_predicted_particles": len(pred_z),
        **distribution_metrics(pred_z, obs_z, pred_dmr, obs_dmr, basin, rng),
    }
    if extra:
        row.update(extra)
    return row


def make_svg(path: Path, metrics: pd.DataFrame):
    rows = metrics.copy()
    rows["leak_order"] = rows["uses_morula_distribution_in_training"].map({"no": 0, "yes_upper_bound": 1}).fillna(2)
    rows = rows.sort_values(["leak_order", "pred_basin_occupancy_q90"])
    width, height = 1050, 470
    left, right, top, bottom = 85, 25, 45, 155
    plot_w = width - left - right
    plot_h = height - top - bottom
    gap = plot_w / max(len(rows), 1)
    bar_w = gap * 0.62
    out = [
        f'<svg xmlns="http://www.w3.org/2000/svg" width="{width}" height="{height}" viewBox="0 0 {width} {height}">',
        '<rect width="100%" height="100%" fill="white"/>',
        f'<text x="{left}" y="28" font-family="Arial" font-size="19" font-weight="700">Non-leaking distribution dynamics: observed morula basin occupancy test</text>',
        f'<line x1="{left}" y1="{height-bottom}" x2="{width-right}" y2="{height-bottom}" stroke="#222"/>',
        f'<line x1="{left}" y1="{top}" x2="{left}" y2="{height-bottom}" stroke="#222"/>',
    ]
    for tick in [0, 0.25, 0.5, 0.75, 0.875, 1.0]:
        y = height - bottom - tick * plot_h
        color = "#b33" if abs(tick - 0.875) < 1e-6 else "#ddd"
        out.append(f'<line x1="{left}" y1="{y:.2f}" x2="{width-right}" y2="{y:.2f}" stroke="{color}" stroke-dasharray="4 4"/>')
        out.append(f'<text x="{left-8}" y="{y+4:.2f}" text-anchor="end" font-family="Arial" font-size="11">{tick:.3g}</text>')
    for i, row in enumerate(rows.itertuples()):
        x = left + i * gap + (gap - bar_w) / 2
        val = float(row.pred_basin_occupancy_q90)
        h = max(0.0, min(1.0, val)) * plot_h
        y = height - bottom - h
        fill = "#2f6f8f" if row.uses_morula_distribution_in_training == "no" else "#b56b2a"
        out.append(f'<rect x="{x:.2f}" y="{y:.2f}" width="{bar_w:.2f}" height="{h:.2f}" fill="{fill}"/>')
        out.append(f'<text x="{x+bar_w/2:.2f}" y="{y-5:.2f}" text-anchor="middle" font-family="Arial" font-size="10">{val:.3f}</text>')
        label = row.model.replace("_", " ")
        out.append(f'<text x="{x+bar_w/2:.2f}" y="{height-bottom+18}" text-anchor="end" transform="rotate(-45 {x+bar_w/2:.2f} {height-bottom+18})" font-family="Arial" font-size="10">{label}</text>')
    out.append(f'<text x="{left}" y="{height-20}" font-family="Arial" font-size="12">Red guide: observed morula q90 occupancy = 0.875. Orange is a morula-leaking upper bound, not a strict model.</text>')
    out.append("</svg>")
    path.write_text("\n".join(out), encoding="utf-8")


def write_doc(path: Path, metrics: pd.DataFrame, validation: pd.DataFrame, extrapolated_center: np.ndarray, observed_center: np.ndarray):
    lines = [
        "# Non-leaking distribution dynamics",
        "",
        "This experiment asks whether morula-like population occupancy can be generated without using morula centroid, morula radius, morula sample distribution, or 8-cell to morula endpoint noise during training. Morula samples are used only for final evaluation.",
        "",
        "Models tested:",
        "- pre-morula velocity ODE/SDE: drift and residual diffusion trained only on transitions ending at or before 8-cell.",
        "- affine Gaussian transition kernel: direct probabilistic kernel z_next | z, tau trained only on pre-morula transitions.",
        "- moment-extrapolated Gaussian: target morula mean/cov extrapolated from pre-morula stage moments.",
        "- validation-tuned OU: attraction strength tuned on 4-cell to 8-cell validation, then applied to an extrapolated morula center.",
        "- morula-calibrated OU upper bound: shown only as a diagnostic ceiling, not a strict model.",
        "",
        f"Pre-morula extrapolated center: {', '.join(f'{v:.4f}' for v in extrapolated_center)}",
        f"Observed morula center, evaluation only: {', '.join(f'{v:.4f}' for v in observed_center)}",
        "",
        "8-cell to morula evaluation:",
    ]
    for row in metrics.sort_values(["uses_morula_distribution_in_training", "pred_basin_occupancy_q90"]).itertuples():
        lines.append(
            f"- {row.model}: occupancy_q90={row.pred_basin_occupancy_q90:.3f}, "
            f"DMR_mean_RMSE={row.dmr_mean_rmse:.4f}, latent_MMD={row.latent_mmd_rbf:.4f}, "
            f"mean_dist={row.pred_mean_distance_to_morula_centroid:.4f}, leakage={row.uses_morula_distribution_in_training}"
        )
    lines.extend(["", "4-cell to 8-cell validation-selected OU parameters:"])
    for row in validation.itertuples():
        lines.append(
            f"- kappa={row.kappa:.4f}, diffusion_scale={row.diffusion_scale:.4f}, "
            f"latent_mean_rmse={row.validation_4cell_to_8cell_latent_mean_rmse:.4f}, cov_error={row.validation_4cell_to_8cell_cov_error:.4f}"
        )
    best_strict = metrics[metrics["uses_morula_distribution_in_training"] == "no"].sort_values("pred_basin_occupancy_q90", ascending=False).iloc[0]
    lines.extend([
        "",
        f"Current non-leaking best occupancy is {best_strict.pred_basin_occupancy_q90:.3f} from {best_strict.model}. This should be compared against observed q90 occupancy 0.875 and the morula-calibrated upper bound, not presented as complete autonomous basin generation unless it approaches the observed target.",
    ])
    path.write_text("\n".join(lines) + "\n", encoding="utf-8")


def main():
    parser = argparse.ArgumentParser()
    parser.add_argument("--q", type=int, default=3)
    parser.add_argument("--lambda", dest="lam", type=float, default=1000.0)
    parser.add_argument("--particles-per-start", type=int, default=200)
    parser.add_argument("--validation-particles", type=int, default=80)
    parser.add_argument("--n-steps", type=int, default=12)
    parser.add_argument("--seed", type=int, default=20260525)
    args = parser.parse_args()

    RESULTS.mkdir(parents=True, exist_ok=True)
    FIGURES.mkdir(parents=True, exist_ok=True)
    DOCS.mkdir(parents=True, exist_ok=True)

    matrix, ann, pairs = load_inputs()
    mu, sd, components, score_df = fit_latent_basis(matrix, args.q)
    obs_ids = stage_ids(ann, "morula")
    obs_z = score_df.loc[obs_ids].to_numpy(dtype=float)
    obs_dmr = matrix.loc[obs_ids].to_numpy(dtype=float)
    observed_basin = basin_definition(obs_z)

    pre_train_stages = set(PRE_MORULA_STAGES)
    coef, velocity_cov, train = fit_operator(score_df, pairs, train_stages=pre_train_stages, lam=args.lam)
    kernel_beta, kernel_cov, kernel_train = fit_affine_kernel(score_df, pairs, train_stages=pre_train_stages, lam=args.lam)
    extrap_center, extrap_cov, moment_df = extrapolate_stage_moments(score_df, ann, args.q, TAU["morula"])
    interp_center, interp_cov, interp_moment_df = extrapolate_stage_moments_from(
        score_df,
        ann,
        args.q,
        TAU["morula"],
        ["MII oocyte", "zygote/PN", "2-cell", "4-cell", "8-cell", "blastocyst"],
        degree=3,
    )
    _, pre_covs = stage_moments(score_df, ann, PRE_MORULA_STAGES)
    obs8_z = score_df.loc[stage_ids(ann, "8-cell")].to_numpy(dtype=float)
    validation = validate_ou_on_4cell_to_8cell(
        score_df,
        ann,
        coef,
        velocity_cov,
        obs8_z.mean(axis=0),
        obs8_z,
        args.seed,
        args.n_steps,
        args.validation_particles,
    )
    upper = calibrate_morula_upper_bound(
        score_df,
        ann,
        coef,
        velocity_cov,
        observed_basin,
        args.seed + 90000,
        args.n_steps,
        args.validation_particles,
    )

    metrics = []
    particles = []
    rng = np.random.default_rng(args.seed)

    def add_model(name, pred_z, meta, leakage, extra=None):
        pred_dmr = decode_latent(pred_z, mu, sd, components)
        row = metric_row("pre_morula_nonleaking", name, leakage, pred_z, obs_z, pred_dmr, obs_dmr, observed_basin, np.random.default_rng(args.seed + len(metrics) + 77), extra)
        metrics.append(row)
        particles.append(particle_table("pre_morula_nonleaking", name, pred_z, meta, observed_basin))

    pred, meta = simulate_from_8cell(score_df, ann, coef, rng, 1, args.n_steps)
    add_model("pre_morula_velocity_deterministic", pred, meta, "no")

    pred, meta = simulate_from_8cell(score_df, ann, coef, np.random.default_rng(args.seed + 1), args.particles_per_start, args.n_steps, cov=velocity_cov, scale=1.0)
    add_model("pre_morula_velocity_residual_sde", pred, meta, "no")

    z8 = score_df.loc[stage_ids(ann, "8-cell")].to_numpy(dtype=float)
    pred = apply_affine_kernel(z8, TAU["8-cell"], kernel_beta, np.random.default_rng(args.seed + 2), kernel_cov, args.particles_per_start)
    meta = pd.DataFrame({
        "particle_id": np.arange(len(pred), dtype=int),
        "start_sample_id": np.repeat(stage_ids(ann, "8-cell"), args.particles_per_start),
        "replicate": np.tile(np.arange(args.particles_per_start, dtype=int), len(stage_ids(ann, "8-cell"))),
    })
    add_model("pre_morula_affine_gaussian_transition_kernel", pred, meta, "no")

    pred = np.random.default_rng(args.seed + 3).multivariate_normal(extrap_center, extrap_cov, size=len(z8) * args.particles_per_start)
    meta = pd.DataFrame({
        "particle_id": np.arange(len(pred), dtype=int),
        "start_sample_id": np.repeat(stage_ids(ann, "8-cell"), args.particles_per_start),
        "replicate": np.tile(np.arange(args.particles_per_start, dtype=int), len(stage_ids(ann, "8-cell"))),
    })
    add_model("pre_morula_moment_extrapolated_gaussian", pred, meta, "no")

    pred, meta = simulate_from_8cell(
        score_df,
        ann,
        coef,
        np.random.default_rng(args.seed + 4),
        args.particles_per_start,
        args.n_steps,
        cov=velocity_cov if float(validation["diffusion_scale"]) > 0 else None,
        scale=float(validation["diffusion_scale"]),
        center=extrap_center,
        kappa=float(validation["kappa"]),
    )
    add_model("pre_morula_validation_tuned_extrapolated_ou", pred, meta, "no", {"validation_tuned_kappa": float(validation["kappa"]), "validation_tuned_diffusion_scale": float(validation["diffusion_scale"])})

    pred = np.random.default_rng(args.seed + 35).multivariate_normal(interp_center, interp_cov, size=len(z8) * args.particles_per_start)
    meta = pd.DataFrame({
        "particle_id": np.arange(len(pred), dtype=int),
        "start_sample_id": np.repeat(stage_ids(ann, "8-cell"), args.particles_per_start),
        "replicate": np.tile(np.arange(args.particles_per_start, dtype=int), len(stage_ids(ann, "8-cell"))),
    })
    add_model("leave_morula_with_blastocyst_interpolated_gaussian", pred, meta, "no_morula_but_uses_future_blastocyst")

    pred, meta = simulate_from_8cell(
        score_df,
        ann,
        coef,
        np.random.default_rng(args.seed + 36),
        args.particles_per_start,
        args.n_steps,
        cov=velocity_cov if float(validation["diffusion_scale"]) > 0 else None,
        scale=float(validation["diffusion_scale"]),
        center=interp_center,
        kappa=float(validation["kappa"]),
    )
    add_model("leave_morula_with_blastocyst_interpolated_ou", pred, meta, "no_morula_but_uses_future_blastocyst", {"validation_tuned_kappa": float(validation["kappa"]), "validation_tuned_diffusion_scale": float(validation["diffusion_scale"])})

    pred, meta = simulate_from_8cell(
        score_df,
        ann,
        coef,
        np.random.default_rng(args.seed + 5),
        args.particles_per_start,
        args.n_steps,
        cov=velocity_cov if float(upper["upper_bound_diffusion_scale"]) > 0 else None,
        scale=float(upper["upper_bound_diffusion_scale"]),
        center=observed_basin["center"],
        kappa=float(upper["upper_bound_kappa"]),
    )
    add_model("morula_calibrated_ou_upper_bound", pred, meta, "yes_upper_bound", upper)

    metric_df = pd.DataFrame(metrics)
    particle_df = pd.concat(particles, ignore_index=True)
    validation_df = pd.DataFrame([validation])
    upper_df = pd.DataFrame([upper])
    moment_df.to_csv(RESULTS / "CSB_TRO_nonleaking_pre_morula_stage_moments.tsv", sep="\t", index=False)
    interp_moment_df.to_csv(RESULTS / "CSB_TRO_nonleaking_no_morula_with_blastocyst_stage_moments.tsv", sep="\t", index=False)
    metric_df.to_csv(RESULTS / "CSB_TRO_nonleaking_distribution_metrics.tsv", sep="\t", index=False)
    particle_df.to_csv(RESULTS / "CSB_TRO_nonleaking_distribution_particles.tsv", sep="\t", index=False)
    validation_df.to_csv(RESULTS / "CSB_TRO_nonleaking_validation_tuning.tsv", sep="\t", index=False)
    upper_df.to_csv(RESULTS / "CSB_TRO_nonleaking_morula_upper_bound_tuning.tsv", sep="\t", index=False)

    pd.DataFrame(kernel_beta, index=["intercept", "tau", *[f"PC{i + 1}" for i in range(args.q)]], columns=[f"next_PC{i + 1}" for i in range(args.q)]).reset_index().rename(columns={"index": "feature"}).to_csv(
        RESULTS / "CSB_TRO_nonleaking_affine_kernel_coefficients.tsv", sep="\t", index=False
    )

    make_svg(FIGURES / "CSB_TRO_nonleaking_distribution_occupancy.svg", metric_df)
    write_doc(DOCS / "CSB_TRO_nonleaking_distribution_dynamics_interpretation.md", metric_df, validation_df, extrap_center, observed_basin["center"])

    manifest = {
        "q": args.q,
        "ridge_lambda": args.lam,
        "particles_per_start": args.particles_per_start,
        "validation_particles": args.validation_particles,
        "n_steps": args.n_steps,
        "seed": args.seed,
        "nonleaking_definition": "training excludes morula centroid, morula radius, morula sample distribution, and 8-cell to morula endpoint noise; morula is evaluation only",
        "n_velocity_training_pairs": int(len(train)),
        "n_kernel_training_pairs": int(len(kernel_train)),
        "extrapolated_morula_center": extrap_center.tolist(),
        "interpolated_morula_center_using_blastocyst_no_morula": interp_center.tolist(),
        "observed_morula_center_evaluation_only": observed_basin["center"].tolist(),
        "outputs": [
            str(RESULTS / "CSB_TRO_nonleaking_distribution_metrics.tsv"),
            str(RESULTS / "CSB_TRO_nonleaking_distribution_particles.tsv"),
            str(RESULTS / "CSB_TRO_nonleaking_validation_tuning.tsv"),
            str(RESULTS / "CSB_TRO_nonleaking_morula_upper_bound_tuning.tsv"),
            str(RESULTS / "CSB_TRO_nonleaking_affine_kernel_coefficients.tsv"),
            str(FIGURES / "CSB_TRO_nonleaking_distribution_occupancy.svg"),
            str(DOCS / "CSB_TRO_nonleaking_distribution_dynamics_interpretation.md"),
        ],
    }
    (RESULTS / "CSB_TRO_nonleaking_distribution_manifest.json").write_text(json.dumps(manifest, indent=2), encoding="utf-8")
    print(json.dumps({
        "manifest": manifest,
        "metrics": metric_df.to_dict(orient="records"),
        "validation": validation_df.to_dict(orient="records"),
        "upper_bound": upper_df.to_dict(orient="records"),
    }, indent=2))


if __name__ == "__main__":
    main()
```
