## Supplemental code for "An entropy-based diagnostic framework for characterizing methylation state dynamics during preimplantation development": run_residual_dmr_biological_annotation.py

```
from __future__ import annotations

import argparse
import json
from pathlib import Path

import numpy as np
import pandas as pd


BASE = Path(r"E:\CSB_TRO_operator_time_DMR_dynamics_2026-05-25")
RESULTS = BASE / "results"
FIGURES = BASE / "figures"
DOCS = BASE / "docs"


def read_tsv(path: Path) -> pd.DataFrame:
    return pd.read_csv(path, sep="\t") if path.exists() else pd.DataFrame()


def parse_track_arg(value: str) -> tuple[str, Path]:
    if "=" not in value:
        path = Path(value)
        return path.stem, path
    label, path = value.split("=", 1)
    return label, Path(path)


def read_bed(path: Path) -> pd.DataFrame:
    cols = ["chr", "start", "end", "name", "score", "strand"]
    df = pd.read_csv(path, sep="\t", header=None, comment="#")
    df = df.iloc[:, : min(df.shape[1], len(cols))]
    df.columns = cols[: df.shape[1]]
    for col in ["start", "end"]:
        df[col] = pd.to_numeric(df[col], errors="coerce")
    return df.dropna(subset=["chr", "start", "end"])


def overlap_regions(dmr: pd.DataFrame, regions: pd.DataFrame, label: str) -> pd.DataFrame:
    rows = []
    regions_by_chr = {c: sub for c, sub in regions.groupby("chr")}
    for row in dmr.itertuples():
        sub = regions_by_chr.get(row.chr)
        if sub is None:
            rows.append({"cluster_name": row.cluster_name, f"{label}_overlap": False, f"{label}_overlap_count": 0, f"{label}_overlap_names": ""})
            continue
        hits = sub[(sub["start"] < row.end) & (sub["end"] > row.start)]
        names = hits["name"].astype(str).tolist() if "name" in hits.columns else []
        rows.append({
            "cluster_name": row.cluster_name,
            f"{label}_overlap": bool(len(hits)),
            f"{label}_overlap_count": int(len(hits)),
            f"{label}_overlap_names": ",".join(names[:20]),
        })
    return pd.DataFrame(rows)


def nearest_gene(dmr: pd.DataFrame, gene_tss: pd.DataFrame) -> pd.DataFrame:
    required = {"chr", "tss", "gene_name"}
    if gene_tss.empty or not required.issubset(set(gene_tss.columns)):
        return pd.DataFrame({
            "cluster_name": dmr["cluster_name"],
            "nearest_gene": "NA_no_gene_tss_file",
            "distance_to_TSS": np.nan,
            "gene_type": "NA",
        })
    gene_tss = gene_tss.copy()
    gene_tss["tss"] = pd.to_numeric(gene_tss["tss"], errors="coerce")
    rows = []
    by_chr = {c: sub.dropna(subset=["tss"]) for c, sub in gene_tss.groupby("chr")}
    for row in dmr.itertuples():
        sub = by_chr.get(row.chr)
        if sub is None or sub.empty:
            rows.append({"cluster_name": row.cluster_name, "nearest_gene": "NA_no_chr_match", "distance_to_TSS": np.nan, "gene_type": "NA"})
            continue
        mid = 0.5 * (float(row.start) + float(row.end))
        dist = np.abs(sub["tss"].to_numpy(dtype=float) - mid)
        i = int(np.argmin(dist))
        hit = sub.iloc[i]
        rows.append({
            "cluster_name": row.cluster_name,
            "nearest_gene": hit["gene_name"],
            "distance_to_TSS": float(hit["tss"] - mid),
            "gene_type": hit["gene_type"] if "gene_type" in hit.index else "NA",
        })
    return pd.DataFrame(rows)


def add_internal_dynamic_annotations(annot: pd.DataFrame) -> pd.DataFrame:
    for name, path in {
        "morula_entry": RESULTS / "CSB_TRO_top100_morula_entry_DMRs.tsv",
        "dynamic_reset": RESULTS / "CSB_TRO_top100_dynamic_reset_DMRs.tsv",
        "blastocyst_exit": RESULTS / "CSB_TRO_top100_blastocyst_exit_DMRs.tsv",
        "latent_loading": RESULTS / "CSB_TRO_latent_loading_DMR_ranking.tsv",
    }.items():
        tab = read_tsv(path)
        if tab.empty or "cluster_name" not in tab.columns:
            annot[f"in_{name}_top100"] = False
            continue
        top = tab.head(100).copy()
        top[f"{name}_rank"] = np.arange(1, len(top) + 1)
        keep_cols = ["cluster_name", f"{name}_rank"]
        extra = [c for c in ["reset_dynamic_score", "morula_entry_velocity", "blastocyst_exit_velocity", "latent_loading_norm"] if c in top.columns]
        top = top[keep_cols + extra]
        annot = annot.merge(top, on="cluster_name", how="left")
        annot[f"in_{name}_top100"] = annot[f"{name}_rank"].notna()
    return annot


def classify_simple_context(row) -> str:
    if pd.notna(row.get("distance_to_TSS", np.nan)):
        d = abs(float(row["distance_to_TSS"]))
        if d <= 1000:
            return "TSS_proximal_1kb"
        if d <= 5000:
            return "TSS_proximal_5kb"
    return "unclassified_without_gene_model"


def make_track_summary(annot: pd.DataFrame, track_labels: list[str]) -> pd.DataFrame:
    rows = []
    sets = {"top25": annot.head(25), "top50": annot.head(50), "top100": annot.head(100)}
    for set_name, sub in sets.items():
        rows.append({
            "feature_set": set_name,
            "n_DMRs": len(sub),
            "mean_abs_residual_delta_beta": float(pd.to_numeric(sub["abs_latent_residual_delta_beta"], errors="coerce").mean()),
            "modules": ",".join(map(str, sorted(sub["module_id"].dropna().unique()))),
        })
        for label in track_labels:
            if f"{label}_overlap" in sub.columns:
                rows[-1][f"{label}_overlap_fraction"] = float(sub[f"{label}_overlap"].mean())
                rows[-1][f"{label}_overlap_count"] = int(sub[f"{label}_overlap"].sum())
    return pd.DataFrame(rows)


def write_doc(path: Path, annot: pd.DataFrame, track_summary: pd.DataFrame, missing: list[str]):
    top = annot.head(10)
    lines = [
        "# Residual DMR biological annotation",
        "",
        "This file converts the mathematical residual DMR ranking into a biological annotation table. With the current local files, annotation is limited to DMR coordinates, modules, CpG/width features, age weights, and overlap with internal dynamic DMR sets.",
        "",
        "External gene, motif, ATAC, histone, CpG island, and repeat annotations were not inferred unless explicit files were supplied to the script.",
        "",
        "Top residual DMRs:",
    ]
    for row in top.itertuples():
        lines.append(
            f"- {row.cluster_name} ({row.module_id}, {row.chr}:{int(row.start)}-{int(row.end)}): "
            f"residual_delta={float(row.latent_residual_delta_beta):.4f}, nearest_gene={getattr(row, 'nearest_gene', 'NA')}"
        )
    if len(track_summary):
        lines.extend(["", "Feature-set summary:"])
        for row in track_summary.itertuples():
            lines.append(f"- {row.feature_set}: n={row.n_DMRs}, modules={row.modules}")
    if missing:
        lines.extend(["", "Missing external inputs for full mechanism validation:"])
        for item in missing:
            lines.append(f"- {item}")
    lines.extend([
        "",
        "Next strict mechanism step: provide RNA/ATAC/histone/motif/gene annotation files, then fit or test whether those external features predict the residual direction without using morula methylation.",
    ])
    path.write_text("\n".join(lines) + "\n", encoding="utf-8")


def main():
    parser = argparse.ArgumentParser()
    parser.add_argument("--gene-tss", type=str, default="")
    parser.add_argument("--bed-track", action="append", default=[], help="Optional label=path BED track. Can be repeated.")
    parser.add_argument("--top-n", type=int, default=100)
    args = parser.parse_args()

    RESULTS.mkdir(parents=True, exist_ok=True)
    DOCS.mkdir(parents=True, exist_ok=True)

    dmr = read_tsv(RESULTS / "CSB_TRO_basin_residual_DMR_ranking.tsv")
    if dmr.empty:
        raise FileNotFoundError("Run run_basin_residual_control_field.py before biological annotation.")
    dmr = dmr.copy()
    for col in ["start", "end", "width", "n_cpg_target", "age_weight_5yr", "abs_latent_residual_delta_beta"]:
        if col in dmr.columns:
            dmr[col] = pd.to_numeric(dmr[col], errors="coerce")
    dmr["cpg_per_100bp"] = dmr["n_cpg_target"] / (dmr["width"] / 100.0)
    dmr["residual_direction_label"] = np.where(dmr["latent_residual_delta_beta"] >= 0, "positive_delta_beta", "negative_delta_beta")

    gene_tss = read_tsv(Path(args.gene_tss)) if args.gene_tss else pd.DataFrame()
    annot = dmr.merge(nearest_gene(dmr, gene_tss), on="cluster_name", how="left")
    annot["simple_genomic_context"] = annot.apply(classify_simple_context, axis=1)
    annot = add_internal_dynamic_annotations(annot)

    track_labels = []
    for spec in args.bed_track:
        label, path = parse_track_arg(spec)
        regions = read_bed(path)
        ov = overlap_regions(annot, regions, label)
        annot = annot.merge(ov, on="cluster_name", how="left")
        track_labels.append(label)

    top_annot = annot.head(args.top_n).copy()
    track_summary = make_track_summary(annot, track_labels)
    top_annot.to_csv(RESULTS / "CSB_TRO_top_residual_DMR_annotation.tsv", sep="\t", index=False)
    annot[["cluster_name", "chr", "start", "end", "nearest_gene", "distance_to_TSS", "gene_type", "simple_genomic_context"]].to_csv(
        RESULTS / "CSB_TRO_residual_DMR_nearest_genes.tsv", sep="\t", index=False
    )
    track_summary.to_csv(RESULTS / "CSB_TRO_residual_DMR_external_track_overlap_summary.tsv", sep="\t", index=False)

    # Placeholder files make the missing mechanism-validation layer explicit and machine-readable.
    placeholder_cols = ["analysis_status", "reason", "required_input"]
    placeholders = {
        "CSB_TRO_residual_DMR_GO_KEGG.tsv": "nearest gene or gene set annotation",
        "CSB_TRO_residual_DMR_motif_enrichment.tsv": "genome FASTA plus motif database or motif-scan output",
        "CSB_TRO_residual_DMR_ATAC_overlap.tsv": "ATAC peak/signal BED files",
        "CSB_TRO_residual_DMR_histone_overlap.tsv": "histone mark peak/signal BED files",
        "CSB_TRO_residual_DMR_RNA_validation.tsv": "stage-resolved RNA expression matrix",
    }
    for filename, req in placeholders.items():
        pd.DataFrame([{"analysis_status": "not_run_missing_external_input", "reason": "No compatible local input file was provided.", "required_input": req}])[placeholder_cols].to_csv(
            RESULTS / filename, sep="\t", index=False
        )

    missing = list(placeholders.values())
    if args.gene_tss:
        missing = [m for m in missing if not m.startswith("nearest gene")]
    if track_labels:
        missing = [m for m in missing if "ATAC" not in m and "histone" not in m]
    write_doc(DOCS / "CSB_TRO_residual_biology_interpretation.md", top_annot, track_summary, missing)
    manifest = {
        "gene_tss_input": args.gene_tss or None,
        "bed_tracks": args.bed_track,
        "top_n": args.top_n,
        "outputs": [
            str(RESULTS / "CSB_TRO_top_residual_DMR_annotation.tsv"),
            str(RESULTS / "CSB_TRO_residual_DMR_nearest_genes.tsv"),
            str(RESULTS / "CSB_TRO_residual_DMR_external_track_overlap_summary.tsv"),
            str(RESULTS / "CSB_TRO_residual_DMR_GO_KEGG.tsv"),
            str(RESULTS / "CSB_TRO_residual_DMR_motif_enrichment.tsv"),
            str(RESULTS / "CSB_TRO_residual_DMR_ATAC_overlap.tsv"),
            str(RESULTS / "CSB_TRO_residual_DMR_histone_overlap.tsv"),
            str(RESULTS / "CSB_TRO_residual_DMR_RNA_validation.tsv"),
            str(DOCS / "CSB_TRO_residual_biology_interpretation.md"),
        ],
    }
    (RESULTS / "CSB_TRO_residual_biology_manifest.json").write_text(json.dumps(manifest, indent=2), encoding="utf-8")
    print(json.dumps({
        "n_annotated": int(len(top_annot)),
        "top_modules": sorted(top_annot["module_id"].dropna().unique().tolist()),
        "external_tracks": track_labels,
        "missing_external_inputs": missing,
    }, indent=2))


if __name__ == "__main__":
    main()
```
