## Supplemental code for "An entropy-based diagnostic framework for characterizing methylation state dynamics during preimplantation development": run_residual_dmr_robustness.py

```
from __future__ import annotations

import argparse
import json
import sys
from pathlib import Path

import numpy as np
import pandas as pd


BASE = Path(r"E:\CSB_TRO_operator_time_DMR_dynamics_2026-05-25")
CODE = BASE / "code"
RESULTS = BASE / "results"
FIGURES = BASE / "figures"
DOCS = BASE / "docs"
sys.path.insert(0, str(CODE))

from run_basin_residual_control_field import (  # noqa: E402
    PRE_MORULA_STAGES,
    build_dmr_residual_table,
    cosine,
    decode_delta_to_dmr,
    encode_delta_to_latent,
    module_summary,
    simulate_strict_pre_morula,
)
from run_morula_basin_sde import (  # noqa: E402
    basin_definition,
    decode_latent,
    distribution_metrics,
    fit_latent_basis,
    fit_operator,
    load_inputs,
    stage_ids,
)


TOPK = [1, 3, 5, 10, 15, 25, 50, 100, 156]


def occupancy(pred_z: np.ndarray, center: np.ndarray, radius: float) -> float:
    return float(np.mean(np.linalg.norm(pred_z - center[None, :], axis=1) <= radius))


def control_from_dmr_set(matrix, sd, components, dmr_table, selected, residual_col="latent_residual_delta_beta"):
    delta = np.zeros(matrix.shape[1], dtype=float)
    by_id = dmr_table.set_index("cluster_name")
    idx = [list(matrix.columns).index(c) for c in selected]
    delta[idx] = by_id.loc[selected, residual_col].to_numpy(dtype=float)
    return encode_delta_to_latent(delta, sd, components)


def evaluate_control(name, pred_z, obs_z, obs_dmr, matrix, mu, sd, components, basin, extra, rng):
    pred_dmr = decode_latent(pred_z, mu, sd, components)
    out = {
        "control_name": name,
        **extra,
        **distribution_metrics(pred_z, obs_z, pred_dmr, obs_dmr, basin, rng),
    }
    return out


def topk_curve(matrix, mu, sd, components, strict_pred_z, obs_z, obs_dmr, basin, dmr_table, rng):
    rows = []
    all_ids = dmr_table["cluster_name"].tolist()
    full_z = control_from_dmr_set(matrix, sd, components, dmr_table, all_ids)
    for k in TOPK:
        selected = dmr_table.head(k)["cluster_name"].tolist()
        z_part = control_from_dmr_set(matrix, sd, components, dmr_table, selected)
        rows.append(evaluate_control(
            f"only_top{k}",
            strict_pred_z + z_part[None, :],
            obs_z,
            obs_dmr,
            matrix,
            mu,
            sd,
            components,
            basin,
            {
                "K": k,
                "control_type": "only_topK",
                "sign": "forward",
                "latent_control_norm": float(np.linalg.norm(z_part)),
                "latent_control_cosine_to_full_residual": cosine(z_part, full_z),
            },
            rng,
        ))
        rows.append(evaluate_control(
            f"sign_flip_top{k}",
            strict_pred_z - z_part[None, :],
            obs_z,
            obs_dmr,
            matrix,
            mu,
            sd,
            components,
            basin,
            {
                "K": k,
                "control_type": "sign_flip_topK",
                "sign": "reverse",
                "latent_control_norm": float(np.linalg.norm(z_part)),
                "latent_control_cosine_to_full_residual": cosine(-z_part, full_z),
            },
            rng,
        ))
        remainder = [c for c in all_ids if c not in set(selected)]
        z_remain = control_from_dmr_set(matrix, sd, components, dmr_table, remainder)
        rows.append(evaluate_control(
            f"remove_top{k}_keep_remainder",
            strict_pred_z + z_remain[None, :],
            obs_z,
            obs_dmr,
            matrix,
            mu,
            sd,
            components,
            basin,
            {
                "K": k,
                "control_type": "remove_topK_keep_remainder",
                "sign": "forward_remainder",
                "latent_control_norm": float(np.linalg.norm(z_remain)),
                "latent_control_cosine_to_full_residual": cosine(z_remain, full_z),
            },
            rng,
        ))
    return pd.DataFrame(rows)


def bootstrap_stability(matrix, ann, score_df, strict_pred_z, components, sd, metadata, modules, n_boot, seed):
    rng = np.random.default_rng(seed)
    morula_ids = stage_ids(ann, "morula")
    selection = {}
    rank_sum = {}
    rank_sq = {}
    contrib = {}
    centers = []
    for b in range(n_boot):
        ids = rng.choice(morula_ids, size=len(morula_ids), replace=True)
        obs_center = score_df.loc[ids].to_numpy(dtype=float).mean(axis=0)
        residual_z = obs_center - strict_pred_z.mean(axis=0)
        obs_dmr = matrix.loc[ids].to_numpy(dtype=float)
        obs_delta = obs_dmr.mean(axis=0) - decode_latent(strict_pred_z, np.zeros(matrix.shape[1]), np.ones(matrix.shape[1]), components).mean(axis=0)
        model_delta = np.zeros(matrix.shape[1])
        tab = build_dmr_residual_table(matrix, metadata, modules, components, sd, residual_z, obs_delta, model_delta)
        centers.append({"bootstrap": b, **{f"center_PC{i + 1}": obs_center[i] for i in range(len(obs_center))}})
        for row in tab.itertuples():
            cid = row.cluster_name
            r = int(row.basin_residual_rank)
            c = float(row.latent_residual_delta_beta)
            selection.setdefault(cid, {"top10": 0, "top25": 0, "top50": 0, "top100": 0})
            if r <= 10:
                selection[cid]["top10"] += 1
            if r <= 25:
                selection[cid]["top25"] += 1
            if r <= 50:
                selection[cid]["top50"] += 1
            if r <= 100:
                selection[cid]["top100"] += 1
            rank_sum[cid] = rank_sum.get(cid, 0.0) + r
            rank_sq[cid] = rank_sq.get(cid, 0.0) + r * r
            contrib.setdefault(cid, []).append(c)
    rows = []
    for cid in matrix.columns:
        vals = np.asarray(contrib.get(cid, [0.0]), dtype=float)
        mean_rank = rank_sum.get(cid, 0.0) / n_boot
        mean_rank_sq = rank_sq.get(cid, 0.0) / n_boot
        rows.append({
            "cluster_name": cid,
            "selection_frequency_top10": selection.get(cid, {}).get("top10", 0) / n_boot,
            "selection_frequency_top25": selection.get(cid, {}).get("top25", 0) / n_boot,
            "selection_frequency_top50": selection.get(cid, {}).get("top50", 0) / n_boot,
            "selection_frequency_top100": selection.get(cid, {}).get("top100", 0) / n_boot,
            "mean_rank": mean_rank,
            "rank_std": float(max(mean_rank_sq - mean_rank * mean_rank, 0.0) ** 0.5),
            "residual_contribution_mean": float(vals.mean()),
            "residual_contribution_q025": float(np.quantile(vals, 0.025)),
            "residual_contribution_q975": float(np.quantile(vals, 0.975)),
        })
    return pd.DataFrame(rows), pd.DataFrame(centers)


def matched_random_controls(matrix, ann, score_df, dmr_table, mu, sd, components, strict_pred_z, obs_z, obs_dmr, basin, n_random, seed):
    rng = np.random.default_rng(seed)
    beta8 = matrix.loc[stage_ids(ann, "8-cell")].mean(axis=0)
    stage_var = []
    for c in matrix.columns:
        means = []
        for stage in ["MII oocyte", "zygote/PN", "2-cell", "4-cell", "8-cell"]:
            means.append(matrix.loc[stage_ids(ann, stage), c].mean())
        stage_var.append(np.var(means))
    covar = dmr_table[["cluster_name", "module_id", "width", "n_cpg_target", "PC_loading_norm"]].copy()
    covar["beta8"] = beta8.loc[covar["cluster_name"]].to_numpy(dtype=float)
    covar["stage_var_pre_morula"] = stage_var
    covar["width"] = pd.to_numeric(covar["width"], errors="coerce").fillna(covar["width"].median())
    covar["n_cpg_target"] = pd.to_numeric(covar["n_cpg_target"], errors="coerce").fillna(covar["n_cpg_target"].median())
    by_id = covar.set_index("cluster_name")
    all_ids = covar["cluster_name"].tolist()
    rows = []
    for k in [10, 25, 50]:
        target = dmr_table.head(k)["cluster_name"].tolist()
        target_rows = by_id.loc[target]
        target_control = control_from_dmr_set(matrix, sd, components, dmr_table, target)
        rows.append(evaluate_control(
            f"observed_top{k}",
            strict_pred_z + target_control[None, :],
            obs_z,
            obs_dmr,
            matrix,
            mu,
            sd,
            components,
            basin,
            {"K": k, "control_type": "observed_topK", "random_iter": -1, "matched": "no"},
            rng,
        ))
        for it in range(n_random):
            chosen = []
            available = set(all_ids) - set(target)
            for target_id, t in target_rows.iterrows():
                candidates = covar[covar["cluster_name"].isin(available)].copy()
                same_module = candidates[candidates["module_id"] == t["module_id"]]
                if len(same_module) >= 3:
                    candidates = same_module
                for col in ["width", "n_cpg_target", "PC_loading_norm", "beta8", "stage_var_pre_morula"]:
                    scale = float(covar[col].std()) + 1e-9
                    candidates[f"d_{col}"] = ((candidates[col].to_numpy(dtype=float) - float(t[col])) / scale) ** 2
                candidates["dist"] = candidates[[f"d_{c}" for c in ["width", "n_cpg_target", "PC_loading_norm", "beta8", "stage_var_pre_morula"]]].sum(axis=1)
                pool = candidates.sort_values("dist").head(min(10, len(candidates)))
                pick = str(pool.sample(1, random_state=int(rng.integers(0, 2**31 - 1))).iloc[0]["cluster_name"])
                chosen.append(pick)
                available.discard(pick)
            z_part = control_from_dmr_set(matrix, sd, components, dmr_table, chosen)
            rows.append(evaluate_control(
                f"matched_random_top{k}",
                strict_pred_z + z_part[None, :],
                obs_z,
                obs_dmr,
                matrix,
                mu,
                sd,
                components,
                basin,
                {
                    "K": k,
                    "control_type": "matched_random",
                    "random_iter": it,
                    "matched": "module,width,cpg,loading,beta8,pre_morula_stage_var",
                    "latent_control_cosine_to_observed_topK": cosine(z_part, target_control),
                },
                rng,
            ))
    return pd.DataFrame(rows)


def module_add_remove(matrix, mu, sd, components, strict_pred_z, obs_z, obs_dmr, basin, dmr_table, residual_z, rng):
    rows = []
    module_df = module_summary(dmr_table, residual_z, sd, components)
    all_ids = dmr_table["cluster_name"].tolist()
    full_z = control_from_dmr_set(matrix, sd, components, dmr_table, all_ids)
    for module_id in module_df["module_id"].tolist():
        ids = dmr_table[dmr_table["module_id"] == module_id]["cluster_name"].tolist()
        z_mod = control_from_dmr_set(matrix, sd, components, dmr_table, ids)
        rows.append(evaluate_control(
            f"only_module_{module_id}",
            strict_pred_z + z_mod[None, :],
            obs_z,
            obs_dmr,
            matrix,
            mu,
            sd,
            components,
            basin,
            {"module_id": module_id, "control_type": "only_module", "n_DMRs": len(ids), "cosine_to_full": cosine(z_mod, full_z)},
            rng,
        ))
        remain = [c for c in all_ids if c not in set(ids)]
        z_remain = control_from_dmr_set(matrix, sd, components, dmr_table, remain)
        rows.append(evaluate_control(
            f"remove_module_{module_id}",
            strict_pred_z + z_remain[None, :],
            obs_z,
            obs_dmr,
            matrix,
            mu,
            sd,
            components,
            basin,
            {"module_id": module_id, "control_type": "remove_module_keep_remainder", "n_DMRs": len(remain), "cosine_to_full": cosine(z_remain, full_z)},
            rng,
        ))
    return pd.DataFrame(rows)


def make_svg(path: Path, topk: pd.DataFrame, random_df: pd.DataFrame):
    rows = topk[topk["control_type"].isin(["only_topK", "sign_flip_topK", "remove_topK_keep_remainder"])].copy()
    width, height = 900, 430
    left, right, top, bottom = 70, 25, 40, 70
    plot_w = width - left - right
    plot_h = height - top - bottom
    ks = sorted(rows["K"].unique())
    colors = {"only_topK": "#2f6f8f", "sign_flip_topK": "#b56b2a", "remove_topK_keep_remainder": "#666666"}
    out = [
        f'<svg xmlns="http://www.w3.org/2000/svg" width="{width}" height="{height}" viewBox="0 0 {width} {height}">',
        '<rect width="100%" height="100%" fill="white"/>',
        f'<text x="{left}" y="26" font-family="Arial" font-size="18" font-weight="700">Residual DMR control robustness: TopK, sign-flip, and remove-TopK</text>',
        f'<line x1="{left}" y1="{height-bottom}" x2="{width-right}" y2="{height-bottom}" stroke="#222"/>',
        f'<line x1="{left}" y1="{top}" x2="{left}" y2="{height-bottom}" stroke="#222"/>',
    ]
    for tick in [0, 0.25, 0.5, 0.75, 0.875, 1.0]:
        y = height - bottom - tick * plot_h
        color = "#b33" if abs(tick - 0.875) < 1e-6 else "#ddd"
        out.append(f'<line x1="{left}" y1="{y:.2f}" x2="{width-right}" y2="{y:.2f}" stroke="{color}" stroke-dasharray="4 4"/>')
        out.append(f'<text x="{left-8}" y="{y+4:.2f}" text-anchor="end" font-family="Arial" font-size="11">{tick:.3g}</text>')
    def xy(k, val):
        x = left + (np.log2(k) / np.log2(max(ks))) * plot_w if k > 0 else left
        y = height - bottom - val * plot_h
        return x, y
    for ctype, sub in rows.groupby("control_type"):
        sub = sub.sort_values("K")
        pts = [xy(float(r.K), float(r.pred_basin_occupancy_q90)) for r in sub.itertuples()]
        d = " ".join([f"{x:.2f},{y:.2f}" for x, y in pts])
        out.append(f'<polyline points="{d}" fill="none" stroke="{colors[ctype]}" stroke-width="2.5"/>')
        for x, y in pts:
            out.append(f'<circle cx="{x:.2f}" cy="{y:.2f}" r="3.5" fill="{colors[ctype]}"/>')
    for k in ks:
        x, _ = xy(float(k), 0)
        out.append(f'<text x="{x:.2f}" y="{height-bottom+18}" text-anchor="middle" font-family="Arial" font-size="10">{int(k)}</text>')
    out.append(f'<text x="{left}" y="{height-20}" font-family="Arial" font-size="12">Blue: only TopK residual; orange: sign-flipped TopK; gray: residual after removing TopK. Red guide: observed occupancy 0.875.</text>')
    out.append("</svg>")
    path.write_text("\n".join(out), encoding="utf-8")


def write_doc(path: Path, summary: dict, topk: pd.DataFrame, boot: pd.DataFrame, random_df: pd.DataFrame, module_df: pd.DataFrame):
    best25 = topk[(topk["control_type"] == "only_topK") & (topk["K"] == 25)].iloc[0]
    flip25 = topk[(topk["control_type"] == "sign_flip_topK") & (topk["K"] == 25)].iloc[0]
    remove25 = topk[(topk["control_type"] == "remove_topK_keep_remainder") & (topk["K"] == 25)].iloc[0]
    rand25 = random_df[(random_df["control_type"] == "matched_random") & (random_df["K"] == 25)]
    stable = boot.sort_values("selection_frequency_top25", ascending=False).head(10)
    lines = [
        "# Residual DMR robustness",
        "",
        "This package tests whether the residual DMR control set is stable, directional, and stronger than matched DMR controls. The residual itself remains diagnostic because it uses observed morula methylation for definition.",
        "",
        f"Observed morula q90 occupancy target: {summary['observed_basin_occupancy_q90']:.3f}",
        f"Strict pre-morula baseline occupancy: {summary['strict_predicted_basin_occupancy_q90']:.3f}",
        "",
        f"Top25 forward residual occupancy: {best25.pred_basin_occupancy_q90:.3f}",
        f"Top25 sign-flip occupancy: {flip25.pred_basin_occupancy_q90:.3f}",
        f"All residual except Top25 occupancy: {remove25.pred_basin_occupancy_q90:.3f}",
        f"Matched random Top25 occupancy mean: {pd.to_numeric(rand25['pred_basin_occupancy_q90']).mean():.3f}",
        f"Matched random Top25 occupancy q95: {pd.to_numeric(rand25['pred_basin_occupancy_q90']).quantile(0.95):.3f}",
        "",
        "Most stable Top25 DMRs by bootstrap selection frequency:",
    ]
    for row in stable.itertuples():
        lines.append(f"- {row.cluster_name}: freq_top25={row.selection_frequency_top25:.3f}, mean_rank={row.mean_rank:.1f}")
    lines.extend(["", "Top module add/remove effects:"])
    for row in module_df.sort_values("pred_basin_occupancy_q90", ascending=False).head(8).itertuples():
        lines.append(f"- {row.control_name}: occupancy_q90={row.pred_basin_occupancy_q90:.3f}, n_DMRs={row.n_DMRs}")
    lines.extend([
        "",
        "Interpretation boundary: strong forward/sign-flip/matched-random separation supports a compact, directional residual DMR control component. It does not by itself identify the external regulatory/chromatin variable that predicts this component without morula methylation.",
    ])
    path.write_text("\n".join(lines) + "\n", encoding="utf-8")


def main():
    parser = argparse.ArgumentParser()
    parser.add_argument("--q", type=int, default=3)
    parser.add_argument("--lambda", dest="lam", type=float, default=1000.0)
    parser.add_argument("--n-steps", type=int, default=12)
    parser.add_argument("--n-bootstrap", type=int, default=500)
    parser.add_argument("--n-random", type=int, default=500)
    parser.add_argument("--seed", type=int, default=20260525)
    args = parser.parse_args()

    RESULTS.mkdir(parents=True, exist_ok=True)
    FIGURES.mkdir(parents=True, exist_ok=True)
    DOCS.mkdir(parents=True, exist_ok=True)

    matrix, ann, pairs = load_inputs()
    mu, sd, components, score_df = fit_latent_basis(matrix, args.q)
    coef, cov, train = fit_operator(score_df, pairs, train_stages=set(PRE_MORULA_STAGES), lam=args.lam)
    strict_pred_z = simulate_strict_pre_morula(score_df, ann, coef, args.n_steps)
    obs_z = score_df.loc[stage_ids(ann, "morula")].to_numpy(dtype=float)
    obs_dmr = matrix.loc[stage_ids(ann, "morula")].to_numpy(dtype=float)
    basin = basin_definition(obs_z)
    residual_z = obs_z.mean(axis=0) - strict_pred_z.mean(axis=0)
    strict_pred_dmr = decode_latent(strict_pred_z, mu, sd, components)
    obs_delta_dmr = obs_dmr.mean(axis=0) - strict_pred_dmr.mean(axis=0)
    dmr8 = matrix.loc[stage_ids(ann, "8-cell")].mean(axis=0).to_numpy(dtype=float)
    model_delta = strict_pred_dmr.mean(axis=0) - dmr8
    metadata = pd.read_csv(RESULTS / "CSB_TRO_DMR_metadata.tsv", sep="\t")
    modules = pd.read_csv(RESULTS / "CSB_TRO_DMR_module_assignments.tsv", sep="\t")
    dmr_table = build_dmr_residual_table(matrix, metadata, modules, components, sd, residual_z, obs_delta_dmr, model_delta)

    rng = np.random.default_rng(args.seed)
    topk = topk_curve(matrix, mu, sd, components, strict_pred_z, obs_z, obs_dmr, basin, dmr_table, rng)
    boot, boot_centers = bootstrap_stability(matrix, ann, score_df, strict_pred_z, components, sd, metadata, modules, args.n_bootstrap, args.seed + 100)
    boot = boot.merge(dmr_table[["cluster_name", "module_id", "basin_residual_rank", "abs_latent_residual_delta_beta"]], on="cluster_name", how="left")
    random_df = matched_random_controls(matrix, ann, score_df, dmr_table, mu, sd, components, strict_pred_z, obs_z, obs_dmr, basin, args.n_random, args.seed + 200)
    mod = module_add_remove(matrix, mu, sd, components, strict_pred_z, obs_z, obs_dmr, basin, dmr_table, residual_z, rng)
    mod = mod.rename(columns={"control_name": "control_name"})

    summary = {
        "q": args.q,
        "ridge_lambda": args.lam,
        "n_bootstrap": args.n_bootstrap,
        "n_random": args.n_random,
        "seed": args.seed,
        "observed_basin_occupancy_q90": float(basin["observed_occupancy_q90"]),
        "strict_predicted_basin_occupancy_q90": occupancy(strict_pred_z, obs_z.mean(axis=0), basin["radius_q90"]),
        "missing_basin_residual_norm": float(np.linalg.norm(residual_z)),
        "n_pre_morula_training_pairs": int(len(train)),
    }

    topk.to_csv(RESULTS / "CSB_TRO_residual_DMR_topK_occupancy_curve.tsv", sep="\t", index=False)
    boot.to_csv(RESULTS / "CSB_TRO_residual_DMR_bootstrap_stability.tsv", sep="\t", index=False)
    boot_centers.to_csv(RESULTS / "CSB_TRO_residual_DMR_bootstrap_centers.tsv", sep="\t", index=False)
    random_df.to_csv(RESULTS / "CSB_TRO_residual_DMR_matched_random_control.tsv", sep="\t", index=False)
    topk[topk["control_type"] == "sign_flip_topK"].to_csv(RESULTS / "CSB_TRO_residual_DMR_sign_flip_control.tsv", sep="\t", index=False)
    topk[topk["control_type"] == "remove_topK_keep_remainder"].to_csv(RESULTS / "CSB_TRO_residual_DMR_remove_topK_necessity.tsv", sep="\t", index=False)
    mod.to_csv(RESULTS / "CSB_TRO_residual_module_add_remove.tsv", sep="\t", index=False)
    pd.DataFrame([summary]).to_csv(RESULTS / "CSB_TRO_residual_DMR_robustness_summary.tsv", sep="\t", index=False)

    make_svg(FIGURES / "CSB_TRO_residual_DMR_topK_occupancy.svg", topk, random_df)
    write_doc(DOCS / "CSB_TRO_residual_DMR_robustness_summary.md", summary, topk, boot, random_df, mod)
    manifest = {
        **summary,
        "outputs": [
            str(RESULTS / "CSB_TRO_residual_DMR_topK_occupancy_curve.tsv"),
            str(RESULTS / "CSB_TRO_residual_DMR_bootstrap_stability.tsv"),
            str(RESULTS / "CSB_TRO_residual_DMR_matched_random_control.tsv"),
            str(RESULTS / "CSB_TRO_residual_DMR_sign_flip_control.tsv"),
            str(RESULTS / "CSB_TRO_residual_DMR_remove_topK_necessity.tsv"),
            str(RESULTS / "CSB_TRO_residual_module_add_remove.tsv"),
            str(FIGURES / "CSB_TRO_residual_DMR_topK_occupancy.svg"),
            str(DOCS / "CSB_TRO_residual_DMR_robustness_summary.md"),
        ],
    }
    (RESULTS / "CSB_TRO_residual_DMR_robustness_manifest.json").write_text(json.dumps(manifest, indent=2), encoding="utf-8")
    print(json.dumps({
        "summary": summary,
        "topK": topk[topk["control_type"] == "only_topK"].to_dict(orient="records"),
        "top_stable": boot.sort_values("selection_frequency_top25", ascending=False).head(10).to_dict(orient="records"),
        "matched_random_top25": random_df[random_df["K"] == 25]["pred_basin_occupancy_q90"].astype(float).describe().to_dict(),
    }, indent=2))


if __name__ == "__main__":
    main()
```
