## Supplemental code for "An entropy-based diagnostic framework for characterizing methylation state dynamics during preimplantation development": run_residual_module_chromatin_gate_control.py

```
from __future__ import annotations

import gzip
import json
import math
import sys
from pathlib import Path

import numpy as np
import pandas as pd


BASE = Path(r"E:\CSB_TRO_operator_time_DMR_dynamics_2026-05-25")
MAIN = Path(r"C:\Users\18068\Desktop\CSB_TRO_Project_2026-05-24")
CODE = BASE / "code"
RESULTS = BASE / "results"
FIGURES = BASE / "figures"
DOCS = BASE / "docs"
EXTERNAL = BASE / "external"
sys.path.insert(0, str(CODE))

from run_basin_residual_control_field import PRE_MORULA_STAGES, cosine, simulate_strict_pre_morula  # noqa: E402
from run_morula_basin_sde import (  # noqa: E402
    basin_definition,
    decode_latent,
    distribution_metrics,
    fit_latent_basis,
    fit_operator,
    load_inputs,
    stage_ids,
)


PRIORITY_MODULES = ["M05", "M01", "M12", "M02", "M10"]

REGION_ANNOT = RESULTS / "CSB_TRO_residual_module_region_annotation.tsv"
MODULE_BASIS = RESULTS / "CSB_TRO_missing_control_term_module_basis.tsv"
MOTIF_FEATURES = RESULTS / "CSB_TRO_module_TF_activity_control_panel_features.tsv"
MATCHED = MAIN / "input_tables" / "GSE81233_matched_non_age_window_regions_n100.tsv"
HISTONE_MANIFEST = EXTERNAL / "histone" / "histone_peak_manifest.tsv"

ATAC_TRACKS = [
    ("ATAC_8cell_2pn", "ATAC", "8-cell", EXTERNAL / "atac" / "GSE101571_8cell_2pn_peaks.bed.gz"),
    ("ATAC_8cell_3pn", "ATAC", "8-cell", EXTERNAL / "atac" / "GSE101571_8cell_3pn_peaks.bed.gz"),
    ("ATAC_ICM_2pn", "ATAC", "ICM", EXTERNAL / "atac" / "GSE101571_icm_2pn_peaks.bed.gz"),
    ("ATAC_ICM_3pn", "ATAC", "ICM", EXTERNAL / "atac" / "GSE101571_icm_3pn_peaks.bed.gz"),
]

OUT_REGION = RESULTS / "CSB_TRO_residual_module_region_composition.tsv"
OUT_OVERLAP = RESULTS / "CSB_TRO_module_chromatin_overlap.tsv"
OUT_STAGE = RESULTS / "CSB_TRO_module_chromatin_stage_delta.tsv"
OUT_FEATURES = RESULTS / "CSB_TRO_chromatin_gated_TF_activity.tsv"
OUT_METRICS = RESULTS / "CSB_TRO_chromatin_gated_control_metrics.tsv"
OUT_ABLATION = RESULTS / "CSB_TRO_chromatin_gated_control_ablation.tsv"
OUT_MANIFEST = RESULTS / "CSB_TRO_chromatin_gated_control_manifest.json"
OUT_SVG = FIGURES / "CSB_TRO_chromatin_gated_control.svg"
OUT_DOC = DOCS / "CSB_TRO_chromatin_gated_control_summary.md"


def read_bed(path: Path) -> pd.DataFrame:
    opener = gzip.open if path.suffix == ".gz" else open
    rows = []
    with opener(path, "rt", encoding="utf-8", errors="replace") as handle:
        for line in handle:
            if not line.strip() or line.startswith(("#", "track", "browser")):
                continue
            parts = line.rstrip("\n").split("\t")
            if len(parts) < 3:
                continue
            try:
                start = int(float(parts[1]))
                end = int(float(parts[2]))
            except ValueError:
                continue
            rows.append({"chr": parts[0], "start": start, "end": end, "name": parts[3] if len(parts) > 3 else f"{parts[0]}:{start}-{end}"})
    return pd.DataFrame(rows)


def overlap_flags(regions: pd.DataFrame, peaks: pd.DataFrame) -> np.ndarray:
    flags = np.zeros(len(regions), dtype=bool)
    if peaks.empty or regions.empty:
        return flags
    peak_by_chr = {c: sub[["start", "end"]].to_numpy(dtype=int) for c, sub in peaks.groupby("chr")}
    for i, row in enumerate(regions.itertuples()):
        arr = peak_by_chr.get(str(row.chr))
        if arr is None:
            continue
        flags[i] = bool(np.any((arr[:, 0] < int(row.end)) & (arr[:, 1] > int(row.start))))
    return flags


def log_comb(n: int, k: int) -> float:
    if k < 0 or k > n:
        return float("-inf")
    return math.lgamma(n + 1) - math.lgamma(k + 1) - math.lgamma(n - k + 1)


def fisher_greater(a: int, b: int, c: int, d: int) -> float:
    # One-sided enrichment p-value for table [[a,b],[c,d]].
    n = a + b + c + d
    row1 = a + b
    col1 = a + c
    lo = max(0, row1 - (n - col1))
    hi = min(row1, col1)
    denom = log_comb(n, row1)
    vals = [log_comb(col1, x) + log_comb(n - col1, row1 - x) - denom for x in range(max(a, lo), hi + 1)]
    if not vals:
        return 1.0
    m = max(vals)
    return float(min(1.0, math.exp(m) * sum(math.exp(v - m) for v in vals)))


def fdr_bh(p: pd.Series) -> pd.Series:
    x = pd.to_numeric(p, errors="coerce").fillna(1.0).to_numpy(dtype=float)
    order = np.argsort(x)
    q = np.empty_like(x)
    prev = 1.0
    n = len(x)
    for rank, idx in enumerate(order[::-1], start=1):
        true_rank = n - rank + 1
        val = x[idx] * n / true_rank
        prev = min(prev, val)
        q[idx] = prev
    return pd.Series(np.minimum(q, 1.0), index=p.index)


def odds_ratio(a: int, b: int, c: int, d: int) -> float:
    return float(((a + 0.5) * (d + 0.5)) / ((b + 0.5) * (c + 0.5)))


def region_composition() -> pd.DataFrame:
    annot = pd.read_csv(REGION_ANNOT, sep="\t")
    annot = annot[annot["module_id"].isin(PRIORITY_MODULES)].copy()
    rows = []
    for module_id, sub in annot.groupby("module_id"):
        rows.append(
            {
                "module_id": module_id,
                "n_DMR": int(len(sub)),
                "promoter_2kb_fraction": float(pd.to_numeric(sub["promoter_2kb"], errors="coerce").fillna(False).astype(bool).mean()),
                "promoter_5kb_fraction": float(pd.to_numeric(sub["promoter_5kb"], errors="coerce").fillna(False).astype(bool).mean()),
                "gene_body_fraction": float(pd.to_numeric(sub["gene_body"], errors="coerce").fillna(False).astype(bool).mean()),
                "intergenic_fraction": float(pd.to_numeric(sub["intergenic"], errors="coerce").fillna(False).astype(bool).mean()),
                "CpG_island_fraction": float(pd.to_numeric(sub["CpG_island"], errors="coerce").fillna(False).astype(bool).mean()) if "CpG_island" in sub else 0.0,
                "CpG_shore_fraction": float(pd.to_numeric(sub["CpG_shore"], errors="coerce").fillna(False).astype(bool).mean()) if "CpG_shore" in sub else 0.0,
                "CpG_shelf_fraction": float(pd.to_numeric(sub["CpG_shelf"], errors="coerce").fillna(False).astype(bool).mean()) if "CpG_shelf" in sub else 0.0,
                "repeat_annotated_fraction": float(sub["repeat_class"].fillna("").astype(str).ne("").mean()) if "repeat_class" in sub else 0.0,
                "mean_abs_residual_contribution": float(pd.to_numeric(sub["residual_contribution"], errors="coerce").abs().mean()),
                "sum_abs_residual_contribution": float(pd.to_numeric(sub["residual_contribution"], errors="coerce").abs().sum()),
                "module_basis_weight": float(pd.to_numeric(sub["module_basis_weight"], errors="coerce").dropna().iloc[0]) if sub["module_basis_weight"].notna().any() else np.nan,
            }
        )
    out = pd.DataFrame(rows).sort_values("sum_abs_residual_contribution", ascending=False)
    out.to_csv(OUT_REGION, sep="\t", index=False)
    return out


def target_regions() -> pd.DataFrame:
    annot = pd.read_csv(REGION_ANNOT, sep="\t")
    return annot[annot["module_id"].isin(PRIORITY_MODULES)][["cluster_name", "module_id", "chr", "start", "end"]].copy()


def matched_background_for_targets(targets: pd.DataFrame, max_sets: int = 20) -> pd.DataFrame:
    bg = pd.read_csv(MATCHED, sep="\t")
    bg = bg[(bg["region_type"] == "matched_non_age_window") & (bg["matched_age_cluster"].isin(targets["cluster_name"]))].copy()
    bg["set_num"] = bg["control_set"].astype(str).str.extract(r"(\d+)").astype(float)
    bg = bg[bg["set_num"].fillna(999999) <= max_sets].copy()
    bg = bg.merge(targets[["cluster_name", "module_id"]].rename(columns={"cluster_name": "matched_age_cluster"}), on="matched_age_cluster", how="left")
    bg["cluster_name"] = bg["control_set"].astype(str) + "|" + bg["matched_age_cluster"].astype(str)
    return bg[["cluster_name", "module_id", "chr", "start", "end", "matched_age_cluster", "control_set"]].copy()


def available_tracks() -> list[tuple[str, str, str, Path, str]]:
    tracks = [(tid, mark, stage, path, "available") for tid, mark, stage, path in ATAC_TRACKS if path.exists()]
    if HISTONE_MANIFEST.exists():
        manifest = pd.read_csv(HISTONE_MANIFEST, sep="\t")
        for rec in manifest.to_dict(orient="records"):
            path = Path(str(rec["resolved_path"]))
            if bool(rec.get("file_exists", False)) and path.exists():
                tracks.append((str(rec["track_id"]), str(rec["mark"]), str(rec["stage"]), path, "available"))
            else:
                tracks.append((str(rec["track_id"]), str(rec["mark"]), str(rec["stage"]), path, "missing_histone_input"))
    return tracks


def overlap_analysis(targets: pd.DataFrame, bg: pd.DataFrame) -> pd.DataFrame:
    rows = []
    for track_id, mark, stage, path, status in available_tracks():
        if status != "available":
            for module_id in PRIORITY_MODULES:
                rows.append(
                    {
                        "track_id": track_id,
                        "mark": mark,
                        "stage": stage,
                        "module_id": module_id,
                        "analysis_status": "not_run_missing_histone_input",
                        "target_n": int((targets["module_id"] == module_id).sum()),
                        "target_overlap_n": np.nan,
                        "target_overlap_fraction": np.nan,
                        "background_n": np.nan,
                        "background_overlap_n": np.nan,
                        "background_overlap_fraction": np.nan,
                        "overlap_odds_ratio": np.nan,
                        "fisher_p_greater": np.nan,
                        "track_path": str(path),
                    }
                )
            continue
        peaks = read_bed(path)
        t_flags = overlap_flags(targets, peaks)
        b_flags = overlap_flags(bg, peaks)
        t = targets.copy()
        b = bg.copy()
        t["overlap"] = t_flags
        b["overlap"] = b_flags
        for module_id in PRIORITY_MODULES:
            ts = t[t["module_id"] == module_id]
            bs = b[b["module_id"] == module_id]
            a = int(ts["overlap"].sum())
            b_no = int(len(ts) - a)
            c = int(bs["overlap"].sum())
            d = int(len(bs) - c)
            rows.append(
                {
                    "track_id": track_id,
                    "mark": mark,
                    "stage": stage,
                    "module_id": module_id,
                    "analysis_status": "completed",
                    "target_n": int(len(ts)),
                    "target_overlap_n": a,
                    "target_overlap_fraction": float(a / len(ts)) if len(ts) else 0.0,
                    "background_n": int(len(bs)),
                    "background_overlap_n": c,
                    "background_overlap_fraction": float(c / len(bs)) if len(bs) else 0.0,
                    "overlap_odds_ratio": odds_ratio(a, b_no, c, d),
                    "fisher_p_greater": fisher_greater(a, b_no, c, d),
                    "track_path": str(path),
                }
            )
    out = pd.DataFrame(rows)
    done = out["analysis_status"] == "completed"
    out["fisher_q_BH"] = np.nan
    if done.any():
        out.loc[done, "fisher_q_BH"] = fdr_bh(out.loc[done, "fisher_p_greater"]).to_numpy()
    out.to_csv(OUT_OVERLAP, sep="\t", index=False)
    return out


def stage_delta(overlap: pd.DataFrame) -> pd.DataFrame:
    rows = []
    done = overlap[overlap["analysis_status"] == "completed"].copy()
    if done.empty:
        out = pd.DataFrame()
        out.to_csv(OUT_STAGE, sep="\t", index=False)
        return out
    for module_id in PRIORITY_MODULES:
        sub = done[done["module_id"] == module_id]
        atac8 = sub[(sub["mark"] == "ATAC") & (sub["stage"] == "8-cell")]["target_overlap_fraction"].mean()
        atacticm = sub[(sub["mark"] == "ATAC") & (sub["stage"] == "ICM")]["target_overlap_fraction"].mean()
        rows.append(
            {
                "module_id": module_id,
                "mark": "ATAC",
                "stage_delta": "8-cell_to_ICM_proxy",
                "mean_signal_start": float(atac8) if np.isfinite(atac8) else np.nan,
                "mean_signal_end": float(atacticm) if np.isfinite(atacticm) else np.nan,
                "delta_end_minus_start": float(atacticm - atac8) if np.isfinite(atac8) and np.isfinite(atacticm) else np.nan,
                "interpretation": "GSE101571 has 8-cell and ICM tracks locally; this is not a direct morula ATAC delta.",
            }
        )
    out = pd.DataFrame(rows)
    out.to_csv(OUT_STAGE, sep="\t", index=False)
    return out


def zscore(x: np.ndarray) -> np.ndarray:
    x = np.asarray(x, dtype=float)
    sd = np.nanstd(x, ddof=1)
    if not np.isfinite(sd) or sd == 0:
        return np.zeros_like(x)
    return (x - np.nanmean(x)) / sd


def control_features(overlap: pd.DataFrame) -> pd.DataFrame:
    basis = pd.read_csv(MODULE_BASIS, sep="\t")
    basis = basis[basis["module_id"].isin(PRIORITY_MODULES)].copy()
    rows = []
    done = overlap[overlap["analysis_status"] == "completed"].copy()
    for track_id, sub in done.groupby("track_id"):
        vals = []
        for module_id in PRIORITY_MODULES:
            rec = sub[sub["module_id"] == module_id].iloc[0]
            vals.append(float(rec["target_overlap_fraction"] - rec["background_overlap_fraction"]))
        zvals = zscore(np.asarray(vals))
        for module_id, val, zv in zip(PRIORITY_MODULES, vals, zvals):
            meta = sub[sub["module_id"] == module_id].iloc[0]
            rows.append(
                {
                    "feature_set": f"{track_id}_chromatin_only",
                    "module_id": module_id,
                    "control_value": val,
                    "control_value_z": zv,
                    "top_TFs": "",
                    "control_modality": "chromatin",
                    "mark": meta["mark"],
                    "stage": meta["stage"],
                    "leakage_status": "methylation_non_leaking_external_chromatin_overlap",
                    "description": "Module target overlap fraction minus matched-background overlap fraction.",
                }
            )

    # Gate the q05 zero-filled motif signal by available ATAC module enrichment.
    if MOTIF_FEATURES.exists() and done[done["mark"] == "ATAC"].shape[0]:
        motif = pd.read_csv(MOTIF_FEATURES, sep="\t")
        motif = motif[motif["feature_set"] == "q05_zero_filled_zscore_rebuilt"].copy()
        if len(motif):
            atac_gate = done[(done["mark"] == "ATAC") & (done["stage"] == "8-cell")].groupby("module_id")[
                ["target_overlap_fraction", "background_overlap_fraction"]
            ].mean()
            for _, m in motif.iterrows():
                module_id = str(m["module_id"])
                if module_id not in atac_gate.index:
                    gate = 0.0
                else:
                    gate = max(0.0, float(atac_gate.loc[module_id, "target_overlap_fraction"] - atac_gate.loc[module_id, "background_overlap_fraction"]))
                rows.append(
                    {
                        "feature_set": "motif_TF_x_ATAC8cell_gate",
                        "module_id": module_id,
                        "control_value": float(m["control_value"]) * gate,
                        "control_value_z": np.nan,
                        "top_TFs": m.get("top_TFs", ""),
                        "control_modality": "chromatin_gated_TF",
                        "mark": "ATAC",
                        "stage": "8-cell",
                        "leakage_status": "methylation_non_leaking_motif_TF_ATAC_gated_no_morula_ATAC",
                        "description": "q<=0.05 motif x TF activity multiplied by positive ATAC 8-cell target-background gate.",
                    }
                )
    feat = pd.DataFrame(rows)
    if feat.empty:
        feat.to_csv(OUT_FEATURES, sep="\t", index=False)
        return feat
    # Re-zscore within feature_set after all control values are assembled, except all-zero sets.
    parts = []
    for _, sub in feat.groupby("feature_set"):
        sub = sub.copy()
        if sub["control_value_z"].isna().any():
            sub["control_value_z"] = zscore(pd.to_numeric(sub["control_value"], errors="coerce").fillna(0.0).to_numpy(dtype=float))
        parts.append(sub)
    feat = pd.concat(parts, ignore_index=True)
    feat = feat.merge(basis[["module_id", "n_DMRs", "latent_control_PC1", "latent_control_PC2", "latent_control_PC3", "latent_control_norm", "ridge_weight"]], on="module_id", how="left")
    feat.to_csv(OUT_FEATURES, sep="\t", index=False)
    return feat


def evaluate_features(features: pd.DataFrame) -> tuple[pd.DataFrame, pd.DataFrame]:
    matrix, ann, pairs = load_inputs()
    mu, sd, components, score_df = fit_latent_basis(matrix, 3)
    coef, _, _ = fit_operator(score_df, pairs, train_stages=set(PRE_MORULA_STAGES), lam=1000.0)
    strict_z = simulate_strict_pre_morula(score_df, ann, coef, n_steps=12)
    obs_z = score_df.loc[stage_ids(ann, "morula")].to_numpy(dtype=float)
    obs_dmr = matrix.loc[stage_ids(ann, "morula")].to_numpy(dtype=float)
    basin = basin_definition(obs_z)
    residual_z = obs_z.mean(axis=0) - strict_z.mean(axis=0)
    rng = np.random.default_rng(20260526)
    rows = []
    ablation = []
    baseline_dmr = decode_latent(strict_z, mu, sd, components)
    rows.append(
        {
            "feature_set": "methylation_only_baseline",
            "control_model": "methylation_only_baseline",
            "validation_status": "baseline",
            "n_features": 0,
            "PC1_control": 0.0,
            "PC2_control": 0.0,
            "PC3_control": 0.0,
            "direction_cosine_to_measured_correction": np.nan,
            "PC3_negative_pull_recovered": 0.0,
            **distribution_metrics(strict_z, obs_z, baseline_dmr, obs_dmr, basin, rng),
        }
    )
    for feature_set, sub in features.groupby("feature_set"):
        dirs = sub[["latent_control_PC1", "latent_control_PC2", "latent_control_PC3"]].to_numpy(dtype=float)
        u = pd.to_numeric(sub["control_value_z"], errors="coerce").fillna(0.0).to_numpy(dtype=float)
        control = (dirs * u[:, None]).sum(axis=0)
        for suffix, vec, status in [("", control, "feature_defined"), ("_sign_flip", -control, "sign_flip_control")]:
            pred = strict_z + vec[None, :]
            pred_dmr = decode_latent(pred, mu, sd, components)
            rows.append(
                {
                    "feature_set": feature_set,
                    "control_model": feature_set + suffix,
                    "validation_status": status,
                    "n_features": int(len(sub)),
                    "PC1_control": float(vec[0]),
                    "PC2_control": float(vec[1]),
                    "PC3_control": float(vec[2]),
                    "direction_cosine_to_measured_correction": cosine(vec, residual_z),
                    "PC3_negative_pull_recovered": float(-vec[2] / (-residual_z[2])) if residual_z[2] < 0 else np.nan,
                    **distribution_metrics(pred, obs_z, pred_dmr, obs_dmr, basin, rng),
                }
            )
        for module_id in PRIORITY_MODULES:
            keep = sub[sub["module_id"] != module_id]
            if keep.empty:
                continue
            vec = (keep[["latent_control_PC1", "latent_control_PC2", "latent_control_PC3"]].to_numpy(dtype=float) * pd.to_numeric(keep["control_value_z"], errors="coerce").fillna(0.0).to_numpy(dtype=float)[:, None]).sum(axis=0)
            pred = strict_z + vec[None, :]
            pred_dmr = decode_latent(pred, mu, sd, components)
            ablation.append(
                {
                    "feature_set": feature_set,
                    "ablation": f"remove_{module_id}",
                    "PC1_control": float(vec[0]),
                    "PC2_control": float(vec[1]),
                    "PC3_control": float(vec[2]),
                    "direction_cosine_to_measured_correction": cosine(vec, residual_z),
                    "PC3_negative_pull_recovered": float(-vec[2] / (-residual_z[2])) if residual_z[2] < 0 else np.nan,
                    **distribution_metrics(pred, obs_z, pred_dmr, obs_dmr, basin, rng),
                }
            )
    metrics = pd.DataFrame(rows)
    abl = pd.DataFrame(ablation)
    metrics.to_csv(OUT_METRICS, sep="\t", index=False)
    abl.to_csv(OUT_ABLATION, sep="\t", index=False)
    return metrics, abl


def make_svg(metrics: pd.DataFrame) -> None:
    rows = metrics[metrics["validation_status"] == "feature_defined"].sort_values("pred_basin_occupancy_q90", ascending=False).head(12)
    width, height = 940, 430
    left, right, top, bottom = 80, 30, 45, 145
    plot_w = width - left - right
    plot_h = height - top - bottom
    gap = plot_w / max(len(rows), 1)
    bar_w = gap * 0.62
    lines = [
        f'<svg xmlns="http://www.w3.org/2000/svg" width="{width}" height="{height}" viewBox="0 0 {width} {height}">',
        '<rect width="100%" height="100%" fill="white"/>',
        f'<text x="{left}" y="28" font-family="Arial" font-size="18" font-weight="700">Chromatin-gated control models</text>',
        f'<line x1="{left}" y1="{height-bottom}" x2="{width-right}" y2="{height-bottom}" stroke="#222"/>',
        f'<line x1="{left}" y1="{top}" x2="{left}" y2="{height-bottom}" stroke="#222"/>',
    ]
    for tick in [0.044, 0.2, 0.3, 0.5, 0.875, 1.0]:
        y = height - bottom - tick * plot_h
        color = "#b23b3b" if tick in [0.044, 0.875] else "#dddddd"
        lines.append(f'<line x1="{left}" y1="{y:.2f}" x2="{width-right}" y2="{y:.2f}" stroke="{color}" stroke-dasharray="4 4"/>')
        lines.append(f'<text x="{left-8}" y="{y+4:.2f}" text-anchor="end" font-family="Arial" font-size="10">{tick:.3g}</text>')
    for i, row in enumerate(rows.itertuples()):
        x = left + i * gap + (gap - bar_w) / 2
        val = float(row.pred_basin_occupancy_q90)
        y = height - bottom - val * plot_h
        h = val * plot_h
        lines.append(f'<rect x="{x:.2f}" y="{y:.2f}" width="{bar_w:.2f}" height="{h:.2f}" fill="#2c6f5a"/>')
        lines.append(f'<text x="{x+bar_w/2:.2f}" y="{y-5:.2f}" text-anchor="middle" font-family="Arial" font-size="10">{val:.3f}</text>')
        label = row.feature_set.replace("_", " ")
        lines.append(f'<text x="{x+bar_w/2:.2f}" y="{height-bottom+18}" text-anchor="end" transform="rotate(-45 {x+bar_w/2:.2f} {height-bottom+18})" font-family="Arial" font-size="9">{label}</text>')
    lines.append(f'<text x="{left}" y="{height-22}" font-family="Arial" font-size="12">Red guides: methylation-only baseline and observed morula occupancy.</text>')
    lines.append("</svg>")
    OUT_SVG.write_text("\n".join(lines), encoding="utf-8")


def write_doc(region: pd.DataFrame, overlap: pd.DataFrame, metrics: pd.DataFrame) -> None:
    completed = overlap[overlap["analysis_status"] == "completed"].copy()
    best_overlap = completed.sort_values("overlap_odds_ratio", ascending=False).head(8)
    feature_defined = metrics[metrics["validation_status"] == "feature_defined"].sort_values("pred_basin_occupancy_q90", ascending=False)
    lines = [
        "# Residual Module Chromatin-Gated Control Analysis",
        "",
        "Status: `completed_with_available_ATAC_and_missing_histone_inputs`",
        "",
        "Goal: test whether M05/M01/M12/M02/M10 residual modules are supported by external chromatin state, and whether chromatin-gated controls explain the measured correction term.",
        "",
        "## Region Composition",
        "",
    ]
    for row in region.itertuples():
        lines.append(
            f"- {row.module_id}: n={row.n_DMR}, promoter2kb={row.promoter_2kb_fraction:.3f}, intergenic={row.intergenic_fraction:.3f}, residual_sum={row.sum_abs_residual_contribution:.3f}"
        )
    lines += ["", "## Strongest Available Chromatin Overlaps", ""]
    if len(best_overlap):
        for row in best_overlap.itertuples():
            lines.append(
                f"- {row.module_id} {row.track_id}: target={row.target_overlap_fraction:.3f}, background={row.background_overlap_fraction:.3f}, OR={row.overlap_odds_ratio:.3f}, q={row.fisher_q_BH:.3g}"
            )
    else:
        lines.append("- No completed external chromatin overlap tracks were available.")
    lines += ["", "## Control Dynamics", ""]
    for row in feature_defined.head(8).itertuples():
        lines.append(
            f"- {row.feature_set}: occupancy={row.pred_basin_occupancy_q90:.3f}, cosine={row.direction_cosine_to_measured_correction:.3f}, PC3_recovery={row.PC3_negative_pull_recovered:.3f}"
        )
    missing_histone = overlap[overlap["analysis_status"] == "not_run_missing_histone_input"]["track_id"].drop_duplicates().tolist()
    lines += [
        "",
        "## Boundary",
        "",
        "The available local chromatin evidence is ATAC 8-cell/ICM only. This is not direct morula chromatin validation.",
        "",
        "Missing histone tracks are input/access boundaries, not negative histone evidence:",
        "",
    ]
    lines += [f"- {x}" for x in missing_histone] if missing_histone else ["- none"]
    OUT_DOC.write_text("\n".join(lines) + "\n", encoding="utf-8")


def main() -> None:
    RESULTS.mkdir(parents=True, exist_ok=True)
    FIGURES.mkdir(parents=True, exist_ok=True)
    DOCS.mkdir(parents=True, exist_ok=True)

    region = region_composition()
    targets = target_regions()
    bg = matched_background_for_targets(targets, max_sets=20)
    overlap = overlap_analysis(targets, bg)
    stage_delta(overlap)
    features = control_features(overlap)
    metrics, ablation = evaluate_features(features) if len(features) else (pd.DataFrame(), pd.DataFrame())
    if len(metrics):
        make_svg(metrics)
    write_doc(region, overlap, metrics if len(metrics) else pd.DataFrame())
    OUT_MANIFEST.write_text(
        json.dumps(
            {
                "status": "completed_with_available_ATAC_and_missing_histone_inputs",
                "n_target_regions": int(len(targets)),
                "n_background_regions": int(len(bg)),
                "outputs": [str(OUT_REGION), str(OUT_OVERLAP), str(OUT_STAGE), str(OUT_FEATURES), str(OUT_METRICS), str(OUT_ABLATION), str(OUT_SVG), str(OUT_DOC)],
            },
            indent=2,
        )
        + "\n",
        encoding="utf-8",
    )
    top_metrics = metrics[metrics["validation_status"] == "feature_defined"].sort_values("pred_basin_occupancy_q90", ascending=False).head(8).to_dict(orient="records") if len(metrics) else []
    print(json.dumps({"status": "completed", "top_metrics": top_metrics}, indent=2))


if __name__ == "__main__":
    main()
```
