## Supplemental code for "An entropy-based diagnostic framework for characterizing methylation state dynamics during preimplantation development": run_residual_module_histone_state_control.py

```
from __future__ import annotations

import gzip
import json
import math
import sys
from pathlib import Path

import numpy as np
import pandas as pd

try:
    import pyBigWig  # type: ignore
except ImportError:  # pragma: no cover - optional runtime dependency
    pyBigWig = None


BASE = Path(r"E:\CSB_TRO_operator_time_DMR_dynamics_2026-05-25")
MAIN = Path(r"C:\Users\18068\Desktop\CSB_TRO_Project_2026-05-24")
CODE = BASE / "code"
RESULTS = BASE / "results"
FIGURES = BASE / "figures"
DOCS = BASE / "docs"
EXTERNAL = BASE / "external"
HISTONE = EXTERNAL / "histone"
sys.path.insert(0, str(CODE))

from run_basin_residual_control_field import PRE_MORULA_STAGES, cosine, simulate_strict_pre_morula  # noqa: E402
from run_morula_basin_sde import (  # noqa: E402
    basin_definition,
    decode_latent,
    distribution_metrics,
    fit_latent_basis,
    fit_operator,
    load_inputs,
    stage_ids,
)


PRIORITY_MODULES = ["M05", "M01", "M12", "M02", "M10"]
STAGES = ["8cell", "morula", "blastocyst"]
MARKS = ["H3K27ac", "H3K4me3", "H3K27me3"]

REGION_ANNOT = RESULTS / "CSB_TRO_residual_module_region_annotation.tsv"
MODULE_BASIS = RESULTS / "CSB_TRO_missing_control_term_module_basis.tsv"
MATCHED = MAIN / "input_tables" / "GSE81233_matched_non_age_window_regions_n100.tsv"
MOTIF_FEATURES = RESULTS / "CSB_TRO_module_TF_activity_control_panel_features.tsv"

OUT_MANIFEST_TSV = HISTONE / "histone_state_peak_manifest.tsv"
OUT_OVERLAP = RESULTS / "CSB_TRO_module_histone_overlap.tsv"
OUT_STAGE = RESULTS / "CSB_TRO_module_histone_stage_delta.tsv"
OUT_FEATURES = RESULTS / "CSB_TRO_histone_gated_TF_activity.tsv"
OUT_METRICS = RESULTS / "CSB_TRO_histone_gated_control_metrics.tsv"
OUT_ABLATION = RESULTS / "CSB_TRO_histone_gated_control_ablation.tsv"
OUT_MANIFEST = RESULTS / "CSB_TRO_histone_state_control_manifest.json"
OUT_SVG = FIGURES / "CSB_TRO_histone_gated_control.svg"
OUT_DOC = DOCS / "CSB_TRO_histone_state_control_summary.md"


def expected_tracks() -> pd.DataFrame:
    rows = []
    for mark in MARKS:
        for stage in STAGES:
            stem = HISTONE / f"{mark}_{stage}.hg19"
            candidates = [
                (Path(str(stem) + ".bed"), "BED_peak_overlap"),
                (Path(str(stem) + ".bed.gz"), "BED_peak_overlap"),
                (Path(str(stem) + ".narrowPeak"), "BED_peak_overlap"),
                (Path(str(stem) + ".narrowPeak.gz"), "BED_peak_overlap"),
                (Path(str(stem) + ".broadPeak"), "BED_peak_overlap"),
                (Path(str(stem) + ".broadPeak.gz"), "BED_peak_overlap"),
                (Path(str(stem) + ".bw"), "bigWig_signal"),
                (Path(str(stem) + ".bigWig"), "bigWig_signal"),
            ]
            resolved, input_type = next(((p, t) for p, t in candidates if p.exists()), candidates[0])
            runnable = resolved.exists() and (input_type != "bigWig_signal" or pyBigWig is not None)
            status = "ready" if runnable else "missing_processed_input" if not resolved.exists() else "missing_pyBigWig_dependency"
            rows.append(
                {
                    "track_id": f"{mark}_{stage}",
                    "mark": mark,
                    "stage": stage,
                    "genome_build": "hg19/GRCh37",
                    "expected_path": str(candidates[0][0]),
                    "resolved_path": str(resolved),
                    "file_exists": bool(resolved.exists()),
                    "input_type": input_type,
                    "analysis_ready": bool(runnable),
                    "analysis_status": status,
                    "size_bytes": int(resolved.stat().st_size) if resolved.exists() else 0,
                    "format": "BED/narrowPeak/broadPeak optionally gzip-compressed, or bigWig signal",
                }
            )
    out = pd.DataFrame(rows)
    HISTONE.mkdir(parents=True, exist_ok=True)
    out.to_csv(OUT_MANIFEST_TSV, sep="\t", index=False)
    return out


def read_bed(path: Path) -> pd.DataFrame:
    opener = gzip.open if path.suffix == ".gz" else open
    rows = []
    with opener(path, "rt", encoding="utf-8", errors="replace") as handle:
        for line in handle:
            if not line.strip() or line.startswith(("#", "track", "browser")):
                continue
            parts = line.rstrip("\n").split("\t")
            if len(parts) < 3:
                continue
            try:
                start = int(float(parts[1]))
                end = int(float(parts[2]))
            except ValueError:
                continue
            rows.append({"chr": parts[0], "start": start, "end": end})
    return pd.DataFrame(rows)


def bigwig_mean_signal(regions: pd.DataFrame, path: Path) -> np.ndarray:
    vals = np.full(len(regions), np.nan, dtype=float)
    if pyBigWig is None:
        return vals
    bw = pyBigWig.open(str(path))
    try:
        chroms = bw.chroms()
        for i, row in enumerate(regions.itertuples()):
            chrom = str(row.chr)
            if chrom not in chroms:
                alt = chrom[3:] if chrom.startswith("chr") else "chr" + chrom
                chrom = alt if alt in chroms else chrom
            if chrom not in chroms:
                continue
            start = max(0, int(row.start))
            end = min(int(row.end), int(chroms[chrom]))
            if end <= start:
                continue
            val = bw.stats(chrom, start, end, type="mean")[0]
            vals[i] = float(val) if val is not None and np.isfinite(val) else 0.0
    finally:
        bw.close()
    return vals


def overlap_flags(regions: pd.DataFrame, peaks: pd.DataFrame) -> np.ndarray:
    flags = np.zeros(len(regions), dtype=bool)
    if peaks.empty or regions.empty:
        return flags
    by_chr = {c: sub[["start", "end"]].to_numpy(dtype=int) for c, sub in peaks.groupby("chr")}
    for i, row in enumerate(regions.itertuples()):
        arr = by_chr.get(str(row.chr))
        if arr is not None:
            flags[i] = bool(np.any((arr[:, 0] < int(row.end)) & (arr[:, 1] > int(row.start))))
    return flags


def log_comb(n: int, k: int) -> float:
    if k < 0 or k > n:
        return float("-inf")
    return math.lgamma(n + 1) - math.lgamma(k + 1) - math.lgamma(n - k + 1)


def fisher_greater(a: int, b: int, c: int, d: int) -> float:
    n = a + b + c + d
    row1 = a + b
    col1 = a + c
    lo = max(0, row1 - (n - col1))
    hi = min(row1, col1)
    denom = log_comb(n, row1)
    vals = [log_comb(col1, x) + log_comb(n - col1, row1 - x) - denom for x in range(max(a, lo), hi + 1)]
    if not vals:
        return 1.0
    m = max(vals)
    return float(min(1.0, math.exp(m) * sum(math.exp(v - m) for v in vals)))


def fdr_bh(p: pd.Series) -> pd.Series:
    x = pd.to_numeric(p, errors="coerce").fillna(1.0).to_numpy(dtype=float)
    order = np.argsort(x)
    q = np.empty_like(x)
    prev = 1.0
    n = len(x)
    for rank, idx in enumerate(order[::-1], start=1):
        true_rank = n - rank + 1
        val = x[idx] * n / true_rank
        prev = min(prev, val)
        q[idx] = prev
    return pd.Series(np.minimum(q, 1.0), index=p.index)


def odds_ratio(a: int, b: int, c: int, d: int) -> float:
    return float(((a + 0.5) * (d + 0.5)) / ((b + 0.5) * (c + 0.5)))


def targets_and_background() -> tuple[pd.DataFrame, pd.DataFrame]:
    annot = pd.read_csv(REGION_ANNOT, sep="\t")
    targets = annot[annot["module_id"].isin(PRIORITY_MODULES)][["cluster_name", "module_id", "chr", "start", "end"]].copy()
    bg = pd.read_csv(MATCHED, sep="\t")
    bg = bg[(bg["region_type"] == "matched_non_age_window") & (bg["matched_age_cluster"].isin(targets["cluster_name"]))].copy()
    bg["set_num"] = bg["control_set"].astype(str).str.extract(r"(\d+)").astype(float)
    bg = bg[bg["set_num"].fillna(999999) <= 20].copy()
    bg = bg.merge(targets[["cluster_name", "module_id"]].rename(columns={"cluster_name": "matched_age_cluster"}), on="matched_age_cluster", how="left")
    bg["cluster_name"] = bg["control_set"].astype(str) + "|" + bg["matched_age_cluster"].astype(str)
    return targets, bg[["cluster_name", "module_id", "chr", "start", "end", "matched_age_cluster", "control_set"]].copy()


def overlap_analysis(manifest: pd.DataFrame, targets: pd.DataFrame, bg: pd.DataFrame) -> pd.DataFrame:
    rows = []
    for rec in manifest.to_dict(orient="records"):
        track_id = str(rec["track_id"])
        mark = str(rec["mark"])
        stage = str(rec["stage"])
        path = Path(str(rec["resolved_path"]))
        input_type = str(rec.get("input_type", "BED_peak_overlap"))
        if not bool(rec.get("analysis_ready", rec.get("file_exists", False))):
            for module_id in PRIORITY_MODULES:
                rows.append(
                    {
                        "track_id": track_id,
                        "mark": mark,
                        "stage": stage,
                        "module_id": module_id,
                        "analysis_status": str(rec.get("analysis_status", "not_run_missing_histone_input")).replace("missing_processed_input", "not_run_missing_histone_input"),
                        "input_type": input_type,
                        "target_n": int((targets["module_id"] == module_id).sum()),
                        "target_overlap_n": np.nan,
                        "target_overlap_fraction": np.nan,
                        "background_n": np.nan,
                        "background_overlap_n": np.nan,
                        "background_overlap_fraction": np.nan,
                        "overlap_odds_ratio": np.nan,
                        "fisher_p_greater": np.nan,
                        "track_path": str(path),
                    }
                )
            continue
        t = targets.copy()
        b = bg.copy()
        if input_type == "bigWig_signal":
            t["signal"] = bigwig_mean_signal(t, path)
            b["signal"] = bigwig_mean_signal(b, path)
        else:
            peaks = read_bed(path)
            t["overlap"] = overlap_flags(t, peaks)
            b["overlap"] = overlap_flags(b, peaks)
        for module_id in PRIORITY_MODULES:
            ts = t[t["module_id"] == module_id]
            bs = b[b["module_id"] == module_id]
            if input_type == "bigWig_signal":
                t_sig = pd.to_numeric(ts["signal"], errors="coerce").fillna(0.0).to_numpy(dtype=float)
                b_sig = pd.to_numeric(bs["signal"], errors="coerce").fillna(0.0).to_numpy(dtype=float)
                target_mean = float(np.nanmean(t_sig)) if len(t_sig) else 0.0
                background_mean = float(np.nanmean(b_sig)) if len(b_sig) else 0.0
                a = int(np.sum(t_sig > 0))
                b_no = int(len(t_sig) - a)
                c = int(np.sum(b_sig > 0))
                d = int(len(b_sig) - c)
                target_value = target_mean
                background_value = background_mean
                effect = target_mean - background_mean
                pval = np.nan
            else:
                a = int(ts["overlap"].sum())
                b_no = int(len(ts) - a)
                c = int(bs["overlap"].sum())
                d = int(len(bs) - c)
                target_value = float(a / len(ts)) if len(ts) else 0.0
                background_value = float(c / len(bs)) if len(bs) else 0.0
                effect = target_value - background_value
                pval = fisher_greater(a, b_no, c, d)
            rows.append(
                {
                    "track_id": track_id,
                    "mark": mark,
                    "stage": stage,
                    "module_id": module_id,
                    "analysis_status": "completed",
                    "input_type": input_type,
                    "target_n": int(len(ts)),
                    "target_overlap_n": a,
                    "target_overlap_fraction": target_value,
                    "background_n": int(len(bs)),
                    "background_overlap_n": c,
                    "background_overlap_fraction": background_value,
                    "target_minus_background_signal": effect,
                    "overlap_odds_ratio": odds_ratio(a, b_no, c, d),
                    "fisher_p_greater": pval,
                    "track_path": str(path),
                }
            )
    out = pd.DataFrame(rows)
    out["fisher_q_BH"] = np.nan
    done = out["analysis_status"] == "completed"
    if done.any():
        out.loc[done, "fisher_q_BH"] = fdr_bh(out.loc[done, "fisher_p_greater"]).to_numpy()
    out.to_csv(OUT_OVERLAP, sep="\t", index=False)
    return out


def stage_delta(overlap: pd.DataFrame) -> pd.DataFrame:
    done = overlap[overlap["analysis_status"] == "completed"].copy()
    rows = []
    for mark in MARKS:
        for module_id in PRIORITY_MODULES:
            sub = done[(done["mark"] == mark) & (done["module_id"] == module_id)]
            vals = {str(r.stage): float(r.target_overlap_fraction - r.background_overlap_fraction) for r in sub.itertuples()}
            for start, end in [("8cell", "morula"), ("morula", "blastocyst")]:
                rows.append(
                    {
                        "module_id": module_id,
                        "mark": mark,
                        "stage_delta": f"{start}_to_{end}",
                        "start_enrichment_fraction": vals.get(start, np.nan),
                        "end_enrichment_fraction": vals.get(end, np.nan),
                        "delta_end_minus_start": vals.get(end, np.nan) - vals.get(start, np.nan) if start in vals and end in vals else np.nan,
                        "analysis_status": "completed" if start in vals and end in vals else "not_run_missing_histone_input",
                    }
                )
    out = pd.DataFrame(rows)
    out.to_csv(OUT_STAGE, sep="\t", index=False)
    return out


def zscore(x: np.ndarray) -> np.ndarray:
    x = np.asarray(x, dtype=float)
    sd = np.nanstd(x, ddof=1)
    if not np.isfinite(sd) or sd == 0:
        return np.zeros_like(x)
    return (x - np.nanmean(x)) / sd


def build_features(overlap: pd.DataFrame, stage: pd.DataFrame) -> pd.DataFrame:
    done = overlap[overlap["analysis_status"] == "completed"].copy()
    basis = pd.read_csv(MODULE_BASIS, sep="\t")
    basis = basis[basis["module_id"].isin(PRIORITY_MODULES)].copy()
    rows = []
    for track_id, sub in done.groupby("track_id"):
        vals = []
        for module_id in PRIORITY_MODULES:
            rec = sub[sub["module_id"] == module_id].iloc[0]
            vals.append(float(rec["target_overlap_fraction"] - rec["background_overlap_fraction"]))
        for module_id, val, zv in zip(PRIORITY_MODULES, vals, zscore(np.asarray(vals))):
            meta = sub[sub["module_id"] == module_id].iloc[0]
            rows.append(
                {
                    "feature_set": f"{track_id}_histone_only",
                    "module_id": module_id,
                    "control_value": val,
                    "control_value_z": zv,
                    "control_modality": "histone",
                    "mark": meta["mark"],
                    "stage": meta["stage"],
                    "leakage_status": "methylation_non_leaking_external_histone_overlap",
                    "description": "Module target histone overlap fraction minus matched background.",
                }
            )
    # Stage-gain features when both stages exist.
    complete_delta = stage[stage["analysis_status"] == "completed"].copy()
    for key, sub in complete_delta.groupby(["mark", "stage_delta"]):
        vals = []
        for module_id in PRIORITY_MODULES:
            r = sub[sub["module_id"] == module_id]
            vals.append(float(r.iloc[0]["delta_end_minus_start"]) if len(r) else 0.0)
        for module_id, val, zv in zip(PRIORITY_MODULES, vals, zscore(np.asarray(vals))):
            rows.append(
                {
                    "feature_set": f"{key[0]}_{key[1]}_histone_delta",
                    "module_id": module_id,
                    "control_value": val,
                    "control_value_z": zv,
                    "control_modality": "histone_delta",
                    "mark": key[0],
                    "stage": key[1],
                    "leakage_status": "methylation_non_leaking_external_histone_stage_delta",
                    "description": "Histone target-background enrichment delta between stages.",
                }
            )
    # Histone-gated motif TF: only possible when histone features exist.
    if MOTIF_FEATURES.exists() and rows:
        motif = pd.read_csv(MOTIF_FEATURES, sep="\t")
        motif = motif[motif["feature_set"] == "q05_zero_filled_zscore_rebuilt"].copy()
        for feature_set in sorted({r["feature_set"] for r in rows if r["control_modality"] == "histone"}):
            hist = pd.DataFrame([r for r in rows if r["feature_set"] == feature_set]).set_index("module_id")
            for rec in motif.to_dict(orient="records"):
                module_id = str(rec["module_id"])
                gate = max(0.0, float(hist.loc[module_id, "control_value"])) if module_id in hist.index else 0.0
                rows.append(
                    {
                        "feature_set": f"motif_TF_x_{feature_set}",
                        "module_id": module_id,
                        "control_value": float(rec["control_value"]) * gate,
                        "control_value_z": np.nan,
                        "control_modality": "histone_gated_TF",
                        "mark": str(hist["mark"].iloc[0]) if len(hist) else "",
                        "stage": str(hist["stage"].iloc[0]) if len(hist) else "",
                        "leakage_status": "methylation_non_leaking_motif_TF_histone_gated",
                        "description": "q<=0.05 motif x TF activity multiplied by positive histone target-background gate.",
                    }
                )
    feat = pd.DataFrame(rows)
    if feat.empty:
        feat = pd.DataFrame(
            [
                {
                    "feature_set": "not_run_missing_histone_input",
                    "module_id": "",
                    "control_value": np.nan,
                    "control_value_z": np.nan,
                    "control_modality": "histone",
                    "mark": "",
                    "stage": "",
                    "leakage_status": "not_run_missing_histone_input",
                    "description": "No local histone peak BED files available.",
                }
            ]
        )
        feat.to_csv(OUT_FEATURES, sep="\t", index=False)
        return feat
    parts = []
    for _, sub in feat.groupby("feature_set"):
        sub = sub.copy()
        if sub["control_value_z"].isna().any():
            sub["control_value_z"] = zscore(pd.to_numeric(sub["control_value"], errors="coerce").fillna(0.0).to_numpy(dtype=float))
        parts.append(sub)
    feat = pd.concat(parts, ignore_index=True)
    feat = feat.merge(basis[["module_id", "n_DMRs", "latent_control_PC1", "latent_control_PC2", "latent_control_PC3", "latent_control_norm", "ridge_weight"]], on="module_id", how="left")
    feat.to_csv(OUT_FEATURES, sep="\t", index=False)
    return feat


def evaluate_features(features: pd.DataFrame) -> tuple[pd.DataFrame, pd.DataFrame]:
    runnable = features[features["leakage_status"] != "not_run_missing_histone_input"].copy()
    if runnable.empty:
        metrics = pd.DataFrame(
            [
                {
                    "feature_set": "not_run_missing_histone_input",
                    "control_model": "not_run_missing_histone_input",
                    "validation_status": "not_run_missing_histone_input",
                    "reason": "No local histone peak BED files were available.",
                }
            ]
        )
        metrics.to_csv(OUT_METRICS, sep="\t", index=False)
        pd.DataFrame().to_csv(OUT_ABLATION, sep="\t", index=False)
        return metrics, pd.DataFrame()
    matrix, ann, pairs = load_inputs()
    mu, sd, components, score_df = fit_latent_basis(matrix, 3)
    coef, _, _ = fit_operator(score_df, pairs, train_stages=set(PRE_MORULA_STAGES), lam=1000.0)
    strict_z = simulate_strict_pre_morula(score_df, ann, coef, n_steps=12)
    obs_z = score_df.loc[stage_ids(ann, "morula")].to_numpy(dtype=float)
    obs_dmr = matrix.loc[stage_ids(ann, "morula")].to_numpy(dtype=float)
    basin = basin_definition(obs_z)
    residual_z = obs_z.mean(axis=0) - strict_z.mean(axis=0)
    rng = np.random.default_rng(20260526)
    rows = []
    abl = []
    baseline_dmr = decode_latent(strict_z, mu, sd, components)
    rows.append({"feature_set": "methylation_only_baseline", "control_model": "methylation_only_baseline", "validation_status": "baseline", **distribution_metrics(strict_z, obs_z, baseline_dmr, obs_dmr, basin, rng)})
    for feature_set, sub in runnable.groupby("feature_set"):
        dirs = sub[["latent_control_PC1", "latent_control_PC2", "latent_control_PC3"]].to_numpy(dtype=float)
        u = pd.to_numeric(sub["control_value_z"], errors="coerce").fillna(0.0).to_numpy(dtype=float)
        control = (dirs * u[:, None]).sum(axis=0)
        for suffix, vec, status in [("", control, "feature_defined"), ("_sign_flip", -control, "sign_flip_control")]:
            pred = strict_z + vec[None, :]
            pred_dmr = decode_latent(pred, mu, sd, components)
            rows.append(
                {
                    "feature_set": feature_set,
                    "control_model": feature_set + suffix,
                    "validation_status": status,
                    "PC1_control": float(vec[0]),
                    "PC2_control": float(vec[1]),
                    "PC3_control": float(vec[2]),
                    "direction_cosine_to_measured_correction": cosine(vec, residual_z),
                    "PC3_negative_pull_recovered": float(-vec[2] / (-residual_z[2])) if residual_z[2] < 0 else np.nan,
                    **distribution_metrics(pred, obs_z, pred_dmr, obs_dmr, basin, rng),
                }
            )
    metrics = pd.DataFrame(rows)
    metrics.to_csv(OUT_METRICS, sep="\t", index=False)
    pd.DataFrame(abl).to_csv(OUT_ABLATION, sep="\t", index=False)
    return metrics, pd.DataFrame(abl)


def make_svg(metrics: pd.DataFrame) -> None:
    if "pred_basin_occupancy_q90" not in metrics.columns:
        return
    rows = metrics[metrics["validation_status"] == "feature_defined"].sort_values("pred_basin_occupancy_q90", ascending=False).head(12)
    if rows.empty:
        return
    width, height = 940, 430
    left, right, top, bottom = 80, 30, 45, 145
    plot_w = width - left - right
    plot_h = height - top - bottom
    gap = plot_w / max(len(rows), 1)
    bar_w = gap * 0.62
    lines = [
        f'<svg xmlns="http://www.w3.org/2000/svg" width="{width}" height="{height}" viewBox="0 0 {width} {height}">',
        '<rect width="100%" height="100%" fill="white"/>',
        f'<text x="{left}" y="28" font-family="Arial" font-size="18" font-weight="700">Histone-state control models</text>',
        f'<line x1="{left}" y1="{height-bottom}" x2="{width-right}" y2="{height-bottom}" stroke="#222"/>',
        f'<line x1="{left}" y1="{top}" x2="{left}" y2="{height-bottom}" stroke="#222"/>',
    ]
    for tick in [0.044, 0.3, 0.5, 0.875, 1.0]:
        y = height - bottom - tick * plot_h
        color = "#b23b3b" if tick in [0.044, 0.875] else "#dddddd"
        lines.append(f'<line x1="{left}" y1="{y:.2f}" x2="{width-right}" y2="{y:.2f}" stroke="{color}" stroke-dasharray="4 4"/>')
        lines.append(f'<text x="{left-8}" y="{y+4:.2f}" text-anchor="end" font-family="Arial" font-size="10">{tick:.3g}</text>')
    for i, row in enumerate(rows.itertuples()):
        x = left + i * gap + (gap - bar_w) / 2
        val = float(row.pred_basin_occupancy_q90)
        y = height - bottom - val * plot_h
        lines.append(f'<rect x="{x:.2f}" y="{y:.2f}" width="{bar_w:.2f}" height="{val * plot_h:.2f}" fill="#6f4d8b"/>')
        lines.append(f'<text x="{x+bar_w/2:.2f}" y="{y-5:.2f}" text-anchor="middle" font-family="Arial" font-size="10">{val:.3f}</text>')
        label = row.feature_set.replace("_", " ")
        lines.append(f'<text x="{x+bar_w/2:.2f}" y="{height-bottom+18}" text-anchor="end" transform="rotate(-45 {x+bar_w/2:.2f} {height-bottom+18})" font-family="Arial" font-size="9">{label}</text>')
    lines.append("</svg>")
    OUT_SVG.write_text("\n".join(lines), encoding="utf-8")


def write_doc(manifest: pd.DataFrame, overlap: pd.DataFrame, metrics: pd.DataFrame) -> None:
    ready = int(manifest["analysis_ready"].sum()) if "analysis_ready" in manifest.columns else int(manifest["file_exists"].sum())
    present = int(manifest["file_exists"].sum())
    missing = manifest.loc[~manifest["analysis_ready"], "track_id"].tolist() if "analysis_ready" in manifest.columns else manifest.loc[~manifest["file_exists"], "track_id"].tolist()
    lines = [
        "# Residual Module Histone-State Control",
        "",
        f"Status: `{'completed_partial_track_mode' if ready else 'not_run_missing_histone_input'}`",
        "",
        "Goal: test whether H3K27ac/H3K4me3/H3K27me3 stage-specific marks explain M05/M01/M12/M02/M10 residual module control.",
        "",
        f"Analysis-ready histone tracks: {ready}/{len(manifest)}",
        "",
        f"Present local histone files: {present}/{len(manifest)}",
        "",
    ]
    if ready == 0:
        lines += [
            "No local histone peak BED files are present. This is an input/access boundary, not negative histone evidence.",
            "",
            "Expected files:",
            "",
        ]
        lines += [f"- `{row.expected_path}`" for row in manifest.itertuples()]
    else:
        best = overlap[overlap["analysis_status"] == "completed"].sort_values("overlap_odds_ratio", ascending=False).head(10)
        lines += ["## Strongest Histone Overlaps", ""]
        for row in best.itertuples():
            lines.append(
                f"- {row.module_id} {row.track_id}: target={row.target_overlap_fraction:.3f}, background={row.background_overlap_fraction:.3f}, OR={row.overlap_odds_ratio:.3f}, q={row.fisher_q_BH:.3g}"
            )
        if "pred_basin_occupancy_q90" in metrics.columns:
            lines += ["", "## Control Dynamics", ""]
            for row in metrics[metrics["validation_status"] == "feature_defined"].sort_values("pred_basin_occupancy_q90", ascending=False).head(8).itertuples():
                lines.append(
                    f"- {row.feature_set}: occupancy={row.pred_basin_occupancy_q90:.3f}, cosine={row.direction_cosine_to_measured_correction:.3f}, PC3_recovery={row.PC3_negative_pull_recovered:.3f}"
                )
    if missing:
        lines += ["", "Missing tracks:", ""]
        lines += [f"- {m}" for m in missing]
    lines += [
        "",
        "Next input priority: processed human early-embryo H3K27ac/H3K4me3/H3K27me3 peak BED or signal tracks for 8-cell, morula, and blastocyst in hg19/GRCh37.",
    ]
    OUT_DOC.write_text("\n".join(lines) + "\n", encoding="utf-8")


def main() -> None:
    RESULTS.mkdir(parents=True, exist_ok=True)
    FIGURES.mkdir(parents=True, exist_ok=True)
    DOCS.mkdir(parents=True, exist_ok=True)
    HISTONE.mkdir(parents=True, exist_ok=True)

    manifest = expected_tracks()
    targets, bg = targets_and_background()
    overlap = overlap_analysis(manifest, targets, bg)
    stage = stage_delta(overlap)
    features = build_features(overlap, stage)
    metrics, ablation = evaluate_features(features)
    make_svg(metrics)
    write_doc(manifest, overlap, metrics)
    OUT_MANIFEST.write_text(
        json.dumps(
            {
                "status": "completed_partial_track_mode" if int(manifest["analysis_ready"].sum()) else "not_run_missing_histone_input",
                "analysis_ready_histone_tracks": int(manifest["analysis_ready"].sum()),
                "present_histone_files": int(manifest["file_exists"].sum()),
                "expected_histone_tracks": int(len(manifest)),
                "partial_track_mode": True,
                "bigwig_supported": pyBigWig is not None,
                "outputs": [str(OUT_MANIFEST_TSV), str(OUT_OVERLAP), str(OUT_STAGE), str(OUT_FEATURES), str(OUT_METRICS), str(OUT_ABLATION), str(OUT_DOC)],
            },
            indent=2,
        )
        + "\n",
        encoding="utf-8",
    )
    print(
        json.dumps(
            {
                "status": "completed_partial_track_mode" if int(manifest["analysis_ready"].sum()) else "not_run_missing_histone_input",
                "analysis_ready_tracks": int(manifest["analysis_ready"].sum()),
                "present_histone_files": int(manifest["file_exists"].sum()),
                "bigwig_supported": pyBigWig is not None,
            },
            indent=2,
        )
    )


if __name__ == "__main__":
    main()
```
