## Supplemental code for "An entropy-based diagnostic framework for characterizing methylation state dynamics during preimplantation development": run_residual_module_motif_tf_activity.py

```
from __future__ import annotations

import json
from pathlib import Path

import numpy as np
import pandas as pd


BASE = Path(r"E:\CSB_TRO_operator_time_DMR_dynamics_2026-05-25")
EXTERNAL = BASE / "external"
RESULTS = BASE / "results"
DOCS = BASE / "docs"

GENE_STAGE = EXTERNAL / "rna" / "gene_stage_matrix.tsv"
MOTIF_ENRICHMENT = EXTERNAL / "motif" / "module_motif_enrichment.tsv"
MODULE_BASIS = RESULTS / "CSB_TRO_missing_control_term_module_basis.tsv"

OUT_TF_DELTA = RESULTS / "CSB_TRO_GSE36552_TF_expression_delta.tsv"
OUT_CANDIDATE_TF = RESULTS / "CSB_TRO_candidate_embryo_TF_expression_delta.tsv"
OUT_TEMPLATE = EXTERNAL / "motif" / "module_motif_enrichment_template.tsv"
OUT_ACTIVITY = RESULTS / "CSB_TRO_residual_module_motif_TF_activity.tsv"
OUT_FEATURES = RESULTS / "CSB_TRO_motif_TF_activity_control_features.tsv"
OUT_DOC = DOCS / "CSB_TRO_motif_TF_activity_interpretation.md"
OUT_MANIFEST = RESULTS / "CSB_TRO_motif_TF_activity_manifest.json"

PRIORITY_MODULES = ["M05", "M01", "M12", "M02", "M10"]
WINDOWS = [
    ("4-cell", "8-cell", "delta_4cell_to_8cell"),
    ("8-cell", "morula", "delta_8cell_to_morula"),
    ("morula", "blastocyst", "delta_morula_to_blastocyst"),
]

EMBRYO_TF_CANDIDATES = [
    "DUX4",
    "DUXA",
    "TPRX1",
    "TPRX2",
    "LEUTX",
    "ZSCAN4",
    "ZSCAN4C",
    "ZSCAN4D",
    "KLF17",
    "TFAP2C",
    "POU5F1",
    "SOX2",
    "NANOG",
    "TEAD4",
    "GATA3",
    "CDX2",
    "KLF4",
    "KLF5",
    "DPPA2",
    "DPPA4",
    "OTX2",
    "PRDM14",
]


def zscore(series: pd.Series) -> pd.Series:
    x = pd.to_numeric(series, errors="coerce").astype(float)
    sd = x.std()
    if not np.isfinite(sd) or sd == 0:
        return x * 0.0
    return (x - x.mean()) / sd


def log_expr(x: pd.Series) -> pd.Series:
    return np.log2(pd.to_numeric(x, errors="coerce").fillna(0.0) + 1.0)


def compute_tf_delta(matrix: pd.DataFrame) -> pd.DataFrame:
    rows = []
    for row in matrix.to_dict(orient="records"):
        rec = {
            "TF": str(row.get("gene_name", "")),
            "gene_id": str(row.get("gene_id", "")),
            "dataset": "GSE36552",
        }
        for stage in ["4-cell", "8-cell", "morula", "blastocyst"]:
            if stage in matrix.columns:
                value = float(row.get(stage, np.nan))
                rec[f"expr_{stage}"] = value
                rec[f"log_expr_{stage}"] = float(np.log2(max(value, 0.0) + 1.0))
                rec[f"detected_{stage}"] = bool(value > 0)
        for start, end, name in WINDOWS:
            if start in row and end in row:
                rec[name] = float(np.log2(float(row[end]) + 1.0) - np.log2(float(row[start]) + 1.0))
        rows.append(rec)
    out = pd.DataFrame(rows)
    for _, _, name in WINDOWS:
        if name in out.columns:
            out[name + "_zscore"] = zscore(out[name])
    return out


def read_motif_enrichment(path: Path) -> pd.DataFrame:
    if not path.exists():
        return pd.DataFrame()
    tab = pd.read_csv(path, sep="\t")
    rename = {}
    for col in tab.columns:
        lower = col.lower()
        if lower in {"tf", "target", "transcription_factor", "motif_name"}:
            rename[col] = "TF"
        elif lower in {"module", "module_id"}:
            rename[col] = "module_id"
        elif lower in {"log_or", "log_odds", "log_odds_ratio", "log2_or"}:
            rename[col] = "log_odds_ratio"
        elif lower in {"q", "qvalue", "q_value", "fdr"}:
            rename[col] = "qvalue"
        elif lower in {"p", "pvalue", "p_value"}:
            rename[col] = "pvalue"
    tab = tab.rename(columns=rename)
    tab = tab.loc[:, ~tab.columns.duplicated()].copy()
    required = {"module_id", "TF"}
    if not required.issubset(tab.columns):
        raise ValueError("Motif enrichment table must contain module_id and TF columns.")
    if "log_odds_ratio" not in tab.columns:
        tab["log_odds_ratio"] = 1.0
    if "qvalue" not in tab.columns:
        tab["qvalue"] = tab["pvalue"] if "pvalue" in tab.columns else 0.05
    tab["TF"] = tab["TF"].astype(str).str.upper()
    return tab


def motif_score(tab: pd.DataFrame) -> pd.Series:
    log_or = pd.to_numeric(tab["log_odds_ratio"], errors="coerce").fillna(0.0)
    q = pd.to_numeric(tab["qvalue"], errors="coerce").fillna(1.0).clip(lower=1e-300, upper=1.0)
    signed_logq = np.sign(log_or) * (-np.log10(q))
    fallback = log_or
    return np.where(np.abs(signed_logq) > 0, signed_logq, fallback)


def build_activity(motif: pd.DataFrame, tf_delta: pd.DataFrame, window_col: str) -> tuple[pd.DataFrame, pd.DataFrame]:
    tf = tf_delta.copy()
    tf["TF"] = tf["TF"].astype(str).str.upper()
    if window_col + "_zscore" not in tf.columns:
        raise ValueError(f"Missing TF delta column: {window_col}_zscore")
    motif = motif[motif["module_id"].isin(PRIORITY_MODULES)].copy()
    motif["motif_score_signed_logq"] = motif_score(motif)
    motif["motif_score_mode"] = np.where(pd.to_numeric(motif["qvalue"], errors="coerce").fillna(1.0) < 1.0, "signed_logq", "log_odds_ratio_fallback")
    joined = motif.merge(tf[["TF", window_col, window_col + "_zscore"]], on="TF", how="inner")
    joined["tf_activity"] = joined["motif_score_signed_logq"] * pd.to_numeric(joined[window_col + "_zscore"], errors="coerce")
    module = (
        joined.groupby("module_id", as_index=False)
        .agg(
            control_value=("tf_activity", "sum"),
            n_TF=("TF", "nunique"),
            top_TFs=("TF", lambda x: ",".join(list(dict.fromkeys(map(str, x)))[:30])),
        )
    )
    module["control_value_z"] = zscore(module["control_value"])
    basis = pd.read_csv(MODULE_BASIS, sep="\t")
    module = module.merge(basis[["module_id", "n_DMRs", "latent_control_PC1", "latent_control_PC2", "latent_control_PC3", "latent_control_norm", "ridge_weight"]], on="module_id", how="left")
    module["candidate_control"] = "motif_TF_activity_" + window_col + "_" + module["module_id"].astype(str)
    module["control_modality"] = "motif_activity"
    module["control_stage_window"] = window_col.replace("delta_", "")
    module["leakage_status"] = "methylation_non_leaking_motif_x_TF_expression"
    module["interpretation"] = "Module-specific motif enrichment multiplied by TF expression change from GSE36552."
    for pc in ["PC1", "PC2", "PC3"]:
        module[f"candidate_control_direction_{pc}"] = module["control_value_z"] * pd.to_numeric(module[f"latent_control_{pc}"], errors="coerce")
    return joined, module


def write_missing_template() -> None:
    rows = []
    for module_id in PRIORITY_MODULES:
        for tf in EMBRYO_TF_CANDIDATES:
            rows.append(
                {
                    "module_id": module_id,
                    "TF": tf,
                    "motif_database": "JASPAR_or_HOCOMOCO_or_CISBP",
                    "log_odds_ratio": np.nan,
                    "pvalue": np.nan,
                    "qvalue": np.nan,
                    "motif_hit_count": np.nan,
                    "background_hit_count": np.nan,
                    "note": "Fill from HOMER/FIMO/module-specific motif enrichment against matched background.",
                }
            )
    pd.DataFrame(rows).to_csv(OUT_TEMPLATE, sep="\t", index=False)


def main() -> None:
    RESULTS.mkdir(parents=True, exist_ok=True)
    DOCS.mkdir(parents=True, exist_ok=True)
    EXTERNAL.joinpath("motif").mkdir(parents=True, exist_ok=True)

    matrix = pd.read_csv(GENE_STAGE, sep="\t")
    tf_delta = compute_tf_delta(matrix)
    tf_delta.to_csv(OUT_TF_DELTA, sep="\t", index=False)
    tf_delta[tf_delta["TF"].str.upper().isin(EMBRYO_TF_CANDIDATES)].to_csv(OUT_CANDIDATE_TF, sep="\t", index=False)

    motif = read_motif_enrichment(MOTIF_ENRICHMENT)
    if motif.empty:
        write_missing_template()
        pd.DataFrame(
            [
                {
                    "analysis_status": "not_run_missing_motif_input",
                    "reason": "No module-specific motif enrichment table was found.",
                    "required_input": str(MOTIF_ENRICHMENT),
                    "template": str(OUT_TEMPLATE),
                }
            ]
        ).to_csv(OUT_ACTIVITY, sep="\t", index=False)
        pd.DataFrame(
            columns=[
                "module_id",
                "candidate_control",
                "control_modality",
                "control_value",
                "control_stage_window",
                "leakage_status",
                "interpretation",
                "control_value_z",
                "latent_control_PC1",
                "latent_control_PC2",
                "latent_control_PC3",
            ]
        ).to_csv(OUT_FEATURES, sep="\t", index=False)
        status = "not_run_missing_motif_input"
        n_activity = 0
    else:
        joined, features = build_activity(motif, tf_delta, "delta_8cell_to_morula")
        joined.to_csv(OUT_ACTIVITY, sep="\t", index=False)
        features.to_csv(OUT_FEATURES, sep="\t", index=False)
        status = "completed"
        n_activity = int(len(joined))

    lines = [
        "# Motif x TF expression activity",
        "",
        f"Status: {status}",
        "",
        "TF expression deltas were computed from GSE36552 gene-stage RPKM using log2(expression + 1).",
        "",
        "Expected motif enrichment input:",
        f"- {MOTIF_ENRICHMENT}",
        "",
        "Required columns: module_id, TF, log_odds_ratio, qvalue. Optional: pvalue, motif_hit_count, background_hit_count, motif_database.",
    ]
    if status != "completed":
        lines.extend(["", f"A fillable motif enrichment template was written to: {OUT_TEMPLATE}"])
    OUT_DOC.write_text("\n".join(lines) + "\n", encoding="utf-8")

    OUT_MANIFEST.write_text(
        json.dumps(
            {
                "gene_stage_matrix": str(GENE_STAGE),
                "motif_enrichment": str(MOTIF_ENRICHMENT) if MOTIF_ENRICHMENT.exists() else None,
                "status": status,
                "n_TF_delta_rows": int(len(tf_delta)),
                "n_activity_rows": n_activity,
                "outputs": [str(OUT_TF_DELTA), str(OUT_CANDIDATE_TF), str(OUT_ACTIVITY), str(OUT_FEATURES), str(OUT_DOC)],
            },
            indent=2,
        )
        + "\n",
        encoding="utf-8",
    )
    print(json.dumps({"status": status, "n_TF_delta_rows": int(len(tf_delta)), "n_activity_rows": n_activity}, indent=2))


if __name__ == "__main__":
    main()
```
