## Supplemental code for "An entropy-based diagnostic framework for characterizing methylation state dynamics during preimplantation development": run_rna_transition_control_amplitude_scan.py

```
from __future__ import annotations

import json
import sys
from pathlib import Path

import numpy as np
import pandas as pd


BASE = Path(r"E:\CSB_TRO_operator_time_DMR_dynamics_2026-05-25")
CODE = BASE / "code"
RESULTS = BASE / "results"
DOCS = BASE / "docs"
sys.path.insert(0, str(CODE))

from run_basin_residual_control_field import PRE_MORULA_STAGES, simulate_strict_pre_morula  # noqa: E402
from run_biological_control_augmented_dynamics import build_control_vector, evaluate_control  # noqa: E402
from run_morula_basin_sde import basin_definition, fit_latent_basis, fit_operator, load_inputs, stage_ids  # noqa: E402


FEATURE_TABLE = RESULTS / "CSB_TRO_RNA_transition_control_features.tsv"
OUT_METRICS = RESULTS / "CSB_TRO_RNA_transition_control_amplitude_scan.tsv"
OUT_DOC = DOCS / "CSB_TRO_RNA_transition_control_amplitude_scan.md"
OUT_MANIFEST = RESULTS / "CSB_TRO_RNA_transition_control_amplitude_scan_manifest.json"


def main() -> None:
    scales = [0.0, 0.25, 0.5, 1.0, 1.5, 2.0, 3.0, 4.0]
    seed = 20260525

    features = pd.read_csv(FEATURE_TABLE, sep="\t")
    matrix, ann, pairs = load_inputs()
    mu, sd, components, score_df = fit_latent_basis(matrix, 3)
    coef, _, _ = fit_operator(score_df, pairs, train_stages=set(PRE_MORULA_STAGES), lam=1000.0)
    strict_pred_z = simulate_strict_pre_morula(score_df, ann, coef, n_steps=12)
    obs_z = score_df.loc[stage_ids(ann, "morula")].to_numpy(dtype=float)
    obs_dmr = matrix.loc[stage_ids(ann, "morula")].to_numpy(dtype=float)
    basin = basin_definition(obs_z)
    residual_z = obs_z.mean(axis=0) - strict_pred_z.mean(axis=0)

    rows = []
    rng = np.random.default_rng(seed)
    for scale in scales:
        sub = features.copy()
        sub["control_value_z"] = pd.to_numeric(sub["control_value_z"], errors="coerce").fillna(0.0) * scale
        control, _ = build_control_vector(sub, "unit", residual_z, 1e-3, rng)
        row = evaluate_control(
            f"RNA_transition_unit_beta_scale_{scale:g}",
            strict_pred_z + control[None, :],
            obs_z,
            obs_dmr,
            mu,
            sd,
            components,
            basin,
            residual_z,
            {
                "control_vector": control,
                "validation_status": "external_feature_fixed_amplitude_scan",
                "n_features": int(len(sub)),
                "beta_mode": "unit_scaled",
                "feature_modality": "RNA",
                "amplitude_scale": scale,
            },
            seed + int(scale * 100),
        )
        rows.append(row)

        flip_row = evaluate_control(
            f"RNA_transition_unit_beta_scale_{scale:g}_sign_flip",
            strict_pred_z - control[None, :],
            obs_z,
            obs_dmr,
            mu,
            sd,
            components,
            basin,
            residual_z,
            {
                "control_vector": -control,
                "validation_status": "sign_flip_control",
                "n_features": int(len(sub)),
                "beta_mode": "unit_scaled_sign_flip",
                "feature_modality": "RNA",
                "amplitude_scale": scale,
            },
            seed + 1000 + int(scale * 100),
        )
        rows.append(flip_row)

    metrics = pd.DataFrame(rows)
    metrics.to_csv(OUT_METRICS, sep="\t", index=False)

    forward = metrics[~metrics["control_model"].str.endswith("_sign_flip")].copy()
    lines = [
        "# RNA transition control amplitude scan",
        "",
        "This scan multiplies the same GSE36552 RNA transition gate by fixed amplitudes. It does not fit beta to the measured morula methylation residual.",
        "",
        "Forward fixed-amplitude results:",
    ]
    for row in forward.itertuples(index=False):
        lines.append(
            f"- scale={row.amplitude_scale:g}: occupancy_q90={row.pred_basin_occupancy_q90:.3f}, "
            f"cosine={row.direction_cosine_to_measured_correction:.3f}, PC3_recovery={row.PC3_negative_pull_recovered:.3f}"
        )
    lines.extend(
        [
            "",
            "Interpretation: increasing rescue under fixed scaling supports direction compatibility, but the scale itself is not independently calibrated by external biology yet.",
        ]
    )
    OUT_DOC.write_text("\n".join(lines) + "\n", encoding="utf-8")
    OUT_MANIFEST.write_text(
        json.dumps(
            {
                "feature_table": str(FEATURE_TABLE),
                "scales": scales,
                "outputs": [str(OUT_METRICS), str(OUT_DOC)],
            },
            indent=2,
        )
        + "\n",
        encoding="utf-8",
    )
    print(json.dumps(forward[["amplitude_scale", "pred_basin_occupancy_q90", "direction_cosine_to_measured_correction", "PC3_negative_pull_recovered"]].to_dict(orient="records"), indent=2))


if __name__ == "__main__":
    main()
```
