## Supplemental code for "An entropy-based diagnostic framework for characterizing methylation state dynamics during preimplantation development": run_rna_transition_control_features.py

```
from __future__ import annotations

import json
from pathlib import Path

import numpy as np
import pandas as pd


BASE = Path(r"E:\CSB_TRO_operator_time_DMR_dynamics_2026-05-25")
MAIN = Path(r"C:\Users\18068\Desktop\CSB_TRO_Project_2026-05-24")
RESULTS = BASE / "results"
DOCS = BASE / "docs"

RNA_BRIDGES = MAIN / "results" / "CSB_TRO_RNA_GSE36552_transition_bridges.tsv"
MODULE_BASIS = RESULTS / "CSB_TRO_missing_control_term_module_basis.tsv"
OUT_FEATURES = RESULTS / "CSB_TRO_RNA_transition_control_features.tsv"
OUT_COMPONENTS = RESULTS / "CSB_TRO_RNA_transition_control_components.tsv"
OUT_DOC = DOCS / "CSB_TRO_RNA_transition_control_interpretation.md"
OUT_MANIFEST = RESULTS / "CSB_TRO_RNA_transition_control_manifest.json"

PRIORITY_MODULES = ["M05", "M01", "M12", "M02", "M10"]


def normalized_target_value(tab: pd.DataFrame, target: pd.Series, col: str, sign: float = 1.0) -> float:
    vals = pd.to_numeric(tab[col], errors="coerce").astype(float)
    denom = float(vals.abs().max())
    if not np.isfinite(denom) or denom == 0:
        return 0.0
    raw = float(target[col]) * sign
    return max(0.0, raw / denom)


def main() -> None:
    RESULTS.mkdir(parents=True, exist_ok=True)
    DOCS.mkdir(parents=True, exist_ok=True)

    bridges = pd.read_csv(RNA_BRIDGES, sep="\t")
    basis = pd.read_csv(MODULE_BASIS, sep="\t")
    target_rows = bridges[(bridges["from_stage"] == "8-cell") & (bridges["to_stage"] == "morula")]
    if target_rows.empty:
        raise ValueError("No GSE36552 8-cell -> morula RNA transition row was found.")
    target = target_rows.iloc[0]

    components = [
        {
            "component": "RNA_potency_loss_8cell_to_morula",
            "raw_value": float(-target["mean_transport_P"]),
            "normalized_positive_activity": normalized_target_value(bridges, target, "mean_transport_P", sign=-1.0),
            "interpretation": "Positive when RNA potency transport decreases from 8-cell to morula.",
        },
        {
            "component": "RNA_entropy_loss_8cell_to_morula",
            "raw_value": float(-target["mean_transport_Hr"]),
            "normalized_positive_activity": normalized_target_value(bridges, target, "mean_transport_Hr", sign=-1.0),
            "interpretation": "Positive when RNA entropy transport decreases from 8-cell to morula.",
        },
        {
            "component": "RNA_transition_displacement_8cell_to_morula",
            "raw_value": float(target["mean_squared_displacement"]),
            "normalized_positive_activity": normalized_target_value(bridges, target, "mean_squared_displacement", sign=1.0),
            "interpretation": "Normalized RNA state displacement for the 8-cell to morula transition.",
        },
        {
            "component": "RNA_potency_penalty_8cell_to_morula",
            "raw_value": float(target["potency_below_threshold_penalty"]),
            "normalized_positive_activity": normalized_target_value(bridges, target, "potency_below_threshold_penalty", sign=1.0),
            "interpretation": "Normalized penalty for falling below the RNA potency threshold.",
        },
    ]
    comp = pd.DataFrame(components)
    activity = float(comp["normalized_positive_activity"].mean())

    rows = []
    priority_basis = basis[basis["module_id"].isin(PRIORITY_MODULES)].copy()
    for row in priority_basis.itertuples(index=False):
        rows.append(
            {
                "module_id": row.module_id,
                "candidate_control": f"RNA_transition_activity_8cell_to_morula_{row.module_id}",
                "control_modality": "RNA",
                "control_value": activity,
                "control_stage_window": "8-cell_to_morula",
                "leakage_status": "methylation_non_leaking_external_RNA_transition_no_gene_link",
                "interpretation": "GSE36552 stage-level RNA transition activity gates the pre-specified residual module direction; not gene-linked yet.",
                "control_value_z": activity,
                "n_DMRs": int(row.n_DMRs),
                "latent_control_PC1": float(row.latent_control_PC1),
                "latent_control_PC2": float(row.latent_control_PC2),
                "latent_control_PC3": float(row.latent_control_PC3),
                "latent_control_norm": float(row.latent_control_norm),
                "ridge_weight": np.nan,
                "candidate_control_direction_PC1": float(activity * row.latent_control_PC1),
                "candidate_control_direction_PC2": float(activity * row.latent_control_PC2),
                "candidate_control_direction_PC3": float(activity * row.latent_control_PC3),
            }
        )

    features = pd.DataFrame(rows)
    features.to_csv(OUT_FEATURES, sep="\t", index=False)
    comp.to_csv(OUT_COMPONENTS, sep="\t", index=False)

    lines = [
        "# RNA transition control features",
        "",
        "This is a first external-RNA control experiment using GSE36552 stage-level transition summaries.",
        "",
        "It does not use morula methylation to define the RNA activity, beta, center, radius, or occupancy. It does use the previously nominated residual module directions M05/M01/M12/M02/M10, so this is a module-gated RNA transition test rather than a gene-linked u_bio mechanism.",
        "",
        f"Composite normalized RNA transition activity: {activity:.6f}",
        "",
        "Components:",
    ]
    for c in comp.itertuples(index=False):
        lines.append(f"- {c.component}: raw={c.raw_value:.6f}, normalized={c.normalized_positive_activity:.6f}")
    lines.extend(
        [
            "",
            "Next stronger test requires gene-linked RNA, TF/motif, ATAC, or histone features per module.",
        ]
    )
    OUT_DOC.write_text("\n".join(lines) + "\n", encoding="utf-8")

    OUT_MANIFEST.write_text(
        json.dumps(
            {
                "rna_transition_input": str(RNA_BRIDGES),
                "module_basis_input": str(MODULE_BASIS),
                "priority_modules": PRIORITY_MODULES,
                "composite_activity": activity,
                "outputs": [str(OUT_FEATURES), str(OUT_COMPONENTS), str(OUT_DOC)],
            },
            indent=2,
        )
        + "\n",
        encoding="utf-8",
    )
    print(f"Wrote {OUT_FEATURES}")
    print(f"Composite RNA transition activity = {activity:.6f}")


if __name__ == "__main__":
    main()
```
