## Supplemental code for "An entropy-based diagnostic framework for characterizing methylation state dynamics during preimplantation development": run_rna_transition_replication_features.py

```
from __future__ import annotations

import json
from pathlib import Path

import numpy as np
import pandas as pd


BASE = Path(r"E:\CSB_TRO_operator_time_DMR_dynamics_2026-05-25")
MAIN = Path(r"C:\Users\18068\Desktop\CSB_TRO_Project_2026-05-24")
RESULTS = BASE / "results"
DOCS = BASE / "docs"

MODULE_BASIS = RESULTS / "CSB_TRO_missing_control_term_module_basis.tsv"
PRIORITY_MODULES = ["M05", "M01", "M12", "M02", "M10"]

DATASETS = [
    {
        "label": "GSE36552",
        "path": MAIN / "results" / "CSB_TRO_RNA_GSE36552_transition_bridges.tsv",
    },
    {
        "label": "GSE44183_external",
        "path": MAIN / "results" / "CSB_TRO_RNA_GSE44183_external_transition_bridges.tsv",
    },
]


def normalized_target_value(tab: pd.DataFrame, target: pd.Series, col: str, sign: float = 1.0) -> float:
    vals = pd.to_numeric(tab[col], errors="coerce").astype(float)
    denom = float(vals.abs().max())
    if not np.isfinite(denom) or denom == 0:
        return 0.0
    raw = float(target[col]) * sign
    return max(0.0, raw / denom)


def dataset_activity(label: str, path: Path) -> tuple[float, pd.DataFrame]:
    tab = pd.read_csv(path, sep="\t")
    target_rows = tab[(tab["from_stage"] == "8-cell") & (tab["to_stage"] == "morula")]
    if target_rows.empty:
        raise ValueError(f"No 8-cell -> morula transition row for {label}: {path}")
    target = target_rows.iloc[0]
    comp = pd.DataFrame(
        [
            {
                "dataset": label,
                "component": "RNA_potency_loss_8cell_to_morula",
                "raw_value": float(-target["mean_transport_P"]),
                "normalized_positive_activity": normalized_target_value(tab, target, "mean_transport_P", sign=-1.0),
            },
            {
                "dataset": label,
                "component": "RNA_entropy_loss_8cell_to_morula",
                "raw_value": float(-target["mean_transport_Hr"]),
                "normalized_positive_activity": normalized_target_value(tab, target, "mean_transport_Hr", sign=-1.0),
            },
            {
                "dataset": label,
                "component": "RNA_transition_displacement_8cell_to_morula",
                "raw_value": float(target["mean_squared_displacement"]),
                "normalized_positive_activity": normalized_target_value(tab, target, "mean_squared_displacement", sign=1.0),
            },
            {
                "dataset": label,
                "component": "RNA_potency_penalty_8cell_to_morula",
                "raw_value": float(target["potency_below_threshold_penalty"]),
                "normalized_positive_activity": normalized_target_value(tab, target, "potency_below_threshold_penalty", sign=1.0),
            },
        ]
    )
    return float(comp["normalized_positive_activity"].mean()), comp


def make_features(label: str, activity: float, basis: pd.DataFrame) -> pd.DataFrame:
    rows = []
    for row in basis[basis["module_id"].isin(PRIORITY_MODULES)].itertuples(index=False):
        rows.append(
            {
                "module_id": row.module_id,
                "candidate_control": f"RNA_transition_activity_{label}_8cell_to_morula_{row.module_id}",
                "control_modality": "RNA",
                "control_value": activity,
                "control_stage_window": "8-cell_to_morula",
                "leakage_status": "methylation_non_leaking_external_RNA_transition_no_gene_link",
                "interpretation": f"{label} stage-level RNA transition activity gates the pre-specified residual module direction; not gene-linked yet.",
                "control_value_z": activity,
                "n_DMRs": int(row.n_DMRs),
                "latent_control_PC1": float(row.latent_control_PC1),
                "latent_control_PC2": float(row.latent_control_PC2),
                "latent_control_PC3": float(row.latent_control_PC3),
                "latent_control_norm": float(row.latent_control_norm),
                "ridge_weight": np.nan,
                "candidate_control_direction_PC1": float(activity * row.latent_control_PC1),
                "candidate_control_direction_PC2": float(activity * row.latent_control_PC2),
                "candidate_control_direction_PC3": float(activity * row.latent_control_PC3),
                "rna_dataset": label,
            }
        )
    return pd.DataFrame(rows)


def main() -> None:
    RESULTS.mkdir(parents=True, exist_ok=True)
    DOCS.mkdir(parents=True, exist_ok=True)
    basis = pd.read_csv(MODULE_BASIS, sep="\t")

    activity_rows = []
    component_parts = []
    feature_tables = {}
    for item in DATASETS:
        activity, comp = dataset_activity(item["label"], item["path"])
        activity_rows.append({"dataset": item["label"], "composite_activity": activity, "transition_table": str(item["path"])})
        component_parts.append(comp)
        feat = make_features(item["label"], activity, basis)
        out = RESULTS / f"CSB_TRO_RNA_transition_{item['label']}_control_features.tsv"
        feat.to_csv(out, sep="\t", index=False)
        feature_tables[item["label"]] = str(out)

    activities = pd.DataFrame(activity_rows)
    components = pd.concat(component_parts, ignore_index=True)
    consensus_activity = float(activities["composite_activity"].mean())
    consensus = make_features("consensus_GSE36552_GSE44183_external", consensus_activity, basis)
    consensus_out = RESULTS / "CSB_TRO_RNA_transition_consensus_control_features.tsv"
    consensus.to_csv(consensus_out, sep="\t", index=False)
    feature_tables["consensus"] = str(consensus_out)

    activities.to_csv(RESULTS / "CSB_TRO_RNA_transition_replication_activities.tsv", sep="\t", index=False)
    components.to_csv(RESULTS / "CSB_TRO_RNA_transition_replication_components.tsv", sep="\t", index=False)

    lines = [
        "# RNA transition replication features",
        "",
        "This builds RNA transition gates from two stage-level RNA datasets and a consensus average.",
        "",
        "Composite activities:",
    ]
    for row in activities.itertuples(index=False):
        lines.append(f"- {row.dataset}: {row.composite_activity:.6f}")
    lines.append(f"- consensus: {consensus_activity:.6f}")
    lines.extend(
        [
            "",
            "These are methylation-non-leaking external RNA transition features, but still not gene-linked or TF/motif-linked.",
        ]
    )
    (DOCS / "CSB_TRO_RNA_transition_replication_features.md").write_text("\n".join(lines) + "\n", encoding="utf-8")
    (RESULTS / "CSB_TRO_RNA_transition_replication_manifest.json").write_text(
        json.dumps(
            {
                "datasets": DATASETS,
                "priority_modules": PRIORITY_MODULES,
                "activities": activity_rows,
                "consensus_activity": consensus_activity,
                "feature_tables": feature_tables,
            },
            indent=2,
            default=str,
        )
        + "\n",
        encoding="utf-8",
    )
    print(json.dumps({"activities": activity_rows, "consensus_activity": consensus_activity, "feature_tables": feature_tables}, indent=2))


if __name__ == "__main__":
    main()
```
