## Supplemental code for "An entropy-based diagnostic framework for characterizing methylation state dynamics during preimplantation development": 5_28_progress__comprehensive_ubio_analysis.py

```
#!/usr/bin/env python
"""
Comprehensive u_bio analysis using all available quantitative signals.

Available signals:
1. Liu2019: 4-stage accessibility (2-cell, 4-cell, 8-cell, morula) - BEST
2. H3K27ac_8cell peaks (score available)
3. H3K27ac_blastocyst peaks (score available)
4. H3K27me3_8cell peaks (score available)
5. H3K4me3_8cell peaks (10-column narrowPeak with signal)

Key innovation:
- Use delta_acc = acc_morula - acc_8cell as the primary u_bio
- Build multi-signal regression to identify the strongest u_bio combination
- Test whether any combination achieves significant global RMSE improvement
- Use DMR-level c_diag analysis stratified by signal strength
"""
from __future__ import annotations
import json
from pathlib import Path
import numpy as np
import pandas as pd
from scipy import stats
from numpy.linalg import solve

OUT = Path("E:/5_28_progress")
H3K27AC_8CELL = Path(
    "E:/CSB_TRO_operator_time_DMR_dynamics_2026-05-25/external/histone/H3K27ac_8cell.hg19.bed.gz"
)
H3K27AC_BLAST = Path(
    "E:/CSB_TRO_operator_time_DMR_dynamics_2026-05-25/external/histone/H3K27ac_blastocyst.hg19.bed.gz"
)
H3K27ME3_8CELL = Path(
    "E:/CSB_TRO_operator_time_DMR_dynamics_2026-05-25/external/histone/H3K27me3_8cell.hg19.bed.gz"
)
H3K4ME3_8CELL = Path(
    "E:/CSB_TRO_operator_time_DMR_dynamics_2026-05-25/external/histone/H3K4me3_8cell.hg19.bed.gz"
)
H3K27ME3_BLAST = Path(
    "E:/CSB_TRO_operator_time_DMR_dynamics_2026-05-25/external/histone/H3K27me3_blastocyst.hg19.bed.gz"
)
H3K4ME3_BLAST = Path(
    "E:/CSB_TRO_operator_time_DMR_dynamics_2026-05-25/external/histone/H3K4me3_blastocyst.hg19.bed.gz"
)

DMR_META = Path(
    "E:/CSB_TRO_operator_time_DMR_dynamics_2026-05-25/results/CSB_TRO_DMR_metadata.tsv"
)
DMR_RESIDUAL = Path(
    "E:/CSB_TRO_operator_time_DMR_dynamics_2026-05-25/results/CSB_TRO_basin_residual_DMR_ranking.tsv"
)
DMR_TRAJ = Path(
    "E:/CSB_TRO_operator_time_DMR_dynamics_2026-05-25/results/CSB_TRO_DMR_stage_mean_trajectory.tsv"
)
CURV = Path("E:/实验进展5_27/CSB_TRO_2026-05-27_entry_exit_curvature.tsv")
MULTISTAGE_ACC = OUT / "CSB_TRO_5_28_dmr_multistage_accessibility.tsv"

SEED = 42
N_BOOT = 2000


def spearman(x, y):
    mask = np.isfinite(x) & np.isfinite(y)
    if mask.sum() < 5:
        return np.nan, np.nan
    r, p = stats.spearmanr(x[mask], y[mask])
    return float(r), float(p)


def perm_test(x, y, n=N_BOOT, seed=SEED):
    rng = np.random.default_rng(seed)
    obs_rho, _ = spearman(x, y)
    if not np.isfinite(obs_rho):
        return np.nan, np.nan
    mask = np.isfinite(x) & np.isfinite(y)
    xv, yv = x[mask], y[mask]
    nulls = [stats.spearmanr(rng.permutation(xv), yv)[0] for _ in range(n)]
    nulls = np.array(nulls)
    return float((nulls >= obs_rho).mean()), float(np.quantile(nulls, 0.95))


def overlap_peaks(peaks_df, dmr_meta, score_col=4):
    """For each DMR, get overlap flag and max peak score."""
    peaks_df["chr"] = peaks_df["chr"].astype(str).str.strip()
    dmr_meta["chr"] = dmr_meta["chr"].astype(str).str.strip()
    by_chr = {c: g for c, g in peaks_df.groupby("chr")}
    rows = []
    for _, dmr in dmr_meta.iterrows():
        chrom, ds, de = dmr["chr"], int(dmr["start"]), int(dmr["end"])
        cluster = dmr["cluster_name"]
        if chrom not in by_chr:
            rows.append({"cluster_name": cluster, "overlap": 0, "score_max": np.nan})
            continue
        sub = by_chr[chrom]
        mask = (sub["start"] < de) & (sub["end"] > ds)
        hits = sub[mask]
        sc = pd.to_numeric(hits.iloc[:, score_col - 1], errors="coerce").dropna() if len(hits) > 0 else pd.Series(dtype=float)
        rows.append({
            "cluster_name": cluster,
            "overlap": int(len(hits) > 0),
            "score_max": float(sc.max()) if len(sc) > 0 else np.nan,
        })
    return pd.DataFrame(rows)


def loocv_rmse(y, X, lam=0.01):
    n = len(y)
    reg = np.diag([lam] * X.shape[1])
    reg[-1, -1] = 0
    errs = []
    for i in range(n):
        tr = [j for j in range(n) if j != i]
        try:
            w = solve(X[tr].T @ X[tr] + reg, X[tr].T @ y[tr])
            errs.append((y[i] - X[i] @ w) ** 2)
        except Exception:
            continue
    return float(np.sqrt(np.mean(errs)))


def bootstrap_improvement(y, X_base, X_bio, lam=0.01, n=N_BOOT, seed=SEED):
    """Permutation test: is improvement > null?"""
    rng = np.random.default_rng(seed)
    obs_base = loocv_rmse(y, X_base, lam)
    obs_bio = loocv_rmse(y, X_bio, lam)
    obs_impr = (obs_base - obs_bio) / obs_base * 100

    # Permute the extra columns in X_bio (not the base columns)
    n_base = X_base.shape[1]
    null_imprs = []
    for _ in range(n):
        X_perm = X_bio.copy()
        for col in range(n_base - 1, X_bio.shape[1] - 1):  # extra u_bio cols
            X_perm[:, col] = rng.permutation(X_bio[:, col])
        rb = loocv_rmse(y, X_perm, lam)
        null_imprs.append((obs_base - rb) / obs_base * 100)

    null_q95 = float(np.quantile(null_imprs, 0.95))
    pp = float((np.array(null_imprs) >= obs_impr).mean())
    return obs_impr, obs_base, obs_bio, null_q95, pp


def standardize(x):
    v = x.copy().astype(float)
    mask = np.isfinite(v)
    if mask.sum() < 2:
        return np.zeros_like(v)
    v[~mask] = 0.0
    v[mask] = (v[mask] - v[mask].mean()) / (v[mask].std() + 1e-8)
    return v


def main():
    print("=" * 70)
    print("Comprehensive u_bio analysis — all available signals")
    print("=" * 70)

    # ── Load core data ─────────────────────────────────────────────────────────
    dmr_meta = pd.read_csv(DMR_META, sep="\t")
    residual = pd.read_csv(DMR_RESIDUAL, sep="\t")
    traj = pd.read_csv(DMR_TRAJ, sep="\t")
    curv = pd.read_csv(CURV, sep="\t")
    multistage = pd.read_csv(MULTISTAGE_ACC, sep="\t")

    stage_means = {}
    for stage, g in traj.groupby("stage"):
        stage_means[stage] = g.set_index("cluster_name")["mean_beta"].to_dict()

    clusters = residual["cluster_name"].tolist()

    # ── Load all histone peak files ────────────────────────────────────────────
    def load_peaks(path, ncols=9):
        cols = ["chr", "start", "end", "name", "score",
                "strand", "fc", "neglog10p", "neglog10q"][:ncols]
        try:
            df = pd.read_csv(path, sep="\t", header=None, compression="gzip",
                             names=cols, usecols=list(range(ncols)))
        except Exception:
            df = pd.read_csv(path, sep="\t", header=None, compression="gzip",
                             usecols=list(range(min(ncols, 5))),
                             names=["chr", "start", "end", "name", "score"])
        return df

    print("Loading histone peaks...")
    p_k27ac_8 = load_peaks(H3K27AC_8CELL)
    p_k27ac_b = load_peaks(H3K27AC_BLAST)
    p_k27me3_8 = load_peaks(H3K27ME3_8CELL)
    p_k4me3_8 = load_peaks(H3K4ME3_8CELL, ncols=10)
    p_k27me3_b = load_peaks(H3K27ME3_BLAST)
    p_k4me3_b = load_peaks(H3K4ME3_BLAST, ncols=10)
    print(f"  k27ac_8cell={len(p_k27ac_8)}, k27ac_blast={len(p_k27ac_b)}")
    print(f"  k27me3_8cell={len(p_k27me3_8)}, k4me3_8cell={len(p_k4me3_8)}")
    print(f"  k27me3_blast={len(p_k27me3_b)}, k4me3_blast={len(p_k4me3_b)}")

    # ── Compute DMR-level histone signals ──────────────────────────────────────
    print("Computing DMR-level histone overlaps...")
    sig = {"cluster_name": clusters}

    for label, peaks in [
        ("k27ac_8cell", p_k27ac_8),
        ("k27ac_blast", p_k27ac_b),
        ("k27me3_8cell", p_k27me3_8),
        ("k4me3_8cell", p_k4me3_8),
        ("k27me3_blast", p_k27me3_b),
        ("k4me3_blast", p_k4me3_b),
    ]:
        ov = overlap_peaks(peaks, dmr_meta)
        ov = ov.set_index("cluster_name")
        sig[f"{label}_ov"] = [int(ov.loc[c, "overlap"]) if c in ov.index else 0 for c in clusters]
        sig[f"{label}_score"] = [ov.loc[c, "score_max"] if c in ov.index else np.nan for c in clusters]

    df = pd.DataFrame(sig)

    # Add accessibility signals
    acc_cols = ["cluster_name", "acc_morula_mean", "acc_8-cell_mean",
                "delta_acc_8cell_to_morula", "acc_2-cell_mean", "acc_4-cell_mean"]
    df = df.merge(multistage[[c for c in acc_cols if c in multistage.columns]],
                  on="cluster_name", how="left")

    # Add residual info
    df = df.merge(
        residual[["cluster_name", "basin_residual_rank",
                  "observed_minus_strict_pred_delta_beta",
                  "latent_residual_delta_beta", "module_id"]],
        on="cluster_name", how="left"
    )
    df = df.merge(curv[["cluster_name", "curvature", "is_inverted_u", "is_u_shape"]],
                  on="cluster_name", how="left")

    # Add methylation
    for st in ["2-cell", "4-cell", "8-cell", "morula", "blastocyst"]:
        safe = st.replace("-", "_")
        df[f"meth_{safe}"] = df["cluster_name"].map(
            lambda c: stage_means.get(st, {}).get(c, np.nan))

    df.to_csv(OUT / "CSB_TRO_5_28_comprehensive_ubio.tsv", sep="\t", index=False)

    # ── Signal coverage summary ────────────────────────────────────────────────
    print("\n--- Signal coverage (DMRs with signal) ---")
    for col in ["acc_morula_mean", "delta_acc_8cell_to_morula",
                "k27ac_8cell_ov", "k27ac_blast_ov", "k27me3_8cell_ov",
                "k4me3_8cell_ov", "k27me3_blast_ov", "k4me3_blast_ov"]:
        if col in df.columns:
            if "ov" in col:
                n = (df[col] > 0).sum()
            else:
                n = df[col].notna().sum()
            print(f"  {col}: {n}/156")

    # ── Correlation analysis: which signals explain c_diag? ───────────────────
    print("\n--- Correlation: signals vs strict_correction ---")
    y_corr = df["observed_minus_strict_pred_delta_beta"].values
    corr_results = {}
    for col in ["acc_morula_mean", "delta_acc_8cell_to_morula",
                "k27ac_8cell_score", "k27ac_blast_score",
                "k27me3_8cell_score", "k4me3_8cell_score",
                "k27me3_blast_score", "k4me3_blast_score"]:
        if col not in df.columns:
            continue
        x = df[col].values
        rho, p = spearman(x, y_corr)
        pp, q95 = perm_test(x, y_corr)
        sig_flag = "***" if rho > q95 else ("**" if p < 0.05 else ("*" if p < 0.1 else ""))
        print(f"  {col:<35}: rho={rho:+.4f}, p={p:.4f}, perm_p={pp:.4f} {sig_flag}")
        corr_results[col] = {"rho": float(rho), "p": float(p), "perm_p": float(pp), "null_q95": float(q95)}

    # ── LOO-CV prediction improvement ─────────────────────────────────────────
    print("\n--- LOO-CV: leave-morula-out prediction with u_bio ---")
    base_valid = df.dropna(subset=["meth_morula", "meth_8_cell"])
    y_base = base_valid["meth_morula"].values
    x_base = base_valid["meth_8_cell"].values
    X_meth = np.column_stack([x_base, np.ones(len(x_base))])
    rmse_base = loocv_rmse(y_base, X_meth)
    print(f"  Baseline (meth-only) RMSE: {rmse_base:.4f}")

    loocv_results = {}
    best_rmse = rmse_base
    best_label = "meth_only"

    for u_col in ["delta_acc_8cell_to_morula", "acc_morula_mean",
                  "k27ac_8cell_score", "k27ac_blast_score",
                  "k4me3_8cell_score", "k27me3_blast_score"]:
        if u_col not in df.columns:
            continue
        valid = base_valid.dropna(subset=[u_col])
        if len(valid) < 30:
            continue
        y_v = valid["meth_morula"].values
        x_v = valid["meth_8_cell"].values
        u_v = standardize(valid[u_col].values)
        X_bio = np.column_stack([x_v, u_v, np.ones(len(x_v))])
        X_m = np.column_stack([x_v, np.ones(len(x_v))])
        rmse_bio = loocv_rmse(y_v, X_bio)
        rmse_m = loocv_rmse(y_v, X_m)
        impr = (rmse_m - rmse_bio) / rmse_m * 100
        print(f"  + {u_col:<35}: RMSE={rmse_bio:.4f}, improvement={impr:+.2f}%")
        if rmse_bio < best_rmse:
            best_rmse = rmse_bio
            best_label = u_col
        loocv_results[u_col] = {"rmse": float(rmse_bio), "improvement_pct": float(impr)}

    # ── Best multi-signal combination ─────────────────────────────────────────
    print("\n--- Best multi-signal u_bio (delta_acc + k27ac_blast) ---")
    valid2 = base_valid.dropna(subset=["delta_acc_8cell_to_morula", "k27ac_blast_score"])
    if len(valid2) >= 20:
        y2 = valid2["meth_morula"].values
        x2 = valid2["meth_8_cell"].values
        u_da = standardize(valid2["delta_acc_8cell_to_morula"].values)
        u_k27 = standardize(valid2["k27ac_blast_score"].values)
        X_m2 = np.column_stack([x2, np.ones(len(x2))])
        X_combo = np.column_stack([x2, u_da, u_k27, np.ones(len(x2))])
        rmse_m2 = loocv_rmse(y2, X_m2)
        rmse_combo = loocv_rmse(y2, X_combo)
        impr_combo = (rmse_m2 - rmse_combo) / rmse_m2 * 100
        print(f"  n={len(valid2)}, baseline={rmse_m2:.4f}, combo={rmse_combo:.4f}, improvement={impr_combo:.2f}%")

        # Bootstrap significance test
        rng = np.random.default_rng(SEED)
        null_combo = []
        for _ in range(N_BOOT):
            X_perm = X_combo.copy()
            X_perm[:, 1] = rng.permutation(X_combo[:, 1])
            X_perm[:, 2] = rng.permutation(X_combo[:, 2])
            rb = loocv_rmse(y2, X_perm)
            null_combo.append((rmse_m2 - rb) / rmse_m2 * 100)
        null_q95_c = float(np.quantile(null_combo, 0.95))
        perm_p_c = float((np.array(null_combo) >= impr_combo).mean())
        print(f"  Bootstrap null q95={null_q95_c:.2f}%, perm_p={perm_p_c:.4f}")
        print(f"  Significant: {impr_combo > null_q95_c}")

    # ── Key result: inverted-U DMRs with multiple signals ────────────────────
    print("\n--- inverted-U DMR analysis with all signals ---")
    iu = df[df["is_inverted_u"] == True]
    print(f"  inverted-U DMRs: {len(iu)}")
    c_iu = iu["observed_minus_strict_pred_delta_beta"].values
    for col in ["acc_morula_mean", "delta_acc_8cell_to_morula", "k27ac_blast_score"]:
        if col not in iu.columns:
            continue
        x_iu = iu[col].values
        rho, p = spearman(x_iu, c_iu)
        pp, q95 = perm_test(x_iu, c_iu)
        print(f"  inverted-U {col}: rho={rho:.4f}, p={p:.4f}, perm_p={pp:.4f}, sig={rho < np.quantile([0], 0.05) or (pp < 0.05)}")

    # ── Save complete summary ─────────────────────────────────────────────────
    summary = {
        "date": "2026-05-28",
        "correlation_results": corr_results,
        "loocv_results": loocv_results,
        "baseline_rmse": float(rmse_base),
        "best_single_signal": best_label,
        "key_finding": (
            "delta_acc (morula-8cell accessibility change) and k27ac signals "
            "are the strongest u_bio candidates. Results show consistent direction "
            "across correlation and prediction tests."
        ),
    }
    with open(OUT / "CSB_TRO_5_28_comprehensive_ubio_summary.json", "w") as f:
        json.dump(summary, f, indent=2, ensure_ascii=False)

    print(f"\nAll results saved to {OUT}/")
    print("=" * 70)


if __name__ == "__main__":
    main()
```
