## Supplemental code for "An entropy-based diagnostic framework for characterizing methylation state dynamics during preimplantation development": 5_28_progress__h3k27me3_ubio_analysis.py

```
#!/usr/bin/env python
"""
H3K27me3 morula quantitative signal as u_bio.

H3K27me3 is a polycomb mark tightly coupled with DNA methylation.
Morula H3K27me3 quantitative bins (100bp resolution) provides
the strongest available quantitative u_bio candidate:
- Stage-matched (morula)
- Quantitative (not just binary overlap)
- Full genome coverage
- From published 2022 dataset

This script:
1. Computes DMR-level H3K27me3 morula signal by averaging bins
2. Also computes 8-cell H3K27me3 signal (from peak overlap)
3. Tests whether H3K27me3 morula signal explains c_diag
4. Tests prediction improvement (LOO-CV)
5. Combines H3K27me3 + accessibility for multi-input u_bio model
"""
from __future__ import annotations
import json
from pathlib import Path
import numpy as np
import pandas as pd
from scipy import stats
from numpy.linalg import solve

OUT = Path("E:/5_28_progress")
H3K27ME3_MORULA = Path(
    "E:/CSB_TRO_operator_time_DMR_dynamics_2026-05-25/external/histone/H3K27me3_morula.hg19.bed"
)
H3K27ME3_8CELL = Path(
    "E:/CSB_TRO_operator_time_DMR_dynamics_2026-05-25/external/histone/H3K27me3_8cell.hg19.bed.gz"
)
H3K27AC_8CELL = Path(
    "E:/CSB_TRO_operator_time_DMR_dynamics_2026-05-25/external/histone/H3K27ac_8cell.hg19.bed.gz"
)
H3K27AC_BLAST = Path(
    "E:/CSB_TRO_operator_time_DMR_dynamics_2026-05-25/external/histone/H3K27ac_blastocyst.hg19.bed.gz"
)
H3K4ME3_8CELL = Path(
    "E:/CSB_TRO_operator_time_DMR_dynamics_2026-05-25/external/histone/H3K4me3_8cell.hg19.bed.gz"
)

DMR_META = Path(
    "E:/CSB_TRO_operator_time_DMR_dynamics_2026-05-25/results/CSB_TRO_DMR_metadata.tsv"
)
DMR_RESIDUAL = Path(
    "E:/CSB_TRO_operator_time_DMR_dynamics_2026-05-25/results/CSB_TRO_basin_residual_DMR_ranking.tsv"
)
DMR_TRAJ = Path(
    "E:/CSB_TRO_operator_time_DMR_dynamics_2026-05-25/results/CSB_TRO_DMR_stage_mean_trajectory.tsv"
)
CURV = Path("E:/实验进展5_27/CSB_TRO_2026-05-27_entry_exit_curvature.tsv")
MULTISTAGE_ACC = OUT / "CSB_TRO_5_28_dmr_multistage_accessibility.tsv"

SEED = 42
N_BOOT = 2000


def spearman(x, y):
    mask = np.isfinite(x) & np.isfinite(y)
    if mask.sum() < 5:
        return np.nan, np.nan
    r, p = stats.spearmanr(x[mask], y[mask])
    return float(r), float(p)


def perm_p(x, y, n=N_BOOT, seed=SEED, one_sided=True):
    rng = np.random.default_rng(seed)
    obs_rho, _ = spearman(x, y)
    if not np.isfinite(obs_rho):
        return np.nan, np.nan
    mask = np.isfinite(x) & np.isfinite(y)
    xv, yv = x[mask], y[mask]
    nulls = []
    for _ in range(n):
        r, _ = stats.spearmanr(rng.permutation(xv), yv)
        nulls.append(r)
    nulls = np.array(nulls)
    q95 = float(np.quantile(nulls, 0.95))
    if one_sided:
        pp = float((nulls >= obs_rho).mean())
    else:
        pp = float((np.abs(nulls) >= np.abs(obs_rho)).mean())
    return pp, q95


def overlap_bins_to_dmr(bins_df, dmr_meta, value_col="signal"):
    """Average bin signal overlapping each DMR."""
    bins_df["chr"] = bins_df["chr"].astype(str).str.strip()
    dmr_meta["chr"] = dmr_meta["chr"].astype(str).str.strip()
    bins_by_chr = {c: g for c, g in bins_df.groupby("chr")}

    rows = []
    for _, dmr in dmr_meta.iterrows():
        chrom, d_s, d_e = dmr["chr"], int(dmr["start"]), int(dmr["end"])
        cluster = dmr["cluster_name"]
        if chrom not in bins_by_chr:
            rows.append({"cluster_name": cluster, "mean_signal": np.nan, "max_signal": np.nan, "n_bins": 0})
            continue
        sub = bins_by_chr[chrom]
        mask = (sub["start"] < d_e) & (sub["end"] > d_s)
        hits = sub[mask][value_col].dropna()
        rows.append({
            "cluster_name": cluster,
            "mean_signal": float(hits.mean()) if len(hits) > 0 else np.nan,
            "max_signal": float(hits.max()) if len(hits) > 0 else np.nan,
            "n_bins": int(len(hits)),
        })
    return pd.DataFrame(rows)


def overlap_peaks_to_dmr(peaks_df, dmr_meta):
    """Binary overlap + score for peak files."""
    peaks_df["chr"] = peaks_df["chr"].astype(str).str.strip()
    dmr_meta["chr"] = dmr_meta["chr"].astype(str).str.strip()
    peaks_by_chr = {c: g for c, g in peaks_df.groupby("chr")}

    rows = []
    for _, dmr in dmr_meta.iterrows():
        chrom, d_s, d_e = dmr["chr"], int(dmr["start"]), int(dmr["end"])
        cluster = dmr["cluster_name"]
        if chrom not in peaks_by_chr:
            rows.append({"cluster_name": cluster, "overlap": 0, "peak_score_max": np.nan})
            continue
        sub = peaks_by_chr[chrom]
        mask = (sub["start"] < d_e) & (sub["end"] > d_s)
        hits = sub[mask]
        score = hits.iloc[:, 4].dropna() if len(hits.columns) > 4 else pd.Series(dtype=float)
        rows.append({
            "cluster_name": cluster,
            "overlap": int(len(hits) > 0),
            "peak_score_max": float(score.max()) if len(score) > 0 else np.nan,
        })
    return pd.DataFrame(rows)


def loocv(y, X, lam=0.01):
    """Ridge LOO-CV, returns per-sample errors."""
    n = len(y)
    errors = []
    reg = np.diag([lam] * X.shape[1])
    reg[-1, -1] = 0  # no penalty on bias
    for i in range(n):
        tr = [j for j in range(n) if j != i]
        Xtr = X[tr]; ytr = y[tr]
        try:
            w = solve(Xtr.T @ Xtr + reg, Xtr.T @ ytr)
            pred = X[i] @ w
            errors.append(abs(y[i] - pred))
        except Exception:
            continue
    return np.array(errors)


def main():
    print("=" * 70)
    print("H3K27me3 morula quantitative signal as u_bio")
    print("=" * 70)

    # ── Load core tables ───────────────────────────────────────────────────────
    dmr_meta = pd.read_csv(DMR_META, sep="\t")
    residual = pd.read_csv(DMR_RESIDUAL, sep="\t")
    traj = pd.read_csv(DMR_TRAJ, sep="\t")
    curv = pd.read_csv(CURV, sep="\t")
    multistage = pd.read_csv(MULTISTAGE_ACC, sep="\t")

    stage_means = {}
    for stage, g in traj.groupby("stage"):
        stage_means[stage] = g.set_index("cluster_name")["mean_beta"].to_dict()

    clusters = residual["cluster_name"].tolist()
    y_morula = np.array([stage_means.get("morula", {}).get(c, np.nan) for c in clusters])
    x_8cell = np.array([stage_means.get("8-cell", {}).get(c, np.nan) for c in clusters])

    c_strict = residual.set_index("cluster_name")["observed_minus_strict_pred_delta_beta"].to_dict()
    strict_corr = np.array([c_strict.get(c, np.nan) for c in clusters])

    # ── Part 1: H3K27me3 morula quantitative bins → DMR signal ────────────────
    print("\n--- Part 1: H3K27me3 morula bins → DMR-level signal ---")
    print("  Loading H3K27me3 morula bins (14.8M rows)... ", end="", flush=True)
    bins = pd.read_csv(H3K27ME3_MORULA, sep="\t", header=None,
                       names=["chr", "start", "end", "signal"],
                       dtype={"chr": str, "start": int, "end": int, "signal": float})
    print(f"done ({len(bins)} bins)")

    k27me3_morula = overlap_bins_to_dmr(bins, dmr_meta, value_col="signal")
    k27me3_morula = k27me3_morula.rename(columns={
        "mean_signal": "k27me3_morula_mean",
        "max_signal": "k27me3_morula_max",
        "n_bins": "k27me3_morula_nbins",
    })

    has_sig = k27me3_morula["k27me3_morula_mean"].notna().sum()
    print(f"  DMRs with H3K27me3 morula signal: {has_sig}/156")

    # ── Part 2: H3K27me3 8-cell peaks → DMR overlap ───────────────────────────
    print("\n--- Part 2: H3K27me3 8-cell peaks → DMR overlap ---")
    peaks_8cell = pd.read_csv(H3K27ME3_8CELL, sep="\t", header=None,
                               compression="gzip",
                               names=["chr","start","end","name","score","strand","fc","neglog10p","neglog10q"],
                               usecols=[0,1,2,3,4])
    k27me3_8cell = overlap_peaks_to_dmr(peaks_8cell, dmr_meta)
    k27me3_8cell = k27me3_8cell.rename(columns={
        "overlap": "k27me3_8cell_overlap",
        "peak_score_max": "k27me3_8cell_score",
    })
    print(f"  8-cell peaks: {len(peaks_8cell)}, DMRs with overlap: {k27me3_8cell['k27me3_8cell_overlap'].sum()}")

    # ── Part 3: H3K27ac peaks (8-cell + blastocyst) ───────────────────────────
    print("\n--- Part 3: H3K27ac peaks ---")
    peaks_k27ac_8cell = pd.read_csv(H3K27AC_8CELL, sep="\t", header=None,
                                     compression="gzip",
                                     names=["chr","start","end","name","score","strand","fc","neglog10p","neglog10q"],
                                     usecols=[0,1,2,3,4])
    k27ac_8cell = overlap_peaks_to_dmr(peaks_k27ac_8cell, dmr_meta).rename(columns={
        "overlap": "k27ac_8cell_overlap", "peak_score_max": "k27ac_8cell_score"})

    peaks_k27ac_blast = pd.read_csv(H3K27AC_BLAST, sep="\t", header=None,
                                     compression="gzip",
                                     names=["chr","start","end","name","score","strand","fc","neglog10p","neglog10q"],
                                     usecols=[0,1,2,3,4])
    k27ac_blast = overlap_peaks_to_dmr(peaks_k27ac_blast, dmr_meta).rename(columns={
        "overlap": "k27ac_blast_overlap", "peak_score_max": "k27ac_blast_score"})

    print(f"  H3K27ac 8-cell peaks: {len(peaks_k27ac_8cell)}")
    print(f"  H3K27ac blastocyst peaks: {len(peaks_k27ac_blast)}")

    # ── Part 4: H3K4me3 8-cell peaks ─────────────────────────────────────────
    peaks_k4me3_8cell = pd.read_csv(H3K4ME3_8CELL, sep="\t", header=None,
                                     compression="gzip",
                                     names=["chr","start","end","name","score","strand","fc","neglog10p","neglog10q"],
                                     usecols=[0,1,2,3,4])
    k4me3_8cell = overlap_peaks_to_dmr(peaks_k4me3_8cell, dmr_meta).rename(columns={
        "overlap": "k4me3_8cell_overlap", "peak_score_max": "k4me3_8cell_score"})

    # ── Merge all signals ─────────────────────────────────────────────────────
    merged = residual[["cluster_name", "basin_residual_rank",
                        "observed_minus_strict_pred_delta_beta",
                        "latent_residual_delta_beta", "module_id"]].copy()
    merged = merged.merge(k27me3_morula, on="cluster_name", how="left")
    merged = merged.merge(k27me3_8cell, on="cluster_name", how="left")
    merged = merged.merge(k27ac_8cell, on="cluster_name", how="left")
    merged = merged.merge(k27ac_blast, on="cluster_name", how="left")
    merged = merged.merge(k4me3_8cell, on="cluster_name", how="left")
    merged = merged.merge(curv[["cluster_name", "curvature", "is_inverted_u", "is_u_shape"]],
                          on="cluster_name", how="left")
    merged = merged.merge(
        multistage[["cluster_name", "acc_morula_mean", "acc_8-cell_mean", "delta_acc_8cell_to_morula"]],
        on="cluster_name", how="left"
    )

    # Add methylation
    for st in ["2-cell", "4-cell", "8-cell", "morula", "blastocyst"]:
        safe = st.replace("-", "_")
        merged[f"meth_{safe}"] = merged["cluster_name"].map(
            lambda c: stage_means.get(st, {}).get(c, np.nan))

    merged.to_csv(OUT / "CSB_TRO_5_28_dmr_full_ubio_signals.tsv", sep="\t", index=False)
    print(f"\n  Full u_bio signal table saved.")

    # ── Part 5: Test H3K27me3 morula as u_bio ─────────────────────────────────
    print("\n--- Part 5: H3K27me3 morula ~ correction term ---")
    u_k27 = merged["k27me3_morula_mean"].values
    c = merged["observed_minus_strict_pred_delta_beta"].values

    rho_k27, p_k27 = spearman(u_k27, c)
    pp_k27, q95_k27 = perm_p(u_k27, c)
    print(f"  H3K27me3 morula ~ strict_correction: rho={rho_k27:.4f}, p={p_k27:.4f}")
    print(f"  Null q95={q95_k27:.4f}, perm_p={pp_k27:.4f}, sig={rho_k27 > q95_k27}")

    # By curvature class
    for grp, mask in [
        ("inverted_u", merged["is_inverted_u"] == True),
        ("u_shape", merged["is_u_shape"] == True),
        ("neg_curv", merged["curvature"] < 0),
        ("top25_residual", merged["basin_residual_rank"] <= 25),
    ]:
        sub = merged[mask].dropna(subset=["k27me3_morula_mean", "observed_minus_strict_pred_delta_beta"])
        if len(sub) < 5:
            continue
        r, p = spearman(sub["k27me3_morula_mean"].values, sub["observed_minus_strict_pred_delta_beta"].values)
        print(f"  {grp} (n={len(sub)}): rho={rho_k27:.4f} -> group rho={r:.4f}, p={p:.4f}")

    # ── Part 6: LOO-CV with H3K27me3 as u_bio ────────────────────────────────
    print("\n--- Part 6: LOO-CV with H3K27me3 morula ---")
    lam = 0.01
    valid = merged.dropna(subset=["meth_morula", "meth_8_cell", "k27me3_morula_mean"])
    print(f"  DMRs with all signals: {len(valid)}")

    y_v = valid["meth_morula"].values
    x_v = valid["meth_8_cell"].values
    u_v = valid["k27me3_morula_mean"].values
    # Standardize u
    u_sc = (u_v - u_v.mean()) / (u_v.std() + 1e-8)

    X_meth = np.column_stack([x_v, np.ones(len(x_v))])
    X_bio = np.column_stack([x_v, u_sc, np.ones(len(x_v))])

    errs_meth = loocv(y_v, X_meth, lam)
    errs_bio = loocv(y_v, X_bio, lam)
    rmse_m = float(np.sqrt(np.mean(errs_meth ** 2)))
    rmse_b = float(np.sqrt(np.mean(errs_bio ** 2)))
    impr = (rmse_m - rmse_b) / rmse_m * 100
    print(f"  RMSE meth-only: {rmse_m:.4f}")
    print(f"  RMSE bio (H3K27me3): {rmse_b:.4f}")
    print(f"  Improvement: {impr:.2f}%")
    print(f"  Bio better: {rmse_b < rmse_m}")

    # Bootstrap perm test
    rng = np.random.default_rng(SEED)
    null_imprs = []
    for _ in range(N_BOOT):
        perm_u = rng.permutation(u_sc)
        X_perm = np.column_stack([x_v, perm_u, np.ones(len(x_v))])
        e_perm = loocv(y_v, X_perm, lam)
        if len(e_perm) == len(errs_meth):
            rm = np.sqrt(np.mean(errs_meth ** 2))
            rb = np.sqrt(np.mean(e_perm ** 2))
            null_imprs.append((rm - rb) / rm * 100)
    null_q95 = float(np.quantile(null_imprs, 0.95))
    pp_loocv = float((np.array(null_imprs) >= impr).mean())
    print(f"  Bootstrap null q95: {null_q95:.2f}%, perm_p={pp_loocv:.4f}")
    print(f"  Improvement > null q95: {impr > null_q95}")

    # ── Part 7: Multi-signal u_bio model ─────────────────────────────────────
    print("\n--- Part 7: Multi-signal u_bio (H3K27me3 + accessibility) ---")
    valid2 = merged.dropna(subset=["meth_morula", "meth_8_cell",
                                    "k27me3_morula_mean", "delta_acc_8cell_to_morula"])
    print(f"  DMRs with H3K27me3 + delta_acc: {len(valid2)}")

    y2 = valid2["meth_morula"].values
    x2 = valid2["meth_8_cell"].values
    u_k = (valid2["k27me3_morula_mean"].values -
           valid2["k27me3_morula_mean"].mean()) / (valid2["k27me3_morula_mean"].std() + 1e-8)
    u_a = (valid2["delta_acc_8cell_to_morula"].values -
           valid2["delta_acc_8cell_to_morula"].mean()) / (valid2["delta_acc_8cell_to_morula"].std() + 1e-8)

    X_m2 = np.column_stack([x2, np.ones(len(x2))])
    X_k = np.column_stack([x2, u_k, np.ones(len(x2))])
    X_a = np.column_stack([x2, u_a, np.ones(len(x2))])
    X_ka = np.column_stack([x2, u_k, u_a, np.ones(len(x2))])

    for label, X in [("meth_only", X_m2), ("H3K27me3", X_k),
                     ("delta_acc", X_a), ("H3K27me3+delta_acc", X_ka)]:
        lam_reg = np.diag([lam] * X.shape[1])
        lam_reg[-1, -1] = 0
        e = loocv(y2, X, lam)
        r = float(np.sqrt(np.mean(e ** 2)))
        base_e = loocv(y2, X_m2, lam)
        base_r = float(np.sqrt(np.mean(base_e ** 2)))
        impr2 = (base_r - r) / base_r * 100 if label != "meth_only" else 0
        print(f"  {label}: RMSE={r:.4f}, improvement={impr2:.2f}%")

    # ── Part 8: Top residual DMR H3K27me3 signal vs random ───────────────────
    print("\n--- Part 8: Top residual DMR H3K27me3 signal ---")
    rng2 = np.random.default_rng(SEED + 1)
    all_k27 = merged["k27me3_morula_mean"].dropna().values
    for k in [25, 50]:
        top = merged[merged["basin_residual_rank"] <= k]
        obs = top["k27me3_morula_mean"].mean()
        null = [rng2.choice(all_k27, size=len(top.dropna(subset=["k27me3_morula_mean"])),
                            replace=False).mean() for _ in range(N_BOOT)]
        q95 = float(np.quantile(null, 0.95))
        print(f"  Top{k}: mean={obs:.3f}, null q95={q95:.3f}, sig={obs > q95}")

    # ── Save summary ──────────────────────────────────────────────────────────
    summary = {
        "date": "2026-05-28",
        "h3k27me3_morula_vs_correction": {
            "rho": float(rho_k27), "p": float(p_k27),
            "perm_p": float(pp_k27), "null_q95": float(q95_k27),
            "significant": bool(rho_k27 > q95_k27),
        },
        "loocv_h3k27me3_ubio": {
            "rmse_meth_only": float(rmse_m),
            "rmse_bio": float(rmse_b),
            "improvement_pct": float(impr),
            "perm_p": float(pp_loocv),
            "null_q95": float(null_q95),
            "significant": bool(impr > null_q95),
        },
    }
    with open(OUT / "CSB_TRO_5_28_h3k27me3_ubio_summary.json", "w") as f:
        json.dump(summary, f, indent=2)

    print(f"\nSaved: {OUT}/CSB_TRO_5_28_h3k27me3_ubio_summary.json")
    print("\n" + "=" * 70)
    print("SUMMARY")
    print(f"H3K27me3 morula ~ correction: rho={rho_k27:.4f}, sig={rho_k27>q95_k27}")
    print(f"LOO-CV improvement: {impr:.2f}%, sig={impr>null_q95}")
    print("=" * 70)


if __name__ == "__main__":
    main()
```
