## Supplemental code for "An entropy-based diagnostic framework for characterizing methylation state dynamics during preimplantation development": 5_28_progress__multistage_accessibility_dynamics.py

```
#!/usr/bin/env python
"""
Stage-specific u_bio dynamics analysis using Liu2019 multi-stage accessibility.

Liu2019 provides quantitative accessibility for 4 stages:
  2-cell, 4-cell, 8-cell, morula

GSE81233 provides methylation for matching stages.

This is the first true paired multi-stage methylation + accessibility
dynamic analysis. Key questions:
1. Does the accessibility trajectory toward morula predict methylation change?
2. Does morula accessibility SPECIFICALLY exceed 8-cell accessibility
   at residual DMRs?
3. Does the stage-specific accessibility trajectory couple to DMR curvature?
4. Can accessibility trajectory improve stage-to-stage methylation prediction?
"""
from __future__ import annotations
import json
from pathlib import Path
import numpy as np
import pandas as pd
from scipy import stats
from numpy.linalg import solve

OUT = Path("E:/5_28_progress")
OUT.mkdir(parents=True, exist_ok=True)

LIU2019 = Path("E:/实验进展5_27/CSB_TRO_2026-05-27_u_bio_rescue_extracted_coordinate_regions.tsv")
DMR_METADATA = Path("E:/CSB_TRO_operator_time_DMR_dynamics_2026-05-25/results/CSB_TRO_DMR_metadata.tsv")
DMR_RESIDUAL = Path("E:/CSB_TRO_operator_time_DMR_dynamics_2026-05-25/results/CSB_TRO_basin_residual_DMR_ranking.tsv")
DMR_TRAJ = Path("E:/CSB_TRO_operator_time_DMR_dynamics_2026-05-25/results/CSB_TRO_DMR_stage_mean_trajectory.tsv")
CURV = Path("E:/实验进展5_27/CSB_TRO_2026-05-27_entry_exit_curvature.tsv")

SEED = 42
N_BOOT = 2000


def cosine(a, b):
    n = np.linalg.norm(a) * np.linalg.norm(b)
    return float(np.dot(a, b) / n) if n > 1e-12 else np.nan


def spearman(x, y):
    mask = np.isfinite(x) & np.isfinite(y)
    if mask.sum() < 5:
        return np.nan, np.nan
    r, p = stats.spearmanr(x[mask], y[mask])
    return float(r), float(p)


def bootstrap_q95(values, size, n_boot=2000, seed=42):
    rng = np.random.default_rng(seed)
    vals = np.array(values, dtype=float)
    vals = vals[np.isfinite(vals)]
    if len(vals) < size:
        return np.nan
    means = [rng.choice(vals, size=size, replace=False).mean() for _ in range(n_boot)]
    return float(np.quantile(means, 0.95))


def main():
    print("=" * 70)
    print("Stage-specific u_bio dynamics: Liu2019 multi-stage accessibility")
    print("=" * 70)

    # ── Load data ──────────────────────────────────────────────────────────────
    liu = pd.read_csv(LIU2019, sep="\t")
    for col in ["Accessibility_2-cell", "Accessibility_4-cell",
                "Accessibility_8-cell", "Accessibility_Morula"]:
        liu[col] = pd.to_numeric(liu[col], errors="coerce")

    dmr_meta = pd.read_csv(DMR_METADATA, sep="\t")
    residual = pd.read_csv(DMR_RESIDUAL, sep="\t")
    traj = pd.read_csv(DMR_TRAJ, sep="\t")
    curv = pd.read_csv(CURV, sep="\t")

    stage_means = {}
    for stage, g in traj.groupby("stage"):
        stage_means[stage] = g.set_index("cluster_name")["mean_beta"].to_dict()

    print(f"Liu2019 regions: {len(liu)}")
    print(f"DMRs: {len(dmr_meta)}")

    # ── Part 1: DMR-level stage-specific accessibility trajectory ──────────────
    print("\n--- Part 1: Build DMR-level multi-stage accessibility ---")

    # Intersect each DMR with Liu2019 regions
    liu["chr"] = liu["chr"].astype(str).str.strip()
    dmr_meta["chr"] = dmr_meta["chr"].astype(str).str.strip()
    liu_by_chr = {c: g for c, g in liu.groupby("chr")}

    acc_stages = {"2-cell": "Accessibility_2-cell",
                  "4-cell": "Accessibility_4-cell",
                  "8-cell": "Accessibility_8-cell",
                  "morula": "Accessibility_Morula"}

    rows = []
    for _, dmr in dmr_meta.iterrows():
        chrom = dmr["chr"]
        d_start, d_end = int(dmr["start"]), int(dmr["end"])
        cluster = dmr["cluster_name"]

        row = {"cluster_name": cluster}
        if chrom in liu_by_chr:
            sub = liu_by_chr[chrom]
            mask = (sub["start"] < d_end) & (sub["end"] > d_start)
            hits = sub[mask]
            n_hits = len(hits)
            for stage, col in acc_stages.items():
                vals = pd.to_numeric(hits[col], errors="coerce").dropna()
                row[f"acc_{stage}_mean"] = float(vals.mean()) if len(vals) > 0 else np.nan
                row[f"acc_{stage}_max"] = float(vals.max()) if len(vals) > 0 else np.nan
            row["n_overlapping_peaks"] = n_hits
        else:
            for stage in acc_stages:
                row[f"acc_{stage}_mean"] = np.nan
                row[f"acc_{stage}_max"] = np.nan
            row["n_overlapping_peaks"] = 0

        rows.append(row)

    dmr_acc = pd.DataFrame(rows)

    # Add methylation stage means
    for mstage in ["2-cell", "4-cell", "8-cell", "morula", "blastocyst"]:
        safe = mstage.replace("-", "_")
        dmr_acc[f"meth_{safe}"] = dmr_acc["cluster_name"].map(
            lambda c: stage_means.get(mstage, {}).get(c, np.nan)
        )

    # Add residual info
    dmr_acc = dmr_acc.merge(
        residual[["cluster_name", "basin_residual_rank",
                  "observed_minus_strict_pred_delta_beta",
                  "latent_residual_delta_beta", "module_id"]],
        on="cluster_name", how="left"
    )
    dmr_acc = dmr_acc.merge(
        curv[["cluster_name", "curvature", "is_inverted_u", "is_u_shape"]],
        on="cluster_name", how="left"
    )

    dmr_acc.to_csv(OUT / "CSB_TRO_5_28_dmr_multistage_accessibility.tsv",
                   sep="\t", index=False)
    print(f"  Saved DMR multi-stage accessibility table")

    has_morula = dmr_acc["acc_morula_mean"].notna().sum()
    has_8cell = dmr_acc["acc_8-cell_mean"].notna().sum()
    print(f"  DMRs with morula acc: {has_morula}/156")
    print(f"  DMRs with 8-cell acc: {has_8cell}/156")

    # ── Part 2: Accessibility TRAJECTORY — key innovation ─────────────────────
    # Define: delta_acc = acc_morula - acc_8cell (stage-specific change)
    # This is morula-specific accessibility GAIN — the true u_bio candidate
    print("\n--- Part 2: Accessibility trajectory (morula - 8cell delta) ---")

    dmr_acc["delta_acc_8cell_to_morula"] = (
        dmr_acc["acc_morula_mean"] - dmr_acc["acc_8-cell_mean"]
    )
    # Also define methylation change
    dmr_acc["delta_meth_8cell_to_morula"] = (
        dmr_acc["meth_morula"] - dmr_acc["meth_8_cell"]
    )
    dmr_acc["strict_correction"] = dmr_acc["observed_minus_strict_pred_delta_beta"]

    valid = dmr_acc.dropna(subset=["delta_acc_8cell_to_morula", "strict_correction"])
    print(f"  DMRs with both delta_acc and strict_correction: {len(valid)}")

    rho_delta, p_delta = spearman(
        valid["delta_acc_8cell_to_morula"].values,
        valid["strict_correction"].values
    )
    print(f"  delta_acc ~ strict_correction: rho={rho_delta:.4f}, p={p_delta:.4f}")

    # This is more powerful than morula accessibility alone because it captures
    # the CHANGE from 8-cell to morula — the stage-specific signal
    rng = np.random.default_rng(SEED)
    null_rhos = []
    u = valid["delta_acc_8cell_to_morula"].values
    c = valid["strict_correction"].values
    for _ in range(N_BOOT):
        perm = rng.permutation(len(u))
        r, _ = stats.spearmanr(u[perm], c)
        null_rhos.append(r)
    null_rhos = np.array(null_rhos)
    perm_p = float((null_rhos >= rho_delta).mean())
    null_q95 = float(np.quantile(null_rhos, 0.95))
    print(f"  Null q95: {null_q95:.4f}, perm_p (one-sided): {perm_p:.4f}")
    print(f"  delta_acc > null q95: {rho_delta > null_q95}")

    # ── Part 3: Stage-specific accessibility as DYNAMIC u_bio ─────────────────
    print("\n--- Part 3: Stage-wise accessibility trajectory vs methylation ---")

    # At each stage, does accessibility predict methylation change to next stage?
    stage_pairs = [
        ("2-cell", "4-cell"),
        ("4-cell", "8-cell"),
        ("8-cell", "morula"),   # key transition
    ]
    traj_results = []
    for s_from, s_to in stage_pairs:
        sf = s_from.replace("-", "_")
        st = s_to.replace("-", "_")
        acc_col = f"acc_{s_to}_mean"  # accessibility at target stage
        meth_from = f"meth_{sf}"
        meth_to = f"meth_{st}"

        sub = dmr_acc.dropna(subset=[acc_col, meth_from, meth_to])
        if len(sub) < 10:
            continue

        delta_meth = sub[meth_to].values - sub[meth_from].values
        acc = sub[acc_col].values

        rho, p = spearman(acc, delta_meth)
        print(f"  {s_from}->{s_to}: acc ~ delta_meth: rho={rho:.4f}, p={p:.4f}, n={len(sub)}")
        traj_results.append({
            "transition": f"{s_from}_to_{s_to}",
            "n": len(sub),
            "rho": rho,
            "p": p,
        })

    # ── Part 4: Leave-morula-out with multi-stage u_bio ───────────────────────
    print("\n--- Part 4: Leave-morula-out with stage-specific u_bio ---")

    # Now use BOTH 8-cell methylation AND delta_acc as predictors
    lam = 0.01
    clusters = dmr_acc["cluster_name"].tolist()
    y_morula = np.array([stage_means.get("morula", {}).get(c, np.nan) for c in clusters])
    x_8cell = np.array([stage_means.get("8-cell", {}).get(c, np.nan) for c in clusters])
    delta_acc = dmr_acc["delta_acc_8cell_to_morula"].values
    acc_morula = dmr_acc["acc_morula_mean"].values

    valid_mask = np.isfinite(y_morula) & np.isfinite(x_8cell) & np.isfinite(delta_acc)
    idx_v = np.where(valid_mask)[0]
    print(f"  DMRs with all 3 signals: {valid_mask.sum()}")

    if valid_mask.sum() >= 20:
        # LOO-CV: meth-only vs meth + delta_acc (the true stage-specific u_bio)
        err_meth, err_bio = [], []
        for i in idx_v:
            tr = idx_v[idx_v != i]

            Xm = np.column_stack([x_8cell[tr], np.ones(len(tr))])
            yt = y_morula[tr]
            reg2 = np.diag([lam, 0.0])
            try:
                cm = solve(Xm.T @ Xm + reg2, Xm.T @ yt)
                pred_m = cm[0] * x_8cell[i] + cm[1]
                err_meth.append(abs(y_morula[i] - pred_m))
            except:
                continue

            Xb = np.column_stack([x_8cell[tr], delta_acc[tr], np.ones(len(tr))])
            reg3 = np.diag([lam, lam, 0.0])
            try:
                cb = solve(Xb.T @ Xb + reg3, Xb.T @ yt)
                pred_b = cb[0] * x_8cell[i] + cb[1] * delta_acc[i] + cb[2]
                err_bio.append(abs(y_morula[i] - pred_b))
            except:
                continue

        rmse_meth = float(np.sqrt(np.mean(np.array(err_meth) ** 2)))
        rmse_bio = float(np.sqrt(np.mean(np.array(err_bio) ** 2)))
        improvement = (rmse_meth - rmse_bio) / rmse_meth * 100
        print(f"  All valid DMRs LOO-CV RMSE meth-only: {rmse_meth:.4f}")
        print(f"  All valid DMRs LOO-CV RMSE bio(delta_acc): {rmse_bio:.4f}")
        print(f"  Improvement: {improvement:.2f}%")
        print(f"  Bio model better: {rmse_bio < rmse_meth}")

        # Bootstrap test
        null_impr = []
        y_v = y_morula[idx_v]
        x_v = x_8cell[idx_v]
        d_v = delta_acc[idx_v]
        for _ in range(N_BOOT):
            perm_d = rng.permutation(d_v)
            e_m, e_b = [], []
            for i in range(len(idx_v)):
                tr = [j for j in range(len(idx_v)) if j != i]
                Xm = np.column_stack([x_v[tr], np.ones(len(tr))])
                yt = y_v[tr]
                try:
                    cm = solve(Xm.T @ Xm + np.diag([lam, 0.0]), Xm.T @ yt)
                    e_m.append(abs(y_v[i] - (cm[0] * x_v[i] + cm[1])))
                    Xb = np.column_stack([x_v[tr], perm_d[tr], np.ones(len(tr))])
                    cb = solve(Xb.T @ Xb + np.diag([lam, lam, 0.0]), Xb.T @ yt)
                    e_b.append(abs(y_v[i] - (cb[0] * x_v[i] + cb[1] * perm_d[i] + cb[2])))
                except:
                    continue
            if e_m and e_b:
                rm = np.sqrt(np.mean(np.array(e_m) ** 2))
                rb = np.sqrt(np.mean(np.array(e_b) ** 2))
                null_impr.append((rm - rb) / rm * 100)

        null_q95_impr = float(np.quantile(null_impr, 0.95))
        perm_p_impr = float((np.array(null_impr) >= improvement).mean())
        print(f"  Bootstrap null q95 improvement: {null_q95_impr:.2f}%")
        print(f"  Bootstrap p (improvement > null): {perm_p_impr:.4f}")
        print(f"  Improvement > null q95: {improvement > null_q95_impr}")
    else:
        rmse_meth = rmse_bio = improvement = np.nan
        null_q95_impr = perm_p_impr = np.nan

    # ── Part 5: The key innovation — accessibility TRAJECTORY is u_bio ────────
    print("\n--- Part 5: Accessibility trajectory as dynamic u_bio ---")
    # Define the accessibility trajectory vector across stages:
    # For each DMR: [acc_2cell, acc_4cell, acc_8cell, acc_morula]
    # This is a time series — does it covary with methylation changes?

    # Stage-wise correlation: at each stage, does accessibility predict
    # methylation at that stage?
    print("  Stage-wise: acc_stage ~ meth_stage")
    for stage in ["2-cell", "4-cell", "8-cell", "morula"]:
        sf = stage.replace("-", "_")
        acc_col = f"acc_{stage}_mean"
        meth_col = f"meth_{sf}"
        sub = dmr_acc.dropna(subset=[acc_col, meth_col])
        rho, p = spearman(sub[acc_col].values, sub[meth_col].values)
        print(f"    {stage}: n={len(sub)}, rho={rho:.4f}, p={p:.4f}")

    # Critical test: does accessibility CHANGE (morula - 8cell) predict
    # METHYLATION CHANGE (morula - 8cell)?
    print("\n  Key test: delta_acc ~ delta_meth (8cell->morula)")
    sub = dmr_acc.dropna(subset=["delta_acc_8cell_to_morula", "delta_meth_8cell_to_morula"])
    rho_key, p_key = spearman(
        sub["delta_acc_8cell_to_morula"].values,
        sub["delta_meth_8cell_to_morula"].values
    )
    print(f"  n={len(sub)}, rho={rho_key:.4f}, p={p_key:.4f}")

    # Permutation test
    null_key = []
    u_k = sub["delta_acc_8cell_to_morula"].values
    c_k = sub["delta_meth_8cell_to_morula"].values
    for _ in range(N_BOOT):
        perm = rng.permutation(len(u_k))
        r, _ = stats.spearmanr(u_k[perm], c_k)
        null_key.append(r)
    null_key = np.array(null_key)
    perm_p_key = float((null_key >= rho_key).mean())
    null_q95_key = float(np.quantile(null_key, 0.95))
    print(f"  Null q95={null_q95_key:.4f}, perm_p={perm_p_key:.4f}")
    print(f"  delta_acc > null q95: {rho_key > null_q95_key}")

    # Curvature-stratified
    print("\n  Curvature-stratified delta_acc ~ delta_meth:")
    for group, mask in [
        ("all", sub.index),
        ("inverted_u", sub[sub["is_inverted_u"] == True].index),
        ("u_shape", sub[sub["is_u_shape"] == True].index),
        ("neg_curvature", sub[sub["curvature"] < 0].index),
    ]:
        g = sub.loc[mask]
        if len(g) < 5:
            continue
        rho_g, p_g = spearman(
            g["delta_acc_8cell_to_morula"].values,
            g["delta_meth_8cell_to_morula"].values
        )
        print(f"    {group} (n={len(g)}): rho={rho_g:.4f}, p={p_g:.4f}")

    # ── Part 6: Top residual DMR accessibility trajectory ─────────────────────
    print("\n--- Part 6: Top residual DMR accessibility trajectory ---")
    rng2 = np.random.default_rng(SEED + 1)
    all_vals_morula = dmr_acc["acc_morula_mean"].dropna().values
    all_vals_delta = dmr_acc["delta_acc_8cell_to_morula"].dropna().values

    for k in [25, 50]:
        top = dmr_acc[dmr_acc["basin_residual_rank"] <= k]
        print(f"  Top{k} residual DMRs:")

        obs_morula = top["acc_morula_mean"].mean()
        q95_morula = bootstrap_q95(all_vals_morula, len(top.dropna(subset=["acc_morula_mean"])), N_BOOT, SEED)
        print(f"    morula_acc mean={obs_morula:.4f}, random q95={q95_morula:.4f}, sig={obs_morula > q95_morula}")

        obs_delta = top["delta_acc_8cell_to_morula"].mean()
        q95_delta = bootstrap_q95(all_vals_delta, len(top.dropna(subset=["delta_acc_8cell_to_morula"])), N_BOOT, SEED + 1)
        print(f"    delta_acc mean={obs_delta:.4f}, random q95={q95_delta:.4f}, sig={obs_delta > q95_delta}")

    # ── Summary ───────────────────────────────────────────────────────────────
    summary = {
        "date": "2026-05-28",
        "key_innovation": (
            "Liu2019 provides 4-stage quantitative accessibility (2-cell, 4-cell, "
            "8-cell, morula). This enables true stage-specific u_bio analysis: "
            "delta_acc = acc_morula - acc_8cell is the morula-specific accessibility gain."
        ),
        "part1_data_coverage": {
            "n_dmrs_with_morula_acc": int(has_morula),
            "n_dmrs_with_8cell_acc": int(has_8cell),
        },
        "part2_delta_acc_vs_correction": {
            "rho": float(rho_delta),
            "p": float(p_delta),
            "perm_p_onesided": float(perm_p),
            "null_q95": float(null_q95),
            "significant": bool(rho_delta > null_q95),
        },
        "part4_loocv_with_delta_acc": {
            "rmse_meth_only": float(rmse_meth) if np.isfinite(rmse_meth) else None,
            "rmse_bio_delta_acc": float(rmse_bio) if np.isfinite(rmse_bio) else None,
            "improvement_pct": float(improvement) if np.isfinite(improvement) else None,
            "bootstrap_perm_p": float(perm_p_impr) if np.isfinite(perm_p_impr) else None,
            "bootstrap_null_q95": float(null_q95_impr) if np.isfinite(null_q95_impr) else None,
            "improvement_gt_null_q95": bool(improvement > null_q95_impr) if np.isfinite(improvement) else None,
        },
        "part5_delta_acc_delta_meth": {
            "rho": float(rho_key),
            "p": float(p_key),
            "perm_p": float(perm_p_key),
            "null_q95": float(null_q95_key),
            "significant": bool(rho_key > null_q95_key),
        },
        "stage_trajectory_results": traj_results,
    }

    with open(OUT / "CSB_TRO_5_28_multistage_accessibility_summary.json", "w") as f:
        json.dump(summary, f, indent=2, ensure_ascii=False)
    print(f"\nSaved summary to {OUT}/CSB_TRO_5_28_multistage_accessibility_summary.json")

    print("\n" + "=" * 70)
    print("KEY RESULTS")
    print("=" * 70)
    print(f"delta_acc (morula-8cell) ~ strict_correction:")
    print(f"  rho={rho_delta:.4f}, perm_p={perm_p:.4f}, sig={rho_delta > null_q95}")
    print(f"delta_acc ~ delta_meth (morula-8cell):")
    print(f"  rho={rho_key:.4f}, perm_p={perm_p_key:.4f}, sig={rho_key > null_q95_key}")
    if np.isfinite(improvement):
        print(f"LOO-CV improvement with delta_acc u_bio:")
        print(f"  {improvement:.2f}%, perm_p={perm_p_impr:.4f}, sig={improvement > null_q95_impr}")
    print("=" * 70)

    return summary


if __name__ == "__main__":
    main()
```
