## Supplemental code for "An entropy-based diagnostic framework for characterizing methylation state dynamics during preimplantation development": 5_28_progress__solve_bottleneck2_redesigned.py

```
#!/usr/bin/env python
"""
Bottleneck 2 - Redesigned.

The first attempt trained the u_bio coefficient on non-morula transitions,
which is wrong: morula is a geometric vertex with different dynamics from
all other transitions. The correct design is:

1. Use morula-specific residual structure directly.
2. Test whether quantitative morula accessibility EXPLAINS the residual
   (i.e., regress c_diag ~ u_bio_continuous).
3. Then test whether u_bio_continuous, when added to the 8-cell prediction,
   moves the prediction toward observed morula in the residual DMRs specifically.
4. Bootstrap to establish whether the improvement is above chance.

Key insight: the question is not "does a global operator generalize"
but "does adding u_bio reduce the specific per-DMR error at top-residual DMRs?"
"""

from __future__ import annotations
import json
from pathlib import Path

import numpy as np
import pandas as pd
from scipy import stats

OUT = Path("E:/5_28_progress")
OUT.mkdir(parents=True, exist_ok=True)

DMR_RESIDUAL = Path(
    "E:/CSB_TRO_operator_time_DMR_dynamics_2026-05-25/results/CSB_TRO_basin_residual_DMR_ranking.tsv"
)
DMR_TRAJ = Path(
    "E:/CSB_TRO_operator_time_DMR_dynamics_2026-05-25/results/CSB_TRO_DMR_stage_mean_trajectory.tsv"
)
FORWARD_PRED = Path(
    "E:/CSB_TRO_operator_time_DMR_dynamics_2026-05-25/results/CSB_TRO_forward_prediction_morula.tsv"
)
ACC_TABLE = OUT / "CSB_TRO_5_28_dmr_quantitative_accessibility.tsv"
CURV_TABLE = Path("E:/实验进展5_27/CSB_TRO_2026-05-27_entry_exit_curvature.tsv")
MODULE_TABLE = Path(
    "E:/CSB_TRO_operator_time_DMR_dynamics_2026-05-25/results/CSB_TRO_DMR_module_assignments.tsv"
)

SEED = 42
N_BOOT = 2000


def main():
    print("=" * 70)
    print("Bottleneck 2 - Redesigned: morula-specific residual vs u_bio")
    print("=" * 70)

    # ── Load tables ────────────────────────────────────────────────────────────
    residual = pd.read_csv(DMR_RESIDUAL, sep="\t")
    traj = pd.read_csv(DMR_TRAJ, sep="\t")
    acc = pd.read_csv(ACC_TABLE, sep="\t")
    curv = pd.read_csv(CURV_TABLE, sep="\t") if CURV_TABLE.exists() else None

    # Stage means
    stage_means = {}
    for stage, g in traj.groupby("stage"):
        stage_means[stage] = g.set_index("cluster_name")["mean_beta"].to_dict()

    clusters = residual["cluster_name"].tolist()

    # Observed morula beta
    y_morula = np.array([stage_means["morula"].get(c, np.nan) for c in clusters])
    # 8-cell beta
    x_8cell = np.array([stage_means["8-cell"].get(c, np.nan) for c in clusters])
    # Methylation-only predicted morula (from pre-computed forward prediction)
    # Use mean predicted beta per DMR from the forward prediction file
    if FORWARD_PRED.exists():
        fp = pd.read_csv(FORWARD_PRED, sep="\t")
        fp_lmo = fp[fp["prediction_label"] == "leave_morula_out"]
        pred_mean = fp_lmo.groupby("cluster_name")["predicted_beta"].mean()
        y_pred_meth = np.array([pred_mean.get(c, np.nan) for c in clusters])
    else:
        # Fallback: use 8-cell as prediction
        y_pred_meth = x_8cell.copy()

    # c_diag = observed - methylation-only predicted (the diagnostic residual)
    c_diag = y_morula - y_pred_meth

    # Residual rank from basin analysis (rank 1 = largest residual)
    rank_map = residual.set_index("cluster_name")["basin_residual_rank"].to_dict()
    abs_res_map = residual.set_index("cluster_name")["abs_latent_residual_delta_beta"].to_dict()

    # Quantitative accessibility
    acc_map = acc.set_index("cluster_name")["morula_acc_mean"].to_dict()
    delta_map = acc.set_index("cluster_name")["morula_minus_8cell_mean"].to_dict()
    max_map = acc.set_index("cluster_name")["morula_acc_max"].to_dict()

    u_morula = np.array([acc_map.get(c, np.nan) for c in clusters])
    u_delta = np.array([delta_map.get(c, np.nan) for c in clusters])
    u_max = np.array([max_map.get(c, np.nan) for c in clusters])
    abs_res = np.array([abs_res_map.get(c, np.nan) for c in clusters])
    res_rank = np.array([rank_map.get(c, np.nan) for c in clusters])

    print(f"\nDMRs loaded: {len(clusters)}")
    print(f"DMRs with u_morula signal: {np.isfinite(u_morula).sum()}")
    print(f"DMRs with c_diag signal: {np.isfinite(c_diag).sum()}")

    # ── TEST 1: Does u_bio (morula accessibility) explain c_diag? ─────────────
    # c_diag = beta * u_morula + epsilon
    # This is the direct diagnostic question: does ATAC explain the residual?
    print("\n--- TEST 1: c_diag ~ u_morula_acc (all DMRs) ---")
    valid = np.isfinite(c_diag) & np.isfinite(u_morula)
    if valid.sum() >= 5:
        rho_all, p_all = stats.spearmanr(u_morula[valid], c_diag[valid])
        r_all, pr_all = stats.pearsonr(u_morula[valid], c_diag[valid])
        print(f"  Spearman rho: {rho_all:.4f}, p={p_all:.4f}")
        print(f"  Pearson r: {r_all:.4f}, p={pr_all:.4f}")
    else:
        rho_all = p_all = r_all = pr_all = np.nan

    # Same test for delta (morula - 8cell accessibility)
    valid_d = np.isfinite(c_diag) & np.isfinite(u_delta)
    if valid_d.sum() >= 5:
        rho_delta, p_delta = stats.spearmanr(u_delta[valid_d], c_diag[valid_d])
        print(f"  Delta (morula-8cell) Spearman rho: {rho_delta:.4f}, p={p_delta:.4f}")
    else:
        rho_delta = p_delta = np.nan

    # ── TEST 2: Top-k residual DMRs – c_diag vs u_morula ─────────────────────
    print("\n--- TEST 2: c_diag ~ u_morula_acc (top-k residual DMRs) ---")
    topk_results = {}
    for k in [25, 50, 100]:
        top_mask = res_rank <= k
        valid_k = top_mask & np.isfinite(c_diag) & np.isfinite(u_morula)
        if valid_k.sum() >= 5:
            rho_k, p_k = stats.spearmanr(u_morula[valid_k], c_diag[valid_k])
            print(f"  Top{k}: Spearman rho={rho_k:.4f}, p={p_k:.4f}, n={valid_k.sum()}")
            topk_results[f"top{k}"] = {"rho": float(rho_k), "p": float(p_k), "n": int(valid_k.sum())}
        else:
            topk_results[f"top{k}"] = {"rho": np.nan, "p": np.nan, "n": int(valid_k.sum())}

    # ── TEST 3: Signed residual direction test ────────────────────────────────
    # The residual in top DMRs has a specific sign (signed_latent_residual_direction = -1)
    # meaning observed morula < predicted (methylation too high, needs to go down)
    # Does accessibility direction align with this correction direction?
    print("\n--- TEST 3: Signed alignment test ---")
    sign_map = residual.set_index("cluster_name")["signed_latent_residual_direction"].to_dict()
    sign_dir = np.array([sign_map.get(c, np.nan) for c in clusters])

    # For top25 residual DMRs: does higher morula accessibility correlate with
    # the NEEDED direction of correction?
    top25_mask = res_rank <= 25
    valid_sign = top25_mask & np.isfinite(u_morula) & np.isfinite(sign_dir)
    if valid_sign.sum() >= 5:
        # When sign_dir == -1, c_diag is negative (need to decrease methylation)
        # morula accessibility should be higher where c_diag is more negative
        rho_sign, p_sign = stats.spearmanr(u_morula[valid_sign], sign_dir[valid_sign])
        print(f"  Top25 u_morula vs sign_dir: Spearman rho={rho_sign:.4f}, p={p_sign:.4f}")
    else:
        rho_sign = p_sign = np.nan

    # ── TEST 4: Bootstrap – does u_morula explain residual better than chance? ─
    print("\n--- TEST 4: Bootstrap permutation test ---")
    rng = np.random.default_rng(SEED)
    valid_all = np.isfinite(c_diag) & np.isfinite(u_morula)
    obs_rho = rho_all if np.isfinite(rho_all) else 0.0

    null_rhos = []
    u_obs = u_morula[valid_all]
    c_obs = c_diag[valid_all]
    for _ in range(N_BOOT):
        perm = rng.permutation(len(u_obs))
        r, _ = stats.spearmanr(u_obs[perm], c_obs)
        null_rhos.append(r)
    null_rhos = np.array(null_rhos)
    perm_p = float((np.abs(null_rhos) >= np.abs(obs_rho)).mean())
    null_q95 = float(np.quantile(null_rhos, 0.95))
    null_q05 = float(np.quantile(null_rhos, 0.05))
    print(f"  Observed rho: {obs_rho:.4f}")
    print(f"  Null q05-q95: {null_q05:.4f} to {null_q95:.4f}")
    print(f"  Permutation p (two-sided): {perm_p:.4f}")
    print(f"  Observed > null q95: {obs_rho > null_q95}")

    # ── TEST 5: Prediction improvement – add u_bio to morula-specific model ───
    # Key redesign: instead of using non-morula transitions for training,
    # use cross-validated leave-one-DMR-out within the morula-specific structure.
    # Model: y_morula_j = a * x_8cell_j + b * u_morula_j + c
    # Train on all other DMRs, predict DMR j.
    print("\n--- TEST 5: Leave-one-DMR-out RMSE with u_bio ---")
    valid_all2 = np.isfinite(y_morula) & np.isfinite(x_8cell) & np.isfinite(u_morula)
    idx_valid = np.where(valid_all2)[0]

    if len(idx_valid) >= 10:
        # Leave-one-DMR-out CV
        lam = 0.01
        errors_meth = []
        errors_bio = []

        for i in idx_valid:
            train_idx = idx_valid[idx_valid != i]

            # Meth-only model on training DMRs
            X_train_m = np.column_stack([x_8cell[train_idx], np.ones(len(train_idx))])
            y_train = y_morula[train_idx]
            reg = np.eye(2) * lam
            reg[1, 1] = 0
            try:
                coef_m = np.linalg.solve(X_train_m.T @ X_train_m + reg, X_train_m.T @ y_train)
            except np.linalg.LinAlgError:
                continue
            pred_m = coef_m[0] * x_8cell[i] + coef_m[1]
            errors_meth.append(abs(y_morula[i] - pred_m))

            # Bio model (meth + u_morula)
            u_train = u_morula[train_idx]
            X_train_b = np.column_stack([x_8cell[train_idx], u_train, np.ones(len(train_idx))])
            reg3 = np.eye(3) * lam
            reg3[2, 2] = 0
            try:
                coef_b = np.linalg.solve(X_train_b.T @ X_train_b + reg3, X_train_b.T @ y_train)
            except np.linalg.LinAlgError:
                continue
            pred_b = coef_b[0] * x_8cell[i] + coef_b[1] * u_morula[i] + coef_b[2]
            errors_bio.append(abs(y_morula[i] - pred_b))

        rmse_loocv_meth = float(np.sqrt(np.mean(np.array(errors_meth) ** 2)))
        rmse_loocv_bio = float(np.sqrt(np.mean(np.array(errors_bio) ** 2)))
        print(f"  Leave-one-DMR-out RMSE (meth only): {rmse_loocv_meth:.4f}")
        print(f"  Leave-one-DMR-out RMSE (meth + u_morula): {rmse_loocv_bio:.4f}")
        print(f"  Improvement: {(rmse_loocv_meth - rmse_loocv_bio) / rmse_loocv_meth * 100:.2f}%")

        # Same for top25 residual DMRs
        top25_valid_idx = idx_valid[res_rank[idx_valid] <= 25]
        if len(top25_valid_idx) >= 5:
            err_m25, err_b25 = [], []
            for i in top25_valid_idx:
                train_idx = idx_valid[idx_valid != i]
                X_train_m = np.column_stack([x_8cell[train_idx], np.ones(len(train_idx))])
                y_train = y_morula[train_idx]
                try:
                    coef_m = np.linalg.solve(X_train_m.T @ X_train_m + reg, X_train_m.T @ y_train)
                    pred_m = coef_m[0] * x_8cell[i] + coef_m[1]
                    err_m25.append(abs(y_morula[i] - pred_m))

                    u_train = u_morula[train_idx]
                    X_train_b = np.column_stack([x_8cell[train_idx], u_train, np.ones(len(train_idx))])
                    coef_b = np.linalg.solve(X_train_b.T @ X_train_b + reg3, X_train_b.T @ y_train)
                    pred_b = coef_b[0] * x_8cell[i] + coef_b[1] * u_morula[i] + coef_b[2]
                    err_b25.append(abs(y_morula[i] - pred_b))
                except np.linalg.LinAlgError:
                    continue

            rmse_top25_meth = float(np.sqrt(np.mean(np.array(err_m25) ** 2)))
            rmse_top25_bio = float(np.sqrt(np.mean(np.array(err_b25) ** 2)))
            print(f"\n  TOP25 RESIDUAL DMRs (leave-one-DMR-out):")
            print(f"    RMSE meth-only: {rmse_top25_meth:.4f}")
            print(f"    RMSE bio model: {rmse_top25_bio:.4f}")
            print(f"    Improvement: {(rmse_top25_meth - rmse_top25_bio) / rmse_top25_meth * 100:.2f}%")
        else:
            rmse_top25_meth = rmse_top25_bio = np.nan
    else:
        rmse_loocv_meth = rmse_loocv_bio = np.nan
        rmse_top25_meth = rmse_top25_bio = np.nan

    # ── TEST 6: Curvature-stratified correlation ───────────────────────────────
    print("\n--- TEST 6: Curvature-stratified c_diag ~ u_morula ---")
    curv_results = {}
    if curv is not None:
        curv_map = curv.set_index("cluster_name")
        for group_name, group_clusters in [
            ("negative_curvature", [c for c in clusters
                                     if c in curv_map.index and curv_map.loc[c, "curvature"] < 0]),
            ("positive_curvature", [c for c in clusters
                                     if c in curv_map.index and curv_map.loc[c, "curvature"] >= 0]),
            ("inverted_u", [c for c in clusters
                             if c in curv_map.index and curv_map.loc[c, "is_inverted_u"]]),
            ("u_shape", [c for c in clusters
                          if c in curv_map.index and curv_map.loc[c, "is_u_shape"]]),
        ]:
            gc_idx = [i for i, c in enumerate(clusters) if c in set(group_clusters)]
            if len(gc_idx) < 5:
                continue
            u_g = u_morula[gc_idx]
            c_g = c_diag[gc_idx]
            valid_g = np.isfinite(u_g) & np.isfinite(c_g)
            if valid_g.sum() < 5:
                continue
            rho_g, p_g = stats.spearmanr(u_g[valid_g], c_g[valid_g])
            print(f"  {group_name} (n={valid_g.sum()}): rho={rho_g:.4f}, p={p_g:.4f}")
            curv_results[group_name] = {"rho": float(rho_g), "p": float(p_g), "n": int(valid_g.sum())}

    # ── Compile all results ────────────────────────────────────────────────────
    summary = {
        "date": "2026-05-28",
        "test1_c_diag_vs_u_morula_all": {
            "spearman_rho": float(rho_all) if np.isfinite(rho_all) else None,
            "spearman_p": float(p_all) if np.isfinite(p_all) else None,
            "pearson_r": float(r_all) if np.isfinite(r_all) else None,
            "pearson_p": float(pr_all) if np.isfinite(pr_all) else None,
        },
        "test1_c_diag_vs_u_delta": {
            "spearman_rho": float(rho_delta) if np.isfinite(rho_delta) else None,
            "spearman_p": float(p_delta) if np.isfinite(p_delta) else None,
        },
        "test2_topk_c_diag_vs_u_morula": topk_results,
        "test3_signed_alignment_top25": {
            "rho": float(rho_sign) if np.isfinite(rho_sign) else None,
            "p": float(p_sign) if np.isfinite(p_sign) else None,
        },
        "test4_permutation_test": {
            "observed_rho": float(obs_rho),
            "null_q05": float(null_q05),
            "null_q95": float(null_q95),
            "permutation_p_two_sided": float(perm_p),
            "observed_gt_null_q95": bool(obs_rho > null_q95),
            "n_permutations": N_BOOT,
        },
        "test5_loocv_rmse": {
            "all_dmr_meth_only": float(rmse_loocv_meth) if np.isfinite(rmse_loocv_meth) else None,
            "all_dmr_bio_model": float(rmse_loocv_bio) if np.isfinite(rmse_loocv_bio) else None,
            "top25_meth_only": float(rmse_top25_meth) if np.isfinite(rmse_top25_meth) else None,
            "top25_bio_model": float(rmse_top25_bio) if np.isfinite(rmse_top25_bio) else None,
        },
        "test6_curvature_stratified": curv_results,
        "interpretation": {
            "primary_finding": (
                "Quantitative morula accessibility (Liu2019) shows Spearman rho correlation "
                "with the diagnostic correction c_diag. The direction and significance of this "
                "correlation (see test1/test2/test4) determines whether u_bio reduces the "
                "residual in a structured way."
            ),
            "level_4_entry_criterion": (
                "Entry to Level 4 requires: (1) significant c_diag ~ u_morula correlation "
                "in top-residual DMRs, OR (2) leave-one-DMR-out RMSE improvement with bio model. "
                "Results: see test2 and test5."
            ),
        },
    }

    out_path = OUT / "CSB_TRO_5_28_bottleneck2_redesigned_summary.json"
    with open(out_path, "w") as f:
        json.dump(summary, f, indent=2)
    print(f"\nSaved: {out_path}")

    # Save per-DMR table
    per_dmr_rows = []
    for i, c in enumerate(clusters):
        per_dmr_rows.append({
            "cluster_name": c,
            "y_morula_observed": float(y_morula[i]) if np.isfinite(y_morula[i]) else None,
            "x_8cell": float(x_8cell[i]) if np.isfinite(x_8cell[i]) else None,
            "y_pred_meth_only": float(y_pred_meth[i]) if np.isfinite(y_pred_meth[i]) else None,
            "c_diag": float(c_diag[i]) if np.isfinite(c_diag[i]) else None,
            "u_morula_acc_mean": float(u_morula[i]) if np.isfinite(u_morula[i]) else None,
            "u_morula_minus_8cell": float(u_delta[i]) if np.isfinite(u_delta[i]) else None,
            "basin_residual_rank": float(res_rank[i]) if np.isfinite(res_rank[i]) else None,
            "abs_residual": float(abs_res[i]) if np.isfinite(abs_res[i]) else None,
        })

    pd.DataFrame(per_dmr_rows).to_csv(
        OUT / "CSB_TRO_5_28_per_dmr_cdiag_vs_ubio.tsv", sep="\t", index=False
    )
    print(f"Saved: {OUT}/CSB_TRO_5_28_per_dmr_cdiag_vs_ubio.tsv")
    return summary


if __name__ == "__main__":
    main()
```
