## Supplemental code for "An entropy-based diagnostic framework for characterizing methylation state dynamics during preimplantation development": 5_29_progress__complete_model_implementation.py

```
#!/usr/bin/env python
"""
Morula-centered gated operator-control dynamics
Complete implementation of all 5 steps:

Step 1: Unified threshold theta=0.02, standard definition table
Step 2: Exit two-part model (logistic re-meth predictor)
Step 3: Module-level impulse J_M,k
Step 4: Minimum control energy E_entry,k and E_exit,k
Step 5: Counterfactual simulator (4 scenarios)
"""
from __future__ import annotations
import json
from pathlib import Path
import numpy as np
import pandas as pd
from scipy import stats
from scipy.special import expit
from numpy.linalg import solve, norm
from sklearn.metrics import roc_auc_score, roc_curve
from sklearn.linear_model import LogisticRegression

OUT = Path("E:/5_29_progress")
OUT.mkdir(parents=True, exist_ok=True)

TRAJ  = Path("E:/CSB_TRO_operator_time_DMR_dynamics_2026-05-25/results/CSB_TRO_DMR_stage_mean_trajectory.tsv")
RESID = Path("E:/CSB_TRO_operator_time_DMR_dynamics_2026-05-25/results/CSB_TRO_basin_residual_DMR_ranking.tsv")
META  = Path("E:/CSB_TRO_operator_time_DMR_dynamics_2026-05-25/results/CSB_TRO_DMR_metadata.tsv")
MS    = Path("E:/5_28_progress/CSB_TRO_5_28_dmr_multistage_accessibility.tsv")
GREEDY= Path("E:/CSB_TRO_operator_time_DMR_dynamics_2026-05-25/results/CSB_TRO_missing_control_term_greedy_modules.tsv")
HIST  = Path("E:/CSB_TRO_operator_time_DMR_dynamics_2026-05-25/external/histone")

SEED  = 42
N_BOOT = 2000
rng   = np.random.default_rng(SEED)

# ── Load core data ─────────────────────────────────────────────────────────────
traj  = pd.read_csv(TRAJ, sep="\t")
resid = pd.read_csv(RESID, sep="\t")
meta  = pd.read_csv(META, sep="\t")
ms    = pd.read_csv(MS, sep="\t")
greedy= pd.read_csv(GREEDY, sep="\t")

stage_means = {}
for s, g in traj.groupby("stage"):
    stage_means[s] = g.set_index("cluster_name")["mean_beta"].to_dict()

clusters  = sorted(resid["cluster_name"].tolist())
stages_ord= ["MII oocyte","zygote/PN","2-cell","4-cell","8-cell","morula","blastocyst"]

def svec(stage):
    return np.array([stage_means.get(stage,{}).get(c, np.nan) for c in clusters])

x_8cell   = svec("8-cell")
x_morula  = svec("morula")
x_blast   = svec("blastocyst")
x_4cell   = svec("4-cell")

meta_map  = meta.set_index("cluster_name")
ms_map    = ms.set_index("cluster_name")
mod_map   = resid.set_index("cluster_name")["module_id"].to_dict()
rank_map  = resid.set_index("cluster_name")["basin_residual_rank"].to_dict()

n_cpg = np.array([meta_map.loc[c,"n_cpg_target"] if c in meta_map.index else np.nan for c in clusters])
width = np.array([meta_map.loc[c,"width"] if c in meta_map.index else np.nan for c in clusters])
acc_morula= np.array([ms_map.loc[c,"acc_morula_mean"] if c in ms_map.index else np.nan for c in clusters])
acc_8cell = np.array([ms_map.loc[c,"acc_8-cell_mean"] if c in ms_map.index else np.nan for c in clusters])


# ══════════════════════════════════════════════════════════════════════════════
# STEP 1: Unified threshold and standard definition table
# ══════════════════════════════════════════════════════════════════════════════
print("="*65)
print("STEP 1: Unified threshold theta=0.02, standard definitions")
print("="*65)

THETA_MO    = 0.02   # morula-zero threshold
THETA_REMETH= 0.05   # re-methylation threshold (blast)
THETA_ENTRY = 0.875  # observed morula basin occupancy

# Standard definitions
is_mzero   = (x_morula <= THETA_MO) & np.isfinite(x_blast)
is_remeth  = is_mzero & (x_blast > THETA_REMETH)
is_stay    = is_mzero & (x_blast <= THETA_REMETH)
is_priority= np.array([mod_map.get(c,"?") in ["M01","M02","M05","M10","M12"] for c in clusters])

# Entry operator
# Train on non-morula transitions
train_pairs = [("MII oocyte","zygote/PN"),("zygote/PN","2-cell"),
               ("2-cell","4-cell"),("4-cell","8-cell")]
X_tr, Y_tr = [], []
for sf, st in train_pairs:
    xf = svec(sf); xt = svec(st)
    v = np.isfinite(xf) & np.isfinite(xt)
    X_tr.append(xf[v].reshape(-1,1)); Y_tr.append(xt[v])
X_tr = np.vstack(X_tr); Y_tr = np.concatenate(Y_tr)
A_aug = np.column_stack([X_tr, np.ones(len(X_tr))])
coef, _, _, _ = np.linalg.lstsq(A_aug, Y_tr, rcond=None)
alpha_op, bias_op = coef[0], coef[1]

# Blastocyst operator (morula as input)
x_blast_pred = alpha_op * x_morula + bias_op
c_diag_entry = x_morula - (alpha_op * x_8cell + bias_op)  # 8cell->morula correction
c_diag_exit  = x_blast - x_blast_pred                      # morula->blast correction

print(f"\nEntry operator: y = {alpha_op:.4f}*x + {bias_op:.4f}")
print(f"Exit operator:  y = {alpha_op:.4f}*x + {bias_op:.4f} (same structure)")
print(f"\nStandard definitions (theta_mo={THETA_MO}, theta_remeth={THETA_REMETH}):")
print(f"  morula-zero DMRs (x_M <= {THETA_MO}): {is_mzero.sum()}/156")
print(f"  re-methylation DMRs: {is_remeth.sum()} ({is_remeth.sum()/is_mzero.sum():.1%} of mzero)")
print(f"  stay-zero DMRs: {is_stay.sum()} ({is_stay.sum()/is_mzero.sum():.1%} of mzero)")

# Histone signal loader
def load_overlap(fname):
    df = pd.read_csv(HIST/fname, sep="\t", header=None, compression="gzip",
                     names=["chr","start","end","name","score"], usecols=[0,1,2,3,4])
    df["chr"] = df["chr"].astype(str).str.strip()
    by_chr = {c:g for c,g in df.groupby("chr")}
    ov, sc = [], []
    for c in clusters:
        if c not in meta_map.index: ov.append(0); sc.append(np.nan); continue
        chrom=str(meta_map.loc[c,"chr"]).strip(); ds=int(meta_map.loc[c,"start"]); de=int(meta_map.loc[c,"end"])
        if chrom not in by_chr: ov.append(0); sc.append(np.nan); continue
        sub=by_chr[chrom]; hits=sub[(sub["start"]<de)&(sub["end"]>ds)]
        sv=pd.to_numeric(hits["score"],errors="coerce").dropna()
        ov.append(int(len(hits)>0)); sc.append(float(sv.max()) if len(sv)>0 else np.nan)
    return np.array(ov), np.array(sc)

k4me3_8cell_ov, k4me3_8cell_sc = load_overlap("H3K4me3_8cell.hg19.bed.gz")
k27me3_blast_ov, k27me3_blast_sc = load_overlap("H3K27me3_blastocyst.hg19.bed.gz")
k27ac_blast_ov, k27ac_blast_sc  = load_overlap("H3K27ac_blastocyst.hg19.bed.gz")
k4me3_blast_ov, k4me3_blast_sc  = load_overlap("H3K4me3_blastocyst.hg19.bed.gz")
k27ac_8cell_ov, k27ac_8cell_sc  = load_overlap("H3K27ac_8cell.hg19.bed.gz")

is_m00 = np.array([mod_map.get(c,"?") == "M00" for c in clusters])

# Standard definition table
def_table = pd.DataFrame({
    "cluster_name": clusters,
    "x_8cell": x_8cell, "x_morula": x_morula, "x_blast": x_blast,
    "x_blast_pred": x_blast_pred,
    "c_diag_entry": c_diag_entry, "c_diag_exit": c_diag_exit,
    "is_mzero": is_mzero.astype(int), "is_remeth": is_remeth.astype(int),
    "is_stay": is_stay.astype(int),
    "module_id": [mod_map.get(c,"?") for c in clusters],
    "is_M00": is_m00.astype(int),
    "is_priority": is_priority.astype(int),
    "n_cpg": n_cpg, "width": width,
    "acc_morula": acc_morula, "acc_8cell": acc_8cell,
    "k4me3_8cell_ov": k4me3_8cell_ov, "k4me3_8cell_sc": k4me3_8cell_sc,
    "k27me3_blast_ov": k27me3_blast_ov, "k4me3_blast_ov": k4me3_blast_ov,
    "k27ac_blast_ov": k27ac_blast_ov, "k27ac_8cell_ov": k27ac_8cell_ov,
    "basin_residual_rank": [rank_map.get(c,np.nan) for c in clusters],
})
def_table.to_csv(OUT/"step1_standard_definitions.tsv", sep="\t", index=False)
print(f"\nSaved: step1_standard_definitions.tsv")


# ══════════════════════════════════════════════════════════════════════════════
# STEP 2: Exit two-part model
# ══════════════════════════════════════════════════════════════════════════════
print("\n" + "="*65)
print("STEP 2: Exit two-part model")
print("="*65)

# Part 1 (classifier): P(y_re=1) = sigma(H3K4me3_8cell + M00 + CpG + width)
# Among morula-zero DMRs

mzero_idx = np.where(is_mzero)[0]
y_bin = is_remeth[mzero_idx].astype(float)

# Features for morula-zero DMRs
def get_features(idx_set):
    k4sc = k4me3_8cell_sc[idx_set]
    k4sc_filled = np.where(np.isfinite(k4sc), k4sc, 0.0)
    m00 = is_m00[idx_set].astype(float)
    cpg = n_cpg[idx_set]; cpg_filled = np.where(np.isfinite(cpg), cpg, np.nanmean(n_cpg))
    w   = width[idx_set]; w_filled   = np.where(np.isfinite(w), w, np.nanmean(width))
    k27me3_b = k27me3_blast_ov[idx_set].astype(float)
    acc_mo = acc_morula[idx_set]; acc_filled = np.where(np.isfinite(acc_mo), acc_mo, 0.0)
    return np.column_stack([k4sc_filled, m00, cpg_filled, w_filled, k27me3_b, acc_filled])

feat_names = ["k4me3_8cell_score","M00","n_cpg","width","k27me3_blast_ov","acc_morula"]
X_feat = get_features(mzero_idx)

print(f"\nFeatures: {feat_names}")
print(f"n={len(y_bin)}, remeth={int(y_bin.sum())}, stay={int((y_bin==0).sum())}")

# Standardize continuous features
X_scaled = X_feat.copy()
feat_means = X_scaled.mean(axis=0)
feat_stds  = X_scaled.std(axis=0) + 1e-8
X_scaled = (X_scaled - feat_means) / feat_stds

# Logistic regression with L2
from sklearn.linear_model import LogisticRegressionCV
lr = LogisticRegressionCV(cv=5, random_state=SEED, max_iter=1000, scoring="roc_auc")
try:
    lr.fit(X_scaled, y_bin)
    proba = lr.predict_proba(X_scaled)[:,1]
    auc_lr = float(roc_auc_score(y_bin, proba))
    print(f"\nLogistic regression (5-fold CV): AUC={auc_lr:.4f}")
    print(f"Coefficients:")
    for name, coef in zip(feat_names, lr.coef_[0]):
        print(f"  {name}: {coef:.4f}")
except Exception as e:
    print(f"LogisticRegressionCV error: {e}")
    # Fallback: simple logistic with L2
    from sklearn.linear_model import LogisticRegression as LR
    lr = LR(C=1.0, random_state=SEED, max_iter=1000)
    lr.fit(X_scaled, y_bin)
    proba = lr.predict_proba(X_scaled)[:,1]
    auc_lr = float(roc_auc_score(y_bin, proba))
    print(f"Logistic regression fallback: AUC={auc_lr:.4f}")

# LOOCV AUC
from sklearn.model_selection import LeaveOneOut
loo_probs = np.zeros(len(y_bin))
loo = LeaveOneOut()
for train_idx, test_idx in loo.split(X_scaled):
    try:
        lr_loo = LogisticRegression(C=1.0, random_state=SEED, max_iter=500)
        lr_loo.fit(X_scaled[train_idx], y_bin[train_idx])
        loo_probs[test_idx] = lr_loo.predict_proba(X_scaled[test_idx])[:,1]
    except:
        loo_probs[test_idx] = 0.5
try:
    auc_loocv = float(roc_auc_score(y_bin, loo_probs))
except:
    auc_loocv = np.nan
print(f"LOOCV AUC: {auc_loocv:.4f}")

# Permutation test
null_aucs = []
for _ in range(N_BOOT):
    y_perm = rng.permutation(y_bin)
    try:
        lr_perm = LogisticRegression(C=1.0, random_state=SEED, max_iter=500)
        lr_perm.fit(X_scaled, y_perm)
        p_perm = lr_perm.predict_proba(X_scaled)[:,1]
        null_aucs.append(float(roc_auc_score(y_perm, p_perm)))
    except:
        null_aucs.append(0.5)
null_aucs = np.array(null_aucs)
perm_p_lr = float((null_aucs >= auc_lr).mean())
q95_lr    = float(np.quantile(null_aucs, 0.95))
print(f"Permutation test: perm_p={perm_p_lr:.4f}, null q95={q95_lr:.4f}, sig={auc_lr>q95_lr}")

# Leave-feature-out
print("\nLeave-feature-out AUC:")
feat_importance = {}
for i, fname in enumerate(feat_names):
    X_loo_feat = np.delete(X_scaled, i, axis=1)
    try:
        lr_lfo = LogisticRegression(C=1.0, random_state=SEED, max_iter=500)
        lr_lfo.fit(X_loo_feat, y_bin)
        p_lfo = lr_lfo.predict_proba(X_loo_feat)[:,1]
        auc_lfo = float(roc_auc_score(y_bin, p_lfo))
        drop = auc_lr - auc_lfo
        feat_importance[fname] = {"auc_without": auc_lfo, "drop": drop}
        print(f"  -{fname}: AUC={auc_lfo:.4f} (drop={drop:.4f})")
    except:
        pass

# Leave-module-out (M00)
m00_mask = is_m00[mzero_idx].astype(bool)
if m00_mask.sum() > 3 and (~m00_mask).sum() > 5:
    X_nom00 = X_scaled[~m00_mask]; y_nom00 = y_bin[~m00_mask]
    try:
        lr_nom00 = LogisticRegression(C=1.0, random_state=SEED, max_iter=500)
        lr_nom00.fit(X_nom00, y_nom00)
        p_test = lr_nom00.predict_proba(X_scaled[m00_mask])[:,1]
        # Apply to M00 DMRs
        if y_bin[m00_mask].sum() > 0 and (y_bin[m00_mask]==0).sum() > 0:
            auc_m00 = float(roc_auc_score(y_bin[m00_mask], p_test))
        else:
            auc_m00 = np.nan
        print(f"\nLeave-M00-out: trained on non-M00, tested on M00 AUC={auc_m00:.4f}")
    except Exception as e:
        print(f"Leave-M00-out: {e}")

# Odds ratios
print("\nOdds ratios (exp(coef)):")
for name, coef in zip(feat_names, lr.coef_[0]):
    print(f"  {name}: OR={np.exp(coef):.3f}")

# Calibration: compare predicted prob vs observed rate
bins = np.linspace(0,1,6)
print("\nCalibration (predicted prob vs observed rate):")
for i in range(len(bins)-1):
    mask = (proba >= bins[i]) & (proba < bins[i+1])
    if mask.sum() == 0: continue
    obs_rate = float(y_bin[mask].mean())
    print(f"  p=[{bins[i]:.1f},{bins[i+1]:.1f}]: n={mask.sum()}, pred_mean={proba[mask].mean():.3f}, obs={obs_rate:.3f}")

# Save predictions
pred_df = pd.DataFrame({
    "cluster_name": [clusters[i] for i in mzero_idx],
    "x_morula": x_morula[mzero_idx], "x_blast": x_blast[mzero_idx],
    "y_remeth": y_bin, "pred_prob_remeth": proba,
    "module_id": [mod_map.get(clusters[i],"?") for i in mzero_idx],
})
pred_df.to_csv(OUT/"step2_exit_model_predictions.tsv", sep="\t", index=False)

step2_results = {
    "auc_train": float(auc_lr), "auc_loocv": float(auc_loocv),
    "perm_p": float(perm_p_lr), "null_q95": float(q95_lr),
    "significant": bool(auc_lr > q95_lr),
    "feature_importance": feat_importance,
    "odds_ratios": {n: float(np.exp(c)) for n, c in zip(feat_names, lr.coef_[0])},
}
print(f"\nStep 2 complete: AUC={auc_lr:.3f}, LOOCV={auc_loocv:.3f}, perm_p={perm_p_lr:.3f}")


# ══════════════════════════════════════════════════════════════════════════════
# STEP 3: Module-level impulse J_M,k
# ══════════════════════════════════════════════════════════════════════════════
print("\n" + "="*65)
print("STEP 3: Module-level impulse J_M,k")
print("="*65)

# J_M,k = ||c_diag restricted to module k||_2 (correction norm per module)
# This measures how much each module contributes to the transition

modules_all = sorted(set(mod_map.values()) - {"?"})

# Entry impulse (8cell->morula correction)
# Exit impulse (morula->blast correction)
impulse_rows = []
for mid in modules_all:
    idx_m = [i for i,c in enumerate(clusters) if mod_map.get(c,"?")==mid]
    if not idx_m: continue

    # Entry: c_diag_entry at module k
    e_entry = c_diag_entry[idx_m]
    e_exit  = c_diag_exit[idx_m]

    j_entry = float(norm(e_entry[np.isfinite(e_entry)]))
    j_exit  = float(norm(e_exit[np.isfinite(e_exit)]))

    # Signed mean direction
    dir_entry = float(np.nanmean(e_entry))
    dir_exit  = float(np.nanmean(e_exit))

    # Fraction in re-meth class
    n_remeth_k = int(is_remeth[idx_m].sum())
    n_mzero_k  = int(is_mzero[idx_m].sum())

    # Cosine alignment: entry and exit impulse directions
    e_en = e_entry[np.isfinite(e_entry) & np.isfinite(e_exit)]
    e_ex = e_exit[np.isfinite(e_entry) & np.isfinite(e_exit)]
    cos_align = float(np.dot(e_en,e_ex)/(norm(e_en)*norm(e_ex)+1e-12)) if len(e_en)>0 else np.nan

    is_priority_m = mid in ["M01","M02","M05","M10","M12"]
    is_m00_m = mid == "M00"

    impulse_rows.append({
        "module": mid, "n_dmr": len(idx_m),
        "J_entry": j_entry, "J_exit": j_exit,
        "dir_entry": dir_entry, "dir_exit": dir_exit,
        "cos_align_entry_exit": cos_align,
        "n_remeth": n_remeth_k, "n_mzero": n_mzero_k,
        "is_priority": is_priority_m, "is_M00": is_m00_m,
        "impulse_ratio": j_exit/j_entry if j_entry>1e-6 else np.nan,
    })
    print(f"  {mid}: n={len(idx_m)}, J_entry={j_entry:.4f}, J_exit={j_exit:.4f}, "
          f"cos={cos_align:.3f}, remeth={n_remeth_k}/{n_mzero_k}")

impulse_df = pd.DataFrame(impulse_rows).sort_values("J_entry", ascending=False)
impulse_df.to_csv(OUT/"step3_module_impulse.tsv", sep="\t", index=False)
print(f"\nTop 5 by J_entry:")
print(impulse_df.nlargest(5,"J_entry")[["module","J_entry","J_exit","cos_align_entry_exit"]].to_string())
print(f"\nTop 5 by J_exit:")
print(impulse_df.nlargest(5,"J_exit")[["module","J_entry","J_exit","cos_align_entry_exit"]].to_string())


# ══════════════════════════════════════════════════════════════════════════════
# STEP 4: Minimum control energy
# ══════════════════════════════════════════════════════════════════════════════
print("\n" + "="*65)
print("STEP 4: Minimum control energy E_entry,k and E_exit,k")
print("="*65)

# Minimum control energy: how much external input is needed to drive the transition?
# E_k = min_u ||u||^2 s.t. x_target = K x_source + B u
# For the DMR-level model: E_entry,k = ||c_diag_entry[module_k]||^2 / ||x_8cell[module_k]||^2
# (normalized by the from-state energy)
# This gives a relative "effort" metric: how much correction relative to source energy

energy_rows = []
for mid in modules_all:
    idx_m = [i for i,c in enumerate(clusters) if mod_map.get(c,"?")==mid]
    if not idx_m: continue

    src_entry = x_8cell[idx_m]; tgt_entry = x_morula[idx_m]
    src_exit  = x_morula[idx_m]; tgt_exit  = x_blast[idx_m]
    corr_entry= c_diag_entry[idx_m]; corr_exit = c_diag_exit[idx_m]

    valid_en = np.isfinite(src_entry) & np.isfinite(corr_entry)
    valid_ex = np.isfinite(src_exit)  & np.isfinite(corr_exit)

    # Control energy = ||correction||^2 / ||source||^2
    src_norm_en = float(norm(src_entry[valid_en]))
    src_norm_ex = float(norm(src_exit[valid_ex]))

    e_entry = float(norm(corr_entry[valid_en])**2 / (src_norm_en**2 + 1e-12))
    e_exit  = float(norm(corr_exit[valid_ex])**2  / (src_norm_ex**2 + 1e-12))

    # Relative energy: E_exit / E_entry
    rel_energy = e_exit / (e_entry + 1e-12)

    # Also compute "control efficiency": ||correction||^2 / ||target - source||^2
    # (how much control is needed relative to total state change)
    delta_entry = tgt_entry[valid_en] - src_entry[valid_en]
    delta_exit  = tgt_exit[valid_ex]  - src_exit[valid_ex]
    eff_entry = float(norm(corr_entry[valid_en])**2 / (norm(delta_entry)**2 + 1e-12))
    eff_exit  = float(norm(corr_exit[valid_ex])**2  / (norm(delta_exit)**2  + 1e-12))

    energy_rows.append({
        "module": mid, "n_dmr": len(idx_m),
        "E_entry": e_entry, "E_exit": e_exit,
        "rel_energy_exit_over_entry": rel_energy,
        "eff_entry": eff_entry, "eff_exit": eff_exit,
        "is_priority": mid in ["M01","M02","M05","M10","M12"],
        "is_M00": mid == "M00",
    })

energy_df = pd.DataFrame(energy_rows).sort_values("E_entry", ascending=False)
energy_df.to_csv(OUT/"step4_control_energy.tsv", sep="\t", index=False)

print("\nControl energy table (top modules):")
priority_energy = energy_df[energy_df["is_priority"]|energy_df["is_M00"]]
print(priority_energy[["module","E_entry","E_exit","rel_energy_exit_over_entry","eff_entry","eff_exit"]].to_string())


# ══════════════════════════════════════════════════════════════════════════════
# STEP 5: Counterfactual simulator
# ══════════════════════════════════════════════════════════════════════════════
print("\n" + "="*65)
print("STEP 5: Counterfactual simulator (4 scenarios)")
print("="*65)

# Basin occupancy for entry
def entry_occupancy(x_pred, x_obs, alpha=0.9):
    valid = np.isfinite(x_pred) & np.isfinite(x_obs)
    dists = np.abs(x_pred[valid] - x_obs[valid])
    radius = np.nanquantile(np.abs(x_obs - np.nanmean(x_obs)), alpha)
    return float((dists <= radius).mean())

# Re-methylation AUC for exit
def exit_remeth_auc(x_pred_blast, x_morula_in=x_morula):
    is_mz = (x_morula_in <= THETA_MO) & np.isfinite(x_pred_blast)
    is_re = is_mz & (x_pred_blast > THETA_REMETH)
    if is_mz.sum() < 5 or is_re.sum() < 2: return np.nan
    y_true = is_re[is_mz].astype(int)
    y_score= x_pred_blast[is_mz]
    try: return float(roc_auc_score(y_true, y_score))
    except: return np.nan

# RMSE for exit
def exit_rmse(x_pred, x_obs):
    v = np.isfinite(x_pred) & np.isfinite(x_obs)
    return float(np.sqrt(np.mean((x_pred[v]-x_obs[v])**2)))

# Pre-compute entry and exit corrections
remeth_idx_b = [i for i,c in enumerate(clusters) if c_diag_exit[i] > 0.05]
demeth_idx_b  = [i for i,c in enumerate(clusters) if c_diag_exit[i] < -0.05]
closure_idx  = [i for i,c in enumerate(clusters) if mod_map.get(c,"?") in ["M01","M05","M12"]]
access_idx   = [i for i,c in enumerate(clusters) if mod_map.get(c,"?") in ["M02","M10"]]

def apply_entry_correction(which_closure, which_access):
    """Apply entry correction with specified branch signs."""
    x_pred = (alpha_op * x_8cell + bias_op).copy()
    for i in closure_idx:
        x_pred[i] += which_closure * abs(c_diag_entry[i])
    for i in access_idx:
        x_pred[i] += which_access  * abs(c_diag_entry[i])
    return x_pred

def apply_exit_correction(which_remeth, which_demeth):
    """Apply exit correction with specified branch signs."""
    x_pred = x_blast_pred.copy()
    for i in remeth_idx_b:
        x_pred[i] += which_remeth * abs(c_diag_exit[i])
    for i in demeth_idx_b:
        x_pred[i] += which_demeth * abs(c_diag_exit[i])
    return x_pred

scenarios = [
    {
        "name": "methylation_only",
        "entry_pred": alpha_op * x_8cell + bias_op,
        "exit_pred":  x_blast_pred,
        "description": "Methylation-only baseline: no u_bio"
    },
    {
        "name": "correct_entry_correct_exit",
        "entry_pred": apply_entry_correction(+1, +1),
        "exit_pred":  apply_exit_correction(+1, -1),
        "description": "Correct entry (closure+access) + correct exit (remeth+demeth)"
    },
    {
        "name": "wrong_entry_correct_exit",
        "entry_pred": apply_entry_correction(-1, +1),   # wrong closure
        "exit_pred":  apply_exit_correction(+1, -1),
        "description": "Wrong entry closure + correct exit"
    },
    {
        "name": "correct_entry_wrong_exit",
        "entry_pred": apply_entry_correction(+1, +1),
        "exit_pred":  apply_exit_correction(+1, +1),   # wrong demeth
        "description": "Correct entry + wrong exit de-methylation direction"
    },
    {
        "name": "wrong_entry_wrong_exit",
        "entry_pred": apply_entry_correction(-1, +1),
        "exit_pred":  apply_exit_correction(-1, +1),   # wrong both
        "description": "Wrong entry + wrong exit"
    },
]

print(f"\n{'Scenario':<35} | {'Entry occ':>10} | {'Exit RMSE':>10} | {'Exit re-AUC':>12} | {'Collapse':>8}")
print("-"*85)

counterfactual_results = []
baseline_entry_occ = entry_occupancy(alpha_op*x_8cell+bias_op, x_morula)
baseline_exit_rmse = exit_rmse(x_blast_pred, x_blast)
# For AUC need a signal that varies; use the correction magnitude as proxy
# We'll compute re-methylation rate instead for morula-zero DMRs
def remeth_rate_pred(x_pred):
    is_mz = is_mzero & np.isfinite(x_pred)
    return float((x_pred[is_mz] > THETA_REMETH).mean()) if is_mz.sum()>0 else np.nan

for sc in scenarios:
    e_occ  = entry_occupancy(sc["entry_pred"], x_morula)
    ex_rmse= exit_rmse(sc["exit_pred"], x_blast)
    ex_rr  = remeth_rate_pred(sc["exit_pred"])
    collapse = (e_occ < 0.2) or (ex_rmse > baseline_exit_rmse * 1.2)

    row = {
        "scenario": sc["name"],
        "entry_occupancy": float(e_occ),
        "exit_rmse": float(ex_rmse),
        "exit_remeth_rate": float(ex_rr) if ex_rr is not None else None,
        "collapse": bool(collapse),
        "description": sc["description"],
    }
    counterfactual_results.append(row)

    name_short = sc["name"][:33]
    print(f"{name_short:<35} | {e_occ:>10.4f} | {ex_rmse:>10.4f} | {ex_rr:>12.4f} | {str(collapse):>8}")

cf_df = pd.DataFrame(counterfactual_results)
cf_df.to_csv(OUT/"step5_counterfactual_table.tsv", sep="\t", index=False)


# ══════════════════════════════════════════════════════════════════════════════
# FINAL: Complete model specification
# ══════════════════════════════════════════════════════════════════════════════
print("\n" + "="*65)
print("COMPLETE MODEL: Morula-centered gated operator-control dynamics")
print("="*65)

print("""
x_8 --[K_entry + B_entry * u_acc]--> x_M --[G_M]--> (q_zero, q_methylated)
     --[K_exit + B_exit * u_K4me3 + C_re * y_re]--> x_B

Where:
  K_entry = K_exit = 0.5611 (shared transition operator)
  u_acc   = morula accessibility (rho=+0.21 with meth_morula, perm_p=0.004)
  G_M     = morula gate (85/156 beta<=0.02, bimodal index=1.564)
  u_K4me3 = H3K4me3 8-cell score (AUC=0.792 for re-meth prediction, perm_p=0.015)
  C_re    = re-methylation correction term (35/85 morula-zero DMRs)
  y_re    = binary re-methylation indicator (predicted by u_K4me3 + M00 + CpG)
""")

# Final summary
final_model = {
    "date": "2026-05-29",
    "model_name": "morula-centered gated operator-control dynamics",
    "operator": {"alpha": float(alpha_op), "bias": float(bias_op),
                 "shared_for_entry_and_exit": True},
    "entry_u_bio": {"signal": "acc_morula", "rho": 0.210, "perm_p": 0.004},
    "morula_gate": {"n_beta_le_002": int(is_mzero.sum()),
                    "bimodal_index": 1.564, "fraction": float(is_mzero.sum()/len(clusters))},
    "exit_u_bio":  {"signal": "k4me3_8cell_score",
                    "auc": float(auc_lr), "loocv_auc": float(auc_loocv), "perm_p": float(perm_p_lr)},
    "exit_remeth_class": {"n": int(is_remeth.sum()), "total_mzero": int(is_mzero.sum()),
                          "rate": float(is_remeth.sum()/is_mzero.sum())},
    "step3_impulse": {r["module"]: {"J_entry":r["J_entry"],"J_exit":r["J_exit"]}
                      for r in impulse_rows},
    "step4_energy":  {r["module"]: {"E_entry":r["E_entry"],"E_exit":r["E_exit"]}
                      for r in energy_rows},
    "step5_counterfactual": counterfactual_results,
}
with open(OUT/"FINAL_COMPLETE_MODEL.json","w",encoding="utf-8") as f:
    json.dump(final_model, f, indent=2, ensure_ascii=False, default=str)

print(f"\nAll outputs saved to {OUT}/")
print(f"Total files: {len(list(OUT.iterdir()))}")
print("\nFiles created:")
for f in sorted(OUT.glob("step*.tsv")) + sorted(OUT.glob("step*.json")) + [OUT/"FINAL_COMPLETE_MODEL.json"]:
    print(f"  {f.name}")
```
