## Supplemental code for "An entropy-based diagnostic framework for characterizing methylation state dynamics during preimplantation development": 5_29_progress__deepest_model.py

```
#!/usr/bin/env python
"""
Deepest possible model with current public data.
Four parallel tracks:
1. Complete gene annotation + functional classification (all 156 DMRs)
2. Cross-species conservation (mouse hg19->mm10 liftover proxy)
3. Improved prediction model (better features, proper CV)
4. TF motif enrichment for re-methylation class
"""
from __future__ import annotations
import json
from pathlib import Path
import numpy as np
import pandas as pd
from scipy import stats
from sklearn.metrics import roc_auc_score
from sklearn.linear_model import LogisticRegression
from sklearn.model_selection import StratifiedKFold, cross_val_score

OUT = Path("E:/5_29_progress")
OUT.mkdir(parents=True, exist_ok=True)

TRAJ  = Path("E:/CSB_TRO_operator_time_DMR_dynamics_2026-05-25/results/CSB_TRO_DMR_stage_mean_trajectory.tsv")
RESID = Path("E:/CSB_TRO_operator_time_DMR_dynamics_2026-05-25/results/CSB_TRO_basin_residual_DMR_ranking.tsv")
META  = Path("E:/CSB_TRO_operator_time_DMR_dynamics_2026-05-25/results/CSB_TRO_DMR_metadata.tsv")
TSS   = Path("E:/CSB_TRO_operator_time_DMR_dynamics_2026-05-25/external/annotations/gene_tss.tsv")
ANN   = Path("E:/CSB_TRO_operator_time_DMR_dynamics_2026-05-25/results/CSB_TRO_residual_module_region_annotation.tsv")
GENE_LINKS = Path("E:/CSB_TRO_operator_time_DMR_dynamics_2026-05-25/results/CSB_TRO_residual_module_gene_links_symbol.tsv")
RNA_EXPR   = Path("E:/CSB_TRO_operator_time_DMR_dynamics_2026-05-25/results/CSB_TRO_GSE36552_gene_stage_expression_long.tsv")
MOTIF      = Path("E:/CSB_TRO_operator_time_DMR_dynamics_2026-05-25/external/motif/module_motif_enrichment.tsv")
MS    = Path("E:/5_28_progress/CSB_TRO_5_28_dmr_multistage_accessibility.tsv")
HIST  = Path("E:/CSB_TRO_operator_time_DMR_dynamics_2026-05-25/external/histone")

SEED = 42
rng  = np.random.default_rng(SEED)
N_BOOT = 2000

# ── Load core ─────────────────────────────────────────────────────────────────
traj  = pd.read_csv(TRAJ, sep="\t")
resid = pd.read_csv(RESID, sep="\t")
meta  = pd.read_csv(META, sep="\t")
tss   = pd.read_csv(TSS, sep="\t")
ann   = pd.read_csv(ANN, sep="\t")
ms    = pd.read_csv(MS, sep="\t")

stage_means = {}
for s, g in traj.groupby("stage"):
    stage_means[s] = g.set_index("cluster_name")["mean_beta"].to_dict()

clusters = sorted(resid["cluster_name"].tolist())
x_morula = np.array([stage_means.get("morula",{}).get(c,np.nan) for c in clusters])
x_blast  = np.array([stage_means.get("blastocyst",{}).get(c,np.nan) for c in clusters])
x_8cell  = np.array([stage_means.get("8-cell",{}).get(c,np.nan) for c in clusters])

meta_map = meta.set_index("cluster_name")
ms_map   = ms.set_index("cluster_name")
mod_map  = resid.set_index("cluster_name")["module_id"].to_dict()
rank_map = resid.set_index("cluster_name")["basin_residual_rank"].to_dict()

is_mzero  = (x_morula <= 0.02) & np.isfinite(x_blast)
is_remeth = is_mzero & (x_blast > 0.05)

# ══════════════════════════════════════════════════════════════════════════════
# TRACK 1: Complete gene annotation for all 156 DMRs
# ══════════════════════════════════════════════════════════════════════════════
print("="*65)
print("TRACK 1: Complete gene annotation (all 156 DMRs)")
print("="*65)

# Build TSS index by chromosome
tss["chr"] = tss["chr"].astype(str).str.strip()
tss_by_chr = {c: g.sort_values("tss") for c, g in tss.groupby("chr")}

def find_nearest_gene(chrom, start, end, tss_by_chr, window=500000):
    """Find nearest gene TSS within window."""
    if chrom not in tss_by_chr:
        return None, None, None, None
    sub = tss_by_chr[chrom]
    mid = (start + end) / 2
    dists = np.abs(sub["tss"].values - mid)
    idx = np.argmin(dists)
    if dists[idx] > window:
        return None, None, None, None
    row = sub.iloc[idx]
    return row["gene_name"], row["gene_id"], row["gene_type"], float(row["tss"] - mid)

def classify_genomic_context(chrom, start, end, tss_by_chr):
    """Classify DMR genomic context."""
    if chrom not in tss_by_chr:
        return "intergenic"
    sub = tss_by_chr[chrom]
    mid = (start + end) / 2
    # Promoter: within 2kb of TSS
    dists = np.abs(sub["tss"].values - mid)
    min_dist = dists.min()
    if min_dist <= 2000:
        return "promoter_2kb"
    elif min_dist <= 5000:
        return "promoter_5kb"
    else:
        return "intergenic"

# Annotate all 156 DMRs
full_annotation = []
for c in clusters:
    if c not in meta_map.index:
        full_annotation.append({"cluster_name": c})
        continue
    chrom = str(meta_map.loc[c,"chr"]).strip()
    start = int(meta_map.loc[c,"start"])
    end   = int(meta_map.loc[c,"end"])
    gene, gene_id, gene_type, dist = find_nearest_gene(chrom, start, end, tss_by_chr)
    context = classify_genomic_context(chrom, start, end, tss_by_chr)
    full_annotation.append({
        "cluster_name": c,
        "chr": chrom, "start": start, "end": end,
        "nearest_gene": gene, "gene_id": gene_id,
        "gene_type": gene_type, "dist_to_tss": dist,
        "genomic_context": context,
        "module_id": mod_map.get(c,"?"),
        "basin_residual_rank": rank_map.get(c,np.nan),
        "x_morula": x_morula[clusters.index(c)],
        "x_blast": x_blast[clusters.index(c)],
        "is_mzero": int(is_mzero[clusters.index(c)]),
        "is_remeth": int(is_remeth[clusters.index(c)]),
    })

full_ann_df = pd.DataFrame(full_annotation)
full_ann_df.to_csv(OUT/"track1_full_gene_annotation.tsv", sep="\t", index=False)

# Summary by module
print("\nGenomic context by module:")
for mid in ["M00","M01","M02","M05","M10","M12"]:
    sub = full_ann_df[full_ann_df["module_id"]==mid]
    if len(sub)==0: continue
    prom = (sub["genomic_context"]=="promoter_2kb").mean()
    interg = (sub["genomic_context"]=="intergenic").mean()
    print(f"  {mid}: n={len(sub)}, promoter_2kb={prom:.2f}, intergenic={interg:.2f}")

# M00 specific genes
m00_genes = full_ann_df[full_ann_df["module_id"]=="M00"][["cluster_name","nearest_gene","gene_type","dist_to_tss","genomic_context"]]
print(f"\nM00 genes (re-methylation class):")
print(m00_genes.to_string())

# Re-methylation class genes
remeth_genes = full_ann_df[full_ann_df["is_remeth"]==1][["cluster_name","nearest_gene","gene_type","dist_to_tss","module_id"]]
print(f"\nRe-methylation DMR genes (n={len(remeth_genes)}):")
print(remeth_genes.to_string())


# ══════════════════════════════════════════════════════════════════════════════
# TRACK 2: Cross-species conservation via mouse methylation data
# ══════════════════════════════════════════════════════════════════════════════
print("\n" + "="*65)
print("TRACK 2: Cross-species conservation (mouse GSE56697)")
print("="*65)

# Strategy: download mouse ICM methylation for the same age-DMR regions
# GSE56697 has: oocyte, 2cell, 4cell, ICM (no morula)
# We can test: do human age-DMRs show similar methylation patterns in mouse?
# This is a conservation test, not a morula-specific test

# Download a small sample of mouse ICM data to test
print("Downloading mouse ICM methylation sample...")
import subprocess, gzip, io

# Download just the first 1M lines of mouse ICM data
url = "https://ftp.ncbi.nlm.nih.gov/geo/series/GSE56nnn/GSE56697/suppl/GSM1386023_ICM_mc_CG_plus.bed.gz"
try:
    result = subprocess.run(
        ["curl", "-s", "--range", "0-5000000", url],
        capture_output=True, timeout=60
    )
    if result.returncode == 0 and len(result.stdout) > 1000:
        # Parse the partial gzip
        try:
            data = gzip.decompress(result.stdout)
            lines = data.decode("utf-8").strip().split("\n")
            mouse_icm = pd.DataFrame([l.split("\t") for l in lines if l],
                                      columns=["chr","start","end","meth","total"])
            mouse_icm["start"] = pd.to_numeric(mouse_icm["start"], errors="coerce")
            mouse_icm["end"]   = pd.to_numeric(mouse_icm["end"], errors="coerce")
            mouse_icm["meth"]  = pd.to_numeric(mouse_icm["meth"], errors="coerce")
            mouse_icm["total"] = pd.to_numeric(mouse_icm["total"], errors="coerce")
            mouse_icm = mouse_icm.dropna()
            mouse_icm["beta"] = mouse_icm["meth"] / (mouse_icm["total"] + 1e-8)
            print(f"  Mouse ICM sample: {len(mouse_icm)} CpGs")
            mouse_available = True
        except Exception as e:
            print(f"  Parse error: {e}")
            mouse_available = False
    else:
        print(f"  Download failed or too small")
        mouse_available = False
except Exception as e:
    print(f"  Download error: {e}")
    mouse_available = False

if not mouse_available:
    print("  Using liftover-based conservation proxy instead")
    # Alternative: use the fact that age-DMRs are defined from GSE102970
    # which is blood methylation aging data
    # The conservation test: are these DMRs in conserved genomic regions?
    # Proxy: distance to nearest gene (conserved genes tend to have conserved regulatory regions)
    # This is a weak proxy but available

    # Better alternative: use the existing mouse data we already have
    # GSE66390 has mouse 2cell/4cell/8cell/ICM methylation (bedGraph format)
    # We can check if human age-DMR regions show similar methylation dynamics in mouse
    print("  Checking GSE66390 mouse methylation...")
    mouse_url = "https://ftp.ncbi.nlm.nih.gov/geo/series/GSE66nnn/GSE66390/suppl/GSM1933930_icm_re1.bedGraph.gz"
    try:
        result2 = subprocess.run(
            ["curl", "-s", "--range", "0-3000000", mouse_url],
            capture_output=True, timeout=60
        )
        if result2.returncode == 0 and len(result2.stdout) > 1000:
            data2 = gzip.decompress(result2.stdout)
            lines2 = data2.decode("utf-8").strip().split("\n")
            mouse_icm2 = pd.DataFrame([l.split("\t") for l in lines2 if l and not l.startswith("track")],
                                       columns=["chr","start","end","beta"])
            mouse_icm2["start"] = pd.to_numeric(mouse_icm2["start"], errors="coerce")
            mouse_icm2["end"]   = pd.to_numeric(mouse_icm2["end"], errors="coerce")
            mouse_icm2["beta"]  = pd.to_numeric(mouse_icm2["beta"], errors="coerce")
            mouse_icm2 = mouse_icm2.dropna()
            print(f"  Mouse ICM (GSE66390): {len(mouse_icm2)} regions")
            mouse_available = True
            mouse_icm = mouse_icm2
        else:
            mouse_available = False
    except Exception as e:
        print(f"  GSE66390 error: {e}")
        mouse_available = False

# Conservation analysis: do human DMR coordinates overlap with mouse methylation?
# Note: human hg19 vs mouse mm9/mm10 - need liftover
# Without liftover, we use gene-based conservation:
# If human DMR is near gene X, and mouse has similar methylation near gene X's ortholog
# This is approximate but valid

print("\n  Gene-based conservation analysis:")
# For each DMR with a nearby gene, check if that gene is known to be developmentally regulated
# Use the RNA expression data we have
try:
    rna = pd.read_csv(RNA_EXPR, sep="\t")
    print(f"  RNA expression data: {len(rna)} rows")
    print(f"  Columns: {rna.columns.tolist()[:8]}")
    # Get genes expressed in morula
    if "stage" in rna.columns and "gene_name" in rna.columns:
        morula_genes = rna[rna["stage"]=="morula"]["gene_name"].unique()
        print(f"  Genes expressed in morula: {len(morula_genes)}")
        # Check overlap with DMR-linked genes
        dmr_genes = set(full_ann_df["nearest_gene"].dropna().tolist())
        overlap = dmr_genes & set(morula_genes)
        print(f"  DMR-linked genes expressed in morula: {len(overlap)}/{len(dmr_genes)}")
except Exception as e:
    print(f"  RNA data error: {e}")


# ══════════════════════════════════════════════════════════════════════════════
# TRACK 3: Improved prediction model
# ══════════════════════════════════════════════════════════════════════════════
print("\n" + "="*65)
print("TRACK 3: Improved prediction model")
print("="*65)

# Load histone signals
def load_overlap(fname):
    df = pd.read_csv(HIST/fname, sep="\t", header=None, compression="gzip",
                     names=["chr","start","end","name","score"], usecols=[0,1,2,3,4])
    df["chr"] = df["chr"].astype(str).str.strip()
    by_chr = {c:g for c,g in df.groupby("chr")}
    ov, sc = [], []
    for c in clusters:
        if c not in meta_map.index: ov.append(0); sc.append(np.nan); continue
        chrom=str(meta_map.loc[c,"chr"]).strip(); ds=int(meta_map.loc[c,"start"]); de=int(meta_map.loc[c,"end"])
        if chrom not in by_chr: ov.append(0); sc.append(np.nan); continue
        sub=by_chr[chrom]; hits=sub[(sub["start"]<de)&(sub["end"]>ds)]
        sv=pd.to_numeric(hits["score"],errors="coerce").dropna()
        ov.append(int(len(hits)>0)); sc.append(float(sv.max()) if len(sv)>0 else np.nan)
    return np.array(ov), np.array(sc)

k4me3_8cell_ov, k4me3_8cell_sc = load_overlap("H3K4me3_8cell.hg19.bed.gz")
k27me3_blast_ov, _ = load_overlap("H3K27me3_blastocyst.hg19.bed.gz")
k27ac_blast_ov, _  = load_overlap("H3K27ac_blastocyst.hg19.bed.gz")
k4me3_blast_ov, _  = load_overlap("H3K4me3_blastocyst.hg19.bed.gz")
k27ac_8cell_ov, _  = load_overlap("H3K27ac_8cell.hg19.bed.gz")

acc_morula = np.array([ms_map.loc[c,"acc_morula_mean"] if c in ms_map.index else np.nan for c in clusters])
n_cpg = np.array([meta_map.loc[c,"n_cpg_target"] if c in meta_map.index else np.nan for c in clusters])
width = np.array([meta_map.loc[c,"width"] if c in meta_map.index else np.nan for c in clusters])

# Add genomic context features
is_promoter = np.array([1 if full_ann_df.set_index("cluster_name").loc[c,"genomic_context"]=="promoter_2kb"
                         else 0 for c in clusters])
is_m00 = np.array([1 if mod_map.get(c,"?")=="M00" else 0 for c in clusters])

# Among morula-zero DMRs
mzero_idx = np.where(is_mzero)[0]
y_bin = is_remeth[mzero_idx].astype(float)

# Build comprehensive feature matrix
def build_features(idx_set):
    k4sc = k4me3_8cell_sc[idx_set]
    k4sc_f = np.where(np.isfinite(k4sc), k4sc, 0.0)
    k4ov = k4me3_8cell_ov[idx_set].astype(float)
    m00  = is_m00[idx_set].astype(float)
    prom = is_promoter[idx_set].astype(float)
    cpg  = n_cpg[idx_set]; cpg_f = np.where(np.isfinite(cpg), cpg, np.nanmean(n_cpg))
    w    = width[idx_set]; w_f   = np.where(np.isfinite(w), w, np.nanmean(width))
    k27me3_b = k27me3_blast_ov[idx_set].astype(float)
    k27ac_b  = k27ac_blast_ov[idx_set].astype(float)
    k4me3_b  = k4me3_blast_ov[idx_set].astype(float)
    k27ac_8  = k27ac_8cell_ov[idx_set].astype(float)
    acc_mo   = acc_morula[idx_set]; acc_f = np.where(np.isfinite(acc_mo), acc_mo, 0.0)
    # Interaction: M00 * k4me3_8cell
    m00_k4 = m00 * k4ov
    # Interaction: promoter * k27me3_blast
    prom_k27 = prom * k27me3_b
    return np.column_stack([k4sc_f, k4ov, m00, prom, cpg_f, w_f,
                             k27me3_b, k27ac_b, k4me3_b, k27ac_8, acc_f,
                             m00_k4, prom_k27])

feat_names_v2 = ["k4me3_8cell_score","k4me3_8cell_ov","M00","promoter_2kb",
                  "n_cpg","width","k27me3_blast","k27ac_blast","k4me3_blast",
                  "k27ac_8cell","acc_morula","M00*k4me3_8cell","promoter*k27me3"]

X_v2 = build_features(mzero_idx)
X_v2_sc = (X_v2 - X_v2.mean(axis=0)) / (X_v2.std(axis=0) + 1e-8)

print(f"\nFeatures v2: {feat_names_v2}")
print(f"n={len(y_bin)}, remeth={int(y_bin.sum())}")

# Stratified 5-fold CV
skf = StratifiedKFold(n_splits=5, shuffle=True, random_state=SEED)
cv_aucs = []
for train_idx, test_idx in skf.split(X_v2_sc, y_bin):
    lr = LogisticRegression(C=0.1, random_state=SEED, max_iter=1000)
    lr.fit(X_v2_sc[train_idx], y_bin[train_idx])
    proba = lr.predict_proba(X_v2_sc[test_idx])[:,1]
    try:
        cv_aucs.append(float(roc_auc_score(y_bin[test_idx], proba)))
    except:
        cv_aucs.append(0.5)

cv_auc_mean = float(np.mean(cv_aucs))
cv_auc_std  = float(np.std(cv_aucs))
print(f"\n5-fold stratified CV AUC: {cv_auc_mean:.4f} ± {cv_auc_std:.4f}")
print(f"Fold AUCs: {[f'{a:.3f}' for a in cv_aucs]}")

# Full model AUC
lr_full = LogisticRegression(C=0.1, random_state=SEED, max_iter=1000)
lr_full.fit(X_v2_sc, y_bin)
proba_full = lr_full.predict_proba(X_v2_sc)[:,1]
auc_full = float(roc_auc_score(y_bin, proba_full))
print(f"Full model AUC: {auc_full:.4f}")

# Permutation test
null_cv_aucs = []
for _ in range(N_BOOT):
    y_perm = rng.permutation(y_bin)
    fold_aucs = []
    for train_idx, test_idx in skf.split(X_v2_sc, y_perm):
        lr_p = LogisticRegression(C=0.1, random_state=SEED, max_iter=500)
        lr_p.fit(X_v2_sc[train_idx], y_perm[train_idx])
        p_p = lr_p.predict_proba(X_v2_sc[test_idx])[:,1]
        try:
            fold_aucs.append(float(roc_auc_score(y_perm[test_idx], p_p)))
        except:
            fold_aucs.append(0.5)
    null_cv_aucs.append(np.mean(fold_aucs))
null_cv_aucs = np.array(null_cv_aucs)
perm_p_v2 = float((null_cv_aucs >= cv_auc_mean).mean())
q95_v2    = float(np.quantile(null_cv_aucs, 0.95))
print(f"Permutation test: perm_p={perm_p_v2:.4f}, null q95={q95_v2:.4f}")
print(f"Significant: {cv_auc_mean > q95_v2}")

# Feature importance
print("\nFeature importance (coefficient magnitude):")
for name, coef in sorted(zip(feat_names_v2, lr_full.coef_[0]),
                          key=lambda x: abs(x[1]), reverse=True):
    print(f"  {name}: {coef:.4f} (OR={np.exp(coef):.3f})")


# ══════════════════════════════════════════════════════════════════════════════
# TRACK 4: TF motif enrichment for re-methylation class
# ══════════════════════════════════════════════════════════════════════════════
print("\n" + "="*65)
print("TRACK 4: TF motif enrichment")
print("="*65)

try:
    motif = pd.read_csv(MOTIF, sep="\t")
    print(f"Motif data: {len(motif)} rows")
    print(f"Columns: {motif.columns.tolist()[:10]}")
    # Look for M00 enrichment
    if "module_id" in motif.columns:
        m00_motif = motif[motif["module_id"]=="M00"]
        print(f"\nM00 motif enrichment: {len(m00_motif)} entries")
        if len(m00_motif) > 0:
            print(m00_motif.head(10).to_string())
except Exception as e:
    print(f"Motif data error: {e}")
    # Try alternative motif file
    try:
        motif2 = pd.read_csv(
            "E:/CSB_TRO_operator_time_DMR_dynamics_2026-05-25/results/CSB_TRO_residual_DMR_motif_enrichment.tsv",
            sep="\t")
        print(f"Alternative motif data: {len(motif2)} rows")
        print(motif2.head(5).to_string())
    except Exception as e2:
        print(f"Alternative motif error: {e2}")


# ══════════════════════════════════════════════════════════════════════════════
# TRACK 5: RNA expression coupling for re-methylation genes
# ══════════════════════════════════════════════════════════════════════════════
print("\n" + "="*65)
print("TRACK 5: RNA expression coupling")
print("="*65)

try:
    rna = pd.read_csv(RNA_EXPR, sep="\t")
    print(f"RNA data: {len(rna)} rows, cols: {rna.columns.tolist()[:8]}")

    # Get genes near re-methylation DMRs
    remeth_ann = full_ann_df[full_ann_df["is_remeth"]==1]
    remeth_genes_list = remeth_ann["nearest_gene"].dropna().tolist()
    print(f"\nRe-methylation DMR nearest genes: {remeth_genes_list}")

    # Check expression of these genes across stages
    if "gene_name" in rna.columns and "stage" in rna.columns:
        for gene in remeth_genes_list[:10]:
            gene_expr = rna[rna["gene_name"]==gene]
            if len(gene_expr) > 0:
                stage_expr = gene_expr.groupby("stage")["expression"].mean() if "expression" in gene_expr.columns else None
                if stage_expr is not None:
                    print(f"  {gene}: {stage_expr.to_dict()}")
except Exception as e:
    print(f"RNA coupling error: {e}")


# ══════════════════════════════════════════════════════════════════════════════
# SAVE ALL RESULTS
# ══════════════════════════════════════════════════════════════════════════════
results = {
    "date": "2026-05-29",
    "track1_gene_annotation": {
        "n_dmrs_annotated": len(full_ann_df),
        "m00_genes": m00_genes["nearest_gene"].dropna().tolist(),
        "remeth_genes": remeth_genes["nearest_gene"].dropna().tolist(),
    },
    "track3_improved_model": {
        "cv_auc_mean": float(cv_auc_mean),
        "cv_auc_std": float(cv_auc_std),
        "full_auc": float(auc_full),
        "perm_p": float(perm_p_v2),
        "null_q95": float(q95_v2),
        "significant": bool(cv_auc_mean > q95_v2),
        "n_features": len(feat_names_v2),
        "top_features": sorted(zip(feat_names_v2, lr_full.coef_[0].tolist()),
                                key=lambda x: abs(x[1]), reverse=True)[:5],
    },
}
with open(OUT/"deepest_model_results.json","w",encoding="utf-8") as f:
    json.dump(results, f, indent=2, ensure_ascii=False, default=str)

print(f"\n{'='*65}")
print("SUMMARY")
print(f"{'='*65}")
print(f"Track 1: {len(full_ann_df)} DMRs annotated with nearest genes")
print(f"Track 3: 5-fold CV AUC={cv_auc_mean:.4f}±{cv_auc_std:.4f}, perm_p={perm_p_v2:.4f}")
print(f"         Significant: {cv_auc_mean > q95_v2}")
print(f"Saved: {OUT}/deepest_model_results.json")
print(f"       {OUT}/track1_full_gene_annotation.tsv")
```
