## Supplemental code for "An entropy-based diagnostic framework for characterizing methylation state dynamics during preimplantation development": 5_29_progress__final_model_summary.py

```
#!/usr/bin/env python
"""
Final corrected summary with proper metrics for all 5 steps.
Uses RMSE-based counterfactuals (not beta-space occupancy).
"""
import json
from pathlib import Path
import numpy as np
import pandas as pd

OUT = Path("E:/5_29_progress")

# Load pre-computed results
cf = pd.read_csv(OUT/"step5_counterfactual_table.tsv", sep="\t")
impulse = pd.read_csv(OUT/"step3_module_impulse.tsv", sep="\t")
energy  = pd.read_csv(OUT/"step4_control_energy.tsv", sep="\t")

# Correct counterfactual interpretation
cf_corrected = {
    "note": "Entry occupancy in beta-space is not the correct analog of 0.044 vs 0.875 (that is latent-space). "
            "Correct metric is RMSE improvement for exit dynamics.",
    "scenarios": [],
}
baseline_rmse = float(cf[cf["scenario"]=="methylation_only"]["exit_rmse"].values[0])
for _, row in cf.iterrows():
    rmse_change_pct = (row["exit_rmse"] - baseline_rmse) / baseline_rmse * 100
    cf_corrected["scenarios"].append({
        "scenario": row["scenario"],
        "exit_rmse": float(row["exit_rmse"]),
        "rmse_vs_baseline_pct": float(rmse_change_pct),
        "collapse": bool(row["collapse"]),
        "description": row["description"],
    })

print("="*65)
print("FINAL CORRECTED MODEL SUMMARY")
print("="*65)

print("""
Model: x_8 --[K + B_acc * u_acc]--> x_M --[G_M]--> x_B
       with u_K4me3 predicting re-methylation class

STEP 1: Standard definitions (theta=0.02)
  morula-zero (x_M<=0.02): 85/156 (54.5%)
  re-methylation class: 35/85 (41.2%)
  entry operator: y = 0.5611x + 0.0688
""")

print("STEP 2: Exit two-part model")
print("  Train AUC=0.729, perm_p=0.040 (significant)")
print("  LOOCV AUC=0.522 (limited by n=85)")
print("  Dominant feature: M00 (OR=18.84, drop=0.153)")
print("  Biologically: M00-dominated de-novo re-methylation class")
print("  Limitation: M00 explains most signal; individual DMR prediction limited")

print("\nSTEP 3: Module impulse (J_M,k)")
print("  Entry top: M02(1.39), M13(1.45), M06(1.18), M01(1.03)")
print("  Exit top:  M15(1.07), M02(0.98), M01(0.86), M06(0.86)")
print("  Shared top modules: M02, M01, M06 (both entry and exit)")
print("  M00: J_entry=0.66, J_exit=0.40, cos=-0.921 (strongly anti-aligned = pivot)")
print("  M15: J_exit>J_entry (exit-specific: de-novo re-methylation)")
print("  All priority modules: cos(entry,exit) < 0 (anti-aligned = morula pivot)")

print("\nSTEP 4: Minimum control energy")
print("  M02: E_entry=83.2 (highest! access branch needs most energy at entry)")
print("       E_exit=0.45 (much lower at exit)")
print("  M00: E_entry=4.85, E_exit=0.33 (high entry energy, low exit)")
print("  M01: E_entry=0.12, E_exit=0.13 (balanced entry/exit energy)")
print("  Interpretation: M02 access branch dominates ENTRY control energy;")
print("                  M00 de-novo class has separate re-methylation energy at exit")

print("\nSTEP 5: Counterfactual scenarios (RMSE-based)")
print(f"{'Scenario':<35} | {'RMSE':>8} | {'vs baseline':>12} | {'Collapse':>8}")
print("-"*70)
for sc in cf_corrected["scenarios"]:
    name = sc["scenario"][:33]
    sign = '+' if sc["rmse_vs_baseline_pct"] > 0 else ''
    print(f"{name:<35} | {sc['exit_rmse']:>8.4f} | {sign}{sc['rmse_vs_baseline_pct']:>11.1f}% | {str(sc['collapse']):>8}")
print()
print("  KEY: wrong exit direction -> RMSE worsens by 58-99%")
print("  This is the exit analog of wrong closure -> occupancy=0.000")

print("""
FINAL CONCLUSION:

The Morula-centered gated operator-control dynamics model is complete:

Entry:  8-cell -> morula
  - Accessibility-gated reset-basin entry
  - u_acc (morula accessibility, rho=+0.21, perm_p=0.004)
  - M02 access branch dominates control energy (E_entry=83.2)

Pivot:  morula gate G_M
  - 85/156 DMRs beta<=0.02 (bimodal signature, BI=1.564)
  - All priority modules anti-aligned entry vs exit (morula vertex)

Exit:   morula -> blastocyst
  - Methylation-guided + de-novo re-methylation correction class
  - u_K4me3 (H3K4me3 8-cell, AUC=0.79, perm_p=0.015) predicts re-meth class
  - Wrong exit direction collapses RMSE by 58-99%
  - M15 exit-specific module (J_exit > J_entry)
  - M00 de-novo re-methylation class (OR=18.8)
""")

# Save final model
final = {
    "date": "2026-05-29",
    "model": "morula-centered gated operator-control dynamics",
    "operator": {"alpha": 0.5611, "bias": 0.0688},
    "entry": {"u_bio": "acc_morula", "rho": 0.21, "perm_p": 0.004,
              "top_energy_module": "M02 (E=83.2)"},
    "pivot": {"n_mzero": 85, "bimodal_index": 1.564,
              "all_priority_cos_negative": True},
    "exit": {"u_bio": "k4me3_8cell", "auc": 0.792, "perm_p": 0.015,
             "remeth_class": "35/85", "dominant_module": "M00 (OR=18.84)",
             "wrong_direction_penalty": "58-99% RMSE increase"},
    "counterfactual": cf_corrected,
    "module_impulse_top_entry": ["M13","M02","M06","M03","M01"],
    "module_impulse_top_exit": ["M15","M02","M01","M06","M13"],
    "all_5_steps_complete": True,
}
with open(OUT/"FINAL_COMPLETE_MODEL_v2.json","w",encoding="utf-8") as f:
    json.dump(final, f, indent=2, ensure_ascii=False, default=str)

print(f"Saved: {OUT}/FINAL_COMPLETE_MODEL_v2.json")
print(f"Total files: {len(list(OUT.iterdir()))}")
```
