## Supplemental code for "An entropy-based diagnostic framework for characterizing methylation state dynamics during preimplantation development": 5_29_progress__go_enrichment.py

```
#!/usr/bin/env python
"""
GO/Pathway enrichment analysis for all DMR gene sets.
Uses Enrichr API via gseapy for GO Biological Process, KEGG, and Reactome.
Gene sets analyzed:
  1. All 156 DMR genes (background)
  2. Re-methylation class genes (35 DMRs)
  3. M00 module genes
  4. Priority residual module genes (M01/M02/M05/M10/M12)
  5. Entry-specific (top25 residual DMR genes)
  6. Blast-activation sub-group (FOXD2, GREM2, DSCAML1...)
  7. Morula-specific sub-group (HAND2, HOXD9, NKX6-2...)
"""
from __future__ import annotations
import json, time
from pathlib import Path
import numpy as np
import pandas as pd
from scipy import stats
import gseapy as gp

OUT = Path("E:/5_29_progress")
OUT.mkdir(parents=True, exist_ok=True)

# ── Load annotation ────────────────────────────────────────────────────────────
ann = pd.read_csv(OUT/"track1_full_gene_annotation.tsv", sep="\t")
rna = pd.read_csv("E:/CSB_TRO_operator_time_DMR_dynamics_2026-05-25/external/rna/gene_stage_matrix.tsv", sep="\t")

# Filter to protein-coding genes only for enrichment
def get_genes(df, filter_coding=True):
    genes = df["nearest_gene"].dropna().tolist()
    if filter_coding:
        coding = df[df["gene_type"]=="protein_coding"]["nearest_gene"].dropna().tolist()
        return list(set(coding))
    return list(set(genes))

# Define gene sets
gene_sets = {
    "all_156_DMR_genes":        get_genes(ann),
    "remeth_class":             get_genes(ann[ann["is_remeth"]==1]),
    "M00_module":               get_genes(ann[ann["module_id"]=="M00"]),
    "priority_modules_M01_M02_M05_M10_M12": get_genes(ann[ann["module_id"].isin(["M01","M02","M05","M10","M12"])]),
    "top25_residual":           get_genes(ann[ann["basin_residual_rank"]<=25]),
    "blast_activation_subgroup": ["FOXD2","GREM2","DSCAML1","SMTNL2"],
    "morula_specific_subgroup":  ["HAND2","HOXD9","NKX6-2"],
    "mzero_stay_genes":         get_genes(ann[(ann["is_mzero"]==1)&(ann["is_remeth"]==0)]),
}

print("Gene set sizes:")
for name, genes in gene_sets.items():
    print(f"  {name}: {len(genes)} genes")
    if len(genes) <= 10:
        print(f"    {genes}")

# ── Enrichr databases to query ─────────────────────────────────────────────────
DATABASES = [
    "GO_Biological_Process_2023",
    "GO_Molecular_Function_2023",
    "KEGG_2021_Human",
    "Reactome_2022",
    "WikiPathway_2023_Human",
    "MSigDB_Hallmark_2020",
    "Human_Phenotype_Ontology",
]

# ── Run enrichment for each gene set ──────────────────────────────────────────
all_results = {}
PVAL_THRESH = 0.05

print("\nRunning Enrichr enrichment analysis...")
for set_name, genes in gene_sets.items():
    if len(genes) < 3:
        print(f"  Skipping {set_name}: too few genes ({len(genes)})")
        continue

    print(f"\n  {set_name} (n={len(genes)}):")
    set_results = {}

    for db in DATABASES:
        try:
            enr = gp.enrichr(
                gene_list=genes,
                gene_sets=db,
                organism="human",
                outdir=None,
                verbose=False,
            )
            if enr.results is not None and len(enr.results) > 0:
                sig = enr.results[enr.results["Adjusted P-value"] < PVAL_THRESH]
                if len(sig) > 0:
                    top = sig.nsmallest(10, "Adjusted P-value")[
                        ["Term","Overlap","P-value","Adjusted P-value","Genes"]
                    ].to_dict("records")
                    set_results[db] = top
                    print(f"    {db}: {len(sig)} significant terms")
                    for r in top[:3]:
                        print(f"      {r['Term'][:60]}: adj_p={r['Adjusted P-value']:.4f}")
                else:
                    print(f"    {db}: no significant terms")
            time.sleep(0.5)  # rate limit
        except Exception as e:
            print(f"    {db}: error - {e}")
            time.sleep(1)

    all_results[set_name] = set_results

# ── Save results ───────────────────────────────────────────────────────────────
with open(OUT/"go_enrichment_results.json","w",encoding="utf-8") as f:
    json.dump(all_results, f, indent=2, ensure_ascii=False, default=str)

# Create summary table
summary_rows = []
for set_name, db_results in all_results.items():
    for db, terms in db_results.items():
        for term in terms:
            summary_rows.append({
                "gene_set": set_name,
                "database": db,
                "term": term["Term"],
                "overlap": term["Overlap"],
                "pvalue": term["P-value"],
                "adj_pvalue": term["Adjusted P-value"],
                "genes": term["Genes"],
            })

if summary_rows:
    summary_df = pd.DataFrame(summary_rows)
    summary_df.to_csv(OUT/"go_enrichment_summary.tsv", sep="\t", index=False)
    print(f"\nSaved {len(summary_df)} enrichment results")
else:
    print("\nNo significant enrichment results found")

# ── Print key findings ─────────────────────────────────────────────────────────
print("\n" + "="*65)
print("KEY ENRICHMENT FINDINGS")
print("="*65)

for set_name in ["remeth_class","M00_module","blast_activation_subgroup","morula_specific_subgroup"]:
    if set_name not in all_results or not all_results[set_name]:
        print(f"\n{set_name}: no significant enrichment")
        continue
    print(f"\n{set_name}:")
    for db, terms in all_results[set_name].items():
        if terms:
            print(f"  {db}:")
            for t in terms[:5]:
                print(f"    {t['Term'][:70]}: adj_p={t['Adjusted P-value']:.4f}, genes={t['Genes'][:60]}")
```
