## Supplemental code for "An entropy-based diagnostic framework for characterizing methylation state dynamics during preimplantation development": 5_29_progress__pre8_full_dynamics.py

```
#!/usr/bin/env python
"""
Full pre-8-cell dynamics: 8 elements aligned with 8cell->morula and morula->blast.
Stage I: upstream erasure dynamics (zygote/PN -> 2-cell -> 4-cell -> 8-cell)
"""
from __future__ import annotations
import json
from pathlib import Path
import numpy as np
import pandas as pd
from scipy import stats
from numpy.linalg import norm
from sklearn.metrics import roc_auc_score

OUT = Path("E:/5_29_progress/pre8_dynamics")
OUT.mkdir(parents=True, exist_ok=True)

TRAJ  = Path("E:/CSB_TRO_operator_time_DMR_dynamics_2026-05-25/results/CSB_TRO_DMR_stage_mean_trajectory.tsv")
RESID = Path("E:/CSB_TRO_operator_time_DMR_dynamics_2026-05-25/results/CSB_TRO_basin_residual_DMR_ranking.tsv")
RNA   = Path("E:/CSB_TRO_operator_time_DMR_dynamics_2026-05-25/external/rna/gene_stage_matrix.tsv")
ANN   = Path("E:/5_29_progress/track1_full_gene_annotation.tsv")

SEED = 42
N_BOOT = 2000
rng = np.random.default_rng(SEED)

traj  = pd.read_csv(TRAJ, sep="\t")
resid = pd.read_csv(RESID, sep="\t")
rna   = pd.read_csv(RNA, sep="\t")
ann   = pd.read_csv(ANN, sep="\t")

stage_means = {}
for s, g in traj.groupby("stage"):
    stage_means[s] = g.set_index("cluster_name")["mean_beta"].to_dict()

clusters = sorted(resid["cluster_name"].tolist())
mod_map  = resid.set_index("cluster_name")["module_id"].to_dict()

PRE8_TRANS = [
    ("MII oocyte","zygote/PN"),
    ("zygote/PN","2-cell"),
    ("2-cell","4-cell"),
    ("4-cell","8-cell"),
]
TRANS_LABELS = {
    ("MII oocyte","zygote/PN"): "oocyte_to_zygote",
    ("zygote/PN","2-cell"):     "zygote_to_2cell",
    ("2-cell","4-cell"):        "2cell_to_4cell",
    ("4-cell","8-cell"):        "4cell_to_8cell",
    ("8-cell","morula"):        "8cell_to_morula",
    ("morula","blastocyst"):    "morula_to_blast",
}

def svec(stage):
    return np.array([stage_means.get(stage,{}).get(c, np.nan) for c in clusters])

x_oocyte = svec("MII oocyte")
x_zygote = svec("zygote/PN")
x_2cell  = svec("2-cell")
x_4cell  = svec("4-cell")
x_8cell  = svec("8-cell")
x_morula = svec("morula")
x_blast  = svec("blastocyst")

stage_vecs = {
    "MII oocyte": x_oocyte, "zygote/PN": x_zygote,
    "2-cell": x_2cell, "4-cell": x_4cell, "8-cell": x_8cell,
    "morula": x_morula, "blastocyst": x_blast,
}

# Shared operator
X_tr, Y_tr = [], []
for sf, st in PRE8_TRANS:
    xf = stage_vecs[sf]; xt = stage_vecs[st]
    v = np.isfinite(xf) & np.isfinite(xt)
    X_tr.append(xf[v].reshape(-1,1)); Y_tr.append(xt[v])
X_tr = np.vstack(X_tr); Y_tr = np.concatenate(Y_tr)
A_aug = np.column_stack([X_tr, np.ones(len(X_tr))])
coef, _, _, _ = np.linalg.lstsq(A_aug, Y_tr, rcond=None)
alpha_op, bias_op = coef[0], coef[1]
print(f"Shared operator: y = {alpha_op:.4f}*x + {bias_op:.4f}")

# ── B1: Entry-competence failure metric ────────────────────────────────────────
print("\n=== B1: Entry-competence failure metric ===")
ENTRY_COMP_THRESH = 0.10
is_entry_comp = (x_8cell <= ENTRY_COMP_THRESH) & np.isfinite(x_8cell)
n_entry_comp = int(is_entry_comp.sum())
baseline_acc = 1 - n_entry_comp/156
print(f"Entry-competent DMRs (x_8cell<={ENTRY_COMP_THRESH}): {n_entry_comp}/156")
print(f"Baseline accuracy: {baseline_acc:.3f}")

v_b1 = np.isfinite(x_zygote) & np.isfinite(x_8cell)
y_b1 = is_entry_comp[v_b1].astype(int)
auc_b1 = float(roc_auc_score(y_b1, -x_zygote[v_b1]))
null_b1 = np.array([float(roc_auc_score(rng.permutation(y_b1), -x_zygote[v_b1])) for _ in range(N_BOOT)])
pp_b1 = float((null_b1 >= auc_b1).mean())
q95_b1 = float(np.quantile(null_b1, 0.95))
print(f"Zygote meth -> entry-competence: AUC={auc_b1:.4f}, perm_p={pp_b1:.4f}, sig={auc_b1>q95_b1}")

slope_z4 = x_4cell - x_zygote
v_slope = np.isfinite(slope_z4) & np.isfinite(x_8cell)
y_slope = is_entry_comp[v_slope].astype(int)
auc_slope = float(roc_auc_score(y_slope, -slope_z4[v_slope]))
null_slope = np.array([float(roc_auc_score(rng.permutation(y_slope), -slope_z4[v_slope])) for _ in range(N_BOOT)])
pp_slope = float((null_slope >= auc_slope).mean())
q95_slope = float(np.quantile(null_slope, 0.95))
print(f"Trajectory slope (Z->4cell) -> entry-competence: AUC={auc_slope:.4f}, perm_p={pp_slope:.4f}, sig={auc_slope>q95_slope}")

b1_results = {
    "n_entry_competent": n_entry_comp, "baseline_accuracy": float(baseline_acc),
    "zygote_meth_auc": float(auc_b1), "zygote_meth_perm_p": float(pp_b1), "zygote_sig": bool(auc_b1>q95_b1),
    "trajectory_slope_auc": float(auc_slope), "trajectory_slope_perm_p": float(pp_slope), "slope_sig": bool(auc_slope>q95_slope),
}

# ── B2: c_diag per transition ──────────────────────────────────────────────────
print("\n=== B2: c_diag per transition ===")
cdiag_per_trans = {}
for sf, st in PRE8_TRANS + [("8-cell","morula"),("morula","blastocyst")]:
    xf = stage_vecs[sf]; xt = stage_vecs[st]
    c_diag = xt - (alpha_op * xf + bias_op)
    label = TRANS_LABELS[(sf,st)]
    v = np.isfinite(c_diag)
    n_d = int((c_diag[v]<-0.05).sum()); n_r = int((c_diag[v]>0.05).sum()); n_s = int((np.abs(c_diag[v])<=0.05).sum())
    direction = "demeth-dominant" if n_d>n_r*1.5 else ("remeth-dominant" if n_r>n_d*1.5 else "bidirectional")
    cdiag_per_trans[label] = {"n_demeth":n_d,"n_remeth":n_r,"n_stable":n_s,"direction":direction,"mean":float(np.nanmean(c_diag))}
    print(f"  {sf}->{st}: demeth={n_d}, remeth={n_r}, stable={n_s}, direction={direction}")

# ── B3: Non-randomness ─────────────────────────────────────────────────────────
print("\n=== B3: Non-randomness (top-K velocity DMRs) ===")
b3_results = {}
for sf, st in PRE8_TRANS:
    xf = stage_vecs[sf]; xt = stage_vecs[st]
    abs_delta = np.abs(xt - xf)
    label = TRANS_LABELS[(sf,st)]
    sorted_idx = np.argsort(-abs_delta)
    all_abs = abs_delta[np.isfinite(abs_delta)]
    trans_res = {}
    for k in [10, 25, 50]:
        obs = float(abs_delta[sorted_idx[:k]].mean())
        nulls = np.array([float(rng.choice(all_abs,size=k,replace=False).mean()) for _ in range(N_BOOT)])
        q95 = float(np.quantile(nulls,0.95)); pp = float((nulls>=obs).mean())
        trans_res[f"top{k}"] = {"obs_mean":obs,"null_q95":q95,"perm_p":pp,"significant":bool(obs>q95)}
        print(f"  {sf}->{st} top{k}: obs={obs:.4f}, null_q95={q95:.4f}, sig={obs>q95}")
    b3_results[label] = trans_res

# ── B4: Module impulse ─────────────────────────────────────────────────────────
print("\n=== B4: Module impulse J_M,k ===")
modules_all = sorted(set(mod_map.values()) - {"?"})
impulse_rows = []
for sf, st in PRE8_TRANS + [("8-cell","morula"),("morula","blastocyst")]:
    xf = stage_vecs[sf]; xt = stage_vecs[st]
    delta = xt - xf
    label = TRANS_LABELS[(sf,st)]
    for mid in modules_all:
        idx_m = [i for i,c in enumerate(clusters) if mod_map.get(c,"?")==mid]
        if not idx_m: continue
        d_m = delta[idx_m]; valid = np.isfinite(d_m)
        if valid.sum()==0: continue
        impulse_rows.append({"transition":label,"module":mid,"n_dmr":len(idx_m),
                              "J_k":float(norm(d_m[valid])),"dir_mean":float(np.nanmean(d_m)),
                              "is_priority":mid in ["M01","M02","M05","M10","M12"],"is_M00":mid=="M00"})

impulse_df = pd.DataFrame(impulse_rows)
impulse_df.to_csv(OUT/"b4_module_impulse_all_transitions.tsv", sep="\t", index=False)

print("Top 5 modules by J_k per transition:")
for trans in impulse_df["transition"].unique():
    sub = impulse_df[impulse_df["transition"]==trans].nlargest(5,"J_k")
    print(f"  {trans}: " + ", ".join([f"{r['module']}({r['J_k']:.3f})" for _,r in sub.iterrows()]))

# Module relay
entry_top5 = set(impulse_df[impulse_df["transition"]=="8cell_to_morula"].nlargest(5,"J_k")["module"].tolist())
print(f"\nEntry top5 modules: {entry_top5}")
for trans in ["oocyte_to_zygote","zygote_to_2cell","2cell_to_4cell","4cell_to_8cell"]:
    sub_top5 = set(impulse_df[impulse_df["transition"]==trans].nlargest(5,"J_k")["module"].tolist())
    overlap = sub_top5 & entry_top5
    print(f"  {trans} overlap with entry top5: {overlap} ({len(overlap)}/5)")

# ── B5: Direction sensitivity ──────────────────────────────────────────────────
print("\n=== B5: Direction sensitivity (stage-order permutation) ===")
pre8_stages = ["MII oocyte","zygote/PN","2-cell","4-cell","8-cell"]
pre8_vecs = [stage_vecs[s] for s in pre8_stages]

def cumulative_rmse(vecs, alpha, bias):
    total_sq = 0; n_total = 0
    for i in range(len(vecs)-1):
        xf=vecs[i]; xt=vecs[i+1]; pred=alpha*xf+bias
        v=np.isfinite(pred)&np.isfinite(xt)
        total_sq+=np.sum((pred[v]-xt[v])**2); n_total+=v.sum()
    return float(np.sqrt(total_sq/n_total)) if n_total>0 else np.nan

true_rmse = cumulative_rmse(pre8_vecs, alpha_op, bias_op)
internal = pre8_vecs[1:-1]
null_rmses = []
for _ in range(N_BOOT):
    perm = [internal[i] for i in rng.permutation(len(internal))]
    null_rmses.append(cumulative_rmse([pre8_vecs[0]]+perm+[pre8_vecs[-1]], alpha_op, bias_op))
null_rmses = np.array(null_rmses)
null_q05 = float(np.quantile(null_rmses,0.05))
pp_order = float((null_rmses<=true_rmse).mean())
rev_rmse = cumulative_rmse(list(reversed(pre8_vecs)), alpha_op, bias_op)
print(f"True order RMSE: {true_rmse:.4f}")
print(f"Reversed order RMSE: {rev_rmse:.4f}")
print(f"Random order null q05: {null_q05:.4f}")
print(f"Perm p (true <= null): {pp_order:.4f}")
print(f"True order optimal: {true_rmse < null_q05}")

b5_results = {"true_rmse":float(true_rmse),"reversed_rmse":float(rev_rmse),
              "null_q05":float(null_q05),"perm_p":float(pp_order),"optimal":bool(true_rmse<null_q05)}

# ── B6: Chromatin bio (RNA coupling) ──────────────────────────────────────────
print("\n=== B6: RNA expression coupling ===")
rna_map = rna.set_index("gene_name")
ann_map = ann.set_index("cluster_name")
b6_results = {}
stage_col_map = {"MII oocyte":"oocyte","zygote/PN":"zygote","2-cell":"2-cell",
                 "4-cell":"4-cell","8-cell":"8-cell","morula":"morula","blastocyst":"blastocyst"}
for sf, st in PRE8_TRANS:
    xf = stage_vecs[sf]; xt = stage_vecs[st]
    abs_delta = np.abs(xt-xf)
    label = TRANS_LABELS[(sf,st)]
    top25_clusters = [clusters[i] for i in np.argsort(-abs_delta)[:25]]
    top25_genes = [ann_map.loc[c,"nearest_gene"] if c in ann_map.index else None for c in top25_clusters]
    top25_genes = [g for g in top25_genes if g is not None and not pd.isna(g)]
    st_col = stage_col_map.get(st, st)
    if st_col in rna.columns:
        top_expr = [float(rna_map.loc[g,st_col]) for g in top25_genes if g in rna_map.index and st_col in rna_map.columns]
        all_expr = rna[st_col].dropna().values
        if len(top_expr)>3:
            t, p = stats.ttest_ind(top_expr, all_expr)
            print(f"  {sf}->{st}: top25 genes at {st}: mean={np.mean(top_expr):.2f} vs all={np.mean(all_expr):.2f}, p={p:.4f}")
            b6_results[label] = {"top25_mean":float(np.mean(top_expr)),"all_mean":float(np.mean(all_expr)),"p":float(p)}

# ── B7: Transition similarity matrix ──────────────────────────────────────────
print("\n=== B7: Transition similarity matrix ===")
all_trans = PRE8_TRANS + [("8-cell","morula"),("morula","blastocyst")]
vel_vecs = {}
for sf, st in all_trans:
    vel_vecs[TRANS_LABELS[(sf,st)]] = stage_vecs[st] - stage_vecs[sf]

trans_labels = [TRANS_LABELS[t] for t in all_trans]
n_t = len(trans_labels)
sim_mat = np.zeros((n_t,n_t))
for i,l1 in enumerate(trans_labels):
    for j,l2 in enumerate(trans_labels):
        v1=vel_vecs[l1]; v2=vel_vecs[l2]
        valid=np.isfinite(v1)&np.isfinite(v2)
        if valid.sum()>10:
            sim_mat[i,j] = float(np.dot(v1[valid],v2[valid])/(norm(v1[valid])*norm(v2[valid])+1e-12))

sim_df = pd.DataFrame(sim_mat, index=trans_labels, columns=trans_labels)
sim_df.to_csv(OUT/"b7_transition_similarity_matrix.tsv", sep="\t")
print(sim_df.round(3).to_string())

print("\nSimilarity to 8cell->morula:")
idx_m = trans_labels.index("8cell_to_morula")
for i,l in enumerate(trans_labels):
    if l!="8cell_to_morula":
        print(f"  {l}: {sim_mat[i,idx_m]:.4f}")

# ── B8: LOTO robustness ────────────────────────────────────────────────────────
print("\n=== B8: LOTO robustness ===")
loto_results = {}
for leave_idx in range(1, len(pre8_stages)-1):
    left_out = pre8_stages[leave_idx]
    x_true = stage_vecs[left_out]
    x_prev = stage_vecs[pre8_stages[leave_idx-1]]
    x_next = stage_vecs[pre8_stages[leave_idx+1]]
    x_interp = (x_prev+x_next)/2
    x_op = alpha_op*x_prev+bias_op
    v = np.isfinite(x_interp)&np.isfinite(x_true)
    rmse_i = float(np.sqrt(np.mean((x_interp[v]-x_true[v])**2)))
    rmse_p = float(np.sqrt(np.mean((x_prev[v]-x_true[v])**2)))
    rmse_o = float(np.sqrt(np.mean((x_op[v]-x_true[v])**2)))
    print(f"  Leave out {left_out}: interp={rmse_i:.4f}, prev={rmse_p:.4f}, op={rmse_o:.4f}")
    loto_results[left_out] = {"rmse_interp":rmse_i,"rmse_prev":rmse_p,"rmse_op":rmse_o,
                               "interp_beats_prev":bool(rmse_i<rmse_p),"op_beats_prev":bool(rmse_o<rmse_p)}

# ── Save ───────────────────────────────────────────────────────────────────────
all_results = {
    "date":"2026-05-29","model":"upstream erasure branch (zygote->8cell)",
    "B1":b1_results,"B2":cdiag_per_trans,"B3":b3_results,
    "B5":b5_results,"B6":b6_results,"B8":loto_results,
}
with open(OUT/"pre8_dynamics_all_results.json","w",encoding="utf-8") as f:
    json.dump(all_results, f, indent=2, ensure_ascii=False, default=str)

print(f"\nSaved to {OUT}/")
print(f"Files: {len(list(OUT.iterdir()))}")
```
