## Supplemental code for "An entropy-based diagnostic framework for characterizing methylation state dynamics during preimplantation development": 5_29_progress__remeth_prediction_framework.py

```
#!/usr/bin/env python
"""
Build unified re-methylation prediction framework for morula->blastocyst.

Goal: Create a basin-occupancy analog for the blastocyst transition.
The key metric: can external signals predict which morula-zero DMRs re-methylate?

Baseline: predict all morula-zero DMRs stay at zero = 58.8% accuracy (50 correct/85)
Target: exceed baseline with external signal-augmented prediction

Signals tested:
1. H3K27me3 blastocyst (Polycomb repression -> de novo methylation recruiter)
2. H3K27me3 8-cell (pre-existing Polycomb state)
3. H3K4me3 blastocyst (active marks - anti-correlated with methylation)
4. H3K27ac blastocyst (active enhancers)
5. H3K4me3 8-cell
6. Liu2019 morula accessibility
7. Multi-signal combination

Framework mirrors 8cell->morula:
  Entry: occupancy 0.044 vs 0.875, top-k matched-random controls
  Exit:  re-methylation accuracy, top-k controls, matched random baseline
"""
from __future__ import annotations
import json
from pathlib import Path
import numpy as np
import pandas as pd
from scipy import stats
from sklearn.metrics import roc_auc_score

OUT = Path("E:/5_29_progress")
OUT.mkdir(parents=True, exist_ok=True)

TRAJ  = Path("E:/CSB_TRO_operator_time_DMR_dynamics_2026-05-25/results/CSB_TRO_DMR_stage_mean_trajectory.tsv")
RESID = Path("E:/CSB_TRO_operator_time_DMR_dynamics_2026-05-25/results/CSB_TRO_basin_residual_DMR_ranking.tsv")
META  = Path("E:/CSB_TRO_operator_time_DMR_dynamics_2026-05-25/results/CSB_TRO_DMR_metadata.tsv")
MS    = Path("E:/5_28_progress/CSB_TRO_5_28_dmr_multistage_accessibility.tsv")

HIST = Path("E:/CSB_TRO_operator_time_DMR_dynamics_2026-05-25/external/histone")

SEED = 42
N_BOOT = 2000

print("Loading data...")
traj  = pd.read_csv(TRAJ, sep="\t")
resid = pd.read_csv(RESID, sep="\t")
meta  = pd.read_csv(META, sep="\t")
ms    = pd.read_csv(MS, sep="\t")

stage_means = {}
for s, g in traj.groupby("stage"):
    stage_means[s] = g.set_index("cluster_name")["mean_beta"].to_dict()

clusters = sorted(resid["cluster_name"].tolist())
x_morula = np.array([stage_means.get("morula",{}).get(c,np.nan) for c in clusters])
x_blast  = np.array([stage_means.get("blastocyst",{}).get(c,np.nan) for c in clusters])
x_8cell  = np.array([stage_means.get("8-cell",{}).get(c,np.nan) for c in clusters])

meta_map = meta.set_index("cluster_name")
ms_map   = ms.set_index("cluster_name")
mod_map  = resid.set_index("cluster_name")["module_id"].to_dict()

# ── Define re-methylation DMR set ─────────────────────────────────────────────
# S_remeth: morula<=0.02 AND blast>0.05
# S_zero_stay: morula<=0.02 AND blast<=0.05
REMETH_THRESH_MO   = 0.02   # morula threshold
REMETH_THRESH_BLAST = 0.05  # blastocyst threshold

is_mzero  = (x_morula <= REMETH_THRESH_MO) & np.isfinite(x_blast)
is_remeth = is_mzero & (x_blast > REMETH_THRESH_BLAST)
is_stay   = is_mzero & (x_blast <= REMETH_THRESH_BLAST)

n_mzero  = int(is_mzero.sum())
n_remeth = int(is_remeth.sum())
n_stay   = int(is_stay.sum())
baseline_acc = n_stay / n_mzero  # predict all stay = baseline

print(f"morula-zero DMRs: {n_mzero}")
print(f"  re-methylate (blast>{REMETH_THRESH_BLAST}): {n_remeth} ({n_remeth/n_mzero:.1%})")
print(f"  stay zero: {n_stay} ({n_stay/n_mzero:.1%})")
print(f"Baseline accuracy (predict all stay): {baseline_acc:.3f}")


# ── Helper: overlap peaks with DMR coords ─────────────────────────────────────
def overlap_peaks(peaks_df, clusters, meta_map, value_col="score"):
    """Return per-DMR mean and binary overlap."""
    peaks_df["chr"] = peaks_df["chr"].astype(str).str.strip()
    by_chr = {c: g for c, g in peaks_df.groupby("chr")}
    rows = []
    for c in clusters:
        if c not in meta_map.index:
            rows.append({"cluster_name": c, "overlap": 0, "score_max": np.nan})
            continue
        chrom = str(meta_map.loc[c, "chr"]).strip()
        ds, de = int(meta_map.loc[c, "start"]), int(meta_map.loc[c, "end"])
        if chrom not in by_chr:
            rows.append({"cluster_name": c, "overlap": 0, "score_max": np.nan})
            continue
        sub = by_chr[chrom]
        hits = sub[(sub["start"] < de) & (sub["end"] > ds)]
        sc = pd.to_numeric(hits[value_col], errors="coerce").dropna()
        rows.append({
            "cluster_name": c,
            "overlap": int(len(hits) > 0),
            "score_max": float(sc.max()) if len(sc) > 0 else np.nan,
        })
    return pd.DataFrame(rows)


print("\nComputing histone overlaps...")
hist_signals = {}
for label, fname, ncols in [
    ("k27me3_blast",  "H3K27me3_blastocyst.hg19.bed.gz", 5),
    ("k27me3_8cell",  "H3K27me3_8cell.hg19.bed.gz",      5),
    ("k4me3_blast",   "H3K4me3_blastocyst.hg19.bed.gz",  5),
    ("k27ac_blast",   "H3K27ac_blastocyst.hg19.bed.gz",  5),
    ("k4me3_8cell",   "H3K4me3_8cell.hg19.bed.gz",       5),
]:
    path = HIST / fname
    try:
        df = pd.read_csv(path, sep="\t", header=None, compression="gzip",
                         names=["chr","start","end","name","score"][:ncols],
                         usecols=list(range(ncols)))
        ov = overlap_peaks(df, clusters, meta_map)
        hist_signals[label] = ov.set_index("cluster_name")
        n_ov = ov["overlap"].sum()
        print(f"  {label}: {len(df)} peaks, {n_ov}/156 DMRs overlap")
    except Exception as e:
        print(f"  {label}: error {e}")

# Accessibility signals
acc_morula = np.array([ms_map.loc[c,"acc_morula_mean"] if c in ms_map.index else np.nan for c in clusters])
acc_delta  = np.array([ms_map.loc[c,"acc_morula_mean"] - ms_map.loc[c,"acc_8-cell_mean"]
                        if c in ms_map.index else np.nan for c in clusters])


# ── Core framework: re-methylation prediction accuracy ──────────────────────────
# For each signal, test: among morula-zero DMRs, does signal predict re-methylation?
# Metric: AUC (continuous signal) or accuracy improvement (binary)
# Bootstrap null: shuffle labels among morula-zero DMRs

rng = np.random.default_rng(SEED)

print("\n" + "="*60)
print("RE-METHYLATION PREDICTION FRAMEWORK")
print("Analogous to basin occupancy in 8cell->morula")
print("="*60)

y_remeth = is_remeth[is_mzero].astype(int)  # labels: 1=remeth, 0=stay
print(f"\nPrediction target: {y_remeth.sum()}/{len(y_remeth)} re-methylate among morula-zero DMRs")
print(f"Baseline accuracy (all-zero prediction): {baseline_acc:.3f}")
print(f"Random AUC expectation: 0.500")

all_signal_results = {}

def evaluate_signal(signal_arr, is_mzero, y_remeth, label, n_boot=N_BOOT):
    """Evaluate signal for re-methylation prediction."""
    x_sig = signal_arr[is_mzero]
    valid = np.isfinite(x_sig)
    if valid.sum() < 5:
        return None
    x_v = x_sig[valid]
    y_v = y_remeth[valid]
    if y_v.sum() < 3 or (y_v == 0).sum() < 3:
        return None

    # AUC
    try:
        auc = float(roc_auc_score(y_v, x_v))
    except:
        auc = np.nan

    # Spearman
    rho, p = stats.spearmanr(x_v, y_v)

    # Bootstrap null AUC
    null_aucs = []
    for _ in range(n_boot):
        y_perm = rng.permutation(y_v)
        try:
            null_aucs.append(float(roc_auc_score(y_perm, x_v)))
        except:
            pass
    null_aucs = np.array(null_aucs)
    auc_q95 = float(np.quantile(null_aucs, 0.95))
    auc_q05 = float(np.quantile(null_aucs, 0.05))
    perm_p_high = float((null_aucs >= auc).mean())
    perm_p_low  = float((null_aucs <= auc).mean())
    perm_p = min(perm_p_high, perm_p_low) * 2  # two-sided
    sig = (auc > auc_q95) or (auc < auc_q05)

    print(f"  {label}: AUC={auc:.4f} (null q05={auc_q05:.4f} q95={auc_q95:.4f}), "
          f"rho={rho:.4f}, p={p:.4f}, sig={sig}")

    return {
        "label": label, "n_valid": int(valid.sum()),
        "auc": float(auc), "null_q05": float(auc_q05), "null_q95": float(auc_q95),
        "perm_p": float(perm_p), "significant": bool(sig),
        "rho": float(rho), "p": float(p),
    }

print("\n--- Histone signals ---")
for sig_label, sig_df in hist_signals.items():
    sig_arr = np.array([sig_df.loc[c,"overlap"] if c in sig_df.index else 0 for c in clusters]).astype(float)
    r = evaluate_signal(sig_arr, is_mzero, y_remeth, sig_label)
    if r: all_signal_results[sig_label] = r

print("\n--- Histone scores (continuous) ---")
for sig_label, sig_df in hist_signals.items():
    sig_arr = np.array([sig_df.loc[c,"score_max"] if c in sig_df.index else np.nan
                         for c in clusters])
    r = evaluate_signal(sig_arr, is_mzero, y_remeth, f"{sig_label}_score")
    if r: all_signal_results[f"{sig_label}_score"] = r

print("\n--- Accessibility signals ---")
for sig_label, sig_arr in [("acc_morula", acc_morula), ("delta_acc", acc_delta)]:
    r = evaluate_signal(sig_arr, is_mzero, y_remeth, sig_label)
    if r: all_signal_results[sig_label] = r

print("\n--- Combined signals ---")
# Key hypothesis: H3K27me3 at blastocyst marks re-methylation sites
# (Polycomb-associated loci get de novo methylated at blastocyst)
# Also test: H3K27me3_blast * (1 - acc_morula) = closed chromatin + Polycomb
if "k27me3_blast" in hist_signals:
    k27me3_blast_arr = np.array([hist_signals["k27me3_blast"].loc[c,"overlap"]
                                  if c in hist_signals["k27me3_blast"].index else 0
                                  for c in clusters]).astype(float)

    # Combined: k27me3 + NOT acc_morula
    acc_standardized = np.where(np.isfinite(acc_morula), acc_morula, 0)
    acc_inv = -acc_standardized  # invert: low acc = good for remeth
    combo = k27me3_blast_arr + 0.5 * acc_inv
    r = evaluate_signal(combo, is_mzero, y_remeth, "k27me3_blast+inv_acc")
    if r: all_signal_results["k27me3_blast+inv_acc"] = r

    # Delta H3K27me3: blast - 8cell (if available)
    if "k27me3_8cell" in hist_signals:
        k27me3_8_arr = np.array([hist_signals["k27me3_8cell"].loc[c,"overlap"]
                                   if c in hist_signals["k27me3_8cell"].index else 0
                                   for c in clusters]).astype(float)
        delta_k27me3 = k27me3_blast_arr - k27me3_8_arr
        r = evaluate_signal(delta_k27me3, is_mzero, y_remeth, "delta_k27me3_blast_minus_8cell")
        if r: all_signal_results["delta_k27me3"] = r

# Also test morula H3K27me3 (quantitative bins) if available
# We know it only covers chr1 up to 72.9Mb, so limited
try:
    morula_k27me3_bins = pd.read_csv(
        "E:/CSB_TRO_operator_time_DMR_dynamics_2026-05-25/external/histone/H3K27me3_morula.hg19.bed.gz",
        sep="\t", header=None, compression="gzip",
        names=["chr","start","end","signal"],
        dtype={"chr":str,"start":int,"end":int,"signal":float})
    mo_by_chr = {c:g for c,g in morula_k27me3_bins.groupby("chr")}
    mo_signals = []
    for c in clusters:
        if c not in meta_map.index:
            mo_signals.append(np.nan); continue
        chrom = str(meta_map.loc[c,"chr"]).strip()
        ds, de = int(meta_map.loc[c,"start"]), int(meta_map.loc[c,"end"])
        if chrom not in mo_by_chr:
            mo_signals.append(np.nan); continue
        sub = mo_by_chr[chrom]
        hits = sub[(sub["start"]<de)&(sub["end"]>ds)]
        mo_signals.append(float(hits["signal"].mean()) if len(hits)>0 else np.nan)
    mo_sig_arr = np.array(mo_signals)
    n_valid_mo = np.isfinite(mo_sig_arr[is_mzero]).sum()
    if n_valid_mo > 5:
        r = evaluate_signal(mo_sig_arr, is_mzero, y_remeth, "k27me3_morula_signal")
        if r: all_signal_results["k27me3_morula_signal"] = r
    else:
        print(f"  k27me3_morula_signal: only {n_valid_mo} valid (chr1 limited)")
except Exception as e:
    print(f"  k27me3_morula: {e}")


# ── Top-K signal test (mirror of top25 residual DMR in 8cell->morula) ─────────
print("\n" + "="*60)
print("TOP-K RE-METHYLATION DMR ACCURACY (mirror of matched-random control)")
print("="*60)

# Find best signal
best_sig = max(all_signal_results.items(), key=lambda x: abs(x[1]["auc"]-0.5))
best_label = best_sig[0]
best_result = best_sig[1]
print(f"\nBest signal: {best_label} (AUC={best_result['auc']:.4f})")

# For the best signal, do top-K analysis
if best_result["significant"]:
    best_arr = None
    if best_label.startswith("k27me3_blast"):
        best_arr = np.array([hist_signals["k27me3_blast"].loc[c,"overlap"]
                              if c in hist_signals["k27me3_blast"].index else 0
                              for c in clusters]).astype(float)
    elif best_label.startswith("k27me3_8cell"):
        best_arr = np.array([hist_signals["k27me3_8cell"].loc[c,"overlap"]
                              if c in hist_signals["k27me3_8cell"].index else 0
                              for c in clusters]).astype(float)

    if best_arr is not None:
        # Among morula-zero DMRs, sort by best_signal
        mzero_idx = np.where(is_mzero)[0]
        sig_vals = best_arr[mzero_idx]
        sorted_idx = mzero_idx[np.argsort(-sig_vals)]  # desc

        for k in [10, 20, 30]:
            if k > len(sorted_idx): continue
            top_k_idx = sorted_idx[:k]
            top_k_labels = is_remeth[top_k_idx]
            top_k_acc = float(top_k_labels.mean())

            # Matched-random baseline
            null_accs = []
            for _ in range(N_BOOT):
                rand_idx = rng.choice(mzero_idx, size=k, replace=False)
                null_accs.append(is_remeth[rand_idx].mean())
            null_accs = np.array(null_accs)
            null_q95 = float(np.quantile(null_accs, 0.95))
            null_max  = float(np.max(null_accs))
            sig_flag = top_k_acc > null_q95

            print(f"  Top{k} by {best_label}: acc={top_k_acc:.3f}, "
                  f"null_q95={null_q95:.3f}, null_max={null_max:.3f}, sig={sig_flag}")


# ── Build occupancy-analog metric ─────────────────────────────────────────────
print("\n" + "="*60)
print("OCCUPANCY-ANALOG METRIC: Re-methylation prediction accuracy")
print("="*60)

# The direct analog of morula basin occupancy for blastocyst:
# Baseline (methylation-only): predict all morula-zero stay at zero
#   accuracy = 50/85 = 58.8%
# Signal-augmented: use H3K27me3_blast to predict which will re-methylate
#   Goal: significantly exceed 58.8%

# Use the continuous signal (H3K27me3 blast score) for threshold-based classification
if "k27me3_blast_score" in all_signal_results:
    score_arr = np.array([hist_signals["k27me3_blast"].loc[c,"score_max"]
                           if c in hist_signals["k27me3_blast"].index else np.nan
                           for c in clusters])
    mzero_scores = score_arr[is_mzero]
    valid_score = np.isfinite(mzero_scores)
    x_sc = mzero_scores[valid_score]
    y_sc = y_remeth[valid_score]

    if len(x_sc) > 10 and y_sc.sum() > 3:
        # Scan thresholds
        print("\nThreshold scan (k27me3_blast_score):")
        for pct in [50, 60, 70, 80, 90]:
            thresh = np.percentile(x_sc[np.isfinite(x_sc)], pct) if np.isfinite(x_sc).sum()>0 else 0
            pred = (x_sc >= thresh).astype(int)
            acc = float((pred == y_sc).mean())
            print(f"  threshold at p{pct}: acc={acc:.3f} vs baseline={baseline_acc:.3f}")

# If no histone score signal is strong, use binary H3K27me3 overlap
print(f"\n--- Binary H3K27me3 blast overlap prediction ---")
k27me3_arr = np.array([hist_signals["k27me3_blast"].loc[c,"overlap"]
                        if c in hist_signals["k27me3_blast"].index else 0
                        for c in clusters]).astype(float) if "k27me3_blast" in hist_signals else np.zeros(len(clusters))

# Among morula-zero DMRs:
mzero_k27 = k27me3_arr[is_mzero]
y_labels  = y_remeth

# Accuracy of predicting re-methylation = H3K27me3 present
pred_binary = (mzero_k27 > 0).astype(int)
acc_k27 = float((pred_binary == y_labels).mean())
tp = int(((pred_binary==1) & (y_labels==1)).sum())
fp = int(((pred_binary==1) & (y_labels==0)).sum())
fn = int(((pred_binary==0) & (y_labels==1)).sum())
tn = int(((pred_binary==0) & (y_labels==0)).sum())
precision = tp/(tp+fp) if (tp+fp)>0 else np.nan
recall    = tp/(tp+fn) if (tp+fn)>0 else np.nan
f1        = 2*precision*recall/(precision+recall) if (precision+recall)>0 else np.nan

print(f"  Accuracy: {acc_k27:.3f} vs baseline {baseline_acc:.3f}")
print(f"  TP={tp}, FP={fp}, FN={fn}, TN={tn}")
print(f"  Precision={precision:.3f}, Recall={recall:.3f}, F1={f1:.3f}")

# Bootstrap null
null_accs = [((rng.permutation(mzero_k27)>0).astype(int) == y_labels).mean() for _ in range(N_BOOT)]
null_q95 = float(np.quantile(null_accs, 0.95))
perm_p_acc = float((np.array(null_accs) >= acc_k27).mean())
print(f"  Bootstrap null q95={null_q95:.3f}, perm_p={perm_p_acc:.4f}")
print(f"  Significant: {acc_k27 > null_q95}")


# ── CRITICAL: Use H3K27me3 blast to define the occupancy-analog ───────────────
# Mirror of 8cell->morula occupancy framework:
# Entry: top25 residual DMR occupancy 0.956 vs random max 0.200
# Exit:  top-K re-methylation DMR re-methylation rate vs random

print("\n" + "="*60)
print("OCCUPANCY FRAMEWORK MIRROR")
print("="*60)
print("8-cell->morula: top25 residual occupancy = 0.956 vs random max = 0.200")
print("morula->blast:  signal-defined DMR re-methylation rate vs random")
print()

# Use k27me3_blast overlap as the primary classifier
# Sort morula-zero DMRs by k27me3 status
has_k27me3 = mzero_k27 > 0
no_k27me3  = mzero_k27 == 0

remeth_rate_k27_yes = float(y_remeth[has_k27me3].mean()) if has_k27me3.sum()>0 else np.nan
remeth_rate_k27_no  = float(y_remeth[no_k27me3].mean()) if no_k27me3.sum()>0 else np.nan

print(f"morula-zero DMRs WITH H3K27me3 at blast: {has_k27me3.sum()} DMRs")
print(f"  Re-methylation rate: {remeth_rate_k27_yes:.3f}")
print(f"morula-zero DMRs WITHOUT H3K27me3 at blast: {no_k27me3.sum()} DMRs")
print(f"  Re-methylation rate: {remeth_rate_k27_no:.3f}")
print(f"Background re-methylation rate: {n_remeth/n_mzero:.3f}")
print()

# Bootstrap significance
null_k27yes_rates = []
for _ in range(N_BOOT):
    k27_perm = rng.permutation(mzero_k27)
    has_perm = k27_perm > 0
    if has_perm.sum() == 0: continue
    null_k27yes_rates.append(y_remeth[has_perm].mean())
null_k27yes_rates = np.array(null_k27yes_rates)
q95_rate = float(np.quantile(null_k27yes_rates, 0.95))
pp_rate  = float((null_k27yes_rates >= remeth_rate_k27_yes).mean())
print(f"Bootstrap: null q95={q95_rate:.3f}, perm_p={pp_rate:.4f}")
print(f"H3K27me3-marked DMRs re-methylate at higher rate: {remeth_rate_k27_yes > q95_rate}")

# Also test: can we combine H3K27me3 blast and acc_morula?
print("\n--- Combined classifier: H3K27me3(blast) + low acc(morula) ---")
# Low morula acc + H3K27me3 at blast = high chance of re-methylation
acc_valid = np.isfinite(acc_morula[is_mzero])
combo_score = mzero_k27 - 0.3 * np.where(np.isfinite(acc_morula[is_mzero]),
                                            acc_morula[is_mzero], 0)
try:
    auc_combo = float(roc_auc_score(y_remeth[acc_valid], combo_score[acc_valid]))
    null_combo = [float(roc_auc_score(rng.permutation(y_remeth[acc_valid]), combo_score[acc_valid]))
                  for _ in range(N_BOOT)]
    null_combo = np.array(null_combo)
    pp_combo = float((null_combo >= auc_combo).mean())
    q95_combo = float(np.quantile(null_combo, 0.95))
    print(f"  AUC={auc_combo:.4f}, null_q95={q95_combo:.4f}, perm_p={pp_combo:.4f}, sig={auc_combo>q95_combo}")
except Exception as e:
    print(f"  Error: {e}")


# ── Save all results ──────────────────────────────────────────────────────────
print("\n" + "="*60)
print("SAVING RESULTS")
print("="*60)

# Per-DMR table with all signals
per_dmr = pd.DataFrame({
    "cluster_name": clusters,
    "x_morula": x_morula,
    "x_blast": x_blast,
    "is_mzero": is_mzero.astype(int),
    "is_remeth": is_remeth.astype(int),
    "acc_morula": acc_morula,
    "module_id": [mod_map.get(c,"?") for c in clusters],
})
for sig_label, sig_df in hist_signals.items():
    per_dmr[f"{sig_label}_ov"] = [
        int(sig_df.loc[c,"overlap"]) if c in sig_df.index else 0 for c in clusters]
per_dmr.to_csv(OUT/"remeth_framework_per_dmr.tsv", sep="\t", index=False)

# Summary JSON
summary = {
    "date": "2026-05-29",
    "re_methylation_framework": {
        "n_morula_zero": n_mzero,
        "n_remeth": n_remeth,
        "n_stay": n_stay,
        "baseline_accuracy": float(baseline_acc),
        "re_methylation_rate": float(n_remeth/n_mzero),
    },
    "signal_results": all_signal_results,
    "h3k27me3_blast_occupancy_analog": {
        "n_k27me3_marked": int(has_k27me3.sum()),
        "remeth_rate_k27_yes": float(remeth_rate_k27_yes) if not np.isnan(remeth_rate_k27_yes) else None,
        "remeth_rate_k27_no": float(remeth_rate_k27_no) if not np.isnan(remeth_rate_k27_no) else None,
        "background_rate": float(n_remeth/n_mzero),
        "bootstrap_q95": float(q95_rate),
        "perm_p": float(pp_rate),
        "significant": bool(remeth_rate_k27_yes > q95_rate) if not np.isnan(remeth_rate_k27_yes) else False,
        "binary_accuracy": float(acc_k27),
        "binary_null_q95": float(null_q95),
        "binary_perm_p": float(perm_p_acc),
        "binary_significant": bool(acc_k27 > null_q95),
    },
}
with open(OUT/"remeth_prediction_framework.json","w",encoding="utf-8") as f:
    json.dump(summary, f, indent=2, ensure_ascii=False, default=str)

print(f"\nSaved: {OUT}/remeth_prediction_framework.json")
print(f"       {OUT}/remeth_framework_per_dmr.tsv")
print()
print("="*60)
print("FRAMEWORK SUMMARY")
print("="*60)
print(f"Baseline (predict all stay): {baseline_acc:.3f}")
print(f"Best AUC signal: {max(all_signal_results.items(), key=lambda x: abs(x[1]['auc']-0.5))[0]}")
print(f"H3K27me3 blast → re-meth rate: {remeth_rate_k27_yes:.3f} vs background {n_remeth/n_mzero:.3f}")
print(f"Significant: {bool(remeth_rate_k27_yes > q95_rate) if not np.isnan(remeth_rate_k27_yes) else False}")
```
