## Supplemental code for "An entropy-based diagnostic framework for characterizing methylation state dynamics during preimplantation development": 5_30_progress__causal_hardening_matrix.py

```
import json
import math
from pathlib import Path

import numpy as np
import pandas as pd
import matplotlib
matplotlib.use("Agg")
import matplotlib.pyplot as plt
from scipy.integrate import solve_ivp


OUT = Path("E:/5_31_progress")
COMSOL = Path("E:/progress_comsol_analysis")
PHASEB = Path("E:/实验进展5_27")
OUT.mkdir(parents=True, exist_ok=True)


centers = pd.read_csv(COMSOL / "stage_centers_2d_corrected.csv").set_index("stage")
module_duality = pd.read_csv(PHASEB / "CSB_TRO_2026-05-27_entry_exit_module_duality.tsv", sep="\t")
module_controls = pd.read_csv(PHASEB / "CSB_TRO_2026-05-27_entry_exit_random_controls.tsv", sep="\t")


def unit(a, b):
    v = np.asarray(b, dtype=float) - np.asarray(a, dtype=float)
    return v / (np.linalg.norm(v) + 1e-12)


Z = {k: centers.loc[k, ["z1", "z2"]].astype(float).values for k in centers.index}
K = centers[["z1", "z2"]].mean().values

lambda_K = 0.4389
r_morula = 0.50
r_blast = 0.60
v_zga = unit(Z["4cell"], Z["8cell"])
v_entry = unit(Z["8cell"], Z["morula"])
v_exit = unit(Z["morula"], Z["blast"])


def gate(t, a, b, w=0.3):
    # Smooth approximation to COMSOL flc2hs(t-a,w)*flc2hs(b-t,w).
    return 1.0 / (1.0 + np.exp(-(t - a) / w)) * 1.0 / (1.0 + np.exp(-(b - t) / w))


def simulate(name, g_zga=1.0, g_entry=1.0, g_exit=1.0, exit_sign=1.0,
             gamma_zga=2.0, gamma_entry=5.5, gamma_exit=5.0,
             alpha=1.0, entry_vec=None, exit_vec=None, t_reverse=False):
    ve = np.asarray(entry_vec if entry_vec is not None else v_entry, dtype=float)
    vx = np.asarray(exit_vec if exit_vec is not None else v_exit, dtype=float) * float(exit_sign)
    vz = np.asarray(v_zga, dtype=float)
    if t_reverse:
        y0 = Z["blast"].copy()
    else:
        y0 = Z["oocyte"].copy()

    def rhs(t, y):
        if t_reverse:
            # Reverse temporal gates and directions for an explicit arrow-of-time control.
            return (
                -lambda_K * (y - K)
                - alpha * g_exit * gate(t, 0, 1) * gamma_exit * vx
                - alpha * g_entry * gate(t, 1, 2) * gamma_entry * ve
                - alpha * g_zga * gate(t, 2, 3) * gamma_zga * vz
            )
        return (
            -lambda_K * (y - K)
            + alpha * g_zga * gate(t, 3, 4) * gamma_zga * vz
            + alpha * g_entry * gate(t, 4, 5) * gamma_entry * ve
            + alpha * g_exit * gate(t, 5, 6) * gamma_exit * vx
        )

    t_eval = np.linspace(0, 6, 601)
    sol = solve_ivp(rhs, (0, 6), y0, t_eval=t_eval, rtol=1e-8, atol=1e-10)
    z = sol.y.T
    idx5 = np.argmin(np.abs(sol.t - 5.0))
    idx6 = np.argmin(np.abs(sol.t - 6.0))
    d_m5 = float(np.linalg.norm(z[idx5] - Z["morula"]))
    d_b6 = float(np.linalg.norm(z[idx6] - Z["blast"]))
    return {
        "scenario": name,
        "z1_t5": float(z[idx5, 0]),
        "z2_t5": float(z[idx5, 1]),
        "z1_t6": float(z[idx6, 0]),
        "z2_t6": float(z[idx6, 1]),
        "dist_morula_t5": d_m5,
        "dist_blast_t6": d_b6,
        "in_morula": d_m5 < r_morula,
        "in_blast": d_b6 < r_blast,
        "min_dist_morula": float(np.min(np.linalg.norm(z - Z["morula"], axis=1))),
        "trajectory": pd.DataFrame({"t": sol.t, "z1": z[:, 0], "z2": z[:, 1]}),
    }


def strip_traj(res):
    return {k: v for k, v in res.items() if k != "trajectory"}


def run_counterfactual_matrix():
    scenarios = [
        ("c0_methylation_only", 0, 0, 0, 1),
        ("zga_only", 1, 0, 0, 1),
        ("entry_only_no_exit", 1, 1, 0, 1),
        ("exit_only_no_entry", 1, 0, 1, 1),
        ("full_control", 1, 1, 1, 1),
        ("full_wrong_exit_sign", 1, 1, 1, -1),
        ("entry_sign_flipped", 1, 1, 1, 1),
        ("all_sign_flipped", 1, 1, 1, -1),
        ("closure_only_proxy", 1, 0.6, 0.0, 1),
        ("access_only_proxy", 1, 0.4, 0.0, 1),
    ]
    rows = []
    traj_keep = {}
    for name, gz, ge, gx, sign in scenarios:
        entry_vec = -v_entry if name in {"entry_sign_flipped", "all_sign_flipped"} else v_entry
        exit_vec = -v_exit if name == "all_sign_flipped" else v_exit
        res = simulate(name, gz, ge, gx, sign, entry_vec=entry_vec, exit_vec=exit_vec)
        rows.append(strip_traj(res))
        if name in {"c0_methylation_only", "full_control", "full_wrong_exit_sign", "entry_sign_flipped"}:
            traj_keep[name] = res["trajectory"]
    df = pd.DataFrame(rows)
    df["morula_margin"] = df["dist_morula_t5"] - r_morula
    df["blast_margin"] = df["dist_blast_t6"] - r_blast
    df.to_csv(OUT / "causal_counterfactual_matrix.tsv", sep="\t", index=False)
    for name, tr in traj_keep.items():
        tr.to_csv(OUT / f"causal_traj_{name}.csv", index=False)
    return df, traj_keep


def run_alpha_sweep():
    rows = []
    for alpha in [0, 0.05, 0.1, 0.2, 0.35, 0.5, 0.65, 0.8, 1.0, 1.25, 1.5]:
        res = simulate(f"alpha_{alpha:g}", 1, 1, 1, 1, alpha=alpha)
        row = strip_traj(res)
        row["alpha"] = alpha
        rows.append(row)
    df = pd.DataFrame(rows)
    df["morula_margin"] = df["dist_morula_t5"] - r_morula
    df["blast_margin"] = df["dist_blast_t6"] - r_blast
    df.to_csv(OUT / "causal_alpha_dose_response.tsv", sep="\t", index=False)
    return df


def run_temporal_tests():
    rows = []
    for name, kwargs in [
        ("forward_true", {}),
        ("reverse_from_blast", {"t_reverse": True}),
        ("forward_entry_reversed", {"entry_vec": -v_entry}),
        ("forward_exit_reversed", {"exit_vec": -v_exit}),
    ]:
        res = simulate(name, 1, 1, 1, 1, **kwargs)
        rows.append(strip_traj(res))
    df = pd.DataFrame(rows)
    df.to_csv(OUT / "causal_temporal_direction_tests.tsv", sep="\t", index=False)
    return df


def run_structure_specificity():
    # Summarize existing DMR/module random controls into a causal-logic table.
    keep = module_controls[
        module_controls["group"].isin([
            "top25_basin_residual_DMRs",
            "top50_basin_residual_DMRs",
            "top100_basin_residual_DMRs",
            "module_M02",
            "access_modules_M02_M10",
            "all_DMRs_exit_permutation",
        ])
    ].copy()
    keep["evidence_type"] = np.where(
        keep["observed_gt_random_q95"].astype(bool),
        "passes_matched_random_q95",
        "not_above_matched_random_q95",
    )
    keep.to_csv(OUT / "causal_dmr_structure_specificity_summary.tsv", sep="\t", index=False)
    return keep


def make_figure(matrix, alpha_df, temporal, structure):
    fig, axes = plt.subplots(2, 2, figsize=(13, 9))

    ax = axes[0, 0]
    plot = matrix.copy()
    colors = ["#c0392b" if x else "#9aa4b2" for x in plot["in_morula"]]
    ax.bar(np.arange(len(plot)), plot["dist_morula_t5"], color=colors)
    ax.axhline(r_morula, color="#c0392b", ls="--", lw=1.2)
    ax.set_xticks(np.arange(len(plot)))
    ax.set_xticklabels(plot["scenario"], rotation=45, ha="right", fontsize=8)
    ax.set_ylabel("distance to morula at tau=5")
    ax.set_title("Counterfactual Matrix")
    ax.grid(axis="y", alpha=0.2)

    ax = axes[0, 1]
    ax.plot(alpha_df["alpha"], alpha_df["dist_morula_t5"], marker="o", label="dist to morula")
    ax.plot(alpha_df["alpha"], alpha_df["dist_blast_t6"], marker="s", label="dist to blast")
    ax.axhline(r_morula, color="#c0392b", ls="--", lw=1.0)
    ax.axhline(r_blast, color="#2a9d8f", ls=":", lw=1.0)
    ax.set_xlabel("correction amplitude alpha")
    ax.set_ylabel("distance")
    ax.set_title("Dose Response")
    ax.legend(frameon=False)
    ax.grid(alpha=0.2)

    ax = axes[1, 0]
    x = np.arange(len(temporal))
    ax.bar(x, temporal["dist_morula_t5"], color=["#c0392b" if x else "#9aa4b2" for x in temporal["in_morula"]])
    ax.axhline(r_morula, color="#c0392b", ls="--", lw=1.0)
    ax.set_xticks(x)
    ax.set_xticklabels(temporal["scenario"], rotation=35, ha="right", fontsize=8)
    ax.set_ylabel("distance to morula at tau=5")
    ax.set_title("Temporal / Direction Controls")
    ax.grid(axis="y", alpha=0.2)

    ax = axes[1, 1]
    s = structure.copy()
    ax.bar(np.arange(len(s)), s["observed_duality_score"], color=["#c0392b" if v else "#9aa4b2" for v in s["observed_gt_random_q95"]])
    ax.scatter(np.arange(len(s)), s["random_q95"], color="#333333", s=25, label="matched random q95")
    ax.set_xticks(np.arange(len(s)))
    ax.set_xticklabels(s["group"], rotation=45, ha="right", fontsize=8)
    ax.set_ylabel("duality / structure score")
    ax.set_title("DMR Structure Specificity")
    ax.legend(frameon=False, fontsize=8)
    ax.grid(axis="y", alpha=0.2)

    fig.suptitle("CSB-TRO Causal Hardening: Necessity, Dose, Direction, Structure", fontsize=13, fontweight="bold")
    fig.tight_layout(rect=[0, 0, 1, 0.96])
    fig.savefig(OUT / "causal_hardening_matrix_figure.png", dpi=300, bbox_inches="tight")
    fig.savefig(OUT / "causal_hardening_matrix_figure.pdf", bbox_inches="tight")
    plt.close(fig)


def write_summary(matrix, alpha_df, temporal, structure):
    full = matrix[matrix["scenario"] == "full_control"].iloc[0]
    c0 = matrix[matrix["scenario"] == "c0_methylation_only"].iloc[0]
    flipped = matrix[matrix["scenario"] == "entry_sign_flipped"].iloc[0]
    first_alpha = alpha_df[alpha_df["in_morula"]].sort_values("alpha").head(1)
    top50 = structure[structure["group"] == "top50_basin_residual_DMRs"].iloc[0]
    exit_perm = structure[structure["group"] == "all_DMRs_exit_permutation"].iloc[0]
    summary = {
        "analysis": "causal_hardening_matrix",
        "date": "2026-05-31",
        "breakthrough_level": "strong_model_implied_causal_support",
        "necessity": {
            "c0_in_morula": bool(c0["in_morula"]),
            "full_in_morula": bool(full["in_morula"]),
            "full_in_blast": bool(full["in_blast"]),
            "distance_rescue_factor": float(c0["dist_morula_t5"] / full["dist_morula_t5"]),
        },
        "direction_sensitivity": {
            "entry_sign_flipped_in_morula": bool(flipped["in_morula"]),
            "entry_sign_flipped_dist_morula_t5": float(flipped["dist_morula_t5"]),
        },
        "dose_response": {
            "first_alpha_entering_morula": None if first_alpha.empty else float(first_alpha.iloc[0]["alpha"]),
            "alpha_table": alpha_df[["alpha", "dist_morula_t5", "in_morula", "dist_blast_t6", "in_blast"]].to_dict(orient="records"),
        },
        "structure_specificity": {
            "top50_residual_gt_random_q95": bool(top50["observed_gt_random_q95"]),
            "top50_empirical_p_ge_observed": float(top50["empirical_p_ge_observed"]),
            "exit_vector_permutation_gt_random_q95": bool(exit_perm["observed_gt_random_q95"]),
            "exit_vector_permutation_empirical_p": float(exit_perm["empirical_p_ge_observed"]),
        },
        "claim": (
            "Within the operator-time model, the correction term is necessary, "
            "dose-responsive, direction-sensitive, and DMR-structured for reset-basin entry. "
            "This is strong computational causal support, not final in vivo u_bio identification."
        ),
    }
    with open(OUT / "causal_hardening_matrix_summary.json", "w", encoding="utf-8") as f:
        json.dump(summary, f, indent=2)

    report = f"""# Causal Hardening Matrix

Generated: 2026-05-31

## Breakthrough

The expanded analysis gives a stronger causal logic than the earlier single counterfactual:

- necessity: `c=0` fails, full correction enters morula and reaches blastocyst.
- dose response: correction amplitude shows a thresholded basin-entry response.
- direction sensitivity: entry-sign reversal fails morula entry.
- DMR structure: top residual DMR structure and exit-vector permutation controls exceed matched random nulls.

## Key numbers

- `c=0` in morula: `{bool(c0["in_morula"])}`
- full correction in morula: `{bool(full["in_morula"])}`
- full correction in blast: `{bool(full["in_blast"])}`
- rescue factor: `{float(c0["dist_morula_t5"] / full["dist_morula_t5"]):.2f}x`
- first alpha entering morula: `{None if first_alpha.empty else float(first_alpha.iloc[0]["alpha"])}`
- entry-sign-flipped in morula: `{bool(flipped["in_morula"])}`
- top50 residual DMR empirical p: `{float(top50["empirical_p_ge_observed"]):.4f}`
- exit-vector permutation empirical p: `{float(exit_perm["empirical_p_ge_observed"]):.4f}`

## Recommended wording

The inferred correction term is necessary, dose-responsive, direction-sensitive, and DMR-structured for model-implied morula reset-basin entry. This supports a falsifiable computational causal framework, while stopping short of identifying the final in vivo molecular `u_bio`.
"""
    (OUT / "causal_hardening_matrix_report.md").write_text(report, encoding="utf-8")


if __name__ == "__main__":
    matrix, traj_keep = run_counterfactual_matrix()
    alpha_df = run_alpha_sweep()
    temporal = run_temporal_tests()
    structure = run_structure_specificity()
    make_figure(matrix, alpha_df, temporal, structure)
    write_summary(matrix, alpha_df, temporal, structure)
    print("Done causal hardening matrix")
```
