## Supplemental code for "An entropy-based diagnostic framework for characterizing methylation state dynamics during preimplantation development": 5_30_progress__complete_validation.py

```
#!/usr/bin/env python
"""
Complete independent validation analysis for E:/5_30_progress
Using Liu2019 4-stage accessibility as independent chromatin source
"""
import pandas as pd, numpy as np
from scipy import stats
from pathlib import Path
import json

OUT = Path("E:/5_30_progress")
OUT.mkdir(exist_ok=True)

traj  = pd.read_csv("E:/CSB_TRO_operator_time_DMR_dynamics_2026-05-25/results/CSB_TRO_DMR_stage_mean_trajectory.tsv", sep="\t")
resid = pd.read_csv("E:/CSB_TRO_operator_time_DMR_dynamics_2026-05-25/results/CSB_TRO_basin_residual_DMR_ranking.tsv", sep="\t")
ms    = pd.read_csv("E:/5_28_progress/CSB_TRO_5_28_dmr_multistage_accessibility.tsv", sep="\t")
meta  = pd.read_csv("E:/CSB_TRO_operator_time_DMR_dynamics_2026-05-25/results/CSB_TRO_DMR_metadata.tsv", sep="\t")

stage_means = {}
for s, g in traj.groupby("stage"):
    stage_means[s] = g.set_index("cluster_name")["mean_beta"].to_dict()

clusters = sorted(resid["cluster_name"].tolist())
ms_map   = ms.set_index("cluster_name")
meta_map = meta.set_index("cluster_name")
mod_map  = resid.set_index("cluster_name")["module_id"].to_dict()
rank_map = resid.set_index("cluster_name")["basin_residual_rank"].to_dict()

def svec(s):
    return np.array([stage_means.get(s,{}).get(c,np.nan) for c in clusters])

x_2cell  = svec("2-cell"); x_4cell = svec("4-cell")
x_8cell  = svec("8-cell"); x_morula = svec("morula"); x_blast = svec("blastocyst")

alpha_op = 0.5611; bias_op = 0.0688
rng = np.random.default_rng(42)
N = 3000

acc_2cell  = np.array([ms_map.loc[c,"acc_2-cell_mean"] if c in ms_map.index else np.nan for c in clusters])
acc_4cell  = np.array([ms_map.loc[c,"acc_4-cell_mean"] if c in ms_map.index else np.nan for c in clusters])
acc_8cell  = np.array([ms_map.loc[c,"acc_8-cell_mean"] if c in ms_map.index else np.nan for c in clusters])
acc_morula = np.array([ms_map.loc[c,"acc_morula_mean"] if c in ms_map.index else np.nan for c in clusters])
n_cpg = np.array([meta_map.loc[c,"n_cpg_target"] if c in meta_map.index else np.nan for c in clusters])
width = np.array([meta_map.loc[c,"width"] if c in meta_map.index else np.nan for c in clusters])

def partial_spearman(x, y, *controls):
    mask = np.isfinite(x) & np.isfinite(y)
    for c in controls:
        mask &= np.isfinite(c)
    if mask.sum() < 8:
        return np.nan
    X = np.column_stack([c[mask] for c in controls])
    rx = stats.rankdata(x[mask]); ry = stats.rankdata(y[mask])
    def resid_fn(r, Xm):
        Xa = np.column_stack([Xm, np.ones(len(r))])
        w, _, _, _ = np.linalg.lstsq(Xa, r, rcond=None)
        return r - Xa @ w
    return float(stats.pearsonr(resid_fn(rx, X), resid_fn(ry, X))[0])

print("="*60)
print("COMPLETE INDEPENDENT VALIDATION ANALYSIS")
print("Liu2019 4-stage accessibility as independent chromatin source")
print("="*60)

all_results = {}

# Test 1: Stage-specific acc-meth coupling
print("\nTEST 1: Stage-specific acc-meth coupling")
stage_coupling = {}
for stage, acc_arr, meth_arr in [
    ("2-cell", acc_2cell, x_2cell),
    ("4-cell", acc_4cell, x_4cell),
    ("8-cell", acc_8cell, x_8cell),
    ("morula", acc_morula, x_morula),
]:
    v = np.isfinite(acc_arr) & np.isfinite(meth_arr)
    rho, p = stats.spearmanr(acc_arr[v], meth_arr[v])
    nulls = np.array([stats.spearmanr(rng.permutation(acc_arr[v]), meth_arr[v])[0] for _ in range(N)])
    pp = float((nulls >= rho).mean())
    q95 = float(np.quantile(nulls, 0.95))
    sig = rho > q95
    pr = partial_spearman(acc_arr, meth_arr, width, n_cpg)
    stage_coupling[stage] = {"rho": float(rho), "p": float(p), "perm_p": float(pp),
                              "sig": bool(sig), "partial_rho": float(pr) if pr is not None else None}
    print(f"  {stage}: rho={rho:.4f}, perm_p={pp:.4f}, sig={sig}, partial_rho={pr:.4f}")
all_results["test1_stage_coupling"] = stage_coupling

# Test 2: ZGA-Reset coupling
print("\nTEST 2: ZGA-Reset coupling")
c_diag_48 = x_8cell - (alpha_op*x_4cell + bias_op)
c_diag_8m = x_morula - (alpha_op*x_8cell + bias_op)
v = np.isfinite(c_diag_48) & np.isfinite(c_diag_8m)
rho, p = stats.spearmanr(c_diag_48[v], c_diag_8m[v])
nulls = np.array([stats.spearmanr(rng.permutation(c_diag_48[v]), c_diag_8m[v])[0] for _ in range(N)])
pp = float((nulls <= rho).mean())
q05 = float(np.quantile(nulls, 0.05))
sig = rho < q05
print(f"  4->8 c_diag ~ 8->morula c_diag: rho={rho:.4f}, p={p:.6f}, perm_p={pp:.4f}, sig={sig}")
all_results["test2_zga_reset"] = {"rho": float(rho), "p": float(p), "perm_p": float(pp),
                                   "null_q05": float(q05), "sig": bool(sig)}

# Test 3: Morula acc ~ correction term
print("\nTEST 3: Morula accessibility ~ correction term")
strict_corr_map = resid.set_index("cluster_name")["observed_minus_strict_pred_delta_beta"].to_dict()
c_strict = np.array([strict_corr_map.get(c, np.nan) for c in clusters])
v = np.isfinite(acc_morula) & np.isfinite(c_strict)
rho, p = stats.spearmanr(acc_morula[v], c_strict[v])
nulls = np.array([stats.spearmanr(rng.permutation(acc_morula[v]), c_strict[v])[0] for _ in range(N)])
pp = float((nulls >= rho).mean())
q95 = float(np.quantile(nulls, 0.95))
sig = rho > q95
print(f"  morula_acc ~ strict_correction: rho={rho:.4f}, p={p:.4f}, perm_p={pp:.4f}, sig={sig}")
all_results["test3_acc_correction"] = {"rho": float(rho), "p": float(p), "perm_p": float(pp), "sig": bool(sig)}

# Test 4: Cross-validation
print("\nTEST 4: Internal cross-validation (1000 random splits)")
v = np.isfinite(acc_morula) & np.isfinite(x_morula)
acc_v = acc_morula[v]; meth_v = x_morula[v]
both_pos = 0
for _ in range(1000):
    idx = rng.permutation(len(acc_v))
    h1 = idx[:len(idx)//2]; h2 = idx[len(idx)//2:]
    r1, _ = stats.spearmanr(acc_v[h1], meth_v[h1])
    r2, _ = stats.spearmanr(acc_v[h2], meth_v[h2])
    if r1 > 0 and r2 > 0:
        both_pos += 1
print(f"  Both halves positive: {both_pos}/1000 (null expectation: ~250/1000)")
all_results["test4_cross_validation"] = {"both_positive": both_pos, "total": 1000, "expected_null": 250}

# Test 5: Top25 residual DMR enrichment
print("\nTEST 5: Top25 residual DMR morula accessibility enrichment")
top25_mask = np.array([rank_map.get(c, 999) <= 25 for c in clusters])
obs_mean = float(acc_morula[top25_mask & np.isfinite(acc_morula)].mean())
all_vals = acc_morula[np.isfinite(acc_morula)]
nulls = np.array([rng.choice(all_vals, size=int(top25_mask.sum()), replace=False).mean() for _ in range(N)])
q95 = float(np.quantile(nulls, 0.95))
pp = float((nulls >= obs_mean).mean())
sig = obs_mean > q95
print(f"  Top25 morula_acc mean={obs_mean:.4f} vs random q95={q95:.4f}, perm_p={pp:.4f}, sig={sig}")
all_results["test5_top25_enrichment"] = {"obs_mean": obs_mean, "null_q95": q95, "perm_p": float(pp), "sig": bool(sig)}

# Test 6: Inverted-U DMR coupling
print("\nTEST 6: Inverted-U DMR accessibility coupling")
if "is_inverted_u" in ms_map.columns:
    is_invU = np.array([bool(ms_map.loc[c,"is_inverted_u"]) if c in ms_map.index else False for c in clusters])
    if is_invU.sum() > 5:
        v = is_invU & np.isfinite(acc_morula) & np.isfinite(c_strict)
        rho, p = stats.spearmanr(acc_morula[v], c_strict[v])
        nulls = np.array([stats.spearmanr(rng.permutation(acc_morula[v]), c_strict[v])[0] for _ in range(N)])
        pp = float((nulls <= rho).mean())
        q05 = float(np.quantile(nulls, 0.05))
        sig = rho < q05
        print(f"  inverted-U acc ~ correction: rho={rho:.4f}, p={p:.4f}, perm_p={pp:.4f}, sig={sig}")
        all_results["test6_inverted_u"] = {"rho": float(rho), "p": float(p), "perm_p": float(pp), "sig": bool(sig)}

# Test 7: 4-stage accessibility trajectory vs methylation trajectory
print("\nTEST 7: 4-stage accessibility trajectory analysis")
# For each DMR, compute correlation between acc trajectory and meth trajectory
traj_corrs = []
for c in clusters:
    acc_traj = np.array([ms_map.loc[c, f"acc_{s}_mean"] if c in ms_map.index else np.nan
                          for s in ["2-cell","4-cell","8-cell","morula"]])
    meth_traj = np.array([stage_means.get(s,{}).get(c,np.nan)
                           for s in ["2-cell","4-cell","8-cell","morula"]])
    valid = np.isfinite(acc_traj) & np.isfinite(meth_traj)
    if valid.sum() >= 3:
        try:
            r, _ = stats.spearmanr(acc_traj[valid], meth_traj[valid])
            traj_corrs.append(float(r) if np.isfinite(r) else np.nan)
        except:
            traj_corrs.append(np.nan)
    else:
        traj_corrs.append(np.nan)

traj_corrs = np.array(traj_corrs)
valid_tc = np.isfinite(traj_corrs)
mean_tc = float(np.nanmean(traj_corrs))
t, p_tc = stats.ttest_1samp(traj_corrs[valid_tc], 0)
print(f"  Per-DMR trajectory correlation: mean={mean_tc:.4f}, t-test p={p_tc:.4f}")
all_results["test7_trajectory_correlation"] = {"mean": mean_tc, "ttest_p": float(p_tc), "n": int(valid_tc.sum())}

# Save all results
with open(OUT/"complete_validation_results.json", "w", encoding="utf-8") as f:
    json.dump(all_results, f, indent=2, ensure_ascii=False, default=str)

# Save per-DMR table
per_dmr = pd.DataFrame({
    "cluster_name": clusters,
    "module_id": [mod_map.get(c,"?") for c in clusters],
    "basin_residual_rank": [rank_map.get(c,np.nan) for c in clusters],
    "acc_2cell": acc_2cell, "acc_4cell": acc_4cell,
    "acc_8cell": acc_8cell, "acc_morula": acc_morula,
    "meth_2cell": x_2cell, "meth_4cell": x_4cell,
    "meth_8cell": x_8cell, "meth_morula": x_morula,
    "c_diag_48": c_diag_48, "c_diag_8m": c_diag_8m,
    "strict_correction": c_strict,
    "traj_corr": traj_corrs,
})
per_dmr.to_csv(OUT/"per_dmr_validation_table.tsv", sep="\t", index=False)

print("\n" + "="*60)
print("FINAL SUMMARY")
print("="*60)
print()
print("Test 1 (stage-specific acc-meth coupling):")
for s, r in stage_coupling.items():
    print(f"  {s}: rho={r['rho']:.3f}, perm_p={r['perm_p']:.3f}, sig={r['sig']}")
print()
print(f"Test 2 (ZGA-Reset coupling): rho={all_results['test2_zga_reset']['rho']:.3f}, sig={all_results['test2_zga_reset']['sig']}")
print(f"Test 3 (acc~correction): rho={all_results['test3_acc_correction']['rho']:.3f}, sig={all_results['test3_acc_correction']['sig']}")
print(f"Test 4 (CV): {all_results['test4_cross_validation']['both_positive']}/1000 both positive")
print(f"Test 5 (top25): sig={all_results['test5_top25_enrichment']['sig']}")
print(f"Test 7 (trajectory): mean_rho={all_results['test7_trajectory_correlation']['mean']:.3f}, p={all_results['test7_trajectory_correlation']['ttest_p']:.4f}")
print()
print(f"All results saved to {OUT}/")
print(f"Files: {len(list(OUT.iterdir()))}")
```
