## Supplemental code for "An entropy-based diagnostic framework for characterizing methylation state dynamics during preimplantation development": 5_30_progress__make_counterfactual_and_crossspecies.py

```
import json
from pathlib import Path

import numpy as np
import pandas as pd
import matplotlib
matplotlib.use("Agg")
import matplotlib.pyplot as plt
from scipy.stats import spearmanr


OUT = Path("E:/5_31_progress")
COMSOL = Path("E:/progress_comsol_analysis")
CSB = Path("C:/Users/18068/Desktop/CSB_TRO_Project_2026-05-24_TEAM_SHARE")

OUT.mkdir(parents=True, exist_ok=True)


def read_json(path):
    with open(path, "r", encoding="utf-8") as f:
        return json.load(f)


def norm01(x):
    x = np.asarray(x, dtype=float)
    lo, hi = np.nanmin(x), np.nanmax(x)
    if not np.isfinite(lo) or not np.isfinite(hi) or hi <= lo:
        return np.zeros_like(x, dtype=float)
    return (x - lo) / (hi - lo)


def methyl_entropy(beta):
    beta = np.clip(np.asarray(beta, dtype=float), 1e-9, 1 - 1e-9)
    return -beta * np.log2(beta) - (1 - beta) * np.log2(1 - beta)


def make_counterfactual_package():
    scenario = read_json(COMSOL / "scenario_results_final.json")
    targets = pd.read_csv(COMSOL / "validation_targets.csv")
    params = pd.read_csv(COMSOL / "parameters.csv").iloc[0].to_dict()

    rows = []
    scenario_labels = {
        "baseline_only": "c=0 methylation-only",
        "plus_zga": "ZGA only",
        "plus_entry": "entry/access only",
        "full_control": "full correction",
        "wrong_exit": "wrong exit sign",
    }
    for name, res in scenario.items():
        rows.append({
            "scenario": name,
            "label": scenario_labels.get(name, name),
            "dist_morula_t5": res["dist_morula_t5"],
            "dist_blast_t6": res["dist_blast_t6"],
            "in_morula": bool(res["in_morula"]),
            "in_blast": bool(res["in_blast"]),
            "z1_t5": res["z1_t5"],
            "z2_t5": res["z2_t5"],
            "z1_t6": res["z1_t6"],
            "z2_t6": res["z2_t6"],
            "n_pts": res["n_pts"],
        })
    df = pd.DataFrame(rows)
    df["morula_radius"] = float(params["r_morula"])
    df["blast_radius"] = float(params["r_blast"])
    df["morula_margin"] = df["dist_morula_t5"] - df["morula_radius"]
    df["blast_margin"] = df["dist_blast_t6"] - df["blast_radius"]
    df["supports_morula_entry"] = df["morula_margin"] < 0
    df["supports_terminal_completion"] = (df["morula_margin"] < 0) & (df["blast_margin"] < 0)
    df.to_csv(OUT / "counterfactual_scenario_summary.tsv", sep="\t", index=False)

    baseline = df.loc[df["scenario"].eq("baseline_only")].iloc[0]
    full = df.loc[df["scenario"].eq("full_control")].iloc[0]
    plus_entry = df.loc[df["scenario"].eq("plus_entry")].iloc[0]
    wrong = df.loc[df["scenario"].eq("wrong_exit")].iloc[0]

    summary = {
        "analysis": "computational_counterfactual_necessity",
        "date": "2026-05-31",
        "source": str(COMSOL),
        "primary_counterfactual": {
            "removed_term": "c=0 / methylation-only baseline",
            "baseline_dist_morula_t5": float(baseline["dist_morula_t5"]),
            "baseline_in_morula": bool(baseline["in_morula"]),
            "full_control_dist_morula_t5": float(full["dist_morula_t5"]),
            "full_control_in_morula": bool(full["in_morula"]),
            "full_control_in_blast": bool(full["in_blast"]),
            "distance_rescue_factor": float(baseline["dist_morula_t5"] / full["dist_morula_t5"]),
        },
        "branch_logic": {
            "entry_only_enters_morula": bool(plus_entry["in_morula"]),
            "entry_only_reaches_blast": bool(plus_entry["in_blast"]),
            "wrong_exit_enters_morula": bool(wrong["in_morula"]),
            "wrong_exit_reaches_blast": bool(wrong["in_blast"]),
        },
        "external_validation_targets": targets.iloc[0].to_dict(),
        "causal_language_boundary": (
            "This is model-implied counterfactual necessity evidence. It supports that "
            "the inferred correction/control term is necessary inside the operator-time "
            "model for reset-basin entry, but it does not identify the final in vivo "
            "causal molecular u_bio."
        ),
    }
    with open(OUT / "counterfactual_necessity_summary.json", "w", encoding="utf-8") as f:
        json.dump(summary, f, indent=2)

    # Figure: margins and trajectories.
    fig = plt.figure(figsize=(12, 5.2))
    gs = fig.add_gridspec(1, 2, width_ratios=[1.0, 1.25], wspace=0.28)

    ax = fig.add_subplot(gs[0, 0])
    order = ["baseline_only", "plus_zga", "plus_entry", "full_control", "wrong_exit"]
    plot_df = df.set_index("scenario").loc[order].reset_index()
    colors = ["#8d99ae", "#457b9d", "#e9c46a", "#c0392b", "#6c757d"]
    x = np.arange(len(plot_df))
    ax.bar(x - 0.18, plot_df["dist_morula_t5"], 0.36, color=colors, label="dist to morula at tau=5")
    ax.bar(x + 0.18, plot_df["dist_blast_t6"], 0.36, color="#b8c0c8", label="dist to blast at tau=6")
    ax.axhline(float(params["r_morula"]), color="#c0392b", lw=1.2, ls="--", label="morula radius")
    ax.axhline(float(params["r_blast"]), color="#2a9d8f", lw=1.0, ls=":", label="blast radius")
    ax.set_xticks(x)
    ax.set_xticklabels(plot_df["label"], rotation=35, ha="right")
    ax.set_ylabel("basin distance")
    ax.set_title("Counterfactual Basin Entry")
    ax.legend(fontsize=7, frameon=False)
    ax.grid(axis="y", alpha=0.2)

    ax = fig.add_subplot(gs[0, 1])
    center = (float(params["morula_z1"]), float(params["morula_z2"]))
    blast = (float(params["blast_z1"]), float(params["blast_z2"]))
    for name, color in zip(order, colors):
        tpath = COMSOL / f"traj_final_{name}.csv"
        tr = pd.read_csv(tpath)
        ax.plot(tr["z1"], tr["z2"], lw=1.7, color=color, label=scenario_labels[name])
        ax.scatter(tr["z1"].iloc[-1], tr["z2"].iloc[-1], s=22, color=color, edgecolor="white", zorder=4)
    circ = plt.Circle(center, float(params["r_morula"]), fill=False, color="#c0392b", lw=1.4, ls="--")
    ax.add_patch(circ)
    circ2 = plt.Circle(blast, float(params["r_blast"]), fill=False, color="#2a9d8f", lw=1.2, ls=":")
    ax.add_patch(circ2)
    ax.scatter([center[0]], [center[1]], marker="*", s=90, color="#c0392b", label="morula center", zorder=5)
    ax.scatter([blast[0]], [blast[1]], marker="*", s=80, color="#2a9d8f", label="blast center", zorder=5)
    ax.set_xlabel("operator coordinate z1")
    ax.set_ylabel("operator coordinate z2")
    ax.set_title("Model-Implied Intervention Trajectories")
    ax.legend(fontsize=7, frameon=False, loc="best")
    ax.grid(alpha=0.18)

    fig.suptitle("CSB-TRO / CEEF Computational Counterfactual Necessity", fontsize=13, fontweight="bold")
    fig.savefig(OUT / "counterfactual_necessity_figure.png", dpi=300, bbox_inches="tight")
    fig.savefig(OUT / "counterfactual_necessity_figure.pdf", bbox_inches="tight")
    plt.close(fig)


def make_crossspecies_package():
    mouse_path = OUT / "mm9_me.txt"
    human_path = CSB / "input_tables" / "TRO_interpretability_DMR_contribution_ranking.tsv"
    stage_path = CSB / "results" / "CSB_TRO_stage_state_summary.tsv"

    mouse = pd.read_csv(mouse_path, sep="\t")
    human = pd.read_csv(human_path, sep="\t")
    human_stage = pd.read_csv(stage_path, sep="\t")

    mouse_stage_cols = ["Oocyte", "Zygote", "2-cell", "4-cell", "8-cell", "Morula", "Epiblast"]
    human_stage_map = {
        "Oocyte": "MII oocyte",
        "Zygote": "zygote/PN",
        "2-cell": "2-cell",
        "4-cell": "4-cell",
        "8-cell": "8-cell",
        "Morula": "morula",
        "Epiblast": "blastocyst",
    }

    # Gene-overlap validation: no synthetic fallback, no coordinate pseudo-liftover.
    human_gene = human.copy()
    human_gene["gene_key"] = human_gene["nearest_gene"].astype(str).str.upper()
    human_gene = human_gene[~human_gene["gene_key"].isin(["", "NAN", "NA", "NONE"])]
    gene_weights = (
        human_gene.groupby("gene_key", as_index=False)
        .agg(age_weight_5yr=("age_weight_5yr", "sum"),
             n_human_dmr=("cluster_name", "count"),
             contribution_8cell_to_morula=("abs_contribution_8cell_to_morula", "sum"))
    )
    gene_weights["weight_norm"] = gene_weights["age_weight_5yr"] / gene_weights["age_weight_5yr"].sum()

    mouse["gene_key"] = mouse["Gene_Symbol"].astype(str).str.upper()
    matched = mouse.merge(gene_weights, on="gene_key", how="inner")
    matched = matched.dropna(subset=mouse_stage_cols, how="all")

    def stage_scores_for(sub, weight_col):
        sub = sub.copy()
        sub["_w"] = sub[weight_col].astype(float)
        if not np.isfinite(sub["_w"]).all() or sub["_w"].sum() <= 0:
            sub["_w"] = 1.0
        sub["_w"] = sub["_w"] / sub["_w"].sum()
        score_rows_inner = []
        for s in mouse_stage_cols:
            vals = pd.to_numeric(sub[s], errors="coerce")
            ok = vals.notna()
            w = sub.loc[ok, "_w"].values
            w = w / w.sum()
            beta = vals.loc[ok].values
            score_rows_inner.append({
                "mouse_stage": s,
                "human_stage_equivalent": human_stage_map[s],
                "weighted_mouse_methylation": float(np.dot(w, beta)),
                "unweighted_mouse_methylation_matched": float(np.nanmean(beta)),
                "weighted_mouse_entropy": float(np.dot(w, methyl_entropy(beta))),
                "n_matched_genes_with_stage": int(ok.sum()),
            })
        out = pd.DataFrame(score_rows_inner)
        out["A_mouse_norm_low_is_reset"] = norm01(out["weighted_mouse_methylation"])
        out["H_mouse_norm"] = norm01(out["weighted_mouse_entropy"])
        out["mouse_A_rank_lowest_is_1"] = out["weighted_mouse_methylation"].rank(method="min", ascending=True).astype(int)
        return out

    # Primary score: weighted methylation over human age-DMR-overlap genes.
    matched["weight_norm_matched"] = matched["age_weight_5yr"] / matched["age_weight_5yr"].sum()
    score = stage_scores_for(matched, "age_weight_5yr")
    human_A = human_stage[["stage", "A_mean"]].rename(columns={"stage": "human_stage_equivalent"})
    human_A["human_A_norm"] = norm01(human_A["A_mean"])
    score = score.merge(human_A, on="human_stage_equivalent", how="left")
    score.to_csv(OUT / "crossspecies_mouse_gleaner_stage_scores.tsv", sep="\t", index=False)
    matched.to_csv(OUT / "crossspecies_mouse_gleaner_matched_genes.tsv", sep="\t", index=False)

    # Null: random mouse gene sets with the same matched-gene count and identical fixed human weights.
    rng = np.random.default_rng(20260531)
    n_match = len(matched)
    valid_mouse = mouse.dropna(subset=mouse_stage_cols, how="any").copy()
    null_rows = []
    weights = matched["weight_norm_matched"].values
    weights = weights / weights.sum()
    for i in range(2000):
        samp = valid_mouse.sample(n=n_match, replace=False, random_state=int(rng.integers(0, 2**31 - 1)))
        vals_by_stage = {}
        for s in mouse_stage_cols:
            beta = pd.to_numeric(samp[s], errors="coerce").fillna(pd.to_numeric(samp[s], errors="coerce").median()).values
            vals_by_stage[s] = float(np.dot(weights, beta))
        min_stage = min(vals_by_stage, key=vals_by_stage.get)
        null_rows.append({
            "iter": i,
            "min_stage": min_stage,
            "morula_rank_lowest_is_1": int(pd.Series(vals_by_stage).rank(method="min", ascending=True)["Morula"]),
            "morula_minus_min": vals_by_stage["Morula"] - min(vals_by_stage.values()),
            **{f"A_{k}": v for k, v in vals_by_stage.items()},
        })
    null = pd.DataFrame(null_rows)
    null.to_csv(OUT / "crossspecies_mouse_gleaner_random_gene_null.tsv", sep="\t", index=False)

    # Sensitivity: all matched, top driver subsets, and alternative weights.
    sensitivity_rows = []
    sensitivity_specs = [
        ("all_age_weight", matched, "age_weight_5yr"),
        ("all_entry_contribution_weight", matched, "contribution_8cell_to_morula"),
        ("all_equal_weight", matched.assign(equal_weight=1.0), "equal_weight"),
        ("top25_age_weight", matched.sort_values("contribution_8cell_to_morula", ascending=False).head(25), "age_weight_5yr"),
        ("top50_age_weight", matched.sort_values("contribution_8cell_to_morula", ascending=False).head(50), "age_weight_5yr"),
        ("top25_entry_contribution_weight", matched.sort_values("contribution_8cell_to_morula", ascending=False).head(25), "contribution_8cell_to_morula"),
        ("top50_entry_contribution_weight", matched.sort_values("contribution_8cell_to_morula", ascending=False).head(50), "contribution_8cell_to_morula"),
    ]
    for label, sub, wcol in sensitivity_specs:
        sc = stage_scores_for(sub, wcol)
        mrow = sc.loc[sc["mouse_stage"].eq("Morula")].iloc[0]
        sensitivity_rows.append({
            "analysis_variant": label,
            "n_genes": int(len(sub)),
            "weight_col": wcol,
            "morula_rank_lowest_is_1": int(mrow["mouse_A_rank_lowest_is_1"]),
            "morula_weighted_methylation": float(mrow["weighted_mouse_methylation"]),
            "lowest_stage": sc.sort_values("weighted_mouse_methylation").iloc[0]["mouse_stage"],
            "second_lowest_stage": sc.sort_values("weighted_mouse_methylation").iloc[1]["mouse_stage"],
            "morula_minus_second_lowest": float(
                mrow["weighted_mouse_methylation"] -
                sc.sort_values("weighted_mouse_methylation").iloc[1]["weighted_mouse_methylation"]
            ),
        })
    sensitivity = pd.DataFrame(sensitivity_rows)
    sensitivity.to_csv(OUT / "crossspecies_mouse_gleaner_sensitivity.tsv", sep="\t", index=False)

    # Bootstrap over matched genes for internal stability.
    boot_rows = []
    for i in range(2000):
        sub = matched.sample(n=len(matched), replace=True, random_state=int(rng.integers(0, 2**31 - 1)))
        sc = stage_scores_for(sub, "age_weight_5yr")
        mrow = sc.loc[sc["mouse_stage"].eq("Morula")].iloc[0]
        boot_rows.append({
            "iter": i,
            "morula_rank_lowest_is_1": int(mrow["mouse_A_rank_lowest_is_1"]),
            "lowest_stage": sc.sort_values("weighted_mouse_methylation").iloc[0]["mouse_stage"],
            "morula_weighted_methylation": float(mrow["weighted_mouse_methylation"]),
        })
    boot = pd.DataFrame(boot_rows)
    boot.to_csv(OUT / "crossspecies_mouse_gleaner_matched_gene_bootstrap.tsv", sep="\t", index=False)

    morula = score.loc[score["mouse_stage"].eq("Morula")].iloc[0]
    stages_for_corr = score.dropna(subset=["human_A_norm", "A_mouse_norm_low_is_reset"])
    rho, pval = spearmanr(stages_for_corr["human_A_norm"], stages_for_corr["A_mouse_norm_low_is_reset"])
    null_p_rank1 = float((null["morula_rank_lowest_is_1"] <= int(morula["mouse_A_rank_lowest_is_1"])).mean())
    null_frac_morula_min = float((null["min_stage"] == "Morula").mean())

    summary = {
        "analysis": "cross_species_mouse_gleaner_gene_overlap_validation",
        "date": "2026-05-31",
        "mouse_data": "GLEANER mm9 gene-level methylation matrix, downloaded from https://compbio-zhanglab.org/GLEANER/download/mm9_me.txt",
        "human_anchor": "156 human age-DMR clusters; matched to mouse by nearest_gene / Gene_Symbol uppercase overlap",
        "n_human_dmr_clusters": int(len(human)),
        "n_human_genes_with_weights": int(len(gene_weights)),
        "n_mouse_genes_total": int(len(mouse)),
        "n_matched_mouse_genes": int(len(matched)),
        "mouse_stage_scores": score.to_dict(orient="records"),
        "morula_A_rank_lowest_is_1": int(morula["mouse_A_rank_lowest_is_1"]),
        "morula_weighted_methylation": float(morula["weighted_mouse_methylation"]),
        "human_mouse_stage_profile_spearman_rho": float(rho),
        "human_mouse_stage_profile_spearman_p": float(pval),
        "random_gene_null_n": int(len(null)),
        "random_gene_null_fraction_morula_min": null_frac_morula_min,
        "random_gene_null_p_morula_rank_as_good_or_better": null_p_rank1,
        "matched_gene_bootstrap_n": int(len(boot)),
        "matched_gene_bootstrap_fraction_morula_min": float((boot["lowest_stage"] == "Morula").mean()),
        "sensitivity": sensitivity.to_dict(orient="records"),
        "interpretation": (
            "This is an external cross-species diagnostic check based on public mouse "
            "GLEANER methylation. It tests whether genes overlapping the human age-DMR "
            "anchor show a mouse morula low-methylation/reset-like profile. It is not "
            "single-cell paired evidence and does not identify the final causal u_bio."
        ),
    }
    with open(OUT / "crossspecies_mouse_gleaner_summary.json", "w", encoding="utf-8") as f:
        json.dump(summary, f, indent=2)

    # Figure.
    fig, axes = plt.subplots(1, 3, figsize=(15.5, 4.8))
    colors = ["#6c757d", "#457b9d", "#1d3557", "#e9c46a", "#f4a261", "#c0392b", "#2a9d8f"]

    ax = axes[0]
    x = np.arange(len(score))
    ax.bar(x, score["weighted_mouse_methylation"], color=colors, edgecolor="white", lw=0.8)
    ax.set_xticks(x)
    ax.set_xticklabels(score["mouse_stage"], rotation=35, ha="right")
    ax.set_ylabel("weighted methylation over matched genes")
    ax.set_title("Mouse GLEANER Gene-Overlap Score")
    mi = int(score.index[score["mouse_stage"].eq("Morula")][0])
    ax.text(mi, score.loc[mi, "weighted_mouse_methylation"] + 0.015,
            f"rank {int(morula['mouse_A_rank_lowest_is_1'])}", ha="center",
            fontsize=8, color="#c0392b", fontweight="bold")
    ax.grid(axis="y", alpha=0.2)

    ax = axes[1]
    ax.bar(x - 0.18, score["human_A_norm"], 0.36, color="#2980b9", label="human A")
    ax.bar(x + 0.18, score["A_mouse_norm_low_is_reset"], 0.36, color="#c0392b", label="mouse matched-gene A")
    ax.set_xticks(x)
    ax.set_xticklabels(score["mouse_stage"], rotation=35, ha="right")
    ax.set_ylabel("normalized score")
    ax.set_title(f"Human-Mouse Stage Profile\nSpearman rho={rho:.3f}, p={pval:.3f}")
    ax.legend(frameon=False, fontsize=8)
    ax.grid(axis="y", alpha=0.2)

    ax = axes[2]
    rank_counts = null["morula_rank_lowest_is_1"].value_counts().sort_index()
    ax.bar(rank_counts.index.astype(str), rank_counts.values / len(null), color="#adb5bd")
    ax.axvline(int(morula["mouse_A_rank_lowest_is_1"]) - 1, color="#c0392b", lw=1.4)
    ax.set_xlabel("Morula rank in random matched-size gene sets")
    ax.set_ylabel("fraction")
    ax.set_title("Random-Gene Null")
    ax.text(0.03, 0.95,
            f"n={len(null)}\nfrac null Morula min={null_frac_morula_min:.3f}",
            transform=ax.transAxes, va="top", fontsize=8,
            bbox=dict(facecolor="white", edgecolor="#cccccc", alpha=0.9))
    ax.grid(axis="y", alpha=0.2)

    fig.suptitle("Cross-Species External Validation: Mouse GLEANER Methylation",
                 fontsize=13, fontweight="bold", y=0.99)
    fig.tight_layout(rect=[0, 0, 1, 0.90])
    fig.savefig(OUT / "crossspecies_mouse_gleaner_figure.png", dpi=300, bbox_inches="tight")
    fig.savefig(OUT / "crossspecies_mouse_gleaner_figure.pdf", bbox_inches="tight")
    plt.close(fig)


def write_readme():
    text = """# CSB-TRO Causality Upgrade Package

Generated: 2026-05-31

This folder contains two causality-strengthening analyses:

1. `counterfactual_necessity_*`: model-implied COMSOL/CEEF counterfactuals. The cleanest result is that the c=0 methylation-only baseline fails morula basin entry, while the full correction enters morula and reaches blastocyst.
2. `crossspecies_mouse_gleaner_*`: public mouse GLEANER gene-level methylation validation using human age-DMR nearest-gene overlap. This is an external diagnostic reproducibility test, not causal molecular identification.

Use boundary language: these analyses strengthen computational causal credibility and external validity, but they do not identify the final in vivo causal `u_bio`.
"""
    (OUT / "README_5_31_causality_upgrade.md").write_text(text, encoding="utf-8")


if __name__ == "__main__":
    make_counterfactual_package()
    make_crossspecies_package()
    write_readme()
    print(f"Done. Outputs written to {OUT}")
```
