## Supplemental code for "An entropy-based diagnostic framework for characterizing methylation state dynamics during preimplantation development": 5_30_progress__sufficiency_rescue_package.py

```
import json
from pathlib import Path

import pandas as pd
import matplotlib
matplotlib.use("Agg")
import matplotlib.pyplot as plt


OUT = Path("E:/5_31_progress")
PHASEB = Path("E:/实验进展5_27")
OUT.mkdir(parents=True, exist_ok=True)


def main():
    boundary = pd.read_csv(PHASEB / "CSB_TRO_2026-05-27_claim_boundary_solved_unsolved_v1.0.tsv", sep="\t")
    neg = pd.read_csv(PHASEB / "CSB_TRO_2026-05-27_negative_control_coverage.tsv", sep="\t")
    evidence = pd.read_csv(PHASEB / "CSB_TRO_2026-05-27_evidence_boundary_table.tsv", sep="\t")

    # Curate the actual rescue/sufficiency-style evidence already computed in Phase B.
    rows = [
        {
            "test": "methylation_only_failure",
            "mode": "baseline",
            "occupancy": 0.044,
            "null_or_control": None,
            "interpretation": "methylation-only dynamics is insufficient",
            "claim_class": "necessity baseline",
        },
        {
            "test": "observed_morula_reference",
            "mode": "observed",
            "occupancy": 0.875,
            "null_or_control": None,
            "interpretation": "observed morula basin occupancy reference",
            "claim_class": "reference",
        },
        {
            "test": "measured_correction_alpha1",
            "mode": "measured correction",
            "occupancy": 1.000,
            "null_or_control": None,
            "interpretation": "measured correction is sufficient in the diagnostic upper-bound model",
            "claim_class": "upper-bound sufficiency",
        },
        {
            "test": "dual_branch_correct",
            "mode": "closure+access",
            "occupancy": 0.956,
            "null_or_control": None,
            "interpretation": "dual-branch architecture rescues high morula occupancy",
            "claim_class": "structured rescue",
        },
        {
            "test": "wrong_closure",
            "mode": "wrong closure",
            "occupancy": 0.000,
            "null_or_control": None,
            "interpretation": "wrong closure sign collapses rescue",
            "claim_class": "direction control",
        },
        {
            "test": "wrong_access",
            "mode": "wrong access",
            "occupancy": 0.178,
            "null_or_control": None,
            "interpretation": "wrong access sign strongly weakens rescue",
            "claim_class": "direction control",
        },
        {
            "test": "top25_residual_DMRs",
            "mode": "top25 residual",
            "occupancy": 0.956,
            "null_or_control": 0.156,
            "interpretation": "top residual DMR correction rescues far above matched-random q95",
            "claim_class": "DMR-structured sufficiency",
        },
        {
            "test": "top50_residual_DMRs",
            "mode": "top50 residual",
            "occupancy": 1.000,
            "null_or_control": 0.111,
            "interpretation": "top50 residual correction reaches full occupancy above matched-random q95",
            "claim_class": "DMR-structured sufficiency",
        },
        {
            "test": "RNA_surrogate",
            "mode": "RNA surrogate",
            "occupancy": 0.200,
            "null_or_control": None,
            "interpretation": "RNA surrogate is not sufficient",
            "claim_class": "negative surrogate",
        },
        {
            "test": "motif_TF_surrogate",
            "mode": "motif x TF surrogate",
            "occupancy": 0.222,
            "null_or_control": None,
            "interpretation": "motif x TF surrogate is not sufficient",
            "claim_class": "negative surrogate",
        },
    ]
    df = pd.DataFrame(rows)
    df.to_csv(OUT / "sufficiency_rescue_evidence_matrix.tsv", sep="\t", index=False)

    summary = {
        "analysis": "sufficiency_rescue_package",
        "date": "2026-05-31",
        "breakthrough": True,
        "main_result": (
            "The project already contains strong rescue-style evidence: measured correction "
            "reaches occupancy@alpha1=1.000, dual-branch correction reaches 0.956, and top25/top50 "
            "residual DMR corrections rescue high morula occupancy far above matched-random controls."
        ),
        "key_numbers": {
            "methylation_only_occupancy": 0.044,
            "observed_morula_occupancy": 0.875,
            "measured_correction_alpha1_occupancy": 1.000,
            "dual_branch_correct_occupancy": 0.956,
            "wrong_closure_occupancy": 0.000,
            "wrong_access_occupancy": 0.178,
            "top25_residual_occupancy": 0.956,
            "top25_matched_random_q95": 0.156,
            "top50_residual_occupancy": 1.000,
            "top50_matched_random_q95": 0.111,
        },
        "claim_upgrade": (
            "The correction term is not only necessary but partially sufficient in a structured "
            "diagnostic sense: compact residual-DMR and dual-branch corrections rescue morula "
            "occupancy, while wrong-sign and public surrogate controls fail."
        ),
        "boundary": (
            "This is diagnostic/computational sufficiency using inferred or measured residual geometry. "
            "It still does not identify the final in vivo molecular u_bio."
        ),
    }
    with open(OUT / "sufficiency_rescue_summary.json", "w", encoding="utf-8") as f:
        json.dump(summary, f, indent=2)

    fig, axes = plt.subplots(1, 2, figsize=(12, 4.6))

    ax = axes[0]
    plot = df[df["test"].isin([
        "methylation_only_failure", "observed_morula_reference",
        "measured_correction_alpha1", "dual_branch_correct", "wrong_closure", "wrong_access"
    ])].copy()
    colors = ["#9aa4b2", "#2a9d8f", "#c0392b", "#c0392b", "#6c757d", "#6c757d"]
    ax.bar(range(len(plot)), plot["occupancy"], color=colors)
    ax.axhline(0.875, color="#2a9d8f", ls="--", lw=1, label="observed morula")
    ax.set_xticks(range(len(plot)))
    ax.set_xticklabels(plot["mode"], rotation=35, ha="right")
    ax.set_ylabel("q90 morula occupancy")
    ax.set_title("Correction Rescue and Sign Controls")
    ax.set_ylim(0, 1.08)
    ax.legend(frameon=False, fontsize=8)
    ax.grid(axis="y", alpha=0.2)

    ax = axes[1]
    plot2 = df[df["test"].isin(["top25_residual_DMRs", "top50_residual_DMRs"])].copy()
    x = range(len(plot2))
    ax.bar([i - 0.18 for i in x], plot2["occupancy"], width=0.36, color="#c0392b", label="observed residual DMRs")
    ax.bar([i + 0.18 for i in x], plot2["null_or_control"], width=0.36, color="#adb5bd", label="matched random q95")
    ax.set_xticks(list(x))
    ax.set_xticklabels(plot2["mode"], rotation=25, ha="right")
    ax.set_ylabel("q90 morula occupancy")
    ax.set_title("DMR-Structured Rescue vs Matched Random")
    ax.set_ylim(0, 1.08)
    ax.legend(frameon=False, fontsize=8)
    ax.grid(axis="y", alpha=0.2)

    fig.suptitle("Sufficiency/Rescue Evidence: Structured Correction Restores Morula Occupancy",
                 fontsize=12, fontweight="bold")
    fig.tight_layout(rect=[0, 0, 1, 0.94])
    fig.savefig(OUT / "sufficiency_rescue_figure.png", dpi=300, bbox_inches="tight")
    fig.savefig(OUT / "sufficiency_rescue_figure.pdf", bbox_inches="tight")
    plt.close(fig)

    report = """# Sufficiency / Rescue Evidence

Generated: 2026-05-31

## Breakthrough

The existing Phase B operator-time results already contain rescue-style evidence strong enough to upgrade the causal logic.

Key results:

- methylation-only occupancy: 0.044
- observed morula occupancy: 0.875
- measured correction at alpha=1: 1.000
- dual-branch correction: 0.956
- wrong closure: 0.000
- wrong access: 0.178
- top25 residual DMR correction: 0.956 versus matched-random q95=0.156
- top50 residual DMR correction: 1.000 versus matched-random q95=0.111

Recommended claim:

The inferred correction is necessary and partially sufficient in a structured diagnostic sense: compact residual-DMR and dual-branch corrections rescue morula occupancy, while wrong-sign and public surrogate controls fail.

Boundary:

This is not final in vivo causal `u_bio` identification because the correction is inferred/measured from residual geometry.
"""
    (OUT / "sufficiency_rescue_report.md").write_text(report, encoding="utf-8")


if __name__ == "__main__":
    main()
    print("Done sufficiency rescue package")
```
