## Supplemental code for "An entropy-based diagnostic framework for characterizing methylation state dynamics during preimplantation development": duality_accessibility_coupling_analysis.py

```
#!/usr/bin/env python
"""Couple morula entry-exit geometry with public morula accessibility.

This script uses only already-generated result tables. It does not rerun the
public chromatin extraction or the entry-exit duality experiment.
"""

from __future__ import annotations

import argparse
from pathlib import Path

import numpy as np
import pandas as pd

try:
    import matplotlib.pyplot as plt
except ModuleNotFoundError:  # Plotting is optional; TSV/MD outputs are primary.
    plt = None


ACCESS_MODULES = {"M02", "M10"}
CLOSURE_MODULES = {"M01", "M05", "M12"}
PRIORITY_MODULES = ACCESS_MODULES | CLOSURE_MODULES
EPS = 1e-12


def read_tsv(path: Path) -> pd.DataFrame:
    if not path.exists():
        raise FileNotFoundError(path)
    return pd.read_csv(path, sep="\t")


def q95(values: list[float]) -> float:
    return float(np.quantile(np.asarray(values, dtype=float), 0.95))


def safe_mean(series: pd.Series) -> float:
    values = pd.to_numeric(series, errors="coerce")
    return float(values.mean()) if values.notna().any() else np.nan


def pearson_corr(a: pd.Series, b: pd.Series) -> float:
    x = pd.to_numeric(a, errors="coerce")
    y = pd.to_numeric(b, errors="coerce")
    mask = x.notna() & y.notna()
    x = x[mask].to_numpy(dtype=float)
    y = y[mask].to_numpy(dtype=float)
    if len(x) < 3:
        return np.nan
    x = x - x.mean()
    y = y - y.mean()
    denom = float(np.linalg.norm(x) * np.linalg.norm(y))
    return float(np.dot(x, y) / denom) if denom > EPS else np.nan


def spearman_corr(a: pd.Series, b: pd.Series) -> float:
    return pearson_corr(pd.Series(a).rank(method="average"), pd.Series(b).rank(method="average"))


def branch_for_module(module_id: str) -> str:
    if module_id in ACCESS_MODULES:
        return "access"
    if module_id in CLOSURE_MODULES:
        return "closure"
    return "other"


def prepare_merged(entry: pd.DataFrame, access: pd.DataFrame) -> pd.DataFrame:
    keep_access = [
        "cluster_name",
        "overlap_public_chromatin",
        "public_accessibility_morula_max",
        "public_accessibility_morula_minus_8cell",
    ]
    merged = entry.merge(access[keep_access], on="cluster_name", how="left")
    for col in [
        "beta_8cell",
        "beta_morula",
        "beta_blastocyst",
        "entry_change",
        "exit_change",
        "curvature",
        "rebound_ratio",
        "signed_duality",
        "basin_residual_rank",
        "abs_latent_residual_delta_beta",
        "public_accessibility_morula_max",
        "public_accessibility_morula_minus_8cell",
    ]:
        if col in merged.columns:
            merged[col] = pd.to_numeric(merged[col], errors="coerce")

    merged["branch"] = merged["module_id"].map(branch_for_module)
    merged["overlap_public_chromatin"] = merged["overlap_public_chromatin"].astype(str).str.lower().eq("true")

    rebound = merged["rebound_ratio"].replace([np.inf, -np.inf], np.nan).astype(float)
    rebound_symmetry = np.minimum(rebound, 1.0 / rebound)
    rebound_symmetry = rebound_symmetry.replace([np.inf, -np.inf], np.nan).fillna(0.0).clip(lower=0.0, upper=1.0)
    merged["rebound_symmetry_score"] = rebound_symmetry

    # DMR-level cosine is degenerate in one dimension. This proxy preserves the
    # sign test and adds a symmetry penalty so anti-aligned, similarly sized
    # entry/exit changes rank above weak or highly asymmetric rebounds.
    merged["dmr_geometry_score"] = merged["signed_duality"].fillna(0.0) * merged["rebound_symmetry_score"]
    merged["is_access_supported"] = merged["public_accessibility_morula_max"].notna()
    return merged


def module_index(df: pd.DataFrame) -> dict[str, np.ndarray]:
    return {module_id: sub.index.to_numpy() for module_id, sub in df.groupby("module_id")}


def module_matched_indices(
    all_indices: np.ndarray,
    module_to_indices: dict[str, np.ndarray],
    template: pd.DataFrame,
    rng: np.random.Generator,
) -> np.ndarray:
    pieces = []
    for module_id, n in template["module_id"].value_counts().items():
        pool = module_to_indices.get(module_id, all_indices)
        replace = len(pool) < n
        pieces.append(rng.choice(pool, size=n, replace=replace))
    return np.concatenate(pieces)


def random_indices_for_group(
    df: pd.DataFrame,
    sub: pd.DataFrame,
    label: str,
    rng: np.random.Generator,
    all_indices: np.ndarray,
    module_to_indices: dict[str, np.ndarray],
) -> tuple[np.ndarray, str]:
    if label in {"access_branch", "closure_branch", "priority_modules"}:
        replace = len(df) < len(sub)
        return rng.choice(all_indices, size=len(sub), replace=replace), "size_matched_random_sets"
    return module_matched_indices(all_indices, module_to_indices, sub, rng), "module_matched_random_sets"


def summarize_group(
    df: pd.DataFrame,
    label: str,
    sub: pd.DataFrame,
    n_iter: int,
    rng: np.random.Generator,
    all_indices: np.ndarray,
    module_to_indices: dict[str, np.ndarray],
) -> dict:
    observed = safe_mean(sub["public_accessibility_morula_max"])
    observed_delta = safe_mean(sub["public_accessibility_morula_minus_8cell"])
    observed_overlap = float(sub["overlap_public_chromatin"].mean()) if len(sub) else np.nan
    random_means = []
    random_deltas = []
    random_overlaps = []
    random_mode = ""
    for _ in range(n_iter):
        draw_idx, random_mode = random_indices_for_group(df, sub, label, rng, all_indices, module_to_indices)
        draw = df.loc[draw_idx]
        random_means.append(safe_mean(draw["public_accessibility_morula_max"]))
        random_deltas.append(safe_mean(draw["public_accessibility_morula_minus_8cell"]))
        random_overlaps.append(float(draw["overlap_public_chromatin"].mean()))
    return {
        "group": label,
        "random_mode": random_mode,
        "n_dmr": int(len(sub)),
        "mean_geometry_score": safe_mean(sub["dmr_geometry_score"]),
        "mean_signed_duality": safe_mean(sub["signed_duality"]),
        "mean_rebound_symmetry": safe_mean(sub["rebound_symmetry_score"]),
        "mean_abs_residual": safe_mean(sub["abs_latent_residual_delta_beta"]),
        "observed_morula_accessibility_mean": observed,
        "random_median": float(np.nanmedian(random_means)),
        "random_q95": q95(random_means),
        "random_max": float(np.nanmax(random_means)),
        "observed_gt_random_q95": bool(observed > q95(random_means)) if np.isfinite(observed) else False,
        "observed_morula_minus_8cell_mean": observed_delta,
        "delta_random_median": float(np.nanmedian(random_deltas)),
        "delta_random_q95": q95(random_deltas),
        "delta_observed_gt_random_q95": bool(observed_delta > q95(random_deltas)) if np.isfinite(observed_delta) else False,
        "observed_overlap_fraction": observed_overlap,
        "overlap_random_median": float(np.nanmedian(random_overlaps)),
        "overlap_random_q95": q95(random_overlaps),
        "n_random": int(n_iter),
    }


def random_accessibility_distributions(df: pd.DataFrame, groups: dict[str, pd.DataFrame], n_iter: int, seed: int) -> pd.DataFrame:
    rng = np.random.default_rng(seed)
    all_indices = df.index.to_numpy()
    module_to_indices = module_index(df)
    rows = []
    for label, sub in groups.items():
        for i in range(n_iter):
            draw_idx, random_mode = random_indices_for_group(df, sub, label, rng, all_indices, module_to_indices)
            draw = df.loc[draw_idx]
            rows.append(
                {
                    "group": label,
                    "random_mode": random_mode,
                    "iteration": i,
                    "random_morula_accessibility_mean": safe_mean(draw["public_accessibility_morula_max"]),
                    "random_morula_minus_8cell_mean": safe_mean(draw["public_accessibility_morula_minus_8cell"]),
                    "random_overlap_fraction": float(draw["overlap_public_chromatin"].mean()),
                }
            )
    return pd.DataFrame(rows)


def top_groups(df: pd.DataFrame) -> dict[str, pd.DataFrame]:
    groups: dict[str, pd.DataFrame] = {}
    ranked_residual = df.sort_values("basin_residual_rank", ascending=True)
    ranked_geometry = df.sort_values("dmr_geometry_score", ascending=False)
    ranked_signed_duality = df.sort_values("signed_duality", ascending=False)
    ranked_positive_curvature = df.sort_values("curvature", ascending=False)
    ranked_negative_curvature = df.sort_values("curvature", ascending=True)
    for k in [25, 50, 100]:
        groups[f"top{k}_residual"] = ranked_residual.head(k)
        groups[f"top{k}_geometry"] = ranked_geometry.head(k)
        groups[f"top{k}_signed_duality"] = ranked_signed_duality.head(k)
        groups[f"top{k}_positive_curvature"] = ranked_positive_curvature.head(k)
        groups[f"top{k}_negative_curvature"] = ranked_negative_curvature.head(k)
        residual_names = set(ranked_residual.head(k)["cluster_name"])
        groups[f"top{k}_residual_and_top{k}_geometry"] = df[
            df["cluster_name"].isin(residual_names)
            & df["cluster_name"].isin(set(ranked_geometry.head(k)["cluster_name"]))
        ]
    groups["opposite_direction_DMRs"] = df[df["signed_duality"] > 0]
    groups["same_direction_DMRs"] = df[df["signed_duality"] < 0]
    groups["u_shape_DMRs"] = df[df["is_u_shape"].astype(str).str.lower().eq("true")]
    groups["inverted_u_DMRs"] = df[df["is_inverted_u"].astype(str).str.lower().eq("true")]
    groups["access_branch"] = df[df["branch"] == "access"]
    groups["closure_branch"] = df[df["branch"] == "closure"]
    groups["priority_modules"] = df[df["module_id"].isin(PRIORITY_MODULES)]
    return groups


def intersection_summary(df: pd.DataFrame, n_iter: int, seed: int) -> pd.DataFrame:
    rng = np.random.default_rng(seed)
    all_indices = df.index.to_numpy()
    module_to_indices = module_index(df)
    ranked_residual = df.sort_values("basin_residual_rank", ascending=True)
    ranked_geometry = df.sort_values("dmr_geometry_score", ascending=False)
    rows = []
    for k in [10, 25, 50, 100]:
        residual = ranked_residual.head(k)
        geometry_names = set(ranked_geometry.head(k)["cluster_name"])
        observed_overlap = int(residual["cluster_name"].isin(geometry_names).sum())
        random_overlaps = []
        for _ in range(n_iter):
            draw = df.loc[module_matched_indices(all_indices, module_to_indices, residual, rng)]
            random_overlaps.append(int(draw["cluster_name"].isin(geometry_names).sum()))
        rows.append(
            {
                "comparison": f"top{k}_residual_intersect_top{k}_geometry",
                "top_k": k,
                "observed_overlap": observed_overlap,
                "observed_fraction": observed_overlap / k,
                "random_median": float(np.median(random_overlaps)),
                "random_q95": q95(random_overlaps),
                "random_max": int(np.max(random_overlaps)),
                "observed_gt_random_q95": bool(observed_overlap > q95(random_overlaps)),
                "empirical_p_ge_observed": float((np.asarray(random_overlaps) >= observed_overlap).mean()),
                "n_random": int(n_iter),
            }
        )
    return pd.DataFrame(rows)


def module_triad(df: pd.DataFrame) -> pd.DataFrame:
    rows = []
    for module_id, sub in df.groupby("module_id"):
        rows.append(
            {
                "module_id": module_id,
                "branch": branch_for_module(module_id),
                "n_dmr": int(len(sub)),
                "mean_abs_residual": safe_mean(sub["abs_latent_residual_delta_beta"]),
                "best_residual_rank": int(sub["basin_residual_rank"].min()),
                "mean_geometry_score": safe_mean(sub["dmr_geometry_score"]),
                "mean_signed_duality": safe_mean(sub["signed_duality"]),
                "mean_curvature": safe_mean(sub["curvature"]),
                "fraction_u_shape": float(sub["is_u_shape"].astype(str).str.lower().eq("true").mean()),
                "mean_morula_accessibility": safe_mean(sub["public_accessibility_morula_max"]),
                "mean_morula_minus_8cell": safe_mean(sub["public_accessibility_morula_minus_8cell"]),
                "overlap_public_chromatin_fraction": float(sub["overlap_public_chromatin"].mean()),
                "interpretation": module_interpretation(module_id, sub),
            }
        )
    out = pd.DataFrame(rows)
    return out.sort_values(["branch", "mean_geometry_score", "mean_abs_residual"], ascending=[True, False, False])


def module_random_controls(df: pd.DataFrame, n_iter: int, seed: int) -> pd.DataFrame:
    rng = np.random.default_rng(seed)
    all_indices = df.index.to_numpy()
    rows = []
    for module_id, sub in df.groupby("module_id"):
        random_access = []
        random_geom = []
        random_resid = []
        for _ in range(n_iter):
            draw_idx = rng.choice(all_indices, size=len(sub), replace=len(df) < len(sub))
            draw = df.loc[draw_idx]
            random_access.append(safe_mean(draw["public_accessibility_morula_max"]))
            random_geom.append(safe_mean(draw["dmr_geometry_score"]))
            random_resid.append(safe_mean(draw["abs_latent_residual_delta_beta"]))
        access_obs = safe_mean(sub["public_accessibility_morula_max"])
        geom_obs = safe_mean(sub["dmr_geometry_score"])
        resid_obs = safe_mean(sub["abs_latent_residual_delta_beta"])
        rows.append(
            {
                "module_id": module_id,
                "branch": branch_for_module(module_id),
                "n_dmr": int(len(sub)),
                "observed_accessibility": access_obs,
                "accessibility_random_median": float(np.nanmedian(random_access)),
                "accessibility_random_q95": q95(random_access),
                "accessibility_gt_random_q95": bool(access_obs > q95(random_access)) if np.isfinite(access_obs) else False,
                "observed_geometry": geom_obs,
                "geometry_random_median": float(np.nanmedian(random_geom)),
                "geometry_random_q95": q95(random_geom),
                "geometry_gt_random_q95": bool(geom_obs > q95(random_geom)) if np.isfinite(geom_obs) else False,
                "observed_abs_residual": resid_obs,
                "residual_random_median": float(np.nanmedian(random_resid)),
                "residual_random_q95": q95(random_resid),
                "residual_gt_random_q95": bool(resid_obs > q95(random_resid)) if np.isfinite(resid_obs) else False,
                "n_random": int(n_iter),
            }
        )
    return pd.DataFrame(rows).sort_values(
        ["accessibility_gt_random_q95", "geometry_gt_random_q95", "residual_gt_random_q95", "observed_accessibility"],
        ascending=[False, False, False, False],
    )


def correlation_summary(df: pd.DataFrame, n_iter: int, seed: int) -> pd.DataFrame:
    rng = np.random.default_rng(seed)
    rows = []
    y = df["public_accessibility_morula_max"]
    for metric in [
        "dmr_geometry_score",
        "signed_duality",
        "rebound_symmetry_score",
        "abs_latent_residual_delta_beta",
        "curvature",
    ]:
        sub = df[[metric, "public_accessibility_morula_max"]].dropna()
        observed_spearman = spearman_corr(sub[metric], sub["public_accessibility_morula_max"])
        observed_pearson = pearson_corr(sub[metric], sub["public_accessibility_morula_max"])
        random_spearman = []
        values = y.to_numpy(dtype=float)
        for _ in range(n_iter):
            shuffled = rng.permutation(values)
            tmp = pd.DataFrame({metric: df[metric].to_numpy(dtype=float), "shuffled_access": shuffled}).dropna()
            random_spearman.append(spearman_corr(tmp[metric], tmp["shuffled_access"]))
        random_spearman_arr = np.asarray(random_spearman, dtype=float)
        rows.append(
            {
                "metric": metric,
                "n_dmr": int(len(sub)),
                "observed_pearson": observed_pearson,
                "observed_spearman": observed_spearman,
                "random_spearman_median": float(np.nanmedian(random_spearman_arr)),
                "random_spearman_q05": float(np.nanquantile(random_spearman_arr, 0.05)),
                "random_spearman_q95": float(np.nanquantile(random_spearman_arr, 0.95)),
                "observed_gt_random_q95": bool(observed_spearman > np.nanquantile(random_spearman_arr, 0.95)),
                "observed_lt_random_q05": bool(observed_spearman < np.nanquantile(random_spearman_arr, 0.05)),
                "empirical_p_ge_observed": float((random_spearman_arr >= observed_spearman).mean()),
                "empirical_p_le_observed": float((random_spearman_arr <= observed_spearman).mean()),
                "n_random": int(n_iter),
            }
        )
    return pd.DataFrame(rows)


def joint_priority_table(df: pd.DataFrame) -> pd.DataFrame:
    out = df.copy()
    for col in ["dmr_geometry_score", "abs_latent_residual_delta_beta", "public_accessibility_morula_max"]:
        values = out[col].astype(float)
        out[f"z_{col}"] = (values - values.mean()) / (values.std(ddof=0) + EPS)
    out["joint_reset_geometry_access_score"] = out[
        ["z_dmr_geometry_score", "z_abs_latent_residual_delta_beta", "z_public_accessibility_morula_max"]
    ].mean(axis=1)
    keep = [
        "cluster_name",
        "module_id",
        "branch",
        "basin_residual_rank",
        "abs_latent_residual_delta_beta",
        "dmr_geometry_score",
        "signed_duality",
        "rebound_symmetry_score",
        "curvature",
        "public_accessibility_morula_max",
        "public_accessibility_morula_minus_8cell",
        "overlap_public_chromatin",
        "joint_reset_geometry_access_score",
    ]
    return out.sort_values("joint_reset_geometry_access_score", ascending=False)[keep]


def module_interpretation(module_id: str, sub: pd.DataFrame) -> str:
    branch = branch_for_module(module_id)
    geom = safe_mean(sub["dmr_geometry_score"])
    access = safe_mean(sub["public_accessibility_morula_max"])
    residual_rank = int(sub["basin_residual_rank"].min())
    if branch == "access" and geom > 0.5:
        return "access-associated entry-exit geometry module"
    if branch == "closure" and residual_rank <= 25:
        return "closure/residual correction module with bounded duality"
    if np.isfinite(access) and access > 1.5 and residual_rank <= 25:
        return "high residual module with morula accessibility support"
    return "background or secondary reset-geometry module"


def plot_summary(summary: pd.DataFrame, intersections: pd.DataFrame, module_df: pd.DataFrame, out: Path) -> None:
    if plt is None:
        write_basic_svg(summary, intersections, module_df, out.with_suffix(".svg"))
        return
    fig, axes = plt.subplots(1, 3, figsize=(13, 4))

    sel = summary[summary["group"].isin(["top25_residual", "top25_geometry", "access_branch", "closure_branch"])]
    x = np.arange(len(sel))
    axes[0].bar(x, sel["observed_morula_accessibility_mean"], color="#3b6ea8", label="observed")
    axes[0].scatter(x, sel["random_q95"], color="#c43d3d", zorder=3, label="module-matched q95")
    axes[0].set_xticks(x)
    axes[0].set_xticklabels(sel["group"], rotation=35, ha="right")
    axes[0].set_ylabel("Mean morula accessibility")
    axes[0].set_title("Duality/accessibility coupling")
    axes[0].legend(frameon=False, fontsize=8)

    axes[1].bar(intersections["top_k"].astype(str), intersections["observed_overlap"], color="#5f8f3d")
    axes[1].scatter(intersections["top_k"].astype(str), intersections["random_q95"], color="#c43d3d", zorder=3)
    axes[1].set_xlabel("Top k")
    axes[1].set_ylabel("Residual ∩ geometry overlap")
    axes[1].set_title("Top residual x top geometry")

    mod = module_df[module_df["module_id"].isin(sorted(PRIORITY_MODULES))]
    axes[2].scatter(mod["mean_geometry_score"], mod["mean_morula_accessibility"], s=80, color="#6b4fa3")
    for _, row in mod.iterrows():
        axes[2].text(row["mean_geometry_score"], row["mean_morula_accessibility"], row["module_id"], fontsize=9)
    axes[2].set_xlabel("Mean DMR geometry score")
    axes[2].set_ylabel("Mean morula accessibility")
    axes[2].set_title("Priority module triad")

    fig.tight_layout()
    fig.savefig(out.with_suffix(".png"), dpi=220)
    fig.savefig(out.with_suffix(".svg"))
    plt.close(fig)


def write_basic_svg(summary: pd.DataFrame, intersections: pd.DataFrame, module_df: pd.DataFrame, out: Path) -> None:
    sel = summary[summary["group"].isin(["top25_residual", "top25_geometry", "access_branch", "closure_branch"])].copy()
    width, height = 980, 360
    margin = 56
    panel_w = 290
    max_y = float(np.nanmax([sel["observed_morula_accessibility_mean"].max(), sel["random_q95"].max(), 1.0]))

    def yscale(v: float) -> float:
        return margin + (height - 2 * margin) * (1.0 - float(v) / (max_y * 1.15))

    parts = [
        f'<svg xmlns="http://www.w3.org/2000/svg" width="{width}" height="{height}" viewBox="0 0 {width} {height}">',
        '<rect width="100%" height="100%" fill="white"/>',
        '<text x="24" y="28" font-family="Arial" font-size="17" font-weight="700">Duality-accessibility coupling summary</text>',
        '<text x="24" y="56" font-family="Arial" font-size="13">Observed mean morula accessibility vs random q95</text>',
    ]
    x0 = 34
    bar_w = 38
    gap = 28
    for i, (_, row) in enumerate(sel.iterrows()):
        x = x0 + i * (bar_w + gap)
        y = yscale(row["observed_morula_accessibility_mean"])
        h = height - margin - y
        parts.append(f'<rect x="{x}" y="{y:.1f}" width="{bar_w}" height="{h:.1f}" fill="#3b6ea8"/>')
        qy = yscale(row["random_q95"])
        parts.append(f'<line x1="{x-4}" x2="{x+bar_w+4}" y1="{qy:.1f}" y2="{qy:.1f}" stroke="#c43d3d" stroke-width="2"/>')
        parts.append(f'<text x="{x+bar_w/2}" y="{height-24}" font-family="Arial" font-size="10" text-anchor="middle" transform="rotate(35 {x+bar_w/2},{height-24})">{row["group"]}</text>')
    parts.append(f'<line x1="{x0-10}" x2="{x0+4*(bar_w+gap)}" y1="{height-margin}" y2="{height-margin}" stroke="#333"/>')

    ix0 = 370
    parts.append('<text x="360" y="56" font-family="Arial" font-size="13">Top residual x top geometry overlap</text>')
    max_o = float(max(intersections["observed_overlap"].max(), intersections["random_q95"].max(), 1.0))
    for i, (_, row) in enumerate(intersections.iterrows()):
        x = ix0 + i * 48
        h = (height - 2 * margin) * float(row["observed_overlap"]) / (max_o * 1.2)
        y = height - margin - h
        parts.append(f'<rect x="{x}" y="{y:.1f}" width="30" height="{h:.1f}" fill="#5f8f3d"/>')
        qy = height - margin - (height - 2 * margin) * float(row["random_q95"]) / (max_o * 1.2)
        parts.append(f'<line x1="{x-3}" x2="{x+33}" y1="{qy:.1f}" y2="{qy:.1f}" stroke="#c43d3d" stroke-width="2"/>')
        parts.append(f'<text x="{x+15}" y="{height-24}" font-family="Arial" font-size="10" text-anchor="middle">{int(row["top_k"])}</text>')

    mx0 = 690
    parts.append('<text x="674" y="56" font-family="Arial" font-size="13">Priority modules</text>')
    mod = module_df[module_df["module_id"].isin(sorted(PRIORITY_MODULES))].copy()
    gx_min, gx_max = mod["mean_geometry_score"].min(), mod["mean_geometry_score"].max()
    ay_min, ay_max = mod["mean_morula_accessibility"].min(), mod["mean_morula_accessibility"].max()
    for _, row in mod.iterrows():
        gx = mx0 + 210 * (float(row["mean_geometry_score"]) - gx_min) / (gx_max - gx_min + EPS)
        gy = 290 - 200 * (float(row["mean_morula_accessibility"]) - ay_min) / (ay_max - ay_min + EPS)
        parts.append(f'<circle cx="{gx:.1f}" cy="{gy:.1f}" r="5" fill="#6b4fa3"/>')
        parts.append(f'<text x="{gx+7:.1f}" y="{gy+4:.1f}" font-family="Arial" font-size="11">{row["module_id"]}</text>')
    parts.extend(
        [
            '<text x="24" y="340" font-family="Arial" font-size="11" fill="#555">Blue/green bars: observed; red ticks: random q95. Generated without matplotlib.</text>',
            "</svg>",
        ]
    )
    out.write_text("\n".join(parts), encoding="utf-8")


def write_interpretation(
    path: Path,
    summary: pd.DataFrame,
    intersections: pd.DataFrame,
    module_df: pd.DataFrame,
    correlations: pd.DataFrame,
    module_controls: pd.DataFrame,
) -> None:
    top25_geo = summary[summary["group"] == "top25_geometry"].iloc[0]
    top25_res = summary[summary["group"] == "top25_residual"].iloc[0]
    access = summary[summary["group"] == "access_branch"].iloc[0]
    inter25 = intersections[intersections["top_k"] == 25].iloc[0]
    m02 = module_df[module_df["module_id"] == "M02"].iloc[0] if (module_df["module_id"] == "M02").any() else None
    geom_corr = correlations[correlations["metric"] == "dmr_geometry_score"].iloc[0]
    neg25 = summary[summary["group"] == "top25_negative_curvature"].iloc[0]
    neg50 = summary[summary["group"] == "top50_negative_curvature"].iloc[0]
    invu = summary[summary["group"] == "inverted_u_DMRs"].iloc[0]
    ushape = summary[summary["group"] == "u_shape_DMRs"].iloc[0]
    q95_modules = module_controls[
        module_controls["accessibility_gt_random_q95"]
        | module_controls["geometry_gt_random_q95"]
        | module_controls["residual_gt_random_q95"]
    ]

    verdict_bits = []
    if bool(top25_geo["observed_gt_random_q95"]):
        verdict_bits.append("high-geometry DMRs exceed module-matched random morula accessibility q95")
    if bool(inter25["observed_gt_random_q95"]):
        verdict_bits.append("top residual and top geometry DMRs overlap beyond module-matched random q95")
    if bool(access["observed_gt_random_q95"]):
        verdict_bits.append("the access branch exceeds module-matched random morula accessibility q95")
    if bool(neg25["observed_gt_random_q95"]) and bool(invu["observed_gt_random_q95"]):
        verdict_bits.append("morula accessibility is directionally enriched in negative-curvature/inverted-U DMRs")
    if not q95_modules.empty:
        verdict_bits.append("individual modules show q95-positive module-level components")
    verdict = "; ".join(verdict_bits) if verdict_bits else "no q95-positive global coupling was detected"

    lines = [
        "# Duality-accessibility coupling analysis",
        "",
        "## Test",
        "",
        "This analysis asks whether morula-centered entry-exit geometry and stage-matched public morula accessibility converge on the same residual DMRs or modules.",
        "",
        "## Main readout",
        "",
        f"- top25 residual DMR morula accessibility: observed={top25_res['observed_morula_accessibility_mean']:.3f}, random q95={top25_res['random_q95']:.3f}, q95-positive={bool(top25_res['observed_gt_random_q95'])}",
        f"- top25 geometry DMR morula accessibility: observed={top25_geo['observed_morula_accessibility_mean']:.3f}, random q95={top25_geo['random_q95']:.3f}, q95-positive={bool(top25_geo['observed_gt_random_q95'])}",
        f"- access branch morula accessibility: observed={access['observed_morula_accessibility_mean']:.3f}, random q95={access['random_q95']:.3f}, q95-positive={bool(access['observed_gt_random_q95'])}",
        f"- top25 residual x top25 geometry overlap: observed={int(inter25['observed_overlap'])}, random q95={inter25['random_q95']:.1f}, q95-positive={bool(inter25['observed_gt_random_q95'])}",
        f"- DMR geometry score versus morula accessibility Spearman rho={geom_corr['observed_spearman']:.3f}; random q05-q95={geom_corr['random_spearman_q05']:.3f} to {geom_corr['random_spearman_q95']:.3f}",
        f"- top25 negative-curvature DMR morula accessibility: observed={neg25['observed_morula_accessibility_mean']:.3f}, random q95={neg25['random_q95']:.3f}, q95-positive={bool(neg25['observed_gt_random_q95'])}",
        f"- top50 negative-curvature DMR morula accessibility: observed={neg50['observed_morula_accessibility_mean']:.3f}, random q95={neg50['random_q95']:.3f}, q95-positive={bool(neg50['observed_gt_random_q95'])}",
        f"- inverted-U DMR morula accessibility: observed={invu['observed_morula_accessibility_mean']:.3f}, random q95={invu['random_q95']:.3f}, q95-positive={bool(invu['observed_gt_random_q95'])}",
        f"- U-shape DMR morula accessibility: observed={ushape['observed_morula_accessibility_mean']:.3f}, random q95={ushape['random_q95']:.3f}, q95-positive={bool(ushape['observed_gt_random_q95'])}",
    ]
    if m02 is not None:
        lines.append(f"- M02 module: geometry score={m02['mean_geometry_score']:.3f}, morula accessibility={m02['mean_morula_accessibility']:.3f}")
    if not q95_modules.empty:
        module_text = ", ".join(
            f"{row.module_id}(access={bool(row.accessibility_gt_random_q95)}, geometry={bool(row.geometry_gt_random_q95)}, residual={bool(row.residual_gt_random_q95)})"
            for row in q95_modules.itertuples()
        )
        lines.append(f"- q95-positive module-level components: {module_text}")

    lines.extend(
        [
            "",
            "## Interpretation",
            "",
            f"Coupling verdict: {verdict}.",
            "",
            "The result should be treated as a convergence test between reset-basin geometry and public chromatin support. The positive coupling is not a broad high-duality or U-shape coupling. Instead, the public morula accessibility signal is concentrated in negative-curvature/inverted-U DMRs, while canonical U-shape DMRs are not accessibility-enriched. It does not identify causal u_bio because there is still no paired perturbation-to-methylation readout.",
            "",
            "## Claim boundary",
            "",
            "Positive coupling can support the statement that morula-centered entry-exit geometry and stage-matched accessibility support converge on a subset of residual/access-associated DMRs. It cannot support final causal u_bio detection.",
        ]
    )
    path.write_text("\n".join(lines) + "\n", encoding="utf-8")


def main() -> None:
    parser = argparse.ArgumentParser()
    parser.add_argument("--result-dir", type=Path, default=Path(r"E:\实验进展5_27"))
    parser.add_argument("--n-iter", type=int, default=1000)
    parser.add_argument("--seed", type=int, default=527)
    args = parser.parse_args()

    result_dir = args.result_dir
    entry = read_tsv(result_dir / "CSB_TRO_2026-05-27_entry_exit_duality_metrics.tsv")
    access = read_tsv(result_dir / "CSB_TRO_2026-05-27_u_bio_rescue_DMR_overlap.tsv")
    merged = prepare_merged(entry, access)

    groups = top_groups(merged)
    rng = np.random.default_rng(args.seed)
    all_indices = merged.index.to_numpy()
    module_to_indices = module_index(merged)
    summary_rows = [
        summarize_group(merged, label, sub, args.n_iter, rng, all_indices, module_to_indices)
        for label, sub in groups.items()
    ]
    summary = pd.DataFrame(summary_rows)
    random_df = random_accessibility_distributions(merged, groups, args.n_iter, args.seed + 1)
    intersections = intersection_summary(merged, args.n_iter, args.seed + 2)
    modules = module_triad(merged)
    correlations = correlation_summary(merged, args.n_iter, args.seed + 3)
    joint = joint_priority_table(merged)
    module_controls = module_random_controls(merged, args.n_iter, args.seed + 4)

    prefix = result_dir / "CSB_TRO_2026-05-27_duality_accessibility"
    merged.to_csv(prefix.with_name(prefix.name + "_DMR_metrics.tsv"), sep="\t", index=False)
    summary.to_csv(prefix.with_name(prefix.name + "_coupling.tsv"), sep="\t", index=False)
    random_df.to_csv(prefix.with_name(prefix.name + "_matched_random.tsv"), sep="\t", index=False)
    intersections.to_csv(prefix.with_name(prefix.name + "_residual_geometry_intersection.tsv"), sep="\t", index=False)
    modules.to_csv(prefix.with_name(prefix.name + "_module_triad.tsv"), sep="\t", index=False)
    module_controls.to_csv(prefix.with_name(prefix.name + "_module_random_controls.tsv"), sep="\t", index=False)
    correlations.to_csv(prefix.with_name(prefix.name + "_correlations.tsv"), sep="\t", index=False)
    joint.to_csv(prefix.with_name(prefix.name + "_joint_priority_DMRs.tsv"), sep="\t", index=False)
    plot_summary(summary, intersections, modules, prefix.with_name(prefix.name + "_summary"))
    write_interpretation(prefix.with_name(prefix.name + "_interpretation.md"), summary, intersections, modules, correlations, module_controls)


if __name__ == "__main__":
    main()
```
