## Supplemental code for "An entropy-based diagnostic framework for characterizing methylation state dynamics during preimplantation development": entry_exit_duality_analysis.py

```
#!/usr/bin/env python
"""Morula-centered entry-exit duality analysis.

Tests whether 8-cell -> morula and morula -> blastocyst changes form a
morula-centered reset-basin entry/exit duality.
"""

from __future__ import annotations

import argparse
from pathlib import Path

import numpy as np
import pandas as pd


ACCESS_MODULES = {"M02", "M10"}
CLOSURE_MODULES = {"M01", "M05", "M12"}
RESIDUAL_MODULES = ACCESS_MODULES | CLOSURE_MODULES
EPS = 1e-12


def read_tsv(path: Path) -> pd.DataFrame:
    if not path.exists():
        raise FileNotFoundError(path)
    return pd.read_csv(path, sep="\t")


def cosine(a: np.ndarray, b: np.ndarray) -> float:
    mask = np.isfinite(a) & np.isfinite(b)
    a = a[mask]
    b = b[mask]
    denom = float(np.linalg.norm(a) * np.linalg.norm(b))
    if len(a) == 0 or denom <= EPS:
        return np.nan
    return float(np.dot(a, b) / denom)


def branch_for_module(module_id: str) -> str:
    if module_id in ACCESS_MODULES:
        return "access"
    if module_id in CLOSURE_MODULES:
        return "closure"
    return "other"


def summarize_group(name: str, df: pd.DataFrame) -> dict:
    entry = df["entry_change"].to_numpy(dtype=float)
    exit_ = df["exit_change"].to_numpy(dtype=float)
    cos = cosine(entry, exit_)
    return {
        "group": name,
        "n_dmr": int(len(df)),
        "mean_beta_8cell": float(df["beta_8cell"].mean()),
        "mean_beta_morula": float(df["beta_morula"].mean()),
        "mean_beta_blastocyst": float(df["beta_blastocyst"].mean()),
        "mean_entry_change": float(df["entry_change"].mean()),
        "mean_exit_change": float(df["exit_change"].mean()),
        "entry_exit_cosine": cos,
        "duality_score_minus_cosine": -cos if np.isfinite(cos) else np.nan,
        "mean_curvature": float(df["curvature"].mean()),
        "median_rebound_ratio": float(df["rebound_ratio"].median()),
        "fraction_u_shape": float(df["is_u_shape"].mean()),
        "fraction_inverted_u": float(df["is_inverted_u"].mean()),
        "fraction_opposite_direction": float((df["entry_change"] * df["exit_change"] < 0).mean()),
    }


def random_duality_controls(dmr: pd.DataFrame, summary: pd.DataFrame, n_iter: int = 1000, seed: int = 27) -> pd.DataFrame:
    rng = np.random.default_rng(seed)
    rows = []

    def score(sub: pd.DataFrame) -> float:
        return -cosine(
            sub["entry_change"].to_numpy(dtype=float),
            sub["exit_change"].to_numpy(dtype=float),
        )

    def module_matched_sample(sub: pd.DataFrame) -> pd.DataFrame:
        pieces = []
        for module_id, count in sub["module_id"].value_counts(dropna=False).items():
            pool = dmr[dmr["module_id"].eq(module_id)]
            replace = len(pool) < count
            if len(pool) == 0:
                pool = dmr
                replace = len(pool) < count
            take = rng.choice(pool.index.to_numpy(), size=int(count), replace=replace)
            pieces.append(dmr.loc[take])
        return pd.concat(pieces, axis=0, ignore_index=True)

    groups = {
        "priority_residual_modules_M01_M02_M05_M10_M12": dmr[dmr["module_id"].isin(RESIDUAL_MODULES)],
        "closure_modules_M01_M05_M12": dmr[dmr["module_id"].isin(CLOSURE_MODULES)],
        "access_modules_M02_M10": dmr[dmr["module_id"].isin(ACCESS_MODULES)],
    }
    ranked = dmr[dmr["basin_residual_rank"].notna()].sort_values("basin_residual_rank")
    for k in [25, 50, 100]:
        groups[f"top{k}_basin_residual_DMRs"] = ranked.head(k)

    module_groups = {
        f"module_{row.module_id}": dmr[dmr["module_id"].eq(row.module_id)]
        for row in dmr[["module_id"]].dropna().drop_duplicates().itertuples(index=False)
    }
    groups.update(module_groups)

    for group_name, sub in groups.items():
        if len(sub) < 2:
            continue
        observed = score(sub)
        mode = "module_matched_random_sets" if group_name.startswith("top") else "size_matched_random_sets"
        vals = []
        for _ in range(n_iter):
            if mode == "module_matched_random_sets":
                rand = module_matched_sample(sub)
            else:
                take = rng.choice(dmr.index.to_numpy(), size=len(sub), replace=False)
                rand = dmr.loc[take]
            vals.append(score(rand))
        arr = np.array(vals, dtype=float)
        rows.append(
            {
                "group": group_name,
                "control_mode": mode,
                "n_dmr": int(len(sub)),
                "observed_duality_score": observed,
                "random_median": float(np.nanmedian(arr)),
                "random_q05": float(np.nanquantile(arr, 0.05)),
                "random_q95": float(np.nanquantile(arr, 0.95)),
                "random_max": float(np.nanmax(arr)),
                "observed_gt_random_q95": bool(observed > np.nanquantile(arr, 0.95)),
                "empirical_p_ge_observed": float((np.sum(arr >= observed) + 1) / (np.sum(np.isfinite(arr)) + 1)),
            }
        )

    # Global null: keep entry vector fixed but permute exit labels across DMRs.
    vals = []
    entry = dmr["entry_change"].to_numpy(dtype=float)
    exit_ = dmr["exit_change"].to_numpy(dtype=float)
    observed = -cosine(entry, exit_)
    for _ in range(n_iter):
        vals.append(-cosine(entry, rng.permutation(exit_)))
    arr = np.array(vals, dtype=float)
    rows.append(
        {
            "group": "all_DMRs_exit_permutation",
            "control_mode": "exit_vector_permutation",
            "n_dmr": int(len(dmr)),
            "observed_duality_score": observed,
            "random_median": float(np.nanmedian(arr)),
            "random_q05": float(np.nanquantile(arr, 0.05)),
            "random_q95": float(np.nanquantile(arr, 0.95)),
            "random_max": float(np.nanmax(arr)),
            "observed_gt_random_q95": bool(observed > np.nanquantile(arr, 0.95)),
            "empirical_p_ge_observed": float((np.sum(arr >= observed) + 1) / (np.sum(np.isfinite(arr)) + 1)),
        }
    )

    return pd.DataFrame(rows)


def build_duality_metrics(root: Path) -> tuple[pd.DataFrame, pd.DataFrame, pd.DataFrame]:
    results = root / "results"
    traj = read_tsv(results / "CSB_TRO_DMR_stage_mean_trajectory.tsv")
    modules = read_tsv(results / "CSB_TRO_DMR_module_assignments.tsv")
    residual = read_tsv(results / "CSB_TRO_basin_residual_DMR_ranking.tsv")

    wide = (
        traj.pivot_table(
            index="cluster_name",
            columns="stage",
            values="mean_beta",
            aggfunc="mean",
        )
        .reset_index()
        .rename_axis(None, axis=1)
    )
    required = ["8-cell", "morula", "blastocyst"]
    missing = [col for col in required if col not in wide.columns]
    if missing:
        raise ValueError(f"Missing stage columns in trajectory table: {missing}")

    dmr = wide[["cluster_name", "8-cell", "morula", "blastocyst"]].rename(
        columns={
            "8-cell": "beta_8cell",
            "morula": "beta_morula",
            "blastocyst": "beta_blastocyst",
        }
    )
    dmr = dmr.merge(modules, on="cluster_name", how="left")
    keep_residual_cols = [
        "cluster_name",
        "basin_residual_rank",
        "latent_residual_delta_beta",
        "abs_latent_residual_delta_beta",
        "signed_latent_residual_direction",
        "chr",
        "start",
        "end",
        "width",
        "n_cpg_target",
    ]
    keep_residual_cols = [c for c in keep_residual_cols if c in residual.columns]
    dmr = dmr.merge(residual[keep_residual_cols], on="cluster_name", how="left")
    dmr["branch"] = dmr["module_id"].map(branch_for_module)

    dmr["entry_change"] = dmr["beta_morula"] - dmr["beta_8cell"]
    dmr["exit_change"] = dmr["beta_blastocyst"] - dmr["beta_morula"]
    dmr["curvature"] = dmr["beta_8cell"] - 2.0 * dmr["beta_morula"] + dmr["beta_blastocyst"]
    dmr["rebound_ratio"] = dmr["exit_change"].abs() / (dmr["entry_change"].abs() + EPS)
    dmr["signed_duality"] = np.where(
        dmr["entry_change"] * dmr["exit_change"] < 0,
        1,
        np.where(dmr["entry_change"] * dmr["exit_change"] > 0, -1, 0),
    )
    dmr["is_u_shape"] = (dmr["beta_8cell"] > dmr["beta_morula"]) & (
        dmr["beta_blastocyst"] > dmr["beta_morula"]
    )
    dmr["is_inverted_u"] = (dmr["beta_8cell"] < dmr["beta_morula"]) & (
        dmr["beta_blastocyst"] < dmr["beta_morula"]
    )
    dmr["is_residual_ranked"] = dmr["basin_residual_rank"].notna()
    dmr["is_priority_residual_module"] = dmr["module_id"].isin(RESIDUAL_MODULES)

    module_rows = []
    for module_id, sub in dmr.dropna(subset=["module_id"]).groupby("module_id"):
        row = summarize_group(module_id, sub)
        row["module_id"] = module_id
        row["branch"] = branch_for_module(module_id)
        row["is_priority_residual_module"] = module_id in RESIDUAL_MODULES
        module_rows.append(row)
    module_summary = pd.DataFrame(module_rows).sort_values(
        ["is_priority_residual_module", "duality_score_minus_cosine", "module_id"],
        ascending=[False, False, True],
    )

    summary_rows = [summarize_group("all_DMRs", dmr)]
    for module_set_name, module_set in [
        ("priority_residual_modules_M01_M02_M05_M10_M12", RESIDUAL_MODULES),
        ("closure_modules_M01_M05_M12", CLOSURE_MODULES),
        ("access_modules_M02_M10", ACCESS_MODULES),
        ("other_modules", set(dmr["module_id"].dropna()) - RESIDUAL_MODULES),
    ]:
        sub = dmr[dmr["module_id"].isin(module_set)]
        if len(sub):
            summary_rows.append(summarize_group(module_set_name, sub))

    for branch, sub in dmr.groupby("branch"):
        summary_rows.append(summarize_group(f"branch_{branch}", sub))

    ranked = dmr[dmr["basin_residual_rank"].notna()].copy()
    if len(ranked):
        ranked = ranked.sort_values("basin_residual_rank")
        for k in [25, 50, 100, 250, 500]:
            sub = ranked.head(k)
            if len(sub):
                row = summarize_group(f"top{k}_basin_residual_DMRs", sub)
                row["rank_cutoff"] = k
                summary_rows.append(row)

    top_entry_path = results / "CSB_TRO_top100_morula_entry_DMRs.tsv"
    top_exit_path = results / "CSB_TRO_top100_blastocyst_exit_DMRs.tsv"
    for name, path in [
        ("top100_morula_entry_dynamic_DMRs", top_entry_path),
        ("top100_blastocyst_exit_dynamic_DMRs", top_exit_path),
    ]:
        if path.exists():
            ids = set(read_tsv(path)["cluster_name"].astype(str))
            sub = dmr[dmr["cluster_name"].isin(ids)]
            if len(sub):
                summary_rows.append(summarize_group(name, sub))

    summary = pd.DataFrame(summary_rows)
    return dmr, module_summary, summary


def write_plot(dmr: pd.DataFrame, module_summary: pd.DataFrame, summary: pd.DataFrame, out_svg: Path, out_png: Path) -> None:
    try:
        import matplotlib.pyplot as plt
    except ModuleNotFoundError:
        write_simple_svg_plot(dmr, module_summary, summary, out_svg)
        write_simple_png_placeholder(summary, out_png)
        return

    fig, axes = plt.subplots(2, 2, figsize=(12, 9))

    selected = summary[
        summary["group"].isin(
            [
                "all_DMRs",
                "top25_basin_residual_DMRs",
                "top50_basin_residual_DMRs",
                "top100_basin_residual_DMRs",
                "priority_residual_modules_M01_M02_M05_M10_M12",
                "closure_modules_M01_M05_M12",
                "access_modules_M02_M10",
            ]
        )
    ].copy()
    selected["label"] = selected["group"].str.replace("_", "\n", regex=False)
    axes[0, 0].bar(selected["label"], selected["duality_score_minus_cosine"], color="#3B6EA8")
    axes[0, 0].axhline(0, color="black", lw=0.8)
    axes[0, 0].set_ylabel("Duality score (-cosine)")
    axes[0, 0].set_title("Entry-exit vector duality")
    axes[0, 0].tick_params(axis="x", labelrotation=45, labelsize=8)

    plot_df = dmr.copy()
    colors = plot_df["branch"].map({"closure": "#B55A30", "access": "#2B8C7E", "other": "#B8B8B8"}).fillna("#B8B8B8")
    axes[0, 1].scatter(plot_df["entry_change"], plot_df["exit_change"], s=8, c=colors, alpha=0.45, linewidths=0)
    ranked = plot_df[plot_df["basin_residual_rank"].notna()].sort_values("basin_residual_rank").head(25)
    axes[0, 1].scatter(ranked["entry_change"], ranked["exit_change"], s=24, facecolors="none", edgecolors="black", linewidths=0.8)
    lim = float(np.nanmax(np.abs(plot_df[["entry_change", "exit_change"]].to_numpy()))) * 1.05
    axes[0, 1].plot([-lim, lim], [lim, -lim], color="black", lw=1.0, ls="--")
    axes[0, 1].axhline(0, color="black", lw=0.6)
    axes[0, 1].axvline(0, color="black", lw=0.6)
    axes[0, 1].set_xlim(-lim, lim)
    axes[0, 1].set_ylim(-lim, lim)
    axes[0, 1].set_xlabel("Entry change: morula - 8-cell")
    axes[0, 1].set_ylabel("Exit change: blastocyst - morula")
    axes[0, 1].set_title("DMR-level entry vs exit")

    mod = module_summary.sort_values("module_id")
    bar_colors = mod["branch"].map({"closure": "#B55A30", "access": "#2B8C7E", "other": "#909090"}).fillna("#909090")
    axes[1, 0].bar(mod["module_id"], mod["duality_score_minus_cosine"], color=bar_colors)
    axes[1, 0].axhline(0, color="black", lw=0.8)
    axes[1, 0].set_ylabel("Duality score (-cosine)")
    axes[1, 0].set_title("Module-level duality")

    axes[1, 1].bar(mod["module_id"], mod["fraction_u_shape"], color=bar_colors)
    axes[1, 1].set_ylim(0, 1)
    axes[1, 1].set_ylabel("Fraction U-shape")
    axes[1, 1].set_title("Morula-centered U-shape fraction")

    fig.suptitle("Morula-centered entry-exit duality analysis", fontsize=14)
    fig.tight_layout(rect=(0, 0, 1, 0.96))
    fig.savefig(out_svg)
    fig.savefig(out_png, dpi=200)
    plt.close(fig)


def _svg_text(x: float, y: float, text: str, size: int = 12, anchor: str = "start", weight: str = "normal") -> str:
    safe = (
        str(text)
        .replace("&", "&amp;")
        .replace("<", "&lt;")
        .replace(">", "&gt;")
        .replace('"', "&quot;")
    )
    return f'<text x="{x:.1f}" y="{y:.1f}" font-size="{size}" text-anchor="{anchor}" font-weight="{weight}" font-family="Arial">{safe}</text>'


def write_simple_svg_plot(dmr: pd.DataFrame, module_summary: pd.DataFrame, summary: pd.DataFrame, out_svg: Path) -> None:
    width, height = 1200, 900
    parts = [
        f'<svg xmlns="http://www.w3.org/2000/svg" width="{width}" height="{height}" viewBox="0 0 {width} {height}">',
        '<rect width="100%" height="100%" fill="white"/>',
        _svg_text(600, 32, "Morula-centered entry-exit duality analysis", 20, "middle", "bold"),
    ]

    selected_groups = [
        "all_DMRs",
        "top25_basin_residual_DMRs",
        "top50_basin_residual_DMRs",
        "top100_basin_residual_DMRs",
        "priority_residual_modules_M01_M02_M05_M10_M12",
        "closure_modules_M01_M05_M12",
        "access_modules_M02_M10",
    ]
    selected = summary[summary["group"].isin(selected_groups)].copy()
    selected["order"] = selected["group"].map({g: i for i, g in enumerate(selected_groups)})
    selected = selected.sort_values("order")

    # Panel A: group duality bars.
    x0, y0, w, h = 65, 80, 500, 300
    parts.append(_svg_text(x0 + w / 2, y0 - 22, "Entry-exit vector duality", 15, "middle", "bold"))
    parts.append(f'<line x1="{x0}" y1="{y0+h}" x2="{x0+w}" y2="{y0+h}" stroke="black" stroke-width="1"/>')
    vals = selected["duality_score_minus_cosine"].astype(float).to_numpy()
    max_abs = max(0.25, float(np.nanmax(np.abs(vals))) if len(vals) else 0.25)
    zero_y = y0 + h / 2
    parts.append(f'<line x1="{x0}" y1="{zero_y}" x2="{x0+w}" y2="{zero_y}" stroke="#555" stroke-width="1"/>')
    bar_w = w / max(1, len(selected)) * 0.65
    for i, row in enumerate(selected.itertuples(index=False)):
        val = float(row.duality_score_minus_cosine)
        cx = x0 + (i + 0.5) * w / len(selected)
        y_val = zero_y - (val / max_abs) * (h / 2 - 22)
        y_top = min(zero_y, y_val)
        bh = abs(y_val - zero_y)
        parts.append(f'<rect x="{cx-bar_w/2:.1f}" y="{y_top:.1f}" width="{bar_w:.1f}" height="{bh:.1f}" fill="#3B6EA8"/>')
        label = str(row.group).replace("_basin_residual_DMRs", "").replace("_modules", "").replace("_", " ")
        parts.append(_svg_text(cx, y0 + h + 18, label[:18], 9, "middle"))
        parts.append(_svg_text(cx, y_top - 4, f"{val:.2f}", 9, "middle"))

    # Panel B: entry vs exit scatter.
    x1, y1, w1, h1 = 665, 80, 470, 300
    parts.append(_svg_text(x1 + w1 / 2, y1 - 22, "DMR-level entry vs exit", 15, "middle", "bold"))
    lim = float(np.nanmax(np.abs(dmr[["entry_change", "exit_change"]].to_numpy())))
    lim = max(lim, 0.01) * 1.05
    parts.append(f'<rect x="{x1}" y="{y1}" width="{w1}" height="{h1}" fill="none" stroke="#333"/>')
    parts.append(f'<line x1="{x1}" y1="{y1+h1/2}" x2="{x1+w1}" y2="{y1+h1/2}" stroke="#777"/>')
    parts.append(f'<line x1="{x1+w1/2}" y1="{y1}" x2="{x1+w1/2}" y2="{y1+h1}" stroke="#777"/>')
    parts.append(f'<line x1="{x1}" y1="{y1}" x2="{x1+w1}" y2="{y1+h1}" stroke="#333" stroke-dasharray="5,5"/>')
    palette = {"closure": "#B55A30", "access": "#2B8C7E", "other": "#B8B8B8"}
    sample = dmr.sample(min(len(dmr), 2500), random_state=1) if len(dmr) > 2500 else dmr
    top25 = set(dmr[dmr["basin_residual_rank"].notna()].sort_values("basin_residual_rank").head(25)["cluster_name"])
    for row in sample.itertuples(index=False):
        px = x1 + ((float(row.entry_change) + lim) / (2 * lim)) * w1
        py = y1 + h1 - ((float(row.exit_change) + lim) / (2 * lim)) * h1
        color = palette.get(row.branch, "#B8B8B8")
        radius = 3.2 if row.cluster_name in top25 else 1.7
        stroke = "black" if row.cluster_name in top25 else "none"
        fill = "none" if row.cluster_name in top25 else color
        parts.append(f'<circle cx="{px:.1f}" cy="{py:.1f}" r="{radius}" fill="{fill}" stroke="{stroke}" stroke-width="0.8" opacity="0.65"/>')
    parts.append(_svg_text(x1 + w1 / 2, y1 + h1 + 28, "Entry: morula - 8-cell", 11, "middle"))
    parts.append(_svg_text(x1 - 8, y1 + h1 / 2, "Exit", 11, "end"))

    # Panel C/D: module duality and U-shape fractions.
    for panel, col, title, ybase in [
        ("C", "duality_score_minus_cosine", "Module-level duality", 470),
        ("D", "fraction_u_shape", "Module U-shape fraction", 470),
    ]:
        px0 = 65 if panel == "C" else 665
        pw, ph = 500 if panel == "C" else 470, 300
        mod = module_summary.sort_values("module_id")
        vals = mod[col].astype(float).to_numpy()
        max_v = max(1.0 if col == "fraction_u_shape" else 0.25, float(np.nanmax(np.abs(vals))) if len(vals) else 1.0)
        zero = ybase + ph if col == "fraction_u_shape" else ybase + ph / 2
        parts.append(_svg_text(px0 + pw / 2, ybase - 22, title, 15, "middle", "bold"))
        parts.append(f'<line x1="{px0}" y1="{zero:.1f}" x2="{px0+pw}" y2="{zero:.1f}" stroke="#555"/>')
        bw = pw / max(1, len(mod)) * 0.7
        for i, row in enumerate(mod.itertuples(index=False)):
            val = float(getattr(row, col))
            cx = px0 + (i + 0.5) * pw / len(mod)
            if col == "fraction_u_shape":
                yv = ybase + ph - (val / max_v) * (ph - 18)
            else:
                yv = zero - (val / max_v) * (ph / 2 - 18)
            yt = min(zero, yv)
            bh = abs(zero - yv)
            color = palette.get(row.branch, "#909090")
            parts.append(f'<rect x="{cx-bw/2:.1f}" y="{yt:.1f}" width="{bw:.1f}" height="{bh:.1f}" fill="{color}"/>')
            parts.append(_svg_text(cx, ybase + ph + 18, row.module_id, 9, "middle"))

    parts.append("</svg>")
    out_svg.write_text("\n".join(parts), encoding="utf-8")


def write_simple_png_placeholder(summary: pd.DataFrame, out_png: Path) -> None:
    from PIL import Image, ImageDraw

    img = Image.new("RGB", (1200, 420), "white")
    draw = ImageDraw.Draw(img)
    draw.text((40, 30), "Morula-centered entry-exit duality analysis", fill="black")
    y = 80
    for _, row in summary.head(12).iterrows():
        text = (
            f"{row['group']}: n={int(row['n_dmr'])}, "
            f"duality={float(row['duality_score_minus_cosine']):.3f}, "
            f"U-shape={float(row['fraction_u_shape']):.3f}"
        )
        draw.text((40, y), text, fill="black")
        y += 26
    img.save(out_png)


def write_interpretation(summary: pd.DataFrame, module_summary: pd.DataFrame, out_md: Path) -> None:
    def md_table(df: pd.DataFrame, cols: list[str]) -> str:
        rows = ["| " + " | ".join(cols) + " |", "| " + " | ".join(["---"] * len(cols)) + " |"]
        for _, r in df[cols].iterrows():
            vals = []
            for c in cols:
                v = r[c]
                if isinstance(v, (float, np.floating)):
                    vals.append(f"{float(v):.3f}")
                else:
                    vals.append(str(v))
            rows.append("| " + " | ".join(vals) + " |")
        return "\n".join(rows)

    def get(group: str, col: str) -> float:
        row = summary.loc[summary["group"] == group]
        if row.empty:
            return np.nan
        return float(row.iloc[0][col])

    top_modules = module_summary.sort_values("duality_score_minus_cosine", ascending=False).head(8)
    priority = module_summary[module_summary["is_priority_residual_module"]].sort_values(
        "duality_score_minus_cosine", ascending=False
    )

    random_path = out_md.with_name("CSB_TRO_2026-05-27_entry_exit_random_controls.tsv")
    random_md = ""
    if random_path.exists():
        random_df = pd.read_csv(random_path, sep="\t")
        key_random = random_df[
            random_df["group"].isin(
                [
                    "all_DMRs_exit_permutation",
                    "top25_basin_residual_DMRs",
                    "top50_basin_residual_DMRs",
                    "top100_basin_residual_DMRs",
                    "access_modules_M02_M10",
                    "closure_modules_M01_M05_M12",
                    "module_M02",
                ]
            )
        ]
        random_md = md_table(
            key_random,
            [
                "group",
                "control_mode",
                "observed_duality_score",
                "random_median",
                "random_q95",
                "observed_gt_random_q95",
                "empirical_p_ge_observed",
            ],
        )

    lines = [
        "# Morula-centered entry-exit duality analysis",
        "",
        "## Test",
        "",
        "This analysis tests whether the 8-cell-to-morula entry vector and morula-to-blastocyst exit vector form a morula-centered duality:",
        "",
        "`Delta_entry = beta_morula - beta_8cell`",
        "",
        "`Delta_exit = beta_blastocyst - beta_morula`",
        "",
        "`duality_score = -cos(Delta_entry, Delta_exit)`",
        "",
        "`curvature = beta_8cell - 2 * beta_morula + beta_blastocyst`",
        "",
        "## Main readout",
        "",
        f"- all DMR duality score: {get('all_DMRs', 'duality_score_minus_cosine'):.3f}",
        f"- all DMR U-shape fraction: {get('all_DMRs', 'fraction_u_shape'):.3f}",
        f"- priority residual-module duality score: {get('priority_residual_modules_M01_M02_M05_M10_M12', 'duality_score_minus_cosine'):.3f}",
        f"- priority residual-module U-shape fraction: {get('priority_residual_modules_M01_M02_M05_M10_M12', 'fraction_u_shape'):.3f}",
        f"- top25 basin residual DMR duality score: {get('top25_basin_residual_DMRs', 'duality_score_minus_cosine'):.3f}",
        f"- top25 basin residual DMR U-shape fraction: {get('top25_basin_residual_DMRs', 'fraction_u_shape'):.3f}",
        "",
        "## Random-control boundary",
        "",
        random_md or "Random-control table was not found.",
        "",
        "## Priority module ranking",
        "",
        md_table(
            priority,
            [
                "module_id",
                "branch",
                "n_dmr",
                "duality_score_minus_cosine",
                "fraction_u_shape",
                "mean_curvature",
                "median_rebound_ratio",
            ],
        ),
        "",
        "## Strongest module-level duality signals",
        "",
        md_table(
            top_modules,
            [
                "module_id",
                "branch",
                "n_dmr",
                "duality_score_minus_cosine",
                "fraction_u_shape",
                "mean_curvature",
                "median_rebound_ratio",
            ],
        ),
        "",
        "## Claim boundary",
        "",
        "A positive score supports morula-centered entry-exit geometry. It does not by itself identify a causal biological input. The result should be written as a methylation-state-space geometry result and, if concentrated in priority residual modules, as module-specific reset-basin entry-exit duality rather than global strict symmetry.",
        "",
    ]
    out_md.write_text("\n".join(lines), encoding="utf-8")


def main() -> None:
    parser = argparse.ArgumentParser()
    parser.add_argument("--root", default=r"E:\CSB_TRO_operator_time_DMR_dynamics_2026-05-25")
    parser.add_argument("--out", default=r"E:\实验进展5_27")
    args = parser.parse_args()

    root = Path(args.root)
    out = Path(args.out)
    out.mkdir(parents=True, exist_ok=True)

    dmr, module_summary, summary = build_duality_metrics(root)
    random_controls = random_duality_controls(dmr, summary, n_iter=1000, seed=27)

    prefix = "CSB_TRO_2026-05-27_entry_exit"
    dmr_path = out / f"{prefix}_duality_metrics.tsv"
    module_path = out / f"{prefix}_module_duality.tsv"
    summary_path = out / f"{prefix}_summary.tsv"
    random_path = out / f"{prefix}_random_controls.tsv"
    curvature_path = out / f"{prefix}_curvature.tsv"
    svg_path = out / f"{prefix}_duality_summary.svg"
    png_path = out / f"{prefix}_duality_summary.png"
    md_path = out / f"{prefix}_duality_interpretation.md"

    dmr.to_csv(dmr_path, sep="\t", index=False)
    module_summary.to_csv(module_path, sep="\t", index=False)
    summary.to_csv(summary_path, sep="\t", index=False)
    random_controls.to_csv(random_path, sep="\t", index=False)
    dmr[
        [
            "cluster_name",
            "module_id",
            "branch",
            "beta_8cell",
            "beta_morula",
            "beta_blastocyst",
            "entry_change",
            "exit_change",
            "curvature",
            "rebound_ratio",
            "is_u_shape",
            "is_inverted_u",
            "basin_residual_rank",
        ]
    ].to_csv(curvature_path, sep="\t", index=False)

    write_plot(dmr, module_summary, summary, svg_path, png_path)
    write_interpretation(summary, module_summary, md_path)

    print(f"Wrote {dmr_path}")
    print(f"Wrote {module_path}")
    print(f"Wrote {summary_path}")
    print(f"Wrote {random_path}")
    print(f"Wrote {curvature_path}")
    print(f"Wrote {svg_path}")
    print(f"Wrote {png_path}")
    print(f"Wrote {md_path}")


if __name__ == "__main__":
    main()
```
