## Supplemental code for "An entropy-based diagnostic framework for characterizing methylation state dynamics during preimplantation development": FINAL_DYNAMICS_PACKAGE_README.md

# CSB/TRO Latest Dynamics Package

This folder freezes the 2026-05-27 project state after the latest morula-centered dynamics updates.

## Core Model State

The current model is best described as:

`perturbation-informed, chromatin-associated, morula-centered diagnostic reset-basin geometry`

Supported:

- methylation-only operator-time propagation fails at morula reset-basin entry;
- the measured correction is non-random, module-specific, direction-sensitive, and organized by closure/access branches;
- entry-exit analysis supports morula as a non-random reset-basin geometric vertex;
- public human morula accessibility partially supports the strongest residual DMRs;
- accessibility further enriches negative-curvature/inverted-U DMRs, suggesting a directional transient chromatin component;
- CBP/p300-HDAC perturbation literature supports the access/closure machinery axis as perturbable.

Not supported:

- final causal `u_bio` detection;
- `do(u_bio) -> Delta c_tau -> Delta x_tau+1`;
- global chromatin replacement;
- genome-wide perfect quadratic rebound.

## Folder Layout

- `code/`: current scripts used for the latest analyses and figure generation.
- `results_all/`: all top-level files from the archived results directory at package time.
- `tables/`: TSV/JSON/HTML outputs and evidence tables.
- `manuscript/`: manuscript drafts, interpretation files, figure plan, and summaries.
- `figures_existing/`: pre-existing generated figures from the result directory.
- `figures_new/`: new publication-style figures generated for this frozen package.
- `downloads_and_second_source/`: lightweight downloaded/audit output directories used for public chromatin and second-source controls.

## New Figure Set

The `figures_new/` folder contains 14 figure stems, each exported as PNG, SVG, and PDF:

1. dynamics roadmap
2. methylation-only occupancy failure
3. entry-exit scatter
4. module duality lollipop
5. duality random controls
6. curvature distribution
7. public chromatin rescue top-k
8. curvature-accessibility coupling
9. residual-geometry intersection
10. module triad bubble
11. joint-priority DMR heatmap
12. evidence boundary matrix
13. integrated reset-basin model
14. joint candidate lollipop

Generated with:

```text
python code_current\make_latest_dynamics_figure_package.py --result-dir <RESULT_DIR> --out-dir <RESULT_DIR>\CSB_TRO_2026-05-27_FINAL_DYNAMICS_PACKAGE\figures_new
```

## Most Recent Dynamics Result

The latest coupling analysis did not show broad high-duality/accessibility coupling. The stronger new signal is directional:

- top25 negative-curvature DMRs: morula accessibility observed 1.793 vs module-matched random q95 1.623;
- top50 negative-curvature DMRs: observed 1.630 vs q95 1.553;
- inverted-U DMRs: observed 1.716 vs q95 1.591;
- canonical U-shape DMRs are not accessibility-enriched.

This supports a bounded statement: public morula accessibility aligns with a negative-curvature/inverted-U transient component of morula-centered reset-basin geometry.
