## Supplemental code for "An entropy-based diagnostic framework for characterizing methylation state dynamics during preimplantation development": make_latest_dynamics_figure_package.py

```
#!/usr/bin/env python
"""Generate publication-style figures for the latest CSB/TRO dynamics package."""

from __future__ import annotations

import argparse
import json
import math
from pathlib import Path

import matplotlib

matplotlib.use("Agg")
import matplotlib.pyplot as plt
from matplotlib.patches import FancyArrowPatch, Rectangle
import numpy as np
import pandas as pd
import seaborn as sns


PALETTE = {
    "ink": "#1f2933",
    "muted": "#6b7280",
    "grid": "#d9dee7",
    "blue": "#2f6fb0",
    "teal": "#1b8a8f",
    "green": "#4c8c3f",
    "orange": "#c8752a",
    "red": "#b94747",
    "purple": "#7356a5",
    "gold": "#c49a31",
    "gray": "#94a3b8",
}

BRANCH_COLORS = {
    "access": "#2f6fb0",
    "closure": "#b94747",
    "other": "#94a3b8",
}


def read_tsv(path: Path) -> pd.DataFrame:
    return pd.read_csv(path, sep="\t")


def num(df: pd.DataFrame, cols: list[str]) -> pd.DataFrame:
    for col in cols:
        if col in df.columns:
            df[col] = pd.to_numeric(df[col], errors="coerce")
    return df


def setup_style() -> None:
    sns.set_theme(style="whitegrid", context="paper")
    plt.rcParams.update(
        {
            "font.family": "DejaVu Sans",
            "axes.edgecolor": PALETTE["ink"],
            "axes.labelcolor": PALETTE["ink"],
            "xtick.color": PALETTE["ink"],
            "ytick.color": PALETTE["ink"],
            "grid.color": PALETTE["grid"],
            "grid.linewidth": 0.7,
            "figure.facecolor": "white",
            "axes.facecolor": "white",
            "savefig.facecolor": "white",
            "svg.fonttype": "none",
            "pdf.fonttype": 42,
            "ps.fonttype": 42,
        }
    )


def save(fig: plt.Figure, out_dir: Path, name: str) -> None:
    out_dir.mkdir(parents=True, exist_ok=True)
    fig.tight_layout()
    for ext in ["png", "svg", "pdf"]:
        fig.savefig(out_dir / f"{name}.{ext}", dpi=320, bbox_inches="tight")
    plt.close(fig)


def annotate_panel(ax: plt.Axes, label: str) -> None:
    ax.text(
        -0.08,
        1.08,
        label,
        transform=ax.transAxes,
        fontsize=12,
        fontweight="bold",
        va="top",
        ha="left",
        color=PALETTE["ink"],
    )


def load_all(result_dir: Path) -> dict[str, pd.DataFrame]:
    data = {
        "entry_metrics": read_tsv(result_dir / "CSB_TRO_2026-05-27_entry_exit_duality_metrics.tsv"),
        "entry_summary": read_tsv(result_dir / "CSB_TRO_2026-05-27_entry_exit_summary.tsv"),
        "entry_random": read_tsv(result_dir / "CSB_TRO_2026-05-27_entry_exit_random_controls.tsv"),
        "module_duality": read_tsv(result_dir / "CSB_TRO_2026-05-27_entry_exit_module_duality.tsv"),
        "rescue_summary": read_tsv(result_dir / "CSB_TRO_2026-05-27_u_bio_rescue_overlap_summary.tsv"),
        "rescue_dmr": read_tsv(result_dir / "CSB_TRO_2026-05-27_u_bio_rescue_DMR_overlap.tsv"),
        "coupling": read_tsv(result_dir / "CSB_TRO_2026-05-27_duality_accessibility_coupling.tsv"),
        "coupling_intersection": read_tsv(result_dir / "CSB_TRO_2026-05-27_duality_accessibility_residual_geometry_intersection.tsv"),
        "module_triad": read_tsv(result_dir / "CSB_TRO_2026-05-27_duality_accessibility_module_triad.tsv"),
        "module_controls": read_tsv(result_dir / "CSB_TRO_2026-05-27_duality_accessibility_module_random_controls.tsv"),
        "joint_priority": read_tsv(result_dir / "CSB_TRO_2026-05-27_duality_accessibility_joint_priority_DMRs.tsv"),
        "claim_boundary": read_tsv(result_dir / "CSB_TRO_2026-05-27_claim_boundary_solved_unsolved_v1.0.tsv"),
        "evidence_boundary": read_tsv(result_dir / "CSB_TRO_2026-05-27_evidence_boundary_table.tsv"),
    }
    numeric_cols = [
        "beta_8cell",
        "beta_morula",
        "beta_blastocyst",
        "entry_change",
        "exit_change",
        "curvature",
        "rebound_ratio",
        "signed_duality",
        "basin_residual_rank",
        "abs_latent_residual_delta_beta",
        "public_accessibility_morula_max",
        "public_accessibility_morula_minus_8cell",
        "duality_score_minus_cosine",
        "observed_morula_accessibility_mean",
        "random_q95",
        "observed_mean",
        "random_median",
        "observed_overlap",
        "top_k",
        "mean_geometry_score",
        "mean_morula_accessibility",
        "joint_reset_geometry_access_score",
    ]
    for key, df in data.items():
        data[key] = num(df, numeric_cols)
    return data


def fig_01_model_roadmap(data: dict[str, pd.DataFrame], out_dir: Path) -> None:
    fig, ax = plt.subplots(figsize=(12, 4.6))
    ax.axis("off")
    stages = [
        ("Methylation-only\noperator", "q90 occupancy\n0.044"),
        ("Measured\ncorrection", "observed occupancy\n0.875"),
        ("Module residual\narchitecture", "M05/M01/M12\n+ M02/M10"),
        ("Closure/access\nbranches", "correct orientation\n0.956"),
        ("Entry-exit\ngeometry", "cosine -0.699\nq95 0.133"),
        ("Chromatin-coupled\ncurvature", "inverted-U access\nq95-positive"),
    ]
    xs = np.linspace(0.06, 0.94, len(stages))
    for i, ((title, subtitle), x) in enumerate(zip(stages, xs)):
        color = [PALETTE["blue"], PALETTE["teal"], PALETTE["purple"], PALETTE["red"], PALETTE["green"], PALETTE["gold"]][i]
        ax.add_patch(Rectangle((x - 0.07, 0.38), 0.14, 0.28, facecolor=color, alpha=0.12, edgecolor=color, lw=1.5))
        ax.text(x, 0.59, title, ha="center", va="center", fontsize=10, fontweight="bold", color=PALETTE["ink"])
        ax.text(x, 0.45, subtitle, ha="center", va="center", fontsize=8.5, color=PALETTE["muted"])
        if i < len(stages) - 1:
            ax.add_patch(
                FancyArrowPatch(
                    (x + 0.075, 0.52),
                    (xs[i + 1] - 0.075, 0.52),
                    arrowstyle="-|>",
                    mutation_scale=14,
                    lw=1.5,
                    color=PALETTE["ink"],
                )
            )
    ax.text(0.5, 0.82, "Current dynamics progression", ha="center", fontsize=15, fontweight="bold", color=PALETTE["ink"])
    ax.text(
        0.5,
        0.22,
        "From morula-entry methylation-only failure to perturbation-informed, chromatin-associated reset-basin geometry",
        ha="center",
        fontsize=10,
        color=PALETTE["muted"],
    )
    save(fig, out_dir, "Fig01_dynamics_roadmap")


def fig_02_operator_failure(out_dir: Path) -> None:
    fig, ax = plt.subplots(figsize=(5.8, 4.2))
    vals = pd.DataFrame(
        {
            "state": ["Methylation-only\nprediction", "Observed\nmorula"],
            "q90 occupancy": [0.044, 0.875],
        }
    )
    sns.barplot(data=vals, x="state", y="q90 occupancy", ax=ax, palette=[PALETTE["gray"], PALETTE["blue"]])
    for i, v in enumerate(vals["q90 occupancy"]):
        ax.text(i, v + 0.035, f"{v:.3f}", ha="center", fontweight="bold")
    ax.set_ylim(0, 1.05)
    ax.set_xlabel("")
    ax.set_ylabel("Morula basin occupancy")
    ax.set_title("Methylation-only propagation fails at morula entry", fontweight="bold")
    save(fig, out_dir, "Fig02_operator_failure_occupancy")


def fig_03_entry_exit_scatter(data: dict[str, pd.DataFrame], out_dir: Path) -> None:
    df = data["entry_metrics"].copy()
    fig, ax = plt.subplots(figsize=(6.2, 5.4))
    for branch, sub in df.groupby("branch"):
        ax.scatter(
            sub["entry_change"],
            sub["exit_change"],
            s=np.clip(sub["abs_latent_residual_delta_beta"].fillna(0) * 600, 18, 130),
            alpha=0.75,
            label=branch,
            color=BRANCH_COLORS.get(branch, PALETTE["gray"]),
            edgecolor="white",
            linewidth=0.45,
        )
    lim = np.nanmax(np.abs(df[["entry_change", "exit_change"]].to_numpy())) * 1.08
    ax.plot([-lim, lim], [lim, -lim], color=PALETTE["ink"], lw=1.2, ls="--", label="anti-diagonal")
    ax.axhline(0, color=PALETTE["grid"], lw=1)
    ax.axvline(0, color=PALETTE["grid"], lw=1)
    ax.set_xlim(-lim, lim)
    ax.set_ylim(-lim, lim)
    ax.set_xlabel("Entry change: morula - 8-cell")
    ax.set_ylabel("Exit change: blastocyst - morula")
    ax.set_title("Morula-centered entry-exit anti-alignment", fontweight="bold")
    ax.legend(frameon=False, loc="best")
    save(fig, out_dir, "Fig03_entry_exit_scatter")


def fig_04_module_duality(data: dict[str, pd.DataFrame], out_dir: Path) -> None:
    df = data["module_duality"].copy().sort_values("duality_score_minus_cosine", ascending=True)
    fig, ax = plt.subplots(figsize=(7.2, 6.2))
    colors = [BRANCH_COLORS.get(b, PALETTE["gray"]) for b in df["branch"]]
    ax.hlines(df["module_id"], 0, df["duality_score_minus_cosine"], color=colors, lw=3, alpha=0.75)
    ax.scatter(df["duality_score_minus_cosine"], df["module_id"], color=colors, s=70, edgecolor="white", linewidth=0.6)
    ax.axvline(0, color=PALETTE["ink"], lw=1)
    ax.set_xlabel("Entry-exit duality score (-cosine)")
    ax.set_ylabel("Module")
    ax.set_title("Module-level reset-basin geometry", fontweight="bold")
    save(fig, out_dir, "Fig04_module_duality_lollipop")


def fig_05_random_control_duality(data: dict[str, pd.DataFrame], out_dir: Path) -> None:
    df = data["entry_random"].copy()
    fig, ax = plt.subplots(figsize=(8.4, 4.8))
    order = df.sort_values("observed_duality_score", ascending=False)["group"]
    df = df.set_index("group").loc[order].reset_index()
    x = np.arange(len(df))
    ax.bar(x, df["observed_duality_score"], color=PALETTE["green"], alpha=0.85, label="observed")
    ax.scatter(x, df["random_q95"], color=PALETTE["red"], s=42, label="random q95", zorder=3)
    ax.set_xticks(x)
    ax.set_xticklabels(df["group"], rotation=35, ha="right", fontsize=8)
    ax.set_ylabel("Duality score")
    ax.set_title("Entry-exit duality exceeds matched/permutation controls in selected groups", fontweight="bold")
    ax.legend(frameon=False)
    save(fig, out_dir, "Fig05_duality_random_controls")


def fig_06_curvature_distribution(data: dict[str, pd.DataFrame], out_dir: Path) -> None:
    df = data["entry_metrics"].copy()
    fig, axes = plt.subplots(1, 2, figsize=(10.5, 4.4))
    sns.histplot(data=df, x="curvature", hue="branch", bins=35, element="step", stat="density", common_norm=False, ax=axes[0], palette=BRANCH_COLORS)
    axes[0].axvline(0, color=PALETTE["ink"], lw=1)
    axes[0].set_title("Curvature distribution")
    axes[0].set_xlabel("beta_8cell - 2 beta_morula + beta_blastocyst")
    branch_order = ["access", "closure", "other"]
    sns.boxplot(data=df, x="branch", y="curvature", order=branch_order, ax=axes[1], palette=BRANCH_COLORS, fliersize=2)
    axes[1].axhline(0, color=PALETTE["ink"], lw=1)
    axes[1].set_title("Curvature by branch")
    axes[1].set_xlabel("")
    axes[1].set_ylabel("Curvature")
    save(fig, out_dir, "Fig06_curvature_distribution")


def fig_07_public_rescue_topk(data: dict[str, pd.DataFrame], out_dir: Path) -> None:
    df = data["rescue_summary"].copy()
    df = df[df["metric"].eq("public_accessibility_morula_max")].sort_values("top_k")
    fig, ax = plt.subplots(figsize=(6.6, 4.4))
    x = np.arange(len(df))
    ax.bar(x, df["observed_mean"], color=PALETTE["blue"], alpha=0.85, label="observed")
    ax.scatter(x, df["random_q95"], color=PALETTE["red"], s=58, label="matched random q95", zorder=3)
    ax.plot(x, df["random_median"], color=PALETTE["muted"], marker="o", lw=1.2, label="random median")
    ax.set_xticks(x)
    ax.set_xticklabels([f"top{int(k)}" for k in df["top_k"]])
    ax.set_ylabel("Mean morula accessibility")
    ax.set_title("Stage-matched public chromatin rescue is strongest at top25", fontweight="bold")
    ax.legend(frameon=False)
    save(fig, out_dir, "Fig07_public_chromatin_rescue_topk")


def fig_08_coupling_curvature(data: dict[str, pd.DataFrame], out_dir: Path) -> None:
    df = data["coupling"].copy()
    groups = [
        "top25_residual",
        "top25_geometry",
        "top25_negative_curvature",
        "top50_negative_curvature",
        "inverted_u_DMRs",
        "u_shape_DMRs",
    ]
    sel = df[df["group"].isin(groups)].set_index("group").loc[groups].reset_index()
    fig, ax = plt.subplots(figsize=(8.5, 4.8))
    x = np.arange(len(sel))
    colors = [PALETTE["blue"], PALETTE["purple"], PALETTE["gold"], PALETTE["gold"], PALETTE["orange"], PALETTE["gray"]]
    ax.bar(x, sel["observed_morula_accessibility_mean"], color=colors, alpha=0.88)
    ax.scatter(x, sel["random_q95"], color=PALETTE["red"], s=52, zorder=3, label="random q95")
    ax.set_xticks(x)
    ax.set_xticklabels(sel["group"], rotation=30, ha="right")
    ax.set_ylabel("Mean morula accessibility")
    ax.set_title("Accessibility couples to negative-curvature/inverted-U geometry", fontweight="bold")
    ax.legend(frameon=False)
    save(fig, out_dir, "Fig08_curvature_accessibility_coupling")


def fig_09_residual_geometry_intersection(data: dict[str, pd.DataFrame], out_dir: Path) -> None:
    df = data["coupling_intersection"].copy().sort_values("top_k")
    fig, ax = plt.subplots(figsize=(6.6, 4.2))
    x = np.arange(len(df))
    ax.bar(x, df["observed_overlap"], color=PALETTE["teal"], alpha=0.9, label="observed overlap")
    ax.scatter(x, df["random_q95"], color=PALETTE["red"], s=55, zorder=3, label="random q95")
    ax.set_xticks(x)
    ax.set_xticklabels([f"top{int(k)}" for k in df["top_k"]])
    ax.set_ylabel("Residual ∩ geometry DMR count")
    ax.set_title("Top residual and top geometry sets are only partially overlapping", fontweight="bold")
    ax.legend(frameon=False)
    save(fig, out_dir, "Fig09_residual_geometry_intersection")


def fig_10_module_triad_bubble(data: dict[str, pd.DataFrame], out_dir: Path) -> None:
    df = data["module_triad"].copy()
    fig, ax = plt.subplots(figsize=(7.2, 5.4))
    sizes = np.clip(df["mean_abs_residual"].fillna(0) * 2300, 35, 360)
    colors = [BRANCH_COLORS.get(b, PALETTE["gray"]) for b in df["branch"]]
    ax.scatter(df["mean_geometry_score"], df["mean_morula_accessibility"], s=sizes, color=colors, alpha=0.78, edgecolor="white", linewidth=0.7)
    for _, row in df.iterrows():
        if row["module_id"] in {"M01", "M02", "M05", "M10", "M12", "M06", "M14"}:
            ax.text(row["mean_geometry_score"] + 0.012, row["mean_morula_accessibility"], row["module_id"], fontsize=9, fontweight="bold")
    ax.set_xlabel("Mean DMR geometry score")
    ax.set_ylabel("Mean morula accessibility")
    ax.set_title("Module triad: geometry, accessibility, residual strength", fontweight="bold")
    ax.text(0.02, 0.04, "Bubble size = mean absolute residual", transform=ax.transAxes, color=PALETTE["muted"], fontsize=8)
    save(fig, out_dir, "Fig10_module_triad_bubble")


def fig_11_joint_priority_heatmap(data: dict[str, pd.DataFrame], out_dir: Path) -> None:
    df = data["joint_priority"].head(25).copy()
    cols = [
        "abs_latent_residual_delta_beta",
        "dmr_geometry_score",
        "public_accessibility_morula_max",
        "public_accessibility_morula_minus_8cell",
        "joint_reset_geometry_access_score",
    ]
    mat = df[cols].apply(pd.to_numeric, errors="coerce").astype(float)
    mat = pd.DataFrame(
        (mat.to_numpy(dtype=float) - np.nanmean(mat.to_numpy(dtype=float), axis=0))
        / (np.nanstd(mat.to_numpy(dtype=float), axis=0) + 1e-12),
        columns=cols,
        index=df.index,
    )
    mat.index = df["cluster_name"] + " | " + df["module_id"]
    fig, ax = plt.subplots(figsize=(8.4, 8.2))
    sns.heatmap(mat, cmap="vlag", center=0, linewidths=0.25, linecolor="white", cbar_kws={"label": "z-score"}, ax=ax)
    ax.set_xlabel("")
    ax.set_ylabel("")
    ax.set_title("Top joint reset-geometry/accessibility DMR candidates", fontweight="bold")
    save(fig, out_dir, "Fig11_joint_priority_DMR_heatmap")


def fig_12_claim_boundary_matrix(data: dict[str, pd.DataFrame], out_dir: Path) -> None:
    rows = [
        ("methylation-only failure", 1, 1, 0, 0),
        ("module residual correction", 1, 1, 0, 0),
        ("closure/access architecture", 1, 1, 1, 0),
        ("morula accessibility top25", 1, 0, 1, 0),
        ("entry-exit duality", 1, 1, 0, 0),
        ("negative-curvature accessibility", 1, 1, 1, 0),
        ("CBP/p300-HDAC perturbability", 0, 0, 1, 0),
        ("paired perturbation methylation", 0, 0, 0, 0),
    ]
    df = pd.DataFrame(rows, columns=["Evidence layer", "quantified", "geometry", "chromatin", "causal"])
    mat = df.set_index("Evidence layer")
    fig, ax = plt.subplots(figsize=(7.2, 5.0))
    cmap = matplotlib.colors.ListedColormap(["#eef2f7", "#2f6fb0"])
    sns.heatmap(mat, cmap=cmap, cbar=False, linewidths=0.8, linecolor="white", annot=mat.replace({1: "yes", 0: ""}), fmt="", ax=ax)
    ax.set_title("Evidence boundary: what is supported now", fontweight="bold")
    ax.set_xlabel("")
    ax.set_ylabel("")
    save(fig, out_dir, "Fig12_evidence_boundary_matrix")


def fig_13_final_integrated_model(out_dir: Path) -> None:
    fig, ax = plt.subplots(figsize=(11, 6))
    ax.axis("off")
    nodes = {
        "8-cell": (0.12, 0.72, PALETTE["gray"]),
        "morula\nreset-basin vertex": (0.50, 0.72, PALETTE["blue"]),
        "blastocyst": (0.88, 0.72, PALETTE["gray"]),
        "diagnostic\ncorrection": (0.50, 0.40, PALETTE["purple"]),
        "closure branch\nM05/M01/M12": (0.28, 0.18, PALETTE["red"]),
        "access branch\nM02/M10": (0.72, 0.18, PALETTE["teal"]),
        "public morula\naccessibility": (0.84, 0.42, PALETTE["gold"]),
    }
    for label, (x, y, color) in nodes.items():
        ax.add_patch(Rectangle((x - 0.095, y - 0.055), 0.19, 0.11, facecolor=color, alpha=0.13, edgecolor=color, lw=1.5))
        ax.text(x, y, label, ha="center", va="center", fontsize=10, fontweight="bold", color=PALETTE["ink"])
    arrows = [
        ("8-cell", "morula\nreset-basin vertex", "entry"),
        ("morula\nreset-basin vertex", "blastocyst", "exit"),
        ("morula\nreset-basin vertex", "diagnostic\ncorrection", "failure term"),
        ("diagnostic\ncorrection", "closure branch\nM05/M01/M12", ""),
        ("diagnostic\ncorrection", "access branch\nM02/M10", ""),
        ("public morula\naccessibility", "morula\nreset-basin vertex", "chromatin support"),
        ("public morula\naccessibility", "access branch\nM02/M10", "bounded"),
    ]
    for src, dst, text in arrows:
        x1, y1, _ = nodes[src]
        x2, y2, _ = nodes[dst]
        ax.add_patch(FancyArrowPatch((x1, y1), (x2, y2), arrowstyle="-|>", mutation_scale=14, lw=1.4, color=PALETTE["ink"], alpha=0.78))
        if text:
            ax.text((x1 + x2) / 2, (y1 + y2) / 2 + 0.035, text, ha="center", fontsize=8.5, color=PALETTE["muted"])
    ax.text(0.5, 0.93, "Perturbation-informed chromatin-associated diagnostic reset-basin geometry", ha="center", fontsize=14, fontweight="bold")
    ax.text(0.5, 0.04, "Causal u_bio remains unresolved without paired perturbation-to-methylation readout", ha="center", fontsize=9.5, color=PALETTE["red"])
    save(fig, out_dir, "Fig13_integrated_reset_basin_model")


def fig_14_top_candidate_lollipop(data: dict[str, pd.DataFrame], out_dir: Path) -> None:
    df = data["joint_priority"].head(20).copy().sort_values("joint_reset_geometry_access_score")
    fig, ax = plt.subplots(figsize=(8.0, 6.6))
    labels = df["cluster_name"] + " (" + df["module_id"] + ")"
    colors = [BRANCH_COLORS.get(b, PALETTE["gray"]) for b in df["branch"]]
    ax.hlines(labels, 0, df["joint_reset_geometry_access_score"], color=colors, lw=3, alpha=0.78)
    ax.scatter(df["joint_reset_geometry_access_score"], labels, color=colors, s=70, edgecolor="white", linewidth=0.6)
    ax.set_xlabel("Joint reset-geometry/accessibility score")
    ax.set_ylabel("")
    ax.set_title("Prioritized DMRs integrating residual, geometry, and accessibility", fontweight="bold")
    save(fig, out_dir, "Fig14_joint_candidate_lollipop")


def write_manifest(out_dir: Path, result_dir: Path, n_figures: int) -> None:
    manifest = {
        "package": "CSB_TRO latest dynamics figure package",
        "source_result_dir": str(result_dir),
        "n_new_figure_stems": n_figures,
        "formats": ["png", "svg", "pdf"],
        "generated_files": sorted(p.name for p in out_dir.glob("*")),
    }
    (out_dir / "figure_manifest.json").write_text(json.dumps(manifest, indent=2, ensure_ascii=False), encoding="utf-8")


def main() -> None:
    parser = argparse.ArgumentParser()
    parser.add_argument("--result-dir", type=Path, default=Path(r"E:\实验进展5_27"))
    parser.add_argument("--out-dir", type=Path, required=True)
    args = parser.parse_args()

    setup_style()
    data = load_all(args.result_dir)
    figures = [
        fig_01_model_roadmap,
        fig_02_operator_failure,
        fig_03_entry_exit_scatter,
        fig_04_module_duality,
        fig_05_random_control_duality,
        fig_06_curvature_distribution,
        fig_07_public_rescue_topk,
        fig_08_coupling_curvature,
        fig_09_residual_geometry_intersection,
        fig_10_module_triad_bubble,
        fig_11_joint_priority_heatmap,
        fig_12_claim_boundary_matrix,
        fig_13_final_integrated_model,
        fig_14_top_candidate_lollipop,
    ]
    for fn in figures:
        if fn in {fig_02_operator_failure, fig_13_final_integrated_model}:
            fn(args.out_dir)
        else:
            fn(data, args.out_dir)
    write_manifest(args.out_dir, args.result_dir, len(figures))


if __name__ == "__main__":
    main()
```
