## Supplemental code for "An entropy-based diagnostic framework for characterizing methylation state dynamics during preimplantation development": second_source_gse101571_atac_overlap.py

```
from __future__ import annotations

import argparse
import gzip
from pathlib import Path

import numpy as np
import pandas as pd


def read_tsv(path: Path) -> pd.DataFrame:
    return pd.read_csv(path, sep="\t")


def read_bed(path: Path) -> pd.DataFrame:
    opener = gzip.open if path.suffix == ".gz" else open
    rows = []
    with opener(path, "rt", encoding="utf-8", errors="replace") as handle:
        for line in handle:
            if not line.strip() or line.startswith(("#", "track", "browser")):
                continue
            parts = line.rstrip("\n").split("\t")
            if len(parts) < 3:
                continue
            try:
                rows.append((str(parts[0]), int(float(parts[1])), int(float(parts[2]))))
            except ValueError:
                continue
    return pd.DataFrame(rows, columns=["chr", "start", "end"]).drop_duplicates()


def add_coordinates(root: Path, dmr: pd.DataFrame) -> pd.DataFrame:
    if {"chr", "start", "end"}.issubset(dmr.columns):
        return dmr
    meta = read_tsv(root / "results" / "CSB_TRO_DMR_metadata.tsv")
    key = next((c for c in ["cluster_name", "dmr_id", "DMR_id"] if c in dmr.columns and c in meta.columns), None)
    if key is None:
        raise ValueError("Could not join DMR ranking to coordinates.")
    return dmr.merge(meta, on=key, how="left", suffixes=("", "_meta"))


def overlap_intervals(query: pd.DataFrame, peaks: pd.DataFrame) -> np.ndarray:
    out = np.zeros(len(query), dtype=bool)
    peak_by_chr = {
        chrom: sub[["start", "end"]].to_numpy(dtype=int)
        for chrom, sub in peaks.groupby("chr", sort=False)
    }
    for i, row in enumerate(query.itertuples(index=False)):
        arr = peak_by_chr.get(row.chr)
        if arr is None:
            continue
        out[i] = bool(np.any((arr[:, 0] < row.end) & (arr[:, 1] > row.start)))
    return out


def matched_random(dmr: pd.DataFrame, top: pd.DataFrame, metric: str, rng: np.random.Generator, n_iter: int) -> list[float]:
    vals = []
    top_ids = set(top["cluster_name"]) if "cluster_name" in top else set(top.index)
    non_top = dmr[~dmr["cluster_name"].isin(top_ids)] if "cluster_name" in dmr else dmr.drop(top.index)
    non_top_metric = non_top[metric].astype(float).to_numpy()
    group_pools = []
    for _ in range(n_iter):
        if not group_pools:
            for _, group in top.groupby(["module_id", "width_bin", "cpg_bin"], dropna=False):
                pool_mask = (
                    (non_top["module_id"].isin(group["module_id"].unique()))
                    & (non_top["width_bin"].isin(group["width_bin"].unique()))
                    & (non_top["cpg_bin"].isin(group["cpg_bin"].unique()))
                ).to_numpy()
                pool_idx = np.flatnonzero(pool_mask)
                if len(pool_idx) < len(group):
                    pool_idx = np.arange(len(non_top), dtype=int)
                if len(pool_idx):
                    group_pools.append((pool_idx, len(group)))
        parts = []
        for pool_idx, group_n in group_pools:
            replace = len(pool_idx) < group_n
            chosen = rng.choice(pool_idx, size=group_n, replace=replace)
            parts.append(non_top_metric[chosen])
        if parts:
            sample_vals = np.concatenate(parts)
            vals.append(float(np.nanmean(sample_vals)))
    return vals


def main() -> None:
    parser = argparse.ArgumentParser()
    parser.add_argument("--root", required=True, type=Path)
    parser.add_argument("--bed-dir", required=True, type=Path)
    parser.add_argument("--out", required=True, type=Path)
    parser.add_argument("--n-iter", type=int, default=1000)
    args = parser.parse_args()
    args.out.mkdir(parents=True, exist_ok=True)

    dmr = read_tsv(args.root / "results" / "CSB_TRO_basin_residual_DMR_ranking.tsv")
    dmr = add_coordinates(args.root, dmr)
    dmr = dmr.dropna(subset=["chr", "start", "end"]).copy()
    dmr["chr"] = dmr["chr"].astype(str)
    dmr["start"] = dmr["start"].astype(int)
    dmr["end"] = dmr["end"].astype(int)
    dmr["width"] = dmr["end"] - dmr["start"]
    rank_col = "abs_latent_residual_delta_beta"
    if rank_col in dmr.columns:
        dmr = dmr.sort_values(rank_col, ascending=False).reset_index(drop=True)

    dmr["width_bin"] = pd.qcut(dmr["width"].rank(method="first"), q=5, labels=False, duplicates="drop")
    if "n_cpg_target" in dmr:
        dmr["cpg_bin"] = pd.qcut(dmr["n_cpg_target"].rank(method="first"), q=4, labels=False, duplicates="drop")
    else:
        dmr["cpg_bin"] = 0

    beds = sorted(args.bed_dir.glob("GSE101571_*_peaks.bed.gz"))
    source_rows = []
    for bed in beds:
        label = bed.name.replace("GSE101571_", "").replace("_peaks.bed.gz", "")
        peaks = read_bed(bed)
        metric = f"overlap_GSE101571_{label}"
        dmr[metric] = overlap_intervals(dmr[["chr", "start", "end"]], peaks)
        source_rows.append({"source": label, "file": str(bed), "n_peaks": int(len(peaks))})

    summary_rows = []
    random_rows = []
    rng = np.random.default_rng(20260527)
    for metric in [c for c in dmr.columns if c.startswith("overlap_GSE101571_")]:
        source = metric.replace("overlap_GSE101571_", "")
        for k in [10, 25, 50, 100]:
            top = dmr.head(k)
            rest = dmr.iloc[k:]
            obs = float(top[metric].mean())
            vals = matched_random(dmr, top, metric, rng, args.n_iter)
            arr = np.array(vals, dtype=float)
            random_median = float(np.nanmedian(arr)) if len(arr) else np.nan
            random_q95 = float(np.nanquantile(arr, 0.95)) if len(arr) else np.nan
            random_max = float(np.nanmax(arr)) if len(arr) else np.nan
            summary_rows.append({
                "analysis": "GSE101571_independent_ATAC_peak_overlap",
                "source": source,
                "stage_matched_to_morula": False,
                "top_k": k,
                "metric": "peak_overlap_fraction",
                "observed_mean": obs,
                "background_mean": float(rest[metric].mean()) if len(rest) else np.nan,
                "random_median": random_median,
                "random_q95": random_q95,
                "random_max": random_max,
                "observed_gt_random_q95": bool(obs > random_q95) if not pd.isna(random_q95) else False,
                "n_random": int(len(arr)),
            })
            for i, val in enumerate(vals):
                random_rows.append({"source": source, "top_k": k, "iteration": i, "random_mean": val})

    dmr.to_csv(args.out / "CSB_TRO_2026-05-27_GSE101571_ATAC_DMR_overlap.tsv", sep="\t", index=False)
    pd.DataFrame(source_rows).to_csv(args.out / "CSB_TRO_2026-05-27_GSE101571_ATAC_source_summary.tsv", sep="\t", index=False)
    pd.DataFrame(summary_rows).to_csv(args.out / "CSB_TRO_2026-05-27_GSE101571_ATAC_overlap_summary.tsv", sep="\t", index=False)
    pd.DataFrame(random_rows).to_csv(args.out / "CSB_TRO_2026-05-27_GSE101571_ATAC_matched_random.tsv", sep="\t", index=False)

    hits = pd.DataFrame(summary_rows)
    positive = hits[hits["observed_gt_random_q95"]]
    lines = [
        "# GSE101571 independent ATAC overlap control",
        "",
        "This is an independent public accessibility boundary control using human 8-cell and ICM ATAC peak BED files from GSE101571.",
        "",
        "It is not a stage-matched morula rescue because GSE101571 does not provide human morula ATAC peaks in the downloaded GEO supplementary files.",
        "",
        "## Result",
        "",
    ]
    if len(positive):
        lines.append("At least one top-k/source comparison exceeded matched-random q95:")
        for row in positive.itertuples(index=False):
            lines.append(
                f"- {row.source} top{row.top_k}: observed={row.observed_mean:.3f}, "
                f"random_median={row.random_median:.3f}, q95={row.random_q95:.3f}, max={row.random_max:.3f}"
            )
    else:
        lines.append("No top-k/source comparison exceeded matched-random q95.")
    lines.extend([
        "",
        "## Boundary",
        "",
        "A positive result here supports independent accessibility association with residual DMRs, but it cannot replicate the Liu2019 morula-stage signal.",
        "A negative result does not refute the Liu2019 morula signal because the stages and assays are different.",
    ])
    (args.out / "CSB_TRO_2026-05-27_GSE101571_ATAC_overlap_interpretation.md").write_text("\n".join(lines) + "\n", encoding="utf-8")


if __name__ == "__main__":
    main()
```
