## Supplemental code for "An entropy-based diagnostic framework for characterizing methylation state dynamics during preimplantation development": summarize_u_bio_rescue_modules.py

```
from __future__ import annotations

import argparse
from pathlib import Path

import numpy as np
import pandas as pd


BRANCH = {
    "M05": "closure",
    "M01": "closure",
    "M12": "closure",
    "M02": "access",
    "M10": "access",
}


def main() -> None:
    parser = argparse.ArgumentParser()
    parser.add_argument("--out", required=True, type=Path)
    args = parser.parse_args()

    overlap_path = args.out / "CSB_TRO_2026-05-27_u_bio_rescue_DMR_overlap.tsv"
    summary_path = args.out / "CSB_TRO_2026-05-27_u_bio_rescue_overlap_summary.tsv"
    evidence_path = args.out / "CSB_TRO_2026-05-27_evidence_boundary_table.tsv"

    dmr = pd.read_csv(overlap_path, sep="\t")
    summary = pd.read_csv(summary_path, sep="\t")

    dmr["branch"] = dmr["module_id"].map(BRANCH).fillna("other")
    rows = []
    for k in [10, 25, 50, 100]:
        top = dmr.head(k).copy()
        for group_col in ["module_id", "branch"]:
            grouped = top.groupby(group_col, dropna=False)
            for group, sub in grouped:
                rows.append({
                    "top_k": k,
                    "group_type": group_col,
                    "group": group,
                    "n_dmr": int(len(sub)),
                    "overlap_fraction": float(sub["overlap_public_chromatin"].mean()),
                    "morula_accessibility_mean": float(sub["public_accessibility_morula_max"].mean(skipna=True)),
                    "morula_minus_8cell_mean": float(sub["public_accessibility_morula_minus_8cell"].mean(skipna=True)),
                    "mean_abs_residual": float(sub["abs_latent_residual_delta_beta"].mean()) if "abs_latent_residual_delta_beta" in sub else np.nan,
                })
    pd.DataFrame(rows).to_csv(args.out / "CSB_TRO_2026-05-27_u_bio_rescue_module_branch_summary.tsv", sep="\t", index=False)

    hit = summary[
        (summary["top_k"] == 25)
        & (summary["metric"] == "public_accessibility_morula_max")
    ].iloc[0]
    top25 = dmr.head(25)
    top25[[
        "cluster_name",
        "module_id",
        "branch",
        "chr",
        "start",
        "end",
        "abs_latent_residual_delta_beta",
        "overlap_public_chromatin",
        "public_accessibility_morula_max",
        "public_accessibility_morula_minus_8cell",
    ]].to_csv(args.out / "CSB_TRO_2026-05-27_u_bio_rescue_top25_DMRs.tsv", sep="\t", index=False)

    if evidence_path.exists():
        evidence = pd.read_csv(evidence_path, sep="\t")
        evidence = evidence[evidence["evidence_layer"] != "human_morula_accessibility_rescue"].copy()
        row = {
            "evidence_layer": "human_morula_accessibility_rescue",
            "result": (
                f"top25 morula accessibility={hit['observed_mean']:.3f}; "
                f"matched random median={hit['random_median']:.3f}; "
                f"q95={hit['random_q95']:.3f}; max={hit['random_max']:.3f}"
            ),
            "interpretation": "Stage-matched public human morula accessibility partially supports the top residual DMR correction signal.",
            "claim_strength": "stage-matched partial support",
            "boundary": "Top25 accessibility-level signal only; overlap fraction and morula-minus-8cell delta are not specific, and this is not causal u_bio.",
            "source": "CSB_TRO_2026-05-27_u_bio_rescue_overlap_summary.tsv",
        }
        pd.concat([evidence, pd.DataFrame([row])], ignore_index=True).to_csv(evidence_path, sep="\t", index=False)

    lines = [
        "# u_bio rescue module/branch interpretation",
        "",
        "## Main rescued signal",
        "",
        (
            f"Top25 residual DMRs show higher human morula accessibility than matched random controls "
            f"(observed={hit['observed_mean']:.3f}, random median={hit['random_median']:.3f}, "
            f"q95={hit['random_q95']:.3f}, max={hit['random_max']:.3f})."
        ),
        "",
        "This upgrades the evidence from purely diagnostic plausibility to stage-matched chromatin partial support for the most extreme residual DMRs.",
        "",
        "## Boundary",
        "",
        "The signal is not a complete u_bio replacement: it is strongest at top25, overlap fraction is not specific, morula-minus-8cell accessibility delta does not exceed random q95, and no perturbation-to-methylation readout is available.",
        "",
        "## Output files",
        "",
        "- CSB_TRO_2026-05-27_u_bio_rescue_module_branch_summary.tsv",
        "- CSB_TRO_2026-05-27_u_bio_rescue_top25_DMRs.tsv",
        "- CSB_TRO_2026-05-27_u_bio_rescue_morula_accessibility_top25.svg/png",
    ]
    (args.out / "CSB_TRO_2026-05-27_u_bio_rescue_module_branch_interpretation.md").write_text("\n".join(lines) + "\n", encoding="utf-8")


if __name__ == "__main__":
    main()
```
