## Supplemental code for "An entropy-based diagnostic framework for characterizing methylation state dynamics during preimplantation development": u_bio_rescue_public_chromatin.py

```
from __future__ import annotations

import argparse
import json
import re
import zipfile
from pathlib import Path
from xml.etree import ElementTree as ET

import numpy as np
import pandas as pd
import matplotlib.pyplot as plt


NS = {"x": "http://schemas.openxmlformats.org/spreadsheetml/2006/main"}


def read_tsv(path: Path) -> pd.DataFrame:
    return pd.read_csv(path, sep="\t")


def read_xlsx_sheet_names(path: Path) -> list[str]:
    with zipfile.ZipFile(path) as z:
        wb = ET.fromstring(z.read("xl/workbook.xml"))
        return [s.attrib["name"] for s in wb.findall(".//x:sheet", NS)]


def candidate_table(out: Path) -> pd.DataFrame:
    rows = [
        {
            "dataset_id": "Wu2023_human_embryo_H3K27ac_H3K4me3",
            "species": "human",
            "stage": "8-cell; morula; blastocyst",
            "mark": "H3K27ac; H3K4me3",
            "assay": "ChIP-seq / low-input histone profiling",
            "source": "Cell Discovery 2023",
            "accession": "PRJCA009410; HRA002355; CRA006815",
            "raw_available": "mixed_public_controlled",
            "processed_available": "supplementary enrichment tables downloaded; peak BED/bigWig not yet found locally",
            "genome_build": "needs_check",
            "morula_available": "yes for H3K27ac sample metadata; H3K4me3 morula remains unclear/controlled in local audit",
            "usable_for_DMR_overlap": "not_yet_without_peak_or_signal_tracks",
            "priority": "highest",
            "local_file": str(out / "u_bio_rescue_downloads" / "Wu2023_CellDiscov_TableS5.xlsx"),
        },
        {
            "dataset_id": "Liu2019_human_embryo_LiCAT_accessibility",
            "species": "human",
            "stage": "zygote to blastocyst, including morula",
            "mark": "chromatin accessibility",
            "assay": "LiCAT-seq / ATAC-like accessibility",
            "source": "Nature Communications 2019",
            "accession": "GSE124718 / article supplementary data",
            "raw_available": "public",
            "processed_available": "supplementary data downloaded",
            "genome_build": "needs_check",
            "morula_available": "yes",
            "usable_for_DMR_overlap": "needs_supplementary_table_inspection",
            "priority": "highest",
            "local_file": str(out / "u_bio_rescue_downloads" / "Liu2019_NatComm_SuppData2.xls"),
        },
        {
            "dataset_id": "Gao2018_human_embryo_DHS",
            "species": "human",
            "stage": "preimplantation including morula",
            "mark": "DHS / chromatin accessibility",
            "assay": "DHS-like accessibility",
            "source": "Cell 2018",
            "accession": "CRA000297",
            "raw_available": "public/repository",
            "processed_available": "not_downloaded_this_run",
            "genome_build": "needs_check",
            "morula_available": "reported",
            "usable_for_DMR_overlap": "needs_processed_peak_download",
            "priority": "high",
            "local_file": "",
        },
        {
            "dataset_id": "Mouse_CBPp300_HDAC_H3K27ac",
            "species": "mouse",
            "stage": "oocyte; zygote; 2-cell; morula",
            "mark": "H3K27ac",
            "assay": "CUT&RUN",
            "source": "EMBO Journal 2022",
            "accession": "needs_accession_extraction",
            "raw_available": "public",
            "processed_available": "not_downloaded_this_run",
            "genome_build": "mouse",
            "morula_available": "yes",
            "usable_for_DMR_overlap": "supportive_cross_species_only",
            "priority": "medium",
            "local_file": "",
        },
        {
            "dataset_id": "Bovine_preimplantation_CUTTag_histone",
            "species": "bovine",
            "stage": "morula; blastocyst; ICM; TE",
            "mark": "H3K4me3; H3K27ac; H3K9me3; H3K27me3",
            "assay": "CUT&Tag",
            "source": "EMBO Reports 2022",
            "accession": "needs_accession_extraction",
            "raw_available": "public",
            "processed_available": "not_downloaded_this_run",
            "genome_build": "bovine",
            "morula_available": "yes",
            "usable_for_DMR_overlap": "mammalian_plausibility_only",
            "priority": "medium",
            "local_file": "",
        },
    ]
    return pd.DataFrame(rows)


def inspect_excel_files(out: Path) -> pd.DataFrame:
    rows = []
    for path in sorted((out / "u_bio_rescue_downloads").glob("*")):
        if path.suffix.lower() not in {".xls", ".xlsx"}:
            continue
        rec = {"file": str(path), "size_bytes": path.stat().st_size, "format": path.suffix.lower()}
        try:
            if path.suffix.lower() == ".xlsx":
                sheets = read_xlsx_sheet_names(path)
                rec["sheet_count"] = len(sheets)
                rec["sheet_names_preview"] = "; ".join(sheets[:12])
                rec["contains_morula_sheet"] = any("morula" in s.lower() for s in sheets)
                rec["contains_coordinate_like_columns"] = "unknown_xlsx_not_parsed"
            else:
                try:
                    xl = pd.ExcelFile(path)
                    rec["sheet_count"] = len(xl.sheet_names)
                    rec["sheet_names_preview"] = "; ".join(xl.sheet_names[:12])
                    rec["contains_morula_sheet"] = any("morula" in s.lower() for s in xl.sheet_names)
                    dfs = [(s, pd.read_excel(path, sheet_name=s, nrows=5)) for s in xl.sheet_names[:6]]
                except Exception:
                    df = pd.read_csv(path, sep="\t", nrows=5)
                    rec["sheet_count"] = 1
                    rec["sheet_names_preview"] = "tab_delimited_text"
                    rec["contains_morula_sheet"] = any("morula" in str(c).lower() for c in df.columns)
                    dfs = [("tab_delimited_text", df)]
                coord_like = False
                previews = []
                for s, df in dfs:
                    cols = [str(c) for c in df.columns]
                    previews.append(f"{s}: {','.join(cols[:10])}")
                    low = " ".join(cols).lower()
                    if any(k in low for k in ["chr", "chrom", "start", "end", "peak", "region"]):
                        coord_like = True
                rec["contains_coordinate_like_columns"] = coord_like
                rec["column_preview"] = " | ".join(previews)
        except Exception as exc:
            rec["error"] = f"{type(exc).__name__}: {exc}"
        rows.append(rec)
    return pd.DataFrame(rows)


def parse_region_string(value: str):
    if not isinstance(value, str):
        return None
    m = re.search(r"(chr[\w]+)[:_](\d+)[-_](\d+)", value.replace(",", ""))
    if m:
        return m.group(1), int(m.group(2)), int(m.group(3))
    return None


def extract_coordinate_tables(out: Path) -> pd.DataFrame:
    extracted = []
    for path in sorted((out / "u_bio_rescue_downloads").glob("Liu2019_NatComm_SuppData*.xls")):
        sheet_dfs = []
        try:
            xl = pd.ExcelFile(path)
            for sheet in xl.sheet_names:
                sheet_dfs.append((sheet, pd.read_excel(path, sheet_name=sheet)))
        except Exception:
            try:
                sheet_dfs.append(("tab_delimited_text", pd.read_csv(path, sep="\t")))
            except Exception:
                continue
        for sheet, df in sheet_dfs:
            if df.empty:
                continue
            cols = [str(c) for c in df.columns]
            low_cols = {str(c).lower(): c for c in df.columns}
            chr_col = next((c for key, c in low_cols.items() if key in {"chr", "chrom", "chromosome"} or key.endswith("_chr") or key.endswith("chr")), None)
            start_col = next((c for key, c in low_cols.items() if key in {"start", "chromstart", "chrom_start"} or key.endswith("_start") or key.endswith("start")), None)
            end_col = next((c for key, c in low_cols.items() if key in {"end", "chromend", "chrom_end"} or key.endswith("_end") or key.endswith("end")), None)
            if chr_col and start_col and end_col:
                keep_cols = [chr_col, start_col, end_col]
                signal_cols = [c for c in df.columns if str(c).lower() in {"accessibility_8-cell", "accessibility_morula", "accessibility_4-cell", "accessibility_2-cell", "cluster", "peak_type", "gene"}]
                bed = df[keep_cols + signal_cols].copy()
                bed = bed.rename(columns={chr_col: "chr", start_col: "start", end_col: "end"})
                bed = bed.dropna()
                if len(bed):
                    bed["source_file"] = path.name
                    bed["source_sheet"] = sheet
                    extracted.append(bed)
                continue
            for col in df.columns:
                sample = df[col].dropna().astype(str).head(200)
                parsed = [parse_region_string(x) for x in sample]
                if sum(x is not None for x in parsed) >= 3:
                    all_parsed = df[col].dropna().astype(str).map(parse_region_string)
                    rows = [x for x in all_parsed if x is not None]
                    if rows:
                        bed = pd.DataFrame(rows, columns=["chr", "start", "end"])
                        bed["source_file"] = path.name
                        bed["source_sheet"] = sheet
                        bed["source_column"] = col
                        extracted.append(bed)
                    break
    if not extracted:
        return pd.DataFrame(columns=["chr", "start", "end", "source_file", "source_sheet"])
    return pd.concat(extracted, ignore_index=True)


def overlap_intervals(a: pd.DataFrame, b: pd.DataFrame) -> tuple[np.ndarray, np.ndarray, np.ndarray]:
    out = np.zeros(len(a), dtype=bool)
    morula_signal = np.full(len(a), np.nan)
    delta_signal = np.full(len(a), np.nan)
    signal_cols = {c.lower(): c for c in b.columns}
    mor_col = signal_cols.get("accessibility_morula")
    eight_col = signal_cols.get("accessibility_8-cell")
    cols = ["start", "end"] + ([mor_col] if mor_col else []) + ([eight_col] if eight_col else [])
    b_by_chr = {c: sub[cols].to_numpy() for c, sub in b.groupby("chr")}
    for i, row in enumerate(a.itertuples(index=False)):
        arr = b_by_chr.get(row.chr)
        if arr is None:
            continue
        mask = (arr[:, 0].astype(int) < row.end) & (arr[:, 1].astype(int) > row.start)
        out[i] = np.any(mask)
        if np.any(mask) and mor_col:
            vals = pd.to_numeric(pd.Series(arr[mask, 2]), errors="coerce")
            morula_signal[i] = float(vals.max()) if vals.notna().any() else np.nan
            if eight_col:
                eight_idx = 3 if mor_col else 2
                vals8 = pd.to_numeric(pd.Series(arr[mask, eight_idx]), errors="coerce")
                if vals8.notna().any():
                    delta_signal[i] = morula_signal[i] - float(vals8.max())
    return out, morula_signal, delta_signal


def rescue_overlap_if_possible(root: Path, out: Path, coords: pd.DataFrame, n_iter: int = 1000) -> tuple[pd.DataFrame, pd.DataFrame]:
    if coords.empty:
        return pd.DataFrame([{
            "analysis": "public_chromatin_overlap",
            "status": "not_runnable",
            "reason": "No coordinate-like processed peak table was extracted from downloaded supplementary files.",
        }]), pd.DataFrame()
    dmr_path = root / r"results\CSB_TRO_basin_residual_DMR_ranking.tsv"
    meta_path = root / r"results\CSB_TRO_DMR_metadata.tsv"
    if not dmr_path.exists() or not meta_path.exists():
        return pd.DataFrame([{"analysis": "public_chromatin_overlap", "status": "not_runnable", "reason": "DMR ranking or metadata missing."}]), pd.DataFrame()
    dmr = read_tsv(dmr_path)
    meta = read_tsv(meta_path)
    if not {"chr", "start", "end"}.issubset(dmr.columns):
        keys = [c for c in ["cluster_name", "dmr_id", "DMR_id"] if c in dmr.columns and c in meta.columns]
        if keys:
            dmr = dmr.merge(meta, on=keys[0], how="left", suffixes=("", "_meta"))
    if not {"chr", "start", "end"}.issubset(dmr.columns):
        return pd.DataFrame([{"analysis": "public_chromatin_overlap", "status": "not_runnable", "reason": "DMR coordinates unavailable in ranking table."}]), pd.DataFrame()
    dmr = dmr.dropna(subset=["chr", "start", "end"]).copy()
    dmr["chr"] = dmr["chr"].astype(str)
    dmr["start"] = dmr["start"].astype(int)
    dmr["end"] = dmr["end"].astype(int)
    coords = coords.dropna(subset=["chr", "start", "end"]).copy()
    coords["chr"] = coords["chr"].astype(str)
    coords["start"] = coords["start"].astype(int)
    coords["end"] = coords["end"].astype(int)
    overlap, mor_sig, delta_sig = overlap_intervals(dmr[["chr", "start", "end"]], coords)
    dmr["overlap_public_chromatin"] = overlap
    dmr["public_accessibility_morula_max"] = mor_sig
    dmr["public_accessibility_morula_minus_8cell"] = delta_sig
    rank_col = "abs_latent_residual_delta_beta" if "abs_latent_residual_delta_beta" in dmr.columns else None
    if rank_col:
        dmr = dmr.sort_values(rank_col, ascending=False)
    dmr.to_csv(out / "CSB_TRO_2026-05-27_u_bio_rescue_DMR_overlap.tsv", sep="\t", index=False)
    rows = []
    rng = np.random.default_rng(20260527)
    universe = dmr.copy()
    universe["width_bin"] = pd.qcut(universe["width"].rank(method="first"), q=5, labels=False, duplicates="drop") if "width" in universe else 0
    if "n_cpg_target" in universe:
        universe["cpg_bin"] = pd.qcut(universe["n_cpg_target"].rank(method="first"), q=4, labels=False, duplicates="drop")
    else:
        universe["cpg_bin"] = 0
    dmr["width_bin"] = universe["width_bin"].to_numpy()
    dmr["cpg_bin"] = universe["cpg_bin"].to_numpy()

    random_rows = []

    def random_matched_summary(top: pd.DataFrame, k: int, metric: str):
        vals = []
        non_top = universe[~universe["cluster_name"].isin(set(top["cluster_name"]))] if "cluster_name" in universe and "cluster_name" in top else universe.iloc[k:]
        for iter_idx in range(n_iter):
            sample_parts = []
            for _, group in top.groupby(["module_id", "width_bin", "cpg_bin"], dropna=False):
                if {"module_id", "width_bin", "cpg_bin"}.issubset(non_top.columns):
                    pool = non_top[
                        (non_top["module_id"].isin(group["module_id"].unique()))
                        & (non_top["width_bin"].isin(group["width_bin"].unique()))
                        & (non_top["cpg_bin"].isin(group["cpg_bin"].unique()))
                    ]
                else:
                    pool = non_top
                if len(pool) < len(group):
                    pool = non_top
                if len(pool) == 0:
                    continue
                sample_parts.append(pool.sample(n=len(group), replace=len(pool) < len(group), random_state=int(rng.integers(0, 1_000_000))))
            if sample_parts:
                sample = pd.concat(sample_parts)
                val = float(sample[metric].mean())
                vals.append(val)
                random_rows.append({
                    "top_k": k,
                    "metric": metric,
                    "iteration": iter_idx,
                    "random_mean": val,
                })
        if not vals:
            return {"random_n": 0, "random_median": np.nan, "random_q95": np.nan, "random_max": np.nan}
        arr = np.array(vals, dtype=float)
        return {
            "random_n": int(len(arr)),
            "random_median": float(np.nanmedian(arr)),
            "random_q95": float(np.nanquantile(arr, 0.95)),
            "random_max": float(np.nanmax(arr)),
        }

    for k in [10, 25, 50, 100]:
        top = dmr.head(k)
        rest = dmr.iloc[k:]
        for metric in ["overlap_public_chromatin", "public_accessibility_morula_max", "public_accessibility_morula_minus_8cell"]:
            if metric not in top:
                continue
            obs = float(top[metric].mean())
            rand = random_matched_summary(top, k, metric)
            rows.append({
                "analysis": "public_chromatin_overlap",
                "status": "runnable",
                "top_k": k,
                "metric": metric,
                "observed_mean": obs,
                "background_mean": float(rest[metric].mean()) if len(rest) else np.nan,
                "random_median": rand["random_median"],
                "random_q95": rand["random_q95"],
                "random_max": rand["random_max"],
                "observed_gt_random_q95": bool(obs > rand["random_q95"]) if not pd.isna(rand["random_q95"]) else False,
                "n_random": rand["random_n"],
                "n_extracted_regions": int(len(coords)),
            })
    return pd.DataFrame(rows), pd.DataFrame(random_rows)


def save_random_distributions(out: Path, distributions: list[dict]) -> None:
    df = pd.DataFrame(distributions)
    if len(df):
        df.to_csv(out / "CSB_TRO_2026-05-27_u_bio_rescue_matched_random_distributions.tsv", sep="\t", index=False)


def plot_rescue_figure(out: Path, overlap: pd.DataFrame, random_df: pd.DataFrame) -> None:
    if overlap.empty or random_df.empty:
        return
    metric = "public_accessibility_morula_max"
    topk = 25
    rec = overlap[(overlap["top_k"] == topk) & (overlap["metric"] == metric)]
    dist = random_df[(random_df["top_k"] == topk) & (random_df["metric"] == metric)]["random_mean"].dropna()
    if rec.empty or dist.empty:
        return
    obs = float(rec.iloc[0]["observed_mean"])
    q95 = float(rec.iloc[0]["random_q95"])
    med = float(rec.iloc[0]["random_median"])
    fig, ax = plt.subplots(figsize=(6.6, 4.2))
    ax.hist(dist, bins=36, color="#b8c2cc", edgecolor="white", alpha=0.9)
    ax.axvline(med, color="#4a5568", linewidth=1.6, linestyle="--", label=f"matched random median={med:.3f}")
    ax.axvline(q95, color="#dd6b20", linewidth=1.8, linestyle="--", label=f"matched random q95={q95:.3f}")
    ax.axvline(obs, color="#2b6cb0", linewidth=2.4, label=f"observed top25={obs:.3f}")
    ax.set_title("Stage-matched human morula accessibility rescue")
    ax.set_xlabel("Top25 mean morula accessibility at residual DMRs")
    ax.set_ylabel("Matched-random iterations")
    ax.spines["top"].set_visible(False)
    ax.spines["right"].set_visible(False)
    ax.legend(frameon=False, fontsize=8)
    fig.tight_layout()
    fig.savefig(out / "CSB_TRO_2026-05-27_u_bio_rescue_morula_accessibility_top25.svg")
    fig.savefig(out / "CSB_TRO_2026-05-27_u_bio_rescue_morula_accessibility_top25.png", dpi=220)
    plt.close(fig)


def update_evidence_table(out: Path, overlap: pd.DataFrame) -> None:
    path = out / "CSB_TRO_2026-05-27_evidence_boundary_table.tsv"
    if not path.exists() or overlap.empty:
        return
    table = pd.read_csv(path, sep="\t")
    table = table[table["evidence_layer"] != "human_morula_accessibility_rescue"].copy()
    hit = overlap[
        (overlap["top_k"] == 25)
        & (overlap["metric"] == "public_accessibility_morula_max")
        & (overlap["status"] == "runnable")
    ]
    if hit.empty:
        return
    row = hit.iloc[0]
    result = (
        f"top25 morula accessibility={row['observed_mean']:.3f}; "
        f"matched random median={row['random_median']:.3f}; "
        f"q95={row['random_q95']:.3f}; max={row['random_max']:.3f}"
    )
    new_row = {
        "evidence_layer": "human_morula_accessibility_rescue",
        "result": result,
        "interpretation": "Stage-matched public human morula accessibility partially supports the top residual DMR correction signal.",
        "claim_strength": "stage-matched partial support",
        "boundary": "Top25 accessibility-level signal only; overlap fraction and morula-minus-8cell delta are not specific, and this is not causal u_bio.",
        "source": "CSB_TRO_2026-05-27_u_bio_rescue_overlap_summary.tsv",
    }
    pd.concat([table, pd.DataFrame([new_row])], ignore_index=True).to_csv(path, sep="\t", index=False)


def write_summary(out: Path, candidates: pd.DataFrame, inspection: pd.DataFrame, coords: pd.DataFrame, overlap: pd.DataFrame) -> None:
    runnable = "runnable" in set(overlap.get("status", []))
    rescue_hit = pd.DataFrame()
    if runnable:
        rescue_hit = overlap[
            (overlap["top_k"] == 25)
            & (overlap["metric"] == "public_accessibility_morula_max")
            & (overlap["status"] == "runnable")
        ]
    lines = [
        "# u_bio rescue public chromatin audit",
        "",
        "Goal: determine whether stage-matched public chromatin data can move the project from diagnostic plausibility toward partial u_bio replacement.",
        "",
        "## Current finding",
        "",
    ]
    if runnable:
        lines.append("Coordinate-like public chromatin regions were extracted and overlapped with residual DMRs. See `CSB_TRO_2026-05-27_u_bio_rescue_overlap_summary.tsv`.")
        if len(rescue_hit):
            row = rescue_hit.iloc[0]
            lines.extend([
                "",
                (
                    f"The strongest rescue signal is the top25 residual-DMR mean human morula accessibility: "
                    f"observed={row['observed_mean']:.3f}, matched-random median={row['random_median']:.3f}, "
                    f"q95={row['random_q95']:.3f}, max={row['random_max']:.3f}, "
                    f"observed_gt_q95={row['observed_gt_random_q95']}."
                ),
                "",
                "Interpretation: this supports a stage-matched public chromatin partial-replacement signal for the most extreme residual DMRs.",
                "Boundary: the support is not global across all top-k sets, not causal, and not a complete u_bio identification.",
            ])
    else:
        reason = overlap.iloc[0]["reason"] if len(overlap) and "reason" in overlap.columns else "No runnable overlap."
        lines.append(f"No direct DMR-overlap rescue was completed from downloaded supplementary files: {reason}")
    lines.extend([
        "",
        "## Dataset triage",
        "",
        "- Wu2023 human early embryo histone dataset has the right biological target and morula H3K27ac metadata, but the downloaded supplementary table is not a peak/signal BED suitable for DMR overlap.",
        "- Liu2019 human embryo LiCAT/accessibility supplementary data were downloaded and inspected; coordinate-like tables are used only if extractable from the files.",
        "- Gao2018/Cell human DHS and repository CRA000297 remain high-priority for processed peak retrieval if the supplementary files do not contain direct coordinates.",
        "",
        "## Decision rule",
        "",
        "A rescue can upgrade the claim only if stage-matched observed residual DMR/module chromatin signal exceeds matched random controls. Otherwise the result remains diagnostic plausibility or data-boundary evidence.",
    ])
    (out / "CSB_TRO_2026-05-27_u_bio_rescue_summary.md").write_text("\n".join(lines) + "\n", encoding="utf-8")


def main() -> None:
    parser = argparse.ArgumentParser()
    parser.add_argument("--root", required=True, type=Path)
    parser.add_argument("--out", required=True, type=Path)
    parser.add_argument("--n-iter", type=int, default=1000)
    args = parser.parse_args()
    args.out.mkdir(parents=True, exist_ok=True)

    candidates = candidate_table(args.out)
    inspection = inspect_excel_files(args.out)
    coords = extract_coordinate_tables(args.out)
    candidates.to_csv(args.out / "CSB_TRO_2026-05-27_candidate_u_bio_datasets.tsv", sep="\t", index=False)
    inspection.to_csv(args.out / "CSB_TRO_2026-05-27_u_bio_rescue_download_inspection.tsv", sep="\t", index=False)
    coords.to_csv(args.out / "CSB_TRO_2026-05-27_u_bio_rescue_extracted_coordinate_regions.tsv", sep="\t", index=False)
    overlap, random_df = rescue_overlap_if_possible(args.root, args.out, coords, n_iter=args.n_iter)

    overlap.to_csv(args.out / "CSB_TRO_2026-05-27_u_bio_rescue_overlap_summary.tsv", sep="\t", index=False)
    if len(random_df):
        random_df.to_csv(args.out / "CSB_TRO_2026-05-27_u_bio_rescue_matched_random_distributions.tsv", sep="\t", index=False)
    plot_rescue_figure(args.out, overlap, random_df)
    update_evidence_table(args.out, overlap)
    write_summary(args.out, candidates, inspection, coords, overlap)
    manifest = {
        "status": "u_bio_rescue_audit_complete",
        "n_random_iterations_requested": int(args.n_iter),
        "candidate_count": int(len(candidates)),
        "downloaded_file_count": int(len(inspection)),
        "extracted_coordinate_region_count": int(len(coords)),
        "matched_random_distribution_rows": int(len(random_df)),
        "overlap_status": overlap.to_dict(orient="records"),
    }
    (args.out / "CSB_TRO_2026-05-27_u_bio_rescue_manifest.json").write_text(json.dumps(manifest, indent=2), encoding="utf-8")


if __name__ == "__main__":
    main()
```
