## Supplemental code for "An entropy-based diagnostic framework for characterizing methylation state dynamics during preimplantation development": assemble_final_figure.py

```
#!/usr/bin/env python
"""
Final publication figure — fully corrected.
Row 1: Vector field panels computed in Python (correction-vector space)
        showing each control operator's contribution at 4 tau snapshots.
Row 2: Trajectory panels in correction-vector space.
"""
import numpy as np
import pandas as pd
import matplotlib
matplotlib.use("Agg")
import matplotlib.pyplot as plt
import matplotlib.patheffects as pe
from matplotlib.patches import Circle, FancyArrowPatch
from matplotlib.colors import LinearSegmentedColormap
from matplotlib.lines import Line2D
from matplotlib.gridspec import GridSpec
from scipy.stats import gaussian_kde
from pathlib import Path
import warnings, json
warnings.filterwarnings("ignore")

OUT = Path("E:/progress_comsol_analysis")
FIG = OUT / "figures_final"
FIG.mkdir(exist_ok=True)

# ── Style ─────────────────────────────────────────────────────────────────────
plt.rcParams.update({
    "font.family":"DejaVu Sans","font.size":9,
    "axes.labelsize":10,"axes.titlesize":10,"axes.titleweight":"bold",
    "axes.linewidth":0.8,"axes.spines.top":False,"axes.spines.right":False,
    "xtick.labelsize":8,"ytick.labelsize":8,
    "xtick.major.size":3,"ytick.major.size":3,
    "xtick.major.width":0.7,"ytick.major.width":0.7,
    "legend.fontsize":7.5,"figure.dpi":300,
    "savefig.dpi":300,"savefig.bbox":"tight","savefig.facecolor":"white",
})

# ── Palettes ──────────────────────────────────────────────────────────────────
STAGE_COLORS = {
    "oocyte":"#F4A261","zygote":"#E76F51","2cell":"#A8DADC",
    "4cell":"#457B9D","8cell":"#1D3557","morula":"#C0392B","blast":"#27AE60",
}
STAGE_LABELS = {
    "oocyte":"Oocyte","zygote":"Zygote","2cell":"2-cell",
    "4cell":"4-cell","8cell":"8-cell","morula":"Morula","blast":"Blast.",
}
STAGE_ORDER = ["oocyte","zygote","2cell","4cell","8cell","morula","blast"]

SCEN_COLORS = {
    "baseline_only":"#7F8C8D","plus_zga":"#2980B9",
    "plus_entry":"#E67E22","full_control":"#C0392B","wrong_exit":"#8E44AD",
}
SCEN_LABELS = {
    "baseline_only":"Methylation-only","plus_zga":"+ZGA",
    "plus_entry":"+Entry","full_control":"Full control","wrong_exit":"Wrong exit",
}

# ── Load data ─────────────────────────────────────────────────────────────────
samples = pd.read_csv(OUT/"sample_trajectories_corrected.csv")
centers = pd.read_csv(OUT/"stage_centers_2d_corrected.csv").set_index("stage")
trajs = {}
for name in ["baseline_only","plus_zga","plus_entry","full_control","wrong_exit"]:
    fp = OUT/f"traj_final_{name}.csv"
    if fp.exists():
        trajs[name] = pd.read_csv(fp)
with open(OUT/"scenario_results_final.json") as f:
    results = json.load(f)

C = {s: centers.loc[s].values for s in centers.index}
r_morula, r_blast, r_8cell = 0.50, 0.60, 0.80

XLIM = (-0.5, 5.8)
YLIM = (-2.15, 2.2)

# ── CEEF vector field in correction-vector space ──────────────────────────────
params = dict(
    z1_oocyte=C["oocyte"][0], z2_oocyte=C["oocyte"][1],
    z1_4cell =C["4cell"][0],  z2_4cell =C["4cell"][1],
    z1_8cell =C["8cell"][0],  z2_8cell =C["8cell"][1],
    z1_morula=C["morula"][0], z2_morula=C["morula"][1],
    z1_blast =C["blast"][0],  z2_blast =C["blast"][1],
    z1_Katt=0.59, z2_Katt=0.04,
    lambda_K=0.4389,
    sigma_M=0.2132, sigma_8=0.7415, sigma_4=0.30,
    gamma_acc=1.0, gamma_closure=1.0,
    gamma_zga=1.0, gamma_re=1.0, gamma_de=1.0,
)
p = params

def smooth_gate(tau, t0, t1, w=0.3):
    """Smooth window function active between t0 and t1."""
    def flc2hs(x, scale):
        # COMSOL flc2hs approximation
        t = x / scale
        t = np.clip(t, -1, 1)
        return 0.5 + t*(0.75 - 0.25*t*t)
    return flc2hs(tau-t0, w) * flc2hs(t1-tau, w)

def compute_field(z1, z2, tau):
    """Compute CEEF vector field at grid points (z1,z2) for given tau."""
    # Gaussian envelopes
    rM2 = (z1-p["z1_morula"])**2 + (z2-p["z2_morula"])**2
    r82 = (z1-p["z1_8cell"])**2  + (z2-p["z2_8cell"])**2
    r42 = (z1-p["z1_4cell"])**2  + (z2-p["z2_4cell"])**2
    gM = np.exp(-rM2/(2*p["sigma_M"]**2))
    g8 = np.exp(-r82/(2*p["sigma_8"]**2))
    g4 = np.exp(-r42/(2*p["sigma_4"]**2))

    # Time gates
    chi_zga   = smooth_gate(tau, 3, 4)
    chi_entry = smooth_gate(tau, 4, 5)
    chi_exit  = smooth_gate(tau, 5, 6)

    # F_K: linear attraction to K-attractor
    FK1 = -p["lambda_K"] * (z1 - p["z1_Katt"])
    FK2 = -p["lambda_K"] * (z2 - p["z2_Katt"])

    # Unit direction vectors
    d_zga   = np.sqrt((p["z1_8cell"]-p["z1_4cell"])**2 + (p["z2_8cell"]-p["z2_4cell"])**2)
    vzga1   = (p["z1_8cell"]-p["z1_4cell"])/d_zga
    vzga2   = (p["z2_8cell"]-p["z2_4cell"])/d_zga

    d_entry = np.sqrt((p["z1_morula"]-p["z1_8cell"])**2 + (p["z2_morula"]-p["z2_8cell"])**2)
    ventry1 = (p["z1_morula"]-p["z1_8cell"])/d_entry
    ventry2 = (p["z2_morula"]-p["z2_8cell"])/d_entry

    d_exit  = np.sqrt((p["z1_blast"]-p["z1_morula"])**2 + (p["z2_blast"]-p["z2_morula"])**2)
    vexit1  = (p["z1_blast"]-p["z1_morula"])/d_exit
    vexit2  = (p["z2_blast"]-p["z2_morula"])/d_exit

    # Component fields
    Fzga1   = chi_zga   * p["gamma_zga"] * g4 * vzga1
    Fzga2   = chi_zga   * p["gamma_zga"] * g4 * vzga2
    Fentry1 = chi_entry * (p["gamma_acc"]+p["gamma_closure"]) * g8 * ventry1
    Fentry2 = chi_entry * (p["gamma_acc"]+p["gamma_closure"]) * g8 * ventry2
    Fexit1  = chi_exit  * (p["gamma_re"]+p["gamma_de"]) * gM * vexit1
    Fexit2  = chi_exit  * (p["gamma_re"]+p["gamma_de"]) * gM * vexit2

    Fu1 = FK1 + Fzga1 + Fentry1 + Fexit1
    Fu2 = FK2 + Fzga2 + Fentry2 + Fexit2
    return Fu1, Fu2, FK1, FK2, Fzga1, Fzga2, Fentry1, Fentry2, Fexit1, Fexit2

# ── Helpers ───────────────────────────────────────────────────────────────────
def kde_bg(ax, alpha=0.28, bw=0.20):
    xmin,xmax = XLIM; ymin,ymax = YLIM
    xx,yy = np.mgrid[xmin:xmax:80j, ymin:ymax:80j]
    pos = np.vstack([xx.ravel(), yy.ravel()])
    stage_map = {"MII oocyte":"oocyte","zygote/PN":"zygote","2-cell":"2cell",
                 "4-cell":"4cell","8-cell":"8cell","morula":"morula","blastocyst":"blast"}
    for s_long, s_short in stage_map.items():
        sub = samples[samples["stage"]==s_long][["z1","z2"]].values
        if len(sub) < 4: continue
        try:
            kde = gaussian_kde(sub.T, bw_method=bw)
            z = kde(pos).reshape(xx.shape); z /= z.max()
            r,g,b = [int(STAGE_COLORS[s_short][1+2*i:3+2*i],16)/255 for i in range(3)]
            cmap = LinearSegmentedColormap.from_list("_",[(1,1,1,0),(r,g,b,alpha)])
            ax.contourf(xx,yy,z,levels=[0.15,0.4,0.7,1.0],cmap=cmap,zorder=1)
        except: pass

def stage_pts(ax, ms=80, fs=8):
    # Precise offsets to avoid overlap in the crowded e1∈[0.2,1.0] region:
    # oocyte(0.97,0.24), zygote(0.47,0.37), 2cell(0.62,0.23),
    # 4cell(0.57,0.56), 8cell(0.22,-0.15), morula(3.63,-0.28), blast(0.61,1.54)
    label_offsets = {
        "oocyte": ( 6,  5),    # right of point
        "zygote": (-48,  4),   # left (avoids oocyte/2cell)
        "2cell":  ( 6, -12),   # below right (avoids oocyte above)
        "4cell":  (-46,  4),   # left (avoids 2cell/oocyte)
        "8cell":  (-44, -12),  # left-below
        "morula": ( 6,   4),   # right (isolated)
        "blast":  ( 6,   4),   # right (isolated)
    }
    for s in STAGE_ORDER:
        c = C[s]
        ax.scatter(c[0],c[1],s=ms,color=STAGE_COLORS[s],
                   edgecolors="white",linewidths=1.4,zorder=6,marker="D")
    for s in STAGE_ORDER:
        c = C[s]; dx,dy = label_offsets[s]
        ax.annotate(STAGE_LABELS[s],(c[0],c[1]),
                    textcoords="offset points",xytext=(dx,dy),
                    fontsize=fs,fontweight="bold",color=STAGE_COLORS[s],
                    path_effects=[pe.withStroke(linewidth=2.5,foreground="white")],
                    zorder=10)

def basin_circle(ax, cx, cy, r, color, alpha=0.10):
    for ri,ai in [(r,alpha),(r*0.55,alpha*0.7)]:
        ax.add_patch(Circle((cx,cy),ri,facecolor=color,alpha=ai,
                            edgecolor=color,linewidth=1.2,linestyle="--",zorder=1))

def grad_traj(ax, x, y, color, lw=2.3, a0=0.2, a1=0.95):
    n = len(x)
    for i in range(n-1):
        a = a0+(a1-a0)*i/(n-1)
        ax.plot(x[i:i+2],y[i:i+2],color=color,lw=lw,alpha=a,
                solid_capstyle="round",zorder=5)

def arrow_mid(ax, x, y, color, frac=0.60, sz=11):
    idx = min(int(len(x)*frac), len(x)-2)
    dx,dy = x[idx+1]-x[idx], y[idx+1]-y[idx]
    ax.annotate("",xy=(x[idx]+dx*2,y[idx]+dy*2),xytext=(x[idx],y[idx]),
                arrowprops=dict(arrowstyle="-|>",color=color,lw=1.5,mutation_scale=sz),zorder=8)

def clean_ax(ax, xl="e1 (entry correction)", yl="e2 (exit correction)"):
    ax.set_xlim(*XLIM); ax.set_ylim(*YLIM)
    ax.set_xlabel(xl,fontsize=9,labelpad=2)
    ax.set_ylabel(yl,fontsize=9,labelpad=2)
    ax.grid(True,alpha=0.15,linewidth=0.5,zorder=0)

def plabel(ax, lbl, x=-0.13, y=1.05):
    ax.text(x,y,lbl,transform=ax.transAxes,fontsize=13,fontweight="bold",va="top",ha="left")

# ── Build grid for field plots ─────────────────────────────────────────────────
NX, NY = 60, 45
x1d = np.linspace(XLIM[0], XLIM[1], NX)
y1d = np.linspace(YLIM[0], YLIM[1], NY)
XX, YY = np.meshgrid(x1d, y1d)

# ══════════════════════════════════════════════════════════════════════════════
# FIGURE: 2-row × 4-col
# ══════════════════════════════════════════════════════════════════════════════
fig = plt.figure(figsize=(22, 12))
gs  = GridSpec(2, 4, figure=fig,
               hspace=0.42, wspace=0.32,
               left=0.05, right=0.98, top=0.93, bottom=0.07)

fig.text(0.5, 0.975,
         "CEEF Phase A: Epigenetic Vector Field Dynamics & Trajectory Control",
         ha="center",va="top",fontsize=14,fontweight="bold",color="#1a1a2e")

# ── Row 1: Vector field panels ────────────────────────────────────────────────
# Color = active operator magnitude (shows WHERE each operator acts)
# Streamlines = TOTAL field (curved, shows actual trajectory direction)
# Trajectory = corresponding scenario overlaid
field_panels = [
    (1.5, "A", "τ=1.5  Methylation-only (F_K)\nBaseline: attraction to K-attractor",
     "YlOrRd", None,    "baseline_only"),
    (3.5, "B", "τ=3.5  +ZGA reconstruction\nF_ZGA: 4-cell → 8-cell push",
     "YlGn",   "zga",   "plus_zga"),
    (4.5, "C", "τ=4.5  +Entry control\nF_entry: 8-cell → Morula push",
     "YlOrBr", "entry", "plus_entry"),
    (5.5, "D", "τ=5.5  Full control (F_exit)\nF_exit: Morula → Blast push",
     "PuRd",   "exit",  "full_control"),
]

for col, (tau, plbl, title, cmap_name, active_op, scen_name) in enumerate(field_panels):
    ax = fig.add_subplot(gs[0, col])

    Fu1, Fu2, FK1, FK2, Fzga1, Fzga2, Fentry1, Fentry2, Fexit1, Fexit2 = \
        compute_field(XX, YY, tau)

    # Pseudocolor: active operator magnitude
    # For FK-only panel: show total field magnitude (hot spot = K-attractor)
    # For control panels: show active operator magnitude (hot spot = where operator acts)
    if active_op == "zga":
        Fcolor = np.sqrt(Fzga1**2 + Fzga2**2)
    elif active_op == "entry":
        Fcolor = np.sqrt(Fentry1**2 + Fentry2**2)
    elif active_op == "exit":
        Fcolor = np.sqrt(Fexit1**2 + Fexit2**2)
    else:
        # FK panel: invert distance to K-attractor so hot spot IS the attractor
        dist_K = np.sqrt((XX - p["z1_Katt"])**2 + (YY - p["z2_Katt"])**2)
        Fcolor = np.exp(-dist_K**2 / (2 * 0.8**2))  # Gaussian centered on K-attractor

    # Streamlines: TOTAL field (curved, biologically meaningful)
    # Add small FK background so streamlines are visible everywhere
    Ftot1 = Fu1 + FK1 * 0.3
    Ftot2 = Fu2 + FK2 * 0.3
    Fspeed = np.sqrt(Ftot1**2 + Ftot2**2) + 1e-9

    im = ax.pcolormesh(XX, YY, Fcolor, cmap=cmap_name,
                       vmin=0, vmax=max(Fcolor.max()*0.85, 0.01),
                       shading="gouraud", zorder=0, rasterized=True)
    plt.colorbar(im, ax=ax, fraction=0.035, pad=0.02, label="|F|")

    # Streamlines colored by total field speed
    ax.streamplot(x1d, y1d, Ftot1, Ftot2,
                  color="white",
                  linewidth=1.2,
                  density=1.4, arrowsize=1.0, arrowstyle="->",
                  zorder=3)

    # Overlay the corresponding trajectory (up to this tau)
    if scen_name in trajs:
        df = trajs[scen_name]
        mask = df["t"] <= tau + 0.05
        x_t, y_t = df.loc[mask, "z1"].values, df.loc[mask, "z2"].values
        if len(x_t) > 1:
            # Gradient trajectory
            n = len(x_t)
            for i in range(n-1):
                a = 0.4 + 0.6*i/(n-1)
                ax.plot(x_t[i:i+2], y_t[i:i+2],
                        color=SCEN_COLORS[scen_name], lw=2.0, alpha=a,
                        solid_capstyle="round", zorder=6)
            # Endpoint star
            ax.scatter(x_t[-1], y_t[-1], s=80,
                       color=SCEN_COLORS[scen_name],
                       edgecolors="white", linewidths=1.2,
                       marker="*", zorder=8)

    # Stage centers
    row1_offsets = {
        "oocyte": ( 5,  4), "zygote": (-44,  4),
        "2cell":  ( 5,-11), "4cell":  (-42,  4),
        "8cell":  (-42,-11),"morula": ( 5,   4), "blast": ( 5, 4),
    }
    for s in STAGE_ORDER:
        c = C[s]
        ax.scatter(c[0], c[1], s=60, color=STAGE_COLORS[s],
                   edgecolors="white", linewidths=1.1, zorder=7, marker="D")
    for s in STAGE_ORDER:
        c = C[s]; dx, dy = row1_offsets[s]
        ax.annotate(STAGE_LABELS[s], (c[0], c[1]),
                    textcoords="offset points", xytext=(dx, dy),
                    fontsize=7, fontweight="bold", color="white",
                    path_effects=[pe.withStroke(linewidth=2, foreground="black")],
                    zorder=10)

    # Basin circles
    for cx, cy, r, color in [
        (C["morula"][0], C["morula"][1], r_morula, "#FF6B6B"),
        (C["blast"][0],  C["blast"][1],  r_blast,  "#69FF69"),
    ]:
        ax.add_patch(Circle((cx, cy), r, fill=False,
                            edgecolor=color, linewidth=1.5, linestyle="--", zorder=4))

    ax.set_xlim(*XLIM); ax.set_ylim(*YLIM)
    ax.set_xlabel("e1",fontsize=8,labelpad=1)
    ax.set_ylabel("e2",fontsize=8,labelpad=1)
    ax.set_title(title,fontsize=9,fontweight="bold",pad=5,color="#1a1a2e")
    ax.text(-0.04,1.04,plbl,transform=ax.transAxes,
            fontsize=13,fontweight="bold",va="top",ha="left")

# ── Row 2, Panel E: All trajectories ─────────────────────────────────────────
ax = fig.add_subplot(gs[1,0])
kde_bg(ax)
basin_circle(ax,C["morula"][0],C["morula"][1],r_morula,STAGE_COLORS["morula"])
basin_circle(ax,C["blast"][0], C["blast"][1], r_blast, STAGE_COLORS["blast"])
basin_circle(ax,C["8cell"][0], C["8cell"][1], r_8cell, STAGE_COLORS["8cell"])
for name,df in trajs.items():
    x,y = df["z1"].values, df["z2"].values
    grad_traj(ax,x,y,SCEN_COLORS[name],lw=2.0)
    arrow_mid(ax,x,y,SCEN_COLORS[name])
    ax.scatter(x[-1],y[-1],s=45,color=SCEN_COLORS[name],
               edgecolors="white",linewidths=0.8,zorder=9)
stage_pts(ax,ms=70,fs=7.5)
handles = [Line2D([0],[0],color=SCEN_COLORS[n],lw=2,label=SCEN_LABELS[n])
           for n in SCEN_COLORS]
ax.legend(handles=handles,loc="upper right",fontsize=6.5,
          framealpha=0.92,edgecolor="#ccc",ncol=1)
clean_ax(ax)
ax.set_title("All Scenario Trajectories",fontsize=10,fontweight="bold",pad=5)
plabel(ax,"E")

# ── Row 2, Panel F: Failure → rescue ─────────────────────────────────────────
ax = fig.add_subplot(gs[1,1])
kde_bg(ax,alpha=0.18)
basin_circle(ax,C["morula"][0],C["morula"][1],r_morula,STAGE_COLORS["morula"])
basin_circle(ax,C["blast"][0], C["blast"][1], r_blast, STAGE_COLORS["blast"])
for name in ["baseline_only","plus_entry","full_control"]:
    if name in trajs:
        df = trajs[name]
        x,y = df["z1"].values, df["z2"].values
        grad_traj(ax,x,y,SCEN_COLORS[name],lw=2.5)
        arrow_mid(ax,x,y,SCEN_COLORS[name])
        ax.scatter(x[-1],y[-1],s=80,color=SCEN_COLORS[name],
                   marker="*",edgecolors="white",linewidths=0.8,zorder=9)
stage_pts(ax,ms=60,fs=7)
for i,(name,lbl) in enumerate([("baseline_only","Baseline: [-][-]"),
                                ("plus_entry",   "+Entry:   [+][-]"),
                                ("full_control", "Full:     [+][+]")]):
    res = results.get(name,{})
    in_m,in_b = res.get("in_morula",False),res.get("in_blast",False)
    col = STAGE_COLORS["blast"] if (in_m and in_b) else \
          STAGE_COLORS["morula"] if in_m else "#E74C3C"
    ax.text(0.03,0.97-i*0.09,lbl,transform=ax.transAxes,fontsize=7.5,
            va="top",color=col,fontweight="bold",
            bbox=dict(boxstyle="round,pad=0.2",facecolor=col,alpha=0.10,
                      edgecolor=col,linewidth=0.8))
clean_ax(ax)
ax.set_title("Failure → Rescue Progression",fontsize=10,fontweight="bold",pad=5)
plabel(ax,"F")

# ── Row 2, Panel G: Morula gate ───────────────────────────────────────────────
ax = fig.add_subplot(gs[1,2])
kde_bg(ax,alpha=0.20)
basin_circle(ax,C["morula"][0],C["morula"][1],r_morula*2.2,
             STAGE_COLORS["morula"],alpha=0.05)
basin_circle(ax,C["morula"][0],C["morula"][1],r_morula,
             STAGE_COLORS["morula"],alpha=0.12)
ax.annotate("",xy=C["morula"],xytext=C["8cell"],
            arrowprops=dict(arrowstyle="-|>",color="#E74C3C",lw=2.8,
                            mutation_scale=20,connectionstyle="arc3,rad=0.08"))
# Entry label: BELOW the arrow (e2=-0.60), clear of all stage labels above
ax.text(1.5, -0.60, "Entry\n(8-cell→Morula)",
        color="#E74C3C", fontsize=8, fontweight="bold", ha="center",
        path_effects=[pe.withStroke(linewidth=2.5, foreground="white")])
ax.annotate("",xy=C["blast"],xytext=C["morula"],
            arrowprops=dict(arrowstyle="-|>",color="#27AE60",lw=2.8,
                            mutation_scale=20,connectionstyle="arc3,rad=-0.1"))
# Exit label: upper right, well clear of entry label
ax.text(3.5, 0.95, "Exit\n(Morula→Blast)",
        color="#27AE60", fontsize=8, fontweight="bold", ha="center",
        path_effects=[pe.withStroke(linewidth=2.5, foreground="white")])
if "full_control" in trajs:
    df = trajs["full_control"]
    grad_traj(ax,df["z1"].values,df["z2"].values,
              SCEN_COLORS["full_control"],lw=1.8,a0=0.15,a1=0.75)
stage_pts(ax,ms=70,fs=7.5)
ax.text(0.97,0.04,"cos(entry,exit) = −0.876\nMorula = turning gate",
        transform=ax.transAxes,fontsize=8,va="bottom",ha="right",
        bbox=dict(boxstyle="round,pad=0.4",facecolor="#FFEAA7",
                  edgecolor="#FDCB6E",linewidth=1,alpha=0.95))
clean_ax(ax)
ax.set_title("Morula as Epigenetic Turning Gate",fontsize=10,fontweight="bold",pad=5)
plabel(ax,"G")

# ── Row 2, Panel H: Basin capture bar chart ───────────────────────────────────
ax = fig.add_subplot(gs[1,3])
scenarios4 = ["baseline_only","plus_zga","plus_entry","full_control","wrong_exit"]
labels4    = ["Baseline","+ZGA","+Entry","Full\nCtrl","Wrong\nExit"]
morula_v = [1 if results.get(n,{}).get("in_morula",False) else 0 for n in scenarios4]
blast_v  = [1 if results.get(n,{}).get("in_blast",False)  else 0 for n in scenarios4]
dist_v   = [results.get(n,{}).get("dist_morula_t5",5.0)   for n in scenarios4]
x = np.arange(len(scenarios4)); bw = 0.32
ax.bar(x-bw/2,morula_v,bw,
       color=[STAGE_COLORS["morula"] if v else "#F5B7B1" for v in morula_v],
       edgecolor="white",linewidth=0.8,zorder=3,label="Morula (τ=5)")
ax.bar(x+bw/2,blast_v,bw,
       color=[STAGE_COLORS["blast"] if v else "#A9DFBF" for v in blast_v],
       edgecolor="white",linewidth=0.8,zorder=3,label="Blast (τ=6)")
ax.set_xticks(x); ax.set_xticklabels(labels4,fontsize=8)
ax.set_ylabel("Basin capture  (1=Yes)",fontsize=9)
ax.set_ylim(0,1.5)
ax.legend(fontsize=7.5,loc="upper right")
ax.grid(True,axis="y",alpha=0.18,linewidth=0.5,zorder=0)
ax.spines["top"].set_visible(False); ax.spines["right"].set_visible(False)
for i,(d,m,b) in enumerate(zip(dist_v,morula_v,blast_v)):
    col = "#27AE60" if (m and b) else "#E67E22" if m else "#E74C3C"
    ax.text(i,max(m,b)+0.08,f"d={d:.2f}",ha="center",
            fontsize=7,color=col,fontweight="bold")
ax.text(0.5,0.02,"Wrong exit → trajectory collapse",
        transform=ax.transAxes,fontsize=7.5,ha="center",va="bottom",
        color="#7F8C8D",style="italic")
ax.set_title("Basin Capture Summary",fontsize=10,fontweight="bold",pad=5)
plabel(ax,"H")

# ── Save ──────────────────────────────────────────────────────────────────────
out_png = FIG/"CEEF_PhaseA_Final_Figure.png"
out_pdf = FIG/"CEEF_PhaseA_Final_Figure.pdf"
plt.savefig(out_png, dpi=300)
try:
    out_pdf.unlink(missing_ok=True)
    plt.savefig(out_pdf)
    print(f"Saved: {out_pdf}")
except Exception as e:
    print(f"PDF skipped: {e}")
plt.close()
print(f"Saved: {out_png}")
```
