## Supplemental code for "An entropy-based diagnostic framework for characterizing methylation state dynamics during preimplantation development": assemble_phase_b_figure.py

```
#!/usr/bin/env python
"""
Phase B final figure — 2-row × 4-col, aligned with Phase A.
Row 1: COMSOL chromatin-gated field panels (A-D)
Row 2: Python analysis panels (E-H)
  E: Entry-Exit scatter (DMR-level)
  F: Module duality lollipop
  G: Curvature distribution (KDE)
  H: Accessibility × duality coupling
"""
import numpy as np
import pandas as pd
import matplotlib
matplotlib.use("Agg")
import matplotlib.pyplot as plt
import matplotlib.patheffects as pe
from matplotlib.lines import Line2D
from matplotlib.gridspec import GridSpec
from scipy.stats import gaussian_kde
from pathlib import Path
from PIL import Image
import warnings, os
warnings.filterwarnings("ignore")

CFIG = Path("E:/progress_comsol_analysis/figures_comsol_b")
BASE = Path("E:/实验进展5_27/CSB_TRO_2026-05-27_FINAL_DYNAMICS_PACKAGE/results_all")
OUT  = Path("E:/progress_comsol_analysis/figures_final")
CORR_VEC = Path("E:/progress_comsol_analysis")

# ── Style — identical to Phase A ─────────────────────────────────────────────
plt.rcParams.update({
    "font.family":"DejaVu Sans","font.size":9,
    "axes.labelsize":10,"axes.titlesize":10,"axes.titleweight":"bold",
    "axes.linewidth":0.8,"axes.spines.top":False,"axes.spines.right":False,
    "xtick.labelsize":8,"ytick.labelsize":8,
    "xtick.major.size":3,"ytick.major.size":3,
    "xtick.major.width":0.7,"ytick.major.width":0.7,
    "legend.fontsize":7.5,"figure.dpi":300,
    "savefig.dpi":300,"savefig.bbox":"tight","savefig.facecolor":"white",
})

# ── Palettes — same as Phase A ────────────────────────────────────────────────
BRANCH_COLORS = {
    "access":  "#C0392B",
    "closure": "#2980B9",
    "other":   "#7F8C8D",
}
BRANCH_LABELS = {
    "access":  "Accessibility (M02, entry)",
    "closure": "Closure (M05, exit)",
    "other":   "Background modules",
}

# Stage colors — same as Phase A
STAGE_COLORS = {
    "oocyte":"#F4A261","zygote":"#E76F51","2cell":"#A8DADC",
    "4cell":"#457B9D","8cell":"#1D3557","morula":"#C0392B","blast":"#27AE60",
}
STAGE_LABELS = {
    "oocyte":"Oocyte","zygote":"Zygote","2cell":"2-cell",
    "4cell":"4-cell","8cell":"8-cell","morula":"Morula","blast":"Blast.",
}
STAGE_ORDER = ["oocyte","zygote","2cell","4cell","8cell","morula","blast"]

def plabel(ax, lbl, x=-0.13, y=1.05):
    ax.text(x, y, lbl, transform=ax.transAxes,
            fontsize=13, fontweight="bold", va="top", ha="left")

# ── Load Phase B data ─────────────────────────────────────────────────────────
df_dmr    = pd.read_csv(BASE/"CSB_TRO_2026-05-27_entry_exit_duality_metrics.tsv", sep="\t")
df_module = pd.read_csv(BASE/"CSB_TRO_2026-05-27_entry_exit_module_duality.tsv", sep="\t")
df_triad  = pd.read_csv(BASE/"CSB_TRO_2026-05-27_duality_accessibility_module_triad.tsv", sep="\t")
df_curv   = pd.read_csv(BASE/"CSB_TRO_2026-05-27_entry_exit_curvature.tsv", sep="\t")
df_corr   = pd.read_csv(BASE/"CSB_TRO_2026-05-27_duality_accessibility_correlations.tsv", sep="\t")

# Load correction-vector space samples (for KDE background in E-G)
samples = pd.read_csv(CORR_VEC/"sample_trajectories_corrected.csv")
centers = pd.read_csv(CORR_VEC/"stage_centers_2d_corrected.csv").set_index("stage")
C = {s: centers.loc[s].values for s in centers.index}
XLIM = (-0.5, 5.8); YLIM = (-2.15, 2.2)

def kde_bg(ax, alpha=0.22, bw=0.20):
    xmin,xmax = XLIM; ymin,ymax = YLIM
    xx,yy = np.mgrid[xmin:xmax:80j, ymin:ymax:80j]
    pos = np.vstack([xx.ravel(), yy.ravel()])
    stage_map = {"MII oocyte":"oocyte","zygote/PN":"zygote","2-cell":"2cell",
                 "4-cell":"4cell","8-cell":"8cell","morula":"morula","blastocyst":"blast"}
    for s_long, s_short in stage_map.items():
        sub = samples[samples["stage"]==s_long][["z1","z2"]].values
        if len(sub) < 4: continue
        try:
            kde = gaussian_kde(sub.T, bw_method=bw)
            z = kde(pos).reshape(xx.shape); z /= z.max()
            r,g,b = [int(STAGE_COLORS[s_short][1+2*i:3+2*i],16)/255 for i in range(3)]
            from matplotlib.colors import LinearSegmentedColormap
            cmap = LinearSegmentedColormap.from_list("_",[(1,1,1,0),(r,g,b,alpha)])
            ax.contourf(xx,yy,z,levels=[0.15,0.4,0.7,1.0],cmap=cmap,zorder=1)
        except: pass

def stage_pts(ax, ms=70, fs=7.5):
    offsets = {
        "oocyte":( 6, 4),"zygote":(-44, 4),"2cell":( 6,-11),
        "4cell":(-42, 4),"8cell":(-42,-11),"morula":( 6, 4),"blast":( 6, 4),
    }
    for s in STAGE_ORDER:
        c = C[s]
        ax.scatter(c[0],c[1],s=ms,color=STAGE_COLORS[s],
                   edgecolors="white",linewidths=1.2,zorder=6,marker="D")
    for s in STAGE_ORDER:
        c = C[s]; dx,dy = offsets[s]
        ax.annotate(STAGE_LABELS[s],(c[0],c[1]),
                    textcoords="offset points",xytext=(dx,dy),
                    fontsize=fs,fontweight="bold",color=STAGE_COLORS[s],
                    path_effects=[pe.withStroke(linewidth=2.5,foreground="white")],
                    zorder=10)

def clean_ax(ax, xl="e1 (entry correction)", yl="e2 (exit correction)"):
    ax.set_xlim(*XLIM); ax.set_ylim(*YLIM)
    ax.set_xlabel(xl,fontsize=9,labelpad=2)
    ax.set_ylabel(yl,fontsize=9,labelpad=2)
    ax.grid(True,alpha=0.15,linewidth=0.5,zorder=0)

# ══════════════════════════════════════════════════════════════════════════════
# FIGURE: 2-row × 4-col
# ══════════════════════════════════════════════════════════════════════════════
fig = plt.figure(figsize=(22, 12))
gs  = GridSpec(2, 4, figure=fig,
               hspace=0.40, wspace=0.32,
               left=0.05, right=0.98, top=0.93, bottom=0.07)

fig.text(0.5, 0.975,
         "CEEF Phase B: Chromatin-Gated Control Operators & DMR Module Duality",
         ha="center", va="top", fontsize=14, fontweight="bold", color="#1a1a2e")

# ── Row 1: COMSOL field panels ────────────────────────────────────────────────
comsol_files = [
    CFIG/"PhaseB_A_M02_access_entry.png",
    CFIG/"PhaseB_B_M05_closure_exit.png",
    CFIG/"PhaseB_C_full_chromatin_control.png",
    CFIG/"PhaseB_D_wrong_exit_collapse.png",
]
comsol_titles = [
    "A  M02 Access-Gated Entry\n(F_entry, γ_acc=1.16)",
    "B  M05 Closure-Gated Exit\n(F_exit, γ_clo=1.89)",
    "C  Full Chromatin Control\n(M02+M05 gated)",
    "D  Wrong-Exit Collapse\n(reversed F_exit)",
]
for col, (fpath, title) in enumerate(zip(comsol_files, comsol_titles)):
    ax = fig.add_subplot(gs[0, col])
    if fpath.exists():
        img = np.array(Image.open(fpath))
        ax.imshow(img, aspect="auto")
    else:
        ax.text(0.5, 0.5, "File not found", transform=ax.transAxes,
                ha="center", va="center", color="red")
    ax.set_title(title, fontsize=9, fontweight="bold", pad=5, color="#1a1a2e")
    ax.axis("off")

# ── Row 2, Panel E: Entry-Exit scatter ───────────────────────────────────────
ax = fig.add_subplot(gs[1, 0])
ax.scatter(df_dmr["entry_change"], df_dmr["exit_change"],
           s=18, color="#BDC3C7", alpha=0.4, zorder=2, linewidths=0)
for branch in ["access", "closure"]:
    sub = df_dmr[df_dmr["branch"]==branch]
    ax.scatter(sub["entry_change"], sub["exit_change"],
               s=40, color=BRANCH_COLORS[branch], alpha=0.85,
               zorder=4, linewidths=0.5, edgecolors="white",
               label=f"{BRANCH_LABELS[branch]} (n={len(sub)})")
t = np.linspace(-0.85, 0.85, 100)
ax.plot(t, -t, color="#E74C3C", lw=1.2, ls="--", alpha=0.6, label="cos=-1")
ax.axhline(0, color="#ccc", lw=0.6, zorder=0)
ax.axvline(0, color="#ccc", lw=0.6, zorder=0)
ax.text(0.97, 0.97, "cos(entry,exit) = -0.876\nMorula = turning gate",
        transform=ax.transAxes, fontsize=8, va="top", ha="right",
        bbox=dict(boxstyle="round,pad=0.4",facecolor="#FFEAA7",
                  edgecolor="#FDCB6E",linewidth=1,alpha=0.95))
ax.set_xlabel("Entry change  (Δβ: 8-cell→Morula)", fontsize=9)
ax.set_ylabel("Exit change  (Δβ: Morula→Blast)", fontsize=9)
ax.set_xlim(-0.9, 1.05); ax.set_ylim(-1.05, 1.05)
ax.legend(loc="lower right", fontsize=7, framealpha=0.9)
ax.grid(True, alpha=0.15, linewidth=0.5, zorder=0)
ax.set_title("DMR Entry-Exit Anti-Alignment", fontsize=10, fontweight="bold", pad=5)
plabel(ax, "E")

# ── Row 2, Panel F: Module duality lollipop ───────────────────────────────────
ax = fig.add_subplot(gs[1, 1])
df_mod_sorted = df_module.sort_values("entry_exit_cosine")
y_pos = np.arange(len(df_mod_sorted))
for i, (_, row) in enumerate(df_mod_sorted.iterrows()):
    color = BRANCH_COLORS.get(row["branch"], "#7F8C8D")
    ax.plot([0, row["entry_exit_cosine"]], [i, i],
            color=color, lw=1.5, alpha=0.8, zorder=2)
    size = max(30, row["n_dmr"] * 4)
    ax.scatter(row["entry_exit_cosine"], i, s=size, color=color,
               zorder=4, edgecolors="white", linewidths=0.8)
    if row["is_priority_residual_module"]:
        ax.text(row["entry_exit_cosine"]-0.02, i, row["group"],
                ha="right", va="center", fontsize=7.5, fontweight="bold",
                color=color,
                path_effects=[pe.withStroke(linewidth=2,foreground="white")])
ax.axvline(0, color="#ccc", lw=0.8, zorder=0)
ax.axvline(-0.876, color="#E74C3C", lw=1.2, ls=":", alpha=0.7)
ax.set_yticks(y_pos)
ax.set_yticklabels(df_mod_sorted["group"].tolist(), fontsize=7.5)
ax.set_xlabel("Entry-Exit cosine similarity", fontsize=9)
ax.set_xlim(-1.05, 0.15)
ax.grid(True, axis="x", alpha=0.15, linewidth=0.5, zorder=0)
handles = [Line2D([0],[0],color=BRANCH_COLORS[b],lw=2,label=BRANCH_LABELS[b])
           for b in ["access","closure","other"]]
handles.append(Line2D([0],[0],color="#E74C3C",lw=1.2,ls=":",label="Overall cos=-0.876"))
ax.legend(handles=handles, loc="lower right", fontsize=6.5, framealpha=0.9)
ax.set_title("Module Entry-Exit Duality\n(dot size = n DMRs, bold = priority)",
             fontsize=10, fontweight="bold", pad=5)
plabel(ax, "F")

# ── Row 2, Panel G: Curvature distribution ────────────────────────────────────
ax = fig.add_subplot(gs[1, 2])
for branch in ["access", "closure", "other"]:
    sub = df_curv[df_curv["branch"]==branch]["curvature"]
    color = BRANCH_COLORS[branch]
    if len(sub) > 3:
        kde = gaussian_kde(sub, bw_method=0.4)
        x_range = np.linspace(df_curv["curvature"].min()-0.05,
                               df_curv["curvature"].max()+0.05, 200)
        y_kde = kde(x_range)
        ax.fill_between(x_range, y_kde, alpha=0.25, color=color)
        ax.plot(x_range, y_kde, color=color, lw=1.8,
                label=f"{BRANCH_LABELS[branch]} (n={len(sub)})")
ax.axvline(0, color="#555", lw=1.0, ls="--", alpha=0.7)
for branch in ["access", "closure"]:
    sub = df_curv[df_curv["branch"]==branch]
    frac_u = sub["is_u_shape"].mean()
    color = BRANCH_COLORS[branch]
    ax.text(0.97, 0.97-(0.12*["access","closure"].index(branch)),
            f"{branch}: {frac_u:.0%} U-shape",
            transform=ax.transAxes, fontsize=8, va="top", ha="right",
            color=color, fontweight="bold",
            path_effects=[pe.withStroke(linewidth=2,foreground="white")])
ax.set_xlabel("Curvature  (exit − entry change)", fontsize=9)
ax.set_ylabel("Density", fontsize=9)
ax.legend(loc="upper left", fontsize=7, framealpha=0.9)
ax.grid(True, alpha=0.15, linewidth=0.5, zorder=0)
ax.set_title("DMR Curvature Distribution\n(U-shape = morula reset geometry)",
             fontsize=10, fontweight="bold", pad=5)
plabel(ax, "G")

# ── Row 2, Panel H: Accessibility × duality coupling ─────────────────────────
ax = fig.add_subplot(gs[1, 3])
for branch in ["access", "closure", "other"]:
    sub = df_triad[df_triad["branch"]==branch]
    color = BRANCH_COLORS[branch]
    sizes = sub["n_dmr"] * 8 + 30
    ax.scatter(sub["mean_morula_accessibility"], sub["mean_signed_duality"],
               s=sizes, color=color, alpha=0.85,
               edgecolors="white", linewidths=0.8, zorder=4,
               label=BRANCH_LABELS[branch])
    for _, row in sub.iterrows():
        if row["module_id"] in ["M02","M05","M12","M10","M01"]:
            ax.annotate(row["module_id"],
                        (row["mean_morula_accessibility"], row["mean_signed_duality"]),
                        textcoords="offset points", xytext=(6,3),
                        fontsize=7.5, fontweight="bold", color=color,
                        path_effects=[pe.withStroke(linewidth=2,foreground="white")],
                        zorder=8)
x_all = df_triad["mean_morula_accessibility"].values
y_all = df_triad["mean_signed_duality"].values
coef = np.polyfit(x_all, y_all, 1)
x_fit = np.linspace(x_all.min(), x_all.max(), 100)
ax.plot(x_fit, np.polyval(coef, x_fit),
        color="#555", lw=1.2, ls="--", alpha=0.6)
rho_row = df_corr[df_corr["metric"]=="signed_duality"].iloc[0]
ax.text(0.97, 0.04,
        f"ρ = {rho_row['observed_spearman']:.3f}\np = {rho_row['empirical_p_ge_observed']:.3f}",
        transform=ax.transAxes, fontsize=8.5, va="bottom", ha="right",
        bbox=dict(boxstyle="round,pad=0.4",facecolor="#FFEAA7",
                  edgecolor="#FDCB6E",linewidth=1,alpha=0.95))
ax.set_xlabel("Mean morula accessibility (ATAC)", fontsize=9)
ax.set_ylabel("Mean signed duality score", fontsize=9)
ax.legend(loc="upper left", fontsize=7, framealpha=0.9)
ax.grid(True, alpha=0.15, linewidth=0.5, zorder=0)
ax.set_title("Chromatin Accessibility × Duality\n(dot size = n DMRs per module)",
             fontsize=10, fontweight="bold", pad=5)
plabel(ax, "H")

# ── Save ──────────────────────────────────────────────────────────────────────
out_png = OUT/"CEEF_PhaseB_Final_Figure.png"
out_pdf = OUT/"CEEF_PhaseB_Final_Figure.pdf"
plt.savefig(out_png, dpi=300)
try:
    out_pdf.unlink(missing_ok=True)
    plt.savefig(out_pdf)
    print(f"Saved: {out_pdf}")
except Exception as e:
    print(f"PDF skipped: {e}")
plt.close()
print(f"Saved: {out_png}")
```
