## Supplemental code for "An entropy-based diagnostic framework for characterizing methylation state dynamics during preimplantation development": build_ceef_model.py

```
#!/usr/bin/env python
"""
Build COMSOL CEEF Model A using mph Python API.
5 scenarios: baseline, +ZGA, +entry, full_control, wrong_exit
"""
import mph
import numpy as np
import pandas as pd
from pathlib import Path
import json

OUT = Path("E:/progress_comsol_analysis")

# Load data
params  = pd.read_csv(OUT/"parameters.csv").iloc[0].to_dict()
centers = pd.read_csv(OUT/"stage_centers_2d.csv").set_index("stage")

print("Starting COMSOL 6.4...")
client = mph.start(cores=2)
print("COMSOL started successfully")

def build_model(scenario_name, g_zga=1.0, g_entry=1.0, g_exit=1.0, wrong_exit=False):
    """Build CEEF Model A for a given scenario."""
    model = client.create(f"CEEF_{scenario_name}")
    java  = model.java

    # ── Parameters ────────────────────────────────────────────────────────────
    java.param().set("alpha_op", "0.5611")
    java.param().set("bias_op",  "0.0688")
    java.param().set("lambda_K", "0.4389")

    # Stage centers
    stage_map = {
        "oocyte":  "MII oocyte",
        "zygote":  "zygote/PN",
        "2cell":   "2-cell",
        "4cell":   "4-cell",
        "8cell":   "8-cell",
        "morula":  "morula",
        "blast":   "blastocyst",
    }
    for short, full in stage_map.items():
        if full in centers.index:
            java.param().set(f"z1_{short}", str(centers.loc[full, "z1"]))
            java.param().set(f"z2_{short}", str(centers.loc[full, "z2"]))

    # K-attractor (mean of all stage centers)
    java.param().set("z1_Katt", str(params["k_att_z1"]))
    java.param().set("z2_Katt", str(params["k_att_z2"]))

    # Basin radii and sigmas
    java.param().set("r_morula", str(params["r_morula"]))
    java.param().set("r_8cell",  str(params["r_8cell"]))
    java.param().set("r_blast",  str(params["r_blast"]))
    java.param().set("sigma_M",  str(params["sigma_M"]))
    java.param().set("sigma_8",  str(params["sigma_8"]))
    java.param().set("sigma_4",  "2.0")

    # Control flags
    java.param().set("g_zga_flag",   str(g_zga))
    java.param().set("g_entry_flag", str(g_entry))
    java.param().set("g_exit_flag",  str(g_exit))
    java.param().set("exit_sign",    "-1.0" if wrong_exit else "1.0")

    # Control strengths
    java.param().set("gamma_acc",     "1.0")
    java.param().set("gamma_closure", "1.0")
    java.param().set("gamma_zga",     "1.0")
    java.param().set("gamma_re",      "1.0")
    java.param().set("gamma_de",      "1.0")

    # ── Component ─────────────────────────────────────────────────────────────
    java.modelNode().create("comp1", True)

    # ── Physics: Global ODEs ──────────────────────────────────────────────────
    java.physics().create("ge", "GlobalEquations", "comp1")
    java.physics("ge").label("DMR State Space ODE")

    ge = java.physics("ge").feature("ge1")
    ge.set("name", ["z1", "z2"])
    ge.set("equation", [
        "z1t - (FK1 + g_zga_flag*Fzga1 + g_entry_flag*Fentry1 + g_exit_flag*Fexit1)",
        "z2t - (FK2 + g_zga_flag*Fzga2 + g_entry_flag*Fentry2 + g_exit_flag*Fexit2)"
    ])
    ge.set("initialValueU",  ["z1_oocyte", "z2_oocyte"])
    ge.set("initialValueUt", ["0", "0"])

    # ── Variables ─────────────────────────────────────────────────────────────
    java.variable().create("var1")
    java.variable("var1").model("comp1")

    v = java.variable("var1")

    # Squared distances
    v.set("rM2", "(z1-z1_morula)^2+(z2-z2_morula)^2")
    v.set("r82", "(z1-z1_8cell)^2+(z2-z2_8cell)^2")
    v.set("r42", "(z1-z1_4cell)^2+(z2-z2_4cell)^2")
    v.set("rB2", "(z1-z1_blast)^2+(z2-z2_blast)^2")

    # Gaussian envelopes
    v.set("gM", "exp(-rM2/(2*sigma_M^2))")
    v.set("g8", "exp(-r82/(2*sigma_8^2))")
    v.set("g4", "exp(-r42/(2*sigma_4^2))")

    # Time gate functions (smooth step)
    v.set("chi_zga",   "flc2hs(tau-3,0.3)*flc2hs(4-tau,0.3)")
    v.set("chi_entry", "flc2hs(tau-4,0.3)*flc2hs(5-tau,0.3)")
    v.set("chi_exit",  "flc2hs(tau-5,0.3)*flc2hs(6-tau,0.3)")

    # F_K: methylation-only baseline field
    v.set("FK1", "-lambda_K*(z1-z1_Katt)")
    v.set("FK2", "-lambda_K*(z2-z2_Katt)")

    # Normalized direction vectors
    v.set("vzga_mag",   "sqrt((z1_8cell-z1_4cell)^2+(z2_8cell-z2_4cell)^2)+1e-12")
    v.set("vzga1",      "(z1_8cell-z1_4cell)/vzga_mag")
    v.set("vzga2",      "(z2_8cell-z2_4cell)/vzga_mag")

    v.set("ventry_mag", "sqrt((z1_morula-z1_8cell)^2+(z2_morula-z2_8cell)^2)+1e-12")
    v.set("ventry1",    "(z1_morula-z1_8cell)/ventry_mag")
    v.set("ventry2",    "(z2_morula-z2_8cell)/ventry_mag")

    v.set("vexit_mag",  "sqrt((z1_blast-z1_morula)^2+(z2_blast-z2_morula)^2)+1e-12")
    v.set("vexit1",     "exit_sign*(z1_blast-z1_morula)/vexit_mag")
    v.set("vexit2",     "exit_sign*(z2_blast-z2_morula)/vexit_mag")

    # Field components
    v.set("Fzga1",   "chi_zga*gamma_zga*g4*vzga1")
    v.set("Fzga2",   "chi_zga*gamma_zga*g4*vzga2")
    v.set("Fentry1", "chi_entry*(gamma_acc+gamma_closure)*g8*ventry1")
    v.set("Fentry2", "chi_entry*(gamma_acc+gamma_closure)*g8*ventry2")
    v.set("Fexit1",  "chi_exit*(gamma_re+gamma_de)*gM*vexit1")
    v.set("Fexit2",  "chi_exit*(gamma_re+gamma_de)*gM*vexit2")

    # Basin occupancy indicators
    v.set("in_morula_basin", "if(sqrt(rM2)<r_morula,1,0)")
    v.set("in_8cell_basin",  "if(sqrt(r82)<r_8cell,1,0)")
    v.set("in_blast_basin",  "if(sqrt(rB2)<r_blast,1,0)")

    # Entry-exit duality cosine
    v.set("entry_exit_cos", "ventry1*vexit1+ventry2*vexit2")

    # Distance to morula center
    v.set("dist_morula", "sqrt(rM2)")
    v.set("dist_blast",  "sqrt(rB2)")

    # ── Study ─────────────────────────────────────────────────────────────────
    java.study().create("std1")
    java.study("std1").label(scenario_name)
    java.study("std1").create("time", "Transient")
    java.study("std1").feature("time").set("tlist", "range(0,0.05,6)")
    java.study("std1").feature("time").set("usertol", True)
    java.study("std1").feature("time").set("rtol", "1e-4")
    java.study("std1").feature("time").set("atol", "1e-6")

    # ── Results ───────────────────────────────────────────────────────────────
    # Trajectory plot
    java.result().create("pg_traj", "PlotGroup1D")
    java.result("pg_traj").label("Trajectory z1 z2 vs tau")
    java.result("pg_traj").create("glob1", "Global")
    java.result("pg_traj").feature("glob1").set("expr", ["z1", "z2"])
    java.result("pg_traj").feature("glob1").set("descr", ["z1(tau)", "z2(tau)"])

    # Basin occupancy
    java.result().create("pg_basin", "PlotGroup1D")
    java.result("pg_basin").label("Basin Occupancy")
    java.result("pg_basin").create("glob1", "Global")
    java.result("pg_basin").feature("glob1").set("expr",
        ["in_morula_basin", "in_8cell_basin", "in_blast_basin"])
    java.result("pg_basin").feature("glob1").set("descr",
        ["Morula basin", "8-cell basin", "Blast basin"])

    # Distance to morula
    java.result().create("pg_dist", "PlotGroup1D")
    java.result("pg_dist").label("Distance to Morula Center")
    java.result("pg_dist").create("glob1", "Global")
    java.result("pg_dist").feature("glob1").set("expr", ["dist_morula", "dist_blast"])
    java.result("pg_dist").feature("glob1").set("descr",
        ["dist to morula", "dist to blast"])

    return model


# ── Build and run all 5 scenarios ─────────────────────────────────────────────
scenarios = [
    ("baseline_only",  0.0, 0.0, 0.0, False),
    ("plus_zga",       1.0, 0.0, 0.0, False),
    ("plus_entry",     1.0, 1.0, 0.0, False),
    ("full_control",   1.0, 1.0, 1.0, False),
    ("wrong_exit",     1.0, 1.0, 1.0, True),
]

results_summary = {}

for name, g_zga, g_entry, g_exit, wrong in scenarios:
    print(f"\n{'='*50}")
    print(f"Building scenario: {name}")
    model = build_model(name, g_zga, g_entry, g_exit, wrong)

    print(f"  Running study...")
    try:
        model.solve("std1")
        print(f"  Solved successfully")

        # Extract results
        java = model.java
        # Get time points and solution
        t_data = java.sol("sol1").getPVals()
        z1_data = java.sol("sol1").getU(0)  # z1
        z2_data = java.sol("sol1").getU(1)  # z2

        # Final position
        z1_final = float(z1_data[-1]) if len(z1_data) > 0 else np.nan
        z2_final = float(z2_data[-1]) if len(z2_data) > 0 else np.nan

        # Check if in morula basin at tau=5
        morula_center = np.array([centers.loc["morula","z1"], centers.loc["morula","z2"]])
        r_morula = float(params["r_morula"])

        # Find tau=5 index
        t_arr = np.array([float(t) for t in t_data])
        idx_morula = np.argmin(np.abs(t_arr - 5.0))
        z1_at_morula = float(z1_data[idx_morula])
        z2_at_morula = float(z2_data[idx_morula])
        dist_at_morula = np.sqrt((z1_at_morula-morula_center[0])**2 +
                                  (z2_at_morula-morula_center[1])**2)
        in_morula = dist_at_morula < r_morula

        # Check if in blast basin at tau=6
        blast_center = np.array([centers.loc["blastocyst","z1"], centers.loc["blastocyst","z2"]])
        r_blast = float(params["r_blast"])
        dist_at_blast = np.sqrt((z1_final-blast_center[0])**2 + (z2_final-blast_center[1])**2)
        in_blast = dist_at_blast < r_blast

        results_summary[name] = {
            "z1_final": z1_final, "z2_final": z2_final,
            "z1_at_morula_tau": z1_at_morula, "z2_at_morula_tau": z2_at_morula,
            "dist_to_morula_at_tau5": float(dist_at_morula),
            "in_morula_basin": bool(in_morula),
            "dist_to_blast_at_tau6": float(dist_at_blast),
            "in_blast_basin": bool(in_blast),
            "n_timepoints": len(t_arr),
        }
        print(f"  tau=5: z1={z1_at_morula:.3f}, z2={z2_at_morula:.3f}, "
              f"dist_morula={dist_at_morula:.3f}, in_morula={in_morula}")
        print(f"  tau=6: z1={z1_final:.3f}, z2={z2_final:.3f}, "
              f"dist_blast={dist_at_blast:.3f}, in_blast={in_blast}")

        # Export trajectory data
        traj_df = pd.DataFrame({
            "tau": [float(t) for t in t_data],
            "z1": [float(x) for x in z1_data],
            "z2": [float(x) for x in z2_data],
        })
        traj_df.to_csv(OUT/f"trajectory_{name}.csv", index=False)
        print(f"  Trajectory saved: trajectory_{name}.csv")

    except Exception as e:
        print(f"  ERROR: {e}")
        results_summary[name] = {"error": str(e)}

    # Save model
    save_path = str(OUT / f"CEEF_{name}.mph")
    model.java.save(save_path)
    print(f"  Model saved: CEEF_{name}.mph")
    client.remove(model)

# Save summary
with open(OUT/"scenario_results_summary.json","w",encoding="utf-8") as f:
    json.dump(results_summary, f, indent=2, ensure_ascii=False, default=str)

print("\n" + "="*60)
print("ALL SCENARIOS COMPLETE")
print("="*60)
for name, res in results_summary.items():
    if "error" not in res:
        print(f"  {name}: morula={res['in_morula_basin']}, blast={res['in_blast_basin']}, "
              f"dist_morula={res['dist_to_morula_at_tau5']:.3f}")
    else:
        print(f"  {name}: ERROR - {res['error'][:50]}")

print(f"\nFiles in {OUT}:")
for f in sorted(OUT.iterdir()):
    print(f"  {f.name}")
```
