## Supplemental code for "An entropy-based diagnostic framework for characterizing methylation state dynamics during preimplantation development": CEEF_FieldModel.java

```
/*
 * CEEF Phase A — 2D Epigenetic Vector Field Model
 * COMSOL 6.4 Java API
 *
 * Builds a 2D spatial field in (z1,z2) = PCA latent space,
 * renders Arrow Surface + Streamline + Surface (|F|) at 4 tau snapshots,
 * exports high-resolution PNG via model.result().export() API.
 *
 * Compile & run:
 *   comsolcompile CEEF_FieldModel.java
 *   comsolbatch -inputfile CEEF_FieldModel.class
 */

import com.comsol.model.*;
import com.comsol.model.util.*;
import java.io.*;

public class CEEF_FieldModel {

    static final String OUT_DIR = "C:\\comsolfigs";
    static final String MPH_OUT = "C:\\comsolfigs\\CEEF_FieldModel.mph";

    static final double[] TAU_SNAPS  = {1.5,  3.5,  4.5,  5.5};
    static final String[] TAU_LABELS = {
        "tau1p5_FK_only",
        "tau3p5_plus_ZGA",
        "tau4p5_plus_entry",
        "tau5p5_full_control"
    };

    public static void main(String[] args) throws Exception {
        new File(OUT_DIR).mkdirs();
        Model model = buildModel();
        System.out.println("Saving model ...");
        model.save(MPH_OUT);
        System.out.println("Saved: " + MPH_OUT);
        exportFigures(model);
        ModelUtil.disconnect();
        System.out.println("All done. Output: " + OUT_DIR);
    }

    // ── Build model ───────────────────────────────────────────────────────────
    static Model buildModel() throws Exception {
        Model model = ModelUtil.create("CEEF_FieldModel");
        model.modelPath("E:\\progress_comsol_analysis");
        model.label("CEEF Phase A — Epigenetic Vector Field");

        // ── Parameters — correction-vector space stage centers ────────────────
        model.param().set("z1_oocyte",  "0.9656");
        model.param().set("z2_oocyte",  "0.2348");
        model.param().set("z1_zygote",  "0.4654");
        model.param().set("z2_zygote",  "0.3707");
        model.param().set("z1_2cell",   "0.6214");
        model.param().set("z2_2cell",   "0.2323");
        model.param().set("z1_4cell",   "0.5680");
        model.param().set("z2_4cell",   "0.5554");
        model.param().set("z1_8cell",   "0.2150");
        model.param().set("z2_8cell",   "-0.1490");
        model.param().set("z1_morula",  "3.6330");
        model.param().set("z2_morula",  "-0.2790");
        model.param().set("z1_blast",   "0.6060");
        model.param().set("z2_blast",   "1.5399");
        // K-attractor: midpoint of oocyte and 8cell
        model.param().set("z1_Katt",    "0.59");
        model.param().set("z2_Katt",    "0.04");
        // Physics parameters (from parameters_corrected.csv)
        model.param().set("lambda_K",   "0.4389");
        model.param().set("sigma_M",    "0.2132",  "1.5 * r_morula");
        model.param().set("sigma_8",    "0.7415",  "1.5 * r_8cell");
        model.param().set("sigma_4",    "0.30");
        model.param().set("gamma_acc",  "1.0");
        model.param().set("gamma_closure", "1.0");
        model.param().set("gamma_zga",  "1.0");
        model.param().set("gamma_re",   "1.0");
        model.param().set("gamma_de",   "1.0");
        model.param().set("tau",        "4.5", "Operator time (swept in export)");

        // ── Component & 2D geometry ───────────────────────────────────────────
        model.modelNode().create("comp1", true);
        model.geom().create("geom1", 2);
        model.geom("geom1").create("r1", "Rectangle");
        // Domain: correction-vector space z1 in [-0.6, 5.8], z2 in [-2.2, 2.2]
        model.geom("geom1").feature("r1").set("pos",  new double[]{-0.6, -2.2});
        model.geom("geom1").feature("r1").set("size", new double[]{6.4, 4.4});
        model.geom("geom1").run("fin");

        // ── Mesh ──────────────────────────────────────────────────────────────
        model.mesh().create("mesh1", "geom1");
        model.mesh("mesh1").create("ftri1", "FreeTri");
        model.mesh("mesh1").feature("size").set("hauto", 2);
        model.mesh("mesh1").run();

        // ── Variables: CEEF vector field F(x,y,tau) ───────────────────────────
        model.variable().create("var1");
        model.variable("var1").model("comp1");

        // Squared distances
        model.variable("var1").set("rM2", "(x-z1_morula)^2+(y-z2_morula)^2");
        model.variable("var1").set("r82", "(x-z1_8cell)^2+(y-z2_8cell)^2");
        model.variable("var1").set("r42", "(x-z1_4cell)^2+(y-z2_4cell)^2");
        model.variable("var1").set("rB2", "(x-z1_blast)^2+(y-z2_blast)^2");

        // Gaussian envelopes
        model.variable("var1").set("gM", "exp(-rM2/(2*sigma_M^2))");
        model.variable("var1").set("g8", "exp(-r82/(2*sigma_8^2))");
        model.variable("var1").set("g4", "exp(-r42/(2*sigma_4^2))");

        // Time gates (smooth Heaviside, built-in flc2hs)
        model.variable("var1").set("chi_zga",
            "flc2hs(tau-3,0.3)*flc2hs(4-tau,0.3)");
        model.variable("var1").set("chi_entry",
            "flc2hs(tau-4,0.3)*flc2hs(5-tau,0.3)");
        model.variable("var1").set("chi_exit",
            "flc2hs(tau-5,0.3)*flc2hs(6-tau,0.3)");

        // F_K: methylation baseline
        model.variable("var1").set("FK1", "-lambda_K*(x-z1_Katt)");
        model.variable("var1").set("FK2", "-lambda_K*(y-z2_Katt)");

        // Unit direction vectors
        model.variable("var1").set("d_zga",
            "sqrt((z1_8cell-z1_4cell)^2+(z2_8cell-z2_4cell)^2)");
        model.variable("var1").set("vzga1", "(z1_8cell-z1_4cell)/d_zga");
        model.variable("var1").set("vzga2", "(z2_8cell-z2_4cell)/d_zga");

        model.variable("var1").set("d_entry",
            "sqrt((z1_morula-z1_8cell)^2+(z2_morula-z2_8cell)^2)");
        model.variable("var1").set("ventry1", "(z1_morula-z1_8cell)/d_entry");
        model.variable("var1").set("ventry2", "(z2_morula-z2_8cell)/d_entry");

        model.variable("var1").set("d_exit",
            "sqrt((z1_blast-z1_morula)^2+(z2_blast-z2_morula)^2)");
        model.variable("var1").set("vexit1", "(z1_blast-z1_morula)/d_exit");
        model.variable("var1").set("vexit2", "(z2_blast-z2_morula)/d_exit");

        // Component fields
        model.variable("var1").set("Fzga1",   "chi_zga*gamma_zga*g4*vzga1");
        model.variable("var1").set("Fzga2",   "chi_zga*gamma_zga*g4*vzga2");
        model.variable("var1").set("Fentry1", "chi_entry*(gamma_acc+gamma_closure)*g8*ventry1");
        model.variable("var1").set("Fentry2", "chi_entry*(gamma_acc+gamma_closure)*g8*ventry2");
        model.variable("var1").set("Fexit1",  "chi_exit*(gamma_re+gamma_de)*gM*vexit1");
        model.variable("var1").set("Fexit2",  "chi_exit*(gamma_re+gamma_de)*gM*vexit2");

        // Total field
        model.variable("var1").set("Fu1", "FK1+Fzga1+Fentry1+Fexit1");
        model.variable("var1").set("Fu2", "FK2+Fzga2+Fentry2+Fexit2");
        model.variable("var1").set("Fmag", "sqrt(Fu1^2+Fu2^2)");
        model.variable("var1").set("Fu1n", "Fu1/(Fmag+1e-9)");
        model.variable("var1").set("Fu2n", "Fu2/(Fmag+1e-9)");

        // ── Physics: trivial Laplace to generate mesh + solution ──────────────
        // CoefficientFormPDE: -div(c*grad(u)) + a*u = f
        // Set c=1, a=0, f=0 => Laplace equation, u=0 on boundary
        model.physics().create("c", "CoefficientFormPDE", "geom1");
        // Do NOT rename the field — keep default "u"
        model.physics("c").feature("cfeq1").set("c", "1");
        model.physics("c").feature("cfeq1").set("a", "0");
        model.physics("c").feature("cfeq1").set("f", "0");
        // Dirichlet BC: u = 0 on all boundaries
        model.physics("c").create("dir1", "DirichletBoundary", 1);
        model.physics("c").feature("dir1").selection().all();
        model.physics("c").feature("dir1").set("r", "0");

        // ── Study: stationary ─────────────────────────────────────────────────
        model.study().create("std1");
        model.study("std1").label("Stationary");
        model.study("std1").create("stat", "Stationary");
        model.study("std1").run();

        // ── Result plot groups ────────────────────────────────────────────────
        buildPlotGroups(model);

        // ── Export features ───────────────────────────────────────────────────
        buildExportFeatures(model);

        return model;
    }

    // ── Plot groups ───────────────────────────────────────────────────────────
    static void buildPlotGroups(Model model) throws Exception {
        for (int i = 0; i < TAU_SNAPS.length; i++) {
            double tau = TAU_SNAPS[i];
            String pg  = "pg" + (i + 1);
            String[][] tauConst = {{"tau", String.valueOf(tau), "Operator time"}};

            model.result().create(pg, "PlotGroup2D");
            model.result(pg).label(TAU_LABELS[i]);
            model.result(pg).set("titletype", "manual");
            model.result(pg).set("title", TAU_LABELS[i].replace("_", " "));
            model.result(pg).set("xlabel", "z1 (PC1)");
            model.result(pg).set("ylabel", "z2 (PC2)");

            // Layer 1: Surface — |F| with Thermal colormap
            model.result(pg).create("surf1", "Surface");
            model.result(pg).feature("surf1").set("expr", "Fmag");
            model.result(pg).feature("surf1").set("const", tauConst);
            model.result(pg).feature("surf1").set("colortable", "Thermal");
            model.result(pg).feature("surf1").set("colorscalemode", "linear");
            model.result(pg).feature("surf1").set("rangecoloractive", "on");
            model.result(pg).feature("surf1").set("rangecolormin", "0");
            model.result(pg).feature("surf1").set("rangecolormax", "0.5");
            model.result(pg).feature("surf1").set("smooth", "internal");

            // Layer 2: Streamline — flow lines (white)
            model.result(pg).create("str1", "Streamline");
            model.result(pg).feature("str1").set("expr", new String[]{"Fu1", "Fu2"});
            model.result(pg).feature("str1").set("const", tauConst);
            model.result(pg).feature("str1").set("posmethod", "start");
            model.result(pg).feature("str1").set("arrowcount", "200");
            model.result(pg).feature("str1").set("color", "white");
            model.result(pg).feature("str1").set("linetype", "tube");
            model.result(pg).feature("str1").set("tuberadiusscaleactive", "on");
            model.result(pg).feature("str1").set("tuberadiusscale", "0.035");
            model.result(pg).feature("str1").set("inheritcolor", "off");

            // Layer 3: Arrow Surface — normalized direction
            model.result(pg).create("arw1", "ArrowSurface");
            model.result(pg).feature("arw1").set("expr", new String[]{"Fu1n", "Fu2n"});
            model.result(pg).feature("arw1").set("const", tauConst);
            model.result(pg).feature("arw1").set("xnumber", "22");
            model.result(pg).feature("arw1").set("ynumber", "18");
            model.result(pg).feature("arw1").set("arrowlength", "normalized");
            model.result(pg).feature("arw1").set("arrowscale", "0.30");
            model.result(pg).feature("arw1").set("color", "gray");
            model.result(pg).feature("arw1").set("inheritcolor", "off");
        }
    }

    // ── Export features ───────────────────────────────────────────────────────
    static void buildExportFeatures(Model model) throws Exception {
        for (int i = 0; i < TAU_SNAPS.length; i++) {
            String expTag  = "exp" + (i + 1);
            String pg      = "pg" + (i + 1);
            String pngPath = OUT_DIR + "\\" + TAU_LABELS[i] + ".png";

            model.result().export().create(expTag, pg, "Image2D");
            model.result().export(expTag).set("sourceobject", pg);
            model.result().export(expTag).set("pngfilename", pngPath);
            model.result().export(expTag).set("imagetype",   "png");
            model.result().export(expTag).set("size",        "current");
            model.result().export(expTag).set("width",       "2400");
            model.result().export(expTag).set("height",      "1800");
            model.result().export(expTag).set("unit",        "px");
            model.result().export(expTag).set("resolution",  "300");
            model.result().export(expTag).set("antialias",   "on");
            model.result().export(expTag).set("zoomextents", "on");
            model.result().export(expTag).set("fontsize",    "22");
            model.result().export(expTag).set("logo2d",      "off");
            model.result().export(expTag).set("axes2d",      "on");
            model.result().export(expTag).set("legend2d",    "on");
            model.result().export(expTag).set("title2d",     "on");
        }
    }

    // ── Run exports ───────────────────────────────────────────────────────────
    static void exportFigures(Model model) throws Exception {
        System.out.println("Running plot groups and exporting ...");
        for (int i = 0; i < TAU_SNAPS.length; i++) {
            String pg     = "pg" + (i + 1);
            String expTag = "exp" + (i + 1);
            String label  = TAU_LABELS[i];
            try {
                System.out.println("  Plotting " + label + " ...");
                model.result(pg).run();
                System.out.println("  Exporting " + label + " ...");
                model.result().export(expTag).run();
                System.out.println("  OK: " + OUT_DIR + "\\" + label + ".png");
            } catch (Exception e) {
                System.out.println("  WARN [" + label + "]: " + e.getMessage());
            }
        }
    }
}
```
