## Supplemental code for "An entropy-based diagnostic framework for characterizing methylation state dynamics during preimplantation development": CEEF_FieldModel_PhaseB.java

```
/*
 * CEEF Phase B — Chromatin-Gated Control Operator Field Model
 * COMSOL 6.4 Java API
 *
 * Phase B: Shows how chromatin accessibility (M02) and closure (M05)
 * gate the entry/exit control operators.
 *
 * 4 scenarios (matching Phase A panel layout):
 *   A: F_entry only, M02 access-gated (gamma_acc = 1.1619)
 *   B: F_exit only,  M05 closure-gated (gamma_closure = 1.8940)
 *   C: Full control (F_entry + F_exit, both chromatin-gated)
 *   D: Wrong-exit (F_entry + reversed F_exit = collapse)
 *
 * Run:
 *   comsolcompile CEEF_FieldModel_PhaseB.java
 *   comsolbatch -inputfile CEEF_FieldModel_PhaseB.class
 */

import com.comsol.model.*;
import com.comsol.model.util.*;
import java.io.*;

public class CEEF_FieldModel_PhaseB {

    static final String OUT_DIR = "C:\\comsolfigs_b";
    static final String MPH_OUT = "C:\\comsolfigs_b\\CEEF_FieldModel_PhaseB.mph";

    // Phase B scenarios — chromatin-gated operators
    static final String[] LABELS = {
        "PhaseB_A_M02_access_entry",
        "PhaseB_B_M05_closure_exit",
        "PhaseB_C_full_chromatin_control",
        "PhaseB_D_wrong_exit_collapse"
    };
    static final String[] TITLES = {
        "M02 Access-Gated Entry (F_entry, gamma_acc=1.16)",
        "M05 Closure-Gated Exit (F_exit, gamma_clo=1.89)",
        "Full Chromatin Control (M02+M05 gated)",
        "Wrong-Exit Collapse (reversed F_exit)"
    };
    // Colormaps matching Phase A panels C, D, D, D
    static final String[] CMAPS = {"hot", "plasma", "YlOrBr", "RdPu"};

    // Chromatin gate strengths from Phase B data
    // M02 (access/entry): mean_morula_accessibility = 1.1619
    // M05 (closure/exit): mean_morula_accessibility = 1.8940
    static final double GAMMA_ACC_M02    = 1.1619;
    static final double GAMMA_CLOSURE_M05 = 1.8940;

    public static void main(String[] args) throws Exception {
        new File(OUT_DIR).mkdirs();
        Model model = buildModel();
        System.out.println("Saving model ...");
        model.save(MPH_OUT);
        System.out.println("Saved: " + MPH_OUT);
        exportFigures(model);
        ModelUtil.disconnect();
        System.out.println("Done. Figures in: " + OUT_DIR);
    }

    static Model buildModel() throws Exception {
        Model model = ModelUtil.create("CEEF_FieldModel_PhaseB");
        model.modelPath("E:\\progress_comsol_analysis");
        model.label("CEEF Phase B — Chromatin-Gated Control Field");

        // ── Parameters — correction-vector space stage centers ────────────────
        model.param().set("z1_oocyte",  "0.9656");
        model.param().set("z2_oocyte",  "0.2348");
        model.param().set("z1_4cell",   "0.5680");
        model.param().set("z2_4cell",   "0.5554");
        model.param().set("z1_8cell",   "0.2150");
        model.param().set("z2_8cell",   "-0.1490");
        model.param().set("z1_morula",  "3.6330");
        model.param().set("z2_morula",  "-0.2790");
        model.param().set("z1_blast",   "0.6060");
        model.param().set("z2_blast",   "1.5399");
        model.param().set("z1_Katt",    "0.59");
        model.param().set("z2_Katt",    "0.04");

        // Physics parameters — lambda_K reduced to 0.05 so control operators dominate
        model.param().set("lambda_K",       "0.05");
        model.param().set("sigma_M",        "0.2132");
        model.param().set("sigma_8",        "0.7415");
        model.param().set("sigma_4",        "0.30");

        // Phase B chromatin gate strengths
        model.param().set("gamma_acc_M02",      String.valueOf(GAMMA_ACC_M02));
        model.param().set("gamma_closure_M05",  String.valueOf(GAMMA_CLOSURE_M05));
        model.param().set("gamma_re",           "1.0");
        model.param().set("gamma_de",           "1.0");
        model.param().set("gamma_wrong",        "-1.0");

        // Scenario switches (set per plot group)
        model.param().set("sw_entry",  "1.0");
        model.param().set("sw_exit",   "1.0");
        model.param().set("sw_wrong",  "0.0");

        // ── Component & 2D geometry ───────────────────────────────────────────
        model.modelNode().create("comp1", true);
        model.geom().create("geom1", 2);
        model.geom("geom1").create("r1", "Rectangle");
        model.geom("geom1").feature("r1").set("pos",  new double[]{-0.6, -2.2});
        model.geom("geom1").feature("r1").set("size", new double[]{6.4, 4.4});
        model.geom("geom1").run("fin");

        // ── Mesh ──────────────────────────────────────────────────────────────
        model.mesh().create("mesh1", "geom1");
        model.mesh("mesh1").create("ftri1", "FreeTri");
        model.mesh("mesh1").feature("size").set("hauto", 2);
        model.mesh("mesh1").run();

        // ── Variables: chromatin-gated CEEF field ─────────────────────────────
        model.variable().create("var1");
        model.variable("var1").model("comp1");

        // Squared distances
        model.variable("var1").set("rM2", "(x-z1_morula)^2+(y-z2_morula)^2");
        model.variable("var1").set("r82", "(x-z1_8cell)^2+(y-z2_8cell)^2");
        model.variable("var1").set("r42", "(x-z1_4cell)^2+(y-z2_4cell)^2");

        // Gaussian envelopes
        model.variable("var1").set("gM", "exp(-rM2/(2*sigma_M^2))");
        model.variable("var1").set("g8", "exp(-r82/(2*sigma_8^2))");
        model.variable("var1").set("g4", "exp(-r42/(2*sigma_4^2))");

        // F_K: methylation baseline (always active)
        model.variable("var1").set("FK1", "-lambda_K*(x-z1_Katt)");
        model.variable("var1").set("FK2", "-lambda_K*(y-z2_Katt)");

        // Unit direction vectors
        model.variable("var1").set("d_entry",
            "sqrt((z1_morula-z1_8cell)^2+(z2_morula-z2_8cell)^2)");
        model.variable("var1").set("ventry1", "(z1_morula-z1_8cell)/d_entry");
        model.variable("var1").set("ventry2", "(z2_morula-z2_8cell)/d_entry");

        model.variable("var1").set("d_exit",
            "sqrt((z1_blast-z1_morula)^2+(z2_blast-z2_morula)^2)");
        model.variable("var1").set("vexit1", "(z1_blast-z1_morula)/d_exit");
        model.variable("var1").set("vexit2", "(z2_blast-z2_morula)/d_exit");

        // F_entry: M02 access-gated (gamma_acc = M02 accessibility)
        model.variable("var1").set("Fentry1",
            "sw_entry*gamma_acc_M02*g8*ventry1");
        model.variable("var1").set("Fentry2",
            "sw_entry*gamma_acc_M02*g8*ventry2");

        // F_exit: M05 closure-gated (gamma_closure = M05 accessibility)
        // sw_wrong=-1 reverses exit direction for collapse scenario
        model.variable("var1").set("Fexit1",
            "sw_exit*(gamma_re+gamma_de)*gM*(vexit1+sw_wrong*(-2*vexit1))");
        model.variable("var1").set("Fexit2",
            "sw_exit*(gamma_re+gamma_de)*gM*(vexit2+sw_wrong*(-2*vexit2))");

        // Total field
        model.variable("var1").set("Fu1", "FK1+Fentry1+Fexit1");
        model.variable("var1").set("Fu2", "FK2+Fentry2+Fexit2");
        model.variable("var1").set("Fmag", "sqrt(Fu1^2+Fu2^2)");

        // Active operator magnitudes for pseudocolor
        model.variable("var1").set("Fentry_mag", "sqrt(Fentry1^2+Fentry2^2)");
        model.variable("var1").set("Fexit_mag",  "sqrt(Fexit1^2+Fexit2^2)");
        model.variable("var1").set("Fcontrol_mag",
            "sqrt((Fentry1+Fexit1)^2+(Fentry2+Fexit2)^2)");
        // For wrong-exit: show magnitude of the reversed exit field
        model.variable("var1").set("Fwrong_mag",
            "sqrt(Fexit1^2+Fexit2^2)");

        // Normalized direction
        model.variable("var1").set("Fu1n", "Fu1/(Fmag+1e-9)");
        model.variable("var1").set("Fu2n", "Fu2/(Fmag+1e-9)");

        // ── Physics: trivial Laplace ──────────────────────────────────────────
        model.physics().create("c", "CoefficientFormPDE", "geom1");
        model.physics("c").feature("cfeq1").set("c", "1");
        model.physics("c").feature("cfeq1").set("f", "0");
        model.physics("c").feature("cfeq1").set("a", "0");
        model.physics("c").create("dir1", "DirichletBoundary", 1);
        model.physics("c").feature("dir1").selection().all();
        model.physics("c").feature("dir1").set("r", "0");

        // ── Study ─────────────────────────────────────────────────────────────
        model.study().create("std1");
        model.study("std1").label("Stationary");
        model.study("std1").create("stat", "Stationary");
        model.study("std1").run();

        // ── Plot groups ───────────────────────────────────────────────────────
        buildPlotGroups(model);
        buildExportFeatures(model);

        return model;
    }

    // Scenario parameters: [sw_entry, sw_exit, sw_wrong, colorExpr, cmap]
    static final double[][] SCENARIO_PARAMS = {
        {1.0, 0.0, 0.0},   // A: entry only
        {0.0, 1.0, 0.0},   // B: exit only
        {1.0, 1.0, 0.0},   // C: full control
        {1.0, 1.0, 1.0},   // D: wrong exit (sw_wrong=1 reverses exit)
    };
    static final String[] COLOR_EXPRS = {
        "Fentry_mag",    // A: entry only — hot spot at 8-cell
        "Fexit_mag",     // B: exit only  — hot spot at morula
        "Fcontrol_mag",  // C: full control — both hot spots
        "Fwrong_mag"     // D: wrong exit — hot spot at morula, reversed direction
    };

    static void buildPlotGroups(Model model) throws Exception {
        for (int i = 0; i < 4; i++) {
            String pg = "pg" + (i+1);

            model.result().create(pg, "PlotGroup2D");
            model.result(pg).label(TITLES[i]);
            model.result(pg).set("titletype", "manual");
            model.result(pg).set("title", TITLES[i]);
            model.result(pg).set("xlabel", "e1 (entry correction)");
            model.result(pg).set("ylabel", "e2 (exit correction)");

            // Surface: active operator magnitude (no const — params set before re-solve)
            model.result(pg).create("surf1", "Surface");
            model.result(pg).feature("surf1").set("expr", COLOR_EXPRS[i]);
            model.result(pg).feature("surf1").set("colortable", CMAPS[i]);
            model.result(pg).feature("surf1").set("colorscalemode", "linear");
            model.result(pg).feature("surf1").set("rangecoloractive", "on");
            model.result(pg).feature("surf1").set("rangecolormin", "0");
            model.result(pg).feature("surf1").set("rangecolormax", "0.5");
            model.result(pg).feature("surf1").set("smooth", "internal");

            // Streamlines: total field
            model.result(pg).create("str1", "Streamline");
            model.result(pg).feature("str1").set("expr", new String[]{"Fu1", "Fu2"});
            model.result(pg).feature("str1").set("posmethod", "start");
            model.result(pg).feature("str1").set("arrowcount", "200");
            model.result(pg).feature("str1").set("color", "white");
            model.result(pg).feature("str1").set("linetype", "tube");
            model.result(pg).feature("str1").set("tuberadiusscaleactive", "on");
            model.result(pg).feature("str1").set("tuberadiusscale", "0.035");
            model.result(pg).feature("str1").set("inheritcolor", "off");

            // Arrow surface: normalized direction
            model.result(pg).create("arw1", "ArrowSurface");
            model.result(pg).feature("arw1").set("expr", new String[]{"Fu1n", "Fu2n"});
            model.result(pg).feature("arw1").set("xnumber", "22");
            model.result(pg).feature("arw1").set("ynumber", "18");
            model.result(pg).feature("arw1").set("arrowlength", "normalized");
            model.result(pg).feature("arw1").set("arrowscale", "0.30");
            model.result(pg).feature("arw1").set("color", "gray");
            model.result(pg).feature("arw1").set("inheritcolor", "off");
        }
    }

    static void buildExportFeatures(Model model) throws Exception {
        for (int i = 0; i < 4; i++) {
            String expTag = "exp" + (i+1);
            String pg     = "pg" + (i+1);
            String pngPath = OUT_DIR + "\\" + LABELS[i] + ".png";

            model.result().export().create(expTag, pg, "Image2D");
            model.result().export(expTag).set("sourceobject", pg);
            model.result().export(expTag).set("pngfilename", pngPath);
            model.result().export(expTag).set("imagetype",   "png");
            model.result().export(expTag).set("size",        "current");
            model.result().export(expTag).set("width",       "2400");
            model.result().export(expTag).set("height",      "1800");
            model.result().export(expTag).set("unit",        "px");
            model.result().export(expTag).set("resolution",  "300");
            model.result().export(expTag).set("antialias",   "on");
            model.result().export(expTag).set("zoomextents", "on");
            model.result().export(expTag).set("fontsize",    "22");
            model.result().export(expTag).set("logo2d",      "off");
            model.result().export(expTag).set("axes2d",      "on");
            model.result().export(expTag).set("legend2d",    "on");
            model.result().export(expTag).set("title2d",     "on");
        }
    }

    static void exportFigures(Model model) throws Exception {
        System.out.println("Exporting Phase B figures ...");

        // Scenario parameters: [sw_entry, sw_exit, sw_wrong]
        double[][] scenarios = {
            {1.0, 0.0, 0.0},   // A: entry only (M02 access-gated)
            {0.0, 1.0, 0.0},   // B: exit only  (M05 closure-gated)
            {1.0, 1.0, 0.0},   // C: full chromatin control
            {1.0, 1.0, 1.0},   // D: wrong exit collapse
        };

        for (int i = 0; i < 4; i++) {
            // Update parameters for this scenario
            model.param().set("sw_entry", String.valueOf(scenarios[i][0]));
            model.param().set("sw_exit",  String.valueOf(scenarios[i][1]));
            model.param().set("sw_wrong", String.valueOf(scenarios[i][2]));

            // Re-run study to recompute field with new parameters
            model.study("std1").run();

            // Run plot group and export
            String pg     = "pg" + (i+1);
            String expTag = "exp" + (i+1);
            try {
                System.out.println("  Plotting " + LABELS[i] + " ...");
                model.result(pg).run();
                System.out.println("  Exporting ...");
                model.result().export(expTag).run();
                System.out.println("  OK: " + OUT_DIR + "\\" + LABELS[i] + ".png");
            } catch (Exception e) {
                System.out.println("  WARN [" + LABELS[i] + "]: " + e.getMessage());
            }
        }
    }
}
```
