## Supplemental code for "An entropy-based diagnostic framework for characterizing methylation state dynamics during preimplantation development": CEEF_ModelA.java

```
/*
 * COMSOL 6.4 Java API Model
 * CEEF Phase A: Global ODE trajectory model
 * DMR operator-time dynamics in 2D latent state space
 *
 * 5 scenarios:
 * 1. baseline_only: F_K only
 * 2. plus_zga: F_K + F_ZGA
 * 3. plus_entry: F_K + F_ZGA + F_entry
 * 4. full_control: F_K + F_ZGA + F_entry + F_exit
 * 5. wrong_exit: F_K + F_ZGA + F_entry + F_wrong_exit
 */

import com.comsol.model.*;
import com.comsol.model.util.*;

public class CEEF_ModelA {

    public static Model run() throws Exception {
        Model model = ModelUtil.create("CEEF_ModelA");
        model.modelPath("E:\\progress_comsol_analysis");
        model.label("CEEF Phase A - DMR Operator-Time Dynamics");

        // ── Parameters ────────────────────────────────────────────────────────
        model.param().set("alpha_op", "0.5611", "Methylation-only operator slope");
        model.param().set("bias_op",  "0.0688", "Methylation-only operator bias");
        model.param().set("lambda_K", "0.4389", "K-field contraction rate");

        // Stage centers in 2D latent space
        model.param().set("z1_oocyte",  "7.035");
        model.param().set("z2_oocyte",  "-0.710");
        model.param().set("z1_zygote",  "1.233");
        model.param().set("z2_zygote",  "3.459");
        model.param().set("z1_2cell",   "-2.151");
        model.param().set("z2_2cell",   "-0.738");
        model.param().set("z1_4cell",   "-4.294");
        model.param().set("z2_4cell",   "-1.834");
        model.param().set("z1_8cell",   "-2.077");
        model.param().set("z2_8cell",   "-0.326");
        model.param().set("z1_morula",  "-3.174");
        model.param().set("z2_morula",  "-0.031");
        model.param().set("z1_blast",   "-4.206");
        model.param().set("z2_blast",   "-0.537");
        model.param().set("z1_Katt",    "-2.5",   "K-attractor z1");
        model.param().set("z2_Katt",    "-0.5",   "K-attractor z2");

        // Basin radii
        model.param().set("r_morula",   "1.421");
        model.param().set("r_8cell",    "2.144");
        model.param().set("r_blast",    "0.852");

        // Gaussian envelope widths
        model.param().set("sigma_M",    "2.132",  "1.5 * r_morula");
        model.param().set("sigma_8",    "3.216",  "1.5 * r_8cell");
        model.param().set("sigma_4",    "2.0");

        // Control strengths
        model.param().set("gamma_acc",      "1.0");
        model.param().set("gamma_closure",  "1.0");
        model.param().set("gamma_zga",      "1.0");
        model.param().set("gamma_re",       "1.0");
        model.param().set("gamma_de",       "1.0");
        model.param().set("gamma_wrong",    "-1.0", "Wrong direction");

        // Operator time
        model.param().set("tau_end",    "6.0");

        // ── Component: Global ODEs ─────────────────────────────────────────────
        model.modelNode().create("comp1", true);

        // Global ODEs and DAEs
        model.physics().create("ge", "GlobalEquations", "comp1");
        model.physics("ge").label("DMR State Space ODE");

        // State variables: z1(tau), z2(tau)
        model.physics("ge").feature("ge1").set("name", new String[]{"z1", "z2"});
        model.physics("ge").feature("ge1").set("equation",
            new String[]{
                "z1t - (FK1 + g_zga*Fzga1 + g_entry*Fentry1 + g_exit*Fexit1)",
                "z2t - (FK2 + g_zga*Fzga2 + g_entry*Fentry2 + g_exit*Fexit2)"
            });
        model.physics("ge").feature("ge1").set("initialValueU",
            new String[]{"z1_oocyte", "z2_oocyte"});
        model.physics("ge").feature("ge1").set("initialValueUt",
            new String[]{"0", "0"});

        // ── Variables ─────────────────────────────────────────────────────────
        model.variable().create("var1");
        model.variable("var1").model("comp1");

        // Gaussian envelopes
        model.variable("var1").set("rM2",
            "(z1-z1_morula)^2 + (z2-z2_morula)^2");
        model.variable("var1").set("r82",
            "(z1-z1_8cell)^2 + (z2-z2_8cell)^2");
        model.variable("var1").set("r42",
            "(z1-z1_4cell)^2 + (z2-z2_4cell)^2");
        model.variable("var1").set("rB2",
            "(z1-z1_blast)^2 + (z2-z2_blast)^2");

        model.variable("var1").set("gM",
            "exp(-rM2/(2*sigma_M^2))");
        model.variable("var1").set("g8",
            "exp(-r82/(2*sigma_8^2))");
        model.variable("var1").set("g4",
            "exp(-r42/(2*sigma_4^2))");
        model.variable("var1").set("gB",
            "exp(-rB2/(2*sigma_M^2))");

        // Time gate functions (smooth Heaviside)
        // ZGA: tau in [3,4] (4-cell to 8-cell)
        model.variable("var1").set("chi_zga",
            "flc2hs(tau-3,0.3)*flc2hs(4-tau,0.3)");
        // Entry: tau in [4,5] (8-cell to morula)
        model.variable("var1").set("chi_entry",
            "flc2hs(tau-4,0.3)*flc2hs(5-tau,0.3)");
        // Exit: tau in [5,6] (morula to blastocyst)
        model.variable("var1").set("chi_exit",
            "flc2hs(tau-5,0.3)*flc2hs(6-tau,0.3)");

        // F_K: methylation-only baseline field
        model.variable("var1").set("FK1",
            "-lambda_K*(z1-z1_Katt)");
        model.variable("var1").set("FK2",
            "-lambda_K*(z2-z2_Katt)");

        // F_ZGA: ZGA reconstruction field (4->8)
        // Direction: 4-cell center -> 8-cell center
        model.variable("var1").set("vzga1",
            "(z1_8cell-z1_4cell)/sqrt((z1_8cell-z1_4cell)^2+(z2_8cell-z2_4cell)^2)");
        model.variable("var1").set("vzga2",
            "(z2_8cell-z2_4cell)/sqrt((z1_8cell-z1_4cell)^2+(z2_8cell-z2_4cell)^2)");
        model.variable("var1").set("Fzga1",
            "chi_zga*gamma_zga*g4*vzga1");
        model.variable("var1").set("Fzga2",
            "chi_zga*gamma_zga*g4*vzga2");

        // F_entry: accessibility-gated reset entry (8->morula)
        // Direction: 8-cell center -> morula center
        model.variable("var1").set("ventry1",
            "(z1_morula-z1_8cell)/sqrt((z1_morula-z1_8cell)^2+(z2_morula-z2_8cell)^2)");
        model.variable("var1").set("ventry2",
            "(z2_morula-z2_8cell)/sqrt((z1_morula-z1_8cell)^2+(z2_morula-z2_8cell)^2)");
        model.variable("var1").set("Fentry1",
            "chi_entry*(gamma_acc+gamma_closure)*g8*ventry1");
        model.variable("var1").set("Fentry2",
            "chi_entry*(gamma_acc+gamma_closure)*g8*ventry2");

        // F_exit: H3K4me3/re-methylation reconstruction exit (morula->blast)
        // Direction: morula center -> blastocyst center
        model.variable("var1").set("vexit1",
            "(z1_blast-z1_morula)/sqrt((z1_blast-z1_morula)^2+(z2_blast-z2_morula)^2)");
        model.variable("var1").set("vexit2",
            "(z2_blast-z2_morula)/sqrt((z1_blast-z1_morula)^2+(z2_blast-z2_morula)^2)");
        model.variable("var1").set("Fexit1",
            "chi_exit*(gamma_re+gamma_de)*gM*vexit1");
        model.variable("var1").set("Fexit2",
            "chi_exit*(gamma_re+gamma_de)*gM*vexit2");

        // Control flags (set per scenario via parameters)
        model.variable("var1").set("g_zga",   "1");
        model.variable("var1").set("g_entry", "1");
        model.variable("var1").set("g_exit",  "1");

        // Basin occupancy indicators
        model.variable("var1").set("in_morula_basin",
            "if(sqrt(rM2)<r_morula,1,0)");
        model.variable("var1").set("in_8cell_basin",
            "if(sqrt(r82)<r_8cell,1,0)");
        model.variable("var1").set("in_blast_basin",
            "if(sqrt(rB2)<r_blast,1,0)");

        // Entry-exit duality (cosine between entry and exit velocity)
        model.variable("var1").set("entry_exit_dot",
            "ventry1*vexit1 + ventry2*vexit2");

        // ── Study ─────────────────────────────────────────────────────────────
        model.study().create("std1");
        model.study("std1").label("Full Control");
        model.study("std1").create("time", "Transient");
        model.study("std1").feature("time").set("tlist", "range(0,0.1,6)");
        model.study("std1").feature("time").set("usertol", true);
        model.study("std1").feature("time").set("rtol", "1e-4");

        // ── Results ───────────────────────────────────────────────────────────
        model.result().create("pg1", "PlotGroup1D");
        model.result("pg1").label("Trajectory z1 vs tau");
        model.result("pg1").create("glob1", "Global");
        model.result("pg1").feature("glob1").set("expr", new String[]{"z1"});
        model.result("pg1").feature("glob1").set("descr", new String[]{"z1(tau)"});

        model.result().create("pg2", "PlotGroup1D");
        model.result("pg2").label("Trajectory z2 vs tau");
        model.result("pg2").create("glob1", "Global");
        model.result("pg2").feature("glob1").set("expr", new String[]{"z2"});

        model.result().create("pg3", "PlotGroup1D");
        model.result("pg3").label("Basin Occupancy vs tau");
        model.result("pg3").create("glob1", "Global");
        model.result("pg3").feature("glob1").set("expr",
            new String[]{"in_morula_basin", "in_8cell_basin", "in_blast_basin"});
        model.result("pg3").feature("glob1").set("descr",
            new String[]{"Morula basin", "8-cell basin", "Blast basin"});

        return model;
    }

    public static void main(String[] args) throws Exception {
        Model model = run();
        model.save("E:\\progress_comsol_analysis\\CEEF_ModelA.mph");
        System.out.println("Model saved: CEEF_ModelA.mph");
        ModelUtil.disconnect();
    }
}
```
