## Supplemental code for "An entropy-based diagnostic framework for characterizing methylation state dynamics during preimplantation development": export_comsol_screenshots.java

```
/*
 * COMSOL 6.4 Java API — Export publication screenshots from CEEF .mph files
 * Run with:
 *   comsolbatch -inputfile export_comsol_screenshots.java
 * or via COMSOL Script runner
 */
import com.comsol.model.*;
import com.comsol.model.util.*;
import java.io.*;

public class export_comsol_screenshots {

    public static void main(String[] args) throws Exception {
        String outDir = "E:\\progress_comsol_analysis\\figures_comsol";
        new File(outDir).mkdirs();

        String[] models = {
            "CEEF_final_baseline_only",
            "CEEF_final_full_control",
            "CEEF_final_wrong_exit"
        };

        for (String mname : models) {
            String mphPath = "E:\\progress_comsol_analysis\\" + mname + ".mph";
            System.out.println("Loading: " + mphPath);

            Model model = ModelUtil.load(mname, mphPath);

            // List available plots
            for (String pg : model.result().toString().split("\n")) {
                System.out.println("  result: " + pg);
            }

            // Try to export each result group
            String[] resultTags = model.result().tags();
            for (String tag : resultTags) {
                try {
                    String pngPath = outDir + "\\" + mname + "_" + tag + ".png";
                    model.result(tag).run();
                    model.result(tag).exportImage(pngPath, "png",
                        new String[]{"size","width","1600",
                                     "size","height","1200",
                                     "antialias","on",
                                     "fontsize","18",
                                     "logo","off"});
                    System.out.println("  Exported: " + pngPath);
                } catch (Exception e) {
                    System.out.println("  Skip " + tag + ": " + e.getMessage());
                }
            }

            ModelUtil.remove(mname);
        }
        System.out.println("Done.");
    }
}
```
