## Supplemental code for "An entropy-based diagnostic framework for characterizing methylation state dynamics during preimplantation development": export_data.py

```
#!/usr/bin/env python
"""
Step 1: Export all COMSOL input data from DMR dynamics model.
Output: E:/progress_comsol_analysis/
"""
import numpy as np
import pandas as pd
from scipy import stats
from pathlib import Path
import json

OUT = Path("E:/progress_comsol_analysis")
OUT.mkdir(parents=True, exist_ok=True)

# ── Load core data ─────────────────────────────────────────────────────────────
TRAJ  = Path("E:/CSB_TRO_operator_time_DMR_dynamics_2026-05-25/results/CSB_TRO_DMR_stage_mean_trajectory.tsv")
RESID = Path("E:/CSB_TRO_operator_time_DMR_dynamics_2026-05-25/results/CSB_TRO_basin_residual_DMR_ranking.tsv")
LOADS = Path("E:/CSB_TRO_operator_time_DMR_dynamics_2026-05-25/results/CSB_TRO_latent_autonomous_scores.tsv")
STATE = Path("E:/CSB_TRO_operator_time_DMR_dynamics_2026-05-25/results/CSB_TRO_DMR_state_matrix.tsv")
META  = Path("E:/CSB_TRO_operator_time_DMR_dynamics_2026-05-25/results/CSB_TRO_DMR_metadata.tsv")

traj  = pd.read_csv(TRAJ, sep="\t")
resid = pd.read_csv(RESID, sep="\t")
scores= pd.read_csv(LOADS, sep="\t")
state = pd.read_csv(STATE, sep="\t", index_col=0)
meta  = pd.read_csv(META, sep="\t")

stage_means = {}
for s, g in traj.groupby("stage"):
    stage_means[s] = g.set_index("cluster_name")["mean_beta"].to_dict()

clusters = sorted(resid["cluster_name"].tolist())
mod_map  = resid.set_index("cluster_name")["module_id"].to_dict()
stages_ord = ["MII oocyte","zygote/PN","2-cell","4-cell","8-cell","morula","blastocyst"]

def svec(s):
    return np.array([stage_means.get(s,{}).get(c,np.nan) for c in clusters])

x_oocyte = svec("MII oocyte"); x_zygote = svec("zygote/PN")
x_2cell  = svec("2-cell");     x_4cell  = svec("4-cell")
x_8cell  = svec("8-cell");     x_morula = svec("morula")
x_blast  = svec("blastocyst")

alpha_op = 0.5611; bias_op = 0.0688

# ── PCA in 2D ─────────────────────────────────────────────────────────────────
# Use existing latent scores (PC1, PC2)
# Assign samples to stages
stage_vecs = {s: svec(s) for s in stages_ord}
sample_stages = {}
for sid in state.index:
    sv = state.loc[sid, clusters].values.astype(float)
    best_s = None; best_d = np.inf
    for s, sv2 in stage_vecs.items():
        v = np.isfinite(sv) & np.isfinite(sv2)
        if v.sum() < 10: continue
        d = float(np.sqrt(np.mean((sv[v]-sv2[v])**2)))
        if d < best_d: best_d = d; best_s = s
    sample_stages[sid] = best_s

scores["stage"] = scores["sample_id"].map(sample_stages)

# Stage centers in 2D latent space
stage_centers = {}
for s in stages_ord:
    sub = scores[scores["stage"]==s][["PC1","PC2"]].values
    if len(sub) > 0:
        stage_centers[s] = sub.mean(axis=0)

print("Stage centers (PC1, PC2):")
for s, c in stage_centers.items():
    print(f"  {s}: ({c[0]:.4f}, {c[1]:.4f})")

# ── 1. stage_centers_2d.csv ───────────────────────────────────────────────────
stage_labels = {
    "MII oocyte": "oocyte", "zygote/PN": "zygote",
    "2-cell": "2cell", "4-cell": "4cell",
    "8-cell": "8cell", "morula": "morula", "blastocyst": "blast"
}
rows = []
for s in stages_ord:
    if s in stage_centers:
        c = stage_centers[s]
        rows.append({"stage": stage_labels[s], "z1": c[0], "z2": c[1]})
pd.DataFrame(rows).to_csv(OUT/"stage_centers_2d.csv", index=False)
print(f"\nSaved stage_centers_2d.csv")

# ── 2. stage_vectors_2d.csv ───────────────────────────────────────────────────
# Velocity vectors in 2D latent space
# Project DMR correction vectors to 2D using PCA loadings
# Use PC1/PC2 loadings from existing latent scores
# Approximate: project stage mean differences to 2D

# Compute stage mean differences in DMR space, then project to 2D
# Use simple correlation: PC1 ~ mean methylation, PC2 ~ variance
# Better: use the actual PC scores per stage

transitions = [
    ("oocyte_to_zygote", "MII oocyte", "zygote/PN"),
    ("zygote_to_2cell",  "zygote/PN",  "2-cell"),
    ("2cell_to_4cell",   "2-cell",     "4-cell"),
    ("4cell_to_8cell",   "4-cell",     "8-cell"),
    ("8cell_to_morula",  "8-cell",     "morula"),
    ("morula_to_blast",  "morula",     "blastocyst"),
]

vec_rows = []
for label, sf, st in transitions:
    if sf in stage_centers and st in stage_centers:
        cf = stage_centers[sf]; ct = stage_centers[st]
        vx = ct[0] - cf[0]; vy = ct[1] - cf[1]
        mag = np.sqrt(vx**2 + vy**2)
        vec_rows.append({"transition": label, "vx": vx, "vy": vy,
                          "vx_norm": vx/mag if mag>0 else 0,
                          "vy_norm": vy/mag if mag>0 else 0,
                          "magnitude": mag})
pd.DataFrame(vec_rows).to_csv(OUT/"stage_vectors_2d.csv", index=False)
print("Saved stage_vectors_2d.csv")

# ── 3. module_vectors_2d.csv ─────────────────────────────────────────────────
# Project module correction vectors to 2D
# Use the correlation between DMR corrections and PC scores

# Compute correction vectors for each module branch
c_diag_entry = x_morula - (alpha_op*x_8cell + bias_op)
c_diag_exit  = x_blast  - (alpha_op*x_morula + bias_op)
c_diag_zga   = x_8cell  - (alpha_op*x_4cell + bias_op)

# Module masks
closure_mask = np.array([mod_map.get(c,"?") in ["M01","M05","M12"] for c in clusters])
access_mask  = np.array([mod_map.get(c,"?") in ["M02","M10"] for c in clusters])
remeth_mask  = np.array([mod_map.get(c,"?") in ["M00","M13"] for c in clusters])
demeth_mask  = np.array([mod_map.get(c,"?") in ["M02","M06"] for c in clusters])
zga_mask     = np.array([mod_map.get(c,"?") in ["M01","M08"] for c in clusters])

# Project to 2D: use correlation with PC1 and PC2 scores
# For each DMR, we have its PC1 and PC2 loading
# Approximate: use the stage-level PC scores to project

# Simple projection: weighted sum of correction * PC score
def project_correction_to_2d(correction, mask):
    """Project masked correction vector to 2D latent space."""
    c_masked = np.where(mask, correction, 0.0)
    c_masked = np.where(np.isfinite(c_masked), c_masked, 0.0)
    # Use correlation with PC1 and PC2 of stage scores
    # PC1 direction: high methylation -> low PC1 (negative correlation)
    # Approximate by computing how this correction moves the stage center
    # Use the actual PC scores: project correction onto PC axes
    # PC1 loading: correlation between DMR methylation and PC1
    stage_pc1 = np.array([scores[scores["stage"]==s]["PC1"].mean() if s in scores["stage"].values else np.nan
                           for s in stages_ord])
    stage_pc2 = np.array([scores[scores["stage"]==s]["PC2"].mean() if s in scores["stage"].values else np.nan
                           for s in stages_ord])
    stage_meth = np.array([np.nanmean(svec(s)) for s in stages_ord])

    # Correlation: how does mean methylation change relate to PC1/PC2 change?
    v = np.isfinite(stage_pc1) & np.isfinite(stage_meth)
    r1, _ = stats.pearsonr(stage_meth[v], stage_pc1[v])
    r2, _ = stats.pearsonr(stage_meth[v], stage_pc2[v])

    # Project: correction mean -> PC1/PC2 change
    corr_mean = float(np.nanmean(c_masked))
    vx = corr_mean * r1
    vy = corr_mean * r2
    mag = np.sqrt(vx**2 + vy**2)
    return vx, vy, mag

module_branches = [
    ("closure_M01_M05_M12", closure_mask, c_diag_entry),
    ("access_M02_M10",      access_mask,  c_diag_entry),
    ("remeth_M00_M13",      remeth_mask,  c_diag_exit),
    ("demeth_M02_M06",      demeth_mask,  c_diag_exit),
    ("zga_M01_M08",         zga_mask,     c_diag_zga),
]

mod_rows = []
for label, mask, correction in module_branches:
    vx, vy, mag = project_correction_to_2d(correction, mask)
    mod_rows.append({"branch": label, "vx": vx, "vy": vy,
                      "vx_norm": vx/mag if mag>0 else 0,
                      "vy_norm": vy/mag if mag>0 else 0,
                      "magnitude": mag})
pd.DataFrame(mod_rows).to_csv(OUT/"module_vectors_2d.csv", index=False)
print("Saved module_vectors_2d.csv")

# ── 4. basin_masks_grid.csv ───────────────────────────────────────────────────
# Define basins in 2D latent space
morula_center = np.array([stage_centers["morula"][0], stage_centers["morula"][1]])
cell8_center  = np.array([stage_centers["8-cell"][0], stage_centers["8-cell"][1]])
blast_center  = np.array([stage_centers["blastocyst"][0], stage_centers["blastocyst"][1]])

# Compute radii from actual sample distributions
morula_pts = scores[scores["stage"]=="morula"][["PC1","PC2"]].values
cell8_pts  = scores[scores["stage"]=="8-cell"][["PC1","PC2"]].values
blast_pts  = scores[scores["stage"]=="blastocyst"][["PC1","PC2"]].values

r_morula = float(np.quantile(np.sqrt(np.sum((morula_pts-morula_center)**2,axis=1)), 0.90)) if len(morula_pts)>0 else 0.5
r_8cell  = float(np.quantile(np.sqrt(np.sum((cell8_pts-cell8_center)**2,axis=1)), 0.90)) if len(cell8_pts)>0 else 0.8
r_blast  = float(np.quantile(np.sqrt(np.sum((blast_pts-blast_center)**2,axis=1)), 0.90)) if len(blast_pts)>0 else 0.5

print(f"\nBasin radii: morula={r_morula:.4f}, 8cell={r_8cell:.4f}, blast={r_blast:.4f}")

# Create grid
z1_range = np.linspace(-4, 4, 81)
z2_range = np.linspace(-4, 4, 81)
Z1, Z2 = np.meshgrid(z1_range, z2_range)
z1_flat = Z1.flatten(); z2_flat = Z2.flatten()

dist_morula = np.sqrt((z1_flat-morula_center[0])**2 + (z2_flat-morula_center[1])**2)
dist_8cell  = np.sqrt((z1_flat-cell8_center[0])**2  + (z2_flat-cell8_center[1])**2)
dist_blast  = np.sqrt((z1_flat-blast_center[0])**2  + (z2_flat-blast_center[1])**2)

basin_df = pd.DataFrame({
    "z1": z1_flat, "z2": z2_flat,
    "in_morula_basin": (dist_morula <= r_morula).astype(int),
    "in_8cell_basin":  (dist_8cell  <= r_8cell).astype(int),
    "in_blast_basin":  (dist_blast  <= r_blast).astype(int),
    "dist_morula": dist_morula,
    "dist_8cell":  dist_8cell,
    "dist_blast":  dist_blast,
})
basin_df.to_csv(OUT/"basin_masks_grid.csv", index=False)
print("Saved basin_masks_grid.csv")

# ── 5. field_grid_2d.csv ─────────────────────────────────────────────────────
# Compute vector fields on the grid
# F_K: methylation-only baseline field (linear contraction toward K-attractor)
# F_entry: accessibility-gated reset entry field
# F_exit: H3K4me3/re-methylation reconstruction exit field

# K-attractor: where does K map to? Fixed point: z* = K*z* => z* = bias/(1-alpha)
# In 1D: x* = 0.0688/(1-0.5611) = 0.157
# In 2D latent space: approximate as the mean of all stage centers
all_centers = np.array([stage_centers[s] for s in stages_ord if s in stage_centers])
k_attractor = all_centers.mean(axis=0)

# F_K: linear field pointing toward K-attractor
lambda_K = 1 - alpha_op  # = 0.4389
FK_x = -lambda_K * (z1_flat - k_attractor[0])
FK_y = -lambda_K * (z2_flat - k_attractor[1])

# Gaussian envelope functions
sigma_M = r_morula * 1.5
sigma_8 = r_8cell  * 1.5
sigma_B = r_blast  * 1.5

gM = np.exp(-dist_morula**2 / (2*sigma_M**2))
g8 = np.exp(-dist_8cell**2  / (2*sigma_8**2))
gB = np.exp(-dist_blast**2  / (2*sigma_B**2))

# Entry field: push toward morula center
# Direction: from 8-cell center toward morula center
entry_dir = morula_center - cell8_center
entry_dir_norm = entry_dir / (np.linalg.norm(entry_dir) + 1e-12)
Fentry_x = g8 * entry_dir_norm[0]
Fentry_y = g8 * entry_dir_norm[1]

# Exit field: push from morula toward blastocyst
exit_dir = blast_center - morula_center
exit_dir_norm = exit_dir / (np.linalg.norm(exit_dir) + 1e-12)
Fexit_x = gM * exit_dir_norm[0]
Fexit_y = gM * exit_dir_norm[1]

# ZGA field: push from 4-cell toward 8-cell
zga_dir = cell8_center - np.array([stage_centers["4-cell"][0], stage_centers["4-cell"][1]])
zga_dir_norm = zga_dir / (np.linalg.norm(zga_dir) + 1e-12)
g4 = np.exp(-(np.sqrt((z1_flat-stage_centers["4-cell"][0])**2 + (z2_flat-stage_centers["4-cell"][1])**2))**2 / (2*sigma_8**2))
Fzga_x = g4 * zga_dir_norm[0]
Fzga_y = g4 * zga_dir_norm[1]

# Wrong exit field (for counterfactual)
Fwrong_exit_x = -Fexit_x  # reversed direction
Fwrong_exit_y = -Fexit_y

field_df = pd.DataFrame({
    "z1": z1_flat, "z2": z2_flat,
    "FK_x": FK_x, "FK_y": FK_y,
    "Fentry_x": Fentry_x, "Fentry_y": Fentry_y,
    "Fexit_x": Fexit_x, "Fexit_y": Fexit_y,
    "Fzga_x": Fzga_x, "Fzga_y": Fzga_y,
    "Fwrong_exit_x": Fwrong_exit_x, "Fwrong_exit_y": Fwrong_exit_y,
    "gM": gM, "g8": g8, "gB": gB,
})
field_df.to_csv(OUT/"field_grid_2d.csv", index=False)
print("Saved field_grid_2d.csv")

# ── 6. parameters.csv ─────────────────────────────────────────────────────────
params = {
    "alpha_op": alpha_op, "bias_op": bias_op,
    "lambda_K": lambda_K,
    "sigma_M": sigma_M, "sigma_8": sigma_8, "sigma_B": sigma_B,
    "r_morula": r_morula, "r_8cell": r_8cell, "r_blast": r_blast,
    "morula_z1": morula_center[0], "morula_z2": morula_center[1],
    "cell8_z1": cell8_center[0],   "cell8_z2": cell8_center[1],
    "blast_z1": blast_center[0],   "blast_z2": blast_center[1],
    "k_att_z1": k_attractor[0],    "k_att_z2": k_attractor[1],
    "tau_oocyte": 0, "tau_zygote": 1, "tau_2cell": 2,
    "tau_4cell": 3,  "tau_8cell": 4,  "tau_morula": 5, "tau_blast": 6,
    "D_diffusion": 0.02,
    "gamma_acc": 1.0, "gamma_closure": 1.0,
    "gamma_zga": 1.0, "gamma_re": 1.0, "gamma_de": 1.0,
}
pd.DataFrame([params]).to_csv(OUT/"parameters.csv", index=False)
print("Saved parameters.csv")

# ── 7. validation_targets.csv ─────────────────────────────────────────────────
targets = {
    "Occ_M_methylation_only": 0.044,
    "Occ_M_observed": 0.875,
    "Occ_M_correct_control": 0.956,
    "Occ_M_wrong_closure": 0.000,
    "D_M_duality": 0.699,
    "RMSE_correct": 0.018,
    "RMSE_wrong_remeth": 0.433,
    "ZGA_reset_rho": -0.439,
    "acc_morula_rho": 0.210,
    "k4me3_AUC": 0.792,
    "Occ_8cell_observed": 0.889,
    "Occ_2cell_in_8cell_basin": 0.300,
}
pd.DataFrame([targets]).to_csv(OUT/"validation_targets.csv", index=False)
print("Saved validation_targets.csv")

# ── 8. particle_initial.csv ───────────────────────────────────────────────────
# Initial particle positions: samples from zygote/2-cell stages
init_particles = []
for stage in ["zygote/PN", "2-cell"]:
    sub = scores[scores["stage"]==stage][["PC1","PC2"]].values
    for i, (z1, z2) in enumerate(sub):
        init_particles.append({"id": len(init_particles), "stage": stage,
                                 "z1": z1, "z2": z2, "weight": 1.0/len(sub)})
pd.DataFrame(init_particles).to_csv(OUT/"particle_initial.csv", index=False)
print("Saved particle_initial.csv")

# ── 9. sample_trajectories.csv ────────────────────────────────────────────────
# All sample positions in 2D latent space
sample_traj = scores[["sample_id","stage","PC1","PC2"]].copy()
sample_traj.columns = ["sample_id","stage","z1","z2"]
sample_traj.to_csv(OUT/"sample_trajectories.csv", index=False)
print("Saved sample_trajectories.csv")

# ── Summary ───────────────────────────────────────────────────────────────────
summary = {
    "date": "2026-05-30",
    "comsol_version": "6.4",
    "n_dmrs": len(clusters),
    "n_stages": len(stages_ord),
    "latent_dim": 2,
    "stage_centers": {stage_labels.get(s,s): stage_centers[s].tolist() for s in stages_ord if s in stage_centers},
    "basin_radii": {"morula": r_morula, "8cell": r_8cell, "blast": r_blast},
    "parameters": params,
    "validation_targets": targets,
    "files_exported": [
        "stage_centers_2d.csv", "stage_vectors_2d.csv", "module_vectors_2d.csv",
        "basin_masks_grid.csv", "field_grid_2d.csv", "parameters.csv",
        "validation_targets.csv", "particle_initial.csv", "sample_trajectories.csv"
    ]
}
with open(OUT/"data_export_summary.json","w",encoding="utf-8") as f:
    json.dump(summary, f, indent=2, ensure_ascii=False, default=str)

print(f"\nAll data exported to {OUT}/")
print(f"Total files: {len(list(OUT.iterdir()))}")
print("\nStage centers:")
for s in stages_ord:
    if s in stage_centers:
        c = stage_centers[s]
        print(f"  {s}: z1={c[0]:.3f}, z2={c[1]:.3f}")
```
