## Supplemental code for "An entropy-based diagnostic framework for characterizing methylation state dynamics during preimplantation development": figures_nature.py

```
#!/usr/bin/env python
"""
Nature/Cell-style publication figures for CEEF Phase A.
Clean white background, gradient trajectories, KDE density,
professional typography and layout.
"""
import numpy as np
import pandas as pd
import matplotlib
matplotlib.use("Agg")
import matplotlib.pyplot as plt
import matplotlib.patheffects as pe
from matplotlib.patches import FancyArrowPatch, Ellipse, FancyBboxPatch
from matplotlib.colors import LinearSegmentedColormap, Normalize
from matplotlib.cm import ScalarMappable
from matplotlib.gridspec import GridSpec, GridSpecFromSubplotSpec
from matplotlib.lines import Line2D
import matplotlib.ticker as ticker
from scipy.stats import gaussian_kde
from pathlib import Path
import json
import warnings
warnings.filterwarnings("ignore")

# ── Paths ──────────────────────────────────────────────────────────────────────
OUT  = Path("E:/progress_comsol_analysis")
DATA = Path("E:/CSB_TRO_operator_time_DMR_dynamics_2026-05-25/results")
FIG  = OUT / "figures_nature"
FIG.mkdir(exist_ok=True)

# ── Typography & style ────────────────────────────────────────────────────────
plt.rcParams.update({
    "font.family":        "DejaVu Sans",
    "font.size":          9,
    "axes.labelsize":     10,
    "axes.titlesize":     11,
    "axes.titleweight":   "bold",
    "axes.linewidth":     0.8,
    "xtick.labelsize":    8,
    "ytick.labelsize":    8,
    "xtick.major.width":  0.7,
    "ytick.major.width":  0.7,
    "xtick.major.size":   3,
    "ytick.major.size":   3,
    "legend.fontsize":    8,
    "legend.frameon":     True,
    "legend.framealpha":  0.9,
    "legend.edgecolor":   "#cccccc",
    "axes.spines.top":    False,
    "axes.spines.right":  False,
    "figure.dpi":         300,
    "savefig.dpi":        300,
    "savefig.bbox":       "tight",
    "savefig.facecolor":  "white",
})

# ── Palette ───────────────────────────────────────────────────────────────────
STAGE_COLORS = {
    "MII oocyte": "#F4A261",
    "zygote/PN":  "#E76F51",
    "2-cell":     "#A8DADC",
    "4-cell":     "#457B9D",
    "8-cell":     "#1D3557",
    "morula":     "#C0392B",
    "blastocyst": "#27AE60",
}
STAGE_LABELS = {
    "MII oocyte": "Oocyte",
    "zygote/PN":  "Zygote",
    "2-cell":     "2-cell",
    "4-cell":     "4-cell",
    "8-cell":     "8-cell",
    "morula":     "Morula",
    "blastocyst": "Blast.",
}
STAGES = ["MII oocyte","zygote/PN","2-cell","4-cell","8-cell","morula","blastocyst"]
STAGE_ORDER = {s: i for i,s in enumerate(STAGES)}

SCEN_COLORS = {
    "baseline_only": "#7F8C8D",
    "plus_zga":      "#2980B9",
    "plus_entry":    "#E67E22",
    "full_control":  "#C0392B",
    "wrong_exit":    "#8E44AD",
}
SCEN_LABELS = {
    "baseline_only": "Methylation-only",
    "plus_zga":      "+ZGA",
    "plus_entry":    "+Entry control",
    "full_control":  "Full control",
    "wrong_exit":    "Wrong exit",
}

# ── Load PCA data ──────────────────────────────────────────────────────────────
scores   = pd.read_csv(DATA/"CSB_TRO_latent_autonomous_scores.tsv", sep="\t")
traj_df  = pd.read_csv(DATA/"CSB_TRO_DMR_stage_mean_trajectory.tsv", sep="\t")
state    = pd.read_csv(DATA/"CSB_TRO_DMR_state_matrix.tsv", sep="\t", index_col=0)

stage_means = {}
for s, g in traj_df.groupby("stage"):
    stage_means[s] = g.set_index("cluster_name")["mean_beta"].to_dict()
clusters = sorted(traj_df["cluster_name"].unique().tolist())
stage_vecs = {s: np.array([stage_means.get(s,{}).get(c,np.nan) for c in clusters]) for s in STAGES}

sample_stages = {}
for sid in state.index:
    sv = state.loc[sid, clusters].values.astype(float)
    best_s, best_d = None, np.inf
    for s, sv2 in stage_vecs.items():
        v = np.isfinite(sv) & np.isfinite(sv2)
        if v.sum() < 10: continue
        d = float(np.sqrt(np.mean((sv[v]-sv2[v])**2)))
        if d < best_d: best_d = d; best_s = s
    sample_stages[sid] = best_s

scores["stage"] = scores["sample_id"].map(sample_stages)

# Clip extreme PC2 outliers (>3 SD from stage mean)
for s in STAGES:
    idx = scores["stage"] == s
    m, sd = scores.loc[idx, "PC2"].mean(), scores.loc[idx, "PC2"].std()
    scores.loc[idx & (np.abs(scores["PC2"]-m) > 3.5*sd), "stage"] = None
scores = scores.dropna(subset=["stage"])

CENTERS = {}
for s in STAGES:
    sub = scores[scores["stage"]==s][["PC1","PC2"]].values
    if len(sub) > 0:
        CENTERS[s] = sub.mean(axis=0)

# ── Map CEEF trajectories to PCA space ────────────────────────────────────────
ceef_centers = pd.read_csv(OUT/"stage_centers_2d_corrected.csv").set_index("stage")
anchor_map = {"8cell":"8-cell", "morula":"morula", "blast":"blastocyst",
              "4cell":"4-cell", "oocyte":"MII oocyte"}
src = np.array([ceef_centers.loc[k,["z1","z2"]].values
                for k in ["oocyte","8cell","morula","blast","4cell"]], dtype=float)
dst = np.array([CENTERS[anchor_map[k]]
                for k in ["oocyte","8cell","morula","blast","4cell"]], dtype=float)
A, _, _, _ = np.linalg.lstsq(np.hstack([src, np.ones((len(src),1))]), dst, rcond=None)

def ceef_to_pca(z1, z2):
    pts = np.column_stack([np.asarray(z1), np.asarray(z2), np.ones(len(np.asarray(z1)))])
    return pts @ A

trajs_raw = {}
for name in ["baseline_only","plus_zga","plus_entry","full_control","wrong_exit"]:
    fp = OUT / f"traj_final_{name}.csv"
    if fp.exists():
        trajs_raw[name] = pd.read_csv(fp)

trajs_pca = {}
for name, df in trajs_raw.items():
    xy = ceef_to_pca(df["z1"].values, df["z2"].values)
    trajs_pca[name] = pd.DataFrame({"t": df["t"].values, "PC1": xy[:,0], "PC2": xy[:,1]})

scale = np.linalg.norm(dst[2]-dst[1]) / np.linalg.norm(src[2]-src[1])
r_morula = 0.50 * scale
r_blast  = 0.60 * scale
r_8cell  = 0.80 * scale
C_morula = CENTERS["morula"]
C_blast  = CENTERS["blastocyst"]
C_8cell  = CENTERS["8-cell"]

with open(OUT/"scenario_results_final.json") as f:
    results = json.load(f)

# ── Helper functions ──────────────────────────────────────────────────────────
def plot_kde_background(ax, scores, stages=None, alpha=0.35, bw=0.4):
    """Per-stage KDE density fill."""
    if stages is None:
        stages = STAGES
    xmin, xmax = ax.get_xlim()
    ymin, ymax = ax.get_ylim()
    xx, yy = np.mgrid[xmin:xmax:80j, ymin:ymax:80j]
    pos = np.vstack([xx.ravel(), yy.ravel()])
    for s in stages:
        sub = scores[scores["stage"]==s][["PC1","PC2"]].values
        if len(sub) < 4: continue
        try:
            kde = gaussian_kde(sub.T, bw_method=bw)
            z = kde(pos).reshape(xx.shape)
            z = z / z.max()
            color = STAGE_COLORS[s]
            r,g,b = int(color[1:3],16)/255, int(color[3:5],16)/255, int(color[5:7],16)/255
            cmap = LinearSegmentedColormap.from_list("_", [(1,1,1,0),(r,g,b,alpha)])
            ax.contourf(xx, yy, z, levels=[0.15, 0.4, 0.7, 1.0],
                       cmap=cmap, zorder=1)
        except Exception:
            pass

def plot_stage_centers(ax, fontsize=8, marker_size=90):
    """Stage centers as diamonds with bold labels."""
    for s in STAGES:
        c = CENTERS[s]
        ax.scatter(c[0], c[1], s=marker_size, color=STAGE_COLORS[s],
                  edgecolors="white", linewidths=1.5, zorder=6, marker="D")
    # Labels with background
    offsets = {
        "MII oocyte": (8, 5), "zygote/PN": (8, -10), "2-cell": (8, 5),
        "4-cell": (-50, -12), "8-cell": (8, 5), "morula": (-52, 5),
        "blastocyst": (8, -12),
    }
    for s in STAGES:
        c = CENTERS[s]
        dx, dy = offsets.get(s, (8, 5))
        ax.annotate(STAGE_LABELS[s], (c[0], c[1]),
                   textcoords="offset points", xytext=(dx, dy),
                   fontsize=fontsize, fontweight="bold", color=STAGE_COLORS[s],
                   path_effects=[pe.withStroke(linewidth=2.5, foreground="white")],
                   zorder=10)

def gradient_trajectory(ax, x, y, color, lw=2.5, alpha_start=0.25, alpha_end=0.95, zorder=5):
    """Draw trajectory with opacity gradient (faint start → vivid end)."""
    n = len(x)
    for i in range(n-1):
        frac = i / (n-1)
        a = alpha_start + (alpha_end - alpha_start) * frac
        ax.plot(x[i:i+2], y[i:i+2], color=color, lw=lw, alpha=a,
               solid_capstyle="round", zorder=zorder)

def draw_arrow_on_traj(ax, x, y, color, frac=0.65, size=12):
    """Arrow in the middle of trajectory."""
    idx = int(len(x) * frac)
    dx = x[idx+1] - x[idx]
    dy = y[idx+1] - y[idx]
    ax.annotate("", xy=(x[idx]+dx*2, y[idx]+dy*2), xytext=(x[idx], y[idx]),
               arrowprops=dict(arrowstyle="-|>", color=color,
                               lw=1.5, mutation_scale=size),
               zorder=8)

def draw_basin_ellipse(ax, cx, cy, r, color, label="", alpha_fill=0.08):
    """Soft ellipse basin."""
    for ri, ai in [(r*1.0, alpha_fill), (r*0.6, alpha_fill*0.7)]:
        ell = Ellipse((cx,cy), width=ri*2, height=ri*2,
                     facecolor=color, alpha=ai, edgecolor=color,
                     linewidth=1.2, linestyle="--", zorder=1)
        ax.add_patch(ell)

def set_clean_axes(ax, xlabel="PC1", ylabel="PC2"):
    ax.set_xlabel(xlabel, fontsize=10, labelpad=3)
    ax.set_ylabel(ylabel, fontsize=10, labelpad=3)
    ax.grid(True, alpha=0.18, linewidth=0.5, zorder=0)
    ax.tick_params(labelsize=8)

def add_panel_label(ax, label, x=-0.12, y=1.04):
    ax.text(x, y, label, transform=ax.transAxes,
           fontsize=13, fontweight="bold", va="top", ha="left")

# ── Figure 1: Landscape overview  ─────────────────────────────────────────────
fig = plt.figure(figsize=(16, 6.5))
gs = GridSpec(1, 2, figure=fig, wspace=0.35, left=0.07, right=0.97,
             top=0.90, bottom=0.12)

fig.text(0.5, 0.97,
         "CEEF Phase A: DMR Operator-Time Dynamics — Preimplantation Epigenetic Landscape",
         ha="center", va="top", fontsize=12, fontweight="bold", color="#1a1a2e")

# ── Panel A: PCA landscape + all trajectories ─────────────────────────────────
ax = fig.add_subplot(gs[0])
ax.set_xlim(-7.5, 10.5); ax.set_ylim(-5.5, 8.0)

plot_kde_background(ax, scores, alpha=0.30, bw=0.35)

draw_basin_ellipse(ax, C_morula[0], C_morula[1], r_morula, STAGE_COLORS["morula"])
draw_basin_ellipse(ax, C_blast[0],  C_blast[1],  r_blast,  STAGE_COLORS["blastocyst"])
draw_basin_ellipse(ax, C_8cell[0],  C_8cell[1],  r_8cell,  STAGE_COLORS["8-cell"])

for name, df in trajs_pca.items():
    x, y = df["PC1"].values, df["PC2"].values
    gradient_trajectory(ax, x, y, SCEN_COLORS[name], lw=2.2)
    draw_arrow_on_traj(ax, x, y, SCEN_COLORS[name], frac=0.55)
    ax.scatter(x[-1], y[-1], s=55, color=SCEN_COLORS[name],
              edgecolors="white", linewidths=1, zorder=9, marker="o")

plot_stage_centers(ax, fontsize=8)

legend_handles = [Line2D([0],[0], color=SCEN_COLORS[n], lw=2.2,
                         label=SCEN_LABELS[n]) for n in SCEN_COLORS]
ax.legend(handles=legend_handles, loc="upper left", fontsize=7.5,
         framealpha=0.92, edgecolor="#cccccc", ncol=1)

set_clean_axes(ax, "PC1 (DMR latent axis 1)", "PC2 (DMR latent axis 2)")
ax.set_title("Epigenetic State Space with CEEF Trajectories", fontsize=11, fontweight="bold", pad=8)
add_panel_label(ax, "A")

# ── Panel B: Distance to morula over time ─────────────────────────────────────
ax2 = fig.add_subplot(gs[1])

ax2.axhspan(0, r_morula, alpha=0.10, color=STAGE_COLORS["morula"], zorder=0)
ax2.axhline(r_morula, color=STAGE_COLORS["morula"], lw=1.2, ls="--", alpha=0.7,
           label=f"Morula basin radius")
ax2.axvline(5.0, color="#999", lw=0.8, ls=":", alpha=0.6)
ax2.text(5.05, ax2.get_ylim()[1] if ax2.get_ylim()[1]>0 else 5,
         "τ=5", fontsize=7.5, color="#777", va="top")

for name, df in trajs_pca.items():
    dm = np.sqrt((df["PC1"]-C_morula[0])**2 + (df["PC2"]-C_morula[1])**2)
    ax2.plot(df["t"], dm, color=SCEN_COLORS[name], lw=2.0,
            label=SCEN_LABELS[name], alpha=0.9)
    # Endpoint dot
    ax2.scatter(df["t"].iloc[-1], dm.iloc[-1], s=40,
               color=SCEN_COLORS[name], zorder=7, edgecolors="white", lw=0.8)

# Re-add the axvline text after plotting
ax2.text(5.05, 0.15, "τ=5\n(morula)", fontsize=7.5, color="#777", va="bottom")

ax2.set_xlim(0, 6.2); ax2.set_ylim(0, None)
ax2.legend(loc="upper right", fontsize=7.5)
set_clean_axes(ax2, "Operator time  τ", "Distance to morula center (PCA)")
ax2.set_title("Morula Basin Convergence by Scenario", fontsize=11, fontweight="bold", pad=8)
add_panel_label(ax2, "B")

plt.savefig(FIG/"Fig1_landscape_overview.png", dpi=300)
(FIG/"Fig1_landscape_overview.pdf").unlink(missing_ok=True)
plt.savefig(FIG/"Fig1_landscape_overview.pdf")
plt.close()
print("Saved Fig1_landscape_overview")

# ── Figure 2: Three-panel failure→rescue ──────────────────────────────────────
fig = plt.figure(figsize=(15, 5.5))
gs = GridSpec(1, 3, figure=fig, wspace=0.30, left=0.06, right=0.97,
             top=0.88, bottom=0.13)

fig.text(0.5, 0.97,
         "Methylation-Only Failure vs Progressive Control Rescue",
         ha="center", va="top", fontsize=12, fontweight="bold", color="#1a1a2e")

scenarios3 = [
    ("baseline_only", "Methylation-only\n(FAILS: no morula entry)", "A"),
    ("plus_entry",    "+Entry control\n(PARTIAL: enters morula)", "B"),
    ("full_control",  "Full control\n(SUCCESS: morula + blast)", "C"),
]

for ax_idx, (name, title, panel_label) in enumerate(scenarios3):
    ax = fig.add_subplot(gs[ax_idx])
    ax.set_xlim(-7.5, 10.5); ax.set_ylim(-5.5, 8.0)

    plot_kde_background(ax, scores, alpha=0.22, bw=0.35)
    draw_basin_ellipse(ax, C_morula[0], C_morula[1], r_morula, STAGE_COLORS["morula"])
    draw_basin_ellipse(ax, C_blast[0],  C_blast[1],  r_blast,  STAGE_COLORS["blastocyst"])

    if name in trajs_pca:
        df = trajs_pca[name]
        x, y = df["PC1"].values, df["PC2"].values
        gradient_trajectory(ax, x, y, SCEN_COLORS[name], lw=2.8)
        draw_arrow_on_traj(ax, x, y, SCEN_COLORS[name], frac=0.52)
        ax.scatter(x[-1], y[-1], s=100, color=SCEN_COLORS[name],
                  edgecolors="white", linewidths=1.5, zorder=9, marker="*")

    plot_stage_centers(ax, fontsize=7.5, marker_size=70)

    res = results.get(name, {})
    in_m = res.get("in_morula", False)
    in_b = res.get("in_blast", False)
    dist  = res.get("dist_morula_t5", 99)

    status_color = "#27AE60" if (in_m and in_b) else ("#E67E22" if in_m else "#E74C3C")
    status_text = ("Morula [+]  Blast [+]" if (in_m and in_b)
                   else "Morula [+]  Blast [-]" if in_m
                   else "Morula [-]  Blast [-]")
    bbox_props = dict(boxstyle="round,pad=0.35", facecolor=status_color,
                     alpha=0.15, edgecolor=status_color, linewidth=1.2)
    ax.text(0.04, 0.97, status_text, transform=ax.transAxes,
           fontsize=8, va="top", color=status_color, fontweight="bold",
           bbox=bbox_props, zorder=11)
    ax.text(0.04, 0.84, f"d_morula = {dist:.2f}", transform=ax.transAxes,
           fontsize=7.5, va="top", color="#555")

    set_clean_axes(ax, "PC1", "PC2")
    ax.set_title(title, fontsize=9.5, fontweight="bold", pad=6, color="#1a1a2e")
    add_panel_label(ax, panel_label)

plt.savefig(FIG/"Fig2_rescue_progression.png", dpi=300)
(FIG/"Fig2_rescue_progression.pdf").unlink(missing_ok=True)
plt.savefig(FIG/"Fig2_rescue_progression.pdf")
plt.close()
print("Saved Fig2_rescue_progression")

# ── Figure 3: Morula as vector-field gate ─────────────────────────────────────
fig = plt.figure(figsize=(9, 7.5))
gs = GridSpec(1, 1, figure=fig, left=0.10, right=0.95, top=0.90, bottom=0.10)
ax = fig.add_subplot(gs[0])
ax.set_xlim(-7.5, 10.5); ax.set_ylim(-5.5, 8.0)

fig.text(0.5, 0.97,
         "Morula as Epigenetic Turning Gate\nEntry–Exit Anti-Alignment (cos = −0.876)",
         ha="center", va="top", fontsize=12, fontweight="bold", color="#1a1a2e")

plot_kde_background(ax, scores, alpha=0.28, bw=0.35)

draw_basin_ellipse(ax, C_morula[0], C_morula[1], r_morula*2.5,
                  STAGE_COLORS["morula"], alpha_fill=0.04)
draw_basin_ellipse(ax, C_morula[0], C_morula[1], r_morula,
                  STAGE_COLORS["morula"], alpha_fill=0.12)

# Entry vector: 8-cell → morula (thick red)
ax.annotate("", xy=C_morula, xytext=C_8cell,
           arrowprops=dict(arrowstyle="-|>", color="#E74C3C", lw=3,
                           mutation_scale=22, connectionstyle="arc3,rad=0.08"))
mid_entry = (C_8cell + C_morula) / 2
ax.text(mid_entry[0]+0.3, mid_entry[1]+0.5,
        "Entry vector\n(8-cell → Morula)",
        color="#E74C3C", fontsize=9, fontweight="bold",
        path_effects=[pe.withStroke(linewidth=2.5, foreground="white")])

# Exit vector: morula → blast (thick green)
ax.annotate("", xy=C_blast, xytext=C_morula,
           arrowprops=dict(arrowstyle="-|>", color="#27AE60", lw=3,
                           mutation_scale=22, connectionstyle="arc3,rad=-0.1"))
mid_exit = (C_morula + C_blast) / 2
ax.text(mid_exit[0]-1.8, mid_exit[1]-0.8,
        "Exit vector\n(Morula → Blast)",
        color="#27AE60", fontsize=9, fontweight="bold",
        path_effects=[pe.withStroke(linewidth=2.5, foreground="white")])

# Full control trajectory
if "full_control" in trajs_pca:
    df = trajs_pca["full_control"]
    gradient_trajectory(ax, df["PC1"].values, df["PC2"].values,
                       SCEN_COLORS["full_control"], lw=2.0, alpha_start=0.2, alpha_end=0.8)

plot_stage_centers(ax, fontsize=9, marker_size=100)

# Annotation box
info_text = ("Entry–Exit cosine = −0.876\n"
             "DMR-space cosine = −0.699\n"
             "Morula = geometric turning gate")
ax.text(0.97, 0.04, info_text, transform=ax.transAxes,
       fontsize=8.5, va="bottom", ha="right",
       bbox=dict(boxstyle="round,pad=0.5", facecolor="#FFEAA7",
                edgecolor="#FDCB6E", linewidth=1.2, alpha=0.95))

add_panel_label(ax, "A")
set_clean_axes(ax, "PC1 (DMR latent axis 1)", "PC2 (DMR latent axis 2)")

plt.savefig(FIG/"Fig3_morula_gate.png", dpi=300)
(FIG/"Fig3_morula_gate.pdf").unlink(missing_ok=True)
plt.savefig(FIG/"Fig3_morula_gate.pdf")
plt.close()
print("Saved Fig3_morula_gate")

# ── Figure 4: Counterfactual collapse ─────────────────────────────────────────
fig = plt.figure(figsize=(14, 5.5))
gs = GridSpec(1, 2, figure=fig, wspace=0.38, left=0.07, right=0.97,
             top=0.88, bottom=0.13)

fig.text(0.5, 0.97,
         "Wrong-Direction Counterfactual: Trajectory Collapse",
         ha="center", va="top", fontsize=12, fontweight="bold", color="#1a1a2e")

# Panel A: trajectories
ax = fig.add_subplot(gs[0])
ax.set_xlim(-7.5, 10.5); ax.set_ylim(-5.5, 8.0)

plot_kde_background(ax, scores, alpha=0.20, bw=0.35)
draw_basin_ellipse(ax, C_morula[0], C_morula[1], r_morula, STAGE_COLORS["morula"])
draw_basin_ellipse(ax, C_blast[0],  C_blast[1],  r_blast,  STAGE_COLORS["blastocyst"])

for name in ["full_control", "wrong_exit"]:
    if name in trajs_pca:
        df = trajs_pca[name]
        x, y = df["PC1"].values, df["PC2"].values
        gradient_trajectory(ax, x, y, SCEN_COLORS[name], lw=2.8)
        draw_arrow_on_traj(ax, x, y, SCEN_COLORS[name], frac=0.60)
        ax.scatter(x[-1], y[-1], s=100, color=SCEN_COLORS[name],
                  edgecolors="white", linewidths=1.5, zorder=9, marker="*")

plot_stage_centers(ax, fontsize=8)

legend_handles2 = [Line2D([0],[0], color=SCEN_COLORS[n], lw=2.5,
                          label=SCEN_LABELS[n])
                  for n in ["full_control","wrong_exit"]]
ax.legend(handles=legend_handles2, loc="upper left", fontsize=8.5, framealpha=0.92)
set_clean_axes(ax, "PC1", "PC2")
ax.set_title("Trajectory Comparison", fontsize=11, fontweight="bold", pad=8)
add_panel_label(ax, "A")

# Panel B: basin capture bar chart
ax2 = fig.add_subplot(gs[1])
scenarios4 = ["baseline_only","plus_zga","plus_entry","full_control","wrong_exit"]
labels4    = ["Baseline","+ZGA","+Entry","Full\nControl","Wrong\nExit"]

morula_vals = [1 if results.get(n,{}).get("in_morula",False) else 0 for n in scenarios4]
blast_vals  = [1 if results.get(n,{}).get("in_blast",False) else 0 for n in scenarios4]
dist_vals   = [results.get(n,{}).get("dist_morula_t5",5.0) for n in scenarios4]

x = np.arange(len(scenarios4))
bar_w = 0.32

b1 = ax2.bar(x - bar_w/2, morula_vals, bar_w,
            color=[STAGE_COLORS["morula"] if v else "#F5B7B1" for v in morula_vals],
            edgecolor="white", linewidth=0.8, zorder=3,
            label="Enters morula basin (τ=5)")
b2 = ax2.bar(x + bar_w/2, blast_vals, bar_w,
            color=[STAGE_COLORS["blastocyst"] if v else "#A9DFBF" for v in blast_vals],
            edgecolor="white", linewidth=0.8, zorder=3,
            label="Enters blast basin (τ=6)")

ax2.set_xticks(x); ax2.set_xticklabels(labels4, fontsize=8.5)
ax2.set_ylabel("Basin capture  (1 = Yes)", fontsize=10)
ax2.set_ylim(0, 1.45)
ax2.legend(fontsize=8, loc="upper right")
ax2.grid(True, axis="y", alpha=0.18, linewidth=0.5, zorder=0)
ax2.tick_params(labelsize=8)

for i, (d, m, b) in enumerate(zip(dist_vals, morula_vals, blast_vals)):
    col = "#27AE60" if (m and b) else ("#E67E22" if m else "#E74C3C")
    ax2.text(i, max(m, b) + 0.07, f"d={d:.2f}", ha="center",
            fontsize=7.5, color=col, fontweight="bold")

ax2.text(0.5, 0.02,
         "Wrong exit: trajectory diverges from blast basin",
         transform=ax2.transAxes, fontsize=8, ha="center", va="bottom", color="#7F8C8D",
         style="italic")

ax2.set_title("Basin Capture Summary", fontsize=11, fontweight="bold", pad=8)
add_panel_label(ax2, "B")
ax2.spines["top"].set_visible(False)
ax2.spines["right"].set_visible(False)

plt.savefig(FIG/"Fig4_counterfactual.png", dpi=300)
(FIG/"Fig4_counterfactual.pdf").unlink(missing_ok=True)
plt.savefig(FIG/"Fig4_counterfactual.pdf")
plt.close()
print("Saved Fig4_counterfactual")

print("\n" + "="*55)
print("ALL FIGURES SAVED TO:", FIG)
for f in sorted(FIG.iterdir()):
    print(f"  {f.name}")
```
