## Supplemental code for "An entropy-based diagnostic framework for characterizing methylation state dynamics during preimplantation development": figures_phase_b.py

```
#!/usr/bin/env python
"""
Phase B publication figures — aligned with Phase A style.
Same palette, layout, font, KDE-style backgrounds, gradient trajectories.

4 panels in a 2×2 composite:
  A: Entry-Exit scatter (DMR-level, cos=-0.876 line)
  B: Module duality lollipop (entry_exit_cosine per module)
  C: Curvature distribution (U-shape vs inverted-U by module)
  D: Accessibility coupling (morula accessibility vs geometry score)
"""
import numpy as np
import pandas as pd
import matplotlib
matplotlib.use("Agg")
import matplotlib.pyplot as plt
import matplotlib.patheffects as pe
from matplotlib.lines import Line2D
from matplotlib.patches import FancyArrowPatch
from matplotlib.gridspec import GridSpec
import warnings
warnings.filterwarnings("ignore")

BASE = "E:/实验进展5_27/CSB_TRO_2026-05-27_FINAL_DYNAMICS_PACKAGE/results_all"
OUT  = "E:/progress_comsol_analysis/figures_final"

# ── Style — identical to Phase A ─────────────────────────────────────────────
plt.rcParams.update({
    "font.family":"DejaVu Sans","font.size":9,
    "axes.labelsize":10,"axes.titlesize":10,"axes.titleweight":"bold",
    "axes.linewidth":0.8,"axes.spines.top":False,"axes.spines.right":False,
    "xtick.labelsize":8,"ytick.labelsize":8,
    "xtick.major.size":3,"ytick.major.size":3,
    "xtick.major.width":0.7,"ytick.major.width":0.7,
    "legend.fontsize":7.5,"figure.dpi":300,
    "savefig.dpi":300,"savefig.bbox":"tight","savefig.facecolor":"white",
})

# ── Palettes — same as Phase A ────────────────────────────────────────────────
BRANCH_COLORS = {
    "access":  "#C0392B",   # red  — accessibility-gated entry
    "closure": "#2980B9",   # blue — closure-gated exit
    "other":   "#7F8C8D",   # gray — background modules
}
BRANCH_LABELS = {
    "access":  "Accessibility (entry)",
    "closure": "Closure (exit)",
    "other":   "Background",
}

MODULE_COLORS = {
    "M02":"#C0392B","M05":"#2980B9","M12":"#8E44AD","M10":"#E67E22",
    "M01":"#27AE60","M06":"#F39C12","M14":"#1ABC9C","M00":"#95A5A6",
    "M13":"#BDC3C7","M07":"#7F8C8D","M09":"#AAB7B8","M04":"#717D7E",
    "M03":"#616A6B","M11":"#515A5A","M08":"#424949","M15":"#2C3E50",
}

def plabel(ax, lbl, x=-0.13, y=1.05):
    ax.text(x, y, lbl, transform=ax.transAxes,
            fontsize=13, fontweight="bold", va="top", ha="left")

# ── Load data ─────────────────────────────────────────────────────────────────
import os
df_dmr    = pd.read_csv(os.path.join(BASE, "CSB_TRO_2026-05-27_entry_exit_duality_metrics.tsv"), sep="\t")
df_module = pd.read_csv(os.path.join(BASE, "CSB_TRO_2026-05-27_entry_exit_module_duality.tsv"), sep="\t")
df_triad  = pd.read_csv(os.path.join(BASE, "CSB_TRO_2026-05-27_duality_accessibility_module_triad.tsv"), sep="\t")
df_curv   = pd.read_csv(os.path.join(BASE, "CSB_TRO_2026-05-27_entry_exit_curvature.tsv"), sep="\t")

# ══════════════════════════════════════════════════════════════════════════════
# FIGURE: 2×2 composite
# ══════════════════════════════════════════════════════════════════════════════
fig = plt.figure(figsize=(14, 12))
gs  = GridSpec(2, 2, figure=fig,
               hspace=0.42, wspace=0.38,
               left=0.09, right=0.97, top=0.93, bottom=0.08)

fig.text(0.5, 0.975,
         "CEEF Phase B: DMR Module Duality & Chromatin Accessibility",
         ha="center", va="top", fontsize=13, fontweight="bold", color="#1a1a2e")

# ── Panel A: Entry-Exit scatter ───────────────────────────────────────────────
ax = fig.add_subplot(gs[0, 0])

# Background: all DMRs in gray
ax.scatter(df_dmr["entry_change"], df_dmr["exit_change"],
           s=18, color="#BDC3C7", alpha=0.4, zorder=2, linewidths=0,
           label="All DMRs (n=156)")

# Highlight priority modules
for branch in ["access", "closure"]:
    sub = df_dmr[df_dmr["branch"] == branch]
    ax.scatter(sub["entry_change"], sub["exit_change"],
               s=40, color=BRANCH_COLORS[branch], alpha=0.85,
               zorder=4, linewidths=0.5, edgecolors="white",
               label=f"{BRANCH_LABELS[branch]} (n={len(sub)})")

# Anti-diagonal line (cos = -1 reference)
lim = 0.85
t = np.linspace(-lim, lim, 100)
ax.plot(t, -t, color="#E74C3C", lw=1.2, ls="--", alpha=0.6,
        label="cos = -1 (perfect anti-alignment)", zorder=1)
ax.axhline(0, color="#ccc", lw=0.6, zorder=0)
ax.axvline(0, color="#ccc", lw=0.6, zorder=0)

# Annotation: overall cosine
ax.text(0.97, 0.97,
        "Overall cos(entry,exit) = -0.876\n(morula = turning gate)",
        transform=ax.transAxes, fontsize=8, va="top", ha="right",
        bbox=dict(boxstyle="round,pad=0.4", facecolor="#FFEAA7",
                  edgecolor="#FDCB6E", linewidth=1, alpha=0.95))

ax.set_xlabel("Entry change  (Δβ: 8-cell → Morula)", fontsize=9)
ax.set_ylabel("Exit change  (Δβ: Morula → Blast)", fontsize=9)
ax.set_xlim(-0.9, 1.05); ax.set_ylim(-1.05, 1.05)
ax.legend(loc="lower right", fontsize=7, framealpha=0.9, edgecolor="#ccc")
ax.grid(True, alpha=0.15, linewidth=0.5, zorder=0)
ax.set_title("DMR Entry-Exit Anti-Alignment", fontsize=10, fontweight="bold", pad=6)
plabel(ax, "A")

# ── Panel B: Module duality lollipop ─────────────────────────────────────────
ax = fig.add_subplot(gs[0, 1])

# Sort by cosine (most negative first)
df_mod_sorted = df_module.sort_values("entry_exit_cosine")
y_pos = np.arange(len(df_mod_sorted))

for i, (_, row) in enumerate(df_mod_sorted.iterrows()):
    color = BRANCH_COLORS.get(row["branch"], "#7F8C8D")
    # Stem
    ax.plot([0, row["entry_exit_cosine"]], [i, i],
            color=color, lw=1.5, alpha=0.8, zorder=2)
    # Dot sized by n_dmr
    size = max(30, row["n_dmr"] * 4)
    ax.scatter(row["entry_exit_cosine"], i,
               s=size, color=color, zorder=4,
               edgecolors="white", linewidths=0.8)
    # Priority label
    if row["is_priority_residual_module"]:
        ax.text(row["entry_exit_cosine"] - 0.02, i,
                row["group"], ha="right", va="center",
                fontsize=7.5, fontweight="bold", color=color,
                path_effects=[pe.withStroke(linewidth=2, foreground="white")])

ax.axvline(0, color="#ccc", lw=0.8, zorder=0)
ax.axvline(-0.876, color="#E74C3C", lw=1.2, ls=":", alpha=0.7,
           label="Overall cos = -0.876")

ax.set_yticks(y_pos)
ax.set_yticklabels(df_mod_sorted["group"].tolist(), fontsize=7.5)
ax.set_xlabel("Entry-Exit cosine similarity", fontsize=9)
ax.set_xlim(-1.05, 0.15)
ax.grid(True, axis="x", alpha=0.15, linewidth=0.5, zorder=0)

# Legend for branches
handles = [Line2D([0],[0], color=BRANCH_COLORS[b], lw=2,
                  label=BRANCH_LABELS[b]) for b in ["access","closure","other"]]
handles.append(Line2D([0],[0], color="#E74C3C", lw=1.2, ls=":",
                       label="Overall cos = -0.876"))
ax.legend(handles=handles, loc="lower right", fontsize=7, framealpha=0.9)
ax.set_title("Module Entry-Exit Duality\n(dot size = n DMRs, bold = priority)",
             fontsize=10, fontweight="bold", pad=6)
plabel(ax, "B")

# ── Panel C: Curvature distribution ──────────────────────────────────────────
ax = fig.add_subplot(gs[1, 0])

# Curvature = exit_change - entry_change (positive = U-shape reset)
# Show distribution by branch
for branch in ["access", "closure", "other"]:
    sub = df_curv[df_curv["branch"] == branch]["curvature"]
    color = BRANCH_COLORS[branch]
    # KDE-style histogram
    from scipy.stats import gaussian_kde
    if len(sub) > 3:
        kde = gaussian_kde(sub, bw_method=0.4)
        x_range = np.linspace(df_curv["curvature"].min()-0.05,
                               df_curv["curvature"].max()+0.05, 200)
        y_kde = kde(x_range)
        ax.fill_between(x_range, y_kde, alpha=0.25, color=color)
        ax.plot(x_range, y_kde, color=color, lw=1.8,
                label=f"{BRANCH_LABELS[branch]} (n={len(sub)})")

ax.axvline(0, color="#555", lw=1.0, ls="--", alpha=0.7, label="curvature = 0")

# Annotate U-shape fraction
for branch in ["access", "closure"]:
    sub = df_curv[df_curv["branch"] == branch]
    frac_u = sub["is_u_shape"].mean()
    color = BRANCH_COLORS[branch]
    ax.text(0.97, 0.97 - (0.12 * ["access","closure"].index(branch)),
            f"{branch}: {frac_u:.0%} U-shape",
            transform=ax.transAxes, fontsize=8, va="top", ha="right",
            color=color, fontweight="bold",
            path_effects=[pe.withStroke(linewidth=2, foreground="white")])

ax.set_xlabel("Curvature  (exit − entry change)", fontsize=9)
ax.set_ylabel("Density", fontsize=9)
ax.legend(loc="upper left", fontsize=7, framealpha=0.9)
ax.grid(True, alpha=0.15, linewidth=0.5, zorder=0)
ax.set_title("DMR Curvature Distribution\n(U-shape = morula reset geometry)",
             fontsize=10, fontweight="bold", pad=6)
plabel(ax, "C")

# ── Panel D: Accessibility coupling ──────────────────────────────────────────
ax = fig.add_subplot(gs[1, 1])

# Scatter: morula accessibility vs signed duality, sized by n_dmr
# colored by branch
for branch in ["access", "closure", "other"]:
    sub = df_triad[df_triad["branch"] == branch]
    color = BRANCH_COLORS[branch]
    sizes = sub["n_dmr"] * 8 + 30
    sc = ax.scatter(sub["mean_morula_accessibility"],
                    sub["mean_signed_duality"],
                    s=sizes, color=color, alpha=0.85,
                    edgecolors="white", linewidths=0.8, zorder=4,
                    label=BRANCH_LABELS[branch])
    # Label priority modules
    for _, row in sub.iterrows():
        if row["module_id"] in ["M02","M05","M12","M10","M01"]:
            ax.annotate(row["module_id"],
                        (row["mean_morula_accessibility"], row["mean_signed_duality"]),
                        textcoords="offset points", xytext=(6, 3),
                        fontsize=7.5, fontweight="bold", color=color,
                        path_effects=[pe.withStroke(linewidth=2, foreground="white")],
                        zorder=8)

# Trend line
from numpy.polynomial import polynomial as P
x_all = df_triad["mean_morula_accessibility"].values
y_all = df_triad["mean_signed_duality"].values
coef = np.polyfit(x_all, y_all, 1)
x_fit = np.linspace(x_all.min(), x_all.max(), 100)
ax.plot(x_fit, np.polyval(coef, x_fit),
        color="#555", lw=1.2, ls="--", alpha=0.6, label="Linear trend")

# Annotation: rho from correlations file
df_corr = pd.read_csv(
    "E:/实验进展5_27/CSB_TRO_2026-05-27_FINAL_DYNAMICS_PACKAGE/results_all/"
    "CSB_TRO_2026-05-27_duality_accessibility_correlations.tsv", sep="\t")
rho_row = df_corr[df_corr["metric"]=="signed_duality"].iloc[0]
ax.text(0.97, 0.04,
        f"ρ = {rho_row['observed_spearman']:.3f}\np = {rho_row['empirical_p_ge_observed']:.3f}",
        transform=ax.transAxes, fontsize=8.5, va="bottom", ha="right",
        bbox=dict(boxstyle="round,pad=0.4", facecolor="#FFEAA7",
                  edgecolor="#FDCB6E", linewidth=1, alpha=0.95))

ax.set_xlabel("Mean morula accessibility (ATAC)", fontsize=9)
ax.set_ylabel("Mean signed duality score", fontsize=9)
ax.legend(loc="upper left", fontsize=7, framealpha=0.9)
ax.grid(True, alpha=0.15, linewidth=0.5, zorder=0)
ax.set_title("Chromatin Accessibility × Duality\n(dot size = n DMRs per module)",
             fontsize=10, fontweight="bold", pad=6)
plabel(ax, "D")

# ── Save ──────────────────────────────────────────────────────────────────────
import os
out_png = os.path.join(OUT, "CEEF_PhaseB_Figure.png")
out_pdf = os.path.join(OUT, "CEEF_PhaseB_Figure.pdf")
plt.savefig(out_png, dpi=300)
try:
    if os.path.exists(out_pdf):
        os.unlink(out_pdf)
    plt.savefig(out_pdf)
    print(f"Saved: {out_pdf}")
except Exception as e:
    print(f"PDF skipped: {e}")
plt.close()
print(f"Saved: {out_png}")
```
