## Supplemental code for "An entropy-based diagnostic framework for characterizing methylation state dynamics during preimplantation development": make_publication_figures.py

```
#!/usr/bin/env python
"""
Publication-quality figures for CEEF Phase A.
Strategy: Use PCA coordinates for natural trajectories,
overlay vector fields and annotations to show dynamics.
"""
import numpy as np
import pandas as pd
import matplotlib
matplotlib.use("Agg")
import matplotlib.pyplot as plt
import matplotlib.patches as mpatches
import matplotlib.patheffects as pe
from matplotlib.patches import FancyArrowPatch, Circle, FancyBboxPatch
from matplotlib.colors import LinearSegmentedColormap
from matplotlib.gridspec import GridSpec
import matplotlib.ticker as ticker
from pathlib import Path
import json

# ── Setup ──────────────────────────────────────────────────────────────────────
OUT = Path("E:/progress_comsol_analysis")
FIG = OUT / "figures_v2"
FIG.mkdir(exist_ok=True)

# Publication style
plt.rcParams.update({
    'font.family': 'Arial',
    'font.size': 10,
    'axes.labelsize': 11,
    'axes.titlesize': 12,
    'axes.titleweight': 'bold',
    'xtick.labelsize': 9,
    'ytick.labelsize': 9,
    'legend.fontsize': 9,
    'figure.dpi': 300,
    'savefig.dpi': 300,
    'savefig.bbox': 'tight',
    'axes.spines.top': False,
    'axes.spines.right': False,
    'axes.linewidth': 0.8,
    'xtick.major.width': 0.8,
    'ytick.major.width': 0.8,
})

# ── Load data ──────────────────────────────────────────────────────────────────
scores = pd.read_csv("E:/CSB_TRO_operator_time_DMR_dynamics_2026-05-25/results/CSB_TRO_latent_autonomous_scores.tsv", sep="\t")
traj = pd.read_csv("E:/CSB_TRO_operator_time_DMR_dynamics_2026-05-25/results/CSB_TRO_DMR_stage_mean_trajectory.tsv", sep="\t")
state = pd.read_csv("E:/CSB_TRO_operator_time_DMR_dynamics_2026-05-25/results/CSB_TRO_DMR_state_matrix.tsv", sep="\t", index_col=0)

stage_means = {}
for s, g in traj.groupby("stage"):
    stage_means[s] = g.set_index("cluster_name")["mean_beta"].to_dict()
clusters = sorted(traj["cluster_name"].unique().tolist())
STAGES = ["MII oocyte","zygote/PN","2-cell","4-cell","8-cell","morula","blastocyst"]
stage_vecs = {s: np.array([stage_means.get(s,{}).get(c,np.nan) for c in clusters]) for s in STAGES}

# Assign samples to stages
sample_stages = {}
for sid in state.index:
    sv = state.loc[sid, clusters].values.astype(float)
    best_s = None; best_d = np.inf
    for s, sv2 in stage_vecs.items():
        v = np.isfinite(sv) & np.isfinite(sv2)
        if v.sum() < 10: continue
        d = float(np.sqrt(np.mean((sv[v]-sv2[v])**2)))
        if d < best_d: best_d = d; best_s = s
    sample_stages[sid] = best_s

scores["stage"] = scores["sample_id"].map(sample_stages)

# Stage centers in PCA space
CENTERS = {}
for s in STAGES:
    sub = scores[scores["stage"]==s][["PC1","PC2"]].values
    if len(sub) > 0:
        CENTERS[s] = sub.mean(axis=0)

# Stage colors - publication quality palette
STAGE_COLORS = {
    "MII oocyte": "#E8A838",
    "zygote/PN":  "#D4A017",
    "2-cell":     "#7CB9E8",
    "4-cell":     "#4A90D9",
    "8-cell":     "#2E6DB4",
    "morula":     "#C0392B",
    "blastocyst": "#27AE60",
}
STAGE_LABELS = {
    "MII oocyte": "Oocyte",
    "zygote/PN":  "Zygote",
    "2-cell":     "2-cell",
    "4-cell":     "4-cell",
    "8-cell":     "8-cell",
    "morula":     "Morula",
    "blastocyst": "Blast.",
}

# Scenario colors
SCEN_COLORS = {
    "baseline_only": "#95A5A6",
    "plus_zga":      "#3498DB",
    "plus_entry":    "#E67E22",
    "full_control":  "#E74C3C",
    "wrong_exit":    "#8E44AD",
}
SCEN_LABELS = {
    "baseline_only": "Methylation-only (K)",
    "plus_zga":      "+ZGA reconstruction",
    "plus_entry":    "+Entry control",
    "full_control":  "Full control",
    "wrong_exit":    "Wrong exit direction",
}

# Load trajectories from corrected scenarios
trajs = {}
for name in ["baseline_only","plus_zga","plus_entry","full_control","wrong_exit"]:
    fp = OUT / ("traj_final_" + name + ".csv")
    if fp.exists():
        df = pd.read_csv(fp)
        # Convert from (e1,e2) back to PCA-like space for visualization
        # We'll use the corrected trajectories but display in a cleaner way
        trajs[name] = df

# Load scenario results
with open(OUT/"scenario_results_final.json") as f:
    results = json.load(f)

print("Data loaded successfully")
print(f"Stages: {list(CENTERS.keys())}")

# ── Map CEEF (e1,e2) trajectories into PCA space ──────────────────────────────
# e1,e2 stage centers from the simulation
ceef_centers = pd.read_csv(OUT/"stage_centers_2d_corrected.csv").set_index("stage")
ceef_key_map = {"8cell": "8-cell", "morula": "morula", "blast": "blastocyst"}
# Build affine from 3 anchor points: 8cell->morula->blast
src = np.array([ceef_centers.loc[k, ["z1","z2"]].values for k in ["8cell","morula","blast"]], dtype=float)
dst = np.array([CENTERS[ceef_key_map[k]] for k in ["8cell","morula","blast"]], dtype=float)
# Least-squares affine: dst = src @ A + b
A, res, rank, sv = np.linalg.lstsq(
    np.hstack([src, np.ones((3,1))]),
    dst, rcond=None)

def ceef_to_pca(z1, z2):
    pts = np.column_stack([np.asarray(z1), np.asarray(z2), np.ones(len(np.asarray(z1)))])
    return pts @ A

trajs_pca = {}
for name, df in trajs.items():
    xy = ceef_to_pca(df["z1"].values, df["z2"].values)
    trajs_pca[name] = pd.DataFrame({"t": df["t"].values, "PC1": xy[:,0], "PC2": xy[:,1]})

# Morula/blast basin radii in PCA space (scale factor from e1 unit to PC1 unit)
scale = float(np.linalg.norm(dst[1]-dst[0]) / np.linalg.norm(src[1]-src[0]))
r_morula_pca = 0.50 * scale
r_blast_pca  = 0.60 * scale
r_8cell_pca  = 0.80 * scale

C_morula = CENTERS["morula"]
C_blast  = CENTERS["blastocyst"]
C_8cell  = CENTERS["8-cell"]

# Results
with open(OUT/"scenario_results_final.json") as f:
    results = json.load(f)

# ── Helper: draw background scatter + stage centers ───────────────────────────
def draw_background(ax, show_scatter=True):
    if show_scatter:
        for s in STAGES:
            sub = scores[scores["stage"]==s][["PC1","PC2"]].values
            ax.scatter(sub[:,0], sub[:,1], s=18, color=STAGE_COLORS[s],
                       alpha=0.35, zorder=2, linewidths=0)
    for s in STAGES:
        c = CENTERS[s]
        ax.scatter(c[0], c[1], s=90, color=STAGE_COLORS[s],
                   edgecolors="white", linewidths=1.2, zorder=5, marker="D")
        ax.annotate(STAGE_LABELS[s], (c[0], c[1]),
                    textcoords="offset points", xytext=(6, 4),
                    fontsize=8, fontweight="bold", color=STAGE_COLORS[s])

def draw_basins(ax):
    for cx, cy, r, color in [
        (C_morula[0], C_morula[1], r_morula_pca, STAGE_COLORS["morula"]),
        (C_blast[0],  C_blast[1],  r_blast_pca,  STAGE_COLORS["blastocyst"]),
        (C_8cell[0],  C_8cell[1],  r_8cell_pca,  STAGE_COLORS["8-cell"]),
    ]:
        circle = Circle((cx, cy), r, fill=True, facecolor=color, alpha=0.13,
                        edgecolor=color, linewidth=1.5, linestyle="--")
        ax.add_patch(circle)

# ── Figure 1: Full trajectory comparison ──────────────────────────────────────
fig, axes = plt.subplots(1, 2, figsize=(14, 6))
fig.suptitle("CEEF Phase A: DMR Operator-Time Dynamics\nFull Preimplantation Trajectory",
             fontsize=13, fontweight="bold")

ax = axes[0]
ax.set_title("Trajectory in DMR PCA Space (PC1, PC2)", fontsize=11)
draw_basins(ax)
draw_background(ax, show_scatter=True)

for name, df in trajs_pca.items():
    ax.plot(df["PC1"], df["PC2"], color=SCEN_COLORS[name], linewidth=2,
            label=SCEN_LABELS[name], alpha=0.85, zorder=4)
    ax.scatter(df["PC1"].iloc[-1], df["PC2"].iloc[-1],
               s=50, color=SCEN_COLORS[name], marker="o", zorder=6)

ax.set_xlabel("PC1 (DMR latent axis 1)", fontsize=10)
ax.set_ylabel("PC2 (DMR latent axis 2)", fontsize=10)
ax.legend(loc="upper right", fontsize=7, framealpha=0.9)
ax.grid(True, alpha=0.25)
ax.set_aspect("equal")

# Right panel: distance to morula over time (in PCA space)
ax2 = axes[1]
ax2.set_title("Distance to Morula Center vs Operator Time", fontsize=11)
ax2.axhline(y=r_morula_pca, color=STAGE_COLORS["morula"], linestyle="--", linewidth=1.5,
            label=f"Morula basin radius (r={r_morula_pca:.2f})")
ax2.axvline(x=5.0, color="gray", linestyle=":", linewidth=1, alpha=0.7)
ax2.text(5.05, 0.05, "τ=5\n(morula)", fontsize=8, color="gray")

for name, df in trajs_pca.items():
    dm = np.sqrt((df["PC1"]-C_morula[0])**2 + (df["PC2"]-C_morula[1])**2)
    ax2.plot(df["t"], dm, color=SCEN_COLORS[name], linewidth=2,
             label=SCEN_LABELS[name], alpha=0.85)

ax2.set_xlabel("Operator time τ", fontsize=10)
ax2.set_ylabel("Distance to morula center (PCA)", fontsize=10)
ax2.legend(loc="upper right", fontsize=7, framealpha=0.9)
ax2.set_xlim(0, 6)
ax2.grid(True, alpha=0.25)

plt.tight_layout()
plt.savefig(FIG/"Fig_C1_full_trajectory.png", dpi=200, bbox_inches="tight")
(FIG/"Fig_C1_full_trajectory.pdf").unlink(missing_ok=True)
plt.savefig(FIG/"Fig_C1_full_trajectory.pdf", bbox_inches="tight")
plt.close()
print("Saved Fig_C1_full_trajectory")

# ── Figure 2: Baseline failure vs control rescue ───────────────────────────────
fig, axes = plt.subplots(1, 3, figsize=(15, 5))
fig.suptitle("CEEF: Methylation-Only Failure vs Control Rescue",
             fontsize=13, fontweight="bold")

for ax_idx, (name, title) in enumerate([
    ("baseline_only", "Methylation-only (K)\nOcc_M = 0 (FAILS)"),
    ("plus_entry",    "+Entry control\nOcc_M > 0 (PARTIAL)"),
    ("full_control",  "Full control\nOcc_M > 0, Occ_B > 0 (SUCCESS)"),
]):
    ax = axes[ax_idx]
    ax.set_title(title, fontsize=10)
    for cx, cy, r, color in [
        (C_morula[0], C_morula[1], r_morula_pca, STAGE_COLORS["morula"]),
        (C_blast[0],  C_blast[1],  r_blast_pca,  STAGE_COLORS["blastocyst"]),
    ]:
        circle = Circle((cx, cy), r, fill=True, facecolor=color, alpha=0.15,
                        edgecolor=color, linewidth=2)
        ax.add_patch(circle)
    draw_background(ax, show_scatter=(ax_idx == 0))

    if name in trajs_pca:
        df = trajs_pca[name]
        ax.plot(df["PC1"], df["PC2"], color=SCEN_COLORS[name], linewidth=2.5, zorder=4)
        mid = len(df)//2
        ax.annotate("", xy=(df["PC1"].iloc[mid+1], df["PC2"].iloc[mid+1]),
                    xytext=(df["PC1"].iloc[mid], df["PC2"].iloc[mid]),
                    arrowprops=dict(arrowstyle="->", color=SCEN_COLORS[name], lw=2))
        ax.scatter(df["PC1"].iloc[-1], df["PC2"].iloc[-1],
                   s=80, color=SCEN_COLORS[name], marker="*", zorder=6)

    res = results.get(name, {})
    if "error" not in res:
        in_m = res.get("in_morula", False)
        in_b = res.get("in_blast", False)
        ax.text(0.05, 0.97, "[+] Morula" if in_m else "[-] Morula",
                transform=ax.transAxes, fontsize=9, va="top",
                color=STAGE_COLORS["morula"] if in_m else "#E74C3C", fontweight="bold")
        ax.text(0.05, 0.89, "[+] Blast" if in_b else "[-] Blast",
                transform=ax.transAxes, fontsize=9, va="top",
                color=STAGE_COLORS["blastocyst"] if in_b else "#E74C3C", fontweight="bold")

    ax.set_xlabel("PC1", fontsize=9)
    ax.set_ylabel("PC2", fontsize=9)
    ax.grid(True, alpha=0.25)
    ax.set_aspect("equal")

plt.tight_layout()
plt.savefig(FIG/"Fig_C2_baseline_vs_rescue.png", dpi=200, bbox_inches="tight")
(FIG/"Fig_C2_baseline_vs_rescue.pdf").unlink(missing_ok=True)
plt.savefig(FIG/"Fig_C2_baseline_vs_rescue.pdf", bbox_inches="tight")
plt.close()
print("Saved Fig_C2_baseline_vs_rescue")

# ── Figure 3: Entry-exit vector reversal ──────────────────────────────────────
fig, ax = plt.subplots(1, 1, figsize=(8, 7))
ax.set_title("Morula as Vector-Field Turning Gate\nEntry-Exit Anti-Alignment (cos = -0.876)",
             fontsize=12, fontweight="bold")

circle = Circle((C_morula[0], C_morula[1]), r_morula_pca, fill=True,
                facecolor=STAGE_COLORS["morula"], alpha=0.2,
                edgecolor=STAGE_COLORS["morula"], linewidth=2)
ax.add_patch(circle)
draw_background(ax, show_scatter=True)

ax.annotate("", xy=(C_morula[0], C_morula[1]),
            xytext=(C_8cell[0], C_8cell[1]),
            arrowprops=dict(arrowstyle="-|>", color="#FF6B6B", lw=3, mutation_scale=20))
ax.text((C_8cell[0]+C_morula[0])/2 + 0.15, (C_8cell[1]+C_morula[1])/2 + 0.15,
        "Entry\n(8-cell→Morula)", color="#FF6B6B", fontsize=9, fontweight="bold")

ax.annotate("", xy=(C_blast[0], C_blast[1]),
            xytext=(C_morula[0], C_morula[1]),
            arrowprops=dict(arrowstyle="-|>", color="#27AE60", lw=3, mutation_scale=20))
ax.text((C_morula[0]+C_blast[0])/2 + 0.15, (C_morula[1]+C_blast[1])/2 - 0.2,
        "Exit\n(Morula→Blast)", color="#27AE60", fontsize=9, fontweight="bold")

if "full_control" in trajs_pca:
    df = trajs_pca["full_control"]
    ax.plot(df["PC1"], df["PC2"], color=SCEN_COLORS["full_control"], linewidth=2,
            alpha=0.7, linestyle="--", label="Full control trajectory")

ax.text(0.05, 0.05,
        "Entry-Exit cosine = -0.876\n(DMR space: -0.699)\nMorula = geometric turning gate",
        transform=ax.transAxes, fontsize=9, va="bottom",
        bbox=dict(boxstyle="round", facecolor="lightyellow", alpha=0.8))

ax.set_xlabel("PC1 (DMR latent axis 1)", fontsize=10)
ax.set_ylabel("PC2 (DMR latent axis 2)", fontsize=10)
ax.grid(True, alpha=0.25)
ax.set_aspect("equal")
ax.legend(fontsize=9)

plt.tight_layout()
plt.savefig(FIG/"Fig_C3_entry_exit_reversal.png", dpi=200, bbox_inches="tight")
(FIG/"Fig_C3_entry_exit_reversal.pdf").unlink(missing_ok=True)
plt.savefig(FIG/"Fig_C3_entry_exit_reversal.pdf", bbox_inches="tight")
plt.close()
print("Saved Fig_C3_entry_exit_reversal")

# ── Figure 4: Wrong-direction counterfactual collapse ─────────────────────────
fig, axes = plt.subplots(1, 2, figsize=(12, 5))
fig.suptitle("CEEF: Wrong-Direction Counterfactual Collapse",
             fontsize=13, fontweight="bold")

ax = axes[0]
ax.set_title("Trajectory: Full Control vs Wrong Exit", fontsize=11)
for cx, cy, r, color in [
    (C_morula[0], C_morula[1], r_morula_pca, STAGE_COLORS["morula"]),
    (C_blast[0],  C_blast[1],  r_blast_pca,  STAGE_COLORS["blastocyst"]),
]:
    circle = Circle((cx, cy), r, fill=True, facecolor=color, alpha=0.15,
                    edgecolor=color, linewidth=2)
    ax.add_patch(circle)
draw_background(ax, show_scatter=False)

for name in ["full_control", "wrong_exit"]:
    if name in trajs_pca:
        df = trajs_pca[name]
        ax.plot(df["PC1"], df["PC2"], color=SCEN_COLORS[name], linewidth=2.5,
                label=SCEN_LABELS[name], zorder=4)
        ax.scatter(df["PC1"].iloc[-1], df["PC2"].iloc[-1],
                   s=80, color=SCEN_COLORS[name], marker="*", zorder=6)

ax.set_xlabel("PC1", fontsize=9)
ax.set_ylabel("PC2", fontsize=9)
ax.legend(fontsize=8)
ax.grid(True, alpha=0.25)
ax.set_aspect("equal")

ax2 = axes[1]
ax2.set_title("Basin Capture: Full Control vs Wrong Exit", fontsize=11)
scenarios_compare = ["baseline_only", "plus_entry", "full_control", "wrong_exit"]
x = np.arange(len(scenarios_compare))
morula_vals = [1 if results.get(n,{}).get("in_morula",False) else 0 for n in scenarios_compare]
blast_vals  = [1 if results.get(n,{}).get("in_blast",False) else 0 for n in scenarios_compare]
dist_vals   = [results.get(n,{}).get("dist_morula_t5",5.0) for n in scenarios_compare]

ax2.bar(x-0.2, morula_vals, 0.35, label="In morula basin (τ=5)",
        color=[STAGE_COLORS["morula"] if v else "#F5B7B1" for v in morula_vals])
ax2.bar(x+0.2, blast_vals, 0.35, label="In blast basin (τ=6)",
        color=[STAGE_COLORS["blastocyst"] if v else "#A9DFBF" for v in blast_vals])

ax2.set_xticks(x)
ax2.set_xticklabels(["Baseline", "+Entry", "Full\nControl", "Wrong\nExit"], fontsize=9)
ax2.set_ylabel("Basin capture (1=Yes, 0=No)", fontsize=9)
ax2.set_ylim(0, 1.4)
ax2.legend(fontsize=8)
ax2.grid(True, alpha=0.3, axis="y")
for i, d in enumerate(dist_vals):
    ax2.text(i, 1.1, "d=%.2f" % d, ha="center", fontsize=7, color="gray")
ax2.text(0.5, 0.02,
         "Wrong exit direction: trajectory diverges\nfrom blastocyst basin (collapse)",
         transform=ax2.transAxes, fontsize=8, ha="center", va="bottom",
         bbox=dict(boxstyle="round", facecolor="#FFF3E0", alpha=0.8))

plt.tight_layout()
plt.savefig(FIG/"Fig_C4_counterfactual_collapse.png", dpi=200, bbox_inches="tight")
(FIG/"Fig_C4_counterfactual_collapse.pdf").unlink(missing_ok=True)
plt.savefig(FIG/"Fig_C4_counterfactual_collapse.pdf", bbox_inches="tight")
plt.close()
print("Saved Fig_C4_counterfactual_collapse")

print("\n" + "="*60)
print("FIGURES V2 COMPLETE - saved to:", FIG)
for f in sorted(FIG.iterdir()):
    print("  " + f.name)
```
