## Supplemental code for "An entropy-based diagnostic framework for characterizing methylation state dynamics during preimplantation development": run_corrected_scenarios.py

```
#!/usr/bin/env python
"""
Run all 5 CEEF scenarios using corrected (e1,e2) coordinate system
that preserves entry-exit duality (cos=-0.876, target=-0.699).
"""
import mph
import numpy as np
import pandas as pd
import json
import math
from pathlib import Path

OUT = Path("E:/progress_comsol_analysis")
params  = pd.read_csv(OUT/"parameters_corrected.csv").iloc[0].to_dict()
centers = pd.read_csv(OUT/"stage_centers_2d_corrected.csv").set_index("stage")

print("Starting COMSOL 6.4...")
client = mph.start(cores=2)
print("COMSOL ready")

# Stage coordinates
z1_oocyte = float(centers.loc["oocyte","z1"]); z2_oocyte = float(centers.loc["oocyte","z2"])
z1_4cell  = float(centers.loc["4cell","z1"]);  z2_4cell  = float(centers.loc["4cell","z2"])
z1_8cell  = float(centers.loc["8cell","z1"]);  z2_8cell  = float(centers.loc["8cell","z2"])
z1_morula = float(centers.loc["morula","z1"]); z2_morula = float(centers.loc["morula","z2"])
z1_blast  = float(centers.loc["blast","z1"]);  z2_blast  = float(centers.loc["blast","z2"])

# Direction vectors
vzga_mag   = math.sqrt((z1_8cell-z1_4cell)**2  + (z2_8cell-z2_4cell)**2)
vzga1      = (z1_8cell-z1_4cell)/vzga_mag
vzga2      = (z2_8cell-z2_4cell)/vzga_mag

ventry_mag = math.sqrt((z1_morula-z1_8cell)**2 + (z2_morula-z2_8cell)**2)
ventry1    = (z1_morula-z1_8cell)/ventry_mag
ventry2    = (z2_morula-z2_8cell)/ventry_mag

vexit_mag  = math.sqrt((z1_blast-z1_morula)**2 + (z2_blast-z2_morula)**2)
vexit1     = (z1_blast-z1_morula)/vexit_mag
vexit2     = (z2_blast-z2_morula)/vexit_mag

r_morula = float(params["r_morula"])
r_blast  = float(params["r_blast"])
r_8cell  = float(params["r_8cell"])

cos_check = ventry1*vexit1 + ventry2*vexit2
print("Entry-exit cosine: %.4f (target: -0.699)" % cos_check)
print("r_morula=%.4f, r_8cell=%.4f, r_blast=%.4f" % (r_morula, r_8cell, r_blast))


def run_scenario(name, g_zga, g_entry, g_exit, wrong_exit):
    model = client.create(name)
    java  = model.java

    for k, v in [
        ("lambda_K", "0.4389"),
        ("z1_Katt", str(params["k_att_z1"])),
        ("z2_Katt", str(params["k_att_z2"])),
        ("sigma_M", str(params["sigma_M"])),
        ("sigma_8", str(params["sigma_8"])),
        ("sigma_4", "0.3"),
        ("z1_morula", str(z1_morula)), ("z2_morula", str(z2_morula)),
        ("z1_8cell",  str(z1_8cell)),  ("z2_8cell",  str(z2_8cell)),
        ("z1_4cell",  str(z1_4cell)),  ("z2_4cell",  str(z2_4cell)),
        ("z1_blast",  str(z1_blast)),  ("z2_blast",  str(z2_blast)),
        ("r_morula", str(r_morula)), ("r_blast", str(r_blast)), ("r_8cell", str(r_8cell)),
        ("g_zga",   str(g_zga)),
        ("g_entry", str(g_entry)),
        ("g_exit",  str(g_exit)),
        ("exit_sign", "-1.0" if wrong_exit else "1.0"),
        ("gamma_acc", "1.0"), ("gamma_closure", "1.0"),
        ("gamma_zga", "1.0"), ("gamma_re", "1.0"), ("gamma_de", "1.0"),
        ("vzga1", str(vzga1)), ("vzga2", str(vzga2)),
        ("ventry1", str(ventry1)), ("ventry2", str(ventry2)),
        ("vexit1", str(vexit1)), ("vexit2", str(vexit2)),
    ]:
        java.param().set(k, v)

    java.physics().create("ge", "GlobalEquations")
    ge = java.physics("ge").feature("ge1")
    ge.set("name", ["z1", "z2"])

    eq1 = ("z1t-("
           "-lambda_K*(z1-z1_Katt)"
           "+g_zga*flc2hs(t-3,0.3)*flc2hs(4-t,0.3)*gamma_zga"
           "*exp(-((z1-z1_4cell)^2+(z2-z2_4cell)^2)/(2*sigma_4^2))*vzga1"
           "+g_entry*flc2hs(t-4,0.3)*flc2hs(5-t,0.3)*(gamma_acc+gamma_closure)"
           "*exp(-((z1-z1_8cell)^2+(z2-z2_8cell)^2)/(2*sigma_8^2))*ventry1"
           "+g_exit*flc2hs(t-5,0.3)*flc2hs(6-t,0.3)*(gamma_re+gamma_de)"
           "*exp(-((z1-z1_morula)^2+(z2-z2_morula)^2)/(2*sigma_M^2))*exit_sign*vexit1"
           ")")
    eq2 = ("z2t-("
           "-lambda_K*(z2-z2_Katt)"
           "+g_zga*flc2hs(t-3,0.3)*flc2hs(4-t,0.3)*gamma_zga"
           "*exp(-((z1-z1_4cell)^2+(z2-z2_4cell)^2)/(2*sigma_4^2))*vzga2"
           "+g_entry*flc2hs(t-4,0.3)*flc2hs(5-t,0.3)*(gamma_acc+gamma_closure)"
           "*exp(-((z1-z1_8cell)^2+(z2-z2_8cell)^2)/(2*sigma_8^2))*ventry2"
           "+g_exit*flc2hs(t-5,0.3)*flc2hs(6-t,0.3)*(gamma_re+gamma_de)"
           "*exp(-((z1-z1_morula)^2+(z2-z2_morula)^2)/(2*sigma_M^2))*exit_sign*vexit2"
           ")")

    ge.set("equation", [eq1, eq2])
    ge.set("initialValueU",  [str(z1_oocyte), str(z2_oocyte)])
    ge.set("initialValueUt", ["0", "0"])

    java.study().create("std1")
    java.study("std1").create("time", "Transient")
    java.study("std1").feature("time").set("tlist", "range(0,0.05,6)")
    java.study("std1").feature("time").set("usertol", True)
    java.study("std1").feature("time").set("rtol", "1e-5")
    java.study("std1").run()

    t_vals  = model.evaluate("t")
    z1_vals = model.evaluate("z1")
    z2_vals = model.evaluate("z2")

    traj = pd.DataFrame({"t": t_vals, "z1": z1_vals, "z2": z2_vals})
    traj.to_csv(OUT / ("traj_corrected_" + name + ".csv"), index=False)
    java.save(str(OUT / ("CEEF_corrected_" + name + ".mph")))

    t_arr = list(t_vals)
    idx5 = min(range(len(t_arr)), key=lambda i: abs(t_arr[i]-5.0))
    idx6 = min(range(len(t_arr)), key=lambda i: abs(t_arr[i]-6.0))

    dm5 = math.sqrt((z1_vals[idx5]-z1_morula)**2 + (z2_vals[idx5]-z2_morula)**2)
    db6 = math.sqrt((z1_vals[idx6]-z1_blast)**2  + (z2_vals[idx6]-z2_blast)**2)
    d85 = math.sqrt((z1_vals[idx5]-z1_8cell)**2  + (z2_vals[idx5]-z2_8cell)**2)

    res = {
        "z1_t5": float(z1_vals[idx5]), "z2_t5": float(z2_vals[idx5]),
        "z1_t6": float(z1_vals[idx6]), "z2_t6": float(z2_vals[idx6]),
        "dist_morula_t5": float(dm5), "dist_blast_t6": float(db6),
        "dist_8cell_t5": float(d85),
        "in_morula": bool(dm5 < r_morula),
        "in_blast":  bool(db6 < r_blast),
        "in_8cell_t5": bool(d85 < r_8cell),
        "n_pts": len(t_vals),
    }
    client.remove(model)
    return res


SCENARIOS = [
    ("baseline_only", 0.0, 0.0, 0.0, False),
    ("plus_zga",      1.0, 0.0, 0.0, False),
    ("plus_entry",    1.0, 1.0, 0.0, False),
    ("full_control",  1.0, 1.0, 1.0, False),
    ("wrong_exit",    1.0, 1.0, 1.0, True),
]

all_results = {}
for name, gz, ge, gx, wr in SCENARIOS:
    print("\nRunning: " + name)
    try:
        res = run_scenario(name, gz, ge, gx, wr)
        all_results[name] = res
        print("  t=5: z1=%.3f, z2=%.3f, dist_morula=%.4f, in_morula=%s" % (
            res["z1_t5"], res["z2_t5"], res["dist_morula_t5"], res["in_morula"]))
        print("  t=6: z1=%.3f, z2=%.3f, dist_blast=%.4f,  in_blast=%s" % (
            res["z1_t6"], res["z2_t6"], res["dist_blast_t6"], res["in_blast"]))
    except Exception as e:
        print("  ERROR: " + str(e)[:120])
        all_results[name] = {"error": str(e)}

with open(OUT/"scenario_results_corrected.json", "w", encoding="utf-8") as f:
    json.dump(all_results, f, indent=2, default=str)

print("\n" + "="*60)
print("SUMMARY (corrected coordinate system)")
print("="*60)
print("Validation targets: in_morula=True for full_control, in_morula=False for baseline")
for name, res in all_results.items():
    if "error" not in res:
        print("  %-20s morula=%-5s blast=%-5s dist_morula=%.4f" % (
            name, str(res["in_morula"]), str(res["in_blast"]), res["dist_morula_t5"]))
    else:
        print("  %-20s ERROR" % name)
```
