## Supplemental code for "An entropy-based diagnostic framework for characterizing methylation state dynamics during preimplantation development": run_final_scenarios.py

```
#!/usr/bin/env python
"""
Final CEEF scenarios with calibrated parameters.
Uses linear field (no Gaussian) and proper basin radii.
"""
import mph
import numpy as np
import pandas as pd
import json
import math
from pathlib import Path

OUT = Path("E:/progress_comsol_analysis")
centers = pd.read_csv(OUT/"stage_centers_2d_corrected.csv").set_index("stage")

print("Starting COMSOL 6.4...")
client = mph.start(cores=2)
print("COMSOL ready")

# Stage coordinates
z1_oocyte = float(centers.loc["oocyte","z1"]); z2_oocyte = float(centers.loc["oocyte","z2"])
z1_4cell  = float(centers.loc["4cell","z1"]);  z2_4cell  = float(centers.loc["4cell","z2"])
z1_8cell  = float(centers.loc["8cell","z1"]);  z2_8cell  = float(centers.loc["8cell","z2"])
z1_morula = float(centers.loc["morula","z1"]); z2_morula = float(centers.loc["morula","z2"])
z1_blast  = float(centers.loc["blast","z1"]);  z2_blast  = float(centers.loc["blast","z2"])
z1_Katt   = float(centers["z1"].mean())
z2_Katt   = float(centers["z2"].mean())

# Direction vectors
vzga_mag   = math.sqrt((z1_8cell-z1_4cell)**2  + (z2_8cell-z2_4cell)**2)
vzga1      = (z1_8cell-z1_4cell)/vzga_mag
vzga2      = (z2_8cell-z2_4cell)/vzga_mag

ventry_mag = math.sqrt((z1_morula-z1_8cell)**2 + (z2_morula-z2_8cell)**2)
ventry1    = (z1_morula-z1_8cell)/ventry_mag
ventry2    = (z2_morula-z2_8cell)/ventry_mag

vexit_mag  = math.sqrt((z1_blast-z1_morula)**2 + (z2_blast-z2_morula)**2)
vexit1     = (z1_blast-z1_morula)/vexit_mag
vexit2     = (z2_blast-z2_morula)/vexit_mag

# Calibrated basin radii (based on stage separation)
# morula is isolated (min dist to other stages = 2.72)
# Use r = 0.5 to capture trajectories reaching within 0.5 of morula center
r_morula = 0.50
r_blast  = 0.60
r_8cell  = 0.80

# Gamma calibrated: need gamma > lambda_K * dist / dt
# dist_8cell_to_morula = 3.42, dt = 1, lambda_K = 0.44
# gamma_entry = 4.0 ensures reaching morula
lambda_K = 0.4389

cos_check = ventry1*vexit1 + ventry2*vexit2
print("Entry-exit cosine: %.4f (target: -0.699)" % cos_check)
print("r_morula=%.3f, r_8cell=%.3f, r_blast=%.3f" % (r_morula, r_8cell, r_blast))
print("gamma_entry=4.0 (calibrated to reach morula in 1 time unit)")


def run_scenario(name, g_zga, g_entry, g_exit, wrong_exit, gamma_entry=4.0, gamma_exit=4.0):
    model = client.create(name)
    java  = model.java

    for k, v in [
        ("lambda_K", str(lambda_K)),
        ("z1_Katt", str(z1_Katt)), ("z2_Katt", str(z2_Katt)),
        ("z1_morula", str(z1_morula)), ("z2_morula", str(z2_morula)),
        ("z1_8cell",  str(z1_8cell)),  ("z2_8cell",  str(z2_8cell)),
        ("z1_4cell",  str(z1_4cell)),  ("z2_4cell",  str(z2_4cell)),
        ("z1_blast",  str(z1_blast)),  ("z2_blast",  str(z2_blast)),
        ("r_morula", str(r_morula)), ("r_blast", str(r_blast)), ("r_8cell", str(r_8cell)),
        ("g_zga",   str(g_zga)),
        ("g_entry", str(g_entry)),
        ("g_exit",  str(g_exit)),
        ("exit_sign", "-1.0" if wrong_exit else "1.0"),
        ("gamma_zga",   "2.0"),
        ("gamma_entry", str(gamma_entry)),
        ("gamma_exit",  str(gamma_exit)),
        ("vzga1", str(vzga1)), ("vzga2", str(vzga2)),
        ("ventry1", str(ventry1)), ("ventry2", str(ventry2)),
        ("vexit1", str(vexit1)), ("vexit2", str(vexit2)),
    ]:
        java.param().set(k, v)

    java.physics().create("ge", "GlobalEquations")
    ge = java.physics("ge").feature("ge1")
    ge.set("name", ["z1", "z2"])

    # Linear field: constant direction * gamma * time_gate
    eq1 = ("z1t-("
           "-lambda_K*(z1-z1_Katt)"
           "+g_zga*flc2hs(t-3,0.3)*flc2hs(4-t,0.3)*gamma_zga*vzga1"
           "+g_entry*flc2hs(t-4,0.3)*flc2hs(5-t,0.3)*gamma_entry*ventry1"
           "+g_exit*flc2hs(t-5,0.3)*flc2hs(6-t,0.3)*gamma_exit*exit_sign*vexit1"
           ")")
    eq2 = ("z2t-("
           "-lambda_K*(z2-z2_Katt)"
           "+g_zga*flc2hs(t-3,0.3)*flc2hs(4-t,0.3)*gamma_zga*vzga2"
           "+g_entry*flc2hs(t-4,0.3)*flc2hs(5-t,0.3)*gamma_entry*ventry2"
           "+g_exit*flc2hs(t-5,0.3)*flc2hs(6-t,0.3)*gamma_exit*exit_sign*vexit2"
           ")")

    ge.set("equation", [eq1, eq2])
    ge.set("initialValueU",  [str(z1_oocyte), str(z2_oocyte)])
    ge.set("initialValueUt", ["0", "0"])

    java.study().create("std1")
    java.study("std1").create("time", "Transient")
    java.study("std1").feature("time").set("tlist", "range(0,0.05,6)")
    java.study("std1").feature("time").set("usertol", True)
    java.study("std1").feature("time").set("rtol", "1e-5")
    java.study("std1").run()

    t_vals  = model.evaluate("t")
    z1_vals = model.evaluate("z1")
    z2_vals = model.evaluate("z2")

    traj = pd.DataFrame({"t": t_vals, "z1": z1_vals, "z2": z2_vals})
    traj.to_csv(OUT / ("traj_final_" + name + ".csv"), index=False)
    java.save(str(OUT / ("CEEF_final_" + name + ".mph")))

    t_arr = list(t_vals)
    idx5 = min(range(len(t_arr)), key=lambda i: abs(t_arr[i]-5.0))
    idx6 = min(range(len(t_arr)), key=lambda i: abs(t_arr[i]-6.0))

    dm5 = math.sqrt((z1_vals[idx5]-z1_morula)**2 + (z2_vals[idx5]-z2_morula)**2)
    db6 = math.sqrt((z1_vals[idx6]-z1_blast)**2  + (z2_vals[idx6]-z2_blast)**2)

    res = {
        "z1_t5": float(z1_vals[idx5]), "z2_t5": float(z2_vals[idx5]),
        "z1_t6": float(z1_vals[idx6]), "z2_t6": float(z2_vals[idx6]),
        "dist_morula_t5": float(dm5), "dist_blast_t6": float(db6),
        "in_morula": bool(dm5 < r_morula),
        "in_blast":  bool(db6 < r_blast),
        "n_pts": len(t_vals),
    }
    client.remove(model)
    return res


SCENARIOS = [
    ("baseline_only", 0.0, 0.0, 0.0, False),
    ("plus_zga",      1.0, 0.0, 0.0, False),
    ("plus_entry",    1.0, 1.0, 0.0, False),
    ("full_control",  1.0, 1.0, 1.0, False),
    ("wrong_exit",    1.0, 1.0, 1.0, True),
]

all_results = {}
for name, gz, ge, gx, wr in SCENARIOS:
    print("\nRunning: " + name)
    try:
        res = run_scenario(name, gz, ge, gx, wr)
        all_results[name] = res
        print("  t=5: z1=%.3f, z2=%.3f, dist_morula=%.4f, in_morula=%s" % (
            res["z1_t5"], res["z2_t5"], res["dist_morula_t5"], res["in_morula"]))
        print("  t=6: z1=%.3f, z2=%.3f, dist_blast=%.4f,  in_blast=%s" % (
            res["z1_t6"], res["z2_t6"], res["dist_blast_t6"], res["in_blast"]))
    except Exception as e:
        print("  ERROR: " + str(e)[:120])
        all_results[name] = {"error": str(e)}

with open(OUT/"scenario_results_final.json", "w", encoding="utf-8") as f:
    json.dump(all_results, f, indent=2, default=str)

print("\n" + "="*60)
print("FINAL SUMMARY")
print("="*60)
print("Validation: baseline=False, full_control=True, wrong_exit=False(blast)")
for name, res in all_results.items():
    if "error" not in res:
        print("  %-20s morula=%-5s blast=%-5s dist_morula=%.4f" % (
            name, str(res["in_morula"]), str(res["in_blast"]), res["dist_morula_t5"]))
    else:
        print("  %-20s ERROR" % name)
```
