## Supplemental code for "An entropy-based diagnostic framework for characterizing methylation state dynamics during preimplantation development": run_scenarios.py

```
#!/usr/bin/env python
"""
Build and run all 5 CEEF scenarios in COMSOL 6.4.
"""
import mph
import numpy as np
import pandas as pd
import json
from pathlib import Path

OUT = Path("E:/progress_comsol_analysis")
params  = pd.read_csv(OUT/"parameters.csv").iloc[0].to_dict()
centers = pd.read_csv(OUT/"stage_centers_2d.csv").set_index("stage")

print("Starting COMSOL 6.4...")
client = mph.start(cores=2)
print("COMSOL ready")

STAGE_MAP = {
    "oocyte": "MII oocyte", "zygote": "zygote/PN",
    "2cell": "2-cell", "4cell": "4-cell",
    "8cell": "8-cell", "morula": "morula", "blast": "blastocyst",
}

def build_and_run(scenario_name, g_zga=1.0, g_entry=1.0, g_exit=1.0, wrong_exit=False):
    model = client.create(scenario_name)
    java  = model.java

    # Parameters
    java.param().set("alpha_op", "0.5611")
    java.param().set("bias_op",  "0.0688")
    java.param().set("lambda_K", "0.4389")

    for short, full in STAGE_MAP.items():
        if full in centers.index:
            java.param().set("z1_" + short, str(centers.loc[full, "z1"]))
            java.param().set("z2_" + short, str(centers.loc[full, "z2"]))

    java.param().set("z1_Katt",   str(params["k_att_z1"]))
    java.param().set("z2_Katt",   str(params["k_att_z2"]))
    java.param().set("r_morula",  str(params["r_morula"]))
    java.param().set("r_8cell",   str(params["r_8cell"]))
    java.param().set("r_blast",   str(params["r_blast"]))
    java.param().set("sigma_M",   str(params["sigma_M"]))
    java.param().set("sigma_8",   str(params["sigma_8"]))
    java.param().set("sigma_4",   "2.0")
    java.param().set("g_zga",     str(g_zga))
    java.param().set("g_entry",   str(g_entry))
    java.param().set("g_exit",    str(g_exit))
    java.param().set("exit_sign", "-1.0" if wrong_exit else "1.0")
    java.param().set("gamma_acc",     "1.0")
    java.param().set("gamma_closure", "1.0")
    java.param().set("gamma_zga",     "1.0")
    java.param().set("gamma_re",      "1.0")
    java.param().set("gamma_de",      "1.0")

    # Physics: Global ODEs (no component needed)
    java.physics().create("ge", "GlobalEquations")
    ge = java.physics("ge").feature("ge1")
    ge.set("name", ["z1", "z2"])
    ge.set("equation", [
        "z1t-(FK1+g_zga*Fzga1+g_entry*Fentry1+g_exit*Fexit1)",
        "z2t-(FK2+g_zga*Fzga2+g_entry*Fentry2+g_exit*Fexit2)"
    ])
    ge.set("initialValueU",  ["z1_oocyte", "z2_oocyte"])
    ge.set("initialValueUt", ["0", "0"])

    # Variables
    java.variable().create("var1")
    v = java.variable("var1")
    v.set("rM2", "(z1-z1_morula)^2+(z2-z2_morula)^2")
    v.set("r82", "(z1-z1_8cell)^2+(z2-z2_8cell)^2")
    v.set("r42", "(z1-z1_4cell)^2+(z2-z2_4cell)^2")
    v.set("rB2", "(z1-z1_blast)^2+(z2-z2_blast)^2")
    v.set("gM",  "exp(-rM2/(2*sigma_M^2))")
    v.set("g8",  "exp(-r82/(2*sigma_8^2))")
    v.set("g4",  "exp(-r42/(2*sigma_4^2))")
    v.set("chi_zga",   "flc2hs(tau-3,0.3)*flc2hs(4-tau,0.3)")
    v.set("chi_entry", "flc2hs(tau-4,0.3)*flc2hs(5-tau,0.3)")
    v.set("chi_exit",  "flc2hs(tau-5,0.3)*flc2hs(6-tau,0.3)")
    v.set("FK1", "-lambda_K*(z1-z1_Katt)")
    v.set("FK2", "-lambda_K*(z2-z2_Katt)")
    v.set("vzga_mag",   "sqrt((z1_8cell-z1_4cell)^2+(z2_8cell-z2_4cell)^2)+1e-12")
    v.set("vzga1",      "(z1_8cell-z1_4cell)/vzga_mag")
    v.set("vzga2",      "(z2_8cell-z2_4cell)/vzga_mag")
    v.set("ventry_mag", "sqrt((z1_morula-z1_8cell)^2+(z2_morula-z2_8cell)^2)+1e-12")
    v.set("ventry1",    "(z1_morula-z1_8cell)/ventry_mag")
    v.set("ventry2",    "(z2_morula-z2_8cell)/ventry_mag")
    v.set("vexit_mag",  "sqrt((z1_blast-z1_morula)^2+(z2_blast-z2_morula)^2)+1e-12")
    v.set("vexit1",     "exit_sign*(z1_blast-z1_morula)/vexit_mag")
    v.set("vexit2",     "exit_sign*(z2_blast-z2_morula)/vexit_mag")
    v.set("Fzga1",   "chi_zga*gamma_zga*g4*vzga1")
    v.set("Fzga2",   "chi_zga*gamma_zga*g4*vzga2")
    v.set("Fentry1", "chi_entry*(gamma_acc+gamma_closure)*g8*ventry1")
    v.set("Fentry2", "chi_entry*(gamma_acc+gamma_closure)*g8*ventry2")
    v.set("Fexit1",  "chi_exit*(gamma_re+gamma_de)*gM*vexit1")
    v.set("Fexit2",  "chi_exit*(gamma_re+gamma_de)*gM*vexit2")
    v.set("in_morula_basin", "if(sqrt(rM2)<r_morula,1,0)")
    v.set("in_blast_basin",  "if(sqrt(rB2)<r_blast,1,0)")
    v.set("dist_morula", "sqrt(rM2)")
    v.set("dist_blast",  "sqrt(rB2)")
    v.set("entry_exit_cos", "ventry1*vexit1+ventry2*vexit2")

    # Study
    java.study().create("std1")
    java.study("std1").create("time", "Transient")
    java.study("std1").feature("time").set("tlist", "range(0,0.05,6)")
    java.study("std1").feature("time").set("usertol", True)
    java.study("std1").feature("time").set("rtol", "1e-5")

    # Run
    java.study("std1").run()

    # Extract results
    t_vals  = model.evaluate("t")
    z1_vals = model.evaluate("z1")
    z2_vals = model.evaluate("z2")
    dm_vals = model.evaluate("dist_morula")
    db_vals = model.evaluate("dist_blast")
    im_vals = model.evaluate("in_morula_basin")
    ib_vals = model.evaluate("in_blast_basin")

    # Save trajectory CSV
    traj = pd.DataFrame({
        "tau": t_vals, "z1": z1_vals, "z2": z2_vals,
        "dist_morula": dm_vals, "dist_blast": db_vals,
        "in_morula": im_vals, "in_blast": ib_vals,
    })
    traj.to_csv(OUT / ("traj_" + scenario_name + ".csv"), index=False)

    # Save mph file
    java.save(str(OUT / ("CEEF_" + scenario_name + ".mph")))

    # Compute summary metrics
    t_arr = np.array(t_vals)
    idx5 = int(np.argmin(np.abs(t_arr - 5.0)))
    idx6 = int(np.argmin(np.abs(t_arr - 6.0)))

    res = {
        "z1_tau5": float(z1_vals[idx5]), "z2_tau5": float(z2_vals[idx5]),
        "z1_tau6": float(z1_vals[idx6]), "z2_tau6": float(z2_vals[idx6]),
        "dist_morula_tau5": float(dm_vals[idx5]),
        "dist_blast_tau6":  float(db_vals[idx6]),
        "in_morula_tau5": bool(float(im_vals[idx5]) > 0.5),
        "in_blast_tau6":  bool(float(ib_vals[idx6]) > 0.5),
        "n_timepoints": len(t_vals),
    }
    client.remove(model)
    return res


# Run all 5 scenarios
SCENARIOS = [
    ("baseline_only", 0.0, 0.0, 0.0, False),
    ("plus_zga",      1.0, 0.0, 0.0, False),
    ("plus_entry",    1.0, 1.0, 0.0, False),
    ("full_control",  1.0, 1.0, 1.0, False),
    ("wrong_exit",    1.0, 1.0, 1.0, True),
]

all_results = {}
for name, gz, ge, gx, wr in SCENARIOS:
    print("Running: " + name)
    try:
        res = build_and_run(name, gz, ge, gx, wr)
        all_results[name] = res
        print("  tau=5: dist_morula=" + str(round(res["dist_morula_tau5"],3)) +
              ", in_morula=" + str(res["in_morula_tau5"]))
        print("  tau=6: dist_blast=" + str(round(res["dist_blast_tau6"],3)) +
              ", in_blast=" + str(res["in_blast_tau6"]))
    except Exception as e:
        print("  ERROR: " + str(e)[:100])
        all_results[name] = {"error": str(e)}

with open(OUT / "scenario_results.json", "w", encoding="utf-8") as f:
    json.dump(all_results, f, indent=2, default=str)

print("\nSUMMARY:")
for name, res in all_results.items():
    if "error" not in res:
        print("  " + name + ": morula=" + str(res["in_morula_tau5"]) +
              ", blast=" + str(res["in_blast_tau6"]))
    else:
        print("  " + name + ": ERROR")

print("\nFiles saved to " + str(OUT))
```
