## Supplemental code for "An entropy-based diagnostic framework for characterizing methylation state dynamics during preimplantation development": 01_prepare_reference.sh

```
#!/usr/bin/env bash
set -euo pipefail

ROOT=/mnt/e/5_31_progress/bismark_full_closure
ENV=$ROOT/env/bismark
MM=$ROOT/tools/micromamba
REF=$ROOT/ref/hg19

mkdir -p "$REF"
cd "$REF"

if [[ ! -s hg19.fa ]]; then
  if [[ ! -s hg19.fa.gz ]]; then
    curl --retry 20 --retry-delay 10 -L \
      https://hgdownload.soe.ucsc.edu/goldenPath/hg19/bigZips/hg19.fa.gz \
      -o hg19.fa.gz
  fi
  gunzip -c hg19.fa.gz > hg19.fa
fi

if [[ ! -d Bisulfite_Genome/CT_conversion || ! -d Bisulfite_Genome/GA_conversion ]]; then
  "$MM" run -p "$ENV" bismark_genome_preparation --bowtie2 "$REF"
fi

echo "Reference ready: $REF"
```
