## Supplemental code for "An entropy-based diagnostic framework for characterizing methylation state dynamics during preimplantation development": 02_download_fastqs.sh

```
#!/usr/bin/env bash
set -euo pipefail

ROOT=${ROOT_OVERRIDE:-/mnt/e/5_31_progress/bismark_full_closure}
SHEET=${1:-$ROOT/samplesheet_E-MTAB-10097_all359.tsv}
LIMIT=${2:-0}
OFFSET=${3:-0}
FASTQ=$ROOT/fastq
mkdir -p "$FASTQ" "$ROOT/logs"

download_resume() {
  local url=$1
  local out=$2
  local remote_size=$3
  local attempt=1
  local max_attempts=${DOWNLOAD_MAX_ATTEMPTS:-120}
  local sleep_seconds=${DOWNLOAD_RETRY_SLEEP:-20}
  local max_seconds=${DOWNLOAD_MAX_SECONDS:-0}
  local start_ts
  start_ts=$(date +%s)

  while [[ "$attempt" -le "$max_attempts" ]]; do
    if [[ "$max_seconds" -gt 0 ]]; then
      local now_ts
      now_ts=$(date +%s)
      if [[ $((now_ts - start_ts)) -ge "$max_seconds" ]]; then
        echo "ERROR: download walltime exceeded ${max_seconds}s for $out" >&2
        return 124
      fi
    fi

    local local_size=0
    if [[ -s "$out" ]]; then
      local_size=$(stat -c%s "$out")
    fi

    echo "$(date -Is) download attempt=$attempt local_size=$local_size remote_size=$remote_size out=$out"
    if [[ "$local_size" -gt 0 ]]; then
      curl -f -L -C - --connect-timeout 60 --speed-time 300 --speed-limit 1024 "$url" -o "$out" && return 0
    else
      curl -f -L --connect-timeout 60 --speed-time 300 --speed-limit 1024 "$url" -o "$out" && return 0
    fi

    local rc=$?
    local after_size=0
    if [[ -s "$out" ]]; then
      after_size=$(stat -c%s "$out")
    fi
    echo "$(date -Is) download failed rc=$rc attempt=$attempt after_size=$after_size; retrying in ${sleep_seconds}s" >&2
    sleep "$sleep_seconds"
    attempt=$((attempt + 1))
  done

  echo "ERROR: download failed after $max_attempts attempts for $out" >&2
  return 1
}

count=0
tail -n +2 "$SHEET" | while IFS= read -r line; do
  sample=$(printf '%s\n' "$line" | cut -f1)
  run=$(printf '%s\n' "$line" | cut -f2)
  fq1=$(printf '%s\n' "$line" | cut -f9)
  fq2=$(printf '%s\n' "$line" | cut -f10)
  count=$((count + 1))
  if [[ "$count" -le "$OFFSET" ]]; then
    continue
  fi
  done_count=$((count - OFFSET))
  if [[ "$LIMIT" != "0" && "$done_count" -gt "$LIMIT" ]]; then
    break
  fi
  mkdir -p "$FASTQ/$run"
  for fq in "$fq1" "$fq2"; do
    out="$FASTQ/$run/$(basename "$fq")"
    remote_size=$(curl -fsSIL "$fq" | awk 'tolower($1)=="content-length:" {gsub("\r","",$2); n=$2} END{print n+0}' || echo 0)
    local_size=0
    if [[ -s "$out" ]]; then
      local_size=$(stat -c%s "$out")
    fi
    if [[ "$remote_size" -gt 0 && "$local_size" -eq "$remote_size" ]]; then
      if gzip -t "$out"; then
        echo "complete: $out ($local_size bytes)"
        continue
      fi
      echo "ERROR: corrupt gzip despite complete size for $out; redownloading" >&2
      rm -f "$out"
      local_size=0
    fi
    if [[ "$remote_size" -eq 0 && "$local_size" -gt 0 ]]; then
      if gzip -t "$out"; then
        echo "complete without remote size: $out ($local_size bytes)"
        continue
      fi
      echo "ERROR: corrupt gzip for $out; redownloading" >&2
      rm -f "$out"
      local_size=0
    fi
    if [[ "$remote_size" -gt 0 && "$local_size" -gt "$remote_size" ]]; then
      rm -f "$out"
      local_size=0
    fi
    download_resume "$fq" "$out" "$remote_size"
    if [[ "$remote_size" -gt 0 ]]; then
      local_size=$(stat -c%s "$out")
      if [[ "$local_size" -ne "$remote_size" ]]; then
        echo "ERROR: incomplete download for $out: got $local_size expected $remote_size" >&2
        exit 1
      fi
    fi
    if ! gzip -t "$out"; then
      echo "ERROR: corrupt gzip after download for $out" >&2
      exit 1
    fi
  done
done

echo "FASTQ downloads complete for $SHEET"
```
