## Supplemental code for "An entropy-based diagnostic framework for characterizing methylation state dynamics during preimplantation development": 03_run_bismark_one.sh

```
#!/usr/bin/env bash
set -euo pipefail

ROOT=${ROOT_OVERRIDE:-/mnt/e/5_31_progress/bismark_full_closure}
ENV=$ROOT/env/bismark
MM=$ROOT/tools/micromamba
REF=$ROOT/ref/hg19
FASTQ=$ROOT/fastq
WORK=$ROOT/work
RESULTS=$ROOT/results

sample=${1:?sample required}
run=${2:?run required}
threads=${3:-4}
cleanup_bam=${4:-1}

mkdir -p "$WORK/$run" "$RESULTS/$run" "$ROOT/logs"
fq1=$(ls "$FASTQ/$run"/*_1.fastq.gz | head -n 1)
fq2=$(ls "$FASTQ/$run"/*_2.fastq.gz | head -n 1)
cd "$WORK/$run"

if ! ls "$RESULTS/$run"/*_bismark_bt2_PE_report.txt >/dev/null 2>&1 || ! grep -q "Sequence pairs analysed in total" "$RESULTS/$run"/*_bismark_bt2_PE_report.txt 2>/dev/null; then
  rm -f "$RESULTS/$run"/*_bismark_bt2_pe.bam "$RESULTS/$run"/*_bismark_bt2_PE_report.txt
  "$MM" run -p "$ENV" bismark \
    --genome "$REF" \
    --bowtie2 \
    --non_directional \
    --multicore "$threads" \
    --output_dir "$RESULTS/$run" \
    -1 "$fq1" -2 "$fq2" \
    > "$ROOT/logs/${run}.bismark.log" 2>&1
fi

bam=$(ls "$RESULTS/$run"/*_bismark_bt2_pe.bam | head -n 1)

if [[ ! -s "${bam%.bam}.deduplicated.bam" ]]; then
  "$MM" run -p "$ENV" deduplicate_bismark \
    --paired \
    --bam "$bam" \
    --output_dir "$RESULTS/$run" \
    > "$ROOT/logs/${run}.dedup.log" 2>&1
fi

dedup=$(ls "$RESULTS/$run"/*deduplicated.bam | head -n 1)

if ! ls "$RESULTS/$run"/*.bismark.cov.gz >/dev/null 2>&1; then
  "$MM" run -p "$ENV" bismark_methylation_extractor \
    --paired-end \
    --bedGraph \
    --gzip \
    --buffer_size 1G \
    --multicore "$threads" \
    --genome_folder "$REF" \
    --output "$RESULTS/$run" \
    "$dedup" \
    > "$ROOT/logs/${run}.extract.log" 2>&1
fi

if [[ "$cleanup_bam" == "1" ]] && ls "$RESULTS/$run"/*.bismark.cov.gz >/dev/null 2>&1; then
  find "$RESULTS/$run" -maxdepth 1 -type f \( -name "*.bam" -o -name "*.bam.bai" \) -delete
fi

echo "$run done"
```
