## Supplemental code for "An entropy-based diagnostic framework for characterizing methylation state dynamics during preimplantation development": 04_run_bismark_batch.sh

```
#!/usr/bin/env bash
set -euo pipefail

ROOT=${ROOT_OVERRIDE:-/mnt/e/5_31_progress/bismark_full_closure}
SHEET=${1:-$ROOT/samplesheet_E-MTAB-10097_priority_order.tsv}
LIMIT=${2:-0}
OFFSET=${3:-0}
THREADS=${4:-4}
CLEANUP_BAM=${5:-1}
count=0

tail -n +2 "$SHEET" | while IFS= read -r line; do
  sample=$(printf '%s\n' "$line" | cut -f1)
  run=$(printf '%s\n' "$line" | cut -f2)
  count=$((count + 1))
  if [[ "$count" -le "$OFFSET" ]]; then
    continue
  fi
  done_count=$((count - OFFSET))
  if [[ "$LIMIT" != "0" && "$done_count" -gt "$LIMIT" ]]; then
    break
  fi
  if ls "$ROOT/results/$run"/*.bismark.cov.gz >/dev/null 2>&1; then
    echo "$run already has cov output; skipping"
    continue
  fi
  bash "$ROOT/scripts/03_run_bismark_one.sh" "$sample" "$run" "$THREADS" "$CLEANUP_BAM"
done
```
