## Supplemental code for "An entropy-based diagnostic framework for characterizing methylation state dynamics during preimplantation development": 05_aggregate_csb_dmrs.py

```
import gzip
import json
import os
from pathlib import Path

import numpy as np
import pandas as pd
from scipy import stats


ROOT = Path(os.environ.get("ROOT_OVERRIDE", "/mnt/e/5_31_progress/bismark_full_closure"))
SHEET = Path(os.environ.get("SHEET_OVERRIDE", ROOT / "samplesheet_E-MTAB-10097_all359.tsv"))
DMR = ROOT / "CSB_TRO_156_residual_DMR_hg19.bed"
RESULTS = ROOT / "results"
OUT = ROOT / "results"


def load_dmrs():
    if DMR.suffix == ".bed":
        d = pd.read_csv(
            DMR,
            sep="\t",
            header=None,
            names=["chr", "start", "end", "cluster_name", "basin_residual_rank", "latent_residual_delta_beta", "module_id"],
        )
        d["abs_latent_residual_delta_beta"] = d["latent_residual_delta_beta"].abs()
    else:
        d = pd.read_csv(DMR, sep="\t")
    d = d.sort_values("basin_residual_rank").reset_index(drop=True)
    d["dmr_index"] = np.arange(len(d))
    return d[["dmr_index", "cluster_name", "chr", "start", "end", "basin_residual_rank", "latent_residual_delta_beta", "abs_latent_residual_delta_beta", "module_id"]].copy()


def aggregate_cov(path, dmr):
    intervals = {}
    for chrom, sub in dmr.groupby("chr"):
        s = sub.sort_values("start")
        intervals[chrom] = (s["start"].to_numpy(), s["end"].to_numpy(), s["dmr_index"].to_numpy())
    meth = np.zeros(len(dmr), dtype=float)
    total = np.zeros(len(dmr), dtype=float)
    opener = gzip.open if str(path).endswith(".gz") else open
    with opener(path, "rt") as f:
        for line in f:
            parts = line.rstrip("\n").split("\t")
            if len(parts) < 6:
                continue
            chrom = parts[0]
            if chrom not in intervals:
                continue
            pos0 = int(parts[1]) - 1 if int(parts[1]) > 0 else int(parts[1])
            m = float(parts[4])
            u = float(parts[5])
            if m + u <= 0:
                continue
            starts, ends, idxs = intervals[chrom]
            j = np.searchsorted(starts, pos0 + 2, side="left")
            if j == 0:
                continue
            hits = np.where(ends[:j] > pos0)[0]
            for h in hits:
                idx = idxs[h]
                meth[idx] += m
                total[idx] += m + u
    return meth, total


def main():
    sheet = pd.read_csv(SHEET, sep="\t")
    dmr = load_dmrs()
    rows = []
    for _, s in sheet.iterrows():
        run = s["run"]
        covs = list((RESULTS / run).glob("*.bismark.cov.gz"))
        if not covs:
            continue
        meth, total = aggregate_cov(covs[0], dmr)
        for i, row in dmr.iterrows():
            rows.append({
                "sample": s["sample"],
                "run": run,
                "condition": s["condition"],
                "lineage": s["lineage"],
                "individual": s["individual"],
                "dmr_index": row["dmr_index"],
                "cluster_name": row["cluster_name"],
                "basin_residual_rank": row["basin_residual_rank"],
                "module_id": row["module_id"],
                "latent_residual_delta_beta": row["latent_residual_delta_beta"],
                "meth": meth[i],
                "total": total[i],
                "beta": np.nan if total[i] == 0 else meth[i] / total[i],
            })
    beta = pd.DataFrame(rows)
    beta.to_csv(OUT / "E-MTAB-10097_full_bismark_CSB_DMR_sample_beta.tsv", sep="\t", index=False)
    if beta.empty:
        raise SystemExit("No cov files found")
    agg = beta.groupby(["condition", "cluster_name"], as_index=False).agg(
        meth=("meth", "sum"),
        total=("total", "sum"),
        n_samples=("sample", "nunique"),
        basin_residual_rank=("basin_residual_rank", "first"),
        latent_residual_delta_beta=("latent_residual_delta_beta", "first"),
        module_id=("module_id", "first"),
    )
    agg["beta"] = agg["meth"] / agg["total"].replace(0, np.nan)
    wide = agg.pivot(index="cluster_name", columns="condition", values="beta").reset_index()
    wcov = agg.pivot(index="cluster_name", columns="condition", values="total").reset_index()
    meta = agg.drop_duplicates("cluster_name")[["cluster_name", "basin_residual_rank", "latent_residual_delta_beta", "module_id"]]
    wide = meta.merge(wide, on="cluster_name", how="left").merge(wcov, on="cluster_name", how="left", suffixes=("", "_calls"))
    if "control" in wide and "dex" in wide:
        wide["dex_minus_control_beta"] = wide["dex"] - wide["control"]
    wide.to_csv(OUT / "E-MTAB-10097_full_bismark_CSB_DMR_condition_delta.tsv", sep="\t", index=False)
    valid = wide.dropna(subset=["dex_minus_control_beta"]) if "dex_minus_control_beta" in wide else wide.iloc[0:0]
    rho = stats.spearmanr(valid["latent_residual_delta_beta"], valid["dex_minus_control_beta"], nan_policy="omit") if len(valid) >= 3 else (np.nan, np.nan)
    sign_p = np.nan
    sign_match = 0
    if len(valid) > 0:
        signed = valid[(valid["latent_residual_delta_beta"] != 0) & (valid["dex_minus_control_beta"] != 0)].copy()
        if len(signed) > 0:
            sign_match = int((np.sign(signed["latent_residual_delta_beta"]) == np.sign(signed["dex_minus_control_beta"])).sum())
            sign_p = stats.binomtest(sign_match, len(signed), 0.5, alternative="greater").pvalue
    summary = {
        "analysis": "E-MTAB-10097_full_Bismark_CSB_DMR_aggregation",
        "processed_runs": int(beta["run"].nunique()),
        "dmrs_with_control_and_dex_beta": int(len(valid)),
        "control_total_cpg_calls": float(agg[agg["condition"].eq("control")]["total"].sum()),
        "dex_total_cpg_calls": float(agg[agg["condition"].eq("dex")]["total"].sum()),
        "spearman_latent_residual_vs_dex_delta_rho": None if np.isnan(rho[0]) else float(rho[0]),
        "spearman_latent_residual_vs_dex_delta_p": None if np.isnan(rho[1]) else float(rho[1]),
        "sign_concordant_dmrs": int(sign_match),
        "sign_concordance_binomial_p_greater": None if np.isnan(sign_p) else float(sign_p),
    }
    with open(OUT / "E-MTAB-10097_full_bismark_CSB_DMR_summary.json", "w") as f:
        json.dump(summary, f, indent=2)
    print(json.dumps(summary, indent=2))


if __name__ == "__main__":
    main()
```
