## Supplemental code for "An entropy-based diagnostic framework for characterizing methylation state dynamics during preimplantation development": 06_drive_closure.sh

```
#!/usr/bin/env bash
set -euo pipefail

ROOT=${ROOT_OVERRIDE:-/home/u8068/bismark_full_closure}
SHEET=${1:-$ROOT/samplesheet_E-MTAB-10097_highdepth_8ctrl_8dex_cap2GB.tsv}
START_OFFSET=${2:-0}
LIMIT=${3:-0}
THREADS=${4:-1}

mkdir -p "$ROOT/logs"
cd "$ROOT"

is_breakthrough() {
  "$ROOT/tools/micromamba" run -p "$ROOT/env/bismark" python - <<'PY'
import json
from pathlib import Path

path = Path("results/E-MTAB-10097_full_bismark_CSB_DMR_summary.json")
if not path.exists():
    raise SystemExit(1)
s = json.loads(path.read_text())
paired = s.get("dmrs_with_control_and_dex_beta", 0) or 0
rho = s.get("spearman_latent_residual_vs_dex_delta_rho")
rho_p = s.get("spearman_latent_residual_vs_dex_delta_p")
sign_p = s.get("sign_concordance_binomial_p_greater")
hit = False
if paired >= 10 and rho is not None and abs(rho) >= 0.30 and rho_p is not None and rho_p <= 0.10:
    hit = True
if paired >= 10 and sign_p is not None and sign_p <= 0.05:
    hit = True
raise SystemExit(0 if hit else 1)
PY
}

count=0
tail -n +2 "$SHEET" | while IFS= read -r line; do
  count=$((count + 1))
  if [[ "$count" -le "$START_OFFSET" ]]; then
    continue
  fi
  done_count=$((count - START_OFFSET))
  if [[ "$LIMIT" != "0" && "$done_count" -gt "$LIMIT" ]]; then
    break
  fi

  sample=$(printf '%s\n' "$line" | cut -f1)
  run=$(printf '%s\n' "$line" | cut -f2)
  echo "$(date -Is) processing $run $sample" | tee -a "$ROOT/logs/closure_driver.log"

  ROOT_OVERRIDE="$ROOT" bash "$ROOT/scripts/02_download_fastqs.sh" "$SHEET" 1 $((count - 1)) >> "$ROOT/logs/closure_driver.log" 2>&1
  ROOT_OVERRIDE="$ROOT" bash "$ROOT/scripts/04_run_bismark_batch.sh" "$SHEET" 1 $((count - 1)) "$THREADS" 1 >> "$ROOT/logs/closure_driver.log" 2>&1
  ROOT_OVERRIDE="$ROOT" "$ROOT/tools/micromamba" run -p "$ROOT/env/bismark" python "$ROOT/scripts/05_aggregate_csb_dmrs.py" >> "$ROOT/logs/closure_driver.log" 2>&1

  if is_breakthrough; then
    cp "$ROOT/results/E-MTAB-10097_full_bismark_CSB_DMR_summary.json" "$ROOT/results/BREAKTHROUGH_summary.json"
    echo "$(date -Is) BREAKTHROUGH after $run" | tee -a "$ROOT/logs/closure_driver.log"
    exit 0
  fi
done

echo "$(date -Is) no breakthrough in requested batch" | tee -a "$ROOT/logs/closure_driver.log"
```
