## Supplemental code for "An entropy-based diagnostic framework for characterizing methylation state dynamics during preimplantation development": 07_wait_and_drive.sh

```
#!/usr/bin/env bash
set -euo pipefail

ROOT=${ROOT_OVERRIDE:-/home/u8068/bismark_full_closure}

while pgrep -fa "(/| )(bismark|bowtie2|bowtie2-align|samtools|deduplicate_bismark|bismark_methylation_extractor)( |$)" | grep -v -E "pgrep|06_drive_closure|07_wait_and_drive" >/dev/null; do
  sleep 300
done

if [[ -d /mnt/e/5_31_progress/bismark_full_closure/scripts ]]; then
  rsync -a /mnt/e/5_31_progress/bismark_full_closure/scripts/ "$ROOT/scripts/"
fi

cd "$ROOT"
ROOT_OVERRIDE="$ROOT" "$ROOT/tools/micromamba" run -p "$ROOT/env/bismark" python "$ROOT/scripts/05_aggregate_csb_dmrs.py" >> "$ROOT/logs/closure_driver.log" 2>&1 || true

offset=0
if [[ -s "$ROOT/results/ERR5354342/ERR5354342_1_bismark_bt2_pe.deduplicated.bismark.cov.gz" ]]; then
  offset=1
fi

ROOT_OVERRIDE="$ROOT" bash "$ROOT/scripts/06_drive_closure.sh" "$ROOT/samplesheet_E-MTAB-10097_highdepth_8ctrl_8dex_cap2GB.tsv" "$offset" 0 1

if [[ -s "$ROOT/results/BREAKTHROUGH_summary.json" ]]; then
  cat "$ROOT/results/BREAKTHROUGH_summary.json"
fi
```
