## Supplemental code for "An entropy-based diagnostic framework for characterizing methylation state dynamics during preimplantation development": 07_watchdog_closure.sh

```
#!/usr/bin/env bash
set -euo pipefail

ROOT=${ROOT_OVERRIDE:-/home/u8068/bismark_full_closure}
SHEET=${1:-$ROOT/samplesheet_E-MTAB-10097_highdepth_8ctrl_8dex_cap2GB.tsv}
THREADS=${2:-1}
SLEEP_SECONDS=${3:-300}

mkdir -p "$ROOT/logs" "$ROOT/results"
cd "$ROOT"

log() {
  echo "$(date -Is) $*" | tee -a "$ROOT/logs/closure_watchdog.log"
}

has_active_bismark() {
  pgrep -af '03_run_bismark_one|04_run_bismark_batch|06_drive_closure|bismark --genome|bowtie2-align|deduplicate_bismark|bismark_methylation_extractor|samtools view' \
    | grep -v '07_watchdog_closure' \
    | grep -v 'pgrep -af' >/dev/null 2>&1
}

aggregate_if_possible() {
  if find "$ROOT/results" -mindepth 2 -maxdepth 2 -name '*.bismark.cov.gz' | grep -q .; then
    ROOT_OVERRIDE="$ROOT" "$ROOT/tools/micromamba" run -p "$ROOT/env/bismark" \
      python "$ROOT/scripts/05_aggregate_csb_dmrs.py" \
      >> "$ROOT/logs/closure_watchdog.log" 2>&1 || true
  fi
}

is_breakthrough() {
  "$ROOT/tools/micromamba" run -p "$ROOT/env/bismark" python - <<'PY'
import json
from pathlib import Path

path = Path("results/E-MTAB-10097_full_bismark_CSB_DMR_summary.json")
if not path.exists():
    raise SystemExit(1)
s = json.loads(path.read_text())
paired = s.get("dmrs_with_control_and_dex_beta", 0) or 0
rho = s.get("spearman_latent_residual_vs_dex_delta_rho")
rho_p = s.get("spearman_latent_residual_vs_dex_delta_p")
sign_p = s.get("sign_concordance_binomial_p_greater")

hit = False
if paired >= 10 and rho is not None and abs(rho) >= 0.30 and rho_p is not None and rho_p <= 0.10:
    hit = True
if paired >= 10 and sign_p is not None and sign_p <= 0.05:
    hit = True
raise SystemExit(0 if hit else 1)
PY
}

first_missing_offset() {
  "$ROOT/tools/micromamba" run -p "$ROOT/env/bismark" python - "$SHEET" "$ROOT" <<'PY'
import sys
import pandas as pd
from pathlib import Path

sheet = pd.read_csv(sys.argv[1], sep="\t")
root = Path(sys.argv[2])
for i, row in sheet.iterrows():
    run = str(row["run"])
    if not list((root / "results" / run).glob("*.bismark.cov.gz")):
        print(i)
        raise SystemExit(0)
print("DONE")
PY
}

log "watchdog started sheet=$SHEET threads=$THREADS sleep=${SLEEP_SECONDS}s"

while true; do
  if has_active_bismark; then
    log "active Bismark/driver detected; waiting"
    sleep "$SLEEP_SECONDS"
    continue
  fi

  aggregate_if_possible

  if is_breakthrough; then
    cp "$ROOT/results/E-MTAB-10097_full_bismark_CSB_DMR_summary.json" "$ROOT/results/BREAKTHROUGH_summary.json"
    date -Is > "$ROOT/results/BREAKTHROUGH.flag"
    log "BREAKTHROUGH detected; stopping watchdog"
    exit 0
  fi

  offset=$(first_missing_offset)
  if [[ "$offset" == "DONE" ]]; then
    log "sheet complete without breakthrough; stopping watchdog"
    exit 0
  fi

  line_no=$((offset + 2))
  run=$(sed -n "${line_no}p" "$SHEET" | cut -f2)
  sample=$(sed -n "${line_no}p" "$SHEET" | cut -f1)
  log "resuming first missing cov offset=$offset run=$run sample=$sample"

  if ! ROOT_OVERRIDE="$ROOT" bash "$ROOT/scripts/02_download_fastqs.sh" "$SHEET" 1 "$offset" \
    >> "$ROOT/logs/closure_watchdog.log" 2>&1; then
    log "download failed for offset=$offset run=$run; retrying after sleep"
    sleep "$SLEEP_SECONDS"
    continue
  fi
  ROOT_OVERRIDE="$ROOT" bash "$ROOT/scripts/04_run_bismark_batch.sh" "$SHEET" 1 "$offset" "$THREADS" 1 \
    >> "$ROOT/logs/closure_watchdog.log" 2>&1 || true
done
```
