## Supplemental code for "An entropy-based diagnostic framework for characterizing methylation state dynamics during preimplantation development": 08_monitor_closure.sh

```
#!/usr/bin/env bash
set -euo pipefail

ROOT=${ROOT_OVERRIDE:-/home/u8068/bismark_full_closure}
INTERVAL=${1:-10}
RUN=${2:-ERR5354290}

summary_json="$ROOT/results/E-MTAB-10097_full_bismark_CSB_DMR_summary.json"

while true; do
  clear
  echo "E-MTAB-10097 Bismark Closure Monitor"
  echo "===================================="
  date
  echo

  echo "[Summary]"
  if [[ -s "$summary_json" ]]; then
    "$ROOT/tools/micromamba" run -p "$ROOT/env/bismark" python - "$summary_json" <<'PY'
import json
import sys
from pathlib import Path

s = json.loads(Path(sys.argv[1]).read_text())
print(f"processed_runs: {s.get('processed_runs')}")
print(f"paired_control_dex_dmrs: {s.get('dmrs_with_control_and_dex_beta')}")
print(f"control_cpg_calls: {s.get('control_total_cpg_calls')}")
print(f"dex_cpg_calls: {s.get('dex_total_cpg_calls')}")
print(f"spearman_rho: {s.get('spearman_latent_residual_vs_dex_delta_rho')}")
print(f"spearman_p: {s.get('spearman_latent_residual_vs_dex_delta_p')}")
print(f"sign_concordant_dmrs: {s.get('sign_concordant_dmrs')}")
print(f"sign_binomial_p: {s.get('sign_concordance_binomial_p_greater')}")
PY
  else
    echo "summary_json: missing"
  fi
  echo

  echo "[Cov Outputs]"
  find "$ROOT/results" -mindepth 2 -maxdepth 2 -name '*.bismark.cov.gz' 2>/dev/null \
    | sed 's#.*/results/##; s#/.*##' \
    | sort -u \
    | awk 'END {print "completed_cov_runs: " NR}'
  echo

  echo "[Disk]"
  df -h "$ROOT" | awk 'NR==1 || NR==2'
  echo

  echo "[Active Processes]"
  ps -eo pid,ppid,etime,pcpu,pmem,args \
    | grep -E '07_watchdog_closure|02_download_fastqs|curl|04_run_bismark_batch|03_run_bismark_one|bismark --genome|bowtie2-align|deduplicate_bismark|bismark_methylation_extractor|samtools view' \
    | grep -v grep \
    | grep -v '08_monitor_closure' \
    || echo "no active closure process"
  echo

  echo "[Current FASTQ: $RUN]"
  ls -lh "$ROOT/fastq/$RUN" 2>/dev/null || echo "no fastq directory for $RUN"
  echo

  echo "[Current Results: $RUN]"
  ls -lh "$ROOT/results/$RUN" 2>/dev/null || echo "no result directory for $RUN"
  echo

  echo "[Recent Watchdog Log]"
  tail -25 "$ROOT/logs/closure_watchdog.log" 2>/dev/null || echo "no watchdog log"
  echo
  echo "Refreshing every ${INTERVAL}s. Press Ctrl+C to exit."
  sleep "$INTERVAL"
done
```
