## Supplemental code for "An entropy-based diagnostic framework for characterizing methylation state dynamics during preimplantation development": 09_complete_remaining_dex.sh

```
#!/usr/bin/env bash
set -euo pipefail

ROOT=${ROOT_OVERRIDE:-/home/u8068/bismark_full_closure}
SHEET=${1:-samplesheet_E-MTAB-10097_highdepth_dex_first.tsv}
START_OFFSET=${2:-3}
END_OFFSET=${3:-7}
THREADS=${4:-1}

cd "$ROOT"
mkdir -p logs results

LOG="logs/robust_dex_completion_$(date +%Y%m%d_%H%M%S).log"
echo "robust_run_start $(date -Is) sheet=$SHEET offsets=${START_OFFSET}-${END_OFFSET}" | tee -a "$LOG"

for offset in $(seq "$START_OFFSET" "$END_OFFSET"); do
  line=$((offset + 2))
  run=$(sed -n "${line}p" "$SHEET" | cut -f2)
  sample=$(sed -n "${line}p" "$SHEET" | cut -f1)

  if [[ -z "$run" ]]; then
    echo "$(date -Is) empty run at offset=$offset; stopping" | tee -a "$LOG"
    break
  fi

  if ls "$ROOT/results/$run"/*.bismark.cov.gz >/dev/null 2>&1; then
    echo "$(date -Is) $run already_done" | tee -a "$LOG"
  else
    echo "$(date -Is) START offset=$offset run=$run sample=$sample" | tee -a "$LOG"
    if ! ROOT_OVERRIDE="$ROOT" DOWNLOAD_MAX_SECONDS="${DOWNLOAD_MAX_SECONDS:-1800}" \
      bash "$ROOT/scripts/02_download_fastqs.sh" "$SHEET" 1 "$offset" >> "$LOG" 2>&1; then
      echo "$(date -Is) SKIP_SLOW_DOWNLOAD offset=$offset run=$run sample=$sample" | tee -a "$LOG"
      continue
    fi
    ROOT_OVERRIDE="$ROOT" bash "$ROOT/scripts/04_run_bismark_batch.sh" "$SHEET" 1 "$offset" "$THREADS" 1 >> "$LOG" 2>&1
  fi

  ROOT_OVERRIDE="$ROOT" "$ROOT/tools/micromamba" run -p "$ROOT/env/bismark" \
    python "$ROOT/scripts/05_aggregate_csb_dmrs.py" >> "$LOG" 2>&1
  cp "$ROOT/results/E-MTAB-10097_full_bismark_CSB_DMR_summary.json" "$ROOT/results/latest_after_${run}.json"
  echo "$(date -Is) SUMMARY_AFTER $run" | tee -a "$LOG"
  cat "$ROOT/results/E-MTAB-10097_full_bismark_CSB_DMR_summary.json" | tee -a "$LOG"
done

echo "robust_run_done $(date -Is)" | tee -a "$LOG"
```
