## Supplemental code for "An entropy-based diagnostic framework for characterizing methylation state dynamics during preimplantation development": 10_autodl_fetch_and_run.ps1

```
$ErrorActionPreference = "Stop"

$Mutex = New-Object System.Threading.Mutex($false, "Global\E_MTAB_10097_AutoDL_Fetch_Run")
if (-not $Mutex.WaitOne(0, $false)) {
    exit 0
}

$Key = Join-Path $env:USERPROFILE ".ssh\autodl_bismark"
$RemoteUserHost = ""
$RemotePort = "23221"
$RemoteRoot = "/root/autodl-tmp/bismark_download"
$WinRoot = "E:\5_31_progress\bismark_full_closure"
$WslRoot = "/home/u8068/bismark_full_closure"
$WslKey = "/home/u8068/.ssh/autodl_bismark"
$Sheet = Join-Path $WinRoot "samplesheet_E-MTAB-10097_balanced_50to500MB.tsv"
$Log = Join-Path $WinRoot "logs\autodl_fetch_and_run.log"

New-Item -ItemType Directory -Force -Path (Join-Path $WinRoot "logs") | Out-Null
New-Item -ItemType Directory -Force -Path (Join-Path $WinRoot "fastq") | Out-Null

function Write-Log {
    param([string]$Message)
    $line = "{0} {1}" -f (Get-Date -Format "s"), $Message
    $line | Tee-Object -FilePath $Log -Append
}

function Invoke-Remote {
    param([string]$Command)
    $encoded = [Convert]::ToBase64String([Text.Encoding]::UTF8.GetBytes($Command))
    Invoke-Wsl "echo '$encoded' | base64 -d | ssh -o BatchMode=yes -o ConnectTimeout=20 -o ServerAliveInterval=15 -o ServerAliveCountMax=2 -i '$WslKey' -p '$RemotePort' '$RemoteUserHost' bash -s"
}

function Invoke-Wsl {
    param([string]$Command)
    & wsl.exe bash -lc $Command
}

function Test-WslCov {
    param([string]$Run)
    & wsl.exe bash -lc "ls '$WslRoot/results/$Run'/*.bismark.cov.gz >/dev/null 2>&1"
    return ($LASTEXITCODE -eq 0)
}

function Test-RemoteFastqReady {
    param([string]$Run)
    $cmd = "test -s '$RemoteRoot/fastq/$Run/${Run}_1.fastq.gz' && test -s '$RemoteRoot/fastq/$Run/${Run}_2.fastq.gz' && grep -Fq 'COMPLETE_RUN $Run' '$RemoteRoot/logs/download_balanced_v2.log' && echo READY || echo NOT_READY"
    $out = Invoke-Remote $cmd
    return (($out | Select-Object -Last 1) -eq "READY")
}

function Test-RemoteDownloaderActive {
    $out = Invoke-Remote "ps -ef | grep -E 'download_fastqs_autodl_v2|curl' | grep -v grep >/dev/null && echo ACTIVE || echo IDLE"
    return (($out | Select-Object -Last 1) -eq "ACTIVE")
}

function Start-RemoteDownloader {
    Write-Log "remote_downloader_restart"
    $cmd = "cd '$RemoteRoot'; screen -S bismark_download_v2 -X quit 2>/dev/null || true; screen -dmS bismark_download_v2 bash -lc 'cd $RemoteRoot; bash scripts/download_fastqs_autodl_v2.sh samplesheet_E-MTAB-10097_balanced_50to500MB.tsv 32 0 >> logs/download_balanced_v2.log 2>&1'; sleep 3; screen -ls"
    Invoke-Remote $cmd | Out-Null
}

function Sync-RunToWsl {
    param([string]$Run)
    Write-Log "scp_start run=$Run route=autodl_to_wsl"
    Invoke-Wsl "mkdir -p '$WslRoot/fastq/$Run'; scp -O -o BatchMode=yes -o ConnectTimeout=20 -o ServerAliveInterval=15 -o ServerAliveCountMax=2 -i '$WslKey' -P '$RemotePort' '${RemoteUserHost}:$RemoteRoot/fastq/$Run/${Run}_1.fastq.gz' '${RemoteUserHost}:$RemoteRoot/fastq/$Run/${Run}_2.fastq.gz' '$WslRoot/fastq/$Run/'"
    if ($LASTEXITCODE -ne 0) {
        throw "direct AutoDL-to-WSL scp failed for $Run"
    }
    Write-Log "scp_done run=$Run route=autodl_to_wsl"
    Invoke-Wsl "gzip -t '$WslRoot/fastq/$Run/${Run}_1.fastq.gz' && gzip -t '$WslRoot/fastq/$Run/${Run}_2.fastq.gz'"
    if ($LASTEXITCODE -ne 0) {
        throw "local gzip validation failed for $Run"
    }
}

function Run-BismarkOne {
    param([string]$Sample, [string]$Run)
    Write-Log "bismark_start run=$Run sample=$Sample"
    Invoke-Wsl "cd '$WslRoot'; ROOT_OVERRIDE='$WslRoot' bash scripts/03_run_bismark_one.sh '$Sample' '$Run' 1 1"
    if ($LASTEXITCODE -ne 0) {
        throw "Bismark failed for $Run"
    }
    Invoke-Wsl "cd '$WslRoot'; ROOT_OVERRIDE='$WslRoot' tools/micromamba run -p env/bismark python scripts/05_aggregate_csb_dmrs.py"
    if ($LASTEXITCODE -ne 0) {
        throw "Aggregation failed after $Run"
    }
    Invoke-Wsl "cd '$WslRoot'; cp results/E-MTAB-10097_full_bismark_CSB_DMR_summary.json results/latest_after_${Run}.json"
    Write-Log "bismark_done run=$Run"
}

$rows = Import-Csv -Path $Sheet -Delimiter "`t"
Write-Log "autodl_fetch_and_run_start rows=$($rows.Count)"

while ($true) {
    $processed = 0
    $madeProgress = $false

    foreach ($row in $rows) {
        $run = $row.run
        $sample = $row.sample

        if (Test-WslCov $run) {
            $processed += 1
            continue
        }

        if (Test-RemoteFastqReady $run) {
            Write-Log "remote_ready run=$run sample=$sample condition=$($row.condition)"
            Sync-RunToWsl $run
            Run-BismarkOne $sample $run
            $madeProgress = $true
            $processed += 1
        }
    }

    Write-Log "loop_status processed_or_existing=$processed total=$($rows.Count)"
    if ($processed -ge $rows.Count) {
        Write-Log "autodl_fetch_and_run_complete"
        break
    }

    if (-not (Test-RemoteDownloaderActive)) {
        Write-Log "remote_idle_detected"
        Start-RemoteDownloader
    }

    Start-Sleep -Seconds 300
}

$Mutex.ReleaseMutex()
```
