## Supplemental code for "An entropy-based diagnostic framework for characterizing methylation state dynamics during preimplantation development": 11_supervisor_watchdog.ps1

```
$ErrorActionPreference = "Continue"

$WinRoot = "E:\5_31_progress\bismark_full_closure"
$WslRoot = "/home/u8068/bismark_full_closure"
$RemoteRoot = "/root/autodl-tmp/bismark_download"
$RemoteUserHost = ""
$RemotePort = "23221"
$Key = Join-Path $env:USERPROFILE ".ssh\autodl_bismark"
$Controller = Join-Path $WinRoot "scripts\10_autodl_fetch_and_run.ps1"
$ControllerLog = Join-Path $WinRoot "logs\autodl_fetch_and_run.log"
$SupervisorLog = Join-Path $WinRoot "logs\supervisor_watchdog.log"
$IntervalSeconds = 600

New-Item -ItemType Directory -Force -Path (Join-Path $WinRoot "logs") | Out-Null

function Write-SupervisorLog {
    param([string]$Message)
    $line = "{0} {1}" -f (Get-Date -Format "s"), $Message
    $line | Tee-Object -FilePath $SupervisorLog -Append
}

function Invoke-WslText {
    param([string]$Command)
    $out = & wsl.exe bash -lc $Command 2>&1
    return ($out -join "`n")
}

function Invoke-RemoteText {
    param([string]$Command)
    $encoded = [Convert]::ToBase64String([Text.Encoding]::UTF8.GetBytes($Command))
    return Invoke-WslText "echo '$encoded' | base64 -d | ssh -o BatchMode=yes -o ConnectTimeout=20 -o ServerAliveInterval=15 -o ServerAliveCountMax=2 -i '/home/u8068/.ssh/autodl_bismark' -p '$RemotePort' '$RemoteUserHost' bash -s"
}

function Ensure-Controller {
    $procs = @(Get-CimInstance Win32_Process | Where-Object {
        $_.CommandLine -like "*10_autodl_fetch_and_run.ps1*" -and $_.CommandLine -notlike "*11_supervisor_watchdog*"
    })
    if ($procs.Count -eq 0) {
        Write-SupervisorLog "controller_absent restart"
        Start-Process powershell.exe -ArgumentList @("-NoProfile","-ExecutionPolicy","Bypass","-File",$Controller) -WindowStyle Hidden | Out-Null
    } elseif ($procs.Count -gt 1) {
        $keep = $procs | Sort-Object CreationDate -Descending | Select-Object -First 1
        $procs | Where-Object { $_.ProcessId -ne $keep.ProcessId } | ForEach-Object {
            Write-SupervisorLog "controller_duplicate_stop pid=$($_.ProcessId)"
            Stop-Process -Id $_.ProcessId -Force -ErrorAction SilentlyContinue
        }
    }
}

function Ensure-RemoteDownloader {
    $active = Invoke-RemoteText "screen -ls | grep -q bismark_download_v2 && ps -ef | grep -E 'download_fastqs_autodl_v2|curl|gzip' | grep -v grep >/dev/null && echo ACTIVE || echo IDLE"
    if ($active -notmatch "ACTIVE") {
        Write-SupervisorLog "remote_downloader_idle restart_from_remaining"
        Invoke-RemoteText "cd '$RemoteRoot'; screen -S bismark_download_v2 -X quit 2>/dev/null || true; screen -dmS bismark_download_v2 bash -lc 'cd $RemoteRoot; MAX_ATTEMPTS=2 DOWNLOAD_MAX_SECONDS=300 bash scripts/download_fastqs_autodl_v2.sh samplesheet_E-MTAB-10097_balanced_50to500MB.tsv 13 19 >> logs/download_balanced_v2.log 2>&1'; screen -ls" | Out-Null
    }
}

function Get-CovCount {
    $txt = Invoke-WslText "find '$WslRoot/results' -name '*.bismark.cov.gz' | wc -l"
    $nums = @($txt -split "`n" | ForEach-Object { $_.Trim() } | Where-Object { $_ -match "^\d+$" })
    if ($nums.Count -gt 0) { return [int]$nums[-1] }
    return -1
}

Write-SupervisorLog "supervisor_start interval=${IntervalSeconds}s"

while ($true) {
    try {
        Ensure-Controller
        Ensure-RemoteDownloader

        $cov = Get-CovCount
        $logAge = 999999
        if (Test-Path $ControllerLog) {
            $logAge = [int]((Get-Date) - (Get-Item $ControllerLog).LastWriteTime).TotalSeconds
        }
        $remoteTail = Invoke-RemoteText "grep -E 'COMPLETE_RUN|SKIP_RUN|FAILED_FILE|START' '$RemoteRoot/logs/download_balanced_v2.log' 2>/dev/null | tail -3"
        $remoteTailOneLine = $remoteTail -replace "`r?`n", " | "
        Write-SupervisorLog "heartbeat cov=$cov controller_log_age_s=$logAge remote_tail=$remoteTailOneLine"

        if ($logAge -gt 3600) {
            Write-SupervisorLog "controller_stale_age_s=$logAge restart"
            Get-CimInstance Win32_Process | Where-Object { $_.CommandLine -like "*10_autodl_fetch_and_run.ps1*" } | ForEach-Object {
                Stop-Process -Id $_.ProcessId -Force -ErrorAction SilentlyContinue
            }
            Start-Process powershell.exe -ArgumentList @("-NoProfile","-ExecutionPolicy","Bypass","-File",$Controller) -WindowStyle Hidden | Out-Null
        }
    } catch {
        Write-SupervisorLog "ERROR $($_.Exception.Message)"
    }

    Start-Sleep -Seconds $IntervalSeconds
}
```
