## Supplemental code for "An entropy-based diagnostic framework for characterizing methylation state dynamics during preimplantation development": 12_lineage_null_analysis.py

```
import json
import math
import os
from pathlib import Path

import numpy as np
import pandas as pd
from scipy import stats


ROOT = Path(os.environ.get("ROOT_OVERRIDE", "/home/u8068/bismark_full_closure"))
OUTDIR = ROOT / "results" / "lineage_null"
OUTDIR.mkdir(parents=True, exist_ok=True)

RNG = np.random.default_rng(20260605)


DATASETS = {
    "full_processed": {
        "sample_beta": ROOT / "results" / "E-MTAB-10097_full_bismark_CSB_DMR_sample_beta.tsv",
        "sheet": ROOT / "samplesheet_E-MTAB-10097_all359.tsv",
    },
    "balanced32_rescue": {
        "sample_beta": ROOT / "results" / "E-MTAB-10097_balanced_32_rescue_sample_beta.tsv",
        "sheet": ROOT / "samplesheet_E-MTAB-10097_balanced_32_rescue.tsv",
    },
}


def clean_lineage(x):
    if pd.isna(x) or str(x).strip() == "":
        return "unknown"
    return str(x).strip()


def build_delta(beta, min_total_per_condition=1):
    if beta.empty:
        return pd.DataFrame()
    agg = beta.groupby(["condition", "cluster_name"], as_index=False).agg(
        meth=("meth", "sum"),
        total=("total", "sum"),
        basin_residual_rank=("basin_residual_rank", "first"),
        module_id=("module_id", "first"),
        latent_residual_delta_beta=("latent_residual_delta_beta", "first"),
    )
    agg["beta"] = agg["meth"] / agg["total"].replace(0, np.nan)
    wide = agg.pivot(index="cluster_name", columns="condition", values="beta").reset_index()
    calls = agg.pivot(index="cluster_name", columns="condition", values="total").reset_index()
    meta = agg.drop_duplicates("cluster_name")[
        ["cluster_name", "basin_residual_rank", "module_id", "latent_residual_delta_beta"]
    ]
    wide = meta.merge(wide, on="cluster_name", how="left").merge(
        calls, on="cluster_name", how="left", suffixes=("", "_calls")
    )
    if "control" not in wide or "dex" not in wide:
        return pd.DataFrame()
    if "control_calls" not in wide:
        wide["control_calls"] = np.nan
    if "dex_calls" not in wide:
        wide["dex_calls"] = np.nan
    valid = wide.dropna(subset=["control", "dex"]).copy()
    valid = valid[
        (valid["control_calls"] >= min_total_per_condition)
        & (valid["dex_calls"] >= min_total_per_condition)
    ].copy()
    valid["dex_minus_control_beta"] = valid["dex"] - valid["control"]
    return valid


def stats_for_delta(delta):
    if delta.empty or "latent_residual_delta_beta" not in delta or "dex_minus_control_beta" not in delta:
        return np.nan, np.nan, 0, np.nan
    if len(delta) >= 3:
        rho, p = stats.spearmanr(
            delta["latent_residual_delta_beta"],
            delta["dex_minus_control_beta"],
            nan_policy="omit",
        )
    else:
        rho, p = np.nan, np.nan
    signed = delta[
        (delta["latent_residual_delta_beta"] != 0)
        & (delta["dex_minus_control_beta"] != 0)
    ]
    sign_match = int(
        (
            np.sign(signed["latent_residual_delta_beta"])
            == np.sign(signed["dex_minus_control_beta"])
        ).sum()
    )
    sign_p = (
        stats.binomtest(sign_match, len(signed), 0.5, alternative="greater").pvalue
        if len(signed) > 0
        else np.nan
    )
    return rho, p, sign_match, sign_p


def permute_condition_p(beta, observed_rho, min_total_per_condition, n_perm=2000):
    samples = beta[["sample", "run", "condition"]].drop_duplicates().reset_index(drop=True)
    if samples["condition"].nunique() != 2 or len(samples) < 4 or math.isnan(observed_rho):
        return np.nan, 0
    labels = samples["condition"].to_numpy().copy()
    runs = samples["run"].to_numpy()
    beta_base = beta.copy()
    abs_obs = abs(observed_rho)
    extreme = 0
    used = 0
    for _ in range(n_perm):
        perm = RNG.permutation(labels)
        label_map = dict(zip(runs, perm))
        beta_base["condition"] = beta_base["run"].map(label_map)
        delta = build_delta(beta_base, min_total_per_condition=min_total_per_condition)
        if len(delta) < 3:
            continue
        rho, _, _, _ = stats_for_delta(delta)
        if math.isnan(rho):
            continue
        used += 1
        if abs(rho) >= abs_obs:
            extreme += 1
    if used == 0:
        return np.nan, 0
    return (extreme + 1) / (used + 1), used


def analyze_dataset(name, cfg):
    beta = pd.read_csv(cfg["sample_beta"], sep="\t")
    sheet = pd.read_csv(cfg["sheet"], sep="\t")
    processed = set(beta["run"].drop_duplicates())
    sheet = sheet[sheet["run"].isin(processed)].copy()
    beta = beta[beta["run"].isin(set(sheet["run"]))].copy()
    beta["lineage_clean"] = beta["lineage"].map(clean_lineage)
    sheet["lineage_clean"] = sheet["lineage"].map(clean_lineage)

    subsets = {"all": beta}
    for lineage in sorted(beta["lineage_clean"].dropna().unique()):
        subsets[f"lineage:{lineage}"] = beta[beta["lineage_clean"].eq(lineage)].copy()

    rows = []
    delta_frames = []
    for subset_name, sub in subsets.items():
        sample_meta = sub[["run", "sample", "condition", "lineage_clean", "individual"]].drop_duplicates()
        n_control = int(sample_meta["condition"].eq("control").sum())
        n_dex = int(sample_meta["condition"].eq("dex").sum())
        for min_calls in [1, 3, 5, 10]:
            delta = build_delta(sub, min_total_per_condition=min_calls)
            rho, p, sign_match, sign_p = stats_for_delta(delta)
            perm_p, n_perm_used = permute_condition_p(
                sub, rho, min_calls, n_perm=2000 if subset_name == "all" else 1000
            )
            rows.append(
                {
                    "dataset": name,
                    "subset": subset_name,
                    "min_total_per_condition": min_calls,
                    "n_samples": int(sample_meta["run"].nunique()),
                    "n_control": n_control,
                    "n_dex": n_dex,
                    "paired_dmrs": int(len(delta)),
                    "rho": None if math.isnan(rho) else float(rho),
                    "spearman_p": None if math.isnan(p) else float(p),
                    "perm_p_two_sided_abs_rho": None if math.isnan(perm_p) else float(perm_p),
                    "n_perm_used": int(n_perm_used),
                    "sign_concordant_dmrs": int(sign_match),
                    "sign_concordance_binomial_p_greater": None
                    if math.isnan(sign_p)
                    else float(sign_p),
                }
            )
            if min_calls == 1 and not delta.empty:
                tmp = delta.copy()
                tmp["dataset"] = name
                tmp["subset"] = subset_name
                delta_frames.append(tmp)

    result = pd.DataFrame(rows)
    result.to_csv(OUTDIR / f"{name}_lineage_null_summary.tsv", sep="\t", index=False)
    if delta_frames:
        pd.concat(delta_frames, ignore_index=True).to_csv(
            OUTDIR / f"{name}_lineage_min1_delta.tsv", sep="\t", index=False
        )
    return result


def main():
    all_results = []
    for name, cfg in DATASETS.items():
        if not cfg["sample_beta"].exists() or not cfg["sheet"].exists():
            continue
        all_results.append(analyze_dataset(name, cfg))
    final = pd.concat(all_results, ignore_index=True)
    final.to_csv(OUTDIR / "lineage_null_summary_all.tsv", sep="\t", index=False)
    best = final.sort_values(["spearman_p", "perm_p_two_sided_abs_rho"], na_position="last").head(20)
    payload = {
        "analysis": "E-MTAB-10097 lineage stratified CSB DMR null tests",
        "n_tests": int(len(final)),
        "best_by_spearman_p": best.to_dict(orient="records"),
    }
    with open(OUTDIR / "lineage_null_summary_all.json", "w") as f:
        json.dump(payload, f, indent=2)
    print(json.dumps(payload, indent=2))


if __name__ == "__main__":
    main()
```
