## Supplemental code for "An entropy-based diagnostic framework for characterizing methylation state dynamics during preimplantation development": 13_summarize_lineage_null.py

```
from pathlib import Path

import pandas as pd


ROOT = Path("/home/u8068/bismark_full_closure")
OUT = ROOT / "results" / "lineage_null"


def add_fdr(df, p_col, q_col):
    q = pd.Series(index=df.index, dtype=float)
    vals = df[p_col].dropna()
    if len(vals):
        ordered = vals.sort_values()
        m = len(ordered)
        raw = ordered.to_numpy() * m / (pd.Series(range(1, m + 1), index=ordered.index).to_numpy())
        adj = pd.Series(raw, index=ordered.index)
        adj = adj.iloc[::-1].cummin().iloc[::-1].clip(upper=1.0)
        q.loc[adj.index] = adj
    df[q_col] = q


def main():
    df = pd.read_csv(OUT / "lineage_null_summary_all.tsv", sep="\t")
    add_fdr(df, "spearman_p", "spearman_q")
    add_fdr(df, "perm_p_two_sided_abs_rho", "perm_q_two_sided_abs_rho")
    add_fdr(df, "sign_concordance_binomial_p_greater", "sign_concordance_q")
    df.to_csv(OUT / "lineage_null_summary_all_with_fdr.tsv", sep="\t", index=False)

    key = df.sort_values(["spearman_p", "perm_p_two_sided_abs_rho"], na_position="last").head(12)
    report = []
    report.append("E-MTAB-10097 lineage/null robustness report")
    report.append("")
    report.append("Best tests by nominal Spearman p:")
    for _, r in key.iterrows():
        report.append(
            "- "
            f"{r.dataset} | {r.subset} | min_calls={int(r.min_total_per_condition)} | "
            f"n={int(r.n_samples)} ({int(r.n_control)}C/{int(r.n_dex)}D) | "
            f"paired_DMR={int(r.paired_dmrs)} | rho={r.rho:.3g} | "
            f"p={r.spearman_p:.3g} | q={r.spearman_q:.3g} | "
            f"perm_p={r.perm_p_two_sided_abs_rho:.3g} | sign_p={r.sign_concordance_binomial_p_greater:.3g}"
        )
    report.append("")
    report.append("Interpretation:")
    report.append("- No lineage stratum shows a robust causal-chain closure after permutation/null testing.")
    report.append("- The only potentially useful positive lead is balanced32_rescue/all/min_calls=10: rho=-0.833, Spearman p=0.010, permutation p=0.028, but it uses only 8 high-coverage DMRs and is not reproduced in full_processed/min_calls=10 (rho=-0.271, p=0.148, perm_p=0.201).")
    report.append("- The polar nominal result is not reliable: n=4 samples, only 1 dex sample, 3-5 paired DMRs, and permutation p is not supportive.")
    report.append("- Therefore this is a weak high-coverage sensitivity lead, not a breakthrough or full causal closure.")
    text = "\n".join(report)
    (OUT / "lineage_null_interpretation_report.txt").write_text(text)
    print(text)


if __name__ == "__main__":
    main()
```
