## Supplemental code for "An entropy-based diagnostic framework for characterizing methylation state dynamics during preimplantation development": download_fastqs_autodl_v2.sh

```
#!/usr/bin/env bash
set -euo pipefail

SHEET=${1:-samplesheet_E-MTAB-10097_balanced_50to500MB.tsv}
LIMIT=${2:-0}
OFFSET=${3:-0}
FASTQ=/root/autodl-tmp/bismark_download/fastq
MAX_ATTEMPTS=${MAX_ATTEMPTS:-5}
DOWNLOAD_MAX_SECONDS=${DOWNLOAD_MAX_SECONDS:-900}
mkdir -p "$FASTQ" logs

remote_size() {
  curl -fsSIL "$1" | awk 'tolower($1)=="content-length:" {gsub("\r","",$2); n=$2} END{print n+0}'
}

download_one() {
  local url=$1
  local out=$2
  local expected=$3
  local attempt=1

  while true; do
    local size=0
    [[ -s "$out" ]] && size=$(stat -c%s "$out")

    if [[ "$expected" -gt 0 && "$size" -eq "$expected" ]]; then
      if gzip -t "$out"; then
        echo "$(date -Is) COMPLETE_FILE $out bytes=$size"
        return 0
      fi
      echo "$(date -Is) CORRUPT_COMPLETE redownload $out bytes=$size expected=$expected" >&2
      rm -f "$out"
      size=0
    fi

    if [[ "$expected" -gt 0 && "$size" -gt "$expected" ]]; then
      echo "$(date -Is) OVERSIZE redownload $out bytes=$size expected=$expected" >&2
      rm -f "$out"
      size=0
    fi

    echo "$(date -Is) DOWNLOAD attempt=$attempt local_size=$size expected=$expected out=$out"
    if [[ "$size" -gt 0 ]]; then
      curl -f -L -C - --max-time "$DOWNLOAD_MAX_SECONDS" --connect-timeout 60 --speed-time 300 --speed-limit 1024 "$url" -o "$out" || true
    else
      curl -f -L --max-time "$DOWNLOAD_MAX_SECONDS" --connect-timeout 60 --speed-time 300 --speed-limit 1024 "$url" -o "$out" || true
    fi

    size=0
    [[ -s "$out" ]] && size=$(stat -c%s "$out")
    if [[ "$expected" -gt 0 && "$size" -eq "$expected" ]] && gzip -t "$out"; then
      echo "$(date -Is) COMPLETE_FILE $out bytes=$size"
      return 0
    fi

    if [[ "$attempt" -ge "$MAX_ATTEMPTS" ]]; then
      echo "$(date -Is) FAILED_FILE $out attempts=$attempt bytes=$size expected=$expected url=$url" >&2
      return 1
    fi
    attempt=$((attempt + 1))
    sleep 20
  done
}

count=0
tail -n +2 "$SHEET" | while IFS= read -r line; do
  sample=$(printf '%s\n' "$line" | cut -f1)
  run=$(printf '%s\n' "$line" | cut -f2)
  condition=$(printf '%s\n' "$line" | cut -f3)
  total_fastq_bytes=$(printf '%s\n' "$line" | cut -f8)
  fq1=$(printf '%s\n' "$line" | cut -f9)
  fq2=$(printf '%s\n' "$line" | cut -f10)
  count=$((count + 1))
  if [[ "$count" -le "$OFFSET" ]]; then
    continue
  fi
  done_count=$((count - OFFSET))
  if [[ "$LIMIT" != "0" && "$done_count" -gt "$LIMIT" ]]; then
    break
  fi

  mkdir -p "$FASTQ/$run"
  echo "$(date -Is) START $run $sample $condition bytes=$total_fastq_bytes"

  run_ok=1
  for fq in "$fq1" "$fq2"; do
    out="$FASTQ/$run/$(basename "$fq")"
    expected=$(remote_size "$fq")
    if ! download_one "$fq" "$out" "$expected"; then
      run_ok=0
      break
    fi
  done

  if [[ "$run_ok" -eq 1 ]]; then
    echo "$(date -Is) COMPLETE_RUN $run"
  else
    echo "$(date -Is) SKIP_RUN $run reason=download_failed"
  fi
done
```
