## Supplemental code for "An entropy-based diagnostic framework for characterizing methylation state dynamics during preimplantation development": prepare_full_bismark_closure.py

```
import pandas as pd
from pathlib import Path


ROOT = Path("/mnt/e/5_31_progress/bismark_full_closure")
META = Path("/mnt/e/5_31_progress/E-MTAB-10096_human_embryo_DEX/E-MTAB-10097.sdrf.txt")
RUNS = Path("/mnt/e/5_31_progress/E-MTAB-10096_human_embryo_DEX/ERP127251_runs.tsv")
DMR = Path("/mnt/e/实验进展5_27/CSB_TRO_2026-05-27_u_bio_rescue_DMR_overlap.tsv")


def main():
    meta = pd.read_csv(META, sep="\t").drop_duplicates("Comment[ENA_RUN]")
    runs = pd.read_csv(RUNS, sep="\t")
    runs["total_fastq_bytes"] = runs["fastq_bytes"].str.split(";").apply(lambda xs: sum(int(x) for x in xs))
    df = meta.merge(
        runs[["run_accession", "fastq_ftp", "fastq_bytes", "read_count", "total_fastq_bytes"]],
        left_on="Comment[ENA_RUN]",
        right_on="run_accession",
        how="left",
    )
    df["condition"] = df["Characteristics[stimulus]"].map({"Ctrl": "control", "Treat": "dex"})
    df["lineage"] = df["Characteristics[inferred lineage]"]
    df["individual"] = df["Characteristics[individual]"]
    df["sex"] = df["Characteristics[sex]"]
    df["sample"] = df["Source Name"]
    df["run"] = df["Comment[ENA_RUN]"]
    df = df[df["condition"].isin(["control", "dex"])].copy()
    df["fastq_1"] = df["fastq_ftp"].str.split(";").str[0].map(lambda x: "https://" + x if x.startswith("ftp.") else x.replace("ftp://", "https://"))
    df["fastq_2"] = df["fastq_ftp"].str.split(";").str[1].map(lambda x: "https://" + x if x.startswith("ftp.") else x.replace("ftp://", "https://"))
    keep = ["sample", "run", "condition", "lineage", "individual", "sex", "read_count", "total_fastq_bytes", "fastq_1", "fastq_2"]
    df[keep].to_csv(ROOT / "samplesheet_E-MTAB-10097_all359.tsv", sep="\t", index=False)
    # Prioritize informative cells first: OK lineage calls and enough reads.
    df["priority"] = 0
    df.loc[df["lineage"].isin(["mural", "polar", "epi", "pe"]), "priority"] += 1
    df.loc[df["read_count"].astype(float) >= 500000, "priority"] += 1
    df.sort_values(["priority", "total_fastq_bytes"], ascending=[False, True])[keep].to_csv(
        ROOT / "samplesheet_E-MTAB-10097_priority_order.tsv", sep="\t", index=False
    )
    dmr = pd.read_csv(DMR, sep="\t")
    dmr[["chr", "start", "end", "cluster_name", "basin_residual_rank", "latent_residual_delta_beta", "module_id"]].to_csv(
        ROOT / "CSB_TRO_156_residual_DMR_hg19.bed",
        sep="\t",
        header=False,
        index=False,
    )
    print(df[keep].shape)
    print(df.groupby(["condition", "lineage"]).size())
    print("total_fastq_GB", df["total_fastq_bytes"].sum() / 1e9)


if __name__ == "__main__":
    main()
```
