## Supplemental code for "An entropy-based diagnostic framework for characterizing methylation state dynamics during preimplantation development": 01_build_integrated_evidence_matrix.py

```
import json
import math
from pathlib import Path

import pandas as pd
from scipy.stats import combine_pvalues, norm


ROOT = Path(r"E:\5_31_progress\CSB_TRO_integrated_evidence")
if Path("/mnt/e").exists():
    ROOT = Path("/mnt/e/5_31_progress/CSB_TRO_integrated_evidence")
OUT = ROOT / "results"
OUT.mkdir(parents=True, exist_ok=True)


ROWS = [
    {
        "dataset": "E-MTAB-10097",
        "contrast": "dex_minus_control",
        "system": "human preimplantation embryo single-cell BS-seq",
        "assay": "BS-seq",
        "methylome": True,
        "perturbation": True,
        "target_specificity": "dex glucocorticoid perturbation; not p300/TET/DNMT-specific",
        "dmr_n": 92,
        "rho": -0.1436,
        "rho_p": 0.172,
        "sign_p": 0.768,
        "evidence_tier": "A: exact system+perturbation+methylome, weak result",
        "usable_claim": "Direct human embryo dex perturbation does not robustly align with CSB residual DMR direction.",
    },
    {
        "dataset": "GSE266195",
        "contrast": "DNMT3L_OE_minus_control",
        "system": "human trophoblast stem cell",
        "assay": "PBAT BS-seq 100-CpG windows",
        "methylome": True,
        "perturbation": True,
        "target_specificity": "DNMT3L methylation machinery perturbation",
        "dmr_n": 153,
        "rho": -0.07730556337367482,
        "rho_p": 0.3422151455759241,
        "sign_p": 0.025994426413430947,
        "evidence_tier": "A-: human hTSC methylation perturbation, global-effect caveat",
        "usable_claim": "DNMT3L OE methylates almost all CSB DMRs but the effect is largely genome-wide rather than CSB-specific.",
    },
    {
        "dataset": "GSE109682",
        "contrast": "SP_minus_CTB",
        "system": "human first-trimester trophoblast populations",
        "assay": "RRBS",
        "methylome": True,
        "perturbation": False,
        "target_specificity": "state comparison",
        "dmr_n": 71,
        "rho": 0.26118323485313877,
        "rho_p": 0.027803618471609587,
        "sign_p": 0.0018449821382700243,
        "evidence_tier": "B: human trophoblast state methylome, not perturbation; matched robustness weak",
        "usable_claim": "A trophoblast subpopulation state axis aligns with CSB residual DMRs in pooled RRBS, but matched-individual robustness weakened it.",
    },
    {
        "dataset": "GSE182015",
        "contrast": "hbdTSC_minus_hESC",
        "system": "human hbdTSC and hESC",
        "assay": "RRBS",
        "methylome": True,
        "perturbation": False,
        "target_specificity": "state comparison",
        "dmr_n": 37,
        "rho": -0.3405466444759841,
        "rho_p": 0.03916317321485477,
        "sign_p": 0.9505641252035275,
        "evidence_tier": "B-: human trophoblast state methylome, low DMR coverage",
        "usable_claim": "hbdTSC-vs-hESC methylation has a weak correlation signal at covered CSB DMRs, but sign test fails and coverage is low.",
    },
    {
        "dataset": "GSE150168",
        "contrast": "hTSC_minus_hESC",
        "system": "human hTSC/tdhTSC and hESC",
        "assay": "bisulfite-seq methylratio",
        "methylome": True,
        "perturbation": False,
        "target_specificity": "state comparison",
        "dmr_n": 111,
        "rho": -0.12237635083253899,
        "rho_p": 0.20070872743370452,
        "sign_p": 0.9357622839571201,
        "evidence_tier": "B-: human trophoblast-like state methylome, not residual-direction aligned",
        "usable_claim": "CSB DMRs are strongly hypomethylated in trophoblast state, but not aligned with latent residual direction.",
    },
    {
        "dataset": "GSE291172",
        "contrast": "SAC_120hr_minus_hPSC",
        "system": "human STAT3-induced embryo model",
        "assay": "WGBS bigWig",
        "methylome": True,
        "perturbation": False,
        "target_specificity": "embryo-model induction state",
        "dmr_n": 152,
        "rho": -0.001727391630096201,
        "rho_p": 0.9831491924369654,
        "sign_p": 0.8541727743327833,
        "evidence_tier": "B: human embryo-model methylome, null residual-direction result",
        "usable_claim": "Human embryo-model induction does not align with CSB residual DMR direction.",
    },
    {
        "dataset": "GSE126958",
        "contrast": "TET_DNMT3_perturbation_abs_delta",
        "system": "human HUES8 ESC",
        "assay": "WGBS",
        "methylome": True,
        "perturbation": True,
        "target_specificity": "TET/DNMT3 methylation machinery perturbation, wrong lineage",
        "dmr_n": 50,
        "rho": None,
        "rho_p": 0.7696230376962304,
        "sign_p": None,
        "evidence_tier": "C: mechanism perturbation in human ESC, not trophoblast",
        "usable_claim": "Human methylation-machinery perturbation did not enrich CSB top DMR methylation sensitivity beyond random CSB DMR sets.",
    },
]


def stouffer(ps, weights=None):
    clean = [(p, 1.0 if weights is None else w) for p, w in zip(ps, weights or [1] * len(ps)) if p is not None and p > 0 and p <= 1]
    if not clean:
        return None, None
    zsum = sum(norm.isf(p) * w for p, w in clean)
    denom = math.sqrt(sum(w * w for _, w in clean))
    z = zsum / denom
    return float(z), float(norm.sf(z))


def main():
    df = pd.DataFrame(ROWS)
    df.to_csv(OUT / "CSB_TRO_integrated_evidence_matrix.tsv", sep="\t", index=False)

    methyl_pert = df[(df["methylome"]) & (df["perturbation"])]
    human_troph_or_embryo = df[df["system"].str.contains("human") & ~df["system"].str.contains("HUES8")]
    all_methyl = df[df["methylome"]]

    summaries = {}
    for name, sub in [
        ("all_methylome_contrasts", all_methyl),
        ("human_trophoblast_or_embryo_contrasts", human_troph_or_embryo),
        ("methylome_perturbation_contrasts", methyl_pert),
    ]:
        pvals = [float(x) for x in sub["rho_p"].dropna()]
        if pvals:
            fisher_stat, fisher_p = combine_pvalues(pvals, method="fisher")
            z, stouffer_p = stouffer(pvals, list(sub["dmr_n"].fillna(1).astype(float)))
        else:
            fisher_stat = fisher_p = z = stouffer_p = None
        summaries[name] = {
            "n_contrasts": int(len(sub)),
            "min_rho_p": float(min(pvals)) if pvals else None,
            "fisher_combined_p_unoriented": float(fisher_p) if pvals else None,
            "weighted_stouffer_p_unoriented": stouffer_p,
            "note": "Unoriented p-value combination only tests whether any association exists across heterogeneous contrasts; it does not prove causal closure.",
        }

    claim = {
        "can_we_construct_a_scientifically_acceptable_dataset": True,
        "what_it_is": "A composite, pre-specified, orthogonal evidence meta-dataset at the CSB-TRO DMR level.",
        "what_it_is_not": "It is not a fabricated paired perturbation dataset and cannot be described as one experiment.",
        "strongest_publishable_claim": (
            "CSB-TRO residual DMRs can be evaluated across independent human trophoblast/embryo-model methylomes "
            "and methylation-machinery perturbation datasets. The integrated evidence supports trophoblast methylome involvement "
            "and DNMT3L responsiveness, but does not yet establish a single-dataset, paired, mechanism-specific causal closure."
        ),
        "not_allowed_claim": (
            "Do not claim that p300/CBP-A485 or TET/DNMT perturbation in human trophoblast directly causes the CSB residual DMR pattern "
            "unless a matching methylome perturbation dataset or new experiment is added."
        ),
        "meta": summaries,
    }
    (OUT / "CSB_TRO_integrated_evidence_summary.json").write_text(json.dumps(claim, indent=2), encoding="utf-8")
    (OUT / "CSB_TRO_integrated_evidence_claim.md").write_text(
        "# CSB-TRO Integrated Evidence Dataset\n\n"
        "This is a scientifically acceptable composite evidence dataset if presented as a pre-specified orthogonal evidence synthesis, not as a new experimental cohort.\n\n"
        "## Strongest Claim\n\n"
        f"{claim['strongest_publishable_claim']}\n\n"
        "## Boundary\n\n"
        f"{claim['not_allowed_claim']}\n\n",
        encoding="utf-8",
    )
    print(json.dumps(claim, indent=2))


if __name__ == "__main__":
    main()
```
