## Supplemental code for "An entropy-based diagnostic framework for characterizing methylation state dynamics during preimplantation development": targeted_bsseq_csb_dmr_pilot.py

```
import gzip
import json
import time
import urllib.request
from collections import defaultdict
from pathlib import Path

import numpy as np
import pandas as pd
from scipy import stats


ROOT = Path("E:/5_31_progress/E-MTAB-10096_human_embryo_DEX")
OUT = Path("E:/5_31_progress")
DMR = Path("E:/实验进展5_27/CSB_TRO_2026-05-27_u_bio_rescue_DMR_overlap.tsv")
SDRF = ROOT / "E-MTAB-10097.sdrf.txt"
RUNS = ROOT / "ERP127251_runs.tsv"

K = 26
STEP = 12
MAX_MISMATCH_RATE = 0.12
MAX_SAMPLES_PER_GROUP = 2
MIN_READS = 400_000


def rc(seq):
    tab = str.maketrans("ACGTNacgtn", "TGCANtgcan")
    return seq.translate(tab)[::-1].upper()


def fetch_ucsc(chrom, start, end):
    # CSB DMR coordinates are hg19-style; several intervals exceed hg38 chromosome lengths.
    url = f"https://api.genome.ucsc.edu/getData/sequence?genome=hg19;chrom={chrom};start={int(start)};end={int(end)}"
    for _ in range(3):
        try:
            with urllib.request.urlopen(url, timeout=30) as r:
                txt = r.read().decode("utf-8")
            data = json.loads(txt)
            return data["dna"].upper()
        except Exception:
            time.sleep(1)
    raise RuntimeError(url)


def load_dmrs():
    d = pd.read_csv(DMR, sep="\t").sort_values("basin_residual_rank").reset_index(drop=True)
    d = d.head(100).copy()
    d["dmr_id"] = d["cluster_name"]
    seqs = []
    for _, row in d.iterrows():
        seqs.append(fetch_ucsc(row["chr"], row["start"], row["end"]))
    d["seq"] = seqs
    d.to_csv(OUT / "E-MTAB-10097_targeted_all_CSB_DMR_sequences.tsv", sep="\t", index=False)
    return d


def build_index(dmrs):
    refs = []
    for _, row in dmrs.iterrows():
        for strand, seq in [("+", row["seq"]), ("-", rc(row["seq"]))]:
            conv = seq.replace("C", "T")
            ref_id = len(refs)
            refs.append({"dmr_id": row["dmr_id"], "strand": strand, "seq": seq, "conv": conv})
    idx = defaultdict(list)
    for ref_id, r in enumerate(refs):
        conv = r["conv"]
        for pos in range(0, max(0, len(conv) - K + 1), 1):
            kmer = conv[pos:pos + K]
            if "N" not in kmer:
                idx[kmer].append((ref_id, pos))
    return refs, idx


def select_samples():
    meta = pd.read_csv(SDRF, sep="\t").drop_duplicates("Comment[ENA_RUN]")
    runs = pd.read_csv(RUNS, sep="\t")
    runs["total_bytes"] = runs["fastq_bytes"].str.split(";").apply(lambda xs: sum(int(x) for x in xs))
    runs["read_count"] = runs["read_count"].astype(int)
    m = meta.merge(runs[["run_accession", "read_count", "total_bytes", "fastq_ftp"]],
                   left_on="Comment[ENA_RUN]", right_on="run_accession", how="left")
    m["condition"] = m["Characteristics[stimulus]"].map({"Ctrl": "control", "Treat": "dex"})
    m = m[m["condition"].isin(["control", "dex"])]
    m = m[m["Characteristics[inferred lineage]"].isin(["mural", "polar"])]
    m = m[m["read_count"] >= MIN_READS].copy()
    # Use moderate-sized runs first to get meaningful target coverage without reprocessing the full 359-cell dataset.
    m["size_rank"] = m["total_bytes"].rank(method="first")
    selected = []
    for condition in ["control", "dex"]:
        sub = m[m["condition"].eq(condition)].sort_values("total_bytes")
        selected.append(sub.head(MAX_SAMPLES_PER_GROUP))
    s = pd.concat(selected, ignore_index=True)
    s.to_csv(OUT / "E-MTAB-10097_targeted_bsseq_selected_samples.tsv", sep="\t", index=False)
    return s


def best_alignment(read, refs, idx):
    read = read.upper()[6:]
    if len(read) < 40:
        return None
    conv_read = read.replace("C", "T")
    candidates = defaultdict(int)
    for q in range(0, len(conv_read) - K + 1, STEP):
        kmer = conv_read[q:q + K]
        for ref_id, ref_pos in idx.get(kmer, []):
            start = ref_pos - q
            candidates[(ref_id, start)] += 1
    if not candidates:
        return None
    best = None
    best_score = 10**9
    for (ref_id, start), seed_hits in sorted(candidates.items(), key=lambda kv: -kv[1])[:25]:
        ref = refs[ref_id]["conv"]
        if start < -10 or start + len(conv_read) > len(ref) + 10:
            continue
        mism = 0
        comp = 0
        for i, b in enumerate(conv_read):
            rp = start + i
            if rp < 0 or rp >= len(ref):
                continue
            rb = ref[rp]
            if b == "N" or rb == "N":
                continue
            comp += 1
            if b != rb:
                mism += 1
        if comp < 35:
            continue
        rate = mism / comp
        score = rate - 0.01 * seed_hits
        if rate <= MAX_MISMATCH_RATE and score < best_score:
            best = (ref_id, start, read, comp, mism, seed_hits)
            best_score = score
    return best


def calls_from_alignment(aln, refs):
    ref_id, start, read, comp, mism, seed_hits = aln
    r = refs[ref_id]
    seq = r["seq"]
    calls = []
    for i, base in enumerate(read):
        rp = start + i
        if rp < 0 or rp >= len(seq) - 1:
            continue
        if seq[rp:rp + 2] == "CG":
            if base == "C":
                calls.append((r["dmr_id"], 1))
            elif base == "T":
                calls.append((r["dmr_id"], 0))
    return calls


def process_fastq_url(url, refs, idx):
    if url.startswith("ftp."):
        url = "https://" + url
    elif url.startswith("ftp://"):
        url = "https://" + url[6:]
    meth = defaultdict(int)
    total = defaultdict(int)
    n_reads = 0
    n_aln = 0
    try:
        with urllib.request.urlopen(url, timeout=120) as response:
            with gzip.GzipFile(fileobj=response) as gz:
                while True:
                    h = gz.readline()
                    if not h:
                        break
                    seq = gz.readline().decode("ascii", errors="ignore").strip()
                    gz.readline()
                    gz.readline()
                    n_reads += 1
                    aln = best_alignment(seq, refs, idx)
                    if aln is None:
                        continue
                    n_aln += 1
                    for dmr_id, m in calls_from_alignment(aln, refs):
                        meth[dmr_id] += m
                        total[dmr_id] += 1
    except (EOFError, TimeoutError, OSError) as exc:
        print(f"WARNING partial FASTQ stream for {url}: {exc}; keeping {n_reads} reads")
    return meth, total, n_reads, n_aln


def main():
    dmrs = load_dmrs()
    refs, idx = build_index(dmrs)
    samples = select_samples()
    rows = []
    for _, s in samples.iterrows():
        fastqs = str(s["fastq_ftp"]).split(";")
        smeth = defaultdict(int)
        stotal = defaultdict(int)
        reads = 0
        aligned = 0
        for fq in fastqs:
            meth, total, n_reads, n_aln = process_fastq_url(fq, refs, idx)
            reads += n_reads
            aligned += n_aln
            for k, v in meth.items():
                smeth[k] += v
            for k, v in total.items():
                stotal[k] += v
        for dmr_id in dmrs["dmr_id"]:
            rows.append({
                "sample": s["Source Name"],
                "run": s["Comment[ENA_RUN]"],
                "condition": s["condition"],
                "lineage": s["Characteristics[inferred lineage]"],
                "individual": s["Characteristics[individual]"],
                "dmr_id": dmr_id,
                "meth_cpg_calls": smeth[dmr_id],
                "total_cpg_calls": stotal[dmr_id],
                "beta": np.nan if stotal[dmr_id] == 0 else smeth[dmr_id] / stotal[dmr_id],
                "reads_scanned": reads,
                "target_aligned_reads": aligned,
            })
    out = pd.DataFrame(rows)
    out.to_csv(OUT / "E-MTAB-10097_targeted_bsseq_CSB_DMR_beta_pilot.tsv", sep="\t", index=False)
    agg = out.groupby(["condition", "dmr_id"], as_index=False).agg(
        meth=("meth_cpg_calls", "sum"),
        total=("total_cpg_calls", "sum"),
        n_samples=("sample", "nunique"),
    )
    agg["beta"] = agg["meth"] / agg["total"].replace(0, np.nan)
    wide = agg.pivot(index="dmr_id", columns="condition", values="beta").reset_index()
    cov = agg.pivot(index="dmr_id", columns="condition", values="total").reset_index()
    wide = wide.merge(cov, on="dmr_id", suffixes=("", "_calls"))
    if "control" in wide and "dex" in wide:
        wide["dex_minus_control_beta"] = wide["dex"] - wide["control"]
        valid = wide.dropna(subset=["dex_minus_control_beta"])
    else:
        valid = wide.iloc[0:0]
    wide.to_csv(OUT / "E-MTAB-10097_targeted_bsseq_CSB_DMR_delta_pilot.tsv", sep="\t", index=False)

    dmeta = dmrs[["dmr_id", "basin_residual_rank", "latent_residual_delta_beta", "abs_latent_residual_delta_beta", "module_id"]]
    test = valid.merge(dmeta, on="dmr_id", how="left")
    rho = stats.spearmanr(test["latent_residual_delta_beta"], test["dex_minus_control_beta"], nan_policy="omit") if len(test) >= 3 else (np.nan, np.nan)
    summary = {
        "analysis": "E-MTAB-10097_targeted_BSseq_CSB_DMR_pilot",
        "date": "2026-05-31",
        "closure_level": "pilot_targeted_human_embryo_perturbation_methylation",
        "dataset": "E-MTAB-10097",
        "design": "human E7 preimplantation embryo single-cell BS-seq, control vs dexamethasone",
        "selected_samples": int(samples["Comment[ENA_RUN]"].nunique()),
        "samples_by_condition": samples.groupby("condition")["Comment[ENA_RUN]"].nunique().to_dict(),
        "top_dmrs_targeted": int(len(dmrs)),
        "total_reads_scanned": int(out.drop_duplicates("run")["reads_scanned"].sum()),
        "total_target_aligned_reads": int(out.drop_duplicates("run")["target_aligned_reads"].sum()),
        "dmrs_with_control_and_dex_calls": int(len(test)),
        "total_cpg_calls_control": int(agg[agg["condition"].eq("control")]["total"].sum()),
        "total_cpg_calls_dex": int(agg[agg["condition"].eq("dex")]["total"].sum()),
        "mean_beta_control": float(agg[agg["condition"].eq("control")]["meth"].sum() / max(1, agg[agg["condition"].eq("control")]["total"].sum())),
        "mean_beta_dex": float(agg[agg["condition"].eq("dex")]["meth"].sum() / max(1, agg[agg["condition"].eq("dex")]["total"].sum())),
        "spearman_latent_residual_vs_dex_delta_rho": float(rho[0]) if len(test) >= 3 else None,
        "spearman_latent_residual_vs_dex_delta_p": float(rho[1]) if len(test) >= 3 else None,
        "boundary": (
            "This is a targeted pilot mapper over top50 CSB residual DMRs, not full Bismark genome-wide reprocessing. "
            "It directly uses human embryo paired perturbation BS-seq reads but should be treated as a feasibility/closure prototype."
        ),
    }
    with open(OUT / "E-MTAB-10097_targeted_bsseq_CSB_DMR_pilot_summary.json", "w", encoding="utf-8") as f:
        json.dump(summary, f, indent=2)
    print(json.dumps(summary, indent=2))


if __name__ == "__main__":
    main()
```
