## Supplemental code for "An entropy-based diagnostic framework for characterizing methylation state dynamics during preimplantation development": 01_pull_gse109682_from_autodl.ps1

```
$ErrorActionPreference = "Stop"

$Key = Join-Path $env:USERPROFILE ".ssh\autodl_bismark"
$Remote = ""
$Port = "23221"
$RemoteRoot = "/root/autodl-tmp/GSE109682"
$LocalRoot = "E:\5_31_progress\GSE109682_TRO_RRBS_closure"

New-Item -ItemType Directory -Force -Path (Join-Path $LocalRoot "raw\cpg_reports") | Out-Null
New-Item -ItemType Directory -Force -Path (Join-Path $LocalRoot "logs") | Out-Null

ssh -o BatchMode=yes -o ConnectTimeout=20 -i $Key -p $Port $Remote "grep -q 'ALL_COMPLETE' $RemoteRoot/logs/download_individual.log && test `$(find $RemoteRoot/cpg_reports -name '*.CpG_report.txt.gz' | wc -l) -eq 12"
if ($LASTEXITCODE -ne 0) {
    throw "Remote GSE109682 individual downloads are not complete yet."
}

scp -O -o BatchMode=yes -o ConnectTimeout=20 -i $Key -P $Port "${Remote}:$RemoteRoot/cpg_reports/*.CpG_report.txt.gz" (Join-Path $LocalRoot "raw\cpg_reports\")
if ($LASTEXITCODE -ne 0) {
    throw "scp failed"
}
Get-ChildItem (Join-Path $LocalRoot "raw\cpg_reports") -Filter "*.CpG_report.txt.gz" | Select-Object Name,Length | Format-Table | Out-String | Set-Content -Path (Join-Path $LocalRoot "logs\cpg_reports_pulled.txt")
```
