## Supplemental code for "An entropy-based diagnostic framework for characterizing methylation state dynamics during preimplantation development": 02_aggregate_gse109682_rrbs.py

```
import gzip
import json
import os
import re
import tarfile
from pathlib import Path

import numpy as np
import pandas as pd
from scipy import stats


ROOT = Path(os.environ.get("GSE109682_ROOT", r"E:\5_31_progress\GSE109682_TRO_RRBS_closure"))
DMR_PATH = Path(os.environ.get("CSB_DMR_PATH", r"E:\5_31_progress\bismark_full_closure\CSB_TRO_156_residual_DMR_hg19.bed"))
SHEET_PATH = ROOT / "samplesheet_GSE109682.tsv"
TAR_PATH = ROOT / "raw" / "GSE109682_RAW.tar"
REPORT_DIR = ROOT / "raw" / "cpg_reports"
OUT = ROOT / "results"
OUT.mkdir(parents=True, exist_ok=True)


def normalize_chrom(chrom):
    chrom = str(chrom).strip()
    if chrom.startswith("chr"):
        return chrom
    return f"chr{chrom}"


def load_dmrs():
    d = pd.read_csv(
        DMR_PATH,
        sep="\t",
        header=None,
        names=["chr", "start", "end", "cluster_name", "basin_residual_rank", "latent_residual_delta_beta", "module_id"],
    )
    d["chr"] = d["chr"].map(normalize_chrom)
    d = d.sort_values("basin_residual_rank").reset_index(drop=True)
    d["dmr_index"] = np.arange(len(d))
    return d


def build_intervals(dmr):
    intervals = {}
    for chrom, sub in dmr.groupby("chr"):
        s = sub.sort_values("start")
        intervals[chrom] = (s["start"].to_numpy(), s["end"].to_numpy(), s["dmr_index"].to_numpy())
    return intervals


def opener_from_tar(member, tar):
    f = tar.extractfile(member)
    if f is None:
        raise RuntimeError(f"Could not read {member.name}")
    if member.name.endswith(".gz"):
        return gzip.open(f, "rt")
    return (line.decode("utf-8", errors="replace") for line in f)


def opener_from_path(path):
    if str(path).endswith(".gz"):
        return gzip.open(path, "rt")
    return open(path, "rt")


def detect_columns(header):
    cols = [c.strip() for c in header.rstrip("\n").split("\t")]
    low = {c.lower(): i for i, c in enumerate(cols)}

    def pick(*names):
        for name in names:
            if name.lower() in low:
                return low[name.lower()]
        return None

    chrom = pick("chr", "chrom", "chromosome")
    start = pick("start", "base", "position", "pos")
    end = pick("end")
    strand = pick("strand")
    coverage = pick("coverage", "cov", "numcs")
    freq_c = pick("freqc", "meth", "percent_methylation", "methylation", "methylation_percentage")
    freq_t = pick("freqt", "unmeth")
    meth = pick("numCs", "meth_count", "methylated", "C_count")
    unmeth = pick("numTs", "unmeth_count", "unmethylated", "T_count")
    return {
        "cols": cols,
        "chrom": chrom,
        "start": start,
        "end": end,
        "strand": strand,
        "coverage": coverage,
        "freq_c": freq_c,
        "freq_t": freq_t,
        "meth": meth,
        "unmeth": unmeth,
    }


def parse_numeric(x):
    if x == "" or x.upper() == "NA":
        return np.nan
    return float(x)


def aggregate_stream(stream_name, fh, dmr):
    intervals = build_intervals(dmr)
    meth_sum = np.zeros(len(dmr), dtype=float)
    total_sum = np.zeros(len(dmr), dtype=float)
    seen_header = False
    col = None
    n_rows = 0
    n_used = 0
    for raw in fh:
        if isinstance(raw, bytes):
            raw = raw.decode("utf-8", errors="replace")
        if not raw.strip() or raw.startswith("#"):
            continue
        parts = raw.rstrip("\n").split("\t")
        if not seen_header:
            if any(re.search(r"[A-Za-z]", p) for p in parts[:4]):
                col = detect_columns(raw)
                seen_header = True
                continue
            is_cpg_report = "CpG_report" in stream_name or "cpg_report" in stream_name.lower()
            col = {
                "chrom": 0,
                "start": 1,
                "end": 2 if len(parts) >= 3 else None,
                "coverage": None,
                "freq_c": None,
                "meth": 3 if is_cpg_report and len(parts) >= 5 else (4 if len(parts) >= 6 else None),
                "unmeth": 4 if is_cpg_report and len(parts) >= 5 else (5 if len(parts) >= 6 else None),
            }
            seen_header = True
        if col["chrom"] is None or col["start"] is None:
            continue
        chrom = normalize_chrom(parts[col["chrom"]])
        if chrom not in intervals:
            continue
        try:
            pos0 = int(float(parts[col["start"]])) - 1
            if col.get("meth") is not None and col.get("unmeth") is not None:
                m = parse_numeric(parts[col["meth"]])
                u = parse_numeric(parts[col["unmeth"]])
                total = m + u
            elif col.get("coverage") is not None and col.get("freq_c") is not None:
                total = parse_numeric(parts[col["coverage"]])
                freq = parse_numeric(parts[col["freq_c"]])
                if freq > 1:
                    freq = freq / 100.0
                m = total * freq
            else:
                continue
        except (ValueError, IndexError):
            continue
        n_rows += 1
        if not np.isfinite(total) or total <= 0:
            continue
        starts, ends, idxs = intervals[chrom]
        j = np.searchsorted(starts, pos0 + 2, side="left")
        if j == 0:
            continue
        hits = np.where(ends[:j] > pos0)[0]
        for h in hits:
            idx = idxs[h]
            meth_sum[idx] += m
            total_sum[idx] += total
            n_used += 1
    return meth_sum, total_sum, {"member": stream_name, "rows_seen": n_rows, "dmr_hits": n_used}


def aggregate_member(tar, member, dmr):
    with opener_from_tar(member, tar) as fh:
        return aggregate_stream(member.name, fh, dmr)


def aggregate_path(path, dmr):
    with opener_from_path(path) as fh:
        return aggregate_stream(path.name, fh, dmr)


def match_members(tar, sheet):
    members = [m for m in tar.getmembers() if m.isfile()]
    mapping = {}
    for _, row in sheet.iterrows():
        candidates = [m for m in members if row["geo_accession"] in m.name or row["sample"].replace("_RRBS", "") in m.name]
        if len(candidates) == 1:
            mapping[row["sample"]] = candidates[0]
        elif len(candidates) > 1:
            txt = [m for m in candidates if re.search(r"\.txt(\.gz)?$", m.name)]
            mapping[row["sample"]] = txt[0] if txt else candidates[0]
    return mapping


def match_paths(sheet):
    paths = list(REPORT_DIR.glob("*.CpG_report.txt.gz")) + list(REPORT_DIR.glob("*.txt.gz")) + list(REPORT_DIR.glob("*.txt"))
    mapping = {}
    for _, row in sheet.iterrows():
        candidates = [p for p in paths if row["geo_accession"] in p.name or row["sample"].replace("_RRBS", "") in p.name]
        if len(candidates) >= 1:
            mapping[row["sample"]] = candidates[0]
    return mapping


def summarize_delta(sample_beta, dmr, left, right):
    sub = sample_beta[sample_beta["population"].isin([left, right])].copy()
    agg = sub.groupby(["population", "cluster_name"], as_index=False).agg(
        meth=("meth", "sum"),
        total=("total", "sum"),
        n_samples=("sample", "nunique"),
    )
    agg["beta"] = agg["meth"] / agg["total"].replace(0, np.nan)
    wide = agg.pivot(index="cluster_name", columns="population", values="beta").reset_index()
    calls = agg.pivot(index="cluster_name", columns="population", values="total").reset_index()
    meta = dmr[["cluster_name", "basin_residual_rank", "latent_residual_delta_beta", "module_id"]]
    wide = meta.merge(wide, on="cluster_name", how="left").merge(calls, on="cluster_name", how="left", suffixes=("", "_calls"))
    delta_col = f"{right}_minus_{left}_beta"
    if left in wide and right in wide:
        wide[delta_col] = wide[right] - wide[left]
    valid = wide.dropna(subset=[delta_col]) if delta_col in wide else wide.iloc[0:0]
    rho, p = (np.nan, np.nan)
    if len(valid) >= 3:
        rho, p = stats.spearmanr(valid["latent_residual_delta_beta"], valid[delta_col])
    sign_p = np.nan
    sign_match = 0
    if len(valid) > 0:
        signed = valid[(valid["latent_residual_delta_beta"] != 0) & (valid[delta_col] != 0)]
        if len(signed) > 0:
            sign_match = int((np.sign(signed["latent_residual_delta_beta"]) == np.sign(signed[delta_col])).sum())
            sign_p = stats.binomtest(sign_match, len(signed), 0.5, alternative="greater").pvalue
    return wide, {
        "contrast": f"{right}_minus_{left}",
        "dmrs_with_paired_beta": int(len(valid)),
        "spearman_rho": None if np.isnan(rho) else float(rho),
        "spearman_p": None if np.isnan(p) else float(p),
        "sign_concordant_dmrs": int(sign_match),
        "sign_concordance_binomial_p_greater": None if np.isnan(sign_p) else float(sign_p),
    }


def main():
    dmr = load_dmrs()
    sheet = pd.read_csv(SHEET_PATH, sep="\t")
    rows = []
    qc = []
    if REPORT_DIR.exists() and list(REPORT_DIR.glob("*.gz")):
        mapping = match_paths(sheet)
        missing = sorted(set(sheet["sample"]) - set(mapping))
        if missing:
            raise SystemExit(f"Could not match report files for samples: {missing}")
        for _, s in sheet.iterrows():
            path = mapping[s["sample"]]
            meth, total, q = aggregate_path(path, dmr)
            q.update(s.to_dict())
            qc.append(q)
            for i, row in dmr.iterrows():
                rows.append({
                    "geo_accession": s["geo_accession"],
                    "sample": s["sample"],
                    "population": s["population"],
                    "individual": s["individual"],
                    "cluster_name": row["cluster_name"],
                    "basin_residual_rank": row["basin_residual_rank"],
                    "latent_residual_delta_beta": row["latent_residual_delta_beta"],
                    "module_id": row["module_id"],
                    "meth": meth[i],
                    "total": total[i],
                    "beta": np.nan if total[i] == 0 else meth[i] / total[i],
                })
    elif TAR_PATH.exists():
        with tarfile.open(TAR_PATH, "r:*") as tar:
            mapping = match_members(tar, sheet)
            missing = sorted(set(sheet["sample"]) - set(mapping))
            if missing:
                raise SystemExit(f"Could not match tar members for samples: {missing}")
            for _, s in sheet.iterrows():
                member = mapping[s["sample"]]
                meth, total, q = aggregate_member(tar, member, dmr)
                q.update(s.to_dict())
                qc.append(q)
                for i, row in dmr.iterrows():
                    rows.append({
                        "geo_accession": s["geo_accession"],
                        "sample": s["sample"],
                        "population": s["population"],
                        "individual": s["individual"],
                        "cluster_name": row["cluster_name"],
                        "basin_residual_rank": row["basin_residual_rank"],
                        "latent_residual_delta_beta": row["latent_residual_delta_beta"],
                        "module_id": row["module_id"],
                        "meth": meth[i],
                        "total": total[i],
                        "beta": np.nan if total[i] == 0 else meth[i] / total[i],
                    })
    else:
        raise SystemExit(f"Missing {REPORT_DIR} reports or {TAR_PATH}")
    sample_beta = pd.DataFrame(rows)
    sample_beta.to_csv(OUT / "GSE109682_CSB_TRO_DMR_sample_beta.tsv", sep="\t", index=False)
    pd.DataFrame(qc).to_csv(OUT / "GSE109682_member_qc.tsv", sep="\t", index=False)

    summaries = []
    for left, right in [("CTB", "EVT"), ("CTB", "SP"), ("SP", "EVT")]:
        wide, summary = summarize_delta(sample_beta, dmr, left, right)
        wide.to_csv(OUT / f"GSE109682_CSB_TRO_DMR_{right}_minus_{left}.tsv", sep="\t", index=False)
        summaries.append(summary)
    out = {
        "analysis": "GSE109682_TRO_RRBS_CSB_residual_DMR_state_direction",
        "processed_samples": int(sample_beta["sample"].nunique()),
        "contrasts": summaries,
    }
    with open(OUT / "GSE109682_CSB_TRO_DMR_summary.json", "w") as f:
        json.dump(out, f, indent=2)
    print(json.dumps(out, indent=2))


if __name__ == "__main__":
    main()
```
