## Supplemental code for "An entropy-based diagnostic framework for characterizing methylation state dynamics during preimplantation development": 03_gse109682_controller.ps1

```
$ErrorActionPreference = "Continue"

$Key = Join-Path $env:USERPROFILE ".ssh\autodl_bismark"
$Remote = ""
$Port = "23221"
$RemoteRoot = "/root/autodl-tmp/GSE109682"
$LocalRoot = "E:\5_31_progress\GSE109682_TRO_RRBS_closure"
$Log = Join-Path $LocalRoot "logs\controller.log"
$DoneFlag = Join-Path $LocalRoot "results\GSE109682_CSB_TRO_DMR_summary.json"
$IntervalSeconds = 600

New-Item -ItemType Directory -Force -Path (Join-Path $LocalRoot "logs") | Out-Null
New-Item -ItemType Directory -Force -Path (Join-Path $LocalRoot "raw") | Out-Null
New-Item -ItemType Directory -Force -Path (Join-Path $LocalRoot "results") | Out-Null

function Write-Log {
    param([string]$Message)
    $line = "{0} {1}" -f (Get-Date -Format "s"), $Message
    $line | Tee-Object -FilePath $Log -Append
}

function Remote {
    param([string]$Command)
    ssh -o BatchMode=yes -o ConnectTimeout=20 -i $Key -p $Port $Remote $Command
}

function Remote-Complete {
    Remote "grep -q 'ALL_COMPLETE' $RemoteRoot/logs/download_individual.log && test `$(find $RemoteRoot/cpg_reports -name '*.CpG_report.txt.gz' | wc -l) -eq 12"
    return ($LASTEXITCODE -eq 0)
}

function Remote-Status {
    $txt = Remote "date; screen -ls | grep gse109682_individual || true; echo complete_marks=`$(grep -E 'COMPLETE |COMPLETE_EXISTING' $RemoteRoot/logs/download_individual.log 2>/dev/null | wc -l); echo files=`$(find $RemoteRoot/cpg_reports -name '*.CpG_report.txt.gz' 2>/dev/null | wc -l); du -sh $RemoteRoot/cpg_reports 2>/dev/null || true; grep -E 'START|COMPLETE|ALL_COMPLETE|curl:' $RemoteRoot/logs/download_individual.log 2>/dev/null | tail -8 || true"
    return ($txt -join " | ")
}

function Pull-And-Analyze {
    Write-Log "pull_start"
    powershell.exe -NoProfile -ExecutionPolicy Bypass -File (Join-Path $LocalRoot "scripts\01_pull_gse109682_from_autodl.ps1")
    if ($LASTEXITCODE -ne 0) {
        throw "pull failed"
    }
    Write-Log "analyze_start"
    wsl.exe bash -lc "cd /mnt/e/5_31_progress/GSE109682_TRO_RRBS_closure; GSE109682_ROOT=/mnt/e/5_31_progress/GSE109682_TRO_RRBS_closure CSB_DMR_PATH=/mnt/e/5_31_progress/bismark_full_closure/CSB_TRO_156_residual_DMR_hg19.bed /home/u8068/bismark_full_closure/tools/micromamba run -p /home/u8068/bismark_full_closure/env/bismark python scripts/02_aggregate_gse109682_rrbs.py > logs/aggregate.log 2>&1"
    if ($LASTEXITCODE -ne 0) {
        throw "analysis failed"
    }
    Write-Log "analyze_done"
}

Write-Log "controller_start"
while ($true) {
    try {
        if (Test-Path $DoneFlag) {
            Write-Log "already_done summary=$DoneFlag"
            break
        }
        if (Remote-Complete) {
            Write-Log "remote_complete"
            Pull-And-Analyze
            break
        }
        $status = Remote-Status
        Write-Log "heartbeat $status"
    } catch {
        Write-Log "ERROR $($_.Exception.Message)"
    }
    Start-Sleep -Seconds $IntervalSeconds
}
```
