## Supplemental code for "An entropy-based diagnostic framework for characterizing methylation state dynamics during preimplantation development": 04_matched_individual_robustness.py

```
import json
import math
import os
from pathlib import Path

import numpy as np
import pandas as pd
from scipy.stats import binomtest, spearmanr


ROOT = Path(os.environ.get("GSE109682_ROOT", r"E:\5_31_progress\GSE109682_TRO_RRBS_closure"))
OUT = ROOT / "results"
SAMPLE_BETA = OUT / "GSE109682_CSB_TRO_DMR_sample_beta.tsv"


def summarize_pair(df, left, right):
    rows = []
    for cluster, sub in df.groupby("cluster_name"):
        latent = sub["latent_residual_delta_beta"].iloc[0]
        module = sub["module_id"].iloc[0]
        rank = sub["basin_residual_rank"].iloc[0]
        deltas = []
        n_pairs = 0
        total_calls = 0.0
        for individual, part in sub.groupby("individual"):
            wide = part.pivot_table(index="individual", columns="population", values="beta", aggfunc="first")
            calls = part.pivot_table(index="individual", columns="population", values="total", aggfunc="first")
            if left not in wide.columns or right not in wide.columns:
                continue
            left_beta = wide[left].iloc[0]
            right_beta = wide[right].iloc[0]
            if not (np.isfinite(left_beta) and np.isfinite(right_beta)):
                continue
            left_calls = calls[left].iloc[0] if left in calls.columns else 0
            right_calls = calls[right].iloc[0] if right in calls.columns else 0
            if left_calls <= 0 or right_calls <= 0:
                continue
            deltas.append(right_beta - left_beta)
            n_pairs += 1
            total_calls += left_calls + right_calls
        if n_pairs:
            rows.append({
                "cluster_name": cluster,
                "module_id": module,
                "basin_residual_rank": rank,
                "latent_residual_delta_beta": latent,
                "n_individual_pairs": n_pairs,
                "total_calls": total_calls,
                f"{right}_minus_{left}_beta_mean_paired": float(np.mean(deltas)),
                f"{right}_minus_{left}_beta_median_paired": float(np.median(deltas)),
            })
    res = pd.DataFrame(rows)
    delta_col = f"{right}_minus_{left}_beta_mean_paired"
    if res.empty:
        return res, {
            "contrast": f"{right}_minus_{left}",
            "dmrs_with_matched_individual_delta": 0,
            "spearman_rho": None,
            "spearman_p": None,
            "sign_concordant_dmrs": 0,
            "sign_concordance_binomial_p_greater": None,
        }
    valid = res[np.isfinite(res[delta_col]) & np.isfinite(res["latent_residual_delta_beta"])]
    rho = p = None
    if len(valid) >= 3:
        rho, p = spearmanr(valid["latent_residual_delta_beta"], valid[delta_col])
        rho = None if math.isnan(rho) else float(rho)
        p = None if math.isnan(p) else float(p)
    concordant = int((np.sign(valid["latent_residual_delta_beta"]) == np.sign(valid[delta_col])).sum())
    binom_p = None
    if len(valid):
        binom_p = float(binomtest(concordant, len(valid), 0.5, alternative="greater").pvalue)
    return res, {
        "contrast": f"{right}_minus_{left}",
        "dmrs_with_matched_individual_delta": int(len(valid)),
        "median_individual_pairs_per_dmr": float(valid["n_individual_pairs"].median()) if len(valid) else None,
        "spearman_rho": rho,
        "spearman_p": p,
        "sign_concordant_dmrs": concordant,
        "sign_concordance_binomial_p_greater": binom_p,
    }


def main():
    df = pd.read_csv(SAMPLE_BETA, sep="\t")
    summaries = []
    for left, right in [("CTB", "EVT"), ("CTB", "SP"), ("SP", "EVT")]:
        res, summary = summarize_pair(df, left, right)
        res.to_csv(OUT / f"GSE109682_CSB_TRO_DMR_matched_{right}_minus_{left}.tsv", sep="\t", index=False)
        summaries.append(summary)
    out = {
        "analysis": "GSE109682_TRO_RRBS_CSB_residual_DMR_matched_individual_robustness",
        "contrasts": summaries,
    }
    (OUT / "GSE109682_CSB_TRO_DMR_matched_summary.json").write_text(json.dumps(out, indent=2), encoding="utf-8")
    print(json.dumps(out, indent=2))


if __name__ == "__main__":
    main()
```
