## Supplemental code for "An entropy-based diagnostic framework for characterizing methylation state dynamics during preimplantation development": download_gse109682_individual_autodl.sh

```
#!/usr/bin/env bash
set -euo pipefail

sheet=${1:-samplesheet_GSE109682_with_urls.tsv}
jobs=${2:-4}
root=/root/autodl-tmp/GSE109682
mkdir -p "$root/cpg_reports" "$root/logs"
cd "$root"

download_one() {
  local gsm="$1"
  local sample="$2"
  local url="$3"
  local filename="$4"
  local out="$root/cpg_reports/$filename"
  echo "$(date -Is) START $gsm $sample $filename"
  if [ -s "$out" ] && gzip -t "$out" >/dev/null 2>&1; then
    echo "$(date -Is) COMPLETE_EXISTING $gsm $sample $filename bytes=$(stat -c%s "$out")"
    return 0
  fi
  curl -fsSL -C - --retry 20 --retry-delay 10 --retry-all-errors "$url" -o "$out"
  gzip -t "$out"
  echo "$(date -Is) COMPLETE $gsm $sample $filename bytes=$(stat -c%s "$out")"
}

export -f download_one
export root

tail -n +2 "$sheet" |
  awk -F '\t' '{gsub(/\r/, ""); print $1 "\t" $2 "\t" $5 "\t" $6}' |
  xargs -P "$jobs" -n 4 bash -c 'download_one "$0" "$1" "$2" "$3"'

echo "$(date -Is) ALL_COMPLETE"
```
