## Supplemental code for "An entropy-based diagnostic framework for characterizing methylation state dynamics during preimplantation development": fix_remote_names_and_restart.sh

```
#!/usr/bin/env bash
set -euo pipefail

root=/root/autodl-tmp/GSE109682
cd "$root"

screen -S gse109682_individual -X quit >/dev/null 2>&1 || true

mkdir -p cpg_reports logs
find cpg_reports -maxdepth 1 -type f -print0 | while IFS= read -r -d '' path; do
  clean=$(printf '%s' "$path" | tr -d '\r')
  if [ "$clean" != "$path" ]; then
    if [ -e "$clean" ] && [ "$(stat -c%s "$clean")" -ge "$(stat -c%s "$path")" ]; then
      rm -f "$path"
    else
      mv -f "$path" "$clean"
    fi
    printf 'cleaned_name %q -> %q\n' "$path" "$clean"
  fi
done

bash "$root/start_gse109682_individual_remote.sh"
```
