## Supplemental code for "An entropy-based diagnostic framework for characterizing methylation state dynamics during preimplantation development": start_gse109682_individual_remote.sh

```
#!/usr/bin/env bash
set -euo pipefail

root=/root/autodl-tmp/GSE109682
cd "$root"

pkill -f GSE109682_RAW.tar 2>/dev/null || true
screen -S gse109682_individual -X quit >/dev/null 2>&1 || true
chmod +x download_gse109682_individual_autodl.sh

if [ -s logs/download_individual.log ]; then
  mv logs/download_individual.log "logs/download_individual.$(date +%Y%m%d_%H%M%S).log"
fi

screen -dmS gse109682_individual bash -lc \
  'cd /root/autodl-tmp/GSE109682; ./download_gse109682_individual_autodl.sh samplesheet_GSE109682_with_urls.tsv 4 >> logs/download_individual.log 2>&1'

screen -ls
sleep 3
tail -30 logs/download_individual.log || true
```
