## Supplemental code for "An entropy-based diagnostic framework for characterizing methylation state dynamics during preimplantation development": start_gse109682_remote.sh

```
#!/usr/bin/env bash
set -euo pipefail

root=/root/autodl-tmp/GSE109682
mkdir -p "$root/raw" "$root/logs"
cd "$root"

pkill -f GSE109682_RAW.tar 2>/dev/null || true

nohup bash -lc '
set -euo pipefail
cd /root/autodl-tmp/GSE109682
curl -fL -C - --retry 20 --retry-delay 10 --retry-all-errors \
  https://ftp.ncbi.nlm.nih.gov/geo/series/GSE109nnn/GSE109682/suppl/GSE109682_RAW.tar \
  -o raw/GSE109682_RAW.tar
rc=$?
echo "download_exit=$rc"
if [ "$rc" -eq 0 ]; then
  tar -tf raw/GSE109682_RAW.tar | head -120 > logs/tar_head.txt
fi
' > "$root/logs/download.log" 2>&1 &

echo "$!" > "$root/logs/download.pid"
echo "started_pid=$(cat "$root/logs/download.pid")"
ps -p "$(cat "$root/logs/download.pid")" -o pid,etime,args
```
