## Supplemental code for "An entropy-based diagnostic framework for characterizing methylation state dynamics during preimplantation development": 01_aggregate_gse150168_csb_dmrs.py

```
import gzip
import json
import math
import os
from collections import defaultdict
from pathlib import Path

import numpy as np
import pandas as pd
from scipy.stats import binomtest, spearmanr


ROOT = Path(os.environ.get("GSE150168_ROOT", r"E:\5_31_progress\GSE150168_naive_hESC_trophoblast_methylome"))
DMR_PATH = Path(os.environ.get(
    "CSB_DMR_PATH", r"E:\5_31_progress\bismark_full_closure\CSB_TRO_156_residual_DMR_hg19.bed"))
if Path("/mnt/e").exists() and "GSE150168_ROOT" not in os.environ:
    ROOT = Path("/mnt/e/5_31_progress/GSE150168_naive_hESC_trophoblast_methylome")
if Path("/mnt/e").exists() and "CSB_DMR_PATH" not in os.environ:
    DMR_PATH = Path("/mnt/e/5_31_progress/bismark_full_closure/CSB_TRO_156_residual_DMR_hg19.bed")

RAW = ROOT / "raw"
OUT = ROOT / "results"
OUT.mkdir(parents=True, exist_ok=True)

SAMPLES = {
    "CT1_hTSC": "GSM4525517_CT1_hTSC_Replicate_2_allchr_methratio_steve_reduced_CG.txt.gz",
    "CT3_hTSC": "GSM4525518_CT3_hTSC_Replicate_2_allchr_methratio_steve_reduced_CG.txt.gz",
    "WIBR3_hESC": "GSM4525519_WIBR3_Primed_hESC_Replicate_2_allchr_methratio_steve_reduced_CG.txt.gz",
    "tdhTSC_L1": "GSM4525520_WIBR3_tdhTSC_Line_1_Replicate_2_allchr_methratio_steve_reduced_CG.txt.gz",
    "tdhTSC_L2": "GSM4525521_WIBR3_tdhTSC_Line_2_Replicate_2_allchr_methratio_steve_reduced_CG.txt.gz",
}
GROUPS = {
    "hTSC": ["CT1_hTSC", "CT3_hTSC"],
    "hESC": ["WIBR3_hESC"],
    "tdhTSC": ["tdhTSC_L1", "tdhTSC_L2"],
}


def normalize_chrom(chrom):
    chrom = str(chrom).strip()
    return chrom if chrom.startswith("chr") else f"chr{chrom}"


def load_dmrs():
    d = pd.read_csv(
        DMR_PATH,
        sep="\t",
        header=None,
        names=["chr", "start", "end", "cluster_name", "basin_residual_rank", "latent_residual_delta_beta", "module_id"],
    )
    d["chr"] = d["chr"].map(normalize_chrom)
    d["dmr_id"] = np.arange(len(d))
    return d


def build_intervals(dmr):
    intervals = defaultdict(list)
    for r in dmr.itertuples(index=False):
        intervals[r.chr].append((int(r.start), int(r.end), int(r.dmr_id)))
    for chrom in intervals:
        intervals[chrom].sort()
    return intervals


def scan_sample(path, intervals):
    meth = defaultdict(int)
    total = defaultdict(int)
    rows_seen = 0
    hits = 0
    with gzip.open(path, "rt") as fh:
        for line in fh:
            rows_seen += 1
            fields = line.rstrip("\n").split("\t")
            if len(fields) < 6:
                continue
            chrom = normalize_chrom(fields[0])
            if chrom not in intervals:
                continue
            try:
                pos = int(fields[1])
                m = int(float(fields[4]))
                t = int(float(fields[5]))
            except ValueError:
                continue
            if t <= 0:
                continue
            # Input positions are 1-based; BED DMRs are 0-based half-open.
            bed_pos = pos - 1
            for start, end, dmr_id in intervals[chrom]:
                if end <= bed_pos:
                    continue
                if start > bed_pos:
                    break
                meth[dmr_id] += m
                total[dmr_id] += t
                hits += 1
    return meth, total, rows_seen, hits


def beta(m, t):
    return float(m) / float(t) if t else np.nan


def group_beta(sample_stats, group, dmr_id):
    m = sum(sample_stats[s]["meth"].get(dmr_id, 0) for s in GROUPS[group])
    t = sum(sample_stats[s]["total"].get(dmr_id, 0) for s in GROUPS[group])
    return beta(m, t), m, t


def summarize(out, delta_col, label):
    valid = out[np.isfinite(out[delta_col]) & np.isfinite(out["latent_residual_delta_beta"])]
    rho = p = None
    if len(valid) >= 3:
        rho, p = spearmanr(valid["latent_residual_delta_beta"], valid[delta_col])
        rho = None if math.isnan(rho) else float(rho)
        p = None if math.isnan(p) else float(p)
    concordant = int((np.sign(valid["latent_residual_delta_beta"]) == np.sign(valid[delta_col])).sum())
    binom_p = float(binomtest(concordant, len(valid), 0.5, alternative="greater").pvalue) if len(valid) else None
    return {
        f"{label}_dmrs": int(len(valid)),
        f"{label}_mean_delta_beta": float(valid[delta_col].mean()) if len(valid) else None,
        f"{label}_median_delta_beta": float(valid[delta_col].median()) if len(valid) else None,
        f"{label}_spearman_rho": rho,
        f"{label}_spearman_p": p,
        f"{label}_sign_concordant_dmrs": concordant,
        f"{label}_sign_concordance_binomial_p_greater": binom_p,
    }


def main():
    dmr = load_dmrs()
    intervals = build_intervals(dmr)
    sample_stats = {}
    qc = []
    for sample, fname in SAMPLES.items():
        meth, total, rows_seen, hits = scan_sample(RAW / fname, intervals)
        sample_stats[sample] = {"meth": meth, "total": total}
        qc.append({"sample": sample, "rows_seen": rows_seen, "dmr_cpg_hits": hits})

    rows = []
    for r in dmr.itertuples(index=False):
        hesc_b, hesc_m, hesc_t = group_beta(sample_stats, "hESC", r.dmr_id)
        htsc_b, htsc_m, htsc_t = group_beta(sample_stats, "hTSC", r.dmr_id)
        tdh_b, tdh_m, tdh_t = group_beta(sample_stats, "tdhTSC", r.dmr_id)
        rows.append({
            "chr": r.chr,
            "start": r.start,
            "end": r.end,
            "cluster_name": r.cluster_name,
            "basin_residual_rank": r.basin_residual_rank,
            "latent_residual_delta_beta": r.latent_residual_delta_beta,
            "module_id": r.module_id,
            "hESC_beta": hesc_b,
            "hESC_total": hesc_t,
            "hTSC_beta": htsc_b,
            "hTSC_total": htsc_t,
            "tdhTSC_beta": tdh_b,
            "tdhTSC_total": tdh_t,
            "hTSC_minus_hESC_beta": htsc_b - hesc_b if np.isfinite(htsc_b) and np.isfinite(hesc_b) else np.nan,
            "tdhTSC_minus_hESC_beta": tdh_b - hesc_b if np.isfinite(tdh_b) and np.isfinite(hesc_b) else np.nan,
        })
    out = pd.DataFrame(rows)
    out.to_csv(OUT / "GSE150168_CSB_TRO_DMR_group_beta.tsv", sep="\t", index=False)
    pd.DataFrame(qc).to_csv(OUT / "GSE150168_member_qc.tsv", sep="\t", index=False)

    summary = {
        "analysis": "GSE150168_naive_hESC_to_trophoblast_like_methylome_CSB_residual_DMR",
        "processed_samples": len(SAMPLES),
        "boundary": "Human hTSC/tdhTSC state comparison, not paired methylome perturbation.",
    }
    summary.update(summarize(out, "hTSC_minus_hESC_beta", "hTSC_minus_hESC"))
    summary.update(summarize(out, "tdhTSC_minus_hESC_beta", "tdhTSC_minus_hESC"))
    (OUT / "GSE150168_CSB_TRO_DMR_summary.json").write_text(json.dumps(summary, indent=2), encoding="utf-8")
    print(json.dumps(summary, indent=2))


if __name__ == "__main__":
    main()
```
