## Supplemental code for "An entropy-based diagnostic framework for characterizing methylation state dynamics during preimplantation development": 01_aggregate_gse182015_csb_dmrs.py

```
import gzip
import json
import math
import os
from collections import defaultdict
from pathlib import Path

import numpy as np
import pandas as pd
from scipy.stats import binomtest, spearmanr


ROOT = Path(os.environ.get("GSE182015_ROOT", r"E:\5_31_progress\GSE182015_hiTSC_RRBS_closure"))
DMR_PATH = Path(os.environ.get(
    "CSB_DMR_PATH", r"E:\5_31_progress\bismark_full_closure\CSB_TRO_156_residual_DMR_hg19.bed"))
if Path("/mnt/e").exists() and "GSE182015_ROOT" not in os.environ:
    ROOT = Path("/mnt/e/5_31_progress/GSE182015_hiTSC_RRBS_closure")
if Path("/mnt/e").exists() and "CSB_DMR_PATH" not in os.environ:
    DMR_PATH = Path("/mnt/e/5_31_progress/bismark_full_closure/CSB_TRO_156_residual_DMR_hg19.bed")

RAW = ROOT / "raw"
OUT = ROOT / "results"
OUT.mkdir(parents=True, exist_ok=True)

GROUPS = {
    "hbdTSC": [
        "GSM5516237_hbdTSC_E2_rep1.CpG.bed.gz",
        "GSM5516238_hbdTSC_E2_rep2.CpG.bed.gz",
        "GSM5516239_hbdTSC_E2_rep3.CpG.bed.gz",
        "GSM5516240_hbdTSC_E9_rep1.CpG.bed.gz",
        "GSM5516241_hbdTSC_E9_rep2.CpG.bed.gz",
        "GSM5516242_hbdTSC_E9_rep3.CpG.bed.gz",
    ],
    "hESC": [
        "GSM5516249_hESCs_rep1.CpG.bed.gz",
        "GSM5516250_hESCs_rep2.CpG.bed.gz",
        "GSM5516251_hESCs_rep3.CpG.bed.gz",
    ],
    "hiTSC": [
        "GSM5516252_KEN_hiTSC_C1_rep1.CpG.bed.gz",
        "GSM5516253_KEN_hiTSC_C1_rep2.CpG.bed.gz",
        "GSM5516254_KEN_hiTSC_C1_rep3.CpG.bed.gz",
        "GSM5516255_KEN_hiTSC_C2_rep1.CpG.bed.gz",
        "GSM5516256_KEN_hiTSC_C2_rep2.CpG.bed.gz",
        "GSM5516257_KEN_hiTSC_C2_rep3.CpG.bed.gz",
        "GSM5516258_KEN_hiTSC_C4_rep1.CpG.bed.gz",
        "GSM5516259_KEN_hiTSC_C4_rep2.CpG.bed.gz",
        "GSM5516260_KEN_hiTSC_C4_rep3.CpG.bed.gz",
        "GSM5516261_PCS_hiTSC_C11_rep1.CpG.bed.gz",
        "GSM5516262_PCS_hiTSC_C11_rep2.CpG.bed.gz",
        "GSM5516263_PCS_hiTSC_C11_rep3.CpG.bed.gz",
        "GSM5516264_GM2_hiTSCs_C16_rep1.CpG.bed.gz",
        "GSM5516265_GM2_hiTSCs_C16_rep2.CpG.bed.gz",
        "GSM5516266_GM2_hiTSCs_C16_rep3.CpG.bed.gz",
    ],
}


def normalize_chrom(chrom):
    chrom = str(chrom).strip()
    return chrom if chrom.startswith("chr") else f"chr{chrom}"


def load_dmrs():
    d = pd.read_csv(
        DMR_PATH,
        sep="\t",
        header=None,
        names=["chr", "start", "end", "cluster_name", "basin_residual_rank", "latent_residual_delta_beta", "module_id"],
    )
    d["chr"] = d["chr"].map(normalize_chrom)
    d["dmr_id"] = np.arange(len(d))
    return d


def build_intervals(dmr):
    intervals = defaultdict(list)
    for r in dmr.itertuples(index=False):
        intervals[r.chr].append((int(r.start), int(r.end), int(r.dmr_id)))
    for chrom in intervals:
        intervals[chrom].sort()
    return intervals


def scan_group(files, intervals):
    meth = defaultdict(int)
    total = defaultdict(int)
    rows_seen = 0
    hits = 0
    for fname in files:
        with gzip.open(RAW / fname, "rt") as fh:
            for line in fh:
                rows_seen += 1
                fields = line.rstrip("\n").split("\t")
                if len(fields) < 8:
                    continue
                chrom = normalize_chrom(fields[0])
                if chrom not in intervals:
                    continue
                try:
                    pos = int(fields[1])
                    m = int(float(fields[6]))
                    t = int(float(fields[7]))
                except ValueError:
                    continue
                if t <= 0:
                    continue
                for start, end, dmr_id in intervals[chrom]:
                    if end <= pos:
                        continue
                    if start > pos:
                        break
                    meth[dmr_id] += m
                    total[dmr_id] += t
                    hits += 1
    return meth, total, rows_seen, hits


def beta(m, t):
    return float(m) / float(t) if t else np.nan


def summarize(out, delta_col, label):
    valid = out[np.isfinite(out[delta_col]) & np.isfinite(out["latent_residual_delta_beta"])]
    rho = p = None
    if len(valid) >= 3:
        rho, p = spearmanr(valid["latent_residual_delta_beta"], valid[delta_col])
        rho = None if math.isnan(rho) else float(rho)
        p = None if math.isnan(p) else float(p)
    concordant = int((np.sign(valid["latent_residual_delta_beta"]) == np.sign(valid[delta_col])).sum())
    binom_p = float(binomtest(concordant, len(valid), 0.5, alternative="greater").pvalue) if len(valid) else None
    return {
        f"{label}_dmrs": int(len(valid)),
        f"{label}_mean_delta_beta": float(valid[delta_col].mean()) if len(valid) else None,
        f"{label}_median_delta_beta": float(valid[delta_col].median()) if len(valid) else None,
        f"{label}_spearman_rho": rho,
        f"{label}_spearman_p": p,
        f"{label}_sign_concordant_dmrs": concordant,
        f"{label}_sign_concordance_binomial_p_greater": binom_p,
    }


def main():
    dmr = load_dmrs()
    intervals = build_intervals(dmr)
    stats = {}
    qc = []
    for group, files in GROUPS.items():
        meth, total, rows_seen, hits = scan_group(files, intervals)
        stats[group] = {"meth": meth, "total": total}
        qc.append({"group": group, "files": len(files), "rows_seen": rows_seen, "dmr_cpg_hits": hits})

    rows = []
    for r in dmr.itertuples(index=False):
        vals = {}
        for group in GROUPS:
            m = stats[group]["meth"].get(r.dmr_id, 0)
            t = stats[group]["total"].get(r.dmr_id, 0)
            vals[f"{group}_beta"] = beta(m, t)
            vals[f"{group}_total"] = t
        rows.append({
            "chr": r.chr,
            "start": r.start,
            "end": r.end,
            "cluster_name": r.cluster_name,
            "basin_residual_rank": r.basin_residual_rank,
            "latent_residual_delta_beta": r.latent_residual_delta_beta,
            "module_id": r.module_id,
            **vals,
            "hbdTSC_minus_hESC_beta": vals["hbdTSC_beta"] - vals["hESC_beta"] if np.isfinite(vals["hbdTSC_beta"]) and np.isfinite(vals["hESC_beta"]) else np.nan,
            "hiTSC_minus_hESC_beta": vals["hiTSC_beta"] - vals["hESC_beta"] if np.isfinite(vals["hiTSC_beta"]) and np.isfinite(vals["hESC_beta"]) else np.nan,
        })
    out = pd.DataFrame(rows)
    out.to_csv(OUT / "GSE182015_CSB_TRO_DMR_group_beta.tsv", sep="\t", index=False)
    pd.DataFrame(qc).to_csv(OUT / "GSE182015_group_qc.tsv", sep="\t", index=False)
    summary = {
        "analysis": "GSE182015_pluripotency_independent_hiTSC_RRBS_CSB_residual_DMR",
        "processed_files": int(sum(len(v) for v in GROUPS.values())),
        "boundary": "Human hbdTSC/hiTSC state comparison, not paired methylome perturbation.",
    }
    summary.update(summarize(out, "hbdTSC_minus_hESC_beta", "hbdTSC_minus_hESC"))
    summary.update(summarize(out, "hiTSC_minus_hESC_beta", "hiTSC_minus_hESC"))
    (OUT / "GSE182015_CSB_TRO_DMR_summary.json").write_text(json.dumps(summary, indent=2), encoding="utf-8")
    print(json.dumps(summary, indent=2))


if __name__ == "__main__":
    main()
```
