## Supplemental code for "An entropy-based diagnostic framework for characterizing methylation state dynamics during preimplantation development": human_naive_akg_a485_csb_gene_perturbation.py

```
import gzip
import json
from pathlib import Path

import numpy as np
import pandas as pd
from scipy import stats
import matplotlib
matplotlib.use("Agg")
import matplotlib.pyplot as plt


ROOT = Path("E:/5_31_progress/GSE247631_AKG_A485")
OUT = Path("E:/5_31_progress")
COUNTS = ROOT / "GSE247631_STAR_counts.txt.gz"
HGNC = ROOT / "hgnc_complete_set.txt"
CSB = OUT / "crossspecies_mouse_gleaner_matched_genes.tsv"


CONTRASTS = {
    "tt2iLGo_A485": (["tt2iLGo_P300i_0_R1_S1", "tt2iLGo_P300i_0_R2_S2"], ["tt2iLGo_P300i_1_R1_S3", "tt2iLGo_P300i_1_R2_S4"]),
    "TSC_A485": (["TSC_P300i_0_R1_S5", "TSC_P300i_0_R2_S6"], ["TSC_P300i_1_R1_S7", "TSC_P300i_1_R2_S8"]),
    "tt2iLGo_dmAKG": (["tt2iLGo_aKG_0_R1_S9", "tt2iLGo_aKG_0_R3_S10"], ["tt2iLGo_aKG_4_R1_S11", "tt2iLGo_aKG_4_R3_S12"]),
    "TSC_dmAKG": (["TSC_aKG_0_R1_S13", "TSC_aKG_0_R3_S14"], ["TSC_aKG_4_R1_S15", "TSC_aKG_4_R3_S16"]),
}


def load_data():
    counts = pd.read_csv(COUNTS, sep="\t")
    counts["ensembl_gene_id"] = counts["gene_id"].astype(str).str.replace(r"\.\d+$", "", regex=True)
    hgnc = pd.read_csv(HGNC, sep="\t", dtype=str, usecols=["symbol", "ensembl_gene_id"])
    hgnc = hgnc.dropna().drop_duplicates("ensembl_gene_id")
    counts = counts.merge(hgnc, on="ensembl_gene_id", how="left")
    counts = counts.dropna(subset=["symbol"]).copy()
    sample_cols = [c for c in counts.columns if c not in {"gene_id", "ensembl_gene_id", "symbol"}]
    counts = counts.groupby("symbol", as_index=False)[sample_cols].sum()
    counts["symbol"] = counts["symbol"].str.upper()

    csb = pd.read_csv(CSB, sep="\t")
    csb["gene"] = csb["gene_key"].astype(str).str.upper()
    csb = csb.groupby("gene", as_index=False).agg(
        n_human_dmr=("n_human_dmr", "sum"),
        age_weight_5yr=("age_weight_5yr", "mean"),
        contribution_8cell_to_morula=("contribution_8cell_to_morula", "mean"),
        weight_norm_matched=("weight_norm_matched", "mean"),
    )
    return counts, csb


def logcpm(counts):
    sample_cols = [c for c in counts.columns if c != "symbol"]
    lib = counts[sample_cols].sum(axis=0)
    x = np.log2(counts[sample_cols].div(lib, axis=1) * 1e6 + 1)
    x.insert(0, "gene", counts["symbol"])
    return x


def compute_effects(x):
    out = x[["gene"]].copy()
    for name, (ctrl, treat) in CONTRASTS.items():
        c = x[ctrl].to_numpy()
        t = x[treat].to_numpy()
        effect = t.mean(axis=1) - c.mean(axis=1)
        tt, p = stats.ttest_ind(t, c, axis=1, equal_var=False)
        out[f"{name}_effect"] = effect
        out[f"{name}_abs_effect"] = np.abs(effect)
        out[f"{name}_p"] = p
    out["A485_mean_abs_effect"] = out[["tt2iLGo_A485_abs_effect", "TSC_A485_abs_effect"]].mean(axis=1)
    out["dmAKG_mean_abs_effect"] = out[["tt2iLGo_dmAKG_abs_effect", "TSC_dmAKG_abs_effect"]].mean(axis=1)
    out["max_abs_effect"] = out[[c for c in out.columns if c.endswith("_abs_effect")]].max(axis=1)
    out["mean_abs_effect_all"] = out[[c for c in out.columns if c.endswith("_abs_effect")]].mean(axis=1)
    out["mean_logcpm"] = x[[c for c in x.columns if c != "gene"]].mean(axis=1)
    return out


def null_test(effects, csb, metric, n_iter=10000):
    merged = effects.merge(csb, on="gene", how="left")
    merged["is_csb"] = merged["n_human_dmr"].notna()
    valid = merged.dropna(subset=[metric, "mean_logcpm"]).copy()
    valid = valid[valid["mean_logcpm"] > 0.5].copy()
    csb_obs = valid[valid["is_csb"]].copy()
    k = len(csb_obs)
    obs = float(csb_obs[metric].mean())
    obs_weighted = float(np.average(csb_obs[metric], weights=np.abs(csb_obs["weight_norm_matched"]) + 1e-6))
    obs_nom = float((csb_obs[[c for c in csb_obs.columns if c.endswith("_p")]].min(axis=1) < 0.05).mean())
    valid["expr_bin"] = pd.qcut(valid["mean_logcpm"].rank(method="first"), q=10, labels=False)
    bin_counts = valid[valid["is_csb"]]["expr_bin"].value_counts().to_dict()
    rng = np.random.default_rng(20260531)
    vals, vals_w, vals_nom = [], [], []
    weights = np.abs(csb_obs["weight_norm_matched"].to_numpy()) + 1e-6
    pcols = [c for c in valid.columns if c.endswith("_p")]
    for _ in range(n_iter):
        picks = []
        for b, n in bin_counts.items():
            pool = valid[(~valid["is_csb"]) & valid["expr_bin"].eq(b)]
            if len(pool) < n:
                pool = valid[~valid["is_csb"]]
            picks.append(pool.sample(n=n, replace=False, random_state=int(rng.integers(0, 2**31 - 1))))
        samp = pd.concat(picks, ignore_index=True)
        vals.append(float(samp[metric].mean()))
        vals_w.append(float(np.average(samp[metric].to_numpy()[:k], weights=weights[:len(samp)])))
        vals_nom.append(float((samp[pcols].min(axis=1) < 0.05).mean()))
    vals = np.asarray(vals)
    vals_w = np.asarray(vals_w)
    vals_nom = np.asarray(vals_nom)
    return merged, csb_obs, {
        "metric": metric,
        "n_csb_genes_tested": int(k),
        "obs_mean": obs,
        "random_median": float(np.median(vals)),
        "random_q95": float(np.quantile(vals, 0.95)),
        "random_q99": float(np.quantile(vals, 0.99)),
        "empirical_p_ge_obs": float((np.sum(vals >= obs) + 1) / (len(vals) + 1)),
        "obs_weighted": obs_weighted,
        "weighted_random_q95": float(np.quantile(vals_w, 0.95)),
        "weighted_empirical_p_ge_obs": float((np.sum(vals_w >= obs_weighted) + 1) / (len(vals_w) + 1)),
        "obs_fraction_any_nominal_p_lt_0_05": obs_nom,
        "nominal_fraction_random_q95": float(np.quantile(vals_nom, 0.95)),
        "nominal_fraction_empirical_p_ge_obs": float((np.sum(vals_nom >= obs_nom) + 1) / (len(vals_nom) + 1)),
    }, pd.DataFrame({f"{metric}_random": vals, f"{metric}_weighted_random": vals_w, f"{metric}_nominal_random": vals_nom})


def main():
    counts, csb = load_data()
    x = logcpm(counts)
    effects = compute_effects(x)
    metrics = ["A485_mean_abs_effect", "dmAKG_mean_abs_effect", "mean_abs_effect_all", "max_abs_effect"]
    summaries, null_frames = [], []
    merged_keep = None
    csb_keep = None
    for metric in metrics:
        merged, csb_obs, s, null = null_test(effects, csb, metric)
        summaries.append(s)
        null_frames.append(null)
        merged_keep = merged
        csb_keep = csb_obs
    summary_df = pd.DataFrame(summaries)
    null_df = pd.concat(null_frames, axis=1)
    merged_keep.to_csv(OUT / "GSE247631_AKG_A485_all_gene_effects_with_CSB.tsv", sep="\t", index=False)
    csb_keep.to_csv(OUT / "GSE247631_AKG_A485_CSB_gene_effects.tsv", sep="\t", index=False)
    summary_df.to_csv(OUT / "GSE247631_AKG_A485_CSB_gene_null_summary.tsv", sep="\t", index=False)
    null_df.to_csv(OUT / "GSE247631_AKG_A485_CSB_gene_random_null.tsv", sep="\t", index=False)

    best = summary_df.sort_values("empirical_p_ge_obs").iloc[0].to_dict()
    corr_a485 = stats.spearmanr(csb_keep["contribution_8cell_to_morula"], csb_keep["A485_mean_abs_effect"], nan_policy="omit")
    corr_akg = stats.spearmanr(csb_keep["contribution_8cell_to_morula"], csb_keep["dmAKG_mean_abs_effect"], nan_policy="omit")
    top = csb_keep.sort_values("A485_mean_abs_effect", ascending=False).head(20)
    result = {
        "analysis": "GSE247631_human_naive_ESC_TSC_dmAKG_A485_RNA_at_CSB_DMR_genes",
        "date": "2026-05-31",
        "dataset": "GSE247631",
        "design": "human naive ESC / induced TSC; paired DMSO vs A-485 and 0mM vs 4mM dm-alphaKG RNA-seq",
        "best_metric": best,
        "directional_coupling": {
            "A485_contribution_vs_effect_rho": float(corr_a485[0]),
            "A485_contribution_vs_effect_p": float(corr_a485[1]),
            "dmAKG_contribution_vs_effect_rho": float(corr_akg[0]),
            "dmAKG_contribution_vs_effect_p": float(corr_akg[1]),
        },
        "top_A485_sensitive_CSB_genes": top[["gene", "A485_mean_abs_effect", "tt2iLGo_A485_effect", "TSC_A485_effect", "contribution_8cell_to_morula"]].to_dict(orient="records"),
        "boundary": "Human in vitro preimplantation-lineage model perturbation, not processed methylation readout and not natural morula.",
    }
    with open(OUT / "GSE247631_AKG_A485_CSB_gene_perturbation_summary.json", "w", encoding="utf-8") as f:
        json.dump(result, f, indent=2)

    fig, axes = plt.subplots(1, 3, figsize=(13, 4.2))
    metric = "A485_mean_abs_effect"
    axes[0].hist(null_df[f"{metric}_random"], bins=40, color="#9aa4b2")
    axes[0].axvline(summary_df.loc[summary_df["metric"].eq(metric), "obs_mean"].iloc[0], color="#c0392b", lw=2)
    axes[0].set_title("A-485 CSB genes vs matched null")
    axes[0].set_xlabel("mean abs effect")
    axes[1].scatter(csb_keep["contribution_8cell_to_morula"], csb_keep["A485_mean_abs_effect"], s=24, color="#2a9d8f")
    axes[1].set_title("A-485 effect vs reset contribution")
    axes[1].set_xlabel("8-cell to morula contribution")
    axes[1].set_ylabel("A-485 mean abs effect")
    axes[2].barh(np.arange(len(top.head(15))), top.head(15)["A485_mean_abs_effect"], color="#c0392b")
    axes[2].set_yticks(np.arange(len(top.head(15))))
    axes[2].set_yticklabels(top.head(15)["gene"], fontsize=8)
    axes[2].invert_yaxis()
    axes[2].set_title("Top A-485-sensitive CSB genes")
    fig.tight_layout()
    fig.savefig(OUT / "GSE247631_AKG_A485_CSB_gene_perturbation_figure.png", dpi=300, bbox_inches="tight")
    fig.savefig(OUT / "GSE247631_AKG_A485_CSB_gene_perturbation_figure.pdf", bbox_inches="tight")
    print(json.dumps(result, indent=2))


if __name__ == "__main__":
    main()
```
