## Supplemental code for "An entropy-based diagnostic framework for characterizing methylation state dynamics during preimplantation development": 01_aggregate_csb_dmrs.py

```
import json
import math
import os
from pathlib import Path

import numpy as np
import pandas as pd
from scipy.stats import binomtest, spearmanr


ROOT = Path(os.environ.get("GSE266195_ROOT", r"E:\5_31_progress\GSE266195_hTSC_DNMT3L_PBAT_closure"))
MATRIX = ROOT / "raw" / "GSE266195_DNAme_100-CpG_windows_10CpGs_reps.txt.gz"
DMR_PATH = Path(os.environ.get(
    "CSB_DMR_PATH", r"E:\5_31_progress\bismark_full_closure\CSB_TRO_156_residual_DMR_hg19.bed"))
OUT = ROOT / "results"
OUT.mkdir(parents=True, exist_ok=True)


CONTROL_COLS = ["BTS11_Control1", "BTS11_Control2", "BTS11_Control3"]
DNMT3L_COLS = ["BTS11_DNMT3L_1", "BTS11_DNMT3L_2", "BTS11_DNMT3L_3"]


def normalize_chrom(chrom):
    chrom = str(chrom).strip()
    return chrom if chrom.startswith("chr") else f"chr{chrom}"


def load_dmrs():
    d = pd.read_csv(
        DMR_PATH,
        sep="\t",
        header=None,
        names=["chr", "start", "end", "cluster_name", "basin_residual_rank", "latent_residual_delta_beta", "module_id"],
    )
    d["chr"] = d["chr"].map(normalize_chrom)
    d["dmr_index"] = np.arange(len(d))
    return d


def build_intervals(dmr):
    intervals = {}
    for chrom, sub in dmr.groupby("chr"):
        s = sub.sort_values("start")
        intervals[chrom] = (s["start"].to_numpy(), s["end"].to_numpy(), s["dmr_index"].to_numpy())
    return intervals


def main():
    dmr = load_dmrs()
    intervals = build_intervals(dmr)
    control_sum = np.zeros(len(dmr), dtype=float)
    dnmt3l_sum = np.zeros(len(dmr), dtype=float)
    weight_sum = np.zeros(len(dmr), dtype=float)
    window_hits = np.zeros(len(dmr), dtype=int)
    rows_seen = 0
    rows_used = 0

    usecols = ["Chromosome", "Start", "End"] + CONTROL_COLS + DNMT3L_COLS
    for chunk in pd.read_csv(MATRIX, sep="\t", usecols=usecols, chunksize=200000, na_values=["NaN", "NA"]):
        rows_seen += len(chunk)
        chunk["chr"] = chunk["Chromosome"].map(normalize_chrom)
        chunk["control_beta"] = chunk[CONTROL_COLS].mean(axis=1, skipna=True) / 100.0
        chunk["dnmt3l_beta"] = chunk[DNMT3L_COLS].mean(axis=1, skipna=True) / 100.0
        for row in chunk.itertuples(index=False):
            chrom = row.chr
            if chrom not in intervals:
                continue
            start = int(row.Start) - 1
            end = int(row.End)
            if not (np.isfinite(row.control_beta) and np.isfinite(row.dnmt3l_beta)):
                continue
            starts, ends, idxs = intervals[chrom]
            j = np.searchsorted(starts, end, side="left")
            if j == 0:
                continue
            hits = np.where(ends[:j] > start)[0]
            for h in hits:
                idx = idxs[h]
                overlap = max(0, min(end, ends[h]) - max(start, starts[h]))
                if overlap <= 0:
                    continue
                control_sum[idx] += row.control_beta * overlap
                dnmt3l_sum[idx] += row.dnmt3l_beta * overlap
                weight_sum[idx] += overlap
                window_hits[idx] += 1
                rows_used += 1

    out = dmr.copy()
    out["overlap_bp"] = weight_sum
    out["window_hits"] = window_hits
    out["control_beta"] = np.divide(control_sum, weight_sum, out=np.full(len(dmr), np.nan), where=weight_sum > 0)
    out["dnmt3l_beta"] = np.divide(dnmt3l_sum, weight_sum, out=np.full(len(dmr), np.nan), where=weight_sum > 0)
    out["DNMT3L_minus_control_beta"] = out["dnmt3l_beta"] - out["control_beta"]
    out.to_csv(OUT / "GSE266195_CSB_TRO_DMR_DNMT3L_minus_control.tsv", sep="\t", index=False)

    valid = out[np.isfinite(out["DNMT3L_minus_control_beta"]) & np.isfinite(out["latent_residual_delta_beta"])]
    rho = p = None
    if len(valid) >= 3:
        rho, p = spearmanr(valid["latent_residual_delta_beta"], valid["DNMT3L_minus_control_beta"])
        rho = None if math.isnan(rho) else float(rho)
        p = None if math.isnan(p) else float(p)
    concordant = int((np.sign(valid["latent_residual_delta_beta"]) == np.sign(valid["DNMT3L_minus_control_beta"])).sum())
    binom_p = float(binomtest(concordant, len(valid), 0.5, alternative="greater").pvalue) if len(valid) else None
    summary = {
        "analysis": "GSE266195_hTSC_DNMT3L_OE_PBAT_CSB_residual_DMR",
        "rows_seen": int(rows_seen),
        "overlap_events": int(rows_used),
        "dmrs_with_DNMT3L_and_control_beta": int(len(valid)),
        "mean_DNMT3L_minus_control_beta": float(valid["DNMT3L_minus_control_beta"].mean()) if len(valid) else None,
        "median_DNMT3L_minus_control_beta": float(valid["DNMT3L_minus_control_beta"].median()) if len(valid) else None,
        "spearman_latent_residual_vs_DNMT3L_delta_rho": rho,
        "spearman_latent_residual_vs_DNMT3L_delta_p": p,
        "sign_concordant_dmrs": concordant,
        "sign_concordance_binomial_p_greater": binom_p,
    }
    (OUT / "GSE266195_CSB_TRO_DMR_summary.json").write_text(json.dumps(summary, indent=2), encoding="utf-8")
    print(json.dumps(summary, indent=2))


if __name__ == "__main__":
    main()
```
