## Supplemental code for "An entropy-based diagnostic framework for characterizing methylation state dynamics during preimplantation development": 01_aggregate_bigwig_csb_dmrs.py

```
import json
import math
import os
from pathlib import Path

import numpy as np
import pandas as pd
import pyBigWig
from scipy.stats import binomtest, spearmanr


ROOT = Path(os.environ.get("GSE280039_ROOT", r"E:\5_31_progress\GSE280039_hTSC_induction_methylome"))
DMR_PATH = Path(os.environ.get(
    "CSB_DMR_PATH", r"E:\5_31_progress\bismark_full_closure\CSB_TRO_156_residual_DMR_hg19.bed"))
RAW = ROOT / "raw"
OUT = ROOT / "results"
OUT.mkdir(parents=True, exist_ok=True)

P0_BW = RAW / "GSE280039_EPSC_to_TSC_P0.bw"
P2_BW = RAW / "GSE280039_EPSC_to_TSC_P2.bw"


def normalize_chrom(chrom):
    chrom = str(chrom).strip()
    return chrom if chrom.startswith("chr") else f"chr{chrom}"


def load_dmrs():
    d = pd.read_csv(
        DMR_PATH,
        sep="\t",
        header=None,
        names=["chr", "start", "end", "cluster_name", "basin_residual_rank", "latent_residual_delta_beta", "module_id"],
    )
    d["chr"] = d["chr"].map(normalize_chrom)
    return d


def bw_mean(bw, chrom, start, end):
    chroms = bw.chroms()
    candidates = [chrom]
    if chrom.startswith("chr"):
        candidates.append(chrom[3:])
    else:
        candidates.append("chr" + chrom)
    for c in candidates:
        if c not in chroms:
            continue
        c_end = chroms[c]
        s = max(0, int(start))
        e = min(int(end), c_end)
        if e <= s:
            return np.nan
        val = bw.stats(c, s, e, type="mean")[0]
        return np.nan if val is None else float(val) / 100.0
    return np.nan


def main():
    dmr = load_dmrs()
    p0 = pyBigWig.open(str(P0_BW))
    p2 = pyBigWig.open(str(P2_BW))
    rows = []
    try:
        for row in dmr.itertuples(index=False):
            p0_beta = bw_mean(p0, row.chr, row.start, row.end)
            p2_beta = bw_mean(p2, row.chr, row.start, row.end)
            rows.append({
                "chr": row.chr,
                "start": row.start,
                "end": row.end,
                "cluster_name": row.cluster_name,
                "basin_residual_rank": row.basin_residual_rank,
                "latent_residual_delta_beta": row.latent_residual_delta_beta,
                "module_id": row.module_id,
                "P0_beta": p0_beta,
                "P2_beta": p2_beta,
                "P2_minus_P0_beta": p2_beta - p0_beta if np.isfinite(p0_beta) and np.isfinite(p2_beta) else np.nan,
            })
    finally:
        p0.close()
        p2.close()
    out = pd.DataFrame(rows)
    out.to_csv(OUT / "GSE280039_CSB_TRO_DMR_P2_minus_P0.tsv", sep="\t", index=False)
    valid = out[np.isfinite(out["P2_minus_P0_beta"]) & np.isfinite(out["latent_residual_delta_beta"])]
    rho = p = None
    if len(valid) >= 3:
        rho, p = spearmanr(valid["latent_residual_delta_beta"], valid["P2_minus_P0_beta"])
        rho = None if math.isnan(rho) else float(rho)
        p = None if math.isnan(p) else float(p)
    concordant = int((np.sign(valid["latent_residual_delta_beta"]) == np.sign(valid["P2_minus_P0_beta"])).sum())
    binom_p = float(binomtest(concordant, len(valid), 0.5, alternative="greater").pvalue) if len(valid) else None
    summary = {
        "analysis": "GSE280039_hEPSC_to_hTSC_P0_P2_bigwig_CSB_residual_DMR",
        "dmrs_with_P0_P2_beta": int(len(valid)),
        "mean_P2_minus_P0_beta": float(valid["P2_minus_P0_beta"].mean()) if len(valid) else None,
        "median_P2_minus_P0_beta": float(valid["P2_minus_P0_beta"].median()) if len(valid) else None,
        "spearman_latent_residual_vs_P2_minus_P0_rho": rho,
        "spearman_latent_residual_vs_P2_minus_P0_p": p,
        "sign_concordant_dmrs": concordant,
        "sign_concordance_binomial_p_greater": binom_p,
    }
    (OUT / "GSE280039_CSB_TRO_DMR_summary.json").write_text(json.dumps(summary, indent=2), encoding="utf-8")
    print(json.dumps(summary, indent=2))


if __name__ == "__main__":
    main()
```
