## Supplemental code for "An entropy-based diagnostic framework for characterizing methylation state dynamics during preimplantation development": 01_aggregate_gse291172_bigwig_csb_dmrs.py

```
import json
import math
import os
from pathlib import Path

import numpy as np
import pandas as pd
import pyBigWig
from scipy.stats import binomtest, spearmanr


ROOT = Path(os.environ.get("GSE291172_ROOT", r"E:\5_31_progress\GSE291172_STAT3_embryo_model_WGBS_closure"))
DMR_PATH = Path(os.environ.get(
    "CSB_DMR_PATH", r"E:\5_31_progress\bismark_full_closure\CSB_TRO_156_residual_DMR_hg19.bed"))
if Path("/mnt/e").exists() and "GSE291172_ROOT" not in os.environ:
    ROOT = Path("/mnt/e/5_31_progress/GSE291172_STAT3_embryo_model_WGBS_closure")
if Path("/mnt/e").exists() and "CSB_DMR_PATH" not in os.environ:
    DMR_PATH = Path("/mnt/e/5_31_progress/bismark_full_closure/CSB_TRO_156_residual_DMR_hg19.bed")
RAW = ROOT / "raw"
OUT = ROOT / "results"
OUT.mkdir(parents=True, exist_ok=True)

SAC_BW = RAW / "GSM8830343_SACD5_WGBS.bw"
HPSC_BW = RAW / "GSM8830344_OG_WGBS.bw"


def normalize_chrom(chrom):
    chrom = str(chrom).strip()
    return chrom if chrom.startswith("chr") else f"chr{chrom}"


def load_dmrs():
    d = pd.read_csv(
        DMR_PATH,
        sep="\t",
        header=None,
        names=["chr", "start", "end", "cluster_name", "basin_residual_rank", "latent_residual_delta_beta", "module_id"],
    )
    d["chr"] = d["chr"].map(normalize_chrom)
    return d


def bw_mean_raw(bw, chrom, start, end):
    chroms = bw.chroms()
    candidates = [chrom]
    if chrom.startswith("chr"):
        candidates.append(chrom[3:])
    else:
        candidates.append("chr" + chrom)
    for c in candidates:
        if c not in chroms:
            continue
        s = max(0, int(start))
        e = min(int(end), chroms[c])
        if e <= s:
            return np.nan
        val = bw.stats(c, s, e, type="mean")[0]
        return np.nan if val is None else float(val)
    return np.nan


def infer_scale(values):
    arr = np.asarray([v for v in values if np.isfinite(v)], dtype=float)
    if arr.size == 0:
        return 1.0
    return 100.0 if np.nanpercentile(arr, 95) > 1.5 else 1.0


def main():
    dmr = load_dmrs()
    sac = pyBigWig.open(str(SAC_BW))
    hpsc = pyBigWig.open(str(HPSC_BW))
    raw_rows = []
    try:
        for row in dmr.itertuples(index=False):
            hpsc_raw = bw_mean_raw(hpsc, row.chr, row.start, row.end)
            sac_raw = bw_mean_raw(sac, row.chr, row.start, row.end)
            raw_rows.append((row, hpsc_raw, sac_raw))
    finally:
        sac.close()
        hpsc.close()

    scale = infer_scale([x for _, h, s in raw_rows for x in (h, s)])
    rows = []
    for row, hpsc_raw, sac_raw in raw_rows:
        hpsc_beta = hpsc_raw / scale if np.isfinite(hpsc_raw) else np.nan
        sac_beta = sac_raw / scale if np.isfinite(sac_raw) else np.nan
        delta = sac_beta - hpsc_beta if np.isfinite(hpsc_beta) and np.isfinite(sac_beta) else np.nan
        rows.append({
            "chr": row.chr,
            "start": row.start,
            "end": row.end,
            "cluster_name": row.cluster_name,
            "basin_residual_rank": row.basin_residual_rank,
            "latent_residual_delta_beta": row.latent_residual_delta_beta,
            "module_id": row.module_id,
            "hPSC_beta": hpsc_beta,
            "SAC_120hr_beta": sac_beta,
            "SAC_minus_hPSC_beta": delta,
        })

    out = pd.DataFrame(rows)
    out.to_csv(OUT / "GSE291172_CSB_TRO_DMR_SAC_minus_hPSC.tsv", sep="\t", index=False)
    valid = out[np.isfinite(out["SAC_minus_hPSC_beta"]) & np.isfinite(out["latent_residual_delta_beta"])]
    rho = p = None
    if len(valid) >= 3:
        rho, p = spearmanr(valid["latent_residual_delta_beta"], valid["SAC_minus_hPSC_beta"])
        rho = None if math.isnan(rho) else float(rho)
        p = None if math.isnan(p) else float(p)
    concordant = int((np.sign(valid["latent_residual_delta_beta"]) == np.sign(valid["SAC_minus_hPSC_beta"])).sum())
    binom_p = float(binomtest(concordant, len(valid), 0.5, alternative="greater").pvalue) if len(valid) else None
    summary = {
        "analysis": "GSE291172_STAT3_induced_human_embryo_model_WGBS_CSB_residual_DMR",
        "comparison": "120hr_SAC_minus_hPSC",
        "bigwig_scale_divisor": scale,
        "dmrs_with_hPSC_and_SAC_beta": int(len(valid)),
        "mean_SAC_minus_hPSC_beta": float(valid["SAC_minus_hPSC_beta"].mean()) if len(valid) else None,
        "median_SAC_minus_hPSC_beta": float(valid["SAC_minus_hPSC_beta"].median()) if len(valid) else None,
        "spearman_latent_residual_vs_SAC_minus_hPSC_rho": rho,
        "spearman_latent_residual_vs_SAC_minus_hPSC_p": p,
        "sign_concordant_dmrs": concordant,
        "sign_concordance_binomial_p_greater": binom_p,
        "boundary": "Human embryo-model WGBS state/induction comparison, not paired p300/TET/DNMT methylome perturbation.",
    }
    (OUT / "GSE291172_CSB_TRO_DMR_summary.json").write_text(json.dumps(summary, indent=2), encoding="utf-8")
    print(json.dumps(summary, indent=2))


if __name__ == "__main__":
    main()
```
