## Supplemental code for "An entropy-based diagnostic framework for characterizing methylation state dynamics during preimplantation development": human_embryo_dex_csb_gene_perturbation.py

```
import gzip
import json
from pathlib import Path

import numpy as np
import pandas as pd
from scipy import stats
import matplotlib
matplotlib.use("Agg")
import matplotlib.pyplot as plt


ROOT = Path("E:/5_31_progress/E-MTAB-10096_human_embryo_DEX")
OUT = Path("E:/5_31_progress")
COUNTS = ROOT / "merge.rpkmforgenes_counts.tsv.gz"
SDRF = ROOT / "E-MTAB-10096.sdrf.txt"
CSB_GENES = OUT / "crossspecies_mouse_gleaner_matched_genes.tsv"


def load_inputs():
    meta = pd.read_csv(SDRF, sep="\t")
    meta = meta.rename(columns={
        "Source Name": "sample",
        "Characteristics[growth condition]": "condition",
        "Characteristics[inferred lineage]": "lineage",
        "Characteristics[individual]": "individual",
        "Characteristics[sex]": "sex",
    })
    meta = meta[["sample", "condition", "lineage", "individual", "sex"]].copy()
    meta["condition"] = meta["condition"].map({"Ctrl": "control", "Treat": "dex"})
    meta = meta[meta["condition"].isin(["control", "dex"])]
    meta = meta[meta["lineage"].isin(["mural", "polar", "epi", "pe"])]

    counts = pd.read_csv(COUNTS, sep="\t")
    counts = counts.rename(columns={counts.columns[0]: "gene"})
    sample_cols = [c for c in counts.columns if c in set(meta["sample"])]
    counts = counts[["gene"] + sample_cols].copy()
    counts = counts.groupby("gene", as_index=False)[sample_cols].sum()

    csb = pd.read_csv(CSB_GENES, sep="\t")
    csb["gene_key"] = csb["gene_key"].astype(str).str.upper()
    csb = csb.groupby("gene_key", as_index=False).agg(
        n_human_dmr=("n_human_dmr", "sum"),
        age_weight_5yr=("age_weight_5yr", "mean"),
        contribution_8cell_to_morula=("contribution_8cell_to_morula", "mean"),
        weight_norm_matched=("weight_norm_matched", "mean"),
    )
    return counts, meta, csb


def normalize_logcpm(counts, meta):
    sample_cols = [c for c in counts.columns if c != "gene"]
    lib = counts[sample_cols].sum(axis=0)
    cpm = counts[sample_cols].div(lib, axis=1) * 1e6
    logcpm = np.log2(cpm + 1.0)
    logcpm.insert(0, "gene", counts["gene"].astype(str).str.upper())
    return logcpm


def lineage_effects(logcpm, meta, min_cells=10):
    rows = []
    for lineage in ["mural", "polar"]:
        sub_meta = meta[meta["lineage"].eq(lineage)].copy()
        ctrl = sub_meta[sub_meta["condition"].eq("control")]["sample"].tolist()
        dex = sub_meta[sub_meta["condition"].eq("dex")]["sample"].tolist()
        ctrl = [c for c in ctrl if c in logcpm.columns]
        dex = [c for c in dex if c in logcpm.columns]
        if len(ctrl) < min_cells or len(dex) < min_cells:
            continue
        c = logcpm[ctrl].to_numpy()
        d = logcpm[dex].to_numpy()
        effect = d.mean(axis=1) - c.mean(axis=1)
        t, p = stats.ttest_ind(d, c, axis=1, equal_var=False, nan_policy="omit")
        rows.append(pd.DataFrame({
            "gene": logcpm["gene"],
            f"{lineage}_dex_minus_control_logcpm": effect,
            f"{lineage}_abs_effect": np.abs(effect),
            f"{lineage}_p": p,
            f"{lineage}_ctrl_n": len(ctrl),
            f"{lineage}_dex_n": len(dex),
            f"{lineage}_mean_logcpm": np.nanmean(np.c_[c, d], axis=1),
        }))
    out = rows[0]
    for r in rows[1:]:
        out = out.merge(r, on="gene", how="outer")
    out["max_abs_effect"] = out[[c for c in out.columns if c.endswith("_abs_effect")]].max(axis=1)
    out["mean_abs_effect"] = out[[c for c in out.columns if c.endswith("_abs_effect")]].mean(axis=1)
    out["min_p"] = out[[c for c in out.columns if c.endswith("_p")]].min(axis=1)
    out["mean_logcpm"] = out[[c for c in out.columns if c.endswith("_mean_logcpm")]].mean(axis=1)
    return out


def random_gene_set_null(effects, csb, n_iter=20000):
    merged = effects.merge(csb, left_on="gene", right_on="gene_key", how="left")
    merged["is_csb"] = merged["gene_key"].notna()
    expressed = merged.dropna(subset=["mean_abs_effect", "mean_logcpm"]).copy()
    csb_obs = expressed[expressed["is_csb"]].copy()
    non = expressed[~expressed["is_csb"]].copy()
    k = len(csb_obs)
    obs_mean = float(csb_obs["mean_abs_effect"].mean())
    obs_top20 = float((csb_obs["min_p"] <= 0.05).mean())
    obs_weighted = float(np.average(csb_obs["mean_abs_effect"], weights=np.abs(csb_obs["weight_norm_matched"]) + 1e-6))

    rng = np.random.default_rng(20260531)
    bins = pd.qcut(expressed["mean_logcpm"].rank(method="first"), q=10, labels=False)
    expressed["expr_bin"] = bins
    csb_bins = expressed.loc[expressed["is_csb"], "expr_bin"].value_counts().to_dict()

    vals_mean, vals_top, vals_weighted = [], [], []
    for _ in range(n_iter):
        picks = []
        for b, n in csb_bins.items():
            pool = expressed[(~expressed["is_csb"]) & (expressed["expr_bin"].eq(b))]
            if len(pool) < n:
                pool = expressed[~expressed["is_csb"]]
            picks.append(pool.sample(n=n, replace=False, random_state=int(rng.integers(0, 2**31 - 1))))
        samp = pd.concat(picks, ignore_index=True)
        vals_mean.append(float(samp["mean_abs_effect"].mean()))
        vals_top.append(float((samp["min_p"] <= 0.05).mean()))
        w = np.abs(csb_obs["weight_norm_matched"].to_numpy()) + 1e-6
        vals_weighted.append(float(np.average(samp["mean_abs_effect"].to_numpy()[:k], weights=w[:len(samp)])))
    vals_mean = np.asarray(vals_mean)
    vals_top = np.asarray(vals_top)
    vals_weighted = np.asarray(vals_weighted)

    # Directional coherence: do genes with stronger morula reset contribution show stronger DEX response?
    corr = stats.spearmanr(csb_obs["contribution_8cell_to_morula"], csb_obs["mean_abs_effect"], nan_policy="omit")
    signed_mural = stats.spearmanr(csb_obs["contribution_8cell_to_morula"], csb_obs.get("mural_dex_minus_control_logcpm"), nan_policy="omit")
    signed_polar = stats.spearmanr(csb_obs["contribution_8cell_to_morula"], csb_obs.get("polar_dex_minus_control_logcpm"), nan_policy="omit")

    summary = {
        "analysis": "E-MTAB-10096_human_embryo_dexamethasone_RNA_at_CSB_DMR_genes",
        "date": "2026-05-31",
        "dataset": {
            "RNA": "E-MTAB-10096",
            "methylation_raw_confirmed": "E-MTAB-10097",
            "title": "Single Cell Multi-Omics of Human Preimplantation Embryos Demonstrates Susceptibility to Excess Glucocorticoid Exposure",
        },
        "sample_counts": {
            "control_cells": int((meta_global["condition"] == "control").sum()),
            "dex_cells": int((meta_global["condition"] == "dex").sum()),
            "mural_control": int(((meta_global["condition"] == "control") & (meta_global["lineage"] == "mural")).sum()),
            "mural_dex": int(((meta_global["condition"] == "dex") & (meta_global["lineage"] == "mural")).sum()),
            "polar_control": int(((meta_global["condition"] == "control") & (meta_global["lineage"] == "polar")).sum()),
            "polar_dex": int(((meta_global["condition"] == "dex") & (meta_global["lineage"] == "polar")).sum()),
        },
        "csb_gene_set": {
            "n_csb_genes_tested": int(k),
            "obs_mean_abs_dex_effect": obs_mean,
            "expr_matched_random_median": float(np.median(vals_mean)),
            "expr_matched_random_q95": float(np.quantile(vals_mean, 0.95)),
            "empirical_p_ge_obs_mean_abs_effect": float((np.sum(vals_mean >= obs_mean) + 1) / (len(vals_mean) + 1)),
            "obs_fraction_nominal_p_lt_0_05": obs_top20,
            "random_fraction_nominal_p_lt_0_05_median": float(np.median(vals_top)),
            "empirical_p_ge_obs_fraction": float((np.sum(vals_top >= obs_top20) + 1) / (len(vals_top) + 1)),
            "obs_weighted_abs_effect": obs_weighted,
            "weighted_random_q95": float(np.quantile(vals_weighted, 0.95)),
            "empirical_p_ge_obs_weighted": float((np.sum(vals_weighted >= obs_weighted) + 1) / (len(vals_weighted) + 1)),
        },
        "directional_coupling": {
            "spearman_contribution_vs_abs_effect_rho": float(corr[0]),
            "spearman_contribution_vs_abs_effect_p": float(corr[1]),
            "spearman_contribution_vs_mural_signed_effect_rho": float(signed_mural[0]),
            "spearman_contribution_vs_mural_signed_effect_p": float(signed_mural[1]),
            "spearman_contribution_vs_polar_signed_effect_rho": float(signed_polar[0]),
            "spearman_contribution_vs_polar_signed_effect_p": float(signed_polar[1]),
        },
        "claim_boundary": (
            "This is a human preimplantation embryo perturbation with RNA processed data and same-study bisulfite raw data, "
            "not a morula-stage methylation-processed perturbation matrix. It can upgrade the causal chain only if the CSB-DMR genes "
            "show expression-matched perturbation enrichment."
        ),
    }
    return merged, csb_obs, pd.DataFrame({
        "random_mean_abs_effect": vals_mean,
        "random_fraction_nominal_p_lt_0_05": vals_top,
        "random_weighted_abs_effect": vals_weighted,
    }), summary


def make_figure(csb_obs, null, summary):
    fig, axes = plt.subplots(1, 3, figsize=(13, 4.2))
    s = summary["csb_gene_set"]
    axes[0].hist(null["random_mean_abs_effect"], bins=40, color="#9aa4b2", alpha=0.85)
    axes[0].axvline(s["obs_mean_abs_dex_effect"], color="#c0392b", lw=2)
    axes[0].set_title("CSB genes vs expression-matched null")
    axes[0].set_xlabel("mean abs DEX effect")
    axes[0].set_ylabel("random sets")

    axes[1].scatter(csb_obs["contribution_8cell_to_morula"], csb_obs["mean_abs_effect"], s=24, color="#2a9d8f", alpha=0.8)
    axes[1].set_title("Reset contribution vs DEX sensitivity")
    axes[1].set_xlabel("8-cell to morula contribution")
    axes[1].set_ylabel("mean abs DEX effect")

    top = csb_obs.sort_values("mean_abs_effect", ascending=False).head(15).copy()
    axes[2].barh(np.arange(len(top)), top["mean_abs_effect"], color="#c0392b")
    axes[2].set_yticks(np.arange(len(top)))
    axes[2].set_yticklabels(top["gene"], fontsize=8)
    axes[2].invert_yaxis()
    axes[2].set_title("Most perturbed CSB-DMR genes")
    axes[2].set_xlabel("mean abs DEX effect")

    fig.suptitle("Human Embryo DEX Perturbation at CSB-TRO DMR Genes", fontsize=12, fontweight="bold")
    fig.tight_layout(rect=[0, 0, 1, 0.92])
    fig.savefig(OUT / "E-MTAB-10096_DEX_CSB_gene_perturbation_figure.png", dpi=300, bbox_inches="tight")
    fig.savefig(OUT / "E-MTAB-10096_DEX_CSB_gene_perturbation_figure.pdf", bbox_inches="tight")
    plt.close(fig)


if __name__ == "__main__":
    counts_global, meta_global, csb_global = load_inputs()
    logcpm_global = normalize_logcpm(counts_global, meta_global)
    effects_global = lineage_effects(logcpm_global, meta_global)
    merged_global, csb_obs_global, null_global, summary_global = random_gene_set_null(
        effects_global, csb_global
    )
    merged_global.to_csv(OUT / "E-MTAB-10096_DEX_all_gene_effects_with_CSB.tsv", sep="\t", index=False)
    csb_obs_global.to_csv(OUT / "E-MTAB-10096_DEX_CSB_gene_effects.tsv", sep="\t", index=False)
    null_global.to_csv(OUT / "E-MTAB-10096_DEX_CSB_expr_matched_null.tsv", sep="\t", index=False)
    with open(OUT / "E-MTAB-10096_DEX_CSB_gene_perturbation_summary.json", "w", encoding="utf-8") as f:
        json.dump(summary_global, f, indent=2)
    make_figure(csb_obs_global, null_global, summary_global)
    print(json.dumps(summary_global, indent=2))
```
