## Supplemental docs for "An entropy-based diagnostic framework for characterizing methylation state dynamics during preimplantation development": CSB_TRO_2026-05-27_diagnostic_evidence_summary.md

# Public-data bounded diagnostic evidence summary

Status: generated from frozen local result files; no original result files were modified.

## Main conclusion

The current evidence supports a biologically structured missing correction term, not a fully identified causal u_bio.

Recommended claim:

Methylation-only operator-time dynamics captures baseline DMR-state propagation but fails at morula reset-basin entry. The measured correction is structured, module-specific, direction-sensitive, and organized by a dual-branch closure/access diagnostic architecture. Public RNA and motif x TF surrogates are weak replacements, while histone-associated evidence supports biological plausibility. Final replacement remains bounded by controlled-access morula H3K27ac/H3K4me3 data.

## Evidence chain

- methylation_only_failure: q90 occupancy=0.044; observed q90 occupancy=0.875 (strong diagnostic)
- measured_correction_upper_bound: alpha_to_observed=0.500; occupancy@alpha1=1.000 (diagnostic upper bound)
- module_specificity: M05/M01/M12 occupancy=0.867; +M02 occupancy=0.956 (strong diagnostic)
- dual_branch_architecture: occupancy=0.956; cosine=0.994; PC3 recovery=0.879 (central result)
- sign_dependence: correct=0.956; wrong closure=0.000; wrong access=0.178 (important control)
- sign_pattern_enumeration: true sign rank=1; percentile=1.000 (important control)
- rna_surrogate: global RNA occupancy=0.200 (boundary)
- motif_tf_surrogate: motif x TF occupancy=0.222 (boundary)
- hesc_histone_proxy: max occupancy=0.511; cosine=0.806 (moderate support)
- public_embryo_histone_diagnostic: max occupancy=0.978; cosine=0.933; PC3 recovery=0.790 (histone-supported diagnostic)
- strict_morula_histone_gap: strict partial histone max occupancy=0.200 (limitation)

## Control status

- dual_branch_sign / wrong_closure_correct_access: 0.000; pass=yes
- dual_branch_sign / correct_closure_wrong_access: 0.178; pass=yes
- dual_branch_sign / wrong_closure_wrong_access: 0.000; pass=yes
- dual_branch_sign / naive_inverse_all_modules: 0.333; pass=yes
- dual_branch_sign / naive_raw_all_modules: 0.000; pass=yes
- dual_branch_random_partition / random 3/2 module partitions: n=1000; median=0.111; q95=0.956; max=0.956; frac>=true=0.096; pass=qualified
- exact_partition_sign / all exact module partitions with branch signs: n=40; median=0.044; q95=0.744; max=0.956; pass=yes
- top_residual_DMR_matched_random / top10 matched random DMRs: n=500; median=0.089; q95=0.133; max=0.156; pass=yes
- top_residual_DMR_matched_random / top25 matched random DMRs: n=500; median=0.089; q95=0.156; max=0.200; pass=yes
- top_residual_DMR_matched_random / top50 matched random DMRs: n=500; median=0.089; q95=0.111; max=0.111; pass=yes
- embryo_histone_matched_random / public embryo histone diagnostic matched random: n=420; median=0.978; q95=0.978; max=1.000; frac>=true=0.705; pass=no

## Immediate manuscript action

1. Keep negative controls in the main figure panels where possible.
2. Use the evidence table as a boundary table, not as proof of causal u_bio.
3. Do not expand into mouse, cross-species, wet-lab, OT, or Schrodinger bridge analyses in the current manuscript.
