## Supplemental docs for "An entropy-based diagnostic framework for characterizing methylation state dynamics during preimplantation development": CSB_TRO_2026-05-27_duality_accessibility_interpretation.md

# Duality-accessibility coupling analysis

## Test

This analysis asks whether morula-centered entry-exit geometry and stage-matched public morula accessibility converge on the same residual DMRs or modules.

## Main readout

- top25 residual DMR morula accessibility: observed=1.660, random q95=1.681, q95-positive=False
- top25 geometry DMR morula accessibility: observed=1.376, random q95=1.613, q95-positive=False
- access branch morula accessibility: observed=1.152, random q95=1.574, q95-positive=False
- top25 residual x top25 geometry overlap: observed=4, random q95=5.0, q95-positive=False
- DMR geometry score versus morula accessibility Spearman rho=0.103; random q05-q95=-0.136 to 0.146
- top25 negative-curvature DMR morula accessibility: observed=1.793, random q95=1.623, q95-positive=True
- top50 negative-curvature DMR morula accessibility: observed=1.630, random q95=1.553, q95-positive=True
- inverted-U DMR morula accessibility: observed=1.716, random q95=1.591, q95-positive=True
- U-shape DMR morula accessibility: observed=1.260, random q95=1.498, q95-positive=False
- M02 module: geometry score=0.295, morula accessibility=1.162
- q95-positive module-level components: M14(access=False, geometry=True, residual=False), M06(access=False, geometry=True, residual=False), M05(access=False, geometry=False, residual=True), M12(access=False, geometry=False, residual=True), M01(access=False, geometry=False, residual=True), M10(access=False, geometry=False, residual=True)

## Interpretation

Coupling verdict: morula accessibility is directionally enriched in negative-curvature/inverted-U DMRs; individual modules show q95-positive module-level components.

The result should be treated as a convergence test between reset-basin geometry and public chromatin support. The positive coupling is not a broad high-duality or U-shape coupling. Instead, the public morula accessibility signal is concentrated in negative-curvature/inverted-U DMRs, while canonical U-shape DMRs are not accessibility-enriched. It does not identify causal u_bio because there is still no paired perturbation-to-methylation readout.

## Claim boundary

Positive coupling can support the statement that morula-centered entry-exit geometry and stage-matched accessibility support converge on a subset of residual/access-associated DMRs. It cannot support final causal u_bio detection.
