## Supplemental docs for "An entropy-based diagnostic framework for characterizing methylation state dynamics during preimplantation development": CSB_TRO_2026-05-27_entry_exit_duality_interpretation.md

# Morula-centered entry-exit duality analysis

## Test

This analysis tests whether the 8-cell-to-morula entry vector and morula-to-blastocyst exit vector form a morula-centered duality:

`Delta_entry = beta_morula - beta_8cell`

`Delta_exit = beta_blastocyst - beta_morula`

`duality_score = -cos(Delta_entry, Delta_exit)`

`curvature = beta_8cell - 2 * beta_morula + beta_blastocyst`

## Main readout

- all DMR duality score: 0.699
- all DMR U-shape fraction: 0.404
- priority residual-module duality score: 0.613
- priority residual-module U-shape fraction: 0.266
- top25 basin residual DMR duality score: 0.587
- top25 basin residual DMR U-shape fraction: 0.360

## Random-control boundary

| group | control_mode | observed_duality_score | random_median | random_q95 | observed_gt_random_q95 | empirical_p_ge_observed |
| --- | --- | --- | --- | --- | --- | --- |
| closure_modules_M01_M05_M12 | size_matched_random_sets | 0.395 | 0.707 | 0.872 | False | 0.957 |
| access_modules_M02_M10 | size_matched_random_sets | 0.872 | 0.704 | 0.871 | True | 0.049 |
| top25_basin_residual_DMRs | module_matched_random_sets | 0.587 | 0.441 | 0.577 | True | 0.039 |
| top50_basin_residual_DMRs | module_matched_random_sets | 0.727 | 0.507 | 0.619 | True | 0.001 |
| top100_basin_residual_DMRs | module_matched_random_sets | 0.735 | 0.654 | 0.714 | True | 0.011 |
| module_M02 | size_matched_random_sets | 0.994 | 0.713 | 0.880 | True | 0.001 |
| all_DMRs_exit_permutation | exit_vector_permutation | 0.699 | 0.007 | 0.133 | True | 0.001 |

## Priority module ranking

| module_id | branch | n_dmr | duality_score_minus_cosine | fraction_u_shape | mean_curvature | median_rebound_ratio |
| --- | --- | --- | --- | --- | --- | --- |
| M02 | access | 30 | 0.994 | 0.333 | -0.138 | 0.172 |
| M05 | closure | 6 | 0.650 | 0.167 | -0.071 | 0.481 |
| M12 | closure | 4 | 0.590 | 0.000 | -0.155 | 0.508 |
| M10 | access | 3 | 0.301 | 0.333 | -0.346 | 0.679 |
| M01 | closure | 21 | 0.263 | 0.238 | 0.069 | 0.979 |

## Strongest module-level duality signals

| module_id | branch | n_dmr | duality_score_minus_cosine | fraction_u_shape | mean_curvature | median_rebound_ratio |
| --- | --- | --- | --- | --- | --- | --- |
| M02 | access | 30 | 0.994 | 0.333 | -0.138 | 0.172 |
| M06 | other | 23 | 0.991 | 0.435 | -0.193 | 1.024 |
| M14 | other | 5 | 0.973 | 0.400 | -0.069 | 1.242 |
| M00 | other | 16 | 0.903 | 0.875 | 0.077 | 2.044 |
| M13 | other | 7 | 0.893 | 0.429 | -0.472 | 4.735 |
| M07 | other | 9 | 0.887 | 0.333 | -0.188 | 0.471 |
| M09 | other | 3 | 0.866 | 0.667 | 0.332 | 0.233 |
| M04 | other | 10 | 0.843 | 0.500 | -0.037 | 0.644 |

## Claim boundary

A positive score supports morula-centered entry-exit geometry. It does not by itself identify a causal biological input. The result should be written as a methylation-state-space geometry result and, if concentrated in priority residual modules, as module-specific reset-basin entry-exit duality rather than global strict symmetry.
