## Supplemental docs for "An entropy-based diagnostic framework for characterizing methylation state dynamics during preimplantation development": CSB_TRO_2026-05-27_main_figure_plan_v1.0.md

# Main figure plan for public-data-bounded diagnostic dynamics v1.0

## Figure 1. Operator-time methylation dynamics and morula-entry failure

Panels:
- DMR state-space/operator-time schematic.
- Methylation-only morula prediction versus observed morula basin.
- q90 occupancy contrast: methylation-only 0.044 versus observed 0.875.
- Measured correction alpha scan showing alpha_to_observed around 0.50.

Claim: methylation-only dynamics is a necessary baseline but is insufficient for morula reset-basin entry.

## Figure 2. Measured correction and residual modules

Panels:
- Definition of c_diag = x_observed,morula - x_predicted,morula.
- Top residual DMR/module contribution plot.
- Greedy module reconstruction: M05, M05/M01, M05/M01/M12, +M02, +M10.
- Matched-random top residual DMR control.

Claim: the correction is non-random and module-specific.

## Figure 3. Dual-branch closure/access diagnostic architecture

Panels:
- Closure branch M05/M01/M12 and access branch M02/M10 schematic.
- Correct closure/access occupancy 0.956.
- Wrong closure/wrong access sign controls.
- Exact sign-pattern and beta-grid robustness summary.

Claim: the correction is direction-sensitive and best organized by a dual-branch diagnostic architecture.

## Figure 4. Public surrogate comparison and data-access boundary

Panels:
- RNA and motif x TF weak surrogate comparison.
- hESC/public embryo histone diagnostic support.
- Public embryo histone matched-random boundary.
- Controlled-access H3K27ac_morula/H3K4me3_morula gap.

Claim: histone evidence supports plausibility, but final biological replacement is data-access limited.

## Figure 5. Morula-centered entry-exit reset-basin geometry

Panels:
- Definition of entry change = beta_morula - beta_8cell and exit change = beta_blastocyst - beta_morula.
- DMR-level entry versus exit scatter with the anti-diagonal reference.
- Duality score comparison: all DMRs, top25/top50/top100 residual DMRs, closure/access branches.
- Random-control boundary: all-DMR exit-permutation q95 and module-matched top residual q95 controls.
- Module-level panel highlighting M02/M10 access and M01/M05/M12 closure behavior.

Claim: morula is supported as a reset-basin geometric vertex with non-random entry-exit anti-alignment and top residual enrichment, but the pattern is partial and module-specific rather than globally symmetric.

Use:
- `CSB_TRO_2026-05-27_entry_exit_duality_summary.svg`
- `CSB_TRO_2026-05-27_entry_exit_summary.tsv`
- `CSB_TRO_2026-05-27_entry_exit_random_controls.tsv`

## Figure 6. Stage-matched public chromatin rescue for top residual DMRs

Panels:
- Extraction/audit schematic: Liu2019 LiCAT/accessibility coordinate-like regions to residual DMR overlap.
- Top25 residual DMR morula accessibility observed mean versus 1000 matched-random distribution.
- Top-k boundary panel showing top25 positive, top50/top100 not q95-positive.
- Boundary inset: overlap fraction and morula-minus-8cell delta are not specific.

Claim: public human morula accessibility partially supports the most extreme residual DMRs, but does not identify complete causal u_bio.

## Table 1. Evidence and boundary table

Use `CSB_TRO_2026-05-27_evidence_boundary_table.tsv`.

## Table 2. What is solved and what is not solved

Use `CSB_TRO_2026-05-27_claim_boundary_solved_unsolved_v1.0.tsv`.

## Table 3. Perturbation-informed machinery support

Use `CSB_TRO_2026-05-27_perturbation_machinery_support.tsv`.

Claim: CBP/p300 and HDAC perturbation literature supports the biological plausibility of a perturbable access/closure chromatin axis, but does not provide paired methylation correction causality.

## Table 4. Causal boundary table

Use `CSB_TRO_2026-05-27_causal_boundary_table_v1.0.tsv`.

Claim: the strongest current framing is perturbation-informed chromatin-associated diagnostic control dynamics; direct causal u_bio detection remains unresolved without paired perturbation methylation readout.
