## Supplemental docs for "An entropy-based diagnostic framework for characterizing methylation state dynamics during preimplantation development": CSB_TRO_2026-05-27_manuscript_v1.0_u_bio_rescue_integrated.md

# Perturbation-informed operator-time methylation dynamics reveals a structured morula-entry correction architecture with partial public chromatin rescue

## Abstract

Early embryonic methylation reset is usually analyzed through stage-wise methylation changes, but this does not directly test whether methylation state propagation is sufficient to generate the morula reset-basin. We constructed a DMR-level operator-time methylation dynamics framework and evaluated strict methylation-only propagation at morula entry. The methylation-only baseline captured part of the DMR-state trajectory but failed to generate the observed morula basin, with q90 occupancy of 0.044 compared with observed occupancy of 0.875. The missing component could be measured as a diagnostic correction term. This correction was non-random, module-specific, and concentrated in M05/M01/M12/M02/M10 residual coordinates. A dual-branch closure/access architecture organized the correction with high occupancy (0.956), high directional alignment, and strong sign dependence; wrong closure orientation collapsed occupancy to 0.000. A new entry-exit analysis further showed that 8-cell-to-morula and morula-to-blastocyst changes are non-randomly anti-aligned in DMR state space (all-DMR duality score=0.699 versus exit-permutation q95=0.133), supporting morula as a reset-basin geometric vertex.

Public RNA and motif x TF surrogates were weak replacements. Histone-associated evidence supported branch plausibility but remained diagnostic because public embryo histone matched-random controls were high and strict morula-entry H3K27ac/H3K4me3 inputs are controlled-access. We therefore performed a public chromatin rescue audit using stage-matched human embryo accessibility data. This identified a limited but positive signal: the top25 residual DMRs showed higher human morula accessibility than 1000 matched-random controls (observed mean=1.660; random median=1.321; random q95=1.566). A non-morula independent ATAC control did not show generic top residual DMR enrichment. Finally, published mouse preimplantation perturbation data connect the inferred access/closure architecture to experimentally perturbable CBP/p300-HDAC H3K27ac machinery, although these data lack methylation readout. Together, these results support a perturbation-informed chromatin-associated diagnostic control framework in which morula-entry failure is structured, biologically constrained, and partly stage-matched chromatin-supported, but not causally identified as u_bio.

## Results

### Methylation-only operator-time dynamics fails at morula basin entry

We first evaluated whether DMR methylation state propagation alone could account for morula reset-basin entry. The methylation-only baseline produced q90 morula basin occupancy of 0.044, whereas observed morula occupancy was 0.875. This established the methylation-only operator as an incomplete but necessary baseline rather than a complete developmental reset model.

### The missing component is measurable as a diagnostic correction

The failure component was defined as the measured correction between the observed morula state and the methylation-only morula prediction. This correction is an upper-bound diagnostic because it uses held-out morula methylation. An alpha scan showed threshold-like basin entry, with alpha_to_observed around 0.50 and occupancy@alpha1 of 1.000.

### The correction is non-random and module-specific

Residual module reconstruction identified a compact correction structure dominated by M05/M01/M12/M02/M10. Greedy reconstruction reached occupancy of 0.867 using M05/M01/M12 and 0.956 after adding M02. Top residual DMR matched-random controls strongly supported non-random structure: top25 observed occupancy was 0.956, while matched-random q95 was 0.156 and max was 0.200.

### A dual-branch closure/access architecture organizes the correction

The strongest diagnostic architecture separated closure-like M05/M01/M12 modules from access-like M02/M10 modules. Correct closure/access orientation reached q90 occupancy of 0.956. Wrong closure with correct access produced occupancy of 0.000, correct closure with wrong access produced 0.178, and both wrong signs produced 0.000. This sign dependence indicates that the branch architecture is not an arbitrary residual fit.

### Morula-centered entry-exit duality reveals reset-basin geometry

We then tested whether morula is only a failed prediction point or whether it acts as a geometric vertex in DMR methylation state space. For each DMR, we defined the entry change as beta_morula minus beta_8cell and the exit change as beta_blastocyst minus beta_morula. A morula-centered entry-exit duality predicts that these vectors should be oppositely oriented, with positive curvature beta_8cell minus 2 beta_morula plus beta_blastocyst for U-shaped DMRs.

Across all 156 DMRs, entry and exit vectors were strongly anti-aligned (cosine=-0.699; duality score=0.699). This exceeded an exit-vector permutation q95 of 0.133, supporting non-random morula-centered geometry. The effect was not a strict global U-shape: 40.4% of DMRs were U-shaped. Top residual DMRs showed module-matched enrichment of this geometry, with top25 duality score=0.587 versus random q95=0.577, top50=0.727 versus q95=0.619, and top100=0.735 versus q95=0.714. The access branch was especially anti-aligned (M02/M10 duality score=0.872; random q95=0.871), driven mainly by M02 (duality score=0.994). Closure modules were weaker than size-matched random background. Thus, the entry-exit result supports a morula-centered reset-basin geometry and top residual enrichment, but it is branch- and module-specific rather than a globally symmetric parabolic rebound.

### Public surrogate layers define the biological boundary

RNA and motif x TF surrogates were weak replacements, with global RNA occupancy around 0.200 and motif x TF occupancy around 0.222. hESC histone and public embryo histone contrasts supported branch plausibility, but public embryo histone matched-random controls were high: true diagnostic occupancy was 0.978 and random median was also 0.978. Therefore, public histone evidence is diagnostic support, not final morula-entry replacement. Strict morula-entry histone replacement remains limited by missing controlled-access H3K27ac_morula and H3K4me3_morula tracks.

### Stage-matched public chromatin partially rescues the top residual DMR signal

To test whether public chromatin data could move the framework beyond diagnostic plausibility, we audited stage-matched human embryo accessibility data. Liu2019 human embryo LiCAT/accessibility supplementary data yielded 19691 coordinate-like public chromatin regions that could be overlapped with residual DMRs.

The strongest result was restricted to the most extreme residual set. For top25 residual DMRs, mean human morula accessibility was 1.660, exceeding the 1000 matched-random q95 of 1.566 (random median=1.321; random max=1.724). This supports a stage-matched public chromatin partial-replacement signal for the top residual DMRs. The effect should not be generalized globally: top50 and top100 did not exceed matched-random q95, overlap fraction was not specific, and morula-minus-8cell accessibility delta did not exceed matched-random q95.

### Known perturbable chromatin machinery supports the access/closure interpretation

We next asked whether the inferred access/closure architecture aligns with known perturbable chromatin machinery. GSE207222 provides mouse preimplantation embryo perturbation data in which CBP/p300 activity was inhibited with A485 and HDAC activity with TSA, followed by RNA-seq and ATAC-seq comparisons. The GEO record and associated study report that CBP/p300 and HDAC activities regulate H3K27ac dynamics, chromatin opening or transition, ZGA, and preimplantation development.

This evidence does not test the methylation correction directly, because the perturbation readouts are RNA and ATAC rather than methylation at residual DMRs. It does, however, support a perturbation-informed machinery interpretation: the inferred access branch is compatible with CBP/p300-H3K27ac chromatin-opening machinery, and the closure branch is compatible with HDAC/deacetylation machinery. This upgrades the biological support from chromatin-associated diagnostic evidence to perturbation-informed chromatin-associated diagnostic evidence, without crossing the causal u_bio detection boundary.

## Discussion

This study establishes a public-data-bounded diagnostic control framework for morula-entry methylation reset. The central result is not that a final causal u_bio has been identified. Instead, the results show that methylation-only dynamics fails in a structured way, and that the measured missing term has module specificity, direction sensitivity, and a dual-branch closure/access organization.

The strongest evidence comes from negative controls. Branch sign controls showed that incorrect closure orientation destroys basin entry, and top residual DMR matched-random controls showed that the high-residual DMR set is far stronger than matched random backgrounds. These controls support the interpretation that the missing correction is not a random mathematical residual.

The new public chromatin rescue result improves the biological footing of the framework. A stage-matched human morula accessibility signal is enriched in the top25 residual DMRs beyond matched-random q95. This is the first current result that directly links the most extreme residual methylation coordinates to a public morula-stage chromatin measurement. However, the signal is narrow: it is accessibility-level support for the top residual set, not a full branch-level, top100-level, or perturbation-validated replacement of u_bio.

An independent public ATAC boundary control further constrains the claim. GSE101571 human 8-cell and ICM ATAC peak BEDs were overlapped with residual DMRs using the same top-k matched-random logic. No top-k/source comparison exceeded matched-random q95. This does not refute the Liu2019 morula signal because the stages are not morula-matched, but it argues against a broad, stage-independent accessibility enrichment explanation.

The perturbation-informed layer further improves interpretability without changing the causal boundary. Published CBP/p300 and HDAC perturbation data in mouse preimplantation embryos show that the access/closure chromatin axis is experimentally perturbable and developmentally relevant. This supports the biological plausibility of the inferred architecture, but it does not show that perturbing the candidate chromatin input changes the methylation correction term or the next methylation state.

The main biological boundary remains paired perturbation-to-methylation readout. Public embryo histone contrasts are compatible with the closure/access architecture, but matched-random histone controls are also high. Strict morula-entry H3K27ac and H3K4me3 tracks are not available in the local public data package and are tied to controlled-access sources. More importantly, no current public layer tests do(u_bio) -> Delta c_tau -> Delta x_tau+1. The correct interpretation is therefore perturbation-informed chromatin-associated diagnostic architecture with partial stage-matched rescue, not final biological replacement.

## Claim boundary

Supported claim:

Methylation-only operator-time dynamics captures baseline DMR-state propagation but fails at morula reset-basin entry. The resulting measured correction is non-random, module-specific, direction-sensitive, and organized by a dual-branch closure/access diagnostic architecture. Public histone-associated evidence supports biological plausibility, while RNA and motif x TF surrogates provide weak replacement signals. Stage-matched public human morula accessibility partially supports the most extreme residual DMRs. Independent non-morula ATAC data do not show generic top residual DMR enrichment, and published CBP/p300-HDAC perturbation studies support the access/closure architecture as a perturbable chromatin machinery axis.

Unsupported claim:

- The final causal biological control input u_bio has been fully identified.
- The available evidence demonstrates do(u_bio) -> Delta c_tau -> Delta x_tau+1.

## Future work

The immediate public-data upgrade is to test whether an independent human embryo accessibility or histone dataset reproduces the top25 residual-DMR morula accessibility signal. A non-morula GSE101571 ATAC peak control did not reproduce the signal, so it should be treated as boundary evidence rather than replication. Gao2018/CRA000297 human morula DNase-seq is the strongest independent stage-matched candidate, but the public directory currently exposes raw paired-end files rather than processed peak/signal tracks. If an independent stage-matched source reproduces the signal, the result can be promoted from supplementary support to a main-text chromatin rescue result. If not, the current Liu2019 signal should remain a bounded supplementary support layer.

The decisive causal upgrade is paired perturbation-to-methylation readout: for example, CBP/p300 inhibition, HDAC perturbation, TF perturbation, or histone-mark perturbation followed by methylation measurement at the residual DMRs and evaluation of the predicted correction change. Longer-term extensions include matched ATAC/histone profiling, perturbation validation, cross-species replication, and continuous-time or optimal-transport formulations. These are future directions and are not required for the current perturbation-informed diagnostic manuscript.
