## Supplemental docs for "An entropy-based diagnostic framework for characterizing methylation state dynamics during preimplantation development": CSB_TRO_2026-05-27_perturbation_informed_summary.md

# Perturbation-informed chromatin machinery support

## Goal

Add a lightweight perturbation-informed evidence layer without overclaiming causal u_bio detection.

## Main result

GSE207222 provides a relevant mouse preimplantation perturbation dataset: CBP/p300 activity was perturbed with A485 and HDAC activity with TSA, followed by RNA-seq and ATAC-seq comparisons. The GEO record also reports H3K27ac CUT&RUN profiling across mouse early embryo stages, including morula.

This supports the biological plausibility of the inferred access/closure architecture:

- CBP/p300 aligns with an access/H3K27ac writer axis.
- HDAC aligns with a closure/deacetylation axis.
- The combined CBP/p300-HDAC axis is a real perturbable chromatin machinery layer in preimplantation embryos.

## Boundary

This does not close the causal u_bio loop because GSE207222 does not provide paired methylation readout after A485/TSA perturbation at the residual DMRs.

Therefore the upgraded claim is:

`stage-matched chromatin-supported, perturbation-informed diagnostic control dynamics`

The unsupported claim remains:

`causal u_bio detected`

## Files

- CSB_TRO_2026-05-27_perturbation_machinery_support.tsv
- CSB_TRO_2026-05-27_causal_boundary_table_v1.0.tsv
- CSB_TRO_2026-05-27_GSE207222_perturbation_audit.tsv
- perturbation_GSE207222_geo.html
