## Supplemental docs for "An entropy-based diagnostic framework for characterizing methylation state dynamics during preimplantation development": CSB_TRO_2026-05-27_second_source_experiment_status.md

# Second-source accessibility experiment status

## Completed

An independent GSE101571 human ATAC peak overlap control was run against residual DMRs with 1000 matched-random iterations.

Result: no top-k/source comparison exceeded matched-random q95.

Most relevant top25 values:

- 8cell_2pn: observed=0.040, random median=0.140, q95=0.280
- 8cell_3pn: observed=0.040, random median=0.160, q95=0.280
- icm_2pn: observed=0.000, random median=0.080, q95=0.200
- icm_3pn: observed=0.000, random median=0.040, q95=0.080

## Interpretation

This does not reproduce the Liu2019 top25 human morula accessibility signal. Because GSE101571 is not morula-stage, the result should be interpreted as a boundary control rather than a failed replication of the stage-matched Liu2019 signal.

Current claim status:

- Liu2019 remains a limited positive stage-matched morula accessibility rescue for top25 residual DMRs.
- GSE101571 does not support a general non-morula ATAC enrichment of top residual DMRs.
- The top25 rescue should not be promoted to a strong main-text result without a true independent morula-stage source.

## Next gate

Continue searching for processed Gao2018/CRA000297 human morula DHS peak/signal files. Raw fastq exists for SAMC013224/SAMC013225, but full raw DNase-seq reprocessing is outside the lightweight replication path unless explicitly chosen.
