## Supplemental docs for "An entropy-based diagnostic framework for characterizing methylation state dynamics during preimplantation development": CSB_TRO_2026-05-27_second_source_replication_triage.md

# Second public source replication triage

## Goal

Test whether an independent public accessibility or histone source reproduces the top25 residual-DMR morula chromatin signal. A reproducible stage-matched signal can be promoted to a stronger main-text result; a non-runnable or negative result keeps Liu2019 as bounded supplementary support.

## Current best second-source candidate

Gao2018 / Cell human early embryo DNase-seq is the highest-priority independent accessibility source because it reports human embryo DHS data across 2-cell, 4-cell, 8-cell, morula, and blastocyst stages. The public repository record is CRA000297 / PRJCA000484, and NGDC BioProject metadata lists human morula DNase-seq replicates SAMC013224 and SAMC013225.

## Why it is suitable

- Independent from Liu2019 LiCAT/accessibility.
- Human embryo rather than cross-species.
- Stage-matched morula accessibility is represented in the metadata.
- Assay is DHS rather than LiCAT/ATAC-like, so replication would not be a same-assay artifact.

## Current blocker

Processed morula peak/signal BED or bigWig files have not yet been located locally. Raw DNase-seq files may be public through NGDC, but full raw reprocessing would be a new analysis rather than a lightweight replication. The next step is to locate processed DHS peak/signal files or supplementary coordinate tables before attempting overlap.

## Decision rule

Use the same top-k and matched-random logic as the Liu2019 audit:

- primary replication: top25 residual DMR morula DHS/accessibility signal exceeds matched-random q95;
- secondary checks: top50/top100, overlap fraction, and morula-minus-8cell contrast;
- upgrade only if the independent stage-matched morula signal is q95-positive under matched random controls;
- otherwise retain Liu2019 as partial supplementary support.

## Status

Triage plus one independent boundary control completed.

GSE101571 human ATAC peak BED files were downloaded and tested as an independent public accessibility source:

- GSE101571_8cell_2pn_peaks.bed.gz: 40426 peaks
- GSE101571_8cell_3pn_peaks.bed.gz: 39498 peaks
- GSE101571_icm_2pn_peaks.bed.gz: 23304 peaks
- GSE101571_icm_3pn_peaks.bed.gz: 8925 peaks

This is not a stage-matched morula rescue because the GEO supplementary files do not provide human morula ATAC peaks. In 1000 matched-random tests, no top-k/source comparison exceeded matched-random q95. Therefore GSE101571 does not replicate or upgrade the Liu2019 top25 morula accessibility signal. It is a boundary control: non-morula independent ATAC peaks do not show the same residual-DMR enrichment.

Output files:

- CSB_TRO_2026-05-27_GSE101571_ATAC_overlap_summary.tsv
- CSB_TRO_2026-05-27_GSE101571_ATAC_matched_random.tsv
- CSB_TRO_2026-05-27_GSE101571_ATAC_DMR_overlap.tsv
- CSB_TRO_2026-05-27_GSE101571_ATAC_overlap_interpretation.md

No second-source morula-stage replication claim should be added until processed Gao2018 morula DHS coordinates or signal tracks are obtained and overlapped.

## Gao2018 / CRA000297 exact morula raw mapping

The NGDC GSA page 2 identifies the stage-matched human morula DNase-seq runs:

- DNase-seq morula rep1: CRX018058 / CRR019758 / SAMC013224
- DNase-seq morula rep2: CRX018059 / CRR019759 / SAMC013225

Public download directories expose raw files only:

- CRR019758_f1.gz: 9725161618 bytes
- CRR019758_r2.gz: 11619481608 bytes
- CRR019759_f1.gz: 8372644189 bytes
- CRR019759_r2.gz: 10123343152 bytes

No processed BED/bigWig/peak file is present in the public CRR019758/CRR019759 directories. Full raw DNase-seq reprocessing would require downloading roughly 39.8 GB compressed data for the two morula replicates, plus alignment, filtering, peak calling, genome-build harmonization, and matched-random overlap. This is a different, heavier experiment than the lightweight public processed-track rescue.
