## Supplemental docs for "An entropy-based diagnostic framework for characterizing methylation state dynamics during preimplantation development": CSB_TRO_advanced_latent_dynamics_interpretation.md

# Advanced latent operator-time dynamics

This package upgrades the prior leave-morula-out point prediction into an autonomous latent-state dynamics analysis. The fitted drift uses only latent state and stage-anchored operator time, so it can be integrated across multiple stages without supplying future-stage summary variables.

- 8-cell to morula deterministic rollout DMR mean RMSE: 0.2471
- 8-cell to morula latent MMD: 1.0362
- 8-cell to morula predicted basin occupancy: 0.0222
- MII to blastocyst deterministic rollout DMR mean RMSE: 0.1537

Jacobian eigenvalues of the autonomous latent drift:
- strict leave-morula autonomous 8-cell to morula DMR mean RMSE: 0.2517
- eigen 1: real=-3.6225, imag=0.0000, class=contracting
- eigen 2: real=-5.8052, imag=0.6561, class=contracting
- eigen 3: real=-5.8052, imag=-0.6561, class=contracting

Largest perturbation effects on morula prediction:
- PC1_increase_1sd_at_8cell: delta RMSE=0.0260
- remove_top50_latent_loading_DMRs_from_score: delta RMSE=0.0115
- PC3_decrease_1sd_at_8cell: delta RMSE=0.0015
- PC2_decrease_1sd_at_8cell: delta RMSE=0.0004
- PC2_increase_1sd_at_8cell: delta RMSE=0.0000
- none: delta RMSE=0.0000

Interpretation boundary: this is still stage-anchored operator time, not real longitudinal tracking of the same embryo. The stochastic residual simulation is a low-dimensional residual diffusion approximation, not a full Fokker-Planck population PDE.
