## Supplemental docs for "An entropy-based diagnostic framework for characterizing methylation state dynamics during preimplantation development": CSB_TRO_basin_residual_control_field_interpretation.md

# Basin residual control-field diagnostic

This analysis treats the strict non-leaking methylation-only failure as a missing control-field problem. It defines the missing morula basin direction in latent space and decodes that direction back to DMR and module space.

Observed morula center: -3.1743, -0.0308, -2.8184
Strict pre-morula predicted center: -2.3829, 0.0031, -0.9009
Missing basin residual vector: -0.7914, -0.0338, -1.9175
Residual norm: 2.0747

Top modules carrying the decoded residual:
- M01: fraction_abs_residual=0.235, cosine=-0.362, top_DMR=cluster_4373
- M05: fraction_abs_residual=0.159, cosine=0.131, top_DMR=cluster_2623
- M10: fraction_abs_residual=0.080, cosine=-0.493, top_DMR=cluster_6262
- M12: fraction_abs_residual=0.062, cosine=-0.141, top_DMR=cluster_6498
- M07: fraction_abs_residual=0.061, cosine=0.004, top_DMR=cluster_2743

Diagnostic control results:
- latent_oracle_alpha_1: occupancy_q90=1.000, DMR_mean_RMSE=0.2383, cosine=1.000
- top50_DMR_restricted_residual_control: occupancy_q90=1.000, DMR_mean_RMSE=0.2393, cosine=0.983
- top100_DMR_restricted_residual_control: occupancy_q90=1.000, DMR_mean_RMSE=0.2386, cosine=0.998
- latent_oracle_alpha_0.75: occupancy_q90=0.978, DMR_mean_RMSE=0.2392, cosine=1.000
- latent_oracle_alpha_1.25: occupancy_q90=0.956, DMR_mean_RMSE=0.2380, cosine=1.000
- top25_DMR_restricted_residual_control: occupancy_q90=0.956, DMR_mean_RMSE=0.2401, cosine=0.968
- latent_oracle_alpha_0.5: occupancy_q90=0.933, DMR_mean_RMSE=0.2406, cosine=1.000
- top10_DMR_restricted_residual_control: occupancy_q90=0.600, DMR_mean_RMSE=0.2415, cosine=0.922

Overlap with existing dynamic DMR sets:
- top100_morula_entry_DMRs: overlap=78/100, expected_random=64.1
- top100_dynamic_reset_DMRs: overlap=76/100, expected_random=64.1
- top100_blastocyst_exit_DMRs: overlap=71/100, expected_random=64.1
- top100_latent_loading_DMRs: overlap=78/100, expected_random=64.1

Best diagnostic control occupancy is 1.000. These controls use the observed morula residual and are therefore not strict predictors. Their purpose is to identify candidate DMR modules and genomic regions that could carry a missing regulatory/chromatin basin-attraction field.

Interpretation boundary: this does not prove that the listed DMRs are causal or that the hidden field is already measured. It provides a prioritized target set for RNA, ATAC, histone-mark, TF motif, DNMT/TET, and nearby-gene annotation.
