## Supplemental docs for "An entropy-based diagnostic framework for characterizing methylation state dynamics during preimplantation development": CSB_TRO_bifurcation_like_basin_entry_summary.md

# Bifurcation-like Basin Entry Scan

Status: `completed`

This analysis treats the measured missing correction term as a diagnostic control parameter and asks whether morula basin entry shows threshold-like behavior.

Important boundary: this supports a bifurcation-like or tipping-like basin-entry interpretation, not a proof of a strict saddle-node bifurcation.

## Full Correction Scan

- Occupancy at alpha=0: 0.044
- Occupancy at alpha=1: 1.000
- First alpha reaching observed q90 occupancy: 0.50
- Steepest local occupancy slope begins near alpha: 0.45

## Module Scan

- M05+M01+M12+M02: occupancy alpha=1 is 0.956; first alpha to observed occupancy is 0.85
- M05+M01+M12+M02+M10: occupancy alpha=1 is 0.956; first alpha to observed occupancy is 0.75
- M05+M01+M12: occupancy alpha=1 is 0.867; first alpha to observed occupancy is 1.05
- M05+M01: occupancy alpha=1 is 0.600; first alpha to observed occupancy is 1.35
- M05: occupancy alpha=1 is 0.422; first alpha to observed occupancy is not_reached

## Jacobian Boundary

The current methylation operator is affine in latent z, and the measured correction is added as a constant control vector. Therefore the diagnostic alpha scan changes endpoint basin entry but does not by itself change the affine methylation-only Jacobian. A future biological u_bio(z,tau) or basin-coupled control model is needed to test true local stability changes.
