## Supplemental docs for "An entropy-based diagnostic framework for characterizing methylation state dynamics during preimplantation development": CSB_TRO_branch_bound_biological_control_summary.md

# Branch-Bound Biological Control Candidates

Status: `completed_with_available_surrogates_no_histone_tracks`

Goal: start replacing the dual-branch proxy with biologically named closure/access variables while explicitly preserving boundaries.

## Top Candidates

- biocontrol_ATAC_TF_RNA_composite: status=ATAC_TF_branch_surrogate; max_occ=0.200; occ@1=0.111; cosine@1=0.415; PC3@1=0.146; signflip_max=0.044
- biocontrol_ATAC_closure_plus_ATAC_access: status=ATAC_branch_proxy; max_occ=0.200; occ@1=0.156; cosine@1=0.386; PC3@1=0.131; signflip_max=0.044
- closure_ATAC_loss_ATAC_8cell_3pn: status=chromatin_proxy_boundary_no_morula_ATAC; max_occ=0.200; occ@1=0.111; cosine@1=0.415; PC3@1=0.146; signflip_max=0.044
- mitochondrial_oxphos_energy_program: status=surrogate_program_not_direct_histone_track; max_occ=0.178; occ@1=0.044; cosine@1=0.910; PC3@1=0.061; signflip_max=0.044
- biocontrol_ATAC_closure_plus_TF_access: status=ATAC_TF_branch_surrogate; max_occ=0.156; occ@1=0.133; cosine@1=0.367; PC3@1=0.161; signflip_max=0.044
- metabolic_onecarbon_sam_program: status=surrogate_program_not_direct_histone_track; max_occ=0.111; occ@1=0.044; cosine@1=0.957; PC3@1=0.034; signflip_max=0.044
- closure_ATAC_loss_ATAC_8cell_2pn: status=chromatin_proxy_boundary_no_morula_ATAC; max_occ=0.067; occ@1=0.044; cosine@1=0.137; PC3@1=0.089; signflip_max=0.044
- access_promoter_ATAC_ATAC_8cell_2pn: status=real_ATAC_promoter_access_proxy_no_morula_stage; max_occ=0.044; occ@1=0.044; cosine@1=0.106; PC3@1=-0.015; signflip_max=0.044
- access_linked_RNA_delta_surrogate: status=weak_surrogate_gene_linked_not_histone; max_occ=0.044; occ@1=0.044; cosine@1=0.106; PC3@1=-0.015; signflip_max=0.044
- closure_histone_repression_program: status=surrogate_program_not_direct_histone_track; max_occ=0.044; occ@1=0.044; cosine@1=-0.957; PC3@1=-0.042; signflip_max=0.156
- access_promoter_activation_program: status=surrogate_program_not_direct_histone_track; max_occ=0.044; occ@1=0.044; cosine@1=-0.910; PC3@1=-0.051; signflip_max=0.178
- access_TF_activity_q05_sparse_flipped_activity_sign: status=promoter_TF_surrogate; max_occ=0.044; occ@1=0.044; cosine@1=-0.106; PC3@1=0.015; signflip_max=0.044

## Boundary

No H3K27ac/H3K4me3/H3K27me3 track is currently analysis-ready, so none of these should be called final histone u_bio. The strongest current candidates are branch-bound ATAC/TF/RNA surrogates. The final replacement still requires direct histone or chromatin-state tracks.
