## Supplemental docs for "An entropy-based diagnostic framework for characterizing methylation state dynamics during preimplantation development": CSB_TRO_chromatin_gated_control_summary.md

# Residual Module Chromatin-Gated Control Analysis

Status: `completed_with_available_ATAC_and_missing_histone_inputs`

Goal: test whether M05/M01/M12/M02/M10 residual modules are supported by external chromatin state, and whether chromatin-gated controls explain the measured correction term.

## Region Composition

- M01: n=21, promoter2kb=0.333, intergenic=0.619, residual_sum=1.260
- M05: n=6, promoter2kb=0.333, intergenic=0.667, residual_sum=0.852
- M10: n=3, promoter2kb=0.667, intergenic=0.333, residual_sum=0.431
- M12: n=4, promoter2kb=0.750, intergenic=0.250, residual_sum=0.333
- M02: n=30, promoter2kb=0.700, intergenic=0.100, residual_sum=0.326

## Strongest Available Chromatin Overlaps

- M10 ATAC_8cell_3pn: target=0.333, background=0.000, OR=76.200, q=0.182
- M02 ATAC_ICM_3pn: target=0.100, background=0.002, OR=53.412, q=0.0022
- M02 ATAC_8cell_3pn: target=0.333, background=0.011, OR=42.580, q=2.37e-09
- M02 ATAC_8cell_2pn: target=0.233, background=0.011, OR=26.532, q=5.35e-06
- M10 ATAC_8cell_2pn: target=0.333, background=0.016, OR=25.000, q=0.298
- M12 ATAC_8cell_2pn: target=0.000, background=0.000, OR=18.778, q=1
- M12 ATAC_ICM_3pn: target=0.000, background=0.000, OR=18.778, q=1
- M12 ATAC_8cell_3pn: target=0.000, background=0.000, OR=18.778, q=1

## Control Dynamics

- motif_TF_x_ATAC8cell_gate: occupancy=0.222, cosine=0.452, PC3_recovery=0.094
- ATAC_8cell_2pn_chromatin_only: occupancy=0.000, cosine=-0.710, PC3_recovery=-0.217
- ATAC_8cell_3pn_chromatin_only: occupancy=0.000, cosine=-0.701, PC3_recovery=-0.211
- ATAC_ICM_2pn_chromatin_only: occupancy=0.000, cosine=-0.496, PC3_recovery=-0.125
- ATAC_ICM_3pn_chromatin_only: occupancy=0.000, cosine=-0.483, PC3_recovery=-0.114

## Boundary

The available local chromatin evidence is ATAC 8-cell/ICM only. This is not direct morula chromatin validation.

Missing histone tracks are input/access boundaries, not negative histone evidence:

- H3K27ac_8cell
- H3K27ac_morula
- H3K27ac_blastocyst
- H3K4me3_8cell
- H3K4me3_morula
- H3K4me3_blastocyst
