## Supplemental docs for "An entropy-based diagnostic framework for characterizing methylation state dynamics during preimplantation development": CSB_TRO_composite_biocontrol_proxy_summary.md

# Composite Biocontrol Proxy Alpha Scan

Status: `completed`

This is route C: it does not replace missing histone-state data, but tests whether available RNA/ATAC/motif/region proxy composites can approximate a module-specific biological control term.

Important boundary: models marked as `orientation_corrected_proxy`, `routeC_proxy`, or `module_hypothesis_upper_bound` are not final non-leaking biological u_bio. They are attack-mode diagnostics for what must be achieved by real histone/chromatin inputs.

## Top Alpha-Scan Results

- ATAC_8cell_2pn_chromatin_only_inverse: max_occ=0.511 at alpha=1.90; occ@1=0.333; cosine@1=0.710; PC3@1=0.217; status=orientation_corrected_proxy_uses_measured_direction_boundary
- available_external_composite_RNA_inverseATAC_motif: max_occ=0.511 at alpha=1.95; occ@1=0.289; cosine@1=0.707; PC3@1=0.189; status=composite_proxy_methylation_non_leaking_inputs_but_orientation_sensitive
- ATAC_8cell_3pn_chromatin_only_inverse: max_occ=0.511 at alpha=1.60; occ@1=0.333; cosine@1=0.701; PC3@1=0.211; status=orientation_corrected_proxy_uses_measured_direction_boundary
- region_state_prior: max_occ=0.378 at alpha=1.35; occ@1=0.311; cosine@1=0.637; PC3@1=0.238; status=static_genomic_proxy_not_external_dynamic
- routeC_chromatin_state_proxy_RNA_inverseATAC_region: max_occ=0.333 at alpha=1.40; occ@1=0.289; cosine@1=0.614; PC3@1=0.205; status=routeC_proxy_not_final_biological_control
- core_module_positive_control_hypothesis: max_occ=0.311 at alpha=1.95; occ@1=0.156; cosine@1=0.751; PC3@1=0.142; status=diagnostic_module_hypothesis_upper_bound_not_external_omics
- ATAC_ICM_2pn_chromatin_only_inverse: max_occ=0.267 at alpha=1.55; occ@1=0.222; cosine@1=0.496; PC3@1=0.125; status=orientation_corrected_proxy_uses_measured_direction_boundary
- ATAC_ICM_3pn_chromatin_only_inverse: max_occ=0.267 at alpha=1.65; occ@1=0.222; cosine@1=0.483; PC3@1=0.114; status=orientation_corrected_proxy_uses_measured_direction_boundary

## Interpretation

If a proxy reaches high occupancy only after orientation correction or module-hypothesis encoding, it does not prove u_bio. It defines the required direction and module pattern for the histone/chromatin data acquisition stage.
