## Supplemental docs for "An entropy-based diagnostic framework for characterizing methylation state dynamics during preimplantation development": CSB_TRO_diagnostic_control_framework_paper_skeleton.md

# Paper Skeleton: Diagnostic Latent Control Framework

Working title:

```text
A diagnostic latent-control framework reveals a missing module-level basin-entry force in human preimplantation methylation dynamics
```

## Central Claim

Methylation-only latent dynamics can recover the morula mean-state trajectory but cannot autonomously generate the morula population basin. A measured missing correction term rescues basin entry, localizes to residual modules M05/M01/M12/M02/M10, and produces threshold-like occupancy behavior. Existing RNA, nearest-gene RNA, ATAC, and simple motif x TF signals do not fully explain this correction term; histone-state inputs are the current data-access boundary for identifying the true biological control variable.

## Figure Plan

### Figure 1: Latent methylation dynamics framework

- DMR state matrix and operator time.
- Stage-anchored latent representation.
- Model form:

```text
dz/dtau = f_meth(z,tau) + B u_bio(tau)
```

### Figure 2: Mean-state prediction works, distribution basin fails

- Strict leave-morula-out latent/module improvement.
- Methylation-only deterministic/SDE occupancy failure.
- Observed morula basin occupancy versus strict prediction.

### Figure 3: Measured missing correction term

- Correction vector:

```text
delta_z = [-0.7914, -0.0338, -1.9175]
```

- Strong PC3-negative and moderate PC1-negative pull.
- Measured correction upper bound rescues occupancy.

### Figure 4: Residual module decomposition

- Greedy module accumulation:

```text
M05 = 0.422
M05+M01 = 0.600
M05+M01+M12 = 0.867
M05+M01+M12+M02 = 0.956
```

- Sign-flip and matched random controls.
- Region composition: M01/M05 distal/intergenic-enriched; M02/M12 more promoter-proximal.

### Figure 5: External biological candidates are partial

- Global RNA direction is strong but coarse.
- Nearest-gene RNA is weak.
- M02-KLF4/KLF5 motif x TF is exploratory and encoding-sensitive.
- ATAC supports M02 accessibility but ATAC-only control direction is opposite to the measured correction.

### Figure 6: Bifurcation-like basin entry

- Full correction alpha scan:

```text
alpha=0 occupancy = 0.044
alpha=1 occupancy = 1.000
first alpha reaching observed occupancy = 0.50
steepest slope near alpha = 0.45
```

- Module-specific alpha scan.
- Wording: threshold-like / bifurcation-like / tipping-like basin entry, not strict saddle-node proof.

### Figure 7: Histone-state boundary and future u_bio

- Histone-state control framework implemented.
- Required inputs: H3K27ac/H3K4me3/H3K27me3 8-cell/morula/blastocyst peak or signal tracks.
- Current status: not_run_missing_histone_input.

## Results Outline

### Result 1

Stage-anchored latent/module methylation dynamics improves strict morula mean-state prediction.

### Result 2

Methylation-only stochastic distribution dynamics fails to generate morula basin occupancy.

### Result 3

A compact measured correction term rescues morula basin entry and defines the target that real u_bio must explain.

### Result 4

Residual decomposition localizes the correction to M05/M01/M12/M02/M10 and shows directional, non-random module structure.

### Result 5

External RNA and motif/ATAC analyses are informative but incomplete: global RNA aligns with direction, nearest-gene RNA is too coarse, M02-KLF4/KLF5 is encoding-sensitive, and ATAC is local/position-support rather than full directional control.

### Result 6

Correction-amplitude and module-specific scans show threshold-like basin entry consistent with a control-induced bifurcation-like transition.

### Result 7

Histone-state gating is the most plausible next biological-control layer, but current local data lack processed H3K27ac/H3K4me3/H3K27me3 peak/signal tracks.

## Wording Boundaries

Do say:

```text
threshold-like basin-entry behavior
bifurcation-like transition
control-induced basin entry
measured missing correction term
diagnostic residual module coordinates
```

Do not say:

```text
strict saddle-node bifurcation proven
M02-KLF4/KLF5 is the true u_bio
ATAC validates the full correction
histone is negative
top residual DMRs are causal drivers
```

## Minimum Publishable Story

Even without histone inputs, the project can be framed as a diagnostic control framework:

```text
1. detect methylation-only model boundary;
2. measure the missing correction term;
3. show threshold-like basin-entry rescue;
4. decompose correction into modules;
5. use external omics to show partial support and identify the missing histone/chromatin boundary.
```

## Stronger Story If Histone Inputs Arrive

If H3K27ac/H3K4me3/H3K27me3 peak or signal tracks are obtained:

```text
1. test M05/M01 distal H3K27ac enhancer-like control;
2. test M02/M12 promoter H3K4me3 control;
3. test H3K27me3 repression/release modules;
4. build histone-gated TF activity;
5. evaluate occupancy/cosine/PC3 recovery/sign-flip/matched-random controls.
```
