## Supplemental docs for "An entropy-based diagnostic framework for characterizing methylation state dynamics during preimplantation development": CSB_TRO_dual_branch_ablation_summary.md

# Dual-Branch Ablation, Sign, and Random Controls

Status: `completed`

This tests whether the M05/M01/M12 closure branch plus the signed M02/M10 access branch is a stable structure rather than a parameter accident.

## Main Model

- full dual-branch occupancy=0.956, cosine=0.998, PC3_recovery=0.680, center_error=-0.074

## Ablation

- closure_branch_only: occupancy=0.600, cosine=0.945, PC1=0.336, PC3=0.399, center_error=0.406
- access_branch_only: occupancy=0.422, cosine=0.917, PC1=0.241, PC3=0.281, center_error=0.629
- remove_M05: occupancy=0.778, cosine=0.968, PC1=0.499, PC3=0.447, center_error=0.305
- remove_M01: occupancy=0.933, cosine=0.987, PC1=0.333, PC3=0.590, center_error=0.110
- remove_M12: occupancy=0.956, cosine=0.998, PC1=0.563, PC3=0.605, center_error=0.027
- remove_M02: occupancy=0.889, cosine=0.951, PC1=0.639, PC3=0.526, center_error=0.166
- remove_M10: occupancy=0.933, cosine=0.981, PC1=0.274, PC3=0.553, center_error=0.182

## Sign Controls

- correct_closure_correct_access: occupancy=0.956, cosine=0.998, PC3=0.680
- wrong_closure_correct_access: occupancy=0.000, cosine=-0.424, PC3=-0.117
- correct_closure_wrong_access: occupancy=0.178, cosine=0.424, PC3=0.117
- wrong_closure_wrong_access: occupancy=0.000, cosine=-0.998, PC3=-0.680
- naive_inverse_all_modules: occupancy=0.333, cosine=0.701, PC3=0.211
- naive_raw_all_modules: occupancy=0.000, cosine=-0.701, PC3=-0.211

## Summary Stats

- full_dual_branch_occupancy: 0.9556 (beta_closure=1.0,beta_access=1.5)
- full_dual_branch_cosine: 0.9982 (direction alignment)
- closure_only_occupancy: 0.6000 (M05/M01/M12 only)
- access_only_occupancy: 0.4222 (M02/M10 only)
- delta_occupancy_remove_M05: 0.1778 (full minus remove_M05)
- delta_occupancy_remove_M01: 0.0222 (full minus remove_M01)
- delta_occupancy_remove_M12: 0.0000 (full minus remove_M12)
- delta_occupancy_remove_M02: 0.0667 (full minus remove_M02)
- delta_occupancy_remove_M10: 0.0222 (full minus remove_M10)
- sign_control_delta_vs_correct_wrong_closure_correct_access: 0.9556 (positive means correct dual-branch sign is better)
- sign_control_delta_vs_correct_correct_closure_wrong_access: 0.7778 (positive means correct dual-branch sign is better)
- sign_control_delta_vs_correct_wrong_closure_wrong_access: 0.9556 (positive means correct dual-branch sign is better)
- sign_control_delta_vs_correct_naive_inverse_all_modules: 0.6222 (positive means correct dual-branch sign is better)
- sign_control_delta_vs_correct_naive_raw_all_modules: 0.9556 (positive means correct dual-branch sign is better)
- true_partition_percentile_pred_basin_occupancy_q90: 1.0000 (empirical p_ge=0.0960; true=0.9556)
- true_partition_percentile_direction_cosine_to_measured_correction: 1.0000 (empirical p_ge=0.0960; true=0.9982)
- true_partition_percentile_PC3_negative_pull_recovered: 1.0000 (empirical p_ge=0.0960; true=0.6799)
- true_exact_partition_sign_percentile_pred_basin_occupancy_q90: 1.0000 (exact branch-sign null p_ge=0.0500; true=0.9556; n=40)
- true_exact_partition_sign_percentile_direction_cosine_to_measured_correction: 1.0000 (exact branch-sign null p_ge=0.0250; true=0.9982; n=40)
- true_exact_partition_sign_percentile_PC3_negative_pull_recovered: 1.0000 (exact branch-sign null p_ge=0.0250; true=0.6799; n=40)

## Interpretation

The critical test is whether correct branch sign beats wrong-sign and random partition controls. If it does, the dual-branch model is not merely an ATAC proxy with tuned amplitude; it is a structured chromatin-state control hypothesis that can be directly replaced by histone tracks when available.
