## Supplemental docs for "An entropy-based diagnostic framework for characterizing methylation state dynamics during preimplantation development": CSB_TRO_dual_branch_chromatin_state_control_summary.md

# Dual-Branch Chromatin-State Control

Status: `completed`

This formalizes the inverse ATAC result into two candidate control branches:

- closure branch: inverse-ATAC signal on M05/M01/M12
- accessibility branch: M02/M10 accessibility signal tested in raw and inverse orientations

## Branch Vectors

- ATAC_8cell_2pn_chromatin_only_inverse / closure_inverse_ATAC_M05_M01_M12: PC1=-0.186, PC2=-0.296, PC3=-0.747, norm=0.825
- ATAC_8cell_2pn_chromatin_only_inverse / access_inverse_ATAC_M02_M10: PC1=0.172, PC2=-0.076, PC3=0.331, norm=0.380
- ATAC_8cell_2pn_chromatin_only_inverse / access_raw_ATAC_M02_M10: PC1=-0.172, PC2=0.076, PC3=-0.331, norm=0.380
- ATAC_8cell_3pn_chromatin_only_inverse / closure_inverse_ATAC_M05_M01_M12: PC1=-0.266, PC2=-0.291, PC3=-0.764, norm=0.860
- ATAC_8cell_3pn_chromatin_only_inverse / access_inverse_ATAC_M02_M10: PC1=0.127, PC2=-0.157, PC3=0.360, norm=0.413
- ATAC_8cell_3pn_chromatin_only_inverse / access_raw_ATAC_M02_M10: PC1=-0.127, PC2=0.157, PC3=-0.360, norm=0.413

## Beta Grid Summary

- ATAC_8cell_3pn_chromatin_only_inverse + inverse_M02_M10: max_occ=1.000 at beta_closure=1.50, beta_access=-1.50; cosine=0.994; PC3=0.879; first_observed_beta_closure=0.60, beta_access=-1.50
- ATAC_8cell_3pn_chromatin_only_inverse + raw_M02_M10: max_occ=1.000 at beta_closure=1.50, beta_access=1.50; cosine=0.994; PC3=0.879; first_observed_beta_closure=0.60, beta_access=1.50
- ATAC_8cell_3pn_chromatin_only_inverse + raw_M02_only: max_occ=1.000 at beta_closure=1.90, beta_access=1.50; cosine=0.984; PC3=0.911; first_observed_beta_closure=1.00, beta_access=1.40
- ATAC_8cell_2pn_chromatin_only_inverse + inverse_M02_M10: max_occ=1.000 at beta_closure=1.70, beta_access=-1.50; cosine=0.979; PC3=0.921; first_observed_beta_closure=0.70, beta_access=-1.50
- ATAC_8cell_2pn_chromatin_only_inverse + raw_M02_M10: max_occ=1.000 at beta_closure=1.70, beta_access=1.50; cosine=0.979; PC3=0.921; first_observed_beta_closure=0.70, beta_access=1.40
- ATAC_8cell_2pn_chromatin_only_inverse + raw_M02_only: max_occ=1.000 at beta_closure=2.30, beta_access=1.50; cosine=0.959; PC3=1.001; first_observed_beta_closure=1.20, beta_access=1.50
- ATAC_8cell_3pn_chromatin_only_inverse + none: max_occ=1.000 at beta_closure=2.40, beta_access=0.00; cosine=0.945; PC3=0.956; first_observed_beta_closure=1.50, beta_access=0.00
- ATAC_8cell_2pn_chromatin_only_inverse + none: max_occ=0.978 at beta_closure=2.50, beta_access=0.00; cosine=0.929; PC3=0.974; first_observed_beta_closure=1.60, beta_access=0.00

## Mechanistic Interpretation

The basin-entry component is dominated by a closure-like M05/M01/M12 branch. M02/M10 accessibility is not a required positive branch for morula basin entry in this model; when included naively with the same inverse-ATAC orientation, it partly cancels the PC3-negative pull. This separates the missing correction into a primary closure/histone-state branch and a secondary promoter-accessibility branch that must be signed and weighted separately.

This result gives a concrete histone target: H3K27ac/H3K4me3/H3K27me3 data should first be asked whether it supports M05/M01/M12 closure or repressive/poised-state gain, not whether it globally matches all residual modules.
