## Supplemental docs for "An entropy-based diagnostic framework for characterizing methylation state dynamics during preimplantation development": CSB_TRO_dual_branch_structure_validation_summary.md

# Dual-Branch Structure Validation

Status: `completed`

This locks down the proxy structure before replacing it with true histone/chromatin/promoter variables.

## Key Results

- beta_grid_points_total: 1681.0000 (beta_c,beta_a in [0,2] step 0.05)
- beta_grid_strong_region_count: 975.0000 (occupancy>=0.875, cosine>=0.9, PC3>=0.4)
- beta_grid_strong_region_fraction: 0.5800 (continuous parameter robustness proxy)
- beta_grid_strong_beta_closure_min: 0.4500 (lower edge of robust region)
- beta_grid_strong_beta_closure_max: 2.0000 (upper edge of robust region)
- beta_grid_strong_beta_access_min: 0.0000 (lower edge of robust region)
- beta_grid_strong_beta_access_max: 2.0000 (upper edge of robust region)
- module_order_top_1: 0.9556 (closure_subset_plus_full_access; M05,M01|M02,M10; cosine=0.998; PC3=0.605)
- module_order_top_2: 0.9556 (closure_subset_plus_full_access; M05,M01,M12|M02,M10; cosine=0.998; PC3=0.680)
- module_order_top_3: 0.9556 (full_closure_plus_access_subset; M05,M01,M12|M02,M10; cosine=0.998; PC3=0.680)
- module_order_top_4: 0.9333 (closure_subset_plus_full_access; M05,M12|M02,M10; cosine=0.987; PC3=0.590)
- module_order_top_5: 0.9333 (full_closure_plus_access_subset; M05,M01,M12|M02; cosine=0.981; PC3=0.553)
- module_order_top_6: 0.8889 (full_closure_plus_access_subset; M05,M01,M12|M10; cosine=0.951; PC3=0.526)
- module_order_top_7: 0.8444 (closure_subset_plus_full_access; M05|M02,M10; cosine=0.990; PC3=0.515)
- module_order_top_8: 0.7778 (closure_subset_plus_full_access; M01,M12|M02,M10; cosine=0.968; PC3=0.447)
- module_order_top_9: 0.6667 (closure_subset_plus_full_access; M01|M02,M10; cosine=0.963; PC3=0.372)
- module_order_top_10: 0.6000 (closure_subset; M05,M01,M12; cosine=0.945; PC3=0.399)
- true_sign_pattern_rank: 1.0000 (M05:+1,M01:+1,M12:+1,M02:-1,M10:-1)
- true_sign_pattern_occupancy: 0.9556 (cosine=0.994; PC3=0.586)
- true_sign_pattern_percentile_occupancy: 1.0000 (32 sign patterns)

## All-Sign Pattern Result

The biologically proposed sign pattern ranked 1/32 by occupancy then cosine.

True sign pattern:

`M05:+1,M01:+1,M12:+1,M02:-1,M10:-1`

## Interpretation

The dual-branch structure is not a single beta setting and not an arbitrary module sign assignment. It occupies a continuous high-performance beta region and the proposed M05/M01/M12 closure sign with opposite M02/M10 access sign is at the top of the exact sign-pattern ranking.
