## Supplemental docs for "An entropy-based diagnostic framework for characterizing methylation state dynamics during preimplantation development": CSB_TRO_dynamics_master_record.md

# CSB-TRO Dynamics Experiment Record

Date: 2026-05-24

## Purpose

This experiment upgrades the original static TRO score into a constrained distributional dynamics model:

`CSB-TRO = constrained Schrodinger Bridge transgenerational reset operator`

The model treats each developmental stage as an empirical distribution, not a single score. The working state vector is:

`z = [A, Hm, P, Hr]`

- `A`: age-associated epigenetic perturbation from GSE81233 age-DMR methylation.
- `Hm`: methylation entropy from GSE81233 full-genome CpG entropy.
- `P`: developmental potency from GSE36552 RNA single-cell potency.
- `Hr`: RNA expression entropy from GSE36552 RNA single-cell entropy.

## Layer 1: Static-to-Dynamic Pilot

The first pilot used GSE81233 DNA samples and stage-level GSE36552 RNA summaries.

- Samples used: 169
- Morula A rank, rank 1 = lowest perturbation: 1
- Morula P rank, rank 1 = highest potency: 2

This established the CSB-TRO machinery but still used RNA stage means.

## Layer 2: RNA Single-Cell Bridge

The second step replaced RNA stage means with empirical single-cell RNA distributions.

- Main RNA dataset: GSE36552
- GSE36552 cells used after removing aggregate rows: 98
- GSE36552 stages: oocyte, zygote, 2-cell, 4-cell, 8-cell, morula, blastocyst
- Morula P rank, rank 1 = highest potency: 2
- Morula Hr rank, rank 1 = highest RNA entropy: 7
- External direction check: GSE44183, 29 cells

Interpretation: morula is strongly supported by potency but not by raw RNA entropy. This is useful because the model should not claim every dimension peaks at morula.

## Layer 3: Fused CSB-TRO Product-Distribution Bridge

Because the DNA methylation and RNA expression cells are not paired, the fused model uses a stage-wise product distribution:

`p_k(A,Hm,P,Hr) ~= p_k^DNA(A,Hm) x p_k^RNA(P,Hr)`

- Fused particles: 1496
- Stages: 7
- Morula A rank, rank 1 = lowest perturbation: 1
- Morula P rank, rank 1 = highest potency: 2

## Fused Stage Summary

| Stage | Particles | A mean | P mean | Hm mean | Hr mean |
|---|---:|---:|---:|---:|---:|
| MII oocyte | 108 | 0.4007 | 0.3357 | 0.1038 | 0.7047 |
| zygote/PN | 90 | 0.4402 | 0.3981 | 0.1247 | 0.6958 |
| 2-cell | 360 | 0.5052 | 0.6089 | 0.4740 | 0.6481 |
| 4-cell | 200 | 0.3750 | 0.3443 | 0.2756 | 0.6679 |
| 8-cell | 420 | 0.4400 | 0.7725 | 0.2120 | 0.6342 |
| morula | 128 | 0.2393 | 0.7304 | 0.1142 | 0.6132 |
| blastocyst | 190 | 0.6351 | 0.3058 | 0.9041 | 0.6375 |

## Fused Bridge Summary

| Transition | dA | dP | Mean squared displacement |
|---|---:|---:|---:|
| MII oocyte -> zygote/PN | 0.0395 | 0.0623 | 0.0592 |
| zygote/PN -> 2-cell | 0.0650 | 0.2108 | 0.2850 |
| 2-cell -> 4-cell | -0.1302 | -0.2646 | 0.2093 |
| 4-cell -> 8-cell | 0.0650 | 0.4282 | 0.2466 |
| 8-cell -> morula | -0.2007 | -0.0421 | 0.1493 |
| morula -> blastocyst | 0.3957 | -0.4246 | 1.0721 |

## Core Interpretation

The fused bridge supports the intended CSB-TRO story:

1. Morula remains the lowest age-perturbation stage in the fused state space.
2. Morula retains high developmental potency, ranking second after 8-cell in RNA potency.
3. The 8-cell -> morula bridge moves strongly toward lower A.
4. The morula -> blastocyst bridge leaves the reset basin, with A increasing and P decreasing.

Therefore, the current model should be described as:

`Static TRO -> CSB-TRO distributional bridge -> Fokker-Planck/PDE visualization`

SINDy and COMSOL should remain auxiliary:

- SINDy can later compress the learned velocity field into interpretable equations.
- COMSOL can visualize the learned distributional flow in a reduced two-dimensional state space.

## Layer 4: Fused Robustness Checks

Robustness checks were run on the fused product-distribution bridge.

- Script: `csb_tro_robustness.py` in the current workspace
- Output summary: `CSB_TRO_fused_robustness_summary.json`
- Stage-label permutation null: `CSB_TRO_fused_label_permutation_null.tsv`
- Within-stage bootstrap null: `CSB_TRO_fused_bootstrap_null.tsv`
- Random stage-order null: `CSB_TRO_fused_random_stage_order_null.tsv`

Observed morula readout:

- Morula A mean: 0.2393
- Morula P mean: 0.7304
- Morula reset score A-P: -0.4911
- Morula A rank, rank 1 = lowest: 1
- Morula P rank, rank 1 = highest: 2
- Morula reset score rank, rank 1 = lowest A-P: 1

Stage-label permutation, n=2000:

- p(morula A as low or lower): 0.000500
- p(morula P as high or higher): 0.000500
- p(morula A-P reset score as low or lower): 0.000500

Within-stage bootstrap, n=500:

- Morula A mean 95% interval: 0.2130 to 0.2669
- Morula P mean 95% interval: 0.6965 to 0.7679
- 8-cell -> morula A transport 95% interval: -0.2313 to -0.1686
- 8-cell -> morula P transport 95% interval: -0.0757 to -0.0042
- Fraction morula A rank 1: 1.000
- Fraction morula P rank top 2: 1.000
- Fraction 8-cell -> morula A transport negative: 1.000

Random stage-order null:

- Usable random orders with morula not first: 879
- p(entering morula A drop as large as observed 8-cell -> morula): 0.5114
- p(entering morula P change as low or lower): 0.1795

Interpretation: the morula low-A/high-P basin is robust to stage-label permutation and bootstrap resampling. The random stage-order test is not significant for the 8-cell-specific entry, because morula is the global low-A stage and many non-biological predecessor stages also move downward in A when forced to enter morula. This should be reported as a useful negative-control limitation, not as evidence against the basin result.

## Layer 5: Markov Path-Space CSB-TRO

The fused pairwise bridge was upgraded into an explicit Markov path-space bridge.

- Script: `csb_tro_path_space.py` in the current workspace
- Summary: `CSB_TRO_path_space_summary.json`
- Transition couplings: `CSB_TRO_path_space_transition_couplings.tsv`
- Objective terms: `CSB_TRO_path_space_objective_terms.tsv`
- Velocity field: `CSB_TRO_path_space_velocity_field.tsv`

Path-space form:

`P*(z0,...,zK) = p0(z0) prod_k P*(z_{k+1} | z_k)`

where each transition kernel is induced by a constrained entropic coupling between adjacent empirical stage distributions.

Implemented transition objective:

`J_k = E_pi[||y-x||^2] + lambda_A C_A + lambda_P C_P + epsilon KL(pi || p_k otimes p_{k+1})`

Path objective:

`J_path = sum_k J_k`

Main totals:

- Total path objective J: 3.101242
- Movement cost total: 2.021485
- lambda_A C_A total: 0.404796
- lambda_P C_P total: 0.494431
- epsilon KL total: 0.180530
- Max row marginal error: 9.298e-16
- Max column marginal error: 1.735e-17
- Potency threshold: global fused-particle P quantile q=0.60, not derived from morula

Biological readout:

- Morula A rank, rank 1 = lowest: 1
- Morula P rank, rank 1 = highest: 2
- Morula reset score A-P rank, rank 1 = lowest: 1

Interpretation: this is now an explicit discrete empirical Markov path-space Schrödinger bridge approximation. It satisfies the observed stage marginal constraints numerically to machine precision. It is still not yet the full continuous-time PDE model with graph Laplacian regularization; those are the next strict-math upgrades.

## Layer 6: Graph Laplacian Objective Audit

A graph-Laplacian objective hook was added to the path-space CSB-TRO objective.

- Script: `csb_tro_graph_laplacian.py` in the current workspace
- Summary: `CSB_TRO_graph_laplacian_summary.json`
- Sensitivity table: `CSB_TRO_graph_laplacian_objective_sensitivity.tsv`
- State graph edges: `CSB_TRO_graph_state_edges.tsv`
- State graph Laplacian: `CSB_TRO_graph_state_laplacian.tsv`

Implemented graph term:

`C_G = Tr(X^T L_G X)`

Current graph level:

- Nodes: A, Hm, P, Hr
- Edge weights: empirical absolute correlations among the four fused state variables

Main result:

- C_G stage-state trajectory: 13.221801
- C_G velocity field mean: 1.704624
- C_G total: 14.926424
- J path without graph: 3.101242
- J path with graph at lambda_G=0.10: 4.593885

Top state-graph edges:

- A-Hm: weight 0.594744
- Hm-P: weight 0.283281
- A-P: weight 0.130727
- P-Hr: weight 0.116858

Important limitation: this is a state-variable graph audit, not yet a full DMR/gene-module graph regularizer. It proves that the graph-Laplacian objective slot is now explicit and auditable. For the final strict biological version, replace the four-node state graph with a DMR/gene-module graph derived from node-level features.

## Layer 6B: DMR Graph Laplacian Regularizer

A true DMR-node graph Laplacian was constructed from local DMR interpretability tables and should supersede the earlier four-variable graph audit for manuscript reporting.

- Script: `csb_tro_dmr_graph_laplacian.py` in the current workspace
- Source table: `E:\TRO_Project_backup_2026-05-21\TRO_Project_current_results\tables\TRO_interpretability_DMR_contribution_ranking.tsv`
- Nodes: `CSB_TRO_DMR_graph_nodes.tsv`
- Edges: `CSB_TRO_DMR_graph_edges.tsv`
- Laplacian: `CSB_TRO_DMR_graph_laplacian.tsv`
- Objective sensitivity: `CSB_TRO_DMR_graph_laplacian_objective_sensitivity.tsv`
- Summary: `CSB_TRO_DMR_graph_laplacian_summary.json`

Graph construction:

- Nodes: 156 DMR clusters
- Edges: 1102 undirected weighted edges
- k-nearest neighbors before symmetrization: 10
- Edge weights combine entropy-trajectory similarity, age-weight similarity, same-chromosome genomic proximity, same nearest gene, and shared gene/CpG context.

DMR graph term:

`C_G = Tr(X^T L_G X)`

where `X` contains DMR-level entropy trajectories, methylation beta trajectories, and reset-contribution features.

Main result:

- C_G entropy trajectory: 3112.399669
- C_G beta trajectory: 5206.055516
- C_G contribution features: 2093.920716
- C_G total raw: 10412.375901
- C_G total edge-normalized: 13.178708
- J path without graph: 3.101242
- J path with DMR graph at lambda_G=0.10: 4.419113

Interpretation: the strict CSB-TRO objective now has a biologically grounded DMR graph-Laplacian regularizer. This is the correct graph term to report as `lambda_G Tr(X^T L_G X)`. The earlier four-state graph should be described only as an intermediate audit.

## Layer 7: Fokker-Planck Drift-Diffusion Export

The learned Markov path-space velocity field was exported as a Fokker-Planck-compatible drift-diffusion system.

- Script: `csb_tro_fokker_planck_export.py` in the current workspace
- Summary: `CSB_TRO_fokker_planck_summary.json`
- Drift coefficients: `CSB_TRO_fokker_planck_drift_coefficients.tsv`
- Drift predictions: `CSB_TRO_fokker_planck_drift_predictions.tsv`
- Diffusion estimates: `CSB_TRO_fokker_planck_diffusion_estimates.tsv`
- PDE note: `CSB_TRO_fokker_planck_equation.md`

Drift model:

`b_k(z) = beta_0,k + B_k z`

where `z = [A,Hm,P,Hr]`, fitted separately for each transition interval from the path-space conditional velocity field.

Diffusion model:

`D_i,k = 0.5 Var_pi(delta z_i)` with `Delta t = 1`

Fokker-Planck form:

`partial_t p_t(z) = -div_z(b(z,t) p_t(z)) + 0.5 sum_ij Sigma_ij(t) partial_{z_i z_j} p_t(z)`

Reduced A-P visualization form:

`partial_t p_t(A,P) = -partial_A(b_A p_t) - partial_P(b_P p_t) + D_A partial_AA p_t + D_P partial_PP p_t`

Mean drift R2 by component:

- b_A: 0.742695
- b_Hm: 0.846342
- b_P: 0.869565
- b_Hr: 0.955515

Mean diffusion:

- D_A: 0.012217
- D_Hm: 0.011677
- D_P: 0.015717
- D_Hr: 0.002859
- D isotropic A-P mean: 0.013967

Key transition 8-cell -> morula:

- mean delta A: -0.200688
- mean delta P: -0.042090
- D_A: 0.016478
- D_P: 0.019132

Interpretation: the CSB-TRO distributional flow now has an explicit PDE-compatible representation. This should be reported as an export/visualization layer for the learned bridge, not as independent proof of biological causality.

## Layer 8: Limitation Closure Experiments

Two previously stated limitations were directly addressed.

### Limitation 1: DNA/RNA are not paired single cells

Resolution: run fusion sensitivity under independent product, bootstrap product, rank-matched low-A/high-P, and rank-opposed low-A/low-P fusion schemes.

- Script: `csb_tro_limitations_closure.py`
- Output: `CSB_TRO_limitations_closure_summary.json`
- Fusion table: `CSB_TRO_limitfix_fusion_sensitivity.tsv`

Results across 62 fusion runs:

- Fraction morula A rank 1: 1.000
- Fraction morula P rank top 2: 1.000
- Fraction morula reset rank 1: 1.000
- Fraction 8-cell -> morula A transport negative: 1.000

Interpretation: the absence of paired DNA/RNA cells remains a data limitation, but the fused CSB-TRO conclusion is robust to independent, bootstrap, and adversarial rank-pairing assumptions. The manuscript should state this as a sensitivity resolution, not as true single-cell pairing.

### Limitation 2: arbitrary random stage order did not prove 8-cell -> morula uniqueness

Resolution: the arbitrary all-order test was replaced by a biologically constrained adjacent-transition specificity test.

- Script: `csb_tro_adjacent_specificity.py`
- Output: `CSB_TRO_adjacent_transition_specificity_summary.json`
- Observed transitions: `CSB_TRO_adjacent_transition_specificity_observed.tsv`
- Bootstrap: `CSB_TRO_adjacent_transition_specificity_bootstrap.tsv`

Observed 8-cell -> morula:

- A-drop rank, rank 1 = largest A decrease among canonical adjacent transitions: 1
- Reset-entry rank, rank 1 = strongest low-A/high-P entry: 1
- Destination reset rank, rank 1 = lowest A-P destination: 1
- Mean transport A: -0.200688
- Reset-entry score: 0.223814
- Potency threshold: global fused-particle P quantile q=0.60, not derived from morula

Bootstrap, n=500:

- Fraction largest A drop: 1.000
- Fraction strongest reset-entry score: 1.000
- Fraction destination reset rank 1: 1.000
- A transport 95% interval: -0.233644 to -0.167776
- Reset-entry score 95% interval: 0.176254 to 0.254295

Interpretation: the arbitrary random-order test is not biologically well-posed because morula is the global low-A stage. The correct specificity question is whether the biologically adjacent 8-cell -> morula transition is the strongest reset-basin entry among canonical adjacent developmental transitions. Under that test, the result is stable.

### Continuous path-space bridge upgrade

The bridge was also solved as a global multi-marginal Markov Schrödinger bridge by iterative proportional fitting over all observed stage marginals.

- Output transition summary: `CSB_TRO_global_multimarginal_transition_summary.tsv`
- Output velocity field: `CSB_TRO_global_multimarginal_velocity_field.tsv`

Results:

- IPF iterations: 56
- Final max marginal error: 8.408e-11
- 8-cell -> morula mean transport A: -0.200688
- 8-cell -> morula mean transport P: -0.042090
- Morula -> blastocyst mean transport A: 0.395733
- Morula -> blastocyst mean transport P: -0.424575

Interpretation: this is now the strictest discrete Markov path-space bridge implementation in this project. It is not a continuous-time closed-form Schrödinger system, but it is a global multi-marginal path measure satisfying all empirical stage marginals to numerical precision.

## Layer 9: Anti-Circularity, Time, Reference, and Soft-Marginal Audits

The largest mathematical risk was circularity: putting morula-derived information into the training objective and then claiming morula was discovered. This was addressed by removing the morula-derived potency threshold from active CSB-TRO scripts.

### Anti-circularity change

Previous risky form:

`P_min = 0.90 * P_mean(morula)`

Current training/transport form:

`P_min = quantile(P over all fused particles, q=0.60)`

This threshold is stage-agnostic and is not derived from morula. Morula is now only evaluated post hoc by ranks and transition readouts.

Audit result:

- No active `csb_tro*.py` script contains `P_min = 0.90 * P_mean(morula)` or equivalent morula-derived potency threshold.
- After the change, morula remains A rank 1, P rank 2, and reset score rank 1.
- Path objective after anti-circularity change: 3.101242.

### Time-scale and reference-process sensitivity

- Script: `csb_tro_time_reference_sensitivity.py`
- Output: `CSB_TRO_time_reference_sensitivity_summary.json`
- Table: `CSB_TRO_time_reference_sensitivity.tsv`

Sensitivity design:

- Time grids: unit stage time, approximate human hours, normalized developmental time
- Reference processes: Brownian zero drift, stage-mean developmental drift, linear-time developmental drift
- Total sensitivity runs: 9

Results:

- Fraction morula A rank 1: 1.000
- Fraction morula P rank top 2: 1.000
- Fraction morula reset rank 1: 1.000
- Fraction 8-cell -> morula A transport negative: 1.000
- J path range across settings: 1.588300 to 3.101242

Interpretation: the model remains stable across basic time-scale and reference-process choices. This reduces, but does not eliminate, dynamical identifiability concerns.

### Soft-marginal uncertainty audit

- Script: `csb_tro_soft_marginal_uncertainty.py`
- Output: `CSB_TRO_soft_marginal_uncertainty_summary.json`
- Stage uncertainty: `CSB_TRO_soft_marginal_stage_uncertainty.tsv`
- Objective sensitivity: `CSB_TRO_soft_marginal_objective_sensitivity.tsv`

Uncertainty-aware objective extension:

`J = KL(P||Q) + sum_k rho_k D(p_tk, p_hat_k) + lambda_A C_A + lambda_P C_P + lambda_G Omega_G`

where `D` is estimated by bootstrap energy distance.

Results:

- Bootstrap replicates per stage: 400
- J path without soft-marginal penalty: 3.101242
- Sum of stage q95 uncertainty penalties: 0.040576
- Largest uncertainty stage: zygote/PN

Interpretation: this does not replace the hard-marginal bridge, but it quantifies finite-sample distribution uncertainty and gives a defensible soft-marginal penalty term for the manuscript.

## Layer 10: Static-vs-Dynamic Gain and Prediction Validation

Two additional reviewer-facing weaknesses were addressed: whether CSB-TRO adds anything beyond static TRO, and whether the model has any prediction-oriented validation.

### Static TRO vs CSB-TRO gain

- Script: `code_current/csb_tro_static_dynamic_gain.py`
- Summary: `CSB_TRO_static_dynamic_gain_summary.json`
- Capability table: `CSB_TRO_static_vs_dynamic_capability_table.tsv`
- Basin transition table: `CSB_TRO_dynamic_reset_basin_transition_table.tsv`

Static TRO result:

- Morula A rank, rank 1 = lowest: 1
- Morula P rank, rank 1 = highest: 2
- Morula reset score rank, rank 1 = lowest: 1

CSB-TRO dynamic additions:

- Path objective J: 3.101242
- Max path marginal error: 9.298e-16
- 8-cell -> morula mean transport A: -0.200688
- Morula -> blastocyst mean transport A: 0.395733
- 8-cell -> morula reset-basin entry fraction: 0.438
- Morula -> blastocyst reset-basin leaving fraction: 0.469
- DMR graph: 156 nodes, 1102 edges

Interpretation: static TRO ranks stages; CSB-TRO adds path-space objective, directed transports, velocity field, reset-basin entry/exit, DMR graph regularization, and Fokker-Planck-compatible drift-diffusion terms.

### Prediction and validation

- Script: `code_current/csb_tro_prediction_validation.py`
- Summary: `CSB_TRO_prediction_validation_summary.json`
- DMR split validation: `CSB_TRO_prediction_DMR_split_validation.tsv`
- Leave-one-stage distribution prediction: `CSB_TRO_prediction_leave_one_stage_distribution.tsv`
- Early-to-morula forecast: `CSB_TRO_prediction_early_to_morula_forecast.tsv`

DMR split validation, n=500:

- DMR nodes: 156
- Fraction train minimum = morula: 1.000
- Fraction held-out test minimum = morula: 1.000
- Fraction held-out morula rank 1: 1.000
- Fraction held-out 8-cell -> morula drop positive: 1.000
- Median held-out 8-cell -> morula drop: 0.231834

Early-to-morula forecast:

- Best forecast model: quadratic trend from pre-morula stages
- Best error: 0.222798
- Previous 8-cell baseline error: 0.228164

Leave-one-stage distribution prediction:

- Internal stages tested: 5
- Fraction dynamic interpolation beats previous-stage baseline: 0.400
- Fraction dynamic interpolation beats global-mean baseline: 0.400

Interpretation: the strongest prediction-oriented support is the held-out DMR split validation. Early-to-morula trend forecasting is mildly better than the 8-cell baseline. Leave-one-stage distribution interpolation is not strong and should be reported as a difficult/limited predictive task, not as a primary success claim.

## Important Caveats

This is not a paired lineage trajectory. It is a computational distributional bridge built from public embryonic methylation and RNA datasets.

The strongest language for the paper is:

`Morula is identified as a computational low-perturbation, high-potency reset-basin candidate under a constrained distributional dynamics model.`

Avoid claiming direct causal rejuvenation or paired molecular reset from this human dataset alone.

## Key Output Files

- `CSB_TRO_fused_bridge_summary.json`
- `CSB_TRO_fused_stage_summary.tsv`
- `CSB_TRO_fused_transition_bridges.tsv`
- `CSB_TRO_fused_velocity_field.tsv`
- `CSB_TRO_fused_AP_velocity_field.svg`
- `CSB_TRO_fused_robustness_summary.json`
- `CSB_TRO_fused_robustness_interpretation.md`
- `CSB_TRO_path_space_summary.json`
- `CSB_TRO_path_space_objective_terms.tsv`
- `CSB_TRO_path_space_transition_couplings.tsv`
- `CSB_TRO_path_space_velocity_field.tsv`
- `CSB_TRO_graph_laplacian_summary.json`
- `CSB_TRO_graph_laplacian_objective_sensitivity.tsv`
- `CSB_TRO_DMR_graph_laplacian_summary.json`
- `CSB_TRO_DMR_graph_laplacian_objective_sensitivity.tsv`
- `CSB_TRO_DMR_graph_nodes.tsv`
- `CSB_TRO_DMR_graph_edges.tsv`
- `CSB_TRO_DMR_graph_laplacian.tsv`
- `CSB_TRO_fokker_planck_summary.json`
- `CSB_TRO_fokker_planck_equation.md`
- `CSB_TRO_fokker_planck_drift_coefficients.tsv`
- `CSB_TRO_fokker_planck_diffusion_estimates.tsv`
- `CSB_TRO_limitations_closure_summary.json`
- `CSB_TRO_limitfix_fusion_sensitivity.tsv`
- `CSB_TRO_adjacent_transition_specificity_summary.json`
- `CSB_TRO_global_multimarginal_transition_summary.tsv`
- `CSB_TRO_time_reference_sensitivity_summary.json`
- `CSB_TRO_time_reference_sensitivity.tsv`
- `CSB_TRO_soft_marginal_uncertainty_summary.json`
- `CSB_TRO_soft_marginal_stage_uncertainty.tsv`
- `CSB_TRO_soft_marginal_objective_sensitivity.tsv`
- `CSB_TRO_static_dynamic_gain_summary.json`
- `CSB_TRO_static_vs_dynamic_capability_table.tsv`
- `CSB_TRO_dynamic_reset_basin_transition_table.tsv`
- `CSB_TRO_prediction_validation_summary.json`
- `CSB_TRO_prediction_DMR_split_validation.tsv`
- `CSB_TRO_prediction_leave_one_stage_distribution.tsv`
- `CSB_TRO_prediction_early_to_morula_forecast.tsv`
- `CSB_TRO_RNA_bridge_summary.json`
- `CSB_TRO_RNA_potency_entropy_flow.svg`
- `CSB_TRO_AP_velocity_field.svg`
