## Supplemental docs for "An entropy-based diagnostic framework for characterizing methylation state dynamics during preimplantation development": CSB_TRO_embryo_histone_dual_branch_summary.md

# Embryo Histone Dual-Branch Replacement

Status: `completed_partial_embryo_track_mode`

Analysis-ready embryo histone tracks: 7/9

This is the branch-resolved embryo histone-state replacement experiment. It replaces the dual-branch chromatin proxy with available real embryo histone peak tracks where possible, while preserving the missing-input boundary for morula H3K27ac/H3K4me3.

## Decision

Public-data resolution: the full strict morula-entry replacement is not closed with currently available public human processed tracks. The public data support a strong histone-state diagnostic in an 8-cell-to-ICM/blastocyst contrast, but the required human morula H3K27ac and morula H3K4me3 processed inputs are still absent/controlled-access.

Therefore the biological-control term is not yet final. The correct conclusion is `strong embryo histone-state diagnostic support, strict morula-entry replacement unresolved by public data`.

## Best Available Histone Control

- feature_set: `embryo_exit_diagnostic_combined_closure_access_specificity_positive_gated`
- status: `blastocyst_ICM_exit_diagnostic_specificity_positive_gated_not_morula_entry`
- max occupancy: 0.978 at alpha=2.50
- cosine at max: 0.933
- PC3 recovery at max: 0.790
- occupancy at alpha=1: 0.511
- sign-flip max occupancy: 0.044
- max matched-random-DMR occupancy: 1.000
- max all-random-control occupancy: 1.000

Matched-random controls are high for the best ICM/blastocyst diagnostic model, so this model must not be used as final DMR-specific u_bio. It shows that the histone-state direction is capable of reproducing the missing correction, but not that the exact target DMR modules are uniquely specified by public tracks.

## Stage-Matched Morula-Entry Result

- best stage-matched/partial morula-entry candidate: `embryo_partial_H3K4me3_8cell_access_positive_gated`; max occupancy=0.200; cosine=0.569; PC3=0.295

## Sign Controls

- correct_closure_correct_access: occupancy=0.978; cosine=0.933; PC3=0.790
- correct_closure_wrong_access: occupancy=0.978; cosine=0.933; PC3=0.790
- naive_all_positive: occupancy=0.978; cosine=0.933; PC3=0.790
- wrong_closure_correct_access: occupancy=0.000; cosine=-0.933; PC3=-0.790
- wrong_closure_wrong_access: occupancy=0.000; cosine=-0.933; PC3=-0.790
- naive_all_negative: occupancy=0.000; cosine=-0.933; PC3=-0.790

## Ablation Note

- full_histone_dual_branch: occupancy=0.978; cosine=0.933; PC3=0.790; included=M05,M01,M12,M02,M10
- closure_branch_only: occupancy=0.978; cosine=0.933; PC3=0.790; included=M05,M01,M12
- remove_access_branch: occupancy=0.978; cosine=0.933; PC3=0.790; included=M05,M01,M12
- remove_M12: occupancy=0.978; cosine=0.933; PC3=0.790; included=M05,M01,M02,M10
- remove_M02: occupancy=0.978; cosine=0.933; PC3=0.790; included=M05,M01,M12,M10

## Boundary

H3K27ac_morula and H3K4me3_morula are still missing as processed local human inputs. Therefore the current result is a real-embryo partial-track replacement, not final closure of the full biological control term.

The highest occupancy currently comes from blastocyst/ICM diagnostic tracks, not the strict 8-cell-to-morula H3K27ac-loss/H3K4me3-morula replacement. Do not promote it to final u_bio.

Sources used: GSE124718 for public human embryo 8-cell/ICM H3K27ac/H3K4me3/H3K27me3 processed peaks; GSE123023 for public human morula H3K27me3 BED from GEO RAW tar. HRA002355/PRJCA009410 is the stage-matched human embryo source for morula H3K27ac/H3K4me3 context, but it is controlled access. The next strict input priority remains H3K27ac_morula and H3K4me3_morula in hg19/GRCh37 BED/bigWig form.
