## Supplemental docs for "An entropy-based diagnostic framework for characterizing methylation state dynamics during preimplantation development": CSB_TRO_external_biocontrol_interpretation.md

# External biocontrol data integration

This stage prepares the coordinate and input layer for real external biological controls.

Declared genome build: hg19/GRCh37

Genome build audit:
- hg19_coordinate_bounds: pass; all_chr_known=True; all_DMRs_within_chr_lengths=True
- hg38_coordinate_bounds: fail; all_chr_known=True; all_DMRs_within_chr_lengths=False
- declared_genome_build: declared; hg19/GRCh37
- chr_naming_style: pass; chr_prefixed
- build_discrimination: hg19_supported_by_bounds; At least one DMR chromosome max coordinate exceeds hg38 but all DMRs fit hg19. Treat coordinates as hg19/GRCh37 unless source metadata proves otherwise.

External input status:
- genome_build: present; E:\CSB_TRO_operator_time_DMR_dynamics_2026-05-25\external\genome_build.txt
- gencode_gtf: present; E:\CSB_TRO_operator_time_DMR_dynamics_2026-05-25\external\annotations\gencode.v19.annotation.gtf.gz
- gene_tss: present; E:\CSB_TRO_operator_time_DMR_dynamics_2026-05-25\external\annotations\gene_tss.tsv
- cpg_islands: missing; E:\CSB_TRO_operator_time_DMR_dynamics_2026-05-25\external\annotations\cpg_islands.bed
- repeatmasker: missing; E:\CSB_TRO_operator_time_DMR_dynamics_2026-05-25\external\annotations\repeatmasker.bed
- rna_gene_matrix: missing; E:\CSB_TRO_operator_time_DMR_dynamics_2026-05-25\external\rna\gene_stage_matrix.tsv
- jaspar_meme: missing; E:\CSB_TRO_operator_time_DMR_dynamics_2026-05-25\external\motif\jaspar_vertebrates.meme
- motif_enrichment: missing; E:\CSB_TRO_operator_time_DMR_dynamics_2026-05-25\external\motif\module_motif_enrichment.tsv
- atac_features: missing; E:\CSB_TRO_operator_time_DMR_dynamics_2026-05-25\external\atac\module_ATAC_features.tsv
- histone_features: missing; E:\CSB_TRO_operator_time_DMR_dynamics_2026-05-25\external\histone\module_histone_features.tsv

Current nearest-gene/TSS links generated: 156

Interpretation boundary: Coordinate bounds support hg19/GRCh37 over hg38/GRCh38 because at least one DMR chromosome maximum exceeds hg38 chromosome length while all DMRs fit hg19. Use hg19/GRCh37 external annotations unless source metadata proves otherwise.
