## Supplemental docs for "An entropy-based diagnostic framework for characterizing methylation state dynamics during preimplantation development": CSB_TRO_figure_legends_and_results_draft.md

# CSB-TRO Figure Legends and Results Draft

Date: 2026-05-24

## Figure Legends

### Figure 1. Static TRO scoring identifies morula as a post hoc low-perturbation, high-potency stage

Static TRO was first evaluated as a stage-level score over the fused embryonic state space, using age-associated epigenetic perturbation (`A`) from GSE81233-derived methylation features and developmental potency (`P`) from GSE36552 single-cell RNA profiles. Bars show the static reset score, defined as `P - A`, for each developmental stage; overlaid lines show the corresponding stage means for `A` and `P`. Under this static readout, morula had the lowest `A` among stages (`A` rank 1), retained high developmental potency (`P` rank 2), and had the strongest reset score (`P - A` rank 1). This analysis identifies morula as a post hoc low-perturbation, high-potency candidate stage, but does not define a path, transition direction, velocity field, or reset-basin entry/exit behavior.

### Figure 2. CSB-TRO velocity field in age-perturbation and potency state space

The constrained Schrodinger bridge transgenerational reset operator (CSB-TRO) was implemented as a discrete empirical Markov path-space approximation over the fused state vector `z = [A, Hm, P, Hr]`, where `Hm` is methylation entropy and `Hr` is RNA expression entropy. The plot shows fused particles projected onto the `A-P` plane, with arrows indicating learned CSB-TRO transport velocities from the global multi-marginal Markov bridge. Stage centroids are overlaid and connected in developmental order. The dashed region denotes the reset basin, defined stage-agnostically as low `A` and high `P` (`A <=` global fused-particle 25th percentile; `P >=` global fused-particle 60th percentile). The 8-cell to morula transition moves toward lower age-associated perturbation while retaining high potency, whereas the morula to blastocyst transition exits this low-`A`/high-`P` region. The potency threshold was computed over all fused particles and was not derived from morula.

### Figure 3. Directional CSB-TRO transport into and out of morula

Mean CSB-TRO transports were computed from the global multi-marginal Markov path-space bridge for the two key adjacent transitions surrounding morula. The 8-cell to morula transition showed a strong decrease in age-associated epigenetic perturbation (`mean transport A = -0.200688`) with a small decrease in potency (`mean transport P = -0.042090`). In contrast, the morula to blastocyst transition showed a marked increase in age-associated perturbation (`mean transport A = +0.395733`) and a decrease in potency (`mean transport P = -0.424575`). These opposing transport directions support an interpretation in which morula is approached as a low-perturbation, high-potency basin and then left during progression to blastocyst.

### Figure 4. Reset-basin entry and exit are dynamic readouts added by CSB-TRO

Reset-basin entry and exit were quantified from source particles under the learned CSB-TRO transition maps. The reset basin was defined independently of stage labels as `B_reset = {z: A <= q25(A), P >= q60(P)}` over all fused particles. The 8-cell to morula transition had the largest reset-basin entry fraction among canonical adjacent transitions (`0.438`), while the morula to blastocyst transition showed a reset-basin leaving fraction of `0.469`. These quantities cannot be obtained from the original static TRO score alone; they require a distributional transport model linking adjacent embryonic stages.

### Figure 5. Held-out DMR split validation supports the morula low-perturbation signal

DMR split validation was used to test whether the morula signal was driven by a small subset of age-associated DMRs. Across 500 random splits of 156 DMR nodes, morula was the minimum-perturbation stage in both training and held-out DMR subsets in all splits (`train minimum morula fraction = 1.000`; `held-out test minimum morula fraction = 1.000`). Morula retained rank 1 in held-out DMRs in all splits, and the held-out 8-cell to morula perturbation drop was positive in all splits, with a median held-out drop of `0.231834`. These results support that the morula low-perturbation readout is distributed across DMR features rather than being explained by a small, unstable DMR subset.

## Results Draft

### A static TRO score identifies morula, but only as a stage-level endpoint

We first evaluated the original TRO concept as a static stage-level score in a fused human early embryo state space. The model combined GSE81233 methylation-derived age-associated epigenetic perturbation (`A`) and methylation entropy (`Hm`) with GSE36552 single-cell RNA-derived developmental potency (`P`) and RNA expression entropy (`Hr`). Because methylation and RNA measurements were not paired in the same cells, we constructed stage-wise product distributions, `p_k(A,Hm,P,Hr) ~= p_k^DNA(A,Hm) x p_k^RNA(P,Hr)`, yielding 1,496 fused particles across seven developmental stages.

In this fused state space, morula had the lowest mean age-associated perturbation (`A = 0.2393`, rank 1) and high developmental potency (`P = 0.7304`, rank 2, second only to 8-cell). A static reset score based on low `A` and high `P` ranked morula first among stages. This result supports morula as a candidate reset-like stage, but the static formulation remains limited: it ranks stages but does not specify a developmental path, transition direction, velocity field, or basin entry/exit process.

### CSB-TRO upgrades the static score to a constrained distributional dynamics model

To move beyond static scoring, we implemented CSB-TRO as a discrete empirical Markov path-space approximation of a constrained Schrodinger bridge over observed developmental distributions. Each stage was treated as an empirical marginal distribution in the four-dimensional state vector `z = [A, Hm, P, Hr]`. Adjacent transition kernels were inferred under entropic transport with biological penalty terms, and the resulting Markov path measure satisfied the observed stage marginals to numerical precision.

The global path objective was `J_path_total = 3.101242`, with movement cost `2.021485`, age-perturbation penalty contribution `0.404796`, potency penalty contribution `0.494431`, and entropic KL contribution `0.180530`. The maximum path marginal error was approximately machine precision in the pairwise path-space implementation (`9.298e-16`) and remained very small in the global multi-marginal implementation (`8.408e-11`). The active potency constraint used a stage-agnostic global fused-particle threshold (`P_min = q60(P) = 0.707284`) and was not derived from morula; morula was evaluated only post hoc.

This dynamic formulation preserved the static morula readout while adding quantities unavailable to static TRO: a path-space objective, directed transition transports, a velocity field, reset-basin entry and exit fractions, DMR graph-Laplacian regularization, and a Fokker-Planck-compatible drift-diffusion export.

### The learned transport identifies entry into and exit from a morula reset basin

The CSB-TRO transport field revealed a directed transition into a low-`A`/high-`P` region at morula followed by exit during blastocyst progression. In the global multi-marginal bridge, the 8-cell to morula transition had a mean age-perturbation transport of `-0.200688`, indicating a substantial decrease in `A`. The same transition had a smaller mean potency transport of `-0.042090`, consistent with morula retaining high potency while reducing age-associated perturbation. By contrast, the morula to blastocyst transition showed a strong increase in `A` (`+0.395733`) and a decrease in `P` (`-0.424575`).

We next quantified these dynamics using a reset basin defined independently of morula as `B_reset = {z: A <= q25(A), P >= q60(P)}` over all fused particles. The 8-cell to morula transition had a reset-basin entry fraction of `0.438`, whereas the morula to blastocyst transition had a basin leaving fraction of `0.469`. Among canonical adjacent transitions, 8-cell to morula ranked first for largest `A` decrease, strongest reset-entry score, and lowest destination reset rank. In 500 bootstrap replicates, this transition remained the largest `A` drop and strongest reset-entry transition in every replicate, with a 95% interval for `A` transport from `-0.233644` to `-0.167776`.

Together, these results support the interpretation that morula is not merely a favorable static stage but a computational reset-basin candidate approached by directed developmental transport and then left as the embryo progresses to blastocyst.

### Robustness analyses reduce circularity and parameter-choice concerns

Several audits were performed to address circularity and sensitivity risks. The principal anti-circularity change was removal of the earlier morula-derived potency threshold. The current CSB-TRO objective uses the global fused-particle 60th percentile of `P`, not `P_mean(morula)`, and morula is assessed only after model fitting. Under this corrected formulation, morula remained `A` rank 1, `P` rank 2, and reset-score rank 1.

The morula readout was stable under label permutation and bootstrap analyses. In 2,000 stage-label permutations, the empirical probabilities of observing morula-like low `A`, high `P`, or reset score by chance were all approximately `0.000500`. In 500 within-stage bootstraps, morula remained `A` rank 1 in all replicates, remained within the top two stages for `P` in all replicates, and the 8-cell to morula `A` transport was negative in all replicates. Across nine time-scale and reference-process settings, morula remained `A` rank 1, `P` top 2, and reset-score rank 1 in every setting, and the 8-cell to morula `A` transport remained negative in every setting.

### DMR graph regularization and split validation support a distributed methylation signal

To make the biological regularization explicit, we constructed a DMR-node graph over 156 DMR clusters with 1,102 weighted undirected edges. Edge weights incorporated entropy-trajectory similarity, age-weight similarity, same-chromosome genomic proximity, shared nearest gene, and shared gene/CpG context. This graph supplied the manuscript-facing graph-Laplacian term, `lambda_G Tr(X^T L_G X)`, replacing an earlier four-state-variable graph audit.

The DMR split validation provided the strongest prediction-oriented support for the methylation component of CSB-TRO. Across 500 DMR splits, morula was the minimum-perturbation stage in all training splits and all held-out test splits. Morula retained held-out rank 1 in all splits, and the held-out 8-cell to morula perturbation drop was positive in all splits, with a median drop of `0.231834`. These findings argue against the morula low-perturbation signal being driven by a small number of unstable DMRs.

### Bayesian bootstrap validation supports posterior stability of the reset-basin call

We next added a posterior-style stability analysis using Bayesian bootstrap resampling. In each of 2,000 iterations, fused stage particles were assigned Dirichlet bootstrap weights and age-DMR features were independently assigned Dirichlet bootstrap weights, without introducing a morula-derived training constraint. The reset candidate was then defined post hoc as the stage with minimum particle-level `A` among stages satisfying the stage-agnostic potency threshold `P >= q60(P)`.

Under this analysis, the posterior-style probability that the particle-level reset candidate was morula was `0.904`. Morula remained `A` rank 1, `P` top 2, and reset-score rank 1 in all bootstrap iterations, and the 8-cell to morula `A` drop was positive in all iterations. At the DMR-feature level, the posterior-style probability that morula was the minimum stage was `1.000`, with a positive 8-cell to morula DMR drop in all iterations. The joint probability that both the particle-level reset candidate and DMR-level minimum were morula was `0.904`.

This analysis strengthens the local generalization claim: the reset-basin call is stable under particle and DMR uncertainty. It should not be presented as broad supervised forecasting; instead, it supports the interpretation of CSB-TRO as an interpretable constrained distributional inference framework with strong local validation.

### Prediction-oriented tests show strong DMR validation but limited distribution forecasting

Prediction-oriented analyses gave a mixed but informative picture. Early-to-morula trend forecasting modestly improved over a previous 8-cell baseline: the best pre-morula quadratic trend forecast error was `0.222798`, compared with a previous-stage 8-cell baseline error of `0.228164`. However, leave-one-stage distribution prediction was more difficult. Dynamic interpolation beat the previous-stage baseline in only `0.400` of internal stage tests and beat the global mean baseline in only `0.400` of tests.

We then implemented a full-stage distribution prediction task rather than only stage-mean forecasting. For each internal held-out stage, an entropic OT/CSB geodesic midpoint was inferred from the neighboring observed stage distributions and used to generate a predicted empirical distribution in `z = [A,Hm,P,Hr]`. Across five held-out internal stages and 80 repeats per stage, the CSB/OT geodesic prediction improved over naive random midpoint interpolation for all stage-level median energy comparisons (`1.000`). However, it remained only partially competitive with simple baselines, beating the previous-stage baseline in `0.400` of stages, the next-stage baseline in `0.600` of stages, and the global-mean baseline in `0.600` of stages.

Therefore, the prediction evidence should be stated narrowly. CSB-TRO now implements full held-out stage distribution prediction and improves over naive midpoint interpolation, but broad distributional forecasting remains limited. The strongest prediction-oriented evidence remains Bayesian/bootstrap posterior stability and held-out DMR split validation, not high-accuracy full-stage forecasting.

### Summary interpretation

Overall, CSB-TRO extends static TRO from a stage-ranking score into a constrained distributional dynamics model. The most defensible conclusion is that morula is identified as a computational low-perturbation, high-potency reset-basin candidate under a constrained distributional dynamics model. The model supports this conclusion through a stage-agnostic path-space bridge, directed 8-cell to morula transport toward lower `A`, reset-basin entry/exit quantification, DMR graph regularization, Bayesian/bootstrap posterior validation, and held-out DMR split validation. CSB-TRO should be interpreted as an interpretable constrained distributional inference framework rather than a high-accuracy supervised forecasting model. The current implementation should be described as a discrete empirical Markov path-space approximation of a constrained Schrodinger bridge, not as a closed-form solution of the full continuous-time Schrodinger bridge PDE/SDE system.
