## Supplemental docs for "An entropy-based diagnostic framework for characterizing methylation state dynamics during preimplantation development": CSB_TRO_final_claim_boundary_summary.md

# Final Claim Boundary Summary

Status: `public_data_bounded_manuscript_ready`

## Frozen Model

```text
dz/dtau = f_meth(z,tau) + beta_c u_closure b_closure + beta_a u_access b_access
closure branch = M05/M01/M12
access branch = M02/M10
```

## Fixed Claims

- Methylation-only q90 occupancy: 0.044.
- Measured correction reaches observed q90 occupancy at alpha=0.50; occupancy@alpha1=1.000.
- Dual-branch proxy max occupancy: 1.000; cosine=0.994; PC3 recovery=0.879.
- Public embryo histone diagnostic max occupancy: 0.978; cosine=0.933; PC3 recovery=0.790.
- Strict morula-entry partial histone max occupancy: 0.200; this is data-access limited.

## Claim Boundary

We identify a robust diagnostic dual-branch control architecture with histone-supported biological plausibility.

We do not claim final identification of u_bio because human H3K27ac_morula and H3K4me3_morula processed tracks are not publicly available locally and are tied to controlled-access HRA002355/PRJCA009410.
