## Supplemental docs for "An entropy-based diagnostic framework for characterizing methylation state dynamics during preimplantation development": CSB_TRO_GSE36552_gene_stage_matrix.md

# GSE36552 gene-stage matrix

Source: Yan et al. 2013 supplementary table 1 known RefSeq gene RPKM matrix.

Genes parsed: 20214
Sample columns matched to local stage metadata: 145

Outputs:
- E:\CSB_TRO_operator_time_DMR_dynamics_2026-05-25\external\rna\gene_stage_matrix.tsv
- E:\CSB_TRO_operator_time_DMR_dynamics_2026-05-25\results\CSB_TRO_residual_module_gene_links_symbol.tsv
- E:\CSB_TRO_operator_time_DMR_dynamics_2026-05-25\results\CSB_TRO_module_linked_RNA_gene_match_summary.tsv
