## Supplemental docs for "An entropy-based diagnostic framework for characterizing methylation state dynamics during preimplantation development": CSB_TRO_hESC_histone_branch_identity_proxy_summary.md

# hESC Histone-State Branch Identity Proxy

Status: `completed_human_hESC_proxy_not_embryo_morula`

These tracks are public human naive/primed hESC histone peak BEDs from GSE52617. They are useful for testing biological branch identity, but they are not embryo morula histone tracks and must not be called final u_bio.

## Top Dynamics

- hESC_proxy_dual_H3K27acloss_H3K4me3: max_occ=0.511; occ@1=0.222; cosine@1=0.806; PC3@1=0.166; signflip=0.044
- hESC_proxy_closure_H3K27ac_low_naive: max_occ=0.089; occ@1=0.067; cosine@1=0.307; PC3@1=0.087; signflip=0.044
- hESC_proxy_closure_H3K27ac_loss_primed_to_naive: max_occ=0.089; occ@1=0.067; cosine@1=0.307; PC3@1=0.087; signflip=0.044
- hESC_proxy_access_H3K4me3_naive: max_occ=0.044; occ@1=0.044; cosine@1=0.569; PC3@1=0.009; signflip=0.044
- hESC_proxy_dual_H3K27aclow_H3K27me3_H3K4me3: max_occ=0.044; occ@1=0.044; cosine@1=0.155; PC3@1=0.123; signflip=0.044
- hESC_proxy_access_H3K27ac_naive: max_occ=0.044; occ@1=0.044; cosine@1=-0.106; PC3@1=0.015; signflip=0.044
- hESC_proxy_closure_H3K27me3_high_naive: max_occ=0.044; occ@1=0.022; cosine@1=-0.307; PC3@1=-0.087; signflip=0.089

## Strongest Module Histone Overlaps

- M02 hESC_naive_H3K27me3: target=0.333, bg=0.002, OR=214.95, q=2.39e-12
- M02 hESC_primed_H3K27me3: target=0.267, bg=0.002, OR=158.54, q=8.96e-10
- M10 hESC_primed_H3K4me3: target=0.333, bg=0.000, OR=76.20, q=0.152
- M02 hESC_primed_H3K4me3: target=0.300, bg=0.006, OR=61.52, q=1.47e-09
- M12 hESC_naive_H3K27me3: target=0.000, bg=0.000, OR=18.78, q=1
- M12 hESC_primed_H3K27ac: target=0.000, bg=0.000, OR=18.78, q=1
- M12 hESC_naive_H3K4me3: target=0.000, bg=0.000, OR=18.78, q=1
- M12 hESC_primed_H3K4me3: target=0.000, bg=0.000, OR=18.78, q=1
- M12 hESC_naive_H3K27ac: target=0.000, bg=0.000, OR=18.78, q=1
- M12 hESC_primed_H3K27me3: target=0.000, bg=0.000, OR=18.78, q=1
- M10 hESC_naive_H3K27me3: target=0.000, bg=0.000, OR=18.14, q=1
- M10 hESC_naive_H3K4me3: target=0.000, bg=0.000, OR=18.14, q=1

## Boundary

This is a biological-state proxy layer. Final biologically interpretable control still requires embryo-stage H3K27ac/H3K4me3/H3K27me3 or equivalent chromatin-state tracks.
