## Supplemental docs for "An entropy-based diagnostic framework for characterizing methylation state dynamics during preimplantation development": CSB_TRO_histone_data_access_and_overlap_plan.md

# Histone Data Access Audit and Overlap Plan

Goal: test whether M02 KLF4/KLF5 residual DMR targets overlap human early-embryo H3K27ac/H3K4me3 peaks at 8-cell, morula, and blastocyst stages.

## Access Boundary

Source/accession note: the human early-embryo histone acetylation study reports H3K27ac/H3K4me3 dynamics and associated accessions `PRJCA009410`, `HRA002355`, and `CRA006815`. Local overlap requires processed BED peaks or raw-data-derived peaks in the same genome build.

This step requires processed peak BED files or controlled-access raw data downloaded outside the script. The audit does not treat missing files as negative biological evidence.

## Local Track Status

- Expected histone tracks: 6
- Available local tracks: 0
- Any overlap runnable: False
- Full 8-cell to morula transition runnable: False

Expected local files are listed in:

`E:\CSB_TRO_operator_time_DMR_dynamics_2026-05-25\external\histone\histone_peak_manifest.tsv`

## Required Next Input

Place BED3+ peak files at the manifest paths, preferably in hg19/GRCh37 coordinates:

- `E:\CSB_TRO_operator_time_DMR_dynamics_2026-05-25\external\histone\H3K27ac_8cell.hg19.bed`
- `E:\CSB_TRO_operator_time_DMR_dynamics_2026-05-25\external\histone\H3K27ac_morula.hg19.bed`
- `E:\CSB_TRO_operator_time_DMR_dynamics_2026-05-25\external\histone\H3K27ac_blastocyst.hg19.bed`
- `E:\CSB_TRO_operator_time_DMR_dynamics_2026-05-25\external\histone\H3K4me3_8cell.hg19.bed`
- `E:\CSB_TRO_operator_time_DMR_dynamics_2026-05-25\external\histone\H3K4me3_morula.hg19.bed`
- `E:\CSB_TRO_operator_time_DMR_dynamics_2026-05-25\external\histone\H3K4me3_blastocyst.hg19.bed`

After files are present, run:

```text
python code\run_histone_overlap_for_m02_klf.py
```

Missing histone files should be reported as `not_run_missing_histone_input`, not as absence of H3K27ac/H3K4me3 support.
