## Supplemental docs for "An entropy-based diagnostic framework for characterizing methylation state dynamics during preimplantation development": CSB_TRO_histone_data_acquisition_plan.md

# Histone Data Acquisition Plan

Goal: obtain processed or signal-level H3K27ac/H3K4me3/H3K27me3 inputs for M05/M01/M12/M02/M10 residual module control analysis.

## Current Local Status

- Expected histone tracks: 9
- Tracks with BED/broadPeak: 0
- Tracks with bigWig signal: 0
- Ready tracks: 0

## Accepted Inputs

Preferred: processed peak files in BED3+, narrowPeak, or broadPeak form.

Alternate: bigWig signal tracks. These can support mean/max signal over DMRs and stage-delta module scores, but need a bigWig summarization tool such as `bigWigAverageOverBed` or `pyBigWig`.

Raw fallback: FASTQ/BAM can be used only after reproducible preprocessing. H3K27ac/H3K4me3 and H3K27me3 should not be peak-called with identical assumptions because H3K27me3 is broad-domain-like.

## Source Status

- Cell Discovery 2023 human early embryo H3K27ac/H3K4me3: highest_priority_human_stage_matched_histone_source; boundary: human raw data appears controlled-access; no local processed BED/bigWig found.
- DevOmics: possible processed/promoter-level epigenomic signal entry point; boundary: need concrete downloadable track/table URLs before use.
- CRA006815: related early embryo histone acetylation raw-data source; boundary: not a direct local human processed BED input; treat as raw-data fallback only.

## Decision Rule

Do not interpret missing local files as negative histone evidence. Once any BED/bigWig is available at the manifest paths, rerun:

```text
python code\run_residual_module_histone_state_control.py
```
