## Supplemental docs for "An entropy-based diagnostic framework for characterizing methylation state dynamics during preimplantation development": CSB_TRO_histone_source_audit_2026-05-26.md

# Histone Source Audit

Date: 2026-05-26

## Question

Can we directly populate the local manifest with processed human early-embryo H3K27ac/H3K4me3 peak BED files for 8-cell, morula, and blastocyst?

## Checked Sources

- Article: Dynamics of histone acetylation during human early embryogenesis, Cell Discovery, 2023.
- Reported accessions: `PRJCA009410`, `HRA002355`, `CRA006815`.
- Downloaded supplementary table:
  `external/histone/source_audit/41421_2022_514_MOESM2_TableS5.xlsx`
- Downloaded supplementary PDF:
  `external/histone/source_audit/41421_2022_514_MOESM1_supplement.pdf`

## Current Finding

`41421_2022_514_MOESM2_TableS5.xlsx` is not a peak-coordinate file. It contains GO enrichment sheets for stage / histone-acetylation categories such as `8cell K27K18K9`, `morula K27 only`, and `blastocyst K27K9`, with columns including `ID`, `Description`, `pvalue`, `p.adjust`, and `Count`.

Therefore, Table S5 cannot be converted into the required BED3+ histone tracks.

The article reports that sequencing data are deposited under `HRA002355` and `CRA006815` in `PRJCA009410`. This does not currently provide local processed peak BED files in the workspace. Missing local peak files remain an input/access boundary, not negative H3K27ac/H3K4me3 biological evidence.

## Decision Boundary

Do not run histone overlap as a biological test until one of these is true:

- processed hg19/GRCh37 peak BED files are obtained and placed at the manifest paths; or
- raw FASTQ/BAM access is available and a reproducible peak-calling workflow is added.

If neither is available, proceed to strengthen the M02-KLF4/KLF5 evidence with motif and TF-expression controls rather than repeatedly querying the same missing histone input.

## Immediate Next Analysis Route

Prioritize motif-control hardening:

- independent motif scanner cross-check if available, e.g. FIMO/HOMER;
- matched-background stability checks;
- random-module controls;
- shuffled-TF controls;
- sign-flip controls;
- confirm whether KLF4/KLF5 remains among the retained candidates.
