## Supplemental docs for "An entropy-based diagnostic framework for characterizing methylation state dynamics during preimplantation development": CSB_TRO_histone_state_control_summary.md

# Residual Module Histone-State Control

Status: `completed_partial_track_mode`

Goal: test whether H3K27ac/H3K4me3/H3K27me3 stage-specific marks explain M05/M01/M12/M02/M10 residual module control.

Analysis-ready histone tracks: 7/9

Present local histone files: 7/9

## Strongest Histone Overlaps

- M10 H3K4me3_blastocyst: target=0.333, background=0.000, OR=76.200, q=0.159
- M10 H3K27me3_blastocyst: target=0.667, background=0.032, OR=41.000, q=0.0479
- M02 H3K4me3_8cell: target=0.433, background=0.019, OR=38.170, q=2.52e-11
- M02 H3K4me3_blastocyst: target=0.200, background=0.008, OR=30.173, q=2.43e-05
- M12 H3K27me3_morula: target=0.000, background=0.000, OR=18.778, q=1
- M12 H3K27me3_8cell: target=0.000, background=0.000, OR=18.778, q=1
- M10 H3K4me3_8cell: target=0.000, background=0.000, OR=18.143, q=1
- M10 H3K27me3_morula: target=0.000, background=0.000, OR=18.143, q=1
- M10 H3K27me3_8cell: target=0.000, background=0.000, OR=18.143, q=1
- M05 H3K27me3_blastocyst: target=0.500, background=0.079, OR=11.095, q=0.0593

## Control Dynamics

- motif_TF_x_H3K27ac_8cell_histone_only: occupancy=0.222, cosine=0.452, PC3_recovery=0.094
- motif_TF_x_H3K27ac_blastocyst_histone_only: occupancy=0.222, cosine=0.452, PC3_recovery=0.094
- motif_TF_x_H3K27me3_blastocyst_histone_only: occupancy=0.222, cosine=0.452, PC3_recovery=0.094
- motif_TF_x_H3K27me3_morula_histone_only: occupancy=0.222, cosine=0.452, PC3_recovery=0.094
- motif_TF_x_H3K4me3_8cell_histone_only: occupancy=0.222, cosine=0.452, PC3_recovery=0.094
- motif_TF_x_H3K4me3_blastocyst_histone_only: occupancy=0.222, cosine=0.452, PC3_recovery=0.094
- H3K27me3_blastocyst_histone_only: occupancy=0.044, cosine=-0.177, PC3_recovery=-0.028
- H3K27me3_morula_to_blastocyst_histone_delta: occupancy=0.044, cosine=0.070, PC3_recovery=-0.006

Missing tracks:

- H3K27ac_morula
- H3K4me3_morula

Next input priority: processed human early-embryo H3K27ac/H3K4me3/H3K27me3 peak BED or signal tracks for 8-cell, morula, and blastocyst in hg19/GRCh37.
