## Supplemental docs for "An entropy-based diagnostic framework for characterizing methylation state dynamics during preimplantation development": CSB_TRO_inverse_ATAC_module_decomposition_summary.md

# Inverse ATAC Module Decomposition

Status: `completed`

Question: whether the inverse ATAC proxy explains the full M05/M01/M12/M02 correction program, or only a local chromatin branch.

## Module-Level Inference

### ATAC_8cell_2pn_chromatin_only_inverse
- M05: inverse_u=0.948, PC3=-0.489, norm=0.582, overlap=0.000, bg=0.024, OR=2.714
- M10: inverse_u=-1.361, PC3=0.197, norm=0.285, overlap=0.333, bg=0.016, OR=25.000
- M02: inverse_u=-0.716, PC3=0.133, norm=0.197, overlap=0.233, bg=0.011, OR=26.532
- M12: inverse_u=0.787, PC3=-0.162, norm=0.166, overlap=0.000, bg=0.000, OR=18.778
- M01: inverse_u=0.342, PC3=-0.096, norm=0.145, overlap=0.095, bg=0.029, OR=4.069

### ATAC_8cell_3pn_chromatin_only_inverse
- M05: inverse_u=0.867, PC3=-0.447, norm=0.532, overlap=0.000, bg=0.032, OR=2.094
- M02: inverse_u=-1.061, PC3=0.197, norm=0.292, overlap=0.333, bg=0.011, OR=42.580
- M01: inverse_u=0.620, PC3=-0.173, norm=0.263, overlap=0.048, bg=0.034, OR=2.013
- M10: inverse_u=-1.121, PC3=0.163, norm=0.235, overlap=0.333, bg=0.000, OR=76.200
- M12: inverse_u=0.694, PC3=-0.143, norm=0.147, overlap=0.000, bg=0.000, OR=18.778

## Dynamics Decomposition

- ATAC_8cell_3pn_chromatin_only_inverse / cumulative_M05+M01+M12: modules=M05,M01,M12; max_occ=1.000 at alpha=2.40; occ@1=0.600; cosine@1=0.945; PC3@1=0.399; random_p@1=0.000
- ATAC_8cell_2pn_chromatin_only_inverse / cumulative_M05+M01+M12: modules=M05,M01,M12; max_occ=0.978 at alpha=2.45; occ@1=0.556; cosine@1=0.929; PC3@1=0.390; random_p@1=0.000
- ATAC_8cell_3pn_chromatin_only_inverse / leave_one_out_remove_M02: modules=M05,M01,M12,M10; max_occ=0.933 at alpha=2.40; occ@1=0.467; cosine@1=0.917; PC3@1=0.314; random_p@1=0.398
- ATAC_8cell_3pn_chromatin_only_inverse / cumulative_M05+M01: modules=M05,M01; max_occ=0.911 at alpha=2.10; occ@1=0.489; cosine@1=0.909; PC3@1=0.324; random_p@1=0.000
- ATAC_8cell_3pn_chromatin_only_inverse / pair_M05+M01: modules=M05,M01; max_occ=0.911 at alpha=2.10; occ@1=0.489; cosine@1=0.909; PC3@1=0.324; random_p@1=0.000
- ATAC_8cell_2pn_chromatin_only_inverse / pair_M05+M12: modules=M05,M12; max_occ=0.911 at alpha=2.50; occ@1=0.467; cosine@1=0.896; PC3@1=0.340; random_p@1=0.198
- ATAC_8cell_3pn_chromatin_only_inverse / pair_M05+M12: modules=M05,M12; max_occ=0.867 at alpha=2.45; occ@1=0.467; cosine@1=0.894; PC3@1=0.308; random_p@1=0.398
- ATAC_8cell_2pn_chromatin_only_inverse / cumulative_M05+M01: modules=M05,M01; max_occ=0.844 at alpha=2.50; occ@1=0.467; cosine@1=0.883; PC3@1=0.305; random_p@1=0.198
- ATAC_8cell_2pn_chromatin_only_inverse / pair_M05+M01: modules=M05,M01; max_occ=0.844 at alpha=2.50; occ@1=0.467; cosine@1=0.883; PC3@1=0.305; random_p@1=0.198
- ATAC_8cell_3pn_chromatin_only_inverse / pair_M01+M12: modules=M01,M12; max_occ=0.800 at alpha=2.50; occ@1=0.311; cosine@1=0.984; PC3@1=0.165; random_p@1=0.398
- ATAC_8cell_2pn_chromatin_only_inverse / leave_one_out_remove_M02: modules=M05,M01,M12,M10; max_occ=0.644 at alpha=2.20; occ@1=0.444; cosine@1=0.855; PC3@1=0.287; random_p@1=0.398
- ATAC_8cell_2pn_chromatin_only_inverse / cumulative_M05+M01+M12+M02: modules=M05,M01,M12,M02; max_occ=0.622 at alpha=1.85; occ@1=0.467; cosine@1=0.835; PC3@1=0.320; random_p@1=0.198

## Mechanistic Boundary

The inverse ATAC proxy is strongest where residual modules are promoter-like/accessibility-linked, especially M02 and M10. It does not supply direct support for the distal/intergenic M05/M01 arm or the M12 promoter-state arm. Therefore inverse ATAC should be treated as a signed chromatin-state benchmark for an accessibility-loss branch, not as the complete biological control term.

Immediate implication: the missing biological control should be split into at least two layers: an accessibility-loss/promoter branch that inverse ATAC approximates, and a histone-state branch needed to explain M05/M01/M12.
