## Supplemental docs for "An entropy-based diagnostic framework for characterizing methylation state dynamics during preimplantation development": CSB_TRO_JASPAR_pwm_matched_background_scan.md

# JASPAR PWM matched-background scan

This scan uses per-DMR matched non-age windows from GSE81233 as background for priority residual modules.

Priority module sequences: 64
Matched background sequences: 640
Matched control sets per DMR: 10
Expressed motifs scanned: 19
Enrichment rows: 95

Boundary: this remains a Python PWM burden approximation. HOMER/FIMO with explicit GC/CpG matching is still preferred for final claims.
