## Supplemental docs for "An entropy-based diagnostic framework for characterizing methylation state dynamics during preimplantation development": CSB_TRO_JASPAR_pwm_module_scan.md

# JASPAR PWM module scan

This is a lightweight Python PWM scan against UCSC hg19 DMR sequences. It is a first-pass motif burden screen, not a replacement for HOMER/FIMO with matched GC/CpG background.

DMR sequences: 156
Expressed JASPAR motifs scanned: 848
Module motif enrichment rows: 4240
