## Supplemental docs for "An entropy-based diagnostic framework for characterizing methylation state dynamics during preimplantation development": CSB_TRO_M02_KLF4_KLF5_ATAC_gated_control_summary.md

# M02 KLF4/KLF5 ATAC-Gated Control Summary

Date: 2026-05-26

## Goal

Test whether the exploratory M02 KLF4/KLF5 motif x TF control has chromatin accessibility support.

## Inputs

Candidate regions:

```text
M02 KLF4/KLF5 residual DMR targets
n = 12
genome build = hg19/GRCh37
```

ATAC data:

```text
GSE101571 8-cell 2PN ATAC peaks
GSE101571 8-cell 3PN ATAC peaks
GSE101571 ICM 2PN ATAC peaks
GSE101571 ICM 3PN ATAC peaks
```

Boundary:

```text
GSE101571 does not provide morula ATAC in this local test.
Therefore this is 8-cell / ICM accessibility support, not direct morula chromatin validation.
```

## Overlap Result

M02 KLF4/KLF5 target DMR overlap:

```text
any 8-cell ATAC overlap = 4 / 12 = 0.333
any ICM ATAC overlap = 3 / 12 = 0.250

8-cell 2PN overlap fraction = 0.250
8-cell 3PN overlap fraction = 0.333
ICM 2PN overlap fraction = 0.250
ICM 3PN overlap fraction = 0.167
```

8-cell ATAC-supported targets:

```text
cluster_3303
cluster_5400
cluster_1851
cluster_2832
```

## Control Dynamics

ATAC-gated M02 KLF4/KLF5 control:

```text
occupancy_q90 = 0.222
direction cosine = 0.452
PC3 recovery = 0.094
DMR mean RMSE = 0.244205
DMR correlation = 0.632164
sign-flip occupancy_q90 = 0.000
```

This matches the ungated q<=0.05 motif x TF result because only M02 is nonzero, and the current augmented-dynamics script uses z-scored control values. Multiplying the only nonzero module by the ATAC overlap fraction does not change the z-scored module pattern.

## Interpretation

Safe conclusion:

```text
One third of M02 KLF4/KLF5 motif-hit residual DMRs overlap 8-cell ATAC peaks, giving partial chromatin accessibility support for the M02 KLF4/KLF5 candidate regulatory signal.
```

Do not claim:

```text
ATAC validates the full missing u_bio.
Morula chromatin accessibility supports the signal.
ATAC gating improves the dynamics beyond motif x TF alone.
```

Current status:

```text
M02 KLF4/KLF5 remains a plausible but modest candidate regulatory control.
It has:
- matched-background motif signal;
- TF expression support from GSE36552;
- partial 8-cell ATAC accessibility support;
- modest control rescue and sign-flip failure.

It lacks:
- direct morula ATAC;
- H3K27ac/H3K4me3 support;
- stronger occupancy/cosine than global RNA transition;
- HOMER/FIMO confirmation.
```

## Next Step

The next best experiment is histone-state validation:

```text
H3K27ac / H3K4me3 8-cell, morula, blastocyst peaks
overlap with M02 KLF4/KLF5 target DMRs
then build histone-gated motif x TF activity
```
