## Supplemental docs for "An entropy-based diagnostic framework for characterizing methylation state dynamics during preimplantation development": CSB_TRO_M02_KLF4_KLF5_ATAC_overlap.md

# M02 KLF4/KLF5 ATAC overlap

GSE101571 ATAC peaks were used as chromatin accessibility support. These data include 8-cell and ICM, not morula, so this is pre-morula/future-accessibility support rather than direct morula ATAC validation.

M02 KLF4/KLF5 target DMRs: 12
Any 8-cell ATAC overlap fraction: 0.333
Any ICM ATAC overlap fraction: 0.250

Track overlap fractions:
- GSE101571_8cell_2pn_ATAC: 0.250
- GSE101571_8cell_3pn_ATAC: 0.333
- GSE101571_icm_2pn_ATAC: 0.250
- GSE101571_icm_3pn_ATAC: 0.167
