## Supplemental docs for "An entropy-based diagnostic framework for characterizing methylation state dynamics during preimplantation development": CSB_TRO_M02_KLF4_KLF5_histone_overlap_summary.md

# M02 KLF4/KLF5 Histone Overlap

Status: `not_run_missing_histone_input`

No local H3K27ac/H3K4me3 peak BED files are present at the manifest paths.

Missing tracks:

- `H3K27ac_8cell`
- `H3K27ac_morula`
- `H3K27ac_blastocyst`
- `H3K4me3_8cell`
- `H3K4me3_morula`
- `H3K4me3_blastocyst`

This is an input availability boundary, not negative H3K27ac/H3K4me3 evidence.
