## Supplemental docs for "An entropy-based diagnostic framework for characterizing methylation state dynamics during preimplantation development": CSB_TRO_M02_KLF4_KLF5_target_regions.md

# M02 KLF4/KLF5 residual DMR targets

These hg19 regions are the M02 residual DMRs with matched-background JASPAR KLF4 and/or KLF5 motif hits.

Target DMRs: 12

Top targets:
- cluster_6406 chr6:40359007-40359130 TFs=KLF4,KLF5 residual_rank=69
- cluster_618 chr1:228604953-228605010 TFs=KLF4,KLF5 residual_rank=95
- cluster_2727 chr15:91447374-91447749 TFs=KLF4,KLF5 residual_rank=111
- cluster_3303 chr17:26553567-26554245 TFs=KLF4,KLF5 residual_rank=116
- cluster_3362 chr17:39685250-39685912 TFs=KLF4,KLF5 residual_rank=124
- cluster_5400 chr4:683148-683772 TFs=KLF4,KLF5 residual_rank=125
- cluster_6871 chr7:27225772-27226329 TFs=KLF4,KLF5 residual_rank=128
- cluster_1851 chr12:56131768-56132083 TFs=KLF4,KLF5 residual_rank=130
- cluster_2832 chr16:2073518-2073792 TFs=KLF4,KLF5 residual_rank=143
- cluster_3336 chr17:37764503-37764802 TFs=KLF4,KLF5 residual_rank=147
- cluster_5307 chr3:147112541-147113092 TFs=KLF4,KLF5 residual_rank=148
- cluster_67 chr1:2559320-2559371 TFs=KLF4 residual_rank=150
