## Supplemental docs for "An entropy-based diagnostic framework for characterizing methylation state dynamics during preimplantation development": CSB_TRO_missing_control_term_interpretation.md

# Missing control-term decomposition

This analysis formalizes the distinction in the current model state: we have not discovered a biological u_bio yet, but we have measured the correction term that such a u_bio must explain.

Model form:

```text
dz_meth/dtau = f_meth(z, tau) + B u_bio(tau)
```

Measured correction vector over 8-cell to morula: -0.7914, -0.0338, -1.9175
Equivalent correction velocity: -4.7485, -0.2031, -11.5051
Baseline strict occupancy: 0.044
Observed occupancy target: 0.875

Closest amplitude to observed occupancy: alpha=0.500, occupancy=0.933

Largest module weights in ridge reconstruction of the correction term:
- M01: weight=1.6309, abs_weight=1.6309, n_DMRs=21
- M02: weight=1.5159, abs_weight=1.5159, n_DMRs=30
- M05: weight=1.1217, abs_weight=1.1217, n_DMRs=6
- M12: weight=1.0939, abs_weight=1.0939, n_DMRs=4
- M10: weight=0.7392, abs_weight=0.7392, n_DMRs=3

Greedy module reconstruction:
- step 1, modules=M05: occupancy=0.422, cosine=0.830
- step 2, modules=M05,M01: occupancy=0.600, cosine=0.920
- step 3, modules=M05,M01,M12: occupancy=0.867, cosine=0.957
- step 4, modules=M05,M01,M12,M02: occupancy=0.956, cosine=0.991
- step 5, modules=M05,M01,M12,M02,M10: occupancy=0.956, cosine=0.993
- step 6, modules=M05,M01,M12,M02,M10,M09: occupancy=0.956, cosine=0.996
- step 7, modules=M05,M01,M12,M02,M10,M09,M08: occupancy=0.978, cosine=0.998
- step 8, modules=M05,M01,M12,M02,M10,M09,M08,M06: occupancy=1.000, cosine=1.000

Interpretation boundary: this is not a non-leaking biological mechanism yet. It defines the correction term that external RNA, ATAC, histone, motif, or chromatin variables must predict in the next stage.
