## Supplemental docs for "An entropy-based diagnostic framework for characterizing methylation state dynamics during preimplantation development": CSB_TRO_module_bio_features_interpretation.md

# Biological control features

This table defines candidate module-level u_bio features for the interpretable control term:

```text
B u_bio(tau) = sum_m beta_m u_m(tau) b_m
```

Current feature modalities:
- RNA: n=30, leakage_status=methylation_non_leaking_external_morula_allowed
- internal_genomic_proxy: n=16, leakage_status=non_morula_methylation_but_not_external_omics
- internal_methylation_prior_proxy: n=16, leakage_status=internal_methylation_prior_not_external_omics
- measured_residual_diagnostic: n=16, leakage_status=uses_morula_methylation_residual

Missing optional external inputs:
- ATAC module activity features
- histone module activity features
- motif enrichment/activity features

Interpretation boundary: internal genomic and measured-residual features are useful for dry-run and upper-bound checks, but they are not proof of a real external biological control variable. RNA/ATAC/histone/motif inputs are required for u_bio identification.
