## Supplemental docs for "An entropy-based diagnostic framework for characterizing methylation state dynamics during preimplantation development": CSB_TRO_module_latent_dynamics_interpretation.md

# CSB-TRO module/latent operator-time dynamics

This run upgrades the DMR-level velocity model from independent single-DMR ridge regressions to module and latent-state dynamics.

- 8-cell baseline leave-morula RMSE: 0.2974
- single-DMR ridge leave-morula RMSE: 0.3113
- DMR-module ridge leave-morula RMSE: 0.2866
- latent PCA ridge leave-morula RMSE: 0.2698

The result supports the diagnosis that morula reset is better captured as a coordinated module/latent transition than as independent per-DMR linear extrapolation. This is still a stage-anchored pseudo-time model, not true longitudinal embryo dynamics.
