## Supplemental docs for "An entropy-based diagnostic framework for characterizing methylation state dynamics during preimplantation development": CSB_TRO_module_latent_validation_interpretation.md

# CSB-TRO module/latent validation interpretation

This validation package keeps the strict leave-morula-out design: transitions involving morula are excluded from training, and morula is predicted from the 8-cell state in stage-anchored developmental operator time.

- 8-cell baseline RMSE: 0.2974
- DMR-module ridge RMSE: 0.2866
- latent PCA ridge RMSE: 0.2698

The paired DMR error tests compare per-DMR prediction errors against the 8-cell baseline using sign-flip paired permutation tests. The bootstrap tables report both DMR bootstrap and stage/sample bootstrap confidence intervals.

Null models test whether comparable performance can be obtained after disrupting operator-time labels, sample-level OT coupling, or module membership. The empirical p-value is the fraction of null RMSE values less than or equal to the observed RMSE, with a +1 correction.

Recommended wording: A DMR-level single-feature velocity model was insufficient for strict leave-morula-out prediction, whereas module-level and latent-state operator-time dynamics improved prediction of the morula methylation reset-basin. The validation package evaluates whether this improvement persists under DMR-level paired errors, bootstrap uncertainty, and randomized time/coupling/module controls.

Caveat: this is a stage-anchored developmental operator-time model, not true longitudinal tracking of the same embryo. In silico sensitivity results should not be described as strong causal evidence.

Key paired tests:
- DMR_module_ridge_k16 absolute_error: mean improvement 0.007586, one-sided p=0.1049
- DMR_module_ridge_k16 squared_error: mean improvement 0.006342, one-sided p=0.03815
- latent_PCA_ridge_q3 absolute_error: mean improvement 0.011379, one-sided p=0.09945
- latent_PCA_ridge_q3 squared_error: mean improvement 0.015653, one-sided p=0.0067

Key null summaries:
- DMR_module_ridge_k16 / random_tau_null: observed RMSE 0.2866, null mean 0.2956, empirical p=0.004975
- latent_PCA_ridge_q3 / random_tau_null: observed RMSE 0.2698, null mean 0.2702, empirical p=0.5124
- DMR_module_ridge_k16 / random_coupling_null: observed RMSE 0.2866, null mean 0.2856, empirical p=0.99
- latent_PCA_ridge_q3 / random_coupling_null: observed RMSE 0.2698, null mean 0.2697, empirical p=0.791
- DMR_module_ridge_k16 / random_module_null: observed RMSE 0.2866, null mean 0.2922, empirical p=0.02985
