## Supplemental docs for "An entropy-based diagnostic framework for characterizing methylation state dynamics during preimplantation development": CSB_TRO_module_linked_RNA_control_interpretation.md

# Biological control-augmented dynamics

This analysis evaluates candidate module-level biological control features in the form:

```text
dz/dtau = f_meth(z,tau) + sum_m beta_m u_m(tau) b_m
```

Current feature modalities and leakage status:
- RNA: n=30, statuses=methylation_non_leaking_external_morula_allowed
- internal_genomic_proxy: n=16, statuses=non_morula_methylation_but_not_external_omics
- internal_methylation_prior_proxy: n=16, statuses=internal_methylation_prior_not_external_omics
- measured_residual_diagnostic: n=16, statuses=uses_morula_methylation_residual

Top model results:
- internal_methylation_prior_proxy_ridge_beta_diagnostic: occupancy=1.000, cosine=1.000, PC3_recovery=0.987, status=uses_morula_methylation_residual_for_beta
- internal_genomic_proxy_ridge_beta_diagnostic: occupancy=1.000, cosine=0.997, PC3_recovery=0.944, status=uses_morula_methylation_residual_for_beta
- combined_external_ridge_beta_diagnostic: occupancy=1.000, cosine=1.000, PC3_recovery=0.983, status=uses_morula_methylation_residual_for_beta
- measured_residual_diagnostic_ridge_beta_diagnostic: occupancy=1.000, cosine=1.000, PC3_recovery=0.998, status=uses_morula_methylation_residual_for_beta
- measured_missing_correction_upper_bound: occupancy=1.000, cosine=1.000, PC3_recovery=1.000, status=measured_residual_upper_bound
- all_available_ridge_beta_diagnostic: occupancy=1.000, cosine=1.000, PC3_recovery=0.999, status=uses_morula_methylation_residual_for_beta
- RNA_ridge_beta_diagnostic: occupancy=1.000, cosine=1.000, PC3_recovery=0.983, status=uses_morula_methylation_residual_for_beta
- measured_residual_diagnostic_unit_beta: occupancy=0.978, cosine=1.000, PC3_recovery=0.764, status=measured_residual_diagnostic_not_nonleaking
- all_available_unit_beta: occupancy=0.956, cosine=0.908, PC3_recovery=0.641, status=includes_internal_proxy_features
- RNA_unit_beta_sign_flip: occupancy=0.356, cosine=0.880, PC3_recovery=0.262, status=sign_flip_control

Interpretation boundary: models whose beta was fit to the measured correction are diagnostic upper bounds. A real u_bio result requires external feature-defined controls that improve occupancy without using morula methylation to define beta or direction.
