## Supplemental docs for "An entropy-based diagnostic framework for characterizing methylation state dynamics during preimplantation development": CSB_TRO_module_linked_RNA_control_summary.md

# Module-Linked RNA Control Summary

Date: 2026-05-26

## Inputs

- Genome build: hg19/GRCh37, supported by coordinate bounds.
- Gene annotation: GENCODE v19 GRCh37.
- RNA matrix: Yan et al. 2013 / GSE36552 supplementary table 1 known RefSeq gene RPKM.
- DMR-gene link: nearest GENCODE TSS by DMR midpoint.
- Priority residual modules: M05, M01, M12, M02, M10.

## Gene Annotation

GENCODE v19 generated:

```text
57,820 gene TSS records
156 residual DMR nearest-TSS links
```

Priority module nearest-gene coverage:

```text
M01: 21 DMRs, 21 nearest genes, 7 promoter_2kb, 8 promoter_5kb
M02: 30 DMRs, 30 nearest genes, 21 promoter_2kb, 27 promoter_5kb
M05: 6 DMRs, 6 nearest genes, 2 promoter_2kb, 2 promoter_5kb
M10: 3 DMRs, 3 nearest genes, 2 promoter_2kb, 2 promoter_5kb
M12: 4 DMRs, 4 nearest genes, 3 promoter_2kb, 3 promoter_5kb
```

GSE36552 RNA symbol match among nearest genes:

```text
M01: 12 / 21 genes matched
M02: 24 / 30 genes matched
M05: 2 / 6 genes matched
M10: 2 / 3 genes matched
M12: 2 / 4 genes matched
```

## Control Results

Reference:

```text
methylation-only strict baseline occupancy_q90 = 0.044
measured correction upper bound occupancy_q90 = 1.000
```

All-module nearest-gene RNA features, including 8-cell to morula and morula to blastocyst windows:

```text
RNA_unit_beta occupancy_q90 = 0.000
RNA_unit_beta cosine = -0.880
RNA_unit_beta_sign_flip occupancy_q90 = 0.356
RNA_ridge_beta_diagnostic occupancy_q90 = 1.000
```

This broad all-window RNA table is not a valid feature-defined external control. It mixes stage windows and contains features whose raw direction opposes the measured correction.

Priority-module 8-cell to morula nearest-gene RNA delta:

```text
RNA_delta_unit_beta occupancy_q90 = 0.200
RNA_delta_unit_beta cosine = 0.448
RNA_delta_unit_beta PC3 recovery = 0.088
RNA_delta_sign_flip occupancy_q90 = 0.000
RNA_delta_ridge_beta_diagnostic occupancy_q90 = 1.000
```

Priority-module RNA repression, defined as negative 8-cell to morula RNA delta:

```text
RNA_repression_unit_beta occupancy_q90 = 0.000
RNA_repression_unit_beta cosine = -0.448
RNA_repression_sign_flip occupancy_q90 = 0.200
```

## Interpretation

The move from global RNA transition to nearest-gene module-linked RNA is informative but not yet decisive.

Safe interpretation:

```text
Nearest-TSS module-linked GSE36552 RNA delta gives modest occupancy rescue and passes sign-flip directionality in the priority-module 8-cell to morula setting, but its direction alignment is much weaker than the global RNA transition gate.
```

This indicates that:

```text
1. Global RNA transition captures the stage-level direction well.
2. Nearest-gene linking is probably too crude for a full u_bio explanation.
3. Several priority modules have incomplete RNA gene-symbol matching.
4. The next biological step should use promoter overlap, distal enhancer links, TF/motif activity, and histone H3K27ac/H3K4me3 support instead of relying only on nearest gene.
```

Do not claim:

```text
nearest-gene RNA explains the missing basin-attraction term.
RNA repression is supported as the control direction.
gene-level u_bio has been identified.
```

## Next Step

Build module-specific motif and TF activity:

```text
u_TF,m = motif_enrichment_TF,m * Delta TF expression_8cell_to_morula
```

This should be tested on M05/M01/M12/M02/M10 separately with matched DMR background controls.
