## Supplemental docs for "An entropy-based diagnostic framework for characterizing methylation state dynamics during preimplantation development": CSB_TRO_module_TF_activity_control_panel_summary.md

# Module TF Activity Control Panel

Status: `completed`

Goal: test whether module-level motif x TF expression activity for M05/M01/M12/M02/M10 robustly explains the measured morula correction direction.

## Main Result

Best feature-defined model: `old_q05_zero_filled_zscore`
- occupancy_q90: 0.222
- cosine: 0.452
- PC3 recovery: 0.094
- shuffled q95 occupancy: 0.222

## Encoding Sensitivity

The panel explicitly separates zero-filled z-scored module encodings from sparse encodings that include only modules with significant motif evidence.
- q05_sparse_flipped_activity_sign: occupancy=0.089, cosine=0.569, PC3_recovery=0.097
- q05_sparse_activity_sign: occupancy=0.044, cosine=-0.569, PC3_recovery=-0.097

Interpretation boundary: if rescue is strong only in zero-filled/z-scored encodings and not sparse significant-motif encodings, M02-KLF4/KLF5 remains exploratory and needs independent FIMO/HOMER plus chromatin-state validation.
