## Supplemental docs for "An entropy-based diagnostic framework for characterizing methylation state dynamics during preimplantation development": CSB_TRO_morula_basin_sde_interpretation.md

# Morula basin SDE analysis

This analysis keeps the developmental axis as stage-anchored operator time. It does not represent true longitudinal tracking of the same embryo.

Morula latent basin center was defined as the observed morula centroid. The q90 radius is 1.5667, giving observed q90 occupancy 0.875.

8-cell to morula distribution-level results:
- full_all_transitions / deterministic: occupancy_q90=0.378, DMR_mean_RMSE=0.2430, latent_MMD=0.6910, mean_dist=1.7126, uses_morula_calibration=no
- full_all_transitions / stochastic_global_residual: occupancy_q90=0.027, DMR_mean_RMSE=0.2434, latent_MMD=0.1246, mean_dist=5.5174, uses_morula_calibration=no
- full_all_transitions / stochastic_stage_conditioned: occupancy_q90=0.161, DMR_mean_RMSE=0.2430, latent_MMD=0.2103, mean_dist=2.6724, uses_morula_calibration=no
- full_all_transitions / basin_ou_calibrated: occupancy_q90=0.892, DMR_mean_RMSE=0.2389, latent_MMD=0.0461, mean_dist=0.9877, uses_morula_calibration=yes
- full_all_transitions / empirical_8cell_morula_noise_fullfit: occupancy_q90=0.253, DMR_mean_RMSE=0.2430, latent_MMD=0.3487, mean_dist=2.0688, uses_morula_calibration=yes
- leave_morula_transitions / deterministic: occupancy_q90=0.044, DMR_mean_RMSE=0.2452, latent_MMD=0.9342, mean_dist=2.1634, uses_morula_calibration=no
- leave_morula_transitions / stochastic_global_residual: occupancy_q90=0.024, DMR_mean_RMSE=0.2459, latent_MMD=0.1462, mean_dist=5.6210, uses_morula_calibration=no
- leave_morula_transitions / stochastic_stage_conditioned: occupancy_q90=0.085, DMR_mean_RMSE=0.2454, latent_MMD=0.2155, mean_dist=3.5436, uses_morula_calibration=no
- leave_morula_transitions / basin_ou_calibrated: occupancy_q90=0.886, DMR_mean_RMSE=0.2390, latent_MMD=0.0728, mean_dist=1.0061, uses_morula_calibration=yes

Local stability summary:
- full_all_transitions / morula_samples: inward_radial=5.4863, tangential=10.0059, contraction_score=0.3818, max_real_eigen=-3.6225
- full_all_transitions / morula_plus_blastocyst_samples: inward_radial=7.9513, tangential=8.8019, contraction_score=0.5628, max_real_eigen=-3.6225
- full_all_transitions / empirical_8cell_to_morula_OT_pairs_fullfit_reference: inward_radial=7.2403, tangential=8.4213, contraction_score=0.5336, max_real_eigen=nan
- leave_morula_transitions / morula_samples: inward_radial=6.8010, tangential=13.0859, contraction_score=0.3780, max_real_eigen=-4.1989
- leave_morula_transitions / morula_plus_blastocyst_samples: inward_radial=8.5648, tangential=10.1064, contraction_score=0.5777, max_real_eigen=-4.1989
- leave_morula_transitions / empirical_8cell_to_morula_OT_pairs_fullfit_reference: inward_radial=7.2403, tangential=8.4213, contraction_score=0.5336, max_real_eigen=nan

Basin correction calibration:
- full_all_transitions: kappa=32.0000, diffusion_scale=1.0000, calibration_occupancy_q90=0.889, objective=0.0157
- leave_morula_transitions: kappa=32.0000, diffusion_scale=0.5000, calibration_occupancy_q90=0.873, objective=0.0025

Interpretation boundary: deterministic and residual/stage-conditioned SDE modes are forward latent simulations under the fitted operator. The basin-OU mode is a calibrated low-dimensional basin transition model when it uses the morula centroid/radius, so it must not be described as strict leave-morula-out extrapolation. In silico perturbations and basin corrections should be described as sensitivity/calibration analyses, not causal mechanism proof.
