## Supplemental docs for "An entropy-based diagnostic framework for characterizing methylation state dynamics during preimplantation development": CSB_TRO_motif_TF_activity_control_dynamics.md

# Biological control-augmented dynamics

This analysis evaluates candidate module-level biological control features in the form:

```text
dz/dtau = f_meth(z,tau) + sum_m beta_m u_m(tau) b_m
```

Current feature modalities and leakage status:
- motif_activity: n=5, statuses=methylation_non_leaking_motif_x_TF_expression

Top model results:
- measured_missing_correction_upper_bound: occupancy=1.000, cosine=1.000, PC3_recovery=1.000, status=measured_residual_upper_bound
- methylation_only_strict_baseline: occupancy=0.044, cosine=nan, PC3_recovery=-0.000, status=baseline
- motif_activity_unit_beta: occupancy=0.044, cosine=nan, PC3_recovery=-0.000, status=external_feature_defined
- motif_activity_unit_beta_sign_flip: occupancy=0.044, cosine=nan, PC3_recovery=0.000, status=sign_flip_control
- motif_activity_ridge_beta_diagnostic: occupancy=0.044, cosine=nan, PC3_recovery=-0.000, status=uses_morula_methylation_residual_for_beta
- combined_external_unit_beta: occupancy=0.044, cosine=nan, PC3_recovery=-0.000, status=external_feature_defined
- combined_external_unit_beta_sign_flip: occupancy=0.044, cosine=nan, PC3_recovery=0.000, status=sign_flip_control
- combined_external_ridge_beta_diagnostic: occupancy=0.044, cosine=nan, PC3_recovery=-0.000, status=uses_morula_methylation_residual_for_beta
- all_available_unit_beta: occupancy=0.044, cosine=nan, PC3_recovery=-0.000, status=includes_internal_proxy_features
- all_available_unit_beta_sign_flip: occupancy=0.044, cosine=nan, PC3_recovery=0.000, status=sign_flip_control

Interpretation boundary: models whose beta was fit to the measured correction are diagnostic upper bounds. A real u_bio result requires external feature-defined controls that improve occupancy without using morula methylation to define beta or direction.
