## Supplemental docs for "An entropy-based diagnostic framework for characterizing methylation state dynamics during preimplantation development": CSB_TRO_motif_TF_activity_control_summary.md

# Motif x TF Activity Control Summary

Date: 2026-05-26

## Goal

Test whether module-specific TF regulatory activity can explain the missing morula basin-attraction correction better than nearest-gene RNA.

The intended variable is:

```text
u_m,t = motif_enrichment_m,t * Delta TF_expression_t
u_m = sum_t u_m,t
```

## Completed Inputs

RNA expression:

```text
GSE36552 gene-stage RPKM matrix
20,214 genes
TF deltas computed for:
- 4-cell -> 8-cell
- 8-cell -> morula
- morula -> blastocyst
```

Motif input:

```text
JASPAR 2024 CORE vertebrates MEME
UCSC hg19 DMR sequences fetched for 156 residual DMRs
848 expressed-TF JASPAR motifs scanned
4,240 module x TF burden rows generated
```

Important boundary:

```text
This was a lightweight Python PWM burden scan, not a full HOMER/FIMO enrichment with matched GC/CpG background.
```

## TF RNA Signal

The RNA side is biologically informative. Examples from 8-cell to morula:

```text
TPRX1 strongly decreases
DUXA strongly decreases
ZSCAN4 strongly decreases
LEUTX decreases
SOX2 decreases
TFAP2C decreases
KLF17 increases
GATA3 increases
CDX2 increases
POU5F1 increases
```

This supports the idea that TF activity around 8-cell to morula is a plausible biological control layer.

## Motif Enrichment Result

Strict signed -log10(q) score:

```text
all qvalue = 1.0
motif score = 0
motif_activity_unit_beta occupancy_q90 = 0.044
```

So the strict JASPAR first-pass burden does not produce a usable enriched motif signal.

Exploratory logOR fallback:

```text
motif_activity_unit_beta:
  occupancy_q90 = 0.000
  direction cosine = -0.734
  PC3 recovery = -0.100

motif_activity_unit_beta_sign_flip:
  occupancy_q90 = 0.222
  direction cosine = 0.734
  PC3 recovery = 0.100

motif_activity_ridge_beta_diagnostic:
  occupancy_q90 = 1.000
  direction cosine = 0.99998
  PC3 recovery = 0.993
```

The diagnostic ridge result only says the module basis can reconstruct the measured correction when beta is fit to the residual. It does not validate the motif x TF activity as a non-leaking external control.

## Interpretation

Safe conclusion:

```text
The TF expression delta layer is available and biologically plausible, but the current lightweight JASPAR PWM burden scan is not sufficient to identify a valid motif x TF control. Under strict q-value scoring it is null; under exploratory logOR fallback its unit-beta direction is opposite to the measured correction.
```

This is not a failure of the TF hypothesis. It is a limitation of the first-pass motif screen:

```text
1. The background was all non-module residual DMRs, not matched GC/CpG/length/background methylation regions.
2. The PWM threshold was generic random-sequence based, not calibrated to local genomic background.
3. Module sizes are small for M05/M10/M12.
4. JASPAR-only motifs may miss early embryo TFs or composite motifs.
5. TF expression alone may need chromatin accessibility or H3K27ac/H3K4me3 state to define activity.
```

## Next Step

Do not claim motif x TF activity explains u_bio yet.

The next stronger route is:

```text
1. Run HOMER or FIMO with hg19 FASTA.
2. Use matched background regions matched for length, GC, CpG density, and age-DMR properties.
3. Compute module-specific motif enrichment for M05/M01/M12/M02/M10.
4. Rebuild motif x TF activity from enriched motifs only.
5. If still weak, integrate H3K27ac/H3K4me3 and ATAC as chromatin-state gates.
```
