## Supplemental docs for "An entropy-based diagnostic framework for characterizing methylation state dynamics during preimplantation development": CSB_TRO_motif_TF_activity_interpretation.md

# Motif x TF expression activity

Status: completed

TF expression deltas were computed from GSE36552 gene-stage RPKM using log2(expression + 1).

Expected motif enrichment input:
- E:\CSB_TRO_operator_time_DMR_dynamics_2026-05-25\external\motif\module_motif_enrichment.tsv

Required columns: module_id, TF, log_odds_ratio, qvalue. Optional: pvalue, motif_hit_count, background_hit_count, motif_database.
