## Supplemental docs for "An entropy-based diagnostic framework for characterizing methylation state dynamics during preimplantation development": CSB_TRO_motif_TF_matched_background_control_summary.md

# Motif x TF Matched-Background Control Summary

Date: 2026-05-26

## Goal

Improve the first-pass JASPAR motif x TF experiment by replacing the all-non-module residual DMR background with matched non-age windows.

## Background Design

Matched regions:

```text
C:\Users\18068\Desktop\CSB_TRO_Project_2026-05-24\input_tables\GSE81233_matched_non_age_window_regions_n100.tsv
```

Each residual age-DMR has 100 matched non-age windows. The pilot used:

```text
priority modules = M05, M01, M12, M02, M10
foreground DMRs = 64
matched background sets per DMR = 10
matched background sequences = 640
total sequences = 704
```

Motif scan:

```text
JASPAR 2024 CORE vertebrates
candidate early-embryo TFs + top 8-cell to morula TF expression-change motifs
19 motifs scanned
module-specific threshold = 95th percentile of matched background motif scores
```

Boundary:

```text
This is still a Python PWM burden pilot. HOMER/FIMO with explicit GC/CpG matched background remains preferred for final claims.
```

## Broad Matched-Background Result

Using all 19 candidate/high-delta motif scores with logOR fallback:

```text
motif_activity_unit_beta occupancy_q90 = 0.000
direction cosine = -0.720
PC3 recovery = -0.131

motif_activity_unit_beta_sign_flip occupancy_q90 = 0.156
direction cosine = 0.720
PC3 recovery = 0.131
```

This broad feature set is not a valid feature-defined control because the unit direction is opposite.

## Strict Significant-Motif Result

Filtering to q <= 0.05 leaves:

```text
M02: KLF4, KLF5
```

Strict q<=0.05 motif x TF activity:

```text
motif_activity_unit_beta occupancy_q90 = 0.222
direction cosine = 0.452
PC3 recovery = 0.094
DMR mean RMSE = 0.244205
DMR correlation = 0.632164

sign-flip occupancy_q90 = 0.000
```

Random shuffled feature controls:

```text
mean occupancy_q90 = 0.095
max occupancy_q90 = 0.222
```

Diagnostic ridge beta:

```text
occupancy_q90 = 1.000
direction cosine = 0.99996
PC3 recovery = 0.996
```

The diagnostic ridge result uses the measured correction to fit beta and is not a non-leaking biological-control model.

## Interpretation

The matched-background strict q<=0.05 motif x TF result is directionally valid but modest:

```text
occupancy improves from 0.044 to 0.222
cosine improves to 0.452
sign-flip fails
signal is localized to M02 KLF4/KLF5 motifs
```

This is slightly stronger than priority-module nearest-gene RNA:

```text
nearest-gene RNA_delta occupancy_q90 = 0.200
nearest-gene RNA_delta cosine = 0.448
nearest-gene RNA_delta PC3 recovery = 0.088
```

But it is not yet strong enough to claim true u_bio identification:

```text
global RNA transition still has much stronger direction cosine = 0.993
motif x TF signal is narrow and M02-dominated
random controls can reach the same occupancy maximum in this small feature setting
```

Safe claim:

```text
A matched-background JASPAR motif x TF pilot nominates an M02-linked KLF4/KLF5 regulatory signal that modestly rescues morula basin occupancy and passes sign-flip control, but the evidence remains exploratory and requires HOMER/FIMO plus chromatin-state validation.
```

## Next Step

Move to chromatin-state validation:

```text
1. test whether M02 and KLF4/KLF5-linked residual DMRs overlap H3K27ac/H3K4me3 or ATAC features;
2. expand matched motif enrichment with more matched sets or HOMER/FIMO;
3. integrate motif x TF with H3K27ac/H3K4me3 as a chromatin gate.
```
