## Supplemental docs for "An entropy-based diagnostic framework for characterizing methylation state dynamics during preimplantation development": CSB_TRO_next_phase_experiment_update_2026-05-26.md

# Next-Phase Experiment Update

Date: 2026-05-26

## Experiments Added

Two experiments were added after the project split into biological-control discovery and complex-dynamics validation.

### 1. Bifurcation-like basin-entry scan

Script:

```text
code/run_bifurcation_like_basin_entry_scan.py
```

Outputs:

```text
results/CSB_TRO_alpha_bifurcation_scan.tsv
results/CSB_TRO_module_specific_bifurcation_scan.tsv
results/CSB_TRO_local_jacobian_eigenvalues.tsv
results/CSB_TRO_bifurcation_like_scan_summary.tsv
figures/CSB_TRO_alpha_bifurcation_scan.svg
figures/CSB_TRO_module_specific_bifurcation_scan.svg
docs/CSB_TRO_bifurcation_like_basin_entry_summary.md
```

Main result:

```text
full measured correction:
alpha=0 occupancy = 0.044
alpha=1 occupancy = 1.000
first alpha reaching observed q90 occupancy = 0.50
steepest local occupancy slope starts near alpha = 0.45
```

Module scan:

```text
M05 alpha=1 occupancy = 0.422
M05+M01 alpha=1 occupancy = 0.600
M05+M01+M12 alpha=1 occupancy = 0.867
M05+M01+M12+M02 alpha=1 occupancy = 0.956
M05+M01+M12+M02+M10 alpha=1 occupancy = 0.956
```

Interpretation:

```text
The measured correction term shows threshold-like basin entry and supports a bifurcation-like / tipping-like interpretation of morula reset.
```

Boundary:

```text
This is not proof of a strict saddle-node bifurcation. The current methylation operator is affine in z, and the measured correction is a constant diagnostic vector, so local Jacobian eigenvalues are unchanged by alpha in this implementation.
```

### 2. Module TF activity control panel

Script:

```text
code/run_module_tf_activity_control_panel.py
```

Outputs:

```text
results/CSB_TRO_module_TF_activity_control_panel_features.tsv
results/CSB_TRO_module_TF_activity_control_panel_metrics.tsv
results/CSB_TRO_module_TF_activity_control_panel_random.tsv
results/CSB_TRO_module_TF_activity_control_panel_summary.tsv
figures/CSB_TRO_module_TF_activity_control_panel.svg
docs/CSB_TRO_module_TF_activity_control_panel_summary.md
```

Main result:

```text
old q<=0.05 zero-filled z-score encoding:
occupancy = 0.222
cosine = 0.452
PC3 recovery = 0.094
shuffled q95 occupancy = 0.222
```

Sparse significant-motif result:

```text
q05 sparse activity sign:
occupancy = 0.044
cosine = -0.569
PC3 recovery = -0.097

q05 sparse flipped activity sign:
occupancy = 0.089
cosine = 0.569
PC3 recovery = 0.097
```

Interpretation:

```text
The M02-KLF4/KLF5 result remains biologically plausible but is encoding-sensitive. The previous occupancy rescue depends on zero-filled z-scored module encoding and is not stronger than the shuffled q95 control. Sparse significant-motif encoding does not rescue occupancy beyond 0.089.
```

This strengthens the caution:

```text
M02-KLF4/KLF5 is not a complete u_bio and should not be claimed as the main biological control term without independent FIMO/HOMER and chromatin-state validation.
```

## Current Decision

The dynamics side is now stronger:

```text
morula reset can be described as a threshold-like, control-induced basin-entry behavior under the measured correction term.
```

The biology side remains unresolved:

```text
current motif x TF evidence nominates M02-KLF4/KLF5 but does not robustly identify u_bio.
```

## Next Best Experiment

Prioritize external chromatin support:

```text
1. obtain processed H3K27ac/H3K4me3/H3K27me3 peaks or signal tracks;
2. overlap M05/M01/M12/M02/M10 residual DMRs against matched background;
3. build histone-gated motif x TF activity;
4. rerun biological-control dynamics.
```

If histone peaks remain unavailable:

```text
1. run independent FIMO/HOMER module motif validation;
2. expand matched background sets;
3. run random module / shuffled TF / motif-label permutation controls;
4. only retain TF candidates that survive sparse encoding and controls.
```
