## Supplemental docs for "An entropy-based diagnostic framework for characterizing methylation state dynamics during preimplantation development": CSB_TRO_nonleaking_distribution_dynamics_interpretation.md

# Non-leaking distribution dynamics

This experiment asks whether morula-like population occupancy can be generated without using morula centroid, morula radius, morula sample distribution, or 8-cell to morula endpoint noise during training. Morula samples are used only for final evaluation.

Models tested:
- pre-morula velocity ODE/SDE: drift and residual diffusion trained only on transitions ending at or before 8-cell.
- affine Gaussian transition kernel: direct probabilistic kernel z_next | z, tau trained only on pre-morula transitions.
- moment-extrapolated Gaussian: target morula mean/cov extrapolated from pre-morula stage moments.
- validation-tuned OU: attraction strength tuned on 4-cell to 8-cell validation, then applied to an extrapolated morula center.
- morula-calibrated OU upper bound: shown only as a diagnostic ceiling, not a strict model.

Pre-morula extrapolated center: 1.4630, -2.4973, -8.4225
Observed morula center, evaluation only: -3.1743, -0.0308, -2.8184

8-cell to morula evaluation:
- pre_morula_moment_extrapolated_gaussian: occupancy_q90=0.000, DMR_mean_RMSE=0.2782, latent_MMD=1.2530, mean_dist=7.7416, leakage=no
- pre_morula_validation_tuned_extrapolated_ou: occupancy_q90=0.000, DMR_mean_RMSE=0.2680, latent_MMD=1.2603, mean_dist=6.1224, leakage=no
- pre_morula_affine_gaussian_transition_kernel: occupancy_q90=0.016, DMR_mean_RMSE=0.2526, latent_MMD=0.4113, mean_dist=4.3093, leakage=no
- pre_morula_velocity_residual_sde: occupancy_q90=0.023, DMR_mean_RMSE=0.2458, latent_MMD=0.1495, mean_dist=5.6173, leakage=no
- pre_morula_velocity_deterministic: occupancy_q90=0.044, DMR_mean_RMSE=0.2452, latent_MMD=0.9342, mean_dist=2.1634, leakage=no
- leave_morula_with_blastocyst_interpolated_gaussian: occupancy_q90=0.023, DMR_mean_RMSE=0.2412, latent_MMD=0.5263, mean_dist=2.2665, leakage=no_morula_but_uses_future_blastocyst
- leave_morula_with_blastocyst_interpolated_ou: occupancy_q90=0.376, DMR_mean_RMSE=0.2409, latent_MMD=0.2897, mean_dist=1.9009, leakage=no_morula_but_uses_future_blastocyst
- morula_calibrated_ou_upper_bound: occupancy_q90=0.848, DMR_mean_RMSE=0.2386, latent_MMD=0.0380, mean_dist=1.0687, leakage=yes_upper_bound

4-cell to 8-cell validation-selected OU parameters:
- kappa=32.0000, diffusion_scale=0.2500, latent_mean_rmse=0.2039, cov_error=0.7464

Current non-leaking best occupancy is 0.044 from pre_morula_velocity_deterministic. This should be compared against observed q90 occupancy 0.875 and the morula-calibrated upper bound, not presented as complete autonomous basin generation unless it approaches the observed target.
