## Supplemental docs for "An entropy-based diagnostic framework for characterizing methylation state dynamics during preimplantation development": CSB_TRO_reproducibility_package_index.md

# Reproducibility Package Index

Status: `public_data_bounded_manuscript_v0.8`

Project root:

`E:\CSB_TRO_operator_time_DMR_dynamics_2026-05-25`

## Primary Reproduction Order

1. `E:\CSB_TRO_operator_time_DMR_dynamics_2026-05-25\code\run_dmr_operator_time_dynamics.py`
2. `E:\CSB_TRO_operator_time_DMR_dynamics_2026-05-25\code\run_module_latent_operator_time_dynamics.py`
3. `E:\CSB_TRO_operator_time_DMR_dynamics_2026-05-25\code\run_module_latent_validation.py`
4. `E:\CSB_TRO_operator_time_DMR_dynamics_2026-05-25\code\run_advanced_latent_dynamics.py`
5. `E:\CSB_TRO_operator_time_DMR_dynamics_2026-05-25\code\run_morula_basin_sde.py`
6. `E:\CSB_TRO_operator_time_DMR_dynamics_2026-05-25\code\run_nonleaking_distribution_dynamics.py`
7. `E:\CSB_TRO_operator_time_DMR_dynamics_2026-05-25\code\run_basin_residual_control_field.py`
8. `E:\CSB_TRO_operator_time_DMR_dynamics_2026-05-25\code\run_missing_control_term_decomposition.py`
9. `E:\CSB_TRO_operator_time_DMR_dynamics_2026-05-25\code\run_bifurcation_like_basin_entry_scan.py`
10. `E:\CSB_TRO_operator_time_DMR_dynamics_2026-05-25\code\run_inverse_ATAC_module_decomposition.py`
11. `E:\CSB_TRO_operator_time_DMR_dynamics_2026-05-25\code\run_dual_branch_chromatin_state_control.py`
12. `E:\CSB_TRO_operator_time_DMR_dynamics_2026-05-25\code\run_dual_branch_robustness_controls.py`
13. `E:\CSB_TRO_operator_time_DMR_dynamics_2026-05-25\code\run_dual_branch_structure_validation.py`
14. `E:\CSB_TRO_operator_time_DMR_dynamics_2026-05-25\code\run_branch_bound_biological_control_candidates.py`
15. `E:\CSB_TRO_operator_time_DMR_dynamics_2026-05-25\code\run_hesc_histone_branch_identity_proxy.py`
16. `E:\CSB_TRO_operator_time_DMR_dynamics_2026-05-25\code\run_embryo_histone_dual_branch_replacement.py`
17. `E:\CSB_TRO_operator_time_DMR_dynamics_2026-05-25\code\run_final_sensitivity_and_claim_boundary.py`

## Core Tables

- `E:\CSB_TRO_operator_time_DMR_dynamics_2026-05-25\results\CSB_TRO_final_model_comparison.tsv`
- `E:\CSB_TRO_operator_time_DMR_dynamics_2026-05-25\results\CSB_TRO_final_sensitivity_summary.tsv`
- `E:\CSB_TRO_operator_time_DMR_dynamics_2026-05-25\results\CSB_TRO_basin_residual_control_metrics.tsv`
- `E:\CSB_TRO_operator_time_DMR_dynamics_2026-05-25\results\CSB_TRO_alpha_bifurcation_scan.tsv`
- `E:\CSB_TRO_operator_time_DMR_dynamics_2026-05-25\results\CSB_TRO_dual_branch_chromatin_state_summary.tsv`
- `E:\CSB_TRO_operator_time_DMR_dynamics_2026-05-25\results\CSB_TRO_dual_branch_sign_control.tsv`
- `E:\CSB_TRO_operator_time_DMR_dynamics_2026-05-25\results\CSB_TRO_dual_branch_random_partition_control.tsv`
- `E:\CSB_TRO_operator_time_DMR_dynamics_2026-05-25\results\CSB_TRO_dual_branch_exact_partition_sign_control.tsv`
- `E:\CSB_TRO_operator_time_DMR_dynamics_2026-05-25\results\CSB_TRO_hESC_histone_branch_summary.tsv`
- `E:\CSB_TRO_operator_time_DMR_dynamics_2026-05-25\results\CSB_TRO_embryo_histone_control_metrics.tsv`
- `E:\CSB_TRO_operator_time_DMR_dynamics_2026-05-25\results\CSB_TRO_embryo_histone_random_controls.tsv`

## Main Figures

- `E:\CSB_TRO_operator_time_DMR_dynamics_2026-05-25\figures\CSB_TRO_final_figure1_concept.svg`
- `E:\CSB_TRO_operator_time_DMR_dynamics_2026-05-25\figures\CSB_TRO_final_figure2_baseline_failure.svg`
- `E:\CSB_TRO_operator_time_DMR_dynamics_2026-05-25\figures\CSB_TRO_final_figure3_threshold_entry.svg`
- `E:\CSB_TRO_operator_time_DMR_dynamics_2026-05-25\figures\CSB_TRO_final_figure4_dual_branch_structure.svg`
- `E:\CSB_TRO_operator_time_DMR_dynamics_2026-05-25\figures\CSB_TRO_final_figure5_external_model_comparison.svg`
- `E:\CSB_TRO_operator_time_DMR_dynamics_2026-05-25\figures\CSB_TRO_final_figure6_data_access_boundary.svg`

## Manuscript Files

- `E:\CSB_TRO_operator_time_DMR_dynamics_2026-05-25\docs\CSB_TRO_public_data_bounded_manuscript_draft.md`
- `E:\CSB_TRO_operator_time_DMR_dynamics_2026-05-25\docs\CSB_TRO_final_claim_boundary_summary.md`
- `E:\CSB_TRO_operator_time_DMR_dynamics_2026-05-25\docs\CSB_TRO_reproducibility_package_index.md`

## Boundary

The public-data package supports the diagnostic dual-branch control framework. It does not include controlled-access H3K27ac_morula or H3K4me3_morula tracks, so it does not claim final u_bio closure.
