## Supplemental docs for "An entropy-based diagnostic framework for characterizing methylation state dynamics during preimplantation development": CSB_TRO_residual_biology_interpretation.md

# Residual DMR biological annotation

This file converts the mathematical residual DMR ranking into a biological annotation table. With the current local files, annotation is limited to DMR coordinates, modules, CpG/width features, age weights, and overlap with internal dynamic DMR sets.

External gene, motif, ATAC, histone, CpG island, and repeat annotations were not inferred unless explicit files were supplied to the script.

Top residual DMRs:
- cluster_2623 (M05, chr15:37190548-37190694): residual_delta=-0.1966, nearest_gene=NA_no_gene_tss_file
- cluster_6262 (M10, chr6:31628935-31629199): residual_delta=-0.1912, nearest_gene=NA_no_gene_tss_file
- cluster_6892 (M05, chr7:37947051-37947061): residual_delta=-0.1798, nearest_gene=NA_no_gene_tss_file
- cluster_5678 (M10, chr4:186425697-186425754): residual_delta=-0.1646, nearest_gene=NA_no_gene_tss_file
- cluster_3275 (M05, chr17:10421635-10422077): residual_delta=-0.1422, nearest_gene=NA_no_gene_tss_file
- cluster_4373 (M01, chr2:49382159-49382401): residual_delta=-0.1362, nearest_gene=NA_no_gene_tss_file
- cluster_4141 (M05, chr19:51858208-51858276): residual_delta=-0.1345, nearest_gene=NA_no_gene_tss_file
- cluster_7042 (M05, chr7:108168130-108168721): residual_delta=-0.1298, nearest_gene=NA_no_gene_tss_file
- cluster_4605 (M01, chr2:227288327-227288518): residual_delta=-0.1246, nearest_gene=NA_no_gene_tss_file
- cluster_6498 (M12, chr6:132931714-132931736): residual_delta=0.1234, nearest_gene=NA_no_gene_tss_file

Feature-set summary:
- top25: n=25, modules=M01,M03,M05,M07,M08,M09,M10,M12
- top50: n=50, modules=M00,M01,M02,M03,M05,M06,M07,M08,M09,M10,M12,M13,M15
- top100: n=100, modules=M00,M01,M02,M03,M04,M05,M06,M07,M08,M09,M10,M11,M12,M13,M14,M15

Missing external inputs for full mechanism validation:
- nearest gene or gene set annotation
- genome FASTA plus motif database or motif-scan output
- ATAC peak/signal BED files
- histone mark peak/signal BED files
- stage-resolved RNA expression matrix

Next strict mechanism step: provide RNA/ATAC/histone/motif/gene annotation files, then fit or test whether those external features predict the residual direction without using morula methylation.
