## Supplemental docs for "An entropy-based diagnostic framework for characterizing methylation state dynamics during preimplantation development": CSB_TRO_residual_DMR_robustness_summary.md

# Residual DMR robustness

This package tests whether the residual DMR control set is stable, directional, and stronger than matched DMR controls. The residual itself remains diagnostic because it uses observed morula methylation for definition.

Observed morula q90 occupancy target: 0.875
Strict pre-morula baseline occupancy: 0.044

Top25 forward residual occupancy: 0.956
Top25 sign-flip occupancy: 0.000
All residual except Top25 occupancy: 0.556
Matched random Top25 occupancy mean: 0.108
Matched random Top25 occupancy q95: 0.156

Most stable Top25 DMRs by bootstrap selection frequency:
- cluster_6655: freq_top25=1.000, mean_rank=14.9
- cluster_2623: freq_top25=1.000, mean_rank=1.4
- cluster_6960: freq_top25=1.000, mean_rank=11.2
- cluster_4904: freq_top25=1.000, mean_rank=20.0
- cluster_4141: freq_top25=1.000, mean_rank=7.0
- cluster_7042: freq_top25=1.000, mean_rank=8.7
- cluster_2743: freq_top25=1.000, mean_rank=11.2
- cluster_6498: freq_top25=1.000, mean_rank=9.7
- cluster_5678: freq_top25=1.000, mean_rank=3.8
- cluster_543: freq_top25=1.000, mean_rank=17.1

Top module add/remove effects:
- remove_module_M11: occupancy_q90=1.000, n_DMRs=152
- remove_module_M07: occupancy_q90=1.000, n_DMRs=147
- remove_module_M08: occupancy_q90=1.000, n_DMRs=149
- remove_module_M00: occupancy_q90=1.000, n_DMRs=140
- remove_module_M02: occupancy_q90=1.000, n_DMRs=126
- remove_module_M09: occupancy_q90=1.000, n_DMRs=153
- remove_module_M06: occupancy_q90=1.000, n_DMRs=133
- remove_module_M13: occupancy_q90=1.000, n_DMRs=149

Interpretation boundary: strong forward/sign-flip/matched-random separation supports a compact, directional residual DMR control component. It does not by itself identify the external regulatory/chromatin variable that predicts this component without morula methylation.
