## Supplemental docs for "An entropy-based diagnostic framework for characterizing methylation state dynamics during preimplantation development": CSB_TRO_RNA_transition_control_amplitude_scan.md

# RNA transition control amplitude scan

This scan multiplies the same GSE36552 RNA transition gate by fixed amplitudes. It does not fit beta to the measured morula methylation residual.

Forward fixed-amplitude results:
- scale=0: occupancy_q90=0.044, cosine=nan, PC3_recovery=-0.000
- scale=0.25: occupancy_q90=0.044, cosine=0.993, PC3_recovery=0.036
- scale=0.5: occupancy_q90=0.044, cosine=0.993, PC3_recovery=0.072
- scale=1: occupancy_q90=0.200, cosine=0.993, PC3_recovery=0.144
- scale=1.5: occupancy_q90=0.378, cosine=0.993, PC3_recovery=0.216
- scale=2: occupancy_q90=0.489, cosine=0.993, PC3_recovery=0.288
- scale=3: occupancy_q90=0.711, cosine=0.993, PC3_recovery=0.432
- scale=4: occupancy_q90=0.956, cosine=0.993, PC3_recovery=0.576

Interpretation: increasing rescue under fixed scaling supports direction compatibility, but the scale itself is not independently calibrated by external biology yet.
