## Supplemental docs for "An entropy-based diagnostic framework for characterizing methylation state dynamics during preimplantation development": CSB_TRO_RNA_transition_control_interpretation.md

# RNA transition control features

This is a first external-RNA control experiment using GSE36552 stage-level transition summaries.

It does not use morula methylation to define the RNA activity, beta, center, radius, or occupancy. It does use the previously nominated residual module directions M05/M01/M12/M02/M10, so this is a module-gated RNA transition test rather than a gene-linked u_bio mechanism.

Composite normalized RNA transition activity: 0.207143

Components:
- RNA_potency_loss_8cell_to_morula: raw=0.037973, normalized=0.089439
- RNA_entropy_loss_8cell_to_morula: raw=0.019792, normalized=0.415306
- RNA_transition_displacement_8cell_to_morula: raw=0.090251, normalized=0.175928
- RNA_potency_penalty_8cell_to_morula: raw=0.053098, normalized=0.147900

Next stronger test requires gene-linked RNA, TF/motif, ATAC, or histone features per module.
