## Supplemental docs for "An entropy-based diagnostic framework for characterizing methylation state dynamics during preimplantation development": CSB_TRO_RNA_transition_replication_control_summary.md

# RNA Transition Replication Control Summary

Date: 2026-05-26

## Question

Can independent external RNA transition summaries provide a methylation-non-leaking control signal compatible with the measured morula basin-attraction correction?

## Inputs

- GSE36552 RNA transition summary:
  `C:\Users\18068\Desktop\CSB_TRO_Project_2026-05-24\results\CSB_TRO_RNA_GSE36552_transition_bridges.tsv`
- GSE44183 external RNA transition summary:
  `C:\Users\18068\Desktop\CSB_TRO_Project_2026-05-24\results\CSB_TRO_RNA_GSE44183_external_transition_bridges.tsv`
- Residual module basis:
  `E:\CSB_TRO_operator_time_DMR_dynamics_2026-05-25\results\CSB_TRO_missing_control_term_module_basis.tsv`

The control uses the pre-specified residual modules:

```text
M05, M01, M12, M02, M10
```

## RNA Activity

Composite 8-cell to morula RNA transition activity:

```text
GSE36552 = 0.207143
GSE44183_external = 0.151959
consensus mean = 0.179551
```

The activity is stage-level RNA transition information. It does not use morula methylation center, radius, occupancy, or residual beta. It is not yet gene-linked or TF/motif-linked.

## Main Results

Reference:

```text
methylation-only strict baseline occupancy_q90 = 0.044444
measured missing correction upper bound occupancy_q90 = 1.000000
```

External RNA unit-beta results:

```text
GSE36552 RNA_unit_beta:
  occupancy_q90 = 0.200000
  direction cosine = 0.992581
  PC3-negative recovery = 0.143994
  DMR mean RMSE = 0.243630
  DMR correlation = 0.631540

GSE44183_external RNA_unit_beta:
  occupancy_q90 = 0.155556
  direction cosine = 0.992581
  PC3-negative recovery = 0.105633
  DMR mean RMSE = 0.244030
  DMR correlation = 0.629897

consensus RNA_unit_beta:
  occupancy_q90 = 0.155556
  direction cosine = 0.992581
  PC3-negative recovery = 0.124814
  DMR mean RMSE = 0.243828
  DMR correlation = 0.630727
```

Sign-flip controls:

```text
GSE36552 sign-flip occupancy_q90 = 0.000000
GSE44183_external sign-flip occupancy_q90 = 0.022222
consensus sign-flip occupancy_q90 = 0.000000
```

Diagnostic ridge-to-measured-correction models reach occupancy_q90 = 1.000000 for all three cases, but those fits use the measured morula methylation residual for beta and are diagnostic upper bounds, not non-leaking external biological models.

## Interpretation

The RNA transition signal replicates across two independent stage-level RNA datasets:

```text
external RNA transition activity has the correct latent direction
strong PC3-negative pull direction is recovered qualitatively
natural unit amplitude gives modest but real occupancy rescue
sign-flip controls fail
```

This is stronger than the prior single-dataset RNA gate, but it is still not a complete biological mechanism. The current RNA feature is stage-level and module-gated by pre-specified residual methylation modules. It does not yet identify which genes, TFs, motifs, chromatin states, or pathways produce the missing methylation correction.

## Boundary

Safe claim:

```text
Independent external RNA transition summaries provide a methylation-non-leaking signal that is directionally aligned with the measured morula basin-attraction correction and modestly rescues morula basin occupancy at natural unit amplitude.
```

Do not claim:

```text
RNA fully explains the morula methylation reset.
The true gene-level u_bio has been identified.
The control strength is independently calibrated.
```

## Next Experiment

The next higher-evidence step is gene/module-linked biology:

```text
1. lock genome build;
2. add GTF/TSS annotation;
3. map M05/M01/M12/M02/M10 residual DMRs to nearby genes;
4. obtain or reconstruct gene-level RNA by stage;
5. test module-linked RNA_delta, TF expression, motif x TF expression, ATAC, and histone controls.
```
