## Supplemental docs for "An entropy-based diagnostic framework for characterizing methylation state dynamics during preimplantation development": CSB_TRO_RNA_transition_replication_features.md

# RNA transition replication features

This builds RNA transition gates from two stage-level RNA datasets and a consensus average.

Composite activities:
- GSE36552: 0.207143
- GSE44183_external: 0.151959
- consensus: 0.179551

These are methylation-non-leaking external RNA transition features, but still not gene-linked or TF/motif-linked.
