## Supplemental docs for "An entropy-based diagnostic framework for characterizing methylation state dynamics during preimplantation development": data_manifest.md

# Data manifest

Date: 2026-05-21

## Project paths

Server project path:

```text
/root/autodl-tmp/TRO_Project
```

Local synchronized result package:

```text
<PROJECT_ROOT>\TRO_Project_current_results
```

## Public datasets

### GSE102970

Role:

Human sperm methylation and male-age-associated DMR source.

Use in this project:

The GEO exposure matrix did not provide usable sample age metadata for direct modeling. The analysis therefore used age-associated DMR / CpG weights extracted from the paper supplementary Table S6.

Local extracted metadata:

```text
metadata/GSE102970_TableS6_age_dmr_weights.tsv
metadata/GSE102970_TableS6_age_cpg_weights.tsv
```

### GSE81233

Role:

Human preimplantation embryo DNA methylome.

Use in this project:

Primary DNA methylation dataset for stage-level methylation entropy and age-associated methylation entropy.

Important exclusion:

```text
GSM2986343_scBS-2C-10-1.Cmet.bed.gz
```

This file repeatedly failed gzip validation and was excluded. Final valid sample count:

```text
204
```

Main result tables:

```text
tables/GSE81233_valid204_stage_epi_age_metrics.tsv
tables/GSE81233_valid204_bootstrap_ground_zero_frequency.tsv
tables/GSE81233_valid204_adjacent_stage_mannwhitney.tsv
tables/GSE81233_valid204_internal_reset_score.tsv
tables/GSE81233_excluded_corrupt_or_missing_files.tsv
```

### GSE36552

Role:

Human preimplantation embryo RNA-seq.

Use in this project:

Primary RNA entropy and potency-marker analysis.

Main result tables:

```text
tables/GSE36552_RNA_entropy_potency_by_stage.tsv
tables/GSE36552_potency_component_by_stage.tsv
tables/GSE36552_potency_pairwise_tests.tsv
```

### GSE44183

Role:

Human and mouse early embryo RNA-seq.

Use in this project:

External RNA potency validation. The human expression matrix supports that 8-cell and morula occupy a high-potency region. It does not include blastocyst in the same way as GSE36552 and should not be treated as direct absolute-value replication.

Main result tables:

```text
tables/GSE44183_external_potency_validation.tsv
tables/GSE44183_external_potency_pairwise_tests.tsv
```

## Stage mapping

DNA to RNA mapping used in the dual-entropy analysis:

```text
MII oocyte -> oocyte
zygote/PN  -> zygote
2-cell     -> 2-cell
4-cell     -> 4-cell
8-cell     -> 8-cell
morula     -> morula
blastocyst -> blastocyst
```

ICM and TE are retained in DNA-only summaries but are not used in the main DNA-RNA stage alignment table.

## Metrics

### S_epi

Generic binary methylation entropy.

### S_epi-age

Age-associated weighted methylation entropy. This is the core DNA damage/perturbation metric.

### S_RNA

Global transcriptome Shannon entropy.

### PotencyScore

Composite potency proxy based on detected gene count and developmental potency-marker activity.

### ResetScore

Internal normalization from MII oocyte = 0 to morula = 1.

### TRO_score

Operational reset score preserving developmental potency:

```text
TRO_score = ResetScore * PotencyPreserve
```
