## Supplemental docs for "An entropy-based diagnostic framework for characterizing methylation state dynamics during preimplantation development": Experiment1B_DNA_robustness_interpretation.md

# Experiment 1B: DNA methylation robustness validation

## Goal

Experiment 1B tests whether the morula-stage minimum of age-associated methylation entropy (`S_epi-age`) is robust to feature coverage, sample imbalance, and DMR/weight choices.

## Main Result

The morula stage remained the minimum `S_epi-age` state across the main robustness settings:

| test | ground zero stage | morula frequency | n regions | conclusion |
|---|---:|---:|---:|---|
| original valid204 | morula | 0.9430 | 56 | pass |
| common DMR | morula | 0.9155 | 54 | pass |
| min_frac 0.1 | morula | NA | 120 | pass |
| min_frac 0.2 | morula | NA | 80 | pass |
| min_frac 0.3 | morula | NA | 56 | pass |
| min_frac 0.4 | morula | NA | 32 | pass |
| min_frac 0.5 | morula | NA | 32 | pass |
| balanced bootstrap n=5 | morula | 0.7675 | 56 | pass |
| balanced bootstrap n=8 | morula | 0.8590 | 56 | pass |

The common-DMR analysis used 54 regions covered across all stages. Morula remained the lowest stage under this fixed feature space:

| stage | S_epi-age |
|---|---:|
| MII oocyte | 0.4558136702 |
| zygote/PN | 0.4395800678 |
| 2-cell | 0.3503776246 |
| 4-cell | 0.2906546125 |
| 8-cell | 0.3670066159 |
| morula | 0.2814881204 |
| blastocyst | 0.3132207991 |
| ICM | 0.3282189436 |
| TE | 0.3069776664 |

Coverage-threshold sensitivity also supported the same conclusion: morula was the minimum at `min_sample_frac` values of 0.1, 0.2, 0.3, 0.4, and 0.5.

## Control Interpretation

Two controls should be interpreted conservatively:

| control | morula frequency | interpretation |
|---|---:|---|
| shuffled weights | 0.8170 | Morula remains frequently lowest after weight shuffling. This suggests that the signal is not exclusively driven by the exact age-weight ordering. |
| random age-DMR subset | 0.6980 | Morula remains frequently lowest across random subsets within the age-DMR feature space. This supports robustness within age-associated DMRs, but is not a non-age genomic random-region control. |

Therefore, the strongest defensible statement is:

> The morula-stage minimum is robust within the age-DMR feature space and across coverage/sample-resampling settings.

Avoid claiming:

> The shuffled-weight and random-subset controls prove strict age-weight specificity.

## Recommended Results Text

Age-associated methylation entropy reached its minimum at the morula stage in the original 204-sample analysis and remained lowest after restricting the analysis to DMRs commonly covered across all developmental stages. The morula minimum was also preserved across multiple minimum sample-coverage thresholds and under balanced bootstrap resampling, indicating that the observed ground-zero candidate is not explained by uneven sample size or stage-specific DMR coverage alone.

## Recommended English Sentence

The morula-stage minimum of age-associated methylation entropy was robust to common-DMR restriction, coverage-threshold variation, and balanced bootstrap resampling, supporting morula as a computational ground-zero candidate within the current age-DMR feature space.

## Files

- `tables/Experiment1B_DNA_robustness_summary.tsv`
- `tables/Experiment1B_common_DMR_stage_metrics.tsv`
- `tables/Experiment1B_coverage_threshold_sensitivity.tsv`
- `tables/Experiment1B_original_bootstrap_ground_zero_frequency.tsv`
- `tables/Experiment1B_balanced_bootstrap_n5_ground_zero_frequency.tsv`
- `tables/Experiment1B_balanced_bootstrap_n8_ground_zero_frequency.tsv`
- `figures/Experiment1B_DNA_robustness_summary.png`
- `figures/Experiment1B_DNA_robustness_summary.pdf`
