## Supplemental docs for "An entropy-based diagnostic framework for characterizing methylation state dynamics during preimplantation development": Experiment2_RNA_dual_state_interpretation.md

# Experiment 2: RNA entropy and potency-marker validation

## Purpose

Experiment 2 tests whether the DNA methylation ground-zero candidate is compatible with an active developmental state. The original working expectation was:

> low `S_epi-age` + high `S_RNA` + high potency.

The data support a more precise model:

> low `S_epi-age` + ordered RNA state + high potency-marker activity.

## Data

- Dataset: GSE36552
- Source: Yan et al. 2013 NSMB Supplementary Table 1 RPKM matrix, with GEO sample metadata.
- RNA cells retained by stage:
  - oocyte: 4
  - zygote: 4
  - 2-cell: 19
  - 4-cell: 9
  - 8-cell: 13
  - morula: 17
  - blastocyst: 39

## Main Result

Morula is not the highest global RNA entropy state. In fact, morula has the lowest mean `S_RNA` among the RNA stages used here:

| stage | S_RNA | PotencyScore |
|---|---:|---:|
| oocyte | 7.449760 | 0.380039 |
| zygote | 7.392980 | 0.414669 |
| 2-cell | 7.071784 | 0.517773 |
| 4-cell | 7.209825 | 0.402878 |
| 8-cell | 7.008319 | 0.635407 |
| morula | 6.871445 | 0.606959 |
| blastocyst | 7.011921 | 0.358131 |

However, morula retains high developmental potency-marker activity:

- Morula `PotencyScore`: 0.606959
- 8-cell `PotencyScore`: 0.635407
- Blastocyst `PotencyScore`: 0.358131

Thus, morula is close to 8-cell in potency proxy and clearly higher than blastocyst.

## Robustness and Statistics

Morula has significantly higher marker score and combined potency score than blastocyst:

| comparison | metric | mean morula | mean blastocyst | BH-adjusted p | Cliff's delta |
|---|---|---:|---:|---:|---:|
| morula vs blastocyst | marker_score | 0.823540 | 0.435908 | 4.018e-05 | 0.773756 |
| morula vs blastocyst | potency_score | 0.606959 | 0.358131 | 4.018e-05 | 0.785822 |

Morula is statistically comparable to 8-cell:

| comparison | metric | mean morula | mean 8-cell | BH-adjusted p |
|---|---|---:|---:|---:|
| morula vs 8-cell | marker_score | 0.823540 | 0.824444 | 0.261804 |
| morula vs 8-cell | potency_score | 0.606959 | 0.635407 | 0.807988 |

Bootstrap confidence intervals for combined potency score:

| stage | mean | 95% CI |
|---|---:|---:|
| 8-cell | 0.635407 | 0.596335-0.691967 |
| morula | 0.606959 | 0.539922-0.669624 |
| blastocyst | 0.358131 | 0.322994-0.396425 |

The morula and blastocyst confidence intervals are well separated, supporting a stable potency difference.

## Marker-Level Interpretation

Morula shows positive stage-level z-scores for several developmental potency markers:

- `POU5F1`: 0.562236
- `NANOG`: 0.776042
- `SOX2`: 1.222802
- `KLF4`: 0.838518
- `KLF17`: 0.813723
- `TFAP2C`: 0.666464
- `GDF3`: 0.594570

This suggests that the morula potency signal is not driven by a single marker alone.

## Fixed Conclusion

Do not write:

> morula RNA entropy is high.

Write:

> Morula represents a computational ground-zero candidate characterized by minimal age-associated methylation entropy and preserved high developmental potency-marker activity.

## Recommended Results Paragraph

RNA-seq analysis did not support a generalized increase in global transcriptomic entropy at the morula stage. Instead, morula showed the lowest mean RNA entropy among the analyzed RNA stages, suggesting a more ordered transcriptomic state. Importantly, this low RNA entropy was accompanied by high developmental potency-marker activity. Morula potency score was significantly higher than blastocyst and statistically comparable to 8-cell, with elevated activity across multiple potency markers including `POU5F1`, `NANOG`, `SOX2`, `KLF4`, `KLF17`, `TFAP2C`, and `GDF3`. Together with the DNA methylation analysis, these results support a refined dual-state model in which the morula ground-zero candidate combines minimal age-associated methylation entropy with preserved developmental potency-marker activity.

## Files

- `tables/GSE36552_RNA_entropy_potency_by_stage.tsv`
- `tables/GSE36552_potency_component_by_stage.tsv`
- `tables/GSE36552_potency_bootstrap_ci.tsv`
- `tables/GSE36552_potency_pairwise_tests.tsv`
- `tables/GSE36552_marker_score_by_stage.tsv`
- `tables/GSE36552_marker_zscore_heatmap_matrix.tsv`
- `tables/dual_entropy_stage_table.tsv`
- `figures/GSE36552_RNA_entropy_by_stage.png`
- `figures/GSE36552_potency_component_by_stage.png`
- `figures/GSE36552_potency_bootstrap_ci.png`
- `figures/GSE36552_marker_heatmap.png`
- `figures/dual_entropy_phase_map.png`
