## Supplemental docs for "An entropy-based diagnostic framework for characterizing methylation state dynamics during preimplantation development": Experiment9_age_DMR_specificity_boundary_control.md

# Experiment 9 age-DMR specificity boundary control

This control asks whether the morula minimum is exclusive to age-DMR weighting or reflects a broader methylation reprogramming minimum.

## Result

- True age-DMR weighted entropy ranks morula as the minimum.
- Generic unweighted methylation entropy on the same age-DMR regions also has a morula-associated low point.
- Shuffled weights and random age-DMR subsets still often select morula, although less strongly than the true age-weighted analysis.

## Interpretation

The result should not be phrased as pure age-DMR specificity. The stable wording is:

```text
Age-DMR weighted entropy strengthens a broader methylation reprogramming minimum at morula.
```

This supports the computational ground-zero model while avoiding an overclaim that morula minimum is exclusively caused by age-DMR identity.

## Key values

```json
{
  "age_DMR_ground_zero_stage": "morula",
  "age_DMR_morula_rank": 1,
  "age_DMR_morula_gap_to_next_non_morula": 0.03493641425126359,
  "generic_S_epi_ground_zero_stage": "morula",
  "generic_S_epi_morula_rank": 1,
  "generic_S_epi_morula_gap_to_next_non_morula": 0.016373633779299235,
  "age_DMR_true_weight_morula_frequency": 0.943,
  "generic_S_epi_bootstrap_morula_frequency": 0.845,
  "shuffled_weight_morula_frequency": 0.817,
  "random_age_DMR_subset_morula_frequency": 0.698,
  "age_vs_shuffled_frequency_ratio": 1.1542227662178703,
  "age_vs_random_subset_frequency_ratio": 1.351002865329513,
  "age_weighted_gap_gain_over_generic_S_epi": 0.018562780471964357,
  "conclusion": "age_weighting_strengthens_a_broader_morula_methylation_reprogramming_minimum",
  "claim_boundary": "The morula minimum is not exclusive proof of age-DMR specificity. It is best interpreted as age-weighted methylation entropy strengthening a broader methylation reprogramming minimum at morula."
}
```
