## Supplemental docs for "An entropy-based diagnostic framework for characterizing methylation state dynamics during preimplantation development": Experiment15_GSE56697_paired_operator_robustness.md

# Experiment 15: GSE56697 paired paternal reset operator robustness

This experiment reruns the paired paternal gamete-to-embryo methylome operator across four genomic bin sizes:

- 100 kb
- 250 kb
- 500 kb
- 1 Mb

The input is the DBA/2J sperm methylome and the outputs are paternal-allele methylomes in 2-cell, 4-cell, ICM, E6.5, and E7.5 embryos from GSE56697.

Main robustness results:

- ground-zero stage stable across bin sizes: True
- best demethylation transition stable across bin sizes: True
- ground-zero calls: ICM paternal, ICM paternal, ICM paternal, ICM paternal
- best-transition calls: sperm -> 2-cell paternal, sperm -> 2-cell paternal, sperm -> 2-cell paternal, sperm -> 2-cell paternal

Interpretation boundary:

This is a paired mouse paternal-genome reset operator validation. It supports the TRO framework as a true gamete-to-embryo operator, but it is not a human paternal-age paired embryo experiment.
