## Supplemental docs for "An entropy-based diagnostic framework for characterizing methylation state dynamics during preimplantation development": Experiment16_GSE56697_maternal_paternal_branch_contrast.md

# Experiment 16: GSE56697 maternal/oocyte versus paternal/sperm branch contrast

This experiment adds the optional parental-allele contrast requested for the paired GSE56697 operator.

It compares:

- paternal branch: sperm -> paternal allele embryo methylomes
- maternal branch: oocyte -> maternal allele embryo methylomes

The purpose is not to force both parental branches to show the same reset pattern. Instead, the analysis tests whether the paternal reset operator is a parental-allele-specific trajectory rather than an artifact of using only embryo stage averages.

| branch | gamete_input | n_common_windows | input_mean_methylation | minimum_embryo_stage | minimum_embryo_mean_methylation | best_demethylation_transition |
| --- | --- | --- | --- | --- | --- | --- |
| paternal | sperm | 3852 | 0.7918591843737672 | ICM paternal | 0.2543141280710788 | sperm -> 2-cell paternal |
| maternal | oocyte | 3989 | 0.5577612371359614 | ICM maternal | 0.2117988555542998 | 4-cell maternal -> ICM maternal |

Claim boundary: this is a mouse parental-allele methylome contrast and does not prove human paternal-age paired reset.
