## Supplemental docs for "An entropy-based diagnostic framework for characterizing methylation state dynamics during preimplantation development": Experiment19_GSE49828_gamete_to_embryo_directional_validation.md

# Experiment19: GSE49828 human gamete-to-embryo directional validation

This experiment adds human sperm RRBS methylomes to the previous GSE49828 independent DNA validation and compares
age-DMR weighted methylation entropy across sperm, MII oocyte, and early embryo stages.

This is not a strict paired parental gamete-to-embryo proof because GSE49828 does not link a specific sperm donor to a
specific embryo trajectory. It is a directional human validation of whether gamete-to-embryo development enters a low
age-associated methylation entropy window near morula or adjacent stages.

```json
{
  "dataset": "GSE49828",
  "validation_type": "human_gamete_to_embryo_directional_RRBS_age_DMR_entropy",
  "strict_pairing": false,
  "sperm_s_epi_age": 0.353853280820592,
  "morula_s_epi_age": 0.30342724336201604,
  "sperm_to_morula_delta_s_epi_age_reduction": 0.050426037458575934,
  "lowest_embryo_stage_by_s_epi_age": "MII oocyte",
  "morula_rank_among_embryo_stages": 3,
  "top3_lowest_embryo_stages": [
    "MII oocyte",
    "4-cell",
    "morula"
  ],
  "supports_morula_or_adjacent_low_age_entropy_window": true,
  "claim_boundary": "GSE49828 includes human gamete and embryo RRBS methylomes but is not a strict matched parental gamete-to-embryo dataset; use as directional human validation only."
}
```
