## Supplemental docs for "An entropy-based diagnostic framework for characterizing methylation state dynamics during preimplantation development": GSE49828_independent_DNA_validation.md

# GSE49828 independent DNA methylation validation

This analysis uses GSE49828 processed RRBS methylation-calling files as an independent DNA methylation validation set.

The validation is directional because GSE49828 is RRBS and has sparse overlap with the GSE102970 age-DMR regions.

```json
{
  "dataset": "GSE49828",
  "validation_type": "independent_human_RRBS_DNA_methylation",
  "ground_zero_stage_by_s_epi_age": "MII oocyte",
  "morula_rank_by_s_epi_age": 3,
  "top3_lowest_s_epi_age_stages": [
    "MII oocyte",
    "4-cell",
    "morula"
  ],
  "supports_morula_or_adjacent_low_age_entropy": true,
  "claim_boundary": "GSE49828 is an independent RRBS dataset with sparse age-DMR overlap; use as directional validation only."
}
```
