## Supplemental docs for "An entropy-based diagnostic framework for characterizing methylation state dynamics during preimplantation development": METHODS_REPRODUCIBILITY.md

# TRO Project Methods and Reproducibility

Date: 2026-05-21

This document records the reproducible analysis scope for the current TRO project result package. It is intended as a pre-submission methods checklist and local rerun guide.

## Project Question

The project tests whether early human preimplantation development contains a computational ground-zero candidate where age-associated epigenetic perturbation is minimized while developmental potency remains preserved.

The final operational object is:

```text
TRO = {E, D, R, C}
```

where `E` is the entropy encoder, `D` is the damage-potency decomposer, `R` is the reset operator, and `C` is the transition cost estimator.

## Data Sources

### GSE102970

Role: human sperm methylation age-associated DMR source.

Use: age-associated DMR and CpG weights were extracted from the paper supplementary Table S6. The GEO exposure matrix did not provide usable sample-level age columns for direct sperm age modeling.

Local files:

```text
metadata/GSE102970_TableS6_age_dmr_weights.tsv
metadata/GSE102970_TableS6_age_cpg_weights.tsv
```

### GSE81233

Role: primary human preimplantation embryo DNA methylation dataset.

Use: stage-level methylation entropy and age-associated methylation entropy.

Final sample count: 204 valid Cmet samples.

Excluded file:

```text
GSM2986343_scBS-2C-10-1.Cmet.bed.gz
```

Reason: repeated gzip validation failure.

### GSE36552

Role: primary human preimplantation embryo RNA dataset.

Use: global RNA entropy, detected-gene potency proxy, developmental potency-marker activity, and stage-level DNA-RNA alignment.

### GSE44183

Role: independent RNA validation dataset.

Use: external validation that 8-cell/morula occupy a high-potency region. Absolute values are not compared directly with GSE36552 because platform and normalization differ.

### GSE49828

Role: independent human RRBS methylation validation dataset.

Use: directional gamete-to-embryo validation using human sperm RRBS methylomes and human preimplantation embryo RRBS methylomes. This dataset supports a low age-DMR entropy window that includes morula/adjacent stages, but it is not a strict matched parental gamete-to-same-embryo dataset.

### GSE56697

Role: paired mouse parental methylome validation dataset.

Use: construction of a true paired-direction gamete-to-embryo methylome reset operator. The paternal branch maps DBA/2J sperm methylation to paternal-allele methylomes in 2-cell, 4-cell, ICM, E6.5, and E7.5 embryos. A maternal branch contrast maps oocyte methylation to matched maternal-allele embryo methylomes.

Local raw files:

```text
data_raw/GSE56697_parental_methylome/
```

Claim boundary: this is a mouse parental-allele methylome validation of the TRO framework. It does not prove a matched human paternal-age sperm-to-embryo reset.

## Stage Mapping

Main DNA-RNA alignment:

```text
MII oocyte -> oocyte
zygote/PN  -> zygote
2-cell     -> 2-cell
4-cell     -> 4-cell
8-cell     -> 8-cell
morula     -> morula
blastocyst -> blastocyst
```

ICM and TE are retained in DNA-only summaries but excluded from the main DNA-RNA aligned table.

## Metrics

### Generic methylation entropy

`S_epi` is binary methylation entropy. It measures methylation-state uncertainty or mixture, not aging by itself.

### Age-associated methylation entropy

`S_epi-age` is the age-associated weighted methylation entropy. This is the primary DNA perturbation metric.

### Internal reset score

`ResetScore` is normalized using:

```text
MII oocyte = 0
morula     = 1
```

This is an internal stage-normalized reset score, not a paternal young/old paired reset index.

### RNA entropy

`S_RNA` is cell-level transcriptomic Shannon entropy summarized by stage.

Important interpretation: morula is not claimed to have the highest global RNA entropy.

### Potency score

`PotencyScore` combines detected-gene signal and developmental potency-marker activity. The marker-based conclusion is tested by leave-one-marker-out robustness and expanded-marker-panel analysis.

### TRO score

```text
PotencyPreserve(g) = PotencyScore(g) / max_g PotencyScore(g)
TRO_score(g) = ResetScore(g) * PotencyPreserve(g)
```

### Ground-zero score

```text
GZ_score(g) = Z[-S_epi-age(g)] + Z[PotencyScore(g)]
```

The expected ground-zero state has low age-associated methylation entropy and preserved high potency-marker activity.

## Statistical Checks

The current result package includes:

- stage-level DNA methylation summaries;
- bootstrap ground-zero frequencies;
- adjacent-stage Mann-Whitney tests with BH correction;
- coverage threshold sensitivity;
- common-DMR analysis;
- shuffled-weight and random-age-DMR controls;
- balanced bootstrap checks;
- RNA potency pairwise tests;
- marker leave-one-out robustness;
- external RNA validation using GSE44183;
- TRO composite score, transition cost, and operator summary;
- unsupervised latent-space validation;
- parental-age residual validation using GSE273723 offspring placenta methylation;
- paired mouse paternal gamete-to-embryo methylome operator validation using GSE56697;
- GSE56697 paired paternal operator robustness across 100 kb, 250 kb, 500 kb, and 1 Mb genomic bins;
- GSE56697 maternal/oocyte versus paternal/sperm branch contrast;
- publication synthesis and g:Profiler enrichment of top reset-driving DMR nearest genes;
- mechanistic DMR-level interpretability of the TRO-defined morula ground-zero state;
- GSE49828 human gamete-to-embryo directional RRBS validation with sperm baseline;
- final claim audit.

## Current Core Results

Primary DNA result:

```text
S_epi-age reaches its minimum at morula.
```

Operational TRO result:

```text
ground_zero_stage = morula
morula_TRO_score = 0.9552284914484476
all_core_checks_pass = True
```

Transition result:

```text
8-cell -> morula is the maximum productive reset transition.
```

Paired mouse operator result:

```text
GSE56697 maps sperm -> paternal embryo allele states directly.
ICM paternal is the lowest paternal methylation state across 100 kb, 250 kb, 500 kb, and 1 Mb bins.
sperm -> 2-cell paternal is the most stable early productive demethylation transition.
```

Publication synthesis result:

```text
Human age-DMR methylation entropy identifies morula as a computational ground-zero candidate.
Paired mouse parental methylome data demonstrate that TRO can be instantiated as a true gamete-to-embryo methylome reset operator.
Top reset-driving DMR nearest genes show exploratory enrichment for developmental and signaling categories, including cadherin/WNT-related terms.
The top 20 reset-driving DMRs account for about 60.1% of positive DMR-level entropy-reset contribution, and the top 50 account for about 91.8%.
```

GSE49828 directional gamete-to-embryo result:

```text
sperm S_epi-age = 0.3539
morula S_epi-age = 0.3034
sperm -> morula S_epi-age reduction = 0.0504
morula rank among embryo stages = 3
```

## Boundaries

Do not claim:

- a matched aged father's sperm was directly reset in a matched embryo;
- `S_epi` alone is aging entropy;
- morula has the highest global RNA entropy;
- the current TRO is a trained neural operator, Koopman operator, neural operator, or Schrodinger Bridge.
- GSE273723 proves a paired sperm-to-preimplantation-embryo reset operator; it is placenta residual validation only.
- GSE56697 proves human paternal-age paired reset; it is a paired mouse parental methylome operator validation.
- GSE49828 proves strict matched parental gamete-to-embryo reset; it is directional human gamete/embryo RRBS validation only.
- g:Profiler enrichment proves a causal developmental mechanism; it is interpretability support for DMR-proximal gene categories.
- nearest-gene or genomic annotation proves that a specific DMR causally controls the morula ground-zero state; these are mechanistic leads.

Use:

- `S_epi-age` as the age-associated methylation perturbation metric;
- `TRO` as an operational, interpretable, data-driven reset operator;
- morula as a computational ground-zero candidate, not a proven universal biological zero point.
- GSE273723 as a boundary test for whether paternal-age sperm DMR CpGs leave detectable residual signal in offspring placenta.
- GSE56697 as the current paired-data instantiation of the TRO framework.
- GSE49828 as directional human gamete-to-embryo methylation support, not paired proof.

## Reproducibility Commands

For local result-package verification:

```powershell
python scripts\check_results_package.py
```

or:

```cmd
run_all_results_only.cmd
```

For full server-side reruns with raw data:

```bash
bash run_all.sh
```

The local E-drive backup is sufficient for writing, result checking, and manuscript figure/table organization. Full raw methylome reruns require the raw Cmet files or a restored raw-data cache.
