## Supplemental docs for "An entropy-based diagnostic framework for characterizing methylation state dynamics during preimplantation development": NEW_SESSION_HANDOFF.md

# CSB-TRO / Methylation Dynamics New Session Handoff

Use this file as the first message in a new conversation. The goal is to preserve the exact project state and prevent the next session from redoing old work or overclaiming the results.

## Project Directories

Main project:

```text
C:\Users\18068\Desktop\CSB_TRO_Project_2026-05-24
```

Current dynamics workspace:

```text
E:\CSB_TRO_operator_time_DMR_dynamics_2026-05-25
```

Current date/context:

```text
2026-05-25
```

## Current Scientific Position

We are doing a CSB-TRO / methylation dynamics project. The current model has advanced beyond ordinary trajectory analysis into:

```text
stage-anchored latent-state methylation operator-time dynamics
```

The model has established:

```text
1. DMR state space
2. module / latent representation
3. velocity/operator-time dynamics
4. strict leave-morula-out validation
5. autonomous latent rollout
6. stochastic diffusion attempts
7. morula basin occupancy and local stability diagnostics
8. measured missing basin-attraction correction term
9. residual DMR/module decomposition
10. scaffolds for biological-control features and augmented dynamics
```

The most accurate current conclusion is:

```text
We have established a stage-anchored latent-state methylation operator-time model that improves mean-state prediction of morula reset and reveals a compact, directional missing basin-attraction correction term. However, uncalibrated methylation-only stochastic dynamics does not autonomously generate the morula population basin. The residual DMR/module results are diagnostic control coordinates, not yet proven biological drivers. The next task is to identify real external biological control variables u_bio and implement a biologically interpretable control term.
```

Chinese concise version:

```text
The remaining question is no longer whether dynamics exist, but what the biological identity of u_bio is. Mathematically, the missing basin-attraction correction term has been estimated and decomposed into a small number of residual DMRs/modules. These modules remain diagnostic coordinates rather than established causal biological drivers. The next step is to explain this correction term using external gene, motif, RNA, ATAC, histone or related evidence.
```

## Core Data

DMR state matrix:

```text
E:\CSB_TRO_operator_time_DMR_dynamics_2026-05-25\results\CSB_TRO_DMR_state_matrix.tsv
```

Size:

```text
169 samples x 156 age-DMRs
```

Sample tau annotation:

```text
E:\CSB_TRO_operator_time_DMR_dynamics_2026-05-25\results\CSB_TRO_sample_tau_annotation.tsv
```

Stages and tau:

```text
MII oocyte = 0
zygote/PN = 0.1667
2-cell = 0.3333
4-cell = 0.5
8-cell = 0.6667
morula = 0.8333
blastocyst = 1
```

OT sample transition couplings:

```text
E:\CSB_TRO_operator_time_DMR_dynamics_2026-05-25\results\CSB_TRO_OT_sample_transition_couplings.tsv
```

DMR-level transition training pairs:

```text
E:\CSB_TRO_operator_time_DMR_dynamics_2026-05-25\results\CSB_TRO_OT_transition_training_pairs.tsv
```

## Completed Results

### 1. Single-DMR Ridge Velocity

Script:

```text
E:\CSB_TRO_operator_time_DMR_dynamics_2026-05-25\code\run_dmr_operator_time_dynamics.py
```

Result:

```text
single-DMR leave-morula-out RMSE = 0.3113
8-cell baseline RMSE = 0.2974
```

Conclusion:

```text
Single-DMR independent ridge does not solve strict morula extrapolation.
Morula reset is not explained by independent linear DMR-wise changes.
```

### 2. Module / Latent Dynamics

Script:

```text
E:\CSB_TRO_operator_time_DMR_dynamics_2026-05-25\code\run_module_latent_operator_time_dynamics.py
```

Result:

```text
8-cell baseline RMSE = 0.2974
single-DMR ridge RMSE = 0.3113
DMR-module ridge RMSE = 0.2866
latent PCA ridge RMSE = 0.2698
```

Conclusion:

```text
Module/latent representation improves strict leave-morula-out prediction relative to both single-DMR ridge and 8-cell baseline.
Morula reset is more consistent with a coordinated module/latent transition than independent DMR drift.
```

### 3. Strict Validation

Script:

```text
E:\CSB_TRO_operator_time_DMR_dynamics_2026-05-25\code\run_module_latent_validation.py
```

Run:

```text
python "E:\CSB_TRO_operator_time_DMR_dynamics_2026-05-25\code\run_module_latent_validation.py" --n-null 200 --n-bootstrap 1000 --seed 20260525
```

Key outputs:

```text
results\CSB_TRO_module_latent_validation_summary.tsv
results\CSB_TRO_module_latent_bootstrap_CI.tsv
results\CSB_TRO_module_latent_null_models.tsv
results\CSB_TRO_module_latent_paired_error_tests.tsv
figures\CSB_TRO_module_latent_validation_rmse.svg
docs\CSB_TRO_module_latent_validation_interpretation.md
```

Key results:

```text
latent DMR bootstrap delta RMSE 95% CI = [-0.0486, -0.0046]
latent stage/sample bootstrap delta RMSE 95% CI = [-0.0351, -0.0217]
latent paired squared-error p = 0.0067
module paired squared-error p = 0.0381
module random tau null p = 0.0050
module random module null p = 0.0299
```

Boundary:

```text
Random coupling null is not clearly worse. Do not claim sample-level OT coupling itself is the key mechanism.
Safe wording: latent/module state representation improves strict morula prediction; coupling-specific mechanism remains unresolved.
```

### 4. Advanced Latent Dynamics

Script:

```text
E:\CSB_TRO_operator_time_DMR_dynamics_2026-05-25\code\run_advanced_latent_dynamics.py
```

Run:

```text
python "E:\CSB_TRO_operator_time_DMR_dynamics_2026-05-25\code\run_advanced_latent_dynamics.py" --q 3 --lambda 1000 --stochastic-n 20 --seed 20260525
```

Key results:

```text
full_all_transitions 8-cell -> morula DMR mean RMSE = 0.2471, correlation = 0.6203
leave_morula_transitions 8-cell -> morula DMR mean RMSE = 0.2517, correlation = 0.5982
early_to_4cell_transitions 8-cell -> morula DMR mean RMSE = 0.2597, correlation = 0.5792
```

Interpretation:

```text
Autonomous latent rollout recovers DMR mean trajectory reasonably well.
But basin occupancy is not reproduced by initial residual diffusion.
Mean-state dynamics is working; autonomous distribution-generating dynamics is not solved.
```

Jacobian:

```text
Eigenvalue real parts all negative:
-3.6225
-5.8052 +/- 0.6561i
```

Perturbation:

```text
PC1 increase 1sd at 8-cell raises morula RMSE by +0.0260
remove top50 latent-loading DMRs raises RMSE by +0.0115
```

### 5. Morula Basin SDE

Script:

```text
E:\CSB_TRO_operator_time_DMR_dynamics_2026-05-25\code\run_morula_basin_sde.py
```

Run:

```text
python "E:\CSB_TRO_operator_time_DMR_dynamics_2026-05-25\code\run_morula_basin_sde.py" --q 3 --lambda 1000 --particles-per-start 200 --calibration-particles-per-start 40 --n-steps 12 --seed 20260525
```

Key result:

```text
observed morula q90 occupancy = 0.875
full_all deterministic occupancy = 0.378
full_all global residual SDE occupancy = 0.027
full_all stage-conditioned SDE occupancy = 0.161
full_all empirical 8-cell -> morula full-fit noise occupancy = 0.253
full_all basin OU calibrated occupancy = 0.892

leave_morula deterministic occupancy = 0.044
leave_morula global residual SDE occupancy = 0.024
leave_morula stage-conditioned SDE occupancy = 0.085
leave_morula basin OU calibrated occupancy = 0.886
```

Interpretation:

```text
Strict uncalibrated methylation-only SDE does not naturally generate morula basin occupancy.
Calibrated basin OU can match occupancy but uses morula information, so it is a diagnostic/upper-bound control, not strict extrapolation.
```

### 6. Non-Leaking Distribution Dynamics

Script:

```text
E:\CSB_TRO_operator_time_DMR_dynamics_2026-05-25\code\run_nonleaking_distribution_dynamics.py
```

Run:

```text
python "E:\CSB_TRO_operator_time_DMR_dynamics_2026-05-25\code\run_nonleaking_distribution_dynamics.py" --q 3 --lambda 1000 --particles-per-start 200 --validation-particles 80 --n-steps 12 --seed 20260525
```

Key results:

```text
pre_morula_velocity_deterministic occupancy = 0.044
residual SDE occupancy = 0.023
affine Gaussian transition kernel occupancy = 0.016
moment extrapolated Gaussian occupancy = 0.000
validation-tuned extrapolated OU occupancy = 0.000
using blastocyst future but no morula interpolated OU occupancy = 0.376
morula-calibrated upper bound occupancy = 0.848
```

Centers:

```text
extrapolated pre-morula center = [1.4630, -2.4973, -8.4225]
observed morula center = [-3.1743, -0.0308, -2.8184]
```

Conclusion:

```text
Pre-morula methylation-only dynamics cannot locate the morula basin distribution.
This motivates hidden biological/regulatory/chromatin control variables.
```

### 7. Basin Residual Control Field

Script:

```text
E:\CSB_TRO_operator_time_DMR_dynamics_2026-05-25\code\run_basin_residual_control_field.py
```

Run:

```text
python "E:\CSB_TRO_operator_time_DMR_dynamics_2026-05-25\code\run_basin_residual_control_field.py" --q 3 --lambda 1000 --n-steps 12 --top-bed 100 --seed 20260525
```

Key correction:

```text
observed morula center = [-3.1743, -0.0308, -2.8184]
strict pre-morula predicted center = [-2.3829, 0.0031, -0.9009]
missing basin residual = [-0.7914, -0.0338, -1.9175]
residual norm = 2.0747
```

Diagnostic controls:

```text
strict occupancy = 0.044
observed occupancy = 0.875
latent oracle alpha=1 occupancy = 1.000
top10 DMR restricted residual control occupancy = 0.600
top25 DMR occupancy = 0.956
top50/top100 DMR occupancy = 1.000
```

Top residual DMR examples:

```text
cluster_2623 M05 chr15:37190548-37190694
cluster_6262 M10 chr6:31628935-31629199
cluster_6892 M05 chr7:37947051-37947061
cluster_5678 M10 chr4:186425697-186425754
cluster_3275 M05 chr17:10421635-10422077
```

Module residual fractions:

```text
M01 23.5%
M05 15.9%
M10 8.0%
M12 6.2%
M07 6.1%
M06 6.1%
M02 6.1%
M08 5.1%
```

Boundary:

```text
These are diagnostic residual control coordinates. Do not call them proven causal biological drivers yet.
```

### 8. Residual DMR Robustness

Script:

```text
E:\CSB_TRO_operator_time_DMR_dynamics_2026-05-25\code\run_residual_dmr_robustness.py
```

Key outputs:

```text
results\CSB_TRO_residual_DMR_topK_occupancy_curve.tsv
results\CSB_TRO_residual_DMR_bootstrap_stability.tsv
results\CSB_TRO_residual_DMR_matched_random_control.tsv
results\CSB_TRO_residual_DMR_sign_flip_control.tsv
results\CSB_TRO_residual_DMR_remove_topK_necessity.tsv
results\CSB_TRO_residual_module_add_remove.tsv
figures\CSB_TRO_residual_DMR_topK_occupancy.svg
docs\CSB_TRO_residual_DMR_robustness_summary.md
```

Key results:

```text
top10 forward occupancy = 0.600
top15 forward occupancy = 0.867
top25 forward occupancy = 0.956
top50 forward occupancy = 1.000

sign-flip top10/top15/top25/top50 occupancy = 0.000

remove top10 keep remainder occupancy = 0.889
remove top15 keep remainder occupancy = 0.822
remove top25 keep remainder occupancy = 0.556
remove top50 keep remainder occupancy = 0.178

matched random top25 mean occupancy = 0.1084
matched random top25 q95 = 0.1556
matched random top25 max = 0.200
```

Interpretation:

```text
Top residual DMRs are compact, directional, stable, and stronger than matched random controls.
But top25 is a core control set, not the entire control signal.
Remaining DMRs still contain distributed auxiliary residual information.
```

### 9. Biological Annotation Scaffold

Script:

```text
E:\CSB_TRO_operator_time_DMR_dynamics_2026-05-25\code\run_residual_dmr_biological_annotation.py
```

Run:

```text
python "E:\CSB_TRO_operator_time_DMR_dynamics_2026-05-25\code\run_residual_dmr_biological_annotation.py" --top-n 100
```

Outputs:

```text
results\CSB_TRO_top_residual_DMR_annotation.tsv
results\CSB_TRO_residual_DMR_nearest_genes.tsv
results\CSB_TRO_residual_DMR_GO_KEGG.tsv
results\CSB_TRO_residual_DMR_motif_enrichment.tsv
results\CSB_TRO_residual_DMR_ATAC_overlap.tsv
results\CSB_TRO_residual_DMR_histone_overlap.tsv
results\CSB_TRO_residual_DMR_RNA_validation.tsv
docs\CSB_TRO_residual_biology_interpretation.md
```

Current status:

```text
External annotation inputs are missing locally.
Outputs correctly report not_run_missing_external_input.
No real biological mechanism is identified yet.
```

### 10. Missing Control Term Decomposition

Script:

```text
E:\CSB_TRO_operator_time_DMR_dynamics_2026-05-25\code\run_missing_control_term_decomposition.py
```

Run:

```text
python "E:\CSB_TRO_operator_time_DMR_dynamics_2026-05-25\code\run_missing_control_term_decomposition.py" --q 3 --lambda 1000 --n-steps 12 --seed 20260525
```

Key outputs:

```text
results\CSB_TRO_missing_control_term_summary.tsv
results\CSB_TRO_missing_control_term_amplitude_scan.tsv
results\CSB_TRO_missing_control_term_module_basis.tsv
results\CSB_TRO_missing_control_term_greedy_modules.tsv
results\CSB_TRO_missing_control_term_reconstruction_metrics.tsv
figures\CSB_TRO_missing_control_term_amplitude_scan.svg
docs\CSB_TRO_missing_control_term_interpretation.md
```

Core result:

```text
missing control delta_z = [-0.7914, -0.0338, -1.9175]
equivalent correction velocity = [-4.7485, -0.2031, -11.5051]
```

Interpretation:

```text
Strong PC3-negative pull
Moderate PC1-negative pull
Weak PC2 correction
```

Amplitude scan:

```text
alpha=0 occupancy = 0.044
alpha=0.1 occupancy = 0.133
alpha=0.25 occupancy = 0.422
alpha=0.5 occupancy = 0.933
alpha=0.75 occupancy = 0.978
alpha=0.875/1.0 occupancy = 1.000
```

Greedy module accumulation:

```text
M05 occupancy = 0.422
M05 + M01 occupancy = 0.600
M05 + M01 + M12 occupancy = 0.867
M05 + M01 + M12 + M02 occupancy = 0.956
M05 + M01 + M12 + M02 + M10 occupancy = 0.956
M05 + M01 + M12 + M02 + M10 + M09 occupancy = 0.956
M05 + M01 + M12 + M02 + M10 + M09 + M08 occupancy = 0.978
M05 + M01 + M12 + M02 + M10 + M09 + M08 + M06 occupancy = 1.000
```

Key distinction:

```text
We have measured B u_bio, but we have not identified true u_bio.
```

### 11. Biological Control Feature Builder

Script:

```text
E:\CSB_TRO_operator_time_DMR_dynamics_2026-05-25\code\run_build_biological_control_features.py
```

Run:

```text
python "E:\CSB_TRO_operator_time_DMR_dynamics_2026-05-25\code\run_build_biological_control_features.py"
```

Outputs:

```text
results\CSB_TRO_module_bio_features.tsv
results\CSB_TRO_module_gene_links.tsv
results\CSB_TRO_module_RNA_activity.tsv
results\CSB_TRO_module_ATAC_activity.tsv
results\CSB_TRO_module_histone_activity.tsv
results\CSB_TRO_module_motif_activity.tsv
docs\CSB_TRO_module_bio_features_interpretation.md
```

Current features:

```text
n_features = 48
modalities:
- internal_genomic_proxy
- internal_methylation_prior_proxy
- measured_residual_diagnostic
```

Important:

```text
No real RNA/ATAC/histone/motif features are available yet because external input files are missing.
This is a scaffold/dry-run, not real u_bio identification.
```

Optional inputs supported:

```text
--gene-links
--rna-matrix
--atac-features
--histone-features
--motif-features
```

### 12. Biological Control Augmented Dynamics

Script:

```text
E:\CSB_TRO_operator_time_DMR_dynamics_2026-05-25\code\run_biological_control_augmented_dynamics.py
```

Run:

```text
python "E:\CSB_TRO_operator_time_DMR_dynamics_2026-05-25\code\run_biological_control_augmented_dynamics.py" --n-random 200 --seed 20260525
```

Outputs:

```text
results\CSB_TRO_bio_control_occupancy_metrics.tsv
results\CSB_TRO_bio_control_direction_alignment.tsv
results\CSB_TRO_biological_control_coefficients.tsv
results\CSB_TRO_bio_control_matched_random_features.tsv
figures\CSB_TRO_bio_control_occupancy.svg
docs\CSB_TRO_bio_control_interpretation.md
```

Dry-run results:

```text
methylation-only strict baseline occupancy = 0.044
measured_missing_correction_upper_bound occupancy = 1.000
internal_genomic_proxy_unit_beta occupancy = 0.089
internal_methylation_prior_proxy_unit_beta occupancy = 0.156
measured_residual_diagnostic_unit_beta occupancy = 0.978
```

Interpretation:

```text
The augmented-control framework works technically.
But current internal proxies are not real external u_bio.
Ridge diagnostic models can reconstruct the correction only because they use the measured morula residual for beta; they are not valid non-leaking biological-control models.
```

## Current Main Problem

The current blocker is not:

```text
more SDE complexity
more latent dimensions
more OU tuning
more particle count
```

The blocker is:

```text
We know the missing correction force B u_bio, but we do not know the real biological variable u_bio.
```

In plain terms:

```text
Mathematically, the missing force, its direction and magnitude, and the modules on which it acts have been identified.
Biologically, it remains unresolved whether this force arises from RNA, ATAC, histone regulation, TF motifs, DNMT/TET activity, the ZGA programme or chromatin remodelling.
```

## Correct Next Strategy

Do not continue expanding the SDE first. The next phase is:

```text
measured correction term
-> residual modules
-> gene / regulatory annotation
-> motif / RNA / ATAC / histone candidate controls
-> module-level u_m(tau)
-> biologically interpretable control term
-> occupancy rescue and direction alignment
```

Recommended priority:

```text
1. Lock genome build and coordinate convention.
2. Add gene annotation / TSS / GTF or GFF3.
3. Annotate M05/M01/M12/M02/M10 DMRs to genes, promoters, introns, exons, intergenic regions.
4. Add CpG island / shore / shelf, repeat, enhancer annotations if available.
5. Run motif scan/enrichment for residual DMRs/modules.
6. Add RNA expression by stage.
7. Build motif x TF expression activity:
   u_TF,module = motif_enrichment_TF,module * Delta TF expression
8. Add ATAC / histone features if available.
9. Run biological-control feature builder with real inputs.
10. Run biological-control augmented dynamics.
```

Model form:

```text
dz_meth/dtau = f_meth(z,tau) + B u_bio(tau)
```

Interpretable module form:

```text
B u_bio(tau) = sum_m beta_m * u_m(tau) * b_m
```

where:

```text
m = residual module, e.g. M05, M01, M12, M02, M10
u_m(tau) = external biological activity score
b_m = module direction in methylation latent space
beta_m = control strength
```

Candidate u_m:

```text
u_RNA,m
u_ATAC,m
u_H3K27ac,m
u_H3K4me3,m
u_H3K27me3,m
u_motif,m
u_TF,m = motif enrichment x TF expression change
```

## Evaluation Criteria for True Candidate u_bio

A real candidate biological control should pass:

```text
1. It does not use morula methylation center / radius / occupancy to define itself.
2. It is linked to residual modules such as M05/M01/M12/M02/M10.
3. It changes around 8-cell -> morula or morula -> blastocyst.
4. Its predicted control direction aligns with measured correction:
   strong PC3-negative pull
   moderate PC1-negative pull
   weak PC2 correction
5. It improves morula occupancy over methylation-only baseline.
6. It reduces center error and improves cosine alignment.
7. Sign-flip control fails.
8. Matched random module/feature controls are weaker.
```

Important metrics:

```text
occupancy rescue
center error reduction
cosine alignment with measured correction
PC3-negative pull recovery
matched random control
sign-flip control
```

Occupancy reference:

```text
methylation-only strict occupancy = 0.044
observed occupancy = 0.875
measured correction upper bound = about 0.956 to 1.000
```

Success target:

```text
If real external biological controls raise occupancy from 0.044 to 0.3-0.6+, while also improving direction alignment and passing controls, that is already strong evidence.
```

## Non-Leaking Levels

Level 1:

```text
methylation-non-leaking
Do not use morula methylation center/radius/occupancy.
May use morula-stage RNA/ATAC/histone as external biological signal.
This tests whether external omics can explain morula methylation correction.
```

Level 2:

```text
fully pre-morula non-leaking
Do not use morula methylation or morula external omics.
Use only pre-morula information to predict morula correction.
This is stronger but much harder and should not be the first target.
```

## Things Not To Overclaim

Do not say:

```text
We have a complete stochastic population dynamics model.
The model autonomously reproduces morula basin.
Top residual DMRs are proven causal drivers.
OT couplings are lineage tracking.
Calibrated basin OU is strict extrapolation.
```

Safe wording:

```text
Latent/module methylation dynamics improves mean-state morula prediction.
Strict methylation-only stochastic dynamics does not autonomously reproduce morula basin occupancy.
Measured residual correction localizes the missing basin-attraction direction to compact, directional, non-random DMR/module coordinates.
These residual modules nominate candidate regulatory/chromatin mechanisms, but real u_bio remains to be identified using external gene, motif, RNA, ATAC, and histone data.
```

## Immediate Next Action For New Session

Start by checking whether external biological input files are available locally. Look for:

```text
GTF/GFF3 or gene TSS table
genome build information
CpG island BED
repeat BED
enhancer BED
motif scan/enrichment table
RNA expression matrix by stage
ATAC peak/signal table by stage
histone mark peak/signal tables by stage
```

If these files are not available, create a manifest/template for required external inputs, then help locate/download or prepare stage-matched public annotation/omics resources.

Once inputs exist, run:

```text
python "E:\CSB_TRO_operator_time_DMR_dynamics_2026-05-25\code\run_residual_dmr_biological_annotation.py" [with external inputs]

python "E:\CSB_TRO_operator_time_DMR_dynamics_2026-05-25\code\run_build_biological_control_features.py" [with external inputs]

python "E:\CSB_TRO_operator_time_DMR_dynamics_2026-05-25\code\run_biological_control_augmented_dynamics.py" --n-random 200 --seed 20260525
```

The next scientific deliverable should be:

```text
CSB_TRO_real_u_bio_input_manifest.tsv
CSB_TRO_residual_module_gene_annotation.tsv
CSB_TRO_residual_module_motif_RNA_activity.tsv
CSB_TRO_biological_control_external_metrics.tsv
CSB_TRO_biologically_interpretable_control_term.md
```

## One-Sentence Handoff

```text
Continue from the completed stage-anchored latent methylation dynamics: mean-state prediction works, strict methylation-only distribution dynamics fails to generate morula basin, the missing correction term is measured as delta_z=[-0.7914,-0.0338,-1.9175] and decomposes mainly through M05/M01/M12/M02/M10 residual modules; now identify true external biological u_bio from gene/motif/RNA/ATAC/histone inputs and test whether an interpretable control term sum beta_m u_m b_m can rescue morula occupancy without using morula methylation.
```
