## Supplemental docs for "An entropy-based diagnostic framework for characterizing methylation state dynamics during preimplantation development": README_DOCS.md

# CSB-TRO Docs

This folder mirrors the most important human-readable project documents.

Start with:

1. README_PROJECT.md
2. CSB_TRO_dynamics_master_record.md

The full machine-readable outputs are in ../results.
The current scripts are in ../code_current.
The copied input tables are in ../input_tables.
