## Supplemental docs for "An entropy-based diagnostic framework for characterizing methylation state dynamics during preimplantation development": README_PROJECT.md

# CSB-TRO Project Snapshot

Date: 2026-05-24

This folder is a standalone CSB-TRO project snapshot moved out of the unrelated antimicrobial peptide workspace.

## Directory Layout

- `code_current/`
  Current CSB-TRO scripts after the anti-circularity fix and robustness upgrades.

- `code_original/`
  Earlier pilot scripts from the archived `AMP Discover Platform v1.0.0` installer directory.

- `results/`
  Full CSB-TRO result directory copied from:
  `<ARCHIVED_RESULTS_ROOT>\csb_tro_dynamics`

- `input_tables/`
  Key local source tables used by the model, including GSE81233 summaries, GSE36552 RNA potency/entropy tables, GSE44183 external RNA validation, DMR contribution tables, and GSE81233 sample genome metrics.

- `savepoints/`
  Previous savepoint made before this project migration.

- `docs/`
  Reserved for manuscript notes and future documentation.

## Current Model Status

The current CSB-TRO version uses a stage-agnostic potency threshold:

`P_min = quantile(P over all fused particles, q=0.60)`

This avoids the main circularity risk: morula is not used to set the training/transport potency threshold. Morula is evaluated post hoc.

Key current results:

- `J_path_total = 3.101242`
- Morula A rank = 1
- Morula P rank = 2
- Morula reset score rank = 1
- 8-cell -> morula A transport remains negative
- DMR graph Laplacian uses 156 DMR nodes and 1102 weighted edges
- Time/reference sensitivity is stable across 9 settings
- Soft-marginal uncertainty audit is included
- Fokker-Planck export now includes full diffusion matrix `Sigma(t)`
- Static-vs-dynamic gain analysis is included
- Prediction validation is included; the strongest support is held-out DMR split validation

## Most Important Files

- `results/CSB_TRO_dynamics_master_record.md`
- `results/CSB_TRO_path_space_summary.json`
- `results/CSB_TRO_fused_robustness_summary.json`
- `results/CSB_TRO_DMR_graph_laplacian_summary.json`
- `results/CSB_TRO_fokker_planck_summary.json`
- `results/CSB_TRO_time_reference_sensitivity_summary.json`
- `results/CSB_TRO_soft_marginal_uncertainty_summary.json`
- `results/CSB_TRO_static_dynamic_gain_summary.json`
- `results/CSB_TRO_prediction_validation_summary.json`

## Raw Data Note

Large original raw Cmet files were not duplicated into this snapshot. Their original local path remains:

`<ARCHIVED_RESULTS_ROOT>\processed\GSE81233_strong_controls\raw_cmet_cache`

This project folder contains derived metrics and result artifacts needed to inspect and reproduce the current CSB-TRO analysis layer.
