## Supplemental docs for "An entropy-based diagnostic framework for characterizing methylation state dynamics during preimplantation development": reproducibility_notes.md

# Reproducibility notes

Date: 2026-05-21

## What this package can reproduce directly

The local synchronized result package can verify final outputs and consistency checks using:

```bash
bash run_all_results_only.sh
```

On Windows CMD, use:

```cmd
run_all_results_only.cmd
```

This checks:

- required result tables exist;
- required figures exist;
- `TRO_operator_summary.json` reports morula as the ground-zero stage;
- all core TRO checks pass;
- morula ranks first by GZ score, TRO score, and BioAgeRank.

## What requires the server project directory

Full reruns require:

```text
/root/autodl-tmp/TRO_Project
```

because large files are not stored in the local result package:

- raw GSE81233 Cmet files;
- intermediate methylation matrices;
- per-sample DMR coverage files;
- cached genome-wide matched-control scans.

Use:

```bash
bash run_all.sh
```

from the server project root.

## Recommended computational environment

Create environment:

```bash
conda env create -f environment.yml
conda activate tro-project
```

Minimum Python packages:

```text
python >= 3.11
numpy
pandas
scipy
matplotlib
openpyxl
requests
```

## Main reproducibility caveats

1. GSE81233 contains one excluded corrupted file:

```text
GSM2986343_scBS-2C-10-1.Cmet.bed.gz
```

It repeatedly failed gzip validation.

2. GSE102970 was not modeled using sample-level age from GEO because the GEO exposure matrix did not provide usable age columns. The project uses Table S6 age-associated DMR weights instead.

3. RNA and DNA datasets are not matched embryos. DNA-RNA integration is stage-level, not paired-sample integration.

4. `S_epi-age` is an age-associated perturbation metric, not a direct epigenetic clock age.

5. Experiment 7 matched non-age-DMR specificity control remains pilot/feasibility unless run with adequate stage coverage and sufficient matched control sets.

## Current final result

The operational TRO result is stored in:

```text
tables/TRO_operator_summary.json
tables/TRO_operator_stage_output.tsv
tables/TRO_operator_transition_output.tsv
```

Current final checks:

```text
ground_zero_stage = morula
GZ_rank(morula) = 1
TRO_rank(morula) = 1
BioAgeRank(morula) = 1
all_core_checks_pass = true
```
