## Supplemental docs for "An entropy-based diagnostic framework for characterizing methylation state dynamics during preimplantation development": TRO_final_results_interpretation_2026-05-20.md

# TRO_Project final results interpretation

Date: 2026-05-20

## Current conclusion

The current public-data analysis supports an operational Transgenerational Reset Operator (TRO) framework in which the morula stage is identified as a computational ground-zero candidate.

The strongest final statement is:

> Morula represents a computational ground-zero candidate characterized by minimal age-associated methylation entropy, preserved high developmental potency-marker activity, and the highest operational TRO score among preimplantation stages.

## Experiment 1: DNA methylation age-entropy reset

Main data:

- GSE102970: sperm age-associated DMR / CpG weights from supplementary Table S6.
- GSE81233: human preimplantation embryo DNA methylation, 204 valid samples after excluding one repeatedly corrupted file.

Key result:

- S_epi-age reaches its minimum at morula: 0.2777817165440405.
- Bootstrap ground-zero frequency for morula: 1886 / 2000 = 94.3%.
- Significant adjacent changes support a reset-like trajectory:
  - 2-cell vs 4-cell: BH p = 0.0091.
  - 8-cell vs morula: BH p = 0.0379.
  - morula vs blastocyst: BH p = 0.0379.

Interpretation:

Age-associated sperm methylation entropy is progressively reduced during preimplantation development and reaches a robust minimum at morula.

## Experiment 1B: DNA robustness

Purpose:

To test whether the morula minimum is caused by coverage, sample imbalance, DMR selection, or weight artifacts.

Key result:

- Common DMR analysis: morula remains the ground-zero stage, bootstrap frequency 0.9155.
- Coverage threshold sensitivity: morula remains the minimum across tested thresholds.
- Balanced bootstrap:
  - n=5: morula frequency 0.7675.
  - n=8: morula frequency 0.8590.
- Shuffled-weight and random-DMR controls reduce the signal, indicating that age-DMR weighting contributes specific information.

Interpretation:

The morula minimum is not a simple coverage or sample-size artifact.

## Experiment 2: RNA entropy and potency

Main data:

- GSE36552: human preimplantation embryo RNA-seq.

Important correction to the original expectation:

Morula does not show globally high RNA entropy. Therefore the correct model is not "morula has highest transcriptome entropy."

The supported model is:

> low age-associated methylation entropy + preserved high potency-marker activity.

Key result:

- Morula potency score: 0.606959.
- 8-cell potency score: 0.635407.
- Blastocyst potency score: 0.358131.
- Morula vs blastocyst:
  - marker score BH p = 4.018e-05.
  - potency score BH p = 4.018e-05.
- Morula vs 8-cell is not significantly different, so the correct statement is that 8-cell and morula form a high-potency region.

Interpretation:

RNA analysis supports a high developmental potency-marker state around 8-cell/morula, followed by potency decline at blastocyst.

## Experiment 3: TRO composite score and marker robustness

Definitions:

- GZ_score = Z[-S_epi-age] + Z[PotencyScore].
- TRO_score = ResetScore x PotencyPreserve.

Key result:

- Morula GZ_rank = 1.
- Morula TRO_rank = 1.
- Morula TRO_score = 0.955228.
- Leave-one-marker-out robustness: all recomputed marker panels still support morula > blastocyst.
- Maximum leave-one-marker-out adjusted p value: 0.0002176868722182.

Interpretation:

The morula ground-zero result does not depend on one single potency marker.

## Experiment 4: External RNA validation

Main data:

- GSE44183 human expression matrix.

Key result:

- Top potency stages:
  - 8-cell rank 1.
  - morula rank 2.
- The external dataset does not prove morula is higher than 8-cell, but it supports that 8-cell/morula occupy a high-potency region.

Interpretation:

External RNA validation supports the high-potency-region conclusion, not a strict morula-only maximum.

## Experiment 5: Transition cost and reset dynamics

Key result:

- The maximum productive reset transition is 8-cell -> morula.
- 8-cell -> morula:
  - damage reduction = 0.105261.
  - reset gain = 0.624380.
  - productive reset gain = 0.729641.
  - reset efficiency = 0.271327, rank 1.
- Morula -> blastocyst:
  - productive reset gain = 0.
  - reset efficiency = 0.
  - interpreted as differentiation/nonproductive transition.
- MII -> morula relative S_epi-age reduction = 37.77%.

Interpretation:

The transition analysis places the strongest productive reset window immediately before morula.

## Experiment 6: Operational TRO operator

The final operational TRO is represented as:

```text
TRO = {E, D, R, C}
```

Where:

- E = Entropy Encoder.
- D = Damage-Potency Decomposer.
- R = Reset Operator.
- C = Cost Estimator.

Final checks:

- GZ_score_morula_rank_1 = true.
- TRO_score_morula_rank_1 = true.
- BioAgeScore_morula_rank_1 = true.
- best_transition_is_8cell_to_morula = true.
- DNA_robustness_common_DMR = true.
- DNA_robustness_balanced_bootstrap = true.
- RNA_potency_morula_gt_blastocyst = true.
- marker_leave_one_out_all_pass = true.
- external_GSE44183_morula_top2 = true.
- all_core_checks_pass = true.

Interpretation:

The project now has a data-driven operational TRO operator, not only a conceptual prototype.

## Claims that are currently supported

Supported:

1. Morula is a computational ground-zero candidate under the current public-data TRO framework.
2. Age-associated methylation entropy is minimized at morula.
3. The morula minimum is robust to common-DMR, coverage, and balanced-bootstrap analyses.
4. Morula retains high developmental potency-marker activity relative to blastocyst.
5. 8-cell/morula form a high-potency region in RNA data.
6. The strongest productive reset transition is 8-cell -> morula.
7. The operational TRO operator ranks morula first.

## Claims that should not be made

Do not claim:

1. Public data prove that one aged father's sperm was reset in one matched embryo.
2. S_epi is aging entropy. S_epi is generic methylation entropy.
3. S_epi-age is a direct biological age clock. It is an age-associated methylation perturbation metric.
4. Morula has the highest global RNA entropy.
5. External RNA validation proves morula is uniquely higher than 8-cell in potency.
6. The current TRO is a trained neural operator or Schrödinger Bridge model. It is an operational, interpretable, data-driven operator.
7. The pilot matched non-age-DMR control proves or disproves age-DMR specificity. The current matched-control run was a feasibility pilot with sparse stage coverage and should not be used as a final scientific claim.

## Experiment 7 pilot status

Experiment 7 was initiated to test age-DMR specificity against matched non-age-DMR genomic windows. The pilot confirmed that the pipeline can generate matched control regions, rescan Cmet files, and produce stage-level summaries. However, the pilot used only one sample per stage and showed sparse coverage in some stages. Therefore it is retained as a feasibility analysis only.

Current interpretation:

> The final manuscript should not rely on Experiment 7 for the main claim unless the matched-control analysis is rerun with adequate sample coverage and multiple matched control sets.

## Recommended manuscript framing

Result 1:

Age-associated methylation entropy identifies morula as a robust computational ground-zero candidate.

Result 2:

Morula preserves high developmental potency-marker activity despite not exhibiting elevated global RNA entropy.

Result 3:

The dual damage-potency phase map separates early reset, ground-zero, and post-morula differentiation states.

Result 4:

An operational TRO operator ranks morula as the optimal reset state and identifies 8-cell -> morula as the maximum productive reset transition.

Final one-sentence abstract conclusion:

> Together, these analyses define an operational Transgenerational Reset Operator in which age-associated methylation perturbation is minimized at morula while developmental potency-marker activity remains high, identifying morula as a computational ground-zero candidate during human preimplantation development.
