## Supplemental docs for "An entropy-based diagnostic framework for characterizing methylation state dynamics during preimplantation development": TRO_interpretability_analysis.md

# TRO interpretability analysis

This analysis explains why morula is selected as the computational ground-zero candidate.

## DMR contribution

For each age-DMR, the main reset-driving contribution was computed as:

```text
abs(age_weight) * (H_8-cell - H_morula)
```

Positive values indicate age-DMR entropy reduction from 8-cell to morula.

Main output:

```text
tables/TRO_interpretability_DMR_contribution_ranking.tsv
tables/TRO_interpretability_top20_reset_driving_DMRs.tsv
tables/TRO_interpretability_top50_reset_driving_DMRs.tsv
```

## Gene and region annotation

DMRs were mapped to hg19 RefGene nearest genes and annotated as promoter, exon, intron, or intergenic, with CpG island/shore/shelf context.

Main output:

```text
tables/TRO_interpretability_DMR_annotation_enrichment.tsv
tables/TRO_interpretability_reset_DMR_gene_function_enrichment.tsv
```

## Potency marker contribution

Morula marker contribution was estimated from positive morula marker z-scores, and interpreted together with leave-one-marker-out robustness.

Main output:

```text
tables/TRO_interpretability_potency_marker_contribution.tsv
tables/marker_leave_one_out_summary.tsv
```

## Summary

```json
{
  "analysis": "TRO interpretability analysis",
  "top_reset_driving_DMR_table": "TRO_interpretability_top20_reset_driving_DMRs.tsv",
  "top_reset_driving_genes": [
    "LOC100506937",
    "PCDH17",
    "LOC101928551",
    "GMDS-DT",
    "FSHR",
    "SLC3A1",
    "CTXN3",
    "MYH1",
    "DSC1",
    "JAKMIP2",
    "ANK2",
    "CCDC146",
    "HAND2-AS1",
    "RASA3",
    "GPANK1",
    "WNT5B",
    "LINC00320",
    "LINC00578",
    "SMTNL2",
    "IRF9"
  ],
  "top_annotation_rows": [
    {
      "annotation_type": "cpg_context",
      "category": "CpG_shelf",
      "top_count": 4,
      "top_fraction": 0.08,
      "background_count": 7,
      "background_fraction": 0.04487179487179487,
      "p_value": 0.14904358363161255,
      "p_adj_BH": 0.5961743345264502
    },
    {
      "annotation_type": "cpg_context",
      "category": "non_CpG_island",
      "top_count": 29,
      "top_fraction": 0.58,
      "background_count": 88,
      "background_fraction": 0.5641025641025641,
      "p_value": 0.46046330876223995,
      "p_adj_BH": 0.8443860487402599
    },
    {
      "annotation_type": "cpg_context",
      "category": "CpG_island",
      "top_count": 2,
      "top_fraction": 0.04,
      "background_count": 7,
      "background_fraction": 0.04487179487179487,
      "p_value": 0.7182139137243789,
      "p_adj_BH": 0.8443860487402599
    },
    {
      "annotation_type": "cpg_context",
      "category": "CpG_shore",
      "top_count": 15,
      "top_fraction": 0.3,
      "background_count": 54,
      "background_fraction": 0.34615384615384615,
      "p_value": 0.8443860487402599,
      "p_adj_BH": 0.8443860487402599
    },
    {
      "annotation_type": "gene_context",
      "category": "intergenic",
      "top_count": 8,
      "top_fraction": 0.16,
      "background_count": 17,
      "background_fraction": 0.10897435897435898,
      "p_value": 0.1303307426721917,
      "p_adj_BH": 0.5961743345264502
    },
    {
      "annotation_type": "gene_context",
      "category": "intron",
      "top_count": 6,
      "top_fraction": 0.12,
      "background_count": 19,
      "background_fraction": 0.12179487179487179,
      "p_value": 0.612501518054413,
      "p_adj_BH": 0.8443860487402599
    },
    {
      "annotation_type": "gene_context",
      "category": "exon",
      "top_count": 7,
      "top_fraction": 0.14,
      "background_count": 23,
      "background_fraction": 0.14743589743589744,
      "p_value": 0.6566216305221799,
      "p_adj_BH": 0.8443860487402599
    },
    {
      "annotation_type": "gene_context",
      "category": "promoter",
      "top_count": 29,
      "top_fraction": 0.58,
      "background_count": 97,
      "background_fraction": 0.6217948717948718,
      "p_value": 0.8204108047527662,
      "p_adj_BH": 0.8443860487402599
    }
  ],
  "top_marker_contributors": [
    {
      "marker": "SOX2",
      "morula_zscore": 1.2228018781783714,
      "positive_fraction_of_morula_marker_signal": 0.1875659185983834
    },
    {
      "marker": "KLF4",
      "morula_zscore": 0.8385183326455721,
      "positive_fraction_of_morula_marker_signal": 0.1286205591690376
    },
    {
      "marker": "KLF17",
      "morula_zscore": 0.8137231243044211,
      "positive_fraction_of_morula_marker_signal": 0.12481721529760477
    },
    {
      "marker": "NANOG",
      "morula_zscore": 0.7760416128221822,
      "positive_fraction_of_morula_marker_signal": 0.11903723782008352
    },
    {
      "marker": "TFAP2C",
      "morula_zscore": 0.6664639623630831,
      "positive_fraction_of_morula_marker_signal": 0.10222909168210761
    }
  ],
  "interpretation": "The morula ground-zero call is explained by explicit DMR-level entropy reductions from 8-cell to morula plus marker-robust preservation of developmental potency. Nearest-gene and annotation enrichment are exploratory because DMR-to-gene mapping is based on hg19 nearest transcript annotation."
}
```

## Claim boundary

This is a biological interpretability analysis, not a black-box SHAP/LIME analysis. It explains the explicit TRO components by DMR-level entropy reduction, genomic annotation, exploratory nearest-gene function, and potency-marker robustness.
