## Supplemental docs for "An entropy-based diagnostic framework for characterizing methylation state dynamics during preimplantation development": TRO_latent_reset_analysis.md

# Experiment 12: Unsupervised latent-space validation

This analysis uses the sample-by-age-DMR methylation beta matrix from GSE81233 and performs PCA/SVD without using TRO scores as input.

## Key result

- Samples analyzed: 196
- Age-DMR features used: 156
- Best latent axis for sample-level S_epi-age: PC1
- Pearson correlation with sample-level S_epi-age: 0.3616
- Spearman correlation with sample-level S_epi-age: 0.1679
- Morula latent low-age rank: 6
- Morula S_epi-age rank: 1
- Largest centroid transition in latent space: MII oocyte -> zygote/PN

## Representation sensitivity

The direct beta-methylation profile PCA does not make morula a unique latent cluster. However, entropy-profile PCA provides a better alignment with sample-level age-associated methylation entropy and places morula in the low age-entropy latent window.

## Interpretation

This provides a light unsupervised validation of the TRO-defined ground-zero state. It asks whether age-DMR-derived profiles can recover a low age-entropy latent region near morula without using the final TRO score as input.

This result should be described as supporting evidence only. It is not a VAE result and should not be used to claim that a neural operator or Schrodinger Bridge was trained.
