## Supplemental docs for "An entropy-based diagnostic framework for characterizing methylation state dynamics during preimplantation development": TRO_operator_schema.md

# Operational Transgenerational Reset Operator schema

Date: 2026-05-20

## Definition

The operational TRO is defined as:

```text
TRO = {E, D, R, C}
```

Where:

- E is the Entropy Encoder.
- D is the Damage-Potency Decomposer.
- R is the Reset Operator.
- C is the Cost Estimator.

## E: Entropy Encoder

Input:

```text
[S_epi, S_epi-age, S_RNA]
```

Operational meaning:

- `S_epi` captures generic methylation-state entropy.
- `S_epi-age` captures age-associated methylation perturbation entropy.
- `S_RNA` captures global transcriptome entropy.

In the current implementation, `S_epi-age` is the primary damage-related methylation metric.

## D: Damage-Potency Decomposer

The current low-dimensional decomposition is:

```text
D_DamageProxy = S_epi-age
D_PotencyProxy = PotencyScore
D_RNAOrderProxy = -S_RNA
```

Interpretation:

- Lower `D_DamageProxy` is better.
- Higher `D_PotencyProxy` is better.
- `D_RNAOrderProxy` is included because morula does not show high global RNA entropy.

## R: Reset Operator

The reset operator is represented by:

```text
R_ResetScore(g) = (S_MII - S_g) / (S_MII - S_morula)
```

where `S` is `S_epi-age`.

Operationally:

- MII oocyte has reset score 0.
- Morula has reset score 1.
- Intermediate stages are placed between these endpoints.

## C: Cost Estimator

Transition cost is computed between neighboring developmental stages using standardized state-vector distances.

The current transition-level outputs include:

- `C_transition_cost`
- `R_damage_reduction`
- `R_potency_change`
- `R_reset_gain`
- `R_productive_reset_gain`
- `R_reset_efficiency`

The maximum productive reset transition is:

```text
8-cell -> morula
```

## Ground-zero decision rule

The operational ground-zero candidate is the stage satisfying:

```text
GZ_rank = 1
TRO_rank = 1
BioAgeRank = 1
```

In the current analysis:

```text
ground_zero_stage = morula
```

## Current validation summary

The core checks all pass:

- Morula ranks first by GZ score.
- Morula ranks first by TRO score.
- Morula ranks first by operational bio-age score.
- 8-cell -> morula is the maximum productive reset transition.
- DNA robustness tests pass.
- RNA potency tests support morula > blastocyst.
- Leave-one-marker-out robustness passes.
- External GSE44183 validation places morula in the top two potency stages.

## Boundary of interpretation

This is an operational, interpretable, data-driven TRO operator. It is not currently a trained neural operator, Schrödinger Bridge, or wet-lab validated causal mechanism.
