## Supplemental docs for "An entropy-based diagnostic framework for characterizing methylation state dynamics during preimplantation development": TRO_Project_current_conclusion_2026-05-20.md

# TRO Project current conclusion, 2026-05-20

## Project Question

This project tests whether early human embryonic development contains a computational ground-zero state for resetting paternal age-associated epigenetic perturbation.

## Experiment 1: DNA methylation age-entropy reset

### Datasets

- GSE102970: human sperm methylation data.
- GSE81233: human preimplantation embryo methylation data.

GSE102970 GEO metadata did not provide usable age labels in the exposure matrix, so age-associated DMR weights were extracted from Table S6 of the associated paper. GSE81233 originally had 205 planned embryo methylation samples; `GSM2986343_scBS-2C-10-1.Cmet.bed.gz` repeatedly failed gzip integrity checks and was excluded. Final valid sample count: 204.

### Core Metrics

- `S_epi`: unweighted binary methylation entropy.
- `S_epi-age`: age-associated weighted methylation entropy; this is the core age-perturbation metric.
- `ResetScore`: internal relative reset score normalized from MII oocyte to morula.

### Main DNA Result

`S_epi-age` reaches its minimum at morula:

| stage | n samples | S_epi-age |
|---|---:|---:|
| MII oocyte | 36 | 0.4463658605 |
| zygote/PN | 30 | 0.4292531168 |
| 2-cell | 20 | 0.3753126443 |
| 4-cell | 25 | 0.3237260110 |
| 8-cell | 48 | 0.3830423615 |
| morula | 8 | 0.2777817165 |
| blastocyst | 5 | 0.3127181308 |
| ICM | 19 | 0.3473458770 |
| TE | 13 | 0.3219834266 |

Bootstrap ground-zero frequency:

- Morula was the minimum in 1886/2000 bootstrap iterations.
- Frequency: 94.3%.

Adjacent-stage sample-level tests:

- 2-cell vs 4-cell: significant decrease, BH p = 0.0091.
- 8-cell vs morula: significant decrease, BH p = 0.0379.
- morula vs blastocyst: significant increase, BH p = 0.0379.

## Experiment 1B: DNA robustness

Experiment 1B tested whether the morula minimum is explained by stage-specific DMR coverage, sample imbalance, or filtering threshold.

Key robustness results:

| test | ground zero stage | morula frequency | n regions | conclusion |
|---|---|---:|---:|---|
| original valid204 | morula | 0.9430 | 56 | pass |
| common DMR | morula | 0.9155 | 54 | pass |
| min_frac 0.1 | morula | NA | 120 | pass |
| min_frac 0.2 | morula | NA | 80 | pass |
| min_frac 0.3 | morula | NA | 56 | pass |
| min_frac 0.4 | morula | NA | 32 | pass |
| min_frac 0.5 | morula | NA | 32 | pass |
| balanced bootstrap n=5 | morula | 0.7675 | 56 | pass |
| balanced bootstrap n=8 | morula | 0.8590 | 56 | pass |

Conclusion:

> The morula-stage minimum of age-associated methylation entropy is robust to common-DMR restriction, coverage-threshold variation, and balanced bootstrap resampling.

Important limitation:

- Shuffled-weight and random age-DMR subset controls still often selected morula as the minimum.
- Therefore, these controls support robustness within the age-DMR feature space, but do not prove strict specificity of the exact age-weight ordering.

## Experiment 2: RNA entropy and potency-marker validation

### Dataset

- GSE36552
- Yan et al. 2013 NSMB Supplementary Table 1 RPKM matrix.

RNA cells retained by stage:

| stage | n cells |
|---|---:|
| oocyte | 4 |
| zygote | 4 |
| 2-cell | 19 |
| 4-cell | 9 |
| 8-cell | 13 |
| morula | 17 |
| blastocyst | 39 |

### Revised RNA Conclusion

The original expectation that morula would show high global RNA entropy was not supported.

Instead:

- Morula has the lowest mean `S_RNA` among the RNA stages analyzed.
- Morula retains high developmental potency-marker activity.

| stage | S_RNA | PotencyScore |
|---|---:|---:|
| oocyte | 7.449760 | 0.380039 |
| zygote | 7.392980 | 0.414669 |
| 2-cell | 7.071784 | 0.517773 |
| 4-cell | 7.209825 | 0.402878 |
| 8-cell | 7.008319 | 0.635407 |
| morula | 6.871445 | 0.606959 |
| blastocyst | 7.011921 | 0.358131 |

### Potency Robustness

Morula has significantly higher marker and potency scores than blastocyst:

| comparison | metric | mean morula | mean blastocyst | BH-adjusted p | Cliff's delta |
|---|---|---:|---:|---:|---:|
| morula vs blastocyst | marker_score | 0.823540 | 0.435908 | 4.018e-05 | 0.773756 |
| morula vs blastocyst | potency_score | 0.606959 | 0.358131 | 4.018e-05 | 0.785822 |

Morula is statistically comparable to 8-cell:

| comparison | metric | mean morula | mean 8-cell | BH-adjusted p |
|---|---|---:|---:|---:|
| morula vs 8-cell | marker_score | 0.823540 | 0.824444 | 0.261804 |
| morula vs 8-cell | potency_score | 0.606959 | 0.635407 | 0.807988 |

Marker-level support at morula includes positive z-scores for:

- `POU5F1`
- `NANOG`
- `SOX2`
- `KLF4`
- `KLF17`
- `TFAP2C`
- `GDF3`

## Fixed Current Conclusion

Do not claim:

> morula RNA entropy is high.

Current fixed conclusion:

> Morula represents a computational ground-zero candidate characterized by minimal age-associated methylation entropy and preserved high developmental potency-marker activity.

## Recommended Paper-Level Sentence

Age-associated methylation entropy was progressively reduced during preimplantation development and reached a robust minimum at the morula stage. RNA-seq analysis did not support elevated global transcriptomic entropy at morula; instead, morula retained high developmental potency-marker activity, with potency scores significantly higher than blastocyst and comparable to 8-cell. These results support a refined dual-state model in which the morula ground-zero candidate combines minimal age-associated methylation entropy with preserved developmental potency-marker activity.

## Cautions

- Do not say public data prove that one older father's sperm was reset in a matched embryo.
- Do not call `S_epi` aging entropy.
- `S_epi-age` is the age-associated perturbation metric.
- `ResetScore` is an internal normalization, not a strict young-vs-old paternal reset index.
- The current RNA potency score is a proxy based on detected genes and selected developmental markers, not CytoTRACE or SCENT.

## Key Output Files

- `notes/Experiment1B_DNA_robustness_interpretation.md`
- `notes/Experiment2_RNA_dual_state_interpretation.md`
- `tables/GSE81233_valid204_stage_epi_age_metrics.tsv`
- `tables/Experiment1B_DNA_robustness_summary.tsv`
- `tables/GSE36552_RNA_entropy_potency_by_stage.tsv`
- `tables/GSE36552_potency_component_by_stage.tsv`
- `tables/GSE36552_potency_pairwise_tests.tsv`
- `tables/dual_entropy_stage_table.tsv`
- `figures/Experiment1B_DNA_robustness_summary.png`
- `figures/GSE36552_marker_heatmap.png`
- `figures/GSE36552_potency_component_by_stage.png`
- `figures/GSE36552_potency_bootstrap_ci.png`
- `figures/dual_entropy_phase_map.png`
