## Supplemental docs for "An entropy-based diagnostic framework for characterizing methylation state dynamics during preimplantation development": TRO_result_file_manifest_2026-05-20.md

# TRO result file manifest

Date: 2026-05-20

## Final core tables

- `tables/GSE81233_valid204_stage_epi_age_metrics.tsv`
- `tables/GSE81233_valid204_bootstrap_ground_zero_frequency.tsv`
- `tables/GSE81233_valid204_adjacent_stage_mannwhitney.tsv`
- `tables/GSE81233_valid204_internal_reset_score.tsv`
- `tables/Experiment1B_DNA_robustness_summary.tsv`
- `tables/GSE36552_RNA_entropy_potency_by_stage.tsv`
- `tables/GSE36552_potency_component_by_stage.tsv`
- `tables/GSE36552_potency_pairwise_tests.tsv`
- `tables/dual_entropy_stage_table.tsv`
- `tables/TRO_composite_score_by_stage.tsv`
- `tables/marker_leave_one_out_summary.tsv`
- `tables/expanded_marker_panel_potency_score.tsv`
- `tables/GSE44183_external_potency_validation.tsv`
- `tables/GSE44183_external_potency_pairwise_tests.tsv`
- `tables/TRO_stage_state_vectors.tsv`
- `tables/TRO_stage_transition_cost.tsv`
- `tables/TRO_reset_depth_summary.tsv`
- `tables/TRO_operator_stage_output.tsv`
- `tables/TRO_operator_transition_output.tsv`
- `tables/TRO_operator_summary.json`

## Final core figures

- `figures/GSE81233_valid204_s_epi_age_by_stage.png`
- `figures/GSE81233_valid204_internal_reset_score.png`
- `figures/GSE81233_valid204_sample_level_s_epi_age_boxplot.png`
- `figures/Experiment1B_DNA_robustness_summary.png`
- `figures/GSE36552_RNA_entropy_by_stage.png`
- `figures/GSE36552_potency_score_by_stage.png`
- `figures/GSE36552_marker_heatmap.png`
- `figures/dual_entropy_phase_map_final.png`
- `figures/TRO_composite_score_by_stage.png`
- `figures/marker_leave_one_out_summary.png`
- `figures/GSE44183_external_potency_validation.png`
- `figures/TRO_damage_potency_state_space.png`
- `figures/TRO_stage_transition_cost.png`
- `figures/TRO_transition_component_changes.png`
- `figures/TRO_operator_diagram.png`

PDF versions are available for most final figures in `figures/`.

## Final notes

- `notes/Experiment1B_DNA_robustness_interpretation.md`
- `notes/Experiment2_RNA_dual_state_interpretation.md`
- `notes/TRO_Project_current_conclusion_2026-05-20.md`
- `notes/TRO_final_results_interpretation_2026-05-20.md`
- `notes/TRO_result_file_manifest_2026-05-20.md`

## Final scripts

- `scripts/12_experiment1b_dna_robustness.py`
- `scripts/13_experiment2a_gse36552_rna_entropy_potency.py`
- `scripts/14_experiment2b_rna_potency_robustness.py`
- `scripts/15_experiment3_tro_composite_and_marker_robustness.py`
- `scripts/16_experiment4_gse44183_external_rna_validation.py`
- `scripts/17_experiment5_transition_reset_cost.py`
- `scripts/18_experiment6_full_operational_tro_operator.py`

## Minimum figure set for a manuscript draft

1. `GSE81233_valid204_s_epi_age_by_stage`
2. `Experiment1B_DNA_robustness_summary`
3. `GSE36552_marker_heatmap`
4. `dual_entropy_phase_map_final`
5. `TRO_composite_score_by_stage`
6. `TRO_stage_transition_cost`
7. `TRO_operator_diagram`

## Recommended main-text figure logic

Figure 1:

DNA methylation age-entropy reset and morula ground-zero candidate.

Figure 2:

RNA potency-marker preservation and dual-entropy phase map.

Figure 3:

TRO composite score and marker robustness.

Figure 4:

Operational TRO operator and reset transition cost.
