## Supplemental environment for "An entropy-based diagnostic framework for characterizing methylation state dynamics during preimplantation development": environment.yml

```
name: csb-tro-supplement
channels:
  - conda-forge
  - defaults
dependencies:
  - python>=3.11
  - pip
  - numpy
  - pandas
  - scipy
  - matplotlib
  - seaborn
  - scikit-learn
  - statsmodels
  - openpyxl
  - requests
  - tqdm
  - pip:
      - gseapy
```
