## Supplemental other for "An entropy-based diagnostic framework for characterizing methylation state dynamics during preimplantation development": CLAIM_TO_FILE_MAP.md

# Manuscript claim-to-file map

Paths are relative to the archive root.

| Results claim | Primary data/result files | Main analysis code |
|---|---|---|
| Morula is the minimum of age-DMR-weighted methylation entropy in the valid GSE81233 cohort | `data/processed_core/GSE81233_valid204_stage_epi_age_metrics.tsv`; `GSE81233_valid204_ground_zero_summary.tsv`; `GSE81233_valid204_stage_sample_counts.tsv` | `code/01_core_stage/10_finalize_gse81233_valid204.py`; `11_stats_and_reset_valid204.py` |
| The morula call is robust to DMR universe, coverage, sample balance and weight controls | `data/processed_core/Experiment1B_*`; `GSE81233_valid204_bootstrap_*` | `code/01_core_stage/12_experiment1b_dna_robustness.py` |
| Morula preserves RNA-derived developmental potency and the finding is not driven by one marker | `data/processed_core/GSE36552_RNA_entropy_potency_by_stage.tsv`; `GSE36552_potency_component_by_stage.tsv`; `marker_leave_one_out_summary.tsv` | `code/01_core_stage/13_experiment2a_gse36552_rna_entropy_potency.py`; `14_experiment2b_rna_potency_robustness.py` |
| Independent RNA data support an 8-cell/morula high-potency window | `data/processed_core/GSE44183_external_potency_validation.tsv` and pairwise tests | `code/01_core_stage/16_experiment4_gse44183_external_rna_validation.py` |
| Morula ranks first by the stage-level ground-zero and TRO scores; 8-cell-to-morula has the highest productive reset efficiency | `data/processed_core/TRO_operator_stage_output.tsv`; `TRO_operator_transition_output.tsv`; `TRO_operator_summary.json` | `code/01_core_stage/17_experiment5_transition_reset_cost.py`; `18_experiment6_full_operational_tro_operator.py` |
| A structured subset of DMRs carries most of the 8-cell-to-morula reset displacement | `data/processed_core/TRO_interpretability_DMR_contribution_ranking.tsv`; `TRO_interpretability_top20_reset_driving_DMRs.tsv`; `TRO_interpretability_top50_reset_driving_DMRs.tsv` | `code/01_core_stage/23_experiment11_tro_interpretability_analysis.py`; `30_experiment18_dmr_mechanistic_interpretability.py` |
| Module/latent representation improves strict morula prediction over independent DMR drift | `data/processed_operator/CSB_TRO_module_latent_model_comparison.tsv`; `CSB_TRO_module_latent_bootstrap_CI.tsv`; `CSB_TRO_module_latent_paired_error_tests.tsv`; `CSB_TRO_module_latent_null_summary.tsv` | `code/03_dmr_operator/run_module_latent_validation.py`; `run_module_latent_operator_time_dynamics.py` |
| Methylation-only distribution dynamics fail to recover the non-leaking morula basin | `data/processed_operator/CSB_TRO_nonleaking_distribution_metrics.tsv`; `CSB_TRO_morula_basin_sde_metrics.tsv` | `code/03_dmr_operator/run_nonleaking_distribution_dynamics.py`; `run_morula_basin_sde.py` |
| The missing correction is concentrated in ranked DMRs/modules and is direction/dose sensitive | `data/processed_operator/CSB_TRO_missing_control_term_*`; `CSB_TRO_residual_DMR_topK_occupancy_curve.tsv`; `CSB_TRO_residual_DMR_remove_topK_necessity.tsv`; `CSB_TRO_residual_DMR_sign_flip_control.tsv`; `CSB_TRO_residual_DMR_matched_random_control.tsv` | `code/03_dmr_operator/run_missing_control_term_decomposition.py`; `run_residual_dmr_robustness.py` |
| Morula has an entry-exit turning-gate geometry and branch-specific exit logic | `data/processed_dynamics/CSB_TRO_2026-05-27_entry_exit_*`; `5_29_progress__partB_*`; `5_29_progress__remeth_*` | `code/04_dynamics_hardening/*entry_exit*`; `5_29_progress__solve_morula_inflection_and_blast.py`; `5_29_progress__remeth_prediction_framework.py` |
| Accessibility associations are stage restricted and provide bounded support for the inferred correction | `data/processed_dynamics/CSB_TRO_2026-05-27_duality_accessibility_*`; `5_28_progress__CSB_TRO_5_28_dmr_multistage_accessibility.tsv`; `5_28_progress__CSB_TRO_5_28_dmr_quantitative_accessibility.tsv` | `code/04_dynamics_hardening/5_28_progress__multistage_accessibility_dynamics.py`; archived 2026-05-27 analysis scripts |
| COMSOL scenarios separate baseline failure, morula entry, successful exit and wrong-direction collapse | `models/comsol/*.mph`; `data/processed_comsol/traj_final_*.csv`; `scenario_results_final.json`; `PHASE_A_FINAL_REPORT.json` | `code/05_comsol/run_all_scenarios.py`; `create_comsol_models.py`; Java model files |
| Independent human GSE49828 provides directional sperm-to-morula support | `data/processed_core/GSE49828_gamete_to_embryo_directional_stage_metrics.tsv`; `GSE49828_gamete_to_embryo_directional_summary.json` | `code/01_core_stage/31_experiment19_gse49828_gamete_to_embryo_directional_validation.py` |
| Paired mouse GSE56697 supports a real gamete-to-embryo operator with stable bin-size sensitivity | `data/processed_core/GSE56697_paired_paternal_operator_*`; `GSE56697_maternal_paternal_branch_*` | `code/01_core_stage/26_experiment14_*`; `27_experiment15_*`; `28_experiment16_*` |
| Gene-matched cross-species analysis gives qualified, weight-dependent mouse morula support | `data/processed_external/crossspecies_mouse_gleaner_*` | `code/06_external_validation/make_counterfactual_and_crossspecies.py` |
| Orthogonal trophoblast/embryo-model contrasts provide heterogeneous, bounded evidence rather than causal closure | `data/processed_external/CSB_TRO_integrated_evidence/results/*`; dataset-specific `results/*.tsv` and `*.json` | `code/06_external_validation/CSB_TRO_integrated_evidence/scripts/01_build_integrated_evidence_matrix.py` and dataset-specific scripts |
