## Supplemental other for "An entropy-based diagnostic framework for characterizing methylation state dynamics during preimplantation development": DATA_AND_CODE_AVAILABILITY.md

# Submission text requiring author completion

## Data Availability

The data analysed in this study were obtained from public repositories under accession codes GSE102970, GSE81233, GSE36552, GSE44183, GSE49828, GSE56697, GSE101571, GSE109682, GSE126958, GSE150168, GSE182015, GSE207222, GSE247631, GSE266195, GSE280039, GSE291172, E-MTAB-10096 and E-MTAB-10097, together with the public mouse GLEANER mm9 methylation matrix. The accession-to-analysis mapping, sample manifests, quality-control records and processed data underlying the reported statistics and figures are provided in the accompanying reproducibility archive and at **[ZENODO OR OTHER DATA-REPOSITORY DOI]**. Large third-party raw sequencing files are not redistributed in the GitHub repository and remain available from their original repositories. One GSE81233 file, GSM2986343_scBS-2C-10-1.Cmet.bed.gz, failed gzip validation and was excluded before analysis. Controlled-access HRA002355/PRJCA009410 records were used only to define a data-access/claim boundary; controlled files are not included in the public archive.

## Code Availability

Custom Python, R, shell and COMSOL scripts used for the analyses are available at **[GITHUB RELEASE URL]** and archived with a persistent identifier at **[ZENODO OR CODE-REPOSITORY DOI]**. The exact publication snapshot includes processed source-data tables, run summaries, deterministic seeds, COMSOL Multiphysics 6.4 model files and SHA-256 checksums. Full raw-data regeneration requires downloading the third-party datasets listed in the archive documentation. COMSOL model regeneration requires a compatible COMSOL Multiphysics licence.
