## Supplemental other for "An entropy-based diagnostic framework for characterizing methylation state dynamics during preimplantation development": DATA_SOURCES.md

# Data sources and redistribution policy

All primary biological inputs were pre-existing public or controlled-access datasets. This package redistributes compact derived tables, sample manifests and analysis outputs, not the large third-party raw sequencing files.

| Source | Role in the manuscript | Access route | Included here |
|---|---|---|---|
| GSE102970 | Published sperm-age DMR/CpG weights | https://www.ncbi.nlm.nih.gov/geo/query/acc.cgi?acc=GSE102970 | Extracted Table S6 weight tables |
| GSE81233 | Primary human preimplantation single-cell methylome | https://www.ncbi.nlm.nih.gov/geo/query/acc.cgi?acc=GSE81233 | Valid-sample manifests, processed DMR/stage tables and QC; raw Cmet files excluded |
| GSE36552 | Primary human preimplantation single-cell transcriptome | https://www.ncbi.nlm.nih.gov/geo/query/acc.cgi?acc=GSE36552 | Processed cell/stage entropy and potency tables |
| GSE44183 | Independent transcriptomic validation | https://www.ncbi.nlm.nih.gov/geo/query/acc.cgi?acc=GSE44183 | Processed within-dataset validation tables |
| GSE49828 | Independent human sperm/embryo RRBS directional validation | https://www.ncbi.nlm.nih.gov/geo/query/acc.cgi?acc=GSE49828 | DMR-level processed values, QC, stage metrics and summaries; raw RRBS files excluded |
| GSE56697 | Paired mouse parental-allele methylome operator | https://www.ncbi.nlm.nih.gov/geo/query/acc.cgi?acc=GSE56697 | Window-level processed matrix, stage/transition metrics and robustness summaries; raw methylomes excluded |
| GSE101571 | Human preimplantation accessibility support | https://www.ncbi.nlm.nih.gov/geo/query/acc.cgi?acc=GSE101571 | DMR overlap and stage-level processed summaries |
| GSE109682 | Human trophoblast RRBS contrast | https://www.ncbi.nlm.nih.gov/geo/query/acc.cgi?acc=GSE109682 | Sample sheets, DMR summaries and QC |
| GSE126958 | TET/DNMT3 machinery perturbation support | https://www.ncbi.nlm.nih.gov/geo/query/acc.cgi?acc=GSE126958 | DMR-level processed table and null summaries |
| GSE150168 | Naive hESC/trophoblast methylome contrast | https://www.ncbi.nlm.nih.gov/geo/query/acc.cgi?acc=GSE150168 | DMR-level processed table and QC |
| GSE182015 | hiTSC/hbdTSC RRBS contrast | https://www.ncbi.nlm.nih.gov/geo/query/acc.cgi?acc=GSE182015 | DMR-level processed table and QC |
| GSE207222 | Early-embryo A-485 accessibility perturbation consistency | https://www.ncbi.nlm.nih.gov/geo/query/acc.cgi?acc=GSE207222 | Processed projection/summary files only |
| GSE247631 | Naive embryo-lineage A-485/2-HG-related transcriptomic perturbation | https://www.ncbi.nlm.nih.gov/geo/query/acc.cgi?acc=GSE247631 | Compact gene-effect and summary tables; large null/all-gene tables excluded |
| GSE266195 | hTSC DNMT3L PBAT methylome perturbation | https://www.ncbi.nlm.nih.gov/geo/query/acc.cgi?acc=GSE266195 | DMR-level contrast and summary |
| GSE280039 | hTSC induction methylome contrast | https://www.ncbi.nlm.nih.gov/geo/query/acc.cgi?acc=GSE280039 | DMR-level contrast and summary |
| GSE291172 | STAT3-induced embryo-model WGBS contrast | https://www.ncbi.nlm.nih.gov/geo/query/acc.cgi?acc=GSE291172 | DMR-level contrast and summary |
| E-MTAB-10096/10097 | Human embryo expression and bisulfite-sequencing source audits | https://www.ebi.ac.uk/biostudies/arrayexpress/studies/E-MTAB-10096 and https://www.ebi.ac.uk/biostudies/arrayexpress/studies/E-MTAB-10097 | Sample/run manifests, scripts and compact derived tables; FASTQ/BAM/reference indices excluded |
| Mouse GLEANER mm9 methylation matrix | Gene-matched cross-species diagnostic | https://compbio-zhanglab.org/GLEANER/download/mm9_me.txt | Matched-gene/stage results and nulls; original third-party matrix not redistributed |
| HRA002355/PRJCA009410 | Controlled-access human embryo histone source audit | Original controlled-access repository record | Access manifests/claim-boundary notes only; controlled files not included |

## Known source exclusions

- `GSM2986343_scBS-2C-10-1.Cmet.bed.gz` failed gzip validation repeatedly and was excluded before the GSE81233 analysis. The final primary stage analysis used 204 technically valid Cmet profiles.
- Raw FASTQ, BAM, SAM, CRAM, bigWig, Cmet/RRBS/WGBS/PBAT files, genome FASTA files, Bismark/Bowtie indices and conda environments are excluded because they are large, third-party data or reproducible from public repositories.
- Large stochastic particle tables and transport-training-pair tables are excluded from the GitHub package. Seeds, compact summaries, coefficients and analysis scripts are retained.

Third-party data remain subject to their original repository terms. No new licence is asserted over third-party source data.
