## Supplemental other for "An entropy-based diagnostic framework for characterizing methylation state dynamics during preimplantation development": FILE_SELECTION.md

# File-selection record

## Included

- Original archived analysis scripts and small command records.
- Age-DMR weights, sample/stage mappings, public accession manifests and QC tables.
- Processed DMR-, sample-, stage- and transition-level data supporting the manuscript.
- Bootstrap/null summaries, paired-error tests, effect tables and claim-boundary records.
- All archived COMSOL `.mph` models and compact trajectory/parameter exports.
- Compact cross-species and orthogonal-validation results.
- Small SVG reference outputs for visual comparison.

## Excluded to keep the package suitable for GitHub and journal review

- Approximately 100 GB of raw/intermediate material from the large development workspace.
- Approximately 31 GB of causal-chain raw/reference/process files.
- Approximately 3.2 GB of GSE49828/GSE56697 raw methylome files.
- Reference genomes, Bismark/Bowtie indices, conda environments, caches and compiled bytecode.
- The 79.5 MB operator-training-pair table, 55.4 MB transport-coupling table and large particle-level simulation tables. Scripts, seeds, coefficients and compact result summaries are retained.
- Duplicate PNG/PDF exports where a compact SVG or the underlying numerical table is available.
- Large third-party all-gene/random-null tables when a compact summary, seed and regeneration script are present.

No source file was modified in place. Files in this archive are copies selected from the existing project workspaces.
