## Supplemental other for "An entropy-based diagnostic framework for characterizing methylation state dynamics during preimplantation development": LICENSE_NOTICE.md

# Licence status

No open-source software licence or derived-data licence was present in the supplied project archives, so none has been invented during packaging.

Before public release, the authors should:

1. choose an OSI-approved licence for original code, subject to institutional policy;
2. choose an appropriate licence for author-generated derived tables;
3. confirm that third-party repository terms permit redistribution of any included metadata or derived content; and
4. add the selected licence files to the DOI-archived release.

Until then, normal copyright applies and the absence of a licence should not be interpreted as permission to reuse the code.
