## Supplemental other for "An entropy-based diagnostic framework for characterizing methylation state dynamics during preimplantation development": NATURE_SUBMISSION_CHECKLIST.md

# Nature-oriented submission checklist

This GitHub reproducibility archive is not a replacement for the journal's formatted Supplementary Information PDF.

- Nature's general SI guidance recommends combining flat Supplementary Methods, Tables, Discussion, Equations and Notes into one PDF, with editable Supplementary Data supplied separately where appropriate.
- Nature prefers no more than ten SI files; single PDFs and audiovisual files may be up to 30 MB each, with a 150 MB cumulative maximum under the general Nature guidance.
- Nature's final-submission guidance may require an `SIGuide.doc` containing a title and a summary of no more than 50 words for each SI file.
- Large datasets should normally be deposited in an appropriate repository rather than embedded in SI.
- A GitHub publication snapshot should be released and archived in a DOI-minting repository such as Zenodo; cite the DOI in the manuscript.
- GitHub browser uploads are limited to 25 MiB per file; command-line Git warns above 50 MiB and blocks files above 100 MiB. Keep the final ZIP below 25 MiB if browser upload is intended.
- Confirm the exact target journal's current author instructions, because individual Nature Portfolio journals can differ.

Official guidance:

- https://www.nature.com/nature/for-authors/supp-info
- https://www.nature.com/nature/for-authors/final-submission
- https://www.nature.com/nature-portfolio/editorial-policies/reporting-standards
- https://docs.github.com/en/repositories/working-with-files/managing-large-files/about-large-files-on-github
