## Supplemental other for "An entropy-based diagnostic framework for characterizing methylation state dynamics during preimplantation development": README.md

# CSB-TRO reproducibility supplement

This archive is the lightweight, GitHub-oriented reproducibility package for the manuscript's transgenerational reset operator (TRO/CSB-TRO) analyses. It contains only files already present in the archived project workspaces: custom code, processed/source-data tables, public-data manifests, COMSOL models and small reference outputs. No experimental values were invented or imputed while assembling this package.

## Start here

1. Read `DATA_SOURCES.md` for the public accessions and files that are intentionally not redistributed.
2. Read `CLAIM_TO_FILE_MAP.md` to locate the data and code underlying each Results claim.
3. Read `REPRODUCIBILITY.md` before attempting a rerun.
4. Run `python verify_package.py` to verify file presence, checksums and GitHub-size constraints.

## Directory layout

```text
code/
  01_core_stage/          Core DNA/RNA entropy, potency and stage-level TRO analyses
  02_stage_transport/     Stage-level transport and distribution analyses
  03_dmr_operator/        DMR-resolved operator-time, latent, basin and control analyses
  04_dynamics_hardening/  Entry/exit, accessibility, remethylation and counterfactual analyses
  05_comsol/              COMSOL model-generation/export scripts
  06_external_validation/ Cross-species and orthogonal public-data analyses
data/
  metadata/               Public-data manifests, sample mappings and age-DMR weights
  processed_*/            Analysis-ready and result tables; raw public sequencing data excluded
models/comsol/             Archived COMSOL Multiphysics 6.4 `.mph` models
docs/source_notes/         Archived interpretation, methods and claim-boundary notes
figures/reference_outputs/ Small SVG reference outputs for result checking
environment/               Conda environment specifications
```

## Reproducibility levels

- **Audit level:** all central manuscript numbers can be traced to bundled TSV/CSV/JSON files and checked without downloading raw sequencing data.
- **Processed-data rerun:** most Python analyses can be rerun after restoring the documented directory layout and updating archived path constants.
- **Full raw-data regeneration:** requires downloading the public GEO/ArrayExpress/GLEANER source files listed in `DATA_SOURCES.md`. COMSOL figures require COMSOL Multiphysics 6.4.

The archived scripts preserve their original analysis-time path constants. They are included unchanged for provenance; users must edit the path block or command-line arguments in the relevant script when running outside the original workstation.

## Scientific interpretation boundary

The package supports a computational morula ground-zero candidate with low age-DMR-weighted methylation entropy, preserved developmental potency and structured operator-time correction. It does not establish a matched human father-sperm-embryo lineage, a unique in-vivo molecular controller, or direct causal rejuvenation. RNA and DNA profiles are integrated at developmental-stage level and are not paired multi-omic measurements from the same embryos.

## Publication deposit

GitHub is suitable for version control and reviewer access, but the publication snapshot should also be archived in a DOI-minting repository such as Zenodo. Replace the placeholders in `DATA_AND_CODE_AVAILABILITY.md` only after the GitHub release URL and archive DOI exist.
