## Supplemental other for "An entropy-based diagnostic framework for characterizing methylation state dynamics during preimplantation development": REPRODUCIBILITY.md

# Reproducibility guide

## 1. Environment

Create the consolidated environment:

```bash
conda env create -f environment/environment.yml
conda activate csb-tro-supplement
python verify_package.py
```

COMSOL `.mph` files were generated with COMSOL Multiphysics 6.4.0.293. They can be inspected without rerunning Python analyses, but regeneration requires a compatible COMSOL installation and licence.

## 2. Package audit without raw data

The compact tables are sufficient to audit manuscript values, sample counts, exclusions, uncertainty summaries and null-control outputs. Use `CLAIM_TO_FILE_MAP.md`, then inspect the referenced TSV/JSON files. `verify_package.py` checks archive integrity but does not re-estimate scientific results.

## 3. Restoring public source data

Download the required datasets from the accession pages in `DATA_SOURCES.md`. Keep raw and processed data separate. Do not rename sample identifiers. For GSE81233, exclude the documented corrupted file before finalization. For the GLEANER analysis, save the downloaded matrix as `mm9_me.txt` beside the cross-species script or update its `OUT`/input path.

## 4. Analysis order

1. `code/01_core_stage`: extract age-DMR weights; aggregate GSE81233; calculate entropy, robustness, RNA potency, stage-level TRO, GSE49828 and GSE56697 validations.
2. `code/02_stage_transport`: build the stage-level transport/distribution summaries from `data/processed_transport` inputs.
3. `code/03_dmr_operator`: create the 156-DMR operator-time state, strict stage-exclusion prediction, module/latent validation, morula-basin diagnostics, residual correction and control analyses.
4. `code/04_dynamics_hardening`: run entry/exit, accessibility, branch-direction, dose-response and counterfactual stress tests.
5. `code/05_comsol`: reconstruct the five COMSOL scenarios and compare exported trajectories with `data/processed_comsol`.
6. `code/06_external_validation`: reproduce the cross-species and heterogeneous orthogonal evidence summaries.

## 5. Archived path constants

The code is an unchanged provenance snapshot from several analysis workspaces. Many scripts contain absolute Windows or Linux paths. Before execution, inspect the path/config block near the top of each script and replace it with paths on the local system. Scripts with command-line arguments should be preferred where available. This package does not claim a single turnkey command for all raw-data analyses.

## 6. Determinism and statistical controls

- The main strict latent validation and morula-basin simulations record seed `20260525`.
- The archived strict validation used 200 null replicates and 1,000 bootstrap resamples.
- Stage uncertainty generally used 1,000 within-stage bootstrap iterations unless a module-specific record states otherwise.
- Random coupling, time/module randomization, matched-random DMR sets, shuffled weights and sign reversals are stored as distinct result tables rather than being merged with observed results.

## 7. Expected boundaries

- DNA and RNA datasets are independent and aligned by stage, not by embryo.
- Optimal transport is used for distribution-aware interpolation, not lineage tracking.
- Target-calibrated morula-basin fits are diagnostic upper bounds.
- COMSOL realizes the archived fitted field; it is not independent evidence of a physical force.
- Controlled-access or unavailable histone tracks are represented only by access/source audits and must not be treated as public packaged data.
