## Supplemental other for "An entropy-based diagnostic framework for characterizing methylation state dynamics during preimplantation development": verify_package.py

```
#!/usr/bin/env python3
"""Verify the packaged archive without modifying scientific data."""

from __future__ import annotations

import hashlib
from pathlib import Path
import sys


ROOT = Path(__file__).resolve().parent
CHECKSUM_FILE = ROOT / "SHA256SUMS.txt"
REQUIRED = [
    "README.md",
    "DATA_SOURCES.md",
    "CLAIM_TO_FILE_MAP.md",
    "REPRODUCIBILITY.md",
    "data/processed_core/GSE81233_valid204_stage_epi_age_metrics.tsv",
    "data/processed_core/TRO_operator_summary.json",
    "data/processed_operator/CSB_TRO_module_latent_model_comparison.tsv",
    "data/processed_operator/CSB_TRO_nonleaking_distribution_metrics.tsv",
    "data/processed_comsol/scenario_results_final.json",
    "data/processed_external/crossspecies_mouse_gleaner_summary.json",
    "data/processed_external/CSB_TRO_integrated_evidence/results/CSB_TRO_integrated_evidence_matrix.tsv",
]


def sha256(path: Path) -> str:
    digest = hashlib.sha256()
    with path.open("rb") as handle:
        for chunk in iter(lambda: handle.read(1024 * 1024), b""):
            digest.update(chunk)
    return digest.hexdigest()


def main() -> int:
    errors: list[str] = []
    for rel in REQUIRED:
        if not (ROOT / rel).is_file():
            errors.append(f"missing required file: {rel}")

    files = [p for p in ROOT.rglob("*") if p.is_file()]
    for path in files:
        rel = path.relative_to(ROOT).as_posix()
        if path.stat().st_size == 0:
            errors.append(f"zero-byte file: {rel}")
        if path.stat().st_size > 50 * 1024 * 1024:
            errors.append(f"file exceeds the package's 50 MiB warning threshold: {rel}")

    if not CHECKSUM_FILE.is_file():
        errors.append("missing SHA256SUMS.txt")
    else:
        for line in CHECKSUM_FILE.read_text(encoding="utf-8").splitlines():
            if not line.strip():
                continue
            expected, rel = line.split("  ", 1)
            target = ROOT / rel
            if not target.is_file():
                errors.append(f"checksum target missing: {rel}")
            elif sha256(target) != expected:
                errors.append(f"checksum mismatch: {rel}")

    total = sum(p.stat().st_size for p in files)
    print(f"files={len(files)} total_MiB={total / (1024 * 1024):.2f}")
    if errors:
        print("FAILED")
        for error in errors:
            print(f"- {error}")
        return 1
    print("OK: required files, size gate and SHA-256 checksums passed")
    return 0


if __name__ == "__main__":
    sys.exit(main())
```
